# Loss of KHSRP Increases Neuronal Growth and Synaptic Transmission and Alters Memory Consolidation Through RNA Stabilization

**DOI:** 10.1101/2020.10.25.354076

**Authors:** Sarah L. Olguin, Priyanka Patel, Michela Dell’Orco, Amy S. Gardiner, Robert Cole, Courtney Buchanan, Anitha Sundara, Joann Mudge, Andrea M. Allan, Pavel Ortinski, Jonathan L. Brigman, Jeffery L. Twiss, Nora I. Perrone-Bizzozero

## Abstract

The KH-type splicing regulatory protein (KHSRP) is an RNA-binding protein linked to decay of AU-rich element containing mRNAs. We have previously shown that KHSRP destabilizes the mRNA encoding the growth-associated protein GAP-43 and decreases neurite growth in cultured embryonic neurons. In contrast, loss of KHSRP stabilizes *Gap43* mRNA and increases neurite growth. Here, we have tested functions of neural KHSRP *in vivo*. We find upregulation of 1460 mRNAs in the neocortex of adult *Khsrp*^−/−^ mice, of which 527 bind to KHSRP with high specificity. These KHSRP targets are involved in pathways for neuronal morphology, axon guidance, neurotransmission and long-term memory. Neocortical neurons show increased axon growth and dendritic spine density in *Khsrp*^−/−^ mice. Analyses of neuronal cultures from embryonic *Khsrp*^−/−^ mice point to a neuron-intrinsic alteration in axonal and dendritic growth and elevations in KHSRP-target mRNAs, including subcellularly localized mRNAs. Hippocampus and infralimbic cortex of *Khsrp*^−/−^ mice show presynaptic elevations in neurotransmission. The *Khsrp*^−/−^ mice have significant deficits in both trace conditioning and attention set-shifting tasks compared *Khsrp*^+/+^ mice, indicating impaired prefrontal- and hippocampal-dependent memory consolidation with loss of KHSRP. Overall, our results indicate that prenatal deletion of KHSRP impairs neuronal development resulting in alterations in neuronal morphology and function by changing post-transcriptional control of neuronal gene expression.

## INTRODUCTION

Post-transcriptional regulation of gene expression plays a critical role in neuronal differentiation and function. Independent from transcription and translation, these mechanisms are especially important in control of specific sets of mRNAs that localize into dendrites and axons (Holt et al., 2019). Stability of mRNAs is also critically important for the regulation of gene expression, as changes in mRNA decay rates can be rapid and precise. mRNA sequences (*cis*-elements) are bound by *trans*-acting factors like RNA binding proteins (RBPs) and miRNAs to effect changes in mRNA decay rates (Bolognani and Perrone-Bizzozero, 2008). Binding by RBPs can stabilize an mRNA by protecting it from nucleases, promote its translation by targeting the mRNA to polysomes, or promote its decay by targeting the bound mRNA to RNA degradation sites in the cell (Szostak and Gebauer, 2013).

The KH-type splicing regulatory protein (KHSRP; also known as KSRP, FUBP2, ZBP2, and MARTA1) is an RBP implicated in decay of AU-rich element (ARE)-containing mRNAs by targeting them to the cytoplasmic exosome complex for degradation (Chen et al., 2001; Gherzi et al., 2004). KHSRP was independently discovered as a single-stranded DNA binding protein, termed Far Upstream Element (FUSE) binding protein 2 (FUBP2), and a KH-homology RBP that enhanced splicing of the neuron-specific *c-Src* N1 exon (Davis-Smyth et al., 1996; Min et al., 1997). KHSRP’s function has been linked to disease conditions including viral infections, diabetes and cancer (Briata et al., 2016), but KHSRP is also highly expressed in neural tissues, including in neurons, and it localizes into both axons and dendrites (Snee et al., 2002). The KHSRP orthologue in rats, MARTA1, was reported to be required for transport of *Map2* mRNA into neuronal dendrites (Rehbein et al., 2002). Similarly, the chicken orthologue, nuclear zipcode binding protein 2 (ZBP2) is involved in targeting nuclear *Actb* mRNA to the cytoplasm (Gu et al., 2002). We previously showed that KHSRP can destabilize the mRNA encoding growth associated protein 43 (*Gap43*) and regulate neurite growth in cultured embryonic hippocampal neurons (Bird et al., 2013). Despite evidence for roles in neurons, KSHRP’s function in the brain has not been systematically defined.

Here, we have used a combination of molecular, cellular, electrophysiological and behavioral approaches to better understand the role of KHSRP in the brain. We find that KHSRP regulates multiple neuronal target mRNAs that are associated with nervous system development and function, including neuronal morphology, axonal growth, and synaptic functions. These gene expression findings are consistent with alterations in neurite growth upon loss of KHSRP, where both the brains and cultured CNS neurons from *Khsrp*^−/−^ mice show increased axon and dendrite growth compared to wild type (*Khsrp*^+/+^) mice. The result of these alterations is an increase in spontaneous neurotransmission and disrupted hippocampal-dependent learning and prefrontal cortex function in the *Khsrp*^−/−^ and *Khsrp*^+/−^ mice. Our findings emphasize the critical role that post-transcriptional modulation of mRNA stability by KHSRP plays in brain development and function.

## RESULTS

### Neuronal KHSRP target mRNAs are upregulated in Khsrp^−/−^ brain

We have previously shown that KHSRP is expressed in cultured embryonic hippocampal neurons where it destabilizes *Gap-43* mRNA and attenuates neurite growth, while loss of KHSRP results in the opposite phenotype to increase both *Gap43* mRNA levels and neurite growth (Bird et al., 2013). The affected neurites were inferred to be ‘axonal’ in those 5 day cultures of cortical neurons based on morphology. Here, we find that levels of KHSRP progressively increase as cultured cortical neurons extend TuJ1-positive neurites that go on to polarize into axons and dendrites (Suppl. Figure S1). Although KHSRP protein expression persists in the adult brain, the function of this protein in neurons remains to be established *in vivo*. We used microarray analyses to systematically test for changes in mRNA levels in the neocortex of *Khsrp*^−/−^ vs. *Khsrp*^+/+^ adult mice. This identified 1460 mRNAs with significantly elevated levels in neocortex upon loss of KHSRP expression (Figure 1A, Suppl. Table S1). We next used RNA co-immunoprecipitation with KHSRP antibodies from neocortex of wild type mice following by next-generation sequencing (RIP-seq) to identify mRNAs bound to KHSRP in brain (Suppl. Table S2). Integrating the RIP-Seq and microarray datasets enabled us to focus our subsequent analyses on a mRNA cohort that is potentially destabilized by direct interactions with KHSRP. From this, we identified 527 mRNAs that are both elevated in *Khsrp*^−/−^ vs. *Khsrp*^+/+^ neocortex and significantly enriched in KHSRP RIP-Seq from *Khsrp*^+/+^ vs. control RIP-Seq using *Khsrp*^−/−^ tissues; we refer to these as ‘KHSRP-target mRNAs’ (Figure 1B; Suppl. Table S4). 444 of these mRNAs have AU-rich elements (AREs) in their 3’ UTRs, suggesting that KHSRP can directly bind to and destabilize those transcripts (Suppl. Table S4). Analysis of the 527 KHSRP-target mRNAs using Ingenuity pathway analyses (IPA) revealed a significant enrichment of their encoded proteins in the control of neuronal morphology, axon development/growth, axonal guidance, long-term memory, neurotransmission, and other brain and neuron structure and function categories (Figure 1C).

**Figure 1:**
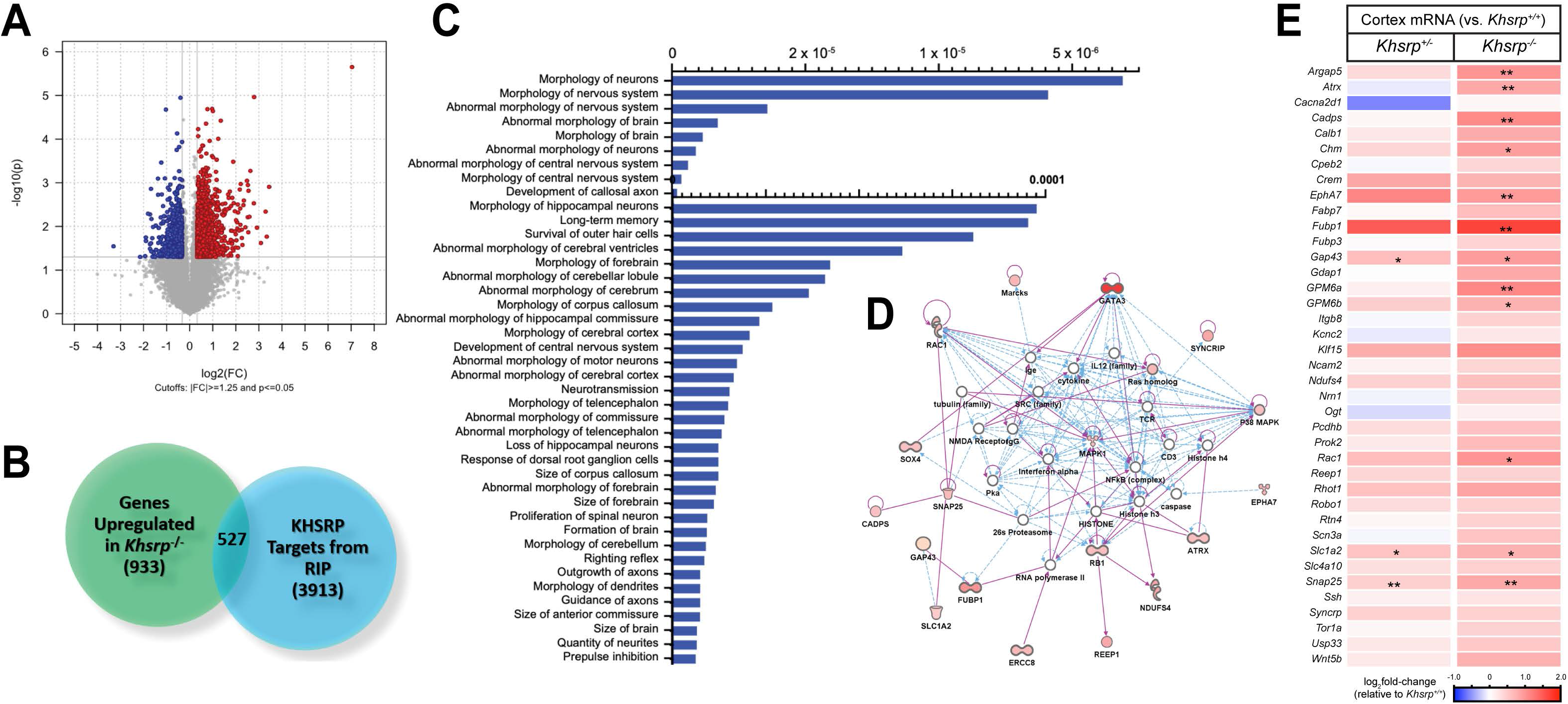
Increased levels of KHSRP-target mRNAs in the neocortex of KHSRP deficient mice. **A)** Volcano Plot for differential expression of mRNAs in neocortex of *Khsrp*^−/−^ vs. *Khsrp*^+/+^ mice from microarray analyses. The vertical grey line correspond to log_2_ fold-chang (1.25 up or down), and the horizontal line represents a p value of 0.05 that were used as as cut off for subsequent analyses. Also see Suppl. Table S1. **B)** Venn diagram for overlap between genes significantly upregulated in *Khsrp*^−/−^ vs. *Khsrp*^+/+^ and those identified as binding to KHSRP by RIP-seq (log_2_ fold-change > 1.4; p < 0.05). See Suppl. Table S2 for RNAs identified by KHSRP RIP-Seq and overlap with data from panel A and Suppl. Table S3 for ARE-containing KHSRP-target mRNAs). **C)** IPA analyses of functional pathways regulated by the 527 KHSRP-target mRNAs from B reveal top pathways related to neuronal morphology. **D)** Top nervous system development and function network identified by IPA. Up-regulated genes are highlighted in red, color intensity is proportional to fold change increase in *Khsrp*^−/−^ cortex. **E)** RTddPCR validation of alterations in levels of KHSRP-target mRNAs from D for *Khsrp^+/−^* and *Khsrp*^−/−^ vs. *Khsrp*^+/+^ neocortex as a heat map based on Log_2_ fold-changes for indicated mRNAs. See Suppl Figure S2 for validation of select KHSRP-target mRNAs in hippocampus and Suppl Table S5 and S6 for mRNA copy number from these analyses (N ≥ 3 per condition; * p ≤ 0.05, ** p ≤ 0.01, and *** p ≤ 0.005 by two-tailed Student’s *t-*test).

The top IPA nervous system development and function network derived from this list of KHSRP-target mRNAs included 58 proteins involved in neuronal morphology and axonal guidance showing different extents of upregulation, including SNAP25, SYNCRIP, MARCKS, RAC1, GAP-43 (Bannai et al., 2004; Hua et al., 2015; Osen-Sand et al., 1993; Xu et al., 2014) (Figure 1D, Suppl. Table S3). Reverse transcriptase coupled droplet digital PCR (RTddPCR) with transcript specific primers validated the unbiased screens used in Figure 1A-D, showing significantly increased levels for many KHSRP-target mRNAs in neocortex from adult *Khsrp*^−/−^ vs. *Khsrp*^+/+^ mice, with *Khsrp^+/−^* samples often showing intermediate levels (Figure 1E; Suppl Table S5). A subset of these mRNAs also showed significantly increased levels in hippocampus of the *Khsrp*^−/−^ and *Khsrp*^+/−^ vs. *Khsrp*^+/+^ mice (Suppl. Figure Figure S2, Suppl. Table S6). Several of the mRNAs validated for upregulation in the cortex and hippocampus have been linked to neuronal differentiation and synaptic function beyond *Gap43* mRNA. For example, *Fubp1* mRNA encodes is a member of the same FUSE protein binding family as KHSRP, and FUBP1 can promote terminal differentiation of neural progenitors (Hwang et al., 2018). *Snap25* encodes a member of the SNARE complex of the synaptic release machinery, but has also been linked to axon growth and synapse development (Bark and Wilson, 1994; Batista et al., 2017; Osen-Sand et al., 1993). The Ephrin receptor EphA7 has been linked to neuronal differentiation, dendritic morphology, and LTP (Beuter et al., 2016; Clifford et al., 2014). Finally, *Scl1a2* encodes the glial high affinity glutamate transporter EAAT2, but there is also evidence for *Slc1a2* expression by neurons (Zhou et al., 2019). Thus, *in vivo* loss of KHSRP protein expression alters levels of a number of different mRNAs whose protein products have clear potential to impact neuronal development and/or function.

### Loss of KHSRP increases axon and dendrite growth in vivo

Given the increase in mRNAs encoding proteins that can affect neuronal morphology and axon growth in the *Khsrp*^−/−^ mice observed above, we asked if loss of KHSRP expression changes neuronal morphology *in vivo*. For this, we crossed Khsrp knockout mice with Thy1-GFP mice that have GFP expression restricted to a subset of pyramidal neurons (Feng et al., 2000). As a measure of axonal growth, we focused on the length of the hippocampal infrapyramidal mossy fiber bundle (IPB), a tract that is pruned late in development normally after postnatal day 20 (Bagri et al., 2003). We had previously shown that adult HuD overexpressing mice have an increase in IPB length (Perrone-Bizzozero et al., 2011). As shown in the panels in Figures 2A and D, IPB length is significantly increased in both *Khsrp*^+/−^ and *Khsrp*^−/−^ compared to *Khsrp*^+/+^ mice. Thus, our findings that the loss of an mRNA destabilizer in this pathway has similar effects to the overexpression of an mRNA stabilizer suggest that the KHSRP to HuD ratios are critical to regulate axonal growth.

**Figure 2:**
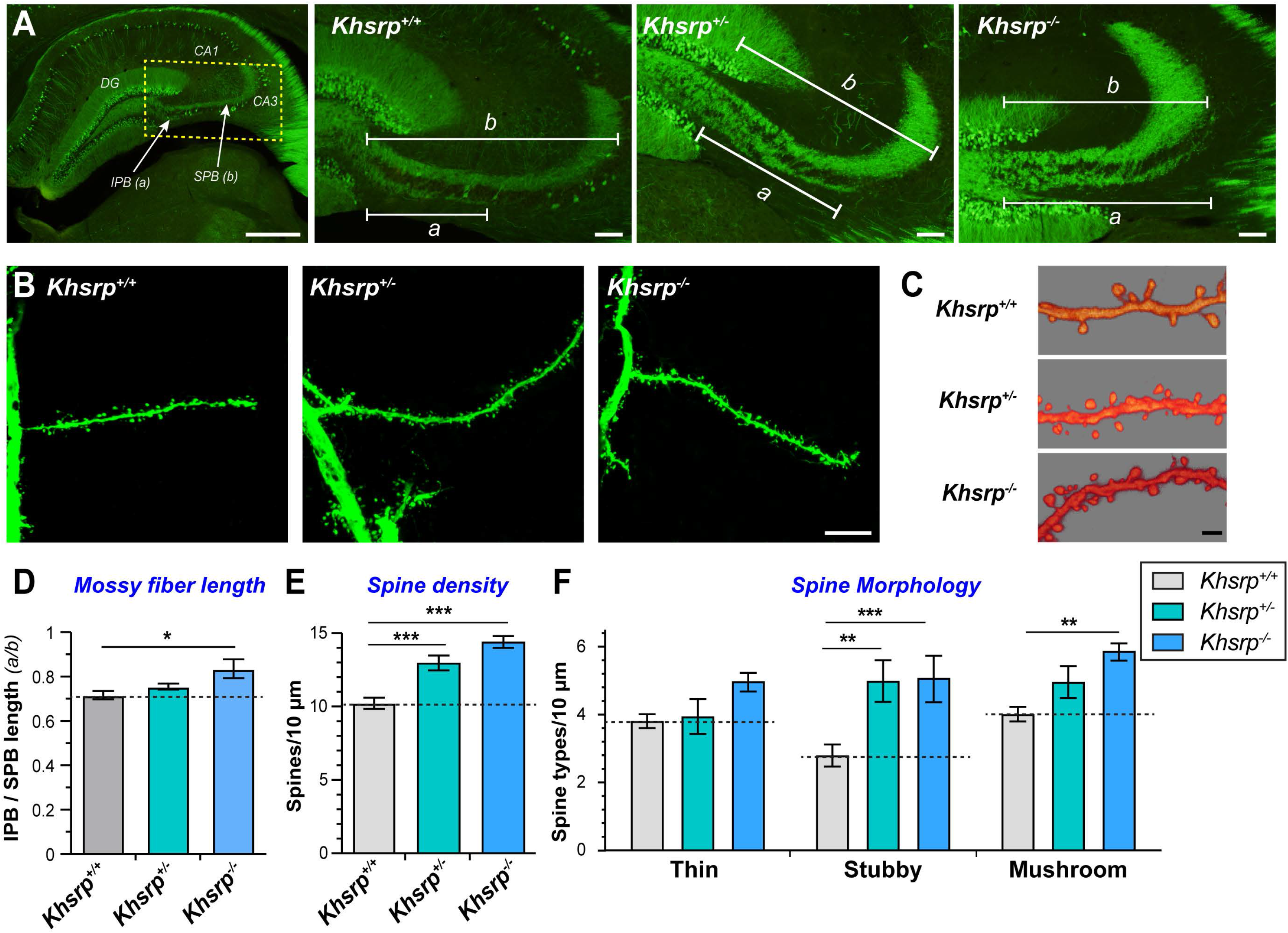
Increased axon and dendrite growth in the brains of KHSRP deficient mice. **A)** Representative images for hippocampi of adult Thy1-GFP mice crossed *with Khsrp^+/−^* and *Khsrp*^−/−^ show altered mossy fiber outgrowth compared to *Khsrp*^+/+^. The low power image at left outlines area for three higher power images to the right. IPB is marked as ‘a’ and SPB as ‘b’ in the three right panels (DG = dentate gyrus; CA1 and CA3 mark corresponding cornu Ammonis subfields). **B-C)** XYZ projections of apical dendrites of layer 5 neurons from Thy1-GFP mice crossed with *Khsrp^+/−^* and *Khsrp*^−/−^ show that loss of KHSRP increases dendritic spine numbers in panel **B**. High magnification three-dimensional rendered XYZ image of segment of second-order apical dendrites are shown in panel **C**. **D-F)** Mean IPB/SPB length (a/b) (**D**) is significantly increased in *Khsrp*^−/−^ compared to *Khsrp*^+/+^ mice and spine density (**E**) is significantly increased in both *Khsrp*^−/−^ and *Khsrp^+/−^* compared to *Khsrp*^+/+^ mice. Density of stubby spines is significantly increased in both *Khsrp*^−/−^ and *Khsrp^+/−^* while the density of mushroom spines is significantly increased in *Khsrp*^−/−^ compared to *Khsrp*^+/+^ mice (**F**). Error bars represent SEM (N = 5; * p < 0.05 by ANOVA) [Scale bars = A, left panel, 500 and right three panels 200 μm; B,10 μm; and C, 2 μm].

To determine if loss of KHSRP affects dendrite development, we examined the number and morphology of spines on apical dendrites of layer 5 pyramidal neurons in the somatosensory cortex from postnatal day 40 *Khsrp*^−/−^, *Khsrp*^+/−^ and *Khsrp*^+/+^ mice. The *Khsrp*^−/−^ and *Khsrp*^+/−^ mice showed significantly increased apical dendrite spine density compared to *Khsrp*^+/+^ mice (Figure 2B,C,E). With *Neurolucida* software, we defined the morphology of dendritic spines as ‘mushroom’, ‘thin’, or ‘stubby’ spines that roughly correspond to their maturity (Ethell and Pasquale, 2005). Significantly greater numbers of stubby and mushroom spines were observed in *Khsrp*^−/−^ mice compared to *Khsrp*^+/+^ (Figure 2F). In addition *Khsrp^+/−^* mice showed significantly more stubby spines compared to *Khsrp*^+/+^ mice, with density of mushroom spines in *Khsrp^+/−^* mice being intermediate between the *Khsrp*^+/+^ and *Khsrp*^−/−^ mice but not reaching statistical significance (Figure 2F). Taken together with the increased length of the IPB axons in *Khsrp*^−/−^ and *Khsrp*^+/−^ mice, these data indicate that KHSRP normally attenuates axonal growth and dendritic spine density as well as controls dendritic spine morphology.

### Elevated neurite growth and KHSRP-target mRNAs with KHSRP deficiency are neuron-intrinsic

Since KHSRP is also expressed in non-neuronal cells in the brain (Snee et al., 2002), it is possible that the *in vivo* neuron morphology and mRNA level changes in the KHSRP knockout mice could result from extrinsic effects on neurons. To explore this possibility, we used primary neuron cultures from single embryonic day 18 (E18) mouse embryos of *Khsrp^+/−^* crosses that included *Khsrp*^−/−^, *Khsrp*^+/−^, and *Khsrp*^+/+^ genotypes in each litter. These cultures contain ≥ 95 % neurons, allowing us to assess neuron-intrinsic growth mechanisms. Low density cultures analyzed at 7 days *in vitro* (DIV) were used to quantify axon and dendrite branching in the neurons by immunostaining using definitive markers for axons and dendrites. Axons were significantly longer in both cortical and hippocampal neurons from *Khsrp*^−/−^ compared to *Khsrp*^+/+^ embryos (Figure 3A,B; Suppl. Figure S2A,B). Cortical and hippocampal neurons from *Khsrp^+/−^* mice showed axon lengths intermediate between *Khsrp*^−/−^ and *Khsrp*^+/+^, but this was only significant in the hippocampal cultures (Figure 3B; Suppl. Figure S3B). *Khsrp*^−/−^ hippocampal neurons showed significantly more branching of their axons than those from *Khsrp*^+/+^ embryos (Suppl. Figure S2B), but this was curiously not the case for the cortical neurons (Figure 3B). In contrast, the cortical neurons showed a greater increase in dendrite length than *Khsrp*^−/−^ hippocampal neurons compared to *Khsrp*^+/+^, with dendrites of *Khsrp^+/−^* cortical neurons showing lengths intermediate between those of *Khsrp*^−/−^ and *Khsrp*^+/+^ neurons (Figure 3B; Suppl. Figure S3B). Both cortical and hippocampal neurons showed significantly increased branching of dendrites for *Khsrp*^−/−^ compared to *Khsrp*^+/+^ neurons (Figure 3C; Suppl. Figure S3C).

**Figure 3:**
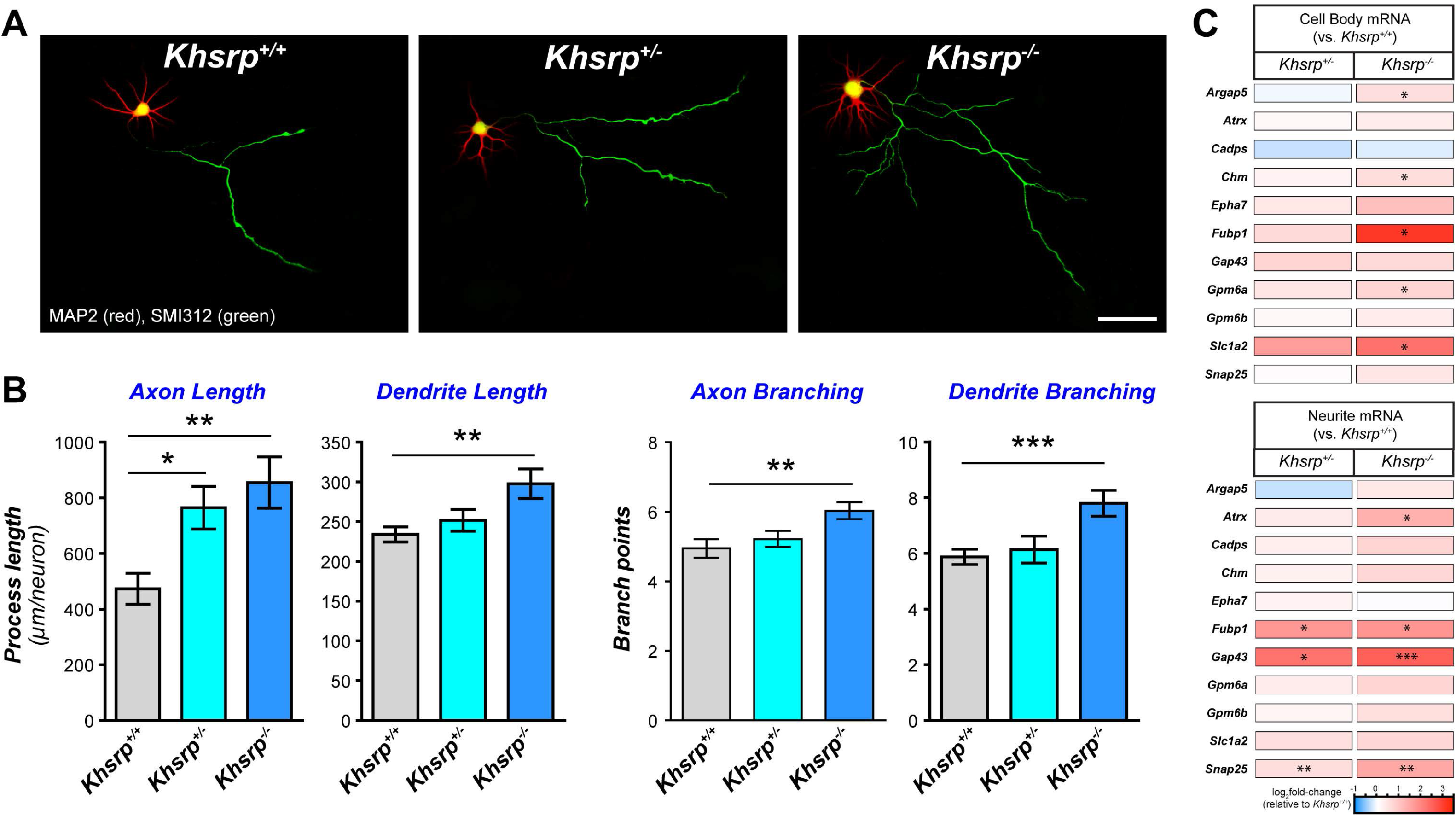
Neurons cultured from KHSRP deficient mice show increased axonal and dendritic growth. **A)** Representative micrographs for cortical neurons from *Khsrp*^+/+^, *Khsrp^+/−^*, and *Khsrp*^−/−^ littermates shown for 7 DIV cultures as indicated. **B)** Axon and dendrite lengths and branching are displayed means ± SEM (N ≥ 75 neurons analyzed over at least three separate cultures preparations; * p ≤ 0.05, ** p ≤ 0.001, and *** p ≤ 0.005 by one-way ANOVA with Tukey post-hoc). **C)** Heat map for levels of KHSRP target mRNAs linked to axon growth and neuronal morphology for cell body and neurite RNA analyses. Data are displayed as Log_2_ fold-change relative to *Khsrp*^+/+^ cortical cultures (N ≥ 4 culture preparations; * p ≤ 0.05, ** p ≤ 0.001, and *** p ≤ 0.005 vs. *Khsrp^+/^* by two-tailed Student’s *t*-test). See Suppl. Figure S3 for parallel analyses in hippocampal neuron cultures and Suppl. Table S5 for mRNA copy number from these analyses [Scale bar = 100 μm].

We used DIV 23 cultures of E18 cortical and hippocampal neurons to measure neuron intrinsic effects of KHSRP on dendritic spine formation. Dendrites and spines were visualized by expression of GFP to fill the neuronal cytoplasm for imaging. Both cortical and hippocampal neurons from *Khsrp*^−/−^ mice showed a modest, but significant increase in dendritic spine density compared to *Khsrp*^+/+^ neurons (Suppl. Figure S4). An increase in mushroom shaped spines in the *Khsrp*^−/−^ cortical neurons accounted for this difference in cortical neurons, while the *Khsrp*^−/−^ hippocampal neurons showed a significant increase in thin spines (Suppl Figure S4B,D). Taken together with the axon and dendritic length and branching with loss of KHSRP shown above, these data indicate that KHSRP likely modulates neuronal growth through neuron-intrinsic mechanisms.

With these changes in axonal and dendritic growth in cultures from mice with partial or complete loss of KHSRP, we asked if the mRNAs showing significant elevations in brain tissues of *Khsrp*^−/−^ mice from Figure 1E might also be altered in cortical neurons cultured from those mice. Further, since several of those KHSRP-target mRNAs are known to be transported into dendrites and/or axons, we separated cell bodies from neurites in these cultures to gain an assessment of overall and neurite-localized mRNA levels. *Argap5, Chm, Gpm6a Fubp1, Gap43,* and *Slc1a2* mRNAs were significantly increased in cell body RNA preparations from *Khsrp*^−/−^ neuron cultures; *Atrx, Fubp1*, *Gap43* and *Snap25* mRNAs were also elevated in *Khsrp*^−/−^ neurites (Figure 3C; Suppl. Table S7). Interestingly, *Atrx* and *Snap25* mRNA showed no significant changes in the cell body RNA preparations, but the mRNAs were significantly elevated in the neurites of the *Khsrp*^−/−^ compared to *Khsrp*^+/+^ neuron cultures (Figure 3C). *Gap43* and *Snap25* mRNAs have previously been reported to localize into axons of cultured neurons (Batista et al., 2017; Hafner et al., 2019; Smith et al., 2004). *Fubp1, Gap43*, and *Snap25* mRNAs were also increased in the *Khsrp^+/−^* vs. *Khsrp*^+/+^ neurites, but none of the mRNAs tested show significant differences between *Khsrp^+/−^* vs. *Khsrp*^+/+^ in the cell body RNA preparations (Figure 3C; Suppl. Table S7). Overall, these data support that the increases in KHSRP-target mRNAs, axon lengths, and dendritic spine growth seen in *Khsrp*^−/−^ mice arise, at least in part, from changes in neuronal gene expression via loss of the neuronal functions of KHSRP.

### Loss of KHSRP elevates excitatory neurotransmission

The changes in neuronal morphology and increased levels of KHSRP target mRNAs encoding synaptic proteins in the *Khsrp*^−/−^ neurons seen above raise the possibility that loss of KHSRP could alter synaptic function. We used slice electrophysiology to compare synaptic function between *Khsrp* knockout and wild type mice. For this, we initially measured mEPSCs in CA3 pyramidal neurons of the dorsal hippocampus of adult *Khsrp*^−/−^, *Khsrp^+/−^* and *Khsrp*^+/+^ mice. mEPSC frequency was significantly increased in CA3 hippocampal neurons from *Khsrp*^−/−^ compared to *Khsrp*^+/+^ and Khsrp^+/−^ mice (Figure 4A,B). Since the analyses of neurite growth and KHSRP-target mRNAs showed differences between hippocampal and cortical neurons, we also assessed synaptic function in infralimbic cortex layer V neurons. *Khsrp*^−/−^ cortical neurons again showed a significant elevation of mEPSC frequency compared to those of *Khsrp*^+/+^ brains (Figures 4C,D). The *Khsrp^+/−^* cortical neurons showed average frequency intermediate between *Khsrp*^−/−^ and *Khsrp*^+/+^ cortices, but this did not reach statistical significance. There was no difference in mEPSC amplitude and mEPSC duration between the genotypes in either hippocampal CA3 or infralimbic cortical neurons (Figures 4B,D). Changes in mEPSC frequency traditionally indicate involvement of pre-synaptic, rather than post-synaptic, mechanisms (Redman, 1990). Thus, the increased mEPSC frequency in the *Khsrp*^−/−^ mice likely reflects increased numbers of functional synapses, consistent with observed increases of axonal and dendritic growth in the mice. Interestingly, post-synaptic sensitivity to glutamate release (*e.g*., number of synaptic AMPA receptors, AMPA receptor subunit composition), as reported by mEPSC amplitude and duration, is not altered by the loss of KHSRP.

**Figure 4:**
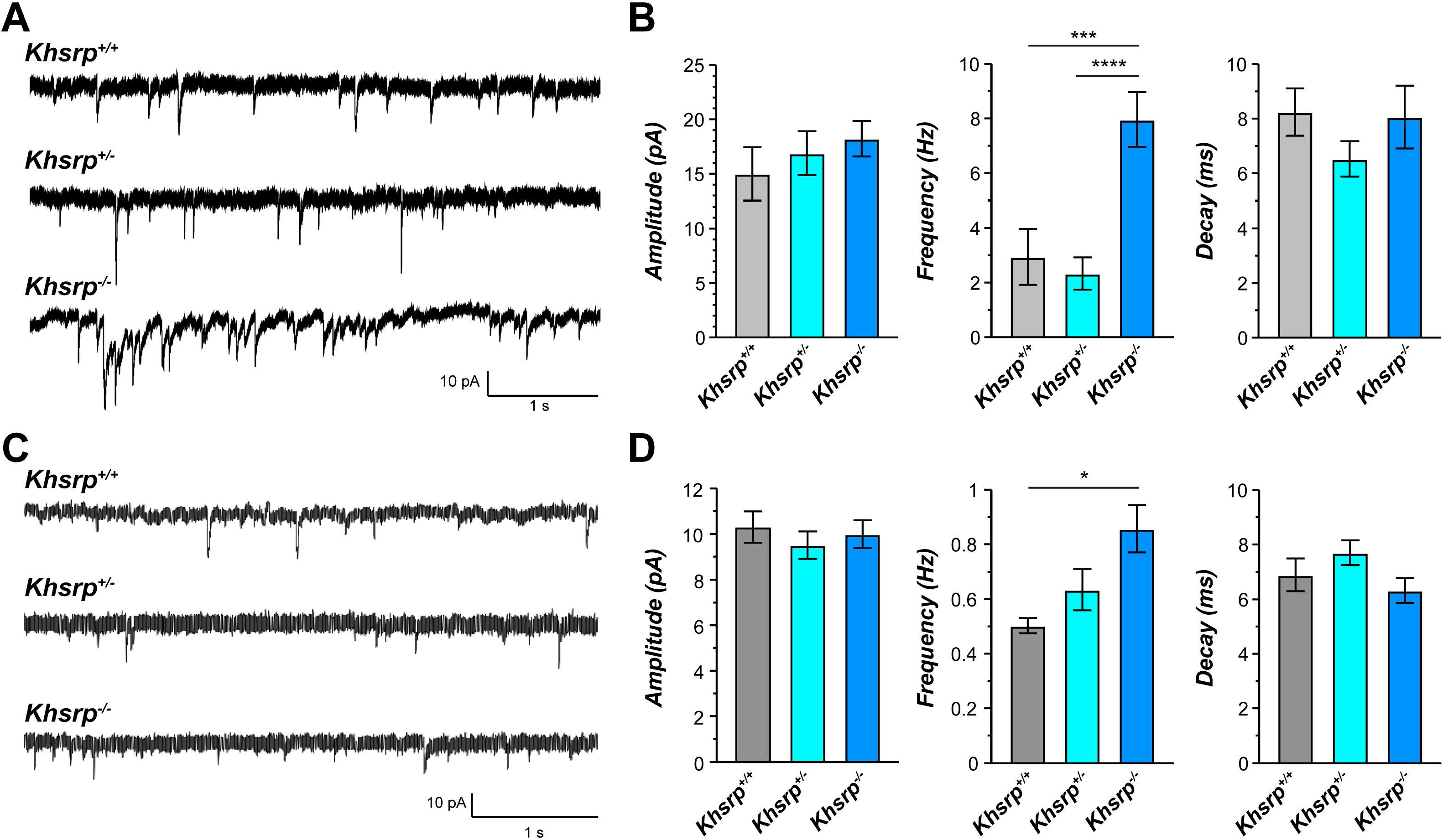
Increased presynaptic activity in KHSRP deficient mice. **A)** Representative traces of AMPA receptor-mediated miniature excitatory post-synaptic currents (mEPSCs) in CA3 pyramidal neurons of the dorsal hippocampus. **B)** mEPSC frequency is significantly increased in *Khsrp*^−/−^ relative to *Khsrp*^+/+^ and *Khsrp*^+/−^ mice. There were no significant differences in mEPSC amplitude or duration (decay time) between genotypes (frequency data analyzed by one-way ANOVA, F(2,28)=11.97, p=0.0002; ***p<0.01 and ****p<0.001 by Tukey’s post-hoc comparisons). **C)** Representative traces of AMPA receptor-mediated mEPSCs in layer 5 pyramidal neurons of the infralimbic cortex. **D)** Infralimbic cortical neurons show increased mEPSC frequency in the *Khsrp*^−/−^ relative to *Khsrp*^+/+^ mice. As in the CA3, no differences in mEPSC amplitude or duration were detected (frequency data analyzed by one-way ANOVA, F(2,31)=5.03, p=0.013; * p<0.05 by Tukey’s post-hoc comparisons),.

### Loss of KHSRP increases locomotor activity and impairs hippocampal- and prefrontal cortex-dependent memory consolidation

Given the alterations in neuronal morphology and function, we asked whether loss of KHSRP expression would affect mouse behavior. Initial behavioral screening was adapted from a subset of tests derived from the Irwin screen for physical health, appearance, sensory utility, motor coordination, and neurological function (Crawley, 1999; Irwin, 1968; Zhao et al., 2006). As shown in Suppl. Table S8, *Khsrp*^+/−^ and *Khsrp*^−/−^ mice exhibit normal physical features including weight, whiskers, eyes, eyelids, teeth, tail, and fur. They also show no differences in normal mouse behaviors including gait, grooming, and rearing. However, *Khsrp*^−/−^ mice display significantly increased circling and spontaneous running (Suppl. Table S8), which may indicate a level of hyperactivity with deletion of the *Khsrp* gene.

We next assessed anxiety-like behavior, locomotion, and exploratory tendencies in these mice using the novel open field and elevated zero maze tests. *Khsrp*^−/−^ and *Khsrp*^+/−^ mice showed no differences in percent of time in the center of the open field (Figure 5A) or in the open arm duration on the zero maze compared to *Khsrp*^+/+^ mice (Suppl. Figure S5), which together indicate no alteration in anxiety-like behaviors with KHSRP loss. However, we found a significant effect of time in both distance and velocity in the open field test (Figure 5B,C), which were due to a significant increase in velocity and distance for the *Khsrp*^−/−^ and *Khsrp*^+/−^ vs. *Khsrp*^+/+^ mice on day 1 compared to all other days. Further examination of day 1 revealed that the first time bin (0-5 min) was significantly different from all other time points for both distance traveled and velocity, which is indicative of novelty-induced locomotion. This difference is driven by the *Khsrp*^+/−^ mice for distance traveled, while the velocity of *Khsrp*^−/−^ and *Khsrp*^+/−^ mice was significantly elevated compared to *Khsrp*^+/+^ controls (Figure 5C,D).

**Figure 5:**
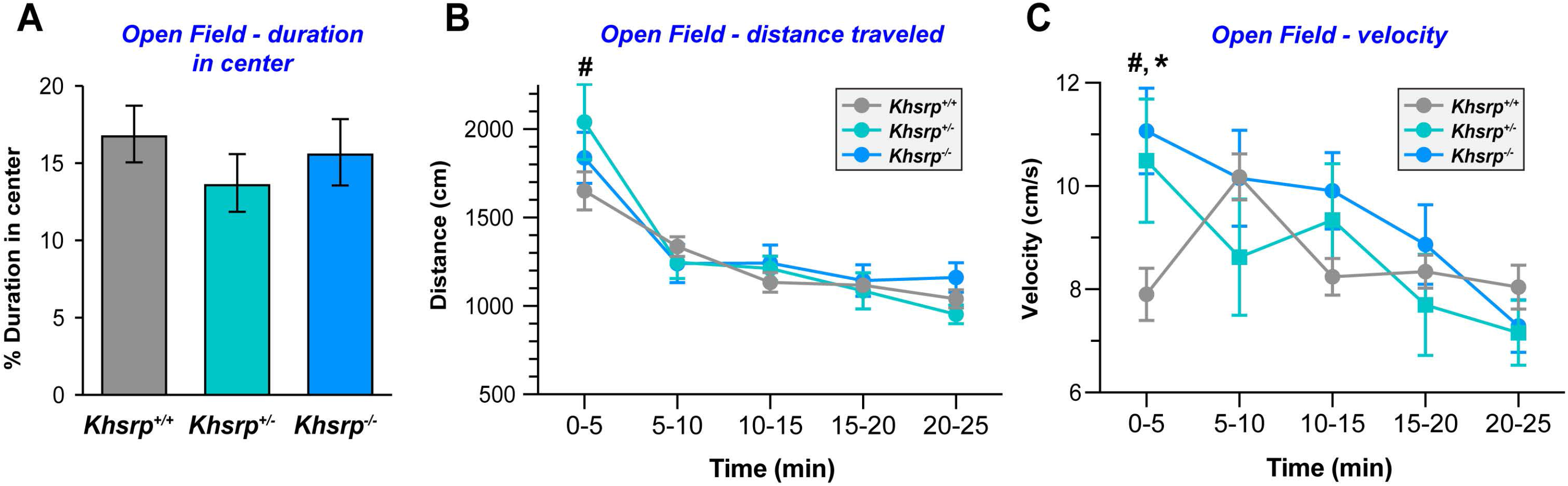
KHSRP deficient mice show increased locomotor activity. **A)** Mice with decreased KHSRP levels display no difference in percentage of time in the center in the Open field test. **B-C)** *Khsrp*^−/−^ and *Khsrp^+/−^* mice have increased locomotor activity in the first 5 min for day 1 as measured by distance traveled (**B**) or velocity (**C**) but decreased locomotor activity over time All values are mean ± SEM (N=28 for *Khsrp*^+/+^ and *Khsrp*^−/−^ and n=10 for *Khsrp^+/−^;* Repeated measures ANOVA show a significant time × genotype interaction p=0.0026. Tukey post hoc tests revealed that the first bin (0-5 min) is significantly different from all other time points for both distance traveled and velocity. This difference is driven by the *Khsrp^+/−^* mice for distance traveled (ANOVA F(2,60)=0.2306, p=0.0449, ^#^p<0.05 *Khsrp^+/−^* vs. *Khsrp*^+/+^), while for velocity *Khsrp^+/−^* and *Khsrp*^−/−^ animals are both significantly elevated compared to *Khsrp*^+/+^ (ANOVA F(2, 60)=1.895, p=0.0064, ^##^p<0.01 *Khsrp^+/−^* vs *Khsrp*^+/+^ and p=0.0442, *p<0.05 *Khsrp*^−/−^ vs. *Khsrp*^+/+^.

Considering the elevations in hippocampal axon/dendrite growth, KHSRP-target mRNAs, and synaptic activity with loss of KHSRP, we next used trace conditioning to test hippocampal function in the *Khsrp*^−/−^ and *Khsrp*^+/−^ mice (McEchron et al., 1998). Trace conditioning tests ability to associate a conditioned stimulus (CS) to an unconditioned stimulus (US) separated by a 30 second trace interval. Training consisted of seven tone (CS) and shock (US) presentations, with initial learning of the task assessed by percentage of freezing during the CS and trace interval. All mice progressively increased freezing to the CS and during the trace interval following each CS presentation during training (Figure 6A-B). There were no main effects of genotype, sex, nor a sex × genotype interaction during the CS and trace portions of training (Figure S6A), so the male and female data for individual genotypes were combined for subsequent analyses. There was no significant time × genotype interaction during the CS (Figure 6A) except for the seventh presentation of the tone, when *Khsrp*^+/−^ mice showed increased freezing compared to the *Khsrp*^+/+^ mice. However, there was a significant effect of time × genotype during the trace responses (Figure 6B) that was observed in both male and female mice (Figure S6A). The *Khsrp*^−/−^ mice showed significantly higher freezing than the *Khsrp*^+/+^ mice at the second, third, and seventh trace periods and than the *Khsrp*^+/−^ at the second and third periods. *Khsrp*^+/−^ mice also showed significantly higher freezing than the *Khsrp*^+/+^ mice at the seventh trace period (Figure 6B). To assess retention of trace fear conditioned responses, freezing behavior of *Khsrp*^+/−^ and *Khsrp*^−/−^ mice was evaluated 24 hours post training (Figure 6B). The CS was delivered in a novel context to examine the response independent of the original fear context. We found no significant effects of genotype on percent freezing to the CS; however, *Khsrp*^−/−^ mice displayed decreased freezing during the trace interval compared to *Khsrp*^+/+^ mice, which was observed in both male and female mice (Figure S6B). Taken together, these changes in trace conditioning suggest that deletion of KHSRP impairs memory consolidation.

**Figure 6:**
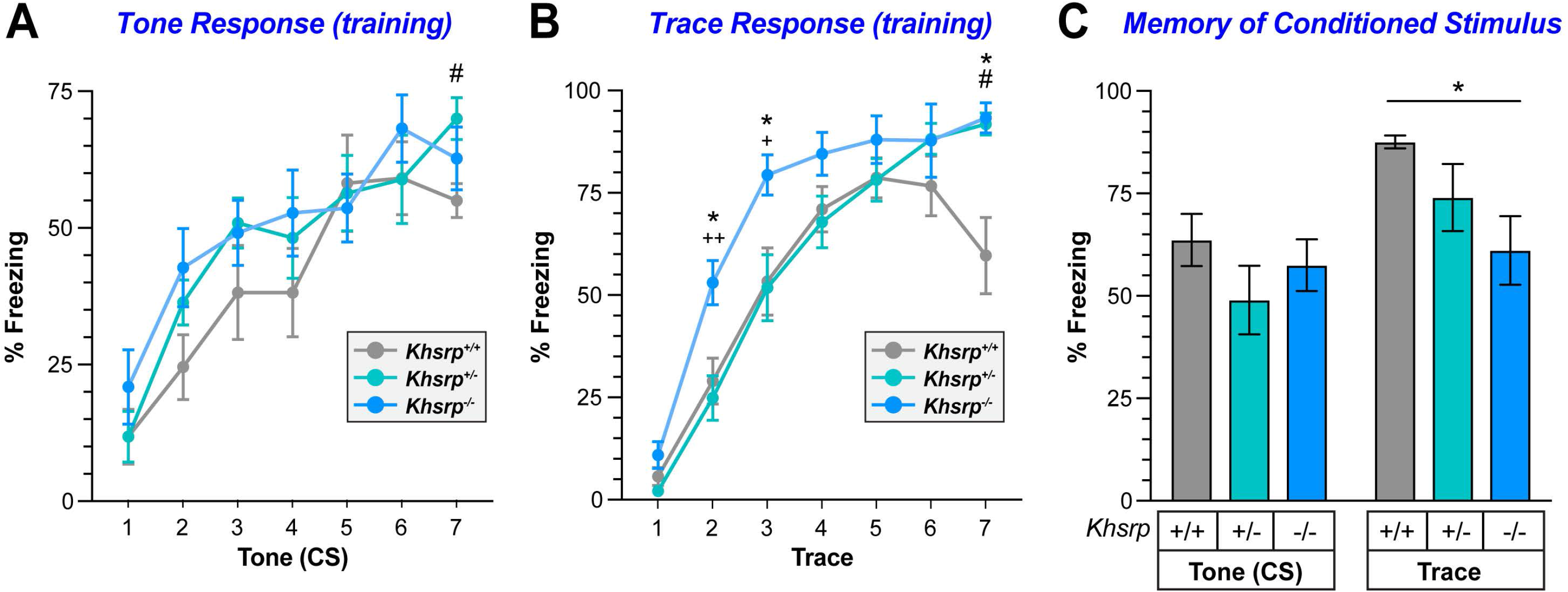
Impaired hippocampal-dependent memory consolidation in KHSRP deficient mice. **A-B**) All mice demonstrate increased freezing during training with increasing presentations of the conditioned stimulus (CS; tone, **A**) and the trace interval following CS (**B**). Notably, *Khsrp^+/−^* mice show increased freezing vs. *Khsrp*^+/+^ mice at the last tone and trace interval presentation. *Khsrp*^−/−^ mice also show increased freezing during the trace period during training vs. *Khsrp*^+/+^ and *Khsrp^+/−^*. Data are shown as mean ± SEM, n=11 (6 male and 5 females/genotype). Repeated measures ANOVA followed by one way ANOVA for each time point. +p<0.05 and ++p<0.01 *Khsrp^+/−^* vs. *Khsrp*^−/−^mice, *p<0.05 *Khsrp*^−/−^ vs. *Khsrp*^+/+^ and (# p<0.05 *Khsrp^+/−^*vs. *Khsrp*^+/+^ **B)** All mice freeze equally during CS presentation approximately 24 hours after training, but *Khsrp*^−/−^ mice display decreased freezing during the trace interval compared to *Khsrp*^+/+^. Data are shown as mean ± SEM, n=11 (6 male and 5 females/genotype; one way-ANOVA (F(2,27)=3.948, p=0.0149, *p<0.05). See Suppl. Figure S6 for sex-specific responses of each genotype.

Since the *Khsrp*^−/−^ mice also showed increased mEPSCs in infralimbic cortex, we used the attentional set shifting task (ASST) to assess the effect of loss of KHSRP on frontal cortex function. The ASST tests executive function by initially exposing animals to a series of problem stages that are used to predict food location (Birrell and Brown, 2000). This includes discrimination-reversal learning within one dimension (odor or tactile discrimination within a platform), an intra-dimensional shift (IDS) with novel exemplars within the previously learned dimension (i.e., odor 1 to odor 2 or platform 1 to platform 2), as well as an extra-dimensional shift (EDS) to the previously unrewarded dimension (i.e., odor to platform). It is well established that reversal learning is mediated by the orbitofrontal cortex (OFC) (Chudasama and Robbins, 2003; Rudebeck et al., 2013; Rudebeck and Murray, 2011) and attentional set shifting is mediated by regions of the ventromedial prefrontal cortex (vmPFC), which includes the infrapyramidal cortex (Birrell and Brown, 2000; Floresco et al., 2008; Floresco and Jentsch, 2011; Hamilton and Brigman, 2015). As shown in Figure 7A, there was a significant main effect of starting dimension during the Simple Discrimination (SD) and Compound Discrimination (CD) stages that was eliminated by the Compound Discrimination Reversal (CDR) stage. It is commonly observed that tactile learning is more difficult to initially acquire than olfactory learning (Thompson et al., 2015), but animals learn to efficiently discriminate tactile differences after repeated exposures (Figure 7A). Reversal learning normally requires more trials to reach criterion than the preceding stage as seen in the *Khsrp*^+/+^ mice, however, this was not true for the *Khsrp*^−/−^ mice (Figure 7B). Interestingly, *Khsrp*^+/−^ mice required increased number of trials to criterion for the first reversal, but this was not sustained on the second reversal (Figure 7B). The *Khsrp*^+/+^ mice formed an attentional set as measured by EDS-IDS, but neither *Khsrp*^+/−^ nor *Khsrp*^−/−^ mice established an attentional set (Figure 7B). Taken together these observations indicate that *Khsrp*^+/−^ and *Khsrp*^−/−^ are able to perform each discrimination, but they do not form an attention shift set so that they approach any changes in the food prediction cues as completely new cues.

**Figure 7:**
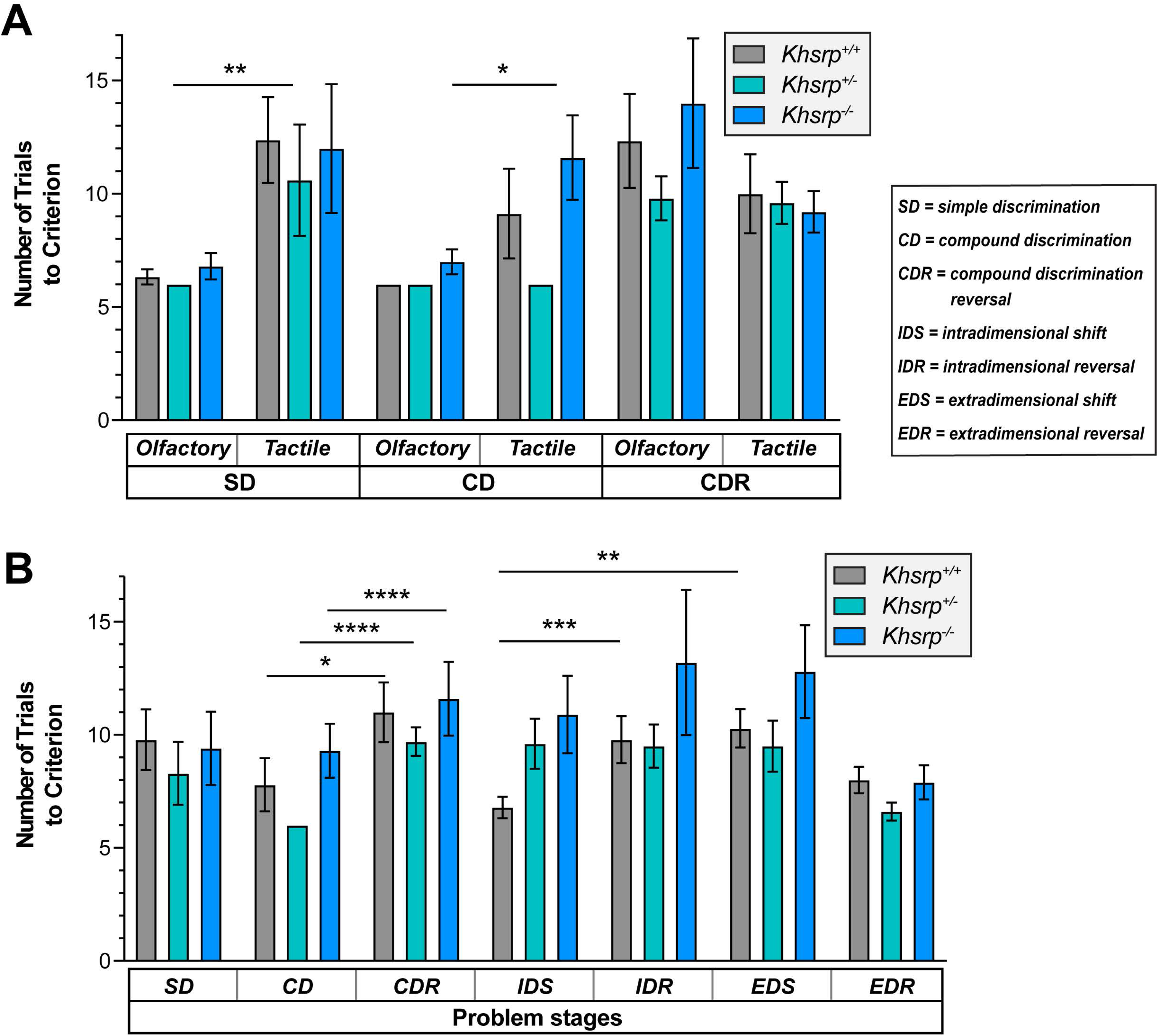
KHSRP deficient mice show decreased prefrontal cortical function. **A)** All mice show increased number of trials to criterion to learn differences in tactile (platform) vs. olfactory (odor) stimuli during CD and CR. See key for specific stages of the ASST task. **B)** All mice were able to perform discrimination (SD, CD, and IDS) and reversals (CDR, IDR, EDR), and showed increased number of trial for CDR vs. CD, although the reversals were no more difficult for *Khsrp+/−* and *Khsrp+/−* mice than the preceding stage for IDR vs IDS. Only *Khsrp+/+* mice formed an attentional set (EDS vs. IDS) while *Khsrp+/−* and *Khsrp+/−* did not (N ≥ 10/genotype; *p<0.05, **p<0.01, ***p<0.001 and ****p<0.0001 by Student’s *t*-tests).

## DISCUSSION

It is increasingly clear that post-transcriptional regulation of mRNAs plays a dynamic role in the regulation of neuronal genes and subsequent changes in behavior (Bolognani et al., 2007b; Bolognani and Perrone-Bizzozero, 2008). Critically, RBPs can regulate the stability of bound mRNAs (Bolognani and Perrone-Bizzozero, 2008; García-Mauriño et al., 2017). Since one mRNA can be translated into protein many times over, regulating the stability of mRNAs can dynamically modify cellular protein content. Here, we show that loss of the RBP KHSRP leads to a unique set of electrophysiological and behavioral changes, resulting in decreased hippocampal- and orbital frontal-dependent learning and memory. Based on increased levels of KHSRP-target mRNAs in brain tissues and parallel changes in KHSRP-target mRNAs plus alterations in axon and dendrite growth *in vivo* and in cultured neurons, the changes in neural activity and behavior seen in the *Khsrp*^−/−^ mice are undoubtedly driven by neuron-intrinsic elevations in KHSRP-target mRNAs when KHSRP expression is decreased.

We previously reported that loss of KHSRP leads to aberrant axonal outgrowth in DIV5 cultured embryonic cortical neurons due to increases in the levels of *Gap43* mRNA (Bird et al., 2013), a mRNA that is post-transcriptionally regulated by changes in its stability (Bolognani et al., 2006; Perrone-Bizzozero et al., 1993). Correct axonal growth regulation is critical for proper development and maintenance of neuronal networks, and the work shows that the axon growth abnormalities are maintained into more mature neurons, with clearly defined axonal and dendritic polarity, as well as impact dendrite growth and spine formation in vitro and in vivo.

Since overexpression of GAP-43 also leads to aberrant axonal sprouting (Fallini et al., 2016; Strittmatter et al., 1995), it is intriguing to speculate that mRNA destabilizing RBPs like KHSRP are needed to control levels of neuronal growth-associated mRNAs as neurons need to slow their growth when they start to make synaptic connections. Interestingly, we found that 444 of the neuronal KHSRP targets identified by RIP-seq were also upregulated in the cortex of *Khsrp*^−/−^ mice and contain 3’ UTR AREs for KHSRP binding (Suppl. Table S3). Thus, binding by KHSRP is predicted to shorten half-lives of those target mRNAs. Many of the KHSRP-target mRNAs identified herein encode proteins involved in neuronal development, axon growth, and synaptic plasticity. Our analyses confirm that absence of KHSRP, with stabilization of these target mRNAs, specifically increases axonal and dendritic growth as well as increases density of dendritic spines. The subtypes of spines that are increased in KHSRP deficient brains include both mature (mushroom) and more immature or plastic (stubby), with density of stubby spines significantly increased in both *Khsrp^+/−^* and *Khsrp*^−/−^ mice. Changes in axonal and dendrite length and branch points were also present in primary neuronal cultures from the *Khsrp^−/−^ mice*, indicating that KHSRP’s regulation of neuronal mRNAs is responsible for these changes.

The increased mEPSC frequency without changes in amplitude or decay seen in the hippocampus and prefrontal cortex of the *Khsrp*^−/−^ mice is consistent with increased numbers of synaptic terminals, increased presynaptic neurotransmitter release, or both (Redman, 1990). The KHSRP-target mRNA *Gap43*, which encodes a well-known growth-promoting protein that localizes to axonal growth cones (Aigner and Caroni, 1993; Van Der Zee et al., 1989), has previously been implicated in hippocampal dependent learning (Rekart et al., 2005). In contrast to our findings, *in vivo* overexpression of GAP-43 enhanced learning (Routtenberg et al., 2000). This suggests that overall increased GAP-43 expression seen with loss of KHSRP cannot alone explain the changes in electrophysiological or behavioral properties of *Khsrp*^−/−^ mice. This is not unexpected, as the phenotype of these mice is undoubtedly driven by the sum of proteome changes resulting from elevations in KHSRP-target mRNAs in these mice. Previous studies suggested that axonal *Gap43* mRNA translation contributes to elongating axonal growth (Donnelly et al., 2013), and loss of KHSRP increases both axon length and branching in the present study. Notably, the transgenic mice used by Routtenberg et al. (2000) for GAP43 overexpression only included a few nucleotides of the 3’UTR (Aigner et al., 1995), and it did not include the ARE that we have previously shown is needed for *Gap43* mRNA’s axonal localization (Yoo et al., 2013). Consequently, increase in locally synthesized GAP-43 could bring a different behavioral phenotype than seen by Routtenberg et al. (2000). Consistent with this, we previously only observed increased axonal growth when the overexpressed *Gap43* mRNA was targeted into axons through its 3’UTR (Donnelly et al., 2013).

Other KHSRP-target mRNAs encode proteins linked to axon growth and synaptogenesis, including ARGAP5, ATRX, CADPS, CHM, EPHA7, FUBP1, FUBP3, GPM6A, GPM6B, RAC1, SLC1A2, and SNAP25 (see Suppl. Table S4), that could affect numbers of synaptic terminals as evidenced by increased dendritic spine density in the brains and cultured neurons of the *Khsrp*^−/−^ mice. Increased neurotransmitter release could also be driven by KHSRP-target mRNA encoded proteins, as mRNA elevations for the SNARE protein SNAP25 seen with loss of KHSRP could elevate the probability of synaptic vesicle release (Hu and Davletov, 2003). Interestingly work from the Hengst lab has linked axonally localizing *Snap25* mRNA to formation of presynaptic terminals (Batista et al., 2017), and we see increased levels of *Snap25* mRNA in neurites of the *Khsrp*^−/−^ and *Khsrp^+/−^* mice. These observations raise the possibility that localized modulation of mRNA survival by KHSRP in axons helps to sculpt synaptic connectivity and contribute to synaptic plasticity in the brain. Consistent with this notion, recent work from the Schuman lab has shown an unexpectedly large population of proteins translated in presynaptic terminals of the adult brain including both *Gap43* and *Snap25* mRNAs (Hafner et al., 2019).

Similar to findings here in KHSRP knockout mice, we previously showed that mice overexpressing the RNA-binding protein HuD have elevated levels of ARE-containing mRNAs as well as altered neuronal morphology and associative learning (Bolognani et al., 2007a; Perrone-Bizzozero et al., 2011). The trace fear conditioning used here requires an intact hippocampus to form an association between the CS and US (McEchron et al., 1998). The *Khsrp*^−/−^ mice are able to initially learn the association between the CS and US during training. However, they display decreased freezing to the CS approximately 24 hours later, indicating that *Khsrp*^−/−^ mice have deficits in temporal processing of information due to the time separation between the training and testing (Meck et al., 2013; Ranganath and Hsieh, 2016). This may indicate a deficit in hippocampal dependent memory consolidation. In addition, mice with a deletion of KHSRP also show deficits in the ASST task, which requires functions of the ventromedial PFC. While *Khsrp*^−/−^ mice can perform reversals, these problems are no more difficult than the preceding discrimination indicating that these mice approach each problem as if it were novel. Thus, global loss of KHSRP impairs attentional set formation of species-appropriate stimuli.

Overall, our results indicate that KHSRP modulates levels of its target mRNAs required for the development of neural connectivity and potentially synaptic plasticity. Loss of KHSRP leads to significant changes in neuronal morphology through neuron-intrinsic mechanisms that persist into adulthood, resulting in impaired glutamatergic transmission and behaviors linked to functions of the hippocampus and prefrontal cortex. Our study emphasizes the importance of post-transcriptional regulation by KHSRP as a driver for brain development and function. The KHSRP-target mRNAs identified here show upregulation upon loss of KHSRP indicating they are targets for destabilization by KHSRP. These observations point to KHSRP is a post-transcriptional master regulator of a mRNA regulon linked to brain development and function.

## MATERIALS AND METHODS

### Animals

All animal studies were conducted in accordance with guidelines for animal use and care established by the University of New Mexico Health Science Center and University of South Carolina Institutional Animal Care and Use Committees (IACUCs). The *Khsrp*^−/−^ animals have deletion of exons 1-13 as described in Lin et al. (2011) and were cross-bred with C57Bl/6 for at least 10 generations. For morphological studies, *Khsrp*^−/−^ mice and were cross-bred with *B6.Cg-Tg(Thy1-EGFP)OJrs/GfngJ* mice (termed Thy1-GFP herein; obtained from Jackson Labs) to eventually generate *Khsrp^−/−^, Khsrp^+/−^* and *Khsrp*^+/+^ with GFP expression in select neurons.

Mouse genotyping was performed using PCR with primers spanning the exon 1 to exon 13 deletion of the mouse Khsrp gene (Lin et al., 2011) or wild type sequence (Transnetyx, TN).

### Primary Neuron Cultures

Primary cortical neuron cultures were prepared from embryonic day 18 (E18) mice. Cortices and hippocampi were dissected in Hibernate E (BrainBits, IL) and dissociated using the *Neural Tissue Dissociation kit* according to manufacturer’s protocol (Miltenyi Biotec, Bergisch Gladbach, Germany). For this, minced cortices or hippocampi were incubated in a pre-warmed Enzyme Mix 1 at 37°C for 15 min; tissues were then triturated with blunted glass pipette and again incubated with Enzyme Mix 2 for 10 min. Triturated tissue was applied to a 40 μm cell strainer. After washing and centrifugation, neurons were seeded on polyethylene-tetrathalate (PET) membrane (1 μm pores; Corning, NY) inserts, glass coverslips or glass-bottomed multiwell plates. All culture substrates were pre-coated with poly-D-lysine (Sigma, MO). *NbActive-1 medium* (BrainBits) supplemented with 100 U/ml of Penicillin-Streptomycin (Life Technologies, MA), 2 mM L-glutamine (Life Technologies), and 1 × N21 supplement (R&D Systems, MN) was used as culture medium. Inserts were seeded at a density of 1.5 × 10^6^ cells per insert and glass-plated cultures were seeded at 15,000 cells per 12 mm coverslip or well of a 24 well plate.

Neurons cultured on glass were used for morphological analyses with durations in culture indicated in the results. Neurons cultured in the PET inserts were used for isolation of neurites from lower membrane surface with cellular material along upper membrane referred to as a cell body preparation as previously described (Willis and Twiss, 2010).

For visualizing dendritic spines in neuron cultures, cortical and hippocampal neurons were transduced with AAV8-GFP (UNC Viral Vector Facility, NC) at 18 days *in vitro* (DIV). Cultures were then fixed as above at DIV23 and analyzed by fluorescent microscopy for GFP.

### RNA isolation and analyses

Total RNA was extracted from the neocortex and hippocampus of male 2-4 month old *Khsrp*^−/−^, *Khsrp^+/−^* and *Khsrp^+/−^* littermate mice using *Trizol* reagent (Invitrogen, CA). For analysis of RNAs from cultured neurons, cell body vs. neurite RNA was isolated using *RNAeasy Microisolation Kit* (Qiagen, CA).

For microarray analyses of transcript levels in cortices from adult males *Khsrp*^+/+^ vs *Khsrp*^−/−^, mRNAs were purified after removal of rRNA (*mRNA-ONLY™ Eukaryotic mRNA Isolation Kit*, Epicentre Biotech., WI). Fluorescently labeled cRNAs derived from these transcripts were used to probe Agilent *Mouse V4.0 LncRNA Array* containing probes for 22,692 mRNAs (ArrayStar, Inc., MD).

RNA-immunoprecipitation (RIP) assays were performed in triplicates using anti-KHSRP antibodies (NBP1-18910, Novus Biologicals, LLC, CO) pre-loaded onto protein G magnetic beads (ThermoFisher) as described in Bolognani et al. (2010). Eight sets of E18 cortices of mixes sexes from *Khsrp*^+/+^ mice were used for these assays and equal number of E18 cortices of *Khsrp*^−/−^ mice were used as controls for the RIP. After washing the beads, RNA was extracted using Trizol and sent for sequencing at the National Center for Genome Resources (NCGR, Santa Fe, NM).

Bioinformatics analyses used the following filters: 1) differentially expressed genes with adjusted p values <0.05 and fold change>1.25 in *Khsrp*^−/−^, 2) RIP-seq data including targets significantly enriched in the KHSRP RIP of *Khsrp*^+/+^ E18 cortex vs. *Khsrp*^−/−^ cortex using a p<0.05 and log_2_ fold-change >1.40 (fold change >2.63). IPA (Qiagen) was used to identify biological pathways and networks enriched in genes within the Nervous system development and function category.

Reverse-transcriptase (RT) coupled PCR was used to validate the KHSRP target mRNAs identified from micro-arrays and RIP-Seq analyses. All RTddPCR analyses were run on RNA isolates from at least 3 mice or culture preparations. For this, RNA yields were normalized across samples prior to reverse transcription based on fluorometric quantification using Ribogreen reagent (Invitrogen). 10-50 ng of RNA from brain samples or 10-25 ng of RNA from cell body and neurite preparations of neuron cultures was reverse transcribed using Sensifast (Bioline, TN). For the neuron cultures, single mouse pup cultures were performed from littermate mice with genotypes tested as above while the neurons were in culture (and analyses performed blinded to genotype). Tissues taken from mouse pups at the time of dissection was used for genotyping as outlined above. cDNA samples were diluted and then processed for extended cycle PCR (to test for neurite purity) or quantitative ddPCR. Extended cycle PCR, with primers for cell body (*cJun*) and glial contamination (*Gfap*) and *Map2* and *Actb* primers as positive control (i.e., neurite localizing mRNAs), was used to assess the purity of neurite RNA preparations. These PCR products were analyzed by agarose gel electrophoresis with ethidium bromide staining. For ddPCR, we used Evagreen reagent (Bio-Rad, CA) with an automated droplet generator; after standard PCR cycles, droplets were analyed using a QX200TM (Bio-Rad). Signals were normalized between reactions/samples using the mitochondrially encoded 12S ribosomal RNA (12S rRNA). Primer sequences are shown in Suppl. Table S9.

### Immunofluorescent staining

All immunofluorescence steps were conducted at room temperature unless specified otherwise. Neuron cultures were fixed with 4% paraformaldehyde (PFA) in phosphate-buffered saline (PBS) for 15 min and washed 3 times in PBS. Samples were permeabilized with 0.3% Triton X-100 in PBS for 15 min and blocked for 1 h in 5 % BSA in PBS + 0.1% Triton X-100 (PBST). Samples were then incubated overnight in humidified chambers at 4°C in the following primary antibodies diluted in blocking buffer: anti-MAP2 (1:700; Ab5392, Abcam, Cambridge, UK), SMI312 (1:250; 837904, BioLegend, CA), anti-HuD (1:400; Ab96474, Abcam), anti-KHSRP (1:500; NBP1-18910, Novus, CO), and Tuj1 (1:500; NB100-1612, Novus). After washes in PBST, coverslips were incubated for 1 h with combination of FITC-conjugated donkey anti-mouse, Cy5-conjugated donkey anti-chicken, and Cy3-conjugated donkey anti-rabbit antibodies (1:500 each; Jackson ImmunoRes., PA) diluted in blocking buffer. Samples were washed 3 times in PBS, rinsed with distilled H2O, and mounted with Prolong Gold Antifade with DAPI (Life Technologies, MA).

### Morphological assessments of axon and dendrite growth in KHSRP deficient mice

We used *Khsrp*^−/−^ mice crossed with Thy1-GFP mice to analyze axon and dendrite growth in vivo. Adult littermates (2-4 months old) consisting of Thy1-GFP/*Khrsp*^+/+^, Thy1-GFP/*Khrsp*^+/−^ and Thy1-GFP/*Khsrp*^−/−^ were perfused intracardially first with phosphate buffered saline (PBS; 37°C), followed by cold 4% paraformaldehyde (PFA, w/v) in PBS. Brains were post-fixed in 4% PFA at 4°C for 4 h and cryoprotected in 30% sucrose (w/v) in PBS for at least 2 d at 4°C. 50 μm thick coronal slices were cut on a freezing microtome. Sections were mounted onto coverslips coated with *Vectashield* (Vector Laboratories). For axonal growth, we assessed of the length of hippocampal mossy fiber IPB. Briefly, the length of GFP-positive mossy fibers in the IPB was measured from the cross section of the hilus at the end of the granule cell layer to the point they cross the pyramidal cell layer. IPB length was divided by the total length of the most medial aspect of the hilus to the apex of the curvature of CA3 as previously described (Perrone-Bizzozero et al., 2011).

For analyses of dendrite morphology in the KHSRP knockout mice, confocal images of apical dendrites of layer V pyramidal neurons in prefrontal cortex (labeled throughout the cell body and the dendritic tree with GFP) were by obtained by Leica SP8X confocal microscope using a 63x/NA 1.4 oil immersion objective at 1 μm Z intervals (1024 × 1024 pixel fields; Wetzlar, Germany). Image stacks consisted of 10-50 optical planes. Second-order dendritic shafts in these images were identified at distance of 100-200 μm from the soma were analyzed using *Neurolucida 360* and *Neurolucida Explorer* software (MBF Bioscience, VT). Spine density was assessed and each spine was categorized based on stalk length and head width as thin, stubby and mushroom using the default software parameters (Harris et al., 1992; Harris and Kater, 1994). These analyses was done blind to the individual genotypes.

### Analysis of neuronal morphology in culture

Cultured hippocampal and cortical neurons from *Khsrp^+/−^* crosses were assessed for axonal and dendritic growth at DIV7 and for dendritic spine density and morphology at DIV23. DIV7 cultures were fixed in 4% PFA and processed for immunofluorescence with MAP2 and SMI-312 to identify dendrites and axons, respectively. Images for neuronal morphology of DIV 7 neurons were captured on Leica DMI6000 epifluorescent microscope with ORCA Flash ER CCD camera (Hamamatsu Photonics, Shizuoka, Japan) with a 40x/NA 1.2 oil immersion objective using Leica LAS AF as tile scans taken randomly across each coverslip/well. For analyses of dendritic spines, GFP expressing DIV23 cultures were fixed with 4% PFA and imaged by confocal microscopy using Leica SP8X as above. For this, GFP-filled spines along 20 μm dendrite segments were imaged as Z stacks using 63X/NA 1.4 oil immersion objective. For both DIV7 and DIV23, neurons were imaged blinded to genotype.

Axon and dendrite morphology in cultured neurons was analyzed from epifluorescent tile scan images using *WIS-NeuroMath* (Rishal et al., 2013) to give average length of each process/neuron and branch density along each axon and dendrite. Dendritic spines in cultured neurons were traced from confocal image stacks using *Neurolucida 360* as above to quantitate spine density and spine type.

### Electrophysiological analyses

Coronal slices (300 μm thick) containing either the infralimbic cortex (PFC) or the dorsal hippocampus were prepared according to published protocols (Takano et al., 2012; Zhou et al., 2018) and maintained in artificial cerebrospinal fluid solution composed of 130 mM NaCl, 3 mM KCl, 1.25 mM NaH_2_PO_4_, 26 mM NaHCO_3_, 10 mM glucose, 1 mM MgCl_2_, and 2 mM CaCl_2_. Slices were continuously perfused with artificial cerebrospinal fluid (aCSF) heated to 32 ± 1°C. Whole-cell patch clamp recordings were done with a K-gluconate based intracellular solution. mEPSCs were recorded at a holding potential of −65 mV in the presence of bath-applied 1 μM tetrodotoxin. Cells from 3-6 mice were analyzed per genotype. Mean mEPSC amplitude and duration were measured from an average trace of 50-100 individual mEPSCs. Duration was computed as a monoexponential fit to the decay phase of the average mEPSC.

### Behavioral studies

Animals were maintained on a reverse 12 h dark/light cycle (lights on at 20:00 hours) in grouped-housed cages. Behavioral testing was conducted using male and female *Khsrp*^−/−^, *Khsrp*^+/−^ and *Khsrp*^+/+^ mice that were age- and sex-matched. Methods for behavioral tests are described in Supplemental Methods. Note that as KHSRP has also been recently identified to interact with the circadian rhythm by targeting PER2 (Chou et al., 2015); therefore, all behavioral measurements were conducted during the dark period between 09:00 and 17:00 in behavioral rooms lit with red lighting.

### Statistical Analyses

All statistical tests were performed using *Prism* software package (version 8.4.0; GraphPad). One-way ANOVAs were used to compare the molecular, morphological, and electrophysiological differences in *Khsrp*^+/−^, *Khsrp*^−/−^ and *Khsrp*^+/+^ mice and post-hoc Tukey tests were used to identify significant changes between two genotypes. Student *t*-tests were used for the analyses of KHSRP-target mRNAs levels in *Khsrp*^−/−^ and *Khsrp^+/−^* vs. *Khsrp*^+/+^ mice. Repeated measures (RM) ANOVA were used for analyses of percent center duration, distance and velocity d 1-5, distances and velocity d 1, 0-5 min through 20-25 in the open field. Tukey post hoc tests were utilized for open field distance and velocity d 1-5, distances and velocity d 1, 0-5 through 20-25 min. Two-way ANOVAs were used to determine if there was a main effect of sex and genotype for the percent open arm duration, distance, and velocity in the zero maze. RM ANOVA was used to determine overall effects of problem stages in ASST and one-way ANOVA was used to compare starting dimension effects of specific stages. To compare each genotype between stages (CDR-CD, IDR-IDS, and EDR-IDS) Student *t*-tests were employed. Lastly, RM ANOVA was utilized for the Tone Training, Trace Training, as well as the Tone-Trace Test followed by individual one-way ANOVAs for each time point in Trace Fear Conditioning.

## ACKNOWLEDGEMENTS

We thank the NM-INBRE Sequencing and Bioinformatics Core (SBC) at the National Center for Genome resources (NCGR) for providing pilot funding and expertise for the KHSRP RIP-seq studies and Ms. Gabriela Perales for her help determining axonal growth in hippocampal slices.

## ADDITIONAL INFORMATION

| **Funder** | **Grant Number** | **Author** |
| --- | --- | --- |
| National Institutes of Health | R01-NS089663 | JL Twiss and<br>N Perrone-Bizzozero |
| NIH-NIGMS IDeA Networks of<br>Biomedical Research Excellence<br>(INBRE) | P20GM103451 | A Sundara and<br>J Mudge |
| Dr. Miriam and Sheldon G. Adelson<br>Medical Research Foundation | unnumbered award | JL Twiss |
| South Carolina EPSCoR Stimulus<br>Research Program | SRP 18-04 | JL Twiss |
| Wings for Life Spinal Cord Injury<br>Research Foundation | WFL-US-09/18 | P Patel |

## AUTHOR CONTRIBUTIONS

Sarah L. Olguin, Priyanka Patel, Michela Dell’Orco, Amy S. Gardiner – experimental design, performed experiments, analyzed data, drafted manuscript.

Robert Cole, Courtney Buchanan, Anitha Sundara, Joann Mudge – performed experiments and analyzed data.

Andrea M. Allan, Pavel Ortinski, Jonathan L. Brigman – experimental design and oversight.

Jeffery L. Twiss, Nora I. Perrone-Bizzozero – experimental design and oversight, project oversight, funded experiments, revised manuscript drafts.

