## Supplemental Materials for "Loss of KHSRP Increases Neuronal Growth and Synaptic Transmission and Alters Memory Consolidation Through RNA Stabilization"

**SUPPLEMENTARY METHODS**

**Animals –** Two cohorts of mice consisting of 28 wild type WT mice (15 male and 13 female), 33 *Khsrp*$^{-/-}$ (17 male and 16 female), and 10 *Khsrp*$^{+/-}$ (5 male and 5 female) were used for behavioral studies as indicated below. Cohort 1 (32 mice) began preliminary behavioral screens between the ages of 9-18 weeks, followed by zero maze at 9-19, open field at 10-21, and Attentional Set Shifting Task (ASST) at ages 17-27 weeks. Cohort 2 (39 mice) began the preliminary behavioral screen between the ages of 10-21 weeks, followed by zero maze at 10-24, open field at 11-24, ASST at 18-29 weeks, and trace fear conditioning at 23-33 weeks.

**Preliminary Behavioral Screen** – All mice were assessed using a subset of tests derived from the Irwin screen as previously described (Irwin 1968, Crawley 1999, Zhao et al. 2006) for physical health, sensory, motor, and neurological function. Exploratory behaviors were observed by placing the mouse in a corner of a clear box (45 x 45 x 22 cm) and recording for 10 min.

**Elevated Zero-Maze Test** – The elevated zero-maze test was conducted as previously described (Díaz-Morán et al. 2014) on a white circular platform (5 cm runway, 60 cm diameter and 50 cm from the floor) consisting of 2 opposing open quadrants with a 0.5 cm raised lip to prevent falling and 2 opposing closed quadrants with 15 cm high walls. The room was illuminated with red fluorescent lights and two single white lights (open arms 90 lux, closed arms 45 lux ). Mice were allowed to freely explore the arena for 5 min. Locomotor activity and time spent in the open vs closed arms was measured using *Ethovision* video tracking system (Noldus Information Technology, VA).

**Open Field Test** – The open field test was conducted as previously described (Wiedholz et al. 2008) in a square arena (40 x 40 x 35 cm) constructed from white Plexiglas. The room was illuminated with red fluorescent lights and two single white lights (center 60 lux, corner 35 lux). Mice were placed in the NW corner of the arena and allowed to freely explore for 30 min per day for 5 consecutive days to establish a baseline of anxiety and locomotor activity. Total distance traveled, velocity, and duration in the (20 x 20 cm) center was measured using *Ethovision*.

**Attentional Set Shifting Task (ASST)** – ASST was conducted as previously described (Young et al. 2010, Marquardt et al. 2014, Thompson et al. 2015). Testing was conducted in an acrylic apparatus (30 x 18 x 12 cm) divided into a start box and 2 choice chambers. Each choice chamber contained a ceramic digging bowl (4.5 x 2.5 cm) placed on an in house manufactured platform (11 x 5 cm) with sandpaper, wood, neoprene, metal wire, tile, or a plastic fiber sponge as textures. Scented medium was made by mixing 150 g of cob bedding with 20 crushed 14 mg dustless precision pellets (#F0568, BioServ, NJ) and 3 g of commercially available powdered spices: nutmeg, ginger, garlic, coriander, thyme, and cinnamon (Kroger Co., OH). Prior to training animals were reduced to 85% free feeding weight and acclimated to food reward in the home cage. Training day 1 consisted of acclimation to the testing chamber and digging in unscented cob medium for reward. Training day 2 introduced the mice to all exemplar combinations encountered during testing (Table 1). A single pellet was placed below the cob in one bowl, randomly assigned between trials, and placement was mimicked in the unrewarded bowl to prevent mice learning experimenter cues. Testing on day 3 was conducted in succession with no inter-session-breaks on 7 discrimination tasks (Table 1). In simple discrimination (SD) mice were trained to discriminate 2 exemplars in either the odor or platform dimension (counterbalanced across genotypes and sex n=34). Upon reaching criterion, mice were moved to compound discrimination (CD) in which the second, non-rewarded dimension, was added. The rewarded exemplar in the initially rewarded dimension was then reversed to form a compound discrimination reversal (CDR). Next, a novel set of exemplars in each dimension were introduced and mice were rewarded for responding to one exemplar in the initially learned dimension (intra-dimensional shift (IDS)). Next, the intra-dimensional reversal (IDR) reversed the correct stimuli within the same dimension. A second novel set of exemplars in both dimensions were introduced in the extra-dimensional shift (EDS) where the rewarded exemplar was in the previously irrelevant dimension. Finally, the correct exemplar within the newly learned dimension was reversed to form an extra-dimensional reversal (EDR).

Trials to criterion, corrects, and errors were recorded for each stage. If a mouse did not dig in either bowl by 2 min the trial was recorded as 'no choice' and the mouse repeated the trial until a choice was made. Criterion was set to 6 consecutive correct responses. Trial latencies to respond were measured from the time the barrier was raised until digging was initiated. A dig was defined as the moment when the mouse's nose or paw broke the surface of the cob-digging medium. Mice were discontinued if they required 60 trials on any 1 problem or 150 trials total.

**Supplementary Methods Table 1**: ***Example problem stages and stimulus combinations for the Attentional Set Shifting Task.*** O = odor and P = Platform, with number indicating different types of odors and platforms.

| **Problem Stage** | **Dimensions** | | **Exemplars** | |
|---|---|---|---|---|
| | **Relevant** | **Irrelevant** | **S+** | **S-** |
| Simple discrimination (SD) | Odor | n/a | O1 | O2 |
| Compound discrimination (CD) | Odor | Platform | O1/P1 | O2/P2 |
| | | | O1/P2 | O2/P1 |
| Compound discrimination reversal (CDR) | Odor | Platform | O2/P2 | O1/P1 |
| | | | O2/P1 | O1/P2 |
| Intradimensional shift (IDS) | Odor | Platform | O3/P4 | O4/P4 |
| | | | O3/P3 | O4/P3 |
| Intradimensional shift reversal (IDR) | Odor | Platform | O4/P4 | O3/P4 |
| | | | O4/P3 | O3/P3 |
| Extradimensional shift (EDS) | Platform | Odor | P5/O5 | P6/O5 |
| | | | P5/O6 | P6/O6 |
| Extradimensional shift reversal (EDR) | Platform | Odor | P6/O5 | P5/O5 |
| | | | P6/O6 | P5/O6 |

**Trace Fear Conditioning –** Studies were conducted between 0900 and 1200 hours under dim red illumination, as previously described (Brady et al. 2012). Briefly, animals were placed into a Coulburn Instruments (Whitehall, PA) Habitest System for 90 seconds of habituation, followed by 7 trials each consisting of the CS (10 seconds, 80 dB 6 Hz clicker), a 30 second trace, the US (1 second, 0.8 mA scrambled foot shock), and a 180 second inter-trial interval. The subject was removed from the chamber 60 seconds following the delivery of the last US.

24 hours later, freezing to the CS in a novel context (a standard, clean mouse cage with minimal bedding) was assessed. The CS was delivered at 180, 310, and 440 seconds. The animal's behavior was

videotaped and the amount of time spent freezing during the tone and during the 30 second trace (40 seconds total duration) was scored for each CS by 2 investigators, one of which was blinded to the genotype. The average for the 3 measures was calculated and expressed as a percentage of time spent freezing.

#### SUPPLEMENTARY METHODS REFERENCES


Brady, M. L., A. M. Allan and K. K. Caldwell (2012). "A limited access mouse model of prenatal alcohol exposure that produces long-lasting deficits in hippocampal-dependent learning and memory." Alcohol Clin Exp Res **36**: 457-466.

Crawley, J. N. (1999). Behavioral phenotyping of transgenic and knockout mice: Experimental design and evaluation of general health, sensory functions, motor abilities, and specific behavioral tests. Brain Res **835:** 18-26.

Díaz-Morán, S., C. Estanislau, T. Cañete, G. Blázquez, A. Ráez, Adolf Tobeña and A. Fernández-Teruel (2014). "Relationships of open-field behaviour with anxiety in the elevated zero-maze test: Focus on freezing and grooming." World J Neurosci **4**: 1-11.

Irwin, S. (1968). Comprehensive observational assessment: Ia. A systematic, quantitative procedure for assessing the behavioral and physiologic state of the mouse. Psychopharmacologia **13:** 222-257.

Marquardt, K., M. Saha, M. Mishina, J. W. Young and J. L. Brigman (2014). "Loss of GluN2A-containing NMDA receptors impairs extra-dimensional set-shifting." Genes Brain Behav **13**: 611-617.

Thompson, S. M., M. Josey, A. Holmes and J. L. Brigman (2015). Conditional loss of GluN2B in cortex and hippocampus impairs attentional set formation. Behav Neurosci **129:** 105-112.

Wiedholz, L. M., W. A. Owens, R. E. Horton, M. Feyder, R. M. Karlsson, K. Hefner, R. Sprengel, T. Celikel, L. C. Daws and A. Holmes (2008). "Mice lacking the AMPA GluR1 receptor exhibit striatal hyperdopaminergia and 'schizophrenia-related' behaviors." Mol Psychiatry **13**: 631-640.

Young, J. W., S. B. Powell, M. A. Geyer, D. V. Jeste and V. B. Risbrough (2010). "The mouse attentional-set-shifting task: a method for assaying successful cognitive aging?" Cogn Affect Behav Neurosci **10**: 243-251.

Zhao, S., J. Edwards, J. Carroll, L. Wiedholz, R. A. Millstein, C. Jaing, D. L. Murphy, T. H. Lanthorn and A. Holmes (2006). Insertion mutation at the C-terminus of the serotonin transporter disrupts brain serotonin function and emotion-related behaviors in mice. Neurosci **140:** 321-334.

#### SUPPLEMENTAL MATERIALS

#### SUPPLEMENTAL TABLES

**Table S1**: ***Analyses of mRNA levels in KHSRP deficient mouse neocortex.***

**Table S2**: ***Identification of KHSRP-target mRNAs through RIP-Seq.***

**Table S3**: ***Overlapping datasets of KHSRP targets***

**Table S4**: ***KHSRP-target mRNAs from integrated expression and RIP-Seq data contain AREs.***

**Table S5**: ***RTddPCR validation of KHSRP-target mRNAs in neocortex.***

**Table S6**: ***RTddPCR validation of KHSRP-target mRNAs in hippocampus.***

**Table S7**: ***RTddPCR validation of KHSRP-target mRNAs in cortical neuron cultures.***

**Table S8**: ***Behavioral analyses of KHSRP deficient mice.***

**Table S9**: ***Primers for RTddPCR analyses.***

#### SUPPLEMENTAL FIGURE LEGENDS

**Figure S1**: ***KHSRP expression progressively increases with maturation of neuronal cultures.***

**A)** Representative epifluorescent images for Tuj1 (cyan), HuD (green) and KHSRP (red) for DIV3-7 cortical neuron cultures from wild type mice are shown [scale bar = 100 µm]. Imaging parameters for individual channels are matched for exposure and gain across each respective column. Arrows indicate neurites where there is a progressive increase in KHSRP and decrease in HuD over days in culture. **B-C)** Quantification of signal intensities from cultures as in A for KHSRP (**B**) and HuD (**C**) in cell bodies and neurites is shown (N > 30 neurons in at least 3 separate cultures; * $p \leq 0.05$, ** $p \leq 0.01$, and *** $p \leq 0.005$ by one-way ANOVA with Tukey post-hoc compared to DIV3 as indicated).

**Figure S2**: ***Elevated KHSRP-target mRNAs in hippocampi of KHSRP deficient mice.***

Heatmap for $Log_2$ fold-change for KHSRP-target mRNAs in *Khsrp*$^{+/-}$ and *Khsrp*$^{-/-}$ vs. *Khsrp*$^{+/+}$ hippocampal RNA isolations is shown (N ≥ 4 per condition; * p ≤ 0.05, ** p ≤ 0.01, and *** p ≤ 0.005 by two-tailed Student's *t*-test).

**Figure S3**: ***KHSRP deficient hippocampal neurons show increased axon and dendrite growth in culture.***

**A)** Representative epifluorescent images for MAP2 (red) and SMI312 (green) immuno-stained hippocampal neurons from *Khsrp*$^{+/+}$, *Khsrp*$^{+/-}$, and *Khsrp*$^{-/-}$ mice at DIV7. **B-C)** Quantification of length and branching for axons and dendrites from DIV7 hippocampal cultures from *Khsrp*$^{+/+}$, *Khsrp*$^{+/-}$, and *Khsrp*$^{-/-}$ mice (N > 30 neurons in at least 3 separate cultures; ** p ≤ 0.01 and *** p ≤ 0.005 by one-way ANOVA with Tukey post-hoc compared to DIV3 as indicated).

**Figure S4**: ***KHSRP deficient neurons show increased dendrite spine density in culture.***

**A-B)** Representative epifluorescent images of distal dendrites for DIV23 cultures of hippocampal (**A**) and cortical (**B**) neurons from *Khsrp*$^{+/+}$ and *Khsrp*$^{-/-}$ mice shown as indicated. **C-D)** Quantification of dendritic spine density and morphology for DIV23 *Khsrp*$^{+/+}$ and *Khsrp*$^{-/-}$ hippocampal (**C**) and cortical (**D**) neurons is shown (N > 30 neurons in at least 3 separate cultures; * p ≤ 0.05 and ** p ≤ 0.01 by Student's *t*-test).

**Figure S5**: **Lack of anxiety like-behavior but increased locomotor activity of KHSRP deficient mice.**

Adult *Khsrp*$^{+/+}$ (n=28) *Khsrp*$^{+/-}$ (n=10) and *Khsrp*$^{-/-}$ (n=25) of both sexes were placed in a Zero Maze apparatus and the percentage time in the open arm (**A**), distance travelled (**B**) and velocity (**C**) were determined both at day 1 and day 5 of training. No changes were observed on percentage time in the open arm but ANOVA analyses showed increased locomotor activity as measured by distance traveled and velocity in the zero maze (ANOVA distance traveled:

$F(2,52)=4.606$ $p=0.0143$; Velocity: $F(2,53)=6.915$, $p=0.0022$; *$p<0.05$ and **$p<0.01$, Dunnett's multiple comparisons test).

**<u>Figure S6</u>: No difference in trace conditioning in male and female mice of any genotype.** The freezing during training (**A**) or testing (**B**) did not differ in males vs. female mice (n=5 for male mice and n=4 for female mice of all genotypes; no significant differences in A by repeated measures ANOVAs and in B by one way ANOVAs for each sex.

### Supplemental Figure S1

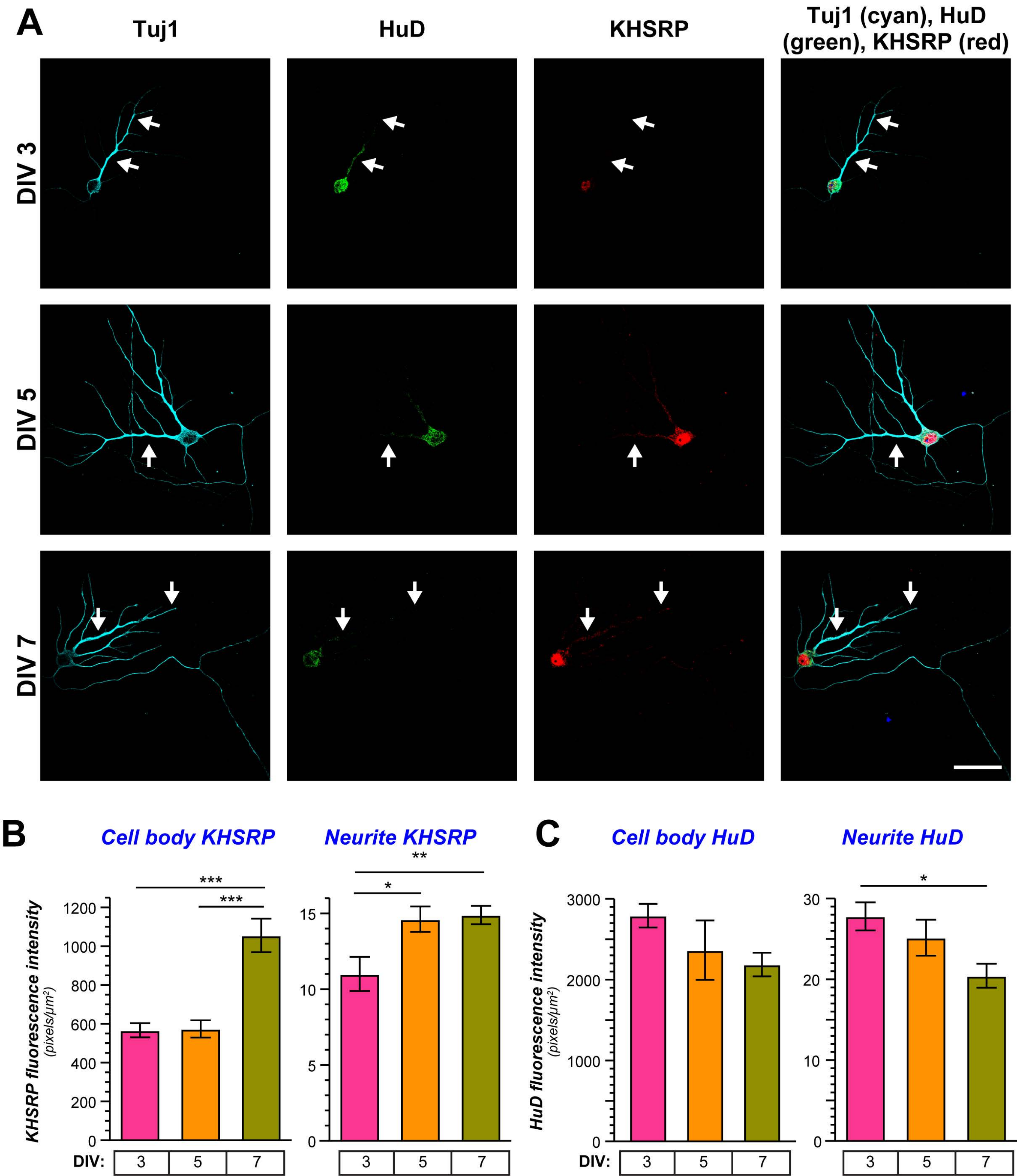

### Supplemental Figure S2

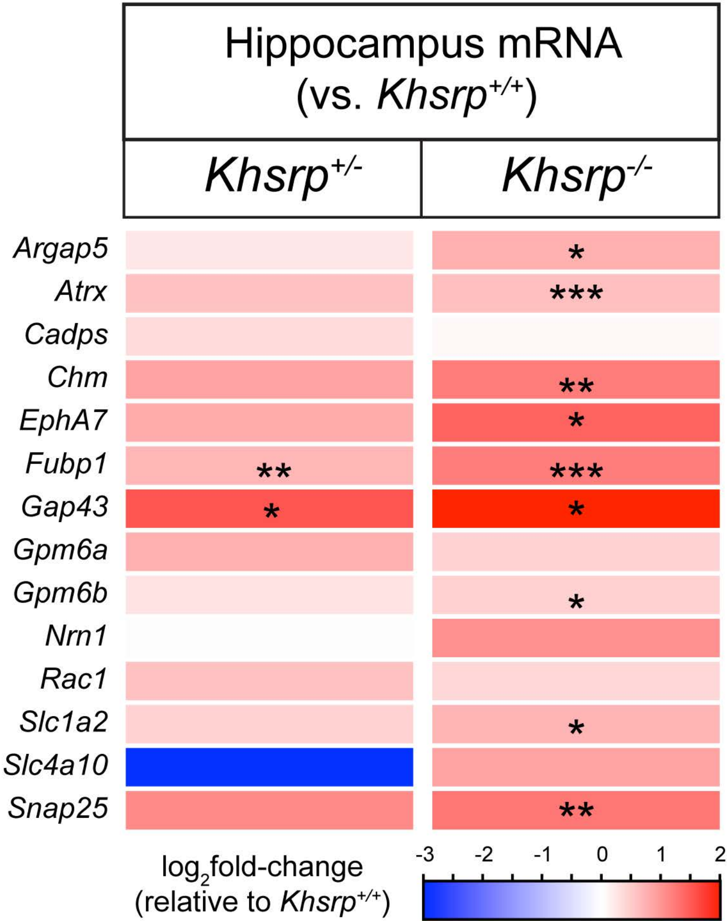

### Supplemental Figure S3

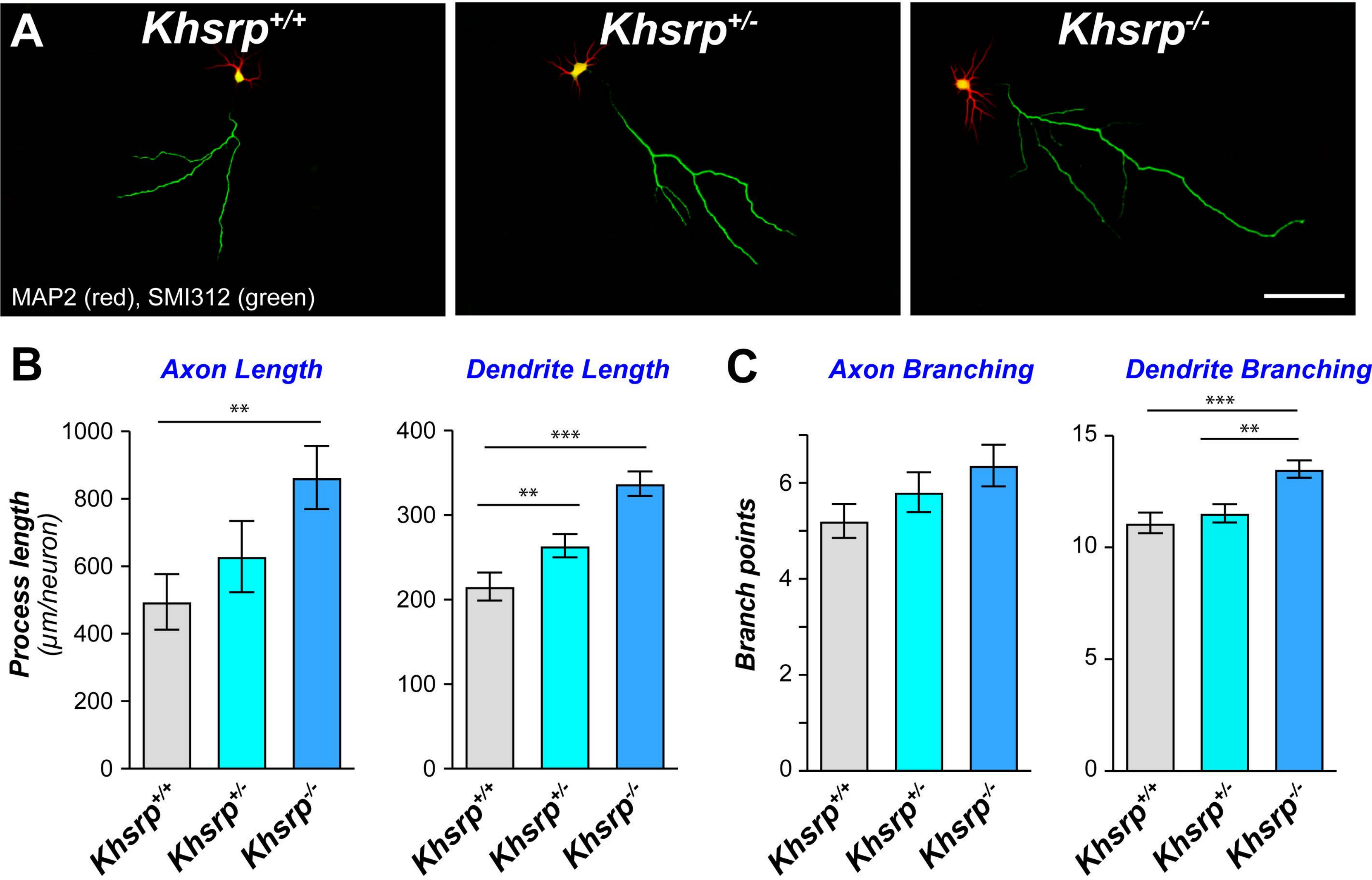

### Supplemental Figure S4

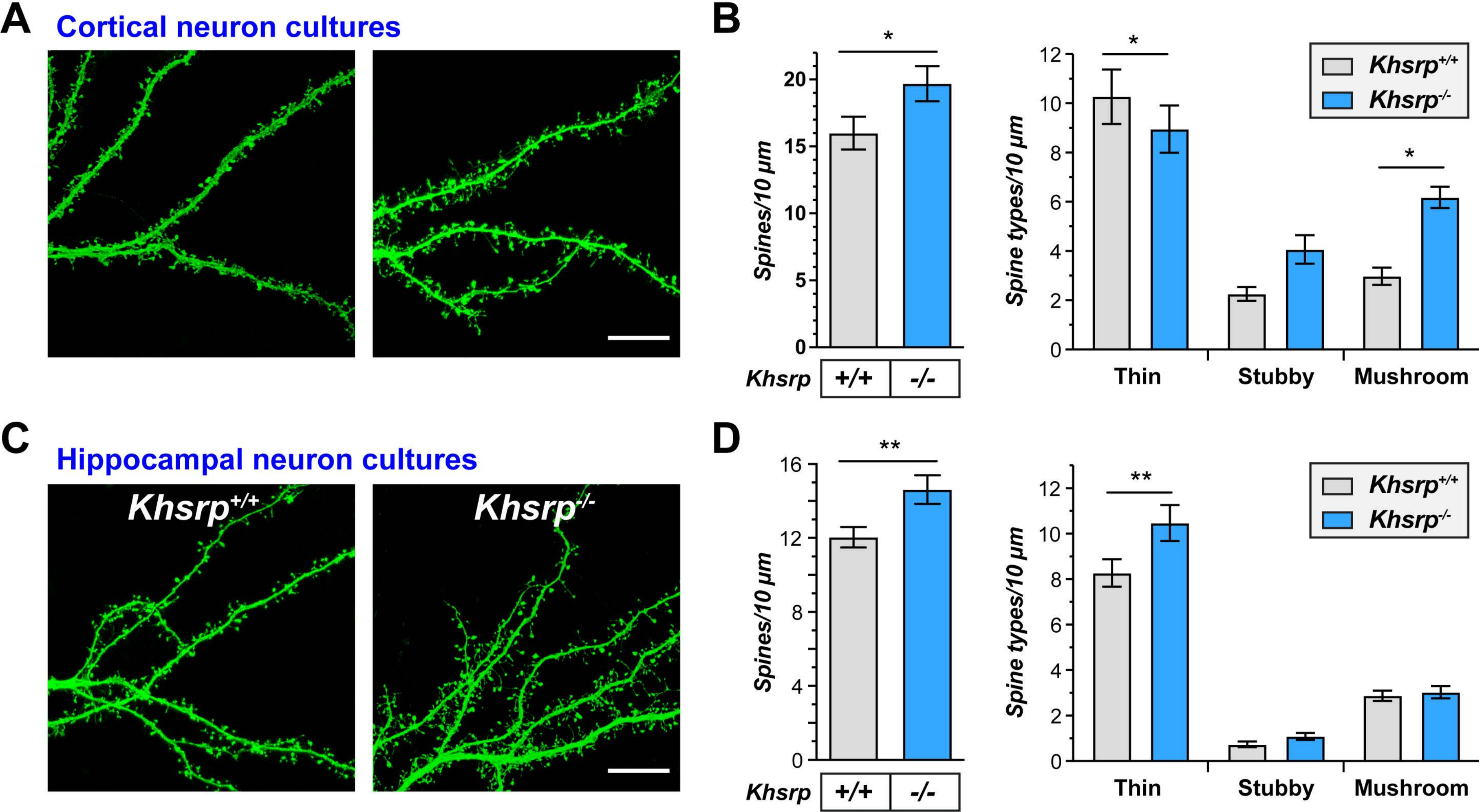

#### Supplemental Figure S5

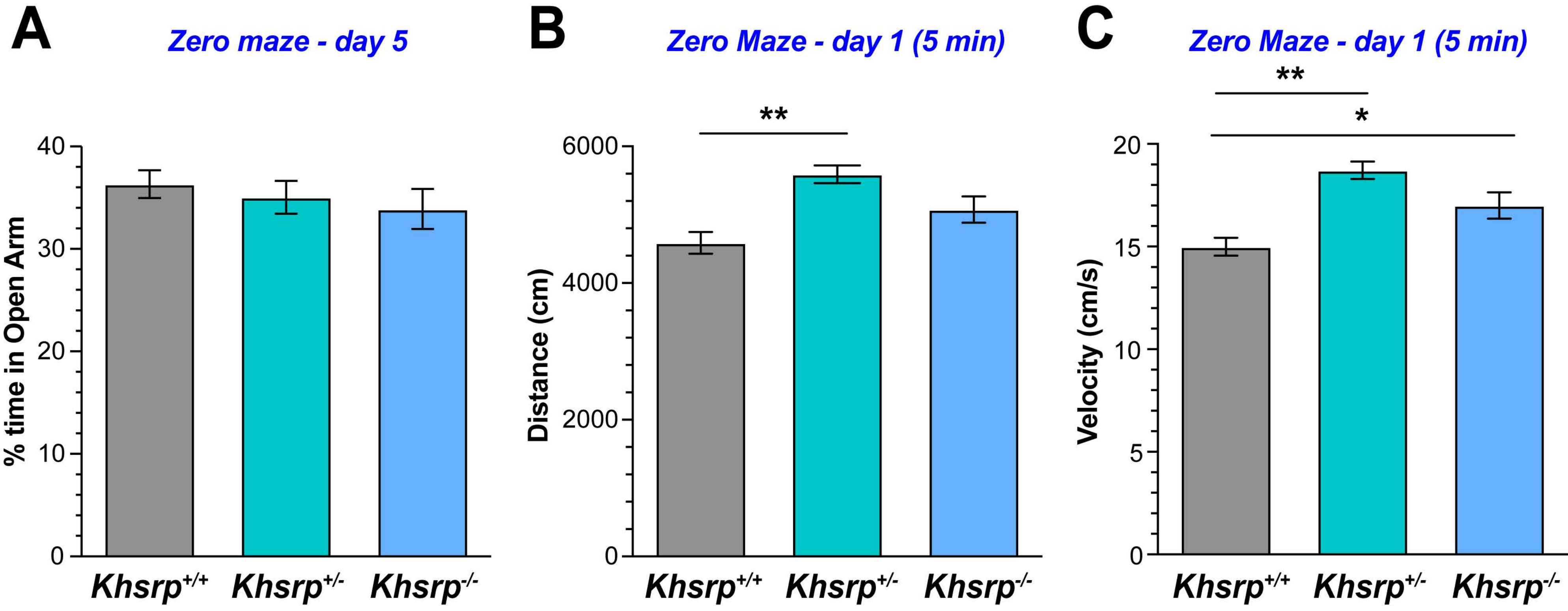

### Supplemental Figure S6

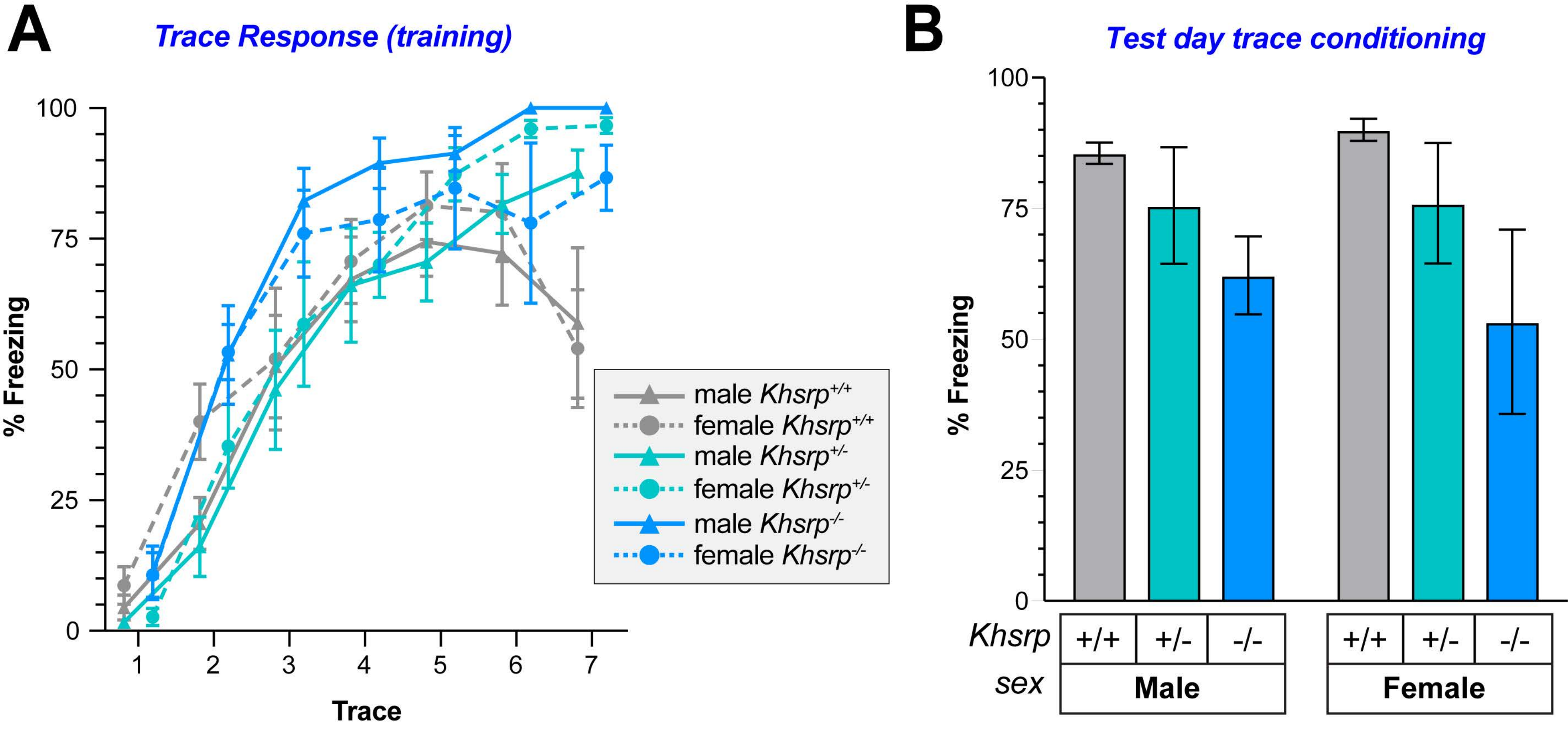
