## Supplemental Tables S1-S9 for "Loss of KHSRP Increases Neuronal Growth and Synaptic Transmission and Alters Memory Consolidation Through RNA Stabilization"

**Table of contents for Supplemental Tables**

**Table S1** : Analyses of mRNA levels in KHSRP deficient mouse neocortex.
**Table S2** : Identification of KHSRP-target mRNAs through RIP-Seq.
**Table S3:** Overlapping datasets of KHSRP targets
**Table S4:** KHSRP-target mRNAs from integrated expression and RIP-Seq data contain AREs.
**Table S5:** RTddPCR validation of KHSRP-target mRNAs in neocortex.
**Table S6:** RTddPCR validation of KHSRP-target mRNAs in hippocampus.
**Table S7:** RTddPCR validation of KHSRP-target mRNAs in cortical neuron cultures.
**Table S8:** Behaviorial analyses of KHSRP deficient mice
**Table S9:** Primer sequences used for RTddPCR.

**Supplemental Table S1 : Analyses of mRNA levels in KHSRP deficient mouse neocortex.**

All Targets Value (Entities where at least 3 out of 12 samples have flags in Present or Marginal)

Samples were probed on Agilent Mouse V4.0 LncRNA Array containing probes for 22,692 mRNAs.

### Column A: ProbeName it represents probe name.
### Column B: P-value, P-value calculated from unpaired t-test.
### Column C: FDR, FDR is calculated from Benjamini Hochberg FDR.
### Column D: Fold Change, the absolute ratio (no log scale) of normalized intensities between two conditions.
### Column E: Regulation, it depicts which group has greater or lower intensity values wrt other group.
### Column F ~ H: Annotations, include seq, GeneSymbol, source, and description.
### Column I,J : Raw Intensity of each group (averaged raw intensities of replicate samples).
### Column K, L: Normalized Intensity of each group (averaged normalized intensities of replicate samples, log2 transformed).
### Column M~R: Raw Intensity of each sample.
### Column S ~ X: Normalized Intensity of each sample (log2 transformed).

| ProbeName | P-value | FDR | Fold Change | Regulation | seqname | GeneSymbol | description | group-KSRP-KO-PFC(raw) | group-Wild-Type-PFC(raw) | group-KSRP-KO-PFC(normalized) | group-Wild-Type-PFC(normalized) | [KSRP-KO-PFC-1, KSRP-KO-PFC](raw) | [KSRP-KO-PFC-2, KSRP-KO-PFC](raw) | [KSRP-KO-PFC-3, KSRP-KO-PFC](raw) | [WT-PFC-1, Wild-Type-PFC](raw) | [WT-PFC-2, Wild-Type-PFC](raw) | [WT-PFC-3, Wild-Type-PFC](raw) | [KSRP-KO-PFC-1, KSRP-KO-PFC](normalized) | [KSRP-KO-PFC-2, KSRP-KO-PFC](normalized) | [KSRP-KO-PFC-3, KSRP-KO-PFC](normalized) | [WT-PFC-1, Wild-Type-PFC](normalized) | [WT-PFC-2, Wild-Type-PFC](normalized) | [WT-PFC-3, Wild-Type-PFC](normalized) |
|---|---|---|---|---|---|---|---|---|---|---|---|---|---|---|---|---|---|---|---|---|---|---|---|
| ASMM10P048 | 0.00000 | 0.0487938 | 131.64418 | up | NM_001037 | Xafl | XIAP associated factor 1 | 6074.027 | 51.587513 | 12.743932 | 5.703432 | 5589.3853 | 5998.2476 | 6634.448 | 50.43867 | 54.3965 | 49.92737 | 12.474701 | 12.802467 | 12.954629 | 5.557416 | 5.6548 | 5.8980813 |
| ASMM10P051 | 0.00001 | 0.0712172 | 6.9397395 | up | NM_177839 | Tnn | tenascin N | 28.735846 | 5 | 5.13425 | 2.339368 | 30.329672 | 24.945026 | 30.93284 | 4.9999995 | 4.9999995 | 4.9999995 | 4.978237 | 5.0988865 | 5.325626 | 2.3250442 | 2.3265402 | 2.3665206 |
| ASMM10P049 | 0.00002 | 0.0712172 | 1.9670746 | up | NM_030696 | Slc16a3 | solute carrier family 16 (monocarboxylic acid transporters), member 3 | 86.183183 | 53.180729 | 6.720394 | 5.744342 | 97.98539 | 80.25874 | 80.30542 | 56.21376 | 56.558704 | 46.769722 | 6.6666107 | 6.7518916 | 6.7426786 | 5.714845 | 5.7096524 | 5.8085284 |
| ASMM10P000 | 0.00002 | 0.0712172 | 1.705975 | up | NM_001081 | Kcnt2 | potassium channel, subfamily T, member 2 | 51.147905 | 36.128377 | 5.965229 | 5.194632 | 60.031315 | 44.467884 | 48.944515 | 39.386616 | 38.453953 | 30.544563 | 5.96757 | 5.9180365 | 6.01008 | 5.207371 | 5.1554384 | 5.2210875 |
| ASMM10P042 | 0.00002 | 0.0712172 | 2.0094855 | up | NM_011154 | Ppp2r3d | protein phosphatase 2 (formerly 2A), regulatory subunit B", delta | 1420.3374 | 840.76714 | 10.692703 | 9.685877 | 1647.1995 | 1314.1704 | 1299.6422 | 828.108 | 884.38007 | 809.81335 | 10.672764 | 10.697886 | 10.707459 | 9.613175 | 9.681167 | 9.763288 |
| ASMM10P047 | 0.00004 | 0.091865 | 2.5483651 | up | NM_026029 | Glod4 | glyoxalase domain containing 4 | 1410.762 | 659.12844 | 10.685909 | 9.336337 | 1559.1185 | 1324.7311 | 1348.4365 | 643.26117 | 698.88196 | 635.2422 | 10.59109 | 10.710099 | 10.756538 | 9.253753 | 9.339611 | 9.415647 |
| ASMM10P000 | 0.00004 | 0.0965186 | 1.6341874 | up | NM_026714 | Ccdc163 | coiled-coil domain containing 163 | 75.486543 | 55.784857 | 6.519073 | 5.8105 | 92.37359 | 65.5182 | 68.56784 | 60.096706 | 59.722446 | 47.53542 | 6.583042 | 6.4654393 | 6.5087376 | 5.8152137 | 5.78424 | 5.832045 |
| ASMM10P006 | 0.00006 | 0.1172169 | 1.2887548 | up | NM_009722 | Atp2a2 | ATPase, Ca++ transporting, cardiac muscle, slow twitch 2 | 17120.918 | 15650.839 | 14.213985 | 13.848007 | 17938.299 | 17175.332 | 16249.124 | 16269.513 | 16458.76 | 14224.244 | 14.199723 | 14.244304 | 14.197929 | 13.86476 | 13.820313 | 13.85895 |
| ASMM10P033 | 0.00009 | 0.1436064 | 1.2904835 | up | NM_008098 | Mtpn | myotrophin | 2891.9995 | 2645.1256 | 11.689122 | 11.32121 | 3310.3872 | 2749.7397 | 2615.8716 | 2645.7063 | 2804.7366 | 2484.9338 | 11.681902 | 11.712051 | 11.673412 | 11.282801 | 11.334611 | 11.346218 |
| ASMM10P027 | 0.00010 | 0.1533694 | 2.3573304 | up | NM_001134 | CK137956 | cDNA sequence CK137956 | 4444.6981 | 2220.9334 | 12.305483 | 11.068329 | 4599.56 | 4131.61 | 4602.9243 | 2242.0532 | 2393.3733 | 2027.3738 | 12.195555 | 12.264744 | 12.456151 | 11.050702 | 11.096836 | 11.05745 |
| ASMM10P014 | 0.00011 | 0.1586373 | 1.8221819 | up | NM_011532 | Tbx1 | T-box 1 | 2270.1471 | 1476.6747 | 11.345299 | 10.479632 | 2660.6772 | 2001.3328 | 2148.4314 | 1426.6234 | 1633.2443 | 1370.1565 | 11.35633 | 11.280354 | 11.399213 | 10.394453 | 10.548532 | 10.495911 |
| ASMM10P020 | 0.00014 | 0.1714836 | 1.4646115 | up | NM_144884 | Tor1a | torsin family 1, member A (torsin A) | 1772.3798 | 1456.3676 | 11.01072 | 10.460202 | 2026.7239 | 1632.1466 | 1658.2689 | 1493.9598 | 1591.7618 | 1283.3813 | 10.977595 | 10.998579 | 11.055985 | 10.461738 | 10.512477 | 10.40639 |
| ASMM10P000 | 0.00014 | 0.1714836 | 1.4950244 | up | NM_145375 | Tm6sf1 | transmembrane 6 superfamily member 1 | 742.32347 | 592.53253 | 9.760545 | 9.180376 | 909.0583 | 658.9665 | 658.9456 | 605.15436 | 648.85443 | 523.5888 | 9.818216 | 9.72489 | 9.738528 | 9.164482 | 9.235774 | 9.140871 |
| ASMM10P033 | 0.00017 | 0.1837477 | 1.4008629 | up | NM_001037 | Gm4925 | predicted gene 4925 | 40.736292 | 34.943032 | 5.63087 | 5.144554 | 49.56256 | 35.173756 | 37.47256 | 37.320927 | 38.186 | 29.32217 | 5.696961 | 5.5850496 | 5.6105995 | 5.1241574 | 5.144876 | 5.1646295 |
| ASMM10P048 | 0.00019 | 0.1912656 | 1.3321141 | up | NM_134013 | Psme4 | proteasome (prosome, macropain) activator subunit 4 | 956.65677 | 853.3246 | 10.119523 | 9.705805 | 1157.0161 | 843.7756 | 869.1786 | 866.69257 | 917.247 | 776.03424 | 10.152063 | 10.066772 | 10.139734 | 9.676638 | 9.734639 | 9.706139 |
| ASMM10P001 | 0.00021 | 0.2026732 | 1.6521897 | up | NM_001142 | Pcdh15 | protocadherin 15 | 25.795125 | 18.690624 | 4.975165 | 4.250786 | 28.626638 | 24.336199 | 24.422539 | 19.910748 | 21.103065 | 15.058059 | 4.8921084 | 5.0643816 | 4.9690056 | 4.2034206 | 4.295677 | 4.25326 |
| ASMM10P018 | 0.00024 | 0.2113654 | 1.9603733 | up | NM_027084 | Prss56 | protease, serine 56 | 8.212015 | 5 | 3.310497 | 2.339368 | 10.798631 | 6.7476473 | 7.0897655 | 4.9999995 | 4.9999995 | 4.9999995 | 3.4341528 | 3.32874 | 3.1685975 | 2.3250442 | 2.3265402 | 2.3665206 |
| ASMM10P007 | 0.00025 | 0.2113654 | 1.413122 | up | NM_017469 | Gucy1b3 | guanylate cyclase 1, soluble, beta 3 | 4909.8504 | 4084.4723 | 12.443467 | 11.944581 | 5385.4165 | 4825.79 | 4518.3447 | 4265.7344 | 4109.55 | 3878.1326 | 12.418829 | 12.486871 | 12.4247 | 11.968344 | 11.877171 | 11.988227 |
| ASMM10P004 | 0.00026 | 0.2113654 | 2.0198417 | up | NM_001109 | Ctnna2 | catenin (cadherin associated protein), alpha 2 | 84.361785 | 50.401801 | 6.681992 | 5.66775 | 100.26538 | 70.208275 | 82.6117 | 56.73153 | 50.882072 | 43.5918 | 6.696957 | 6.564891 | 6.7841287 | 5.730877 | 5.561 | 5.711373 |
| ASMM10P044 | 0.00030 | 0.2233886 | 2.2499115 | up | NM_025660 | Ribc1 | RIB43A domain with coiled-coils 1 | 328.22765 | 177.14879 | 8.628704 | 7.458835 | 363.5714 | 293.9581 | 327.15344 | 172.50511 | 202.85187 | 156.08939 | 8.513245 | 8.595202 | 8.777664 | 7.350386 | 7.5622745 | 7.4638457 |
| ASMM10P037 | 0.00033 | 0.232541 | 3.6982485 | up | NM_019459 | Nphs1 | nephrosis 1, nephrin | 33.074149 | 10.652915 | 5.330769 | 3.443927 | 35.529716 | 27.228264 | 36.464466 | 10.498419 | 11.694398 | 9.765928 | 5.207635 | 5.215173 | 5.5694995 | 3.2560549 | 3.428305 | 3.6474211 |
| ASMM10P021 | 0.00045 | 0.2817691 | 1.6332149 | up | NM_001159 | Dusp15 | dual specificity phosphatase-like 15 | 9857.9057 | 7094.207 | 13.433303 | 12.725589 | 10405.773 | 9195.375 | 9972.569 | 7559.0806 | 7675.936 | 6047.6045 | 13.392041 | 13.380363 | 13.527506 | 12.778343 | 12.767568 | 12.630855 |
| ASMM10P032 | 0.00046 | 0.2817691 | 1.7286159 | up | NM_181410 | Gtf2h3 | general transcription factor IIH, polypeptide 3 | 175.99549 | 124.81076 | 7.747954 | 6.958336 | 198.634 | 171.20844 | 158.14403 | 135.69478 | 124.92262 | 113.81488 | 7.662759 | 7.837457 | 7.743645 | 6.9969563 | 6.8485107 | 7.029542 |
| ASMM10P014 | 0.00050 | 0.281935 | 1.480959 | up | NM_009655 | Alcam | activated leukocyte cell adhesion molecule | 7312.529 | 5809.184 | 13.018643 | 12.452111 | 7627.798 | 7033.697 | 7276.092 | 5825.662 | 5977.4907 | 5624.3994 | 12.949982 | 13.02079 | 13.085156 | 12.414587 | 12.412505 | 12.529241 |
| ASMM10P052 | 0.00052 | 0.281935 | 1.5689434 | up | NM_172301 | Ccnb1 | cyclin B1 | 243.21232 | 188.8168 | 8.204593 | 7.5548 | 287.5295 | 222.29745 | 219.81 | 189.61858 | 194.35193 | 182.4799 | 8.182077 | 8.208882 | 8.222821 | 7.486711 | 7.4981494 | 7.6795397 |
| ASMM10P046 | 0.00052 | 0.281935 | 1.6503849 | up | NM_178884 | Obsl1 | obscurin-like 1 | 1597.5973 | 1159.8977 | 10.866592 | 10.14379 | 1747.6438 | 1509.5314 | 1535.6166 | 1210.0432 | 1154.917 | 1114.7329 | 10.768207 | 10.883433 | 10.948137 | 10.162165 | 10.054091 | 10.215114 |
| ASMM10P006 | 0.00052 | 0.281935 | 1.5465095 | up | NM_009062 | Rgs4 | regulator of G-protein signaling 4 | 2636.2556 | 2012.6374 | 11.560853 | 10.931837 | 2805.2593 | 2557.4543 | 2546.0532 | 2046.5696 | 2122.6826 | 1868.6599 | 11.438886 | 11.611599 | 11.632074 | 10.920152 | 10.930893 | 10.944467 |
| ASMM10P031 | 0.00053 | 0.281935 | 6.2062008 | up | NM_001008 | 1700001J03 | RIKEN cDNA 1700001J03 gene | 624.01035 | 116.74725 | 9.497458 | 6.863748 | 684.6645 | 440.64035 | 746.7262 | 115.83711 | 112.26162 | 122.14301 | 9.4035845 | 9.167673 | 9.921117 | 6.762958 | 6.6980724 | 7.130213 |
| ASMM10P026 | 0.00056 | 0.282713 | 2.2033444 | up | NM_057172 | Fubp1 | far upstream element (FUSE) binding protein 1 | 13248.106 | 7066.2549 | 13.848054 | 12.708359 | 14616.98 | 11900.381 | 13226.957 | 7558.3213 | 8106.8315 | 5533.612 | 13.894822 | 13.741781 | 13.907559 | 12.777508 | 12.837453 | 12.510116 |
| ASMM10P049 | 0.00057 | 0.282713 | 1.393227 | up | NM_011289 | Rpl27 | ribosomal protein L27 | 365.64137 | 318.204 | 8.774893 | 8.296463 | 433.8301 | 316.60507 | 346.48895 | 328.79596 | 341.57004 | 284.246 | 8.763604 | 8.699083 | 8.861992 | 8.281303 | 8.307602 | 8.300483 |
| ASMM10P008 | 0.00059 | 0.2854821 | 1.6195715 | up | NM_145984 | Prepl | prolyl endopeptidase-like | 52.811441 | 39.839138 | 6.015337 | 5.319724 | 61.436607 | 47.81804 | 49.179676 | 46.804565 | 42.210205 | 30.502645 | 6.0026407 | 6.0261354 | 6.017234 | 5.4543095 | 5.28618 | 5.218684 |
| ASMM10P005 | 0.00063 | 0.296778 | 1.5211423 | up | NM_001166 | Rtel1 | regulator of telomere elongation helicase 1 | 166.94481 | 133.8457 | 7.662285 | 7.05713 | 205.6988 | 144.72499 | 150.41064 | 146.94592 | 134.3178 | 120.27337 | 7.714186 | 7.598261 | 7.6744075 | 7.1134286 | 6.9526496 | 7.105311 |
| ASMM10P038 | 0.00065 | 0.296778 | 1.6548735 | up | NM_001040 | Sytl2 | synaptotagmin-like 2 | 37.308386 | 26.794673 | 5.495965 | 4.769244 | 46.292507 | 30.857082 | 34.77557 | 27.350191 | 30.620201 | 22.413628 | 5.5959377 | 5.396691 | 5.4952664 | 4.673538 | 4.8357863 | 4.798408 |
| ASMM10P055 | 0.00067 | 0.296778 | 1.485868 | up | NM_201617 | Gnas | GNAS (guanine nucleotide binding protein, alpha stimulating) complex locus | 82.577307 | 67.83064 | 6.662743 | 6.091437 | 92.953476 | 74.146416 | 80.63203 | 70.6831 | 72.063484 | 60.745335 | 6.5924244 | 6.645508 | 6.750298 | 6.049622 | 6.0579834 | 6.166707 |
| ASMM10P052 | 0.00068 | 0.296778 | 1.8307418 | up | NM_010887 | Ndufs4 | NADH dehydrogenase (ubiquinone) Fe-S protein 4 | 7460.2165 | 4793.6975 | 13.044015 | 12.171586 | 7470.3325 | 7368.331 | 7541.986 | 4792.784 | 4779.2373 | 4809.0713 | 12.916757 | 13.074373 | 13.140914 | 12.130677 | 12.088416 | 12.295666 |
| ASMM10P041 | 0.00071 | 0.296778 | 1.6833249 | up | NM_001080 | Fat3 | FAT tumor suppressor homolog 3 (Drosophila) | 597.92865 | 425.75774 | 9.465768 | 8.714455 | 648.4591 | 561.8595 | 583.46735 | 426.5594 | 454.85886 | 395.85495 | 9.32056 | 9.510037 | 9.566708 | 8.660164 | 8.724685 | 8.758515 |
| ASMM10P007 | 0.00074 | 0.296778 | 1.2897851 | up | NM_016796 | Vamp4 | vesicle-associated membrane protein 4 | 374.06488 | 353.58821 | 8.810827 | 8.443696 | 438.18124 | 348.20804 | 335.80536 | 358.60928 | 393.43924 | 308.7161 | 8.779332 | 8.836699 | 8.81645 | 8.407706 | 8.514879 | 8.408504 |
| ASMM10P055 | 0.00083 | 0.296778 | 1.5147262 | up | NM_020593 | Fbxo3 | F-box protein 3 | 10832.584 | 8303.2423 | 13.558052 | 12.958995 | 10993.565 | 10133.251 | 11370.937 | 8569.859 | 8793.364 | 7546.504 | 13.466198 | 13.521234 | 13.686725 | 12.965391 | 12.958969 | 12.952626 |
| ASMM10P034 | 0.00084 | 0.296778 | 1.3223154 | up | NM_172538 | Vezt | vezatin, adherens junctions transmembrane protein | 11430.159 | 10127.022 | 13.633168 | 13.230101 | 12946.766 | 10335.042 | 11008.67 | 10365.002 | 10733.447 | 9282.617 | 13.705673 | 13.552667 | 13.641163 | 13.224733 | 13.223566 | 13.242005 |
| ASMM10P015 | 0.00086 | 0.296778 | 2.2469979 | up | NM_019413 | Robo1 | roundabout homolog 1 (Drosophila) | 182.01027 | 97.538791 | 7.775602 | 6.607604 | 232.64473 | 159.23082 | 154.15527 | 106.40353 | 90.53136 | 95.68148 | 7.887202 | 7.730493 | 7.709112 | 6.645738 | 6.3873796 | 6.789693 |
| ASMM10P027 | 0.00086 | 0.296778 | 1.2989768 | up | NM_011682 | Utrn | utrophin | 21575.754 | 19359.584 | 14.544888 | 14.167513 | 22509.81 | 20423.543 | 21793.908 | 19159.988 | 20771.113 | 18147.652 | 14.551407 | 14.4908 | 14.592458 | 14.113695 | 14.171173 | 14.21767 |
| ASMM10P009 | 0.00090 | 0.296778 | 5.5989208 | up | NM_207258 | Fam71f1 | family with sequence similarity 71, member F1 | 23.545488 | 5 | 4.824517 | 2.339368 | 19.162556 | 22.356258 | 29.11765 | 4.9999995 | 4.9999995 | 4.9999995 | 4.2901683 | 4.9458127 | 5.2375703 | 2.3250442 | 2.3265402 | 2.3665206 |
| ASMM10P023 | 0.00090 | 0.296778 | 1.793916 | up | NM_001205 | Casc4 | cancer susceptibility candidate 4 | 128.35797 | 88.515035 | 7.302656 | 6.459544 | 136.87439 | 123.97526 | 124.22427 | 90.128845 | 100.62629 | 74.78997 | 7.131203 | 7.384733 | 7.392033 | 6.393628 | 6.535817 | 6.449187 |
| ASMM10P026 | 0.00091 | 0.296778 | 1.456289 | up | NM_172865 | Manea | mannosidase, endo-alpha | 5464.1829 | 4452.468 | 12.60616 | 12.063864 | 5801.1987 | 5320.517 | 5270.833 | 4320.1943 | 4682.192 | 4355.0176 | 12.532209 | 12.637436 | 12.648836 | 11.982994 | 12.056399 | 12.152198 |
| ASMM10P053 | 0.00094 | 0.2969371 | 1.6457668 | up | NM_027300 | Spata3 | spermatogenesis associated 3 | 263.4084 | 194.77801 | 8.313729 | 7.594969 | 309.88824 | 239.01062 | 241.32635 | 183.0182 | 216.99292 | 184.3229 | 8.2855425 | 8.308055 | 8.3475895 | 7.4361906 | 7.656248 | 7.6924686 |
| ASMM10P026 | 0.00100 | 0.2972282 | 1.2881233 | up | NM_001048 | Ccl27a | chemokine (C-C motif) ligand 27A | 6658.789 | 6085.0211 | 12.883993 | 12.518723 | 7348.1113 | 6019.8687 | 6608.387 | 6254.2554 | 6343.622 | 5657.186 | 12.894512 | 12.808033 | 12.949435 | 12.51651 | 12.501795 | 12.537863 |
| ASMM10P015 | 0.00104 | 0.2972282 | 1.6362207 | up | NM_010581 | Cd47 | CD47 antigen (Rh-related antigen, integrin-associated signal transducer) | 626.50062 | 457.48755 | 9.527001 | 8.816634 | 744.4196 | 548.7166 | 586.36566 | 473.36505 | 441.53577 | 457.56183 | 9.530155 | 9.476907 | 9.573942 | 8.812786 | 8.682839 | 8.954277 |
| ASMM10P003 | 0.00105 | 0.2972282 | 1.4648007 | up | NM_172779 | Ddx26b | DEAD/H (Asp-Glu-Ala-Asp/His) box polypeptide 26B | 42.325302 | 34.603424 | 5.680781 | 5.130076 | 52.895176 | 35.68115 | 38.39958 | 35.832363 | 39.30639 | 28.67152 | 5.7880626 | 5.6067452 | 5.647534 | 5.0682073 | 5.186333 | 5.1356883 |
| ASMM10P000 | 0.00107 | 0.2972282 | 4.6359733 | up | NM_001102 | Rhox3c | reproductive homeobox 3C | 20.903267 | 5.582828 | 4.654543 | 2.44167 | 17.81035 | 19.72736 | 25.172092 | 6.7484856 | 4.9999995 | 4.9999995 | 4.190746 | 4.759955 | 5.0129266 | 2.63195 | 2.3265402 | 2.3665206 |
| ASMM10P006 | 0.00110 | 0.2972282 | 1.2704888 | up | NM_001252 | Cldn25 | claudin 25 | 377.32551 | 361.44592 | 8.823906 | 8.478522 | 430.50537 | 359.00064 | 342.47052 | 384.829 | 376.45972 | 323.04904 | 8.751668 | 8.876427 | 8.843622 | 8.511479 | 8.453562 | 8.470525 |
| ASMM10P039 | 0.00111 | 0.2972282 | 1.3430764 | up | NM_010681 | Lama4 | laminin, alpha 4 | 568.5275 | 507.38872 | 9.389911 | 8.96437 | 685.971 | 503.33636 | 516.27515 | 508.4179 | 520.62756 | 493.1207 | 9.407742 | 9.355521 | 9.406471 | 8.917555 | 8.915734 | 9.059821 |
| ASMM10P040 | 0.00111 | 0.2972282 | 1.4780492 | up | NM_010589 | Jak3 | Janus kinase 3 | 402.82642 | 330.24206 | 8.91105 | 8.347356 | 474.55905 | 397.80527 | 336.11493 | 356.44937 | 350.82922 | 283.4476 | 8.891425 | 9.023967 | 8.817758 | 8.399714 | 8.345399 | 8.296954 |
| ASMM10P008 | 0.00113 | 0.2972282 | 5.7525693 | up | NM_133209 | Pilrb1 | paired immunoglobin-like type 2 receptor beta 1 | 62.566644 | 13.563843 | 6.256402 | 3.732195 | 58.982655 | 53.844772 | 74.872505 | 19.530453 | 12.06715 | 9.093926 | 5.9396067 | 6.190355 | 6.6392436 | 4.174787 | 3.4699936 | 3.5518053 |
| ASMM10P027 | 0.00113 | 0.2972282 | 1.2857251 | up | NM_001033 | Zyg11b | zyg-ll family member B, cell cycle regulator | 5677.8647 | 5217.3617 | 12.655239 | 12.292657 | 6146.203 | 5630.8794 | 5256.5117 | 5496.7935 | 5270.0527 | 4885.239 | 12.613534 | 12.707874 | 12.644309 | 12.331523 | 12.226135 | 12.320313 |
| ASMM10P050 | 0.00116 | 0.2972282 | 1.3019491 | up | NM_001033 | 5730507C01 | RIKEN cDNA 5730507C01 gene | 699.38872 | 641.12347 | 9.677907 | 9.297234 | 842.0956 | 664.0266 | 592.04395 | 664.9785 | 678.1163 | 580.2756 | 9.709086 | 9.736639 | 9.587995 | 9.306874 | 9.297114 | 9.287713 |
| ASMM10P042 | 0.00117 | 0.2972282 | 1.4884055 | up | NM_011734 | Siae | sialic acid acetylesterase | 409.47055 | 333.88588 | 8.939445 | 8.365678 | 464.14288 | 392.31158 | 371.9572 | 339.12784 | 338.83563 | 323.69418 | 8.857056 | 9.001258 | 8.960022 | 8.328913 | 8.29516 | 8.47296 |
| ASMM10P052 | 0.00124 | 0.2972282 | 10.888679 | up | NM_134220 | Vmn1r198 | vomeronasal 1 receptor 198 | 54.118401 | 5.121866 | 5.946034 | 2.501277 | 56.189816 | 75.98622 | 30.179167 | 5.3148813 | 4.9999995 | 5.0507174 | 5.869562 | 6.677461 | 5.2910795 | 2.4159396 | 2.3265402 | 2.7613516 |
| ASMM10P015 | 0.00125 | 0.2972282 | 1.7214498 | up | NM_023631 | Aox4 | aldehyde oxidase 4 | 37.370899 | 26.089483 | 5.517206 | 4.733581 | 40.04553 | 32.86429 | 39.202877 | 27.871334 | 27.632767 | 22.764347 | 5.385912 | 5.4852796 | 5.680425 | 4.69989 | 4.682719 | 4.8181353 |
| ASMM10P018 | 0.00125 | 0.2972282 | 1.3768223 | up | NM_053141 | Pcdhb16 | protocadherin beta 16 | 3595.9569 | 3084.8236 | 12.002144 | 11.540801 | 3872.845 | 3314.1008 | 3600.9248 | 3225.4524 | 3166.4355 | 2862.583 | 11.92656 | 11.9725 | 12.107371 | 11.567971 | 11.509961 | 11.544472 |
| ASMM10P042 | 0.00126 | 0.2972282 | 1.5571278 | up | NM_027207 | 1600002K03 | RIKEN cDNA 1600002K03 gene | 528.75996 | 406.13264 | 9.28701 | 8.648122 | 604.67896 | 459.38297 | 522.21796 | 413.6015 | 416.28986 | 388.50656 | 9.217085 | 9.224799 | 9.419145 | 8.616859 | 8.594419 | 8.733089 |
| ASMM10P032 | 0.00128 | 0.2972282 | 1.3471987 | up | NM_001081 | Mn1 | meningioma 1 | 105.91032 | 95.496488 | 7.004091 | 6.574128 | 127.96582 | 100.14622 | 89.618904 | 100.39269 | 102.66308 | 83.4337 | 7.0354877 | 7.0741267 | 6.9026575 | 6.5588975 | 6.566114 | 6.5973725 |
| ASMM10P038 | 0.00131 | 0.2972282 | 1.4569213 | up | NM_027164 | Lrrc27 | leucine rich repeat containing 27 | 101.49157 | 84.253553 | 6.932573 | 6.38965 | 129.24538 | 85.071556 | 90.157776 | 91.37848 | 92.76494 | 68.61724 | 7.0464425 | 6.8385715 | 6.9127064 | 6.4140525 | 6.424042 | 6.330857 |
| ASMM10P041 | 0.00132 | 0.2972282 | 1.4450077 | up | NM_022029 | Nrgn | neurogranin | 2588.3289 | 2113.2814 | 11.5269 | 10.995823 | 3062.0806 | 2472.6414 | 2230.2646 | 2007.6387 | 2351.8591 | 1980.3464 | 11.564549 | 11.56246 | 11.453692 | 10.88863 | 11.073391 | 11.025448 |
| ASMM10P054 | 0.00134 | 0.2972282 | 1.3939644 | up | NM_010431 | Hif1a | hypoxia inducible factor 1, alpha subunit | 427.22247 | 373.24617 | 9.000434 | 8.521241 | 482.64218 | 395.18326 | 403.84198 | 378.47534 | 416.42197 | 324.8412 | 8.913184 | 9.011945 | 9.076174 | 8.490153 | 8.594982 | 8.478587 |
| ASMM10P033 | 0.00136 | 0.2972282 | 1.3353781 | up | NM_001197 | Foxp1 | forkhead box P1 | 2056.3696 | 1831.8685 | 11.21159 | 10.794342 | 2275.0715 | 1888.8085 | 2005.2288 | 1854.1124 | 1991.8025 | 1649.6906 | 11.142753 | 11.187262 | 11.304756 | 10.777337 | 10.835226 | 10.770463 |
| ASMM10P032 | 0.00140 | 0.2972282 | 2.1538658 | up | NM_001134 | Ccdc62 | coiled-coil domain containing 62 | 41.80118 | 23.489097 | 5.680873 | 4.573944 | 42.73645 | 35.9353 | 46.73179 | 25.008104 | 26.74742 | 18.711767 | 5.4818206 | 5.6176014 | 5.943196 | 4.533748 | 4.635472 | 4.552613 |
| ASMM10P029 | 0.00143 | 0.2972282 | 2.2537618 | up | NM_001145 | Ttc39a | tetratricopeptide repeat domain 39A | 56.275617 | 30.059929 | 6.093047 | 4.920712 | 66.327805 | 42.603413 | 59.895634 | 35.361095 | 32.41912 | 22.399572 | 6.112055 | 5.8581643 | 6.3089232 | 5.0490084 | 4.91624 | 4.796889 |
| ASMM10P027 | 0.00143 | 0.2972282 | 4.5085015 | up | NM_007443 | Ambp | alpha 1 microglobulin/bikunin | 21.208305 | 5.535125 | 4.622848 | 2.4502 | 29.992342 | 20.047863 | 13.584711 | 4.9999995 | 6.605377 | 4.9999995 | 4.9629993 | 4.784816 | 4.1207285 | 2.3250442 | 2.659035 | 2.3665206 |

| | | | | | | | | | | | | | | | | | | | | | | | |
|---|---|---|---|---|---|---|---|---|---|---|---|---|---|---|---|---|---|---|---|---|---|---|---|
| ASMM10P012 | 0.00151 | 0.2972282 | 1.7583265 | up | NM_183028 | Pcmtd1 | protein-L-isoaspartate (D-aspartate) O-methyltransferase domain containing 1 | 922.51615 | 627.09318 | 10.079304 | 9.265101 | 999.713 | 825.46106 | 942.3744 | 629.52234 | 639.252 | 612.5052 | 9.939534 | 10.0409 | 10.257479 | 9.223994 | 9.210445 | 9.360865 |
| ASMM10P042 | 0.00152 | 0.2972282 | 5.8607286 | up | NM_001103 | Vmn2r80 | vomeronasal 2, receptor 80 | 28.145749 | 5.995386 | 5.061223 | 2.510143 | 27.05799 | 36.42551 | 20.953747 | 7.9861593 | 4.9999995 | 4.9999995 | 4.8074117 | 5.634746 | 4.741511 | 2.8373678 | 2.3265402 | 2.3665206 |
| ASMM10P006 | 0.00155 | 0.2972282 | 1.5642975 | up | NM_001252 | 4930453H2 | RIKEN cDNA 4930453H23 gene | 33.638162 | 25.874476 | 5.350284 | 4.704769 | 40.750378 | 27.989914 | 32.174194 | 30.124744 | 28.461613 | 19.037071 | 5.411871 | 5.2568455 | 5.3821344 | 4.8111186 | 4.7287655 | 4.574422 |
| ASMM10P036 | 0.00156 | 0.2972282 | 1.7585345 | up | NM_170669 | Rps15a | ribosomal protein S15A | 19720.233 | 13118.84 | 14.414187 | 13.599813 | 19476.023 | 18741.871 | 20942.805 | 12959.018 | 12997.35 | 13400.153 | 14.332533 | 14.373349 | 14.536678 | 13.539103 | 13.490668 | 13.769668 |
| ASMM10P006 | 0.00156 | 0.2972282 | 2.3371275 | up | NM_001205 | Clec9a | C-type lectin domain family 9, member a | 47.92285 | 24.363963 | 5.854965 | 4.630229 | 57.310642 | 49.03065 | 37.427258 | 28.361818 | 23.615828 | 21.114243 | 5.8978863 | 6.058152 | 5.6088576 | 4.7285085 | 4.448114 | 4.714064 |
| ASMM10P043 | 0.00157 | 0.2972282 | 2.4461547 | up | NM_007738 | Col7a1 | collagen, type VII, alpha 1 | 260.77646 | 126.67257 | 8.271645 | 6.981129 | 342.43613 | 189.75194 | 250.1413 | 121.8493 | 133.78001 | 124.38841 | 8.429735 | 7.9903255 | 8.394874 | 6.8410726 | 6.9464846 | 7.1558304 |
| ASMM10P012 | 0.00158 | 0.2972282 | 1.6344637 | up | ENSMUST0 | Adra1a | adrenergic receptor, alpha 1a | 4150.4559 | 2994.8282 | 12.205559 | 11.496742 | 4874.7417 | 3441.1553 | 4135.4707 | 3062.7952 | 3109.962 | 2811.7275 | 12.278399 | 12.022576 | 12.315703 | 11.487201 | 11.484627 | 11.518398 |
| ASMM10P036 | 0.00158 | 0.2972282 | 3.348182 | up | NM_027053 | Lrrc51 | leucine rich repeat containing 51 | 82.982965 | 29.867384 | 6.668585 | 4.925207 | 78.60207 | 94.053764 | 76.29306 | 33.804848 | 27.203753 | 28.59355 | 6.3551717 | 6.9825706 | 6.668014 | 4.983248 | 4.6603775 | 5.131997 |
| ASMM10P034 | 0.00159 | 0.2972282 | 1.3256005 | up | NM_133737 | Lancl2 | LanC (bacterial lantibiotic synthetase component C)-like 2 | 6753.8659 | 6001.8579 | 12.905563 | 12.498917 | 7402.6523 | 6468.314 | 6390.6313 | 6051.9834 | 6025.5854 | 5928.005 | 12.906913 | 12.909271 | 12.900506 | 12.468753 | 12.425522 | 12.602477 |
| ASMM10P031 | 0.00169 | 0.2972282 | 1.2857897 | up | NM_029659 | Styxl1 | serine/threonine/tyrosine interacting-like 1 | 256.24302 | 241.71129 | 8.265703 | 7.903048 | 323.37894 | 228.9219 | 216.42822 | 258.49838 | 258.15463 | 208.48085 | 8.349663 | 8.246717 | 8.200728 | 7.937464 | 7.903117 | 7.868563 |
| ASMM10P019 | 0.00169 | 0.2972282 | 1.3756009 | up | NM_001177 | Gm7102 | predicted gene 7102 | 3806.5259 | 3274.5039 | 12.086285 | 11.626223 | 4037.4587 | 3671.5671 | 3710.5518 | 3263.3872 | 3379.449 | 3180.6755 | 11.985905 | 12.113518 | 12.159433 | 11.585722 | 11.602215 | 11.690733 |
| ASMM10P025 | 0.00169 | 0.2972282 | 1.6755167 | up | NM_001040 | Rap1gds1 | RAP1, GTP-GDP dissociation stimulator 1 | 180.7355 | 132.37107 | 7.786732 | 7.042126 | 203.7894 | 169.41872 | 168.99837 | 125.71079 | 143.53459 | 127.86782 | 7.698898 | 7.8213143 | 7.8399844 | 6.885124 | 7.048587 | 7.1926675 |
| ASMM10P006 | 0.00169 | 0.2972282 | 1.5060138 | up | NM_009830 | Ccne2 | cyclin E2 | 448.43376 | 358.53303 | 9.058836 | 8.468101 | 531.03735 | 393.65204 | 420.61188 | 365.43225 | 356.13943 | 354.0274 | 9.039504 | 9.00697 | 9.130035 | 8.434494 | 8.367817 | 8.601993 |
| ASMM10P013 | 0.00173 | 0.2972282 | 1.6646898 | up | NM_001160 | A830018L1 | RIKEN cDNA A830018L16 gene | 24142.696 | 16999.131 | 14.706773 | 13.971519 | 23820.531 | 22494.697 | 26112.861 | 16660.303 | 17902.867 | 16434.223 | 14.629401 | 14.617664 | 14.873253 | 13.899465 | 13.940889 | 14.074204 |
| ASMM10P008 | 0.00173 | 0.2972282 | 1.5957202 | up | NM_027482 | 5730508B0 | RIKEN cDNA 5730508B09 gene | 542.92486 | 408.62426 | 9.330719 | 8.656511 | 598.0013 | 508.98218 | 521.7911 | 398.83374 | 426.57715 | 400.46188 | 9.200845 | 9.373373 | 9.417938 | 8.564761 | 8.63125 | 8.773522 |
| ASMM10P041 | 0.00178 | 0.2972282 | 1.5243466 | up | NM_016812 | Banp | BTG3 associated nuclear protein | 4919.666 | 3800.4235 | 12.452931 | 11.84474 | 5072.7715 | 5094.783 | 4591.4434 | 3711.6362 | 4000.5361 | 3689.0981 | 12.331909 | 12.573972 | 12.452913 | 11.772268 | 11.839799 | 11.922154 |
| ASMM10P004 | 0.00180 | 0.2972282 | 1.4131601 | up | NM_001112 | Atp5c1 | ATP synthase, H+ transporting, mitochondrial F1 complex, gamma polypeptide 1 | 297.2915 | 255.69604 | 8.487456 | 7.988531 | 339.24915 | 262.9429 | 289.68246 | 263.53384 | 266.18982 | 237.36447 | 8.415321 | 8.442743 | 8.604304 | 7.9632683 | 7.948205 | 8.05412 |
| ASMM10P048 | 0.00182 | 0.2972282 | 2.1388409 | up | NM_021294 | Dbil5 | diazepam binding inhibitor-like 5 | 989.97924 | 557.65346 | 10.186215 | 9.089386 | 1027.2196 | 967.76843 | 974.9497 | 493.31827 | 639.5407 | 540.1014 | 9.981733 | 10.266508 | 10.310405 | 8.8741865 | 9.212294 | 9.181678 |
| ASMM10P017 | 0.00183 | 0.2972282 | 1.7138436 | up | NM_173423 | Fem1c | fem-1 homolog c (C.elegans) | 429.47762 | 302.95358 | 9.004906 | 8.22767 | 479.4822 | 382.85092 | 426.09973 | 285.98444 | 327.45117 | 295.42514 | 8.905405 | 8.964369 | 9.144943 | 8.084812 | 8.246924 | 8.3512745 |
| ASMM10P034 | 0.00185 | 0.2972282 | 3.1746248 | up | NM_012013 | Figla | folliculogenesis specific basic helix-loop-helix | 55.778556 | 20.752081 | 6.073328 | 4.406742 | 59.199406 | 40.443962 | 67.6923 | 21.99127 | 21.409496 | 18.855478 | 5.9457 | 5.783677 | 6.4906073 | 4.342863 | 4.315336 | 4.562027 |
| ASMM10P004 | 0.00187 | 0.2973645 | 1.6805204 | up | NM_001024 | Zbtb45 | zinc finger and BTB domain containing 45 | 37.290345 | 26.369989 | 5.498802 | 4.749894 | 45.544018 | 33.247936 | 33.079082 | 27.923044 | 26.414724 | 24.7722 | 5.5718403 | 5.502612 | 5.4219527 | 4.703028 | 4.6175804 | 4.9290724 |
| ASMM10P023 | 0.00191 | 0.3002908 | 1.3097777 | up | NM_146130 | Hnrnpa3 | heterogeneous nuclear ribonucleoprotein A3 | 3738.4302 | 3375.4169 | 12.058253 | 11.668931 | 4087.5798 | 3554.5427 | 3573.168 | 3529.238 | 3326.086 | 3270.9268 | 12.005629 | 12.069665 | 12.099464 | 11.69367 | 11.579931 | 11.733191 |
| ASMM10P005 | 0.00202 | 0.3038459 | 1.4977459 | up | NM_001166 | Rtel1 | regulator of telomere elongation helicase 1 | 253.76495 | 203.68878 | 8.246329 | 7.663536 | 329.56537 | 208.24196 | 223.48752 | 211.31485 | 211.22939 | 188.52211 | 8.377485 | 8.117444 | 8.244058 | 7.6454926 | 7.6199074 | 7.7252083 |
| ASMM10P028 | 0.00203 | 0.3038459 | 1.3207695 | up | NM_001199 | Ccl27b | chemokine (C-C motif) ligand 27b | 5836.7227 | 5218.2104 | 12.695941 | 12.294563 | 6552.5215 | 5245.692 | 5711.9546 | 5332.732 | 5282.497 | 5039.4023 | 12.711228 | 12.617348 | 12.759248 | 12.282685 | 12.236103 | 12.3649 |
| ASMM10P045 | 0.00204 | 0.3038459 | 1.9480745 | up | NM_025379 | Cox7b | cytochrome c oxidase subunit VIIb | 1611.2895 | 986.67835 | 10.877624 | 9.915575 | 1681.7136 | 1493.3684 | 1658.7864 | 945.7043 | 993.35876 | 1020.972 | 10.708385 | 10.868162 | 11.056324 | 9.8071995 | 9.845722 | 10.093803 |
| ASMM10P000 | 0.00205 | 0.3038459 | 1.6319059 | up | NM_145150 | Prc1 | protein regulator of cytokinesis 1 | 28.354282 | 20.901443 | 5.112266 | 4.405708 | 31.91107 | 26.032438 | 27.119337 | 25.442322 | 20.253239 | 17.008768 | 5.053479 | 5.1551266 | 5.1281924 | 4.560242 | 4.2345185 | 4.422364 |
| ASMM10P044 | 0.00206 | 0.3038459 | 1.5097176 | up | NM_001085 | Gm14501 | predicted gene 14501 | 39.313783 | 30.937875 | 5.57447 | 4.980192 | 48.71611 | 33.267414 | 35.957825 | 32.67375 | 31.93902 | 28.200855 | 5.670803 | 5.5040464 | 5.548562 | 4.932836 | 4.8953614 | 5.112378 |
| ASMM10P045 | 0.00207 | 0.3038459 | 1.3806288 | up | NM_022322 | Tnmd | tenomodulin | 71.864907 | 63.592725 | 6.457949 | 5.992623 | 82.81644 | 64.29239 | 68.48589 | 63.22682 | 74.08199 | 53.469364 | 6.4258747 | 6.440973 | 6.5069985 | 5.8878736 | 6.09754 | 5.992456 |
| ASMM10P034 | 0.00211 | 0.3038459 | 2.279389 | up | NM_001191 | Dync1i1 | dynein cytoplasmic 1 intermediate chain 1 | 68.73819 | 36.939423 | 6.403436 | 5.214789 | 71.48314 | 65.760345 | 68.971085 | 46.315998 | 33.34251 | 31.15976 | 6.222201 | 6.4712996 | 6.5168076 | 5.4407845 | 4.956131 | 5.2474513 |
| ASMM10P031 | 0.00211 | 0.3038459 | 2.0116501 | up | NM_019447 | Hgfac | hepatocyte growth factor activator | 77.498585 | 46.952372 | 6.565002 | 5.556622 | 89.566574 | 73.89613 | 69.03305 | 56.92057 | 42.21767 | 41.718876 | 6.5378942 | 6.639692 | 6.5174184 | 5.73602 | 5.2865005 | 5.647346 |
| ASMM10P033 | 0.00212 | 0.3038459 | 1.3986206 | up | NM_053202 | Foxp1 | forkhead box P1 | 6997.994 | 5880.1741 | 12.953277 | 12.469273 | 7257.5977 | 6532.0444 | 7204.34 | 5822.182 | 6288.897 | 5529.4434 | 12.864183 | 12.923206 | 13.072443 | 12.411833 | 12.487612 | 12.508373 |
| ASMM10P000 | 0.00213 | 0.3038459 | 1.3104808 | up | NM_001198 | Dync1i2 | dynein cytoplasmic 1 intermediate chain 2 | 526.39188 | 482.14408 | 9.279248 | 8.889152 | 639.4435 | 497.56665 | 442.1655 | 515.6281 | 487.32697 | 443.47717 | 9.299101 | 9.341376 | 9.1972685 | 8.936032 | 8.820643 | 8.910782 |
| ASMM10P055 | 0.00221 | 0.304465 | 3.5959545 | up | NM_008806 | Pde6b | phosphodiesterase 6B, cGMP, rod receptor, beta polypeptide | 52.465211 | 17.343403 | 5.995897 | 4.149522 | 55.32124 | 59.85198 | 42.222412 | 16.368917 | 17.020163 | 18.64113 | 5.848234 | 6.3407383 | 5.7987185 | 3.9147985 | 3.986467 | 4.547301 |
| ASMM10P033 | 0.00222 | 0.304465 | 3.2284883 | up | NM_001168 | Jazf1 | JAZF zinc finger 1 | 37.414559 | 13.886816 | 5.519231 | 3.828372 | 35.441788 | 38.67133 | 38.130558 | 14.26042 | 13.098379 | 14.30165 | 5.2046094 | 5.717422 | 5.635661 | 3.7086325 | 3.5891461 | 4.1873374 |
| ASMM10P042 | 0.00223 | 0.304465 | 1.5071162 | up | NM_133984 | Hemk1 | HemK methyltransferase family member 1 | 3758.4321 | 2948.5197 | 12.068983 | 11.477192 | 4109.832 | 3319.003 | 3846.4612 | 2926.5498 | 3036.6863 | 2882.323 | 12.015678 | 11.974247 | 12.217024 | 11.425459 | 11.450476 | 11.555642 |
| ASMM10P001 | 0.00225 | 0.304465 | 1.3288823 | up | NM_010767 | Masp2 | mannan-binding lectin serine peptidase 2 | 116.82861 | 107.00536 | 7.148681 | 6.738468 | 141.40999 | 107.015 | 102.06083 | 105.86388 | 117.47681 | 97.6754 | 7.176751 | 7.1700974 | 7.099195 | 6.6372814 | 6.758887 | 6.819235 |
| ASMM10P048 | 0.00226 | 0.304465 | 1.6167908 | up | NM_019933 | Ptpn4 | protein tyrosine phosphatase, non-receptor type 4 | 267.1008 | 202.19802 | 8.336636 | 7.643503 | 301.54587 | 238.54364 | 261.2129 | 195.10051 | 237.86891 | 173.62465 | 8.246013 | 8.304701 | 8.459195 | 7.527315 | 7.7889977 | 7.6141973 |
| ASMM10P022 | 0.00233 | 0.3095849 | 1.5598826 | up | NM_001001 | Thnsl1 | threonine synthase-like 1 (bacterial) | 107.4892 | 83.259925 | 7.015364 | 6.373927 | 136.2204 | 85.379776 | 100.86741 | 92.875565 | 88.26817 | 68.63604 | 7.122889 | 6.843059 | 7.0801454 | 6.4395447 | 6.3507853 | 6.331451 |
| ASMM10P005 | 0.00233 | 0.3095849 | 1.3237228 | up | NM_001177 | Cpeb2 | cytoplasmic polyadenylation element binding protein 2 | 268.02487 | 247.4706 | 8.335361 | 7.93076 | 323.10822 | 250.39996 | 230.56642 | 258.7283 | 279.66092 | 204.02258 | 8.348367 | 8.372307 | 8.285408 | 7.938943 | 8.017774 | 7.835562 |
| ASMM10P000 | 0.00234 | 0.3095849 | 1.7145828 | up | NM_026856 | Zfp644 | zinc finger protein 644 | 274.9202 | 195.00962 | 8.370217 | 7.592359 | 333.74155 | 249.69334 | 241.32571 | 174.51698 | 225.12944 | 185.38245 | 8.39432 | 8.368926 | 8.347405 | 7.3678927 | 7.7084475 | 7.700738 |
| ASMM10P040 | 0.00237 | 0.3115739 | 1.5696649 | up | NM_007975 | F2rl3 | coagulation factor II (thrombin) receptor-like 3 | 65.975473 | 50.591546 | 6.327392 | 5.676936 | 80.77561 | 55.58423 | 61.56658 | 52.285923 | 51.61006 | 47.878654 | 6.393409 | 6.2365627 | 6.3522058 | 5.6088576 | 5.580931 | 5.841019 |
| ASMM10P054 | 0.00246 | 0.3149252 | 1.3977815 | up | NM_001039 | Triobp | TRIO and F-actin binding protein | 69.355158 | 60.361122 | 6.407821 | 5.924682 | 80.36166 | 61.499935 | 66.20388 | 63.792633 | 61.402546 | 55.888187 | 6.3857512 | 6.3788457 | 6.458867 | 5.901291 | 5.8231344 | 6.049622 |
| ASMM10P021 | 0.00250 | 0.3154602 | 1.5002694 | up | NM_178762 | Pced1a | PC-esterase domain containing 1A | 201.07495 | 165.02806 | 7.939522 | 7.354301 | 223.75397 | 204.46637 | 175.0045 | 172.99129 | 184.55261 | 137.54027 | 7.832907 | 8.092313 | 7.893347 | 7.3542743 | 7.418119 | 7.290509 |
| ASMM10P000 | 0.00253 | 0.3156142 | 1.3781464 | up | NM_009543 | Rnf103 | ring finger protein 103 | 291.49772 | 257.67472 | 8.460648 | 7.997918 | 334.88406 | 278.92078 | 260.68832 | 251.71426 | 277.08566 | 244.22424 | 8.398663 | 8.526375 | 8.456905 | 7.8956714 | 8.001811 | 8.096273 |
| ASMM10P049 | 0.00255 | 0.3156142 | 5.0986831 | up | NM_148953 | Asb16 | ankyrin repeat and SOCS box-containing 16 | 28.440107 | 6.206835 | 5.063266 | 2.713142 | 39.983753 | 20.09873 | 25.237839 | 5.951515 | 4.9999995 | 7.66899 | 5.383355 | 4.789049 | 5.0173945 | 2.5009575 | 2.3265402 | 3.3119268 |
| ASMM10P035 | 0.00255 | 0.3156142 | 1.7499083 | up | NM_023440 | Tmem86b | transmembrane protein 86B | 2966.203 | 2008.5951 | 11.72732 | 10.92004 | 3035.1147 | 2843.3018 | 3020.1926 | 2220.645 | 2138.3179 | 1666.8224 | 11.54995 | 11.755353 | 11.876656 | 11.035245 | 10.940352 | 10.784524 |
| ASMM10P008 | 0.00256 | 0.3156142 | 1.9053509 | up | NM_145839 | Rasgef1b | RasGEF domain family, member 1B | 70.945087 | 44.854047 | 6.412684 | 5.482627 | 92.35741 | 64.38178 | 56.09607 | 53.70319 | 47.248455 | 33.610497 | 6.5825396 | 6.442067 | 6.2134457 | 5.647346 | 5.4499855 | 5.3505507 |
| ASMM10P009 | 0.00258 | 0.3162314 | 1.3401848 | up | NM_177640 | D030056L2 | RIKEN cDNA D030056L22 gene | 477.99029 | 429.28792 | 9.150221 | 8.727789 | 555.01636 | 466.479 | 412.47552 | 434.4856 | 441.8692 | 411.50897 | 9.098622 | 9.246786 | 9.105256 | 8.690863 | 8.683893 | 8.808612 |
| ASMM10P004 | 0.00261 | 0.3168118 | 1.4482683 | up | NM_001033 | A230046K0 | RIKEN cDNA A230046K03 gene | 79.103497 | 66.936681 | 6.596403 | 6.062074 | 91.17882 | 69.208 | 76.92367 | 78.102135 | 67.26802 | 55.439888 | 6.565931 | 6.5440345 | 6.6792426 | 6.190355 | 5.9583664 | 6.0375 |
| ASMM10P018 | 0.00262 | 0.3168118 | 1.7137779 | up | NM_001122 | Spata3 | spermatogenesis associated 3 | 468.28163 | 332.21826 | 9.132947 | 8.355766 | 492.03836 | 453.29022 | 459.5163 | 317.77063 | 363.81302 | 315.07114 | 8.937922 | 9.208595 | 9.252323 | 8.232891 | 8.3984995 | 8.435909 |
| ASMM10P013 | 0.00263 | 0.3168118 | 1.5950338 | up | NM_031163 | Col2a1 | collagen, type II, alpha 1 | 923.50192 | 697.01597 | 10.087027 | 9.41344 | 971.5078 | 888.62396 | 910.374 | 703.4939 | 774.78357 | 612.77045 | 9.905212 | 10.146767 | 10.209102 | 9.3864 | 9.492443 | 9.361477 |
| ASMM10P035 | 0.00265 | 0.3170857 | 1.9985077 | up | NM_001033 | Zfp575 | zinc finger protein 575 | 413.43212 | 251.80439 | 8.958217 | 7.959294 | 421.69373 | 438.41592 | 380.1867 | 291.89703 | 249.38744 | 214.1287 | 8.724685 | 9.160299 | 8.989666 | 8.11757 | 7.853793 | 7.9065175 |
| ASMM10P004 | 0.00267 | 0.3170857 | 2.0112947 | up | NM_001039 | Gbp10 | guanylate-binding protein 10 | 137.38131 | 84.976811 | 7.40408 | 6.395956 | 139.948 | 133.41537 | 138.78055 | 82.91413 | 103.09959 | 68.91671 | 7.1628494 | 7.490644 | 7.558747 | 6.277602 | 6.572613 | 6.337652 |
| ASMM10P036 | 0.00269 | 0.3170857 | 1.6574267 | up | NM_001162 | Lrrc51 | leucine rich repeat containing 51 | 121.00502 | 89.565352 | 7.203197 | 6.474252 | 142.1923 | 114.56834 | 106.25442 | 82.679825 | 107.08021 | 78.93602 | 7.183768 | 7.267495 | 7.1583276 | 6.2734523 | 6.6273212 | 6.521982 |
| ASMM10P053 | 0.00273 | 0.3172304 | 1.448743 | up | NM_001177 | Zfp964 | zinc finger protein 964 | 28.199379 | 23.317748 | 5.105179 | 4.570377 | 31.019148 | 26.356295 | 27.222694 | 25.492592 | 23.816141 | 20.64451 | 5.01036 | 5.170969 | 5.1342072 | 4.5629125 | 4.461943 | 4.6862755 |
| ASMM10P030 | 0.00275 | 0.3172304 | 2.323853 | up | NM_033541 | Oas1c | 2'-5' oligoadenylate synthetase 1C | 1844.5208 | 934.96699 | 11.052644 | 9.836125 | 1864.1592 | 1521.0132 | 2148.3901 | 886.2258 | 1036.109 | 882.56616 | 10.862854 | 10.896404 | 11.398675 | 9.7083435 | 9.908559 | 9.891474 |
| ASMM10P003 | 0.00277 | 0.3172304 | 1.3224441 | up | NM_010461 | Hoxb8 | homeobox B8 | 135.47774 | 125.85918 | 7.374009 | 6.970802 | 155.86276 | 125.93215 | 124.6383 | 127.2781 | 132.14261 | 118.15682 | 7.3175006 | 7.407629 | 7.3968964 | 6.9048414 | 6.9289894 | 7.078575 |
| ASMM10P000 | 0.00281 | 0.3172304 | 2.2698514 | up | NM_001198 | Postn | periostin, osteoblast specific factor | 32.303033 | 16.827427 | 5.290391 | 4.107793 | 39.073547 | 29.858873 | 27.976679 | 16.788631 | 16.389088 | 17.304562 | 5.350442 | 5.348772 | 5.171958 | 3.9491515 | 3.9256544 | 4.4485726 |
| ASMM10P020 | 0.00283 | 0.3172304 | 2.0431994 | up | NM_080854 | Slc34a3 | solute carrier family 34 (sodium phosphate), member 3 | 271.9752 | 163.9239 | 8.363312 | 7.332482 | 294.67212 | 246.31094 | 274.94254 | 155.06212 | 206.73775 | 129.97183 | 8.21204 | 8.350178 | 8.527717 | 7.192529 | 7.58934 | 7.215576 |
| ASMM10P019 | 0.00284 | 0.3172304 | 1.3965253 | up | NM_001163 | Tnks2 | tankyrase, TRF1-interacting ankyrin-related ADP-ribose polymerase 2 | 184.90634 | 162.44213 | 7.819307 | 7.337465 | 210.91559 | 166.38809 | 177.41533 | 166.5334 | 166.52686 | 154.26613 | 7.7503467 | 7.796151 | 7.9114227 | 7.2991095 | 7.26525 | 7.4480357 |
| ASMM10P055 | 0.00288 | 0.3172304 | 2.0748901 | up | NM_001033 | Gm5535 | predicted gene 5535 | 63.141951 | 35.947499 | 6.243302 | 5.190267 | 83.56601 | 50.884544 | 54.9753 | 37.281155 | 34.68077 | 35.880573 | 6.438807 | 6.1075163 | 6.1835823 | 5.1228175 | 5.0118513 | 5.436132 |
| ASMM10P036 | 0.00288 | 0.3172304 | 1.3195196 | up | NM_028230 | Shmt2 | serine hydroxymethyltransferase 2 (mitochondrial) | 1366.2797 | 1236.0211 | 10.634919 | 10.234906 | 1556.004 | 1259.7599 | 1283.0751 | 1248.463 | 1241.8297 | 1217.7706 | 10.585989 | 10.631703 | 10.687066 | 10.205524 | 10.159423 | 10.339773 |
| ASMM10P029 | 0.00293 | 0.3172304 | 3.7551115 | up | NM_025786 | Rnf186 | ring finger protein 186 | 18.027819 | 6.068094 | 4.444362 | 2.535506 | 16.701937 | 21.108213 | 16.273306 | 4.9999995 | 8.204284 | 4.9999995 | 4.090512 | 4.86257 | 4.380003 | 2.3250442 | 2.914953 | 2.3665206 |
| ASMM10P010 | 0.00295 | 0.3172304 | 6.8853629 | up | NM_138312 | Fam172a | family with sequence similarity 172, member A | 37.540327 | 5.685595 | 5.404314 | 2.620781 | 54.015015 | 38.389465 | 20.216501 | 5.5309806 | 4.9999995 | 6.5258036 | 5.8160043 | 5.706497 | 4.69044 | 2.442414 | 2.3265402 | 3.093389 |
| ASMM10P029 | 0.00298 | 0.3172304 | 2.9160297 | up | NM_001008 | Asap3 | ArfGAP with SH3 domain, ankyrin repeat and PH domain 3 | 43.548127 | 17.415213 | 5.687778 | 4.143772 | 58.56566 | 40.9039 | 31.17482 | 15.447089 | 21.087831 | 15.71072 | 5.928541 | 5.7987185 | 5.336073 | 3.8272169 | 4.2937646 | 4.3103347 |
| ASMM10P012 | 0.00300 | 0.3172304 | 1.3693559 | up | NM_025622 | Lgals2 | lectin, galactose-binding, soluble 2 | 165.19777 | 148.03803 | 7.656788 | 7.203291 | 186.8865 | 152.0954 | 156.61142 | 151.5551 | 152.85814 | 139.70084 | 7.576403 | 7.6636467 | 7.7303143 | 7.1570463 | 7.1408124 | 7.312013 |
| ASMM10P023 | 0.00300 | 0.3172304 | 1.7982214 | up | NM_172666 | Agps | alkylglycerone phosphate synthase | 6743.242 | 4392.636 | 12.893536 | 12.046966 | 6470.2124 | 6370.643 | 7388.8706 | 4315.245 | 4523.031 | 4339.632 | 12.689649 | 12.88235 | 13.10861 | 11.980622 | 12.012109 | 12.148166 |
| ASMM10P011 | 0.00303 | 0.3172304 | 2.3498275 | up | NM_207558 | Olfr750 | olfactory receptor 750 | 51.616244 | 26.959655 | 5.986769 | 4.754214 | 55.48014 | 45.30623 | 54.062363 | 35.25667 | 23.751202 | 21.871094 | 5.8521056 | 5.9494057 | 6.158796 | 5.0445027 | 4.457869 | 4.760271 |
| ASMM10P004 | 0.00303 | 0.3172304 | 1.4023151 | up | NM_001042 | Rabl3 | RAB, member of RAS oncogene family-like 3 | 98.598972 | 86.821195 | 6.917753 | 6.429943 | 109.16797 | 89.28148 | 97.347466 | 92.95455 | 98.125755 | 69.38328 | 6.817916 | 6.907095 | 7.0282483 | 6.4401345 | 6.503048 | 6.346645 |
| ASMM10P013 | 0.00304 | 0.3172304 | 1.6842024 | up | NM_001130 | Oxr1 | oxidation resistance 1 | 1426.9975 | 1011.3883 | 10.702842 | 9.950777 | 1567.5441 | 1364.3217 | 1349.1266 | 1114.5509 | 920.35425 | 999.25977 | 10.599345 | 10.751214 | 10.757968 | 10.048232 | 9.7404375 | 10.063661 |
| ASMM10P016 | 0.00306 | 0.3172304 | 1.2718112 | up | NM_133771 | Memo1 | mediator of cell motility 1 | 604.26438 | 570.52706 | 9.477843 | 9.130958 | 699.07275 | 567.533 | 546.1874 | 585.64166 | 572.03925 | 553.90027 | 9.438674 | 9.519102 | 9.475752 | 9.119357 | 9.0525465 | 9.220971 |
| ASMM10P011 | 0.00308 | 0.3172304 | 2.6777236 | up | NM_010024 | Dct | dopachrome tautomerase | 242.9495 | 110.80156 | 8.197244 | 6.776237 | 246.46437 | 199.82027 | 282.56387 | 98.72394 | 133.8291 | 99.85163 | 7.9700174 | 8.059979 | 8.561737 | 6.533129 | 6.9471807 | 6.8484025 |
| ASMM10P048 | 0.00308 | 0.3172304 | 1.8384581 | up | NM_176982 | Fbxo48 | F-box protein 48 | 21.063538 | 13.952325 | 4.676964 | 3.798468 | 23.783089 | 19.8223 | 19.585224 | 14.96848 | 17.56147 | 9.327024 | 4.6150217 | 4.767899 | 4.647972 | 3.7812326 | 4.0307837 | 3.5833874 |
| ASMM10P044 | 0.00310 | 0.3172304 | 1.5097716 | up | NM_018818 | Chm | choroidermia | 340.11328 | 274.64744 | 8.682642 | 8.088312 | 371.38205 | 330.3355 | 318.62228 | 273.31293 | 283.23804 | 267.39136 | 8.546617 | 8.75964 | 8.74167 | 8.01641 | 8.033621 | 8.214905 |
| ASMM10P000 | 0.00311 | 0.3172304 | 2.1025641 | up | NM_198934 | Pou2f1 | POU domain, class 2, transcription factor 1 | 85.754588 | 49.78968 | 6.72116 | 5.64901 | 80.93946 | 89.141655 | 87.18265 | 55.479298 | 51.072212 | 42.81753 | 6.396286 | 6.9044747 | 6.8627186 | 5.694889 | 5.565352 | 5.686789 |
| ASMM10P055 | 0.00314 | 0.3172304 | 4.2466177 | up | NM_001005 | Inadl | InaD-like (Drosophila) | 40.79127 | 11.601165 | 5.626902 | 3.540588 | 49.885864 | 31.178057 | 41.30989 | 13.526272 | 8.513834 | 12.763389 | 5.7063694 | 5.411067 | 5.763271 | 3.6312304 | 2.9651318 | 4.0254025 |
| ASMM10P011 | 0.00319 | 0.3172304 | 2.3563715 | up | NM_053105 | Klhl1 | kelch-like 1 | 1045.319 | 532.71575 | 10.258979 | 9.022412 | 1109.143 | 896.08765 | 1130.7262 | 449.77444 | 596.2101 | 552.1627 | 10.097075 | 10.158331 | 10.521531 | 8.741406 | 9.111055 | 9.214775 |
| ASMM10P034 | 0.00320 | 0.3172304 | 1.3389208 | up | NM_178873 | Adck2 | aarF domain containing kinase 2 | 150.48276 | 138.11905 | 7.525626 | 7.104555 | 169.38255 | 143.1241 | 138.94164 | 146.94737 | 140.68819 | 126.72158 | 7.4322534 | 7.5838346 | 7.5607896 | 7.1135406 | 7.018996 | 7.181129 |
| ASMM10P049 | 0.00323 | 0.3172304 | 1.8083325 | up | NM_172802 | Fscn2 | fascin homolog 2, actin-bundling protein, retinal (Strongylocentrotus purpuratus) | 147.52206 | 100.32113 | 7.503243 | 6.648584 | 156.91069 | 138.88756 | 146.76793 | 99.22832 | 102.03088 | 99.704185 | 7.326432 | 7.545729 | 7.6375694 | 6.541355 | 6.5577135 | 6.846682 |
| ASMM10P014 | 0.00323 | 0.3172304 | 1.3661505 | up | NM_019388 | Cd86 | CD86 antigen | 75.813077 | 67.714567 | 6.538725 | 6.088609 | 84.23868 | 69.2269 | 73.97365 | 67.62364 | 75.131256 | 60.388805 | 6.450621 | 6.544857 | 6.620698 | 5.9878564 | 6.1189938 | 6.1589766 |
| ASMM10P003 | 0.00324 | 0.3172304 | 1.2672702 | up | NM_021377 | Sorcs1 | VPS10 domain receptor protein SORCS 1 | 600.5029 | 572.77288 | 9.470124 | 9.1284 | 697.59375 | 565.5687 | 538.34625 | 594.5072 | 637.76587 | 486.04556 | 9.435235 | 9.515596 | 9.459541 | 9.1385565 | 9.20723 | 9.039413 |
| ASMM10P039 | 0.00325 | 0.3172304 | 5.0281904 | up | NM_201227 | Dand5 | DAN domain family, member 5 | 39.304893 | 9.033514 | 5.524125 | 3.194086 | 46.73191 | 22.666744 | 48.516026 | 7.7571225 | 10.39004 | 8.953379 | 5.610106 | 4.964973 | 5.997297 | 2.7938476 | 3.258851 | 3.5295596 |
| ASMM10P011 | 0.00327 | 0.3172304 | 1.5901439 | up | NM_028032 | Ppp2r2a | protein phosphatase 2, regulatory subunit B, alpha | 946.0045 | 710.6032 | 10.116862 | 9.447704 | 1027.0757 | 934.10767 | 876.83014 | 676.542 | 735.0959 | 720.1717 | 9.98028 | 10.215346 | 10.154959 | 9.329809 | 9.413707 | 9.599597 |
| ASMM10P054 | 0.00328 | 0.3172304 | 1.3285038 | up | NM_144838 | Sgtb | small glutamine-rich tetratricopeptide repeat (TPR)-containing, beta | 1136.8611 | 1012.4841 | 10.365261 | 9.955458 | 1425.5148 | 953.02795 | 1032.0405 | 1047.5619 | 1066.8016 | 923.08875 | 10.465334 | 10.243004 | 10.387444 | 9.962681 | 9.94902 | 9.954674 |
| ASMM10P041 | 0.00329 | 0.3172304 | 1.6038909 | up | NM_001081 | Phip | pleckstrin homology domain interacting protein | 1270.6587 | 949.25913 | 10.538428 | 9.856852 | 1426.9034 | 1138.0532 | 1247.0195 | 907.3287 | 954.661 | 985.7877 | 10.466407 | 10.501253 | 10.647625 | 9.743659 | 9.784963 | 10.041935 |

| | | | | | | | | | | | | | | | | | | | | | | | |
|---|---|---|---|---|---|---|---|---|---|---|---|---|---|---|---|---|---|---|---|---|---|---|---|
| ASMM10P052 | 0.00335 | 0.3172304 | 1.4129326 | up | NM_008112 | Gdi2 | guanosine diphosphate (GDP) dissociation inhibitor 2 | 1710.6171 | 1457.5548 | 10.965768 | 10.467075 | 1836.8955 | 1644.1394 | 1650.8163 | 1439.4772 | 1505.2108 | 1427.9763 | 10.839516 | 11.008377 | 11.049411 | 10.407826 | 10.432557 | 10.560843 |
| ASMM10P006 | 0.00335 | 0.3172304 | 1.3678143 | up | NM_001271 | Pcdh9 | protocadherin 9 | 826.98138 | 727.46528 | 9.929306 | 9.477433 | 891.02563 | 793.6229 | 796.2956 | 755.01965 | 778.5149 | 648.8613 | 9.789702 | 9.988575 | 10.00964 | 9.490458 | 9.500169 | 9.441673 |
| ASMM10P052 | 0.00337 | 0.3172304 | 1.3790085 | up | NM_029879 | Rgs7bp | regulator of G-protein signalling 7 binding protein | 4160.9545 | 3566.3788 | 12.214458 | 11.750826 | 4315.8755 | 4333.48 | 3833.508 | 3594.9053 | 3650.848 | 3453.383 | 12.098231 | 12.331523 | 12.213619 | 11.721389 | 11.717366 | 11.813724 |
| ASMM10P005 | 0.00339 | 0.3172304 | 1.4615066 | up | NM_001161 | Gucy1b3 | guanylate cyclase 1, soluble, beta 3 | 1601.0239 | 1305.0561 | 10.854122 | 10.306666 | 1941.2285 | 1495.7465 | 1366.0968 | 1384.7754 | 1240.1677 | 1290.2253 | 10.917515 | 10.870308 | 10.774544 | 10.351173 | 10.156393 | 10.412432 |
| ASMM10P005 | 0.00341 | 0.3175536 | 3.4606573 | up | NM_001162 | Susd2 | sushi domain containing 2 | 41.608608 | 13.821033 | 5.613484 | 3.822438 | 52.43968 | 45.67755 | 26.708593 | 14.948392 | 13.446538 | 13.06817 | 5.7758274 | 5.959531 | 5.105094 | 3.7785928 | 3.631034 | 4.0576873 |
| ASMM10P018 | 0.00343 | 0.3175536 | 1.868154 | up | NM_053139 | Pcdhb14 | protocadherin beta 14 | 55.936 | 36.31538 | 6.104113 | 5.2025 | 57.221287 | 52.429832 | 58.15688 | 41.339516 | 34.73695 | 32.869675 | 5.895576 | 6.1495337 | 6.26723 | 5.2757587 | 5.0133243 | 5.3184166 |
| ASMM10P022 | 0.00345 | 0.3175536 | 1.6809543 | up | NM_022995 | Pmepa1 | prostate transmembrane protein, androgen induced 1 | 15112.021 | 10521.259 | 14.032212 | 13.282932 | 14946.7 | 13739.749 | 16649.613 | 9897.157 | 11196.18 | 10470.439 | 13.929114 | 13.935967 | 14.231556 | 13.164616 | 13.282619 | 13.401561 |
| ASMM10P053 | 0.00347 | 0.3180999 | 2.2452999 | up | NM_001271 | Aire | autoimmune regulator (autoimmune polyendocrinopathy candidiasis ectodermal dystrophy) | 10.363888 | 5.575329 | 3.623043 | 2.456135 | 14.29523 | 6.8258104 | 9.970625 | 4.9999995 | 6.7259884 | 4.9999995 | 3.8649077 | 3.341191 | 3.6630306 | 2.3250442 | 2.67684 | 2.3665206 |
| ASMM10P024 | 0.00348 | 0.3180999 | 1.3923041 | up | NM_028136 | Dhx36 | DEAH (Asp-Glu-Ala-His) box polypeptide 36 | 1311.7298 | 1128.8524 | 10.584108 | 10.106634 | 1445.8695 | 1215.3765 | 1273.9434 | 1110.904 | 1162.0796 | 1113.5736 | 10.486636 | 10.587534 | 10.678154 | 10.0409 | 10.065434 | 10.213567 |
| ASMM10P004 | 0.00352 | 0.3192422 | 1.3983841 | up | NM_001098 | Pcdh10 | protocadherin 10 | 10234.017 | 8523.203 | 13.482629 | 12.998868 | 10399.027 | 9910.009 | 10393.015 | 8509.542 | 8641.605 | 8418.462 | 13.38888 | 13.484298 | 13.574709 | 12.951906 | 12.931803 | 13.112896 |
| ASMM10P008 | 0.00353 | 0.3192422 | 1.5992357 | up | NM_029909 | C330018D2( | RIKEN cDNA C330018D20 gene | 111.57407 | 85.38385 | 7.094569 | 6.417186 | 124.31348 | 104.05602 | 106.35269 | 81.034805 | 92.392365 | 82.72438 | 6.995348 | 7.128779 | 7.1595798 | 6.247802 | 6.417138 | 6.586619 |
| ASMM10P053 | 0.00355 | 0.3192422 | 1.2935413 | up | NM_001163 | AU041133 | expressed sequence AU041133 | 734.08138 | 681.82373 | 9.755848 | 9.384522 | 806.29224 | 683.6858 | 712.2661 | 710.4759 | 715.4949 | 619.5004 | 9.6449375 | 9.773568 | 9.849039 | 9.403881 | 9.371151 | 9.378534 |
| ASMM10P007 | 0.00355 | 0.3192422 | 1.3631157 | up | NM_011336 | Ccl27a | chemokine (C-C motif) ligand 27A | 6170.5168 | 5324.0853 | 12.769268 | 12.32236 | 7045.7217 | 5406.211 | 6059.6177 | 5245.5845 | 5505.2554 | 5221.416 | 12.82129 | 12.657883 | 12.82863 | 12.258705 | 12.295012 | 12.413362 |
| ASMM10P001 | 0.00357 | 0.3194283 | 1.4215972 | up | NM_199038 | Casc4 | cancer susceptibility candidate 4 | 104.86451 | 89.280073 | 6.989673 | 6.48216 | 127.85049 | 89.967445 | 96.7756 | 89.08896 | 93.78193 | 84.96933 | 7.033943 | 6.917701 | 7.0173736 | 6.3785872 | 6.4417233 | 6.626169 |
| ASMM10P011 | 0.00363 | 0.3212348 | 1.3429637 | up | NM_026598 | Ebpl | emopamil binding protein-like | 699.90507 | 629.19903 | 9.688442 | 9.263022 | 785.3808 | 671.174 | 643.1604 | 673.36176 | 685.91473 | 528.3206 | 9.608289 | 9.750898 | 9.706139 | 9.322004 | 9.314891 | 9.15217 |
| ASMM10P003 | 0.00370 | 0.3220281 | 1.6374953 | up | NM_172409 | Fmnl2 | formin-like 2 | 81.064405 | 60.251824 | 6.631878 | 5.920387 | 90.60275 | 79.949326 | 72.64114 | 67.929535 | 57.0451 | 55.780838 | 6.5543284 | 6.7466664 | 6.594638 | 5.9945493 | 5.7199397 | 6.0466714 |
| ASMM10P039 | 0.00371 | 0.3220281 | 1.6930552 | up | NM_026067 | Eri1 | exoribonuclease 1 | 249.82339 | 180.63352 | 8.242557 | 7.482928 | 275.26843 | 210.34369 | 263.85806 | 201.49452 | 189.27513 | 151.13092 | 8.122816 | 8.130447 | 8.474409 | 7.5740476 | 7.456944 | 7.4177933 |
| ASMM10P031 | 0.00372 | 0.3220281 | 1.7796197 | up | NM_172707 | Ppp1cb | protein phosphatase 1, catalytic subunit, beta isoform | 9211.809 | 6034.4939 | 13.331039 | 12.49947 | 9491.532 | 8632.076 | 9511.819 | 5661.387 | 5910.331 | 6531.7637 | 13.24708 | 13.27984 | 13.466198 | 12.367944 | 12.393829 | 12.736638 |
| ASMM10P023 | 0.00374 | 0.3220281 | 1.2735632 | up | NM_183112 | 1700029I15I | RIKEN cDNA 1700029I15 gene | 1975.5 | 1850.527 | 11.155656 | 10.806786 | 2220.4146 | 1768.2365 | 1937.849 | 1953.1464 | 1906.1895 | 1692.2451 | 11.103003 | 11.10354 | 11.260427 | 10.849144 | 10.766831 | 10.804383 |
| ASMM10P027 | 0.00379 | 0.3220281 | 2.3504609 | up | NM_018754 | Sfn | stratifin | 3173.4961 | 1579.4884 | 11.816813 | 10.583869 | 2775.32 | 3550.6467 | 3194.5215 | 1563.0936 | 1636.4615 | 1538.91 | 11.423893 | 12.067732 | 11.958814 | 10.525721 | 10.551782 | 10.674105 |
| ASMM10P042 | 0.00384 | 0.3246558 | 3.2757932 | up | NM_026915 | Lyzl4 | lysozyme-like 4 | 34.553018 | 12.290432 | 5.330508 | 3.618664 | 50.387417 | 22.447702 | 30.823935 | 10.94632 | 16.50127 | 9.423706 | 5.7192473 | 4.951931 | 5.320346 | 3.3197172 | 3.9371533 | 3.599121 |
| ASMM10P051 | 0.00389 | 0.3255979 | 4.5603995 | up | NM_175214 | Kif27 | kinesin family member 27 | 27.151525 | 7.013575 | 5.002105 | 2.812945 | 32.584846 | 17.09423 | 31.7755 | 10.75019 | 4.9999995 | 5.2905354 | 5.086925 | 4.5579176 | 5.3614736 | 3.2911968 | 2.3265402 | 2.8210986 |
| ASMM10P006 | 0.00396 | 0.3279542 | 1.6829555 | up | NM_001271 | Milr1 | mast cell immunoglobulin like receptor 1 | 39.867345 | 28.591631 | 5.616862 | 4.865865 | 40.94506 | 38.111156 | 40.54582 | 28.441818 | 31.234186 | 26.098888 | 5.4196825 | 5.6968217 | 5.734083 | 4.732956 | 4.8633375 | 5.0013027 |
| ASMM10P002 | 0.00398 | 0.3279542 | 1.4288231 | up | NM_008073 | Gabrg2 | gamma-aminobutyric acid (GABA) A receptor, subunit gamma 2 | 2137.0839 | 1774.4119 | 11.265015 | 10.750187 | 2444.75 | 1958.68 | 2007.8218 | 1772.965 | 1727.5044 | 1822.7664 | 11.245081 | 11.243325 | 11.306638 | 10.707459 | 10.631093 | 10.91201 |
| ASMM10P009 | 0.00404 | 0.3279542 | 3.0765181 | up | NM_153786 | Vgll2 | vestigial like 2 homolog (Drosophila) | 58.795509 | 22.658727 | 6.150514 | 4.529215 | 62.46383 | 42.908768 | 71.01393 | 22.39903 | 21.701962 | 23.875189 | 6.025999 | 5.8678284 | 6.5577135 | 4.3718357 | 4.3346252 | 4.8811846 |
| ASMM10P005 | 0.00404 | 0.3279542 | 1.3679772 | up | NM_001177 | Gpm6b | glycoprotein m6b | 7105.0957 | 6134.7234 | 12.972221 | 12.520177 | 8043.5283 | 6539.8687 | 6731.89 | 5735.2344 | 6960.7407 | 5708.195 | 13.016997 | 12.924893 | 12.974774 | 12.383039 | 12.623021 | 12.554472 |
| ASMM10P047 | 0.00405 | 0.3279542 | 2.921756 | up | NM_001081 | Gpr179 | G protein-coupled receptor 179 | 241.80616 | 101.45033 | 8.193997 | 6.647161 | 222.86583 | 288.3783 | 214.17435 | 87.43769 | 123.34139 | 93.571915 | 7.828252 | 8.5684595 | 8.18528 | 6.351027 | 6.8303814 | 6.760076 |
| ASMM10P035 | 0.00412 | 0.3279542 | 2.1974847 | up | NM_053165 | Clec2h | C-type lectin domain family 2, member h | 54.083796 | 29.526898 | 6.047542 | 4.911689 | 54.354767 | 60.604355 | 47.292267 | 28.127874 | 32.86155 | 27.59127 | 5.823749 | 6.3579783 | 5.960899 | 4.7157393 | 4.9366956 | 5.082632 |
| ASMM10P007 | 0.00414 | 0.3279542 | 1.4975822 | up | NM_025534 | Ccdc82 | coiled-coil domain containing 82 | 72.145272 | 57.938941 | 6.443665 | 5.86103 | 91.00934 | 67.98616 | 57.440315 | 62.934265 | 63.116203 | 47.766354 | 6.562363 | 6.51954 | 6.2490926 | 5.8809204 | 5.8633585 | 5.838811 |
| ASMM10P000 | 0.00423 | 0.3279542 | 1.2778093 | up | NM_080428 | Fbxw7 | F-box and WD-40 domain protein 7 | 15402.238 | 14261.323 | 14.063453 | 13.70978 | 15627.898 | 14678.077 | 15900.739 | 15337.479 | 14705.795 | 12740.696 | 13.99735 | 14.030775 | 14.162233 | 13.774043 | 13.662663 | 13.692635 |
| ASMM10P043 | 0.00423 | 0.3279542 | 2.3346397 | up | NM_010620 | Kif15 | kinesin family member 15 | 37.974085 | 19.48725 | 5.534254 | 4.311054 | 43.667736 | 36.20777 | 34.04675 | 17.373394 | 20.364222 | 20.724134 | 5.5142903 | 5.6265817 | 5.461889 | 3.9978032 | 4.244046 | 4.691312 |
| ASMM10P005 | 0.00424 | 0.3279542 | 1.5793167 | up | NM_001159 | Tcea1 | transcription elongation factor A (SII) 1 | 300.19936 | 231.56629 | 8.502361 | 7.843061 | 321.36642 | 297.7781 | 281.45355 | 220.83589 | 250.95052 | 222.91246 | 8.337575 | 8.611768 | 8.557741 | 7.7046666 | 7.8618245 | 7.9626913 |
| ASMM10P051 | 0.00425 | 0.3279542 | 2.4226778 | up | NM_011705 | Vrk1 | vaccinia related kinase 1 | 50.776942 | 25.270436 | 5.954269 | 4.677667 | 60.416756 | 44.953342 | 46.960728 | 22.481062 | 25.3665 | 27.963745 | 5.9781775 | 5.935224 | 5.9494057 | 4.377207 | 4.555034 | 5.1007586 |
| ASMM10P012 | 0.00432 | 0.3281053 | 1.8598183 | up | NM_019396 | Cyhr1 | cysteine and histidine rich 1 | 451.27061 | 293.81395 | 9.080599 | 8.185437 | 448.3001 | 480.7565 | 424.75522 | 285.05322 | 315.22906 | 281.15958 | 8.810049 | 9.289786 | 9.141961 | 8.080562 | 8.1913185 | 8.2844305 |
| ASMM10P051 | 0.00432 | 0.3281053 | 1.3627346 | up | NM_001024 | Diras2 | DIRAS family, GTP-binding RAS-like 2 | 86990.175 | 83929.312 | 16.704473 | 16.257968 | 84726.15 | 87404.07 | 88840.305 | 85275.445 | 95706.13 | 70806.36 | 16.58938 | 16.691517 | 16.832521 | 16.240944 | 16.31638 | 16.21658 |
| ASMM10P049 | 0.00434 | 0.3281053 | 2.7979194 | up | NM_019645 | Pkp1 | plakophilin 1 | 424.16562 | 188.21275 | 8.985798 | 7.501444 | 447.51193 | 358.32303 | 466.6619 | 161.81725 | 260.54413 | 142.27687 | 8.807112 | 8.874052 | 9.276231 | 7.252856 | 7.9160805 | 7.3353953 |
| ASMM10P001 | 0.00434 | 0.3281053 | 1.5536943 | up | NM_010298 | Glrb | glycine receptor, beta subunit | 1395.4242 | 1065.0013 | 10.664352 | 10.028649 | 1554.8473 | 1197.7391 | 1433.6863 | 1040.6558 | 1087.7854 | 1066.5626 | 10.584365 | 10.567496 | 10.841194 | 9.952463 | 9.977713 | 10.155771 |
| ASMM10P042 | 0.00438 | 0.3284821 | 1.2653466 | up | NM_001114 | U2surp | U2 snRNP-associated SURP domain containing | 2354.4526 | 2222.4125 | 11.408647 | 11.069114 | 2536.1477 | 2238.601 | 2288.6091 | 2270.6619 | 2333.4958 | 2063.0798 | 11.294745 | 11.444334 | 11.486861 | 11.069208 | 11.06029 | 11.077844 |
| ASMM10P033 | 0.00441 | 0.3292405 | 1.5264063 | up | NM_019715 | Kcmf1 | potassium channel modulatory factor 1 | 6577.5143 | 5063.009 | 12.860076 | 12.249937 | 6715.303 | 5938.9478 | 7078.292 | 4893.8623 | 5428.9517 | 4866.213 | 12.744011 | 12.788603 | 13.047614 | 12.161049 | 12.273059 | 12.315703 |
| ASMM10P004 | 0.00446 | 0.3292405 | 1.3381438 | up | NM_001102 | Kbtbd8 | kelch repeat and BTB (POZ) domain containing 8 | 24.513403 | 21.943918 | 4.901532 | 4.481298 | 27.106478 | 21.805656 | 24.628075 | 22.484688 | 24.648596 | 18.69847 | 4.809439 | 4.914209 | 4.980947 | 4.377697 | 4.5143228 | 4.5518756 |
| ASMM10P005 | 0.00448 | 0.3292405 | 3.8896383 | up | NM_001166 | Tbc1d25 | TBC1 domain family, member 25 | 50.156181 | 15.621302 | 5.923338 | 3.963702 | 63.214973 | 39.87585 | 47.37772 | 17.523869 | 11.21631 | 18.123726 | 6.0440965 | 5.7624574 | 5.9634614 | 4.010867 | 3.3713493 | 4.508891 |
| ASMM10P049 | 0.00449 | 0.3292405 | 1.691638 | up | NM_008890 | Pnmt | phenylethanolamine-N-methyltransferase | 52.560076 | 37.668993 | 6.012534 | 5.254114 | 57.221096 | 49.491978 | 50.967155 | 36.82852 | 38.93033 | 37.24813 | 5.895465 | 6.072576 | 6.0695624 | 5.104418 | 5.1714487 | 5.486474 |
| ASMM10P004 | 0.00457 | 0.3292405 | 1.5239804 | up | NM_130885 | Oxr1 | oxidation resistance 1 | 1804.7057 | 1414.2092 | 11.02826 | 10.420415 | 2046.5371 | 1558.7373 | 1808.8428 | 1492.8739 | 1348.5337 | 1401.2201 | 10.9899 | 10.92743 | 11.167449 | 10.459789 | 10.270343 | 10.531114 |
| ASMM10P048 | 0.00458 | 0.3292405 | 1.3248379 | up | NM_001190 | Usp34 | ubiquitin specific peptidase 34 | 210.79863 | 195.06687 | 8.002419 | 7.596604 | 243.19144 | 184.45338 | 204.75108 | 217.22757 | 200.69475 | 167.2783 | 7.95008 | 7.9441423 | 8.113036 | 7.6843734 | 7.5454354 | 7.560002 |
| ASMM10P005 | 0.00458 | 0.3292405 | 9.7886601 | up | NM_001161 | Zscan4e | zinc finger and SCAN domain containing 4E | 83.728367 | 10.241623 | 6.588069 | 3.296958 | 114.81049 | 88.57647 | 47.798145 | 14.722059 | 4.9999995 | 11.00281 | 6.891194 | 6.896792 | 5.9762225 | 3.7566733 | 2.3265402 | 3.8076608 |
| ASMM10P021 | 0.00463 | 0.3292405 | 2.9120818 | up | NM_198630 | 1810024B03 | RIKEN cDNA 1810024B03 gene | 337.20015 | 140.65787 | 8.660099 | 7.118048 | 340.25104 | 277.599 | 393.7504 | 121.0228 | 143.91351 | 157.0373 | 8.420333 | 8.5179405 | 9.042023 | 6.8301983 | 7.051695 | 7.4722505 |
| ASMM10P022 | 0.00466 | 0.3292405 | 1.681582 | up | NM_001165 | Dcaf17 | DDB1 and CUL4 associated factor 17 | 564.35237 | 402.57671 | 9.382483 | 8.632664 | 610.1565 | 575.29706 | 507.60355 | 387.39923 | 403.79742 | 416.53348 | 9.228702 | 9.534712 | 9.384036 | 8.5203705 | 8.551378 | 8.826244 |
| ASMM10P027 | 0.00467 | 0.3292405 | 1.931099 | up | NM_011732 | Ybx1 | Y box protein 1 | 399.9792 | 251.31778 | 8.911829 | 7.962407 | 413.23068 | 422.3083 | 364.39862 | 264.8753 | 232.89978 | 256.17825 | 8.696415 | 9.10988 | 8.929192 | 7.971104 | 7.7578096 | 8.158307 |
| ASMM10P005 | 0.00468 | 0.3292405 | 1.7816702 | up | NM_001161 | Cbfb | core binding factor beta | 458.17349 | 308.7701 | 9.085631 | 8.252401 | 497.51105 | 372.55322 | 504.4562 | 328.60388 | 328.67294 | 269.03348 | 8.954277 | 8.926969 | 9.375648 | 8.280957 | 8.252679 | 8.223567 |
| ASMM10P004 | 0.00469 | 0.3292405 | 1.7249787 | up | NM_001043 | Fmn1 | formin 1 | 108.99481 | 78.598049 | 7.062387 | 6.275809 | 119.30332 | 98.21833 | 109.46279 | 79.82282 | 97.752846 | 58.21848 | 6.9428234 | 7.0450153 | 7.1993227 | 6.2233663 | 6.498041 | 6.1060185 |
| ASMM10P042 | 0.00472 | 0.3292405 | 1.2683416 | up | NM_007502 | Atp1b3 | ATPase, Na+/K+ transporting, beta 3 polypeptide | 24511.333 | 22799.927 | 14.735505 | 14.392562 | 23933.746 | 24643.355 | 24956.898 | 22329.936 | 25349.76 | 20720.084 | 14.637787 | 14.759909 | 14.80882 | 14.327823 | 14.427638 | 14.422225 |
| ASMM10P040 | 0.00475 | 0.3292405 | 1.4418796 | up | NM_183035 | Defb34 | defensin beta 34 | 1463.7135 | 1214.305 | 10.731884 | 10.203933 | 1729.8516 | 1212.6737 | 1448.6152 | 1330.9834 | 1220.8544 | 1091.0771 | 10.753559 | 10.584577 | 10.857516 | 10.296002 | 10.133341 | 10.182457 |
| ASMM10P039 | 0.00476 | 0.3292405 | 1.4710993 | up | NM_023182 | Ctrl | chymotrypsin-like | 761.44592 | 618.90697 | 9.802159 | 9.245264 | 865.07904 | 633.35913 | 785.8996 | 627.9385 | 658.4675 | 570.3149 | 9.746489 | 9.668567 | 9.99142 | 9.21951 | 9.253289 | 9.262993 |
| ASMM10P013 | 0.00477 | 0.3292405 | 1.3636929 | up | NM_178920 | Mal2 | mal, T cell differentiation protein 2 | 919.74502 | 808.30564 | 10.073712 | 9.626193 | 1014.0428 | 877.75916 | 867.4331 | 892.6179 | 793.76935 | 738.52966 | 9.961746 | 10.124461 | 10.13493 | 9.7191725 | 9.527095 | 9.632313 |
| ASMM10P015 | 0.00479 | 0.3292405 | 4.2861453 | up | NM_026588 | Stx19 | syntaxin 19 | 19.950008 | 5 | 4.439049 | 2.339368 | 34.104683 | 10.701241 | 15.0441 | 4.9999995 | 4.9999995 | 4.9999995 | 5.1500616 | 3.9027317 | 4.264354 | 2.3250442 | 2.3265402 | 2.3665206 |
| ASMM10P017 | 0.00482 | 0.3292405 | 1.4135518 | up | NM_008651 | Mybl1 | myeloblastosis oncogene-like 1 | 138.58219 | 119.86105 | 7.399758 | 6.900434 | 156.75456 | 141.8155 | 117.17651 | 127.03417 | 126.6193 | 105.92968 | 7.3242145 | 7.5720415 | 7.3030195 | 6.9026575 | 6.8676147 | 6.931029 |
| ASMM10P054 | 0.00484 | 0.3292405 | 1.9285723 | up | NM_030597 | Lsm2 | LSM2 homolog, U6 small nuclear RNA associated (S. cerevisiae) | 41.072693 | 25.822035 | 5.653831 | 4.706297 | 44.504158 | 41.821205 | 36.892715 | 31.851322 | 23.252628 | 22.362156 | 5.5407977 | 5.8307633 | 5.5899305 | 4.8957815 | 4.4286847 | 4.7944255 |
| ASMM10P040 | 0.00485 | 0.3292405 | 1.729228 | up | NM_013834 | Sfrp1 | secreted frizzled-related protein 1 | 239.64973 | 169.76501 | 8.190109 | 7.399981 | 245.30855 | 260.4255 | 213.21513 | 171.95345 | 181.49309 | 155.8485 | 7.9613423 | 8.4315 | 8.177485 | 7.345556 | 7.392947 | 7.46144 |
| ASMM10P010 | 0.00488 | 0.3292405 | 1.3371939 | up | NM_001037 | Edil3 | EGF-like repeats and discoidin I-like domains 3 | 590.01607 | 530.39812 | 9.444832 | 9.025623 | 674.98706 | 538.073 | 556.98816 | 508.7894 | 557.2025 | 525.20245 | 9.383146 | 9.44706 | 9.504289 | 8.918589 | 9.013337 | 9.144943 |
| ASMM10P006 | 0.00491 | 0.3292405 | 1.6097215 | up | NM_001253 | Mbnl1 | muscleblind-like 1 (Drosophila) | 206.86644 | 157.25253 | 7.975591 | 7.28878 | 236.20378 | 176.04259 | 208.35294 | 159.49112 | 156.28432 | 155.98216 | 7.9087496 | 7.875823 | 8.1422 | 7.2334247 | 7.1704206 | 7.462494 |
| ASMM10P047 | 0.00493 | 0.3292405 | 1.6509606 | up | NM_001005 | Ypel2 | yippee-like 2 (Drosophila) | 49256.512 | 36292.454 | 15.793012 | 15.069706 | 43199.285 | 53431.41 | 51138.84 | 36795.75 | 38042.5 | 34039.113 | 15.540427 | 15.917434 | 15.921174 | 15.048478 | 15.041697 | 15.118943 |
| ASMM10P005 | 0.00493 | 0.3292405 | 1.6740367 | up | NM_001159 | Slc44a1 | solute carrier family 44, member 1 | 20.184178 | 14.339319 | 4.614158 | 3.870827 | 22.90082 | 19.646011 | 18.005703 | 13.670619 | 16.89206 | 12.455279 | 4.5625935 | 4.7547646 | 4.525116 | 3.6478431 | 3.9753656 | 3.989272 |
| ASMM10P014 | 0.00494 | 0.3292405 | 1.4710004 | up | NM_001110 | Dzip3 | DAZ interacting protein 3, zinc finger | 1166.4594 | 949.93777 | 10.420826 | 9.864029 | 1243.5375 | 1184.6969 | 1071.1439 | 998.6087 | 938.5765 | 912.6281 | 10.263813 | 10.55364 | 10.445026 | 9.888218 | 9.76466 | 9.939208 |
| ASMM10P036 | 0.00498 | 0.3292405 | 1.8254471 | up | NM_001011 | Olfr543 | olfactory receptor 543 | 90.72067 | 60.940346 | 6.787808 | 5.919558 | 102.5103 | 72.518005 | 97.133705 | 62.499847 | 74.252144 | 46.069046 | 6.728525 | 6.6103287 | 7.0245695 | 5.869608 | 6.1013765 | 5.787689 |
| ASMM10P029 | 0.00503 | 0.3292405 | 1.9067746 | up | NM_019732 | Runx3 | runt related transcription factor 3 | 33.510137 | 21.02076 | 5.358341 | 4.427207 | 34.460815 | 33.54005 | 32.529545 | 20.821987 | 21.590195 | 20.650099 | 5.1633472 | 5.5135403 | 5.398135 | 4.2671003 | 4.3278418 | 4.6866775 |
| ASMM10P039 | 0.00508 | 0.3292405 | 1.4426257 | up | NM_198007 | Ascc3 | activating signal cointegrator 1 complex subunit 3 | 788.3552 | 655.99235 | 9.858848 | 9.330151 | 880.11224 | 746.38617 | 738.5672 | 637.5294 | 658.46967 | 671.97797 | 9.770346 | 9.902553 | 9.903645 | 9.239142 | 9.253501 | 9.49781 |
| ASMM10P012 | 0.00514 | 0.3292405 | 1.4025167 | up | NM_178399 | 3110035E14 | RIKEN cDNA 3110035E14 gene | 206.38339 | 180.73203 | 7.979982 | 7.491964 | 223.08922 | 194.78163 | 201.27933 | 180.96538 | 193.50227 | 167.72845 | 7.8288655 | 8.02487 | 8.086209 | 7.4210835 | 7.4911957 | 7.5636115 |
| ASMM10P005 | 0.00515 | 0.3292405 | 1.4267544 | up | NM_001199 | Ghitm | growth hormone inducible transmembrane protein | 1487.939 | 1253.5573 | 10.764742 | 10.252005 | 1592.5975 | 1369.4674 | 1501.7522 | 1255.9567 | 1290.954 | 1213.7611 | 10.62539 | 10.756212 | 10.912624 | 10.214561 | 10.208733 | 10.332721 |
| ASMM10P051 | 0.00515 | 0.3292405 | 1.9923416 | up | NM_009396 | Tnfaip2 | tumor necrosis factor, alpha-induced protein 2 | 696.00305 | 418.34963 | 9.680131 | 8.685666 | 657.7828 | 676.9988 | 753.22754 | 417.7128 | 466.08044 | 371.25565 | 9.34462 | 9.762159 | 9.933615 | 8.629802 | 8.75964 | 8.667557 |
| ASMM10P025 | 0.00515 | 0.3292405 | 1.3905732 | up | NM_011020 | Hspa4l | heat shock protein 4 like | 34173.851 | 29397.584 | 15.240197 | 14.764517 | 33096.87 | 34183.223 | 35241.46 | 29694.855 | 29561.54 | 28936.357 | 15.130871 | 15.250319 | 15.3394 | 14.73917 | 14.674238 | 14.880143 |
| ASMM10P002 | 0.00517 | 0.3292405 | 1.3460923 | up | NM_177408 | Gabrg2 | gamma-aminobutyric acid (GABA) A receptor, subunit gamma 2 | 2226.2199 | 1968.1076 | 11.323999 | 10.895222 | 2503.55 | 2078.5728 | 2096.5369 | 2132.2498 | 1887.8785 | 1884.1945 | 11.272217 | 11.332123 | 11.367658 | 10.978521 | 10.751548 | 10.955597 |
| ASMM10P009 | 0.00525 | 0.3292405 | 1.2834316 | up | NM_153195 | Fbxo7 | F-box protein 7 | 56.257078 | 53.45794 | 6.10516 | 5.745153 | 64.17664 | 55.23504 | 49.359554 | 57.138054 | 59.63999 | 43.595776 | 6.06575 | 6.226495 | 6.0232344 | 5.7415996 | 5.7821984 | 5.7116623 |
| ASMM10P011 | 0.00526 | 0.3292405 | 1.4714328 | up | NM_011200 | Ptp4a1 | protein tyrosine phosphatase 4a1 | 16519.812 | 13157.584 | 14.156511 | 13.599289 | 15755.993 | 15922.858 | 17880.584 | 12860.725 | 14136.655 | 12475.373 | 14.009912 | 14.133593 | 14.326028 | 13.523999 | 13.606384 | 13.667485 |
| ASMM10P026 | 0.00527 | 0.3292405 | 2.6507554 | up | NM_029937 | Nup210l | nucleoporin 210-like | 49.760557 | 22.087506 | 5.898507 | 4.492103 | 61.573624 | 34.236576 | 53.47147 | 21.90079 | 21.022131 | 23.339598 | 6.005441 | 5.5460105 | 6.1440687 | 4.33563 | 4.2898555 | 4.850824 |
| ASMM10P039 | 0.00528 | 0.3292405 | 2.8485186 | up | NM_001009 | Slc38a8 | solute carrier family 38, member 8 | 107.41704 | 45.363427 | 7.026409 | 5.516197 | 93.43025 | 135.08249 | 93.73839 | 50.73748 | 44.760616 | 40.592186 | 6.599154 | 7.508708 | 6.9713655 | 5.567083 | 5.372816 | 5.608693 |
| ASMM10P019 | 0.00529 | 0.3292405 | 1.4044811 | up | NM_183195 | Marveld1 | MARVEL (membrane-associating) domain containing 1 | 8393.9757 | 6992.1982 | 13.196959 | 12.706922 | 9277.391 | 7368.701 | 8535.835 | 6693.1577 | 7942.884 | 6340.553 | 13.204515 | 13.075252 | 13.311111 | 12.615654 | 12.811841 | 12.693271 |
| ASMM10P050 | 0.00531 | 0.3292405 | 1.7785179 | up | NM_177290 | Itgb8 | integrin beta 8 | 187.63949 | 129.11992 | 7.829815 | 6.99914 | 226.97803 | 173.95596 | 161.98447 | 110.91731 | 149.6847 | 126.75774 | 7.8520803 | 7.859416 | 7.77795 | 6.7046914 | 7.1106257 | 7.1821027 |
| ASMM10P010 | 0.00533 | 0.3292405 | 1.71677 | up | NM_001177 | Gm10767 | predicted gene 10767 | 63.406542 | 44.820082 | 6.283244 | 5.503548 | 70.67111 | 61.28975 | 58.258766 | 44.60994 | 44.345097 | 45.50521 | 6.2055407 | 6.3746967 | 6.2694955 | 5.383946 | 5.3576646 | 5.769032 |
| ASMM10P002 | 0.00534 | 0.3292405 | 1.4840987 | up | NM_008194 | Gyk | glycerol kinase | 611.25942 | 494.93838 | 9.496092 | 8.926505 | 640.11206 | 607.8329 | 585.8333 | 521.1451 | 502.1403 | 461.52975 | 9.300665 | 9.615288 | 9.572322 | 8.94965 | 8.862991 | 8.966873 |
| ASMM10P040 | 0.00535 | 0.3292405 | 1.5162812 | up | NM_009506 | Vegfc | vascular endothelial growth factor C | 287.84814 | 231.7104 | 8.444906 | 7.844368 | 315.6365 | 276.00653 | 271.9014 | 217.22751 | 251.67583 | 226.22787 | 8.311854 | 8.510186 | 8.512677 | 7.684173 | 7.866776 | 7.982156 |
| ASMM10P006 | 0.00536 | 0.3292405 | 1.3799227 | up | NM_001253 | Mbnl1 | muscleblind-like 1 (Drosophila) | 214.68606 | 190.93823 | 8.029867 | 7.565279 | 248.41054 | 198.50294 | 197.14471 | 179.62257 | 220.62302 | 172.56909 | 7.9803867 | 8.051698 | 8.057516 | 7.4094133 | 7.680914 | 7.605511 |
| ASMM10P043 | 0.00542 | 0.3292405 | 2.0284076 | up | NM_001191 | Ddx43 | DEAD (Asp-Glu-Ala-Asp) box polypeptide 43 | 96.000248 | 57.5922 | 6.873316 | 5.852969 | 95.61512 | 80.47044 | 111.91518 | 59.50714 | 65.16978 | 48.09968 | 6.6303663 | 6.756158 | 7.2334247 | 5.80081 | 5.9110093 | 5.847087 |
| ASMM10P012 | 0.00544 | 0.3292405 | 1.3421783 | up | NM_001085 | Tspyl5 | testis-specific protein, Y-encoded-like 5 | 47001.085 | 42391.942 | 15.716255 | 15.291678 | 45402.34 | 46773.227 | 48827.688 | 41894.055 | 45145.53 | 40136.24 | 15.613987 | 15.701617 | 15.83316 | 15.227434 | 15.270768 | 15.376833 |
| ASMM10P015 | 0.00544 | 0.3292405 | 1.2653174 | up | NM_013918 | Usp25 | ubiquitin specific peptidase 25 | 1470.751 | 1398.4142 | 10.741528 | 10.402029 | 1601.4939 | 1470.6029 | 1340.1562 | 1411.268 | 1478.8423 | 1305.1323 | 10.634052 | 10.843617 | 10.746915 | 10.378347 | 10.401962 | 10.425777 |
| ASMM10P020 | 0.00545 | 0.3292405 | 1.4722449 | up | NM_001081 | Ttc30a2 | tetratricopeptide repeat domain 30A2 | 402.828 | 334.47153 | 8.925037 | 8.367019 | 421.80078 | 399.82602 | 386.8572 | 352.9273 | 346.2641 | 304.22318 | 8.72509 | 9.031986 | 9.018034 | 8.384083 | 8.326403 | 8.390571 |

| | | | | | | | | | | | | | | | | | | | | | | |
|---|---|---|---|---|---|---|---|---|---|---|---|---|---|---|---|---|---|---|---|---|---|---|
| ASMM10P016 | 0.00551 | 0.3292405 | 1.4600049 | up | NM_023294 | Ndc80 | NDC80 homolog, kinetochore complex component (S. cerevisiae) | 306.6542 | 256.40693 | 8.536305 | 7.990332 | 325.18774 | 301.705 | 293.06985 | 252.74376 | 279.88416 | 236.59288 | 8.357038 | 8.6315975 | 8.62028 | 7.9027658 | 8.018909 | 8.049321 |
| ASMM10P040 | 0.00555 | 0.3292405 | 1.5416031 | up | NM_178395 | Zdhhc2 | zinc finger, DHHC domain containing 2 | 12912.498 | 9750.667 | 13.806628 | 13.182197 | 12578.435 | 13262.767 | 12896.293 | 9802.598 | 9351.993 | 10097.41 | 13.665814 | 13.892315 | 13.861755 | 13.150675 | 13.043987 | 13.351928 |
| ASMM10P007 | 0.00556 | 0.3292405 | 1.3597903 | up | NM_011848 | Nek3 | NIMA (never in mitosis gene a)-related expressed kinase 3 | 18.591131 | 16.454584 | 4.493484 | 4.0501 | 21.93517 | 17.180252 | 16.65797 | 18.679592 | 18.872742 | 11.81142 | 4.501157 | 4.5643644 | 4.4149313 | 4.1058164 | 4.131818 | 3.9126658 |
| ASMM10P024 | 0.00558 | 0.3292405 | 2.7557886 | up | NM_001012 | Wfdc13 | WAP four-disulfide core domain 13 | 15.930968 | 6.607477 | 4.268 | 2.805535 | 18.404903 | 15.557511 | 13.83049 | 5.2254744 | 7.0862527 | 7.510703 | 4.237528 | 4.4227595 | 4.1437125 | 2.4033093 | 2.730752 | 3.282543 |
| ASMM10P010 | 0.00564 | 0.3292405 | 2.2756254 | up | NM_053112 | Ear10 | eosinophil-associated, ribonuclease A family, member 10 | 16.79255 | 8.851254 | 4.339026 | 3.152763 | 15.277378 | 15.31029 | 19.789982 | 10.790289 | 9.441186 | 6.3222857 | 3.9558127 | 4.3998876 | 4.6613774 | 3.298198 | 3.109694 | 3.0503964 |
| ASMM10P020 | 0.00565 | 0.3292405 | 1.2600704 | up | NM_001001 | Ccdc148 | coiled-coil domain containing 148 | 691.45293 | 656.26185 | 9.664172 | 9.330667 | 836.1593 | 618.7396 | 619.4599 | 688.15375 | 642.6572 | 637.9746 | 9.700266 | 9.640085 | 9.652164 | 9.351094 | 9.220101 | 9.420807 |
| ASMM10P054 | 0.00567 | 0.3292405 | 1.4410769 | up | NM_153166 | Cpne5 | copine V | 9063.6692 | 7297.1994 | 13.304052 | 12.776904 | 9318.953 | 9831.099 | 8040.9556 | 7259.023 | 7486.6265 | 7145.9487 | 13.213637 | 13.472699 | 13.225819 | 12.724278 | 12.731988 | 12.874447 |
| ASMM10P027 | 0.00569 | 0.3292405 | 1.2910367 | up | NM_198026 | Iqcc | IQ motif containing C | 448.62823 | 419.64409 | 9.064304 | 8.695774 | 514.72943 | 399.3371 | 431.81815 | 433.59988 | 423.17987 | 402.15253 | 8.999694 | 9.029748 | 9.163469 | 8.686471 | 8.621036 | 8.779814 |
| ASMM10P009 | 0.00571 | 0.3292405 | 1.6443384 | up | NM_181731 | Cabyr | calcium-binding tyrosine-(Y)-phosphorylation regulated (fibrousheathin 2) | 49.375482 | 36.386127 | 5.920127 | 5.20262 | 56.018143 | 45.34858 | 46.759724 | 33.719425 | 39.721985 | 35.716972 | 5.8649645 | 5.951013 | 5.944404 | 4.978237 | 5.2008924 | 5.4287305 |
| ASMM10P038 | 0.00571 | 0.3292405 | 1.5746092 | up | NM_175023 | Rbbp6 | retinoblastoma binding protein 6 | 19136.761 | 14202.016 | 14.367527 | 13.712533 | 17873.672 | 19217.729 | 20318.883 | 13720.688 | 14563.587 | 14321.773 | 14.191037 | 14.409229 | 14.502316 | 13.622457 | 13.64528 | 13.869864 |
| ASMM10P050 | 0.00572 | 0.3292405 | 1.363721 | up | NM_172578 | Mis18bp1 | MIS18 binding protein 1 | 48.037475 | 42.473639 | 5.870113 | 5.422565 | 56.25359 | 39.324074 | 48.53476 | 43.358913 | 47.43303 | 36.628975 | 5.8709373 | 5.741796 | 5.9976068 | 5.346955 | 5.4565372 | 5.4642024 |
| ASMM10P008 | 0.00575 | 0.3292405 | 1.4594942 | up | NM_029553 | Ttc8 | tetratricopeptide repeat domain 8 | 97.759752 | 81.34753 | 6.894164 | 6.348695 | 115.26782 | 80.3475 | 97.66394 | 86.45748 | 82.073 | 75.51211 | 6.8966866 | 6.7536287 | 7.0321765 | 6.3350167 | 6.2495747 | 6.461495 |
| ASMM10P011 | 0.00575 | 0.3292405 | 1.2982984 | up | NM_020506 | Xpo4 | exportin 4 | 142.17032 | 134.42184 | 7.442247 | 7.065625 | 163.6275 | 132.0147 | 130.86877 | 141.73122 | 135.00598 | 126.52831 | 7.384153 | 7.472509 | 7.47008 | 7.058977 | 6.9588995 | 7.1789994 |
| ASMM10P004 | 0.00580 | 0.3292405 | 1.9058937 | up | NM_001080 | Tmpo | thymopoietin | 471.50203 | 290.47248 | 9.096217 | 8.165749 | 605.1156 | 331.39914 | 477.99136 | 302.452 | 326.04312 | 242.92232 | 9.218575 | 8.764902 | 9.305173 | 8.167581 | 8.240774 | 8.088892 |
| ASMM10P031 | 0.00583 | 0.3292405 | 1.3492095 | up | NM_028975 | Tmem33 | transmembrane protein 33 | 22551.571 | 19471.061 | 14.60807 | 14.175955 | 21869.01 | 22819.139 | 22966.564 | 18928.193 | 20333.637 | 19151.354 | 14.505935 | 14.639724 | 14.67855 | 14.092805 | 14.139514 | 14.295547 |
| ASMM10P014 | 0.00588 | 0.3292405 | 1.4075089 | up | NM_028295 | Pdia5 | protein disulfide isomerase associated 5 | 3465.9412 | 2894.9536 | 11.940901 | 11.447757 | 4372.7036 | 3055.9797 | 2969.1404 | 2961.993 | 3125.5344 | 2597.3333 | 12.118761 | 11.852505 | 11.851438 | 11.438886 | 11.493271 | 11.411115 |
| ASMM10P041 | 0.00590 | 0.3292405 | 1.4153862 | up | NM_013669 | Snap91 | synaptosomal-associated protein 91 | 166.34936 | 144.29369 | 7.666661 | 7.165465 | 190.6729 | 155.16429 | 153.21089 | 154.89937 | 139.0423 | 138.93939 | 7.604784 | 7.6945944 | 7.7006054 | 7.190689 | 7.0012035 | 7.304504 |
| ASMM10P004 | 0.00592 | 0.3292405 | 1.6373911 | up | NM_001080 | Tmpo | thymopoietin | 492.52592 | 363.09313 | 9.188593 | 8.477194 | 537.65204 | 416.61993 | 523.3058 | 362.16196 | 419.33234 | 307.7851 | 9.055319 | 9.090022 | 9.420437 | 8.421291 | 8.60634 | 8.40395 |
| ASMM10P054 | 0.00595 | 0.3292405 | 1.4264119 | up | NM_001170 | Wdr33 | WD repeat domain 33 | 787.79365 | 663.54821 | 9.859972 | 9.347581 | 849.95526 | 760.9959 | 752.4298 | 676.4785 | 653.9679 | 660.19824 | 9.720259 | 9.927621 | 9.932036 | 9.329653 | 9.243446 | 9.469645 |
| ASMM10P020 | 0.00598 | 0.3292405 | 1.4886318 | up | NM_026176 | Pdcl | phosducin-like | 10892.123 | 8559.2612 | 13.57204 | 12.998053 | 10686.424 | 10733.113 | 11256.832 | 8020.411 | 9066.1455 | 8591.227 | 13.430421 | 13.611572 | 13.674128 | 12.864183 | 12.991664 | 13.138313 |
| ASMM10P027 | 0.00599 | 0.3292405 | 6.0272349 | up | NM_080458 | Bsnd | Bartter syndrome, infantile, with sensorineural deafness (Barttin) | 28.623679 | 5 | 4.930865 | 2.339368 | 44.95138 | 11.248029 | 29.671629 | 4.9999995 | 4.9999995 | 4.9999995 | 5.555047 | 3.9723592 | 5.2651877 | 2.3250442 | 2.3265402 | 2.3665206 |
| ASMM10P022 | 0.00602 | 0.3292405 | 1.9115828 | up | NM_177383 | Gpr21 | G protein-coupled receptor 21 | 908.3212 | 563.53343 | 10.042256 | 9.107488 | 1006.7739 | 695.02673 | 1023.163 | 603.7415 | 598.8213 | 488.03748 | 9.949565 | 9.800636 | 10.376566 | 9.159672 | 9.117824 | 9.044968 |
| ASMM10P005 | 0.00602 | 0.3292405 | 1.4242815 | up | NM_022309 | Cbfb | core binding factor beta | 596.28895 | 501.52711 | 9.45535 | 8.945116 | 690.00354 | 493.12622 | 605.7371 | 542.2887 | 501.81546 | 460.47717 | 9.417631 | 9.329653 | 9.618766 | 9.010175 | 8.862518 | 8.962654 |
| ASMM10P055 | 0.00605 | 0.3292405 | 1.8177555 | up | NM_001145 | 6430550D23 | RIKEN cDNA 6430550D23 gene | 114.8606 | 78.083517 | 7.144974 | 6.282816 | 110.45449 | 116.67989 | 117.44742 | 78.99678 | 89.93298 | 65.32079 | 6.8359566 | 7.2915826 | 7.3073826 | 6.2075453 | 6.376886 | 6.264016 |
| ASMM10P049 | 0.00608 | 0.3292405 | 1.5056095 | up | NM_001163 | Rhot1 | ras homolog gene family, member T1 | 689.7285 | 551.84245 | 9.672715 | 9.082367 | 720.17737 | 657.56537 | 691.44275 | 558.47876 | 553.4201 | 543.6285 | 9.487539 | 9.721472 | 9.809133 | 9.051252 | 9.004032 | 9.191817 |
| ASMM10P006 | 0.00610 | 0.3292405 | 1.3814749 | up | NM_008072 | Gabrd | gamma-aminobutyric acid (GABA) A receptor, subunit delta | 633.45437 | 552.95733 | 9.545734 | 9.079525 | 684.8831 | 636.7107 | 578.7693 | 595.6086 | 588.26025 | 475.00314 | 9.404467 | 9.676295 | 9.556441 | 9.141961 | 9.090124 | 9.00649 |
| ASMM10P034 | 0.00612 | 0.3292405 | 1.7508531 | up | NM_178608 | Reep1 | receptor accessory protein 1 | 133.18383 | 92.607577 | 7.339552 | 6.531494 | 161.47356 | 120.11383 | 117.9641 | 96.34879 | 84.33429 | 97.13965 | 7.367454 | 7.3358727 | 7.315329 | 6.4949565 | 6.2885165 | 6.8110085 |
| ASMM10P026 | 0.00613 | 0.3292405 | 1.4559713 | up | NM_145711 | Tox | thymocyte selection-associated high mobility group box | 205.91386 | 175.12301 | 7.977073 | 7.435091 | 223.8735 | 200.14996 | 193.71811 | 195.2856 | 192.46558 | 137.61784 | 7.834324 | 8.063429 | 8.033465 | 7.529308 | 7.4846263 | 7.291338 |
| ASMM10P021 | 0.00615 | 0.3292405 | 5.7359318 | up | NM_001271 | Trp53tg5 | transformation related protein 53 target 5 | 54.568443 | 9.958743 | 5.859309 | 3.339281 | 88.20833 | 51.620895 | 23.876104 | 11.468539 | 10.456589 | 7.9511023 | 6.514963 | 6.128293 | 4.9346714 | 3.3812447 | 3.2715187 | 3.3650804 |
| ASMM10P031 | 0.00621 | 0.3292405 | 1.524307 | up | NM_001042 | Prss57 | protease, serine 57 | 103.22873 | 82.101752 | 6.970995 | 6.362841 | 124.04965 | 90.83114 | 94.805405 | 82.520065 | 80.90662 | 82.87857 | 6.99225 | 6.9320846 | 6.9886503 | 6.270583 | 6.228912 | 6.5890293 |
| ASMM10P005 | 0.00621 | 0.3292405 | 1.4376769 | up | NM_001163 | Fam172a | family with sequence similarity 172, member A | 95.310115 | 80.948732 | 6.864237 | 6.340498 | 106.23862 | 98.61772 | 81.074005 | 86.24987 | 84.62753 | 71.968796 | 6.779909 | 7.05326 | 6.7595425 | 6.331606 | 6.293307 | 6.39658 |
| ASMM10P036 | 0.00625 | 0.3292405 | 2.2116391 | up | NM_053080 | Aldh1a3 | aldehyde dehydrogenase family 1, subfamily A3 | 904.52893 | 493.01821 | 10.05313 | 8.908014 | 852.99414 | 844.02435 | 1016.5683 | 459.54697 | 606.1617 | 413.34595 | 9.725688 | 10.067304 | 10.366397 | 8.772125 | 9.135466 | 8.81645 |
| ASMM10P026 | 0.00627 | 0.3292405 | 1.6274127 | up | NM_025663 | Gpatch4 | G patch domain containing 4 | 60.081105 | 44.360939 | 6.191099 | 5.488519 | 73.69272 | 52.487034 | 54.06356 | 47.120823 | 41.97575 | 43.986244 | 6.2620378 | 6.1521497 | 6.159109 | 5.4648194 | 5.2781587 | 5.722578 |
| ASMM10P044 | 0.00627 | 0.3292405 | 1.3186031 | up | NM_009530 | Atrx | alpha thalassemia/mental retardation syndrome X-linked homolog (human) | 25739.256 | 22877.442 | 14.800703 | 14.401693 | 24356.12 | 25857.457 | 27004.191 | 23314.867 | 24358.02 | 20959.438 | 14.664965 | 14.81877 | 14.918375 | 14.381764 | 14.385359 | 14.437956 |
| ASMM10P036 | 0.00629 | 0.3292405 | 1.3076472 | up | NM_001101 | Gsg1l | GSG1-like | 48137.406 | 44923.808 | 15.762061 | 15.375088 | 45616.742 | 49222.895 | 49572.582 | 45878.78 | 48096.75 | 40795.895 | 15.622331 | 15.801127 | 15.862726 | 15.349952 | 15.372552 | 15.40276 |
| ASMM10P036 | 0.00630 | 0.3292405 | 1.5839012 | up | NM_201374 | Ccdc155 | coiled-coil domain containing 155 | 1930.9357 | 1443.0694 | 11.112835 | 10.449353 | 2379.4438 | 1534.4203 | 1878.9431 | 1419.2527 | 1446.0808 | 1463.8748 | 11.205961 | 10.906086 | 11.226458 | 10.386277 | 10.367843 | 10.593938 |
| ASMM10P042 | 0.00633 | 0.3292405 | 1.2925477 | up | NM_011352 | Sema7a | sema domain, immunoglobulin domain (Ig), and GPI membrane anchor, (semaphorin) 7A | 121.86638 | 114.71759 | 7.20875 | 6.838532 | 148.62988 | 103.24298 | 113.72629 | 113.83878 | 124.46989 | 105.84411 | 7.248963 | 7.1174555 | 7.2598314 | 6.7430563 | 6.8433104 | 6.9292307 |
| ASMM10P031 | 0.00636 | 0.3292405 | 1.9035135 | up | NM_007925 | Eln | elastin | 1482.4228 | 927.58517 | 10.749028 | 9.820363 | 1545.0392 | 1273.4087 | 1628.8206 | 824.5427 | 1081.6869 | 876.5259 | 10.576723 | 10.645571 | 11.024791 | 9.6064205 | 9.970396 | 9.884274 |
| ASMM10P015 | 0.00638 | 0.3292405 | 2.0993301 | up | NM_028091 | Osgepl1 | O-sialoglycoprotein endopeptidase-like 1 | 70.340603 | 40.617724 | 6.434002 | 5.364073 | 66.193855 | 66.311775 | 78.51618 | 39.791523 | 42.482853 | 39.578796 | 6.1099324 | 6.48398 | 6.7080936 | 5.2231374 | 5.295524 | 5.5735574 |
| ASMM10P015 | 0.00642 | 0.3292405 | 1.4907097 | up | NM_008997 | Rab11b | RAB11B, member RAS oncogene family | 4796.3137 | 3778.0881 | 12.408329 | 11.83233 | 5278.7554 | 4570.591 | 4539.5947 | 3387.1492 | 4001.0361 | 3946.079 | 12.385596 | 12.408667 | 12.430724 | 11.638681 | 11.840403 | 12.017905 |
| ASMM10P023 | 0.00643 | 0.3292405 | 1.8182846 | up | NM_029475 | Adal | adenosine deaminase-like | 316.58848 | 211.879 | 8.582394 | 7.719816 | 320.60553 | 325.51355 | 303.64636 | 218.30794 | 201.18715 | 216.1419 | 8.334626 | 8.738901 | 8.673654 | 7.6905866 | 7.5494576 | 7.9194026 |
| ASMM10P020 | 0.00644 | 0.3292405 | 1.6319108 | up | NM_145526 | P2rx3 | purinergic receptor P2X, ligand-gated ion channel, 3 | 175.9922 | 132.309 | 7.750263 | 7.043701 | 183.93239 | 161.38087 | 182.66335 | 138.4709 | 132.24489 | 126.21121 | 7.550624 | 7.7503467 | 7.9498177 | 7.0260954 | 6.9302564 | 7.17475 |
| ASMM10P036 | 0.00644 | 0.3292405 | 1.5316909 | up | NM_177001 | 9130023H24 | RIKEN cDNA 9130023H24 gene | 3122.0823 | 2405.1403 | 11.799659 | 11.184534 | 3171.876 | 2888.875 | 3305.4958 | 2331.2356 | 2636.1292 | 2248.0562 | 11.619994 | 11.775877 | 12.003106 | 11.10938 | 11.240716 | 11.203506 |
| ASMM10P005 | 0.00648 | 0.3300405 | 1.5335197 | up | NM_001172 | Cuedc1 | CUE domain containing 1 | 151.29974 | 120.70911 | 7.528692 | 6.911845 | 163.03761 | 160.7374 | 130.1242 | 123.34457 | 127.54707 | 111.23568 | 7.379432 | 7.7440205 | 7.4626236 | 6.8583846 | 6.879818 | 6.9973335 |
| ASMM10P044 | 0.00665 | 0.3337726 | 1.3949048 | up | NM_212444 | Gyk | glycerol kinase | 526.41615 | 455.46263 | 9.290751 | 8.810584 | 576.30646 | 504.0339 | 498.9081 | 500.65213 | 443.23508 | 422.50067 | 9.152825 | 9.357399 | 9.362029 | 8.89603 | 8.689594 | 8.846129 |
| ASMM10P008 | 0.00667 | 0.3337726 | 1.4254822 | up | NM_146239 | Cdk17 | cyclin-dependent kinase 17 | 903.48473 | 763.21535 | 10.054604 | 9.543154 | 956.6092 | 876.3689 | 877.4761 | 838.5588 | 755.509 | 695.57825 | 9.885958 | 10.122084 | 10.155771 | 9.629228 | 9.452948 | 9.547287 |
| ASMM10P041 | 0.00670 | 0.3337726 | 1.7552739 | up | NM_028908 | Map10 | microtubule-associated protein 10 | 1487.7033 | 1010.1281 | 10.761188 | 9.949491 | 1469.0648 | 1391.4646 | 1602.5804 | 955.00055 | 1078.735 | 996.64874 | 10.509743 | 10.775518 | 10.998302 | 9.821985 | 9.966147 | 10.060342 |
| ASMM10P029 | 0.00671 | 0.3337726 | 1.5267856 | up | NM_023537 | Rab3b | RAB3B, member RAS oncogene family | 4843.4772 | 3724.0568 | 12.427762 | 11.817264 | 4851.8906 | 4701.309 | 4977.232 | 3694.9553 | 3627.3503 | 3849.8647 | 12.271359 | 12.447731 | 12.564195 | 11.76523 | 11.708176 | 11.978386 |
| ASMM10P055 | 0.00671 | 0.3337726 | 1.30648 | up | NM_001159 | Fip1l1 | FIP1 like 1 (S. cerevisiae) | 195.02161 | 183.68251 | 7.896126 | 7.510441 | 219.97607 | 194.70015 | 170.3886 | 190.42545 | 205.31007 | 155.31201 | 7.8112 | 8.024225 | 7.852954 | 7.492798 | 7.5809426 | 7.4575834 |
| ASMM10P008 | 0.00675 | 0.3337726 | 1.2881813 | up | NM_029773 | Spopl | speckle-type POZ protein-like | 180.17606 | 171.47991 | 7.781253 | 7.415918 | 207.38365 | 163.09232 | 170.0522 | 173.29791 | 176.39111 | 164.75072 | 7.725311 | 7.7682266 | 7.8502226 | 7.3572054 | 7.353121 | 7.537427 |
| ASMM10P035 | 0.00677 | 0.3337726 | 1.3613701 | up | NM_026815 | Upk1a | uroplakin 1A | 271.74046 | 245.66846 | 8.35881 | 7.913751 | 314.69815 | 252.51755 | 248.00568 | 260.6039 | 283.66544 | 192.73605 | 8.308747 | 8.384083 | 8.383601 | 7.9493465 | 8.036007 | 7.7558994 |
| ASMM10P022 | 0.00678 | 0.3337726 | 1.611073 | up | NM_020283 | B3galt1 | UDP-Gal:betaGlcNAc beta 1,3-galactosyltransferase, polypeptide 1 | 2380.317 | 1750.4501 | 11.418901 | 10.730879 | 2491.3533 | 2142.9492 | 2506.6484 | 1729.656 | 1707.9448 | 1813.7495 | 11.264808 | 11.384131 | 11.607764 | 10.672441 | 10.615802 | 10.904394 |
| ASMM10P027 | 0.00679 | 0.3337726 | 1.6196347 | up | NM_146254 | Wdr78 | WD repeat domain 78 | 52.892865 | 39.967581 | 6.025206 | 5.329537 | 53.886646 | 51.61112 | 53.18083 | 47.58503 | 39.400703 | 32.91701 | 5.8124666 | 6.12811 | 6.13504 | 5.4786496 | 5.189257 | 5.3207045 |
| ASMM10P032 | 0.00679 | 0.3337726 | 1.8625135 | up | NM_008318 | Ibsp | integrin binding sialoprotein | 88.328225 | 56.736052 | 6.735048 | 5.837797 | 113.25938 | 71.93032 | 79.79497 | 57.488026 | 53.09835 | 59.62178 | 6.87253 | 6.5982633 | 6.7343493 | 5.7514286 | 5.622062 | 6.1398993 |
| ASMM10P007 | 0.00680 | 0.3337726 | 1.3358852 | up | NM_026004 | Nt5c3 | 5'-nucleotidase, cytosolic III | 692.59442 | 623.0576 | 9.67436 | 9.256564 | 755.03516 | 656.8503 | 665.8978 | 630.47314 | 626.94305 | 611.7566 | 9.551844 | 9.719964 | 9.751271 | 9.22656 | 9.184034 | 9.359097 |
| ASMM10P055 | 0.00685 | 0.3349872 | 1.2751069 | up | NM_028130 | Zfp157 | zinc finger protein 157 | 315.18828 | 300.3552 | 8.565124 | 8.214506 | 374.46918 | 265.13403 | 305.96164 | 305.8949 | 326.76355 | 268.40714 | 8.559191 | 8.454033 | 8.682148 | 8.1811075 | 8.243692 | 8.218718 |
| ASMM10P020 | 0.00697 | 0.3350276 | 1.6383259 | up | NM_027133 | Lnp | limb and neural patterns | 18420.875 | 13073.4 | 14.301788 | 13.589566 | 16893.727 | 17182.01 | 21186.887 | 12974.498 | 13525.161 | 12720.541 | 14.107682 | 14.248745 | 14.548937 | 13.541182 | 13.536951 | 13.690564 |
| ASMM10P027 | 0.00703 | 0.3350276 | 1.7664375 | up | NM_025962 | Mmachc | methylmalonic aciduria cblC type, with homocystinuria | 6816.3682 | 4524.3088 | 12.908647 | 12.087804 | 6976.02 | 5990.525 | 7482.5596 | 4343.2275 | 4331.309 | 4898.39 | 12.800589 | 12.799667 | 13.125684 | 11.99366 | 11.945095 | 12.324657 |
| ASMM10P020 | 0.00703 | 0.3350276 | 1.3171919 | up | NM_001047 | Ttc21b | tetratricopeptide repeat domain 21B | 651.43543 | 594.95257 | 9.588881 | 9.191416 | 705.7193 | 621.8818 | 626.7052 | 595.8768 | 617.25055 | 571.73035 | 9.454136 | 9.6449375 | 9.66757 | 9.14322 | 9.162518 | 9.268509 |
| ASMM10P031 | 0.00705 | 0.3350276 | 1.9786561 | up | NM_138301 | Trpm2 | transient receptor potential cation channel, subfamily M, member 2 | 273.5684 | 169.28483 | 8.377454 | 7.392933 | 265.99774 | 278.16162 | 276.54584 | 181.74341 | 156.10854 | 170.00253 | 8.075192 | 8.521853 | 8.535317 | 7.426914 | 7.1686277 | 7.5832577 |
| ASMM10P053 | 0.00715 | 0.3350276 | 1.3260658 | up | NM_001177 | Irak1 | interleukin-1 receptor-associated kinase 1 | 6936.121 | 6177.5145 | 12.939464 | 12.532312 | 7003.9795 | 6685.366 | 7119.0176 | 6071.542 | 6859.9136 | 5601.088 | 12.810042 | 12.951906 | 13.056445 | 12.4732 | 12.600172 | 12.523564 |
| ASMM10P000 | 0.00717 | 0.3350276 | 1.6497648 | up | NM_001077 | Git2 | G protein-coupled receptor kinase-interactor 2 | 78.255658 | 58.433282 | 6.588375 | 5.866114 | 79.446724 | 80.62699 | 74.69326 | 69.858475 | 58.917267 | 46.524105 | 6.369509 | 6.7595425 | 6.636072 | 6.033341 | 5.763594 | 5.8014073 |
| ASMM10P015 | 0.00717 | 0.3350276 | 1.4738963 | up | NM_013921 | Prss30 | protease, serine 30 | 1795.5397 | 1462.7468 | 11.027401 | 10.467766 | 1903.1803 | 1576.8059 | 1906.633 | 1489.8573 | 1527.7646 | 1370.6185 | 10.88912 | 10.949186 | 11.243897 | 10.454713 | 10.452243 | 10.496342 |
| ASMM10P012 | 0.00720 | 0.3350276 | 1.7225916 | up | NM_001111 | Kctd9 | potassium channel tetramerisation domain containing 9 | 469.72658 | 330.48609 | 9.133427 | 8.348847 | 472.95312 | 423.4947 | 512.73193 | 323.3659 | 364.9386 | 303.15378 | 8.886593 | 9.114554 | 9.399135 | 8.257583 | 8.402886 | 8.386071 |
| ASMM10P016 | 0.00721 | 0.3350276 | 4.0293389 | up | NM_001039 | Acsbg2 | acyl-CoA synthetase bubblegum family member 2 | 46.81918 | 14.279837 | 5.839911 | 3.829368 | 47.186 | 52.567657 | 40.703884 | 12.98317 | 11.210741 | 18.6456 | 5.6242123 | 6.1547084 | 5.740813 | 3.5700662 | 3.37045 | 4.547588 |
| ASMM10P032 | 0.00721 | 0.3350276 | 2.0011388 | up | NM_011537 | Tbx5 | T-box 5 | 112.6912 | 69.117875 | 7.113766 | 6.112944 | 102.239 | 123.1413 | 112.6933 | 74.392624 | 74.71748 | 58.24352 | 6.7249775 | 7.3726287 | 7.2436905 | 6.1225266 | 6.1095333 | 6.106773 |
| ASMM10P010 | 0.00728 | 0.3350276 | 1.5131168 | up | NM_026725 | Dusp23 | dual specificity phosphatase 23 | 5073.1216 | 3914.9758 | 12.490657 | 11.893134 | 5031.0015 | 4856.213 | 5332.1504 | 3925.933 | 3877.4045 | 3941.5898 | 12.316403 | 12.49355 | 12.662018 | 11.86405 | 11.799673 | 12.015678 |
| ASMM10P041 | 0.00729 | 0.3350276 | 1.268507 | up | NM_028347 | Neil1 | nei endonuclease VIII-like 1 (E. coli) | 303.73818 | 288.36029 | 8.502151 | 8.159019 | 389.0077 | 261.79977 | 260.40707 | 295.7505 | 297.87024 | 271.46014 | 8.613321 | 8.438061 | 8.4550705 | 8.135335 | 8.106432 | 8.235291 |
| ASMM10P004 | 0.00733 | 0.3350276 | 1.3482562 | up | NM_001076 | Usp33 | ubiquitin specific peptidase 33 | 3314.4908 | 2896.9994 | 11.883149 | 11.452054 | 3558.443 | 3034.0718 | 3350.9575 | 2918.1465 | 2901.0613 | 2871.7905 | 11.791402 | 11.839272 | 12.018772 | 11.420532 | 11.384616 | 11.551014 |
| ASMM10P011 | 0.00734 | 0.3350276 | 1.9304252 | up | NM_013632 | Pnp | purine-nucleoside phosphorylase | 178.16786 | 114.95073 | 7.770869 | 6.82195 | 179.11882 | 184.39285 | 170.9919 | 103.7664 | 145.80962 | 95.276184 | 7.5111585 | 7.943649 | 7.8577995 | 6.609382 | 7.071562 | 6.784907 |
| ASMM10P052 | 0.00735 | 0.3350276 | 1.3341131 | up | NM_024473 | BC005537 | cDNA sequence BC005537 | 2214.7749 | 1978.7532 | 11.321395 | 10.905514 | 2400.04 | 2262.541 | 1981.7438 | 1919.1451 | 2116.0615 | 1901.0529 | 11.219369 | 11.457283 | 11.287533 | 10.823653 | 10.924257 | 10.968632 |
| ASMM10P044 | 0.00743 | 0.3350276 | 1.7425564 | up | NM_183319 | Xkrx | X Kell blood group precursor related X linked | 598.87939 | 413.14641 | 9.464872 | 8.663667 | 648.253 | 531.0444 | 617.34076 | 478.19775 | 366.47754 | 394.76395 | 9.320284 | 9.428184 | 9.646149 | 8.825819 | 8.411036 | 8.754146 |
| ASMM10P000 | 0.00744 | 0.3350276 | 1.452811 | up | NM_025421 | Acyp1 | acylphosphatase 1, erythrocyte (common) type | 1416.099 | 1162.618 | 10.688357 | 10.14951 | 1595.316 | 1278.253 | 1374.728 | 1131.3379 | 1137.0562 | 1219.4598 | 10.629254 | 10.652854 | 10.782963 | 10.069938 | 10.037238 | 10.341354 |
| ASMM10P023 | 0.00752 | 0.3350276 | 1.3379127 | up | NM_029530 | Lamp5 | lysosomal-associated membrane protein family, member 5 | 39.68232 | 35.747619 | 5.593445 | 5.173461 | 47.423264 | 32.87236 | 38.751335 | 41.670975 | 36.095154 | 29.476728 | 5.631755 | 5.4862704 | 5.662309 | 5.2854743 | 5.0631495 | 5.1717587 |
| ASMM10P015 | 0.00754 | 0.3350276 | 1.3126334 | up | NM_025483 | Senp7 | SUMO1/sentrin specific peptidase 7 | 683.72848 | 626.25362 | 9.655693 | 9.263229 | 740.15564 | 684.2745 | 626.7553 | 659.3334 | 636.5095 | 582.91797 | 9.523616 | 9.775282 | 9.668182 | 9.292477 | 9.204329 | 9.292882 |
| ASMM10P042 | 0.00758 | 0.3350276 | 1.4872557 | up | NM_019814 | Higd1a | HIG1 domain family, member 1A | 29580.236 | 23525.311 | 15.014061 | 14.441409 | 29387.86 | 27624.234 | 31728.615 | 23627.45 | 23214.322 | 23734.162 | 14.947822 | 14.918375 | 15.175987 | 14.397904 | 14.327823 | 14.598499 |
| ASMM10P008 | 0.00764 | 0.3350276 | 1.2803478 | up | NM_031257 | Plekha2 | pleckstrin homology domain-containing, family A (phosphoinositide binding specific) memb | 57.286482 | 54.714369 | 6.133231 | 5.776696 | 64.464424 | 56.57957 | 50.815453 | 61.27235 | 58.427807 | 44.44295 | 6.0730414 | 6.2613497 | 6.065303 | 5.8404574 | 5.751725 | 5.7379045 |
| ASMM10P004 | 0.00764 | 0.3350276 | 1.8768549 | up | NM_001039 | Cd44 | CD44 antigen | 87.709457 | 58.018043 | 6.753633 | 5.845316 | 84.62879 | 89.84359 | 88.65599 | 70.11607 | 61.58545 | 42.352608 | 6.457788 | 6.9158034 | 6.887307 | 6.038737 | 5.827397 | 5.669813 |
| ASMM10P038 | 0.00765 | 0.3350276 | 1.3232922 | up | NM_018811 | Abhd2 | abhydrolase domain containing 2 | 175.93077 | 161.8432 | 7.732219 | 7.328088 | 217.06795 | 168.35306 | 142.37129 | 160.78175 | 180.25697 | 144.49089 | 7.7908316 | 7.811362 | 7.5944643 | 7.2440314 | 7.384327 | 7.3559046 |
| ASMM10P007 | 0.00765 | 0.3350276 | 1.4108279 | up | NM_010199 | Fgf12 | fibroblast growth factor 12 | 780.94332 | 667.37309 | 9.839594 | 9.343052 | 909.38824 | 670.21857 | 763.22314 | 746.8244 | 710.1402 | 545.15466 | 9.8196 | 9.748317 | 9.950866 | 9.472547 | 9.360865 | 9.195745 |
| ASMM10P046 | 0.00770 | 0.3350276 | 1.706423 | up | NM_138949 | Zfp286 | zinc finger protein 286 | 2197.1301 | 1532.8224 | 11.31066 | 10.539684 | 2127.982 | 2139.1323 | 2324.2761 | 1602.9757 | 1486.2394 | 1509.252 | 11.042272 | 11.381444 | 11.508263 | 10.563686 | 10.40958 | 10.645787 |
| ASMM10P006 | 0.00770 | 0.3350276 | 1.6721403 | up | NM_008504 | Gzmm | granzyme M (lymphocyte met-ase 1) | 40.725602 | 29.514426 | 5.641438 | 4.899742 | 40.95099 | 44.103596 | 37.12222 | 31.394642 | 34.079376 | 23.06926 | 5.4198337 | 5.907352 | 5.597128 | 4.873867 | 4.987754 | 4.837605 |
| ASMM10P054 | 0.00771 | 0.3350276 | 1.3123779 | up | NM_001160 | Smek1 | SMEK homolog 1, suppressor of mek1 (Dictyostelium) | 49.750941 | 45.940757 | 5.926518 | 5.534334 | 56.9613 | 49.532906 | 42.758617 | 49.343033 | 48.96283 | 39.516407 | 5.8884125 | 6.0739627 | 5.8171773 | 5.527452 | 5.503797 | 5.571754 |
| ASMM10P018 | 0.00772 | 0.3350276 | 1.4018847 | up | NM_146089 | Haus1 | HAUS augmin-like complex, subunit 1 | 5672.9389 | 4743.9059 | 12.643355 | 12.155987 | 6053.784 | 4940.12 | 6024.9126 | 4606.123 | 5046.3696 | 4579.225 | 12.593333 | 12.521125 | 12.815607 | 12.075969 | 12.170946 | 12.221047 |
| ASMM10P002 | 0.00772 | 0.3350276 | 1.7963252 | up | NM_019834 | Git2 | G protein-coupled receptor kinase-interactor 2 | 141.11645 | 95.828037 | 7.428262 | 6.583214 | 156.18538 | 115.57641 | 151.58757 | 103.29759 | 90.21336 | 93.97316 | 7.3196135 | 7.279358 | 7.685816 | 6.6022844 | 6.382029 | 6.7653284 |

| | | | | | | | | | | | | | | | | | | | | | | | |
|---|---|---|---|---|---|---|---|---|---|---|---|---|---|---|---|---|---|---|---|---|---|---|---|
| ASMM10P035 | 0.00778 | 0.3353115 | 1.5928657 | up | NM_027059 | Smco2 | single-pass membrane protein with coiled-coil domains 2 | 180.70313 | 139.64769 | 7.779444 | 7.10782 | 200.15147 | 151.8422 | 190.11572 | 161.31944 | 147.26263 | 110.36099 | 7.6728005 | 7.661343 | 8.004189 | 7.248444 | 7.0880823 | 6.9869323 |
| ASMM10P026 | 0.00779 | 0.3353115 | 1.600514 | up | NM_025343 | Rmnd1 | required for meiotic nuclear division 1 homolog (S. cerevisiae) | 3333.9903 | 2457.383 | 11.894174 | 11.215639 | 3213.3137 | 3621.064 | 3167.5933 | 2575.6858 | 2505.8533 | 2290.61 | 11.63752 | 12.096787 | 11.948215 | 11.248507 | 11.171951 | 11.226458 |
| ASMM10P025 | 0.00779 | 0.3353115 | 2.4058577 | up | NM_133871 | Ifi44 | interferon-induced protein 44 | 575.65665 | 287.05259 | 9.404944 | 8.138393 | 495.2231 | 568.1963 | 663.55054 | 283.48996 | 345.47968 | 232.18813 | 8.946704 | 9.520545 | 9.747583 | 8.072174 | 8.323631 | 8.019373 |
| ASMM10P011 | 0.00784 | 0.3354865 | 1.2576063 | up | NM_009794 | Capn2 | calpain 2 | 45663.96 | 43784.141 | 15.669133 | 15.338452 | 44576.523 | 44812.227 | 47603.13 | 43574.183 | 48681.273 | 39129.266 | 15.591228 | 15.628379 | 15.787791 | 15.28689 | 15.387956 | 15.340511 |
| ASMM10P029 | 0.00801 | 0.3381178 | 2.2715723 | up | NM_025572 | 2610528J11I | RIKEN cDNA 2610528J11 gene | 200.31877 | 105.29606 | 7.90027 | 6.716579 | 241.58612 | 136.04555 | 223.32463 | 105.74019 | 102.4977 | 107.65029 | 7.939962 | 7.518885 | 8.241963 | 6.6348057 | 6.563312 | 6.9516187 |
| ASMM10P005 | 0.00803 | 0.3381178 | 4.155185 | up | NM_001164 | Ifih1 | interferon induced with helicase C domain 1 | 37.690914 | 10.474478 | 5.427864 | 3.372952 | 58.206337 | 29.023718 | 25.842688 | 7.3023157 | 13.163459 | 10.957659 | 5.919781 | 5.3118234 | 5.0519886 | 2.7212975 | 3.5981574 | 3.7994 |
| ASMM10P014 | 0.00808 | 0.3381178 | 1.3307126 | up | NM_172440 | Stxbp5l | syntaxin binding protein 5-like | 20497.378 | 17930.14 | 14.470063 | 14.057864 | 19937.256 | 20047.064 | 21507.814 | 17907.162 | 18242.213 | 17641.045 | 14.370598 | 14.465874 | 14.573717 | 14.01684 | 13.979941 | 14.176811 |
| ASMM10P015 | 0.00809 | 0.3381178 | 1.275557 | up | NM_009555 | Zfp40 | zinc finger protein 40 | 406.77028 | 385.6579 | 8.922134 | 8.571007 | 489.8549 | 350.03745 | 380.4185 | 370.16614 | 426.3471 | 360.46045 | 8.932877 | 8.843103 | 8.990423 | 8.455896 | 8.630695 | 8.62643 |
| ASMM10P019 | 0.00809 | 0.3381178 | 1.2877117 | up | NM_029682 | Stambpl1 | STAM binding protein like 1 | 85.168148 | 80.594978 | 6.702042 | 6.337233 | 98.30089 | 78.146545 | 79.05701 | 80.3887 | 84.9191 | 76.477135 | 6.671984 | 6.715404 | 6.7187386 | 6.2353587 | 6.2974815 | 6.4788575 |
| ASMM10P050 | 0.00810 | 0.3381178 | 1.3555156 | up | NM_001025 | Gm527 | predicted gene 527 | 377.99739 | 339.69414 | 8.826817 | 8.387976 | 413.66522 | 392.01556 | 328.3114 | 354.21793 | 362.46323 | 302.40125 | 8.69772 | 8.999288 | 8.783444 | 8.388689 | 8.391778 | 8.38346 |
| ASMM10P055 | 0.00812 | 0.3381178 | 1.3372405 | up | NM_010861 | Myl2 | myosin, light polypeptide 2, regulatory, cardiac, slow | 319.53989 | 289.78059 | 8.584326 | 8.165067 | 372.84787 | 314.65112 | 271.12067 | 276.9754 | 317.38437 | 274.982 | 8.552321 | 8.691839 | 8.508819 | 8.037327 | 8.202581 | 8.255294 |
| ASMM10P041 | 0.00814 | 0.3381178 | 1.4364516 | up | NM_023483 | 1110032A03 | RIKEN cDNA 1110032A03 gene | 356.92057 | 302.55818 | 8.745136 | 8.222627 | 412.62662 | 334.5323 | 323.6028 | 275.18546 | 330.69492 | 301.79416 | 8.693323 | 8.780691 | 8.761394 | 8.027064 | 8.260826 | 8.37999 |
| ASMM10P007 | 0.00816 | 0.3381178 | 1.443728 | up | NM_018764 | Pcdh7 | protocadherin 7 | 118.13304 | 99.194902 | 7.155999 | 6.6262 | 147.99529 | 95.51692 | 110.88692 | 94.84267 | 113.96143 | 88.78061 | 7.2425966 | 7.0045567 | 7.220843 | 6.4720674 | 6.717958 | 6.688574 |
| ASMM10P046 | 0.00816 | 0.3381178 | 2.4093183 | up | NM_009635 | Avil | advillin | 236.89213 | 120.61129 | 8.169907 | 6.901282 | 231.01648 | 269.31088 | 210.34904 | 124.54173 | 103.70112 | 133.59103 | 7.87715 | 8.476307 | 8.156265 | 6.8708677 | 6.5815725 | 7.2514067 |
| ASMM10P005 | 0.00823 | 0.3381393 | 1.9584897 | up | NM_001159 | Pdc | phosducin | 22.604896 | 13.760813 | 4.780894 | 3.811153 | 24.36651 | 20.118822 | 23.329355 | 14.961557 | 12.371211 | 13.94967 | 4.652299 | 4.7904515 | 4.899933 | 3.7794206 | 3.506296 | 4.1477423 |
| ASMM10P001 | 0.00824 | 0.3381393 | 1.3506264 | up | NM_013520 | Flt3l | FMS-like tyrosine kinase 3 ligand | 514.06601 | 453.93422 | 9.236051 | 8.802422 | 662.7488 | 409.79663 | 469.6526 | 482.60773 | 473.99722 | 405.19772 | 9.356933 | 9.06674 | 9.284479 | 8.838915 | 8.780507 | 8.787844 |
| ASMM10P012 | 0.00825 | 0.3381393 | 1.5042705 | up | NM_029561 | Ndfip2 | Nedd4 family interacting protein 2 | 15092.549 | 11759.275 | 14.025157 | 13.436093 | 13855.664 | 14920.136 | 16501.846 | 12446.855 | 12151.869 | 10679.101 | 13.806691 | 14.051109 | 14.21767 | 13.472699 | 13.401561 | 13.434018 |
| ASMM10P005 | 0.00827 | 0.3381393 | 1.3293553 | up | NM_001199 | Tti2 | TELO2 interacting protein 2 | 76.406613 | 70.315623 | 6.545115 | 6.134388 | 87.563 | 73.52971 | 68.12713 | 81.889366 | 69.6992 | 59.358303 | 6.504358 | 6.6318536 | 6.499133 | 6.2607336 | 6.009683 | 6.1327477 |
| ASMM10P055 | 0.00835 | 0.3381393 | 1.4014967 | up | NM_019800 | Rpl5 | ribosomal protein L5 | 1930.8307 | 1635.9303 | 11.119311 | 10.632343 | 2321.819 | 1690.587 | 1780.0862 | 1625.5979 | 1582.4037 | 1699.7892 | 11.170172 | 11.044163 | 11.143598 | 10.582758 | 10.502151 | 10.812119 |
| ASMM10P053 | 0.00837 | 0.3381393 | 1.3388447 | up | NM_025859 | Arl1 | ADP-ribosylation factor-like 1 | 6069.5052 | 5345.077 | 12.748206 | 12.327218 | 6185.453 | 6086.752 | 5936.3105 | 5435.106 | 5276.9375 | 5323.1875 | 12.623462 | 12.819338 | 12.801819 | 12.312797 | 12.23074 | 12.438116 |
| ASMM10P049 | 0.00838 | 0.3381393 | 1.3720653 | up | NM_021361 | Nova1 | neuro-oncological ventral antigen 1 | 202.87492 | 181.33047 | 7.949323 | 7.492974 | 229.16295 | 176.13048 | 203.33134 | 177.07016 | 204.155 | 162.76625 | 7.867516 | 7.87715 | 8.103303 | 7.3884435 | 7.570381 | 7.520097 |
| ASMM10P014 | 0.00839 | 0.3381393 | 1.3474747 | up | NM_198626 | Xxylt1 | xyloside xylosyltransferase 1 | 523.98847 | 468.89966 | 9.27715 | 8.846892 | 590.8546 | 536.6358 | 444.475 | 492.7922 | 510.97485 | 402.93192 | 9.185223 | 9.4421015 | 9.204127 | 8.871859 | 8.886593 | 8.782225 |
| ASMM10P026 | 0.00839 | 0.3381393 | 1.5932026 | up | NM_194343 | Trim45 | tripartite motif-containing 45 | 79.687533 | 61.396969 | 6.612076 | 5.940146 | 80.87956 | 84.98076 | 73.20228 | 70.694336 | 63.669765 | 49.826805 | 6.395184 | 6.8359566 | 6.605087 | 6.0501146 | 5.8747478 | 5.895576 |
| ASMM10P000 | 0.00846 | 0.3381393 | 1.9145724 | up | NM_001077 | Git2 | G protein-coupled receptor kinase-interactor 2 | 130.71968 | 83.153773 | 7.313012 | 6.37599 | 146.57489 | 103.30316 | 142.281 | 89.83394 | 76.378845 | 83.248535 | 7.2272644 | 7.118464 | 7.5933075 | 6.389894 | 6.1435738 | 6.5945015 |
| ASMM10P008 | 0.00849 | 0.3381393 | 1.3460581 | up | NM_054070 | Afg3l1 | AFG3(ATPase family gene 3)-like 1 (yeast) | 633.48051 | 561.55808 | 9.536683 | 9.107942 | 778.2746 | 534.30707 | 587.85986 | 549.9847 | 567.861 | 566.82855 | 9.596158 | 9.435235 | 9.578656 | 9.02906 | 9.041478 | 9.253289 |
| ASMM10P003 | 0.00850 | 0.3381393 | 1.2881513 | up | NM_001164 | Serpinb6a | serine (or cysteine) peptidase inhibitor, clade B, member 6a | 196.14496 | 186.72688 | 7.904871 | 7.539568 | 219.14052 | 178.25061 | 191.04375 | 185.4066 | 201.31764 | 173.4564 | 7.8052225 | 7.895543 | 8.013846 | 7.4555583 | 7.5505013 | 7.6126456 |
| ASMM10P004 | 0.00856 | 0.3381393 | 1.3366305 | up | NM_001081 | Rimbp2 | RIMS binding protein 2 | 944.64616 | 834.39853 | 10.093478 | 9.674878 | 1161.7893 | 924.31793 | 747.83124 | 859.7777 | 874.4546 | 768.9633 | 10.156094 | 10.201524 | 9.922817 | 9.664721 | 9.664431 | 9.695481 |
| ASMM10P008 | 0.00857 | 0.3381393 | 1.2647387 | up | NM_145544 | Rap1gds1 | RAP1, GTP-GDP dissociation stimulator 1 | 466.46586 | 446.70671 | 9.12087 | 8.782031 | 525.2306 | 460.81943 | 413.32056 | 479.73618 | 448.25238 | 412.13156 | 9.025659 | 9.227941 | 9.109011 | 8.830469 | 8.705096 | 8.810528 |
| ASMM10P005 | 0.00858 | 0.3381393 | 1.5885214 | up | NM_001195 | Nek10 | NIMA (never in mitosis gene a)- related kinase 10 | 34.46599 | 26.3699 | 5.39941 | 4.731725 | 34.92191 | 34.61297 | 33.86309 | 28.526548 | 31.35762 | 19.225533 | 5.183075 | 5.5627027 | 5.4524517 | 4.737068 | 4.8691034 | 4.5890045 |
| ASMM10P000 | 0.00859 | 0.3381393 | 1.3159692 | up | NM_177462 | Zmym6 | zinc finger, MYM-type 6 | 205.51852 | 192.84502 | 7.975104 | 7.578978 | 222.97363 | 190.94281 | 202.63911 | 202.07236 | 211.91959 | 164.54312 | 7.8284535 | 8.000811 | 8.096047 | 7.577486 | 7.62386 | 7.5355883 |
| ASMM10P006 | 0.00859 | 0.3381393 | 1.3371184 | up | NM_001243 | Cacfd1 | calcium channel flower domain containing 1 | 293.67421 | 267.04907 | 8.471367 | 8.05224 | 327.02975 | 293.42517 | 260.56772 | 284.08313 | 265.0976 | 251.96648 | 8.365775 | 8.592431 | 8.455896 | 8.075409 | 7.9438925 | 8.137419 |
| ASMM10P036 | 0.00864 | 0.3381393 | 1.9918316 | up | NM_030226 | 9130019O22 | RIKEN cDNA 9130019O22 gene | 486.47516 | 292.10858 | 9.164853 | 8.170757 | 486.97693 | 396.60248 | 575.84607 | 289.48376 | 342.77878 | 244.06319 | 8.925609 | 9.018795 | 9.550155 | 8.104944 | 8.312615 | 8.094713 |
| ASMM10P013 | 0.00866 | 0.3381393 | 1.8096671 | up | NM_001163 | Map3k12 | mitogen-activated protein kinase kinase kinase 12 | 2043.5492 | 1337.4412 | 11.196542 | 10.340818 | 2211.909 | 1655.846 | 2262.8926 | 1252.0382 | 1328.8805 | 1431.405 | 11.097257 | 11.021171 | 11.471199 | 10.209102 | 10.250419 | 10.562933 |
| ASMM10P032 | 0.00869 | 0.3381393 | 1.6441627 | up | NM_001005 | Zcwpw1 | zinc finger, CW type with PWWP domain 1 | 19.42274 | 13.894897 | 4.541699 | 3.824346 | 23.921848 | 18.994232 | 15.352139 | 13.848908 | 15.860971 | 11.974811 | 4.6239753 | 4.7059507 | 4.295172 | 3.6646967 | 3.8753536 | 3.9329886 |
| ASMM10P038 | 0.00873 | 0.3381393 | 1.5865808 | up | NM_147093 | Olfr558 | olfactory receptor 558 | 89.657915 | 69.050577 | 6.78197 | 6.116049 | 89.73467 | 96.331406 | 82.90767 | 72.589806 | 74.24176 | 60.320164 | 6.5397067 | 7.0161448 | 6.7900596 | 6.0903454 | 6.1011157 | 6.156687 |
| ASMM10P003 | 0.00881 | 0.3381393 | 1.2534687 | up | NM_028454 | Tm7sf2 | transmembrane 7 superfamily member 2 | 1147.0183 | 1092.1983 | 10.386205 | 10.060279 | 1384.3232 | 972.7866 | 1083.9452 | 1077.1858 | 1153.6055 | 1045.8037 | 10.423178 | 10.274994 | 10.460442 | 9.998352 | 10.052782 | 10.129702 |
| ASMM10P026 | 0.00884 | 0.3381393 | 2.2082717 | up | NM_001177 | Asph | aspartate-beta-hydroxylase | 1762.8331 | 929.09673 | 10.971226 | 9.828308 | 1763.5426 | 1309.8168 | 2215.14 | 918.3433 | 968.30914 | 900.63776 | 10.781472 | 10.693729 | 11.438477 | 9.762593 | 9.803879 | 9.918453 |
| ASMM10P013 | 0.00890 | 0.3381393 | 1.4967509 | up | NM_001271 | Kcns2 | K+ voltage-gated channel, subfamily S, 2 | 3614.43 | 2853.3985 | 12.012035 | 11.430201 | 3644.2534 | 3449.5 | 3749.5366 | 2912.7156 | 2757.4282 | 2890.0518 | 11.833179 | 12.026758 | 12.176169 | 11.418308 | 11.313102 | 11.559193 |
| ASMM10P028 | 0.00891 | 0.3381393 | 1.4660375 | up | NM_008538 | Marcks | myristoylated alanine rich protein kinase C substrate | 9175.3372 | 7310.7271 | 13.325925 | 12.774003 | 9634.6875 | 8568.761 | 9322.563 | 7499.4253 | 6889.534 | 7543.222 | 13.270088 | 13.263793 | 13.443895 | 12.765536 | 12.605516 | 12.950958 |
| ASMM10P047 | 0.00891 | 0.3381393 | 1.5226348 | up | NM_026561 | Nt5c3b | 5'-nucleotidase, cytosolic IIIB | 98.555903 | 78.819887 | 6.909631 | 6.303061 | 115.47869 | 83.03387 | 97.15515 | 83.70785 | 75.07511 | 77.6767 | 6.899478 | 6.804097 | 7.0253186 | 6.2918243 | 6.118089 | 6.4992704 |
| ASMM10P013 | 0.00892 | 0.3381393 | 1.517966 | up | NM_133227 | Nup155 | nucleoporin 155 | 95.88646 | 76.37813 | 6.862269 | 6.260129 | 112.65871 | 98.05771 | 76.94296 | 79.84019 | 77.28593 | 72.00827 | 6.8645306 | 7.0425367 | 6.6797385 | 6.2238083 | 6.1593275 | 6.3972516 |
| ASMM10P046 | 0.00895 | 0.3381393 | 1.3786198 | up | NM_010904 | Nefh | neurofilament, heavy polypeptide | 38301.951 | 33288.896 | 15.395773 | 14.932549 | 39495.21 | 35251.74 | 40158.902 | 30681.508 | 37374.285 | 31810.896 | 15.377383 | 15.294621 | 15.515316 | 14.786232 | 15.003913 | 15.007501 |
| ASMM10P018 | 0.00898 | 0.3381393 | 1.2701143 | up | NM_001081 | Dmxl1 | Dmx-like 1 | 383.54007 | 365.55871 | 8.841361 | 8.496403 | 457.84412 | 333.63873 | 359.13736 | 363.09705 | 374.8989 | 358.68018 | 8.838458 | 8.775524 | 8.910102 | 8.424199 | 8.44653 | 8.61848 |
| ASMM10P020 | 0.00899 | 0.3381393 | 1.484134 | up | NM_173443 | Vcpip1 | valosin containing protein (p97)/p47 complex interacting protein 1 | 1746.4781 | 1416.0392 | 10.990991 | 10.42137 | 1741.7527 | 1768.1102 | 1729.5714 | 1407.746 | 1467.5332 | 1372.8383 | 10.764073 | 11.103003 | 11.105898 | 10.374837 | 10.390345 | 10.498928 |
| ASMM10P021 | 0.00901 | 0.3381393 | 1.5443469 | up | NM_001128 | Ano3 | anoctamin 3 | 39448.365 | 30462.899 | 15.438142 | 14.811145 | 35962.074 | 40965.145 | 41417.875 | 28875.977 | 31445.959 | 31066.762 | 15.237135 | 15.518014 | 15.559276 | 14.699581 | 14.755796 | 14.978058 |
| ASMM10P006 | 0.00903 | 0.3381393 | 1.967601 | up | NM_001276 | Acy1 | aminoacylase 1 | 111.14876 | 70.801313 | 7.098359 | 6.121921 | 110.137 | 108.80148 | 114.5078 | 92.39044 | 66.2511 | 53.762398 | 6.8315053 | 7.1941075 | 7.2694635 | 6.4305854 | 5.9365363 | 5.9986415 |
| ASMM10P007 | 0.00903 | 0.3381393 | 1.3498359 | up | NM_025510 | Adprm | ADP-ribose/CDP-alcohol diphosphatase, manganese dependent | 1402.2677 | 1248.927 | 10.678652 | 10.245868 | 1511.5314 | 1377.8595 | 1317.4121 | 1339.9507 | 1208.9463 | 1197.884 | 10.547798 | 10.761713 | 10.726444 | 10.30458 | 10.120265 | 10.312758 |
| ASMM10P008 | 0.00903 | 0.3381393 | 1.5533705 | up | NM_026536 | Atp5s | ATP synthase, H+ transporting, mitochondrial F0 complex, subunit S | 416.54906 | 324.77975 | 8.958847 | 8.323445 | 488.87183 | 385.62656 | 375.1488 | 310.83978 | 312.6888 | 350.81067 | 8.930405 | 8.975818 | 8.970318 | 8.202116 | 8.178586 | 8.589633 |
| ASMM10P019 | 0.00904 | 0.3381393 | 1.2888884 | up | NM_028319 | Zfp518a | zinc finger protein 518A | 2114.5293 | 1961.244 | 11.254775 | 10.888648 | 2276.9673 | 2041.8197 | 2024.8009 | 1928.0321 | 2218.2126 | 1737.4874 | 11.143896 | 11.303769 | 11.316661 | 10.829307 | 10.995742 | 10.840895 |
| ASMM10P033 | 0.00905 | 0.3381393 | 1.3415079 | up | NM_008377 | Lrig1 | leucine-rich repeats and immunoglobulin-like domains 1 | 1293.3341 | 1165.2102 | 10.566653 | 10.142798 | 1389.6721 | 1321.1194 | 1169.2109 | 1212.1631 | 1284.4778 | 998.9896 | 10.430531 | 10.705743 | 10.563686 | 10.165121 | 10.200103 | 10.06317 |
| ASMM10P041 | 0.00906 | 0.3381393 | 1.7320395 | up | NM_023814 | Tbx18 | T-box18 | 376.80929 | 266.35218 | 8.826904 | 8.034432 | 407.5579 | 361.70233 | 361.16763 | 227.21988 | 298.0534 | 273.78326 | 8.6762085 | 8.887517 | 8.916985 | 7.747326 | 8.107952 | 8.248017 |
| ASMM10P007 | 0.00910 | 0.3384515 | 1.50153 | up | NM_026210 | Tvp23b | trans-golgi network vesicle protein 23B | 1729.1207 | 1386.5458 | 10.977315 | 10.390882 | 1727.8082 | 1735.6416 | 1723.9122 | 1365.408 | 1429.3694 | 1364.8601 | 10.751214 | 11.078674 | 11.102058 | 10.330217 | 10.35157 | 10.490859 |
| ASMM10P004 | 0.00912 | 0.3387874 | 1.3729285 | up | NM_001111 | Uba3 | ubiquitin-like modifier activating enzyme 3 | 272.44557 | 241.49413 | 8.362231 | 7.904974 | 306.5797 | 233.4093 | 277.3477 | 242.09825 | 258.70248 | 223.68166 | 8.270497 | 8.275197 | 8.5409975 | 7.8401403 | 7.9066586 | 7.968123 |
| ASMM10P034 | 0.00915 | 0.3389229 | 4.5986903 | up | NM_001007 | Noto | notochord homolog (Xenopus laevis) | 37.894818 | 9.605401 | 5.461335 | 3.260112 | 38.122658 | 51.557514 | 24.004282 | 6.830988 | 12.343548 | 9.641668 | 5.314577 | 6.126035 | 4.943394 | 2.647482 | 3.5020628 | 3.6307921 |
| ASMM10P052 | 0.00916 | 0.3389229 | 1.3295739 | up | NM_019499 | Mad2l1 | MAD2 mitotic arrest deficient-like 1 | 296.56373 | 271.43111 | 8.484988 | 8.074024 | 321.0749 | 297.59668 | 271.01962 | 284.48398 | 278.90344 | 250.9059 | 8.336238 | 8.610486 | 8.508239 | 8.077225 | 8.013195 | 8.131651 |
| ASMM10P024 | 0.00925 | 0.3391552 | 1.3221561 | up | NM_080793 | Setd7 | SET domain containing (lysine methyltransferase) 7 | 38505.08 | 34696.889 | 15.400791 | 14.997899 | 36365.383 | 39919.297 | 39230.56 | 33524.26 | 37788.008 | 32778.4 | 15.254581 | 15.462257 | 15.485536 | 14.912525 | 15.022463 | 15.058708 |
| ASMM10P027 | 0.00925 | 0.3391552 | 1.8649018 | up | NM_001004 | Ppp3r2 | protein phosphatase 3, regulatory subunit B, alpha isoform (calcineurin B, type II) | 105.98745 | 69.950545 | 7.019268 | 6.120168 | 106.58949 | 89.27864 | 122.09421 | 77.488014 | 79.38761 | 52.97601 | 6.7840014 | 6.907037 | 7.3667655 | 6.180723 | 6.2008486 | 5.9789333 |
| ASMM10P004 | 0.00926 | 0.3391552 | 1.3830515 | up | NM_001098 | Pcdh10 | protocadherin 10 | 1706.4272 | 1483.3437 | 10.958205 | 10.49035 | 1872.2803 | 1692.7777 | 1554.2235 | 1479.8573 | 1450.1808 | 1519.993 | 10.868475 | 11.046665 | 10.959476 | 10.444422 | 10.37148 | 10.655149 |
| ASMM10P006 | 0.00927 | 0.3391552 | 1.4657781 | up | NM_001271 | Adra1a | adrenergic receptor, alpha 1a | 67.398914 | 55.513426 | 6.345602 | 5.793935 | 83.32765 | 65.75628 | 53.112812 | 59.516563 | 63.484573 | 43.539143 | 6.4340615 | 6.4711585 | 6.131585 | 5.800915 | 5.871131 | 5.709759 |
| ASMM10P016 | 0.00932 | 0.3391552 | 1.3295531 | up | NM_001172 | Trim40 | tripartite motif-containing 40 | 53.812576 | 49.188791 | 6.046508 | 5.635566 | 58.131435 | 50.249348 | 53.056946 | 52.708443 | 50.242504 | 44.615425 | 5.9170413 | 6.093137 | 6.1293445 | 5.620617 | 5.5436397 | 5.742442 |
| ASMM10P034 | 0.00939 | 0.3391552 | 1.2806104 | up | NM_026168 | Ergic2 | ERGIC and golgi 2 | 1812.4823 | 1698.1225 | 11.042444 | 10.685612 | 1946.4802 | 1684.4038 | 1806.5629 | 1811.6477 | 1771.5791 | 1511.1407 | 10.920537 | 11.04109 | 11.165705 | 10.740295 | 10.668518 | 10.648024 |
| ASMM10P044 | 0.00942 | 0.3391552 | 1.3519052 | up | NM_012060 | Bcap31 | B cell receptor associated protein 31 | 15705.052 | 13584.762 | 14.084564 | 13.64957 | 15258.772 | 16159.553 | 15696.831 | 13214.084 | 14057.577 | 13482.626 | 13.955977 | 14.154931 | 14.142783 | 13.570165 | 13.595945 | 13.782599 |
| ASMM10P024 | 0.00943 | 0.3391552 | 1.3161047 | up | NM_021409 | Pard6b | par-6 family cell polarity regulator beta | 41.926342 | 38.776737 | 5.680578 | 5.284304 | 47.40491 | 40.65865 | 37.715466 | 45.425117 | 40.656635 | 30.24846 | 5.6312156 | 5.791215 | 5.6193047 | 5.411067 | 5.235841 | 5.2060046 |
| ASMM10P013 | 0.00946 | 0.3391552 | 1.5443984 | up | NM_177151 | Vps13b | vacuolar protein sorting 13B (yeast) | 325.80672 | 257.39449 | 8.62177 | 7.994725 | 350.7395 | 320.8256 | 305.85507 | 251.13829 | 254.5217 | 266.52347 | 8.463086 | 8.720272 | 8.681952 | 7.8923216 | 7.8822956 | 8.209558 |
| ASMM10P012 | 0.00947 | 0.3391552 | 1.370059 | up | NM_008866 | Lypla1 | lysophospholipase 1 | 17300.916 | 14811.752 | 14.225815 | 13.771577 | 16466.736 | 17027.03 | 18408.982 | 14490.202 | 15884.057 | 14060.997 | 14.07185 | 14.227746 | 14.377848 | 13.700881 | 13.773121 | 13.840728 |
| ASMM10P009 | 0.00951 | 0.3391552 | 1.9201874 | up | NM_175369 | Ccdc122 | coiled-coil domain containing 122 | 238.62008 | 153.85508 | 8.186385 | 7.245138 | 256.28226 | 230.09618 | 229.48181 | 127.4231 | 168.3879 | 165.75424 | 8.027064 | 8.252512 | 8.27958 | 6.9079046 | 7.282159 | 7.545351 |
| ASMM10P005 | 0.00955 | 0.3391552 | 1.5244692 | up | NM_001159 | Rbp4 | retinol binding protein 4, plasma | 192.97405 | 155.65872 | 7.879992 | 7.271685 | 215.73688 | 171.85527 | 191.33 | 171.7845 | 144.18848 | 151.00319 | 7.7830505 | 7.841744 | 8.015181 | 7.343387 | 7.055338 | 7.4163294 |
| ASMM10P034 | 0.00955 | 0.3391552 | 1.4852794 | up | NM_001168 | Phf14 | PHD finger protein 14 | 9586.8167 | 7497.7999 | 13.38485 | 12.814116 | 8900.372 | 9904.37 | 9955.708 | 7786.8423 | 7690.8174 | 7015.74 | 13.145327 | 13.483209 | 13.526014 | 12.823549 | 12.772706 | 12.846092 |
| ASMM10P002 | 0.00955 | 0.3391552 | 2.6191173 | up | NM_010899 | Nfatc2 | nuclear factor of activated T cells, cytoplasmic, calcineurin dependent 2 | 151.61446 | 69.241536 | 7.502765 | 6.113685 | 147.13785 | 110.56158 | 197.14394 | 80.35794 | 64.50807 | 62.858597 | 7.2337003 | 7.217441 | 8.057155 | 6.2345457 | 5.8969316 | 6.209577 |
| ASMM10P026 | 0.00958 | 0.3391552 | 1.5393026 | up | NM_172683 | Pogz | pogo transposable element with ZNF domain | 242.78397 | 192.88095 | 8.20709 | 7.584813 | 255.71974 | 215.66544 | 256.96674 | 200.198 | 193.85786 | 184.587 | 8.022979 | 8.165712 | 8.432578 | 7.565893 | 7.4944563 | 7.694089 |
| ASMM10P003 | 0.00961 | 0.3391552 | 1.477325 | up | NM_177266 | Gfm2 | G elongation factor, mitochondrial 2 | 35.995833 | 29.047619 | 5.451771 | 4.888784 | 41.874428 | 36.062622 | 30.05045 | 31.012835 | 29.580935 | 26.549088 | 5.448286 | 5.6216726 | 5.285355 | 4.855696 | 4.783933 | 5.026723 |
| ASMM10P050 | 0.00965 | 0.3391552 | 2.0322023 | up | NM_027438 | Pnma1 | paraneoplastic antigen MA1 | 4075.9792 | 2364.137 | 12.174008 | 11.150964 | 3999.357 | 3729.9946 | 4498.586 | 2398.2764 | 2010.4873 | 2683.6472 | 11.971499 | 12.129087 | 12.42144 | 11.144903 | 10.849144 | 11.458846 |
| ASMM10P047 | 0.00966 | 0.3391552 | 1.4579898 | up | NM_001111 | Stau2 | staufen (RNA binding protein) homolog 2 (Drosophila) | 1815.6776 | 1495.9914 | 11.047656 | 10.503676 | 1839.4667 | 1763.1708 | 1844.3954 | 1511.1357 | 1513.1089 | 1463.7296 | 10.841194 | 11.099581 | 11.202194 | 10.478224 | 10.43915 | 10.593653 |
| ASMM10P051 | 0.00969 | 0.3391552 | 1.5338871 | up | NM_020567 | Gmnn | geminin | 1696.4554 | 1329.4401 | 10.94875 | 10.331557 | 1802.5989 | 1533.7155 | 1753.0518 | 1334.4456 | 1265.5265 | 1388.3481 | 10.81738 | 10.905578 | 11.123291 | 10.300194 | 10.181651 | 10.512827 |
| ASMM10P049 | 0.00970 | 0.3391552 | 1.3263247 | up | NM_199196 | Suz12 | suppressor of zeste 12 homolog (Drosophila) | 310.12009 | 285.12144 | 8.549268 | 8.141834 | 349.84106 | 281.4634 | 299.0558 | 280.61258 | 295.4402 | 279.31155 | 8.459195 | 8.537439 | 8.651171 | 8.054495 | 8.095811 | 8.275197 |
| ASMM10P027 | 0.00972 | 0.3391552 | 1.5509978 | up | NM_001081 | Rab42 | RAB42, member RAS oncogene family | 1822.9424 | 1391.9394 | 11.03056 | 10.397363 | 2174.9553 | 1403.1147 | 1890.7573 | 1359.8854 | 1492.5355 | 1323.3972 | 11.07092 | 10.785721 | 11.235038 | 10.325522 | 10.417677 | 10.44889 |
| ASMM10P005 | 0.00972 | 0.3391552 | 1.4660842 | up | NM_001177 | Phka2 | phosphorylase kinase alpha 2 | 173.51667 | 144.99632 | 7.723964 | 7.171996 | 195.34602 | 180.06088 | 145.1431 | 158.34921 | 141.61316 | 135.0266 | 7.639657 | 7.911627 | 7.6206074 | 7.222494 | 7.028376 | 7.2651176 |
| ASMM10P009 | 0.00978 | 0.339228 | 1.5047408 | up | NM_176836 | Fam76b | family with sequence similarity 76, member B | 334.87551 | 271.38697 | 8.65509 | 8.065575 | 367.83548 | 280.3834 | 356.40765 | 282.54202 | 301.8893 | 229.7296 | 8.533107 | 8.532953 | 8.89921 | 8.064036 | 8.128366 | 8.004323 |
| ASMM10P033 | 0.00980 | 0.339228 | 1.4309914 | up | NM_023670 | Igf2bp3 | insulin-like growth factor 2 mRNA binding protein 3 | 302.97666 | 254.65443 | 8.499594 | 7.982579 | 374.438 | 237.67857 | 296.8134 | 254.80663 | 270.76892 | 238.38774 | 8.559027 | 8.298854 | 8.6409 | 7.9151764 | 7.9705625 | 8.061997 |
| ASMM10P031 | 0.00981 | 0.339228 | 1.2548135 | up | NM_023525 | Cad | carbamoyl-phosphate synthetase 2, aspartate transcarbamylase, and dihydroorotase | 11417.498 | 10719.185 | 13.631621 | 13.304148 | 11721.601 | 11309.729 | 11221.164 | 11139.741 | 10523.599 | 10494.214 | 13.557728 | 13.667485 | 13.66965 | 13.312899 | 13.195585 | 13.40396 |
| ASMM10P033 | 0.00983 | 0.339228 | 1.3607149 | up | NM_023160 | Cml1 | camello-like 1 | 5700.7619 | 4996.0534 | 12.66474 | 12.220375 | 5909.393 | 5614.9897 | 5577.903 | 4770.9673 | 5849.75 | 4367.443 | 12.560913 | 12.703863 | 12.729445 | 12.124437 | 12.380209 | 12.15648 |
| ASMM10P047 | 0.00985 | 0.339228 | 1.291361 | up | NM_172658 | Slco4c1 | solute carrier organic anion transporter family, member 4C1 | 786.66398 | 730.62317 | 9.85112 | 9.482228 | 896.2962 | 679.10675 | 784.589 | 733.3617 | 812.6061 | 645.9017 | 9.797721 | 9.766832 | 9.988808 | 9.449995 | 9.560415 | 9.436274 |
| ASMM10P054 | 0.00997 | 0.3427251 | 1.3629519 | up | NM_019912 | Ube2d2a | ubiquitin-conjugating enzyme E2D 2A | 6001.6737 | 5190.486 | 12.732662 | 12.285928 | 6142.625 | 6500.3887 | 5362.0073 | 5389.747 | 5277.904 | 4903.807 | 12.612045 | 12.915071 | 12.670871 | 12.298325 | 12.232108 | 12.32735 |
| ASMM10P034 | 0.01002 | 0.3427251 | 1.6057729 | up | NM_052993 | C1galt1 | core 1 synthase, glycoprotein-N-acetylgalactosamine 3-beta-galactosyltransferase, 1 | 118.24333 | 89.287212 | 7.162307 | 6.479039 | 144.38916 | 109.73321 | 100.60761 | 79.891815 | 97.65206 | 90.31776 | 7.2048254 | 7.206104 | 7.0759907 | 6.2245502 | 6.4968634 | 6.715703 |
| ASMM10P051 | 0.01007 | 0.3427251 | 2.0480473 | up | NM_001160 | Tmem196 | transmembrane protein 196 | 94.265308 | 55.159348 | 6.828272 | 5.794023 | 114.75957 | 69.315315 | 98.72104 | 55.390526 | 51.040184 | 59.047333 | 6.8905096 | 6.546934 | 7.0473723 | 5.6919537 | 5.564597 | 6.125518 |

| | | | | | | | | | | | | | | | | | | | | | | | |
|---|---|---|---|---|---|---|---|---|---|---|---|---|---|---|---|---|---|---|---|---|---|---|---|
| ASMM10P054 | 0.01010 | 0.3427251 | 2.2008504 | up | NM_172589 | Lhfpl2 | lipoma HMGIC fusion partner-like 2 | 49.764744 | 26.502737 | 5.88389 | 4.745829 | 65.203476 | 32.910583 | 51.180172 | 31.353102 | 23.62932 | 24.52579 | 6.0878334 | 5.487816 | 6.076021 | 4.8720703 | 4.4488688 | 4.916548 |
| ASMM10P014 | 0.01013 | 0.3427251 | 1.9285152 | up | NM_009435 | Tssk1 | testis-specific serine kinase 1 | 235.1737 | 148.80686 | 8.15395 | 7.206459 | 243.3718 | 191.2128 | 270.9365 | 140.00679 | 156.45467 | 149.95912 | 7.950845 | 8.003233 | 8.507772 | 7.0434084 | 7.172013 | 7.403957 |
| ASMM10P015 | 0.01013 | 0.3427251 | 1.4180439 | up | NM_146108 | Hibch | 3-hydroxyisobutyryl-Coenzyme A hydrolase | 1406.1244 | 1188.5773 | 10.681009 | 10.177107 | 1494.9122 | 1292.8533 | 1430.6077 | 1124.3298 | 1266.4897 | 1174.9125 | 10.533594 | 10.67133 | 10.838104 | 10.060342 | 10.183333 | 10.287646 |
| ASMM10P034 | 0.01015 | 0.3427251 | 1.4508882 | up | NM_026414 | Asprv1 | aspartic peptidase, retroviral-like 1 | 2246.7867 | 1834.6931 | 11.331975 | 10.795039 | 2445.1023 | 1937.1049 | 2358.153 | 1979.9564 | 1771.428 | 1752.695 | 11.245762 | 11.222021 | 11.528142 | 10.866746 | 10.668081 | 10.850289 |
| ASMM10P001 | 0.01020 | 0.3427251 | 1.3571835 | up | NM_023292 | Pus3 | pseudouridine synthase 3 | 714.33568 | 634.54273 | 9.72123 | 9.280615 | 745.0163 | 691.787 | 706.20374 | 645.31104 | 669.19415 | 589.123 | 9.5327635 | 9.793718 | 9.83721 | 9.257697 | 9.275307 | 9.30884 |
| ASMM10P000 | 0.01024 | 0.3427251 | 1.3803509 | up | NM_001013 | Ercc6l2 | excision repair cross-complementing rodent repair deficiency, complementation group 6 like | 102.14441 | 90.427503 | 6.96119 | 6.496155 | 118.87849 | 97.63251 | 89.922226 | 85.44747 | 99.8892 | 85.94584 | 6.9376855 | 7.0367827 | 6.909103 | 6.3199825 | 6.527192 | 6.6412916 |
| ASMM10P052 | 0.01026 | 0.3427251 | 1.4409249 | up | NM_001163 | Fam151b | family with sequence similarity 151, member B | 360.54616 | 304.61493 | 8.760281 | 8.233286 | 411.53864 | 344.1326 | 325.96725 | 305.29755 | 291.7674 | 316.77985 | 8.689331 | 8.820643 | 8.770869 | 8.1790495 | 8.07792 | 8.442888 |
| ASMM10P044 | 0.01030 | 0.3427251 | 1.4138176 | up | NM_177429 | Ofd1 | oral-facial-digital syndrome 1 gene homolog (human) | 197.23367 | 172.06133 | 7.917129 | 7.417533 | 205.94283 | 186.41235 | 199.34584 | 176.14735 | 191.17804 | 148.8586 | 7.7161837 | 7.9620967 | 8.073107 | 7.3808613 | 7.475326 | 7.396412 |
| ASMM10P024 | 0.01042 | 0.3427251 | 1.3984006 | up | NM_021896 | Gucy1a3 | guanylate cyclase 1, soluble, alpha 3 | 26445.445 | 22101.998 | 14.838288 | 14.354511 | 27895.3 | 23900.672 | 27540.363 | 23352.58 | 21261.455 | 21691.96 | 14.861311 | 14.715014 | 14.93854 | 14.383767 | 14.1971 | 14.482665 |
| ASMM10P008 | 0.01042 | 0.3427251 | 1.2563287 | up | NM_026842 | Ubqln1 | ubiquilin 1 | 697.29308 | 665.124 | 9.676958 | 9.347744 | 807.58685 | 685.6908 | 598.6016 | 644.10114 | 710.73425 | 640.5366 | 9.648261 | 9.778796 | 9.603818 | 9.255091 | 9.362462 | 9.42568 |
| ASMM10P025 | 0.01042 | 0.3427251 | 2.4972265 | up | NM_022427 | Gpr88 | G-protein coupled receptor 88 | 35778.757 | 16543.249 | 15.254033 | 13.933707 | 28726.03 | 49487.65 | 29122.592 | 18115.758 | 16626.57 | 14887.418 | 14.902094 | 15.817528 | 15.042478 | 14.034113 | 13.837893 | 13.929114 |
| ASMM10P021 | 0.01045 | 0.3427251 | 5.3418349 | up | NM_146450 | Olfr1314 | olfactory receptor 1314 | 31.798199 | 7.154363 | 5.257483 | 2.840147 | 39.23885 | 23.040272 | 33.115475 | 4.9999995 | 4.9999995 | 11.463089 | 5.3582597 | 4.990063 | 5.424125 | 2.3250442 | 2.3265402 | 3.8688571 |
| ASMM10P006 | 0.01048 | 0.3427251 | 1.822763 | up | NM_001204 | Myadml2 | myeloid-associated differentiation marker-like 2 | 293.3682 | 194.06203 | 8.456178 | 7.590051 | 350.84476 | 230.45778 | 298.80206 | 184.22832 | 185.68956 | 212.2682 | 8.463548 | 8.254928 | 8.650058 | 7.4457273 | 7.428978 | 7.8954477 |
| ASMM10P036 | 0.01049 | 0.3427251 | 1.8908919 | up | NM_001024 | Mctp2 | multiple C2 domains, transmembrane 2 | 364.6992 | 233.28018 | 8.776631 | 7.857564 | 352.31915 | 327.68176 | 414.09668 | 233.43646 | 234.48286 | 231.92122 | 8.470121 | 8.748219 | 9.111553 | 7.7873826 | 7.7670817 | 8.018228 |
| ASMM10P012 | 0.01050 | 0.3427251 | 1.9521474 | up | NM_001034 | Zbtbd6 | kelch repeat and BTB (POZ) domain containing 6 | 456.25119 | 282.51739 | 9.092424 | 8.127362 | 428.75677 | 442.44003 | 497.55676 | 277.83157 | 275.02435 | 294.69626 | 8.746828 | 9.171849 | 9.358596 | 8.042128 | 7.991592 | 8.348367 |
| ASMM10P012 | 0.01057 | 0.3427251 | 1.6060806 | up | NM_011541 | Tcea1 | transcription elongation factor A (SII) 1 | 1372.1796 | 1021.3271 | 10.644808 | 9.961264 | 1325.693 | 1446.0598 | 1344.7859 | 989.3109 | 1116.2821 | 958.38837 | 10.359431 | 10.822187 | 10.752807 | 9.872004 | 10.011168 | 10.00062 |
| ASMM10P041 | 0.01057 | 0.3427251 | 1.61866 | up | NM_016919 | Col5a3 | collagen, type V, alpha 3 | 443.17662 | 330.19665 | 9.042537 | 8.347737 | 472.2015 | 368.95786 | 488.3705 | 326.19315 | 346.62366 | 317.77313 | 8.884235 | 8.912826 | 9.330549 | 8.268538 | 8.328674 | 8.445998 |
| ASMM10P012 | 0.01057 | 0.3427251 | 1.3087776 | up | NM_026401 | Mrp63 | mitochondrial ribosomal protein 63 | 525.9995 | 485.81782 | 9.290719 | 8.902499 | 566.76556 | 506.89975 | 504.3332 | 502.96982 | 497.41275 | 457.0709 | 9.12977 | 9.367271 | 9.375115 | 8.903595 | 8.851099 | 8.952802 |
| ASMM10P033 | 0.01059 | 0.3427251 | 1.3518287 | up | NM_175414 | Tspan9 | tetraspanin 9 | 16981.185 | 14749.12 | 14.201487 | 13.766575 | 17837.164 | 16279.335 | 16827.055 | 13453.229 | 15953.876 | 14840.255 | 14.189349 | 14.167316 | 14.247798 | 13.59899 | 13.780499 | 13.920237 |
| ASMM10P043 | 0.01060 | 0.3427251 | 1.3380765 | up | NM_001177 | Rpgr | retinitis pigmentosa GTPase regulator | 90.400532 | 83.725865 | 6.789649 | 6.369488 | 103.14659 | 85.10809 | 82.946915 | 87.556175 | 100.08401 | 63.53741 | 6.7392035 | 6.8391695 | 6.790574 | 6.3531938 | 6.5297403 | 6.225531 |
| ASMM10P036 | 0.01067 | 0.3427251 | 1.4643278 | up | NM_009943 | Cox6a2 | cytochrome c oxidase subunit VIa polypeptide 2 | 12989.298 | 10406.668 | 13.815235 | 13.264997 | 13976.123 | 10828.522 | 14163.248 | 10014.498 | 11677.571 | 9527.936 | 13.820313 | 13.618794 | 14.006599 | 13.177245 | 13.341826 | 13.275919 |
| ASMM10P008 | 0.01067 | 0.3427251 | 1.3461078 | up | NM_139297 | Ugp2 | UDP-glucose pyrophosphorylase 2 | 1911.7928 | 1709.9121 | 11.117124 | 10.68833 | 2074.1025 | 1772.9169 | 1888.3591 | 1594.8679 | 1937.2444 | 1597.624 | 11.010903 | 11.106177 | 11.234293 | 10.554514 | 10.787205 | 10.723272 |
| ASMM10P015 | 0.01068 | 0.3427251 | 7.5598536 | up | NM_001082 | Gm5483 | predicted gene 5483 | 156.47027 | 21.110616 | 7.340451 | 4.422093 | 86.40466 | 290.00952 | 92.99662 | 19.48727 | 24.94219 | 18.902388 | 6.4845743 | 8.577694 | 6.9590855 | 4.1708145 | 4.5312934 | 4.564171 |
| ASMM10P007 | 0.01069 | 0.3427251 | 1.2782735 | up | NM_011274 | Uri1 | URI1, prefoldin-like chaperone | 123.21819 | 118.42165 | 7.238234 | 6.884037 | 134.82079 | 119.40546 | 115.42832 | 121.59222 | 124.81912 | 108.85361 | 7.107948 | 7.3266344 | 7.2801194 | 6.8380995 | 6.8470507 | 6.966962 |
| ASMM10P006 | 0.01070 | 0.3427251 | 1.2944137 | up | NM_001252 | Tle2 | transducin-like enhancer of split 2, homolog of Drosophila E(spl) | 405.96722 | 382.78836 | 8.924958 | 8.552659 | 473.0782 | 397.03317 | 347.79028 | 414.76813 | 422.3091 | 311.28784 | 8.88706 | 9.020949 | 8.866865 | 8.622489 | 8.616362 | 8.4191265 |
| ASMM10P007 | 0.01072 | 0.3427251 | 1.3910645 | up | NM_018784 | St3gal6 | ST3 beta-galactoside alpha-2,3-sialyltransferase 6 | 220.48961 | 192.37974 | 8.056809 | 7.58062 | 272.68402 | 195.67274 | 193.11208 | 182.96474 | 198.4386 | 195.73587 | 8.10988 | 8.032348 | 8.0282 | 7.4356747 | 7.528422 | 7.7777634 |
| ASMM10P037 | 0.01075 | 0.3427251 | 1.8186673 | up | NM_025533 | Nosip | nitric oxide synthase interacting protein | 4708.528 | 3033.7576 | 12.378151 | 11.515269 | 4234.1323 | 4528.3916 | 5363.06 | 3102.0571 | 2901.4988 | 3097.717 | 12.067732 | 12.394819 | 12.671902 | 11.505035 | 11.385072 | 11.655701 |
| ASMM10P008 | 0.01078 | 0.3427251 | 1.3706737 | up | NM_029398 | Tmem14a | transmembrane protein 14A | 2176.4784 | 1882.5485 | 11.288167 | 10.833282 | 2442.5806 | 1862.447 | 2224.4077 | 1820.4352 | 1958.7617 | 1868.4487 | 11.244742 | 11.171653 | 11.448105 | 10.748405 | 10.807421 | 10.944019 |
| ASMM10P009 | 0.01083 | 0.3427251 | 1.5235335 | up | NM_175098 | Ccdc126 | coiled-coil domain containing 126 | 190.70213 | 153.61393 | 7.86313 | 7.255709 | 208.9244 | 168.11418 | 195.06781 | 160.05939 | 148.0282 | 152.75421 | 7.7363715 | 7.8094506 | 8.043568 | 7.238486 | 7.094927 | 7.4337134 |
| ASMM10P006 | 0.01086 | 0.3427251 | 1.3177943 | up | NM_008996 | Rab1 | RAB1, member RAS oncogene family | 7616.2362 | 6748.7929 | 13.064166 | 12.666041 | 8229.935 | 7618.5757 | 7000.1978 | 6806.3296 | 6520.103 | 6919.946 | 13.048001 | 13.118188 | 13.02631 | 12.637011 | 12.537066 | 12.824047 |
| ASMM10P048 | 0.01089 | 0.3427251 | 2.3273739 | up | NM_030679 | Myh1 | myosin, heavy polypeptide 1, skeletal muscle, adult | 103.76657 | 52.218751 | 6.9332 | 5.714497 | 129.5218 | 64.708786 | 117.06912 | 49.21402 | 59.319588 | 48.122646 | 7.049591 | 6.448921 | 7.301087 | 5.522324 | 5.773385 | 5.847781 |
| ASMM10P031 | 0.01089 | 0.3427251 | 1.3632459 | up | NM_007832 | Dck | deoxycytidine kinase | 224.99716 | 201.2301 | 8.094367 | 7.647321 | 268.67572 | 189.16544 | 217.15033 | 198.16666 | 207.03801 | 198.48563 | 8.090049 | 7.9865813 | 8.20647 | 7.551925 | 7.5911627 | 7.798875 |
| ASMM10P022 | 0.01089 | 0.3427251 | 1.3136758 | up | NM_010726 | Phyh | phytanoyl-CoA hydroxylase | 4569.2068 | 4079.232 | 12.340698 | 11.947089 | 5092.147 | 4211.1875 | 4404.286 | 3894.734 | 4115.212 | 4227.75 | 12.338333 | 12.287672 | 12.396089 | 11.850815 | 11.880595 | 12.109856 |
| ASMM10P038 | 0.01092 | 0.3427251 | 1.4023243 | up | NM_146190 | Tubgcp5 | tubulin, gamma complex associated protein 5 | 2687.0038 | 2275.2095 | 11.586771 | 11.098951 | 2760.5688 | 2500.3003 | 2800.1423 | 2418.4023 | 2415.3079 | 1991.9183 | 11.414064 | 11.577255 | 11.768994 | 11.155006 | 11.109901 | 11.031946 |
| ASMM10P046 | 0.01092 | 0.3427251 | 1.3420884 | up | NM_001277 | Grip1 | glutamate receptor interacting protein 1 | 818.2377 | 726.98447 | 9.904003 | 9.479523 | 906.3887 | 845.65735 | 702.66705 | 749.4371 | 730.0253 | 701.491 | 9.813129 | 10.070452 | 9.828427 | 9.478194 | 9.402751 | 9.557624 |
| ASMM10P006 | 0.01095 | 0.3427251 | 1.4328104 | up | NM_007464 | Birc3 | baculoviral IAP repeat-containing 3 | 106.63761 | 92.040781 | 7.033762 | 6.514914 | 110.84251 | 110.42828 | 98.64204 | 99.04964 | 102.85667 | 74.21603 | 6.8398323 | 7.215576 | 7.0458775 | 6.5380535 | 6.5691977 | 6.4374914 |
| ASMM10P011 | 0.01099 | 0.3427251 | 1.8781554 | up | NM_001037 | Gm6878 | predicted gene 6878 | 163.04445 | 105.7586 | 7.630371 | 6.721054 | 186.5234 | 127.68415 | 174.9258 | 104.7972 | 99.86982 | 112.60878 | 7.5721793 | 7.4264593 | 7.8924737 | 6.622115 | 6.526819 | 7.014229 |
| ASMM10P012 | 0.01099 | 0.3427251 | 1.695165 | up | NM_026346 | Fbxo32 | F-box protein 32 | 8004.1953 | 5595.0724 | 13.140146 | 12.37872 | 7749.495 | 7097.483 | 9165.608 | 5406.0356 | 6682.212 | 4696.9697 | 12.970642 | 13.033848 | 13.415947 | 12.301069 | 12.572733 | 12.262358 |
| ASMM10P039 | 0.01104 | 0.3427251 | 1.2988324 | up | NM_178591 | Nrg1 | neuregulin 1 | 497.21418 | 459.55795 | 9.197686 | 8.820471 | 601.5087 | 489.28952 | 400.84433 | 504.02423 | 474.42166 | 400.22797 | 9.210244 | 9.316348 | 9.066467 | 8.906708 | 8.781621 | 8.773084 |
| ASMM10P005 | 0.01104 | 0.3427251 | 1.3465899 | up | NM_001166 | Zfml | zinc finger, matrin-like | 306.56123 | 277.01991 | 8.528645 | 8.099335 | 350.16125 | 301.36356 | 268.15887 | 264.1474 | 293.54935 | 273.36298 | 8.460345 | 8.630263 | 8.495328 | 7.966946 | 8.08533 | 8.2457285 |
| ASMM10P014 | 0.01109 | 0.3427251 | 1.4274318 | up | NM_001256 | Gm933 | predicted gene 933 | 85.701348 | 74.438425 | 6.719555 | 6.206134 | 90.362305 | 83.63528 | 83.10646 | 79.84818 | 86.959755 | 56.50734 | 6.550082 | 6.816291 | 6.7922935 | 6.2239847 | 6.330163 | 6.0642533 |
| ASMM10P031 | 0.01112 | 0.3427251 | 1.507275 | up | NM_009933 | Col6a1 | collagen, type VI, alpha 1 | 11730.183 | 9073.1926 | 13.660043 | 13.0681 | 12708.38 | 9704.673 | 12777.495 | 9425.332 | 10313.462 | 7480.7837 | 13.678284 | 13.454915 | 13.84693 | 13.100376 | 13.167183 | 12.936742 |
| ASMM10P049 | 0.01114 | 0.3427251 | 2.7530127 | up | NM_025766 | Fam187a | family with sequence similarity 187, member A | 57.636308 | 25.449254 | 6.128955 | 4.667944 | 68.198364 | 44.189377 | 60.521183 | 21.450605 | 23.20413 | 31.693026 | 6.1511507 | 5.9094887 | 6.326226 | 4.3064103 | 4.42468 | 5.2727413 |
| ASMM10P027 | 0.01115 | 0.3427251 | 1.2642267 | up | NM_001045 | Akna | AT-hook transcription factor | 29.427753 | 28.016062 | 5.169865 | 4.83161 | 32.359898 | 25.834122 | 30.08924 | 28.801252 | 32.056644 | 23.19029 | 5.076238 | 5.1457305 | 5.287626 | 4.7517304 | 4.9007554 | 4.8423433 |
| ASMM10P030 | 0.01116 | 0.3427251 | 4.1848841 | up | NM_021274 | Cxcl10 | chemokine (C-X-C motif) ligand 10 | 710.86071 | 206.48789 | 9.667171 | 7.601983 | 850.4375 | 455.86633 | 826.2783 | 120.70371 | 254.38133 | 244.37863 | 9.721472 | 9.21638 | 10.063661 | 6.827309 | 7.881901 | 8.09674 |
| ASMM10P009 | 0.01120 | 0.3427251 | 1.314358 | up | NM_172946 | Krt222 | keratin 222 | 79.52005 | 74.734597 | 6.606398 | 6.21204 | 88.29811 | 76.33319 | 73.92885 | 80.12863 | 87.00426 | 57.0709 | 6.5168076 | 6.682492 | 6.6198945 | 6.2288384 | 6.33126 | 6.076021 |
| ASMM10P049 | 0.01120 | 0.3427251 | 1.9265309 | up | NM_025727 | Klhl10 | kelch-like 10 | 232.10995 | 147.99831 | 8.145367 | 7.199361 | 209.75607 | 260.21194 | 226.36183 | 166.3241 | 153.39731 | 124.27351 | 7.743645 | 8.43009 | 8.262365 | 7.297012 | 7.146242 | 7.15483 |
| ASMM10P030 | 0.01122 | 0.3427251 | 1.4778768 | up | NM_031869 | Prkab1 | protein kinase, AMP-activated, beta 1 non-catalytic subunit | 16604.712 | 13137.57 | 14.163943 | 13.600417 | 17489.08 | 13949.495 | 18375.56 | 13610.005 | 13572.675 | 12230.029 | 14.166205 | 13.950502 | 14.375122 | 13.614338 | 13.540442 | 13.646471 |
| ASMM10P013 | 0.01122 | 0.3427251 | 1.4754759 | up | NM_027122 | Mfsd3 | major facilitator superfamily domain containing 3 | 1154.628 | 931.4858 | 10.395396 | 9.834216 | 1344.7211 | 1174.4745 | 944.68835 | 928.7835 | 894.0062 | 971.6677 | 10.382798 | 10.542448 | 10.260943 | 9.781123 | 9.70074 | 10.020785 |
| ASMM10P007 | 0.01124 | 0.3427386 | 1.4174833 | up | NM_026250 | Gin1 | gypsy retrotransposon integrase 1 | 147.12569 | 127.98721 | 7.499496 | 6.996164 | 155.5419 | 138.74129 | 147.09389 | 132.9579 | 130.47278 | 120.53096 | 7.314233 | 7.544598 | 7.639657 | 6.9688067 | 6.9108295 | 7.1088567 |
| ASMM10P014 | 0.01127 | 0.3427419 | 1.2726819 | up | NM_177357 | Kalrn | kalirin, RhoGEF kinase | 14552.608 | 13459.144 | 13.98033 | 13.632458 | 15356.683 | 13074.975 | 15226.167 | 13111.551 | 14930.031 | 12335.85 | 13.966332 | 13.871449 | 14.103208 | 13.555245 | 13.688677 | 13.653451 |
| ASMM10P034 | 0.01143 | 0.343714 | 1.4633334 | up | NM_001109 | Cntn4 | contactin 4 | 11365.223 | 9001.4017 | 13.623588 | 13.074329 | 10797.894 | 10500.736 | 12797.038 | 9289.118 | 9188.297 | 8526.79 | 13.443296 | 13.578546 | 13.848922 | 13.079804 | 13.012727 | 13.130457 |
| ASMM10P030 | 0.01153 | 0.343714 | 1.3209755 | up | NM_011890 | Sgcb | sarcoglycan, beta (dystrophin-associated glycoprotein) | 18017.437 | 15990.716 | 14.282511 | 13.880907 | 17622.432 | 17049.635 | 19380.244 | 15700.048 | 17221.125 | 15050.974 | 14.17508 | 14.230017 | 14.442435 | 13.80963 | 13.884319 | 13.948772 |
| ASMM10P040 | 0.01164 | 0.343714 | 2.3111426 | up | NM_019938 | Pmfbp1 | polyamine modulated factor 1 binding protein 1 | 13.457247 | 6.781441 | 4.017287 | 2.808681 | 11.734931 | 15.111639 | 13.525172 | 8.802742 | 6.004568 | 5.537012 | 3.5573802 | 4.3820662 | 4.112415 | 2.9857993 | 2.5607247 | 2.8795185 |
| ASMM10P002 | 0.01164 | 0.343714 | 1.3296351 | up | NM_027478 | Tmem230 | transmembrane protein 230 | 609.69556 | 551.55273 | 9.492891 | 9.08186 | 679.59937 | 550.5536 | 598.9337 | 562.14594 | 541.7501 | 550.76215 | 9.392183 | 9.481126 | 9.605363 | 9.061119 | 8.972552 | 9.21191 |
| ASMM10P018 | 0.01167 | 0.343714 | 1.5663884 | up | NM_053129 | Pcdhb4 | protocadherin beta 4 | 310.34922 | 241.01573 | 8.549491 | 7.902049 | 352.61157 | 272.86435 | 305.57175 | 233.86252 | 231.8914 | 257.29327 | 8.471155 | 8.496289 | 8.68103 | 7.790614 | 7.751797 | 8.163737 |
| ASMM10P007 | 0.01171 | 0.343714 | 1.8516387 | up | NM_010954 | Ncam2 | neural cell adhesion molecule 2 | 158.30036 | 105.84725 | 7.598725 | 6.709922 | 147.09467 | 152.34872 | 175.4577 | 115.87272 | 118.02701 | 83.64202 | 7.2331142 | 7.666091 | 7.896969 | 6.7638917 | 6.765222 | 6.6006527 |
| ASMM10P038 | 0.01172 | 0.343714 | 1.572148 | up | NM_145151 | Crebzf | CREB/ATF bZIP transcription factor | 1243.8867 | 950.32197 | 10.500097 | 9.84736 | 1399.4641 | 1018.3608 | 1313.8352 | 974.5547 | 1112.7369 | 763.6743 | 10.438995 | 10.338608 | 10.722687 | 9.851313 | 10.007926 | 9.68284 |
| ASMM10P011 | 0.01173 | 0.343714 | 1.5194742 | up | NM_028039 | Esco2 | establishment of cohesion 1 homolog 2 (S. cerevisiae) | 26.094368 | 20.701589 | 4.994845 | 4.391272 | 26.369965 | 23.982498 | 27.930641 | 24.80138 | 20.872108 | 16.43128 | 4.7690516 | 5.045317 | 5.170165 | 4.518795 | 4.2802534 | 4.3747687 |
| ASMM10P040 | 0.01175 | 0.343714 | 1.3421298 | up | NM_011240 | Ranbp2 | RAN binding protein 2 | 9440.807 | 8227.4223 | 13.366329 | 12.941805 | 9125.165 | 9402.134 | 9795.122 | 8609.356 | 8591.766 | 7481.145 | 13.183679 | 13.409159 | 13.506149 | 12.969161 | 12.919093 | 12.937161 |
| ASMM10P004 | 0.01178 | 0.343714 | 1.2671932 | up | NM_001102 | Aplp2 | amyloid beta (A4) precursor-like protein 2 | 1605.7134 | 1536.2199 | 10.874325 | 10.532689 | 1743.393 | 1578.485 | 1495.2621 | 1562.8374 | 1726.6185 | 1319.2039 | 10.764925 | 10.950473 | 10.907578 | 10.525224 | 10.62962 | 10.443222 |
| ASMM10P044 | 0.01178 | 0.343714 | 1.6690054 | up | NM_019768 | Morf4l2 | mortality factor 4 like 2 | 19.273289 | 14.009345 | 4.549914 | 3.810925 | 19.548141 | 19.039576 | 19.23215 | 17.909918 | 13.65304 | 10.465078 | 4.3190994 | 4.710459 | 4.6201825 | 4.0456853 | 3.6524842 | 3.7346056 |
| ASMM10P027 | 0.01183 | 0.343714 | 1.3336437 | up | NM_023223 | Cdc20 | cell division cycle 20 | 946.69168 | 853.70095 | 10.120336 | 9.704962 | 997.55963 | 912.3796 | 930.1358 | 880.7554 | 919.6342 | 760.71326 | 9.936719 | 10.186639 | 10.237649 | 9.699325 | 9.738737 | 9.676825 |
| ASMM10P038 | 0.01187 | 0.343714 | 4.7740229 | up | NM_146925 | Olfr481 | olfactory receptor 481 | 27.962244 | 6.91862 | 5.063868 | 2.808663 | 33.52786 | 29.577375 | 20.781498 | 4.9999995 | 4.9999995 | 10.75586 | 5.125467 | 5.337788 | 4.7283497 | 2.3250442 | 2.3265402 | 3.7744038 |
| ASMM10P000 | 0.01194 | 0.343714 | 2.6078777 | up | NM_001163 | Tnnt3 | troponin T3, skeletal, fast | 41.4207 | 19.400027 | 5.666361 | 4.283485 | 44.55454 | 41.7686 | 37.93896 | 17.147343 | 16.95479 | 24.097948 | 5.5423465 | 5.8292317 | 5.6275053 | 3.9779758 | 3.9798481 | 4.892631 |
| ASMM10P045 | 0.01194 | 0.343714 | 1.3554842 | up | NM_053201 | Magee1 | melanoma antigen, family E, 1 | 95670.94 | 92456.617 | 16.858636 | 16.419828 | 93050.59 | 94395.48 | 99566.75 | 96592.38 | 93836.92 | 86940.55 | 16.770472 | 16.822634 | 16.982803 | 16.403906 | 16.294453 | 16.561125 |
| ASMM10P055 | 0.01195 | 0.343714 | 1.3337701 | up | NM_001077 | Slc1a2 | solute carrier family 1 (glial high affinity glutamate transporter), member 2 | 16664.316 | 14735.112 | 14.174293 | 13.758783 | 16554.43 | 16684.34 | 16754.178 | 13462.855 | 17107.21 | 13635.272 | 14.081491 | 14.202172 | 14.239217 | 13.599884 | 13.875091 | 13.801375 |
| ASMM10P040 | 0.01197 | 0.343714 | 1.5643336 | up | NM_020561 | Smpdl3a | sphingomyelin phosphodiesterase, acid-like 3A | 1742.2831 | 1345.7493 | 10.98931 | 10.343762 | 1890.6123 | 1632.792 | 1703.445 | 1138.3873 | 1502.2566 | 1396.6039 | 10.882587 | 10.998822 | 11.086521 | 10.07847 | 10.429232 | 10.523583 |
| ASMM10P007 | 0.01197 | 0.343714 | 1.7789809 | up | NM_016900 | Cav2 | caveolin 2 | 595.10062 | 399.65011 | 9.452664 | 8.621613 | 591.78687 | 527.3032 | 666.2118 | 366.80838 | 433.20566 | 398.9363 | 9.187222 | 9.41887 | 9.751899 | 8.440197 | 8.655076 | 8.769565 |
| ASMM10P005 | 0.01198 | 0.343714 | 1.2855217 | up | NM_001167 | Fam45a | family with sequence similarity 45, member A | 340.3457 | 322.98559 | 8.679847 | 8.317493 | 386.41403 | 331.69086 | 302.93222 | 310.45578 | 343.3809 | 315.1201 | 8.603832 | 8.766186 | 8.669524 | 8.200519 | 8.315139 | 8.436822 |
| ASMM10P047 | 0.01200 | 0.343714 | 1.4939186 | up | NM_001159 | St8sia4 | ST8 alpha-N-acetyl-neuraminide alpha-2,8-sialyltransferase 4 | 56.164659 | 45.71919 | 6.10714 | 5.528038 | 59.844162 | 47.260094 | 61.38972 | 45.75059 | 51.898224 | 39.508755 | 5.9635997 | 6.0087686 | 6.3490515 | 5.4230237 | 5.5899305 | 5.571161 |
| ASMM10P031 | 0.01200 | 0.343714 | 1.5455656 | up | NM_172126 | Adam1a | a disintegrin and metallopeptidase domain 1a | 218.1913 | 172.97775 | 8.04696 | 7.418825 | 237.11778 | 239.68668 | 177.76945 | 194.79979 | 182.06367 | 142.06978 | 7.913754 | 8.311652 | 7.915473 | 7.525501 | 7.3972354 | 7.333738 |
| ASMM10P053 | 0.01201 | 0.343714 | 1.7814901 | up | NM_001033 | Ffar3 | free fatty acid receptor 3 | 12.384259 | 8.364022 | 3.901493 | 3.068408 | 12.723241 | 9.943736 | 14.485799 | 10.620341 | 8.487047 | 5.984677 | 3.6790257 | 3.8122778 | 4.213175 | 3.2747104 | 2.9599218 | 2.970593 |
| ASMM10P040 | 0.01202 | 0.343714 | 1.4218951 | up | NM_026532 | Nutf2 | nuclear transport factor 2 | 30432.535 | 25492.969 | 15.05383 | 14.546015 | 28385.578 | 30554.08 | 32357.947 | 25119.48 | 26056.994 | 25302.432 | 14.882991 | 15.069186 | 15.209313 | 14.493057 | 14.464293 | 14.680695 |
| ASMM10P034 | 0.01206 | 0.343714 | 1.5879607 | up | NM_007532 | Bcat1 | branched chain aminotransferase 1, cytosolic | 12223.347 | 8984.629 | 13.729881 | 13.062706 | 12694.342 | 10571.471 | 13404.229 | 8232.748 | 9111.639 | 9609.5 | 13.677524 | 13.588463 | 13.923656 | 12.90166 | 12.999598 | 13.28686 |
| ASMM10P024 | 0.01206 | 0.343714 | 1.4597165 | up | NM_019510 | Trpc3 | transient receptor potential cation channel, subfamily C, member 3 | 562.38157 | 469.56271 | 9.38055 | 8.834862 | 631.13153 | 545.86725 | 510.14594 | 526.15454 | 526.2781 | 356.2555 | 9.281944 | 9.467753 | 9.3919525 | 8.962453 | 8.932492 | 8.60964 |
| ASMM10P043 | 0.01208 | 0.343714 | 1.5504421 | up | NM_009237 | Sox3 | SRY (sex determining region Y)-box 3 | 756.28073 | 590.01485 | 9.80129 | 9.168611 | 789.8041 | 688.0705 | 790.9676 | 543.76074 | 696.6449 | 529.6389 | 9.616927 | 9.784963 | 10.001981 | 9.014271 | 9.335664 | 9.155897 |
| ASMM10P025 | 0.01212 | 0.343714 | 1.5882818 | up | NM_177586 | Eif5a2 | eukaryotic translation initiation factor 5A2 | 146.82484 | 112.53334 | 7.480224 | 6.812757 | 165.46599 | 117.29593 | 157.7126 | 121.2142 | 110.34889 | 106.03694 | 7.4003515 | 7.300212 | 7.740107 | 6.8336368 | 6.671896 | 6.932737 |
| ASMM10P007 | 0.01217 | 0.343714 | 2.305527 | up | NM_013459 | Cfd | complement factor D (adipsin) | 27.827176 | 14.266403 | 5.069573 | 3.864477 | 29.84511 | 31.354708 | 22.28171 | 14.04829 | 13.32867 | 15.42225 | 4.9568596 | 5.4181123 | 4.833748 | 3.687456 | 3.6184213 | 4.287553 |
| ASMM10P041 | 0.01218 | 0.343714 | 1.5517755 | up | NM_027823 | Arhgap42 | Rho GTPase activating protein 42 | 187.01272 | 148.08465 | 7.830781 | 7.196861 | 211.6743 | 161.21954 | 188.14433 | 131.38132 | 174.09602 | 138.7766 | 7.7553425 | 7.7483974 | 7.988603 | 6.9523807 | 7.335579 | 7.3026237 |
| ASMM10P048 | 0.01224 | 0.343714 | 1.3013364 | up | NM_016961 | Mapk9 | mitogen-activated protein kinase 9 | 236.08979 | 222.30187 | 8.167556 | 7.787562 | 264.70938 | 212.46509 | 231.09491 | 238.56691 | 219.58424 | 208.75446 | 8.067529 | 8.146253 | 8.288886 | 7.8188877 | 7.6736383 | 7.87016 |
| ASMM10P024 | 0.01225 | 0.343714 | 1.2827227 | up | NM_001243 | Gm14322 | predicted gene 14322 | 14730.053 | 13610.862 | 13.999687 | 13.640477 | 15712.227 | 13237.024 | 15240.909 | 13421.571 | 15629.934 | 11781.08 | 14.004014 | 13.888179 | 14.106867 | 13.591778 | 13.747293 | 13.582361 |
| ASMM10P045 | 0.01226 | 0.343714 | 1.3811442 | up | NM_139144 | Ogt | O-linked N-acetylglucosamine (GlcNAc) transferase (UDP-N-acetylglucosamine:polypeptide-N | 70414.523 | 65583.36 | 16.367692 | 15.901828 | 69083.05 | 66167.71 | 75992.81 | 67180.87 | 70442.03 | 59127.18 | 16.250591 | 16.275463 | 16.577023 | 15.871997 | 15.886818 | 15.94667 |
| ASMM10P005 | 0.01227 | 0.343714 | 1.306306 | up | NM_001145 | Ythdf3 | YTH domain family 3 | 155.03625 | 146.18659 | 7.566824 | 7.181331 | 176.3594 | 133.77405 | 154.97531 | 144.5701 | 164.51277 | 129.47691 | 7.4891624 | 7.49457 | 7.71674 | 7.0880823 | 7.2452135 | 7.210698 |
| ASMM10P008 | 0.01229 | 0.343714 | 1.462804 | up | NM_145483 | Zfp160 | zinc finger protein 160 | 123.61381 | 103.10372 | 7.236848 | 6.688111 | 139.98561 | 101.01691 | 129.8389 | 105.80773 | 107.2555 | 96.24793 | 7.1633816 | 7.086921 | 7.4602404 | 6.636072 | 6.6297073 | 6.7985544 |

| | | | | | | | | | | | | | | | | | | | | | | | |
|---|---|---|---|---|---|---|---|---|---|---|---|---|---|---|---|---|---|---|---|---|---|---|---|
| ASMM10P033 | 0.01230 | 0.343714 | 1.4593902 | up | NM_025574 | Pyurf | Pigy upstream reading frame | 809.57176 | 668.71303 | 9.900289 | 9.354923 | 821.02155 | 770.6518 | 837.04193 | 657.5836 | 728.8261 | 619.7294 | 9.672052 | 9.945703 | 10.083111 | 9.285805 | 9.399811 | 9.379153 |
| ASMM10P006 | 0.01230 | 0.343714 | 1.3968359 | up | NM_007485 | Rhod | ras homolog gene family, member D | 27.064539 | 23.06884 | 5.037447 | 4.555284 | 31.134745 | 26.30186 | 23.757011 | 24.221865 | 23.667587 | 21.317068 | 5.0173945 | 5.1683884 | 4.9265575 | 4.486923 | 4.4527655 | 4.7261643 |
| ASMM10P027 | 0.01231 | 0.343714 | 1.3050704 | up | NM_019967 | Brinp1 | bone morphogenic protein/retinoic acid inducible neural specific 1 | 10350.978 | 9256.9073 | 13.497421 | 13.113294 | 10313.414 | 10015.988 | 10723.532 | 9339.261 | 9277.532 | 9153.929 | 13.374543 | 13.506149 | 13.611572 | 13.086104 | 13.032496 | 13.221281 |
| ASMM10P002 | 0.01234 | 0.343714 | 1.3916866 | up | NM_019643 | Fam60a | family with sequence similarity 60, member A | 81.560448 | 72.18141 | 6.64433 | 6.167495 | 87.390854 | 72.44927 | 84.84122 | 82.73113 | 76.428894 | 57.384205 | 6.500682 | 6.609382 | 6.822925 | 6.2744293 | 6.144242 | 6.0838146 |
| ASMM10P008 | 0.01235 | 0.343714 | 1.3651242 | up | NM_146260 | Tmie | transmembrane inner ear | 22.643969 | 19.563882 | 4.774431 | 4.325399 | 26.58008 | 18.732187 | 22.61964 | 20.265593 | 20.310333 | 18.115719 | 4.78241 | 4.685991 | 4.8548923 | 4.2281427 | 4.2398753 | 4.5081787 |
| ASMM10P024 | 0.01236 | 0.343714 | 1.9836683 | up | NM_022994 | Dap3 | death associated protein 3 | 1035.5357 | 620.21048 | 10.230955 | 9.242784 | 1213.0441 | 795.6416 | 1097.9215 | 545.9026 | 595.17957 | 719.54926 | 10.22203 | 9.991559 | 10.479276 | 9.020949 | 9.109011 | 9.5983925 |
| ASMM10P032 | 0.01236 | 0.343714 | 1.4061601 | up | NM_001083 | Zfp655 | zinc finger protein 655 | 85.093013 | 72.998802 | 6.68861 | 6.196849 | 105.15742 | 77.59115 | 72.53047 | 72.35406 | 74.56007 | 72.082275 | 6.7674675 | 6.706211 | 6.5921507 | 6.085144 | 6.1062756 | 6.399127 |
| ASMM10P015 | 0.01241 | 0.343714 | 1.3123722 | up | NM_027355 | Rnf168 | ring finger protein 168 | 117.51443 | 110.74524 | 7.163378 | 6.771201 | 136.911 | 107.30326 | 108.32902 | 113.1722 | 133.25381 | 85.80971 | 7.1322584 | 7.1733456 | 7.1845293 | 6.7329545 | 6.9414043 | 6.6392436 |
| ASMM10P000 | 0.01242 | 0.343714 | 1.3102855 | up | NM_026437 | Fam45a | family with sequence similarity 45, member A | 282.53682 | 262.11151 | 8.413937 | 8.024056 | 329.2342 | 263.4351 | 254.94115 | 256.44968 | 265.57608 | 264.30878 | 8.376127 | 8.445351 | 8.420333 | 7.92657 | 7.9461193 | 8.199478 |
| ASMM10P004 | 0.01242 | 0.343714 | 4.417067 | up | NM_001030 | Enpp7 | ectonucleotide pyrophosphatase/phosphodiesterase 7 | 25.778576 | 6.341243 | 4.879884 | 2.736795 | 27.21987 | 13.935831 | 36.180027 | 5.9749594 | 4.9999995 | 8.04877 | 4.815003 | 4.2672324 | 5.557416 | 2.5039337 | 2.3265402 | 3.3799114 |
| ASMM10P039 | 0.01250 | 0.3446001 | 3.4697819 | up | NM_212447 | Marveld3 | MARVEL (membrane-associating) domain containing 3 | 41.480595 | 14.976677 | 5.672541 | 3.877695 | 45.630775 | 37.213654 | 41.597355 | 15.01952 | 10.111051 | 19.799461 | 5.575168 | 5.668265 | 5.7741885 | 3.786636 | 3.2139995 | 4.632451 |
| ASMM10P023 | 0.01259 | 0.3456972 | 1.6208887 | up | NM_021275 | Kcna4 | potassium voltage-gated channel, shaker-related subfamily, member 4 | 11151.515 | 7969.4632 | 13.596895 | 12.90011 | 10676.461 | 10339.989 | 12438.096 | 8060.625 | 7395.8975 | 8451.867 | 13.428569 | 13.553411 | 13.808704 | 12.869012 | 12.714933 | 13.116384 |
| ASMM10P000 | 0.01263 | 0.3464988 | 3.5843498 | up | ENSMUST | Ano5 | anoctamin 5 | 24.705704 | 9.39282 | 4.901846 | 3.060134 | 20.61821 | 26.326466 | 27.172436 | 12.116879 | 11.061581 | 4.9999995 | 4.404285 | 5.169255 | 5.131997 | 3.4630399 | 3.3508422 | 2.3665206 |
| ASMM10P005 | 0.01265 | 0.3465103 | 1.2841384 | up | NM_023122 | Gpm6b | glycoprotein m6b | 8152.7157 | 7501.7909 | 13.159993 | 12.799192 | 8616.529 | 7499.217 | 8342.401 | 7415.6753 | 8579.444 | 6510.2534 | 13.100376 | 13.099762 | 13.27984 | 12.751566 | 12.916012 | 12.729998 |
| ASMM10P036 | 0.01274 | 0.3480001 | 1.4643428 | up | NM_013886 | Hdgfrp3 | hepatoma-derived growth factor, related protein 3 | 33239.924 | 27155.907 | 15.189077 | 14.638823 | 29803.191 | 32936.63 | 36979.95 | 27447.691 | 28343.055 | 25676.975 | 14.978058 | 15.177829 | 15.411343 | 14.620057 | 14.60283 | 14.693583 |
| ASMM10P039 | 0.01281 | 0.3480001 | 1.9351898 | up | NM_015802 | Dlc1 | deleted in liver cancer 1 | 40.194464 | 23.734119 | 5.548713 | 4.596237 | 58.409946 | 27.676083 | 34.497364 | 24.539314 | 24.414186 | 22.248857 | 5.925019 | 5.2383127 | 5.482806 | 4.5039144 | 4.498359 | 4.786439 |
| ASMM10P017 | 0.01284 | 0.3480001 | 1.3465916 | up | NM_010197 | Fgf1 | fibroblast growth factor 1 | 26466.42 | 23222.841 | 14.842697 | 14.413385 | 25998.46 | 26522.465 | 26878.336 | 21376.281 | 28006.012 | 20286.23 | 14.758791 | 14.858587 | 14.910713 | 14.274707 | 14.58168 | 14.383767 |
| ASMM10P030 | 0.01287 | 0.3480001 | 1.4260419 | up | NM_027901 | Gtf3c2 | general transcription factor IIIC, polypeptide 2, beta | 2037.5739 | 1699.634 | 11.197888 | 10.685871 | 2088.637 | 1840.4786 | 2183.606 | 1696.9757 | 1805.5206 | 1596.4058 | 11.018631 | 11.155475 | 11.419557 | 10.64655 | 10.68882 | 10.722244 |
| ASMM10P044 | 0.01288 | 0.3480001 | 1.4005291 | up | NM_026662 | Prps2 | phosphoribosyl pyrophosphate synthetase 2 | 11854.065 | 9908.25 | 13.686112 | 13.20014 | 12024.562 | 10999.046 | 12538.587 | 9178.808 | 10277.793 | 10268.149 | 13.597673 | 13.640349 | 13.820313 | 13.063999 | 13.16396 | 13.37246 |
| ASMM10P011 | 0.01290 | 0.3482364 | 1.5502089 | up | NM_011849 | Nek4 | NIMA (never in mitosis gene a)-related expressed kinase 4 | 138.21637 | 109.72715 | 7.408117 | 6.775655 | 149.94188 | 128.26875 | 136.43848 | 111.17158 | 105.0882 | 112.92167 | 7.2615275 | 7.43215 | 7.5306745 | 6.707712 | 6.6006527 | 7.0185995 |
| ASMM10P042 | 0.01294 | 0.3485711 | 1.274235 | up | NM_175025 | Atp2c1 | ATPase, Ca++-sequestering | 20565.71 | 18836.39 | 14.473838 | 14.124206 | 19623.957 | 19951.113 | 22122.059 | 19118.352 | 19794.787 | 17596.031 | 14.344764 | 14.460258 | 14.616491 | 14.109428 | 14.095006 | 14.168185 |
| ASMM10P055 | 0.01295 | 0.3485711 | 1.6516614 | up | NM_183088 | 2410018M0 | RIKEN cDNA 2410018M08 gene | 51.575454 | 37.806128 | 5.979964 | 5.256046 | 57.017147 | 52.139824 | 45.56939 | 34.171253 | 41.193672 | 38.05346 | 5.8895745 | 6.142364 | 5.9079523 | 5.0006065 | 5.251802 | 5.5157285 |
| ASMM10P006 | 0.01299 | 0.3486364 | 1.7052855 | up | NM_009329 | Zfp354a | zinc finger protein 354A | 259.10068 | 183.84936 | 8.287275 | 7.517262 | 274.71454 | 206.43991 | 296.1476 | 188.40613 | 190.33685 | 172.8051 | 8.119049 | 8.106046 | 8.63673 | 7.4762607 | 7.4681044 | 7.60742 |
| ASMM10P046 | 0.01300 | 0.3486364 | 3.8688663 | up | NM_001243 | A630095N1 | RIKEN cDNA A630095N17 gene | 44.166071 | 12.462324 | 5.623447 | 3.671536 | 56.398018 | 20.025572 | 56.074623 | 12.155472 | 12.951978 | 12.279521 | 5.8744955 | 4.783028 | 6.212816 | 3.4691696 | 3.5747993 | 3.970638 |
| ASMM10P050 | 0.01302 | 0.3486364 | 1.4000687 | up | NM_001033 | Fbxo33 | F-box protein 33 | 476.03997 | 411.44714 | 9.152329 | 8.666832 | 501.26178 | 436.0292 | 490.82892 | 421.1522 | 422.99194 | 390.19727 | 8.965098 | 9.152396 | 9.339494 | 8.643165 | 8.619963 | 8.737367 |
| ASMM10P054 | 0.01307 | 0.3486364 | 1.8620131 | up | NM_021322 | Wdr4 | WD repeat domain 4 | 34.57432 | 22.154819 | 5.38799 | 4.491127 | 36.361732 | 39.0417 | 28.319527 | 27.355108 | 20.546331 | 18.563017 | 5.2403207 | 5.730088 | 5.193562 | 4.673713 | 4.259212 | 4.540456 |
| ASMM10P004 | 0.01317 | 0.349529 | 1.6187558 | up | NM_001114 | Iqsec2 | IQ motif and Sec7 domain 2 | 20750.005 | 14965.204 | 14.48241 | 13.787525 | 20268.7 | 17466.043 | 24515.273 | 15813.454 | 15041.733 | 14040.425 | 14.38905 | 14.268154 | 14.790026 | 13.825289 | 13.701304 | 13.835981 |
| ASMM10P031 | 0.01319 | 0.349529 | 1.4815028 | up | NM_026448 | Klhl7 | kelch-like 7 | 1623.5139 | 1319.6239 | 10.88846 | 10.321398 | 1622.4224 | 1615.9393 | 1632.18 | 1395.1017 | 1283.8734 | 1279.8967 | 10.653192 | 10.98455 | 11.027638 | 10.361826 | 10.199778 | 10.402591 |
| ASMM10P049 | 0.01319 | 0.349529 | 2.6326429 | up | NM_177331 | Gen1 | Gen homolog 1, endonuclease (Drosophila) | 28.133645 | 12.150691 | 5.021932 | 3.62542 | 41.647747 | 18.566587 | 24.1866 | 10.284982 | 13.1208 | 13.046291 | 5.4406815 | 4.6712074 | 4.953908 | 3.2265804 | 3.593333 | 4.056348 |
| ASMM10P030 | 0.01332 | 0.3509438 | 1.850921 | up | NM_194336 | Gbp6 | guanylate binding protein 6 | 990.75163 | 648.30905 | 10.190092 | 9.301848 | 944.217 | 1042.6658 | 985.3701 | 553.0946 | 768.34546 | 623.4871 | 9.869142 | 10.374072 | 10.327061 | 9.0382805 | 9.4795475 | 9.387717 |
| ASMM10P055 | 0.01333 | 0.3509438 | 1.3495876 | up | NM_146137 | Amigo1 | adhesion molecule with Ig like domain 1 | 562.41413 | 501.9403 | 9.378344 | 8.945825 | 602.72986 | 507.32648 | 577.18604 | 505.77377 | 552.04266 | 448.00446 | 9.212996 | 9.369264 | 9.552772 | 8.91034 | 9.001527 | 8.925609 |
| ASMM10P000 | 0.01338 | 0.3509438 | 1.3105348 | up | NM_001271 | Crem | cAMP responsive element modulator | 126.18122 | 118.37559 | 7.269877 | 6.879721 | 141.3066 | 126.00114 | 111.2359 | 115.84561 | 131.1143 | 108.16687 | 7.1757984 | 7.4083123 | 7.22552 | 6.7634234 | 6.91803 | 6.9577103 |
| ASMM10P011 | 0.01340 | 0.3509438 | 2.3534975 | up | NM_172648 | Ifi205 | interferon activated gene 205 | 22.739066 | 11.387162 | 4.769848 | 3.535042 | 26.567678 | 16.883831 | 24.76569 | 10.640978 | 10.451141 | 13.069368 | 4.7812724 | 4.538916 | 4.989357 | 3.2772174 | 3.269857 | 4.058052 |
| ASMM10P054 | 0.01341 | 0.3509438 | 1.3651119 | up | NM_172402 | Slc25a32 | solute carrier family 25, member 32 | 207.05971 | 186.52961 | 7.986073 | 7.537054 | 220.66925 | 206.01805 | 194.49182 | 183.53032 | 197.32646 | 178.73206 | 7.815703 | 8.102162 | 8.040355 | 7.4407954 | 7.520419 | 7.649948 |
| ASMM10P011 | 0.01343 | 0.3509438 | 1.274937 | up | NM_007422 | Adss | adenylosuccinate synthetase, non muscle | 3052.1192 | 2828.4582 | 11.768949 | 11.418523 | 3177.3975 | 2889.6008 | 3089.3594 | 2893.3281 | 2975.457 | 2616.5894 | 11.622007 | 11.776209 | 11.908632 | 11.409164 | 11.422513 | 11.423893 |
| ASMM10P018 | 0.01344 | 0.3509438 | 1.4373225 | up | NM_033595 | Pcdhga12 | protocadherin gamma subfamily A, 12 | 112.3574 | 93.92767 | 7.078241 | 6.554857 | 144.12111 | 97.991875 | 94.9592 | 93.79709 | 93.85021 | 94.13571 | 7.2024045 | 7.041606 | 6.9907126 | 6.454996 | 6.4423623 | 6.7672133 |
| ASMM10P042 | 0.01345 | 0.3509438 | 1.2787365 | up | NM_008829 | Pgr | progesterone receptor | 213.9041 | 205.41854 | 8.028779 | 7.67406 | 232.88292 | 212.57234 | 196.25703 | 219.19211 | 208.67767 | 188.38583 | 7.8882 | 8.147303 | 8.050835 | 7.69564 | 7.602638 | 7.723903 |
| ASMM10P042 | 0.01353 | 0.3509756 | 1.3166684 | up | NM_011992 | Rcn2 | reticulocalbin 2 | 4345.8961 | 3903.099 | 12.273705 | 11.876813 | 4403.5845 | 4258.1787 | 4375.925 | 3694.2031 | 4277.3765 | 3737.7175 | 12.129577 | 12.307284 | 12.384254 | 11.764179 | 11.927929 | 11.938331 |
| ASMM10P029 | 0.01355 | 0.3509756 | 1.2999158 | up | NM_178376 | Rraga | Ras-related GTP binding A | 19773.786 | 17955.591 | 14.423839 | 14.04542 | 21545.525 | 18032.494 | 19743.338 | 18910.922 | 20363.633 | 14592.217 | 14.485637 | 14.318127 | 14.467752 | 14.091131 | 14.141611 | 13.903519 |
| ASMM10P024 | 0.01355 | 0.3509756 | 1.4596893 | up | NM_008227 | Hcn3 | hyperpolarization-activated, cyclic nucleotide-gated K+ 3 | 12328.99 | 9871.9353 | 13.740072 | 13.19441 | 12081.832 | 12474.103 | 12431.035 | 10185.539 | 9113.355 | 10316.912 | 13.603125 | 13.80963 | 13.80746 | 13.203011 | 12.999857 | 13.380363 |
| ASMM10P053 | 0.01360 | 0.3509756 | 5.8267376 | up | NM_145603 | Ces2c | carboxylesterase 2C | 35.576952 | 7.189796 | 5.387405 | 2.844716 | 37.75538 | 21.626812 | 47.348663 | 4.9999995 | 4.9999995 | 11.569389 | 5.2983756 | 4.901455 | 5.9623833 | 2.3250442 | 2.3265402 | 3.8825645 |
| ASMM10P047 | 0.01362 | 0.3509756 | 1.4970897 | up | NM_008339 | Cd79b | CD79B antigen | 68.306862 | 54.846212 | 6.37324 | 5.79108 | 82.35772 | 53.612617 | 68.95025 | 55.598118 | 55.726055 | 53.214462 | 6.419485 | 6.184037 | 6.5161986 | 5.698187 | 5.6898766 | 5.985175 |
| ASMM10P020 | 0.01363 | 0.3509756 | 1.5356561 | up | NM_030188 | Ttc30a1 | tetratricopeptide repeat domain 30A1 | 1736.7225 | 1359.3098 | 10.981377 | 10.362522 | 1708.8088 | 1730.9691 | 1770.3896 | 1322.1145 | 1365.8322 | 1389.9827 | 10.732855 | 11.074058 | 11.137218 | 10.284124 | 10.287646 | 10.515795 |
| ASMM10P026 | 0.01363 | 0.3509756 | 1.4351483 | up | NM_001081 | Dpy19l4 | dpy-19-like 4 (C. elegans) | 141.63909 | 121.99713 | 7.444209 | 6.923009 | 149.23798 | 131.77519 | 143.9041 | 117.67197 | 135.20912 | 113.11031 | 7.254848 | 7.469847 | 7.607931 | 6.788719 | 6.960443 | 7.0198646 |
| ASMM10P012 | 0.01363 | 0.3509756 | 2.4750538 | up | NM_029407 | Fam227a | family with sequence similarity 227, member A | 41.763455 | 19.880116 | 5.650078 | 4.342618 | 42.016937 | 30.15445 | 53.118977 | 20.280167 | 18.09868 | 21.261501 | 5.4553227 | 5.362873 | 6.132037 | 4.229187 | 4.076953 | 4.721713 |
| ASMM10P042 | 0.01365 | 0.3509756 | 1.4300038 | up | NM_175535 | Arhgap20 | Rho GTPase activating protein 20 | 2016.9017 | 1672.4831 | 11.180687 | 10.664668 | 2337.635 | 1652.6003 | 2060.4697 | 1688.9182 | 1628.9745 | 1699.5566 | 11.178984 | 11.018066 | 11.345011 | 10.639455 | 10.543155 | 10.811394 |
| ASMM10P004 | 0.01375 | 0.3513929 | 1.9865573 | up | NM_001025 | Kcnc2 | potassium voltage gated channel, Shaw-related subfamily, member 2 | 91.074508 | 56.458231 | 6.804433 | 5.814163 | 83.56272 | 86.061195 | 103.59961 | 51.22888 | 70.16531 | 47.980503 | 6.438672 | 6.8540154 | 7.120612 | 5.5794735 | 6.0195823 | 5.8434324 |
| ASMM10P001 | 0.01377 | 0.3513929 | 1.2733401 | up | NM_011353 | Serf1 | small EDRK-rich factor 1 | 2463.3523 | 2288.8395 | 11.463792 | 11.115174 | 2622.3728 | 2524.9285 | 2242.7556 | 2294.1682 | 2349.3171 | 2223.0332 | 11.335889 | 11.593734 | 11.461752 | 11.08472 | 11.07196 | 11.188842 |
| ASMM10P041 | 0.01385 | 0.3513929 | 1.3111236 | up | NM_024245 | Kif23 | kinesin family member 23 | 129.93065 | 121.6498 | 7.302736 | 6.911932 | 156.34088 | 106.4264 | 127.02468 | 130.84102 | 137.14642 | 96.96197 | 7.321474 | 7.160848 | 7.425886 | 6.9466457 | 6.9807243 | 6.808427 |
| ASMM10P039 | 0.01386 | 0.3513929 | 1.4778555 | up | NM_026385 | Pllp | plasma membrane proteolipid | 6762.2213 | 5399.1443 | 12.899243 | 12.335738 | 6982.825 | 6192.535 | 7111.304 | 4751.071 | 6037.531 | 5408.831 | 12.804639 | 12.839263 | 13.053826 | 12.116962 | 12.42898 | 12.461271 |
| ASMM10P038 | 0.01390 | 0.3513929 | 1.8494787 | up | NM_153591 | Nars2 | asparaginyl-tRNA synthetase 2 (mitochondrial)(putative) | 523.43935 | 342.00251 | 9.280767 | 8.393648 | 517.3632 | 488.96198 | 563.99286 | 307.49124 | 341.7785 | 376.7378 | 9.005371 | 9.315785 | 9.521144 | 8.187337 | 8.307957 | 8.68565 |
| ASMM10P037 | 0.01391 | 0.3513929 | 1.3682084 | up | NM_025898 | Napa | N-ethylmaleimide sensitive fusion protein attachment protein alpha | 35732.902 | 31234.786 | 15.299015 | 14.846727 | 36689.516 | 33900.52 | 36608.67 | 30854.672 | 30427.121 | 32422.566 | 15.266886 | 15.236032 | 15.394126 | 14.795547 | 14.710682 | 15.033951 |
| ASMM10P008 | 0.01396 | 0.3513929 | 1.2565392 | up | NM_026420 | Paip2 | polyadenylate-binding protein-interacting protein 2 | 4740.9259 | 4452.6145 | 12.394618 | 12.065162 | 5015.1265 | 4574.6987 | 4632.9526 | 4430.614 | 4435.2524 | 4491.977 | 12.311946 | 12.410017 | 12.46189 | 12.017905 | 11.982994 | 12.194587 |
| ASMM10P027 | 0.01397 | 0.3513929 | 1.7530836 | up | NM_019961 | Pex3 | peroxisomal biogenesis factor 3 | 3747.7063 | 2499.926 | 12.048989 | 11.239094 | 3313.4763 | 3738.528 | 4191.1147 | 2589.7864 | 2697.3936 | 2212.598 | 11.683373 | 12.131685 | 12.331909 | 11.258327 | 11.277621 | 11.181335 |
| ASMM10P004 | 0.01402 | 0.3513929 | 1.5130519 | up | NM_001033 | Gm1673 | predicted gene 1673 | 498.09636 | 399.07647 | 9.219312 | 8.62185 | 499.29874 | 529.2385 | 465.75183 | 418.4067 | 400.74048 | 378.08224 | 8.960605 | 9.424408 | 9.2729225 | 8.63269 | 8.541998 | 8.690863 |
| ASMM10P021 | 0.01403 | 0.3513929 | 1.3254554 | up | NM_012032 | Serinc3 | serine incorporator 3 | 13719.653 | 12177.509 | 13.894959 | 13.488471 | 13190.163 | 13218.549 | 14750.248 | 12800.481 | 12533.498 | 11198.548 | 13.737468 | 13.883266 | 14.064144 | 13.516985 | 13.440908 | 13.507521 |
| ASMM10P027 | 0.01404 | 0.3513929 | 2.6070136 | up | NM_001081 | Fam154a | family with sequence similarity 154, member A | 90.183838 | 43.094205 | 6.791769 | 5.409371 | 96.34615 | 85.59007 | 88.615295 | 50.147575 | 29.756344 | 49.378696 | 6.6422305 | 6.84679 | 6.886286 | 5.550103 | 4.7933326 | 5.8846765 |
| ASMM10P029 | 0.01405 | 0.3513929 | 1.3013175 | up | NM_027250 | Sel1l | Sel1 repeat containing 1 | 4653.0357 | 4230.649 | 12.367291 | 11.987318 | 4859.1094 | 4227.1406 | 4872.857 | 4330.164 | 4667.902 | 3693.8809 | 12.274147 | 12.293816 | 12.53391 | 11.987251 | 12.050707 | 11.923996 |
| ASMM10P037 | 0.01407 | 0.3513929 | 1.8002507 | up | NM_001103 | Vmn2r87 | vomeronasal 2, receptor 87 | 39.816415 | 26.175366 | 5.577486 | 4.729289 | 51.633335 | 29.309584 | 38.506325 | 23.167063 | 30.333591 | 25.025444 | 5.7541356 | 5.3253803 | 5.652943 | 4.4230485 | 4.82127 | 4.943547 |
| ASMM10P008 | 0.01410 | 0.3514898 | 1.3283466 | up | NM_026899 | Ssu72 | Ssu72 RNA polymerase II CTD phosphatase homolog (yeast) | 8023.4291 | 7097.2743 | 13.146495 | 12.736863 | 8383.864 | 7852.8784 | 7833.545 | 7176.8994 | 6778.572 | 7336.3516 | 13.073825 | 13.171769 | 13.193891 | 12.713816 | 12.591695 | 12.905079 |
| ASMM10P009 | 0.01419 | 0.3519262 | 1.4105264 | up | NM_212433 | Fbxo3 | F-box protein 3 | 9010.7437 | 7513.1908 | 13.298857 | 12.802624 | 9139.085 | 7966.32 | 9926.826 | 7728.8833 | 8328.3955 | 6482.2935 | 13.186669 | 13.189199 | 13.520704 | 12.809367 | 12.875552 | 12.722952 |
| ASMM10P053 | 0.01421 | 0.3519262 | 1.4148884 | up | NM_001164 | Zfp809 | zinc finger protein 809 | 128.90886 | 111.52538 | 7.299667 | 6.798978 | 145.24876 | 109.38359 | 132.09422 | 109.41447 | 117.77138 | 107.3903 | 7.213898 | 7.202013 | 7.483089 | 6.6859236 | 6.7621627 | 6.948849 |
| ASMM10P032 | 0.01428 | 0.3523875 | 1.4524319 | up | NM_001164 | Thsd7a | thrombospondin, type I, domain containing 7A | 16585.155 | 13361.904 | 14.162437 | 13.623966 | 15110.777 | 17751.943 | 16892.744 | 13442.713 | 13339.675 | 13303.324 | 13.942359 | 14.28966 | 14.255292 | 13.59701 | 13.521791 | 13.753099 |
| ASMM10P003 | 0.01428 | 0.3523875 | 1.4785171 | up | NM_177338 | Hmbox1 | homeobox containing 1 | 216.02824 | 175.55068 | 8.013647 | 7.449496 | 273.1161 | 212.9057 | 162.06293 | 174.86174 | 181.55093 | 170.23936 | 8.111767 | 8.149865 | 7.7793097 | 7.369871 | 7.393451 | 7.585167 |
| ASMM10P000 | 0.01433 | 0.3526764 | 1.3170601 | up | NM_001160 | Il1rapl1 | interleukin 1 receptor accessory protein-like 1 | 76.511298 | 71.715103 | 6.554391 | 6.15707 | 82.9787 | 70.83395 | 75.721245 | 83.86803 | 73.982185 | 57.295094 | 6.428956 | 6.5776486 | 6.656568 | 6.2945113 | 6.0953083 | 6.0813894 |
| ASMM10P005 | 0.01434 | 0.3526764 | 1.369764 | up | NM_001177 | Irak1 | interleukin-1 receptor-associated kinase 1 | 5693.3888 | 4904.2537 | 12.654022 | 12.200095 | 5545.344 | 5612.2734 | 5922.549 | 4724.553 | 5422.005 | 4566.203 | 12.463982 | 12.702277 | 12.795807 | 12.110691 | 12.272041 | 12.217552 |
| ASMM10P008 | 0.01437 | 0.3526764 | 1.4427693 | up | NM_029814 | Chmp5 | charged multivesicular body protein 5 | 422.07709 | 356.53871 | 8.988685 | 8.459844 | 435.97415 | 408.71722 | 421.5399 | 354.9056 | 364.25772 | 350.45282 | 8.771173 | 9.063083 | 9.131798 | 8.391413 | 8.400397 | 8.587722 |
| ASMM10P020 | 0.01440 | 0.3526764 | 1.5595864 | up | NM_146845 | Olfr1098 | olfactory receptor 1098 | 276.54131 | 213.47252 | 8.366752 | 7.725588 | 335.5575 | 209.2878 | 284.77863 | 241.85487 | 203.40428 | 195.15842 | 8.400992 | 8.124148 | 8.575115 | 7.8383594 | 7.5653987 | 7.7730064 |
| ASMM10P037 | 0.01442 | 0.3526764 | 1.929199 | up | NM_009944 | Cox7a1 | cytochrome c oxidase subunit VIIa 1 | 1103.8458 | 691.15509 | 10.339189 | 9.391187 | 1147.728 | 1006.2808 | 1157.5287 | 564.186 | 710.181 | 799.09827 | 10.142027 | 10.323757 | 10.551782 | 9.067465 | 9.361047 | 9.745048 |
| ASMM10P029 | 0.01447 | 0.3533238 | 1.5242547 | up | NM_001111 | Grik2 | glutamate receptor, ionotropic, kainate 2 (beta 2) | 63.041022 | 50.744461 | 6.28018 | 5.672076 | 61.351727 | 63.26723 | 64.50411 | 51.385555 | 57.719597 | 43.12823 | 6.0003834 | 6.417697 | 6.42246 | 5.5837655 | 5.736183 | 5.69628 |
| ASMM10P031 | 0.01448 | 0.3533688 | 1.4486299 | up | NM_153510 | Pilra | paired immunoglobin-like type 2 receptor alpha | 119.47603 | 101.97047 | 7.186186 | 6.651497 | 138.37323 | 98.41337 | 121.6415 | 109.99618 | 119.47517 | 76.44006 | 7.1467185 | 7.049424 | 7.3624144 | 6.693676 | 6.782957 | 6.4778566 |
| ASMM10P043 | 0.01454 | 0.3543959 | 1.5246622 | up | NM_001017 | Lamp2 | lysosomal-associated membrane protein 2 | 8685.0651 | 6676.3279 | 13.250995 | 12.642505 | 9181.46 | 7964.7983 | 8908.937 | 5764.528 | 7192.791 | 7071.6646 | 13.193891 | 13.188172 | 13.370921 | 12.393342 | 12.678219 | 12.855954 |
| ASMM10P035 | 0.01458 | 0.3543961 | 1.3235695 | up | NM_175529 | Leng9 | leukocyte receptor cluster (LRC) member 9 | 6186.3367 | 5476.1463 | 12.768541 | 12.364107 | 6333.1055 | 5710.769 | 6515.1357 | 5469.508 | 5519.087 | 5439.8438 | 12.653848 | 12.725709 | 12.926066 | 12.320734 | 12.298875 | 12.472712 |
| ASMM10P054 | 0.01462 | 0.3543961 | 1.9945797 | up | NM_144829 | Aarsd1 | alanyl-tRNA synthetase domain containing 1 | 18.180939 | 10.698 | 4.445056 | 3.448971 | 17.095078 | 14.703299 | 22.74444 | 12.089939 | 10.4122 | 9.59186 | 4.127954 | 4.3440347 | 4.8631783 | 3.4610496 | 3.2625587 | 3.6233044 |
| ASMM10P023 | 0.01470 | 0.3543961 | 1.2849001 | up | NM_028664 | Ankrd45 | ankyrin repeat domain 45 | 44450.748 | 41522.926 | 15.622235 | 15.260579 | 42027.19 | 44420.33 | 46904.723 | 40951.117 | 45041.47 | 38576.19 | 15.485536 | 15.617755 | 15.763415 | 15.195766 | 15.265999 | 15.319973 |
| ASMM10P033 | 0.01472 | 0.3543961 | 2.1164969 | up | NM_008461 | Klra14 | killer cell lectin-like receptor subfamily A, member 14, pseudogene | 105.08217 | 59.557245 | 6.986927 | 5.905248 | 100.57831 | 80.246704 | 134.4215 | 63.50044 | 62.565365 | 52.60593 | 6.701234 | 6.751761 | 7.5077853 | 5.894614 | 5.8516135 | 5.9695177 |
| ASMM10P041 | 0.01475 | 0.3543961 | 1.5261901 | up | NM_001198 | Ccpg1os | cell cycle progression 1, opposite strand | 4749.5368 | 3653.0081 | 12.39861 | 11.788675 | 4389.2397 | 4970.459 | 4888.9116 | 3586.351 | 3735.5242 | 3637.149 | 12.124437 | 12.531491 | 12.539901 | 11.717366 | 11.74754 | 11.901119 |
| ASMM10P053 | 0.01480 | 0.3543961 | 1.4079054 | up | NM_009405 | Tnni2 | troponin I, skeletal, fast 2 | 262.28142 | 228.22363 | 8.317704 | 7.824154 | 272.4073 | 246.24867 | 268.1883 | 230.16805 | 240.8485 | 213.65434 | 8.107952 | 8.349503 | 8.495657 | 7.766021 | 7.803055 | 7.9033847 |
| ASMM10P018 | 0.01486 | 0.3543961 | 1.5338946 | up | NM_053142 | Pcdhb17 | protocadherin beta 17 | 9802.588 | 7419.5795 | 13.418127 | 12.800928 | 9550.17 | 9517.612 | 10339.982 | 7022.0454 | 7201.269 | 8035.424 | 13.257479 | 13.427152 | 13.56975 | 12.679777 | 12.681104 | 13.041902 |
| ASMM10P050 | 0.01487 | 0.3543961 | 1.8158314 | up | NM_019784 | Tex21 | testis expressed gene 21 | 28.134626 | 18.353546 | 5.07947 | 4.218839 | 29.590107 | 20.48749 | 34.32628 | 18.964102 | 21.797909 | 14.298628 | 4.9437184 | 4.820769 | 5.4739213 | 4.1296573 | 4.3396745 | 4.1871862 |
| ASMM10P054 | 0.01490 | 0.3543961 | 1.764027 | up | NM_001080 | B230217C1 | RIKEN cDNA B230217C12 gene | 172.5542 | 120.42773 | 7.715939 | 6.897066 | 186.5361 | 190.41122 | 140.71529 | 103.9963 | 141.7106 | 115.57629 | 7.5723157 | 7.9961414 | 7.57936 | 6.6121244 | 7.0297794 | 7.0492954 |
| ASMM10P004 | 0.01495 | 0.3543961 | 1.6307882 | up | NM_001128 | Cecr2 | cat eye syndrome chromosome region, candidate 2 | 98.269977 | 71.453775 | 6.863647 | 6.158077 | 132.42702 | 70.48888 | 91.89403 | 68.951775 | 84.93322 | 60.47633 | 7.079789 | 6.5706882 | 6.940463 | 6.0150676 | 6.2975783 | 6.161586 |
| ASMM10P046 | 0.01496 | 0.3543961 | 1.2727432 | up | NM_054056 | Pawr | PRKC, apoptosis, WT1, regulator | 114.5124 | 110.23964 | 7.124276 | 6.776334 | 130.4949 | 114.52816 | 98.51415 | 110.68158 | 126.08321 | 93.95412 | 7.0618687 | 7.266367 | 7.044591 | 6.702015 | 6.861872 | 6.7651157 |
| ASMM10P033 | 0.01499 | 0.3543961 | 1.7126268 | up | NM_009525 | Wnt5b | wingless-related MMTV integration site 5B | 82.012377 | 58.824857 | 6.657143 | 5.880932 | 78.35788 | 81.73091 | 85.94834 | 54.143425 | 68.95704 | 53.374107 | 6.351027 | 6.7789316 | 6.841469 | 5.6592283 | 5.9940248 | 5.989542 |
| ASMM10P021 | 0.01502 | 0.3543961 | 1.6672095 | up | NM_023645 | Kdelc1 | KDEL (Lys-Asp-Glu-Leu) containing 1 | 180.27189 | 132.28717 | 7.774909 | 7.037473 | 183.84523 | 150.36938 | 206.60106 | 138.11461 | 146.93198 | 111.81492 | 7.5495696 | 7.64691 | 8.128246 | 7.022599 | 7.084094 | 7.0057263 |

| | | | | | | | | | | | | | | | | | | | | | | | |
|---|---|---|---|---|---|---|---|---|---|---|---|---|---|---|---|---|---|---|---|---|---|---|---|
| ASMM10P005 | 0.01503 | 0.3543961 | 1.2751433 | up | NM_001267 | Pex2 | peroxisomal biogenesis factor 2 | 691.6897 | 652.40719 | 9.671285 | 9.320626 | 782.0683 | 603.1496 | 689.8512 | 626.8386 | 705.074 | 625.30896 | 9.602449 | 9.606019 | 9.805388 | 9.21638 | 9.351242 | 9.394256 |
| ASMM10P002 | 0.01507 | 0.3543961 | 2.14552 | up | NM_175465 | Sestd1 | SEC14 and spectrin domains 1 | 51.760802 | 28.215365 | 5.931985 | 4.830657 | 73.14117 | 36.255577 | 45.88566 | 33.93905 | 23.315357 | 27.391687 | 6.2505207 | 5.628392 | 5.9170413 | 4.9889464 | 4.4311395 | 5.071886 |
| ASMM10P042 | 0.01507 | 0.3543961 | 3.9880092 | up | NM_130455 | Grin3b | glutamate receptor, ionotropic, NMDA3B | 27.376242 | 8.237562 | 5.051568 | 3.055899 | 22.562334 | 28.0975 | 31.468891 | 5.246089 | 8.280715 | 11.185882 | 4.542477 | 5.2631626 | 5.349064 | 2.4066072 | 2.928198 | 3.8328922 |
| ASMM10P017 | 0.01515 | 0.3543961 | 2.1864458 | up | NM_176963 | Galm | galactose mutarotase | 81.401668 | 45.199987 | 6.628815 | 5.500227 | 95.439 | 64.39115 | 84.374855 | 40.669 | 40.69218 | 54.23878 | 6.6285257 | 6.442284 | 6.815635 | 5.2535815 | 5.236858 | 6.0102425 |
| ASMM10P014 | 0.01516 | 0.3543961 | 1.3969018 | up | NM_019631 | Tmem45a | transmembrane protein 45a | 176.44236 | 155.03825 | 7.746736 | 7.264505 | 197.14648 | 184.75322 | 147.42738 | 151.71869 | 179.09438 | 134.30168 | 7.6505527 | 7.9471173 | 7.6425366 | 7.1585917 | 7.3763514 | 7.2585716 |
| ASMM10P039 | 0.01526 | 0.3543961 | 2.0507442 | up | NM_146208 | Neil3 | nei like 3 (E. coli) | 94.697701 | 54.579098 | 6.814379 | 5.778231 | 124.3591 | 65.791306 | 93.942696 | 52.129517 | 51.35827 | 60.249508 | 6.9959903 | 6.472204 | 6.9749427 | 5.6054773 | 5.574118 | 6.155099 |
| ASMM10P009 | 0.01526 | 0.3543961 | 1.2971275 | up | NM_171826 | Cldn25 | claudin 25 | 401.91664 | 378.28538 | 8.916967 | 8.541647 | 433.88718 | 371.01547 | 400.84726 | 408.5816 | 388.61523 | 337.6593 | 8.764261 | 8.920068 | 9.066572 | 8.598423 | 8.496666 | 8.529851 |
| ASMM10P027 | 0.01531 | 0.3543961 | 1.708435 | up | NM_001004 | AA415398 | expressed sequence AA415398 | 7088.2002 | 4841.2774 | 12.957584 | 12.184909 | 6183.5483 | 6950.258 | 8130.7944 | 4905.039 | 4812.932 | 4805.8613 | 12.6225 | 13.006462 | 13.243791 | 12.16441 | 12.095655 | 12.294662 |
| ASMM10P014 | 0.01532 | 0.3543961 | 1.2868287 | up | NM_176928 | Brwd1 | bromodomain and WD repeat domain containing 1 | 19251.66 | 17554.926 | 14.383982 | 14.020162 | 19452.453 | 18175.541 | 20126.986 | 16796.727 | 18457.834 | 17410.217 | 14.331066 | 14.327823 | 14.493057 | 13.909639 | 13.998117 | 14.15273 |
| ASMM10P020 | 0.01534 | 0.3543961 | 1.5332734 | up | NM_028235 | Ttc30b | tetratricopeptide repeat domain 30B | 7265.8073 | 5543.3903 | 12.998981 | 12.382366 | 6860.9204 | 6895.1475 | 8041.354 | 5261.8306 | 5747.394 | 5620.9463 | 12.775207 | 12.994654 | 13.227083 | 12.262358 | 12.356859 | 12.527882 |
| ASMM10P027 | 0.01539 | 0.3543961 | 1.9512716 | up | NM_175517 | Fam221b | family with sequence similarity 221, member B | 149.0421 | 93.74698 | 7.510669 | 6.546254 | 151.54729 | 127.53771 | 168.0413 | 90.69329 | 88.24335 | 102.3043 | 7.27586 | 7.4241085 | 7.8320374 | 6.403908 | 6.350239 | 6.884615 |
| ASMM10P027 | 0.01545 | 0.3543961 | 1.7363121 | up | NM_015828 | Gne | glucosamine (UDP-N-acetyl)-2-epimerase/N-acetylmannosamine kinase | 936.80124 | 649.77328 | 10.11148 | 9.315454 | 896.3353 | 945.97626 | 968.09216 | 639.8976 | 614.70325 | 694.719 | 9.798479 | 10.235538 | 10.300423 | 9.244269 | 9.157486 | 9.544606 |
| ASMM10P005 | 0.01550 | 0.3543961 | 2.1642689 | up | NM_001166 | Slc39a11 | solute carrier family 39 (metal ion transporter), member 11 | 88.580092 | 49.167127 | 6.726059 | 5.612179 | 117.31349 | 77.94003 | 70.486755 | 55.925327 | 37.929935 | 53.64612 | 6.9186606 | 6.711896 | 6.547621 | 5.7074223 | 5.1335545 | 5.9955616 |
| ASMM10P042 | 0.01552 | 0.3543961 | 1.3641321 | up | NM_001002 | Bfsp2 | beaded filament structural protein 2, phakinin | 74.648712 | 66.220917 | 6.498999 | 6.051016 | 90.75516 | 57.92162 | 75.269356 | 73.68602 | 70.77155 | 54.20518 | 6.557079 | 6.292629 | 6.64729 | 6.110614 | 6.032751 | 6.009683 |
| ASMM10P041 | 0.01553 | 0.3543961 | 1.253159 | up | NM_019797 | Trip4 | thyroid hormone receptor interactor 4 | 658.10257 | 632.5283 | 9.601171 | 9.275602 | 707.4849 | 660.36975 | 606.45306 | 641.37726 | 668.38513 | 587.8225 | 9.456259 | 9.726884 | 9.62037 | 9.248057 | 9.2729225 | 9.305825 |
| ASMM10P041 | 0.01555 | 0.3543961 | 1.3789572 | up | NM_178906 | AI593442 | expressed sequence AI593442 | 59783.636 | 53682.245 | 16.085494 | 15.621917 | 58185.805 | 55549.344 | 65615.76 | 54456.574 | 61716.156 | 44874.004 | 15.979592 | 15.981073 | 16.295818 | 15.592535 | 15.711377 | 15.561838 |
| ASMM10P051 | 0.01556 | 0.3543961 | 1.5108688 | up | NM_027335 | 2210016F16 | RIKEN cDNA 2210016F16 gene | 176.36137 | 144.25304 | 7.750796 | 7.155417 | 195.10153 | 182.25768 | 151.7249 | 131.00558 | 176.40482 | 125.34871 | 7.63785 | 7.927444 | 7.6870933 | 6.948415 | 7.3532524 | 7.1645846 |
| ASMM10P028 | 0.01562 | 0.3543961 | 1.6732448 | up | NM_029100 | Sepn1 | selenoprotein N, 1 | 1714.8193 | 1230.612 | 10.951012 | 10.208364 | 1916.9644 | 1328.6688 | 1898.8248 | 1302.0947 | 1423.7574 | 965.98395 | 10.899404 | 10.713628 | 11.240006 | 10.265333 | 10.346462 | 10.013297 |
| ASMM10P008 | 0.01570 | 0.3543961 | 1.7350878 | up | NM_144553 | Dlgap5 | discs, large (Drosophila) homolog-associated protein 5 | 2149.1003 | 1480.1927 | 11.275499 | 10.48049 | 1941.0785 | 2093.0833 | 2413.139 | 1614.832 | 1555.642 | 1270.104 | 10.917203 | 11.344194 | 11.565099 | 10.572378 | 10.476898 | 10.392194 |
| ASMM10P037 | 0.01573 | 0.3543961 | 1.5553068 | up | NM_025378 | Ifitm3 | interferon induced transmembrane protein 3 | 5793.7225 | 4411.0951 | 12.683264 | 12.046065 | 5998.585 | 6214.433 | 5168.1494 | 3723.2742 | 4952.568 | 4557.443 | 12.581356 | 12.846516 | 12.621921 | 11.779093 | 12.145798 | 12.213305 |
| ASMM10P029 | 0.01574 | 0.3543961 | 1.6872923 | up | NM_010100 | Edar | ectodysplasin-A receptor | 163.19612 | 117.94687 | 7.628669 | 6.87396 | 182.2402 | 126.1366 | 181.21155 | 108.97319 | 134.6166 | 110.25082 | 7.53584 | 7.4095664 | 7.940602 | 6.6806073 | 6.955845 | 6.9854264 |
| ASMM10P028 | 0.01580 | 0.3543961 | 1.3082853 | up | NM_028264 | Tmem55a | transmembrane protein 55A | 664.33116 | 608.23677 | 9.610851 | 9.223174 | 734.77234 | 687.305 | 570.91614 | 617.9646 | 618.419 | 588.3267 | 9.512966 | 9.782671 | 9.536916 | 9.195494 | 9.1665325 | 9.307495 |
| ASMM10P008 | 0.01583 | 0.3543961 | 1.3803616 | up | NM_139154 | Rab40c | Rab40c, member RAS oncogene family | 113.21777 | 101.53956 | 7.122519 | 6.657473 | 117.31551 | 106.498 | 115.83979 | 112.72862 | 109.2064 | 82.683655 | 6.9187603 | 7.1619906 | 7.286807 | 6.7279067 | 6.658843 | 6.5856695 |
| ASMM10P054 | 0.01585 | 0.3543961 | 2.2809089 | up | NM_027658 | Hexim2 | hexamethylene bis-acetamide inducible 2 | 40.598896 | 21.415736 | 5.620072 | 4.430463 | 45.59352 | 44.01923 | 32.183937 | 24.85164 | 16.671167 | 22.7244 | 5.5734005 | 5.9042287 | 5.382587 | 4.5231495 | 3.9523098 | 4.8159304 |
| ASMM10P052 | 0.01590 | 0.3543961 | 1.4522844 | up | NM_010476 | Hsd17b7 | hydroxysteroid (17-beta) dehydrogenase 7 | 4794.6319 | 3895.5214 | 12.412139 | 11.873815 | 4883.366 | 4381.951 | 5118.5786 | 3554.5051 | 4200.5864 | 3931.4727 | 12.279591 | 12.349241 | 12.607585 | 11.705319 | 11.905905 | 12.010221 |
| ASMM10P026 | 0.01592 | 0.3543961 | 1.4542516 | up | NM_001079 | Syne1 | spectrin repeat containing, nuclear envelope 1 | 156.91097 | 130.13493 | 7.558397 | 7.01812 | 192.43199 | 159.8857 | 118.41521 | 139.6542 | 131.12068 | 119.62991 | 7.61841 | 7.7363715 | 7.320409 | 7.0385323 | 6.9182916 | 7.097536 |
| ASMM10P011 | 0.01598 | 0.3543961 | 1.5241009 | up | NM_028516 | 1700056E22 | RIKEN cDNA 1700056E22 gene | 187.16084 | 151.22108 | 7.835943 | 7.227984 | 204.41629 | 164.73242 | 192.33382 | 135.9641 | 169.19371 | 148.50542 | 7.7048264 | 7.780673 | 8.022329 | 7.0003886 | 7.291059 | 7.3925056 |
| ASMM10P028 | 0.01604 | 0.3543961 | 1.4770771 | up | NM_007745 | Cort | cortistatin | 7021.9945 | 5489.1513 | 12.926284 | 12.363538 | 9175.351 | 5858.6504 | 6031.982 | 5410.912 | 5715.516 | 5341.026 | 13.193239 | 12.766274 | 12.819338 | 12.30447 | 12.343886 | 12.442259 |
| ASMM10P011 | 0.01608 | 0.3543961 | 1.8006482 | up | NM_001037 | Samd4 | sterile alpha motif domain containing 4 | 463.22509 | 303.99142 | 9.079032 | 8.230515 | 531.4651 | 331.58826 | 526.6219 | 329.42337 | 287.85034 | 294.70056 | 9.040746 | 8.765703 | 9.430646 | 8.28385 | 8.059094 | 8.348602 |
| ASMM10P027 | 0.01612 | 0.3543961 | 1.3226509 | up | NM_011902 | Tekt2 | tektin 2 | 302.8925 | 279.7544 | 8.52014 | 8.116708 | 325.77188 | 281.07855 | 301.82706 | 287.81323 | 285.06546 | 266.38452 | 8.359764 | 8.535622 | 8.665035 | 8.096273 | 8.044969 | 8.208882 |
| ASMM10P044 | 0.01613 | 0.3543961 | 1.3913316 | up | NM_138742 | Nap1l3 | nucleosome assembly protein 1-like 3 | 33503.304 | 28740.033 | 15.206231 | 14.729765 | 29955.475 | 34666.05 | 35888.387 | 28876.932 | 29569.377 | 27773.791 | 14.990087 | 15.268338 | 15.360269 | 14.700542 | 14.675291 | 14.813462 |
| ASMM10P014 | 0.01616 | 0.3543961 | 2.168865 | up | NM_033474 | Arvcf | armadillo repeat gene deleted in velo-cardio-facial syndrome | 44.207098 | 24.149992 | 5.708803 | 4.591862 | 53.2675 | 50.21561 | 29.138184 | 29.63803 | 26.519234 | 16.292713 | 5.7951193 | 6.092228 | 5.2390604 | 4.789852 | 4.6239753 | 4.3617597 |
| ASMM10P004 | 0.01617 | 0.3543961 | 1.3757575 | up | NM_001109 | C2cd5 | C2 calcium-dependent domain containing 5 | 50.178601 | 44.685371 | 5.94117 | 5.480944 | 55.26444 | 51.475937 | 43.795425 | 52.60847 | 46.81332 | 34.634323 | 5.8466206 | 6.124322 | 5.8525686 | 5.617773 | 5.4363527 | 5.388707 |
| ASMM10P007 | 0.01629 | 0.3543961 | 1.9036659 | up | NM_019684 | Srpk3 | serine/arginine-rich protein specific kinase 3 | 296.47698 | 187.60653 | 8.464861 | 7.536081 | 304.86093 | 223.86285 | 360.70715 | 208.31093 | 202.34663 | 152.16202 | 8.260718 | 8.218131 | 8.915734 | 7.621958 | 7.558241 | 7.428043 |
| ASMM10P032 | 0.01634 | 0.3543961 | 1.7932785 | up | NM_133906 | Zkscan1 | zinc finger with KRAB and SCAN domains 1 | 62.097015 | 42.993677 | 6.246885 | 5.404285 | 63.562195 | 69.16503 | 53.56382 | 52.482037 | 48.163204 | 28.335789 | 6.050709 | 6.5436096 | 6.1463356 | 5.614682 | 5.479689 | 5.1184845 |
| ASMM10P016 | 0.01634 | 0.3543961 | 1.3819317 | up | NM_026514 | Cdc42ep3 | CDC42 effector protein (Rho GTPase binding) 3 | 3347.2814 | 2856.4147 | 11.899919 | 11.433232 | 3347.9573 | 3413.1812 | 3280.7058 | 2846.5496 | 2848.6936 | 2874.001 | 11.699139 | 12.00979 | 11.990827 | 11.38675 | 11.360499 | 11.552448 |
| ASMM10P030 | 0.01637 | 0.3543961 | 1.4188314 | up | NM_009170 | Shh | sonic hedgehog | 1732.8164 | 1465.9525 | 10.976339 | 10.471636 | 1833.763 | 1559.3588 | 1805.3273 | 1370.7988 | 1541.332 | 1485.7267 | 10.836844 | 10.929187 | 11.162988 | 10.335207 | 10.462644 | 10.617058 |
| ASMM10P000 | 0.01638 | 0.3543961 | 1.3323839 | up | NM_173374 | Srsf1 | serine/arginine-rich splicing factor 1 | 1378.8432 | 1239.715 | 10.648159 | 10.234149 | 1533.7743 | 1156.948 | 1445.8074 | 1286.3787 | 1310.4333 | 1122.333 | 10.566714 | 10.523583 | 10.854179 | 10.246851 | 10.232265 | 10.223331 |
| ASMM10P024 | 0.01644 | 0.3543961 | 1.2618773 | up | NM_001039 | Tmco1 | transmembrane and coiled-coil domains 1 | 4068.154 | 3767.7634 | 12.169357 | 11.833785 | 4897.2764 | 3792.5544 | 3514.6313 | 3685.1716 | 3858.2644 | 3759.8542 | 12.284821 | 12.146749 | 12.0765 | 11.759217 | 11.793923 | 11.948215 |
| ASMM10P047 | 0.01644 | 0.3543961 | 1.5532355 | up | NM_009423 | Traf4 | TNF receptor associated factor 4 | 67.12472 | 52.43704 | 6.358979 | 5.723702 | 74.57002 | 55.90949 | 70.89465 | 59.236378 | 48.159767 | 49.914974 | 6.2777143 | 6.244074 | 6.5551476 | 5.7940784 | 5.4795237 | 5.897504 |
| ASMM10P046 | 0.01645 | 0.3543961 | 1.7804949 | up | NM_027031 | Efcab9 | EF-hand calcium binding domain 9 | 376.42221 | 256.69391 | 8.824401 | 7.992122 | 344.80478 | 378.8265 | 405.63535 | 246.54942 | 275.74924 | 247.78307 | 8.439458 | 8.950037 | 9.083707 | 7.864928 | 7.995068 | 8.116371 |
| ASMM10P054 | 0.01650 | 0.3543961 | 2.0796086 | up | NM_001114 | Ppp4r1 | protein phosphatase 4, regulatory subunit 1 | 39.612379 | 22.803045 | 5.593043 | 4.536731 | 34.97167 | 46.039043 | 37.826424 | 21.681034 | 24.380922 | 22.34718 | 5.185012 | 5.9710746 | 5.623043 | 4.3211255 | 4.496043 | 4.793025 |
| ASMM10P035 | 0.01650 | 0.3543961 | 1.5533185 | up | NM_027994 | Cand1 | cullin associated and neddylation disassociated 1 | 1412.3855 | 1086.3446 | 10.687711 | 10.052357 | 1478.855 | 1278.1584 | 1480.143 | 1033.2091 | 1049.4526 | 1176.372 | 10.521531 | 10.652481 | 10.88912 | 9.941056 | 9.927105 | 10.28891 |
| ASMM10P005 | 0.01651 | 0.3543961 | 1.3114293 | up | NM_001267 | Pex2 | peroxisomal biogenesis factor 2 | 622.63062 | 571.09896 | 9.521956 | 9.130816 | 697.3595 | 546.2596 | 624.27277 | 540.141 | 613.63104 | 559.52484 | 9.434688 | 9.468548 | 9.662632 | 9.002721 | 9.154154 | 9.235573 |
| ASMM10P010 | 0.01651 | 0.3543961 | 1.5265272 | up | NM_009785 | Cacna2d3 | calcium channel, voltage-dependent, alpha2/delta subunit 3 | 32784.9 | 25889.527 | 15.181535 | 14.571281 | 30191.225 | 33685.715 | 34477.76 | 25601.74 | 24427.758 | 27639.084 | 15.006234 | 15.226351 | 15.312019 | 14.517377 | 14.38905 | 14.807417 |
| ASMM10P015 | 0.01652 | 0.3543961 | 1.4686434 | up | NM_011638 | Tfrc | transferrin receptor | 229.95931 | 190.80485 | 8.122634 | 7.568149 | 260.8347 | 187.46613 | 241.5771 | 180.34702 | 209.5948 | 182.47273 | 8.047863 | 7.9712486 | 8.348789 | 7.4157186 | 7.6094275 | 7.679302 |
| ASMM10P052 | 0.01653 | 0.3543961 | 1.4378593 | up | NM_026184 | Ero1lb | ERO1-like beta (S. cerevisiae) | 8546.6292 | 6947.8464 | 13.230995 | 12.707072 | 8245.813 | 7885.8325 | 9508.242 | 6906.3506 | 7111.3135 | 6825.875 | 13.050242 | 13.178522 | 13.46422 | 12.655005 | 12.66033 | 12.805882 |
| ASMM10P009 | 0.01657 | 0.3543961 | 1.4438426 | up | NM_176917 | Mettl4 | methyltransferase like 4 | 65.076862 | 55.253792 | 6.318199 | 5.788286 | 72.34262 | 64.509865 | 58.3781 | 68.11461 | 52.054295 | 45.59247 | 6.2369976 | 6.4455075 | 6.272093 | 5.998541 | 5.5943727 | 5.771944 |
| ASMM10P033 | 0.01658 | 0.3543961 | 1.3856967 | up | NM_007881 | Atn1 | atrophin 1 | 92.145725 | 79.855827 | 6.79473 | 6.324119 | 117.47801 | 81.85034 | 77.108826 | 81.936874 | 79.142426 | 78.48818 | 6.9205236 | 6.7808623 | 6.6828055 | 6.2619486 | 6.196953 | 6.5134554 |
| ASMM10P023 | 0.01664 | 0.3543961 | 1.6091733 | up | NM_177253 | Cstf3 | cleavage stimulation factor, 3' pre-RNA, subunit 3 | 1890.1677 | 1413.8503 | 11.096567 | 10.410248 | 1786.7246 | 1865.0305 | 2018.748 | 1514.9915 | 1582.8976 | 1143.6619 | 10.80257 | 11.173615 | 11.313517 | 10.481971 | 10.502479 | 10.246293 |
| ASMM10P004 | 0.01666 | 0.3543961 | 1.3431649 | up | NM_001083 | Mpzl1 | myelin protein zero-like 1 | 154.3866 | 142.29958 | 7.56218 | 7.136543 | 168.48672 | 148.44368 | 146.2294 | 137.48882 | 169.31633 | 120.09358 | 7.423042 | 7.6307597 | 7.6327376 | 7.01462 | 7.292228 | 7.102782 |
| ASMM10P009 | 0.01667 | 0.3543961 | 2.0622859 | up | NM_170671 | Mycbpap | MYCBP associated protein | 36.906043 | 21.282635 | 5.487545 | 4.4433 | 32.276836 | 32.552284 | 45.889008 | 21.893204 | 22.021542 | 19.93316 | 5.071583 | 5.4739213 | 5.9171295 | 4.3344593 | 4.353632 | 4.6418095 |
| ASMM10P007 | 0.01672 | 0.3543961 | 2.1685044 | up | NM_011976 | Sema4g | sema domain, immunoglobulin domain (Ig), transmembrane domain (TM) and short cytopla | 41.307453 | 22.231685 | 5.613848 | 4.497148 | 49.072025 | 47.341545 | 27.50879 | 20.405058 | 26.5581 | 19.731897 | 5.6809034 | 6.011325 | 5.1493163 | 4.2394457 | 4.6250715 | 4.6269264 |
| ASMM10P005 | 0.01674 | 0.3543961 | 1.3572552 | up | NM_001163 | Kif9 | kinesin family member 9 | 166.7928 | 151.32668 | 7.672296 | 7.231604 | 186.65897 | 159.75337 | 153.96606 | 144.64732 | 157.63896 | 151.69377 | 7.574326 | 7.7350426 | 7.7075205 | 7.08918 | 7.1818237 | 7.4238095 |
| ASMM10P025 | 0.01675 | 0.3543961 | 1.505004 | up | NM_008839 | Pik3ca | phosphatidylinositol 3-kinase, catalytic, alpha polypeptide | 1238.1259 | 981.64212 | 10.498734 | 9.908967 | 1250.5981 | 1091.9935 | 1371.786 | 1009.6997 | 1056.1511 | 879.07556 | 10.273668 | 10.443539 | 10.778996 | 9.904531 | 9.935101 | 9.887269 |
| ASMM10P007 | 0.01677 | 0.3543961 | 1.4685898 | up | NM_207686 | Elavl2 | ELAV (embryonic lethal, abnormal vision, Drosophila)-like 2 (Hu antigen B) | 95.074707 | 78.573053 | 6.852545 | 6.298114 | 108.81929 | 73.59543 | 102.8094 | 85.55212 | 82.59764 | 67.5694 | 6.813748 | 6.63383 | 7.1100574 | 6.32212 | 6.25867 | 6.313551 |
| ASMM10P037 | 0.01682 | 0.3543961 | 1.3066022 | up | NM_021549 | Pnkp | polynucleotide kinase 3'- phosphatase | 3523.582 | 3179.9653 | 11.971902 | 11.586082 | 3588.6062 | 3311.3152 | 3670.8245 | 3216.3972 | 3327.6365 | 2995.8623 | 11.805685 | 11.970742 | 12.139278 | 11.560284 | 11.581577 | 11.616384 |
| ASMM10P036 | 0.01689 | 0.3543961 | 1.2556564 | up | NM_020599 | Rlbp1 | retinaldehyde binding protein 1 | 23.758718 | 22.62993 | 4.854687 | 4.526246 | 26.452433 | 22.86277 | 21.96095 | 24.77021 | 23.415628 | 19.703953 | 4.7733803 | 4.978395 | 4.8122864 | 4.516344 | 4.4370446 | 4.625348 |
| ASMM10P028 | 0.01691 | 0.3543961 | 1.3007588 | up | NM_011060 | Padi3 | peptidyl arginine deiminase, type III | 63.616666 | 58.986989 | 6.265668 | 5.886315 | 79.82834 | 59.18657 | 51.835087 | 66.807 | 61.442863 | 48.711105 | 6.3766074 | 6.324015 | 6.0963817 | 5.5691086 | 5.8240776 | 5.8657575 |
| ASMM10P007 | 0.01697 | 0.3543961 | 1.5793183 | up | NM_019706 | Rnf138 | ring finger protein 138 | 833.2586 | 634.30513 | 9.937327 | 9.278025 | 844.71875 | 870.10754 | 784.9495 | 566.869 | 668.8227 | 667.2237 | 9.711983 | 10.110583 | 9.989415 | 9.074223 | 9.274434 | 9.485418 |
| ASMM10P028 | 0.01698 | 0.3543961 | 1.2767213 | up | NM_175364 | Lyrm2 | LYR motif containing 2 | 1122.2974 | 1052.184 | 10.364362 | 10.011919 | 1253.9856 | 1068.0612 | 1044.8453 | 1040.5499 | 1041.1615 | 1074.8407 | 10.277739 | 10.408267 | 10.407081 | 9.952086 | 9.916512 | 10.167158 |
| ASMM10P026 | 0.01703 | 0.3543961 | 1.3057288 | up | NM_001198 | Arhgef2 | rho/rac guanine nucleotide exchange factor (GEF) 2 | 56.299773 | 52.716956 | 6.112501 | 5.727646 | 59.542202 | 50.25888 | 59.098236 | 55.706367 | 57.902153 | 44.542347 | 5.955062 | 6.0932646 | 6.2891774 | 5.701656 | 5.741283 | 5.7399993 |
| ASMM10P008 | 0.01704 | 0.3543961 | 1.5640842 | up | NM_146265 | Olfr1506 | olfactory receptor 1506 | 78.991457 | 61.318077 | 6.58499 | 5.939672 | 84.632256 | 88.99211 | 63.350006 | 67.09724 | 68.14175 | 48.71524 | 6.4579062 | 6.902537 | 6.394526 | 5.975467 | 5.977558 | 5.8659897 |
| ASMM10P048 | 0.01706 | 0.3543961 | 1.5973716 | up | NM_028336 | Tmem107 | transmembrane protein 107 | 197.86618 | 153.99612 | 7.919208 | 7.243508 | 206.10918 | 212.18332 | 175.36603 | 147.9785 | 192.23322 | 121.77663 | 7.7175603 | 8.143856 | 7.896207 | 7.1227026 | 7.482135 | 7.1256857 |
| ASMM10P050 | 0.01711 | 0.3543961 | 1.5646076 | up | NM_001161 | Grhl1 | grainyhead-like 1 (Drosophila) | 66.818068 | 51.295762 | 6.342516 | 5.696715 | 76.16555 | 51.053123 | 73.23553 | 52.14923 | 53.276535 | 48.46152 | 6.3089232 | 6.1129665 | 6.6056585 | 5.60605 | 5.6265817 | 5.857514 |
| ASMM10P008 | 0.01711 | 0.3543961 | 1.4898386 | up | NM_146136 | Slc16a4 | solute carrier family 16 (monocarboxylic acid transporters), member 4 | 114.94878 | 94.564443 | 7.130367 | 6.555211 | 134.23853 | 105.2782 | 105.32961 | 82.55348 | 108.1282 | 93.01165 | 7.100308 | 7.146242 | 7.144551 | 6.2710543 | 6.642123 | 6.7524557 |
| ASMM10P000 | 0.01713 | 0.3543961 | 10.145863 | up | NM_133211 | Tlr7 | toll-like receptor 7 | 254.56453 | 22.727466 | 7.845259 | 4.50244 | 142.35521 | 530.6215 | 90.71959 | 17.05949 | 24.592268 | 26.53064 | 7.186096 | 9.427152 | 6.9225297 | 3.9709933 | 4.510692 | 5.0256333 |
| ASMM10P021 | 0.01716 | 0.3543961 | 1.6119767 | up | NM_001039 | Defb25 | defensin beta 25 | 54.519849 | 40.702651 | 6.051429 | 5.362598 | 65.21095 | 53.119186 | 45.229412 | 37.41661 | 42.718796 | 41.972546 | 6.088086 | 6.1706243 | 5.895576 | 5.128375 | 5.3028655 | 5.656553 |
| ASMM10P055 | 0.01716 | 0.3543961 | 2.86811 | up | NM_001146 | Acot7 | acyl-CoA thioesterase 7 | 38.846388 | 16.246398 | 5.549963 | 4.029863 | 38.531532 | 28.695316 | 49.312317 | 12.770519 | 16.6354 | 19.333275 | 5.331597 | 5.2961836 | 6.0221095 | 3.5440414 | 3.948425 | 4.5971227 |
| ASMM10P040 | 0.01717 | 0.3543961 | 1.4018542 | up | NM_011503 | Stxbp2 | syntaxin binding protein 2 | 1034.9301 | 886.22085 | 10.250409 | 9.763073 | 1073.953 | 1037.3715 | 993.4658 | 849.9042 | 923.40674 | 885.3516 | 10.048489 | 10.366397 | 10.336341 | 9.648813 | 9.745048 | 9.895357 |
| ASMM10P053 | 0.01717 | 0.3543961 | 1.4066067 | up | NM_019738 | Nupr1 | nuclear protein transcription regulator 1 | 669.3161 | 571.18851 | 9.61277 | 9.120551 | 832.2237 | 582.2167 | 593.5079 | 498.85138 | 671.86914 | 542.845 | 9.692422 | 9.553853 | 9.592036 | 8.889763 | 9.281858 | 9.190033 |
| ASMM10P035 | 0.01718 | 0.3543961 | 1.7351968 | up | NM_025620 | Rep15 | RAB15 effector protein | 1346.9654 | 913.84194 | 10.60274 | 9.80764 | 1467.0841 | 1020.0599 | 1553.7521 | 900.11993 | 905.3091 | 936.0968 | 10.507673 | 10.341354 | 10.959192 | 9.731449 | 9.717097 | 9.974375 |
| ASMM10P007 | 0.01728 | 0.3543961 | 1.7443772 | up | NM_010892 | Nek2 | NIMA (never in mitosis gene a)-related expressed kinase 2 | 3055.384 | 2041.0567 | 11.74979 | 10.947078 | 3530.1316 | 2197.8733 | 3438.147 | 2186.7607 | 1898.698 | 2037.7113 | 11.779667 | 11.420532 | 12.049171 | 11.015479 | 10.761304 | 11.064451 |
| ASMM10P025 | 0.01733 | 0.3543961 | 1.2604294 | up | NM_008991 | Abcd3 | ATP-binding cassette, sub-family D (ALD), member 3 | 5738.1582 | 5388.1061 | 12.669733 | 12.335817 | 6151.198 | 5724.4775 | 5338.799 | 5051.71 | 5665.352 | 5447.2563 | 12.616281 | 12.72806 | 12.664857 | 12.203539 | 12.32735 | 12.476563 |
| ASMM10P014 | 0.01735 | 0.3543961 | 1.572306 | up | NM_001013 | Dtx3l | deltex 3-like (Drosophila) | 1856.4314 | 1414.1129 | 11.0724 | 10.419518 | 1797.4823 | 1773.9873 | 1997.8247 | 1307.7179 | 1487.9607 | 1446.6602 | 10.811113 | 11.107376 | 11.29871 | 10.271016 | 10.412432 | 10.575105 |
| ASMM10P035 | 0.01739 | 0.3543961 | 1.669674 | up | NM_013538 | Cdca3 | cell division cycle associated 3 | 254.25717 | 186.92327 | 8.274475 | 7.534909 | 245.1825 | 251.84575 | 265.74326 | 170.71452 | 210.90123 | 179.15405 | 7.9601383 | 8.380205 | 8.483083 | 7.334021 | 7.61816 | 7.652546 |
| ASMM10P030 | 0.01740 | 0.3543961 | 1.4839428 | up | NM_030052 | Cox7b2 | cytochrome c oxidase subunit VIIb2 | 6283.6059 | 4979.3202 | 12.786775 | 12.217339 | 6128.142 | 5661.185 | 7061.4907 | 5435.505 | 5339.7305 | 4162.725 | 12.608399 | 12.713429 | 13.038497 | 12.313156 | 12.249974 | 12.088888 |
| ASMM10P004 | 0.01744 | 0.3543961 | 1.4025532 | up | NM_001113 | Gatad2a | GATA zinc finger domain containing 2A | 40.028398 | 34.059344 | 5.600558 | 5.112503 | 49.577374 | 35.98032 | 34.5275 | 32.909203 | 36.48942 | 32.77941 | 5.697567 | 5.61896 | 5.485148 | 4.94324 | 5.0796914 | 5.314577 |
| ASMM10P005 | 0.01744 | 0.3543961 | 1.2847844 | up | NM_001163 | Rabggtb | RAB geranylgeranyl transferase, b subunit | 685.32177 | 641.41809 | 9.657704 | 9.296177 | 749.47375 | 689.25775 | 617.2338 | 695.9651 | 625.89667 | 602.3925 | 9.539597 | 9.788043 | 9.645471 | 9.369018 | 9.181224 | 9.33829 |
| ASMM10P006 | 0.01749 | 0.3543961 | 1.4135631 | up | NM_001253 | Sept6 | septin 6 | 3029.547 | 2522.2974 | 11.749594 | 11.250258 | 3593.7908 | 2401.6492 | 3093.201 | 2750.7524 | 2462.7039 | 2353.436 | 11.809191 | 11.528686 | 11.910905 | 11.338708 | 11.142449 | 11.269616 |
| ASMM10P019 | 0.01751 | 0.3543961 | 1.396898 | up | NM_011750 | Sf1 | splicing factor 1 | 2219.3453 | 1893.6052 | 11.322208 | 10.839982 | 2228.4358 | 2059.7302 | 2369.87 | 1937.8285 | 1986.69 | 1756.2971 | 11.110196 | 11.318869 | 11.53756 | 10.836432 | 10.829709 | 10.853804 |
| ASMM10P009 | 0.01760 | 0.3543961 | 2.4268725 | up | NM_177765 | Ttll13 | tubulin tyrosine ligase-like family, member 13 | 60.931298 | 31.136773 | 6.220993 | 4.941894 | 65.76734 | 64.053925 | 52.97263 | 22.218237 | 35.818523 | 35.373558 | 6.101254 | 6.4353056 | 6.126418 | 4.3585205 | 5.0506907 | 5.4164715 |
| ASMM10P050 | 0.01761 | 0.3543961 | 1.3696048 | up | NM_009706 | Arhgap5 | Rho GTPase activating protein 5 | 1714.3065 | 1519.605 | 10.968338 | 10.514578 | 1768.2843 | 1718.8145 | 1655.8207 | 1480.6842 | 1763.6102 | 1314.531 | 10.78647 | 11.064877 | 11.053667 | 10.445605 | 10.65996 | 10.43817 |
| ASMM10P039 | 0.01761 | 0.3543961 | 1.2739076 | up | NM_019708 | Scoc | short coiled-coil protein | 43702.452 | 40876.568 | 15.590069 | 15.240808 | 41211.33 | 45691.496 | 44204.53 | 42006.918 | 41504.977 | 39117.81 | 15.448338 | 15.659248 | 15.662621 | 15.233477 | 15.149548 | 15.3394 |
| ASMM10P013 | 0.01765 | 0.3543961 | 1.6773472 | up | NM_011994 | Abcd2 | ATP-binding cassette, sub-family D (ALD), member 2 | 7039.919 | 4979.2617 | 12.952497 | 12.206316 | 7704.21 | 5692.369 | 7723.178 | 4306.0127 | 6187.7803 | 4443.992 | 12.958969 | 12.721277 | 13.177245 | 11.979735 | 12.460065 | 12.179147 |

| | | | | | | | | | | | | | | | | | | | | | | | |
|---|---|---|---|---|---|---|---|---|---|---|---|---|---|---|---|---|---|---|---|---|---|---|---|
| ASMM10P032 | 0.01767 | 0.3543961 | 1.6814762 | up | NM_031998 | Cep41 | centrosomal protein 41 | 3716.941 | 2601.646 | 12.044523 | 11.294794 | 3550.351 | 3427.0784 | 4173.3936 | 2439.3584 | 2553.1772 | 2812.4023 | 11.787951 | 12.017243 | 12.328374 | 11.169924 | 11.195738 | 11.518721 |
| ASMM10P036 | 0.01767 | 0.3543961 | 1.4063178 | up | NM_023258 | Pycard | PYD and CARD domain containing | 41.772338 | 35.926236 | 5.671645 | 5.179722 | 46.190243 | 33.634132 | 45.492638 | 41.13684 | 37.94986 | 28.692007 | 5.5924053 | 5.5179853 | 5.904544 | 5.2682133 | 5.1342072 | 5.136746 |
| ASMM10P053 | 0.01767 | 0.3543961 | 1.4461899 | up | NM_021323 | Usp29 | ubiquitin specific peptidase 29 | 309.35468 | 259.43626 | 8.538891 | 8.006634 | 370.57257 | 261.76602 | 295.72546 | 255.10678 | 248.77568 | 274.42633 | 8.543702 | 8.437956 | 8.635014 | 7.9173503 | 7.8502226 | 8.252328 |
| ASMM10P029 | 0.01772 | 0.3543961 | 1.2558534 | up | NM_025909 | Oma1 | OMA1 homolog, zinc metallopeptidase (S. cerevisiae) | 213.16192 | 207.8109 | 8.019563 | 7.690895 | 239.94873 | 211.21751 | 188.31952 | 205.0948 | 220.0912 | 198.2467 | 7.9307013 | 8.137995 | 7.9899917 | 7.597865 | 7.6778164 | 7.7970023 |
| ASMM10P042 | 0.01773 | 0.3543961 | 1.6009557 | up | NM_001166 | Armc8 | armadillo repeat containing 8 | 234.6639 | 178.73013 | 8.145824 | 7.46689 | 290.03885 | 199.7685 | 214.18434 | 171.1935 | 165.94647 | 199.05042 | 8.192574 | 8.059223 | 8.185674 | 7.3386555 | 7.2598314 | 7.802184 |
| ASMM10P041 | 0.01780 | 0.3543961 | 1.306736 | up | NM_133718 | Tmem30a | transmembrane protein 30A | 26779.639 | 24196.705 | 14.861994 | 14.476026 | 27897.883 | 26509.564 | 25931.47 | 23949.842 | 23596.514 | 25043.76 | 14.863755 | 14.85645 | 14.865776 | 14.417122 | 14.341092 | 14.669864 |
| ASMM10P021 | 0.01785 | 0.3543961 | 1.2711343 | up | NM_133838 | Ehd4 | EH-domain containing 4 | 7045.2079 | 6508.1916 | 12.959712 | 12.613595 | 7089.343 | 7027.7666 | 7018.514 | 6338.3364 | 6658.9033 | 6527.335 | 12.829868 | 13.019301 | 13.029966 | 12.538548 | 12.567541 | 12.734697 |
| ASMM10P037 | 0.01787 | 0.3543961 | 1.8630254 | up | NM_001081 | Mki67 | antigen identified by monoclonal antibody Ki 67 | 236.17733 | 154.10814 | 8.149003 | 7.251356 | 275.44354 | 180.78287 | 252.30557 | 142.44429 | 145.82268 | 174.05746 | 8.123474 | 7.9164524 | 8.407084 | 7.0656123 | 7.071884 | 7.616572 |
| ASMM10P017 | 0.01792 | 0.3543961 | 1.3365792 | up | NM_013819 | H2-M3 | histocompatibility 2, M region locus 3 | 736.17912 | 668.37858 | 9.761484 | 9.342938 | 811.9129 | 700.2406 | 696.38385 | 695.35254 | 774.6858 | 535.0974 | 9.656192 | 9.810714 | 9.817545 | 9.368057 | 9.492264 | 9.168494 |
| ASMM10P023 | 0.01798 | 0.3543961 | 1.9290515 | up | NM_133851 | Nusap1 | nucleolar and spindle associated protein 1 | 49.290686 | 31.69081 | 5.92364 | 4.975748 | 52.379326 | 48.303177 | 47.189556 | 24.715769 | 40.866386 | 29.490274 | 5.7744536 | 6.038437 | 5.958028 | 4.5132737 | 5.2415714 | 5.1723986 |
| ASMM10P007 | 0.01799 | 0.3543961 | 1.2734711 | up | NM_010709 | Eif2d | eukaryotic translation initiation factor 2D | 2599.5822 | 2414.8659 | 11.538787 | 11.190021 | 2758.947 | 2705.9194 | 2333.8801 | 2465.147 | 2436.0322 | 2343.4185 | 11.413186 | 11.68929 | 11.513886 | 11.185957 | 11.122582 | 11.261524 |
| ASMM10P006 | 0.01805 | 0.3543961 | 1.3743861 | up | NM_009939 | Cops2 | COP9 (constitutive photomorphogenic) homolog, subunit 2 (Arabidopsis thaliana) | 2477.1514 | 2138.0855 | 11.476356 | 11.017569 | 2520.2827 | 2349.904 | 2561.2676 | 2125.9363 | 2152.5198 | 2135.8005 | 11.283235 | 11.505035 | 11.640799 | 10.97451 | 10.948137 | 11.13006 |
| ASMM10P044 | 0.01808 | 0.3543961 | 1.4616731 | up | NM_175180 | Wdr44 | WD repeat domain 44 | 358.24642 | 298.94398 | 8.755888 | 8.208267 | 365.37912 | 371.01376 | 338.34637 | 282.6567 | 326.40518 | 287.77005 | 8.521377 | 8.919855 | 8.826431 | 8.065 | 8.242169 | 8.317632 |
| ASMM10P009 | 0.01810 | 0.3543961 | 1.3661321 | up | NM_194069 | Ifi27 | interferon, alpha-inducible protein 27 | 1139.5917 | 1002.2473 | 10.387273 | 9.937176 | 1216.4456 | 1141.5002 | 1060.8292 | 933.4725 | 1070.3873 | 1002.8821 | 10.226887 | 10.5057 | 10.429232 | 9.788043 | 9.954394 | 10.069091 |
| ASMM10P055 | 0.01815 | 0.3543961 | 1.7623144 | up | NM_023813 | Camk2d | calcium/calmodulin-dependent protein kinase II, delta | 1091.1824 | 749.87798 | 10.323933 | 9.506462 | 1161.615 | 1057.2261 | 1054.7062 | 827.4355 | 604.15155 | 818.0469 | 10.155771 | 10.39559 | 10.420438 | 9.611653 | 9.129948 | 9.777784 |
| ASMM10P014 | 0.01816 | 0.3543961 | 1.6354053 | up | NM_023737 | Ehhadh | enoyl-Coenzyme A, hydratase/3-hydroxyacyl Coenzyme A dehydrogenase | 250.71789 | 189.24742 | 8.255145 | 7.545497 | 242.50436 | 272.41776 | 237.23154 | 183.21643 | 226.51158 | 158.01424 | 7.9459677 | 8.493475 | 8.325993 | 7.437996 | 7.7178617 | 7.4806333 |
| ASMM10P018 | 0.01817 | 0.3543961 | 1.349809 | up | NM_033588 | Pcdhga5 | protocadherin gamma subfamily A, 5 | 129.5981 | 119.44193 | 7.315706 | 6.88295 | 139.31163 | 127.60804 | 121.87463 | 130.83821 | 136.29538 | 91.1922 | 7.1565175 | 7.4256387 | 7.364961 | 6.9462495 | 6.972966 | 6.729636 |
| ASMM10P041 | 0.01817 | 0.3543961 | 1.4805101 | up | NM_001005 | Mreg | melanoregulin | 6031.6704 | 4769.4487 | 12.730263 | 12.164169 | 6192.916 | 5173.8887 | 6728.2065 | 4551.1206 | 4860.1694 | 4897.056 | 12.625803 | 12.591695 | 12.973291 | 12.055811 | 12.112625 | 12.32407 |
| ASMM10P051 | 0.01818 | 0.3543961 | 2.1969441 | up | NM_028878 | Slc6a19 | solute carrier family 6 (neurotransmitter transporter), member 19 | 25.433339 | 13.81109 | 4.948996 | 3.813498 | 24.845251 | 28.053783 | 23.400982 | 12.480532 | 13.503209 | 15.44953 | 4.681714 | 5.2605696 | 4.904705 | 3.5125747 | 3.6372094 | 4.29071 |
| ASMM10P035 | 0.01819 | 0.3543961 | 1.3839356 | up | NM_024457 | Rap1b | RAS related protein 1b | 3618.9297 | 3079.3564 | 12.004925 | 11.536148 | 3575.3323 | 3797.318 | 3484.1387 | 2991.05 | 3151.1426 | 3095.8767 | 11.80072 | 12.149197 | 12.064858 | 11.451026 | 11.502443 | 11.654975 |
| ASMM10P028 | 0.01821 | 0.3543961 | 1.4830012 | up | NM_028604 | Trmt11 | tRNA methyltransferase 11 | 169.24857 | 139.88813 | 7.687569 | 7.119049 | 197.41766 | 141.14182 | 169.18622 | 132.26962 | 142.23878 | 145.156 | 7.6538467 | 7.5671167 | 7.841744 | 6.9612913 | 7.0340614 | 7.3617954 |
| ASMM10P055 | 0.01824 | 0.3543961 | 1.4201697 | up | NM_001253 | Anp32e | acidic (leucine-rich) nuclear phosphoprotein 32 family, member E | 10478.419 | 8568.541 | 13.505531 | 12.999468 | 9936.766 | 9848.862 | 11649.63 | 8076.167 | 9416.912 | 8212.544 | 13.3242 | 13.475632 | 13.716762 | 12.870753 | 13.053826 | 13.073825 |
| ASMM10P000 | 0.01824 | 0.3543961 | 1.8098995 | up | NM_001272 | Arvcf | armadillo repeat gene deleted in velo-cardio-facial syndrome | 280.85872 | 187.50129 | 8.389756 | 7.533846 | 321.77628 | 207.56429 | 313.2356 | 162.87587 | 214.80907 | 184.81894 | 8.339866 | 8.113726 | 8.715675 | 7.263508 | 7.6423903 | 7.69564 |
| ASMM10P013 | 0.01825 | 0.3543961 | 1.4104385 | up | NM_009634 | Adsl | adenylosuccinate lyase | 285.88074 | 246.94665 | 8.435873 | 7.93973 | 309.14288 | 257.2443 | 291.25504 | 252.53934 | 242.69456 | 245.60605 | 8.281534 | 8.414032 | 8.612054 | 7.901553 | 7.8139896 | 8.103646 |
| ASMM10P051 | 0.01831 | 0.3543961 | 1.2608943 | up | NM_026503 | 1110058L19 | RIKEN cDNA 1110058L19 gene | 20688.862 | 19071.331 | 14.482864 | 14.148416 | 20694.2 | 20022.47 | 21349.916 | 18670.719 | 19347.129 | 19196.145 | 14.422225 | 14.465212 | 14.561154 | 14.080812 | 14.063246 | 14.301191 |
| ASMM10P027 | 0.01832 | 0.3543961 | 1.8691046 | up | NM_026560 | Cdca8 | cell division cycle associated 8 | 1278.8399 | 811.6312 | 10.538796 | 9.636448 | 1177.6464 | 1221.8976 | 1436.9758 | 784.7072 | 765.87646 | 884.30994 | 10.176058 | 10.596248 | 10.844081 | 9.540898 | 9.474419 | 9.894028 |
| ASMM10P011 | 0.01834 | 0.3543961 | 1.7408388 | up | NM_001110 | Cdca2 | cell division cycle associated 2 | 74.616391 | 52.456243 | 6.521461 | 5.721678 | 76.06323 | 70.966934 | 76.81901 | 46.05469 | 55.77787 | 55.53617 | 6.306963 | 6.5798163 | 6.677603 | 5.433234 | 5.6917114 | 6.040089 |
| ASMM10P031 | 0.01836 | 0.3543961 | 4.7953978 | up | NM_025951 | Pi4k2b | phosphatidylinositol 4-kinase type 2 beta | 33.826667 | 8.270193 | 5.297415 | 3.035764 | 38.403057 | 42.641083 | 20.44186 | 4.9999995 | 7.6124997 | 12.198081 | 5.326563 | 5.8595185 | 4.7061634 | 2.3250442 | 2.8208594 | 3.9613898 |
| ASMM10P022 | 0.01838 | 0.3543961 | 1.3365391 | up | NM_015730 | Chrna4 | cholinergic receptor, nicotinic, alpha polypeptide 4 | 4055.3316 | 3615.3622 | 12.177304 | 11.758802 | 4275.0786 | 3570.9531 | 4319.963 | 3643.0203 | 4052.0386 | 3151.0278 | 12.08545 | 12.07756 | 12.368902 | 11.74206 | 11.855416 | 11.67893 |
| ASMM10P049 | 0.01841 | 0.3543961 | 1.6747958 | up | NM_008425 | Kcnj2 | potassium inwardly-rectifying channel, subfamily J, member 2 | 18751.92 | 13075.664 | 14.322134 | 13.578149 | 18619.854 | 14906.236 | 22729.67 | 14406.966 | 14099.689 | 10720.336 | 14.257833 | 14.048358 | 14.660212 | 13.691268 | 13.599884 | 13.443296 |
| ASMM10P051 | 0.01843 | 0.3543961 | 1.7019486 | up | NM_001042 | Fancc | Fanconi anemia, complementation group C | 914.79805 | 634.93957 | 10.048588 | 9.2814 | 995.6044 | 684.56036 | 1064.2294 | 641.9291 | 664.0038 | 598.8858 | 9.9345255 | 9.776878 | 10.434361 | 9.249663 | 9.2647295 | 9.329809 |
| ASMM10P009 | 0.01844 | 0.3543961 | 2.7542178 | up | NM_198634 | Tigd3 | tigger transposable element derived 3 | 137.56225 | 59.496938 | 7.361165 | 5.899522 | 113.70927 | 109.91361 | 189.06386 | 58.34261 | 54.638855 | 65.50935 | 6.878348 | 7.208867 | 7.9962797 | 5.7704973 | 5.6606975 | 6.267372 |
| ASMM10P031 | 0.01846 | 0.3543961 | 1.3469952 | up | NM_001110 | Atp2a2 | ATPase, Ca++ transporting, cardiac muscle, slow twitch 2 | 64448.951 | 60141.008 | 16.214437 | 15.784693 | 60893.113 | 66228.09 | 66225.65 | 57231.453 | 65808.625 | 57382.945 | 16.04459 | 16.277786 | 16.320936 | 15.653617 | 15.799418 | 15.901043 |
| ASMM10P013 | 0.01848 | 0.3543961 | 1.49593 | up | NM_153533 | Tenc1 | tensin like C1 domain-containing phosphatase | 169.91907 | 141.95902 | 7.700216 | 7.119173 | 183.36485 | 169.4317 | 156.96065 | 145.50131 | 176.03601 | 104.33974 | 7.545729 | 7.8217483 | 7.7331705 | 7.0983353 | 7.3499384 | 6.909246 |
| ASMM10P031 | 0.01850 | 0.3543961 | 1.2858556 | up | NM_011263 | Rest | RE1-silencing transcription factor | 48.495896 | 46.30114 | 5.898544 | 5.535815 | 52.974136 | 46.13858 | 46.374973 | 54.435005 | 45.828182 | 38.640232 | 5.7895327 | 5.973045 | 5.933053 | 5.6671195 | 5.4026213 | 5.537704 |
| ASMM10P054 | 0.01851 | 0.3543961 | 1.3584543 | up | NM_001081 | Vps8 | vacuolar protein sorting 8 homolog (S. cerevisiae) | 67.472772 | 60.945687 | 6.375237 | 5.933271 | 72.713615 | 64.39949 | 65.30521 | 57.261055 | 70.696976 | 54.87903 | 6.242858 | 6.4424844 | 6.44037 | 5.7447953 | 6.0312057 | 6.0238132 |
| ASMM10P015 | 0.01852 | 0.3543961 | 1.3883773 | up | NM_023844 | Jam2 | junction adhesion molecule 2 | 1576.166 | 1364.676 | 10.841697 | 10.368297 | 1769.9414 | 1319.0416 | 1639.5149 | 1273.1982 | 1469.7379 | 1351.0918 | 10.787932 | 10.702763 | 11.034396 | 10.230872 | 10.393952 | 10.480068 |
| ASMM10P039 | 0.01854 | 0.3543961 | 1.2799922 | up | NM_001252 | Csgalnact1 | chondroitin sulfate N-acetylgalactosaminyltransferase 1 | 4738.1823 | 4358.1016 | 12.39098 | 12.034845 | 4955.915 | 4229.097 | 5029.535 | 4440.932 | 4583.289 | 4050.0837 | 12.30179 | 12.295666 | 12.575484 | 12.021966 | 12.026758 | 12.055811 |
| ASMM10P026 | 0.01857 | 0.3543961 | 1.5151521 | up | NM_177377 | Fam166b | family with sequence similarity 166, member B | 94.478503 | 76.506077 | 6.860569 | 6.261107 | 92.69236 | 86.81746 | 103.92569 | 80.51753 | 82.70527 | 66.29543 | 6.5876684 | 6.86679 | 7.12725 | 6.2374754 | 6.2613497 | 6.2844954 |
| ASMM10P005 | 0.01858 | 0.3543961 | 1.2640966 | up | NM_001198 | App | amyloid beta (A4) precursor protein | 12645.115 | 11774.664 | 13.774366 | 13.436259 | 13108.96 | 13635.562 | 11190.822 | 12768.73 | 11960.46 | 10594.802 | 13.73014 | 13.926475 | 13.666482 | 13.513229 | 13.374543 | 13.421005 |
| ASMM10P006 | 0.01858 | 0.3543961 | 1.2596829 | up | NM_001253 | Kcnma1 | potassium large conductance calcium-activated channel, subfamily M, alpha member 1 | 4116.124 | 3808.838 | 12.187795 | 11.854734 | 5055.9126 | 3638.8035 | 3653.6558 | 3907.69 | 3770.447 | 3748.377 | 12.326469 | 12.103809 | 12.133106 | 11.856749 | 11.764179 | 11.943274 |
| ASMM10P054 | 0.01858 | 0.3543961 | 2.1212148 | up | NM_001110 | Crem | cAMP responsive element modulator | 65.218479 | 36.93378 | 6.31246 | 5.227569 | 55.558624 | 59.90775 | 80.189064 | 37.86465 | 36.316967 | 36.619724 | 5.854155 | 6.3421926 | 6.741032 | 5.145926 | 5.073096 | 5.4636855 |
| ASMM10P029 | 0.01862 | 0.3543961 | 1.3624645 | up | NM_013868 | Hspb7 | heat shock protein family, member 7 (cardiovascular) | 65.861308 | 59.123 | 6.334589 | 5.88837 | 72.29814 | 55.904133 | 69.38165 | 57.95247 | 69.03036 | 50.38617 | 6.2365627 | 6.2440033 | 6.5231996 | 5.7611566 | 5.995437 | 5.908516 |
| ASMM10P001 | 0.01863 | 0.3543961 | 2.1047163 | up | NM_001136 | Ikbkg | inhibitor of kappaB kinase gamma | 16.512721 | 8.874104 | 4.232306 | 3.158681 | 25.74486 | 10.04741 | 13.745892 | 9.248258 | 10.90771 | 6.466345 | 4.735291 | 3.8252718 | 4.1363564 | 3.065251 | 3.3291626 | 3.0816283 |
| ASMM10P007 | 0.01864 | 0.3543961 | 1.4317385 | up | NM_010212 | Fhl2 | four and a half LIM domains 2 | 300.07403 | 255.34282 | 8.501116 | 7.983348 | 335.17596 | 293.85123 | 271.1949 | 274.3117 | 232.29605 | 259.4207 | 8.399551 | 8.594591 | 8.509205 | 8.021386 | 7.754047 | 8.17461 |
| ASMM10P003 | 0.01867 | 0.3543961 | 1.4155431 | up | NM_001163 | Prom1 | prominin 1 | 386.82781 | 335.40584 | 8.863566 | 8.362211 | 410.9767 | 386.2493 | 363.25742 | 320.4657 | 403.99393 | 281.7579 | 8.687926 | 8.978472 | 8.924301 | 8.245613 | 8.552493 | 8.288526 |
| ASMM10P020 | 0.01868 | 0.3543961 | 1.4082059 | up | NM_018732 | Scn3a | sodium channel, voltage-gated, type III, alpha | 456.87729 | 394.11746 | 9.099057 | 8.605199 | 470.70374 | 438.53564 | 461.3925 | 404.5602 | 393.11298 | 384.6792 | 8.877385 | 9.160816 | 9.258971 | 8.585569 | 8.512886 | 8.717142 |
| ASMM10P025 | 0.01870 | 0.3543961 | 1.5759776 | up | NM_001113 | Fxr1 | fragile X mental retardation gene 1, autosomal homolog | 288.92111 | 222.60855 | 8.447344 | 7.791097 | 321.6272 | 248.8121 | 296.32404 | 213.47801 | 213.8745 | 240.47313 | 8.339349 | 8.364371 | 8.638313 | 7.661875 | 7.636008 | 8.075409 |
| ASMM10P039 | 0.01877 | 0.3543961 | 1.7796389 | up | NM_007426 | Angpt2 | angiopoietin 2 | 109.33309 | 74.923698 | 7.060423 | 6.228838 | 119.5984 | 88.17836 | 120.2225 | 69.2371 | 74.846924 | 80.68707 | 6.9460816 | 6.8893795 | 7.345808 | 6.0221095 | 6.1127253 | 6.5516806 |
| ASMM10P032 | 0.01881 | 0.3543961 | 1.3723957 | up | NM_178625 | Tmem209 | transmembrane protein 209 | 5983.0064 | 5133.1179 | 12.724016 | 12.267319 | 6325.6665 | 5663.963 | 5959.3896 | 5098.8027 | 4841.731 | 5458.82 | 12.653177 | 12.714232 | 12.804639 | 12.216154 | 12.105394 | 12.48041 |
| ASMM10P007 | 0.01885 | 0.3543961 | 1.9274274 | up | NM_025565 | Spc25 | SPC25, NDC80 kinetochore complex component, homolog (S. cerevisiae) | 365.5284 | 226.49267 | 8.758304 | 7.811628 | 382.58325 | 269.47256 | 444.5294 | 259.47986 | 209.29173 | 210.70642 | 8.592688 | 8.477622 | 9.204602 | 7.94239 | 7.607807 | 7.8846855 |
| ASMM10P026 | 0.01890 | 0.3543961 | 1.3095868 | up | NM_001171 | Bag1 | BCL2-associated athanogene 1 | 76584.193 | 74826.58 | 16.494727 | 16.105615 | 75807.875 | 75940.79 | 78003.914 | 74138.17 | 77440.41 | 72901.16 | 16.398306 | 16.472843 | 16.613031 | 16.020136 | 16.030523 | 16.266186 |
| ASMM10P008 | 0.01893 | 0.3543961 | 1.2908227 | up | NM_145542 | Ahcyl1 | S-adenosylhomocysteine hydrolase-like 1 | 261.18426 | 249.16242 | 8.310397 | 7.942106 | 282.01675 | 241.6294 | 259.90662 | 250.32831 | 283.38785 | 213.7711 | 8.158461 | 8.321976 | 8.450754 | 7.887344 | 8.035136 | 7.9038386 |
| ASMM10P015 | 0.01897 | 0.3543961 | 1.3963638 | up | NM_026554 | Ncbp2 | nuclear cap binding protein subunit 2 | 4576.118 | 3843.4056 | 12.346108 | 11.864433 | 4467.768 | 4464.4673 | 4796.1187 | 3823.9182 | 3788.2698 | 3918.0283 | 12.152694 | 12.374476 | 12.511155 | 11.820265 | 11.768994 | 12.004041 |
| ASMM10P006 | 0.01898 | 0.3543961 | 1.9601208 | up | NM_009689 | Birc5 | baculoviral IAP repeat-containing 5 | 102.03268 | 62.528019 | 6.944104 | 5.973162 | 108.10021 | 120.9511 | 77.046715 | 67.9528 | 56.058556 | 63.5727 | 6.8038597 | 7.346313 | 6.6821404 | 5.9948807 | 5.698334 | 6.2262707 |
| ASMM10P011 | 0.01904 | 0.3543961 | 1.3049519 | up | NM_008133 | Glud1 | glutamate dehydrogenase 1 | 1028.9043 | 951.69932 | 10.237491 | 9.853494 | 1139.6688 | 941.02856 | 1006.0154 | 907.15295 | 1111.0458 | 836.8992 | 10.131729 | 10.228585 | 10.352158 | 9.743052 | 10.004711 | 9.812719 |
| ASMM10P035 | 0.01905 | 0.3543961 | 1.4428511 | up | NM_001039 | Gm7092 | predicted gene 7092 | 111.79158 | 95.496145 | 7.105905 | 6.576982 | 112.13112 | 106.61038 | 116.63324 | 97.42865 | 101.11037 | 87.94941 | 6.85775 | 7.1637278 | 7.296236 | 6.51226 | 6.5437355 | 6.674951 |
| ASMM10P026 | 0.01907 | 0.3543961 | 1.3658604 | up | NM_183423 | Pla2g12a | phospholipase A2, group XIIA | 65515.812 | 59783.125 | 16.232088 | 15.782278 | 67935.32 | 60198.562 | 68413.555 | 59034.035 | 61002.094 | 59313.246 | 16.208199 | 16.103659 | 16.384407 | 15.69634 | 15.697496 | 15.952999 |
| ASMM10P025 | 0.01911 | 0.3543961 | 1.3110763 | up | NM_173181 | Zc2hc1a | zinc finger, C2HC-type containing 1A | 3411.6766 | 3089.6881 | 11.932134 | 11.541382 | 3707.193 | 3326.352 | 3201.4849 | 3207.1724 | 2864.0215 | 3197.8704 | 11.858699 | 11.977082 | 11.96062 | 11.556799 | 11.368042 | 11.699305 |
| ASMM10P008 | 0.01914 | 0.3543961 | 1.9096018 | up | NM_026307 | Cuta | cutA divalent cation tolerance homolog (E. coli) | 27.656594 | 17.341715 | 5.074991 | 4.141719 | 24.311085 | 28.635609 | 30.023088 | 16.197731 | 20.771713 | 15.0557 | 4.6487484 | 5.2925897 | 5.2836356 | 3.8973572 | 4.2748356 | 4.2529655 |
| ASMM10P006 | 0.01916 | 0.3543961 | 1.2665257 | up | NM_009211 | Smarcc1 | SWI/SNF related, matrix associated, actin dependent regulator of chromatin, subfamily c, me | 517.84186 | 494.51759 | 9.265711 | 8.924834 | 586.30634 | 501.7398 | 465.47943 | 481.2627 | 504.63132 | 497.65875 | 9.174801 | 9.351242 | 9.271089 | 8.835083 | 8.86804 | 9.07138 |
| ASMM10P052 | 0.01919 | 0.3543961 | 1.6898131 | up | NM_015786 | Hist1h1c | histone cluster 1, H1c | 16435.628 | 11406.828 | 14.147265 | 13.390401 | 14331.49 | 18516.688 | 16458.705 | 9569.089 | 13069.625 | 11581.769 | 13.867761 | 14.359719 | 14.214314 | 13.120595 | 13.49601 | 13.554598 |
| ASMM10P012 | 0.01924 | 0.3543961 | 1.4877628 | up | NM_001168 | Arhgap39 | Rho GTPase activating protein 39 | 4455.1976 | 3512.9533 | 12.300333 | 11.727188 | 4503.0454 | 3847.079 | 5015.4683 | 3855.065 | 3699.45 | 2984.345 | 12.162653 | 12.165613 | 12.572733 | 11.837131 | 11.733999 | 11.610436 |
| ASMM10P025 | 0.01925 | 0.3543961 | 1.3321621 | up | NM_011197 | Ptgfrn | prostaglandin F2 receptor negative regulator | 22359.941 | 19676.783 | 14.596747 | 14.182978 | 23749.182 | 19523.24 | 23807.402 | 19067.049 | 22848.422 | 17114.879 | 14.624562 | 14.433249 | 14.732431 | 14.104968 | 14.3126 | 14.131365 |
| ASMM10P052 | 0.01929 | 0.3543961 | 1.4312291 | up | NM_027992 | Tmem106b | transmembrane protein 106B | 981.47157 | 822.42835 | 10.173572 | 9.656318 | 1014.2647 | 959.31506 | 970.835 | 760.3035 | 882.69336 | 824.2882 | 9.962132 | 10.254744 | 10.303841 | 9.498761 | 9.678845 | 9.791347 |
| ASMM10P017 | 0.01930 | 0.3543961 | 1.4027732 | up | NM_019421 | Cd320 | CD320 antigen | 98.351987 | 85.656618 | 6.909766 | 6.421484 | 106.98151 | 102.22196 | 85.85249 | 94.518875 | 82.5346 | 79.91638 | 6.788476 | 7.101288 | 6.8395333 | 6.4666443 | 6.2569337 | 6.540874 |
| ASMM10P020 | 0.01933 | 0.3543961 | 1.4234385 | up | NM_009399 | Tnfrsf11a | tumor necrosis factor receptor superfamily, member 11a | 114.77111 | 99.833225 | 7.13159 | 6.622209 | 126.28261 | 117.3245 | 100.70622 | 119.38088 | 104.8432 | 75.2756 | 7.016541 | 7.3007627 | 7.077465 | 6.812098 | 6.5969343 | 6.457596 |
| ASMM10P000 | 0.01935 | 0.3543961 | 1.3916797 | up | NM_021461 | Mknk1 | MAP kinase-interacting serine/threonine kinase 1 | 198.17061 | 171.91027 | 7.896999 | 7.420172 | 251.4333 | 187.10214 | 155.9764 | 174.75373 | 169.46582 | 171.52925 | 7.998241 | 7.968266 | 7.7244906 | 7.369287 | 7.293563 | 7.597666 |
| ASMM10P053 | 0.01935 | 0.3543961 | 1.4850408 | up | NM_025747 | 4933411K20 | RIKEN cDNA 4933411K20 gene | 296.57917 | 243.64633 | 8.489119 | 7.918617 | 295.13196 | 298.15866 | 296.4469 | 238.89926 | 257.6214 | 234.41832 | 8.214905 | 8.613442 | 8.639011 | 7.8213143 | 7.90025 | 8.034286 |
| ASMM10P001 | 0.01937 | 0.3543961 | 1.2709771 | up | NM_001172 | Wdr17 | WD repeat domain 17 | 411.95834 | 395.16181 | 8.951158 | 8.60522 | 453.34296 | 391.7961 | 390.73596 | 446.2824 | 392.90643 | 346.2966 | 8.824697 | 8.998777 | 9.03 | 8.7314 | 8.51227 | 8.57199 |
| ASMM10P050 | 0.01938 | 0.3543961 | 1.265179 | up | NM_172805 | Kcnh5 | potassium voltage-gated channel, subfamily H (eag-related), member 5 | 5569.8769 | 5207.9407 | 12.621242 | 12.281901 | 6532.5107 | 4765.149 | 5411.971 | 5251.097 | 5828.0737 | 4544.6514 | 12.703863 | 12.471118 | 12.688746 | 12.260505 | 12.375271 | 12.209926 |
| ASMM10P015 | 0.01940 | 0.3543961 | 1.3278468 | up | NM_145410 | Fam173a | family with sequence similarity 173, member A | 35655.315 | 32268.644 | 15.300398 | 14.891309 | 33389.707 | 35396.465 | 38179.773 | 32806.97 | 32349.863 | 31649.098 | 15.143812 | 15.298544 | 15.458837 | 14.873253 | 14.800152 | 15.000522 |
| ASMM10P024 | 0.01945 | 0.3543961 | 1.809415 | up | NM_001037 | Defb28 | defensin beta 28 | 14.180652 | 9.249676 | 4.056159 | 3.200636 | 19.913017 | 11.900998 | 10.727941 | 12.439701 | 8.913395 | 6.395931 | 4.347442 | 4.047154 | 3.7738812 | 3.5083277 | 3.0262406 | 3.067339 |
| ASMM10P005 | 0.01945 | 0.3543961 | 1.7408855 | up | NM_022880 | Slc29a1 | solute carrier family 29 (nucleoside transporters), member 1 | 651.18297 | 444.47209 | 9.5724 | 8.772579 | 664.247 | 511.1924 | 778.1095 | 501.35458 | 447.15 | 384.91168 | 9.360085 | 9.380752 | 9.976364 | 8.89816 | 8.701524 | 8.718053 |
| ASMM10P054 | 0.01947 | 0.3543961 | 1.742084 | up | NM_176931 | Syt15 | synaptotagmin XV | 15.403776 | 10.818668 | 4.201535 | 3.40072 | 19.882668 | 13.552419 | 12.77624 | 12.830599 | 13.433629 | 6.1917768 | 4.345517 | 4.2296805 | 4.029406 | 3.551204 | 3.629175 | 3.021782 |
| ASMM10P000 | 0.01947 | 0.3543961 | 1.9667012 | up | ENSMUST( | Acy1 | aminoacylase 1 | 67.157511 | 41.523114 | 6.334256 | 5.358478 | 76.89069 | 45.97208 | 78.609764 | 46.103745 | 50.92846 | 27.537136 | 6.3241134 | 5.968849 | 6.7098045 | 5.434374 | 5.5623646 | 5.078695 |
| ASMM10P001 | 0.01948 | 0.3543961 | 1.5115913 | up | NM_001001 | Atp11c | ATPase, class VI, type 11C | 205.50309 | 168.7328 | 7.978665 | 7.382597 | 206.27382 | 202.23164 | 208.00381 | 163.16667 | 201.61543 | 141.41629 | 7.718321 | 8.07792 | 8.139754 | 7.265934 | 7.5536766 | 7.32818 |
| ASMM10P055 | 0.01952 | 0.3543961 | 1.4689931 | up | NM_028376 | Pfn4 | profilin family, member 4 | 1732.8818 | 1414.1408 | 10.975881 | 10.421053 | 1703.9551 | 1857.1262 | 1637.5642 | 1371.9492 | 1447.1327 | 1423.3406 | 10.727158 | 11.167449 | 11.033035 | 10.33791 | 10.369313 | 10.555936 |
| ASMM10P036 | 0.01962 | 0.3543961 | 1.3601493 | up | NM_026858 | Xrcc6bp1 | XRCC6 binding protein 1 | 456.79187 | 405.78135 | 9.087421 | 8.643656 | 530.0753 | 407.47708 | 432.82324 | 404.6696 | 386.174 | 426.50046 | 9.037604 | 9.058125 | 9.1665325 | 8.585888 | 8.4873705 | 8.857708 |
| ASMM10P010 | 0.01968 | 0.3543961 | 1.3051959 | up | NM_001039 | Zfp808 | zinc finger protein 80 | 1362.9718 | 1252.2579 | 10.630746 | 10.246479 | 1571.4532 | 1110.4847 | 1406.9775 | 1266.5349 | 1347.4733 | 1142.7655 | 10.60307 | 10.470932 | 10.818235 | 10.224575 | 10.269638 | 10.245225 |
| ASMM10P026 | 0.01970 | 0.3543961 | 1.2501143 | up | NM_027147 | Enho | energy homeostasis associated | 69821.897 | 70734.912 | 16.337856 | 16.015796 | 68659.92 | 71019.65 | 69786.12 | 68198.625 | 78511.08 | 65495.03 | 16.22723 | 16.378803 | 16.407534 | 15.889488 | 16.05424 | 16.103659 |
| ASMM10P006 | 0.01971 | 0.3543961 | 1.2930837 | up | NM_001271 | Arrb2 | arrestin, beta 2 | 5927.3061 | 5379.8643 | 12.711612 | 12.340796 | 6171.367 | 5549.2754 | 6061.276 | 5357.323 | 5297.1206 | 5485.1494 | 12.62082 | 12.684147 | 12.829868 | 12.290426 | 12.240567 | 12.491395 |
| ASMM10P017 | 0.01976 | 0.3543961 | 1.3246262 | up | NM_025887 | Rab5a | RAB5A, member RAS oncogene family | 4626.2684 | 4097.3784 | 12.35714 | 11.951554 | 5083.325 | 4190.3833 | 4605.097 | 4068.7039 | 3926.512 | 4296.9194 | 12.335 | 12.279591 | 12.456828 | 11.907302 | 11.814938 | 12.132423 |
| ASMM10P007 | 0.01977 | 0.3543961 | 1.2774094 | up | NM_019637 | Styx | serine/threonine/tyrosine interaction protein | 1298.4013 | 1223.2868 | 10.570329 | 10.217108 | 1396.529 | 1318.8359 | 1179.839 | 1181.8678 | 1286.9894 | 1201.0033 | 10.435003 | 10.702308 | 10.573677 | 10.132123 | 10.203155 | 10.316047 |
| ASMM10P038 | 0.01979 | 0.3543961 | 1.4185605 | up | NM_175533 | 5830411N06 | RIKEN cDNA 5830411N06 gene | 869.42383 | 737.3965 | 9.996295 | 9.491867 | 949.7206 | 908.5518 | 749.9991 | 841.94696 | 685.15344 | 685.0891 | 9.877243 | 10.183333 | 9.928309 | 9.636601 | 9.312622 | 9.526379 |

| | | | | | | | | | | | | | | | | | | | | | | | |
|---|---|---|---|---|---|---|---|---|---|---|---|---|---|---|---|---|---|---|---|---|---|---|---|
| ASMM10P003 | 0.01979 | 0.3543961 | 1.3198601 | up | NM_001170 | Trip4 | thyroid hormone receptor interactor 4 | 419.45555 | 386.40127 | 8.977062 | 8.576677 | 472.11536 | 397.6646 | 388.5867 | 391.2477 | 371.93655 | 396.01956 | 8.883963 | 9.023504 | 9.023719 | 8.534482 | 8.435909 | 8.75964 |
| ASMM10P004 | 0.01983 | 0.3543961 | 1.5336889 | up | NM_001081 | Mdn1 | midasin homolog (yeast) | 65.967842 | 53.111457 | 6.343534 | 5.726528 | 63.72524 | 69.81901 | 64.359276 | 61.460636 | 57.22066 | 40.653076 | 6.05445 | 6.556986 | 6.4191647 | 5.8440127 | 5.7239604 | 5.61161 |
| ASMM10P017 | 0.01984 | 0.3543961 | 2.101901 | up | NM_206958 | Ltbp1 | latent transforming growth factor beta binding protein 1 | 18.486312 | 10.321701 | 4.446489 | 3.374795 | 19.056019 | 12.53923 | 23.863688 | 9.570955 | 13.24518 | 8.148969 | 4.283046 | 4.12304 | 4.9333816 | 3.1200266 | 3.6088047 | 3.3955522 |
| ASMM10P022 | 0.01986 | 0.3543961 | 1.2987236 | up | NM_177784 | Klhl23 | kelch-like 23 | 223.15432 | 210.17981 | 8.08636 | 7.709265 | 259.69366 | 197.1442 | 212.6251 | 216.84064 | 203.94632 | 209.75247 | 8.041641 | 8.043123 | 8.1743145 | 7.6816645 | 7.568528 | 7.8776026 |
| ASMM10P051 | 0.01992 | 0.3543961 | 1.5557267 | up | NM_172645 | Suco | SUN domain containing ossification factor | 1417.9534 | 1086.7843 | 10.691717 | 10.054128 | 1364.2458 | 1452.515 | 1437.0994 | 1121.4067 | 1028.9363 | 1110.0099 | 10.403616 | 10.827041 | 10.844493 | 10.056829 | 9.896822 | 10.208733 |
| ASMM10P028 | 0.01992 | 0.3543961 | 1.4498296 | up | NM_181401 | Tmem64 | transmembrane protein 64 | 324.23815 | 273.21778 | 8.617934 | 8.08205 | 328.00186 | 336.89392 | 307.81866 | 285.91034 | 270.4378 | 263.3052 | 8.370283 | 8.792755 | 8.690763 | 8.084154 | 7.968677 | 8.19332 |
| ASMM10P012 | 0.01993 | 0.3543961 | 1.6940882 | up | NM_145852 | Ropn1l | ropporin 1-like | 667.35324 | 477.42669 | 9.624054 | 8.863545 | 700.78424 | 610.76587 | 690.5096 | 406.5857 | 484.21347 | 541.4809 | 9.443162 | 9.622267 | 9.806734 | 8.592688 | 8.811245 | 9.186703 |
| ASMM10P053 | 0.01996 | 0.3543961 | 1.3257073 | up | NM_027498 | Sik3 | SIK family kinase 3 | 412.24358 | 378.86668 | 8.953301 | 8.546538 | 454.02313 | 388.51917 | 394.18845 | 400.26474 | 360.5012 | 375.8341 | 8.828101 | 8.986833 | 9.044968 | 8.571104 | 8.38505 | 8.683461 |
| ASMM10P008 | 0.02003 | 0.3543961 | 1.2587455 | up | NM_028314 | 2700097O09 | RIKEN cDNA 2700097O09 gene | 495.07072 | 477.51169 | 9.207342 | 8.875356 | 538.1413 | 489.64542 | 457.42545 | 479.21573 | 493.43536 | 459.88397 | 9.0576315 | 9.317823 | 9.2465725 | 8.828704 | 8.836538 | 8.960825 |
| ASMM10P049 | 0.02004 | 0.3543961 | 1.303013 | up | NM_029545 | Dtd2 | D-tyrosyl-tRNA deacylase 2 | 253.90722 | 238.08701 | 8.263627 | 7.881776 | 285.3558 | 265.5652 | 210.80067 | 241.22595 | 264.0095 | 209.02559 | 8.173449 | 8.458288 | 8.159144 | 7.8348126 | 7.9379416 | 7.8725724 |
| ASMM10P005 | 0.02004 | 0.3543961 | 1.3399514 | up | NM_001145 | Smim20 | small integral membrane protein 20 | 1002.3571 | 897.06633 | 10.200215 | 9.778034 | 1071.629 | 1035.048 | 900.3943 | 859.8232 | 949.1281 | 882.2477 | 10.04494 | 10.363197 | 10.192507 | 9.665147 | 9.777784 | 9.891171 |
| ASMM10P000 | 0.02006 | 0.3543961 | 1.8657137 | up | NM_007654 | Cd72 | CD72 antigen | 69.236969 | 44.060837 | 6.380514 | 5.480787 | 73.81675 | 82.437355 | 51.456802 | 43.834915 | 45.00695 | 43.340645 | 6.2642627 | 6.7934113 | 6.0838695 | 5.35899 | 5.3798637 | 5.703507 |
| ASMM10P050 | 0.02007 | 0.3543961 | 1.2645377 | up | NM_145991 | Cdc73 | cell division cycle 73, Paf1/RNA polymerase II complex component | 554.58661 | 528.5504 | 9.361246 | 9.022636 | 605.92896 | 550.28033 | 507.55054 | 551.396 | 526.3536 | 507.9016 | 9.219897 | 9.480315 | 9.383525 | 9.033437 | 8.932736 | 9.101734 |
| ASMM10P010 | 0.02008 | 0.3543961 | 1.8370408 | up | NM_178683 | Depdc1b | DEP domain containing 1B | 407.07389 | 269.57497 | 8.933533 | 8.05615 | 364.47208 | 439.2331 | 417.5165 | 242.46878 | 305.87952 | 260.37662 | 8.51757 | 9.162518 | 9.120512 | 7.841952 | 8.146047 | 8.18045 |
| ASMM10P041 | 0.02010 | 0.3543961 | 1.427649 | up | NM_016742 | Cdc37 | cell division cycle 37 | 1577.6402 | 1328.08 | 10.845224 | 10.331583 | 1574.5134 | 1510.8102 | 1647.597 | 1332.5248 | 1344.4597 | 1307.2555 | 10.606026 | 10.884959 | 11.044688 | 10.297456 | 10.268742 | 10.428551 |
| ASMM10P012 | 0.02012 | 0.3543961 | 1.5032296 | up | NM_175212 | Tmem65 | transmembrane protein 65 | 5572.332 | 4318.5229 | 12.61196 | 12.023895 | 5386.673 | 4878.275 | 6452.048 | 4205.1123 | 4744.9033 | 4005.5532 | 12.419299 | 12.503021 | 12.913561 | 11.951872 | 12.077121 | 12.042692 |
| ASMM10P011 | 0.02018 | 0.3543961 | 1.8337403 | up | NM_008125 | Gjb2 | gap junction protein, beta 2 | 5970.9211 | 3843.1553 | 12.72194 | 11.847151 | 5643.8145 | 5850.2524 | 6418.6963 | 3285.1328 | 3615.259 | 4629.074 | 12.49355 | 12.764405 | 12.907866 | 11.595033 | 11.703956 | 12.242464 |
| ASMM10P053 | 0.02020 | 0.3543961 | 2.0707085 | up | NM_023608 | Gdpd2 | glycerophosphodiester phosphodiesterase domain containing 2 | 134.55987 | 78.189903 | 7.342824 | 6.2927 | 120.2504 | 171.12392 | 112.3053 | 75.07886 | 81.93011 | 77.56074 | 6.953613 | 7.8362694 | 7.238591 | 6.134941 | 6.2469583 | 6.4962006 |
| ASMM10P035 | 0.02021 | 0.3543961 | 3.4354303 | up | NM_206872 | Vmn1r177 | vomeronasal 1 receptor 177 | 106.40701 | 34.497277 | 6.872705 | 5.092214 | 147.80888 | 46.024364 | 125.38779 | 42.361485 | 38.451805 | 22.67854 | 7.239897 | 5.9707575 | 7.407461 | 5.3088465 | 5.1551266 | 4.81267 |
| ASMM10P012 | 0.02021 | 0.3543961 | 2.0472333 | up | NM_011105 | Pkdrej | polycystic kidney disease (polycystin) and REJ (sperm receptor for egg jelly homolog, sea ur | 1121.9048 | 634.70755 | 10.313393 | 9.279718 | 1159.5121 | 763.7541 | 1442.4481 | 648.6516 | 686.2304 | 569.24066 | 10.153679 | 9.93535 | 10.851151 | 9.263586 | 9.315583 | 9.259984 |
| ASMM10P048 | 0.02022 | 0.3543961 | 1.5421043 | up | NM_177369 | Myh8 | myosin, heavy polypeptide 8, skeletal muscle, perinatal | 106.74926 | 82.80732 | 6.99399 | 6.36909 | 142.46835 | 87.07031 | 90.70912 | 90.48876 | 75.16621 | 82.76699 | 7.187169 | 6.87253 | 6.9222717 | 6.4001937 | 6.1197376 | 6.5873384 |
| ASMM10P004 | 0.02022 | 0.3543961 | 1.2519463 | up | NM_001024 | Nufip2 | nuclear fragile X mental retardation protein interacting protein 2 | 601.00584 | 577.79278 | 9.471384 | 9.147211 | 671.59515 | 617.15717 | 514.2652 | 582.66833 | 604.6042 | 546.1058 | 9.376621 | 9.636385 | 9.401145 | 9.112299 | 9.131125 | 9.198209 |
| ASMM10P045 | 0.02028 | 0.3543961 | 1.8142572 | up | NM_029316 | 1700020N15 | RIKEN cDNA 1700020N15 gene | 405.1992 | 265.16958 | 8.898552 | 8.039173 | 440.17426 | 294.2017 | 481.22165 | 273.81747 | 258.503 | 263.18826 | 8.785291 | 8.596535 | 9.313831 | 8.018909 | 7.905666 | 8.192945 |
| ASMM10P049 | 0.02035 | 0.3543961 | 1.3382936 | up | NM_001110 | 1810043H04 | RIKEN cDNA 1810043H04 gene | 884.48097 | 787.22005 | 10.014647 | 9.594252 | 1008.2077 | 823.2269 | 822.0083 | 735.2961 | 792.3345 | 834.02954 | 9.950866 | 10.038399 | 10.054676 | 9.453562 | 9.522461 | 9.806734 |
| ASMM10P047 | 0.02036 | 0.3543961 | 1.3222186 | up | NM_080420 | Lpo | lactoperoxidase | 75.762063 | 69.28869 | 6.525745 | 6.122785 | 89.96137 | 74.32146 | 63.00336 | 72.71414 | 69.61512 | 65.53681 | 6.543164 | 6.648406 | 6.3856664 | 6.092496 | 6.007912 | 6.2679462 |
| ASMM10P007 | 0.02039 | 0.3543961 | 1.4665731 | up | NM_020594 | Zc3h8 | zinc finger CCCH type containing 8 | 606.72726 | 497.09049 | 9.483324 | 8.930875 | 672.8696 | 536.80505 | 610.50714 | 511.4787 | 450.0817 | 529.71106 | 9.38026 | 9.442763 | 9.626949 | 8.925415 | 8.711084 | 9.156127 |
| ASMM10P048 | 0.02040 | 0.3543961 | 1.2599812 | up | NM_024226 | Rtn4 | reticulon 4 | 1547.6451 | 1481.2293 | 10.817919 | 10.484516 | 1631.92 | 1527.1791 | 1483.8363 | 1597.1404 | 1486.1471 | 1360.4005 | 10.660594 | 10.899404 | 10.893758 | 10.55687 | 10.409053 | 10.487627 |
| ASMM10P005 | 0.02041 | 0.3543961 | 1.3074258 | up | NM_001165 | Spopl | speckle-type POZ protein-like | 93.068933 | 86.815962 | 6.827152 | 6.440423 | 107.18572 | 79.72263 | 92.29845 | 85.10972 | 91.83097 | 83.507195 | 6.7923703 | 6.7426786 | 6.946407 | 6.3129973 | 6.4097834 | 6.598488 |
| ASMM10P026 | 0.02046 | 0.3543961 | 1.9491257 | up | NM_001025 | Camk2d | calcium/calmodulin-dependent protein kinase II, delta | 376.46624 | 237.7935 | 8.819333 | 7.856506 | 443.12817 | 342.01443 | 344.25613 | 245.12076 | 183.30959 | 284.95016 | 8.794854 | 8.811444 | 8.8517 | 7.8568926 | 7.409081 | 8.303543 |
| ASMM10P009 | 0.02050 | 0.3543961 | 1.4108328 | up | NM_172507 | Sh3bgrl2 | SH3 domain binding glutamic acid-rich protein like 2 | 74.41494 | 64.399008 | 6.515363 | 6.018816 | 79.23117 | 72.692635 | 71.321014 | 64.77631 | 64.34507 | 64.075645 | 6.365856 | 6.6148095 | 6.565423 | 5.925384 | 5.893588 | 6.2374754 |
| ASMM10P006 | 0.02051 | 0.3543961 | 1.3262585 | up | NM_001242 | Slc4a10 | solute carrier family 4, sodium bicarbonate cotransporter-like, member 10 | 728.16473 | 656.15898 | 9.737555 | 9.330193 | 856.05743 | 610.55273 | 717.88403 | 687.1087 | 622.1935 | 659.17474 | 9.732068 | 9.620942 | 9.859654 | 9.349046 | 9.174211 | 9.467321 |
| ASMM10P053 | 0.02059 | 0.3543961 | 1.584025 | up | NM_172774 | Atrip | ATR interacting protein | 38.168029 | 28.942241 | 5.541169 | 4.877574 | 44.171665 | 37.02128 | 33.311142 | 26.030142 | 31.391695 | 29.404886 | 5.529583 | 5.660926 | 5.4329977 | 4.5935035 | 4.870696 | 5.168522 |
| ASMM10P003 | 0.02060 | 0.3543961 | 1.4161794 | up | NM_025826 | Acadsb | acyl-Coenzyme A dehydrogenase, short/branched chain | 590.31278 | 500.32292 | 9.445542 | 8.943538 | 600.8574 | 575.0013 | 595.07965 | 492.19907 | 513.0645 | 495.7052 | 9.208184 | 9.533732 | 9.59471 | 8.869918 | 8.893231 | 9.067465 |
| ASMM10P040 | 0.02062 | 0.3543961 | 1.7200346 | up | NM_001159 | Mcf2l | mcf.2 transforming sequence-like | 27.453264 | 19.138213 | 5.059914 | 4.277477 | 24.971565 | 30.951414 | 26.436813 | 20.474857 | 22.237043 | 14.70274 | 4.6896787 | 5.400589 | 5.0894756 | 4.2442913 | 4.367923 | 4.2202163 |
| ASMM10P013 | 0.02062 | 0.3543961 | 1.2735624 | up | NM_183256 | Cox14 | cytochrome c oxidase assembly protein 14 | 24722.724 | 22636.891 | 14.741189 | 14.392319 | 23305.238 | 26937.459 | 23925.475 | 23621.594 | 22707.264 | 21581.816 | 14.602119 | 14.880143 | 14.741305 | 14.397306 | 14.3052 | 14.474452 |
| ASMM10P008 | 0.02069 | 0.3543961 | 2.0208534 | up | NM_144872 | Eml3 | echinoderm microtubule associated protein like 3 | 79.364918 | 47.458029 | 6.59721 | 5.582245 | 74.496414 | 67.11689 | 96.48145 | 45.29181 | 46.556377 | 50.5259 | 6.275993 | 6.502254 | 7.013382 | 5.406651 | 5.427237 | 5.912847 |
| ASMM10P039 | 0.02070 | 0.3543961 | 1.3438263 | up | NM_198095 | Bst2 | bone marrow stromal cell antigen 2 | 1312.3661 | 1167.4334 | 10.577752 | 10.151406 | 1540.5273 | 1207.9629 | 1188.608 | 1041.2538 | 1232.5295 | 1228.5168 | 10.573432 | 10.578289 | 10.581536 | 9.953295 | 10.149352 | 10.35157 |
| ASMM10P016 | 0.02071 | 0.3543961 | 1.3594009 | up | NM_028887 | Smchd1 | SMC hinge domain containing 1 | 331.13151 | 298.62398 | 8.64693 | 8.203959 | 340.99814 | 319.79086 | 332.60553 | 324.41202 | 305.02682 | 266.4331 | 8.423836 | 8.715238 | 8.801715 | 8.261651 | 8.141195 | 8.20903 |
| ASMM10P019 | 0.02072 | 0.3543961 | 1.7505743 | up | NM_001001 | Cyp2c44 | cytochrome P450, family 2, subfamily c, polypeptide 44 | 244.0131 | 172.86066 | 8.218163 | 7.410334 | 245.86725 | 241.1927 | 244.97934 | 204.14069 | 144.2443 | 170.19699 | 7.966026 | 8.319886 | 8.368576 | 7.5907774 | 7.0558558 | 7.58437 |
| ASMM10P021 | 0.02076 | 0.3543961 | 1.3584609 | up | NM_010434 | Hipk3 | homeodomain interacting protein kinase 3 | 47563.764 | 42673.648 | 15.740563 | 15.29859 | 44331.27 | 46354.953 | 52005.07 | 42723.94 | 48745.45 | 36551.555 | 15.581576 | 15.689105 | 15.951008 | 15.256401 | 15.39225 | 15.247119 |
| ASMM10P017 | 0.02076 | 0.3543961 | 1.393717 | up | NM_026165 | Slc25a46 | solute carrier family 25, member 46 | 798.79583 | 688.94378 | 9.877988 | 9.39905 | 841.6359 | 707.145 | 847.6066 | 658.2917 | 749.55304 | 658.9866 | 9.707783 | 9.824647 | 10.101534 | 9.288792 | 9.442323 | 9.466036 |
| ASMM10P012 | 0.02078 | 0.3543961 | 1.5767553 | up | NM_020490 | Ltb4r2 | leukotriene B4 receptor 2 | 97.537766 | 73.98703 | 6.870994 | 6.214035 | 121.41087 | 70.450005 | 100.75243 | 70.72544 | 81.62821 | 69.60744 | 6.9652195 | 6.5694575 | 7.078304 | 6.050709 | 6.240713 | 6.3506827 |
| ASMM10P007 | 0.02079 | 0.3543961 | 1.3261714 | up | NM_011356 | Frzb | frizzled-related protein | 117.59064 | 108.7282 | 7.167495 | 6.760228 | 132.39323 | 105.90059 | 114.4781 | 102.54392 | 118.65247 | 104.98822 | 7.0796137 | 7.1538763 | 7.2689962 | 6.5890293 | 6.773625 | 6.91803 |
| ASMM10P047 | 0.02080 | 0.3543961 | 1.2858322 | up | NM_025560 | Oxld1 | oxidoreductase like domain containing 1 | 9237.7253 | 8428.8997 | 13.339931 | 12.977229 | 9912.485 | 8717.311 | 9083.38 | 8192.13 | 8315.533 | 8779.036 | 13.320627 | 13.29982 | 13.399346 | 12.895895 | 12.872714 | 13.163077 |
| ASMM10P037 | 0.02084 | 0.3543961 | 4.0742334 | up | NM_001166 | Vmn1r168 | vomeronasal 1 receptor 168 | 151.09677 | 44.392979 | 7.394553 | 5.368025 | 195.77948 | 69.638596 | 187.87222 | 43.256927 | 67.50582 | 22.461191 | 7.643431 | 6.5530796 | 7.987149 | 5.3427167 | 5.962844 | 4.798513 |
| ASMM10P054 | 0.02085 | 0.3543961 | 1.441453 | up | NM_026712 | Zfp414 | zinc finger protein 414 | 15428.171 | 12549.91 | 14.062278 | 13.534754 | 16488.314 | 13850.051 | 15946.147 | 12259.047 | 11905.165 | 13485.522 | 14.075224 | 13.944293 | 14.167316 | 13.451289 | 13.368575 | 13.784397 |
| ASMM10P000 | 0.02087 | 0.3543961 | 1.738378 | up | NM_001167 | Zfp568 | zinc finger protein 568 | 462.5844 | 316.96571 | 9.089237 | 8.291495 | 452.2125 | 374.2336 | 561.3071 | 332.82922 | 332.31082 | 285.7571 | 8.819979 | 8.932736 | 9.514996 | 8.299577 | 8.267462 | 8.3074465 |
| ASMM10P005 | 0.02090 | 0.3543961 | 1.2862997 | up | NM_001177 | Wbp1l | WW domain binding protein 1 like | 33.208941 | 31.173241 | 5.345632 | 4.982405 | 34.450066 | 33.046516 | 32.13024 | 34.179443 | 34.08582 | 25.254461 | 5.162702 | 5.4951053 | 5.3790874 | 5.0013027 | 4.9883766 | 4.957535 |
| ASMM10P022 | 0.02094 | 0.3543961 | 1.43728 | up | NM_175551 | Dido1 | death inducer-obliterator 1 | 20959.836 | 17087.465 | 14.502775 | 13.979434 | 21828.93 | 18939.996 | 22110.582 | 16544.738 | 16473.998 | 18243.658 | 14.504127 | 14.38905 | 14.61515 | 13.889693 | 13.820864 | 14.227746 |
| ASMM10P017 | 0.02095 | 0.3543961 | 1.4745676 | up | NM_207530 | Osbpl1a | oxysterol binding protein-like 1A | 10790.146 | 8593.5496 | 13.543104 | 12.982812 | 10895.938 | 9536.638 | 11937.861 | 9879.169 | 9194.907 | 6706.5728 | 13.454062 | 13.430421 | 13.74483 | 13.161145 | 13.014097 | 12.773195 |
| ASMM10P034 | 0.02097 | 0.3543961 | 1.4553477 | up | NM_025591 | Fam136a | family with sequence similarity 136, member A | 14042.361 | 11270.108 | 13.92396 | 13.382597 | 13755.762 | 13384.493 | 14986.829 | 11252.42 | 10593.268 | 11964.635 | 13.789908 | 13.902191 | 14.079783 | 13.329792 | 13.206426 | 13.611572 |
| ASMM10P054 | 0.02098 | 0.3543961 | 1.2627875 | up | NM_144788 | Hectd1 | HECT domain containing 1 | 368.44024 | 357.87637 | 8.796439 | 8.459827 | 403.57434 | 337.54907 | 364.1973 | 383.0838 | 393.46655 | 297.07877 | 8.6657295 | 8.795416 | 8.928171 | 8.505676 | 8.515267 | 8.358538 |
| ASMM10P033 | 0.02103 | 0.3543961 | 2.0445191 | up | NM_207210 | Dyrk4 | dual-specificity tyrosine-(Y)-phosphorylation regulated kinase 4 | 231.21217 | 136.22914 | 8.114493 | 7.082731 | 209.30287 | 185.54164 | 298.792 | 149.74258 | 136.8696 | 122.07524 | 7.7398734 | 7.953716 | 8.649889 | 7.1402783 | 6.9785404 | 7.129375 |
| ASMM10P015 | 0.02104 | 0.3543961 | 1.443685 | up | NM_011581 | Thbs2 | thrombospondin 2 | 3089.7495 | 2526.6027 | 11.784233 | 11.254477 | 3210.9949 | 2749.9905 | 3308.2632 | 2298.309 | 2713.803 | 2567.696 | 11.63574 | 11.712593 | 12.004366 | 11.086521 | 11.284704 | 11.392206 |
| ASMM10P005 | 0.02107 | 0.3543961 | 1.2851149 | up | NM_001160 | Cldn10 | claudin 10 | 476.7073 | 450.78966 | 9.157278 | 8.795381 | 506.22867 | 448.91898 | 474.97424 | 449.33167 | 481.02542 | 422.0119 | 8.97992 | 9.195179 | 9.296736 | 8.740173 | 8.801902 | 8.844068 |
| ASMM10P039 | 0.02112 | 0.3543961 | 1.7095656 | up | NM_001039 | Scoc | short coiled-coil protein | 94.606238 | 67.794654 | 6.853485 | 6.079855 | 104.29099 | 78.82355 | 100.70418 | 56.772083 | 79.10048 | 67.5114 | 6.7562547 | 6.7268577 | 7.077342 | 5.731434 | 6.195678 | 6.312453 |
| ASMM10P015 | 0.02113 | 0.3543961 | 1.7204231 | up | NM_026400 | Dnajb11 | DnaJ (Hsp40) homolog, subfamily B, member 11 | 90.289988 | 63.490618 | 6.774549 | 5.991785 | 101.73959 | 68.9921 | 100.13828 | 73.82073 | 56.26222 | 60.388905 | 6.7173085 | 6.538764 | 7.0675735 | 6.1132917 | 5.7029552 | 6.159109 |
| ASMM10P015 | 0.02115 | 0.3543961 | 1.51707 | up | NM_145517 | Ormdl1 | ORM1-like 1 (S. cerevisiae) | 290.10619 | 232.56821 | 8.455194 | 7.853906 | 294.62906 | 297.219 | 278.47052 | 235.49193 | 224.02681 | 238.18588 | 8.211769 | 8.608981 | 8.544832 | 7.800453 | 7.700738 | 8.060528 |
| ASMM10P038 | 0.02121 | 0.3543961 | 2.864969 | up | NM_146926 | Olfr477 | olfactory receptor 477 | 41.527337 | 16.001873 | 5.543469 | 4.02495 | 54.62672 | 19.51425 | 50.44104 | 18.809 | 15.65395 | 13.542669 | 5.830451 | 4.7464104 | 6.0535464 | 4.116441 | 3.8539822 | 4.104426 |
| ASMM10P042 | 0.02122 | 0.3543961 | 1.3309151 | up | NM_001033 | Gucy1a2 | guanylate cyclase 1, soluble, alpha 2 | 285.75815 | 260.42435 | 8.424391 | 8.011972 | 345.74286 | 269.828 | 241.7036 | 282.8612 | 241.72574 | 256.6861 | 8.44441 | 8.47926 | 8.349503 | 8.065834 | 7.8089514 | 8.161132 |
| ASMM10P014 | 0.02127 | 0.3543961 | 1.393685 | up | NM_008365 | Il18r1 | interleukin 18 receptor 1 | 146.7885 | 129.33014 | 7.491259 | 7.012354 | 156.38306 | 148.24025 | 135.74219 | 129.1741 | 130.3281 | 128.48822 | 7.32206 | 7.6292 | 7.522516 | 6.927053 | 6.9096384 | 7.200371 |
| ASMM10P010 | 0.02130 | 0.3543961 | 1.5514985 | up | NM_001029 | Fcrlb | Fc receptor-like B | 99.271467 | 77.486027 | 6.909266 | 6.275603 | 111.26991 | 110.19788 | 76.34661 | 75.86205 | 88.10559 | 68.49044 | 6.8456283 | 7.213261 | 6.668908 | 6.149243 | 6.3486795 | 6.328888 |
| ASMM10P004 | 0.02137 | 0.3543961 | 1.5413035 | up | NM_001034 | Tnfsf12Tnfs | Tnfsf12-Tnfsf13 readthrough transcript | 192.03953 | 153.35476 | 7.871115 | 7.246964 | 201.08769 | 162.77863 | 212.25227 | 147.56009 | 180.0734 | 132.43079 | 7.678534 | 7.7638655 | 8.170944 | 7.118909 | 7.3828726 | 7.239109 |
| ASMM10P008 | 0.02137 | 0.3543961 | 1.6255944 | up | NM_028967 | Batf2 | basic leucine zipper transcription factor, ATF-like 2 | 4439.1952 | 3196.7552 | 12.293252 | 11.592284 | 4628.7373 | 3495.2737 | 5193.5747 | 3347.132 | 3028.1648 | 3214.9688 | 12.204494 | 12.045038 | 12.630223 | 11.622612 | 11.446065 | 11.708176 |
| ASMM10P043 | 0.02140 | 0.3543961 | 1.3076414 | up | NM_010498 | Ids | iduronate 2-sulfatase | 33799.872 | 30680.336 | 15.2171 | 14.830133 | 33961.574 | 31157.562 | 36280.48 | 29481.898 | 32455.89 | 30103.22 | 15.166219 | 15.10083 | 15.384251 | 14.7359 | 14.80882 | 14.945679 |
| ASMM10P052 | 0.02147 | 0.3543961 | 1.418724 | up | NM_134071 | Ankrd32 | ankyrin repeat domain 32 | 8497.7409 | 7005.0608 | 13.219189 | 12.714595 | 8124.7715 | 7765.9873 | 9602.464 | 6922.1377 | 7383.9976 | 6709.047 | 13.031441 | 13.14887 | 13.477257 | 12.65663 | 12.713089 | 12.774067 |
| ASMM10P016 | 0.02150 | 0.3543961 | 1.3377568 | up | NM_001008 | Zfp760 | zinc finger protein 760 | 113.84837 | 104.55063 | 7.107302 | 6.687486 | 138.6245 | 107.4736 | 95.447014 | 104.963 | 127.85033 | 80.83856 | 7.1489997 | 7.175462 | 6.997443 | 6.6239166 | 6.883955 | 6.5545855 |
| ASMM10P015 | 0.02152 | 0.3543961 | 1.7007365 | up | NM_153390 | Pxt1 | peroxisomal, testis specific 1 | 23.875326 | 17.514179 | 4.859235 | 4.093076 | 27.081198 | 20.941181 | 23.6036 | 22.863821 | 19.475737 | 10.202979 | 4.8081927 | 4.8512764 | 4.9182367 | 4.4022126 | 4.176166 | 3.7008483 |
| ASMM10P006 | 0.02156 | 0.3543961 | 2.6148067 | up | NM_009988 | Cxadr | coxsackie virus and adenovirus receptor | 163.99245 | 74.037182 | 7.5843 | 6.197596 | 201.90144 | 98.69725 | 191.37865 | 63.007465 | 69.77529 | 89.32879 | 7.683285 | 7.0541043 | 8.0155115 | 5.883075 | 6.0116405 | 6.6980724 |
| ASMM10P030 | 0.02159 | 0.3543961 | 1.4568274 | up | NM_001033 | Dhrsx | dehydrogenase/reductase (SDR family) X chromosome | 259.71259 | 217.823 | 8.302906 | 7.760076 | 264.79846 | 243.24336 | 271.09595 | 215.4641 | 223.42038 | 214.58453 | 8.067951 | 8.332133 | 8.508634 | 7.673496 | 7.697585 | 7.909147 |
| ASMM10P005 | 0.02159 | 0.3543961 | 1.4402519 | up | NM_001190 | Snx7 | sorting nexin 7 | 1082.416 | 902.65979 | 10.315241 | 9.78892 | 1092.452 | 1051.107 | 1103.6891 | 972.3996 | 863.1483 | 872.43146 | 10.073992 | 10.385095 | 10.486636 | 9.847542 | 9.644516 | 9.8747015 |
| ASMM10P048 | 0.02160 | 0.3543961 | 1.3896845 | up | NM_145429 | Arrb2 | arrestin, beta 2 | 6395.8279 | 5408.5101 | 12.817296 | 12.342539 | 6180.5527 | 6119.3457 | 6887.5854 | 5589.2266 | 5253.642 | 5382.6616 | 12.621921 | 12.824047 | 13.00592 | 12.350552 | 12.222055 | 12.455009 |
| ASMM10P047 | 0.02161 | 0.3543961 | 1.4260384 | up | NM_173755 | Ube2o | ubiquitin-conjugating enzyme E2O | 1129.3185 | 931.0888 | 10.34555 | 9.833538 | 1457.1841 | 861.5374 | 1069.2339 | 1026.2261 | 904.2725 | 862.7678 | 10.497503 | 10.097604 | 10.441544 | 9.9286375 | 9.714658 | 9.857317 |
| ASMM10P000 | 0.02161 | 0.3543961 | 3.6184094 | up | NM_001136 | Klrc1 | killer cell lectin-like receptor subfamily C, member 1 | 20.466406 | 5.563328 | 4.449399 | 2.594044 | 35.206482 | 9.140041 | 17.052696 | 4.9999995 | 4.9999995 | 6.689984 | 5.196107 | 3.7050989 | 4.446992 | 2.3250442 | 2.3265402 | 3.1305466 |
| ASMM10P039 | 0.02164 | 0.3543961 | 1.8037768 | up | NM_001080 | Rab11fip1 | RAB11 family interacting protein 1 (class I) | 275.64494 | 186.98067 | 8.376608 | 7.525588 | 266.13123 | 231.98938 | 328.8142 | 211.86671 | 206.54153 | 142.53377 | 8.076109 | 8.266939 | 8.786777 | 7.649806 | 7.5883775 | 7.338579 |
| ASMM10P004 | 0.02166 | 0.3543961 | 1.404662 | up | NM_001080 | Pdpk1 | 3-phosphoinositide dependent protein kinase 1 | 32576.476 | 27668.629 | 15.162006 | 14.671783 | 30189.19 | 32153.086 | 35387.152 | 26198.586 | 28422.305 | 28384.996 | 15.005062 | 15.137671 | 15.343285 | 14.552685 | 14.611919 | 14.850745 |
| ASMM10P007 | 0.02169 | 0.3543961 | 1.289898 | up | NM_010251 | Gabra4 | gamma-aminobutyric acid (GABA) A receptor, subunit alpha 4 | 699.95873 | 653.7022 | 9.692817 | 9.32556 | 732.7187 | 689.497 | 677.6605 | 678.82446 | 658.01105 | 624.2711 | 9.508163 | 9.788678 | 9.781611 | 9.333996 | 9.253061 | 9.389624 |
| ASMM10P030 | 0.02176 | 0.3543961 | 1.5333557 | up | NM_145211 | Oas1a | 2'-5' oligoadenylate synthetase 1A | 271.81516 | 219.73517 | 8.362142 | 7.745449 | 298.28088 | 279.2091 | 237.95549 | 196.50471 | 280.9422 | 181.7586 | 8.229629 | 8.528122 | 8.328674 | 7.5389237 | 8.023384 | 7.67404 |
| ASMM10P041 | 0.02178 | 0.3543961 | 1.7566578 | up | NM_179796 | Syncrip | synaptotagmin binding, cytoplasmic RNA interacting protein | 98.066997 | 67.64525 | 6.89214 | 6.079307 | 102.73321 | 114.99502 | 76.47276 | 61.835896 | 80.531006 | 60.568848 | 6.7329545 | 7.272711 | 6.670755 | 5.8531246 | 6.2217264 | 6.16307 |
| ASMM10P009 | 0.02182 | 0.3543961 | 1.4439052 | up | NM_178916 | Rfesd | Rieske (Fe-S) domain containing | 664.25736 | 553.23518 | 9.609623 | 9.079647 | 774.47943 | 633.05695 | 585.2357 | 526.42834 | 520.0521 | 613.2251 | 9.589977 | 9.668182 | 9.570709 | 8.962654 | 8.913632 | 9.362654 |
| ASMM10P021 | 0.02186 | 0.3543961 | 1.6588432 | up | NM_028119 | Ddb2 | damage specific DNA binding protein 2 | 60.034708 | 44.882412 | 6.206633 | 5.476455 | 56.88826 | 58.019238 | 65.196625 | 52.98725 | 50.338993 | 31.320993 | 5.8867936 | 6.2954693 | 6.4376354 | 5.6287613 | 5.5458064 | 5.254798 |
| ASMM10P004 | 0.02188 | 0.3543961 | 1.2759333 | up | NM_001081 | Taf1 | TAF1 RNA polymerase II, TATA box binding protein (TBP)-associated factor | 241.60837 | 234.20056 | 8.202258 | 7.850705 | 263.0985 | 236.35982 | 225.36679 | 259.75635 | 253.95416 | 188.89116 | 8.058613 | 8.291762 | 8.256399 | 7.9438925 | 7.879071 | 7.7291517 |
| ASMM10P014 | 0.02189 | 0.3543961 | 1.5595771 | up | NM_011782 | Adamts5 | a disintegrin-like and metallopeptidase (reprolysin type) with thrombospondin type 1 motif, : | 73.902233 | 57.871866 | 6.505138 | 5.863983 | 79.18253 | 68.68951 | 73.83466 | 56.879425 | 55.218082 | 61.51809 | 6.3653116 | 6.532179 | 6.617923 | 5.7353024 | 5.6754255 | 6.181221 |
| ASMM10P025 | 0.02190 | 0.3543961 | 1.4724186 | up | NM_011828 | Hs2st1 | heparan sulfate 2-O-sulfotransferase 1 | 4055.1722 | 3241.8607 | 12.16755 | 11.609363 | 4068.841 | 3512.1416 | 4584.534 | 3137.4287 | 3366.528 | 3221.6255 | 11.997952 | 12.053563 | 12.451137 | 11.519593 | 11.596938 | 11.711557 |
| ASMM10P003 | 0.02191 | 0.3543961 | 1.2539432 | up | NM_018860 | Rpl41 | ribosomal protein L41 | 24498.512 | 22942.386 | 14.734422 | 14.40795 | 25701.275 | 23247.754 | 24546.506 | 21939.146 | 23716.236 | 23171.777 | 14.743409 | 14.668524 | 14.791334 | 14.304398 | 14.34916 | 14.570293 |

| | | | | | | | | | | | | | | | | | | | | | | | |
|---|---|---|---|---|---|---|---|---|---|---|---|---|---|---|---|---|---|---|---|---|---|---|---|
| ASMM10P043 | 0.02195 | 0.3543961 | 1.5102112 | up | NM_028173 | Tram1 | translocating chain-associating membrane protein 1 | 10897.555 | 8379.5519 | 13.559964 | 12.965214 | 10598.654 | 9751.572 | 12342.44 | 7638.1616 | 8668.816 | 8831.678 | 13.417438 | 13.463314 | 13.79914 | 12.790612 | 12.935203 | 13.169826 |
| ASMM10P030 | 0.02196 | 0.3543961 | 1.4333002 | up | NM_001042 | Htra3 | HtrA serine peptidase 3 | 260.86823 | 222.20346 | 8.30578 | 7.786439 | 272.6249 | 231.19415 | 278.78564 | 214.5966 | 241.87119 | 210.14258 | 8.109163 | 8.26108 | 8.547096 | 7.668798 | 7.810083 | 7.8804355 |
| ASMM10P007 | 0.02198 | 0.3543961 | 1.4543023 | up | NM_013703 | Vldlr | very low density lipoprotein receptor | 731.14462 | 610.52322 | 9.755133 | 9.214805 | 773.8157 | 726.37177 | 693.2464 | 533.23694 | 727.26636 | 571.06635 | 9.587995 | 9.865174 | 9.812229 | 8.9834795 | 9.396393 | 9.264544 |
| ASMM10P041 | 0.02201 | 0.3543961 | 1.7983953 | up | NM_175206 | Fbxl22 | F-box and leucine-rich repeat protein 22 | 88.803594 | 57.716281 | 6.7092 | 5.862489 | 122.71583 | 60.241642 | 83.45331 | 57.446922 | 56.15352 | 59.5484 | 6.9797564 | 6.3494215 | 6.798421 | 5.7502384 | 5.699974 | 6.137256 |
| ASMM10P045 | 0.02203 | 0.3543961 | 1.7650083 | up | NM_029536 | Gpr165 | G protein-coupled receptor 165 | 128.39692 | 89.659408 | 7.296565 | 6.47689 | 142.31502 | 110.60222 | 132.27351 | 74.896065 | 94.94796 | 99.1342 | 7.1857414 | 7.2181206 | 7.485834 | 6.132103 | 6.4592547 | 6.8393135 |
| ASMM10P037 | 0.02204 | 0.3543961 | 1.3243765 | up | NM_028166 | 1600014C1( | RIKEN cDNA 1600014C10 gene | 18781.414 | 16676.421 | 14.350558 | 13.945245 | 18060.637 | 19070.969 | 19212.635 | 16606.492 | 16488.336 | 16934.436 | 14.215691 | 14.401163 | 14.434821 | 13.893158 | 13.825289 | 14.117288 |
| ASMM10P040 | 0.02205 | 0.3543961 | 1.4873491 | up | NM_001277 | Ank1 | ankyrin 1, erythroid | 108.81886 | 90.133662 | 7.054977 | 6.482233 | 113.49862 | 120.7834 | 92.17455 | 95.51183 | 103.88898 | 71.000175 | 6.8758063 | 7.344372 | 6.9447517 | 6.4837384 | 6.583754 | 6.3792076 |
| ASMM10P021 | 0.02208 | 0.3543961 | 1.4361957 | up | NM_029025 | Tmem81 | transmembrane protein 81 | 1423.166 | 1189.0782 | 10.70259 | 10.180337 | 1435.0431 | 1325.565 | 1508.8899 | 1179.6729 | 1199.7139 | 1187.8478 | 10.473128 | 10.710884 | 10.923757 | 10.129436 | 10.109902 | 10.301674 |
| ASMM10P032 | 0.02208 | 0.3543961 | 1.29983 | up | NM_010338 | Gpr37 | G protein-coupled receptor 37 | 34450.466 | 31745.925 | 15.246679 | 14.868356 | 34308.797 | 33238.902 | 35803.7 | 28794.53 | 35721.168 | 30722.078 | 15.181187 | 15.20217 | 15.356681 | 14.692411 | 14.945679 | 14.966979 |
| ASMM10P024 | 0.02213 | 0.3543961 | 1.3513888 | up | NM_001161 | Myo1b | myosin IB | 189.79509 | 172.57695 | 7.854548 | 7.420105 | 213.73648 | 163.93898 | 191.70981 | 189.32336 | 164.8393 | 163.56818 | 7.770328 | 7.7749743 | 8.018342 | 7.484952 | 7.2474875 | 7.5278764 |
| ASMM10P008 | 0.02218 | 0.3543961 | 1.2810665 | up | NM_145363 | Tspan10 | tetraspanin 10 | 2759.9555 | 2537.8566 | 11.613592 | 11.256247 | 3524.0789 | 2333.3726 | 2422.415 | 2785.7324 | 2550.7498 | 2277.0876 | 11.777259 | 11.493603 | 11.569914 | 11.355502 | 11.194128 | 11.21911 |
| ASMM10P024 | 0.02220 | 0.3543961 | 1.9550622 | up | NM_009047 | Rem1 | rad and gem related GTP binding protein 1 | 13.812537 | 8.38703 | 4.067114 | 3.099899 | 13.264761 | 13.67797 | 14.49488 | 8.700802 | 7.5942197 | 8.866067 | 3.7465394 | 4.241072 | 4.2137303 | 2.9652212 | 2.8174348 | 3.5170422 |
| ASMM10P010 | 0.02220 | 0.3543961 | 1.2616059 | up | NM_001253 | Kcnma1 | potassium large conductance calcium-activated channel, subfamily M, alpha member 1 | 2013.1932 | 1913.4273 | 11.18811 | 10.852849 | 2163.7056 | 2010.8435 | 1865.0305 | 2111.1492 | 1870.1725 | 1758.9602 | 11.06328 | 11.283709 | 11.217341 | 10.963904 | 10.737452 | 10.85719 |
| ASMM10P022 | 0.02232 | 0.3546919 | 1.4337539 | up | NM_001164 | Tank | TRAF family member-associated Nf-kappa B activator | 37.557437 | 31.35924 | 5.518156 | 4.998359 | 40.87182 | 39.21335 | 32.58714 | 30.998251 | 33.59672 | 29.48275 | 5.4170113 | 5.736868 | 5.400589 | 4.8548923 | 4.9680996 | 5.1720843 |
| ASMM10P053 | 0.02233 | 0.3546919 | 1.4397583 | up | NM_145618 | Narg2 | NMDA receptor-regulated gene 2 | 231.57583 | 197.13255 | 8.142163 | 7.616336 | 239.2633 | 245.36174 | 210.10245 | 194.8332 | 200.86928 | 195.69518 | 7.926479 | 8.345088 | 8.154921 | 7.5257506 | 7.5469136 | 7.776344 |
| ASMM10P051 | 0.02234 | 0.3546919 | 1.4108514 | up | NM_175340 | Nhlrc1 | NHL repeat containing 1 | 12977.181 | 10816.056 | 13.8171 | 13.320534 | 12336.771 | 12747.292 | 13847.479 | 11049.124 | 10291.628 | 11107.417 | 13.638166 | 13.838521 | 13.974612 | 13.29982 | 13.165199 | 13.496582 |
| ASMM10P052 | 0.02237 | 0.3546919 | 1.6488841 | up | NM_001169 | F2r | coagulation factor II (thrombin) receptor | 2044.0096 | 1476.9172 | 11.204978 | 10.483488 | 1928.0856 | 1958.0483 | 2245.895 | 1309.3372 | 1577.9393 | 1543.4752 | 10.909031 | 11.242344 | 11.463558 | 10.273308 | 10.498269 | 10.678886 |
| ASMM10P042 | 0.02239 | 0.3548333 | 1.3769444 | up | NM_026962 | Kbtbd3 | kelch repeat and BTB (POZ) domain containing 3 | 653.08275 | 569.89112 | 9.591597 | 9.130126 | 669 | 632.9875 | 657.26074 | 560.25806 | 582.77 | 566.6453 | 9.37185 | 9.667402 | 9.735538 | 9.056965 | 9.08062 | 9.252794 |
| ASMM10P024 | 0.02250 | 0.3552887 | 1.3231024 | up | NM_001039 | Asxl1 | additional sex combs like 1 | 73.882945 | 67.129913 | 6.47953 | 6.075605 | 91.43815 | 72.2004 | 58.010284 | 71.208466 | 71.01203 | 59.169243 | 6.569827 | 6.6052365 | 6.2635255 | 6.0612454 | 6.0375834 | 6.127986 |
| ASMM10P008 | 0.02253 | 0.3552887 | 1.3695642 | up | NM_026515 | 2810417H13 | RIKEN cDNA 2810417H13 gene | 240.86251 | 216.36861 | 8.195349 | 7.741632 | 255.4219 | 211.22128 | 255.94434 | 233.88841 | 233.8864 | 181.33102 | 8.021638 | 8.13817 | 8.42624 | 7.7907023 | 7.7636547 | 7.6705403 |
| ASMM10P000 | 0.02253 | 0.3552887 | 1.5143758 | up | NM_001146 | Homer3 | homer homolog 3 (Drosophila) | 61.586523 | 48.743792 | 6.220416 | 5.621693 | 70.51591 | 66.957054 | 47.286606 | 52.65711 | 49.64342 | 43.930847 | 6.201374 | 6.499362 | 5.9605117 | 5.6190963 | 5.525023 | 5.7209587 |
| ASMM10P004 | 0.02260 | 0.3552894 | 1.6123389 | up | NM_001025 | Zfp943 | zinc finger prtoein 943 | 106.86125 | 80.571287 | 7.023593 | 6.334438 | 115.04121 | 83.09051 | 122.45203 | 78.81974 | 87.2023 | 75.69182 | 6.8942323 | 6.8056364 | 7.370911 | 6.2038527 | 6.334741 | 6.464721 |
| ASMM10P024 | 0.02263 | 0.3552894 | 1.6079903 | up | NM_010863 | Myo1b | myosin IB | 5684.1205 | 4079.0678 | 12.629019 | 11.94376 | 5776.887 | 4467.7505 | 6807.724 | 3926.8416 | 4548.6455 | 3761.7163 | 12.523109 | 12.375271 | 12.988677 | 11.86442 | 12.016829 | 11.950032 |
| ASMM10P020 | 0.02267 | 0.3552894 | 1.2948297 | up | NM_133207 | Kcnh7 | potassium voltage-gated channel, subfamily H (eag-related), member 7 | 115.97993 | 107.8492 | 7.120816 | 6.748054 | 150.10219 | 90.21527 | 107.62233 | 116.61482 | 115.41879 | 91.51401 | 7.2642245 | 6.9222717 | 7.175953 | 6.7757607 | 6.735108 | 6.7332935 |
| ASMM10P031 | 0.02272 | 0.3552894 | 1.5616413 | up | NM_183000 | Asic3 | acid-sensing (proton-gated) ion channel 3 | 75.703253 | 59.718835 | 6.541074 | 5.898011 | 70.84418 | 80.45885 | 75.80673 | 65.2072 | 68.06283 | 45.886475 | 6.2085767 | 6.755877 | 6.658768 | 5.935506 | 5.9756355 | 5.782891 |
| ASMM10P049 | 0.02278 | 0.3552894 | 1.6837518 | up | NM_172575 | Zfp277 | zinc finger protein 277 | 5374.234 | 3778.3571 | 12.578023 | 11.826343 | 5112.0996 | 5299.369 | 5711.2334 | 3690.2883 | 3307.288 | 4337.495 | 12.345634 | 12.630223 | 12.758212 | 11.762367 | 11.569914 | 12.146749 |
| ASMM10P003 | 0.02280 | 0.3552894 | 1.279525 | up | NM_026817 | Rabl2 | RAB, member of RAS oncogene family-like 2 | 274.17364 | 259.76934 | 8.366733 | 8.011125 | 327.66055 | 271.1904 | 223.66997 | 255.47256 | 269.544 | 254.29147 | 8.368022 | 8.48715 | 8.245027 | 7.920238 | 7.964636 | 8.1485 |
| ASMM10P023 | 0.02283 | 0.3552894 | 1.7119806 | up | NM_139303 | Kif18a | kinesin family member 18A | 121.97389 | 87.931975 | 7.229622 | 6.453955 | 123.66178 | 117.67841 | 124.57249 | 77.55048 | 90.861435 | 95.38401 | 6.9892373 | 7.303842 | 7.3957853 | 6.181704 | 6.3938627 | 6.7862988 |
| ASMM10P038 | 0.02285 | 0.3552894 | 1.6445816 | up | NM_027298 | Ccdc89 | coiled-coil domain containing 89 | 149.33463 | 111.78277 | 7.516314 | 6.798594 | 141.79222 | 137.25166 | 168.96 | 109.10102 | 127.04521 | 99.20208 | 7.1795015 | 7.529782 | 7.8396597 | 6.682238 | 6.873711 | 6.8398323 |
| ASMM10P051 | 0.02288 | 0.3552894 | 1.5802996 | up | NM_028377 | Ccdc176 | coiled-coil domain containing 176 | 110.88753 | 85.008318 | 7.071795 | 6.411596 | 128.39342 | 87.1904 | 117.07876 | 85.68144 | 81.2427 | 88.100815 | 7.0399127 | 6.874178 | 7.301293 | 6.3236322 | 6.2340693 | 6.677088 |
| ASMM10P007 | 0.02294 | 0.3552894 | 1.3431543 | up | NM_011961 | Plod2 | procollagen lysine, 2-oxoglutarate 5-dioxygenase 2 | 145.08444 | 133.11258 | 7.474846 | 7.049221 | 159.66031 | 127.99809 | 147.59491 | 126.1748 | 149.61748 | 123.54547 | 7.350294 | 7.4294944 | 7.6447487 | 6.8918285 | 7.1094413 | 7.1463923 |
| ASMM10P047 | 0.02294 | 0.3552894 | 1.4510317 | up | NM_010661 | Krt12 | keratin 12 | 14506.617 | 11681.594 | 13.969641 | 13.432562 | 16477.299 | 11844.722 | 15197.83 | 11647.428 | 11425.336 | 11972.017 | 14.073231 | 13.735439 | 14.100254 | 13.37907 | 13.305536 | 13.613081 |
| ASMM10P004 | 0.02295 | 0.3552894 | 3.3700613 | up | NM_001014 | Abi3bp | ABI gene family, member 3 (NESH) binding protein | 32.063508 | 11.622744 | 5.275118 | 3.522343 | 24.887577 | 35.174755 | 36.128193 | 11.673151 | 8.201421 | 14.993661 | 4.6850324 | 5.585164 | 5.555157 | 3.4064155 | 2.9144 | 4.2462134 |
| ASMM10P035 | 0.02296 | 0.3552894 | 1.3877321 | up | NM_001037 | Pinlyp | phospholipase A2 inhibitor and LY6/PLAUR domain containing | 55.750021 | 48.834667 | 6.091197 | 5.618468 | 64.06046 | 54.141884 | 49.047718 | 57.196743 | 44.999172 | 44.308086 | 6.062712 | 6.197742 | 6.0131364 | 5.743388 | 5.379438 | 5.732577 |
| ASMM10P018 | 0.02298 | 0.3552894 | 1.5648898 | up | NM_001100 | Taf4b | TAF4B RNA polymerase II, TATA box binding protein (TBP)-associated factor | 214.20964 | 168.85717 | 8.037106 | 7.391045 | 207.8092 | 212.90225 | 221.91748 | 174.70241 | 164.89449 | 166.9746 | 7.7287474 | 8.149681 | 8.232891 | 7.3686786 | 7.2479963 | 7.5564613 |
| ASMM10P032 | 0.02308 | 0.3552894 | 1.3469329 | up | NM_027356 | Ufsp1 | UFM1-specific peptidase 1 | 13194.939 | 11514.315 | 13.842291 | 13.412613 | 13867.283 | 12611.096 | 13106.438 | 11308.847 | 11021.244 | 12212.854 | 13.810246 | 13.827466 | 13.889162 | 13.338698 | 13.256449 | 13.642693 |
| ASMM10P024 | 0.02309 | 0.3552894 | 1.4406508 | up | NM_009641 | Angpt4 | angiopoietin 4 | 26.646593 | 21.612491 | 4.984739 | 4.458018 | 36.31872 | 21.62684 | 21.99422 | 21.77183 | 24.132772 | 18.93287 | 5.238142 | 4.9015713 | 4.8145037 | 4.327155 | 4.479836 | 4.567064 |
| ASMM10P029 | 0.02310 | 0.3552894 | 1.3043411 | up | NM_019563 | Cited4 | Cbp/p300-interacting transactivator, with Glu/Asp-rich carboxy-terminal domain, 4 | 6363.1604 | 5724.0059 | 12.810923 | 12.427602 | 7296.736 | 5250.8267 | 6541.9185 | 5994.3 | 5698.7617 | 5478.956 | 12.878289 | 12.619967 | 12.934513 | 12.455586 | 12.338717 | 12.488503 |
| ASMM10P011 | 0.02314 | 0.3555446 | 1.3391191 | up | NM_009029 | Rb1 | retinoblastoma 1 | 102.22837 | 94.20264 | 6.966119 | 6.544834 | 112.83843 | 105.26244 | 88.58423 | 92.576225 | 111.5695 | 78.4622 | 6.86679 | 7.146116 | 6.88545 | 6.4330463 | 6.688782 | 6.512675 |
| ASMM10P029 | 0.02316 | 0.3556094 | 2.158447 | up | NM_001081 | Clhc1 | clathrin heavy chain linker domain containing 1 | 41.109206 | 22.037995 | 5.589711 | 4.479717 | 58.21919 | 27.804646 | 37.303783 | 23.558847 | 18.116701 | 24.438438 | 5.920232 | 5.244953 | 5.6039476 | 4.449329 | 4.0783825 | 4.91144 |
| ASMM10P051 | 0.02322 | 0.355857 | 1.2984614 | up | NM_153538 | Zcchc6 | zinc finger, CCHC domain containing 6 | 257.11601 | 242.37107 | 8.269409 | 7.892606 | 327.69952 | 226.51877 | 217.12973 | 257.33408 | 282.45126 | 187.32788 | 8.368461 | 8.233501 | 8.206264 | 7.93154 | 8.029989 | 7.7162876 |
| ASMM10P017 | 0.02325 | 0.3559498 | 1.3399726 | up | NM_021559 | Zfp191 | zinc finger protein 191 | 2254.3238 | 2008.5611 | 11.347427 | 10.925223 | 2272.9294 | 2133.9941 | 2356.048 | 1956.6212 | 2180.7744 | 1888.3478 | 11.14113 | 11.373805 | 11.527346 | 10.852226 | 10.964252 | 10.959192 |
| ASMM10P004 | 0.02326 | 0.3559498 | 1.9371354 | up | NM_001114 | 9930111J21 | RIKEN cDNA 9930111J21 gene 1 | 46.503345 | 27.181466 | 5.747992 | 4.794068 | 67.8501 | 42.058186 | 29.601748 | 29.112251 | 27.850027 | 24.582119 | 6.1437387 | 5.8384614 | 5.261777 | 4.7668176 | 4.6963625 | 4.9190226 |
| ASMM10P055 | 0.02337 | 0.3570764 | 1.304215 | up | NM_001190 | Gne | glucosamine (UDP-N-acetyl)-2-epimerase/N-acetylmannosamine kinase | 64.900847 | 60.082063 | 6.301536 | 5.918354 | 80.114334 | 55.430523 | 59.157684 | 61.581867 | 60.19674 | 58.467583 | 6.3814816 | 6.23184 | 6.2912855 | 5.8473673 | 5.7954173 | 6.1122775 |
| ASMM10P007 | 0.02351 | 0.3572388 | 1.2885912 | up | NM_012061 | Cadps | Ca2+-dependent secretion activator | 380.21457 | 357.84964 | 8.830068 | 8.464273 | 438.43652 | 377.43448 | 324.7727 | 367.53845 | 346.53705 | 359.47342 | 8.780907 | 8.943852 | 8.765444 | 8.443246 | 8.328126 | 8.621447 |
| ASMM10P033 | 0.02352 | 0.3572388 | 1.8530454 | up | NM_027564 | 4921507P07 | RIKEN cDNA 4921507P07 gene | 86.54583 | 56.825754 | 6.719051 | 5.829152 | 98.00842 | 68.82457 | 92.8045 | 51.170227 | 54.16474 | 65.142296 | 6.667155 | 6.5348654 | 6.9551315 | 5.5780463 | 5.6486773 | 6.2607336 |
| ASMM10P040 | 0.02354 | 0.3572388 | 1.624062 | up | NM_024191 | Arl2bp | ADP-ribosylation factor-like 2 binding protein | 610.25986 | 446.30699 | 9.481331 | 8.781724 | 616.35645 | 500.74118 | 713.68195 | 468.993 | 439.55463 | 430.37335 | 9.243446 | 9.348543 | 9.852003 | 8.799358 | 8.674598 | 8.871216 |
| ASMM10P055 | 0.02360 | 0.3572388 | 1.4522724 | up | NM_026916 | Nupr1l | nuclear protein transcriptional regulator 1 like | 198.72929 | 169.00723 | 7.929896 | 7.391584 | 194.44626 | 196.66566 | 205.07596 | 173.44409 | 185.3253 | 148.2523 | 7.633149 | 8.040025 | 8.116513 | 7.3588266 | 7.4257574 | 7.3901668 |
| ASMM10P020 | 0.02361 | 0.3572388 | 1.5075996 | up | NM_026430 | Uxs1 | UDP-glucuronate decarboxylase 1 | 467.21881 | 376.69135 | 9.122575 | 8.530321 | 534.1023 | 452.53397 | 415.02017 | 353.94595 | 352.55865 | 423.56946 | 9.048422 | 9.205546 | 9.113756 | 8.388259 | 8.352937 | 8.849768 |
| ASMM10P005 | 0.02364 | 0.3572388 | 1.4307279 | up | NM_001164 | Tmem42 | transmembrane protein 42 | 4297.2794 | 3555.4231 | 12.260408 | 11.743659 | 4745.769 | 3969.9407 | 4176.1284 | 3538.3762 | 3203.4622 | 3924.431 | 12.236964 | 12.214848 | 12.329412 | 11.697865 | 11.525645 | 12.007466 |
| ASMM10P004 | 0.02373 | 0.3578164 | 3.2020873 | up | NM_001039 | Seh1l | SEH1-like (S. cerevisiae | 62.021031 | 24.298053 | 6.240466 | 4.561453 | 55.653 | 73.58101 | 56.829082 | 19.498396 | 18.68124 | 34.714523 | 5.8559613 | 6.6330676 | 6.232369 | 4.1715136 | 4.1201777 | 5.3926687 |
| ASMM10P023 | 0.02378 | 0.3578164 | 1.4057893 | up | NM_011428 | Snap25 | synaptosomal-associated protein 25 | 18830.835 | 15737.307 | 14.336621 | 13.84524 | 18183.82 | 16485.959 | 21822.727 | 16331.254 | 17580.65 | 13300.018 | 14.226206 | 14.188185 | 14.595471 | 13.873421 | 13.910131 | 13.752169 |
| ASMM10P011 | 0.02378 | 0.3578164 | 1.3484134 | up | NM_008466 | Kpna3 | karyopherin (importin) alpha 3 | 32246.704 | 28522.466 | 15.148974 | 14.717711 | 30401.588 | 31773.111 | 34565.414 | 27375.537 | 29180.71 | 29011.15 | 15.011757 | 15.121584 | 15.313581 | 14.61443 | 14.655712 | 14.882991 |
| ASMM10P014 | 0.02392 | 0.3582227 | 8.5132475 | up | NM_001033 | Csta | cystatin A | 156.99632 | 18.727495 | 7.244856 | 4.155147 | 82.37501 | 314.53073 | 74.08321 | 22.826075 | 10.149779 | 23.20663 | 6.42023 | 8.69036 | 6.6239786 | 4.4000354 | 3.2220855 | 4.843319 |
| ASMM10P008 | 0.02394 | 0.3582227 | 1.3395248 | up | NM_144931 | Nae1 | NEDD8 activating enzyme E1 subunit 1 | 552.99696 | 497.89623 | 9.354938 | 8.933217 | 634.6273 | 505.37708 | 518.9865 | 490.0654 | 474.16113 | 529.46216 | 9.290606 | 9.362462 | 9.411747 | 8.863155 | 8.780907 | 9.155589 |
| ASMM10P021 | 0.02397 | 0.3582227 | 2.2653642 | up | NM_172521 | Nutm1 | NUT midline carcinoma, family member 1 | 139.82471 | 73.51217 | 7.356727 | 6.176984 | 161.53825 | 83.551895 | 174.38397 | 80.091034 | 88.79311 | 51.652367 | 7.368121 | 6.814493 | 7.8875675 | 6.227697 | 6.3587375 | 5.944518 |
| ASMM10P018 | 0.02406 | 0.3582227 | 1.4329262 | up | NM_007505 | Atp5a1 | ATP synthase, H+ transporting, mitochondrial F1 complex, alpha subunit 1 | 844.12837 | 708.45595 | 9.962807 | 9.443842 | 828.57776 | 819.3945 | 884.41284 | 721.93097 | 727.8892 | 675.54767 | 9.68753 | 10.03268 | 10.16821 | 9.427152 | 9.398177 | 9.506198 |
| ASMM10P047 | 0.02408 | 0.3582227 | 1.6672165 | up | NM_028963 | 4933427D14 | RIKEN cDNA 4933427D14 gene | 43.797602 | 30.845522 | 5.704592 | 4.96715 | 58.633953 | 31.146622 | 41.612232 | 36.185875 | 27.744114 | 28.606577 | 5.9304895 | 5.40842 | 5.774866 | 5.0796914 | 4.689362 | 5.1323977 |
| ASMM10P014 | 0.02410 | 0.3582227 | 1.2597491 | up | NM_027341 | Dzip3 | DAZ interacting protein 3, zinc finger | 166.52453 | 162.97995 | 7.672527 | 7.339391 | 177.61897 | 156.18352 | 165.7711 | 169.5595 | 172.21849 | 147.16187 | 7.4997234 | 7.704153 | 7.8137045 | 7.323597 | 7.314917 | 7.3796577 |
| ASMM10P019 | 0.02415 | 0.3582227 | 1.7632869 | up | NM_030703 | Cpn1 | carboxypeptidase N, polypeptide 1 | 224.06981 | 158.13508 | 8.100737 | 7.282469 | 218.45561 | 232.98454 | 220.76927 | 129.54501 | 187.9347 | 156.92552 | 7.800966 | 8.273291 | 8.227953 | 6.9307184 | 7.4462194 | 7.4704704 |
| ASMM10P008 | 0.02416 | 0.3582227 | 1.7965053 | up | NM_029227 | Tomm20l | translocase of outer mitochondrial membrane 20 homolog (yeast)-like | 204.35963 | 137.73993 | 7.93527 | 7.090077 | 214.28679 | 149.79085 | 249.00124 | 157.91718 | 143.39105 | 111.91156 | 7.7738843 | 7.6423903 | 8.389536 | 7.2176023 | 7.0461707 | 7.006458 |
| ASMM10P036 | 0.02420 | 0.3582227 | 1.4601764 | up | NM_029078 | Pcf11 | cleavage and polyadenylation factor subunit homolog (S. cerevisiae) | 833.96502 | 669.6401 | 9.90604 | 9.359897 | 997.30646 | 893.4007 | 611.1879 | 663.0003 | 717.4741 | 628.4459 | 9.936494 | 10.152397 | 9.629228 | 9.302642 | 9.375115 | 9.401934 |
| ASMM10P034 | 0.02422 | 0.3582227 | 2.2043207 | up | NM_001167 | D6Ertd527e | DNA segment, Chr 6, ERATO Doi 527, expressed | 92.838972 | 49.576233 | 6.782703 | 5.642369 | 103.63427 | 58.62285 | 116.2598 | 44.020336 | 54.791126 | 49.917236 | 6.7464075 | 6.3102455 | 7.291456 | 5.365053 | 5.6643977 | 5.897656 |
| ASMM10P026 | 0.02424 | 0.3582227 | 1.253969 | up | NM_008417 | Kcna2 | potassium voltage-gated channel, shaker-related subfamily, member 2 | 68416.083 | 69450.546 | 16.311967 | 15.985465 | 68308.445 | 65640.75 | 71299.055 | 70314.65 | 72994.805 | 65042.184 | 16.21797 | 16.259827 | 16.458103 | 15.930034 | 15.930034 | 16.096327 |
| ASMM10P014 | 0.02427 | 0.3582227 | 1.2836679 | up | NM_017404 | Mrpl39 | mitochondrial ribosomal protein L39 | 211.40001 | 202.16901 | 8.01352 | 7.653248 | 225.5917 | 211.99245 | 196.61588 | 209.73686 | 204.79602 | 191.97415 | 7.844801 | 8.142349 | 8.053411 | 7.633682 | 7.5754857 | 7.7505774 |
| ASMM10P011 | 0.02442 | 0.3590043 | 1.3287448 | up | NM_028643 | Micu2 | mitochondrial calcium uptake 2 | 5801.3228 | 5137.0995 | 12.680479 | 12.270415 | 5902.5103 | 5520.4336 | 5981.0244 | 4997.243 | 5037.5605 | 5376.495 | 12.557732 | 12.678354 | 12.807317 | 12.189873 | 12.168459 | 12.452913 |
| ASMM10P015 | 0.02447 | 0.3590043 | 2.3085074 | up | NM_011395 | Slc22a3 | solute carrier family 22 (organic cation transporter), member 3 | 873.9348 | 455.77751 | 9.991132 | 8.784171 | 735.29364 | 1062.2607 | 824.25006 | 556.8218 | 347.92297 | 462.58777 | 9.514385 | 10.399142 | 10.059868 | 9.048567 | 8.333856 | 8.970091 |
| ASMM10P027 | 0.02449 | 0.3590043 | 1.8165962 | up | NM_029198 | 4930538K18 | RIKEN cDNA 4930538K18 gene | 129.67298 | 85.23389 | 7.272312 | 6.411074 | 138.12799 | 158.47481 | 92.41615 | 98.5536 | 83.585396 | 73.562675 | 7.1436157 | 7.725311 | 6.9480085 | 6.530432 | 6.276357 | 6.426433 |
| ASMM10P042 | 0.02455 | 0.3590043 | 1.34937 | up | NM_023831 | Ift46 | intraflagellar transport 46 | 118.92422 | 107.97032 | 7.182798 | 6.750512 | 134.0213 | 114.91097 | 107.84039 | 117.56032 | 100.91003 | 105.44061 | 7.0979652 | 7.271707 | 7.1787205 | 6.786988 | 6.5404406 | 6.924106 |
| ASMM10P037 | 0.02456 | 0.3590043 | 1.6262295 | up | NM_008437 | Napsa | napsin A aspartic peptidase | 62.803393 | 47.01933 | 6.273691 | 5.572161 | 60.68188 | 60.47947 | 67.24883 | 43.301113 | 51.148937 | 46.60794 | 5.98436 | 6.355692 | 6.4810224 | 5.344807 | 5.5674477 | 5.804227 |
| ASMM10P034 | 0.02457 | 0.3590043 | 1.5026 | up | NM_173428 | Sspo | SCO-spondin | 184.02572 | 150.35143 | 7.798978 | 7.211517 | 217.31131 | 192.37787 | 142.38799 | 183.92989 | 146.14262 | 120.98177 | 7.79261 | 8.009689 | 7.5946345 | 7.443404 | 7.0757027 | 7.1154437 |
| ASMM10P008 | 0.02458 | 0.3590043 | 1.4293749 | up | NM_028248 | Tmem87b | transmembrane protein 87B | 80.617951 | 68.854266 | 6.625001 | 6.109617 | 91.271774 | 76.79799 | 73.78409 | 62.964912 | 72.99469 | 70.603195 | 6.5670233 | 6.6917024 | 6.6162777 | 5.8819747 | 6.07607 | 6.3708057 |
| ASMM10P036 | 0.02462 | 0.3593271 | 1.641976 | up | NM_016720 | Neu3 | neuraminidase 3 | 883.44275 | 648.20935 | 10.018434 | 9.303001 | 897.7976 | 745.5937 | 1006.937 | 574.43066 | 764.24243 | 605.95496 | 9.800847 | 9.900377 | 10.354078 | 9.091473 | 9.471565 | 9.345965 |
| ASMM10P015 | 0.02467 | 0.3595223 | 1.5836494 | up | NM_172826 | Dact2 | dapper homolog 2, antagonist of beta-catenin (xenopus) | 49990.045 | 37776.94 | 15.789326 | 15.126073 | 59440.965 | 38629.5 | 51899.67 | 37522.242 | 40400.297 | 35408.28 | 16.005611 | 15.41772 | 15.944647 | 15.084926 | 15.110092 | 15.183201 |
| ASMM10P018 | 0.02473 | 0.3601435 | 1.3248139 | up | NM_178377 | Commd10 | COMM domain containing 10 | 304.84597 | 278.90722 | 8.517076 | 8.111286 | 343.2681 | 320.16946 | 251.10036 | 278.97757 | 288.323 | 269.4211 | 8.434129 | 8.717142 | 8.399957 | 8.047144 | 8.061171 | 8.225544 |
| ASMM10P042 | 0.02476 | 0.3601534 | 1.5843078 | up | NM_001033 | 1190002N15 | RIKEN cDNA 1190002N15 gene | 14823.638 | 10934.923 | 13.991862 | 13.328009 | 16027.34 | 11210.458 | 17201.117 | 11933.932 | 10818.448 | 10052.39 | 14.033317 | 13.658104 | 14.284165 | 13.411757 | 13.231181 | 13.34109 |
| ASMM10P028 | 0.02486 | 0.3601534 | 1.9658557 | up | NM_027450 | Glipr2 | GLI pathogenesis-related 2 | 56.426605 | 32.630079 | 6.031442 | 5.056285 | 81.83203 | 34.31717 | 53.130615 | 35.0304 | 31.383852 | 31.475985 | 6.4115644 | 5.5494814 | 6.1332817 | 5.0349913 | 4.870138 | 5.263726 |
| ASMM10P047 | 0.02487 | 0.3601534 | 1.3250361 | up | NM_025303 | Stau2 | staufen (RNA binding protein) homolog 2 (Drosophila) | 13315.193 | 11795.895 | 13.852488 | 13.446457 | 13334.853 | 12177.633 | 14433.093 | 11473.229 | 12107.624 | 11806.831 | 13.748633 | 13.774719 | 14.034113 | 13.357664 | 13.393243 | 13.588463 |
| ASMM10P043 | 0.02488 | 0.3601534 | 1.9591309 | up | NM_019538 | Placl | placental specific protein 1 | 29.51524 | 18.380673 | 5.147971 | 4.177757 | 31.594933 | 34.449417 | 22.501371 | 18.47456 | 24.8002 | 11.867259 | 5.041199 | 5.555047 | 4.8476677 | 4.092209 | 4.5226684 | 3.918395 |
| ASMM10P001 | 0.02489 | 0.3601534 | 1.8683145 | up | NM_001205 | Jmjd4 | jumonji domain containing 4 | 284.08584 | 182.71814 | 8.405644 | 7.503906 | 268.79422 | 222.88867 | 360.57462 | 197.12595 | 191.71495 | 159.31352 | 8.090752 | 8.211607 | 8.914572 | 7.543155 | 7.4788604 | 7.4897037 |
| ASMM10P033 | 0.02490 | 0.3601534 | 1.4830929 | up | NM_172893 | Parp12 | poly (ADP-ribose) polymerase family, member 12 | 6141.6683 | 4848.8538 | 12.753854 | 12.185245 | 5907.9326 | 5539.5996 | 6977.4727 | 4425.6025 | 5334.6113 | 4786.3477 | 12.558878 | 12.681104 | 13.02158 | 12.016829 | 12.248919 | 12.289987 |
| ASMM10P050 | 0.02491 | 0.3601534 | 2.3032739 | up | NM_029803 | Ifi27l2a | interferon, alpha-inducible protein 27 like 2A | 1404.6349 | 746.87365 | 10.675592 | 9.471906 | 1333.5664 | 1587.1063 | 1293.2319 | 494.67215 | 892.6321 | 853.3167 | 10.369313 | 10.957678 | 10.699784 | 8.879767 | 9.697153 | 9.838797 |
| ASMM10P021 | 0.02494 | 0.3601534 | 1.8682996 | up | NM_181589 | Ckap2l | cytoskeleton associated protein 2-like | 190.66631 | 123.56156 | 7.840274 | 6.938548 | 224.84502 | 135.44322 | 211.71068 | 113.51666 | 119.99118 | 137.17683 | 7.840312 | 7.5125766 | 8.167932 | 6.7383947 | 6.7897615 | 7.287487 |
| ASMM10P004 | 0.02496 | 0.3601534 | 1.6199795 | up | NM_001113 | Vmn2r29 | vomeronasal 2, receptor 29 | 113.8796 | 86.694785 | 7.130715 | 6.434739 | 118.44441 | 107.4313 | 115.76308 | 93.10626 | 76.06758 | 90.910515 | 6.9321985 | 7.174867 | 7.285079 | 6.4423623 | 6.1373806 | 6.724475 |
| ASMM10P052 | 0.02497 | 0.3601534 | 1.6984656 | up | NM_134213 | Vmn1r199 | vomeronasal 1 receptor 199 | 706.65832 | 491.44061 | 9.677771 | 8.913539 | 779.6381 | 500.531 | 839.80585 | 516.581 | 530.2926 | 427.44824 | 9.598073 | 9.34823 | 10.087011 | 8.938583 | 8.941302 | 8.860733 |

| | | | | | | | | | | | | | | | | | | | | | | | |
|---|---|---|---|---|---|---|---|---|---|---|---|---|---|---|---|---|---|---|---|---|---|---|---|
| ASMM10P023 | 0.02499 | 0.3601534 | 1.3483397 | up | NM_018828 | Fnbp4 | formin binding protein 4 | 362.61115 | 329.38157 | 8.761445 | 8.330261 | 433.26636 | 312.5497 | 342.0174 | 389.2608 | 348.29022 | 250.5937 | 8.761815 | 8.681268 | 8.841253 | 8.5256815 | 8.3355255 | 8.129577 |
| ASMM10P029 | 0.02501 | 0.3601534 | 1.720854 | up | NM_008725 | Nppa | natriuretic peptide type A | 19.976674 | 13.561883 | 4.573237 | 3.790112 | 23.55465 | 22.15854 | 14.216832 | 14.103721 | 15.019879 | 11.56205 | 4.602195 | 4.93433 | 4.1831846 | 3.6929436 | 3.7957206 | 3.8816714 |
| ASMM10P003 | 0.02509 | 0.3602197 | 3.5731179 | up | NM_001271 | Tk1 | thymidine kinase 1 | 45.653032 | 16.684862 | 5.812941 | 3.975757 | 45.40017 | 44.542255 | 47.01667 | 10.246071 | 14.64133 | 25.167185 | 5.5672407 | 5.9203506 | 5.9512315 | 3.2206564 | 3.7554388 | 4.951177 |
| ASMM10P054 | 0.02509 | 0.3602197 | 1.5568692 | up | NM_029001 | Elovl7 | ELOVL family member 7, elongation of long chain fatty acids (yeast) | 135.37193 | 106.59038 | 7.36408 | 6.725432 | 141.36012 | 150.07956 | 114.67612 | 96.46784 | 124.03748 | 99.26581 | 7.1762867 | 7.644492 | 7.27146 | 6.496967 | 6.8385715 | 6.840757 |
| ASMM10P002 | 0.02512 | 0.360309 | 1.382491 | up | NM_011234 | Rad51 | RAD51 homolog | 78.955668 | 69.766933 | 6.596272 | 6.129002 | 88.570435 | 72.792625 | 75.503944 | 74.67065 | 65.00138 | 69.62877 | 6.5212245 | 6.6164737 | 6.651117 | 6.12811 | 5.907536 | 6.351359 |
| ASMM10P035 | 0.02516 | 0.3607266 | 1.4319672 | up | NM_009568 | Zfp94 | zinc finger protein 94 | 210.91523 | 180.67256 | 8.008612 | 7.490614 | 215.79492 | 225.56487 | 191.38591 | 185.67648 | 179.64175 | 176.69945 | 7.783571 | 8.226537 | 8.015729 | 7.456944 | 7.3801274 | 7.6347704 |
| ASMM10P009 | 0.02520 | 0.3608139 | 1.4475244 | up | NM_175418 | Mybpc1 | myosin binding protein C, slow-type | 258.45463 | 218.70351 | 8.295662 | 7.762075 | 280.64075 | 235.05933 | 259.66382 | 236.17662 | 198.041 | 221.89291 | 8.150529 | 8.286865 | 8.449593 | 7.8039455 | 7.5254126 | 7.956866 |
| ASMM10P023 | 0.02527 | 0.3611075 | 1.3210914 | up | NM_172653 | Slc39a10 | solute carrier family 39 (zinc transporter), member 10 | 63335.319 | 59779.3 | 16.178922 | 15.777191 | 59935.27 | 62252.297 | 67818.39 | 62500.023 | 60159.297 | 56678.58 | 16.01558 | 16.158913 | 16.362272 | 15.774575 | 15.680805 | 15.876194 |
| ASMM10P034 | 0.02534 | 0.3611486 | 1.5206534 | up | NM_175035 | Gimap5 | GTPase, IMAP family member 5 | 256.34984 | 207.02086 | 8.287343 | 7.682651 | 255.98155 | 245.00356 | 268.0644 | 189.78957 | 225.4418 | 205.83122 | 8.025091 | 8.3422575 | 8.49468 | 7.487875 | 7.711437 | 7.8486423 |
| ASMM10P030 | 0.02538 | 0.3611486 | 1.2777483 | up | NM_172405 | Fam175a | family with sequence similarity 175, member A | 631.44112 | 592.13697 | 9.535584 | 9.18198 | 741.3034 | 508.1004 | 644.91956 | 587.65393 | 641.0619 | 547.69507 | 9.52576 | 9.371689 | 9.709302 | 9.12549 | 9.216121 | 9.204329 |
| ASMM10P012 | 0.02543 | 0.3611486 | 1.3555841 | up | NM_145960 | Mtrf1 | mitochondrial translational release factor 1 | 2270.3435 | 1993.4115 | 11.350574 | 10.91166 | 2473.2642 | 2017.9769 | 2319.7893 | 1980.4446 | 1904.464 | 2095.326 | 11.256927 | 11.289761 | 11.505035 | 10.867216 | 10.766043 | 11.10172 |
| ASMM10P014 | 0.02545 | 0.3611486 | 1.6466078 | up | NM_001081 | Marf1 | meiosis arrest female 1 | 2408.967 | 1735.8752 | 11.433467 | 10.71397 | 2239.3276 | 2393.886 | 2593.6873 | 1552.9491 | 1770.6951 | 1883.9814 | 11.117604 | 11.524024 | 11.658773 | 10.519234 | 10.667267 | 10.955409 |
| ASMM10P054 | 0.02548 | 0.3611486 | 1.855758 | up | NM_001038 | Gm4787 | predicted gene 4787 | 88.550315 | 58.155922 | 6.760285 | 5.868277 | 75.197716 | 89.87861 | 100.57462 | 68.001686 | 55.77024 | 50.69584 | 6.2892756 | 6.916012 | 7.0755687 | 5.9958315 | 5.691386 | 5.917613 |
| ASMM10P006 | 0.02551 | 0.3611486 | 1.508645 | up | NM_001205 | Sept7 | septin 7 | 2110.6315 | 1668.3418 | 11.251416 | 10.658163 | 2111.9795 | 2017.3367 | 2202.5784 | 1653.1838 | 1551.2018 | 1800.6398 | 11.032662 | 11.288837 | 11.432749 | 10.607777 | 10.472285 | 10.894426 |
| ASMM10P021 | 0.02552 | 0.3611486 | 1.533946 | up | NM_001077 | Nkx2-2 | NK2 homeobox 2 | 369.72311 | 294.68077 | 8.799953 | 8.182706 | 363.64206 | 342.31186 | 403.2154 | 286.0304 | 346.49582 | 251.51608 | 8.513539 | 8.813314 | 9.073007 | 8.08533 | 8.327743 | 8.135044 |
| ASMM10P005 | 0.02558 | 0.3611486 | 1.5583223 | up | NM_001161 | Klhl7 | kelch-like 7 | 1228.8939 | 937.71478 | 10.48618 | 9.846186 | 1176.8961 | 1149.9371 | 1359.8485 | 979.90344 | 909.76764 | 923.47327 | 10.174991 | 10.515795 | 10.767754 | 9.858746 | 9.724032 | 9.955781 |
| ASMM10P054 | 0.02561 | 0.3611486 | 1.7423709 | up | NM_001102 | Kng1 | kininogen 1 | 105.30746 | 75.395651 | 7.017504 | 6.216452 | 97.394295 | 101.0088 | 117.5193 | 95.06147 | 74.18299 | 56.942493 | 6.6578164 | 7.0865912 | 7.3081036 | 6.475956 | 6.099655 | 6.073745 |
| ASMM10P006 | 0.02567 | 0.3611486 | 1.5508162 | up | NM_009483 | Kdm6a | lysine (K)-specific demethylase 6A | 1900.3876 | 1450.3779 | 11.089744 | 10.456716 | 2020.1805 | 1507.7732 | 2173.2092 | 1483.4921 | 1402.1393 | 1465.5023 | 10.973385 | 10.880276 | 11.415571 | 10.448158 | 10.326048 | 10.595943 |
| ASMM10P052 | 0.02570 | 0.3611486 | 1.8305663 | up | NM_001081 | S100z | S100 calcium binding protein, zeta | 837.35437 | 534.20566 | 9.905344 | 9.033054 | 988.9657 | 551.5196 | 971.5778 | 561.2474 | 507.64282 | 533.72675 | 9.927621 | 9.483831 | 10.30458 | 9.058942 | 8.875738 | 9.164482 |
| ASMM10P005 | 0.02570 | 0.3611486 | 1.3879535 | up | NM_001159 | Cep97 | centrosomal protein 97 | 217.87901 | 191.62242 | 8.04853 | 7.57557 | 237.69366 | 180.59564 | 235.34773 | 197.48991 | 200.22466 | 177.1527 | 7.9164524 | 7.9148765 | 8.31426 | 7.546372 | 7.5422583 | 7.638081 |
| ASMM10P055 | 0.02577 | 0.3611486 | 1.5186134 | up | NM_026582 | Wls | wntless homolog (Drosophila) | 1959.6834 | 1531.8158 | 11.142566 | 10.539811 | 2007.6904 | 1648.7498 | 2222.61 | 1555.2842 | 1484.7802 | 1555.3829 | 10.964794 | 11.015479 | 11.447425 | 10.52034 | 10.407081 | 10.692013 |
| ASMM10P013 | 0.02579 | 0.3611486 | 1.3300996 | up | NM_001033 | Slc2a13 | solute carrier family 2 (facilitated glucose transporter), member 13 | 40058.229 | 36052.297 | 15.461562 | 15.050027 | 35872.707 | 41432.17 | 42869.81 | 37197.438 | 39195.098 | 31764.354 | 15.23217 | 15.538528 | 15.613987 | 15.071283 | 15.073737 | 15.005062 |
| ASMM10P025 | 0.02579 | 0.3611486 | 1.340431 | up | NM_133869 | Cept1 | choline/ethanolaminephosphotransferase 1 | 1190.2218 | 1064.8622 | 10.450508 | 10.027811 | 1240.1587 | 1191.0518 | 1139.4548 | 1080.352 | 1037.9099 | 1076.3246 | 10.259006 | 10.558923 | 10.533594 | 10.002848 | 9.9114 | 10.169184 |
| ASMM10P050 | 0.02580 | 0.3611486 | 1.2679446 | up | NM_026347 | Iah1 | isoamyl acetate-hydrolyzing esterase 1 homolog (S. cerevisiae) | 3373.2945 | 3140.3801 | 11.909385 | 11.566894 | 3546.9238 | 3549.1755 | 3023.7842 | 3213.3477 | 3093.4219 | 3114.3706 | 11.784453 | 12.065815 | 11.877888 | 11.559193 | 11.476928 | 11.66456 |
| ASMM10P019 | 0.02581 | 0.3611486 | 2.2529605 | up | NM_207268 | Ccdc87 | coiled-coil domain containing 87 | 1135.9089 | 612.07906 | 10.368601 | 9.196779 | 1128.2661 | 967.4793 | 1311.9814 | 551.28485 | 481.95258 | 802.99976 | 10.118801 | 10.265638 | 10.721364 | 9.032936 | 8.804163 | 9.753238 |
| ASMM10P017 | 0.02582 | 0.3611486 | 1.961419 | up | NM_001081 | Mpp7 | membrane protein, palmitoylated 7 (MAGUK p55 subfamily member 7) | 143.85987 | 91.170943 | 7.461929 | 6.490031 | 156.77368 | 123.48542 | 151.3205 | 72.85793 | 91.31118 | 109.34372 | 7.32442 | 7.3775578 | 7.6838083 | 6.0949154 | 6.4012475 | 6.97393 |
| ASMM10P011 | 0.02584 | 0.3611956 | 1.524363 | up | NM_028222 | Cdkn3 | cyclin-dependent kinase inhibitor 3 | 141.7398 | 114.94592 | 7.446387 | 6.83818 | 147.29126 | 133.65494 | 144.2732 | 125.95738 | 103.67161 | 115.20878 | 7.2351265 | 7.4929967 | 7.6110373 | 6.8887315 | 6.5807943 | 7.0450153 |
| ASMM10P039 | 0.02588 | 0.3612074 | 1.2545694 | up | NM_011721 | Wrn | Werner syndrome homolog (human) | 208.98525 | 199.99164 | 7.963108 | 7.635916 | 278.16898 | 172.9874 | 175.79936 | 206.8169 | 207.66714 | 185.49088 | 8.138274 | 7.8512697 | 7.8997817 | 7.610143 | 7.596191 | 7.7014146 |
| ASMM10P042 | 0.02589 | 0.3612074 | 1.5163788 | up | NM_019779 | Cyp11a1 | cytochrome P450, family 11, subfamily a, polypeptide 1 | 65.448259 | 50.903002 | 6.287096 | 5.686466 | 87.13192 | 59.72065 | 49.492207 | 52.05469 | 50.83971 | 49.814606 | 6.4974346 | 6.3377175 | 6.0261354 | 5.603993 | 5.5603395 | 5.8950644 |
| ASMM10P014 | 0.02597 | 0.3617306 | 1.2981088 | up | NM_024273 | Timmdc1 | translocase of inner mitochondrial membrane domain containing 1 | 4967.2635 | 4526.9838 | 12.465777 | 12.089365 | 4794.7363 | 5051.3726 | 5055.6816 | 4662.641 | 4928.267 | 3990.0435 | 12.256031 | 12.557732 | 12.583567 | 12.092659 | 12.137395 | 12.038042 |
| ASMM10P031 | 0.02604 | 0.3621027 | 1.3804698 | up | NM_009007 | Rac1 | RAS-related C3 botulinum substrate 1 | 88653.478 | 82696.11 | 16.724708 | 16.259549 | 81816.52 | 91697.47 | 92446.445 | 83902.17 | 84056.72 | 80129.44 | 16.525383 | 16.770472 | 16.87827 | 16.211964 | 16.141523 | 16.42516 |
| ASMM10P033 | 0.02610 | 0.3622928 | 1.3244236 | up | NM_177003 | 9630033F20 | RIKEN cDNA 9630033F20 gene | 14365.551 | 12700.586 | 13.951889 | 13.546525 | 13718.983 | 13490.757 | 15886.912 | 12370.113 | 13872.17 | 11859.475 | 13.78553 | 13.910131 | 14.160007 | 13.464834 | 13.57773 | 13.59701 |
| ASMM10P024 | 0.02613 | 0.3622928 | 1.2509736 | up | NM_009673 | Anxa5 | annexin A5 | 26928.337 | 25321.439 | 14.864742 | 14.54169 | 27625.348 | 25497.164 | 27662.498 | 23237.305 | 29048.607 | 23678.406 | 14.846998 | 14.799405 | 14.947822 | 14.378993 | 14.650607 | 14.595471 |
| ASMM10P015 | 0.02619 | 0.3622928 | 1.3356034 | up | NM_010587 | Itsn1 | intersectin 1 (SH3 domain protein 1A) | 12725.838 | 11220.038 | 13.790663 | 13.373171 | 12496.494 | 12089.158 | 13591.861 | 10672.149 | 11509.79 | 11478.175 | 13.65712 | 13.76249 | 13.952379 | 13.26312 | 13.315952 | 13.540442 |
| ASMM10P028 | 0.02621 | 0.3622928 | 2.0644124 | up | NM_031872 | Tas1r3 | taste receptor, type 1, member 3 | 56.779371 | 33.458854 | 6.106095 | 5.060364 | 45.00984 | 68.38275 | 56.945522 | 36.36773 | 40.17821 | 23.830622 | 5.5566525 | 6.5265546 | 6.235079 | 5.0858583 | 5.216617 | 4.8786173 |
| ASMM10P012 | 0.02626 | 0.3622928 | 1.3240592 | up | NM_133817 | Zfp451 | zinc finger protein 451 | 3387.6557 | 3011.8932 | 11.909526 | 11.504559 | 3443.565 | 3087.6035 | 3631.7986 | 3201.214 | 3191.5664 | 2642.8992 | 11.740312 | 11.86555 | 12.122717 | 11.553767 | 11.521432 | 11.438477 |
| ASMM10P048 | 0.02630 | 0.3622928 | 1.4565885 | up | NM_013529 | Gfpt2 | glutamine fructose-6-phosphate transaminase 2 | 121.57615 | 103.69192 | 7.202922 | 6.660329 | 150.20645 | 111.89839 | 102.6236 | 111.73198 | 128.43315 | 70.910614 | 7.2653484 | 7.2351265 | 7.1082916 | 6.714881 | 6.8889294 | 6.377176 |
| ASMM10P019 | 0.02633 | 0.3622928 | 1.2622932 | up | NM_021791 | Doc2g | double C2, gamma | 543.29907 | 520.59228 | 9.322919 | 8.986872 | 667.94434 | 468.8687 | 493.08417 | 548.8055 | 600.41864 | 412.5527 | 9.369018 | 9.253997 | 9.345743 | 9.025659 | 9.12184 | 8.813118 |
| ASMM10P009 | 0.02634 | 0.3622928 | 1.3182451 | up | NM_153391 | Wdr19 | WD repeat domain 19 | 349.45611 | 322.01868 | 8.711098 | 8.312479 | 414.81146 | 310.3696 | 323.18726 | 302.4932 | 324.6703 | 338.89255 | 8.702427 | 8.670938 | 8.759929 | 8.167932 | 8.234354 | 8.535152 |
| ASMM10P010 | 0.02639 | 0.3622928 | 1.3468498 | up | NM_025557 | Pcp4l1 | Purkinje cell protein 4-like 1 | 30286.063 | 26746.836 | 15.045936 | 14.616347 | 30129.021 | 33010.934 | 27718.234 | 23991.023 | 29247.47 | 27002.014 | 15.000522 | 15.184723 | 14.952563 | 14.419968 | 14.660212 | 14.768861 |
| ASMM10P006 | 0.02640 | 0.3622928 | 1.8156981 | up | NM_001277 | Bmpr1b | bone morphogenetic protein receptor, type 1B | 932.35343 | 607.15322 | 10.077687 | 9.217163 | 945.8418 | 706.6945 | 1144.524 | 659.21576 | 561.72876 | 600.51514 | 9.87166 | 9.822331 | 10.53907 | 9.291833 | 9.025659 | 9.333996 |
| ASMM10P048 | 0.02641 | 0.3622928 | 1.4512456 | up | NM_019400 | Rabep1 | rabaptin, RAB GTPase binding effector protein 1 | 9382.0102 | 7470.4837 | 13.34712 | 12.809828 | 8848.4375 | 8874.733 | 10422.86 | 7260.4526 | 7403.5576 | 7747.441 | 13.13663 | 13.324612 | 13.580118 | 12.724928 | 12.71651 | 12.988047 |
| ASMM10P004 | 0.02644 | 0.3622928 | 1.4337142 | up | NM_001033 | Tatdn2 | TatD DNase domain containing 2 | 71.950107 | 59.396669 | 6.421292 | 5.901535 | 97.5736 | 52.53976 | 65.73696 | 63.1373 | 62.390377 | 52.66233 | 6.6597996 | 6.1533113 | 6.4507656 | 5.8859577 | 5.8480015 | 5.970645 |
| ASMM10P012 | 0.02646 | 0.3622928 | 1.2537256 | up | NM_007798 | Ctsb | cathepsin B | 1817.0627 | 1712.5515 | 11.024517 | 10.698295 | 2189.7349 | 1822.1844 | 1439.2688 | 1775.8751 | 1742.5692 | 1619.2103 | 11.083814 | 11.142753 | 10.846983 | 10.709658 | 10.642784 | 10.742443 |
| ASMM10P006 | 0.02651 | 0.3622928 | 1.3277553 | up | NM_007853 | Degs1 | degenerative spermatocyte homolog 1 (Drosophila) | 1771.9703 | 1600.2156 | 11.007999 | 10.599009 | 1924.1552 | 1764.8453 | 1626.9105 | 1524.1132 | 1578.0833 | 1698.4504 | 10.904612 | 11.100452 | 11.018932 | 10.489866 | 10.49862 | 10.808542 |
| ASMM10P009 | 0.02654 | 0.3622928 | 1.3408195 | up | NM_175312 | B630005N1 | RIKEN cDNA B630005N14 gene | 514.00897 | 464.75754 | 9.260289 | 8.837174 | 530.892 | 498.64597 | 512.48895 | 447.56223 | 503.0922 | 443.6182 | 9.039413 | 9.343277 | 9.398177 | 8.735362 | 8.864961 | 8.911199 |
| ASMM10P021 | 0.02655 | 0.3622928 | 1.2761934 | up | NM_026030 | Eif2s2 | eukaryotic translation initiation factor 2, subunit 2 (beta) | 826.82007 | 780.12007 | 9.930736 | 9.578889 | 857.24945 | 794.0144 | 829.19635 | 812.7941 | 805.2275 | 722.3386 | 9.733793 | 9.988808 | 10.069606 | 9.58624 | 9.546608 | 9.603818 |
| ASMM10P003 | 0.02658 | 0.3622928 | 1.3326956 | up | NM_027009 | Rfc3 | replication factor C (activator 1) 3 | 587.79458 | 530.18992 | 9.437432 | 9.023084 | 677.22797 | 534.9234 | 551.23236 | 524.66943 | 500.39703 | 565.5033 | 9.387515 | 9.437729 | 9.487051 | 8.959685 | 8.859 | 9.250568 |
| ASMM10P017 | 0.02658 | 0.3622928 | 4.8892885 | up | NM_001271 | Crem | cAMP responsive element modulator | 38.824383 | 9.631075 | 5.484583 | 3.194959 | 49.95447 | 20.605762 | 45.912918 | 5.147094 | 7.9602923 | 15.78584 | 5.7080975 | 4.8272767 | 5.918376 | 2.3942375 | 2.8726068 | 4.3180323 |
| ASMM10P006 | 0.02660 | 0.3623384 | 1.385765 | up | NM_008106 | Opn1mw | opsin 1 (cone pigments), medium-wave-sensitive (color blindness, deutan) | 38.046803 | 33.038433 | 5.531809 | 5.061126 | 42.43236 | 40.3755 | 31.332548 | 34.153347 | 38.27945 | 26.682503 | 5.470556 | 5.78077 | 5.3441005 | 4.9997115 | 5.149012 | 5.034655 |
| ASMM10P012 | 0.02668 | 0.3629418 | 1.494667 | up | NM_033325 | Loxl2 | lysyl oxidase-like 2 | 123.83369 | 102.65467 | 7.253509 | 6.673685 | 121.92334 | 129.8795 | 119.69822 | 96.674164 | 119.65328 | 91.63656 | 6.9709816 | 7.4508147 | 7.3387322 | 6.5009484 | 6.7853227 | 6.734785 |
| ASMM10P051 | 0.02671 | 0.363089 | 1.3638535 | up | NM_172646 | Dnm3 | dynamin 3 | 30451.427 | 26665.582 | 15.059257 | 14.611569 | 29724.12 | 28107.982 | 33522.18 | 26215.043 | 26741.678 | 27040.025 | 14.972752 | 14.940432 | 15.264588 | 14.553996 | 14.510149 | 14.770561 |
| ASMM10P051 | 0.02672 | 0.363089 | 1.6061648 | up | NM_026604 | Fam135a | family with sequence similarity 135, member A | 236.17116 | 173.21989 | 8.111646 | 7.428026 | 323.94678 | 157.60869 | 226.95801 | 177.7412 | 189.50356 | 152.4149 | 8.352144 | 7.7170763 | 8.2657175 | 7.3943024 | 7.4597607 | 7.430015 |
| ASMM10P050 | 0.02689 | 0.3634869 | 1.2576054 | up | NM_025522 | Dhrs7 | dehydrogenase/reductase (SDR family) member 7 | 16436.783 | 15393.508 | 14.154966 | 13.824287 | 15694.631 | 16236.787 | 17378.93 | 15112.815 | 16368.855 | 14698.855 | 14.002977 | 14.165382 | 14.296539 | 13.750289 | 13.81244 | 13.910131 |
| ASMM10P039 | 0.02697 | 0.3634869 | 2.3502579 | up | NM_001195 | Gm4975 | predicted gene 4975 | 27.34676 | 13.623784 | 5.022747 | 3.789928 | 28.04645 | 33.97886 | 20.01497 | 14.712562 | 11.230981 | 14.927809 | 4.861328 | 5.5309076 | 4.6760063 | 3.7547314 | 3.3734562 | 4.241597 |
| ASMM10P009 | 0.02698 | 0.3634869 | 1.3443353 | up | NM_175246 | Snip1 | Smad nuclear interacting protein 1 | 545.94189 | 482.35848 | 9.318948 | 8.892055 | 661.9458 | 419.3889 | 556.49097 | 499.53476 | 486.41745 | 461.12323 | 9.355751 | 9.099158 | 9.501934 | 8.892406 | 8.818461 | 8.965297 |
| ASMM10P053 | 0.02699 | 0.3634869 | 1.4163806 | up | NM_001136 | Eif2d | eukaryotic translation initiation factor 2D | 10287.978 | 8473.131 | 13.482193 | 12.979984 | 10563.668 | 9296.834 | 11003.432 | 8086.37 | 8107.839 | 9225.184 | 13.411757 | 13.39531 | 13.639511 | 12.873499 | 12.838615 | 13.227837 |
| ASMM10P011 | 0.02699 | 0.3634869 | 1.7533674 | up | NM_133684 | Marc2 | mitochondrial amidoxime reducing component 2 | 269.66011 | 186.40449 | 8.336692 | 7.526564 | 310.19794 | 206.55464 | 292.22775 | 179.47261 | 168.3565 | 211.38437 | 8.287202 | 8.107264 | 8.61561 | 7.408483 | 7.2818274 | 7.8893805 |
| ASMM10P033 | 0.02711 | 0.3634869 | 1.6793584 | up | NM_009013 | Rad51ap1 | RAD51 associated protein 1 | 610.42748 | 433.93614 | 9.485782 | 8.737872 | 606.5317 | 516.29285 | 708.4579 | 400.26895 | 439.50146 | 462.038 | 9.22176 | 9.39446 | 9.8411255 | 8.571213 | 8.674206 | 8.968196 |
| ASMM10P025 | 0.02714 | 0.3634869 | 1.4892705 | up | NM_029655 | Snx7 | sorting nexin 7 | 968.135 | 779.89916 | 10.150906 | 9.576301 | 1050.784 | 937.1128 | 916.5082 | 771.038 | 700.04205 | 868.61743 | 10.015553 | 10.219657 | 10.21751 | 9.519753 | 9.341376 | 9.867773 |
| ASMM10P029 | 0.02716 | 0.3634869 | 1.6154583 | up | NM_019660 | Mycbp | c-myc binding protein | 438.80663 | 328.87213 | 9.033476 | 8.341533 | 440.4892 | 386.41147 | 489.51923 | 301.41116 | 336.1685 | 349.03674 | 8.786777 | 8.9791975 | 9.334454 | 8.161312 | 8.282095 | 8.581191 |
| ASMM10P007 | 0.02723 | 0.3634869 | 1.2645068 | up | NM_178050 | Atl2 | atlastin GTPase 2 | 1001.8793 | 951.44417 | 10.202181 | 9.863606 | 1075.1996 | 969.9075 | 960.5308 | 905.7484 | 1023.37 | 925.2141 | 10.050505 | 10.269959 | 10.286079 | 9.741313 | 9.890386 | 9.959119 |
| ASMM10P054 | 0.02728 | 0.3634869 | 1.6588367 | up | NM_030137 | Cstad | CSA-conditional, T cell activation-dependent protein | 43.47291 | 31.258727 | 5.707901 | 4.977729 | 52.663025 | 30.75451 | 47.001194 | 27.99777 | 38.2573 | 27.521112 | 5.7816114 | 5.3914437 | 5.9506483 | 4.70722 | 5.1480412 | 5.0779266 |
| ASMM10P008 | 0.02732 | 0.3634869 | 1.3046759 | up | NM_146096 | Abhd17b | abhydrolase domain containing 17B | 363.16993 | 339.09521 | 8.770403 | 8.386711 | 410.44186 | 355.8796 | 323.18832 | 316.74133 | 353.88467 | 346.65964 | 8.686212 | 8.864739 | 8.760257 | 8.2289095 | 8.358289 | 8.572935 |
| ASMM10P025 | 0.02733 | 0.3634869 | 1.6496885 | up | NM_027206 | Tnfaip8l2 | tumor necrosis factor, alpha-induced protein 8-like 2 | 1983.5383 | 1420.223 | 11.146592 | 10.424399 | 2002.6218 | 1558.8784 | 2389.1147 | 1493.4493 | 1377.1229 | 1390.0967 | 10.961595 | 10.927929 | 11.550253 | 10.460442 | 10.296642 | 10.516112 |
| ASMM10P051 | 0.02733 | 0.3634869 | 1.2695117 | up | NM_009238 | Sox4 | SRY (sex determining region Y)-box 4 | 1108.6368 | 1043.5898 | 10.338787 | 9.994513 | 1209.2523 | 1167.8229 | 948.8352 | 1078.541 | 1129.4062 | 922.82214 | 10.216383 | 10.534644 | 10.265333 | 10.001292 | 10.028412 | 9.953835 |
| ASMM10P015 | 0.02734 | 0.3634869 | 2.7733805 | up | NM_001164 | Spats2l | spermatogenesis associated, serine-rich 2-like | 40.067494 | 16.470511 | 5.539605 | 4.067959 | 28.11251 | 31.573229 | 60.516743 | 16.298693 | 19.26801 | 13.844829 | 4.865133 | 5.4275885 | 6.326093 | 3.90602 | 4.1603804 | 4.1374774 |
| ASMM10P055 | 0.02735 | 0.3634869 | 1.5195646 | up | NM_199306 | Wdtc1 | WD and tetratricopeptide repeats 1 | 240.53495 | 193.66397 | 8.194717 | 7.591059 | 234.88443 | 227.8717 | 258.84872 | 185.24084 | 206.90598 | 188.8451 | 7.8987455 | 8.241963 | 8.443443 | 7.453788 | 7.5907774 | 7.728612 |
| ASMM10P013 | 0.02737 | 0.3634869 | 1.2737581 | up | NM_028003 | Rpap3 | RNA polymerase II associated protein 3 | 17480.775 | 16088.766 | 14.243325 | 13.894234 | 16618.725 | 17119.564 | 18704.035 | 16323.731 | 16496.773 | 15445.795 | 14.087436 | 14.238433 | 14.404107 | 13.872461 | 13.826068 | 13.984173 |
| ASMM10P022 | 0.02739 | 0.3634869 | 1.2817759 | up | NM_001004 | Gpr158 | G protein-coupled receptor 158 | 202.5148 | 192.50932 | 7.939789 | 7.581644 | 246.39717 | 191.59877 | 169.54845 | 195.78893 | 188.25262 | 193.4864 | 7.96968 | 8.004746 | 7.8449397 | 7.5340304 | 7.448549 | 7.762354 |
| ASMM10P007 | 0.02741 | 0.3634869 | 1.436287 | up | NM_010774 | Mbd4 | methyl-CpG binding domain protein 4 | 140.93774 | 120.29841 | 7.423457 | 6.901113 | 156.09547 | 112.8155 | 153.90225 | 136.78644 | 122.757 | 101.35178 | 7.318761 | 7.2444587 | 7.7071514 | 7.008737 | 6.82275 | 6.871852 |
| ASMM10P053 | 0.02742 | 0.3634869 | 1.5225722 | up | NM_172472 | Tfe3 | transcription factor E3 | 90.113753 | 72.383065 | 6.792122 | 6.185612 | 86.711784 | 87.47096 | 96.158516 | 72.275085 | 74.96353 | 69.91058 | 6.4885707 | 6.8781147 | 7.009681 | 6.0835476 | 6.116044 | 6.357243 |
| ASMM10P010 | 0.02743 | 0.3634869 | 1.4539671 | up | NM_028042 | Ercc8 | excision repaiross-complementing rodent repair deficiency, complementation group 8 | 55.035723 | 45.490682 | 6.065883 | 5.525888 | 64.38353 | 42.896957 | 57.826683 | 46.141293 | 45.49536 | 44.835392 | 6.0720882 | 5.8674903 | 6.2580695 | 5.435794 | 5.3929296 | 5.7489405 |
| ASMM10P008 | 0.02743 | 0.3634869 | 1.3717969 | up | NM_029349 | 2310035C2 | RIKEN cDNA 2310035C23 gene | 157.75585 | 142.4481 | 7.595853 | 7.139787 | 163.32309 | 153.94281 | 156.00166 | 166.37479 | 142.39246 | 118.57704 | 7.3817782 | 7.6811123 | 7.72467 | 7.297679 | 7.035666 | 7.0860147 |
| ASMM10P017 | 0.02750 | 0.3634869 | 1.5729175 | up | NM_001159 | Slc25a2 | solute carrier family 25 (mitochondrial carrier, ornithine transporter) member 2 | 197.9047 | 155.11412 | 7.921953 | 7.26851 | 187.40814 | 185.93694 | 220.36903 | 167.3606 | 156.84935 | 141.1324 | 7.581915 | 7.9576073 | 8.226337 | 7.305805 | 7.1753054 | 7.32442 |
| ASMM10P044 | 0.02750 | 0.3634869 | 1.3513622 | up | NM_019496 | Ammecr1 | Alport syndrome, mental retardation, midface hypoplasia and elliptocytosis chromosomal reg | 92.509447 | 84.894763 | 6.823313 | 6.388899 | 104.67468 | 81.71341 | 91.14025 | 103.63759 | 88.174164 | 62.872536 | 6.761752 | 6.778579 | 6.9296083 | 6.6074767 | 6.3492827 | 6.2099366 |
| ASMM10P029 | 0.02751 | 0.3634869 | 1.7371946 | up | NM_172696 | Inadl | InaD-like (Drosophila) | 2283.5828 | 1550.0115 | 11.352048 | 10.555289 | 2013.6289 | 2116.3706 | 2720.749 | 1428.371 | 1661.6079 | 1560.0557 | 10.96995 | 11.359316 | 11.726879 | 10.396976 | 10.574756 | 10.694135 |
| ASMM10P007 | 0.02763 | 0.3641171 | 1.4636013 | up | NM_011852 | Oas1g | 2'-5' oligoadenylate synthetase 1G | 89.358013 | 74.649342 | 6.771068 | 6.221545 | 101.82598 | 80.53028 | 85.71778 | 65.43298 | 81.41417 | 77.100876 | 6.7186484 | 6.7575808 | 6.836975 | 5.9396067 | 6.2369976 | 6.488032 |
| ASMM10P033 | 0.02764 | 0.3641171 | 1.2843342 | up | NM_010595 | Kcna1 | potassium voltage-gated channel, shaker-related subfamily, member 1 | 17350.396 | 15899.225 | 14.232306 | 13.871285 | 18843.076 | 15520.521 | 17687.59 | 14580.068 | 17551.898 | 15565.708 | 14.282898 | 14.099273 | 14.314747 | 13.712879 | 13.908446 | 13.992531 |
| ASMM10P031 | 0.02765 | 0.3641171 | 1.6615384 | up | NM_178799 | Acad12 | acyl-Coenzyme A dehydrogenase family, member 12 | 231.20718 | 171.62236 | 8.140161 | 7.407642 | 220.53137 | 219.79489 | 253.29527 | 151.03455 | 199.8336 | 163.99892 | 7.814513 | 8.193683 | 8.412288 | 7.1521378 | 7.5399327 | 7.5308547 |
| ASMM10P025 | 0.02768 | 0.3642519 | 1.54472 | up | NM_173347 | Prune | prune homolog (Drosophila) | 12104.612 | 9137.816 | 13.712041 | 13.084695 | 11566.245 | 11251.624 | 13495.966 | 8487.296 | 8670.341 | 10255.811 | 13.536951 | 13.661297 | 13.937874 | 12.947591 | 12.936742 | 13.369753 |
| ASMM10P052 | 0.02776 | 0.3647826 | 1.6350042 | up | NM_134214 | Vmn1r214 | vomeronasal 1 receptor 214 | 481.63173 | 350.3076 | 9.141996 | 8.432702 | 493.4781 | 372.6983 | 578.7188 | 377.57175 | 356.66455 | 316.6865 | 8.942534 | 8.927576 | 9.555878 | 8.486571 | 8.369218 | 8.442316 |
| ASMM10P051 | 0.02777 | 0.3647826 | 1.7843829 | up | NM_178911 | Pld4 | phospholipase D family, member 4 | 80.507785 | 53.390572 | 6.587942 | 5.752517 | 92.81689 | 53.360596 | 95.34587 | 59.077965 | 52.77622 | 48.31753 | 6.5902424 | 6.1774125 | 6.996172 | 5.7899776 | 5.6131654 | 5.8544083 |
| ASMM10P031 | 0.02791 | 0.3659022 | 1.2724431 | up | NM_019875 | Abcb9 | ATP-binding cassette, sub-family B (MDR/TAP), member 9 | 36432.329 | 34098.995 | 15.31773 | 14.970129 | 40474.02 | 31853.064 | 36969.902 | 33449.156 | 36600.81 | 32247.02 | 15.418987 | 15.124649 | 15.409554 | 14.905529 | 14.9758 | 15.029058 |
| ASMM10P027 | 0.02792 | 0.3659022 | 1.3660604 | up | NM_026344 | Dph2 | DPH2 homolog (S. cerevisiae) | 67.331915 | 58.823482 | 6.337665 | 5.887644 | 88.921394 | 56.355682 | 56.71867 | 60.912468 | 58.808296 | 56.749683 | 6.5280805 | 6.255343 | 6.229571 | 5.8323092 | 5.7614155 | 6.069206 |

| | | | | | | | | | | | | | | | | | | | | | | | |
|---|---|---|---|---|---|---|---|---|---|---|---|---|---|---|---|---|---|---|---|---|---|---|---|
| ASMM10P033 | 0.02792 | 0.3659022 | 2.0590816 | up | NM_183183 | Gprin3 | GPRIN family member 3 | 2172.617 | 1216.5773 | 11.254051 | 10.21205 | 1727.1802 | 2910.6504 | 1880.0204 | 1225.9396 | 1187.2535 | 1236.5388 | 10.750139 | 11.783896 | 11.228119 | 10.181651 | 10.094448 | 10.360052 |
| ASMM10P050 | 0.02797 | 0.3661464 | 1.3906682 | up | NM_020557 | Cmpk2 | cytidine monophosphate (UMP-CMP) kinase 2, mitochondrial | 176.54259 | 155.84177 | 7.748968 | 7.27319 | 202.95427 | 152.01755 | 174.65594 | 150.59335 | 154.58687 | 162.34508 | 7.69279 | 7.663319 | 7.890796 | 7.148298 | 7.15483 | 7.5164423 |
| ASMM10P019 | 0.02798 | 0.3661464 | 1.7410923 | up | NM_054042 | Cd248 | CD248 antigen, endosialin | 808.55692 | 564.07846 | 9.895473 | 9.09548 | 868.40155 | 741.7244 | 815.5448 | 490.59406 | 509.72122 | 691.9201 | 9.752144 | 9.890884 | 10.04339 | 8.865335 | 8.882126 | 9.538979 |
| ASMM10P018 | 0.02799 | 0.3661464 | 1.3782996 | up | NM_053145 | Pcdhb20 | protocadherin beta 20 | 8710.2425 | 7389.5382 | 13.254896 | 12.792006 | 7923.805 | 9197.2295 | 9009.693 | 7833.364 | 7310.4517 | 7024.799 | 12.997403 | 13.381302 | 13.385983 | 12.83147 | 12.697255 | 12.847294 |
| ASMM10P027 | 0.02810 | 0.3661846 | 1.3524201 | up | NM_026368 | Caap1 | caspase activity and apoptosis inhibitor 1 | 5253.1627 | 4593.1897 | 12.543884 | 12.108341 | 4955.62 | 5485.907 | 5317.961 | 4642.156 | 4647.0806 | 4490.3325 | 12.301069 | 12.670871 | 12.659713 | 12.087215 | 12.044144 | 12.193664 |
| ASMM10P012 | 0.02812 | 0.3661846 | 1.580278 | up | NM_027435 | Atad2 | ATPase family, AAA domain containing 2 | 1387.7737 | 1022.5806 | 10.62929 | 9.969112 | 1746.3665 | 958.6095 | 1458.3452 | 1015.138 | 1055.495 | 997.1087 | 10.766831 | 10.252877 | 10.868162 | 9.911907 | 9.934175 | 10.061253 |
| ASMM10P000 | 0.02814 | 0.3661846 | 2.28837 | up | NM_001003 | Kdm2b | lysine (K)-specific demethylase 2B | 715.749 | 362.80823 | 9.673529 | 8.479209 | 589.74054 | 552.9696 | 1004.5369 | 406.48734 | 379.5509 | 302.38644 | 9.182539 | 9.488079 | 10.349969 | 8.592123 | 8.462256 | 8.383247 |
| ASMM10P044 | 0.02816 | 0.3661846 | 1.8660643 | up | NM_053185 | Col4a6 | collagen, type IV, alpha 6 | 540.36006 | 348.61414 | 9.31132 | 8.411321 | 565.7216 | 443.99167 | 611.3669 | 295.8324 | 331.51593 | 418.4941 | 9.126543 | 9.177603 | 9.629814 | 8.136044 | 8.264367 | 8.833553 |
| ASMM10P017 | 0.02817 | 0.3661846 | 1.3298367 | up | NM_030141 | 1700061G19 | RIKEN cDNA 1700061G19 gene | 161.87422 | 150.00501 | 7.632095 | 7.220845 | 166.81464 | 169.84825 | 148.95978 | 157.24619 | 158.83171 | 133.93712 | 7.4094133 | 7.8251166 | 7.6617537 | 7.2134833 | 7.1945744 | 7.2544785 |
| ASMM10P014 | 0.02818 | 0.3661846 | 1.6679253 | up | ENSMUST( | Zbtb21 | zinc finger and BTB domain containing 21 | 812.26307 | 580.63602 | 9.890194 | 9.152139 | 798.5517 | 656.2227 | 982.0148 | 627.66 | 602.29614 | 511.95193 | 9.631134 | 9.718691 | 10.320757 | 9.218719 | 9.125712 | 9.111987 |
| ASMM10P025 | 0.02825 | 0.3664222 | 1.269473 | up | NM_025316 | Ndufb5 | NADH dehydrogenase (ubiquinone) 1 beta subcomplex, 5 | 14688.015 | 13561.937 | 13.988959 | 13.64473 | 14128.985 | 14346.504 | 15588.557 | 13644.46 | 13753.934 | 13287.416 | 13.835981 | 13.996813 | 14.134084 | 13.617939 | 13.56515 | 13.7511 |
| ASMM10P015 | 0.02828 | 0.3664222 | 1.5987328 | up | NM_133731 | Prss22 | protease, serine 22 | 520.62627 | 387.97706 | 9.240997 | 8.564068 | 623.82104 | 382.15356 | 555.9042 | 455.35583 | 419.25668 | 289.31866 | 9.2609005 | 8.961662 | 9.500429 | 8.761042 | 8.606095 | 8.325068 |
| ASMM10P054 | 0.02837 | 0.3664222 | 1.5289064 | up | NM_177340 | Synpo | synaptopodin | 29.420332 | 23.275595 | 5.169297 | 4.556797 | 32.437107 | 26.39343 | 29.430458 | 19.986069 | 28.000185 | 21.84053 | 5.0798917 | 5.173689 | 5.2543106 | 4.2075796 | 4.7040443 | 4.758767 |
| ASMM10P011 | 0.02839 | 0.3664222 | 1.3672785 | up | NM_013869 | Tnfrsf19 | tumor necrosis factor receptor superfamily, member 19 | 25547.843 | 21874.438 | 14.789113 | 14.337806 | 24954.918 | 23321.959 | 28366.652 | 19971.725 | 23992.182 | 21659.408 | 14.705977 | 14.674238 | 14.987124 | 14.169287 | 14.364858 | 14.479272 |
| ASMM10P026 | 0.02843 | 0.3664222 | 1.3251279 | up | NM_144896 | Pet112 | PET112 homolog (S. cerevisiae) | 1280.2294 | 1161.5392 | 10.550946 | 10.144814 | 1303.0771 | 1247.0402 | 1290.5708 | 1244.3672 | 1143.7518 | 1096.4985 | 10.335505 | 10.620921 | 10.696412 | 10.202234 | 10.043679 | 10.18853 |
| ASMM10P027 | 0.02849 | 0.3664222 | 1.9530297 | up | NM_011660 | Txn1 | thioredoxin 1 | 51.489285 | 32.331076 | 5.988218 | 5.022504 | 52.713074 | 48.3022 | 53.45258 | 29.113857 | 28.733603 | 39.145767 | 5.7831793 | 6.038182 | 6.1432934 | 4.7670727 | 4.743268 | 5.5571723 |
| ASMM10P050 | 0.02850 | 0.3664222 | 1.3984897 | up | NM_001037 | Prps1l3 | phosphoribosyl pyrophosphate synthetase 1-like 3 | 3374.5177 | 2846.8041 | 11.910962 | 11.427092 | 3232.3296 | 3517.3972 | 3373.8264 | 2786.401 | 2889.6943 | 2864.317 | 11.648514 | 12.055811 | 12.02856 | 11.35633 | 11.379467 | 11.545479 |
| ASMM10P007 | 0.02858 | 0.3664222 | 1.259282 | up | NM_026158 | Isoc2b | isochorismatase domain containing 2b | 151.55487 | 147.02798 | 7.524072 | 7.191471 | 187.70818 | 133.25192 | 133.70451 | 159.50815 | 139.94035 | 141.63544 | 7.5838346 | 7.4884477 | 7.4999337 | 7.2337813 | 7.010557 | 7.3300734 |
| ASMM10P035 | 0.02866 | 0.3664222 | 1.4285793 | up | NM_011980 | Zfp146 | zinc finger protein 146 | 152.93895 | 131.92466 | 7.552774 | 7.038193 | 150.16318 | 151.92427 | 156.72939 | 137.64099 | 135.61299 | 122.52 | 7.2646017 | 7.662196 | 7.7315254 | 7.0159917 | 6.9652195 | 7.1333685 |
| ASMM10P001 | 0.02875 | 0.3664222 | 1.743304 | up | NM_001252 | Cizl | CDKN1A interacting zinc finger protein 1 | 139.95238 | 92.85362 | 7.337409 | 6.535585 | 211.87364 | 103.93022 | 104.05328 | 106.60489 | 85.15874 | 86.79723 | 7.7563396 | 7.127108 | 7.128779 | 6.648801 | 6.301996 | 6.655957 |
| ASMM10P043 | 0.02877 | 0.3664222 | 1.2955072 | up | NM_023873 | Cep70 | centrosomal protein 70 | 680.30714 | 631.67076 | 9.646168 | 9.272651 | 737.6153 | 584.86053 | 718.4456 | 675.8014 | 663.43823 | 555.77264 | 9.518422 | 9.559469 | 9.860614 | 9.327262 | 9.263203 | 9.227489 |
| ASMM10P009 | 0.02878 | 0.3664222 | 1.414568 | up | NM_172517 | Rbbp5 | retinoblastoma binding protein 5 | 78.272223 | 67.620615 | 6.587741 | 6.08738 | 80.38081 | 80.55848 | 73.87738 | 69.67106 | 66.69917 | 66.491615 | 6.386116 | 6.758269 | 6.618839 | 6.0289993 | 5.9452944 | 6.2878456 |
| ASMM10P008 | 0.02881 | 0.3664222 | 1.5386607 | up | NM_026465 | Prorsd1 | prolyl-tRNA synthetase domain containing 1 | 196.11997 | 157.88646 | 7.908569 | 7.286894 | 207.64807 | 198.38686 | 182.32497 | 137.65718 | 167.58998 | 168.41222 | 7.727498 | 8.051012 | 7.9471984 | 7.0161448 | 7.2752943 | 7.569244 |
| ASMM10P039 | 0.02881 | 0.3664222 | 1.308338 | up | NM_172473 | Hace1 | HECT domain and ankyrin repeat containing, E3 ubiquitin protein ligase 1 | 608.77007 | 559.62868 | 9.490082 | 9.102346 | 624.8644 | 614.2186 | 587.2272 | 574.26855 | 576.3793 | 528.2382 | 9.265683 | 9.628239 | 9.576323 | 9.09067 | 9.065044 | 9.151325 |
| ASMM10P035 | 0.02881 | 0.3664222 | 1.4162982 | up | NM_011909 | Usp18 | ubiquitin specific peptidase 18 | 164.50316 | 142.0205 | 7.63673 | 7.134605 | 198.51895 | 162.8321 | 132.15842 | 125.53657 | 166.15091 | 134.37401 | 7.66121 | 7.7646894 | 7.484291 | 6.8825397 | 7.2623076 | 7.258968 |
| ASMM10P053 | 0.02883 | 0.3664222 | 1.2664587 | up | NM_001077 | Tsks | testis-specific serine kinase substrate | 115.7245 | 110.73506 | 7.130256 | 6.789456 | 140.96848 | 93.10204 | 113.10298 | 114.72172 | 113.67999 | 103.80348 | 7.1725054 | 6.968063 | 7.2501993 | 6.7506247 | 6.7147317 | 6.9030113 |
| ASMM10P011 | 0.02888 | 0.3664222 | 1.3635816 | up | NM_025697 | Spryd7 | SPRY domain containing 7 | 2607.7528 | 2262.302 | 11.546006 | 11.098605 | 2720.6465 | 2537.5847 | 2565.0273 | 2267.8284 | 2120.3462 | 2398.7314 | 11.389653 | 11.604231 | 11.644135 | 11.067108 | 10.928868 | 11.29984 |
| ASMM10P031 | 0.02889 | 0.3664222 | 1.7939946 | up | NM_198621 | Cct8l1 | chaperonin containing TCP1, subunit 8 (theta)-like 1 | 119.46963 | 81.399419 | 7.183912 | 6.340736 | 103.27677 | 145.12943 | 110.00269 | 92.06096 | 85.737816 | 66.39948 | 6.740936 | 7.60226 | 7.208539 | 6.4252367 | 6.3105927 | 6.286379 |
| ASMM10P043 | 0.02892 | 0.3664222 | 1.4440176 | up | NM_001039 | Gtf2a2 | general transcription factor II A, 2 | 873.22255 | 727.43033 | 10.007456 | 9.477367 | 855.98895 | 812.9166 | 950.7621 | 780.6009 | 735.144 | 666.5461 | 9.731449 | 10.023241 | 10.267677 | 9.534174 | 9.414097 | 9.483831 |
| ASMM10P008 | 0.02894 | 0.3664222 | 1.4324026 | up | NM_028097 | Tmem68 | transmembrane protein 68 | 1445.3063 | 1208.9931 | 10.719771 | 10.201334 | 1432.6016 | 1483.2969 | 1420.0204 | 1201.3208 | 1165.3898 | 1260.2688 | 10.471392 | 10.856802 | 10.83112 | 10.152063 | 10.070452 | 10.381488 |
| ASMM10P029 | 0.02897 | 0.3664222 | 1.3737977 | up | NM_027830 | Mcm9 | minichromosome maintenance complex component 9 | 18.701198 | 16.054874 | 4.497289 | 4.039119 | 21.773903 | 15.17986 | 19.14983 | 15.33688 | 17.99268 | 14.835061 | 4.4900174 | 4.3878965 | 4.6139517 | 3.8168828 | 4.067992 | 4.232482 |
| ASMM10P007 | 0.02898 | 0.3664222 | 1.5242515 | up | NM_019586 | Ube2j1 | ubiquitin-conjugating enzyme E2J 1 | 583.61875 | 459.19267 | 9.42633 | 8.818229 | 634.00757 | 631.82153 | 485.02716 | 452.72305 | 423.20532 | 501.64963 | 9.289786 | 9.665683 | 9.32352 | 8.751216 | 8.621225 | 9.082245 |
| ASMM10P025 | 0.02900 | 0.3664222 | 1.9829913 | up | NM_054045 | Hist2h3c2 | histone cluster 2, H3c2 | 2155.4354 | 1279.446 | 11.263204 | 10.275526 | 1763.5625 | 2577.8696 | 2124.874 | 1179.3533 | 1196.696 | 1462.2886 | 10.781957 | 11.624657 | 11.382999 | 10.128771 | 10.106328 | 10.591479 |
| ASMM10P054 | 0.02900 | 0.3664222 | 1.7394037 | up | NM_172567 | Mettl2 | methyltransferase like 2 | 103.17102 | 72.714893 | 6.982676 | 6.184083 | 88.24023 | 104.1133 | 117.15951 | 79.09938 | 79.0295 | 60.0158 | 6.515586 | 7.129666 | 7.302775 | 6.2092075 | 6.194813 | 6.148228 |
| ASMM10P004 | 0.02912 | 0.3664222 | 1.4245231 | up | NM_001081 | Sh2b1 | SH2B adaptor protein 1 | 108.30306 | 92.799095 | 7.043928 | 6.533449 | 127.41662 | 99.77532 | 97.71725 | 85.404724 | 94.51675 | 98.47581 | 7.029177 | 7.0691233 | 7.033483 | 6.319078 | 6.4520216 | 6.8292465 |
| ASMM10P008 | 0.02920 | 0.3664222 | 1.4942922 | up | NM_134068 | Dusp22 | dual specificity phosphatase 22 | 1203.67 | 961.1616 | 10.458958 | 9.879496 | 1152.9719 | 1123.7777 | 1334.2604 | 968.52295 | 1006.9336 | 908.02826 | 10.147413 | 10.487889 | 10.741572 | 9.842802 | 9.864244 | 9.931441 |
| ASMM10P035 | 0.02921 | 0.3664222 | 1.3849533 | up | NM_010908 | Nfkbib | nuclear factor of kappa light polypeptide gene enhancer in B cells inhibitor, beta | 440.45097 | 384.70043 | 9.039325 | 8.569488 | 471.92972 | 378.4188 | 471.0044 | 384.71786 | 389.35577 | 380.02765 | 8.883188 | 8.946877 | 9.287911 | 8.511271 | 8.498834 | 8.698359 |
| ASMM10P045 | 0.02927 | 0.3664222 | 2.5602277 | up | NM_011667 | Uba1y | ubiquitin-activating enzyme, Chr Y | 37.775444 | 16.038969 | 5.348933 | 3.992661 | 65.90791 | 20.831032 | 26.58739 | 17.606796 | 19.958233 | 10.551878 | 6.104303 | 4.8440013 | 5.0984945 | 4.0186462 | 4.2137303 | 3.745606 |
| ASMM10P000 | 0.02927 | 0.3664222 | 2.3945626 | up | NM_001271 | Dnmt3b | DNA methyltransferase 3B | 131.49917 | 65.394946 | 7.273456 | 6.013694 | 166.87639 | 82.137634 | 145.48347 | 64.96281 | 51.219467 | 80.00256 | 7.4100785 | 6.7862988 | 7.6239905 | 5.928973 | 5.5701923 | 6.541916 |
| ASMM10P012 | 0.02948 | 0.3664222 | 1.3584405 | up | NM_080556 | Tm9sf2 | transmembrane 9 superfamily member 2 | 787.05058 | 696.57199 | 9.859666 | 9.417715 | 844.3073 | 750.14685 | 766.6976 | 705.46606 | 652.4717 | 731.7782 | 9.71167 | 9.909701 | 9.957628 | 9.391146 | 9.241921 | 9.620078 |
| ASMM10P041 | 0.02952 | 0.3664222 | 1.2982339 | up | NM_178027 | Vps26b | vacuolar protein sorting 26 homolog B (yeast) | 6567.4491 | 5947.3539 | 12.854478 | 12.477928 | 6409.7754 | 6185.347 | 7107.225 | 6107.3027 | 6468.429 | 5266.33 | 12.674223 | 12.837453 | 13.051758 | 12.480977 | 12.529811 | 12.422995 |
| ASMM10P041 | 0.02953 | 0.3664222 | 1.3316091 | up | NM_001113 | Gria4 | glutamate receptor, ionotropic, AMPA4 (alpha 4) | 3141.1066 | 2779.7312 | 11.806964 | 11.393793 | 3184.7024 | 2867.7158 | 3370.9016 | 2842.272 | 2790.0486 | 2706.873 | 11.625422 | 11.768315 | 12.027154 | 11.384131 | 11.327374 | 11.469875 |
| ASMM10P044 | 0.02955 | 0.3664222 | 1.6747609 | up | NM_021487 | Kcne1l | potassium voltage-gated channel, Isk-related family, member 1-like, pseudogene | 2819.509 | 1977.6465 | 11.645552 | 10.901597 | 2508.437 | 3226.9048 | 2723.1853 | 1831.2555 | 1977.8281 | 2123.856 | 11.274275 | 11.932582 | 11.729799 | 10.758719 | 10.823957 | 11.122115 |
| ASMM10P038 | 0.02955 | 0.3664222 | 1.6604522 | up | NM_001136 | Lsp1 | lymphocyte specific 1 | 24.744817 | 17.519677 | 4.886126 | 4.154549 | 27.41307 | 28.619032 | 18.20235 | 18.78734 | 19.933214 | 13.838478 | 4.825365 | 5.291852 | 4.5411596 | 4.1151595 | 4.211777 | 4.1367116 |
| ASMM10P015 | 0.02956 | 0.3664222 | 1.2629641 | up | NM_010515 | Igf2r | insulin-like growth factor 2 receptor | 22619.241 | 20861.389 | 14.614418 | 14.277605 | 20976.832 | 23836.266 | 23044.625 | 21194.828 | 21069.582 | 20319.756 | 14.449593 | 14.710682 | 14.68298 | 14.256933 | 14.187379 | 14.388502 |
| ASMM10P037 | 0.02958 | 0.3664222 | 1.4237002 | up | NM_001037 | Nell1 | NEL-like 1 | 472.32347 | 404.40788 | 9.14479 | 8.635144 | 507.14676 | 450.42368 | 459.39996 | 368.25903 | 409.64362 | 435.321 | 8.983066 | 9.199505 | 9.251798 | 8.448085 | 8.572256 | 8.885092 |
| ASMM10P006 | 0.02959 | 0.3664222 | 1.3216582 | up | NM_001271 | Rtn3 | reticulon 3 | 285.79338 | 261.95473 | 8.421587 | 8.019238 | 349.14545 | 273.1847 | 235.05 | 301.92224 | 241.84625 | 242.09569 | 8.456905 | 8.497894 | 8.309962 | 8.163801 | 7.8097587 | 8.084154 |
| ASMM10P055 | 0.02966 | 0.3664222 | 1.3835968 | up | NM_026654 | Toe1 | target of EGR1, member 1 (nuclear) | 201.59848 | 179.91956 | 7.950269 | 7.481846 | 211.01321 | 190.11183 | 203.67041 | 193.64635 | 170.65749 | 175.45485 | 7.7513113 | 7.993576 | 8.105921 | 7.517535 | 7.3018637 | 7.626139 |
| ASMM10P040 | 0.02966 | 0.3664222 | 1.3491778 | up | NM_001024 | Pdp2 | pyruvate dehyrogenase phosphatase catalytic subunit 2 | 5376.5737 | 4698.457 | 12.575626 | 12.143546 | 5773.875 | 4771.797 | 5584.049 | 4237.397 | 4970.8105 | 4887.1636 | 12.522379 | 12.47388 | 12.730619 | 11.96062 | 12.148829 | 12.321188 |
| ASMM10P004 | 0.02967 | 0.3664222 | 1.2856411 | up | NM_001007 | Slc7a6os | solute carrier family 7, member 6 opposite strand | 573.72303 | 537.41905 | 9.406534 | 9.044046 | 605.9926 | 577.133 | 538.0435 | 528.71265 | 558.1167 | 525.4278 | 9.220259 | 9.540898 | 9.458445 | 8.970318 | 9.016169 | 9.145651 |
| ASMM10P019 | 0.02968 | 0.3664222 | 1.6039567 | up | NM_026553 | Yif1a | Yip1 interacting factor homolog A (S. cerevisiae) | 714.71233 | 540.31548 | 9.723632 | 9.041997 | 724.0008 | 733.4897 | 686.6465 | 477.5369 | 524.56195 | 618.8476 | 9.493964 | 9.877799 | 9.799133 | 8.82393 | 8.9262705 | 9.37579 |
| ASMM10P032 | 0.02969 | 0.3664222 | 2.3466354 | up | NM_029363 | 2010109A12 | RIKEN cDNA 2010109A12 gene | 36.532051 | 19.105313 | 5.468371 | 4.237777 | 41.304432 | 27.627745 | 40.663975 | 13.118129 | 22.67752 | 21.52029 | 5.4309306 | 5.235619 | 5.738562 | 3.5840118 | 4.3910403 | 4.7382784 |
| ASMM10P020 | 0.02973 | 0.3664222 | 1.3875198 | up | NM_182994 | Arl5a | ADP-ribosylation factor-like 5A | 545.83417 | 473.997 | 9.336902 | 8.864394 | 585.6428 | 583.1924 | 468.6673 | 521.36426 | 444.02512 | 456.60162 | 9.173832 | 9.555507 | 9.281368 | 8.950889 | 8.691192 | 8.951101 |
| ASMM10P019 | 0.02978 | 0.3664222 | 1.2801972 | up | NM_153597 | Trpt1 | tRNA phosphotransferase 1 | 215.57037 | 205.32017 | 8.028088 | 7.671722 | 246.47903 | 226.9655 | 173.26659 | 212.84076 | 221.98952 | 181.13023 | 7.9701686 | 8.236178 | 7.8779173 | 7.657613 | 7.689458 | 7.6680946 |
| ASMM10P004 | 0.02981 | 0.3664222 | 1.4667849 | up | NM_001077 | Acn9 | ACN9 homolog (S. cerevisiae) | 229.38106 | 192.22441 | 8.131905 | 7.579248 | 229.49434 | 242.38208 | 216.26675 | 188.51582 | 192.9661 | 195.19131 | 7.8696136 | 8.326403 | 8.199699 | 7.4769177 | 7.487421 | 7.773405 |
| ASMM10P025 | 0.02985 | 0.3664222 | 1.3926529 | up | NM_012012 | Exo1 | exonuclease 1 | 38.266888 | 32.837715 | 5.533013 | 5.055177 | 47.552532 | 34.956623 | 32.291508 | 30.226685 | 37.504826 | 30.781633 | 5.6358585 | 5.5760865 | 5.3870935 | 4.816178 | 5.119154 | 5.2301993 |
| ASMM10P008 | 0.02985 | 0.3664222 | 2.0362448 | up | NM_027210 | Ceacam13 | carcinoembryonic antigen-related cell adhesion molecule 13 | 153.34258 | 89.432597 | 7.509554 | 6.483643 | 162.03346 | 102.94943 | 195.04486 | 93.33972 | 82.71635 | 92.24172 | 7.371918 | 7.1133127 | 8.043431 | 6.446543 | 6.2615037 | 6.742882 |
| ASMM10P025 | 0.02986 | 0.3664222 | 1.8021055 | up | NM_009264 | Sprr1a | small proline-rich protein 1A | 87.970023 | 58.728266 | 6.737445 | 5.887761 | 85.3046 | 106.55438 | 72.051094 | 60.182297 | 55.599865 | 60.402637 | 6.46856 | 7.1629796 | 6.5807943 | 5.8174615 | 5.6864944 | 6.1593275 |
| ASMM10P004 | 0.03000 | 0.3664222 | 1.3404283 | up | NM_001252 | Usp33 | ubiquitin specific peptidase 33 | 804.05132 | 723.87423 | 9.890031 | 9.467337 | 807.6226 | 786.3455 | 818.18585 | 783.0263 | 755.3846 | 633.2118 | 9.648431 | 9.973431 | 10.048232 | 9.537503 | 9.452762 | 9.411747 |
| ASMM10P008 | 0.03004 | 0.3664222 | 1.2728252 | up | NM_028475 | Ubap2l | ubiquitin associated protein 2-like | 2230.3391 | 2090.562 | 11.33166 | 10.983626 | 2314.3872 | 2093.9482 | 2282.682 | 2001.4955 | 2238.897 | 2031.2936 | 11.166616 | 11.345811 | 11.482554 | 10.882896 | 11.006908 | 11.061074 |
| ASMM10P034 | 0.03012 | 0.3664222 | 1.2941579 | up | NM_026011 | Arl8b | ADP-ribosylation factor-like 8B | 2213.8977 | 2033.7402 | 11.31712 | 10.945106 | 2281.101 | 2400.197 | 1960.3951 | 2135.5803 | 2142.7764 | 1822.864 | 11.148036 | 11.527058 | 11.276266 | 10.979011 | 10.944019 | 10.912289 |
| ASMM10P043 | 0.03012 | 0.3664222 | 1.4249825 | up | NM_053216 | 1700102P08 | RIKEN cDNA 1700102P08 gene | 80.79097 | 69.020884 | 6.620776 | 6.109832 | 89.32953 | 86.407005 | 66.636375 | 80.62835 | 65.51366 | 60.920643 | 6.533921 | 6.8594346 | 6.468973 | 6.239787 | 5.919266 | 6.170443 |
| ASMM10P007 | 0.03018 | 0.3664222 | 1.7623916 | up | NM_021272 | Fabp7 | fatty acid binding protein 7, brain | 6213.2991 | 4113.0424 | 12.763991 | 11.946456 | 5506.249 | 7544.6733 | 5588.975 | 3592.1023 | 4070.8428 | 4676.182 | 12.452913 | 13.107071 | 12.731988 | 11.719958 | 11.862524 | 12.256887 |
| ASMM10P035 | 0.03021 | 0.3664222 | 3.9878877 | up | NM_175628 | A2m | alpha-2-macroglobulin | 1662.0481 | 521.46646 | 10.845724 | 8.8501 | 2006.3889 | 963.6465 | 2016.1089 | 265.80933 | 477.7172 | 820.87286 | 10.963547 | 10.261699 | 11.311927 | 7.9771967 | 8.790431 | 9.782671 |
| ASMM10P013 | 0.03022 | 0.3664222 | 1.4225529 | up | NM_028829 | Paqr8 | progestin and adipoQ receptor family member VIII | 38.652491 | 32.92339 | 5.571167 | 5.062684 | 40.154476 | 35.75355 | 40.049446 | 30.806925 | 36.42604 | 31.537205 | 5.3905315 | 5.610106 | 5.712863 | 4.844898 | 5.07679 | 5.2663655 |
| ASMM10P000 | 0.03024 | 0.3664222 | 1.3159597 | up | ENSMUST( | Rsrc2 | arginine/serine-rich coiled-coil 2 | 1118.4302 | 1016.7429 | 10.352922 | 9.956807 | 1187.3225 | 982.1485 | 1185.8197 | 1092.2607 | 1061.9673 | 896.0006 | 10.192046 | 10.288431 | 10.578289 | 10.018448 | 9.941659 | 9.910313 |
| ASMM10P033 | 0.03025 | 0.3664222 | 1.3437328 | up | NM_026345 | Mansc1 | MANSC domain containing 1 | 4304.4268 | 3755.0554 | 12.245024 | 11.818777 | 5490.518 | 3448.7864 | 3973.976 | 3622.3901 | 4112.514 | 3530.2622 | 12.447731 | 12.026247 | 12.261093 | 11.733502 | 11.87963 | 11.8432 |
| ASMM10P007 | 0.03025 | 0.3664222 | 1.5939062 | up | NM_011331 | Ccl12 | chemokine (C-C motif) ligand 12 | 31.428647 | 23.704134 | 5.264463 | 4.591897 | 30.115635 | 29.255888 | 34.914417 | 22.203817 | 26.251263 | 22.657322 | 4.9680996 | 5.3233695 | 5.501921 | 4.357137 | 4.607801 | 4.810752 |
| ASMM10P012 | 0.03034 | 0.3664222 | 1.3849658 | up | NM_001024 | Kbtbd7 | kelch repeat and BTB (POZ) domain containing 7 | 15887.228 | 13421.425 | 14.101475 | 13.631625 | 14267.827 | 16387.705 | 17006.152 | 13025.062 | 13813.249 | 13425.964 | 13.85895 | 14.181718 | 14.263757 | 13.546188 | 13.573967 | 13.774719 |
| ASMM10P026 | 0.03036 | 0.3664222 | 1.3916134 | up | NM_178214 | Hist2h2be | histone cluster 2, H2be | 3785.08 | 3226.9445 | 12.077166 | 11.600408 | 3648.2617 | 3690.0151 | 4016.9631 | 3028.5393 | 3538.533 | 3113.7612 | 11.835768 | 12.118279 | 12.277452 | 11.468038 | 11.669195 | 11.66399 |
| ASMM10P023 | 0.03037 | 0.3664222 | 1.4524209 | up | NM_178404 | Zc3h6 | zinc finger CCCH type containing 6 | 93.859764 | 79.171764 | 6.847098 | 6.308638 | 102.60131 | 90.241936 | 88.736046 | 74.70447 | 78.559166 | 84.251656 | 6.729636 | 6.9231625 | 6.8884954 | 6.1288075 | 6.1847396 | 6.612368 |
| ASMM10P016 | 0.03038 | 0.3664222 | 1.7081349 | up | NM_011553 | Tcp10a | t-complex protein 10a | 117.59588 | 84.358588 | 7.158199 | 6.385777 | 141.7433 | 101.20031 | 109.84403 | 82.81671 | 71.621155 | 98.6379 | 7.1791863 | 7.088627 | 7.2067842 | 6.276357 | 6.0491714 | 6.8318033 |
| ASMM10P028 | 0.03039 | 0.3664222 | 1.3776299 | up | NM_009788 | Calb1 | calbindin 1 | 6374.9788 | 5429.9589 | 12.815811 | 12.353622 | 6289.654 | 6753.485 | 6081.7974 | 5495.8887 | 5108.313 | 5685.675 | 12.64481 | 12.967768 | 12.834854 | 12.330054 | 12.184675 | 12.546138 |
| ASMM10P047 | 0.03041 | 0.3664222 | 1.5189984 | up | NM_027136 | Ovca2 | candidate tumor suppressor in ovarian cancer 2 | 1122.2758 | 888.41561 | 10.368173 | 9.765053 | 1087.5948 | 1117.445 | 1161.7877 | 868.2316 | 861.07623 | 935.939 | 10.069938 | 10.479795 | 10.554787 | 9.678845 | 9.642149 | 9.974165 |
| ASMM10P001 | 0.03046 | 0.3664222 | 1.7855712 | up | NM_009377 | Th | tyrosine hydroxylase | 15.955353 | 10.55427 | 4.262783 | 3.426397 | 13.912831 | 14.796029 | 19.1572 | 10.321561 | 12.04565 | 9.295598 | 3.8208635 | 4.3527327 | 4.6147532 | 3.232391 | 3.468371 | 3.5784304 |
| ASMM10P048 | 0.03052 | 0.3664222 | 1.3976505 | up | NM_008143 | Gnb2l1 | guanine nucleotide binding protein (G protein), beta polypeptide 2 like 1 | 467.83419 | 406.97692 | 9.132366 | 8.649363 | 479.66452 | 462.43796 | 461.4001 | 394.6797 | 406.68137 | 419.5697 | 8.905516 | 9.23237 | 9.259213 | 8.550568 | 8.560982 | 8.836538 |
| ASMM10P005 | 0.03053 | 0.3664222 | 1.5266829 | up | NM_001177 | Nfkb2 | nuclear factor of kappa light polypeptide gene enhancer in B cells 2, p49/p100 | 127.28439 | 102.99572 | 7.290415 | 6.680014 | 134.1787 | 124.00928 | 123.66518 | 96.21479 | 101.47127 | 111.3011 | 7.0997243 | 7.3852077 | 7.3863125 | 6.492861 | 6.5490313 | 6.998151 |
| ASMM10P015 | 0.03059 | 0.3664222 | 1.9430181 | up | NM_172450 | 4930539E08 | RIKEN cDNA 4930539E08 gene | 3789.2376 | 2316.0476 | 12.063394 | 11.105095 | 3806.738 | 3132.5193 | 4428.4556 | 2100.1475 | 1938.7279 | 2909.2673 | 11.901119 | 11.884955 | 12.404108 | 10.955879 | 10.788891 | 11.570514 |
| ASMM10P008 | 0.03060 | 0.3664222 | 1.39832 | up | NM_133739 | Tmem123 | transmembrane protein 123 | 77.524018 | 67.863161 | 6.5701 | 6.086405 | 86.05218 | 74.65258 | 71.867294 | 60.418453 | 76.60573 | 66.5653 | 6.478671 | 6.6545277 | 6.5771003 | 5.821756 | 6.147499 | 6.2899604 |
| ASMM10P037 | 0.03061 | 0.3664222 | 1.2978085 | up | NM_001111 | Nav2 | neuron navigator 2 | 31.733564 | 29.713359 | 5.275187 | 4.899109 | 34.92971 | 27.940428 | 32.330555 | 29.45833 | 36.873417 | 22.80833 | 5.1833754 | 5.253345 | 5.3888397 | 4.782861 | 5.0931964 | 4.82127 |
| ASMM10P006 | 0.03065 | 0.3664222 | 2.0086869 | up | NM_007935 | Epc1 | enhancer of polycomb homolog 1 (Drosophila) | 34.720045 | 21.119646 | 5.399291 | 4.393038 | 30.345102 | 40.414272 | 33.40076 | 29.311113 | 18.349005 | 15.69882 | 4.9793205 | 5.7825856 | 5.435966 | 4.774874 | 4.094513 | 4.309727 |
| ASMM10P006 | 0.03065 | 0.3664222 | 1.471245 | up | NM_001253 | Gpm6a | glycoprotein m6a | 7832.5407 | 6240.5976 | 13.108009 | 12.550971 | 7635.415 | 6766.39 | 9095.817 | 6406.56 | 5951.7915 | 6363.4414 | 12.951433 | 12.969605 | 13.402988 | 12.552611 | 12.403657 | 12.696645 |
| ASMM10P012 | 0.03066 | 0.3664222 | 1.5171538 | up | NM_145469 | Nipal2 | NIPA-like domain containing 2 | 464.63397 | 370.73499 | 9.112961 | 8.511594 | 447.6057 | 419.6113 | 526.6849 | 378.44806 | 412.5674 | 321.1895 | 8.80776 | 9.100339 | 9.430784 | 8.489898 | 8.582983 | 8.4619 |
| ASMM10P007 | 0.03068 | 0.3664222 | 1.2880108 | up | NM_019665 | Arl6 | ADP-ribosylation factor-like 6 | 3058.9933 | 2806.6135 | 11.772585 | 11.407441 | 3242.752 | 3042.4487 | 2891.7793 | 2807.2017 | 2683.0193 | 2929.6196 | 11.653684 | 11.844795 | 11.819277 | 11.366635 | 11.268755 | 11.586932 |
| ASMM10P017 | 0.03071 | 0.3664222 | 1.3538933 | up | NM_021522 | Usp14 | ubiquitin specific peptidase 14 | 3720.1477 | 3257.0099 | 12.052626 | 11.615512 | 3936.817 | 3561.4834 | 3662.1428 | 3332.2236 | 2946.1677 | 3492.6384 | 11.948633 | 12.073008 | 12.136238 | 11.615039 | 11.40502 | 11.826478 |

| | | | | | | | | | | | | | | | | | | | | | | | |
|---|---|---|---|---|---|---|---|---|---|---|---|---|---|---|---|---|---|---|---|---|---|---|---|
| ASMM10P024 | 0.03071 | 0.3664222 | 1.4083442 | up | NM_001099 | Gm14305 | predicted gene 14305 | 3688.1697 | 3140.1344 | 12.043515 | 11.549515 | 3779.5378 | 3447.0337 | 3837.9375 | 3052.5757 | 3811.6406 | 2556.1868 | 11.89028 | 12.025417 | 12.214848 | 11.482125 | 11.779093 | 11.387327 |
| ASMM10P030 | 0.03075 | 0.3664222 | 1.2905959 | up | NM_010305 | Gnai1 | guanine nucleotide binding protein (G protein), alpha inhibiting 1 | 69160.31 | 68138.601 | 16.332848 | 15.964811 | 69116.664 | 64384.605 | 73979.66 | 67943.13 | 72795.51 | 63677.164 | 16.25273 | 16.220432 | 16.525383 | 15.888452 | 15.928571 | 16.07741 |
| ASMM10P025 | 0.03076 | 0.3664222 | 4.2767608 | up | NM_178697 | Clca5 | chloride channel calcium activated 5 | 24.045978 | 5.34835 | 4.642852 | 2.546334 | 33.788048 | 29.809235 | 8.540652 | 4.9999995 | 4.9999995 | 6.0450525 | 5.137015 | 5.3461847 | 3.4453566 | 2.3250442 | 2.3265402 | 2.9874163 |
| ASMM10P000 | 0.03076 | 0.3664222 | 1.3340673 | up | NM_001081 | Ubr3 | ubiquitin protein ligase E3 component n-recognin 3 | 349.27627 | 319.19095 | 8.714309 | 8.298477 | 397.66647 | 350.781 | 299.38135 | 328.97205 | 300.08533 | 328.51547 | 8.643469 | 8.84679 | 8.652667 | 8.2819195 | 8.119424 | 8.494088 |
| ASMM10P030 | 0.03077 | 0.3664222 | 1.2772205 | up | NM_011341 | Sdf4 | stromal cell derived factor 4 | 14589.736 | 13431.575 | 13.985634 | 13.632627 | 16225.896 | 13677.937 | 13865.376 | 13199.199 | 13109.774 | 13985.753 | 14.050079 | 13.930041 | 13.976783 | 13.566154 | 13.49862 | 13.833106 |
| ASMM10P026 | 0.03081 | 0.3664222 | 1.7027203 | up | NM_007592 | Car8 | carbonic anhydrase 8 | 700.54457 | 496.76229 | 9.667595 | 8.899753 | 866.4377 | 512.2861 | 722.9099 | 433.9758 | 664.11176 | 392.1993 | 9.749269 | 9.382561 | 9.8709545 | 8.688948 | 9.264974 | 8.745338 |
| ASMM10P018 | 0.03085 | 0.3664222 | 1.4992719 | up | NM_146681 | Olfr1424 | olfactory receptor 1424 | 12.012604 | 9.796446 | 3.86476 | 3.280498 | 12.646261 | 11.73888 | 11.652671 | 12.969968 | 9.846472 | 6.5728993 | 3.6709993 | 4.028729 | 3.8945503 | 3.5679526 | 3.172361 | 3.101179 |
| ASMM10P055 | 0.03086 | 0.3664222 | 2.2406698 | up | NM_026100 | Tctex1d1 | Tctex1 domain containing 1 | 12.306185 | 7.120055 | 3.880197 | 2.716267 | 12.73914 | 8.918831 | 15.260583 | 10.794541 | 5.5656238 | 4.9999995 | 3.681833 | 3.6739674 | 4.2847915 | 3.2986822 | 2.483599 | 2.3665206 |
| ASMM10P053 | 0.03091 | 0.3664222 | 1.397543 | up | NM_001048 | Azi2 | 5-azacytidine induced gene 2 | 454.59197 | 391.12489 | 9.075951 | 8.593058 | 510.9083 | 495.43857 | 357.42905 | 390.40652 | 395.67365 | 387.2945 | 8.990204 | 9.33468 | 8.902968 | 8.529851 | 8.522299 | 8.727024 |
| ASMM10P038 | 0.03098 | 0.3664222 | 1.3061045 | up | NM_025515 | Ccdc90b | coiled-coil domain containing 90B | 951.11946 | 873.19036 | 10.126698 | 9.741428 | 1000.9608 | 886.76733 | 965.63025 | 844.736 | 917.58844 | 857.24664 | 9.942591 | 10.142942 | 10.294562 | 9.641509 | 9.735233 | 9.847542 |
| ASMM10P006 | 0.03100 | 0.3664222 | 1.5062616 | up | NM_001252 | Spock3 | sparc/osteonectin, cwcv and kazal-like domains proteoglycan 3 | 387.30048 | 311.97247 | 8.861346 | 8.270374 | 381.25632 | 444.97714 | 335.66797 | 318.56354 | 316.35547 | 300.9984 | 8.58727 | 9.180807 | 8.815961 | 8.237401 | 8.197441 | 8.376279 |
| ASMM10P033 | 0.03104 | 0.3664222 | 1.3227955 | up | NM_001164 | Cops7a | COP9 (constitutive photomorphogenic) homolog, subunit 7a (Arabidopsis thaliana) | 2341.7672 | 2085.951 | 11.384766 | 10.981176 | 2558.7004 | 1955.8453 | 2510.756 | 2103.681 | 2100.907 | 2053.265 | 11.303769 | 11.240716 | 11.609814 | 10.957678 | 10.91201 | 11.073841 |
| ASMM10P006 | 0.03108 | 0.3664222 | 1.2861768 | up | NM_008048 | Igfbp7 | insulin-like growth factor binding protein 7 | 1122.3704 | 1058.1028 | 10.364148 | 10.001059 | 1239.4563 | 1125.9467 | 1001.7081 | 1018.3608 | 1260.4437 | 895.5038 | 10.257055 | 10.489866 | 10.345522 | 9.916358 | 10.177665 | 9.909153 |
| ASMM10P010 | 0.03109 | 0.3664222 | 1.5453483 | up | NM_001166 | Gcnt4 | glucosaminyl (N-acetyl) transferase 4, core 2 (beta-1,6-N-acetylglucosaminyltransferase) | 5864.1318 | 4451.189 | 12.675949 | 12.048017 | 6434.436 | 4514.998 | 6642.9614 | 4494.576 | 5271.8125 | 3587.1785 | 12.680293 | 12.390642 | 12.956912 | 12.040914 | 12.227967 | 11.87517 |
| ASMM10P041 | 0.03111 | 0.3664222 | 1.317196 | up | NM_146223 | Cplx3 | complexin 3 | 585.39008 | 532.6438 | 9.428057 | 9.030587 | 685.47644 | 486.5631 | 584.1307 | 535.1699 | 511.956 | 550.8055 | 9.407043 | 9.308261 | 9.568868 | 8.990204 | 8.889264 | 9.212294 |
| ASMM10P050 | 0.03113 | 0.3664222 | 1.4003195 | up | NM_023409 | Npc2 | Niemann Pick type C2 | 4496.2853 | 3778.9227 | 12.316526 | 11.83077 | 4351.0063 | 4238.187 | 4899.6626 | 3479.227 | 4106.2393 | 3751.3018 | 12.109489 | 12.298875 | 12.541213 | 11.673412 | 11.873802 | 11.945095 |
| ASMM10P007 | 0.03114 | 0.3664222 | 1.5339844 | up | NM_023266 | Zfp120 | zinc finger protein 120 | 187.28969 | 150.19828 | 7.840515 | 7.223231 | 183.10757 | 178.88649 | 199.87502 | 149.58815 | 148.20963 | 152.79707 | 7.5440564 | 7.8999968 | 8.077491 | 7.1383815 | 7.096814 | 7.4344974 |
| ASMM10P032 | 0.03124 | 0.3664222 | 1.7156761 | up | NM_029408 | Iqcd | IQ motif containing D | 19.347132 | 13.58287 | 4.54795 | 3.769173 | 22.07358 | 19.902063 | 16.065752 | 11.34407 | 17.798828 | 11.605711 | 4.5098815 | 4.774268 | 4.359701 | 3.367592 | 4.0521817 | 3.8877451 |
| ASMM10P005 | 0.03126 | 0.3664222 | 1.7834132 | up | NM_001161 | Kif26b | kinesin family member 26B | 1097.1484 | 720.22043 | 10.298955 | 9.464314 | 1155.0842 | 804.1939 | 1332.1672 | 741.55725 | 655.9157 | 763.18835 | 10.14995 | 10.007423 | 10.739491 | 9.462951 | 9.248823 | 9.681167 |
| ASMM10P006 | 0.03126 | 0.3664222 | 2.5344636 | up | NM_009596 | Scgb1b27 | secretoglobin, family 1B, member 27 | 79.850178 | 34.145015 | 6.457388 | 5.115708 | 127.13599 | 39.364388 | 73.050156 | 32.79972 | 33.423244 | 36.21208 | 7.0262337 | 5.744157 | 6.6017747 | 4.9382668 | 4.959784 | 5.4490733 |
| ASMM10P015 | 0.03128 | 0.3664222 | 1.2515226 | up | NM_011062 | Pdpk1 | 3-phosphoinositide dependent protein kinase 1 | 27849.107 | 26333.097 | 14.918044 | 14.594359 | 25877.65 | 28053.13 | 29616.54 | 25987.705 | 27775.002 | 25236.584 | 14.752267 | 14.93854 | 15.063324 | 14.540614 | 14.566196 | 14.676268 |
| ASMM10P027 | 0.03140 | 0.3664222 | 2.9995569 | up | NM_183428 | Epb4.1 | erythrocyte protein band 4.1 | 55.274066 | 19.745795 | 5.849186 | 4.264437 | 101.53971 | 31.476141 | 32.806347 | 19.5695 | 27.761276 | 11.906611 | 6.714274 | 5.4236455 | 5.4096384 | 4.177598 | 4.69083 | 3.9248817 |
| ASMM10P013 | 0.03144 | 0.3666296 | 1.3321095 | up | NM_026746 | Nsmce2 | non-SMC element 2 homolog (MMS21, S. cerevisiae) | 2440.8819 | 2149.7334 | 11.438713 | 11.025001 | 2857.3254 | 1977.2241 | 2488.0962 | 2130.241 | 2105.0593 | 2213.9 | 11.461752 | 11.257958 | 11.59643 | 10.976742 | 10.915356 | 11.182904 |
| ASMM10P007 | 0.03155 | 0.367344 | 1.4964489 | up | NM_019777 | Ikbke | inhibitor of kappaB kinase epsilon | 135.48769 | 113.81416 | 7.38284 | 6.801297 | 135.6784 | 133.62212 | 137.16255 | 126.78635 | 133.3284 | 81.32773 | 7.1174555 | 7.4925876 | 7.5384774 | 6.8991575 | 6.9417715 | 6.5629625 |
| ASMM10P051 | 0.03158 | 0.367344 | 3.1072955 | up | NM_001146 | Gpr52 | G protein-coupled receptor 52 | 30.439245 | 11.127588 | 5.110175 | 3.474515 | 24.332018 | 46.853798 | 20.13192 | 14.5233 | 8.701177 | 10.158288 | 4.649979 | 5.996739 | 4.6838055 | 3.7353652 | 2.994458 | 3.693722 |
| ASMM10P036 | 0.03158 | 0.367344 | 1.728185 | up | NM_172908 | Ovch2 | ovochymase 2 | 38.91402 | 26.52983 | 5.540661 | 4.751403 | 46.87172 | 25.945158 | 43.925182 | 24.58181 | 30.136993 | 24.870687 | 5.614799 | 5.1500616 | 5.857121 | 4.5070305 | 4.812065 | 4.935113 |
| ASMM10P027 | 0.03162 | 0.36737 | 1.2853256 | up | NM_080442 | Tssk3 | testis-specific serine kinase 3 | 52.911388 | 50.943872 | 6.01661 | 5.654476 | 62.132053 | 47.53143 | 49.07068 | 53.192818 | 62.825134 | 36.813663 | 6.018085 | 6.017936 | 6.013809 | 5.6345634 | 5.857121 | 5.471744 |
| ASMM10P006 | 0.03176 | 0.3682963 | 1.4152215 | up | NM_207624 | Ace | angiotensin I converting enzyme (peptidyl-dipeptidase A) 1 | 73.507663 | 63.624887 | 6.496634 | 5.995606 | 78.724686 | 72.2764 | 69.521904 | 57.36839 | 70.22501 | 63.28126 | 6.357736 | 6.6062346 | 6.525932 | 5.7480335 | 6.020343 | 6.2184424 |
| ASMM10P005 | 0.03177 | 0.3682963 | 1.2684934 | up | NM_001163 | Tsr3 | TSR3 20S rRNA accumulation | 238.9909 | 230.233 | 8.182354 | 7.839238 | 276.31744 | 226.2221 | 214.43317 | 227.01547 | 228.11322 | 235.57031 | 8.129051 | 8.230673 | 8.187337 | 7.746028 | 7.729014 | 8.042671 |
| ASMM10P025 | 0.03183 | 0.3682963 | 1.3175355 | up | NM_009250 | Serpini1 | serine (or cysteine) peptidase inhibitor, clade I, member 1 | 6683.9116 | 5963.9524 | 12.885021 | 12.487179 | 7357.1807 | 5810.114 | 6884.44 | 6374.9946 | 5502.825 | 6014.0376 | 12.899056 | 12.752115 | 13.003893 | 12.547247 | 12.293816 | 12.620475 |
| ASMM10P055 | 0.03187 | 0.3682963 | 3.6090098 | up | NM_146849 | Olfr1157 | olfactory receptor 1157 | 27.051231 | 7.664248 | 4.826713 | 2.97511 | 48.401173 | 19.705822 | 13.046699 | 5.646322 | 7.891865 | 9.454556 | 5.6625867 | 4.758767 | 4.058785 | 2.4582467 | 2.863719 | 3.6033638 |
| ASMM10P004 | 0.03187 | 0.3682963 | 1.3198121 | up | NM_001110 | Trim9 | tripartite motif-containing 9 | 925.83279 | 841.18375 | 10.087447 | 9.687114 | 955.2519 | 973.31537 | 848.9311 | 829.0824 | 874.7656 | 819.70325 | 9.882913 | 10.275589 | 10.103839 | 9.6154995 | 9.664721 | 9.781123 |
| ASMM10P009 | 0.03192 | 0.3682963 | 1.2812399 | up | NM_175009 | Eny2 | enhancer of yellow 2 homolog (Drosophila) | 2815.6707 | 2597.0559 | 11.65299 | 11.295449 | 2886.5596 | 2840.662 | 2719.7905 | 2709.6174 | 2498.871 | 2582.6794 | 11.480631 | 11.754215 | 11.724124 | 11.317926 | 11.166616 | 11.401806 |
| ASMM10P044 | 0.03197 | 0.3682963 | 1.3382281 | up | NM_025921 | 2610002M0( | RIKEN cDNA 2610002M06 gene | 20731.048 | 17948.695 | 14.473649 | 14.053325 | 18529.115 | 21616.512 | 22047.516 | 18560.547 | 17837.129 | 17448.41 | 14.249907 | 14.562206 | 14.608833 | 14.069829 | 13.933264 | 14.156881 |
| ASMM10P004 | 0.03207 | 0.3682963 | 1.4108975 | up | NM_001077 | Stag2 | stromal antigen 2 | 786.51833 | 670.09807 | 9.855808 | 9.359195 | 822.0012 | 706.2439 | 831.3099 | 632.9009 | 675.3289 | 702.0644 | 9.674585 | 9.820714 | 10.072125 | 9.23039 | 9.288792 | 9.558402 |
| ASMM10P014 | 0.03212 | 0.3687007 | 1.3169871 | up | NM_028815 | Cep97 | centrosomal protein 97 | 5880.1951 | 5260.4944 | 12.701176 | 12.303935 | 5544.2812 | 5882.3105 | 6213.9937 | 5289.3076 | 5581.222 | 4910.9536 | 12.463475 | 12.772706 | 12.867347 | 12.271359 | 12.310392 | 12.330054 |
| ASMM10P030 | 0.03214 | 0.3687007 | 1.6351544 | up | NM_001163 | Lcorl | ligand dependent nuclear receptor corepressor-like | 754.88285 | 558.0915 | 9.799046 | 9.089619 | 751.5465 | 712.13855 | 800.9635 | 509.89645 | 520.51575 | 643.8623 | 9.5427685 | 9.836445 | 10.017924 | 8.92233 | 8.91493 | 9.431597 |
| ASMM10P013 | 0.03217 | 0.3687545 | 1.4669795 | up | NM_008304 | Sdc2 | syndecan 2 | 1262.8348 | 1024.4597 | 10.521223 | 9.968374 | 1231.4968 | 1457.0165 | 1099.9912 | 1028.5609 | 1125.7339 | 919.0842 | 10.249564 | 10.831943 | 10.482162 | 9.933904 | 10.023241 | 9.947978 |
| ASMM10P041 | 0.03230 | 0.3687704 | 1.715117 | up | NM_008559 | Mc1r | melanocortin 1 receptor | 68.620645 | 48.315783 | 6.386559 | 5.608252 | 59.043705 | 80.517296 | 66.300934 | 52.513615 | 49.929085 | 42.50465 | 5.9416614 | 6.7569723 | 6.4610424 | 5.6155243 | 5.5342507 | 5.67498 |
| ASMM10P005 | 0.03234 | 0.3687704 | 1.4121139 | up | NM_001163 | Pgm3 | phosphoglucomutase 3 | 620.40066 | 529.4791 | 9.518352 | 9.020496 | 629.7596 | 598.13727 | 633.3051 | 537.4474 | 506.67496 | 544.31494 | 9.277493 | 9.59259 | 9.684974 | 8.995434 | 8.872738 | 9.1933155 |
| ASMM10P030 | 0.03242 | 0.3687704 | 1.3814522 | up | NM_001253 | Tet1 | tet methylcytosine dioxygenase 1 | 1946.1086 | 1676.5672 | 11.136467 | 10.670281 | 1861.9092 | 1900.549 | 2075.8677 | 1708.2238 | 1776.1803 | 1545.2975 | 10.861347 | 11.195738 | 11.352316 | 10.656437 | 10.672441 | 10.681966 |
| ASMM10P028 | 0.03265 | 0.3689783 | 1.3606687 | up | NM_001256 | Fsbp | fibrinogen silencer binding protein | 72.45917 | 66.211738 | 6.477228 | 6.032912 | 76.73923 | 70.16005 | 70.47823 | 72.759315 | 78.96548 | 46.91042 | 6.3204064 | 6.5639024 | 6.5473747 | 6.0936236 | 6.1929984 | 5.812114 |
| ASMM10P011 | 0.03268 | 0.3689783 | 1.343735 | up | NM_009351 | Tep1 | telomerase associated protein 1 | 2217.6054 | 1959.302 | 11.313432 | 10.887183 | 2424.7092 | 1876.3379 | 2351.769 | 1798.2905 | 2176.1648 | 1903.4508 | 11.235312 | 11.179852 | 11.525131 | 10.729733 | 10.961595 | 10.970221 |
| ASMM10P044 | 0.03276 | 0.3689783 | 1.3149391 | up | NM_029730 | Mospd2 | motile sperm domain containing 2 | 513.98348 | 475.00318 | 9.260891 | 8.865895 | 523.5369 | 519.9205 | 498.49304 | 483.53537 | 521.14856 | 420.3256 | 9.020767 | 9.400429 | 9.361477 | 8.841785 | 8.916985 | 8.838915 |
| ASMM10P052 | 0.03279 | 0.3689783 | 1.4528686 | up | NM_009256 | Serpinb9 | serine (or cysteine) peptidase inhibitor, clade B, member 9 | 36.433237 | 30.151501 | 5.478004 | 4.9391 | 39.789394 | 36.922348 | 32.58797 | 27.694841 | 33.067833 | 29.691828 | 5.376674 | 5.6566296 | 5.400709 | 4.6896787 | 4.9447737 | 5.1828475 |
| ASMM10P025 | 0.03281 | 0.3689783 | 2.4363982 | up | NM_001243 | Gm8298 | predicted gene 8298 | 26.704828 | 12.827144 | 4.993445 | 3.708695 | 19.55596 | 28.766994 | 31.79153 | 11.89853 | 12.737931 | 13.84497 | 4.319856 | 5.2985077 | 5.361972 | 3.435716 | 3.5526922 | 4.1376777 |
| ASMM10P007 | 0.03297 | 0.3689783 | 1.2828863 | up | NM_025339 | Tmem42 | transmembrane protein 42 | 434.45831 | 407.71972 | 9.010874 | 8.651481 | 533.34125 | 400.69174 | 369.34195 | 441.8432 | 373.11093 | 408.20502 | 9.047137 | 9.03486 | 8.950625 | 8.716481 | 8.440197 | 8.797764 |
| ASMM10P036 | 0.03299 | 0.3689783 | 1.4108859 | up | NM_080638 | Mvp | major vault protein | 8615.5147 | 7101.4144 | 13.23249 | 12.735889 | 8966.227 | 7246.241 | 9634.076 | 6508.9497 | 7652.934 | 7142.3594 | 13.15706 | 13.058079 | 13.482332 | 12.57219 | 12.762763 | 12.872714 |
| ASMM10P004 | 0.03300 | 0.3689783 | 1.290144 | up | NM_001029 | Specc1 | sperm antigen with calponin homology and coiled-coil domains 1 | 115.53555 | 108.80376 | 7.132318 | 6.764786 | 139.76051 | 102.10912 | 104.73702 | 108.1767 | 107.49072 | 110.74386 | 7.1604347 | 7.0989313 | 7.1375875 | 6.6699033 | 6.6335363 | 6.9909177 |
| ASMM10P044 | 0.03303 | 0.3689783 | 1.2822205 | up | NM_026070 | Ccdc53 | coiled-coil domain containing 53 | 1855.6654 | 1727.9042 | 11.071636 | 10.712991 | 2048.4834 | 1717.0599 | 1801.4529 | 1672.1538 | 1684.9484 | 1826.6105 | 10.991956 | 11.062954 | 11.159997 | 10.627097 | 10.597412 | 10.914465 |
| ASMM10P052 | 0.03307 | 0.3689783 | 1.6016039 | up | NM_145513 | Tiprl | TIP41, TOR signalling pathway regulator-like (S. cerevisiae) | 366.96793 | 275.52665 | 8.773301 | 8.093784 | 380.72162 | 289.54614 | 430.63602 | 280.79404 | 269.06158 | 276.72433 | 8.584883 | 8.575686 | 9.159334 | 8.055002 | 7.9620967 | 8.264252 |
| ASMM10P020 | 0.03309 | 0.3689783 | 1.2592139 | up | NM_019667 | Stam2 | signal transducing adaptor molecule (SH3 domain and ITAM motif) 2 | 11412.784 | 10656.773 | 13.629778 | 13.297254 | 10674.958 | 12088.663 | 11474.73 | 10718.543 | 11282.839 | 9968.938 | 13.427832 | 13.76062 | 13.700881 | 13.269149 | 13.290142 | 13.332472 |
| ASMM10P005 | 0.03315 | 0.3689783 | 1.2860391 | up | NM_001164 | Cpsf1 | cleavage and polyadenylation specific factor 1 | 9017.2713 | 8169.2247 | 13.300888 | 12.937954 | 9321.898 | 8419.208 | 9310.708 | 8287.108 | 7850.911 | 8369.655 | 13.214678 | 13.24708 | 13.440908 | 12.913933 | 12.797043 | 13.102886 |
| ASMM10P048 | 0.03320 | 0.3689783 | 1.3254568 | up | NM_001277 | Gas7 | growth arrest specific 7 | 50538.343 | 46322.56 | 15.827598 | 15.421109 | 50564.973 | 47348.297 | 53701.76 | 45771.668 | 46241.47 | 46954.543 | 15.759643 | 15.722624 | 16.000528 | 15.347412 | 15.310144 | 15.60577 |
| ASMM10P008 | 0.03323 | 0.3689783 | 1.4598857 | up | NM_145967 | Vstm2a | V-set and transmembrane domain containing 2A | 296.37902 | 247.73491 | 8.480074 | 7.934219 | 350.1453 | 271.284 | 267.70776 | 228.6478 | 235.04108 | 279.51584 | 8.460215 | 8.487708 | 8.4923 | 7.7565217 | 7.770328 | 8.275807 |
| ASMM10P029 | 0.03330 | 0.3689783 | 1.3435251 | up | NM_001161 | Gm13272 | predicted gene 13272 | 103.67773 | 95.264963 | 6.996005 | 6.569982 | 104.8195 | 95.15106 | 111.06261 | 103.08569 | 100.715 | 81.9942 | 6.7645054 | 7.000031 | 7.223478 | 6.597871 | 6.537575 | 6.5744987 |
| ASMM10P006 | 0.03331 | 0.3689783 | 1.6250883 | up | NM_009828 | Ccna2 | cyclin A2 | 10.139962 | 7.43 | 3.615336 | 2.914818 | 11.855489 | 7.545596 | 11.0188 | 6.163247 | 9.892415 | 6.234337 | 3.5725958 | 3.460714 | 3.8126993 | 2.537105 | 3.1783915 | 3.0289583 |
| ASMM10P005 | 0.03344 | 0.3689783 | 1.3468159 | up | NM_021386 | Cldn10 | claudin 10 | 551.03497 | 494.13262 | 9.352042 | 8.92249 | 565.627 | 528.23334 | 559.24457 | 464.25854 | 541.5425 | 476.59683 | 9.126165 | 9.421631 | 9.508331 | 8.785291 | 8.971791 | 9.010387 |
| ASMM10P029 | 0.03345 | 0.3689783 | 1.9820463 | up | NM_001122 | Lepr | leptin receptor | 558.19829 | 347.91879 | 9.37154 | 8.38455 | 571.47986 | 555.9005 | 547.2145 | 272.32898 | 314.98785 | 456.43954 | 9.140871 | 9.495388 | 9.478362 | 8.012028 | 8.190732 | 8.950889 |
| ASMM10P053 | 0.03348 | 0.3689783 | 1.2803968 | up | NM_001161 | Cbfb | core binding factor beta | 74.017883 | 70.915315 | 6.508729 | 6.152138 | 76.07241 | 68.8851 | 77.09614 | 73.202545 | 78.33906 | 61.20434 | 6.307206 | 6.5364885 | 6.682492 | 6.101962 | 6.1790824 | 6.1753693 |
| ASMM10P054 | 0.03348 | 0.3689783 | 1.3642799 | up | ENSMUST( | Cartpt | CART prepropeptide | 650.0992 | 569.88109 | 9.573368 | 9.125229 | 776.2215 | 651.4873 | 522.5888 | 510.7702 | 606.1501 | 592.72296 | 9.592365 | 9.707783 | 9.419957 | 8.924114 | 9.135224 | 9.316348 |
| ASMM10P005 | 0.03348 | 0.3689783 | 1.9416553 | up | NM_001172 | Rbms3 | RNA binding motif, single stranded interacting protein | 47.430589 | 30.027575 | 5.863302 | 4.906015 | 53.166046 | 44.782413 | 44.343307 | 28.23304 | 24.52142 | 37.328266 | 5.793136 | 5.9273534 | 5.8694158 | 4.721795 | 4.5065928 | 5.489656 |
| ASMM10P023 | 0.03350 | 0.3689783 | 1.4941814 | up | NM_011931 | Rfwd2 | ring finger and WD repeat domain 2 | 6258.9798 | 4923.1033 | 12.78079 | 12.201435 | 5613.4087 | 5887.7803 | 7275.7505 | 4821.772 | 5584.6396 | 4362.8984 | 12.483136 | 12.774827 | 13.084407 | 12.137953 | 12.311102 | 12.155249 |
| ASMM10P011 | 0.03350 | 0.3689783 | 1.5687556 | up | NM_146051 | 3830406C13 | RIKEN cDNA 3830406C13 gene | 308.96006 | 244.01443 | 8.543925 | 7.894304 | 319.30838 | 267.7674 | 339.8044 | 287.42975 | 269.04202 | 175.57152 | 8.329054 | 8.470525 | 8.832196 | 8.093684 | 7.961878 | 7.6273513 |
| ASMM10P000 | 0.03352 | 0.3689783 | 1.5968883 | up | NM_010653 | Klrc2 | killer cell lectin-like receptor subfamily C, member 2 | 83.270741 | 63.707189 | 6.675596 | 6.000332 | 73.667564 | 83.09553 | 93.04913 | 67.70968 | 68.372375 | 55.039513 | 6.2613497 | 6.8058143 | 6.9596233 | 5.9892244 | 5.9827733 | 6.0289993 |
| ASMM10P000 | 0.03358 | 0.3689783 | 1.535621 | up | NM_133885 | Osbpl9 | oxysterol binding protein-like 9 | 103.60318 | 81.564853 | 6.970278 | 6.351456 | 121.81406 | 80.19998 | 108.79548 | 80.329765 | 78.181145 | 86.18365 | 6.9697475 | 6.750921 | 7.1901655 | 6.2335815 | 6.175783 | 6.645003 |
| ASMM10P053 | 0.03359 | 0.3689783 | 1.4340244 | up | NM_178709 | Rnf214 | ring finger protein 214 | 263.68902 | 224.61606 | 8.319805 | 7.799735 | 292.71304 | 246.55048 | 251.80354 | 200.60687 | 230.6761 | 242.5652 | 8.204498 | 8.351727 | 8.403189 | 7.568184 | 7.7437983 | 8.087223 |
| ASMM10P039 | 0.03361 | 0.3689783 | 1.3029749 | up | NM_025436 | Sc4mol | sterol-C4-methyl oxidase-like | 21478.165 | 19324.132 | 14.537381 | 14.155572 | 21068.898 | 20948.521 | 22417.076 | 17274.46 | 21274.88 | 19423.055 | 14.452809 | 14.522394 | 14.63694 | 13.951581 | 14.198804 | 14.31633 |
| ASMM10P023 | 0.03364 | 0.3689783 | 2.3417996 | up | NM_001142 | Lmo2 | LIM domain only 2 | 11.913191 | 5.458315 | 3.79972 | 2.572102 | 13.147982 | 14.719231 | 7.8723598 | 4.9999995 | 4.9999995 | 6.3749447 | 3.7333345 | 4.3454237 | 3.3204021 | 2.3250442 | 2.3265402 | 3.064723 |
| ASMM10P047 | 0.03369 | 0.3689783 | 1.5513584 | up | NM_010880 | Ncl | nucleolin | 805.14544 | 625.50368 | 9.89183 | 9.258298 | 822.25885 | 720.6312 | 872.54626 | 576.96246 | 603.52924 | 696.01935 | 9.676295 | 9.853567 | 10.145627 | 9.097994 | 9.128721 | 9.548178 |
| ASMM10P040 | 0.03374 | 0.3689783 | 1.3743881 | up | NM_148952 | E2f4 | E2F transcription factor 4 | 50.944438 | 44.277475 | 5.94415 | 5.485361 | 62.0409 | 50.761784 | 40.03063 | 47.586884 | 44.05138 | 41.19416 | 6.015704 | 6.10498 | 5.711766 | 5.478858 | 5.3466086 | 5.630615 |
| ASMM10P045 | 0.03377 | 0.3689783 | 2.6793653 | up | NM_146261 | Fam199x | family with sequence similarity 199, X-linked | 31.867002 | 14.160726 | 5.231212 | 3.809321 | 39.976723 | 19.858498 | 35.765785 | 10.930661 | 12.70873 | 18.842787 | 5.3830695 | 4.7705345 | 5.5400333 | 3.3174734 | 3.549539 | 4.560951 |
| ASMM10P017 | 0.03377 | 0.3689783 | 1.6435443 | up | NM_020603 | Wdr46 | WD repeat domain 46 | 8346.6356 | 5884.5529 | 13.176538 | 12.459728 | 7442.1494 | 7076.5093 | 10521.248 | 6703.927 | 5930.5327 | 5019.199 | 12.910901 | 13.025228 | 13.593486 | 12.619401 | 12.398792 | 12.360991 |
| ASMM10P046 | 0.03377 | 0.3689783 | 1.3091617 | up | NM_172439 | Inpp5j | inositol polyphosphate 5-phosphatase J | 2429.6854 | 2205.0187 | 11.450432 | 11.061789 | 2412.3335 | 2361.493 | 2515.2297 | 2227.514 | 2234.0022 | 2153.54 | 11.227402 | 11.51169 | 11.612204 | 11.040554 | 11.002795 | 11.142017 |
| ASMM10P047 | 0.03382 | 0.3689783 | 1.4385322 | up | NM_029658 | Fam101b | family with sequence similarity 101, member B | 4473.2335 | 3666.3159 | 12.315912 | 11.791314 | 4486.2856 | 4233.5195 | 4699.8955 | 3290.5032 | 3675.4136 | 4033.0308 | 12.160128 | 12.297786 | 12.489821 | 11.597448 | 11.725506 | 12.050988 |
| ASMM10P039 | 0.03383 | 0.3689783 | 1.7366165 | up | NM_027998 | Cldn23 | claudin 23 | 282.56619 | 200.4546 | 8.416347 | 7.620067 | 307.82272 | 246.2879 | 293.58795 | 209.61421 | 161.49805 | 230.25153 | 8.27603 | 8.349663 | 8.623347 | 7.6328883 | 7.219247 | 8.008067 |
| ASMM10P009 | 0.03389 | 0.3689783 | 1.3089092 | up | NM_178650 | Tbc1d10c | TBC1 domain family, member 10c | 27.422862 | 24.854965 | 5.051135 | 4.66277 | 33.864456 | 23.250507 | 25.153622 | 24.973223 | 25.540123 | 24.05155 | 5.1404743 | 5.0017586 | 5.0111723 | 4.5314097 | 4.566889 | 4.8900113 |
| ASMM10P002 | 0.03389 | 0.3689783 | 1.5830103 | up | NM_001167 | Zfp568 | zinc finger protein 568 | 401.04455 | 302.4039 | 8.889592 | 8.226921 | 424.6623 | 304.23846 | 474.2329 | 312.61545 | 307.82904 | 286.7672 | 8.731653 | 8.642904 | 9.294218 | 8.211415 | 8.155694 | 8.313654 |
| ASMM10P033 | 0.03397 | 0.3689783 | 1.5172064 | up | NM_023042 | Recql | RecQ protein-like | 2812.9625 | 2177.5285 | 11.64572 | 11.044302 | 2631.4648 | 2594.4744 | 3212.9482 | 2129.9297 | 2209.511 | 2193.1448 | 11.34164 | 11.632633 | 11.962886 | 10.97647 | 10.987882 | 11.168555 |
| ASMM10P008 | 0.03398 | 0.3689783 | 1.298473 | up | NM_030108 | Tmem33 | transmembrane protein 33 | 1651.5164 | 1526.8204 | 10.910923 | 10.534107 | 1698.0519 | 1750.1675 | 1506.3297 | 1536.0664 | 1542.6276 | 1501.7671 | 10.721364 | 11.090565 | 10.920839 | 10.500891 | 10.464063 | 10.637366 |
| ASMM10P008 | 0.03407 | 0.3689783 | 1.2554412 | up | NM_026417 | Yipf4 | Yip1 domain family, member 4 | 205.49255 | 199.38943 | 7.960348 | 7.632153 | 249.30502 | 185.25578 | 181.91685 | 200.31667 | 194.04929 | 203.80232 | 7.986466 | 7.95008 | 7.944497 | 7.5666757 | 7.495905 | 7.833879 |
| ASMM10P001 | 0.03410 | 0.3689783 | 1.3069907 | up | NM_018854 | Ift20 | intraflagellar transport 20 | 1226.4812 | 1122.4465 | 10.486372 | 10.100123 | 1315.7046 | 1074.8961 | 1288.8428 | 1120.5376 | 1127.0757 | 1119.7261 | 10.347223 | 10.417377 | 10.694516 | 10.054884 | 10.025318 | 10.220167 |
| ASMM10P007 | 0.03420 | 0.3689783 | 1.3185028 | up | NM_018864 | Impa1 | inositol (myo)-1(or 4)-monophosphatase 1 | 2724.9375 | 2443.2446 | 11.609623 | 11.210722 | 2740.61 | 2542.8743 | 2891.3281 | 2451.722 | 2492.2031 | 2385.8086 | 11.404153 | 11.605924 | 11.818791 | 11.178747 | 11.162988 | 11.290431 |
| ASMM10P033 | 0.03423 | 0.3689783 | 1.3644914 | up | NM_028832 | Mterfd3 | MTERF domain containing 3 | 7416.3707 | 6351.0676 | 13.02931 | 12.580946 | 7035.7056 | 6927.0215 | 8286.385 | 6351.418 | 6382.5083 | 6319.2764 | 12.817522 | 13.002055 | 13.268352 | 12.542833 | 12.510625 | 12.689381 |
| ASMM10P010 | 0.03425 | 0.3689783 | 1.468433 | up | NM_030711 | Erap1 | endoplasmic reticulum aminopeptidase 1 | 359.87986 | 301.04071 | 8.762541 | 8.208264 | 402.11502 | 339.74872 | 337.77585 | 257.08942 | 314.09732 | 331.9354 | 8.659678 | 8.804016 | 8.82393 | 7.9296336 | 8.185829 | 8.509329 |

| | | | | | | | | | | | | | | | | | | | | | | | |
|---|---|---|---|---|---|---|---|---|---|---|---|---|---|---|---|---|---|---|---|---|---|---|---|
| ASMM10P042 | 0.03427 | 0.3689783 | 1.3868494 | up | NM_001033 | Pls1 | plastin 1 (I-isoform) | 7019.6383 | 5944.041 | 12.94692 | 12.475109 | 6724.8184 | 6327.7866 | 8006.31 | 5828.4663 | 6666.7563 | 5336.9004 | 12.746779 | 12.873499 | 13.220484 | 12.415621 | 12.569589 | 12.440118 |
| ASMM10P048 | 0.03431 | 0.3689783 | 1.4064355 | up | NM_029348 | Zbtb4 | zinc finger and BTB domain containing 4 | 264.09657 | 228.32829 | 8.302242 | 7.810199 | 335.7676 | 247.15842 | 209.3637 | 198.26172 | 278.1243 | 208.59886 | 8.401985 | 8.355689 | 8.149053 | 7.5522857 | 8.009024 | 7.869287 |
| ASMM10P036 | 0.03443 | 0.3692783 | 1.4174896 | up | NM_027420 | 2610034B18 | RIKEN cDNA 2610034B18 gene | 297.45091 | 255.462 | 8.48994 | 7.986602 | 317.07736 | 256.0472 | 319.22818 | 253.84532 | 253.58377 | 258.9569 | 8.31953 | 8.406596 | 8.743694 | 7.9097996 | 7.87715 | 8.172856 |
| ASMM10P009 | 0.03447 | 0.3692783 | 1.3291281 | up | NM_176839 | Abcc5 | ATP-binding cassette, sub-family C (CFTR/MRP), member 5 | 1009.326 | 920.12143 | 10.204118 | 9.793638 | 1175.3273 | 917.4811 | 935.1697 | 1087.1068 | 966.6751 | 706.5824 | 10.173871 | 10.191632 | 10.246851 | 10.011703 | 9.8020115 | 9.567199 |
| ASMM10P028 | 0.03449 | 0.3692783 | 1.4106445 | up | NM_016658 | Galt | galactose-1-phosphate uridyl transferase | 2663.7277 | 2232.3218 | 11.571771 | 11.075417 | 2679.875 | 2320.7463 | 2990.5618 | 2271.2654 | 2506.6501 | 1919.0499 | 11.369677 | 11.485991 | 11.859646 | 11.070169 | 11.172329 | 10.983753 |
| ASMM10P018 | 0.03456 | 0.369388 | 1.2560127 | up | NM_001146 | Wac | WW domain containing adaptor with coiled-coil | 2674.1118 | 2521.5881 | 11.57821 | 11.249359 | 3047.8857 | 2186.1582 | 2788.2915 | 2671.2034 | 2584.0474 | 2309.5134 | 11.560284 | 11.411115 | 11.76323 | 11.292009 | 11.217058 | 11.239009 |
| ASMM10P005 | 0.03459 | 0.369388 | 1.2854502 | up | NM_001164 | Psmd10 | proteasome (prosome, macropain) 26S subunit, non-ATPase, 10 | 466.79268 | 440.29159 | 9.123154 | 8.76088 | 521.0773 | 454.6887 | 424.61203 | 460.91083 | 412.0267 | 447.93723 | 9.015039 | 9.212754 | 9.141669 | 8.775877 | 8.581508 | 8.925256 |
| ASMM10P027 | 0.03467 | 0.369388 | 1.3716851 | up | NM_026039 | Med18 | mediator complex subunit 18 | 2456.878 | 2126.1156 | 11.463039 | 11.00709 | 2371.517 | 2402.9373 | 2596.1797 | 2039.0736 | 2298.7192 | 2040.5541 | 11.199675 | 11.52899 | 11.660452 | 10.913891 | 11.04027 | 11.067108 |
| ASMM10P026 | 0.03472 | 0.3694719 | 1.4880624 | up | NM_025995 | Fbxo5 | F-box protein 5 | 449.48281 | 363.89129 | 9.055258 | 8.481823 | 486.01312 | 351.74716 | 510.68814 | 372.41803 | 418.02353 | 301.2323 | 8.922547 | 8.849768 | 9.393459 | 8.466266 | 8.60153 | 8.377673 |
| ASMM10P044 | 0.03478 | 0.3694719 | 1.7847805 | up | NM_001001 | Gja6 | gap junction protein, alpha 6 | 213.4298 | 142.53368 | 7.983329 | 7.147583 | 235.12958 | 143.55 | 261.60983 | 153.48799 | 139.76915 | 134.3439 | 7.901124 | 7.5874968 | 8.461367 | 7.175707 | 7.008266 | 7.2587748 |
| ASMM10P031 | 0.03483 | 0.3694719 | 1.5062739 | up | NM_026735 | Mob1b | MOB kinase activator 1B | 107.0606 | 87.10223 | 7.028685 | 6.437701 | 124.67089 | 100.27911 | 96.231804 | 91.292595 | 75.579994 | 94.4341 | 6.9985557 | 7.077231 | 7.010269 | 6.4130473 | 6.1275835 | 6.7724724 |
| ASMM10P043 | 0.03489 | 0.3694719 | 1.4502132 | up | NM_177275 | Amigo3 | adhesion molecule with Ig like domain 3 | 1012.0178 | 837.37195 | 10.214925 | 9.67866 | 994.2956 | 942.97943 | 1098.7783 | 889.3456 | 810.10834 | 812.6619 | 9.932944 | 10.231535 | 10.480295 | 9.714099 | 9.55357 | 9.76831 |
| ASMM10P050 | 0.03489 | 0.3694719 | 1.6835246 | up | NM_020287 | Insm2 | insulinoma-associated 2 | 154.45595 | 113.46686 | 7.565229 | 6.813744 | 159.3213 | 135.272 | 168.77454 | 125.74657 | 94.5875 | 120.06651 | 7.3467436 | 7.5107546 | 7.8381886 | 6.886116 | 6.452829 | 7.1022873 |
| ASMM10P042 | 0.03490 | 0.3694719 | 1.3046321 | up | NM_023431 | Mum1 | melanoma associated antigen (mutated) 1 | 1533.4774 | 1395.8556 | 10.783827 | 10.400184 | 1972.737 | 1291.6582 | 1336.037 | 1358.0599 | 1374.0922 | 1455.4147 | 10.938724 | 10.669754 | 10.743003 | 10.322771 | 10.293989 | 10.583792 |
| ASMM10P036 | 0.03493 | 0.3694719 | 1.7702787 | up | NM_031252 | Il23a | interleukin 23, alpha subunit p19 | 92.89885 | 63.560506 | 6.822531 | 5.998554 | 83.7658 | 112.08162 | 82.84913 | 63.949337 | 62.13797 | 64.59421 | 6.441814 | 7.2373734 | 6.7884045 | 5.904813 | 5.8424435 | 6.248406 |
| ASMM10P044 | 0.03498 | 0.3694719 | 1.3340567 | up | NM_012010 | Eif2s3x | eukaryotic translation initiation factor 2, subunit 3, structural gene X-linked | 8949.8271 | 7906.8093 | 13.289538 | 12.873718 | 10075.114 | 7345.0034 | 9429.364 | 8238.502 | 8908.606 | 6573.32 | 13.34109 | 13.071597 | 13.455926 | 12.903851 | 12.973291 | 12.744011 |
| ASMM10P045 | 0.03499 | 0.3694719 | 1.392149 | up | NM_001034 | Ccdc160 | coiled-coil domain containing 160 | 1360.2962 | 1169.2277 | 10.634449 | 10.157136 | 1363.1714 | 1280.0592 | 1437.658 | 1175.7283 | 1133.7388 | 1198.216 | 10.402779 | 10.655563 | 10.845006 | 10.124687 | 10.033755 | 10.312965 |
| ASMM10P007 | 0.03508 | 0.3694719 | 1.6287223 | up | NM_021325 | Cd200r1 | CD200 receptor 1 | 56.080343 | 40.830041 | 6.060585 | 5.356845 | 74.62932 | 38.865223 | 54.746487 | 35.753765 | 49.49245 | 37.243908 | 6.2786264 | 5.725268 | 6.177862 | 5.06429 | 5.519974 | 5.4862704 |
| ASMM10P008 | 0.03511 | 0.3694719 | 2.8075059 | up | NM_027613 | Mmrn1 | multimerin 1 | 41.060745 | 16.830009 | 5.555022 | 4.065733 | 38.00891 | 60.032463 | 25.140863 | 22.743109 | 12.94969 | 14.797227 | 5.3094025 | 6.345084 | 5.01058 | 4.394448 | 3.5746086 | 4.2281427 |
| ASMM10P042 | 0.03513 | 0.3694719 | 1.3711548 | up | NM_024266 | Rps25 | ribosomal protein S25 | 2268.2546 | 1966.9025 | 11.352247 | 10.896856 | 2170.678 | 2381.0488 | 2253.0369 | 2004.5223 | 2016.586 | 1879.5991 | 11.068865 | 11.519593 | 11.468283 | 10.884959 | 10.853399 | 10.952209 |
| ASMM10P040 | 0.03513 | 0.3694719 | 1.5587689 | up | NM_173189 | Mcph1 | microcephaly, primary autosomal recessive 1 | 129.66972 | 102.36752 | 7.310248 | 6.669841 | 126.4437 | 112.25527 | 150.3102 | 118.15413 | 101.65909 | 87.28933 | 7.0188828 | 7.238887 | 7.6729746 | 6.7941246 | 6.552059 | 6.6633396 |
| ASMM10P041 | 0.03514 | 0.3694719 | 1.6605467 | up | NM_146745 | Olfr957 | olfactory receptor 957 | 89.973908 | 64.341883 | 6.740524 | 6.008866 | 117.73561 | 61.027695 | 91.15842 | 74.8075 | 57.4175 | 60.80065 | 6.92437 | 6.367465 | 6.9297376 | 6.1304975 | 5.728288 | 6.1678123 |
| ASMM10P016 | 0.03516 | 0.3694719 | 1.3465191 | up | NM_198006 | Coa5 | cytochrome C oxidase assembly factor 5 | 22198.394 | 19221.995 | 14.584881 | 14.155646 | 20597.738 | 20604.396 | 25393.047 | 19061.465 | 20102.453 | 18502.066 | 14.413924 | 14.502316 | 14.838402 | 14.103208 | 14.117288 | 14.246442 |
| ASMM10P032 | 0.03517 | 0.3694719 | 1.3476377 | up | NM_007475 | Rplp0 | ribosomal protein, large, P0 | 1372.35 | 1224.9466 | 10.649352 | 10.218919 | 1414.4396 | 1326.4994 | 1376.111 | 1162.9349 | 1229.7007 | 1282.2043 | 10.451896 | 10.711925 | 10.784234 | 10.108349 | 10.143806 | 10.404602 |
| ASMM10P000 | 0.03525 | 0.3694719 | 1.4361821 | up | NM_025675 | Dph6 | diphthamine biosynthesis 6 | 409.72471 | 348.90506 | 8.948634 | 8.426395 | 419.63486 | 419.49646 | 390.0428 | 353.73404 | 326.64648 | 366.33466 | 8.717415 | 9.099842 | 9.028645 | 8.387089 | 8.243326 | 8.648771 |
| ASMM10P035 | 0.03534 | 0.3694719 | 1.4135064 | up | NM_001105 | Vmn2r51 | vomeronasal 2, receptor 51 | 83.813953 | 71.786493 | 6.666681 | 6.167402 | 103.54108 | 69.162926 | 78.737854 | 76.33624 | 65.42706 | 73.59618 | 6.744987 | 6.543494 | 6.7115607 | 6.1573124 | 5.917848 | 6.427046 |
| ASMM10P025 | 0.03547 | 0.3694719 | 1.4184539 | up | NM_175515 | Intu | inturned planar cell polarity effector homolog (Drosophila) | 45.843588 | 39.353031 | 5.819971 | 5.315652 | 47.24265 | 44.549515 | 45.7386 | 37.171047 | 41.706165 | 39.18188 | 5.625842 | 5.9204907 | 5.9135814 | 5.1186504 | 5.269521 | 5.558785 |
| ASMM10P054 | 0.03548 | 0.3694719 | 1.2869452 | up | NM_011190 | Psme2 | proteasome (prosome, macropain) activator subunit 2 (PA28 beta) | 249.2394 | 239.92243 | 8.24601 | 7.882059 | 262.82108 | 241.16904 | 243.72807 | 250.05798 | 276.2177 | 193.4916 | 8.057965 | 8.31953 | 8.360535 | 7.8859024 | 7.9977617 | 7.762514 |
| ASMM10P009 | 0.03553 | 0.3694719 | 1.4495157 | up | NM_212484 | Cnot6 | CCR4-NOT transcription complex, subunit 6 | 687.14611 | 570.14 | 9.662531 | 9.12696 | 659.89545 | 668.15497 | 733.3879 | 542.5906 | 627.0678 | 540.7616 | 9.350583 | 9.74436 | 9.892649 | 9.011945 | 9.184467 | 9.184467 |
| ASMM10P052 | 0.03553 | 0.3694719 | 1.2675838 | up | NM_019551 | Tdp2 | tyrosyl-DNA phosphodiesterase 2 | 453.67914 | 434.40519 | 9.085697 | 8.743615 | 490.48477 | 404.37946 | 466.1732 | 463.07822 | 432.44174 | 407.6956 | 8.934866 | 9.04779 | 9.274434 | 8.782135 | 8.6532955 | 8.795416 |
| ASMM10P007 | 0.03557 | 0.3694719 | 1.3526365 | up | NM_013787 | Skp2 | S-phase kinase-associated protein 2 (p45) | 49.904254 | 44.900783 | 5.939475 | 5.5037 | 49.15161 | 52.47959 | 48.081562 | 47.547424 | 47.89686 | 39.258064 | 5.6840496 | 6.151739 | 5.982635 | 5.477604 | 5.472216 | 5.561281 |
| ASMM10P039 | 0.03558 | 0.3694719 | 2.2291834 | up | NM_011925 | Cd97 | CD97 antigen | 49.235646 | 25.247801 | 5.835834 | 4.679318 | 59.798134 | 27.17423 | 60.734573 | 22.3132 | 28.230324 | 25.19988 | 5.9621367 | 5.213139 | 6.3322253 | 4.365945 | 4.718102 | 4.953908 |
| ASMM10P043 | 0.03558 | 0.3694719 | 1.5476172 | up | NM_177352 | Gk5 | glycerol kinase 5 (putative) | 391.17268 | 306.06327 | 8.873446 | 8.243397 | 370.6823 | 348.84323 | 453.99252 | 312.37457 | 323.36697 | 282.44827 | 8.544504 | 8.839124 | 9.23671 | 8.210069 | 8.229027 | 8.291096 |
| ASMM10P032 | 0.03561 | 0.3694719 | 1.584765 | up | NM_148951 | Gipc3 | GIPC PDZ domain containing family, member 3 | 97.106828 | 75.120987 | 6.894655 | 6.230386 | 93.88431 | 109.52048 | 87.915695 | 85.75048 | 68.40371 | 71.20877 | 6.606015 | 7.2035975 | 6.8743525 | 6.3246956 | 5.983394 | 6.3830686 |
| ASMM10P038 | 0.03562 | 0.3694719 | 3.7142245 | up | NM_007424 | Acan | aggrecan | 48.540665 | 13.877995 | 5.704078 | 3.811017 | 55.881187 | 69.32819 | 20.412619 | 13.814891 | 11.58409 | 16.235003 | 5.8604436 | 6.547268 | 4.704523 | 3.660508 | 3.4149501 | 4.3575935 |
| ASMM10P050 | 0.03565 | 0.3694719 | 1.3363399 | up | NM_027828 | Fam110c | family with sequence similarity 110, member C | 493.90344 | 445.30346 | 9.183051 | 8.764764 | 624.50305 | 444.2453 | 412.96198 | 387.14935 | 530.2849 | 418.47614 | 9.263586 | 9.178523 | 9.107044 | 8.519802 | 8.941089 | 8.833401 |
| ASMM10P055 | 0.03568 | 0.3694719 | 2.2778835 | up | NM_010474 | Hs3st1 | heparan sulfate (glucosamine) 3-O-sulfotransferase 1 | 165.77911 | 87.183112 | 7.631612 | 6.443918 | 114.43118 | 176.72964 | 206.1765 | 96.570305 | 88.98287 | 75.99616 | 6.88687 | 7.8824883 | 8.125477 | 6.4990115 | 6.362057 | 6.470685 |
| ASMM10P012 | 0.03580 | 0.3694719 | 1.259481 | up | NM_025412 | Pycrl | pyrroline-5-carboxylate reductase-like | 17633.985 | 16387.434 | 14.254129 | 13.921299 | 16919.732 | 18368.7 | 17613.523 | 16362.797 | 16374.516 | 16424.99 | 14.109428 | 14.344764 | 14.308194 | 13.87635 | 13.814317 | 14.073231 |
| ASMM10P009 | 0.03581 | 0.3694719 | 2.9107686 | up | NM_199465 | Nexn | nexilin | 844.27905 | 315.24108 | 9.819791 | 8.278391 | 590.99835 | 1385.2139 | 556.6249 | 283.38803 | 366.58707 | 295.74814 | 9.185896 | 10.770463 | 9.5030155 | 8.071115 | 8.411619 | 8.35244 |
| ASMM10P043 | 0.03585 | 0.3694719 | 1.3019879 | up | NM_172928 | Dclk3 | doublecortin-like kinase 3 | 1184.7378 | 1082.2082 | 10.430411 | 10.049695 | 1331.6478 | 1256.528 | 966.0376 | 1148.2751 | 1040.7466 | 1057.6028 | 10.366735 | 10.628497 | 10.296002 | 10.08831 | 9.915747 | 10.145029 |
| ASMM10P000 | 0.03593 | 0.3694719 | 1.2541004 | up | NM_001252 | Cadps2 | Ca2+-dependent activator protein for secretion 2 | 356.94127 | 346.64424 | 8.742886 | 8.416233 | 409.1078 | 306.1174 | 355.5986 | 366.11096 | 336.40967 | 337.4121 | 8.681458 | 8.651171 | 8.89603 | 8.436944 | 8.2834425 | 8.528314 |
| ASMM10P036 | 0.03593 | 0.3694719 | 1.4933876 | up | NM_011746 | Mkrn3 | makorin, ring finger protein, 3 | 611.57562 | 493.68696 | 9.495526 | 8.916938 | 578.0604 | 598.9737 | 657.69275 | 477.88205 | 567.18536 | 435.99347 | 9.155281 | 9.59517 | 9.736128 | 8.82452 | 9.038963 | 8.88733 |
| ASMM10P006 | 0.03603 | 0.3694719 | 1.2808309 | up | NM_008497 | Lhb | luteinizing hormone beta | 1620.7803 | 1525.8075 | 10.886795 | 10.529715 | 1647.3063 | 1672.4324 | 1542.6022 | 1587.0696 | 1537.7102 | 1452.6427 | 10.673313 | 11.034396 | 10.952676 | 10.54731 | 10.461082 | 10.580753 |
| ASMM10P011 | 0.03611 | 0.3694719 | 1.9775588 | up | NM_001033 | 4922505E12 | RIKEN cDNA 4922505E12 gene | 26.395581 | 16.106199 | 5.006563 | 4.022842 | 26.676304 | 28.1536 | 24.35684 | 15.23011 | 13.88603 | 19.202456 | 4.7877703 | 5.266945 | 4.964973 | 3.805037 | 3.6765585 | 4.586931 |
| ASMM10P011 | 0.03612 | 0.3694719 | 1.4131305 | up | NM_146106 | Lyplal1 | lysophospholipase-like 1 | 3056.8076 | 2567.0903 | 11.771013 | 11.272118 | 3246.986 | 2900.6797 | 3022.757 | 2443.275 | 2361.051 | 2896.945 | 11.655701 | 11.780166 | 11.877171 | 11.171653 | 11.082241 | 11.56246 |
| ASMM10P049 | 0.03618 | 0.3694719 | 1.3260448 | up | NM_029788 | Rnft1 | ring finger protein, transmembrane 1 | 1515.5397 | 1376.2601 | 10.786661 | 10.379532 | 1535.8623 | 1422.8665 | 1587.8903 | 1335.6873 | 1549.9893 | 1243.1036 | 10.567496 | 10.804605 | 10.987882 | 10.301291 | 10.470569 | 10.366735 |
| ASMM10P021 | 0.03619 | 0.3694719 | 1.6478684 | up | NM_175093 | Trib3 | tribbles homolog 3 (Drosophila) | 987.02327 | 711.45566 | 10.15511 | 9.434509 | 1026.608 | 749.9723 | 1184.4895 | 724.61743 | 837.885 | 571.86456 | 9.97974 | 9.909153 | 10.576437 | 9.432239 | 9.601804 | 9.269484 |
| ASMM10P018 | 0.03624 | 0.3694719 | 1.3200156 | up | NM_010316 | Gng3 | guanine nucleotide binding protein (G protein), gamma 3 | 5680.4787 | 5039.3449 | 12.644569 | 12.244014 | 6084.588 | 4796.677 | 6160.171 | 5030.772 | 4986.252 | 5101.0107 | 12.600172 | 12.480977 | 12.852557 | 12.197024 | 12.155919 | 12.379098 |
| ASMM10P004 | 0.03631 | 0.3696581 | 2.0528931 | up | NM_001033 | Lsmem1 | leucine-rich single-pass membrane protein 1 | 13.674554 | 7.822117 | 4.042574 | 3.004916 | 12.64035 | 12.35583 | 16.027481 | 10.22165 | 5.501031 | 7.7436714 | 3.6700187 | 4.1006923 | 4.3570123 | 3.2151742 | 2.47202 | 3.3275535 |
| ASMM10P013 | 0.03634 | 0.3697467 | 1.7355438 | up | NM_001024 | C1ql4 | complement component 1, q subcomponent-like 4 | 63.607033 | 41.577371 | 6.182351 | 5.386964 | 99.56759 | 43.06722 | 48.18629 | 44.102036 | 47.179966 | 33.45011 | 6.688128 | 5.87325 | 5.9856763 | 5.368286 | 5.447798 | 5.344807 |
| ASMM10P021 | 0.03637 | 0.3698236 | 1.7492433 | up | NM_027759 | Fsip1 | fibrous sheath-interacting protein 1 | 188.80199 | 134.74303 | 7.853412 | 7.046681 | 194.14276 | 184.00572 | 188.25748 | 109.6968 | 135.11928 | 159.41301 | 7.6310205 | 7.939831 | 7.9893837 | 6.689775 | 6.9596233 | 7.490644 |
| ASMM10P008 | 0.03643 | 0.3698934 | 1.6651519 | up | NM_028326 | Zfp618 | zinc finger protein 618 | 73.1854 | 53.5298 | 6.483696 | 5.748043 | 76.87967 | 79.90003 | 62.7765 | 52.38826 | 48.649494 | 59.551647 | 6.323764 | 6.7461176 | 6.3812075 | 5.611844 | 5.494903 | 6.1373806 |
| ASMM10P022 | 0.03644 | 0.3698934 | 1.7919276 | up | NM_007557 | Bmp7 | bone morphogenetic protein 7 | 10488.593 | 6705.605 | 13.487511 | 12.645998 | 9386.36 | 9258.46 | 12820.958 | 5888.687 | 6231.809 | 7996.319 | 13.221281 | 13.388276 | 13.852975 | 12.430334 | 12.471118 | 13.036543 |
| ASMM10P029 | 0.03648 | 0.3699809 | 1.4630476 | up | NM_001122 | Nfia | nuclear factor I/A | 14.029596 | 11.371886 | 4.06103 | 3.512053 | 17.323698 | 9.450022 | 15.315068 | 13.261979 | 12.834151 | 8.019528 | 4.1481566 | 3.7446542 | 4.290279 | 3.599121 | 3.5619462 | 3.3750925 |
| ASMM10P002 | 0.03651 | 0.3701092 | 1.4594 | up | NM_153387 | Tubgcp4 | tubulin, gamma complex associated protein 4 | 70.022093 | 57.630439 | 6.396008 | 5.850633 | 85.960785 | 71.024185 | 53.08131 | 69.8937 | 54.14106 | 48.856556 | 6.4769144 | 6.581001 | 6.1301084 | 6.0343614 | 5.6478825 | 5.8696537 |
| ASMM10P041 | 0.03661 | 0.3707032 | 1.4655373 | up | NM_145612 | Zfp810 | zinc finger protein 810 | 4399.7097 | 3532.5155 | 12.287491 | 11.736061 | 4905.6743 | 3621.339 | 4672.1157 | 3377.1846 | 3284.0212 | 3936.3406 | 12.286824 | 12.097562 | 12.478086 | 11.63574 | 11.558526 | 12.013917 |
| ASMM10P039 | 0.03666 | 0.3710573 | 1.355387 | up | NM_008054 | Fyn | Fyn proto-oncogene | 5511.4164 | 4793.0406 | 12.60746 | 12.168756 | 5480.9595 | 5150.8066 | 5902.483 | 4579.7275 | 4729.1274 | 5070.267 | 12.443678 | 12.587443 | 12.79126 | 12.064858 | 12.072507 | 12.368902 |
| ASMM10P006 | 0.03673 | 0.3711082 | 1.3518 | up | NM_007713 | Clk3 | CDC-like kinase 3 | 920.56433 | 818.95503 | 10.086236 | 9.651355 | 935.6934 | 905.3557 | 920.6439 | 791.0617 | 828.5717 | 837.2317 | 9.858473 | 10.175661 | 10.224575 | 9.553042 | 9.587212 | 9.81381 |
| ASMM10P031 | 0.03678 | 0.3711082 | 1.270913 | up | NM_011583 | Theg | testicular haploid expressed gene | 39.328916 | 37.100663 | 5.578137 | 5.232272 | 47.877975 | 32.54796 | 37.560814 | 36.714 | 38.952698 | 35.635292 | 5.6476917 | 5.4738007 | 5.612918 | 5.0988865 | 5.1727977 | 5.4251304 |
| ASMM10P023 | 0.03683 | 0.3711082 | 1.3992006 | up | NM_024254 | Katnbl1 | katanin p80 subunit B like 1 | 221.61396 | 195.01348 | 8.085674 | 7.601071 | 224.25365 | 215.07256 | 225.51567 | 190.8138 | 197.4591 | 196.76755 | 7.8372226 | 8.16205 | 8.257749 | 7.4966164 | 7.5215178 | 7.785079 |
| ASMM10P018 | 0.03684 | 0.3711082 | 1.5254977 | up | NM_146682 | Olfr76 | olfactory receptor 76 | 74.307086 | 59.511288 | 6.499592 | 5.890312 | 78.09597 | 58.068447 | 86.75684 | 59.49035 | 72.17015 | 46.873363 | 6.345811 | 6.2974815 | 6.855484 | 5.7999315 | 6.059969 | 5.811036 |
| ASMM10P036 | 0.03693 | 0.3711082 | 1.6133474 | up | NM_008035 | Folr2 | folate receptor 2 (fetal) | 207.75874 | 159.25422 | 7.992363 | 7.302306 | 193.37958 | 212.25319 | 217.64346 | 142.85962 | 173.80653 | 161.09651 | 7.624343 | 8.14503 | 8.207717 | 7.0698586 | 7.331894 | 7.505166 |
| ASMM10P010 | 0.03694 | 0.3711082 | 1.3862394 | up | NM_001081 | Slc9a3 | solute carrier family 9 (sodium/hydrogen exchanger), member 3 | 90.061149 | 79.721307 | 6.791387 | 6.320211 | 92.654076 | 85.46318 | 92.06619 | 83.134346 | 76.824394 | 79.20518 | 6.5868745 | 6.844413 | 6.942875 | 6.2822905 | 6.1511507 | 6.527192 |
| ASMM10P012 | 0.03697 | 0.3711082 | 1.3122695 | up | NM_009009 | Rad21 | RAD21 homolog (S. pombe) | 58952.728 | 55655.388 | 16.06138 | 15.669316 | 54801.766 | 56613.535 | 65442.883 | 54700.68 | 63734.773 | 48530.71 | 15.886818 | 16.00969 | 16.287632 | 15.599271 | 15.757243 | 15.651434 |
| ASMM10P002 | 0.03706 | 0.3717152 | 1.6564832 | up | NM_028354 | Tdp1 | tyrosyl-DNA phosphodiesterase 1 | 19.730094 | 14.161097 | 4.578234 | 3.85011 | 21.143139 | 16.63059 | 21.416552 | 14.93555 | 12.491642 | 15.056098 | 4.4436994 | 4.5174856 | 4.7735167 | 3.7761521 | 3.5210404 | 4.2531385 |
| ASMM10P006 | 0.03711 | 0.3719771 | 1.3090471 | up | NM_008637 | Nudt1 | nudix (nucleoside diphosphate linked moiety X)-type motif 1 | 28.974521 | 26.246043 | 5.115993 | 4.727476 | 38.28208 | 23.306719 | 25.334763 | 31.288214 | 27.82648 | 19.623434 | 5.3207045 | 5.0046053 | 5.0226693 | 4.868471 | 4.695276 | 4.618681 |
| ASMM10P046 | 0.03715 | 0.3720131 | 1.3618282 | up | NM_175489 | Osbpl8 | oxysterol binding protein-like 8 | 456.46278 | 403.64196 | 9.08492 | 8.639375 | 502.83917 | 500.4278 | 366.12137 | 397.4452 | 417.69482 | 395.78586 | 8.972174 | 9.347522 | 8.935063 | 8.560411 | 8.599555 | 8.758159 |
| ASMM10P010 | 0.03717 | 0.3720131 | 1.3528301 | up | NM_028281 | Pcbd2 | pterin 4 alpha carbinolamine dehydratase/dimerization cofactor of hepatocyte nuclear factor 1 a | 2940.3932 | 2570.2429 | 11.711029 | 11.275049 | 3134.456 | 3039.8977 | 2646.826 | 2439.3438 | 2456.3079 | 2815.077 | 11.599471 | 11.843632 | 11.689985 | 11.169648 | 11.135905 | 11.519593 |
| ASMM10P000 | 0.03726 | 0.3720131 | 1.7553373 | up | NM_001166 | Bcas3 | breast carcinoma amplified sequence 3 | 14.761218 | 9.804121 | 4.126956 | 3.315208 | 18.488726 | 9.585342 | 16.209587 | 8.684594 | 11.270182 | 9.457586 | 4.2442913 | 3.7626827 | 4.3738947 | 2.9627013 | 3.3788128 | 3.6041098 |
| ASMM10P031 | 0.03745 | 0.3720131 | 1.5943386 | up | NM_172274 | Cc2d2a | coiled-coil and C2 domain containing 2A | 227.39955 | 175.37252 | 8.109759 | 7.436801 | 215.24216 | 199.22734 | 267.72916 | 198.95018 | 187.88799 | 139.2794 | 7.7803984 | 8.056464 | 8.492415 | 7.5572934 | 7.4457273 | 7.3073826 |
| ASMM10P033 | 0.03749 | 0.3720131 | 1.4252716 | up | NM_001270 | Mical3 | microtubule associated monooxygenase, calponin and LIM domain containing 3 | 254.89941 | 217.86319 | 8.269773 | 7.758536 | 272.14758 | 214.00998 | 278.54068 | 209.253 | 224.91034 | 219.42622 | 8.106936 | 8.15671 | 8.545673 | 7.629881 | 7.707297 | 7.9384303 |
| ASMM10P020 | 0.03753 | 0.3720131 | 1.3173424 | up | NM_030091 | Ola1 | Obg-like ATPase 1 | 340.77488 | 316.45029 | 8.686955 | 8.289325 | 351.4462 | 331.4835 | 339.39493 | 315.91788 | 322.10974 | 311.32324 | 8.46538 | 8.765444 | 8.830042 | 8.225176 | 8.223455 | 8.419344 |
| ASMM10P017 | 0.03774 | 0.3720131 | 1.4404111 | up | NM_027072 | Plac8l1 | PLAC8-like 1 | 107.1277 | 89.495405 | 7.010183 | 6.483702 | 127.4437 | 78.26359 | 115.6758 | 96.52388 | 90.409355 | 81.55298 | 7.029542 | 6.7173085 | 7.2836986 | 6.4979177 | 6.3854904 | 6.567699 |
| ASMM10P041 | 0.03786 | 0.3720131 | 1.3446761 | up | NM_008884 | Pml | promyelocytic leukemia | 2231.0159 | 1960.3612 | 11.319037 | 10.891779 | 2286.2913 | 1931.078 | 2475.6785 | 1984.2828 | 2026.093 | 1870.7079 | 11.149354 | 11.217058 | 11.5907 | 10.869985 | 10.859718 | 10.945633 |
| ASMM10P052 | 0.03790 | 0.3720131 | 1.4384506 | up | NM_026512 | Bphl | biphenyl hydrolase-like (serine hydrolase, breast epithelial mucin-associated antigen) | 2364.2349 | 1959.9734 | 11.410772 | 10.886256 | 2375.9358 | 2397.4443 | 2319.3247 | 1875.9336 | 1832.7777 | 2171.209 | 11.202194 | 11.525645 | 11.504477 | 10.795786 | 10.708674 | 11.154309 |
| ASMM10P007 | 0.03794 | 0.3720131 | 1.3560706 | up | NM_015822 | Fbxl3 | F-box and leucine-rich repeat protein 3 | 273.55842 | 245.83412 | 8.370533 | 7.931101 | 292.4968 | 237.72366 | 290.4548 | 268.11703 | 236.69269 | 232.69264 | 8.203749 | 8.299494 | 8.6083555 | 7.9899917 | 7.779739 | 8.023571 |
| ASMM10P040 | 0.03799 | 0.3720131 | 1.7309627 | up | NM_007525 | Bard1 | BRCA1 associated RING domain 1 | 268.64703 | 189.12206 | 8.342533 | 7.550959 | 260.03757 | 231.70424 | 314.19928 | 181.30003 | 175.78397 | 210.28217 | 8.044323 | 8.264367 | 8.71891 | 7.423505 | 7.34747 | 7.881901 |
| ASMM10P005 | 0.03804 | 0.3720131 | 1.3429592 | up | NM_001162 | Fnip2 | folliculin interacting protein 2 | 84.983899 | 77.607188 | 6.707048 | 6.281633 | 86.46763 | 82.16843 | 86.315636 | 83.715675 | 75.83184 | 73.27405 | 6.48608 | 6.786887 | 6.8481774 | 6.291905 | 6.132488 | 6.420505 |
| ASMM10P004 | 0.03806 | 0.3720131 | 1.4619789 | up | NM_001077 | Gimap8 | GTPase, IMAP family member 8 | 15.228875 | 12.38499 | 4.209353 | 3.661431 | 15.326578 | 14.71895 | 15.641098 | 12.19078 | 13.17135 | 11.79284 | 3.9613898 | 4.345283 | 4.3213873 | 3.4737964 | 3.5996253 | 3.910871 |
| ASMM10P000 | 0.03809 | 0.3720131 | 1.279404 | up | NM_001013 | Lipo1 | lipase, member O1 | 153.46008 | 147.84515 | 7.551245 | 7.195773 | 176.64742 | 142.02232 | 141.71051 | 133.1934 | 169.0206 | 141.32144 | 7.4911957 | 7.574326 | 7.5882144 | 6.9716086 | 7.2890773 | 7.3266344 |
| ASMM10P055 | 0.03811 | 0.3720131 | 1.7315881 | up | ENSMUST( | Wdr65 | WD repeat domain 65 | 194.47017 | 139.50675 | 7.883854 | 7.091759 | 231.73395 | 187.75844 | 163.91812 | 110.33422 | 139.81013 | 168.37589 | 7.8820987 | 7.974589 | 7.7948756 | 6.6982164 | 7.0085316 | 7.568528 |
| ASMM10P012 | 0.03813 | 0.3720131 | 1.677254 | up | NM_027174 | Col22a1 | collagen, type XXII, alpha 1 | 1757.7414 | 1259.5127 | 10.992843 | 10.246742 | 1791.6862 | 1518.9243 | 1962.6136 | 1333.9128 | 1017.1888 | 1427.4365 | 10.807421 | 10.893076 | 11.278032 | 10.298906 | 9.881609 | 10.559711 |
| ASMM10P050 | 0.03817 | 0.3720131 | 2.0715404 | up | NM_181815 | Cep128 | centrosomal protein 128 | 25.710425 | 14.450819 | 4.925177 | 3.874473 | 19.879684 | 21.516706 | 35.734886 | 16.126587 | 15.83824 | 11.387631 | 4.345085 | 4.8922944 | 5.5381503 | 3.891195 | 3.8730438 | 3.859179 |
| ASMM10P034 | 0.03817 | 0.3720131 | 1.3085688 | up | NM_212468 | Ssbp1 | single-stranded DNA binding protein 1 | 302.59108 | 285.36417 | 8.51859 | 8.1306 | 319.28708 | 289.04663 | 299.43954 | 306.61966 | 325.4602 | 224.01266 | 8.328913 | 8.573689 | 8.653168 | 8.184497 | 8.2375555 | 7.969748 |
| ASMM10P023 | 0.03820 | 0.3720131 | 1.3982759 | up | NM_001142 | Lmo2 | LIM domain only 2 | 236.41184 | 207.75319 | 8.173499 | 7.68985 | 229.68944 | 246.00104 | 233.54504 | 226.22032 | 208.25153 | 188.78773 | 7.87016 | 8.348602 | 8.301735 | 7.7419252 | 7.59997 | 7.7276545 |

| | | | | | | | | | | | | | | | | | | | | | | |
|---|---|---|---|---|---|---|---|---|---|---|---|---|---|---|---|---|---|---|---|---|---|---|
| ASMM10P004 0.03821 | 0.3720131 | 1.3327203 | up | NM_001128 | Gm2382 | predicted gene 2382 | 386.66338 | 354.71471 | 8.862922 | 8.448548 | 426.16086 | 347.57333 | 386.25595 | 321.7584 | 381.61157 | 360.77417 | 8.738565 | 8.835083 | 9.015119 | 8.24983 | 8.468937 | 8.626878 |
| ASMM10P000 0.03822 | 0.3720131 | 1.434453 | up | NM_023906 | Asb3 | ankyrin repeat and SOCS box-containing 3 | 268.54995 | 228.3255 | 8.068711 | 7.826728 | 277.18878 | 241.9737 | 286.48737 | 226.64983 | 222.36066 | 235.96602 | 8.133556 | 8.323421 | 8.584709 | 7.743645 | 7.69171 | 8.044829 |
| ASMM10P049 0.03823 | 0.3720131 | 1.3508162 | up | NM_019679 | Fmnl1 | formin-like 1 | 396.5007 | 357.9766 | 8.896917 | 8.463085 | 400.06317 | 364.46606 | 424.97287 | 376.37784 | 383.5825 | 313.96945 | 8.651937 | 8.896619 | 9.142194 | 8.482081 | 8.477085 | 8.43009 |
| ASMM10P006 0.03827 | 0.3720131 | 1.9019008 | up | NM_009109 | Ryr1 | ryanodine receptor 1, skeletal muscle | 232.28159 | 145.5209 | 8.094468 | 7.167026 | 285.37808 | 151.11418 | 260.3525 | 133.7096 | 135.79497 | 167.05812 | 8.1735935 | 7.6553984 | 8.4544115 | 6.976355 | 6.967429 | 7.5572934 |
| ASMM10P008 0.03838 | 0.3720131 | 2.6213357 | up | NM_028857 | Ccdc30 | coiled-coil domain containing 30 | 15.773089 | 6.548798 | 4.138695 | 2.748393 | 20.150013 | 6.920263 | 20.24899 | 9.150742 | 4.9999995 | 5.4956536 | 4.366068 | 3.3565423 | 4.6934743 | 3.0495691 | 2.3265402 | 2.869069 |
| ASMM10P003 0.03845 | 0.3720131 | 1.9172833 | up | NM_020331 | Gtf2ird1 | general transcription factor II I repeat domain-containing 1 | 28.943703 | 17.990603 | 5.127012 | 4.187949 | 32.474953 | 21.380135 | 32.97602 | 15.667639 | 17.195908 | 21.108263 | 5.0815725 | 4.8818564 | 5.4176083 | 3.8484962 | 4.001621 | 4.7137294 |
| ASMM10P024 0.03846 | 0.3720131 | 2.3542496 | up | NM_001195 | Kcnmb3 | potassium large conductance calcium-activated channel, subfamily M, beta member 3 | 39.481499 | 18.902098 | 5.506484 | 4.271217 | 37.825226 | 22.43765 | 58.18162 | 18.260891 | 21.574816 | 16.870588 | 5.301051 | 4.9509344 | 6.267467 | 4.0745935 | 4.326582 | 4.412475 |
| ASMM10P000 0.03850 | 0.3720131 | 1.4239632 | up | NM_175683 | Dclre1c | DNA cross-link repair 1C, PSO2 homolog (S. cerevisiae) | 242.46259 | 203.35831 | 8.170474 | 7.660562 | 312.05673 | 177.01451 | 238.31653 | 215.74443 | 194.83643 | 199.49406 | 8.296954 | 7.8843536 | 8.330113 | 7.6749415 | 7.502664 | 7.8040795 |
| ASMM10P040 0.03850 | 0.3720131 | 1.4470462 | up | NM_008278 | Hpgd | hydroxyprostaglandin dehydrogenase 15 (NAD) | 259.64556 | 221.48809 | 8.304338 | 7.771227 | 268.01584 | 245.03894 | 265.8819 | 192.68689 | 262.3689 | 209.40848 | 8.085852 | 8.342939 | 8.484224 | 7.5105944 | 7.927608 | 7.8754797 |
| ASMM10P005 0.03851 | 0.3720131 | 1.483304 | up | NM_001163 | Lrrcc1 | leucine rich repeat and coiled-coil domain containing 1 | 3040.8731 | 2386.4037 | 11.743204 | 11.17439 | 3119.9382 | 2400.6826 | 3601.9985 | 2319.2756 | 2563.361 | 2276.5745 | 11.593206 | 11.527743 | 12.108665 | 11.101876 | 11.203506 | 11.217789 |
| ASMM10P004 0.03855 | 0.3720131 | 1.4133665 | up | NM_001025 | Dnm1l | dynamin 1-like | 265.96535 | 230.48973 | 8.339403 | 7.840267 | 267.21237 | 262.47748 | 268.2062 | 234.0266 | 223.20937 | 234.23322 | 8.081198 | 8.441183 | 8.495828 | 7.791541 | 7.69564 | 8.033621 |
| ASMM10P043 0.03870 | 0.3720131 | 1.2653792 | up | NM_178740 | Slitrk4 | SLIT and NTRK-like family, member 4 | 10246.659 | 9420.4643 | 13.475652 | 13.136082 | 10899.476 | 9003.579 | 10836.922 | 9498.323 | 9185.108 | 9577.962 | 13.454915 | 13.350657 | 13.621385 | 13.112029 | 13.011931 | 13.284287 |
| ASMM10P012 0.03877 | 0.3720131 | 1.3262983 | up | NM_180588 | Reep4 | receptor accessory protein 4 | 216.9499 | 200.71397 | 8.048966 | 7.641561 | 221.33075 | 231.45273 | 198.06621 | 207.47038 | 206.88512 | 187.78642 | 7.8191414 | 8.263548 | 8.064208 | 7.6147356 | 7.590614 | 7.719332 |
| ASMM10P014 0.03878 | 0.3720131 | 1.4023102 | up | NM_023175 | Nit2 | nitrilase family, member 2 | 1136.1155 | 972.81817 | 10.38394 | 9.896135 | 1097.4629 | 1146.8126 | 1164.0709 | 949.40485 | 1014.7509 | 954.2988 | 10.081117 | 10.512827 | 10.557877 | 9.814453 | 9.8781185 | 9.995833 |
| ASMM10P009 0.03878 | 0.3720131 | 1.3760906 | up | NM_207525 | Opa3 | optic atrophy 3 | 27.655336 | 23.933526 | 5.04853 | 4.587955 | 35.862152 | 24.985054 | 22.118801 | 25.545265 | 29.118958 | 17.136356 | 5.2222214 | 5.100693 | 4.8226757 | 4.565204 | 4.7635856 | 4.435074 |
| ASMM10P037 0.03879 | 0.3720131 | 2.5443351 | up | NM_001038 | Gm4745 | predicted gene 4745 | 138.33865 | 63.76759 | 7.287753 | 5.940465 | 184.34575 | 68.6443 | 162.02591 | 70.02846 | 83.330795 | 37.943516 | 7.55311 | 6.531497 | 7.778653 | 6.037056 | 6.2724476 | 5.5118904 |
| ASMM10P028 0.03890 | 0.3720131 | 1.2823957 | up | NM_001098 | Sdcbp | syndecan binding protein | 11308.074 | 10289.049 | 13.617069 | 13.258227 | 11385.423 | 11368.524 | 11170.276 | 10030.909 | 9942.719 | 10893.519 | 13.510799 | 13.676548 | 13.663859 | 13.179604 | 13.127193 | 13.467885 |
| ASMM10P052 0.03894 | 0.3720131 | 1.9331258 | up | NM_001033 | Dcp1b | DCP1 decapping enzyme homolog B (S. cerevisiae) | 907.97727 | 556.7363 | 10.026888 | 9.075952 | 918.6572 | 1150.0679 | 655.2067 | 434.54935 | 639.73474 | 595.9248 | 9.834962 | 10.516112 | 9.7295885 | 8.691373 | 9.212754 | 9.323729 |
| ASMM10P050 0.03896 | 0.3720131 | 1.8132628 | up | NM_001271 | Dnmt3a | DNA methyltransferase 3A | 15.614373 | 10.089061 | 4.224434 | 3.365846 | 12.86999 | 15.191999 | 18.781132 | 11.20691 | 10.148301 | 8.911973 | 3.6965978 | 4.389773 | 4.586931 | 3.3519824 | 3.2217379 | 3.5238173 |
| ASMM10P006 0.03897 | 0.3720131 | 1.3043409 | up | NM_001277 | Zmat4 | zinc finger, matrin type 4 | 118.74679 | 111.12902 | 7.172144 | 6.788823 | 143.06238 | 104.75827 | 108.41971 | 127.7753 | 100.72129 | 104.89048 | 7.192529 | 7.1379232 | 7.18598 | 6.911962 | 6.5377493 | 6.916758 |
| ASMM10P009 0.03898 | 0.3720131 | 1.8247955 | up | NM_175333 | Slc25a41 | solute carrier family 25, member 41 | 31.123759 | 21.05851 | 5.251274 | 4.38354 | 31.264076 | 33.05521 | 29.051992 | 29.64527 | 16.346497 | 17.183762 | 5.024273 | 5.4953756 | 5.2341743 | 4.7901506 | 3.9210525 | 4.4394155 |
| ASMM10P035 0.03898 | 0.3720131 | 1.3790733 | up | NM_001033 | Atxn7l3b | ataxin 7-like 3B | 822.6618 | 715.7811 | 9.921295 | 9.457595 | 861.0284 | 827.6012 | 779.3558 | 709.1016 | 671.1203 | 767.1214 | 9.7404375 | 10.044509 | 9.978937 | 9.400429 | 9.280146 | 9.692211 |
| ASMM10P041 0.03906 | 0.3720131 | 1.5944106 | up | NM_001024 | Adamts15 | a disintegrin-like and metallopeptidase (reprolysin type) with thrombospondin type 1 motif, | 1017.6633 | 754.4919 | 10.199434 | 9.52641 | 1222.0753 | 745.3207 | 1085.5938 | 681.58655 | 736.2849 | 845.60425 | 10.234622 | 9.900075 | 10.463604 | 9.33829 | 9.415874 | 9.8250675 |
| ASMM10P052 0.03919 | 0.3720131 | 1.256953 | up | NM_001122 | Gpbp1 | GC-rich promoter binding protein 1 | 356.34063 | 347.9747 | 8.750149 | 8.420218 | 373.3579 | 354.96732 | 340.69666 | 367.20844 | 386.61255 | 290.1031 | 8.553774 | 8.861079 | 8.835593 | 8.441797 | 8.489349 | 8.329508 |
| ASMM10P030 0.03920 | 0.3720131 | 1.4086907 | up | NM_001114 | Scfd2 | Sec1 family domain containing 2 | 927.57202 | 785.88855 | 10.086288 | 9.591933 | 992.4189 | 791.32007 | 998.9771 | 742.90356 | 790.6346 | 824.1275 | 9.931698 | 9.984414 | 10.342751 | 9.465393 | 9.519552 | 9.7908535 |
| ASMM10P007 0.03923 | 0.3720131 | 1.5803598 | up | NM_024255 | Hsdl2 | hydroxysteroid dehydrogenase like 2 | 68.880487 | 53.876908 | 6.404788 | 5.744535 | 68.47098 | 68.49853 | 69.67195 | 44.744007 | 67.76398 | 49.122738 | 6.156687 | 6.528503 | 6.5291743 | 5.3881254 | 5.9687057 | 5.876774 |
| ASMM10P036 0.03924 | 0.3720131 | 1.4400022 | up | NM_001115 | Fancf | Fanconi anemia, complementation group F | 5205.4517 | 4206.4667 | 12.513188 | 11.987117 | 4958.766 | 4498.306 | 6159.283 | 4198.0713 | 4357.67 | 4063.6587 | 12.302759 | 12.386082 | 12.850723 | 11.950032 | 11.951416 | 12.059903 |
| ASMM10P010 0.03924 | 0.3720131 | 1.7989017 | up | NM_008533 | Cd180 | CD180 antigen | 42.23334 | 28.381266 | 5.676048 | 4.828931 | 52.949734 | 35.508736 | 38.24155 | 22.225588 | 30.260351 | 32.65786 | 5.78893 | 5.5996194 | 5.6395936 | 4.359432 | 4.816519 | 5.310843 |
| ASMM10P039 0.03927 | 0.3720131 | 1.3139137 | up | NM_177793 | Mettl24 | methyltransferase like 24 | 130.06561 | 121.59788 | 7.30413 | 6.910259 | 156.34131 | 106.06702 | 127.78851 | 113.21133 | 147.30084 | 104.28147 | 7.3216734 | 7.15609 | 7.434626 | 6.73382 | 7.0887604 | 6.9081974 |
| ASMM10P009 0.03931 | 0.3720131 | 1.259451 | up | NM_177610 | Duox2 | dual oxidase 2 | 530.30596 | 501.79882 | 9.280252 | 8.947457 | 685.2531 | 470.10513 | 435.55966 | 491.19308 | 497.87228 | 516.3311 | 9.405667 | 9.25821 | 9.176879 | 8.867141 | 8.852594 | 9.122636 |
| ASMM10P043 0.03934 | 0.3720131 | 1.3125053 | up | NM_134163 | Mbnl3 | muscleblind-like 3 (Drosophila) | 253.23949 | 233.56676 | 8.251372 | 7.859049 | 298.38696 | 202.37172 | 258.9598 | 233.08821 | 237.15948 | 230.45259 | 8.230275 | 8.078813 | 8.445028 | 7.7844214 | 7.783484 | 8.009241 |
| ASMM10P029 0.03935 | 0.3720131 | 1.4242322 | up | NM_133889 | Bsdc1 | BSD domain containing 1 | 2737.5609 | 2297.6723 | 11.613022 | 11.102837 | 2916.0486 | 2276.4675 | 3020.1665 | 2641.7795 | 2428.2927 | 1822.9447 | 11.496356 | 11.466665 | 11.876044 | 11.279495 | 11.116393 | 10.912624 |
| ASMM10P036 0.03936 | 0.3720131 | 1.5272327 | up | NM_028177 | Ndufab1 | NADH dehydrogenase (ubiquinone) 1, alpha/beta subcomplex, 1 | 266.77849 | 212.39346 | 8.333274 | 7.722354 | 252.45555 | 311.68402 | 236.19589 | 210.69849 | 223.10234 | 203.37955 | 8.003855 | 8.677392 | 8.318574 | 7.6407924 | 7.6951227 | 7.8311462 |
| ASMM10P018 0.03938 | 0.3720131 | 1.3165452 | up | NM_009793 | Camk4 | calcium/calmodulin-dependent protein kinase IV | 470.93686 | 430.20954 | 9.124177 | 8.72742 | 544.80774 | 494.7503 | 373.25253 | 471.66522 | 411.80997 | 407.15344 | 9.074223 | 9.33321 | 8.965098 | 8.807448 | 8.580517 | 8.794295 |
| ASMM10P053 0.03940 | 0.3720131 | 1.4500515 | up | NM_025418 | Vta1 | Vps20-associated 1 homolog (S. cerevisiae) | 186.54688 | 158.13688 | 7.833879 | 7.297775 | 186.56947 | 204.34618 | 168.725 | 166.42572 | 151.87819 | 156.10672 | 7.572834 | 8.091057 | 7.837747 | 7.298205 | 7.1309085 | 7.464212 |
| ASMM10P005 0.03942 | 0.3720131 | 2.2184774 | up | NM_001167 | Nme8 | NME/NM23 family member 8 | 23.987344 | 12.519599 | 4.81291 | 3.66334 | 22.555035 | 15.95954 | 33.447456 | 10.326879 | 15.172499 | 12.059419 | 4.541649 | 4.459899 | 5.4371824 | 3.233961 | 3.8118925 | 3.9441674 |
| ASMM10P047 0.03947 | 0.3720131 | 1.2671357 | up | NM_033562 | Derl2 | Der1-like domain family, member 2 | 3950.2164 | 3701.6253 | 12.136292 | 11.794721 | 3923.9639 | 3912.2415 | 4014.4438 | 3528.636 | 4104.2935 | 3471.9465 | 11.942134 | 12.189873 | 12.27687 | 11.693198 | 11.871689 | 11.819277 |
| ASMM10P000 0.03950 | 0.3720131 | 1.8517856 | up | NM_175933 | Pex5 | peroxisomal biogenesis factor 5 | 33.708548 | 22.310227 | 5.357594 | 4.468677 | 29.695213 | 29.908625 | 41.521805 | 20.858356 | 30.300924 | 15.771401 | 4.9501405 | 5.3512506 | 5.7713923 | 4.2699285 | 4.819375 | 4.3167286 |
| ASMM10P046 0.03953 | 0.3720131 | 1.3043322 | up | NM_153069 | Leap2 | liver-expressed antimicrobial peptide 2 | 27.135053 | 24.804996 | 5.026343 | 4.643032 | 34.356277 | 21.01078 | 26.038101 | 25.337349 | 30.40114 | 18.676498 | 5.159517 | 4.8568764 | 5.0626354 | 4.5538673 | 4.824664 | 4.5505633 |
| ASMM10P034 0.03953 | 0.3720131 | 1.2997132 | up | NM_001164 | Nagk | N-acetylglucosamine kinase | 426.58744 | 397.41044 | 8.994352 | 8.616158 | 476.22174 | 368.80936 | 434.73123 | 378.52185 | 409.91125 | 403.79822 | 8.896619 | 8.912225 | 9.174211 | 8.490425 | 8.573374 | 8.784676 |
| ASMM10P009 0.03959 | 0.3720131 | 1.5202518 | up | NM_172522 | Megf11 | multiple EGF-like-domains 11 | 296.98097 | 239.20634 | 8.487502 | 7.883192 | 321.87796 | 305.7641 | 263.30084 | 211.26248 | 234.84344 | 271.5131 | 8.340923 | 8.650058 | 8.471525 | 7.64518 | 7.768899 | 8.235496 |
| ASMM10P025 0.03959 | 0.3720131 | 1.5872265 | up | NM_001142 | Fam46c | family with sequence similarity 46, member C | 431.55557 | 330.46491 | 9.016174 | 8.349666 | 386.13187 | 441.9513 | 466.58353 | 315.66843 | 350.69846 | 325.02783 | 8.60281 | 9.170189 | 9.275522 | 8.224445 | 8.345088 | 8.479464 |
| ASMM10P014 0.03964 | 0.3720131 | 2.7148566 | up | NM_009930 | Col3a1 | collagen, type III, alpha 1 | 115.93795 | 49.255821 | 7.054818 | 5.613942 | 113.80589 | 166.37825 | 67.62972 | 40.039944 | 47.116135 | 60.611385 | 6.87927 | 7.795934 | 6.489251 | 5.2321687 | 5.445427 | 6.1642313 |
| ASMM10P009 0.03969 | 0.3720131 | 1.3668201 | up | NM_177567 | BC049762 | cDNA sequence BC049762 | 736.72158 | 646.60925 | 9.752032 | 9.301209 | 830.90173 | 583.98474 | 795.27826 | 635.8316 | 746.67285 | 557.3233 | 9.69051 | 9.557847 | 10.007739 | 9.235135 | 9.43791 | 9.230581 |
| ASMM10P011 0.03972 | 0.3720131 | 1.3773431 | up | NM_134079 | Adk | adenosine kinase | 4498.8956 | 3834.6444 | 12.317306 | 11.855418 | 4750.2793 | 3877.5686 | 4868.839 | 3546.2065 | 3875.5312 | 4082.1956 | 12.239964 | 12.179147 | 12.532807 | 11.702554 | 11.799293 | 12.064407 |
| ASMM10P043 0.03975 | 0.3720131 | 1.416421 | up | NM_007807 | Cybb | cytochrome b-245, beta polypeptide | 62.115023 | 53.600299 | 6.256147 | 5.753897 | 63.061695 | 56.097813 | 67.18556 | 50.69904 | 58.50638 | 51.595478 | 6.040302 | 6.248406 | 6.479733 | 5.565892 | 5.7531476 | 5.942651 |
| ASMM10P027 0.03976 | 0.3720131 | 1.287829 | up | NM_008822 | Pex7 | peroxisomal biogenesis factor 7 | 1029.6418 | 953.31774 | 10.231911 | 9.86697 | 1183.03 | 883.7168 | 1022.1787 | 936.1575 | 927.51715 | 996.27856 | 10.184706 | 10.136191 | 10.374837 | 9.791549 | 9.749494 | 10.059868 |
| ASMM10P008 0.03982 | 0.3720131 | 1.6898878 | up | NM_028478 | Rassf6 | Ras association (RalGDS/AF-6) domain family member 6 | 79.955029 | 55.830243 | 6.564251 | 5.807323 | 111.17732 | 58.677803 | 70.00997 | 51.9419 | 52.88775 | 62.66108 | 6.844413 | 6.3114285 | 6.536911 | 5.599963 | 5.6164665 | 6.2055407 |
| ASMM10P025 0.03983 | 0.3720131 | 1.2757009 | up | NM_008212 | Hadh | hydroxyacyl-Coenzyme A dehydrogenase | 28397.436 | 26456.169 | 14.948073 | 14.596782 | 27904.855 | 27438.639 | 29848.814 | 23646.621 | 29752.504 | 25969.383 | 14.864944 | 14.907991 | 15.071283 | 14.399956 | 14.68298 | 14.707411 |
| ASMM10P041 0.03983 | 0.3720131 | 1.560179 | up | NM_028974 | Kbtbd13 | kelch repeat and BTB (POZ) domain containing 13 | 307.92597 | 240.33938 | 8.540472 | 7.89876 | 279.035 | 303.5398 | 341.2076 | 259.75464 | 238.83199 | 222.4315 | 8.142797 | 8.639958 | 8.838661 | 7.943649 | 7.79261 | 7.9600224 |
| ASMM10P012 0.03986 | 0.3720131 | 1.2905446 | up | NM_009437 | Tst | thiosulfate sulfurtransferase, mitochondrial | 10811.981 | 9829.4093 | 13.552004 | 13.184024 | 11356.814 | 10071.506 | 11007.622 | 8588.251 | 10770.477 | 10129.5 | 13.504865 | 13.510799 | 13.640349 | 12.967768 | 13.227837 | 13.356468 |
| ASMM10P025 0.03991 | 0.3720772 | 1.3712314 | up | NM_007382 | Acadm | acyl-Coenzyme A dehydrogenase, medium chain | 12070.744 | 10301.03 | 13.706255 | 13.250783 | 10804.455 | 11971.55 | 13436.228 | 9731.03 | 11303.196 | 9868.863 | 13.443895 | 13.74483 | 13.930041 | 13.14263 | 13.292971 | 13.316749 |
| ASMM10P039 0.03998 | 0.3721001 | 1.2879114 | up | NM_133765 | Fbxo31 | F-box protein 31 | 70399.312 | 69730.268 | 16.363778 | 15.998745 | 70714.49 | 65335.555 | 75147.89 | 67139.58 | 75934.06 | 66117.164 | 16.289722 | 16.243767 | 16.557846 | 15.870261 | 16.005611 | 16.120363 |
| ASMM10P040 0.04004 | 0.3722978 | 2.3385398 | up | NM_009864 | Cdh1 | cadherin 1 | 1741.9015 | 930.77355 | 10.982989 | 9.757381 | 1609.7684 | 1752.4817 | 1863.4545 | 754.2704 | 677.14246 | 1360.9078 | 10.641051 | 11.091812 | 11.216104 | 9.489465 | 9.294556 | 10.488122 |
| ASMM10P014 0.04012 | 0.3726643 | 1.2820104 | up | NM_001033 | Slc9a2 | solute carrier family 9 (sodium/hydrogen exchanger), member 2 | 62767.609 | 60912.133 | 16.159605 | 15.801197 | 64953.457 | 55515.02 | 67834.35 | 65133.87 | 62324.723 | 55277.805 | 16.141523 | 15.973813 | 16.36348 | 15.829924 | 15.722624 | 15.851044 |
| ASMM10P047 0.04013 | 0.3726643 | 1.6297309 | up | NM_023913 | Ern1 | endoplasmic reticulum (ER) to nucleus signalling 1 | 77.436213 | 56.881009 | 6.544612 | 5.839978 | 84.72266 | 55.25989 | 92.32609 | 56.364456 | 58.64926 | 55.62931 | 6.4595947 | 6.2273903 | 6.946851 | 5.7201304 | 5.7569976 | 6.0428066 |
| ASMM10P025 0.04018 | 0.3730427 | 1.4909928 | up | NM_001163 | Fam102b | family with sequence similarity 102, member B | 13750.43 | 10731.294 | 13.888506 | 13.312232 | 12186.241 | 14246.574 | 14818.425 | 10815.445 | 9751.617 | 11626.819 | 13.616282 | 13.98202 | 14.067215 | 13.277399 | 13.099762 | 13.559536 |
| ASMM10P004 0.04021 | 0.3730634 | 1.2735002 | up | NM_009637 | Aebp2 | AE binding protein 2 | 205.18814 | 198.16608 | 7.972155 | 7.623356 | 212.21405 | 210.8132 | 192.53716 | 212.0614 | 201.56805 | 180.8688 | 7.759178 | 8.133241 | 8.024047 | 7.6507826 | 7.5533147 | 7.665971 |
| ASMM10P027 0.04024 | 0.3730634 | 1.3984664 | up | NM_001003 | Cyp4x1 | cytochrome P450, family 4, subfamily x, polypeptide 1 | 65.364638 | 56.544156 | 6.314403 | 5.830557 | 79.566505 | 56.33278 | 60.19463 | 53.995613 | 55.73368 | 59.903175 | 6.3714952 | 6.25517 | 6.316544 | 5.6551538 | 5.6902905 | 6.146228 |
| ASMM10P001 0.04026 | 0.3730634 | 1.7948812 | up | NM_001039 | Trim63 | tripartite motif-containing 63 | 271.91883 | 182.87304 | 8.343037 | 7.499148 | 245.65346 | 218.35072 | 351.7523 | 206.33922 | 191.82048 | 150.45941 | 7.964219 | 8.1846 | 8.880291 | 7.607604 | 7.4794993 | 7.4103417 |
| ASMM10P040 0.04031 | 0.3732779 | 1.4276558 | up | NM_199446 | Phkb | phosphorylase kinase beta | 39.89171 | 34.32487 | 5.616185 | 5.102537 | 38.17418 | 41.281322 | 40.219627 | 36.941963 | 40.725388 | 25.30726 | 5.31688 | 5.811544 | 5.7201304 | 5.10893 | 5.2378287 | 4.960851 |
| ASMM10P007 0.04031 | 0.3732779 | 1.2826884 | up | NM_021486 | Bcmo1 | beta-carotene 15,15'-monooxygenase | 13.642743 | 12.791024 | 4.048936 | 3.689765 | 14.767219 | 13.522398 | 12.638611 | 15.55454 | 13.068178 | 9.750354 | 3.9093103 | 4.2256527 | 4.011845 | 3.8375483 | 3.5868516 | 3.6448958 |
| ASMM10P018 0.04034 | 0.3732779 | 1.280553 | up | NM_018821 | Socs6 | suppressor of cytokine signaling 6 | 146.37414 | 141.62077 | 7.492919 | 7.136152 | 151.59012 | 137.268 | 150.2643 | 155.12737 | 149.06972 | 120.66521 | 7.2762756 | 7.530063 | 7.6724186 | 7.1930923 | 7.104377 | 7.110987 |
| ASMM10P016 0.04043 | 0.3732779 | 1.6308071 | up | ENSMUST0 | Runx2 | runt related transcription factor 2 | 213.54323 | 160.47094 | 8.023362 | 7.317776 | 185.18942 | 230.01337 | 225.42691 | 158.5129 | 174.82181 | 148.07811 | 7.561335 | 8.252132 | 8.256619 | 7.2237344 | 7.340776 | 7.3888173 |
| ASMM10P021 0.04045 | 0.3732779 | 1.293838 | up | NM_175466 | Zfp770 | zinc finger protein 770 | 9775.122 | 8799.7788 | 13.411366 | 13.039709 | 9369.096 | 9690.801 | 10265.469 | 8552.062 | 8906.6045 | 8940.67 | 13.220484 | 13.453089 | 13.560525 | 12.961115 | 12.971343 | 13.186669 |
| ASMM10P039 0.04048 | 0.3732779 | 1.8378822 | up | NM_001081 | Neto2 | neuropilin (NRP) and tolloid (TLL)-like 2 | 25.188437 | 16.334446 | 4.93441 | 4.056365 | 23.453892 | 27.503645 | 24.607775 | 14.39037 | 16.675318 | 17.937649 | 4.5948257 | 5.228581 | 4.979822 | 3.721516 | 3.9531317 | 4.494448 |
| ASMM10P025 0.04052 | 0.3732779 | 1.7190857 | up | NM_001163 | 4930579F01 | RIKEN cDNA 4930579F01 gene | 55.718865 | 39.862744 | 6.076764 | 5.295122 | 58.82867 | 64.77963 | 43.548294 | 36.08932 | 54.436462 | 29.06245 | 5.935876 | 6.4502945 | 5.84412 | 5.076238 | 5.655471 | 5.153657 |
| ASMM10P050 0.04055 | 0.3733011 | 1.5229589 | up | NM_001142 | AK010878 | cDNA sequence AK010878 | 1416.4562 | 1119.2127 | 10.686432 | 10.079555 | 1318.8315 | 1600.5361 | 1330.001 | 1278.2285 | 1180.9595 | 898.4501 | 10.351173 | 10.970524 | 10.737599 | 10.237649 | 10.086708 | 9.914308 |
| ASMM10P043 0.04057 | 0.3733204 | 1.294202 | up | NM_176979 | Topbp1 | topoisomerase (DNA) II binding protein 1 | 750.00536 | 696.33335 | 9.790095 | 9.418033 | 781.37787 | 764.7857 | 703.8525 | 730.7162 | 669.5344 | 688.74945 | 9.6009445 | 9.936957 | 9.832385 | 9.444343 | 9.276231 | 9.533524 |
| ASMM10P007 0.04060 | 0.3733665 | 1.5033463 | up | NM_012037 | Vat1 | vesicle amine transport protein 1 homolog (T californica) | 2677.8992 | 2062.5365 | 11.555002 | 10.966825 | 3190.0015 | 1897.2615 | 2946.4346 | 2189.2368 | 2042.38 | 1955.9928 | 11.627917 | 11.193197 | 11.843893 | 11.016554 | 10.871968 | 11.011953 |
| ASMM10P019 0.04083 | 0.3745456 | 1.3589063 | up | NM_145501 | Pi4k2a | phosphatidylinositol 4-kinase type 2 alpha | 1177.0603 | 1030.7559 | 10.425399 | 9.982953 | 1214.0524 | 1051.08 | 1266.0486 | 1046.7188 | 1009.6303 | 1035.9187 | 10.224049 | 10.38488 | 10.667267 | 9.960259 | 9.869798 | 10.118801 |
| ASMM10P011 0.04094 | 0.3746733 | 1.4572337 | up | NM_001025 | Sema3g | sema domain, immunoglobulin domain (Ig), short basic domain, secreted, (semaphorin) 3G | 131.41705 | 110.31594 | 7.321687 | 6.778454 | 156.27641 | 108.08532 | 129.88942 | 99.15806 | 112.61448 | 119.17528 | 7.320253 | 7.184043 | 7.460764 | 6.5400186 | 6.7021055 | 7.093239 |
| ASMM10P048 0.04098 | 0.3746733 | 1.308759 | up | NM_145962 | Pank3 | pantothenate kinase 3 | 845.99212 | 774.84124 | 9.959795 | 9.571596 | 849.0304 | 893.1113 | 795.83466 | 769.81525 | 825.33777 | 729.3707 | 9.7191725 | 10.151515 | 10.008698 | 9.516168 | 9.582352 | 9.616267 |
| ASMM10P033 0.04100 | 0.3746733 | 1.3339792 | up | NM_175523 | Ppm1k | protein phosphatase 1K (PP2C domain containing) | 48279.612 | 43929.182 | 15.754558 | 15.338822 | 44505.477 | 45411.555 | 54921.805 | 45932.066 | 45727.785 | 40127.695 | 15.586971 | 15.649504 | 16.0272 | 15.351208 | 15.290175 | 15.375084 |
| ASMM10P009 0.04104 | 0.3746733 | 1.5538136 | up | NM_173786 | Apol9a | apolipoprotein L 9a | 17.73887 | 13.651214 | 4.419922 | 3.784108 | 16.00542 | 16.660141 | 20.55105 | 16.272581 | 14.1797 | 10.50136 | 4.0247955 | 4.520783 | 4.7141867 | 3.9031584 | 3.7113287 | 3.3378378 |
| ASMM10P004 0.04113 | 0.3746733 | 1.4852873 | up | NM_001081 | Bod1l | biorientation of chromosomes in cell division 1-like | 42.814604 | 33.345469 | 5.653402 | 5.08266 | 61.1714 | 31.456516 | 35.815895 | 37.80714 | 32.58435 | 29.644917 | 5.9958315 | 5.422516 | 5.5418587 | 5.1430044 | 4.9239206 | 5.181055 |
| ASMM10P036 0.04114 | 0.3746733 | 1.7357925 | up | NM_007438 | Aldoa | aldolase A, fructose-bisphosphate | 76.24611 | 53.444909 | 6.539437 | 5.743843 | 62.4577 | 76.10209 | 90.17854 | 59.874157 | 57.561756 | 42.898815 | 6.025693 | 6.679447 | 6.9131722 | 5.8089747 | 5.7326775 | 5.6898766 |
| ASMM10P053 0.04117 | 0.3746733 | 2.103528 | up | NM_001163 | Plekhb1 | pleckstrin homology domain containing, family B (evectins) member 1 | 45.313714 | 24.931293 | 5.726469 | 4.653658 | 61.123055 | 27.307547 | 47.51054 | 25.080757 | 20.823929 | 28.889193 | 5.9940248 | 5.2184176 | 5.5669647 | 4.539079 | 4.2773957 | 5.1444993 |
| ASMM10P007 0.04122 | 0.3747757 | 1.4966188 | up | NM_021431 | Nudt11 | nudix (nucleoside diphosphate linked moiety X)-type motif 11 | 589.73737 | 473.47338 | 9.441433 | 8.859726 | 619.6704 | 641.5723 | 507.96942 | 449.83777 | 442.54416 | 528.0382 | 9.249663 | 9.690073 | 9.3845625 | 8.741919 | 8.686471 | 9.150788 |
| ASMM10P002 0.04125 | 0.3747757 | 1.3662801 | up | NM_017368 | Celf1 | CUGBP, Elav-like family member 1 | 40.818274 | 36.896144 | 5.638923 | 5.18867 | 46.76416 | 39.649475 | 36.041187 | 44.570442 | 41.544937 | 24.573053 | 5.6108475 | 5.754242 | 5.5516806 | 5.382587 | 5.264964 | 4.9184594 |
| ASMM10P015 0.04127 | 0.3747757 | 1.3596216 | up | NM_207653 | Cflar | CASP8 and FADD-like apoptosis regulator | 3999.1565 | 3475.7462 | 12.150722 | 11.707517 | 3759.0566 | 4249.804 | 3988.609 | 3257.598 | 3888.5728 | 3281.0679 | 11.882149 | 12.303609 | 12.266409 | 11.583833 | 11.801472 | 11.737247 |
| ASMM10P033 0.04131 | 0.3748763 | 1.7600719 | up | NM_053096 | Cml2 | camello-like 2 | 27.904249 | 18.871274 | 5.052054 | 4.23642 | 35.60089 | 27.696829 | 20.415028 | 15.443789 | 25.14117 | 16.028864 | 5.2118006 | 5.239642 | 4.7047205 | 3.8265626 | 4.542477 | 4.3402205 |
| ASMM10P016 0.04138 | 0.3748763 | 1.504985 | up | NM_011765 | Zfp97 | zinc finger protein 97 | 38.89904 | 31.455299 | 5.579426 | 4.989677 | 40.87999 | 34.583874 | 41.233257 | 37.7363 | 26.95598 | 29.673618 | 5.417123 | 5.561 | 5.760155 | 5.140918 | 4.6458616 | 5.182251 |
| ASMM10P007 0.04139 | 0.3748763 | 1.4555489 | up | NM_013835 | Trove2 | TROVE domain family, member 2 | 487.10287 | 405.11559 | 9.182302 | 8.640739 | 459.11368 | 472.85526 | 529.33966 | 399.18402 | 454.76776 | 361.395 | 8.842401 | 9.267308 | 9.437197 | 8.566498 | 8.724294 | 8.631424 |
| ASMM10P007 0.04139 | 0.3748763 | 1.3397297 | up | NM_016804 | Mtx2 | metaxin 2 | 1219.9983 | 1094.5723 | 10.483671 | 10.061729 | 1330.8787 | 1112.69 | 1216.4263 | 1082.0101 | 1017.2957 | 1184.4111 | 10.365423 | 10.473589 | 10.612 | 10.00589 | 9.88184 | 10.297456 |

| | | | | | | | | | | | | | | | | | | | | | | | |
|---|---|---|---|---|---|---|---|---|---|---|---|---|---|---|---|---|---|---|---|---|---|---|---|
| ASMM10P032 | 0.04147 | 0.3750299 | 1.4164911 | up | NM_133925 | Rbm28 | RNA binding motif protein 28 | 627.03727 | 536.83642 | 9.530906 | 9.028584 | 629.82996 | 590.89636 | 660.3855 | 608.08685 | 584.2894 | 418.133 | 9.277801 | 9.573325 | 9.7415905 | 9.171361 | 9.083167 | 8.831224 |
| ASMM10P009 | 0.04153 | 0.3750299 | 1.5361474 | up | NM_178879 | B3gnt9 | UDP-GlcNAc:betaGal beta-1,3-N-acetylglucosaminyltransferase 9 | 111.22185 | 87.817438 | 7.068289 | 6.448972 | 140.34863 | 92.20542 | 101.11151 | 78.27619 | 85.75573 | 99.420395 | 7.1674075 | 6.953613 | 7.0838456 | 6.193166 | 6.311194 | 6.842556 |
| ASMM10P002 | 0.04154 | 0.3750299 | 1.2758557 | up | NM_001163 | Eme2 | essential meiotic endonuclease 1 homolog 2 (S. pombe) | 194.85588 | 187.93323 | 7.899198 | 7.547733 | 202.60817 | 192.92822 | 189.03125 | 190.44035 | 193.81706 | 179.54227 | 7.6881204 | 8.013531 | 7.995942 | 7.493371 | 7.493827 | 7.656 |
| ASMM10P004 | 0.04161 | 0.3750299 | 3.4115552 | up | NM_001081 | Cped1 | cadherin-like and PC-esterase domain containing 1 | 43.805712 | 15.773088 | 5.662786 | 3.892357 | 58.44349 | 24.22297 | 48.750675 | 12.778218 | 10.15694 | 24.384106 | 5.925804 | 5.0591507 | 6.003404 | 3.5455687 | 3.2236433 | 4.907858 |
| ASMM10P053 | 0.04162 | 0.3750299 | 1.342784 | up | NM_001159 | Dnajb2 | DnaJ (Hsp40) homolog, subfamily B, member 2 | 73.868293 | 67.417637 | 6.507817 | 6.082589 | 73.50473 | 72.27838 | 75.82177 | 71.668564 | 67.78371 | 62.800636 | 6.257869 | 6.606307 | 6.6592736 | 6.0703163 | 5.9691086 | 6.208343 |
| ASMM10P044 | 0.04167 | 0.3750299 | 1.389731 | up | NM_001098 | Yy2 | Yy2 transcription factor | 740.96957 | 642.35485 | 9.77225 | 9.297444 | 727.70636 | 772.72284 | 722.4795 | 595.74243 | 698.54803 | 632.7741 | 9.497932 | 9.94902 | 9.869798 | 9.142347 | 9.339076 | 9.41091 |
| ASMM10P037 | 0.04167 | 0.3750299 | 1.4832997 | up | NM_133770 | Adck4 | aarF domain containing kinase 4 | 111.12889 | 89.172273 | 7.04743 | 6.47862 | 149.89648 | 84.706696 | 98.78348 | 80.61423 | 94.26732 | 92.63527 | 7.261138 | 6.832565 | 7.048587 | 6.2396717 | 6.448921 | 6.747267 |
| ASMM10P006 | 0.04170 | 0.3750299 | 1.3620878 | up | NM_009418 | Tpp2 | tripeptidyl peptidase II | 999.43388 | 879.69506 | 10.197003 | 9.751184 | 1048.0024 | 900.80145 | 1049.4978 | 881.966 | 846.65967 | 910.4595 | 10.01142 | 10.167158 | 10.412432 | 9.70074 | 9.616927 | 9.935884 |
| ASMM10P026 | 0.04174 | 0.3750299 | 1.5914371 | up | NM_008256 | Hmgcs2 | 3-hydroxy-3-methylglutaryl-Coenzyme A synthase 2 | 20.843916 | 16.084836 | 4.663739 | 3.993409 | 20.029291 | 22.211166 | 20.29129 | 19.593103 | 18.529345 | 10.13206 | 4.357421 | 4.9374347 | 4.6963625 | 4.1795454 | 4.108765 | 3.6919174 |
| ASMM10P045 | 0.04177 | 0.3750299 | 1.6376647 | up | NM_001017 | Gm21943 | predicted gene, 21943 | 2400.5827 | 1688.5946 | 11.38891 | 10.67727 | 2929.3813 | 1583.8081 | 2688.5586 | 1707.8162 | 1626.346 | 1731.6215 | 11.502443 | 10.955597 | 11.708689 | 10.655563 | 10.53907 | 10.837176 |
| ASMM10P025 | 0.04184 | 0.3750524 | 1.252595 | up | NM_010763 | Man1a2 | mannosidase, alpha, class 1A, member 2 | 17463.174 | 16307.623 | 14.238911 | 13.913991 | 17121.432 | 17103.299 | 18164.791 | 16663.432 | 15820.041 | 16439.395 | 14.129369 | 14.235747 | 14.351616 | 13.900181 | 13.766567 | 14.075224 |
| ASMM10P005 | 0.04185 | 0.3750524 | 1.573514 | up | NM_001170 | Slc35b3 | solute carrier family 35, member B3 | 111.70851 | 87.326933 | 7.10391 | 6.44992 | 107.62233 | 105.19399 | 122.3092 | 84.36105 | 85.43694 | 92.18281 | 6.797144 | 7.145298 | 7.369287 | 6.3016253 | 6.3060217 | 6.742112 |
| ASMM10P004 | 0.04187 | 0.3750524 | 1.2919757 | up | NM_001114 | Eml4 | echinoderm microtubule associated protein like 4 | 96.571072 | 92.66676 | 6.88064 | 6.511061 | 113.24322 | 82.96715 | 93.502846 | 107.1921 | 104.57703 | 66.23115 | 6.872163 | 6.802328 | 6.967429 | 6.6572747 | 6.5932217 | 6.2826867 |
| ASMM10P014 | 0.04194 | 0.3750524 | 2.6782254 | up | NM_001111 | Runx1 | runt related transcription factor 1 | 37.140533 | 13.838736 | 5.245118 | 3.823841 | 70.16926 | 19.80742 | 21.44492 | 14.01759 | 15.18218 | 12.316439 | 6.1937447 | 4.7657886 | 4.7758207 | 3.6847057 | 3.8122778 | 3.9745383 |
| ASMM10P013 | 0.04196 | 0.3750524 | 1.2971656 | up | NM_026002 | Mtdh | metadherin | 3365.8009 | 3068.8715 | 11.907946 | 11.532584 | 3253.421 | 3508.5007 | 3335.481 | 3080.7676 | 3205.2224 | 2920.6245 | 11.659215 | 12.050707 | 12.013917 | 11.493271 | 11.526381 | 11.578099 |
| ASMM10P029 | 0.04202 | 0.3750524 | 1.3613759 | up | NM_001038 | Ccdc23 | coiled-coil domain containing 23 | 101.12621 | 91.909272 | 6.959945 | 6.514879 | 98.79477 | 96.98816 | 107.59569 | 99.90303 | 100.84728 | 74.97751 | 6.677461 | 7.0266657 | 7.175707 | 6.551939 | 6.5400186 | 6.4526796 |
| ASMM10P008 | 0.04204 | 0.3750524 | 1.890641 | up | NM_029801 | Tsacc | TSSK6 activating co-chaperone | 71.402177 | 46.057427 | 6.442263 | 5.523387 | 72.52512 | 82.19244 | 59.48897 | 42.371468 | 40.330353 | 55.47046 | 6.2401805 | 6.787585 | 6.299022 | 5.3092027 | 5.2222214 | 6.038737 |
| ASMM10P038 | 0.04204 | 0.3750524 | 1.5246207 | up | NM_027363 | Chp2 | calcineurin-like EF hand protein 2 | 98.30393 | 78.901833 | 6.905601 | 6.297151 | 93.24342 | 116.07948 | 85.58889 | 82.18793 | 90.19106 | 64.32651 | 6.596502 | 7.2852354 | 6.8350654 | 6.2662272 | 6.381348 | 6.2438765 |
| ASMM10P052 | 0.04212 | 0.3750524 | 1.6519819 | up | NM_025415 | Cks2 | CDC28 protein kinase regulatory subunit 2 | 60.72927 | 44.092013 | 6.206235 | 5.482037 | 58.67315 | 48.43164 | 75.08302 | 43.320683 | 45.904346 | 43.05101 | 5.9324017 | 6.04265 | 6.643654 | 5.345479 | 5.406411 | 5.694222 |
| ASMM10P009 | 0.04214 | 0.3750524 | 1.3737146 | up | NM_153581 | Gpm6a | glycoprotein m6a | 3003.9037 | 2586.6632 | 11.742813 | 11.28473 | 3037.3608 | 2743.2854 | 3231.065 | 2685.5142 | 2416.4001 | 2658.0754 | 11.5513 | 11.708176 | 11.968962 | 11.299377 | 11.11048 | 11.444334 |
| ASMM10P033 | 0.04218 | 0.3750524 | 1.4227778 | up | NM_001113 | Sema4f | sema domain, immunoglobulin domain (Ig), TM domain, and short cytoplasmic domain | 170.50101 | 148.22073 | 7.703789 | 7.195078 | 175.65149 | 187.53455 | 148.317 | 140.6474 | 179.305 | 124.70979 | 7.4836264 | 7.9727345 | 7.655005 | 7.04922 | 7.377423 | 7.1585917 |
| ASMM10P054 | 0.04223 | 0.3750524 | 1.5484352 | up | NM_009393 | Tnnc1 | troponin C, cardiac/slow skeletal | 1237.6453 | 934.38621 | 10.463852 | 9.833041 | 1541.5176 | 862.7308 | 1308.6874 | 874.68414 | 948.05273 | 980.42175 | 10.574337 | 10.098611 | 10.718607 | 9.688197 | 9.776009 | 10.034916 |
| ASMM10P031 | 0.04234 | 0.3750524 | 1.4622101 | up | NM_011738 | Ywhah | tyrosine 3-monooxygenase/tryptophan 5-monooxygenase activation protein, eta polypeptide | 664.51409 | 548.02237 | 9.616301 | 9.06815 | 670.93353 | 632.6704 | 689.93835 | 573.31915 | 487.6708 | 583.07715 | 9.37602 | 9.667188 | 9.805694 | 9.089241 | 8.821457 | 9.293752 |
| ASMM10P014 | 0.04236 | 0.3750524 | 1.3456218 | up | NM_183390 | Klhl6 | kelch-like 6 | 1062.6014 | 949.11043 | 10.282061 | 9.853788 | 1120.731 | 911.805 | 1155.2683 | 952.2678 | 1073.8422 | 821.2213 | 10.110929 | 10.18616 | 10.549093 | 9.818216 | 9.959789 | 9.783358 |
| ASMM10P016 | 0.04240 | 0.3750524 | 1.3322775 | up | NM_144802 | Hnrnpll | heterogeneous nuclear ribonucleoprotein L-like | 11510.676 | 10077.251 | 13.638541 | 13.224646 | 10341.336 | 12732.387 | 11458.306 | 10103.5 | 10448.336 | 9679.917 | 13.37907 | 13.837035 | 13.699517 | 13.190439 | 13.183679 | 13.29982 |
| ASMM10P014 | 0.04248 | 0.3750524 | 1.3256635 | up | NM_025708 | Tmem186 | transmembrane protein 186 | 1756.7584 | 1584.2356 | 10.99354 | 10.586825 | 1798.8243 | 1539.7537 | 1931.6971 | 1616.0996 | 1626.2782 | 1510.3289 | 10.812497 | 10.911195 | 11.256927 | 10.574756 | 10.538804 | 10.646915 |
| ASMM10P000 | 0.04250 | 0.3750524 | 1.2771751 | up | NM_011351 | Sema6c | sema domain, transmembrane domain (TM), and cytoplasmic domain, (semaphorin) 6C | 257.97638 | 245.92798 | 8.284622 | 7.931665 | 301.49088 | 227.28477 | 245.15349 | 246.02861 | 236.89993 | 254.8554 | 8.2457285 | 8.238615 | 8.369521 | 7.861974 | 7.7814326 | 8.151589 |
| ASMM10P011 | 0.04250 | 0.3750524 | 1.3177419 | up | NM_021504 | Ngly1 | N-glycanase 1 | 273.70462 | 253.51498 | 8.369958 | 7.97189 | 318.8962 | 256.72717 | 245.49048 | 244.83946 | 243.88191 | 271.82358 | 8.3273 | 8.411036 | 8.371538 | 7.855959 | 7.8218565 | 8.237855 |
| ASMM10P024 | 0.04256 | 0.3750524 | 1.5281738 | up | NM_021280 | Plcg1 | phospholipase C, gamma 1 | 167.62163 | 138.63325 | 7.683187 | 7.071379 | 172.76506 | 159.37065 | 170.72919 | 145.27666 | 177.0433 | 93.5798 | 7.462357 | 7.7315254 | 7.85568 | 7.0958 | 7.3581676 | 6.760169 |
| ASMM10P011 | 0.04258 | 0.3750524 | 1.6354886 | up | NM_011825 | Grem2 | gremlin 2 homolog, cysteine knot superfamily (Xenopus laevis) | 9167.4423 | 6595.2381 | 13.323794 | 12.614072 | 8656.173 | 9315.018 | 9531.136 | 6963.645 | 5243.7817 | 7578.2876 | 13.104461 | 13.398166 | 13.468754 | 12.662596 | 12.220651 | 12.958969 |
| ASMM10P020 | 0.04260 | 0.3750524 | 1.4093862 | up | NM_009261 | Strbp | spermatid perinuclear RNA binding protein | 45258.041 | 38658.281 | 15.658615 | 15.163548 | 44720.402 | 38260.066 | 52793.656 | 38859.797 | 41186.9 | 35928.145 | 15.595149 | 15.406883 | 15.973813 | 15.14149 | 15.134304 | 15.21485 |
| ASMM10P053 | 0.04273 | 0.3750524 | 1.415385 | up | NM_001161 | Pex7 | peroxisomal biogenesis factor 7 | 60.727261 | 50.42048 | 6.173732 | 5.672537 | 82.946556 | 43.727646 | 55.50758 | 51.90721 | 52.249832 | 47.104397 | 6.4286017 | 5.8950644 | 6.19753 | 5.5989027 | 5.5997763 | 5.8189335 |
| ASMM10P000 | 0.04275 | 0.3750524 | 1.5440283 | up | NM_010486 | Elavl2 | ELAV (embryonic lethal, abnormal vision, Drosophila)-like 2 (Hu antigen B) | 325.24691 | 251.5313 | 8.594282 | 7.967583 | 355.9429 | 238.72774 | 381.0701 | 256.6386 | 256.98425 | 240.97104 | 8.482917 | 8.305958 | 8.993971 | 7.9271803 | 7.896969 | 8.078599 |
| ASMM10P023 | 0.04293 | 0.3750524 | 1.7376542 | up | NM_011354 | Serf2 | small EDRK-rich factor 2 | 87.195998 | 60.553483 | 6.725835 | 5.928694 | 77.99861 | 72.48007 | 111.10931 | 66.14408 | 59.57078 | 55.94559 | 6.3439093 | 6.6097126 | 7.2238817 | 5.9549103 | 5.7804613 | 6.050709 |
| ASMM10P051 | 0.04295 | 0.3750524 | 1.4968815 | up | NM_201358 | Lyrm4 | LYR motif containing 4 | 623.46076 | 502.17225 | 9.523689 | 8.941729 | 633.76746 | 629.3984 | 607.21643 | 473.58475 | 463.735 | 569.197 | 9.288564 | 9.660737 | 9.621765 | 8.813722 | 8.751819 | 9.259645 |
| ASMM10P004 | 0.04300 | 0.3750524 | 1.2734966 | up | NM_001080 | Sp2 | Sp2 transcription factor | 53.362056 | 50.089316 | 6.011243 | 5.662447 | 68.37857 | 46.184547 | 45.523052 | 52.818237 | 49.59501 | 47.854702 | 6.1540737 | 5.974493 | 5.905161 | 5.623659 | 5.523691 | 5.8399925 |
| ASMM10P026 | 0.04303 | 0.3750524 | 1.5990342 | up | NM_026677 | Rab13 | RAB13, member RAS oncogene family | 46.375331 | 33.961779 | 5.779993 | 5.102792 | 57.232822 | 50.62431 | 31.268862 | 34.59484 | 38.940094 | 28.350403 | 5.896393 | 6.101845 | 5.341741 | 5.017166 | 5.1721873 | 5.1190233 |
| ASMM10P023 | 0.04305 | 0.3750524 | 1.2863871 | up | NM_024181 | Dnajc10 | DnaJ (Hsp40) homolog, subfamily C, member 10 | 1560.0327 | 1456.8829 | 10.828662 | 10.465337 | 1590.5134 | 1650.9047 | 1438.68 | 1446.0846 | 1493.6123 | 1430.9519 | 10.622499 | 11.017304 | 10.846183 | 10.414331 | 10.41907 | 10.562611 |
| ASMM10P030 | 0.04306 | 0.3750524 | 1.4497971 | up | NM_001005 | Ado | 2-aminoethanethiol (cysteamine) dioxygenase | 13568.204 | 10862.429 | 13.864236 | 13.328385 | 11864.046 | 12957.254 | 15883.311 | 10272.02 | 11451.914 | 10863.353 | 13.573967 | 13.860089 | 14.158652 | 13.216236 | 13.308159 | 13.46076 |
| ASMM10P045 | 0.04307 | 0.3750524 | 1.4908471 | up | NM_001039 | Ccdc108 | coiled-coil domain containing 108 | 806.7036 | 641.23248 | 9.864473 | 9.288341 | 1026.4608 | 636.0801 | 757.5699 | 537.5781 | 669.57513 | 716.5442 | 9.979143 | 9.674388 | 9.939888 | 8.996123 | 9.276534 | 9.592365 |
| ASMM10P018 | 0.04320 | 0.3750524 | 1.3212636 | up | NM_053146 | Pcdhb21 | protocadherin beta 21 | 5984.1648 | 5307.89 | 12.717436 | 12.315518 | 5996.844 | 5124.4487 | 6831.2017 | 5466.916 | 5594.3125 | 4862.4414 | 12.5801 | 12.579604 | 12.992605 | 12.319164 | 12.312797 | 12.314593 |
| ASMM10P032 | 0.04328 | 0.3750524 | 1.2905564 | up | NM_023142 | Arpc1b | actin related protein 2/3 complex, subunit 1B | 4062.1838 | 3743.2546 | 12.173208 | 11.805214 | 4049.744 | 4392.7544 | 3744.053 | 3618.7021 | 4336.979 | 3274.0828 | 11.992301 | 12.352774 | 12.174548 | 11.732589 | 11.947496 | 11.735559 |
| ASMM10P040 | 0.04330 | 0.3750524 | 1.4551579 | up | NM_030110 | Micu3 | mitochondrial calcium uptake family, member 3 | 456.53053 | 382.91271 | 9.09719 | 8.556014 | 471.94656 | 423.04144 | 474.6036 | 367.59677 | 361.1199 | 420.02145 | 8.883468 | 9.111987 | 9.296114 | 8.443581 | 8.386577 | 8.837884 |
| ASMM10P005 | 0.04331 | 0.3750524 | 1.3389962 | up | NM_001163 | Frmd3 | FERM domain containing 3 | 362.72288 | 323.811 | 8.739256 | 8.318104 | 482.81195 | 305.37747 | 299.97922 | 344.56805 | 297.38388 | 329.48108 | 8.913832 | 8.648372 | 8.655564 | 8.3512745 | 8.104589 | 8.498449 |
| ASMM10P042 | 0.04334 | 0.3750524 | 1.2795957 | up | NM_133986 | Tcta | T cell leukemia translocation altered gene | 10460.497 | 9508.8017 | 13.502753 | 13.147065 | 11489.555 | 9061.46 | 10830.477 | 9113.584 | 9378.868 | 10033.953 | 13.526014 | 13.362178 | 13.620066 | 13.053826 | 13.04867 | 13.338698 |
| ASMM10P037 | 0.04334 | 0.3750524 | 1.3346224 | up | NM_001136 | Ate1 | arginyltransferase 1 | 174.46417 | 157.66909 | 7.708622 | 7.29219 | 224.61617 | 128.5498 | 170.22653 | 159.90208 | 166.44151 | 146.66367 | 7.8386817 | 7.4360847 | 7.8510985 | 7.237292 | 7.2639194 | 7.3753586 |
| ASMM10P021 | 0.04336 | 0.3750524 | 1.4613914 | up | NM_172858 | Pak7 | p21 protein (Cdc42/Rac)-activated kinase 7 | 5830.3857 | 4615.8716 | 12.664327 | 12.116984 | 6731.334 | 4352.695 | 6407.128 | 4417.7163 | 4691.7905 | 4738.108 | 12.750311 | 12.337591 | 12.905079 | 12.013464 | 12.061422 | 12.276067 |
| ASMM10P038 | 0.04343 | 0.3750524 | 1.4791609 | up | NM_027347 | Med23 | mediator complex subunit 23 | 723.6891 | 588.33203 | 9.733789 | 9.16901 | 773.9303 | 604.2246 | 792.9124 | 557.0018 | 557.0541 | 650.9402 | 9.588693 | 9.608289 | 10.004385 | 9.048695 | 9.013073 | 9.445262 |
| ASMM10P009 | 0.04353 | 0.3750524 | 1.3874996 | up | NM_198645 | Ccdc58 | coiled-coil domain containing 58 | 462.18902 | 406.06875 | 9.113742 | 8.641254 | 481.48257 | 464.1312 | 440.95328 | 390.85968 | 389.89896 | 437.4476 | 8.909382 | 9.238071 | 9.193772 | 8.532352 | 8.50028 | 8.891131 |
| ASMM10P007 | 0.04364 | 0.3750524 | 2.0886218 | up | NM_024412 | Clcnka | chloride channel Ka | 36.128199 | 19.828786 | 5.374994 | 4.312442 | 54.850266 | 28.426292 | 25.108038 | 24.146519 | 14.869291 | 20.470549 | 5.8359694 | 5.280961 | 5.0080504 | 4.4830494 | 3.7799895 | 4.674288 |
| ASMM10P052 | 0.04371 | 0.3750524 | 1.7712328 | up | NM_177322 | Agtr1a | angiotensin II receptor, type 1a | 270.38001 | 187.94376 | 8.355434 | 7.53068 | 241.87967 | 293.96664 | 275.29373 | 151.58562 | 212.64381 | 199.60184 | 7.9422693 | 8.59534 | 8.528692 | 7.157949 | 7.629017 | 7.8050737 |
| ASMM10P011 | 0.04371 | 0.3750524 | 1.5069597 | up | NM_017388 | Ear3 | eosinophil-associated, ribonuclease A family, member 3 | 27.856689 | 21.880058 | 5.069147 | 4.477506 | 30.743362 | 20.60229 | 32.224415 | 25.901718 | 20.308836 | 19.429619 | 4.99645 | 4.826866 | 5.3841243 | 4.587169 | 4.2397733 | 4.6055756 |
| ASMM10P006 | 0.04382 | 0.3750524 | 1.7609538 | up | NM_009376 | Ift88 | intraflagellar transport 88 | 124.41068 | 86.684858 | 7.248476 | 6.432119 | 103.24718 | 120.11319 | 149.87167 | 101.21848 | 87.02681 | 71.80929 | 6.740636 | 7.3357387 | 7.6690545 | 6.5709124 | 6.331726 | 6.3937197 |
| ASMM10P029 | 0.04391 | 0.3750524 | 1.2873504 | up | NM_181585 | Pik3r3 | phosphatidylinositol 3 kinase, regulatory subunit, polypeptide 3 (p55) | 138.71741 | 133.45894 | 7.414851 | 7.050446 | 149.88133 | 126.6404 | 139.6305 | 124.71211 | 152.48509 | 123.17961 | 7.2609615 | 7.415064 | 7.568528 | 6.8732176 | 7.1368766 | 7.141245 |
| ASMM10P034 | 0.04392 | 0.3750524 | 1.3338417 | up | NM_001127 | Pparg | peroxisome proliferator activated receptor gamma | 67.073432 | 61.960027 | 6.364699 | 5.949111 | 68.229996 | 71.286064 | 61.704235 | 71.66738 | 66.65929 | 47.55341 | 6.1518717 | 6.586619 | 6.355605 | 6.0701804 | 5.944404 | 5.832749 |
| ASMM10P012 | 0.04401 | 0.3750524 | 1.370577 | up | NM_175498 | Pnma2 | paraneoplastic antigen MA2 | 22761.273 | 19399.077 | 14.622497 | 14.167713 | 20968.154 | 23346.422 | 23969.244 | 19787.107 | 17994.83 | 20415.295 | 14.447813 | 14.676268 | 14.743409 | 14.154931 | 13.951581 | 14.396628 |
| ASMM10P042 | 0.04403 | 0.3750524 | 1.849057 | up | NM_029247 | 4930550C14 | RIKEN cDNA 4930550C14 gene | 66.00513 | 44.071882 | 6.339627 | 5.452837 | 65.17766 | 57.58248 | 75.25525 | 33.02338 | 52.83605 | 46.356216 | 6.0872607 | 6.2844954 | 6.647124 | 4.9477925 | 5.614799 | 5.7959194 |
| ASMM10P010 | 0.04421 | 0.3750524 | 1.5501146 | up | NM_008329 | Ifi204 | interferon activated gene 204 | 138.91179 | 111.98728 | 7.416278 | 6.783903 | 148.5522 | 131.0806 | 137.10258 | 89.84641 | 130.98051 | 115.13493 | 7.2483582 | 7.462778 | 7.5376987 | 6.3900714 | 6.917701 | 7.043938 |
| ASMM10P013 | 0.04423 | 0.3750524 | 1.528593 | up | NM_030720 | Gpr84 | G protein-coupled receptor 84 | 728.76993 | 567.3024 | 9.720102 | 9.107897 | 865.14307 | 510.62503 | 810.5417 | 603.91595 | 648.776 | 449.21524 | 9.746662 | 9.378727 | 10.034916 | 9.160606 | 9.235375 | 8.927711 |
| ASMM10P047 | 0.04428 | 0.3750524 | 1.6557996 | up | NM_001195 | Kat7 | K(lysine) acetyltransferase 7 | 179.17647 | 133.47458 | 7.766386 | 7.038858 | 194.0813 | 145.19621 | 198.25189 | 107.6973 | 148.4687 | 144.25774 | 7.6307597 | 7.6030855 | 8.065314 | 6.6631417 | 7.0994134 | 7.35402 |
| ASMM10P008 | 0.04432 | 0.3750524 | 1.3519936 | up | NM_026392 | Tmem70 | transmembrane protein 70 | 1408.0498 | 1255.826 | 10.684165 | 10.249077 | 1540.1099 | 1334.6855 | 1349.354 | 1239.8939 | 1139.2681 | 1388.316 | 10.573086 | 10.720981 | 10.758429 | 10.195494 | 10.03926 | 10.512477 |
| ASMM10P036 | 0.04436 | 0.3750524 | 1.2542259 | up | NM_146193 | Btbd1 | BTB (POZ) domain containing 1 | 43192.875 | 41069.356 | 15.585766 | 15.258969 | 43060.496 | 40690.938 | 45827.19 | 39317.61 | 42402.184 | 41488.273 | 15.534709 | 15.504787 | 15.717803 | 15.155029 | 15.181187 | 15.440691 |
| ASMM10P041 | 0.04436 | 0.3750524 | 1.5365815 | up | NM_010242 | Fut4 | fucosyltransferase 4 | 55.574762 | 43.869878 | 6.089596 | 5.469872 | 58.528408 | 57.874847 | 50.32103 | 42.54828 | 41.280792 | 47.780563 | 5.9274755 | 6.2916074 | 6.0497046 | 5.31523 | 5.2552414 | 5.8391433 |
| ASMM10P014 | 0.04437 | 0.3750524 | 1.2849979 | up | NM_139145 | Hlcs | holocarboxylase synthetase (biotin- [propriony-Coenzyme A-carboxylase (ATP-hydrolysing)] | 4329.7277 | 3971.8633 | 12.26928 | 11.907514 | 4449.618 | 4074.5015 | 4465.0635 | 3778.586 | 3934.0566 | 4202.9473 | 12.14703 | 12.247448 | 12.413362 | 11.801065 | 11.818334 | 12.103143 |
| ASMM10P004 | 0.04439 | 0.3750524 | 1.3706257 | up | NM_001081 | Cartpt | CART prepropeptide | 1347.7384 | 1162.0102 | 10.59956 | 10.144725 | 1558.2819 | 1471.6512 | 1013.2822 | 1085.4302 | 1250.0162 | 1150.5842 | 10.590476 | 10.845006 | 10.363197 | 10.007926 | 10.168603 | 10.257646 |
| ASMM10P000 | 0.04442 | 0.3750524 | 2.4381109 | up | NM_178886 | Ldlrad3 | low density lipoprotein receptor class A domain containing 3 | 110.2649 | 54.515996 | 7.027172 | 5.741408 | 74.447655 | 101.2916 | 155.05545 | 54.857735 | 70.679474 | 38.01078 | 6.2749047 | 7.089432 | 7.7171783 | 5.679312 | 6.0306215 | 5.5142903 |
| ASMM10P020 | 0.04446 | 0.3750524 | 1.2836904 | up | NM_001190 | Gpr155 | G protein-coupled receptor 155 | 19635.131 | 17742.846 | 14.400326 | 14.040028 | 17431.613 | 21269.734 | 20204.047 | 17855.615 | 18609.727 | 16763.195 | 14.160007 | 14.543257 | 14.497713 | 14.011051 | 14.00878 | 14.100254 |
| ASMM10P052 | 0.04446 | 0.3750524 | 1.5384005 | up | NM_183310 | Ccser1 | coiled-coil serine rich 1 | 31.558784 | 24.770885 | 5.262069 | 4.640638 | 35.903458 | 31.440758 | 27.332136 | 31.715576 | 20.428446 | 22.168634 | 5.224076 | 5.4218473 | 5.140285 | 4.889566 | 4.249488 | 4.782861 |
| ASMM10P013 | 0.04453 | 0.3751845 | 1.5828636 | up | NM_009267 | Spt1 | salivary protein 1 | 183.9047 | 139.42151 | 7.77596 | 7.113423 | 218.13742 | 123.19052 | 210.38617 | 153.99522 | 144.83199 | 119.43731 | 7.7979274 | 7.373431 | 8.156521 | 7.182818 | 7.061429 | 7.0960217 |
| ASMM10P043 | 0.04453 | 0.3751845 | 1.2758685 | up | NM_025975 | Dynlt3 | dynein light chain Tctex-type 3 | 22991.521 | 21007.809 | 14.641029 | 14.289549 | 24224.021 | 20109.205 | 24641.336 | 20462.582 | 21492.309 | 21068.537 | 14.655712 | 14.473791 | 14.793584 | 14.203714 | 14.218026 | 14.446908 |
| ASMM10P031 | 0.04455 | 0.3751845 | 1.8948517 | up | NM_009903 | Cldn4 | claudin 4 | 21.291424 | 13.08857 | 4.648045 | 3.72596 | 18.26373 | 15.797899 | 29.812643 | 14.781839 | 14.456571 | 10.027301 | 4.2285156 | 4.44389 | 5.27173 | 3.7626827 | 3.7389402 | 3.6762578 |
| ASMM10P045 | 0.04456 | 0.3751845 | 1.2925795 | up | NM_011515 | Vamp7 | vesicle-associated membrane protein 7 | 18511.851 | 16819.084 | 14.32694 | 13.956687 | 17758.375 | 19387.277 | 18389.9 | 16373.064 | 16517.932 | 17566.256 | 14.183426 | 14.421164 | 14.376229 | 13.877212 | 13.827466 | 14.165382 |
| ASMM10P047 | 0.04458 | 0.375269 | 1.2694451 | up | NM_177564 | Dhrs11 | dehydrogenase/reductase (SDR family) member 11 | 10869.485 | 10038.629 | 13.555214 | 13.211016 | 11702.431 | 9340.428 | 11565.596 | 9692.877 | 11914.124 | 8508.885 | 13.556559 | 13.40084 | 13.708244 | 13.138313 | 13.369753 | 13.124983 |
| ASMM10P036 | 0.04463 | 0.375491 | 1.3637287 | up | NM_028343 | Tmem135 | transmembrane protein 135 | 424.38884 | 379.95229 | 8.997577 | 8.55002 | 439.3106 | 430.13644 | 403.71948 | 356.07516 | 386.3972 | 397.3845 | 8.782545 | 9.134929 | 9.075257 | 8.397879 | 8.487921 | 8.764261 |
| ASMM10P038 | 0.04466 | 0.3756522 | 1.9813494 | up | NM_177249 | Usp47 | ubiquitin specific peptidase 47 | 456.35756 | 271.4694 | 9.042188 | 8.055704 | 482.92813 | 306.61298 | 579.53156 | 281.838 | 221.86292 | 310.70728 | 8.914406 | 8.653168 | 9.558989 | 8.061171 | 7.688696 | 8.417246 |
| ASMM10P053 | 0.04486 | 0.3758018 | 1.3301763 | up | NM_001142 | Pcdh15 | protocadherin 15 | 3122.8042 | 2797.966 | 11.798558 | 11.38694 | 3167.3105 | 2853.0317 | 3348.0703 | 3018.2485 | 3089.2327 | 2286.4167 | 11.617054 | 11.76097 | 12.017649 | 11.461752 | 11.475079 | 11.22399 |
| ASMM10P050 | 0.04498 | 0.3758018 | 1.3405123 | up | NM_178744 | Zbtb1 | zinc finger and BTB domain containing 1 | 82.295575 | 75.436978 | 6.660786 | 6.238001 | 84.861465 | 74.6021 | 87.42316 | 71.277534 | 87.762695 | 67.270706 | 6.461495 | 6.6537256 | 6.867136 | 6.062458 | 6.3439093 | 6.307636 |
| ASMM10P025 | 0.04511 | 0.3763361 | 1.6352233 | up | NM_028943 | Sgms2 | sphingomyelin synthase 2 | 1076.6978 | 787.37359 | 10.283678 | 9.57419 | 1201.575 | 835.1655 | 1193.353 | 618.40063 | 924.9646 | 818.75555 | 10.206525 | 10.056593 | 10.587915 | 9.196442 | 9.746662 | 9.779466 |
| ASMM10P000 | 0.04515 | 0.3763361 | 1.415477 | up | ENSMUST( | Prl2c1 | Prolactin family 2, subfamily c, member 1 | 311.93017 | 264.70769 | 8.539277 | 8.037989 | 367.9899 | 232.79411 | 335.0065 | 284.64594 | 255.75119 | 253.72595 | 8.533708 | 8.271337 | 8.812786 | 8.078364 | 7.88916 | 8.146442 |
| ASMM10P033 | 0.04519 | 0.3763361 | 1.4271355 | up | NM_028335 | Zfp248 | zinc finger protein 248 | 6513.5474 | 5340.5342 | 12.840635 | 12.327512 | 6100.7056 | 6137.3525 | 7302.584 | 5223.5845 | 5120.02 | 5677.998 | 12.60393 | 12.826499 | 13.091476 | 12.249974 | 12.189053 | 12.54351 |
| ASMM10P019 | 0.04525 | 0.3764826 | 2.2833969 | up | NM_011416 | Smarca2 | SWI/SNF related, matrix associated, actin dependent regulator of chromatin, subfamily a, me | 26.897025 | 13.9831 | 4.993212 | 3.802031 | 20.51315 | 22.908329 | 37.269596 | 19.061272 | 11.251431 | 11.636598 | 4.39563 | 4.9815025 | 5.602504 | 4.1376777 | 3.3767157 | 3.8916981 |
| ASMM10P053 | 0.04530 | 0.3765184 | 1.62005 | up | NM_021350 | Chml | choroideremia-like | 25.459796 | 18.785354 | 4.948919 | 4.25288 | 25.689093 | 28.487286 | 22.20301 | 16.331596 | 22.523216 | 17.50125 | 4.732147 | 5.2846246 | 4.8299847 | 3.9111795 | 4.384973 | 4.4624887 |
| ASMM10P002 | 0.04550 | 0.3771497 | 1.2558893 | up | NM_031251 | Ctns | cystinosis, nephropathic | 47.706711 | 46.198493 | 5.851439 | 5.52273 | 60.755013 | 39.854584 | 42.510536 | 49.859756 | 55.046463 | 33.68926 | 5.9856763 | 5.7614155 | 5.8072257 | 5.5418587 | 5.6714725 | 5.3548584 |
| ASMM10P022 | 0.04554 | 0.3771497 | 1.9687115 | up | NM_029309 | Morn5 | MORN repeat containing 5 | 198.90571 | 128.01142 | 7.92113 | 6.943878 | 217.54192 | 211.28252 | 167.89268 | 88.87897 | 135.4169 | 159.73839 | 7.7946196 | 8.1385145 | 7.830256 | 6.375606 | 6.962967 | 7.493062 |
| ASMM10P047 | 0.04556 | 0.3771497 | 1.6841484 | up | NM_207225 | Hdac4 | histone deacetylase 4 | 57.618267 | 41.789986 | 6.137179 | 5.385159 | 65.04075 | 46.04011 | 61.77394 | 49.87985 | 32.52693 | 42.963177 | 6.0833282 | 5.971159 | 6.3570485 | 5.5425224 | 4.921382 | 5.6915736 |

| | | | | | | | | | | | | | | | | | | | | | | | |
|---|---|---|---|---|---|---|---|---|---|---|---|---|---|---|---|---|---|---|---|---|---|---|---|
| ASMM10P041 | 0.04561 | 0.3771497 | 1.3586063 | up | NM_008408 | Stt3a | STT3, subunit of the oligosaccharyltransferase complex, homolog A (S. cerevisiae) | 2125.4199 | 1868.1392 | 11.264795 | 10.822668 | 2018.6387 | 2272.0537 | 2085.5674 | 1940.0817 | 1882.4602 | 1781.8757 | 10.972925 | 11.462145 | 11.359316 | 10.838805 | 10.746612 | 10.882587 |
| ASMM10P053 | 0.04563 | 0.3771497 | 2.7745792 | up | NM_001007 | Tceanc | transcription elongation factor A (SII) N-terminal and central domain containing | 17.003585 | 6.762744 | 4.290931 | 2.818662 | 13.524159 | 24.971846 | 12.514749 | 4.9999995 | 7.1583047 | 8.129928 | 3.7751245 | 5.099753 | 3.9979148 | 2.3250442 | 2.7405214 | 3.3904197 |
| ASMM10P050 | 0.04573 | 0.3771497 | 1.3253014 | up | NM_134042 | Aldh6a1 | aldehyde dehydrogenase family 6, subfamily A1 | 22045.733 | 19581.498 | 14.583593 | 14.177273 | 21373.172 | 20191.19 | 24572.838 | 17596.531 | 22167.145 | 18980.818 | 14.476174 | 14.481899 | 14.792707 | 13.986067 | 14.260641 | 14.28511 |
| ASMM10P046 | 0.04574 | 0.3771497 | 1.3544278 | up | NM_023907 | Foxi1 | forkhead box I1 | 149.539 | 136.0373 | 7.520563 | 7.082879 | 150.32938 | 152.63062 | 145.65701 | 147.62062 | 131.49261 | 128.99867 | 7.266216 | 7.6690545 | 7.6264186 | 7.119333 | 6.9229145 | 7.206391 |
| ASMM10P038 | 0.04576 | 0.3771497 | 1.3099643 | up | NM_181517 | Ipo7 | importin 7 | 155.58494 | 146.24248 | 7.573526 | 7.183999 | 164.1837 | 133.87321 | 168.6979 | 153.27522 | 155.88724 | 129.56497 | 7.3884435 | 7.495125 | 7.8370104 | 7.1739573 | 7.166314 | 7.211725 |
| ASMM10P019 | 0.04578 | 0.3771497 | 1.3415576 | up | NM_020032 | Poll | polymerase (DNA directed), lambda | 4664.7073 | 4077.6008 | 12.365668 | 11.941759 | 5213.963 | 3916.946 | 4863.213 | 3748.3347 | 4111.1772 | 4373.2905 | 12.372374 | 12.193664 | 12.530967 | 11.787951 | 11.878876 | 12.158451 |
| ASMM10P032 | 0.04586 | 0.3771497 | 1.5857458 | up | NM_054073 | Tsga13 | testis specific gene A13 | 95.836832 | 72.158675 | 6.842454 | 6.177293 | 117.51588 | 66.338455 | 103.65616 | 67.42617 | 74.65118 | 74.398674 | 6.9209805 | 6.484844 | 7.121539 | 5.9827733 | 6.1079426 | 6.441163 |
| ASMM10P010 | 0.04591 | 0.3771497 | 1.6207208 | up | NM_001164 | Fcrl6 | Fc receptor-like 6 | 29.471976 | 21.305928 | 5.138726 | 4.44209 | 34.30754 | 32.353397 | 21.754992 | 20.22386 | 22.38733 | 21.306593 | 5.1576624 | 5.4632435 | 4.7952724 | 4.224004 | 4.37661 | 4.7256575 |
| ASMM10P025 | 0.04591 | 0.3771497 | 1.6023082 | up | NM_001013 | Ttf2 | transcription termination factor, RNA polymerase II | 951.64417 | 714.19758 | 10.126447 | 9.446295 | 923.3805 | 855.0858 | 1076.4662 | 614.6106 | 736.4952 | 791.48694 | 9.841332 | 10.086112 | 10.451896 | 9.188332 | 9.416215 | 9.734338 |
| ASMM10P028 | 0.04601 | 0.3771497 | 1.5584303 | up | NM_145078 | 2610305D13 | RIKEN cDNA 2610305D13 gene | 335.78548 | 265.01482 | 8.662852 | 8.022758 | 321.96072 | 314.99118 | 370.40454 | 231.35933 | 326.5813 | 237.10384 | 8.341438 | 8.692454 | 8.954663 | 7.773405 | 8.243052 | 8.051817 |
| ASMM10P039 | 0.04605 | 0.3771497 | 1.5826864 | up | NM_029751 | Rpl18a | ribosomal protein L18A | 114.8758 | 85.725138 | 7.084471 | 6.422095 | 149.47581 | 117.07999 | 78.07158 | 79.319214 | 90.70064 | 87.15556 | 7.256716 | 7.295746 | 6.7009506 | 6.2142725 | 6.3913884 | 6.6606255 |
| ASMM10P022 | 0.04608 | 0.3771497 | 1.2875651 | up | NM_020575 | March7 | membrane-associated ring finger (C3HC4) 7 | 2843.3839 | 2615.0113 | 11.667369 | 11.302724 | 2797.1655 | 2717.0789 | 3015.9072 | 2683.1045 | 2714.3413 | 2447.588 | 11.434331 | 11.695756 | 11.87202 | 11.297546 | 11.286263 | 11.324362 |
| ASMM10P007 | 0.04613 | 0.3771497 | 1.3990195 | up | NM_011643 | Trpc1 | transient receptor potential cation channel, subfamily C, member 1 | 298.3055 | 258.33819 | 8.475409 | 7.990993 | 348.12842 | 223.04019 | 323.7479 | 282.91974 | 282.73334 | 209.36148 | 8.451836 | 8.212381 | 8.762009 | 8.066499 | 8.031421 | 7.8750577 |
| ASMM10P041 | 0.04619 | 0.3771497 | 1.2564887 | up | NM_025360 | Tmed3 | transmembrane emp24 domain containing 3 | 22628.521 | 20972.919 | 14.613708 | 14.28431 | 22140.082 | 22343.713 | 23401.768 | 20632.91 | 20543.7 | 21742.148 | 14.521608 | 14.608833 | 14.710682 | 14.214314 | 14.154012 | 14.484604 |
| ASMM10P051 | 0.04620 | 0.3771497 | 1.2754371 | up | NM_153529 | Nrn1 | neuritin 1 | 201.62039 | 195.50542 | 7.943643 | 7.592652 | 224.76967 | 203.82684 | 176.26465 | 231.89693 | 187.5814 | 167.03792 | 7.8399844 | 8.0880165 | 7.902929 | 7.777145 | 7.4436717 | 7.557138 |
| ASMM10P027 | 0.04630 | 0.3771497 | 3.1824272 | up | NM_024198 | Gpx7 | glutathione peroxidase 7 | 27.096689 | 10.329939 | 4.93009 | 3.259963 | 41.954475 | 15.030251 | 24.30534 | 17.60216 | 4.9999995 | 8.387658 | 5.4528465 | 4.3761 | 4.9613237 | 4.0176315 | 2.3265402 | 3.435716 |
| ASMM10P002 | 0.04631 | 0.3771497 | 1.2663656 | up | NM_031843 | Dpp7 | dipeptidylpeptidase 7 | 1049.2767 | 992.61455 | 10.265819 | 9.925125 | 1187.6421 | 992.2457 | 967.94244 | 996.268 | 929.16486 | 1052.4108 | 10.192507 | 10.305668 | 10.299282 | 9.883815 | 9.752144 | 10.139416 |
| ASMM10P019 | 0.04634 | 0.3771497 | 1.3819056 | up | NM_019437 | Rfk | riboflavin kinase | 87.810593 | 77.602468 | 6.746763 | 6.280104 | 96.80407 | 73.25216 | 93.37555 | 71.063354 | 85.3374 | 76.40665 | 6.6483417 | 6.626169 | 6.965777 | 6.0586205 | 6.304405 | 6.477285 |
| ASMM10P027 | 0.04636 | 0.3771497 | 3.485804 | up | NM_028698 | Ccdc30 | coiled-coil domain containing 30 | 32.787271 | 12.213965 | 5.284822 | 3.48333 | 41.359196 | 33.567787 | 23.43483 | 9.283303 | 6.957135 | 20.401457 | 5.4327726 | 5.514857 | 4.906836 | 3.072404 | 2.7077398 | 4.6698475 |
| ASMM10P004 | 0.04640 | 0.3771497 | 1.8530975 | up | NM_001012 | Zfp708 | zinc finger protein 708 | 42.891387 | 27.271236 | 5.686796 | 4.796858 | 39.074284 | 54.96954 | 34.630337 | 26.51734 | 27.103378 | 28.19299 | 5.3506584 | 6.2196956 | 5.490035 | 4.6239753 | 4.6546893 | 5.111908 |
| ASMM10P002 | 0.04642 | 0.377216 | 1.9479791 | up | NM_138752 | Plekhg2 | pleckstrin homology domain containing, family G (with RhoGef domain) member 2 | 112.30437 | 70.563609 | 7.070896 | 6.108918 | 96.7581 | 149.02843 | 91.12658 | 73.02512 | 91.33696 | 47.328747 | 6.647814 | 7.635644 | 6.9292307 | 6.0982475 | 6.4020677 | 5.826439 |
| ASMM10P012 | 0.04650 | 0.3773855 | 1.496964 | up | NM_177755 | Klhl38 | kelch-like 38 | 71.07265 | 58.022304 | 6.43473 | 5.852691 | 84.06162 | 58.1362 | 71.02013 | 71.87611 | 47.964283 | 54.22652 | 6.4468017 | 6.299522 | 6.5578666 | 6.0739627 | 5.474092 | 6.010017 |
| ASMM10P028 | 0.04651 | 0.3773855 | 1.5449396 | up | NM_001122 | Epha7 | Eph receptor A7 | 56.649415 | 44.074876 | 6.107884 | 5.480334 | 54.719475 | 66.76749 | 48.46128 | 44.964603 | 44.974415 | 42.28561 | 5.8324738 | 6.495999 | 5.99518 | 5.3952117 | 5.3790874 | 5.6667023 |
| ASMM10P043 | 0.04663 | 0.3778835 | 1.3376392 | up | NM_021548 | Arpp19 | cAMP-regulated phosphoprotein 19 | 46023.764 | 41900.181 | 15.686765 | 15.267076 | 45248.547 | 40438.184 | 52384.562 | 43355.28 | 46764.383 | 35580.88 | 15.609929 | 15.48782 | 15.962545 | 15.28092 | 15.326235 | 15.194072 |
| ASMM10P010 | 0.04672 | 0.3781644 | 1.3891949 | up | NM_026965 | Comtd1 | catechol-O-methyltransferase domain containing 1 | 625.18861 | 542.68842 | 9.528647 | 9.054398 | 651.903 | 594.2553 | 629.40753 | 524.07117 | 512.5025 | 591.4916 | 9.330263 | 9.582554 | 9.673123 | 8.957395 | 8.891131 | 9.314667 |
| ASMM10P022 | 0.04677 | 0.3781644 | 1.6161147 | up | NM_028846 | Usp20 | ubiquitin specific peptidase 20 | 45.117579 | 34.296896 | 5.792333 | 5.099803 | 40.672173 | 49.5377 | 45.142864 | 30.898518 | 43.742924 | 28.249245 | 5.4089146 | 6.0742364 | 5.893847 | 4.8485684 | 5.3367567 | 5.1140842 |
| ASMM10P003 | 0.04688 | 0.3781644 | 1.2806912 | up | NM_172706 | 9330182L06 | RIKEN cDNA 9330182L06 gene | 342.48226 | 328.04618 | 8.693019 | 8.336096 | 374.56747 | 332.45364 | 320.42566 | 354.95886 | 302.43362 | 326.74606 | 8.559554 | 8.770444 | 8.749059 | 8.392013 | 8.130447 | 8.485829 |
| ASMM10P008 | 0.04688 | 0.3781644 | 1.3563862 | up | NM_133905 | Papd4 | PAP associated domain containing 4 | 235.86599 | 213.09642 | 8.16538 | 7.725612 | 260.7807 | 236.47664 | 210.34062 | 200.8124 | 209.22043 | 229.25644 | 8.047545 | 8.292515 | 8.156081 | 7.5702834 | 7.6063285 | 8.000225 |
| ASMM10P023 | 0.04690 | 0.3781644 | 1.657326 | up | NM_029617 | Casc5 | cancer susceptibility candidate 5 | 24.41847 | 17.34253 | 4.861893 | 4.133036 | 30.731892 | 17.353256 | 25.170263 | 21.050854 | 14.182039 | 16.794697 | 4.9956975 | 4.5771847 | 5.0127983 | 4.282072 | 3.7118893 | 4.405147 |
| ASMM10P039 | 0.04691 | 0.3781644 | 1.3527698 | up | NM_026818 | Cilp2 | cartilage intermediate layer protein 2 | 371.02399 | 329.72107 | 8.780046 | 8.34413 | 478.02582 | 330.78937 | 304.25677 | 320.26556 | 310.27484 | 358.6228 | 8.902044 | 8.762009 | 8.676086 | 8.244882 | 8.169419 | 8.618089 |
| ASMM10P051 | 0.04693 | 0.3781644 | 1.4056782 | up | NM_025588 | Exoc2 | exocyst complex component 2 | 15298.732 | 12696.009 | 14.037871 | 13.546605 | 14155.245 | 13929.638 | 17811.312 | 12417.509 | 12515.177 | 13155.341 | 13.842809 | 13.948772 | 14.322032 | 13.470238 | 13.436111 | 13.733465 |
| ASMM10P041 | 0.04697 | 0.3781644 | 1.4045594 | up | NM_027448 | Lca5 | Leber congenital amaurosis 5 (human) | 48.247448 | 41.995737 | 5.883019 | 5.392901 | 50.031075 | 53.671745 | 41.039524 | 50.429905 | 42.64868 | 32.908627 | 5.710149 | 6.1856666 | 5.7532415 | 5.5571723 | 5.3014636 | 5.3200684 |
| ASMM10P033 | 0.04710 | 0.3785633 | 1.5952196 | up | NM_015768 | Prok2 | prokineticin 2 | 1600.2954 | 1192.3544 | 10.847031 | 10.173276 | 1477.0427 | 1354.0853 | 1969.7582 | 1338.2467 | 1246.9347 | 991.8817 | 10.51865 | 10.741572 | 11.280872 | 10.303841 | 10.163206 | 10.052782 |
| ASMM10P041 | 0.04722 | 0.3785633 | 1.684081 | up | NM_173433 | Kdm4d | lysine (K)-specific demethylase 4D | 216.51173 | 156.25481 | 8.02678 | 7.274819 | 222.0847 | 167.99188 | 259.4586 | 153.66779 | 144.38448 | 170.71216 | 7.823436 | 7.8085346 | 8.44837 | 7.178195 | 7.056573 | 7.589688 |
| ASMM10P036 | 0.04723 | 0.3785633 | 1.5064464 | up | NM_008486 | Anpep | alanyl (membrane) aminopeptidase | 4375.6332 | 3444.7865 | 12.284288 | 11.693138 | 4275.262 | 4429.4746 | 4422.163 | 3147.5034 | 3128.271 | 4058.585 | 12.086116 | 12.364277 | 12.40247 | 11.524864 | 11.495107 | 12.059444 |
| ASMM10P028 | 0.04723 | 0.3785633 | 1.5679342 | up | NM_172869 | Frmd3 | FERM domain containing 3 | 742.35904 | 566.11654 | 9.759824 | 9.110959 | 740.86786 | 583.93036 | 902.2789 | 628.5793 | 600.5624 | 469.20792 | 9.52457 | 9.557442 | 10.197461 | 9.221341 | 9.122068 | 8.989469 |
| ASMM10P045 | 0.04726 | 0.3785633 | 1.2538346 | up | NM_001033 | A830080D0 | RIKEN cDNA A830080D01 gene | 461.87965 | 448.12754 | 9.110526 | 8.784179 | 508.1632 | 439.52527 | 437.95047 | 503.16138 | 415.84787 | 425.37338 | 8.984753 | 9.163094 | 9.183731 | 8.903881 | 8.593447 | 8.855209 |
| ASMM10P014 | 0.04737 | 0.3785633 | 1.6472371 | up | NM_010465 | Hoxc6 | homeobox C6 | 38.681287 | 28.168486 | 5.558897 | 4.838848 | 34.202736 | 45.04819 | 36.792934 | 32.540234 | 26.335135 | 25.63009 | 5.152472 | 5.939168 | 5.5850496 | 4.9246516 | 4.612876 | 4.9790173 |
| ASMM10P054 | 0.04740 | 0.3785633 | 1.9778283 | up | NM_175491 | Smcr8 | Smith-Magenis syndrome chromosome region, candidate 8 homolog (human) | 37.812058 | 22.263575 | 5.480643 | 4.496726 | 31.474777 | 27.793777 | 54.16762 | 25.389233 | 23.525621 | 17.875872 | 5.034919 | 5.244512 | 6.162498 | 4.5567465 | 4.44389 | 4.489541 |
| ASMM10P005 | 0.04744 | 0.3785633 | 1.2948946 | up | NM_021388 | Extl2 | exostoses (multiple)-like 2 | 438.16913 | 411.70634 | 9.034286 | 8.661452 | 498.2068 | 397.3614 | 418.93918 | 380.02112 | 414.4401 | 440.6578 | 8.956758 | 9.022037 | 9.124064 | 8.495328 | 8.588509 | 8.900518 |
| ASMM10P050 | 0.04745 | 0.3785633 | 1.5296402 | up | NM_053122 | Immp2l | IMP2 inner mitochondrial membrane peptidase-like (S. cerevisiae) | 931.98331 | 733.43337 | 10.101797 | 9.488605 | 887.4869 | 1002.6846 | 905.7784 | 717.11475 | 672.6281 | 810.55725 | 9.783613 | 10.319544 | 10.202234 | 9.417938 | 9.283216 | 9.76466 |
| ASMM10P002 | 0.04751 | 0.3785633 | 1.2947913 | up | NM_001166 | Rtel1 | regulator of telomere elongation helicase 1 | 103.88214 | 95.20658 | 6.942397 | 6.569677 | 143.25392 | 87.80256 | 80.58993 | 100.42007 | 103.135 | 82.06467 | 7.1941075 | 6.884117 | 6.7489657 | 6.55959 | 6.573825 | 6.5756164 |
| ASMM10P003 | 0.04751 | 0.3785633 | 1.7058131 | up | NM_028450 | Gulp1 | GULP, engulfment adaptor PTB domain containing 1 | 72.628343 | 52.544273 | 6.467308 | 5.696848 | 85.88266 | 61.335224 | 70.667145 | 38.621704 | 62.31188 | 56.699234 | 6.475956 | 6.375521 | 6.550446 | 5.176731 | 5.8459783 | 6.067835 |
| ASMM10P055 | 0.04756 | 0.3785633 | 1.5224164 | up | NM_001110 | Samd11 | sterile alpha motif domain containing 11 | 4478.7989 | 3455.3766 | 12.301034 | 11.694671 | 4213.8164 | 3873.1802 | 5349.4 | 3952.2349 | 3636.3652 | 2777.5298 | 12.059903 | 12.176844 | 12.666356 | 11.87202 | 11.711131 | 11.500863 |
| ASMM10P013 | 0.04756 | 0.3785633 | 1.3758444 | up | NM_010267 | Gdap1 | ganglioside-induced differentiation-associated-protein 1 | 406.16498 | 360.51173 | 8.93289 | 8.472573 | 397.49353 | 395.32227 | 425.67914 | 391.0499 | 350.99252 | 339.49277 | 8.642273 | 9.012651 | 9.143747 | 8.533501 | 8.346218 | 8.538 |
| ASMM10P007 | 0.04757 | 0.3785633 | 2.1787635 | up | NM_011243 | Rarb | retinoic acid receptor, beta | 139.6791 | 74.06663 | 7.332928 | 6.209419 | 124.08743 | 204.97446 | 89.97541 | 86.02459 | 68.76361 | 67.41169 | 6.992764 | 8.095811 | 6.91021 | 6.32775 | 5.9900637 | 6.3104424 |
| ASMM10P007 | 0.04758 | 0.3785633 | 1.4009772 | up | NM_010217 | Ctgf | connective tissue growth factor | 924.04219 | 798.59382 | 10.091149 | 9.604716 | 934.1552 | 931.0247 | 906.94666 | 876.60004 | 707.55133 | 811.6301 | 9.857032 | 10.212046 | 10.20437 | 9.691286 | 9.3560295 | 9.766832 |
| ASMM10P041 | 0.04758 | 0.3785633 | 1.277968 | up | NM_001164 | Tpm1 | tropomyosin 1, alpha | 59256.425 | 57334.148 | 16.078219 | 15.724368 | 54702.31 | 60659.336 | 62407.63 | 58125.95 | 57639.938 | 56236.555 | 15.88351 | 16.120363 | 16.230785 | 15.682177 | 15.626099 | 15.864827 |
| ASMM10P036 | 0.04759 | 0.3785633 | 1.5217559 | up | NM_009099 | Trim30a | tripartite motif-containing 30A | 556.12417 | 445.10224 | 9.366355 | 8.760618 | 527.1931 | 562.1315 | 579.0479 | 393.4811 | 552.9758 | 388.84982 | 9.031325 | 9.510299 | 9.557442 | 8.544504 | 9.003456 | 8.733895 |
| ASMM10P053 | 0.04763 | 0.3785633 | 1.5570806 | up | NM_026276 | Aasdhppt | aminoadipate-semialdehyde dehydrogenase-phosphopantetheinyl transferase | 163.22342 | 128.16834 | 7.637528 | 6.998685 | 148.99391 | 153.8543 | 186.82204 | 135.19849 | 126.0024 | 123.30413 | 7.252572 | 7.6802573 | 7.9797554 | 6.9917755 | 6.8610206 | 7.1432576 |
| ASMM10P031 | 0.04764 | 0.3785633 | 1.6522863 | up | NM_025725 | Ccdc96 | coiled-coil domain containing 96 | 156.81257 | 117.0415 | 7.581899 | 6.857435 | 136.73508 | 176.28073 | 157.4219 | 105.53252 | 138.2928 | 107.29919 | 7.1291046 | 7.878583 | 7.738009 | 6.6316776 | 6.9928646 | 6.9477634 |
| ASMM10P006 | 0.04765 | 0.3785633 | 4.2416436 | up | NM_008091 | Gata3 | GATA binding protein 3 | 30.759844 | 8.641773 | 5.110517 | 3.025893 | 43.052986 | 14.933651 | 34.292896 | 4.9999995 | 5.363958 | 15.561361 | 5.492697 | 4.3666368 | 5.472216 | 2.3250442 | 2.453433 | 4.2992024 |
| ASMM10P008 | 0.04781 | 0.3787201 | 1.6572008 | up | NM_027892 | Ppp1r12a | protein phosphatase 1, regulatory (inhibitor) subunit 12A | 308.21781 | 221.12911 | 8.506632 | 7.777884 | 382.0856 | 209.40799 | 333.15985 | 225.19472 | 197.22142 | 240.97119 | 8.590341 | 8.125017 | 8.804539 | 7.7355537 | 7.519285 | 8.078813 |
| ASMM10P046 | 0.04783 | 0.3787201 | 1.8413706 | up | NM_001168 | R3hdm2 | R3H domain containing 2 | 4611.0844 | 2844.6231 | 12.300034 | 11.419254 | 3668.4595 | 3854.2546 | 6310.539 | 3097.4507 | 2921.5996 | 2514.819 | 11.84189 | 12.168786 | 12.889426 | 11.50207 | 11.394186 | 11.361506 |
| ASMM10P007 | 0.04785 | 0.3787201 | 1.6837046 | up | NM_025386 | Fbxo36 | F-box protein 36 | 38.228941 | 27.33437 | 5.533332 | 4.781693 | 47.12104 | 30.583395 | 36.982388 | 24.686941 | 25.105265 | 32.210903 | 5.6224537 | 5.3848624 | 5.592679 | 4.512165 | 4.5396852 | 5.2932277 |
| ASMM10P048 | 0.04786 | 0.3787201 | 1.3112614 | up | NM_177445 | Dars | aspartyl-tRNA synthetase | 2354.0601 | 2140.1929 | 11.40327 | 11.012314 | 2645.729 | 2249.9978 | 2166.4534 | 1942.0641 | 2106.847 | 2371.6677 | 11.35017 | 11.449719 | 11.40992 | 10.84016 | 10.916883 | 11.2799 |
| ASMM10P013 | 0.04791 | 0.3789557 | 1.2779355 | up | NM_029236 | Bcdin3d | BCDIN3 domain containing | 2252.3292 | 2080.6477 | 11.331856 | 10.978041 | 2491.9182 | 1928.4323 | 2336.6372 | 2060.6904 | 2027.7183 | 2153.5344 | 11.265204 | 11.214623 | 11.515741 | 10.930141 | 10.862122 | 11.14186 |
| ASMM10P027 | 0.04805 | 0.3793178 | 1.2529426 | up | NM_008120 | Gja4 | gap junction protein, alpha 4 | 3380.6848 | 3183.5128 | 11.905755 | 11.580435 | 3698.8972 | 2895.2202 | 3547.937 | 3023.5703 | 3614.37 | 2912.5981 | 11.85389 | 11.777259 | 12.086116 | 11.464935 | 11.702879 | 11.57349 |
| ASMM10P009 | 0.04807 | 0.3793309 | 1.3756988 | up | NM_175016 | Alkbh2 | alkB, alkylation repair homolog 2 (E. coli) | 57.458702 | 51.070738 | 6.140982 | 5.680817 | 56.396996 | 52.153576 | 63.825535 | 50.835087 | 58.419483 | 43.957645 | 5.874278 | 6.1429296 | 6.4057384 | 5.569286 | 5.751559 | 5.721607 |
| ASMM10P032 | 0.04810 | 0.3793309 | 1.2944771 | up | NM_028990 | Tmem168 | transmembrane protein 168 | 26.823715 | 24.749242 | 5.02724 | 4.654871 | 28.935225 | 27.305326 | 24.230593 | 24.032932 | 28.188164 | 22.02663 | 4.9070306 | 5.2183003 | 4.956389 | 4.477055 | 4.7152047 | 4.772352 |
| ASMM10P046 | 0.04819 | 0.3793309 | 1.2707991 | up | NM_023651 | Pex13 | peroxisomal biogenesis factor 13 | 652.77978 | 618.14638 | 9.591353 | 9.245617 | 668.22253 | 615.7769 | 674.3399 | 616.80927 | 647.97705 | 589.65283 | 9.369586 | 9.632313 | 9.77216 | 9.192572 | 9.234077 | 9.310202 |
| ASMM10P001 | 0.04822 | 0.3793309 | 1.7078639 | up | NM_138676 | Shkbp1 | Sh3kbp1 binding protein 1 | 381.68276 | 268.52255 | 8.827721 | 8.055528 | 317.61795 | 368.35034 | 459.08 | 295.0405 | 274.24606 | 236.28108 | 8.32179 | 8.91034 | 9.251033 | 8.131868 | 7.988068 | 8.046648 |
| ASMM10P048 | 0.04827 | 0.3793309 | 1.271978 | up | NM_178645 | Blmh | bleomycin hydrolase | 38708.515 | 36213.087 | 15.410613 | 15.063539 | 36518.348 | 36162.746 | 43444.45 | 36766.965 | 37116.875 | 34755.42 | 15.261946 | 15.333442 | 15.63645 | 15.046139 | 14.99493 | 15.149548 |
| ASMM10P014 | 0.04832 | 0.3793309 | 1.3606547 | up | NM_007929 | Emp2 | epithelial membrane protein 2 | 9589.1443 | 8191.2593 | 13.382962 | 12.938661 | 8922.238 | 9665.997 | 10179.198 | 7446.859 | 8747.864 | 8379.055 | 13.147073 | 13.449712 | 13.552102 | 12.756655 | 12.953506 | 13.105824 |
| ASMM10P033 | 0.04832 | 0.3793309 | 1.5150844 | up | NM_001081 | Lmod3 | leiomodin 3 (fetal) | 89.774495 | 73.268585 | 6.764957 | 6.165559 | 103.84837 | 67.38558 | 98.08954 | 80.1253 | 90.36255 | 49.317905 | 6.750298 | 6.5069985 | 7.0375743 | 6.2286177 | 6.3849835 | 5.883075 |
| ASMM10P050 | 0.04852 | 0.3801484 | 1.4709694 | up | NM_144524 | Angel1 | angel homolog 1 (Drosophila) | 161.93545 | 138.54102 | 7.622384 | 7.065617 | 192.02829 | 153.48819 | 140.28987 | 130.24568 | 188.47302 | 96.904366 | 7.6149664 | 7.676553 | 7.575634 | 6.938886 | 7.4506903 | 6.8072753 |
| ASMM10P053 | 0.04860 | 0.3802121 | 1.4603259 | up | NM_021895 | Actn4 | actinin alpha 4 | 38.044259 | 31.177428 | 5.537066 | 4.990775 | 40.22787 | 30.306553 | 43.598354 | 30.986317 | 31.822006 | 30.72396 | 5.39417 | 5.3715663 | 5.845461 | 4.8538637 | 4.8909 | 5.2275624 |
| ASMM10P005 | 0.04883 | 0.3808696 | 1.420256 | up | NM_001177 | Tnfaip8 | tumor necrosis factor, alpha-induced protein 8 | 456.51557 | 396.27544 | 9.095239 | 8.589088 | 492.89548 | 439.69034 | 436.9609 | 328.45422 | 492.56555 | 367.80655 | 8.9401455 | 9.163703 | 9.1818695 | 8.279491 | 8.833553 | 8.654221 |
| ASMM10P039 | 0.04886 | 0.3809879 | 1.6784327 | up | NM_021491 | Smpd3 | sphingomyelin phosphodiesterase 3, neutral | 925.59084 | 659.46746 | 10.081757 | 9.334642 | 799.66046 | 1090.3828 | 886.72925 | 589.59827 | 691.0863 | 697.7178 | 9.632461 | 10.440824 | 10.171985 | 9.130035 | 9.32352 | 9.550371 |
| ASMM10P034 | 0.04888 | 0.38101 | 1.2755116 | up | NM_023538 | Agk | acylglycerol kinase | 28570.564 | 26660.481 | 14.959004 | 14.607928 | 26545.04 | 29215.434 | 29951.217 | 26823.205 | 25910.02 | 27248.219 | 14.801826 | 14.996682 | 15.078504 | 14.589238 | 14.453724 | 14.780822 |
| ASMM10P009 | 0.04890 | 0.38101 | 1.9416246 | up | NM_175104 | Fam53c | family with sequence similarity 53, member C | 40.562363 | 24.002487 | 5.554334 | 4.59707 | 59.581867 | 25.827625 | 36.277596 | 19.624018 | 25.515478 | 26.867964 | 5.9563155 | 5.1453056 | 5.561382 | 4.1819153 | 4.565204 | 5.044091 |
| ASMM10P015 | 0.04904 | 0.3811128 | 1.2574575 | up | NM_026782 | Ccdc167 | coiled-coil domain containing 167 | 240.55774 | 234.35227 | 8.192127 | 7.861617 | 269.36163 | 241.17142 | 211.14018 | 219.49892 | 249.02527 | 234.53262 | 8.093991 | 8.319686 | 8.162704 | 7.697743 | 7.85185 | 8.035259 |
| ASMM10P016 | 0.04906 | 0.3811128 | 1.5904833 | up | NM_020568 | Plin4 | perilipin 4 | 48.550748 | 35.333124 | 5.828654 | 5.159189 | 70.44954 | 39.20114 | 36.001564 | 42.105476 | 31.48766 | 32.406235 | 6.199444 | 5.7359176 | 5.5506015 | 5.301051 | 4.875375 | 5.3011413 |
| ASMM10P047 | 0.04909 | 0.3811128 | 1.2612116 | up | NM_013851 | Abca8b | ATP-binding cassette, sub-family A (ABC1), member 8b | 539.5265 | 517.58908 | 9.324512 | 8.989701 | 574.00323 | 537.1274 | 507.44888 | 564.4594 | 492.71567 | 495.59216 | 9.145844 | 9.444545 | 9.383146 | 9.068353 | 8.834011 | 9.06674 |
| ASMM10P021 | 0.04915 | 0.3811128 | 1.264617 | up | NM_178411 | Zfp334 | zinc finger protein 334 | 3545.6676 | 3319.2958 | 11.981853 | 11.643152 | 3739.8728 | 3167.8403 | 3729.2896 | 3138.2957 | 3434.9849 | 3384.6067 | 11.873154 | 11.903945 | 12.168459 | 11.520562 | 11.623899 | 11.784996 |
| ASMM10P042 | 0.04921 | 0.3811128 | 1.3100928 | up | NM_009382 | Thy1 | thymus cell antigen 1, theta | 2895.5678 | 2601.3602 | 11.686349 | 11.29668 | 3341.0088 | 2258.1707 | 3087.524 | 2698.23 | 2566.2622 | 2539.5884 | 11.695756 | 11.455537 | 11.907753 | 11.310044 | 11.205658 | 11.374337 |
| ASMM10P043 | 0.04934 | 0.3811128 | 1.5903357 | up | NM_009757 | Bmp15 | bone morphogenetic protein 15 | 80.248243 | 61.565723 | 6.612956 | 5.943625 | 87.32113 | 65.5847 | 87.8389 | 56.302475 | 59.062073 | 69.33262 | 6.4990115 | 6.467017 | 6.8728395 | 5.7184753 | 5.7667294 | 6.3456693 |
| ASMM10P045 | 0.04935 | 0.3811128 | 1.3083737 | up | NM_207633 | Yipf6 | Yip1 domain family, member 6 | 4241.6623 | 3803.2046 | 12.231458 | 11.843683 | 3994.3027 | 4411.2324 | 4319.4517 | 3705.2405 | 3983.1758 | 3721.1975 | 11.970056 | 12.355743 | 12.368575 | 11.768994 | 11.830885 | 11.931171 |
| ASMM10P017 | 0.04938 | 0.3811128 | 1.3694547 | up | NM_177595 | Mkx | mohawk homeobox | 456.70278 | 401.14176 | 9.074155 | 8.620554 | 522.1786 | 352.99496 | 494.93478 | 394.70447 | 467.02048 | 341.70032 | 9.0175295 | 8.853843 | 9.351094 | 8.550983 | 8.762697 | 8.547982 |
| ASMM10P008 | 0.04943 | 0.3811128 | 1.3270196 | up | NM_027415 | Tmem70 | transmembrane protein 70 | 1824.0527 | 1647.3029 | 11.049979 | 10.64179 | 1948.1962 | 1815.2932 | 1708.6687 | 1652.6976 | 1506.725 | 1782.486 | 10.921075 | 11.137913 | 11.09095 | 10.606901 | 10.434738 | 10.88373 |
| ASMM10P001 | 0.04944 | 0.3811128 | 1.7400772 | up | NM_027678 | Zranb3 | zinc finger, RAN-binding domain containing 3 | 612.50291 | 408.15437 | 9.451243 | 8.652091 | 711.652 | 387.3037 | 738.55304 | 409.43787 | 388.6178 | 426.40744 | 9.467753 | 8.982571 | 9.903404 | 8.602139 | 8.496774 | 8.857361 |
| ASMM10P006 | 0.04946 | 0.3811128 | 1.548067 | up | NM_001242 | Nit1 | nitrilase 1 | 73.325801 | 57.403845 | 6.484589 | 5.854121 | 66.4184 | 86.10771 | 67.451294 | 61.11265 | 57.748642 | 53.350243 | 6.1138673 | 6.8553243 | 6.4845743 | 5.8363943 | 5.7370906 | 5.9888773 |
| ASMM10P043 | 0.04947 | 0.3811128 | 1.4264354 | up | NM_177911 | Tgm4 | transglutaminase 4 (prostate) | 60.387297 | 50.989538 | 6.200324 | 5.68791 | 67.11942 | 64.77043 | 49.27204 | 49.47243 | 51.66944 | 51.826744 | 6.129807 | 6.450095 | 6.021071 | 5.5317907 | 5.5832295 | 5.9487095 |
| ASMM10P032 | 0.04952 | 0.3811128 | 1.3141535 | up | NM_013895 | Timm13 | translocase of inner mitochondrial membrane 13 | 998.45835 | 908.49453 | 10.195097 | 9.800963 | 1004.9042 | 939.19214 | 1051.2787 | 926.2397 | 924.8876 | 874.3563 | 9.946719 | 10.224242 | 10.414331 | 9.778017 | 9.746489 | 9.878384 |
| ASMM10P014 | 0.04952 | 0.3811128 | 1.3214001 | up | NM_011949 | Mapk1 | mitogen-activated protein kinase 1 | 62768.957 | 58608.178 | 16.165922 | 15.763854 | 61767.46 | 60360.42 | 66178.99 | 58049.5 | 56849.793 | 60925.242 | 16.069048 | 16.112337 | 16.31638 | 15.677713 | 15.608239 | 16.005611 |

| | | | | | | | | | | | | | | | | | | | | | | | |
|---|---|---|---|---|---|---|---|---|---|---|---|---|---|---|---|---|---|---|---|---|---|---|---|
| ASMM10P008 | 0.04953 | 0.3811128 | 1.3550526 | up | NM_027409 | Mospd1 | motile sperm domain containing 1 | 455.38874 | 410.98752 | 9.095534 | 8.657185 | 447.6143 | 451.9471 | 466.60483 | 401.51343 | 471.92722 | 359.5219 | 8.807926 | 9.202677 | 9.275999 | 8.5752325 | 8.774757 | 8.621566 |
| ASMM10P004 | 0.04955 | 0.3811128 | 1.3107658 | up | NM_001025 | Ccdc141 | coiled-coil domain containing 141 | 80.208036 | 74.694112 | 6.61737 | 6.22696 | 84.919464 | 67.88276 | 87.821884 | 73.94559 | 83.82352 | 66.313225 | 6.462496 | 6.516926 | 6.8726873 | 6.1155877 | 6.2806077 | 6.284684 |
| ASMM10P019 | 0.04956 | 0.3811128 | 1.3335525 | up | NM_013806 | Abcc2 | ATP-binding cassette, sub-family C (CFTR/MRP), member 2 | 55.138113 | 49.096943 | 6.042003 | 5.626728 | 74.63201 | 48.111526 | 42.670803 | 56.676636 | 49.036728 | 41.577465 | 6.278755 | 6.033341 | 5.813913 | 5.729348 | 5.506685 | 5.644152 |
| ASMM10P047 | 0.04976 | 0.3811128 | 1.8856034 | up | NM_008798 | Pdcd1 | programmed cell death 1 | 24.826006 | 15.649739 | 4.891369 | 3.976342 | 24.757147 | 30.561064 | 19.159807 | 12.754242 | 20.131626 | 14.063349 | 4.675598 | 5.383355 | 4.615153 | 3.5424073 | 4.225822 | 4.160798 |
| ASMM10P036 | 0.04977 | 0.3811128 | 1.3291187 | up | NM_018880 | Trim3 | tripartite motif-containing 3 | 2295.3354 | 2052.0139 | 11.367205 | 10.956735 | 2286.935 | 2090.4783 | 2508.5928 | 2030.743 | 2055.1836 | 2070.1152 | 11.14966 | 11.342868 | 11.609086 | 10.908467 | 10.878893 | 11.082844 |
| ASMM10P006 | 0.04978 | 0.3811128 | 1.2550501 | up | NM_001271 | Cnot7 | CCR4-NOT transcription complex, subunit 7 | 288.36826 | 275.8221 | 8.422328 | 8.094583 | 381.32663 | 258.07135 | 225.70679 | 267.58356 | 290.36383 | 269.5189 | 8.587476 | 8.42044 | 8.259068 | 7.9867992 | 8.071062 | 8.225888 |
| ASMM10P033 | 0.04990 | 0.3811128 | 1.3622585 | up | NM_178045 | Rassf4 | Ras association (RalGDS/AF-6) domain family member 4 | 1796.7706 | 1590.8015 | 11.025213 | 10.579212 | 2010.8542 | 1536.8352 | 1842.6223 | 1348.5479 | 1859.3251 | 1564.5316 | 10.967649 | 10.907966 | 11.200024 | 10.312758 | 10.728065 | 10.696814 |
| ASMM10P016 | 0.04990 | 0.3811128 | 1.2833162 | up | NM_153551 | Dennd1c | DENN/MADD domain containing 1C | 993.51553 | 929.57008 | 10.188016 | 9.828139 | 1026.3352 | 899.9813 | 1054.2301 | 941.28925 | 1010.316 | 837.105 | 9.978937 | 10.165658 | 10.419452 | 9.799907 | 9.871381 | 9.813129 |
| ASMM10P020 | 0.04993 | 0.3811128 | 1.3258914 | up | NM_153125 | Sec16a | SEC16 homolog A (S. cerevisiae) | 300.54954 | 277.47563 | 8.508607 | 8.101644 | 302.9899 | 301.69363 | 296.9651 | 311.08997 | 269.17633 | 252.16058 | 8.253001 | 8.631424 | 8.641396 | 8.202581 | 7.9632683 | 8.139084 |
| ASMM10P051 | 0.04995 | 0.3811128 | 2.1820916 | up | NM_010076 | Drd1 | dopamine receptor D1A | 5984.4957 | 2980.6077 | 12.618422 | 11.49271 | 4482.002 | 9432.868 | 4038.6172 | 3108.9937 | 2847.9536 | 2984.8757 | 12.15792 | 13.412525 | 12.284821 | 11.507025 | 11.360197 | 11.610909 |

**Supplemental Table S2: Identification of KHSRP-target mRNAs through RIP-Seq.**

| id | ENSEMBL gene ID | Gene Symbol | Gene Info | log2FoldChange | pval | padj |
|---|---|---|---|---|---|---|
| 18183 | ENSMUSG00000054091 | 1810037I17Rik; Gm2 | 67704 [Gene Symbol: 1810037I17Rik] [Locus Tag: ] [Chromosome: 3] [Map Location: 3 | 67.68240451 | 2.87E-45 | 5.95E-42 |
| 15509 | ENSMUSG00000007670 | Khsrp | 16549 [Gene Symbol: Khsrp] [Locus Tag: ] [Chromosome: 17] [Map Location: 17 D\|17 2 | 34.82793612 | 5.60E-50 | 2.61E-46 |
| 16596 | ENSMUSG00000039221 | Rpl22l1 | 68028 [Gene Symbol: Rpl22l1] [Locus Tag: ] [Chromosome: 3] [Map Location: 3\|3 A3] [ | 32.20174299 | 4.76E-05 | 0.000159 |
| 4606 | ENSMUSG00000012483 | Rpa3 | 68240 [Gene Symbol: Rpa3] [Locus Tag: ] [Chromosome: 6] [Map Location: 6\|6 A1] [Des | 29.28898208 | 1.45E-21 | 6.93E-20 |
| 8641 | ENSMUSG00000028248 | Pnisr | 66625 [Gene Symbol: Pnisr] [Locus Tag: ] [Chromosome: 4] [Map Location: 4\|4 A3] [Des | 26.35381375 | 9.24E-32 | 2.29E-29 |
| 2604 | ENSMUSG00000027782 | Kpna4 | 16649 [Gene Symbol: Kpna4] [Locus Tag: ] [Chromosome: 3] [Map Location: 3\|3 E2] [De | 25.33717099 | 9.22E-33 | 2.96E-30 |
| 8372 | ENSMUSG00000028044 | Cks1b | 54124 [Gene Symbol: Cks1b] [Locus Tag: ] [Chromosome: 3] [Map Location: 3\|3 F1] [De | 24.67545336 | 6.99E-15 | 1.14E-13 |
| 14663 | ENSMUSG00000024614 | Tmx3 | 67988 [Gene Symbol: Tmx3] [Locus Tag: ] [Chromosome: 18] [Map Location: 18\|18 E4] | 23.8038627 | 2.80E-41 | 3.47E-38 |
| 11479 | ENSMUSG00000074781 | Ube2n | 93765 [Gene Symbol: Ube2n] [Locus Tag: ] [Chromosome: 10] [Map Location: 10\|10 C2 | 23.3243694 | 1.91E-32 | 5.46E-30 |
| 3745 | ENSMUSG00000043421 | Hilpda | 69573 [Gene Symbol: Hilpda] [Locus Tag: ] [Chromosome: 6] [Map Location: 6\|6 A3.3] [ | 23.17045516 | 9.46E-34 | 3.60E-31 |
| 7565 | ENSMUSG00000063480 | Nhp2l1; LOC1008624 | 20826 [Gene Symbol: Nhp2l1] [Locus Tag: ] [Chromosome: 15] [Map Location: 15 E1\|1 | 22.89018525 | 2.11E-28 | 2.89E-26 |
| 13686 | ENSMUSG00000044786 | Zfp36 | 22695 [Gene Symbol: Zfp36] [Locus Tag: ] [Chromosome: 7] [Map Location: 7 A3\|7 16. | 21.98038883 | 4.79E-24 | 3.40E-22 |
| 12664 | ENSMUSG00000019890 | Nts | 67405 [Gene Symbol: Nts] [Locus Tag: ] [Chromosome: 10] [Map Location: 10\|10 D1] [D | 21.8840858 | 5.42E-19 | 1.68E-17 |
| 10310 | ENSMUSG00000038975 | Rabggtb | 19352 [Gene Symbol: Rabggtb] [Locus Tag: ] [Chromosome: 3] [Map Location: 3 H3\|3 7 | 21.34802076 | 1.94E-32 | 5.46E-30 |
| 16245 | ENSMUSG00000021537 | Cetn3 | 12626 [Gene Symbol: Cetn3] [Locus Tag: ] [Chromosome: 13] [Map Location: 13\|13 C3 | 20.99102163 | 1.45E-32 | 4.29E-30 |
| 18047 | ENSMUSG00000028706 | Nsun4 | 72181 [Gene Symbol: Nsun4] [Locus Tag: ] [Chromosome: 4] [Map Location: 4\|4 D1] [D | 20.51032582 | 8.65E-31 | 1.81E-28 |
| 15778 | ENSMUSG00000020644 | Id2 | 15902 [Gene Symbol: Id2] [Locus Tag: ] [Chromosome: 12] [Map Location: 12 8.57 cM\| | 20.28043985 | 3.70E-35 | 1.72E-32 |
| 11588 | ENSMUSG00000038598 | AI481877 | 100155 [Gene Symbol: AI481877] [Locus Tag: ] [Chromosome: 4] [Map Location: 4\|4 B | 20.25004483 | 1.44E-27 | 1.68E-25 |
| 9024 | ENSMUSG00000024317 | Rnf138 | 56515 [Gene Symbol: Rnf138] [Locus Tag: ] [Chromosome: 18] [Map Location: 18\|18 A | 20.20556112 | 1.50E-39 | 1.55E-36 |
| 14984 | ENSMUSG00000037266 | Rsrp1 | 27981 [Gene Symbol: Rsrp1] [Locus Tag: ] [Chromosome: 4] [Map Location: 4 D3\|4 67. | 20.09950547 | 1.16E-15 | 2.16E-14 |
| 17612 | ENSMUSG00000022674 | Ube2v2 | 70620 [Gene Symbol: Ube2v2] [Locus Tag: ] [Chromosome: 16] [Map Location: 16\|16 A | 19.98765843 | 1.18E-36 | 8.11E-34 |
| 16198 | ENSMUSG00000066637 | Ttc32 | 75516 [Gene Symbol: Ttc32] [Locus Tag: ] [Chromosome: 12] [Map Location: 12\|12 A1. | 19.86412176 | 1.28E-13 | 1.71E-12 |
| 17793 | ENSMUSG00000015672 | Mrpl32 | 75398 [Gene Symbol: Mrpl32] [Locus Tag: ] [Chromosome: 13] [Map Location: 13\|13 A | 19.32873421 | 3.84E-27 | 4.16E-25 |
| 6763 | ENSMUSG00000033578 | Tmem35 | 67564 [Gene Symbol: Tmem35] [Locus Tag: ] [Chromosome: X] [Map Location: X\|X E3] [ | 18.10869121 | 1.49E-27 | 1.71E-25 |
| 3263 | ENSMUSG00000026960 | Arl6ip6 | 65103 [Gene Symbol: Arl6ip6] [Locus Tag: ] [Chromosome: 2] [Map Location: 2\|2 C1] [ | 18.09881797 | 8.82E-36 | 4.69E-33 |
| 1132 | ENSMUSG00000014905 | Dnajb9 | 27362 [Gene Symbol: Dnajb9] [Locus Tag: ] [Chromosome: 12] [Map Location: 12\|12 B | 17.60835057 | 1.02E-32 | 3.23E-30 |
| 15176 | ENSMUSG00000025040 | Fundc1 | 72018 [Gene Symbol: Fundc1] [Locus Tag: ] [Chromosome: X] [Map Location: X\|X A2] [ | 17.55331318 | 2.54E-32 | 6.86E-30 |
| 1646 | ENSMUSG00000073131 | Vma21 | 67048 [Gene Symbol: Vma21] [Locus Tag: ] [Chromosome: X] [Map Location: X\|X] [Desc | 17.53347423 | 1.60E-35 | 7.62E-33 |
| 272 | ENSMUSG00000022205 | Sub1 | 20024 [Gene Symbol: Sub1] [Locus Tag: ] [Chromosome: 15] [Map Location: 15\|15 A2] | 17.32264375 | 7.83E-31 | 1.66E-28 |
| 16657 | ENSMUSG00000036022 | Fam122b | 78755 [Gene Symbol: Fam122b] [Locus Tag: ] [Chromosome: X] [Map Location: X\|X A5] | 17.16848298 | 1.94E-18 | 5.54E-17 |
| 16954 | ENSMUSG00000043832 | Clec4a3 | 73149 [Gene Symbol: Clec4a3] [Locus Tag: ] [Chromosome: 6] [Map Location: 6\|6 F2] [ | 16.97217269 | 0.014284 | 0.028618 |
| 16434 | ENSMUSG00000028232 | Tmem68 | 72098 [Gene Symbol: Tmem68] [Locus Tag: ] [Chromosome: 4] [Map Location: 4\|4 A1] | 16.72720166 | 4.41E-36 | 2.65E-33 |
| 4850 | ENSMUSG00000024480 | Ap3s1 | 11777 [Gene Symbol: Ap3s1] [Locus Tag: ] [Chromosome: 18] [Map Location: 18\|18 C] | 16.68165629 | 6.64E-26 | 6.31E-24 |
| 12674 | ENSMUSG00000045996 | Polr2k; LOC1008624 | 17749 [Gene Symbol: Polr2k] [Locus Tag: ] [Chromosome: 15] [Map Location: 15\|15 A2 | 16.6535678 | 9.29E-11 | 7.81E-10 |
| 5581 | ENSMUSG00000019818 | Cd164 | 53599 [Gene Symbol: Cd164] [Locus Tag: ] [Chromosome: 10] [Map Location: 10 B2\|10 | 16.62658697 | 1.72E-29 | 2.87E-27 |
| 6153 | ENSMUSG00000071337 | Tia1 | 21841 [Gene Symbol: Tia1] [Locus Tag: ] [Chromosome: 6] [Map Location: 6\|6 D2] [Des | 16.61413432 | 2.75E-25 | 2.34E-23 |
| 9876 | ENSMUSG00000085793 | Lin52; Gm7020 | 217708 [Gene Symbol: Lin52] [Locus Tag: ] [Chromosome: 12] [Map Location: 12\|12 D | 16.54243212 | 4.44E-29 | 6.77E-27 |
| 14097 | ENSMUSG00000029819 | Npy | 109648 [Gene Symbol: Npy] [Locus Tag: ] [Chromosome: 6] [Map Location: 6 24.04 cM\| | 16.37065547 | 2.24E-37 | 1.74E-34 |
| 16711 | ENSMUSG00000036764 | Dnajc12 | 30045 [Gene Symbol: Dnajc12] [Locus Tag: ] [Chromosome: 10] [Map Location: 10\|10 | 16.27737164 | 7.33E-14 | 1.02E-12 |
| 2628 | ENSMUSG00000020390 | Ube2b | 22210 [Gene Symbol: Ube2b] [Locus Tag: ] [Chromosome: 11] [Map Location: 11\|11 B1 | 16.22847577 | 5.73E-37 | 4.10E-34 |
| 1837 | ENSMUSG00000055210 | Foxd2 | 17301 [Gene Symbol: Foxd2] [Locus Tag: ] [Chromosome: 4] [Map Location: 4 D1\|4 52. | 16.0538771 | 1.56E-05 | 5.76E-05 |
| 517 | ENSMUSG00000024097 | Srsf7 | 225027 [Gene Symbol: Srsf7] [Locus Tag: ] [Chromosome: 17] [Map Location: 17\|17 E3 | 15.99770724 | 9.69E-25 | 7.67E-23 |
| 1341 | ENSMUSG00000034855 | Cxcl10 | 15945 [Gene Symbol: Cxcl10] [Locus Tag: ] [Chromosome: 5] [Map Location: 5 E2\|5 46. | 15.85728444 | 0.024121 | 0.045559 |
| 6852 | ENSMUSG00000029373 | Pf4 | 56744 [Gene Symbol: Pf4] [Locus Tag: ] [Chromosome: 5] [Map Location: 5\|5 E1] [Descr | 15.77294746 | 1.21E-06 | 5.39E-06 |
| 53 | ENSMUSG00000033186 | Mzt1 | 76789 [Gene Symbol: Mzt1] [Locus Tag: ] [Chromosome: 14] [Map Location: 14\|14 E2. | 15.76920395 | 6.13E-36 | 3.46E-33 |
| 15472 | ENSMUSG00000025666 | Tmem47 | 192216 [Gene Symbol: Tmem47] [Locus Tag: ] [Chromosome: X] [Map Location: X\|X A7 | 15.67654302 | 1.23E-34 | 5.32E-32 |
| 17978 | ENSMUSG00000044155 | Lsm8 | 76522 [Gene Symbol: Lsm8] [Locus Tag: ] [Chromosome: 6] [Map Location: 6\|6 A2] [Des | 15.67163025 | 7.79E-21 | 3.34E-19 |
| 395 | ENSMUSG00000001774 | Chordc1 | 66917 [Gene Symbol: Chordc1] [Locus Tag: ] [Chromosome: 9] [Map Location: 9\|9 A3] [ | 15.57569918 | 1.87E-22 | 1.02E-20 |
| 10951 | ENSMUSG00000021098 | 4930447C04Rik | 75801 [Gene Symbol: 4930447C04Rik] [Locus Tag: ] [Chromosome: 12] [Map Location: | 15.54090998 | 1.81E-13 | 2.34E-12 |
| 12522 | ENSMUSG00000002732 | Fkbp7 | 14231 [Gene Symbol: Fkbp7] [Locus Tag: ] [Chromosome: 2] [Map Location: 2 C3\|2 45. | 15.28501442 | 7.99E-14 | 1.10E-12 |
| 8231 | ENSMUSG00000021134 | Srsf5 | 20384 [Gene Symbol: Srsf5] [Locus Tag: ] [Chromosome: 12] [Map Location: 12\|12 D2] | 15.26068007 | 2.72E-23 | 1.73E-21 |
| 4730 | ENSMUSG00000079317 | Trappc2 | 66226 [Gene Symbol: Trappc2] [Locus Tag: ] [Chromosome: X] [Map Location: X\|X F5] [ | 15.1555029 | 2.32E-17 | 5.57E-16 |
| 8209 | ENSMUSG00000054752 | Fsd1l | 319636 [Gene Symbol: Fsd1l] [Locus Tag: ] [Chromosome: 4] [Map Location: 4\|4 B2] [D | 15.01255472 | 2.12E-36 | 1.37E-33 |
| 10375 | ENSMUSG00000001707 | Eef1e1 | 66143 [Gene Symbol: Eef1e1] [Locus Tag: ] [Chromosome: 13] [Map Location: 13\|13 A5 | 14.88028828 | 2.44E-23 | 1.57E-21 |
| 3844 | ENSMUSG00000021643 | Serf1 | 20365 [Gene Symbol: Serf1] [Locus Tag: ] [Chromosome: 13] [Map Location: 13 D1\|13 | 14.87023811 | 9.85E-25 | 7.77E-23 |
| 2175 | ENSMUSG00000031197 | Vbp1 | 22327 [Gene Symbol: Vbp1] [Locus Tag: ] [Chromosome: X] [Map Location: X A7.3\|X 38 | 14.77712586 | 7.09E-32 | 1.78E-29 |
| 9197 | ENSMUSG00000026097 | Ormdl1 | 227102 [Gene Symbol: Ormdl1] [Locus Tag: ] [Chromosome: 1] [Map Location: 1\|1 C1. | 14.76050789 | 1.52E-21 | 7.26E-20 |
| 15111 | ENSMUSG00000026034 | Clk1 | 12747 [Gene Symbol: Clk1] [Locus Tag: ] [Chromosome: 1] [Map Location: 1 C1.3\|1 29. | 14.69940108 | 5.08E-22 | 2.60E-20 |
| 8253 | ENSMUSG00000017677 | Wsb1 | 78889 [Gene Symbol: Wsb1] [Locus Tag: ] [Chromosome: 11] [Map Location: 11\|11 B5] | 14.58008155 | 4.14E-34 | 1.64E-31 |
| 7055 | ENSMUSG00000018239 | Zcchc10 | 67966 [Gene Symbol: Zcchc10] [Locus Tag: ] [Chromosome: 11] [Map Location: 11 B1.3 | 14.49112728 | 1.56E-18 | 4.52E-17 |
| 6098 | ENSMUSG00000014177 | Tvp23b | 67510 [Gene Symbol: Tvp23b] [Locus Tag: ] [Chromosome: 11] [Map Location: 11\|11 B | 14.42885616 | 1.23E-19 | 4.31E-18 |
| 13403 | ENSMUSG00000037722 | Gnpnat1 | 54342 [Gene Symbol: Gnpnat1] [Locus Tag: ] [Chromosome: 14] [Map Location: 14\|14 | 14.28126207 | 4.44E-19 | 1.41E-17 |
| 5528 | ENSMUSG00000062198 | 2700097O09Rik | 72658 [Gene Symbol: 2700097O09Rik] [Locus Tag: ] [Chromosome: 12] [Map Location: | 14.26460867 | 6.55E-20 | 2.40E-18 |
| 13335 | ENSMUSG00000096696 | Zfp960 | 449000 [Gene Symbol: Zfp960] [Locus Tag: ] [Chromosome: 17] [Map Location: 17\|17 | 14.2123026 | 5.30E-05 | 0.000175 |
| 13274 | ENSMUSG00000026429 | Ube2t | 67196 [Gene Symbol: Ube2t] [Locus Tag: ] [Chromosome: 1] [Map Location: 1\|1 E4] [De | 14.06776438 | 6.77E-14 | 9.48E-13 |
| 10172 | ENSMUSG00000051234 | Rnf7 | 19823 [Gene Symbol: Rnf7] [Locus Tag: ] [Chromosome: 9] [Map Location: 9\|9 D2.3] [D | 13.85184161 | 5.68E-32 | 1.49E-29 |
| 5751 | ENSMUSG00000017404 | Rpl19 | 19921 [Gene Symbol: Rpl19] [Locus Tag: ] [Chromosome: 11] [Map Location: 11\|11 D] | 13.59130376 | 6.48E-08 | 3.54E-07 |
| 17802 | ENSMUSG00000029817 | Tra2a | 101214 [Gene Symbol: Tra2a] [Locus Tag: ] [Chromosome: 6] [Map Location: 6\|6 B2.3] | 13.58899686 | 2.20E-21 | 1.02E-19 |
| 8724 | ENSMUSG00000028563 | Tm2d1 | 94043 [Gene Symbol: Tm2d1] [Locus Tag: ] [Chromosome: 4] [Map Location: 4\|4 C6] [D | 13.54891492 | 8.23E-23 | 4.80E-21 |
| 9402 | ENSMUSG00000027832 | Ptx3 | 19288 [Gene Symbol: Ptx3] [Locus Tag: ] [Chromosome: 3] [Map Location: 3 E1\|3 30.2 | 13.51749735 | 6.18E-07 | 2.89E-06 |
| 17429 | ENSMUSG00000030759 | Far1 | 67420 [Gene Symbol: Far1] [Locus Tag: ] [Chromosome: 7] [Map Location: 7\|7 F2] [Desc | 13.51528129 | 6.44E-23 | 3.88E-21 |
| 325 | ENSMUSG00000031200 | Mtcp1 | 17763 [Gene Symbol: Mtcp1] [Locus Tag: ] [Chromosome: X] [Map Location: X\|X A7.3] [ | 13.49935641 | 2.02E-20 | 7.92E-19 |
| 10181 | ENSMUSG00000009207 | Lnp | 69605 [Gene Symbol: Lnp] [Locus Tag: ] [Chromosome: 2] [Map Location: 2\|2 C3] [Desc | 13.48053444 | 4.35E-33 | 1.45E-30 |
| 118 | ENSMUSG00000031327 | Chic1 | 12212 [Gene Symbol: Chic1] [Locus Tag: ] [Chromosome: X] [Map Location: X D\|X 42.3 | 13.38804219 | 1.88E-31 | 4.32E-29 |
| 11422 | ENSMUSG00000007613 | Tgfbr1 | 21812 [Gene Symbol: Tgfbr1] [Locus Tag: ] [Chromosome: 4] [Map Location: 4 B1\|4 26. | 13.37650041 | 6.86E-29 | 1.02E-26 |
| 4604 | ENSMUSG00000032551 | 1110059G10Rik | 66202 [Gene Symbol: 1110059G10Rik] [Locus Tag: ] [Chromosome: 9] [Map Location: | 13.32148885 | 3.25E-21 | 1.47E-19 |
| 15462 | ENSMUSG00000071172 | Srsf3 | 20383 [Gene Symbol: Srsf3] [Locus Tag: ] [Chromosome: 17] [Map Location: 17 A3.3\|17 | 13.27518028 | 2.34E-32 | 6.48E-30 |
| 17481 | ENSMUSG00000027677 | Ttc14 | 67120 [Gene Symbol: Ttc14] [Locus Tag: ] [Chromosome: 3] [Map Location: 3\|3 B] [Desc | 12.91953604 | 3.57E-20 | 1.36E-18 |
| 2711 | ENSMUSG00000060743 | Gm12657; H3f3c; H3 | 667250 [Gene Symbol: Gm12657] [Locus Tag: ] [Chromosome: 4] [Map Location: 4\|4 C | 12.77724795 | 3.93E-19 | 1.26E-17 |
| 12235 | ENSMUSG00000090110 | Cmc4; Mtcp1 | 105886298 [Gene Symbol: Cmc4] [Locus Tag: ] [Chromosome: X] [Map Location: ] [Desc | 12.66485425 | 7.72E-23 | 4.54E-21 |
| 12969 | ENSMUSG00000048040 | Arxes2 | 76976 [Gene Symbol: Arxes2] [Locus Tag: ] [Chromosome: X] [Map Location: X\|X F1] [De | 12.54005064 | 2.14E-29 | 3.47E-27 |

| | | | | | | |
|---|---|---|---|---|---|---|
| 3327 | ENSMUSG00000026435 | Slc45a3 | 212980 [Gene Symbol: Slc45a3] [Locus Tag: ] [Chromosome: 1] [Map Location: 1\|1 E4] | 12.46746908 | 2.18E-12 | 2.36E-11 |
| 12800 | ENSMUSG00000020570 | Sypl | 19027 [Gene Symbol: Sypl] [Locus Tag: ] [Chromosome: 12] [Map Location: 12 13.87 cM | 12.42491569 | 9.22E-29 | 1.32E-26 |
| 16220 | ENSMUSG00000014075 | Tctex1d2 | 66061 [Gene Symbol: Tctex1d2] [Locus Tag: ] [Chromosome: 16] [Map Location: 16\|16 | 12.33431279 | 4.83E-18 | 1.30E-16 |
| 17341 | ENSMUSG00000071722 | Spin4 | 270624 [Gene Symbol: Spin4] [Locus Tag: ] [Chromosome: X] [Map Location: X\|X C3] [D | 12.33287238 | 7.25E-16 | 1.39E-14 |
| 7146 | ENSMUSG00000022941 | Ripply3 | 170765 [Gene Symbol: Ripply3] [Locus Tag: ] [Chromosome: 16] [Map Location: 16\|16 | 12.29442368 | 6.87E-05 | 0.000222 |
| 5481 | ENSMUSG00000054717 | Hmgb2 | 97165 [Gene Symbol: Hmgb2] [Locus Tag: ] [Chromosome: 8] [Map Location: 8 B2\|8 29 | 12.23147022 | 2.92E-12 | 3.10E-11 |
| 18225 | ENSMUSG00000020993 | Trappc6b | 78232 [Gene Symbol: Trappc6b] [Locus Tag: ] [Chromosome: 12] [Map Location: 12\|12 | 12.2287733 | 1.56E-19 | 5.33E-18 |
| 5776 | ENSMUSG00000040459 | Arglu1 | 234023 [Gene Symbol: Arglu1] [Locus Tag: ] [Chromosome: 8] [Map Location: 8\|8 A1.1 | 12.17767452 | 7.09E-21 | 3.05E-19 |
| 726 | ENSMUSG00000075266 | Cenpw | 66311 [Gene Symbol: Cenpw] [Locus Tag: ] [Chromosome: 10] [Map Location: 10\|10 A4 | 12.16025372 | 3.97E-11 | 3.54E-10 |
| 13412 | ENSMUSG00000045382 | Cxcr4 | 12767 [Gene Symbol: Cxcr4] [Locus Tag: ] [Chromosome: 1] [Map Location: 1 E4\|1 56.4 | 12.14303339 | 3.15E-20 | 1.20E-18 |
| 516 | ENSMUSG00000020149 | Rab1a | 19324 [Gene Symbol: Rab1a] [Locus Tag: ] [Chromosome: 11] [Map Location: 11 A3.1\|1 | 12.12652558 | 2.86E-33 | 9.86E-31 |
| 1497 | ENSMUSG00000034334 | Fam151b | 73942 [Gene Symbol: Fam151b] [Locus Tag: ] [Chromosome: 13] [Map Location: 13\|13 | 12.11495936 | 8.13E-16 | 1.55E-14 |
| 12305 | ENSMUSG00000032381 | Fam96a | 68250 [Gene Symbol: Fam96a] [Locus Tag: ] [Chromosome: 9] [Map Location: 9\|9 D] [D | 12.10469805 | 3.71E-23 | 2.32E-21 |
| 845 | ENSMUSG00000070493 | Gm13202; Chchd2 | 433806 [Gene Symbol: Gm13202] [Locus Tag: ] [Chromosome: 4] [Map Location: 4\|4 E | 12.05850154 | 4.53E-24 | 3.25E-22 |
| 18580 | ENSMUSG00000022676 | Snai2 | 20583 [Gene Symbol: Snai2] [Locus Tag: ] [Chromosome: 16] [Map Location: 16 A1\|16 | 11.86557042 | 4.36E-09 | 2.87E-08 |
| 1968 | ENSMUSG00000021028 | Mbip | 217588 [Gene Symbol: Mbip] [Locus Tag: ] [Chromosome: 12] [Map Location: 12\|12 C1 | 11.86423657 | 1.49E-19 | 5.11E-18 |
| 2366 | ENSMUSG00000042271 | Nxt2 | 237082 [Gene Symbol: Nxt2] [Locus Tag: ] [Chromosome: X] [Map Location: X\|X F2] [De | 11.81944392 | 9.76E-23 | 5.62E-21 |
| 8625 | ENSMUSG00000002728 | Naa20 | 67877 [Gene Symbol: Naa20] [Locus Tag: ] [Chromosome: 2] [Map Location: 2 71.28 cM | 11.79813081 | 1.85E-21 | 8.73E-20 |
| 17362 | ENSMUSG00000031246 | Sh3bgrl | 56726 [Gene Symbol: Sh3bgrl] [Locus Tag: ] [Chromosome: X] [Map Location: X\|X D] [D | 11.79137615 | 2.19E-31 | 4.98E-29 |
| 12216 | ENSMUSG00000053178 | Mterf1b | 208595 [Gene Symbol: Mterf1b] [Locus Tag: ] [Chromosome: 5] [Map Location: 5 A1\|5] | 11.78542412 | 2.16E-09 | 1.48E-08 |
| 18500 | ENSMUSG00000049929 | Lpar4 | 78134 [Gene Symbol: Lpar4] [Locus Tag: ] [Chromosome: X] [Map Location: X\|X D] [Des | 11.77721821 | 1.07E-16 | 2.36E-15 |
| 12102 | ENSMUSG00000071180 | LOC102642895; LOC | 102642895 [Gene Symbol: LOC102642895] [Locus Tag: ] [Chromosome: 4] [Map Locati | 11.77428941 | 5.66E-19 | 1.75E-17 |
| 6668 | ENSMUSG00000073542 | Cep76 | 225659 [Gene Symbol: Cep76] [Locus Tag: ] [Chromosome: 18] [Map Location: 18\|18 E | 11.69685588 | 1.40E-24 | 1.08E-22 |
| 12873 | ENSMUSG00000028382 | Ptbp3 | 230257 [Gene Symbol: Ptbp3] [Locus Tag: ] [Chromosome: 4] [Map Location: 4\|4 B3] [D | 11.66040369 | 1.30E-31 | 3.09E-29 |
| 2612 | ENSMUSG00000050192 | Eif5a2 | 208691 [Gene Symbol: Eif5a2] [Locus Tag: ] [Chromosome: 3] [Map Location: 3\|3 A3] [D | 11.60846202 | 1.72E-28 | 2.45E-26 |
| 5501 | ENSMUSG00000007850 | Hnrnph1 | 59013 [Gene Symbol: Hnrnph1] [Locus Tag: ] [Chromosome: 11] [Map Location: 11\|11 | 11.58931504 | 7.20E-23 | 4.30E-21 |
| 10686 | ENSMUSG00000004885 | Crabp2 | 12904 [Gene Symbol: Crabp2] [Locus Tag: ] [Chromosome: 3] [Map Location: 3F1\|3] [D | 11.42473648 | 7.32E-11 | 6.27E-10 |
| 1656 | ENSMUSG00000021765 | Fst | 14313 [Gene Symbol: Fst] [Locus Tag: ] [Chromosome: 13] [Map Location: 13\|13 D2.2] | 11.35785323 | 2.07E-23 | 1.35E-21 |
| 9083 | ENSMUSG00000040520 | Manea | 242362 [Gene Symbol: Manea] [Locus Tag: ] [Chromosome: 4] [Map Location: 4\|4 A3] [ | 11.32584551 | 2.59E-30 | 5.01E-28 |
| 9349 | ENSMUSG00000026843 | Fubp3 | 320267 [Gene Symbol: Fubp3] [Locus Tag: ] [Chromosome: 2] [Map Location: 2\|2 B] [De | 11.32448305 | 4.94E-30 | 9.29E-28 |
| 1444 | ENSMUSG00000024646 | Cyb5a | 109672 [Gene Symbol: Cyb5a] [Locus Tag: ] [Chromosome: 18] [Map Location: 18 E4\|1 | 11.27763436 | 1.45E-25 | 1.29E-23 |
| 11989 | ENSMUSG00000063804 | Lin28b | 380669 [Gene Symbol: Lin28b] [Locus Tag: ] [Chromosome: 10] [Map Location: 10\|10 | 11.24207171 | 6.48E-20 | 2.38E-18 |
| 5290 | ENSMUSG00000028676 | Gm5446; Srsf10; LOC | 432730 [Gene Symbol: Gm5446] [Locus Tag: ] [Chromosome: 13] [Map Location: 13\|1 | 11.21016185 | 7.51E-27 | 7.82E-25 |
| 194 | ENSMUSG00000027641 | Rbl1 | 19650 [Gene Symbol: Rbl1] [Locus Tag: ] [Chromosome: 2] [Map Location: 2 H1\|2 78.0 | 11.19784649 | 1.17E-14 | 1.84E-13 |
| 299 | ENSMUSG00000050786 | Ccdc126 | 57895 [Gene Symbol: Ccdc126] [Locus Tag: ] [Chromosome: 6] [Map Location: 6\|6 B2.3 | 11.19560396 | 3.62E-22 | 1.89E-20 |
| 14392 | ENSMUSG00000023330 | Dtwd1 | 69185 [Gene Symbol: Dtwd1] [Locus Tag: ] [Chromosome: 2] [Map Location: 2\|2 F2] [D | 11.18407359 | 1.57E-22 | 8.85E-21 |
| 3107 | ENSMUSG00000030865 | Chp2 | 70261 [Gene Symbol: Chp2] [Locus Tag: ] [Chromosome: 7] [Map Location: 7\|7 F3] [Des | 11.13615155 | 2.85E-05 | 9.98E-05 |
| 744 | ENSMUSG00000019791 | Hint3 | 66847 [Gene Symbol: Hint3] [Locus Tag: ] [Chromosome: 10] [Map Location: 10\|10 A4] | 11.12154882 | 1.69E-17 | 4.12E-16 |
| 11849 | ENSMUSG00000073678 | Pgap1 | 241062 [Gene Symbol: Pgap1] [Locus Tag: ] [Chromosome: 1] [Map Location: 1\|1 C1.1] | 11.10340465 | 7.47E-31 | 1.60E-28 |
| 15353 | ENSMUSG00000022884 | Eif4a2 | 13682 [Gene Symbol: Eif4a2] [Locus Tag: ] [Chromosome: 16] [Map Location: 16 B1\|16 | 11.08610592 | 5.41E-28 | 6.81E-26 |
| 8152 | ENSMUSG00000021072 | Tmx1 | 72736 [Gene Symbol: Tmx1] [Locus Tag: ] [Chromosome: 12] [Map Location: 12\|12 C2] | 11.02323648 | 2.98E-27 | 3.27E-25 |
| 16691 | ENSMUSG00000035293 | G2e3 | 217558 [Gene Symbol: G2e3] [Locus Tag: ] [Chromosome: 12] [Map Location: 12\|12 C1 | 10.85730969 | 1.77E-18 | 5.10E-17 |
| 3178 | ENSMUSG00000054196 | Cthrc1 | 68588 [Gene Symbol: Cthrc1] [Locus Tag: ] [Chromosome: 15] [Map Location: 15\|15 C] | 10.84897834 | 2.72E-12 | 2.91E-11 |
| 3302 | ENSMUSG00000021832 | Psmc6 | 67089 [Gene Symbol: Psmc6] [Locus Tag: ] [Chromosome: 14] [Map Location: 14\|14 C1 | 10.79941823 | 3.55E-29 | 5.51E-27 |
| 5258 | ENSMUSG00000028229 | Rmdn1 | 66302 [Gene Symbol: Rmdn1] [Locus Tag: ] [Chromosome: 4] [Map Location: 4\|4 A3] [D | 10.79939098 | 1.24E-20 | 5.09E-19 |
| 4778 | ENSMUSG00000042834 | Nrep | 27528 [Gene Symbol: Nrep] [Locus Tag: ] [Chromosome: 18] [Map Location: 18\|18 B1] | 10.79916177 | 1.13E-31 | 2.73E-29 |
| 4676 | ENSMUSG00000089911 | Hiat1 | 15247 [Gene Symbol: Hiat1] [Locus Tag: ] [Chromosome: 3] [Map Location: 3\|3 G2] [De | 10.79534962 | 2.42E-30 | 4.85E-28 |
| 14428 | ENSMUSG00000029366 | Dck | 13178 [Gene Symbol: Dck] [Locus Tag: ] [Chromosome: 5] [Map Location: 5\|5 E2] [Desc | 10.7660235 | 4.38E-18 | 1.18E-16 |
| 964 | ENSMUSG00000017418 | Arl5b | 75869 [Gene Symbol: Arl5b] [Locus Tag: ] [Chromosome: 2] [Map Location: 2\|2 A2] [De | 10.74868613 | 1.21E-20 | 4.99E-19 |
| 1106 | ENSMUSG00000018846 | Pank3 | 211347 [Gene Symbol: Pank3] [Locus Tag: ] [Chromosome: 11] [Map Location: 11\|11 A | 10.72990028 | 3.18E-31 | 6.95E-29 |
| 8238 | ENSMUSG00000029385 | Ccng2 | 12452 [Gene Symbol: Ccng2] [Locus Tag: ] [Chromosome: 5] [Map Location: 5\|5 E3.3-F | 10.70769879 | 1.56E-19 | 5.33E-18 |
| 11619 | ENSMUSG00000021252 | 0610007P14Rik | 58520 [Gene Symbol: 0610007P14Rik] [Locus Tag: ] [Chromosome: 12] [Map Location: | 10.70524156 | 4.99E-27 | 5.28E-25 |
| 738 | ENSMUSG00000043542 | Zc2hc1a | 67306 [Gene Symbol: Zc2hc1a] [Locus Tag: ] [Chromosome: 3] [Map Location: 3\|3 A1] [ | 10.69573601 | 2.52E-30 | 4.94E-28 |
| 11441 | ENSMUSG00000027162 | Lin7c | 22343 [Gene Symbol: Lin7c] [Locus Tag: ] [Chromosome: 2] [Map Location: 2 E3\|2 56.6 | 10.67652291 | 5.36E-28 | 6.79E-26 |
| 6356 | ENSMUSG00000051147 | Nat2 | 17961 [Gene Symbol: Nat2] [Locus Tag: ] [Chromosome: 8] [Map Location: 8 33.38 cM\| | 10.64929309 | 0.000323 | 0.000925 |
| 13265 | ENSMUSG00000026154 | Sdhaf4 | 68002 [Gene Symbol: Sdhaf4] [Locus Tag: ] [Chromosome: 1] [Map Location: 1\|1 A5] [D | 10.64777535 | 2.01E-13 | 2.58E-12 |
| 2212 | ENSMUSG00000042682 | Selk | 80795 [Gene Symbol: Selk] [Locus Tag: ] [Chromosome: 14] [Map Location: 14\|14 B] [D | 10.58704523 | 9.76E-24 | 6.73E-22 |
| 11593 | ENSMUSG00000000581 | C1d | 57316 [Gene Symbol: C1d] [Locus Tag: ] [Chromosome: 11] [Map Location: 11\|11 A2] [ | 10.57545587 | 6.35E-26 | 6.06E-24 |
| 6274 | ENSMUSG00000000058 | Cav2 | 12390 [Gene Symbol: Cav2] [Locus Tag: ] [Chromosome: 6] [Map Location: 6\|6 A2] [Des | 10.57502948 | 4.95E-10 | 3.73E-09 |
| 7901 | ENSMUSG00000027811 | 4930579G24Rik | 75939 [Gene Symbol: 4930579G24Rik] [Locus Tag: ] [Chromosome: 3] [Map Location: | 10.52799529 | 2.00E-10 | 1.60E-09 |
| 4487 | ENSMUSG00000095990 | Zfp97 | 22759 [Gene Symbol: Zfp97] [Locus Tag: ] [Chromosome: 17] [Map Location: 17\|17 A3. | 10.48871493 | 5.33E-05 | 0.000176 |
| 17947 | ENSMUSG00000039097 | Rln1 | 19773 [Gene Symbol: Rln1] [Locus Tag: ] [Chromosome: 19] [Map Location: 19 C1\|19 2 | 10.48039939 | 1.80E-05 | 6.55E-05 |
| 16509 | ENSMUSG00000055531 | Cpsf6 | 432508 [Gene Symbol: Cpsf6] [Locus Tag: ] [Chromosome: 10] [Map Location: 10\|10 D | 10.4739419 | 3.17E-31 | 6.95E-29 |
| 10227 | ENSMUSG00000027132 | Katnbl1 | 72425 [Gene Symbol: Katnbl1] [Locus Tag: ] [Chromosome: 2] [Map Location: 2\|2 E4] [ | 10.46074094 | 2.14E-25 | 1.86E-23 |
| 1319 | ENSMUSG00000030042 | Pole4 | 66979 [Gene Symbol: Pole4] [Locus Tag: ] [Chromosome: 6] [Map Location: 6\|6 D1] [De | 10.43399886 | 8.11E-27 | 8.39E-25 |
| 4880 | ENSMUSG00000047694 | Yipf6 | 77929 [Gene Symbol: Yipf6] [Locus Tag: ] [Chromosome: X] [Map Location: X\|X C2] [Des | 10.43020907 | 2.41E-22 | 1.30E-20 |
| 16662 | ENSMUSG00000025939 | Ube2w | 66799 [Gene Symbol: Ube2w] [Locus Tag: ] [Chromosome: 1] [Map Location: 1\|1 A3] [D | 10.41795204 | 1.05E-27 | 1.24E-25 |
| 3428 | ENSMUSG00000042167 | Papd4 | 100715 [Gene Symbol: Papd4] [Locus Tag: ] [Chromosome: 13] [Map Location: 13\|13 C | 10.39064751 | 9.80E-23 | 5.63E-21 |
| 18467 | ENSMUSG00000018442 | Derl2 | 116891 [Gene Symbol: Derl2] [Locus Tag: ] [Chromosome: 11] [Map Location: 11 B4\|11 | 10.38525855 | 8.32E-26 | 7.76E-24 |
| 247 | ENSMUSG00000074754 | Gm561 | 228715 [Gene Symbol: Gm561] [Locus Tag: ] [Chromosome: 2] [Map Location: 2\|2 G1] | 10.32906519 | 1.17E-07 | 6.15E-07 |
| 10968 | ENSMUSG00000020948 | Klhl28 | 66689 [Gene Symbol: Klhl28] [Locus Tag: ] [Chromosome: 12] [Map Location: 12\|12 C1 | 10.31426376 | 1.86E-28 | 2.58E-26 |
| 3033 | ENSMUSG00000022338 | Eny2 | 223527 [Gene Symbol: Eny2] [Locus Tag: ] [Chromosome: 15] [Map Location: 15\|15 B3 | 10.29605869 | 2.46E-27 | 2.73E-25 |
| 8065 | ENSMUSG00000027957 | Slc35a3 | 229782 [Gene Symbol: Slc35a3] [Locus Tag: ] [Chromosome: 3] [Map Location: 3\|3 G1] | 10.28855464 | 5.55E-25 | 4.61E-23 |
| 5047 | ENSMUSG00000014813 | Stc1 | 20855 [Gene Symbol: Stc1] [Locus Tag: ] [Chromosome: 14] [Map Location: 14\|14 D2] [ | 10.27631763 | 3.69E-17 | 8.67E-16 |
| 1431 | ENSMUSG00000024795 | Kif20b | 240641 [Gene Symbol: Kif20b] [Locus Tag: ] [Chromosome: 19] [Map Location: 19\|19 C | 10.22053582 | 5.83E-13 | 6.95E-12 |
| 8290 | ENSMUSG00000039016 | Timm8b | 30057 [Gene Symbol: Timm8b] [Locus Tag: ] [Chromosome: 9] [Map Location: 9\|9 B] [D | 10.21050106 | 1.13E-17 | 2.82E-16 |
| 6529 | ENSMUSG00000040464 | Gtpbp10 | 207704 [Gene Symbol: Gtpbp10] [Locus Tag: ] [Chromosome: 5] [Map Location: 5\|5 A1 | 10.15153069 | 2.62E-15 | 4.55E-14 |
| 14253 | ENSMUSG00000036169 | Sostdc1 | 66042 [Gene Symbol: Sostdc1] [Locus Tag: ] [Chromosome: 12] [Map Location: 12\|12 B | 10.13371402 | 6.26E-13 | 7.45E-12 |
| 14707 | ENSMUSG00000091537 | Tma7 | 66167 [Gene Symbol: Tma7] [Locus Tag: ] [Chromosome: 9] [Map Location: 9\|9 F2] [Des | 10.07318606 | 3.25E-16 | 6.61E-15 |
| 7417 | ENSMUSG00000032293 | Ireb2 | 64602 [Gene Symbol: Ireb2] [Locus Tag: ] [Chromosome: 9] [Map Location: 9 B\|9 29.81 | 10.05921918 | 1.05E-29 | 1.84E-27 |
| 154 | ENSMUSG00000046032 | Snx12 | 55988 [Gene Symbol: Snx12] [Locus Tag: ] [Chromosome: X] [Map Location: X\|X C2] [De | 10.05758107 | 7.74E-28 | 9.60E-26 |
| 9476 | ENSMUSG00000033411 | Ctdspl2 | 329506 [Gene Symbol: Ctdspl2] [Locus Tag: ] [Chromosome: 2] [Map Location: 2 E5\|2 6 | 10.04854986 | 1.85E-28 | 2.58E-26 |
| 10496 | ENSMUSG00000028175 | Depdc1a | 76131 [Gene Symbol: Depdc1a] [Locus Tag: ] [Chromosome: 3] [Map Location: 3\|3 H4] | 10.03706703 | 2.09E-06 | 8.92E-06 |
| 6775 | ENSMUSG00000037361 | Sf3b6 | 66055 [Gene Symbol: Sf3b6] [Locus Tag: ] [Chromosome: 12] [Map Location: 12\|12 A1. | 10.00970233 | 1.05E-24 | 8.23E-23 |

| | | | | | | |
|---|---|---|---|---|---|---|
| 11914 | ENSMUSG00000026049 | Tex30 | 75623 [Gene Symbol: Tex30] [Locus Tag: ] [Chromosome: 1] [Map Location: 1\|1 C1] [De | 10.00075831 | 8.01E-12 | 7.99E-11 |
| 11662 | ENSMUSG00000020137 | Thap2 | 66816 [Gene Symbol: Thap2] [Locus Tag: ] [Chromosome: 10] [Map Location: 10\|10 D2 | 9.958409213 | 4.16E-17 | 9.72E-16 |
| 8150 | ENSMUSG00000094595 | Fsbp | 100503583 [Gene Symbol: Fsbp] [Locus Tag: ] [Chromosome: 4] [Map Location: 4\|4 5.1 | 9.915237073 | 9.46E-13 | 1.09E-11 |
| 12518 | ENSMUSG00000021113 | Snapc1 | 75627 [Gene Symbol: Snapc1] [Locus Tag: ] [Chromosome: 12] [Map Location: 12\|12 C | 9.896108393 | 4.97E-25 | 4.17E-23 |
| 2822 | ENSMUSG00000021189 | Atxn3 | 110616 [Gene Symbol: Atxn3] [Locus Tag: ] [Chromosome: 12] [Map Location: 12\|12 E] | 9.877274192 | 9.31E-27 | 9.53E-25 |
| 5218 | ENSMUSG00000021431 | Snrnp48 | 67797 [Gene Symbol: Snrnp48] [Locus Tag: ] [Chromosome: 13] [Map Location: 13\|13 | 9.874913633 | 4.34E-24 | 3.16E-22 |
| 12628 | ENSMUSG00000052681 | Rap1b | 215449 [Gene Symbol: Rap1b] [Locus Tag: ] [Chromosome: 10] [Map Location: 10\|10 D | 9.869312088 | 2.32E-25 | 2.00E-23 |
| 16876 | ENSMUSG00000040455 | Usp45 | 77593 [Gene Symbol: Usp45] [Locus Tag: ] [Chromosome: 4] [Map Location: 4\|4 A3] [De | 9.857655691 | 1.87E-27 | 2.11E-25 |
| 5808 | ENSMUSG00000031517 | Gpm6a | 234267 [Gene Symbol: Gpm6a] [Locus Tag: ] [Chromosome: 8] [Map Location: 8\|8 B3.2 | 9.854122862 | 6.24E-30 | 1.13E-27 |
| 15923 | ENSMUSG00000040123 | Zmym5 | 219105 [Gene Symbol: Zmym5] [Locus Tag: ] [Chromosome: 14] [Map Location: 14\|14 | 9.814592781 | 1.56E-25 | 1.37E-23 |
| 13662 | ENSMUSG00000022664 | Slc35a5 | 74102 [Gene Symbol: Slc35a5] [Locus Tag: ] [Chromosome: 16] [Map Location: 16 B5\|1 | 9.784769664 | 1.40E-25 | 1.25E-23 |
| 6752 | ENSMUSG00000037720 | Tmem33 | 67878 [Gene Symbol: Tmem33] [Locus Tag: ] [Chromosome: 5] [Map Location: 5\|5 D] [ | 9.77622943 | 1.46E-27 | 1.69E-25 |
| 293 | ENSMUSG00000073639 | Rab18 | 19330 [Gene Symbol: Rab18] [Locus Tag: ] [Chromosome: 18] [Map Location: 18 A1\|18 | 9.774714248 | 7.24E-29 | 1.07E-26 |
| 15374 | ENSMUSG00000057113 | Npm1 | 18148 [Gene Symbol: Npm1] [Locus Tag: ] [Chromosome: 11] [Map Location: 11\|11 A4 | 9.753850052 | 7.02E-28 | 8.77E-26 |
| 7345 | ENSMUSG00000039105 | Atp6v1g1 | 66290 [Gene Symbol: Atp6v1g1] [Locus Tag: ] [Chromosome: 4] [Map Location: 4\|4 C1] | 9.746075642 | 2.96E-21 | 1.35E-19 |
| 17468 | ENSMUSG00000026723 | Trdmt1 | 13434 [Gene Symbol: Trdmt1] [Locus Tag: ] [Chromosome: 2] [Map Location: 2\|2 A1] [ | 9.714606222 | 6.28E-19 | 1.92E-17 |
| 12937 | ENSMUSG00000035898 | Uba6 | 231380 [Gene Symbol: Uba6] [Locus Tag: ] [Chromosome: 5] [Map Location: 5\|5 E1] [De | 9.710210634 | 4.61E-27 | 4.94E-25 |
| 13215 | ENSMUSG00000016559 | H3f3b; LOC10264204 | 15081 [Gene Symbol: H3f3b] [Locus Tag: ] [Chromosome: 11] [Map Location: 11\|11 E2 | 9.701366173 | 5.47E-23 | 3.34E-21 |
| 17024 | ENSMUSG00000005813 | Metap1 | 75624 [Gene Symbol: Metap1] [Locus Tag: ] [Chromosome: 3] [Map Location: 3\|3 G3] [ | 9.700227861 | 1.79E-28 | 2.53E-26 |
| 15884 | ENSMUSG00000039361 | Picalm | 233489 [Gene Symbol: Picalm] [Locus Tag: ] [Chromosome: 7] [Map Location: 7 E1\|7 5 | 9.69264851 | 1.15E-20 | 4.77E-19 |
| 14909 | ENSMUSG00000027201 | Myef2 | 17876 [Gene Symbol: Myef2] [Locus Tag: ] [Chromosome: 2] [Map Location: 2 F1\|2 61. | 9.687263656 | 1.90E-29 | 3.10E-27 |
| 7101 | ENSMUSG00000079084 | Ccdc82 | 66396 [Gene Symbol: Ccdc82] [Locus Tag: ] [Chromosome: 9] [Map Location: 9\|9] [Des | 9.680153171 | 4.00E-19 | 1.28E-17 |
| 3175 | ENSMUSG00000006941 | Eif1b | 68969 [Gene Symbol: Eif1b] [Locus Tag: ] [Chromosome: 9] [Map Location: 9\|9 F4] [Des | 9.672647299 | 7.79E-29 | 1.13E-26 |
| 8304 | ENSMUSG00000056763 | Cspp1 | 211660 [Gene Symbol: Cspp1] [Locus Tag: ] [Chromosome: 1] [Map Location: 1\|1 A2] [ | 9.658341385 | 3.68E-18 | 1.00E-16 |
| 8977 | ENSMUSG00000036810 | Cnep1r1 | 382030 [Gene Symbol: Cnep1r1] [Locus Tag: ] [Chromosome: 8] [Map Location: 8\|8 C4 | 9.644345346 | 1.10E-25 | 9.97E-24 |
| 16697 | ENSMUSG00000062248 | Cks2 | 66197 [Gene Symbol: Cks2] [Locus Tag: ] [Chromosome: 13] [Map Location: 13\|13 A5] [ | 9.636671834 | 8.69E-11 | 7.34E-10 |
| 8116 | ENSMUSG00000041703 | Zic5 | 65100 [Gene Symbol: Zic5] [Locus Tag: ] [Chromosome: 14] [Map Location: 14\|14 E5] [ | 9.631133031 | 2.52E-23 | 1.62E-21 |
| 4925 | ENSMUSG00000020492 | Ska2 | 66140 [Gene Symbol: Ska2] [Locus Tag: ] [Chromosome: 11] [Map Location: 11\|11 C] [ | 9.604270437 | 1.13E-17 | 2.82E-16 |
| 1226 | ENSMUSG00000033790 | Tubgcp5 | 233276 [Gene Symbol: Tubgcp5] [Locus Tag: ] [Chromosome: 7] [Map Location: 7\|7 B5 | 9.600363488 | 6.08E-17 | 1.39E-15 |
| 8166 | ENSMUSG00000038607 | Gng10 | 14700 [Gene Symbol: Gng10] [Locus Tag: ] [Chromosome: 4] [Map Location: 4 B3\|4 32. | 9.538352988 | 4.79E-20 | 1.79E-18 |
| 15076 | ENSMUSG00000030790 | Adm | 11535 [Gene Symbol: Adm] [Locus Tag: ] [Chromosome: 7] [Map Location: 7 F1\|7 57.7 | 9.533055488 | 1.55E-08 | 9.39E-08 |
| 8886 | ENSMUSG00000078453 | Abracl | 73112 [Gene Symbol: Abracl] [Locus Tag: ] [Chromosome: 10] [Map Location: 10\|10 A3 | 9.514334939 | 2.64E-14 | 3.95E-13 |
| 13272 | ENSMUSG00000048834 | Vstm2a | 211739 [Gene Symbol: Vstm2a] [Locus Tag: ] [Chromosome: 11] [Map Location: 11\|11 | 9.479501703 | 2.50E-25 | 2.15E-23 |
| 12132 | ENSMUSG00000026753 | Ppp6c | 67857 [Gene Symbol: Ppp6c] [Locus Tag: ] [Chromosome: 2] [Map Location: 2\|2 B] [Des | 9.465681792 | 1.46E-26 | 1.47E-24 |
| 16259 | ENSMUSG00000001260 | Gabrg1 | 14405 [Gene Symbol: Gabrg1] [Locus Tag: ] [Chromosome: 5] [Map Location: 5 C3.1\|5 | 9.460625974 | 3.92E-16 | 7.84E-15 |
| 7637 | ENSMUSG00000000355 | Mcts1 | 68995 [Gene Symbol: Mcts1] [Locus Tag: ] [Chromosome: X] [Map Location: X\|X A2] [De | 9.421885419 | 2.38E-15 | 4.15E-14 |
| 13534 | ENSMUSG00000019942 | Cdk1 | 12534 [Gene Symbol: Cdk1] [Locus Tag: ] [Chromosome: 10] [Map Location: 10 B5.3\|1 | 9.409566228 | 1.10E-09 | 7.85E-09 |
| 9854 | ENSMUSG00000038760 | Trhr | 22045 [Gene Symbol: Trhr] [Locus Tag: ] [Chromosome: 15] [Map Location: 15 B3.2\|15 | 9.387726725 | 4.11E-13 | 5.00E-12 |
| 2522 | ENSMUSG00000027099 | Mtx2 | 53375 [Gene Symbol: Mtx2] [Locus Tag: ] [Chromosome: 2] [Map Location: 2\|2 D] [Desc | 9.385817899 | 7.08E-26 | 6.69E-24 |
| 14592 | ENSMUSG00000006262 | Mob1b | 68473 [Gene Symbol: Mob1b] [Locus Tag: ] [Chromosome: 5] [Map Location: 5\|5 E2] [D | 9.367759784 | 2.36E-27 | 2.65E-25 |
| 17557 | ENSMUSG00000028134 | Ptbp2 | 56195 [Gene Symbol: Ptbp2] [Locus Tag: ] [Chromosome: 3] [Map Location: 3\|3 G1] [De | 9.318406297 | 7.37E-25 | 5.96E-23 |
| 17050 | ENSMUSG00000038174 | Fam126b | 213056 [Gene Symbol: Fam126b] [Locus Tag: ] [Chromosome: 1] [Map Location: 1 C1.3 | 9.302765515 | 2.45E-27 | 2.73E-25 |
| 6316 | ENSMUSG00000034998 | Foxn2 | 14236 [Gene Symbol: Foxn2] [Locus Tag: ] [Chromosome: 17] [Map Location: 17\|17 E5 | 9.277462095 | 2.80E-27 | 3.09E-25 |
| 10246 | ENSMUSG00000038717 | Atp5l | 27425 [Gene Symbol: Atp5l] [Locus Tag: ] [Chromosome: 9] [Map Location: 9\|9 A5.2] [ | 9.273722739 | 2.97E-10 | 2.32E-09 |
| 15365 | ENSMUSG00000028211 | Trp53inp1 | 60599 [Gene Symbol: Trp53inp1] [Locus Tag: ] [Chromosome: 4] [Map Location: 4\|4 A1 | 9.2657394 | 8.11E-12 | 8.07E-11 |
| 17003 | ENSMUSG00000031290 | Lrch2 | 210297 [Gene Symbol: Lrch2] [Locus Tag: ] [Chromosome: X] [Map Location: X\|X F2] [D | 9.219995512 | 3.38E-17 | 7.97E-16 |
| 8944 | ENSMUSG00000030613 | Ccdc90b | 66365 [Gene Symbol: Ccdc90b] [Locus Tag: ] [Chromosome: 7] [Map Location: 7\|7 E1] [ | 9.219065342 | 6.94E-18 | 1.80E-16 |
| 14417 | ENSMUSG00000019936 | Epyc | 13516 [Gene Symbol: Epyc] [Locus Tag: ] [Chromosome: 10] [Map Location: 10 50.35 c | 9.164618829 | 0.000491 | 0.001356 |
| 10500 | ENSMUSG00000027676 | Ccdc39 | 51938 [Gene Symbol: Ccdc39] [Locus Tag: ] [Chromosome: 3] [Map Location: 3\|3 A3] [ | 9.163343178 | 2.67E-15 | 4.63E-14 |
| 3701 | ENSMUSG00000001999 | Blvra | 109778 [Gene Symbol: Blvra] [Locus Tag: ] [Chromosome: 2] [Map Location: 2 61.76 cM | 9.134714847 | 3.38E-16 | 6.83E-15 |
| 1672 | ENSMUSG00000046873 | Mbtps2 | 270669 [Gene Symbol: Mbtps2] [Locus Tag: ] [Chromosome: X] [Map Location: X\|X F4] | 9.119402996 | 2.03E-26 | 2.02E-24 |
| 6215 | ENSMUSG00000047139 | Cd24a | 12484 [Gene Symbol: Cd24a] [Locus Tag: ] [Chromosome: 10] [Map Location: 10 B2\|10 | 9.114814268 | 1.24E-23 | 8.36E-22 |
| 5489 | ENSMUSG00000049536 | Tceal1 | 237052 [Gene Symbol: Tceal1] [Locus Tag: ] [Chromosome: X] [Map Location: X\|X F1] [ | 9.097042936 | 2.31E-19 | 7.70E-18 |
| 6627 | ENSMUSG00000063406 | Tmed5 | 73130 [Gene Symbol: Tmed5] [Locus Tag: ] [Chromosome: 5] [Map Location: 5\|5 E5] [D | 9.09165881 | 2.03E-19 | 6.85E-18 |
| 4976 | ENSMUSG00000032328 | Tmem30a | 69981 [Gene Symbol: Tmem30a] [Locus Tag: ] [Chromosome: 9] [Map Location: 9 E1\|9 | 9.074090824 | 8.28E-28 | 1.01E-25 |
| 6390 | ENSMUSG00000051579 | Tceal8 | 66684 [Gene Symbol: Tceal8] [Locus Tag: ] [Chromosome: X] [Map Location: X\|X F1] [De | 9.050084688 | 1.12E-20 | 4.67E-19 |
| 16045 | ENSMUSG00000038535 | Zfp280d | 235469 [Gene Symbol: Zfp280d] [Locus Tag: ] [Chromosome: 9] [Map Location: 9\|9 D] [ | 9.042874422 | 9.38E-25 | 7.46E-23 |
| 11865 | ENSMUSG00000071359 | Tbpl1 | 237336 [Gene Symbol: Tbpl1] [Locus Tag: ] [Chromosome: 10] [Map Location: 10\|10 A | 9.02858407 | 5.32E-28 | 6.78E-26 |
| 10861 | ENSMUSG00000028256 | Odf2l | 52184 [Gene Symbol: Odf2l] [Locus Tag: ] [Chromosome: 3] [Map Location: 3 69.36 cM | 9.026972061 | 4.56E-15 | 7.59E-14 |
| 11210 | ENSMUSG00000021700 | Rab3c | 67295 [Gene Symbol: Rab3c] [Locus Tag: ] [Chromosome: 13] [Map Location: 13\|13 D2 | 9.026695201 | 7.64E-26 | 7.19E-24 |
| 12843 | ENSMUSG00000045205 | Dpy19l4 | 381510 [Gene Symbol: Dpy19l4] [Locus Tag: ] [Chromosome: 4] [Map Location: 4 A1\|4 | 8.999748847 | 6.80E-25 | 5.53E-23 |
| 7415 | ENSMUSG00000046434 | Hnrnpa1; Gm5803 | 15382 [Gene Symbol: Hnrnpa1] [Locus Tag: ] [Chromosome: 15] [Map Location: 15 F3\| | 8.999527616 | 4.71E-16 | 9.29E-15 |
| 9843 | ENSMUSG00000028986 | Klhl7 | 52323 [Gene Symbol: Klhl7] [Locus Tag: ] [Chromosome: 5] [Map Location: 5 A3\|5 10.6 | 8.969168266 | 4.17E-27 | 4.48E-25 |
| 3031 | ENSMUSG00000000600 | Krit1 | 79264 [Gene Symbol: Krit1] [Locus Tag: ] [Chromosome: 5] [Map Location: 5 A1\|5 2.26 | 8.960236833 | 9.64E-19 | 2.86E-17 |
| 9196 | ENSMUSG00000027822 | Slc33a1 | 11416 [Gene Symbol: Slc33a1] [Locus Tag: ] [Chromosome: 3] [Map Location: 3\|3 E1-E | 8.956163142 | 1.03E-23 | 7.11E-22 |
| 3533 | ENSMUSG00000027828 | Ssr3 | 67437 [Gene Symbol: Ssr3] [Locus Tag: ] [Chromosome: 3] [Map Location: 3\|3 E1] [Desc | 8.955563228 | 9.68E-27 | 9.85E-25 |
| 11455 | ENSMUSG00000020415 | Pttg1 | 30939 [Gene Symbol: Pttg1] [Locus Tag: ] [Chromosome: 11] [Map Location: 11\|11 A5] | 8.955263318 | 5.54E-11 | 4.84E-10 |
| 11838 | ENSMUSG00000031433 | Rbm41 | 237073 [Gene Symbol: Rbm41] [Locus Tag: ] [Chromosome: X] [Map Location: X\|X F1] [ | 8.95457006 | 2.01E-24 | 1.53E-22 |
| 16408 | ENSMUSG00000001127 | Araf | 11836 [Gene Symbol: Araf] [Locus Tag: ] [Chromosome: X] [Map Location: X 16.3 cM\|X | 8.933623063 | 3.46E-22 | 1.82E-20 |
| 11998 | ENSMUSG00000024645 | Timm21 | 67105 [Gene Symbol: Timm21] [Locus Tag: ] [Chromosome: 18] [Map Location: 18\|18 | 8.929430863 | 6.72E-16 | 1.30E-14 |
| 5164 | ENSMUSG00000090553 | Snrpe | 20643 [Gene Symbol: Snrpe] [Locus Tag: ] [Chromosome: 1] [Map Location: 1\|1 E4] [De | 8.922911534 | 1.47E-13 | 1.94E-12 |
| 8743 | ENSMUSG00000029283 | Cdc7 | 12545 [Gene Symbol: Cdc7] [Locus Tag: ] [Chromosome: 5] [Map Location: 5\|5 E] [Desc | 8.918969267 | 2.42E-24 | 1.80E-22 |
| 14310 | ENSMUSG00000038217 | Tlcd2 | 380712 [Gene Symbol: Tlcd2] [Locus Tag: ] [Chromosome: 11] [Map Location: 11\|11 B | 8.887819631 | 4.79E-14 | 6.87E-13 |
| 13260 | ENSMUSG00000057278 | Snrpg | 68011 [Gene Symbol: Snrpg] [Locus Tag: ] [Chromosome: 6] [Map Location: 6\|6 B1] [De | 8.864727354 | 6.22E-12 | 6.31E-11 |
| 2389 | ENSMUSG00000000560 | Gabra2 | 14395 [Gene Symbol: Gabra2] [Locus Tag: ] [Chromosome: 5] [Map Location: 5 C3.1\|5 | 8.845112976 | 3.66E-16 | 7.37E-15 |
| 3478 | ENSMUSG00000042742 | B630005N14Rik | 101148 [Gene Symbol: B630005N14Rik] [Locus Tag: ] [Chromosome: 6] [Map Location: | 8.843518033 | 1.20E-25 | 1.08E-23 |
| 9309 | ENSMUSG00000012405 | Rpl15 | 66480 [Gene Symbol: Rpl15] [Locus Tag: ] [Chromosome: 14] [Map Location: 14 A2\|14 | 8.790828953 | 2.24E-24 | 1.69E-22 |
| 5538 | ENSMUSG00000039899 | Fgl2 | 14190 [Gene Symbol: Fgl2] [Locus Tag: ] [Chromosome: 5] [Map Location: 5 A3\|5 9.83 | 8.787885223 | 6.34E-08 | 3.47E-07 |
| 17902 | ENSMUSG00000063253 | Scoc | 56367 [Gene Symbol: Scoc] [Locus Tag: ] [Chromosome: 8] [Map Location: 8\|8 C2] [Des | 8.783233744 | 1.07E-21 | 5.20E-20 |
| 10070 | ENSMUSG00000037490 | Slc2a12 | 353169 [Gene Symbol: Slc2a12] [Locus Tag: ] [Chromosome: 10] [Map Location: 10\|10 | 8.77632861 | 4.89E-22 | 2.51E-20 |
| 9853 | ENSMUSG00000021520 | Uqcrb | 67530 [Gene Symbol: Uqcrb] [Locus Tag: ] [Chromosome: 13] [Map Location: 13\|13 B3 | 8.775842386 | 3.25E-14 | 4.79E-13 |
| 15098 | ENSMUSG00000009549 | LOC102642637; Srp1 | 102642637 [Gene Symbol: LOC102642637] [Locus Tag: ] [Chromosome: 2] [Map Locati | 8.771396218 | 9.73E-22 | 4.76E-20 |
| 14443 | ENSMUSG00000020362 | Cnot6 | 104625 [Gene Symbol: Cnot6] [Locus Tag: ] [Chromosome: 11] [Map Location: 11\|11 B | 8.726123628 | 4.79E-19 | 1.50E-17 |
| 16922 | ENSMUSG00000059878 | Zfp422 | 67255 [Gene Symbol: Zfp422] [Locus Tag: ] [Chromosome: 6] [Map Location: 6\|6 E3] [D | 8.724128513 | 7.75E-19 | 2.34E-17 |

| | | | | | | |
|---|---|---|---|---|---|---|
| 10802 | ENSMUSG00000063145 | Bbs5 | 72569 [Gene Symbol: Bbs5] [Locus Tag: ] [Chromosome: 2] [Map Location: 2\|2 C3] [Des | 8.722105972 | 9.61E-15 | 1.53E-13 |
| 4847 | ENSMUSG00000073007 | Fam46d | 213449 [Gene Symbol: Fam46d] [Locus Tag: ] [Chromosome: X] [Map Location: X\|X D] [ | 8.70578602 | 1.17E-15 | 2.17E-14 |
| 88 | ENSMUSG00000024261 | Syt4 | 20983 [Gene Symbol: Syt4] [Locus Tag: ] [Chromosome: 18] [Map Location: 18 B1\|18 1 | 8.693844625 | 1.33E-27 | 1.56E-25 |
| 15826 | ENSMUSG00000098557 | Kctd12 | 239217 [Gene Symbol: Kctd12] [Locus Tag: ] [Chromosome: 14] [Map Location: 14\|14 | 8.682293974 | 3.97E-28 | 5.17E-26 |
| 2406 | ENSMUSG00000029920 | Smarcad1 | 13990 [Gene Symbol: Smarcad1] [Locus Tag: ] [Chromosome: 6] [Map Location: 6 C1\|6 | 8.662409253 | 1.67E-26 | 1.67E-24 |
| 1061 | ENSMUSG00000020863 | Luc7l3 | 67684 [Gene Symbol: Luc7l3] [Locus Tag: ] [Chromosome: 11] [Map Location: 11\|11 C] | 8.657947527 | 1.00E-18 | 2.97E-17 |
| 8557 | ENSMUSG00000038622 | Med30 | 69790 [Gene Symbol: Med30] [Locus Tag: ] [Chromosome: 15] [Map Location: 15\|15 C] | 8.643193202 | 2.31E-22 | 1.25E-20 |
| 9982 | ENSMUSG00000031232 | Magt1 | 67075 [Gene Symbol: Magt1] [Locus Tag: ] [Chromosome: X] [Map Location: X\|X D] [De | 8.642758869 | 3.29E-16 | 6.66E-15 |
| 1050 | ENSMUSG00000028293 | Slc35a1 | 24060 [Gene Symbol: Slc35a1] [Locus Tag: ] [Chromosome: 4] [Map Location: 4\|4 A5] [ | 8.631856365 | 5.13E-22 | 2.61E-20 |
| 2264 | ENSMUSG00000075229 | Ccdc58 | 381045 [Gene Symbol: Ccdc58] [Locus Tag: ] [Chromosome: 16] [Map Location: 16\|16 | 8.621578883 | 3.60E-18 | 9.85E-17 |
| 184 | ENSMUSG00000006715 | Gmnn | 57441 [Gene Symbol: Gmnn] [Locus Tag: ] [Chromosome: 13] [Map Location: 13\|13 A3 | 8.620349734 | 7.49E-05 | 0.000241 |
| 9207 | ENSMUSG00000027829 | Ccnl1 | 56706 [Gene Symbol: Ccnl1] [Locus Tag: ] [Chromosome: 3] [Map Location: 3\|3 E1] [De | 8.603651123 | 3.58E-24 | 2.61E-22 |
| 10772 | ENSMUSG00000021712 | Trim23 | 81003 [Gene Symbol: Trim23] [Locus Tag: ] [Chromosome: 13] [Map Location: 13\|13 D | 8.598204453 | 3.06E-20 | 1.17E-18 |
| 14171 | ENSMUSG00000021831 | Ero1l | 50527 [Gene Symbol: Ero1l] [Locus Tag: ] [Chromosome: 14] [Map Location: 14\|14 C-D | 8.586668842 | 8.54E-24 | 5.91E-22 |
| 7404 | ENSMUSG00000035762 | Tmem161b | 72745 [Gene Symbol: Tmem161b] [Locus Tag: ] [Chromosome: 13] [Map Location: 13\| | 8.577857665 | 1.55E-23 | 1.03E-21 |
| 18509 | ENSMUSG00000033805 | Ephx4 | 384214 [Gene Symbol: Ephx4] [Locus Tag: ] [Chromosome: 5] [Map Location: 5\|5 E5] [D | 8.559406715 | 1.65E-10 | 1.34E-09 |
| 13668 | ENSMUSG00000029755 | Dlx5 | 13395 [Gene Symbol: Dlx5] [Locus Tag: ] [Chromosome: 6] [Map Location: 6 A1\|6 2.83 | 8.553401592 | 1.37E-15 | 2.50E-14 |
| 5056 | ENSMUSG00000020423 | Btg2 | 12227 [Gene Symbol: Btg2] [Locus Tag: ] [Chromosome: 1] [Map Location: 1 E4\|1 58.1 | 8.552385051 | 4.94E-11 | 4.35E-10 |
| 18072 | ENSMUSG00000032449 | Slc25a36 | 192287 [Gene Symbol: Slc25a36] [Locus Tag: ] [Chromosome: 9] [Map Location: 9\|9 E3 | 8.543810849 | 2.97E-26 | 2.92E-24 |
| 1749 | ENSMUSG00000047227 | Gm527 | 217648 [Gene Symbol: Gm527] [Locus Tag: ] [Chromosome: 12] [Map Location: 12\|12 | 8.524318832 | 2.60E-13 | 3.27E-12 |
| 6225 | ENSMUSG00000029823 | Luc7l2 | 192196 [Gene Symbol: Luc7l2] [Locus Tag: ] [Chromosome: 6] [Map Location: 6\|6 B1] [ | 8.49981209 | 2.50E-19 | 8.26E-18 |
| 9192 | ENSMUSG00000030291 | Med21 | 108098 [Gene Symbol: Med21] [Locus Tag: ] [Chromosome: 6] [Map Location: 6 G3\|6 7 | 8.495387935 | 5.93E-15 | 9.79E-14 |
| 12089 | ENSMUSG00000053475 | Tnfaip6 | 21930 [Gene Symbol: Tnfaip6] [Locus Tag: ] [Chromosome: 2] [Map Location: 2\|2 C1.1] | 8.48608576 | 4.53E-07 | 2.16E-06 |
| 17273 | ENSMUSG00000059005 | Hnrnpa3 | 229279 [Gene Symbol: Hnrnpa3] [Locus Tag: ] [Chromosome: 2] [Map Location: 2\|2 A7 | 8.484432345 | 9.67E-23 | 5.59E-21 |
| 8095 | ENSMUSG00000017778 | LOC102642884; Cox | 102642884 [Gene Symbol: LOC102642884] [Locus Tag: ] [Chromosome: 2] [Map Locati | 8.473733877 | 4.85E-08 | 2.71E-07 |
| 15889 | ENSMUSG00000072623 | Zfp9 | 22750 [Gene Symbol: Zfp9] [Locus Tag: ] [Chromosome: 6] [Map Location: 6\|6 F1] [Desc | 8.460389991 | 5.69E-18 | 1.49E-16 |
| 3694 | ENSMUSG00000025626 | Phf6 | 70998 [Gene Symbol: Phf6] [Locus Tag: ] [Chromosome: X] [Map Location: X\|X A4] [Des | 8.458584492 | 1.62E-20 | 6.53E-19 |
| 17387 | ENSMUSG00000031242 | Chmp1b; 2610002M | 67064 [Gene Symbol: Chmp1b] [Locus Tag: ] [Chromosome: 18] [Map Location: 18\|18 | 8.446013357 | 4.92E-18 | 1.32E-16 |
| 1342 | ENSMUSG00000033792 | Atp7a | 11977 [Gene Symbol: Atp7a] [Locus Tag: ] [Chromosome: X] [Map Location: X D\|X 47.3 | 8.441332027 | 2.90E-18 | 8.04E-17 |
| 16997 | ENSMUSG00000072980 | LOC100862454; LOC | 100862454 [Gene Symbol: LOC100862454] [Locus Tag: ] [Chromosome: 17] [Map Loca | 8.424312132 | 1.97E-06 | 8.44E-06 |
| 16056 | ENSMUSG00000031367 | Ap1s2 | 108012 [Gene Symbol: Ap1s2] [Locus Tag: ] [Chromosome: X] [Map Location: X\|X F5] [D | 8.393600319 | 1.91E-17 | 4.63E-16 |
| 14039 | ENSMUSG00000036676 | Tmtc3 | 237500 [Gene Symbol: Tmtc3] [Locus Tag: ] [Chromosome: 10] [Map Location: 10\|10 D | 8.387982398 | 3.50E-14 | 5.12E-13 |
| 17820 | ENSMUSG00000027387 | Zc3h8 | 57432 [Gene Symbol: Zc3h8] [Locus Tag: ] [Chromosome: 2] [Map Location: 2\|2 F1] [De | 8.373647579 | 6.78E-13 | 8.03E-12 |
| 6640 | ENSMUSG00000048970 | C1galt1c1 | 59048 [Gene Symbol: C1galt1c1] [Locus Tag: ] [Chromosome: X] [Map Location: X\|X A2 | 8.337663661 | 1.97E-14 | 3.00E-13 |
| 5785 | ENSMUSG00000022974 | Paxbp1 | 67367 [Gene Symbol: Paxbp1] [Locus Tag: ] [Chromosome: 16] [Map Location: 16\|16 C | 8.284979288 | 4.44E-19 | 1.41E-17 |
| 3892 | ENSMUSG00000037710 | Cisd1 | 52637 [Gene Symbol: Cisd1] [Locus Tag: ] [Chromosome: 10] [Map Location: 10 B5.3\|1 | 8.277980686 | 7.48E-22 | 3.74E-20 |
| 105 | ENSMUSG00000035183 | Slc24a5 | 317750 [Gene Symbol: Slc24a5] [Locus Tag: ] [Chromosome: 2] [Map Location: 2\|2 F1] | 8.267804585 | 1.49E-25 | 1.32E-23 |
| 14238 | ENSMUSG00000021704 | Mtx3 | 382793 [Gene Symbol: Mtx3] [Locus Tag: ] [Chromosome: 13] [Map Location: 13\|13 C3 | 8.266684803 | 6.57E-25 | 5.39E-23 |
| 10033 | ENSMUSG00000061762 | Tac1 | 21333 [Gene Symbol: Tac1] [Locus Tag: ] [Chromosome: 6] [Map Location: 6 A1\|6 3.31 | 8.259470277 | 1.06E-14 | 1.67E-13 |
| 13705 | ENSMUSG00000019917 | Sept10 | 103080 [Gene Symbol: Sept10] [Locus Tag: ] [Chromosome: 10] [Map Location: 10\|10 | 8.255787037 | 6.21E-23 | 3.75E-21 |
| 10543 | ENSMUSG00000024383 | Map3k2 | 26405 [Gene Symbol: Map3k2] [Locus Tag: ] [Chromosome: 18] [Map Location: 18\|18 | 8.248297712 | 9.07E-27 | 9.33E-25 |
| 14664 | ENSMUSG00000028572 | Hook1 | 77963 [Gene Symbol: Hook1] [Locus Tag: ] [Chromosome: 4] [Map Location: 4 C5\|4 44. | 8.246659747 | 4.54E-16 | 8.99E-15 |
| 3093 | ENSMUSG00000020988 | L2hgdh | 217666 [Gene Symbol: L2hgdh] [Locus Tag: ] [Chromosome: 12] [Map Location: 12\|12 | 8.239915379 | 3.23E-19 | 1.05E-17 |
| 16060 | ENSMUSG00000016409 | Nkap | 67050 [Gene Symbol: Nkap] [Locus Tag: ] [Chromosome: X] [Map Location: X\|X A3.3] [D | 8.194549638 | 4.56E-24 | 3.26E-22 |
| 18277 | ENSMUSG00000044966 | Fbxo48 | 319701 [Gene Symbol: Fbxo48] [Locus Tag: ] [Chromosome: 11] [Map Location: 11\|11 | 8.178915628 | 5.13E-05 | 0.00017 |
| 17364 | ENSMUSG00000003721 | Insig2 | 72999 [Gene Symbol: Insig2] [Locus Tag: ] [Chromosome: 1] [Map Location: 1\|1 E2] [De | 8.169931064 | 1.39E-23 | 9.31E-22 |
| 387 | ENSMUSG00000037458 | Azin1 | 54375 [Gene Symbol: Azin1] [Locus Tag: ] [Chromosome: 15] [Map Location: 15\|15 C] [ | 8.143542242 | 4.74E-27 | 5.04E-25 |
| 7187 | ENSMUSG00000050912 | Tmem123 | 71929 [Gene Symbol: Tmem123] [Locus Tag: ] [Chromosome: 9] [Map Location: 9\|9 A1 | 8.121200612 | 2.41E-13 | 3.04E-12 |
| 10326 | ENSMUSG00000020849 | Ywhae | 22627 [Gene Symbol: Ywhae] [Locus Tag: ] [Chromosome: 11] [Map Location: 11 45.92 | 8.117131518 | 1.07E-26 | 1.08E-24 |
| 16558 | ENSMUSG00000021733 | Slc4a7 | 218756 [Gene Symbol: Slc4a7] [Locus Tag: ] [Chromosome: 14] [Map Location: 14\|14 | 8.114650667 | 3.57E-25 | 3.02E-23 |
| 181 | ENSMUSG00000025860 | Xiap | 11798 [Gene Symbol: Xiap] [Locus Tag: ] [Chromosome: X] [Map Location: X\|X A3-A5] [ | 8.113598757 | 7.93E-24 | 5.51E-22 |
| 14644 | ENSMUSG00000025894 | Aasdhppt | 67618 [Gene Symbol: Aasdhppt] [Locus Tag: ] [Chromosome: 9] [Map Location: 9\|9 A1] | 8.111485788 | 3.59E-22 | 1.88E-20 |
| 18337 | ENSMUSG00000069049 | Eif2s3y | 26908 [Gene Symbol: Eif2s3y] [Locus Tag: ] [Chromosome: Y] [Map Location: Y\|Y A1] [De | 8.099601871 | 1.88E-11 | 1.77E-10 |
| 16652 | ENSMUSG00000025979 | Mob4 | 19070 [Gene Symbol: Mob4] [Locus Tag: ] [Chromosome: 1] [Map Location: 1\|1 C1] [De | 8.089348562 | 1.71E-22 | 9.43E-21 |
| 14907 | ENSMUSG00000028420 | Tmem38b | 52076 [Gene Symbol: Tmem38b] [Locus Tag: ] [Chromosome: 4] [Map Location: 4 B2\|4 | 8.086678487 | 1.86E-19 | 6.28E-18 |
| 7705 | ENSMUSG00000024072 | Yipf4 | 67864 [Gene Symbol: Yipf4] [Locus Tag: ] [Chromosome: 17] [Map Location: 17\|17 E3] | 8.0704627 | 9.23E-13 | 1.07E-11 |
| 1622 | ENSMUSG00000074794 | Arrdc3 | 105171 [Gene Symbol: Arrdc3] [Locus Tag: ] [Chromosome: 13] [Map Location: 13\|13 | 8.055527314 | 2.25E-25 | 1.95E-23 |
| 14908 | ENSMUSG00000028243 | Ubxn2b | 68053 [Gene Symbol: Ubxn2b] [Locus Tag: ] [Chromosome: 4] [Map Location: 4\|4 A1] [ | 8.048566029 | 3.02E-21 | 1.37E-19 |
| 7093 | ENSMUSG00000041685 | Fcho2 | 218503 [Gene Symbol: Fcho2] [Locus Tag: ] [Chromosome: 13] [Map Location: 13\|13 D | 8.043737866 | 1.09E-17 | 2.74E-16 |
| 6004 | ENSMUSG00000025544 | Tm9sf2 | 68059 [Gene Symbol: Tm9sf2] [Locus Tag: ] [Chromosome: 14] [Map Location: 14 E5\|1 | 8.037164043 | 5.27E-24 | 3.71E-22 |
| 5945 | ENSMUSG00000009470 | Tnpo1 | 238799 [Gene Symbol: Tnpo1] [Locus Tag: ] [Chromosome: 13] [Map Location: 13 D1\|1 | 8.029010698 | 6.31E-26 | 6.06E-24 |
| 14904 | ENSMUSG00000047996 | Prrg1 | 546336 [Gene Symbol: Prrg1] [Locus Tag: ] [Chromosome: X] [Map Location: X\|X B] [De | 8.01662516 | 7.08E-16 | 1.36E-14 |
| 9070 | ENSMUSG00000028979 | Masp2 | 17175 [Gene Symbol: Masp2] [Locus Tag: ] [Chromosome: 4] [Map Location: 4\|4 E1] [De | 7.967997337 | 7.86E-22 | 3.91E-20 |
| 12474 | ENSMUSG00000045730 | Adrb2 | 11555 [Gene Symbol: Adrb2] [Locus Tag: ] [Chromosome: 18] [Map Location: 18 E1\|18 | 7.966877882 | 2.45E-06 | 1.03E-05 |
| 6066 | ENSMUSG00000090862 | LOC102642137; Rps1 | 102642137 [Gene Symbol: LOC102642137] [Locus Tag: ] [Chromosome: Un] [Map Loca | 7.962684531 | 0.000626 | 0.001695 |
| 11518 | ENSMUSG00000034732 | Pabpc5 | 93728 [Gene Symbol: Pabpc5] [Locus Tag: ] [Chromosome: X] [Map Location: X\|X D] [De | 7.960767185 | 6.86E-14 | 9.58E-13 |
| 8268 | ENSMUSG00000000869 | Il4 | 16189 [Gene Symbol: Il4] [Locus Tag: ] [Chromosome: 11] [Map Location: 11 B1.3\|11 3 | 7.95864581 | 4.39E-21 | 1.95E-19 |
| 2419 | ENSMUSG00000033543 | Gtf2a2 | 235459 [Gene Symbol: Gtf2a2] [Locus Tag: ] [Chromosome: 9] [Map Location: 9\|9 D] [D | 7.956372979 | 4.01E-21 | 1.79E-19 |
| 8644 | ENSMUSG00000042043 | Tbca | 21371 [Gene Symbol: Tbca] [Locus Tag: ] [Chromosome: 13] [Map Location: 13\|13 D1] | 7.940834211 | 3.70E-14 | 5.39E-13 |
| 16893 | ENSMUSG00000003929 | Zfp81 | 224694 [Gene Symbol: Zfp81] [Locus Tag: ] [Chromosome: 17] [Map Location: 17\|17 B | 7.94013619 | 2.45E-19 | 8.13E-18 |
| 7624 | ENSMUSG00000004642 | Slbp | 20492 [Gene Symbol: Slbp] [Locus Tag: ] [Chromosome: 5] [Map Location: 5\|5 B2] [Des | 7.931386535 | 3.42E-19 | 1.11E-17 |
| 4538 | ENSMUSG00000047466 | 8030462N17Rik | 212163 [Gene Symbol: 8030462N17Rik] [Locus Tag: ] [Chromosome: 18] [Map Locatio | 7.925825451 | 5.84E-18 | 1.53E-16 |
| 13938 | ENSMUSG00000040331 | Nsmce4a | 67872 [Gene Symbol: Nsmce4a] [Locus Tag: ] [Chromosome: 7] [Map Location: 7\|7 F3] | 7.896657246 | 7.57E-23 | 4.49E-21 |
| 2380 | ENSMUSG00000028156 | Eif4e | 13684 [Gene Symbol: Eif4e] [Locus Tag: ] [Chromosome: 3] [Map Location: 3 G3-H1\|3 6 | 7.879751357 | 3.17E-24 | 2.33E-22 |
| 16646 | ENSMUSG00000026771 | Spopl | 76857 [Gene Symbol: Spopl] [Locus Tag: ] [Chromosome: 2] [Map Location: 2\|2 A3] [De | 7.866171861 | 7.73E-22 | 3.86E-20 |
| 12171 | ENSMUSG00000003923 | Tfam | 21780 [Gene Symbol: Tfam] [Locus Tag: ] [Chromosome: 10] [Map Location: 10 B5\|10 3 | 7.864823435 | 5.12E-22 | 2.61E-20 |
| 6613 | ENSMUSG00000091736 | Yy2 | 100073351 [Gene Symbol: Yy2] [Locus Tag: ] [Chromosome: X] [Map Location: X F4\|X] [ | 7.841862454 | 1.28E-12 | 1.44E-11 |
| 8172 | ENSMUSG00000087651 | 1500009L16Rik | 69784 [Gene Symbol: 1500009L16Rik] [Locus Tag: ] [Chromosome: 10] [Map Location: | 7.833107395 | 3.06E-13 | 3.79E-12 |
| 12454 | ENSMUSG00000007872 | Id3 | 15903 [Gene Symbol: Id3] [Locus Tag: ] [Chromosome: 4] [Map Location: 4 D3\|4 68.34 | 7.829573386 | 2.23E-09 | 1.53E-08 |
| 17665 | ENSMUSG00000064061 | Dzip3 | 224170 [Gene Symbol: Dzip3] [Locus Tag: ] [Chromosome: 16] [Map Location: 16\|16 B | 7.79263513 | 5.85E-17 | 1.34E-15 |
| 9483 | ENSMUSG00000074093 | Svip | 75744 [Gene Symbol: Svip] [Locus Tag: ] [Chromosome: 7] [Map Location: 7\|7 B4] [Desc | 7.78684891 | 1.25E-20 | 5.15E-19 |
| 10759 | ENSMUSG00000033157 | Abhd10 | 213012 [Gene Symbol: Abhd10] [Locus Tag: ] [Chromosome: 16] [Map Location: 16\|16 | 7.785445762 | 9.22E-21 | 3.89E-19 |
| 11594 | ENSMUSG00000041180 | Hectd2 | 226098 [Gene Symbol: Hectd2] [Locus Tag: ] [Chromosome: 19] [Map Location: 19\|19 | 7.770793742 | 2.10E-23 | 1.37E-21 |
| 332 | ENSMUSG00000025815 | Dhtkd1 | 209692 [Gene Symbol: Dhtkd1] [Locus Tag: ] [Chromosome: 2] [Map Location: 2\|2 A1] | 7.767257556 | 1.22E-21 | 5.87E-20 |

| | | | | | | |
|---|---|---|---|---|---|---|
| 6278 | ENSMUSG00000047554 | Tmem41b | 233724 [Gene Symbol: Tmem41b] [Locus Tag: ] [Chromosome: 7] [Map Location: 7 F1 | | 7.766877105 | 1.13E-17 | 2.82E-16 |
| 14939 | ENSMUSG00000042138 | Msantd2 | 235184 [Gene Symbol: Msantd2] [Locus Tag: ] [Chromosome: 9] [Map Location: 9 \| 9 A4 | 7.758760069 | 1.16E-16 | 2.53E-15 |
| 14990 | ENSMUSG00000024256 | Adcyap1 | 11516 [Gene Symbol: Adcyap1] [Locus Tag: ] [Chromosome: 17] [Map Location: 17 \| 17 | 7.736971395 | 3.73E-15 | 6.29E-14 |
| 13669 | ENSMUSG00000022978 | Mis18a | 66578 [Gene Symbol: Mis18a] [Locus Tag: ] [Chromosome: 16] [Map Location: 16 \| 16 C | 7.735594377 | 1.26E-15 | 2.32E-14 |
| 6531 | ENSMUSG00000099689 | Zfp383 | 73729 [Gene Symbol: Zfp383] [Locus Tag: ] [Chromosome: 7] [Map Location: 7 \| 7 B1] [D | 7.731493521 | 2.06E-13 | 2.65E-12 |
| 11225 | ENSMUSG00000027808 | Serp1 | 28146 [Gene Symbol: Serp1] [Locus Tag: ] [Chromosome: 3] [Map Location: 3 D \| 3 28.5 | 7.727881025 | 1.28E-23 | 8.61E-22 |
| 9745 | ENSMUSG00000091512 | Lamtor3 | 56692 [Gene Symbol: Lamtor3] [Locus Tag: ] [Chromosome: 3] [Map Location: 3 \| 3 G3] | 7.727180487 | 3.22E-16 | 6.56E-15 |
| 17409 | ENSMUSG00000025245 | Lztfl1 | 93730 [Gene Symbol: Lztfl1] [Locus Tag: ] [Chromosome: 9] [Map Location: 9 74.36 cM | 7.712081562 | 1.60E-22 | 8.96E-21 |
| 1936 | ENSMUSG00000028675 | Pnrc2 | 52830 [Gene Symbol: Pnrc2] [Locus Tag: ] [Chromosome: 4] [Map Location: 4 D3 \| 4 68. | 7.67272937 | 1.47E-13 | 1.95E-12 |
| 15844 | ENSMUSG00000022109 | Med4 | 67381 [Gene Symbol: Med4] [Locus Tag: ] [Chromosome: 14] [Map Location: 14 \| 14 D3 | 7.664639354 | 1.39E-17 | 3.42E-16 |
| 3964 | ENSMUSG00000054702 | Ap1s3 | 252903 [Gene Symbol: Ap1s3] [Locus Tag: ] [Chromosome: 1] [Map Location: 1 \| 1 C4] [D | 7.660144609 | 1.09E-07 | 5.76E-07 |
| 3692 | ENSMUSG00000054405 | Dnajc8 | 68598 [Gene Symbol: Dnajc8] [Locus Tag: ] [Chromosome: 4] [Map Location: 4 \| 4 D2.3] | 7.65152463 | 5.15E-24 | 3.64E-22 |
| 1212 | ENSMUSG00000032423 | Syncrip | 56403 [Gene Symbol: Syncrip] [Locus Tag: ] [Chromosome: 9] [Map Location: 9 \| 9 E3.2] | 7.645817965 | 1.80E-22 | 9.86E-21 |
| 7722 | ENSMUSG00000021079 | Timm9 | 30056 [Gene Symbol: Timm9] [Locus Tag: ] [Chromosome: 12] [Map Location: 12 C3 \| 1 | 7.631424893 | 1.26E-15 | 2.32E-14 |
| 9582 | ENSMUSG00000036916 | Zfp280c | 208968 [Gene Symbol: Zfp280c] [Locus Tag: ] [Chromosome: X] [Map Location: X \| X A4] | 7.620761208 | 8.76E-22 | 4.32E-20 |
| 13392 | ENSMUSG00000026349 | Ccnt2 | 72949 [Gene Symbol: Ccnt2] [Locus Tag: ] [Chromosome: 1] [Map Location: 1 \| 1 E3] [De | 7.604764302 | 1.24E-19 | 4.35E-18 |
| 1665 | ENSMUSG00000060073 | Psma3 | 19167 [Gene Symbol: Psma3] [Locus Tag: ] [Chromosome: 12] [Map Location: 12 \| 12 C3 | 7.601054561 | 2.02E-20 | 7.92E-19 |
| 14913 | ENSMUSG00000014504 | Srp19 | 66384 [Gene Symbol: Srp19] [Locus Tag: ] [Chromosome: 18] [Map Location: 18 \| 18 B3 | 7.587464797 | 4.27E-16 | 8.48E-15 |
| 3285 | ENSMUSG00000059142 | Zfp945 | 240041 [Gene Symbol: Zfp945] [Locus Tag: ] [Chromosome: 17] [Map Location: 17 \| 17 | 7.583919398 | 5.53E-20 | 2.05E-18 |
| 4905 | ENSMUSG00000047115 | Fam221a | 231946 [Gene Symbol: Fam221a] [Locus Tag: ] [Chromosome: 6] [Map Location: 6 \| 6 B2 | 7.576917166 | 2.16E-08 | 1.28E-07 |
| 17935 | ENSMUSG00000020962 | Gtf2a1 | 83602 [Gene Symbol: Gtf2a1] [Locus Tag: ] [Chromosome: 12] [Map Location: 12 \| 12 E] | 7.5684501 | 6.23E-25 | 5.13E-23 |
| 12164 | ENSMUSG00000058729 | Lin9 | 72568 [Gene Symbol: Lin9] [Locus Tag: ] [Chromosome: 1] [Map Location: 1 \| 1 H4] [Des | 7.56617699 | 1.40E-19 | 4.86E-18 |
| 13184 | ENSMUSG00000034391 | Fbxo15 | 50764 [Gene Symbol: Fbxo15] [Locus Tag: ] [Chromosome: 18] [Map Location: 18 \| 18 E | 7.530562342 | 1.23E-15 | 2.27E-14 |
| 4468 | ENSMUSG00000028546 | Elavl4 | 15572 [Gene Symbol: Elavl4] [Locus Tag: ] [Chromosome: 4] [Map Location: 4 C7 \| 4 51. | 7.527986809 | 2.03E-19 | 6.85E-18 |
| 10838 | ENSMUSG00000037234 | Hook3 | 320191 [Gene Symbol: Hook3] [Locus Tag: ] [Chromosome: 8] [Map Location: 8 \| 8 A2] [ | 7.513243105 | 2.74E-25 | 2.34E-23 |
| 3118 | ENSMUSG00000023919 | Cenpq | 83815 [Gene Symbol: Cenpq] [Locus Tag: ] [Chromosome: 17] [Map Location: 17 \| 17 B2 | 7.489040059 | 6.23E-14 | 8.78E-13 |
| 17946 | ENSMUSG00000058267 | Mrps14 | 64659 [Gene Symbol: Mrps14] [Locus Tag: ] [Chromosome: 1] [Map Location: 1 \| 1 H2.1 | 7.472871733 | 1.37E-11 | 1.31E-10 |
| 4118 | ENSMUSG00000021661 | Ankra2 | 68558 [Gene Symbol: Ankra2] [Locus Tag: ] [Chromosome: 13] [Map Location: 13 \| 13 D | 7.468723227 | 8.81E-18 | 2.25E-16 |
| 13936 | ENSMUSG00000033439 | Trmt13 | 229780 [Gene Symbol: Trmt13] [Locus Tag: ] [Chromosome: 3] [Map Location: 3 \| 3 G1] | 7.460308504 | 6.80E-13 | 8.05E-12 |
| 18598 | ENSMUSG00000035958 | Tdp2 | 56196 [Gene Symbol: Tdp2] [Locus Tag: ] [Chromosome: 13] [Map Location: 13 A3.1 \| 1 | 7.456962009 | 9.72E-18 | 2.46E-16 |
| 6310 | ENSMUSG00000057842 | Zfp595 | 218314 [Gene Symbol: Zfp595] [Locus Tag: ] [Chromosome: 13] [Map Location: 13 \| 13 | 7.453058713 | 5.19E-13 | 6.24E-12 |
| 9407 | ENSMUSG00000028249 | Sdcbp | 53378 [Gene Symbol: Sdcbp] [Locus Tag: ] [Chromosome: 4] [Map Location: 4 \| 4 A1] [De | 7.43715819 | 5.43E-23 | 3.33E-21 |
| 1347 | ENSMUSG00000024976 | Shoc2 | 56392 [Gene Symbol: Shoc2] [Locus Tag: ] [Chromosome: 19] [Map Location: 19 \| 19 D2 | 7.436765761 | 1.93E-20 | 7.62E-19 |
| 5062 | ENSMUSG00000060090 | Rp2 | 19889 [Gene Symbol: Rp2] [Locus Tag: ] [Chromosome: X] [Map Location: X \| X A2] [Desc | 7.428796281 | 1.90E-20 | 7.52E-19 |
| 16456 | ENSMUSG00000038633 | Degs1 | 13244 [Gene Symbol: Degs1] [Locus Tag: ] [Chromosome: 1] [Map Location: 1 \| 1 H5] [De | 7.425609007 | 8.77E-18 | 2.24E-16 |
| 6803 | ENSMUSG00000033813 | Tcea1 | 21399 [Gene Symbol: Tcea1] [Locus Tag: ] [Chromosome: 1] [Map Location: 1 \| 1 A1] [De | 7.425003295 | 2.22E-19 | 7.41E-18 |
| 652 | ENSMUSG00000051185 | Fam174a | 67698 [Gene Symbol: Fam174a] [Locus Tag: ] [Chromosome: 1] [Map Location: 1 \| 1 D] [ | 7.421453238 | 6.55E-17 | 1.49E-15 |
| 4314 | ENSMUSG00000029208 | Guf1 | 231279 [Gene Symbol: Guf1] [Locus Tag: ] [Chromosome: 5] [Map Location: 5 \| 5 C3.1] [ | 7.417033134 | 3.59E-15 | 6.05E-14 |
| 7103 | ENSMUSG00000018379 | Srsf1 | 110809 [Gene Symbol: Srsf1] [Locus Tag: ] [Chromosome: 11] [Map Location: 11 C \| 11 5 | 7.41147231 | 1.28E-24 | 9.98E-23 |
| 15153 | ENSMUSG00000047238 | Mageh1 | 75625 [Gene Symbol: Mageh1] [Locus Tag: ] [Chromosome: X] [Map Location: X \| X F2] [D | 7.395526706 | 1.61E-13 | 2.11E-12 |
| 1653 | ENSMUSG00000028693 | Nasp | 50927 [Gene Symbol: Nasp] [Locus Tag: ] [Chromosome: 4] [Map Location: 4 D1 \| 4 53.2 | 7.393953612 | 2.25E-17 | 5.42E-16 |
| 15368 | ENSMUSG00000027245 | Hypk | 67693 [Gene Symbol: Hypk] [Locus Tag: ] [Chromosome: 2] [Map Location: 2 \| 2 F1] [Des | 7.390351159 | 7.91E-15 | 1.28E-13 |
| 6729 | ENSMUSG00000059811 | Atl2 | 56298 [Gene Symbol: Atl2] [Locus Tag: ] [Chromosome: 17] [Map Location: 17 \| 17 E3] [ | 7.36683037 | 1.74E-20 | 6.97E-19 |
| 7453 | ENSMUSG00000037808 | Fam76b | 72826 [Gene Symbol: Fam76b] [Locus Tag: ] [Chromosome: 9] [Map Location: 9 \| 9 A1] [ | 7.362662821 | 8.88E-22 | 4.36E-20 |
| 10276 | ENSMUSG00000026878 | Rab14 | 68365 [Gene Symbol: Rab14] [Locus Tag: ] [Chromosome: 2] [Map Location: 2 \| 2 B] [Des | 7.342859649 | 3.01E-20 | 1.16E-18 |
| 2099 | ENSMUSG00000031521 | Aga | 11593 [Gene Symbol: Aga] [Locus Tag: ] [Chromosome: 8] [Map Location: 8 \| 8 B3] [Descr | 7.34127427 | 2.01E-08 | 1.20E-07 |
| 6880 | ENSMUSG00000019998 | Stx7 | 53331 [Gene Symbol: Stx7] [Locus Tag: ] [Chromosome: 10] [Map Location: 10 \| 10 A3] [ | 7.335159681 | 9.35E-25 | 7.46E-23 |
| 1729 | ENSMUSG00000007656 | Arpp19 | 59046 [Gene Symbol: Arpp19] [Locus Tag: ] [Chromosome: 9] [Map Location: 9 \| 9 E1] [D | 7.330666065 | 8.74E-19 | 2.62E-17 |
| 7914 | ENSMUSG00000026696 | Vamp4 | 53330 [Gene Symbol: Vamp4] [Locus Tag: ] [Chromosome: 1] [Map Location: 1 70.29 c | 7.325007057 | 1.53E-22 | 8.64E-21 |
| 12644 | ENSMUSG00000022661 | Cd200 | 17470 [Gene Symbol: Cd200] [Locus Tag: ] [Chromosome: 16] [Map Location: 16 29.53 | 7.321951784 | 2.73E-24 | 2.02E-22 |
| 12626 | ENSMUSG00000040681 | Hmgn1; LOC1026418 | 15312 [Gene Symbol: Hmgn1] [Locus Tag: ] [Chromosome: 16] [Map Location: 16 C4 \| 1 | 7.319735654 | 4.92E-14 | 7.04E-13 |
| 18103 | ENSMUSG00000028419 | Chmp5 | 76959 [Gene Symbol: Chmp5] [Locus Tag: ] [Chromosome: 4] [Map Location: 4 \| 4 A5] [D | 7.311121223 | 1.76E-20 | 7.02E-19 |
| 480 | ENSMUSG00000022111 | Uchl3 | 50933 [Gene Symbol: Uchl3] [Locus Tag: ] [Chromosome: 14] [Map Location: 14 E2.3 \| 1 | 7.309076018 | 6.79E-14 | 9.50E-13 |
| 17605 | ENSMUSG00000019775 | Rgs17 | 56533 [Gene Symbol: Rgs17] [Locus Tag: ] [Chromosome: 10] [Map Location: 10 \| 10 A1 | 7.307258882 | 4.76E-19 | 1.50E-17 |
| 600 | ENSMUSG00000035151 | Elmod2 | 244548 [Gene Symbol: Elmod2] [Locus Tag: ] [Chromosome: 8] [Map Location: 8 \| 8 C2] | 7.300867491 | 5.45E-16 | 1.06E-14 |
| 6974 | ENSMUSG00000027285 | Haus2 | 66296 [Gene Symbol: Haus2] [Locus Tag: ] [Chromosome: 2] [Map Location: 2 \| 2 F1] [De | 7.294165832 | 2.89E-18 | 8.02E-17 |
| 3206 | ENSMUSG00000031201 | Brcc3 | 210766 [Gene Symbol: Brcc3] [Locus Tag: ] [Chromosome: X] [Map Location: X \| X A7.3] | 7.291455575 | 5.00E-19 | 1.56E-17 |
| 8550 | ENSMUSG00000021514 | Zfp369 | 170936 [Gene Symbol: Zfp369] [Locus Tag: ] [Chromosome: 13] [Map Location: 13 \| 13 | 7.282628166 | 4.08E-18 | 1.10E-16 |
| 1863 | ENSMUSG00000061273 | Mmgt1 | 236792 [Gene Symbol: Mmgt1] [Locus Tag: ] [Chromosome: X] [Map Location: X \| X A5] [ | 7.279963858 | 9.62E-21 | 4.02E-19 |
| 16837 | ENSMUSG00000025199 | Chuk | 12675 [Gene Symbol: Chuk] [Locus Tag: ] [Chromosome: 19] [Map Location: 19 C3 \| 19 | 7.275172142 | 3.83E-22 | 1.98E-20 |
| 14679 | ENSMUSG00000038127 | Ccdc50 | 67501 [Gene Symbol: Ccdc50] [Locus Tag: ] [Chromosome: 16] [Map Location: 16 B2 \| 1 | 7.26378005 | 2.10E-22 | 1.14E-20 |
| 7205 | ENSMUSG00000026017 | Carf | 241066 [Gene Symbol: Carf] [Locus Tag: ] [Chromosome: 1] [Map Location: 1 \| 1 C2] [Des | 7.252957967 | 9.90E-16 | 1.86E-14 |
| 16380 | ENSMUSG00000005897 | Nr2c1 | 22025 [Gene Symbol: Nr2c1] [Locus Tag: ] [Chromosome: 10] [Map Location: 10 C2 \| 10 | 7.249570457 | 9.22E-16 | 1.74E-14 |
| 3008 | ENSMUSG00000028212 | Ccne2 | 12448 [Gene Symbol: Ccne2] [Locus Tag: ] [Chromosome: 4] [Map Location: 4 \| 4 A2] [De | 7.248152959 | 2.54E-18 | 7.11E-17 |
| 6173 | ENSMUSG00000036943 | Rab8b | 235442 [Gene Symbol: Rab8b] [Locus Tag: ] [Chromosome: 9] [Map Location: 9 \| 9 C] [De | 7.235876513 | 2.03E-20 | 7.94E-19 |
| 7080 | ENSMUSG00000028945 | Rheb | 19744 [Gene Symbol: Rheb] [Locus Tag: ] [Chromosome: 5] [Map Location: 5 \| 5 A3] [Des | 7.23512631 | 1.62E-16 | 3.48E-15 |
| 3106 | ENSMUSG00000059554 | Ccdc28a | 215814 [Gene Symbol: Ccdc28a] [Locus Tag: ] [Chromosome: 10] [Map Location: 10 \| 1 | 7.23173892 | 2.17E-09 | 1.49E-08 |
| 6377 | ENSMUSG00000029655 | N4bp2l2 | 381695 [Gene Symbol: N4bp2l2] [Locus Tag: ] [Chromosome: 5] [Map Location: 5 \| 5 G3 | 7.213472415 | 1.60E-23 | 1.05E-21 |
| 9641 | ENSMUSG00000028180 | Zranb2 | 53861 [Gene Symbol: Zranb2] [Locus Tag: ] [Chromosome: 3] [Map Location: 3 \| 3 H4] [D | 7.206222189 | 1.34E-19 | 4.66E-18 |
| 14938 | ENSMUSG00000032621 | Srek1 | 218543 [Gene Symbol: Srek1] [Locus Tag: ] [Chromosome: 13] [Map Location: 13 \| 13 D | 7.202144931 | 1.18E-16 | 2.58E-15 |
| 18019 | ENSMUSG00000027708 | Dcun1d1 | 114893 [Gene Symbol: Dcun1d1] [Locus Tag: ] [Chromosome: 3] [Map Location: 3 \| 3 B] | 7.189866088 | 5.65E-22 | 2.86E-20 |
| 4999 | ENSMUSG00000025001 | Hells | 15201 [Gene Symbol: Hells] [Locus Tag: ] [Chromosome: 19] [Map Location: 19 \| 19 C3- | 7.187082364 | 3.71E-12 | 3.88E-11 |
| 17733 | ENSMUSG00000028165 | Cisd2 | 67006 [Gene Symbol: Cisd2] [Locus Tag: ] [Chromosome: 3] [Map Location: 3 \| 3 G3] [De | 7.180681351 | 2.59E-21 | 1.19E-19 |
| 13814 | ENSMUSG00000030647 | Ndufc2; LOC1026413 | 68197 [Gene Symbol: Ndufc2] [Locus Tag: ] [Chromosome: 7] [Map Location: 7 \| 7 E3] [D | 7.167073153 | 2.91E-15 | 4.99E-14 |
| 12232 | ENSMUSG00000008489 | Elavl2 | 15569 [Gene Symbol: Elavl2] [Locus Tag: ] [Chromosome: 4] [Map Location: 4 C5 \| 4 42. | 7.155370706 | 6.54E-23 | 3.93E-21 |
| 52 | ENSMUSG00000043013 | Onecut1 | 15379 [Gene Symbol: Onecut1] [Locus Tag: ] [Chromosome: 9] [Map Location: 9 D \| 9 41 | 7.151435882 | 3.81E-12 | 3.98E-11 |
| 11723 | ENSMUSG00000022911 | Arl13b | 68146 [Gene Symbol: Arl13b] [Locus Tag: ] [Chromosome: 16] [Map Location: 16 \| 16 C1 | 7.133015852 | 2.12E-20 | 8.27E-19 |
| 16371 | ENSMUSG00000022820 | Ndufb4; Gm3873; Gn | 68194 [Gene Symbol: Ndufb4] [Locus Tag: ] [Chromosome: 16] [Map Location: 16 \| 16 B | 7.124138376 | 1.04E-07 | 5.54E-07 |
| 18392 | ENSMUSG00000025537 | Phkg1 | 18682 [Gene Symbol: Phkg1] [Locus Tag: ] [Chromosome: 5] [Map Location: 5 G1.3 \| 5 6 | 7.119940114 | 3.74E-18 | 1.02E-16 |
| 609 | ENSMUSG00000050050 | Ccdc158 | 320696 [Gene Symbol: Ccdc158] [Locus Tag: ] [Chromosome: 5] [Map Location: 5 \| 5 E2 | 7.116541937 | 3.33E-07 | 1.62E-06 |
| 13245 | ENSMUSG00000029413 | Naaa | 67111 [Gene Symbol: Naaa] [Locus Tag: ] [Chromosome: 5] [Map Location: 5 \| 5 E3] [Des | 7.113689782 | 7.75E-14 | 1.08E-12 |
| 7839 | ENSMUSG00000019966 | Kitl | 17311 [Gene Symbol: Kitl] [Locus Tag: ] [Chromosome: 10] [Map Location: 10 D1 \| 10 51 | 7.112816238 | 1.68E-20 | 6.73E-19 |
| 14695 | ENSMUSG00000071551 | Akr1c19 | 432720 [Gene Symbol: Akr1c19] [Locus Tag: ] [Chromosome: 13] [Map Location: 13 \| 13 | 7.112695771 | 7.18E-10 | 5.29E-09 |
| 10226 | ENSMUSG00000019948 | Actr6 | 67019 [Gene Symbol: Actr6] [Locus Tag: ] [Chromosome: 10] [Map Location: 10 \| 10 C2] | 7.111713476 | 3.00E-16 | 6.16E-15 |

| | | | | | | |
|---|---|---|---|---|---|---|
| 3805 | ENSMUSG00000060935 | Tmem263 | 103266 [Gene Symbol: Tmem263] [Locus Tag: ] [Chromosome: 10] [Map Location: 10\|1 | 7.101954839 | 1.82E-22 | 9.98E-21 |
| 934 | ENSMUSG00000029267 | Mtf2 | 17765 [Gene Symbol: Mtf2] [Locus Tag: ] [Chromosome: 5] [Map Location: 5\|5 F] [Descr | 7.09419758 | 1.05E-17 | 2.63E-16 |
| 18021 | ENSMUSG00000031130 | Brs3 | 12209 [Gene Symbol: Brs3] [Locus Tag: ] [Chromosome: X] [Map Location: X\|X A7.1-A7. | 7.093122337 | 0.023985 | 0.045336 |
| 10949 | ENSMUSG00000074264 | Amy1 | 11722 [Gene Symbol: Amy1] [Locus Tag: ] [Chromosome: 3] [Map Location: 3 F3\|3 49.3 | 7.079987483 | 1.46E-08 | 8.87E-08 |
| 2847 | ENSMUSG00000044164 | Rnf182 | 328234 [Gene Symbol: Rnf182] [Locus Tag: ] [Chromosome: 13] [Map Location: 13\|13 / | 7.059908777 | 7.46E-07 | 3.44E-06 |
| 583 | ENSMUSG00000030543 | Mesp2 | 17293 [Gene Symbol: Mesp2] [Locus Tag: ] [Chromosome: 7] [Map Location: 7 D3\|7 45. | 7.05901032 | 1.75E-07 | 8.94E-07 |
| 1120 | ENSMUSG00000037475 | Thoc2 | 331401 [Gene Symbol: Thoc2] [Locus Tag: ] [Chromosome: X] [Map Location: X\|X A4] [D | 7.050650233 | 2.34E-21 | 1.08E-19 |
| 11553 | ENSMUSG00000022837 | Iqcb1 | 320299 [Gene Symbol: Iqcb1] [Locus Tag: ] [Chromosome: 16] [Map Location: 16\|16 B3 | 7.039363625 | 2.10E-14 | 3.19E-13 |
| 17280 | ENSMUSG00000029569 | Tmem168 | 101118 [Gene Symbol: Tmem168] [Locus Tag: ] [Chromosome: 6] [Map Location: 6\|6 A | 7.036612611 | 2.45E-16 | 5.07E-15 |
| 2815 | ENSMUSG00000038646 | Fam103a1 | 67148 [Gene Symbol: Fam103a1] [Locus Tag: ] [Chromosome: 7] [Map Location: 7\|7 D3 | 7.036163764 | 1.72E-13 | 2.24E-12 |
| 16788 | ENSMUSG00000029415 | Sdad1 | 231452 [Gene Symbol: Sdad1] [Locus Tag: ] [Chromosome: 5] [Map Location: 5\|5 E2] [D | 7.029176867 | 2.38E-21 | 1.10E-19 |
| 13885 | ENSMUSG00000005233 | Spc25 | 66442 [Gene Symbol: Spc25] [Locus Tag: ] [Chromosome: 2] [Map Location: 2\|2 C3] [De | 7.01149508 | 2.02E-08 | 1.20E-07 |
| 574 | ENSMUSG00000060771 | Tsga10 | 211484 [Gene Symbol: Tsga10] [Locus Tag: ] [Chromosome: 1] [Map Location: 1\|1 B] [D | 7.004431407 | 2.10E-14 | 3.20E-13 |
| 3971 | ENSMUSG00000015247 | Nipsnap3b | 66536 [Gene Symbol: Nipsnap3b] [Locus Tag: ] [Chromosome: 4] [Map Location: 4\|4 B3 | 7.00351834 | 6.49E-19 | 1.98E-17 |
| 5499 | ENSMUSG00000019927 | Ube2d1 | 216080 [Gene Symbol: Ube2d1] [Locus Tag: ] [Chromosome: 10] [Map Location: 10\|10 | 7.002919221 | 6.78E-20 | 2.48E-18 |
| 18450 | ENSMUSG00000091625 | Lsm5; LOC10263625 | 66373 [Gene Symbol: Lsm5] [Locus Tag: ] [Chromosome: 6] [Map Location: 6\|6 C1] [Des | 7.002497669 | 7.98E-06 | 3.08E-05 |
| 18272 | ENSMUSG00000035236 | Scai | 320271 [Gene Symbol: Scai] [Locus Tag: ] [Chromosome: 2] [Map Location: 2\|2 B] [Desc | 6.998597748 | 1.17E-23 | 8.00E-22 |
| 18384 | ENSMUSG00000030393 | Zik1 | 22775 [Gene Symbol: Zik1] [Locus Tag: ] [Chromosome: 7] [Map Location: 7\|7 A1] [Desc | 6.992075875 | 8.81E-14 | 1.21E-12 |
| 18142 | ENSMUSG00000040774 | Cept1 | 99712 [Gene Symbol: Cept1] [Locus Tag: ] [Chromosome: 3] [Map Location: 3\|3 F2.3] [D | 6.989903534 | 5.38E-19 | 1.67E-17 |
| 10998 | ENSMUSG00000019876 | Pkib | 18768 [Gene Symbol: Pkib] [Locus Tag: ] [Chromosome: 10] [Map Location: 10 B4\|10 2 | 6.985102257 | 1.46E-09 | 1.03E-08 |
| 13842 | ENSMUSG00000019792 | Trmt11 | 73681 [Gene Symbol: Trmt11] [Locus Tag: ] [Chromosome: 10] [Map Location: 10\|10 A | 6.981211238 | 3.01E-13 | 3.74E-12 |
| 7317 | ENSMUSG00000019689 | 1110001J03Rik | 66117 [Gene Symbol: 1110001J03Rik] [Locus Tag: ] [Chromosome: 6] [Map Location: 6 | 6.980743409 | 2.56E-07 | 1.27E-06 |
| 7153 | ENSMUSG00000074826 | Gm10767 | 100038538 [Gene Symbol: Gm10767] [Locus Tag: ] [Chromosome: 13] [Map Location: | 6.970780212 | 5.64E-06 | 2.23E-05 |
| 4275 | ENSMUSG00000034120 | Srsf2 | 20382 [Gene Symbol: Srsf2] [Locus Tag: ] [Chromosome: 11] [Map Location: 11 E2\|11 8 | 6.9635243 | 1.14E-22 | 6.49E-21 |
| 12285 | ENSMUSG00000032113 | Chek1 | 12649 [Gene Symbol: Chek1] [Locus Tag: ] [Chromosome: 9] [Map Location: 9\|9 A5.3] [ | 6.953640698 | 8.60E-11 | 7.28E-10 |
| 12897 | ENSMUSG00000045827 | Serpinb9 | 20723 [Gene Symbol: Serpinb9] [Locus Tag: ] [Chromosome: 13] [Map Location: 13 A3. | 6.94814698 | 0.006665 | 0.014422 |
| 9978 | ENSMUSG00000075470 | Alg10b | 380959 [Gene Symbol: Alg10b] [Locus Tag: ] [Chromosome: 15] [Map Location: 15\|15 | 6.945671759 | 5.77E-23 | 3.50E-21 |
| 5067 | ENSMUSG00000056260 | Lrif1 | 321000 [Gene Symbol: Lrif1] [Locus Tag: ] [Chromosome: 3] [Map Location: 3\|3 F2.3] [D | 6.940780761 | 3.93E-14 | 5.73E-13 |
| 9894 | ENSMUSG00000045140 | Pigw | 70325 [Gene Symbol: Pigw] [Locus Tag: ] [Chromosome: 11] [Map Location: 11\|11 B5] | 6.937464988 | 3.21E-15 | 5.47E-14 |
| 14341 | ENSMUSG00000028003 | Lrat | 79235 [Gene Symbol: Lrat] [Locus Tag: ] [Chromosome: 3] [Map Location: 3\|3 E3] [Desc | 6.934839759 | 0.008366 | 0.017729 |
| 4333 | ENSMUSG00000047749 | Zc3hav1l | 209032 [Gene Symbol: Zc3hav1l] [Locus Tag: ] [Chromosome: 6] [Map Location: 6\|6 B1 | 6.934706746 | 6.45E-14 | 9.06E-13 |
| 1714 | ENSMUSG00000029911 | Ssbp1 | 381760 [Gene Symbol: Ssbp1] [Locus Tag: ] [Chromosome: 6] [Map Location: 6\|6 B2] [D | 6.930806313 | 3.32E-17 | 7.85E-16 |
| 4169 | ENSMUSG00000067780 | Pi15 | 94227 [Gene Symbol: Pi15] [Locus Tag: ] [Chromosome: 1] [Map Location: 1\|1 A4] [Des | 6.929714516 | 6.88E-06 | 2.69E-05 |
| 2560 | ENSMUSG00000022309 | Angpt1 | 11600 [Gene Symbol: Angpt1] [Locus Tag: ] [Chromosome: 15] [Map Location: 15 B3.1\| | 6.928246276 | 1.09E-20 | 4.54E-19 |
| 16058 | ENSMUSG00000033111 | 3830406C13Rik | 218734 [Gene Symbol: 3830406C13Rik] [Locus Tag: ] [Chromosome: 14] [Map Locatior | 6.915636116 | 3.22E-14 | 4.76E-13 |
| 7228 | ENSMUSG00000026078 | Pdcl3 | 68833 [Gene Symbol: Pdcl3] [Locus Tag: ] [Chromosome: 1] [Map Location: 1\|1 B] [Desc | 6.914154914 | 3.69E-19 | 1.19E-17 |
| 18051 | ENSMUSG00000047215 | Rpl9 | 20005 [Gene Symbol: Rpl9] [Locus Tag: ] [Chromosome: 5] [Map Location: 5\|5 C3.1] [D | 6.907557059 | 1.13E-06 | 5.05E-06 |
| 5948 | ENSMUSG00000028403 | Zdhhc21 | 68268 [Gene Symbol: Zdhhc21] [Locus Tag: ] [Chromosome: 4] [Map Location: 4 39.4 cl | 6.904008983 | 3.59E-23 | 2.25E-21 |
| 14575 | ENSMUSG00000022744 | Cldnd1 | 224250 [Gene Symbol: Cldnd1] [Locus Tag: ] [Chromosome: 16] [Map Location: 16\|16 | 6.884352371 | 4.20E-20 | 1.59E-18 |
| 5678 | ENSMUSG00000038683 | Pak1ip1 | 68083 [Gene Symbol: Pak1ip1] [Locus Tag: ] [Chromosome: 13] [Map Location: 13\|13 / | 6.86184992 | 4.32E-17 | 1.00E-15 |
| 14801 | ENSMUSG00000020225 | Tmbim4 | 68212 [Gene Symbol: Tmbim4] [Locus Tag: ] [Chromosome: 10] [Map Location: 10\|10 | 6.855055263 | 3.97E-12 | 4.13E-11 |
| 6465 | ENSMUSG00000027076 | Timm10 | 30059 [Gene Symbol: Timm10] [Locus Tag: ] [Chromosome: 2] [Map Location: 2\|2 E1] [ | 6.843237903 | 5.60E-18 | 1.47E-16 |
| 14458 | ENSMUSG00000035234 | Fam175a | 70681 [Gene Symbol: Fam175a] [Locus Tag: ] [Chromosome: 5] [Map Location: 5\|5 E4] | 6.833923584 | 4.97E-10 | 3.74E-09 |
| 12215 | ENSMUSG00000052676 | Zmat1 | 215693 [Gene Symbol: Zmat1] [Locus Tag: ] [Chromosome: X] [Map Location: X\|X E3] [D | 6.833381318 | 1.85E-13 | 2.39E-12 |
| 13287 | ENSMUSG00000029209 | Gnpda2 | 67980 [Gene Symbol: Gnpda2] [Locus Tag: ] [Chromosome: 5] [Map Location: 5\|5 D] [D | 6.824606926 | 2.67E-15 | 4.62E-14 |
| 8658 | ENSMUSG00000074513 | Arfip1 | 99889 [Gene Symbol: Arfip1] [Locus Tag: ] [Chromosome: 3] [Map Location: 3\|3 F1] [De | 6.816168043 | 1.62E-19 | 5.52E-18 |
| 7496 | ENSMUSG00000040651 | Fam208a | 218850 [Gene Symbol: Fam208a] [Locus Tag: ] [Chromosome: 14] [Map Location: 14\|1 | 6.814260948 | 2.26E-18 | 6.38E-17 |
| 3913 | ENSMUSG00000046699 | Slitrk4 | 245446 [Gene Symbol: Slitrk4] [Locus Tag: ] [Chromosome: X] [Map Location: X\|X A5] [ | 6.802723012 | 2.16E-15 | 3.78E-14 |
| 1086 | ENSMUSG00000021013 | Ttc8 | 76260 [Gene Symbol: Ttc8] [Locus Tag: ] [Chromosome: 12] [Map Location: 12\|12 E] [D | 6.799361622 | 1.50E-15 | 2.71E-14 |
| 4939 | ENSMUSG00000020630 | Rnaseh1 | 19819 [Gene Symbol: Rnaseh1] [Locus Tag: ] [Chromosome: 12] [Map Location: 12\|12 / | 6.797941246 | 9.62E-13 | 1.11E-11 |
| 4345 | ENSMUSG00000078622 | Ccdc47 | 67163 [Gene Symbol: Ccdc47] [Locus Tag: ] [Chromosome: 11] [Map Location: 11\|11 E | 6.795557503 | 2.30E-22 | 1.25E-20 |
| 7300 | ENSMUSG00000090919 | Pabpc4l | 241989 [Gene Symbol: Pabpc4l] [Locus Tag: ] [Chromosome: 3] [Map Location: 3\|3 B] [ | 6.795200764 | 1.07E-09 | 7.65E-09 |
| 3495 | ENSMUSG00000031438 | Rnf128 | 66889 [Gene Symbol: Rnf128] [Locus Tag: ] [Chromosome: X] [Map Location: X\|X F1] [D | 6.791364865 | 9.54E-09 | 5.99E-08 |
| 2232 | ENSMUSG00000024592 | C330018D20Rik | 77422 [Gene Symbol: C330018D20Rik] [Locus Tag: ] [Chromosome: 18] [Map Location: | 6.78972535 | 1.47E-10 | 1.20E-09 |
| 17817 | ENSMUSG00000090935 | Synj2bp | 24071 [Gene Symbol: Synj2bp] [Locus Tag: ] [Chromosome: 12] [Map Location: 12 D2\| | 6.781217167 | 5.70E-21 | 2.50E-19 |
| 2923 | ENSMUSG00000021840 | Mapk1ip1l | 218975 [Gene Symbol: Mapk1ip1l] [Locus Tag: ] [Chromosome: 14] [Map Location: 14\| | 6.772346319 | 1.96E-21 | 9.18E-20 |
| 15119 | ENSMUSG00000042670 | Immp1l | 66541 [Gene Symbol: Immp1l] [Locus Tag: ] [Chromosome: 2] [Map Location: 2\|2 E3] [ | 6.770154616 | 7.80E-19 | 2.35E-17 |
| 13743 | ENSMUSG00000025066 | Sfr1 | 67788 [Gene Symbol: Sfr1] [Locus Tag: ] [Chromosome: 19] [Map Location: 19\|19 D2] [ | 6.7658169 | 1.50E-14 | 2.32E-13 |
| 18010 | ENSMUSG00000028060 | 2810403A07Rik | 74200 [Gene Symbol: 2810403A07Rik] [Locus Tag: ] [Chromosome: 3] [Map Location: 3 | 6.751997423 | 2.22E-17 | 5.36E-16 |
| 9420 | ENSMUSG00000027835 | Pdcd10 | 56426 [Gene Symbol: Pdcd10] [Locus Tag: ] [Chromosome: 3] [Map Location: 3\|3 E3] [D | 6.742810599 | 2.06E-16 | 4.34E-15 |
| 13247 | ENSMUSG00000039704 | Lmbrd2 | 320506 [Gene Symbol: Lmbrd2] [Locus Tag: ] [Chromosome: 15] [Map Location: 15\|15 | 6.739083021 | 1.37E-21 | 6.60E-20 |
| 5134 | ENSMUSG00000022838 | Eaf2 | 106389 [Gene Symbol: Eaf2] [Locus Tag: ] [Chromosome: 16] [Map Location: 16\|16 B3] | 6.735279781 | 5.43E-14 | 7.73E-13 |
| 5568 | ENSMUSG00000024759 | Atl3 | 109168 [Gene Symbol: Atl3] [Locus Tag: ] [Chromosome: 19] [Map Location: 19\|19 A] [ | 6.725540106 | 9.85E-21 | 4.11E-19 |
| 10476 | ENSMUSG00000031878 | Nae1 | 234664 [Gene Symbol: Nae1] [Locus Tag: ] [Chromosome: 8] [Map Location: 8\|8 D3] [D | 6.7219412 | 2.75E-18 | 7.69E-17 |
| 4279 | ENSMUSG00000013662 | Atad1 | 67979 [Gene Symbol: Atad1] [Locus Tag: ] [Chromosome: 19] [Map Location: 19\|19 C3 | 6.718649079 | 2.64E-21 | 1.21E-19 |
| 11198 | ENSMUSG00000057143 | Trim12c | 319236 [Gene Symbol: Trim12c] [Locus Tag: ] [Chromosome: 7] [Map Location: 7\|7 E3] | 6.717641154 | 7.37E-07 | 3.40E-06 |
| 18091 | ENSMUSG00000044814 | Olfr543 | 257947 [Gene Symbol: Olfr543] [Locus Tag: ] [Chromosome: 7] [Map Location: 7\|7 E3] | 6.717299092 | 0.008314 | 0.01763 |
| 11250 | ENSMUSG00000029918 | Gm12540; Mrps33 | 100041328 [Gene Symbol: Gm12540] [Locus Tag: ] [Chromosome: 3] [Map Location: 3 | 6.716677923 | 1.20E-07 | 6.33E-07 |
| 825 | ENSMUSG00000071266 | Zfp946 | 74149 [Gene Symbol: Zfp946] [Locus Tag: ] [Chromosome: 17] [Map Location: 17\|17 A | 6.701177013 | 4.72E-17 | 1.09E-15 |
| 15532 | ENSMUSG00000028133 | Rwdd3 | 66568 [Gene Symbol: Rwdd3] [Locus Tag: ] [Chromosome: 3] [Map Location: 3\|3 G1] [D | 6.691715749 | 1.67E-10 | 1.35E-09 |
| 14464 | ENSMUSG00000037725 | Ckap2 | 80986 [Gene Symbol: Ckap2] [Locus Tag: ] [Chromosome: 8] [Map Location: 8\|8 A2] [De | 6.69070039 | 5.00E-07 | 2.37E-06 |
| 13031 | ENSMUSG00000010803 | Gabra1 | 14394 [Gene Symbol: Gabra1] [Locus Tag: ] [Chromosome: 11] [Map Location: 11 A5\|1 | 6.688518979 | 1.30E-08 | 7.99E-08 |
| 11262 | ENSMUSG00000020328 | Nudcd2 | 52653 [Gene Symbol: Nudcd2] [Locus Tag: ] [Chromosome: 11] [Map Location: 11 A5\|1 | 6.68283932 | 2.72E-15 | 4.70E-14 |
| 5127 | ENSMUSG00000054690 | Emcn | 59308 [Gene Symbol: Emcn] [Locus Tag: ] [Chromosome: 3] [Map Location: 3\|3 G3] [De | 6.679783908 | 3.01E-10 | 2.34E-09 |
| 17124 | ENSMUSG00000019773 | Fbxo5 | 67141 [Gene Symbol: Fbxo5] [Locus Tag: ] [Chromosome: 10] [Map Location: 10\|10 A1 | 6.673651081 | 3.40E-09 | 2.28E-08 |
| 5226 | ENSMUSG00000035351 | Nup37 | 69736 [Gene Symbol: Nup37] [Locus Tag: ] [Chromosome: 10] [Map Location: 10\|10 C2 | 6.666779815 | 1.22E-11 | 1.18E-10 |
| 14531 | ENSMUSG00000026064 | LOC102643131; Ptp4 | 102643131 [Gene Symbol: LOC102643131] [Locus Tag: ] [Chromosome: 6] [Map Locati | 6.663582018 | 9.94E-15 | 1.58E-13 |
| 11773 | ENSMUSG00000026150 | Mff | 75734 [Gene Symbol: Mff] [Locus Tag: ] [Chromosome: 1] [Map Location: 1\|1 C5] [Descr | 6.663446519 | 7.61E-23 | 4.50E-21 |
| 417 | ENSMUSG00000057388 | Mrpl18 | 67681 [Gene Symbol: Mrpl18] [Locus Tag: ] [Chromosome: 17] [Map Location: 17\|17 A | 6.661493952 | 7.10E-16 | 1.36E-14 |
| 9850 | ENSMUSG00000030304 | Ergic2 | 67456 [Gene Symbol: Ergic2] [Locus Tag: ] [Chromosome: 6] [Map Location: 6\|6 G3] [De | 6.660782271 | 7.39E-18 | 1.91E-16 |
| 3867 | ENSMUSG00000059897 | Zfp930 | 234358 [Gene Symbol: Zfp930] [Locus Tag: ] [Chromosome: 8] [Map Location: 8\|8 B3.3 | 6.660292673 | 7.43E-07 | 3.43E-06 |
| 10832 | ENSMUSG00000043004 | Gng2 | 14702 [Gene Symbol: Gng2] [Locus Tag: ] [Chromosome: 14] [Map Location: 14 A3\|14 | 6.652444203 | 1.23E-23 | 8.35E-22 |
| 13949 | ENSMUSG00000007646 | Rad51c | 114714 [Gene Symbol: Rad51c] [Locus Tag: ] [Chromosome: 11] [Map Location: 11 C\|1 | 6.650026182 | 1.91E-11 | 1.79E-10 |
| 10797 | ENSMUSG00000038212 | Hiatl1 | 66631 [Gene Symbol: Hiatl1] [Locus Tag: ] [Chromosome: 13] [Map Location: 13\|13 B3 | 6.628720501 | 3.42E-20 | 1.30E-18 |

| | | | | | | |
|---|---|---|---|---|---|---|
| 15044 | ENSMUSG00000047632 | Fgfbp3 | 72514 [Gene Symbol: Fgfbp3] [Locus Tag: ] [Chromosome: 19] [Map Location: 19\|19 C2 | 6.62784796 | 7.15E-14 | 9.97E-13 |
| 12466 | ENSMUSG00000053604 | Rpia | 19895 [Gene Symbol: Rpia] [Locus Tag: ] [Chromosome: 6] [Map Location: 6 C1\|6 32.06 | 6.625918161 | 4.67E-19 | 1.48E-17 |
| 5429 | ENSMUSG00000071528 | Gm15151; Usmg5; LC | 102637290 [Gene Symbol: Gm15151] [Locus Tag: ] [Chromosome: X] [Map Location: X | 6.622585031 | 1.33E-11 | 1.28E-10 |
| 1331 | ENSMUSG00000071716 | Apol7e | 666348 [Gene Symbol: Apol7e] [Locus Tag: ] [Chromosome: 15] [Map Location: 15\|15 E | 6.61424435 | 0.000181 | 0.000542 |
| 6868 | ENSMUSG00000052962 | Mrpl35 | 66223 [Gene Symbol: Mrpl35] [Locus Tag: ] [Chromosome: 6] [Map Location: 6\|6 C3] [D | 6.610117337 | 1.11E-16 | 2.44E-15 |
| 5086 | ENSMUSG00000046404 | Yod1 | 226418 [Gene Symbol: Yod1] [Locus Tag: ] [Chromosome: 1] [Map Location: 1\|1 E4] [De | 6.605820558 | 1.92E-21 | 8.99E-20 |
| 17412 | ENSMUSG00000078773 | Rad54b | 623474 [Gene Symbol: Rad54b] [Locus Tag: ] [Chromosome: 4] [Map Location: 4\|4 A1] | 6.604759566 | 5.18E-13 | 6.23E-12 |
| 1365 | ENSMUSG00000026787 | Gad2 | 14417 [Gene Symbol: Gad2] [Locus Tag: ] [Chromosome: 2] [Map Location: 2 A3\|2 15.1 | 6.597093456 | 2.13E-15 | 3.74E-14 |
| 9610 | ENSMUSG00000069237 | Fam8a1 | 97863 [Gene Symbol: Fam8a1] [Locus Tag: ] [Chromosome: 13] [Map Location: 13\|13 A | 6.570581336 | 6.37E-22 | 3.22E-20 |
| 17246 | ENSMUSG00000019923 | Zwint | 52696 [Gene Symbol: Zwint] [Locus Tag: ] [Chromosome: 10] [Map Location: 10 B5.3\|1 | 6.569355551 | 1.03E-17 | 2.59E-16 |
| 97 | ENSMUSG00000073008 | Gpr174 | 213439 [Gene Symbol: Gpr174] [Locus Tag: ] [Chromosome: X] [Map Location: X\|X D] [I | 6.566854288 | 0.003122 | 0.007293 |
| 7096 | ENSMUSG00000026107 | Nabp1 | 109019 [Gene Symbol: Nabp1] [Locus Tag: ] [Chromosome: 1] [Map Location: 1\|1 C1] [I | 6.56552318 | 1.07E-09 | 7.65E-09 |
| 14079 | ENSMUSG00000045284 | Dcaf12l1 | 245404 [Gene Symbol: Dcaf12l1] [Locus Tag: ] [Chromosome: X] [Map Location: X\|X A4 | 6.559693312 | 6.89E-15 | 1.12E-13 |
| 7592 | ENSMUSG00000031226 | Pbdc1 | 67683 [Gene Symbol: Pbdc1] [Locus Tag: ] [Chromosome: X] [Map Location: X\|X C3] [De | 6.557318516 | 1.14E-20 | 4.73E-19 |
| 6351 | ENSMUSG00000034724 | Cnot6l | 231464 [Gene Symbol: Cnot6l] [Locus Tag: ] [Chromosome: 5] [Map Location: 5\|5 E3] [I | 6.552280972 | 5.75E-23 | 3.50E-21 |
| 4851 | ENSMUSG00000028397 | Kdm4c | 76804 [Gene Symbol: Kdm4c] [Locus Tag: ] [Chromosome: 4] [Map Location: 4\|4 C3] [D | 6.549345973 | 9.08E-18 | 2.31E-16 |
| 17008 | ENSMUSG00000031167 | Rbm3 | 19652 [Gene Symbol: Rbm3] [Locus Tag: ] [Chromosome: X] [Map Location: X A1.1\|X 3. | 6.538532082 | 1.84E-13 | 2.38E-12 |
| 9939 | ENSMUSG00000015733 | Capza2 | 12343 [Gene Symbol: Capza2] [Locus Tag: ] [Chromosome: 6] [Map Location: 6 A2\|6 7.9 | 6.528780821 | 4.59E-22 | 2.37E-20 |
| 14790 | ENSMUSG00000023505 | Cdca3 | 14793 [Gene Symbol: Cdca3] [Locus Tag: ] [Chromosome: 6] [Map Location: 6 F2\|6 59.1 | 6.515563852 | 2.34E-05 | 8.34E-05 |
| 14047 | ENSMUSG00000025940 | Tmem70 | 70397 [Gene Symbol: Tmem70] [Locus Tag: ] [Chromosome: 1] [Map Location: 1\|1 A3] | 6.515505634 | 1.72E-14 | 2.65E-13 |
| 5723 | ENSMUSG00000031297 | Slc7a3 | 11989 [Gene Symbol: Slc7a3] [Locus Tag: ] [Chromosome: X] [Map Location: X\|X C3] [De | 6.505710868 | 6.81E-05 | 0.000221 |
| 18222 | ENSMUSG00000019975 | Ikbip | 67454 [Gene Symbol: Ikbip] [Locus Tag: ] [Chromosome: 10] [Map Location: 10\|10 C2] | 6.489273106 | 4.79E-13 | 5.79E-12 |
| 17407 | ENSMUSG00000029530 | Ccr9 | 12769 [Gene Symbol: Ccr9] [Locus Tag: ] [Chromosome: 9] [Map Location: 9 74.33 cM\| | 6.486625017 | 2.79E-20 | 1.08E-18 |
| 10088 | ENSMUSG00000042133 | Ppig | 228005 [Gene Symbol: Ppig] [Locus Tag: ] [Chromosome: 2] [Map Location: 2\|2 C2] [De | 6.485786977 | 1.04E-14 | 1.64E-13 |
| 12735 | ENSMUSG00000044906 | 4930503L19Rik | 269033 [Gene Symbol: 4930503L19Rik] [Locus Tag: ] [Chromosome: 18] [Map Locatior | 6.485505079 | 2.12E-10 | 1.69E-09 |
| 8632 | ENSMUSG00000037826 | Ppm1k | 243382 [Gene Symbol: Ppm1k] [Locus Tag: ] [Chromosome: 6] [Map Location: 6\|6 B3] [ | 6.483448318 | 2.18E-19 | 7.28E-18 |
| 13332 | ENSMUSG00000027346 | Gpcpd1 | 74182 [Gene Symbol: Gpcpd1] [Locus Tag: ] [Chromosome: 2] [Map Location: 2\|2 F3] [I | 6.477108166 | 2.79E-15 | 4.80E-14 |
| 13337 | ENSMUSG00000096039 | D830030K20Rik | 320333 [Gene Symbol: D830030K20Rik] [Locus Tag: ] [Chromosome: 14] [Map Locatior | 6.476998059 | 0.011861 | 0.024263 |
| 3627 | ENSMUSG00000032757 | Bet1 | 12068 [Gene Symbol: Bet1] [Locus Tag: ] [Chromosome: 6] [Map Location: 6\|6 A1] [Des | 6.462745105 | 1.12E-13 | 1.51E-12 |
| 14271 | ENSMUSG00000066324 | Impad1 | 242291 [Gene Symbol: Impad1] [Locus Tag: ] [Chromosome: 4] [Map Location: 4\|4 A1] | 6.461497908 | 3.01E-22 | 1.58E-20 |
| 8158 | ENSMUSG00000071253 | Slc25a16 | 73132 [Gene Symbol: Slc25a16] [Locus Tag: ] [Chromosome: 10] [Map Location: 10\|10 | 6.453523078 | 1.43E-19 | 4.95E-18 |
| 17813 | ENSMUSG00000042712 | Wbp5 | 22381 [Gene Symbol: Wbp5] [Locus Tag: ] [Chromosome: X] [Map Location: X\|X F1] [De | 6.445151717 | 7.84E-14 | 1.09E-12 |
| 15439 | ENSMUSG00000037857 | Nufip2 | 68564 [Gene Symbol: Nufip2] [Locus Tag: ] [Chromosome: 11] [Map Location: 11\|11 B5 | 6.443645876 | 4.17E-20 | 1.58E-18 |
| 603 | ENSMUSG00000029254 | Stap1 | 56792 [Gene Symbol: Stap1] [Locus Tag: ] [Chromosome: 5] [Map Location: 5\|5 E1] [Des | 6.433188394 | 0.000184 | 0.000552 |
| 5423 | ENSMUSG00000025059 | Gk | 14933 [Gene Symbol: Gk] [Locus Tag: ] [Chromosome: X] [Map Location: X 39.32 cM\|X ( | 6.428536478 | 4.10E-12 | 4.25E-11 |
| 3588 | ENSMUSG00000027552 | E2f5 | 13559 [Gene Symbol: E2f5] [Locus Tag: ] [Chromosome: 3] [Map Location: 3\|3 A1] [Desc | 6.416873829 | 4.16E-17 | 9.73E-16 |
| 774 | ENSMUSG00000014956 | Ppp1cb | 19046 [Gene Symbol: Ppp1cb] [Locus Tag: ] [Chromosome: 5] [Map Location: 5\|5 B1] [I | 6.415965441 | 1.64E-22 | 9.15E-21 |
| 7904 | ENSMUSG00000020171 | Yeats4 | 64050 [Gene Symbol: Yeats4] [Locus Tag: ] [Chromosome: 10] [Map Location: 10\|10 D2 | 6.401101317 | 1.21E-21 | 5.83E-20 |
| 8995 | ENSMUSG00000090946 | Ccdc71l | 72123 [Gene Symbol: Ccdc71l] [Locus Tag: ] [Chromosome: 12] [Map Location: 12\|12 A | 6.401078679 | 2.37E-18 | 6.68E-17 |
| 9505 | ENSMUSG00000075700 | Selt | 69227 [Gene Symbol: Selt] [Locus Tag: ] [Chromosome: 3] [Map Location: 3\|3 D] [Descri | 6.395106916 | 4.24E-21 | 1.89E-19 |
| 17918 | ENSMUSG00000030471 | Zdhhc13 | 243983 [Gene Symbol: Zdhhc13] [Locus Tag: ] [Chromosome: 7] [Map Location: 7\|7 B4 | 6.384982637 | 5.07E-19 | 1.58E-17 |
| 4296 | ENSMUSG00000048540 | Nhlh2 | 18072 [Gene Symbol: Nhlh2] [Locus Tag: ] [Chromosome: 3] [Map Location: 3\|3 F2.2] [I | 6.383113235 | 1.61E-10 | 1.31E-09 |
| 15967 | ENSMUSG00000018199 | Trove2 | 20822 [Gene Symbol: Trove2] [Locus Tag: ] [Chromosome: 1] [Map Location: 1 F\|1 62.5 | 6.376878033 | 6.57E-22 | 3.30E-20 |
| 17485 | ENSMUSG00000026103 | Gls | 14660 [Gene Symbol: Gls] [Locus Tag: ] [Chromosome: 1] [Map Location: 1 C1.1\|1 26.8 | 6.365799921 | 9.31E-14 | 1.27E-12 |
| 6290 | ENSMUSG00000028884 | Rpa2 | 19891 [Gene Symbol: Rpa2] [Locus Tag: ] [Chromosome: 4] [Map Location: 4\|4 D2.3] [D | 6.360157505 | 7.97E-10 | 5.83E-09 |
| 1650 | ENSMUSG00000018341 | Il12rb2 | 16162 [Gene Symbol: Il12rb2] [Locus Tag: ] [Chromosome: 6] [Map Location: 6 C1\|6 30 | 6.358171348 | 2.37E-19 | 7.88E-18 |
| 2172 | ENSMUSG00000026032 | Ndufb3 | 66495 [Gene Symbol: Ndufb3] [Locus Tag: ] [Chromosome: 1] [Map Location: 1\|1 C1.3] | 6.351631643 | 1.47E-13 | 1.95E-12 |
| 1076 | ENSMUSG00000021639 | Gtf2h2 | 23894 [Gene Symbol: Gtf2h2] [Locus Tag: ] [Chromosome: 13] [Map Location: 13 D1\|1: | 6.348176802 | 1.53E-12 | 1.70E-11 |
| 245 | ENSMUSG00000022604 | Cep97 | 74201 [Gene Symbol: Cep97] [Locus Tag: ] [Chromosome: 16] [Map Location: 16\|16 C1 | 6.338746726 | 1.15E-12 | 1.31E-11 |
| 1720 | ENSMUSG00000060703 | Cd302 | 66205 [Gene Symbol: Cd302] [Locus Tag: ] [Chromosome: 2] [Map Location: 2\|2 C1.1] [ | 6.33667427 | 9.95E-11 | 8.33E-10 |
| 6148 | ENSMUSG00000021938 | Pspc1 | 66645 [Gene Symbol: Pspc1] [Locus Tag: ] [Chromosome: 14] [Map Location: 14\|14 C3] | 6.335161691 | 5.89E-19 | 1.81E-17 |
| 4927 | ENSMUSG00000046096 | BC030336 | 233812 [Gene Symbol: BC030336] [Locus Tag: ] [Chromosome: 7] [Map Location: 7\|7 F | 6.334957495 | 5.14E-21 | 2.26E-19 |
| 15470 | ENSMUSG00000027896 | Slc16a4 | 229699 [Gene Symbol: Slc16a4] [Locus Tag: ] [Chromosome: 3] [Map Location: 3\|3 F2.3 | 6.328770685 | 1.59E-12 | 1.76E-11 |
| 18431 | ENSMUSG00000035212 | Leprot | 230514 [Gene Symbol: Leprot] [Locus Tag: ] [Chromosome: 4] [Map Location: 4\|4 C6] [I | 6.328724552 | 4.32E-17 | 1.00E-15 |
| 3681 | ENSMUSG00000020246 | Hcfc2 | 67933 [Gene Symbol: Hcfc2] [Locus Tag: ] [Chromosome: 10] [Map Location: 10\|10 C1] | 6.324807196 | 7.70E-20 | 2.79E-18 |
| 679 | ENSMUSG00000021590 | Spata9 | 75571 [Gene Symbol: Spata9] [Locus Tag: ] [Chromosome: 13] [Map Location: 13\|13 C1 | 6.323549135 | 9.60E-14 | 1.31E-12 |
| 16507 | ENSMUSG00000004127 | Trmt10a | 108943 [Gene Symbol: Trmt10a] [Locus Tag: ] [Chromosome: 3] [Map Location: 3\|3 G3 | 6.322091617 | 9.04E-13 | 1.05E-11 |
| 8108 | ENSMUSG00000044349 | Snhg11 | 319317 [Gene Symbol: Snhg11] [Locus Tag: ] [Chromosome: 2] [Map Location: 2\|2 H1] | 6.316748481 | 8.77E-10 | 6.38E-09 |
| 10931 | ENSMUSG00000021714 | Cenpk | 60411 [Gene Symbol: Cenpk] [Locus Tag: ] [Chromosome: 13] [Map Location: 13\|13 D1 | 6.305919048 | 2.43E-08 | 1.43E-07 |
| 13787 | ENSMUSG00000066456 | Hmgn3 | 94353 [Gene Symbol: Hmgn3] [Locus Tag: ] [Chromosome: 9] [Map Location: 9\|9 E3.1] | 6.30036137 | 1.68E-13 | 2.18E-12 |
| 17639 | ENSMUSG00000052428 | Tmco1 | 68944 [Gene Symbol: Tmco1] [Locus Tag: ] [Chromosome: 1] [Map Location: 1\|1 H2.3] | 6.294143336 | 1.53E-15 | 2.76E-14 |
| 3742 | ENSMUSG00000028252 | Ccnc | 51813 [Gene Symbol: Ccnc] [Locus Tag: ] [Chromosome: 4] [Map Location: 4\|4 A3] [Des | 6.29325898 | 4.86E-14 | 6.97E-13 |
| 11199 | ENSMUSG00000021969 | Zdhhc20 | 75965 [Gene Symbol: Zdhhc20] [Locus Tag: ] [Chromosome: 14] [Map Location: 14\|14 | 6.283105429 | 1.95E-20 | 7.67E-19 |
| 3713 | ENSMUSG00000063889 | Crem | 12916 [Gene Symbol: Crem] [Locus Tag: ] [Chromosome: 18] [Map Location: 18\|18 A] [I | 6.278268473 | 1.24E-14 | 1.93E-13 |
| 18370 | ENSMUSG00000040139 | 9430038I01Rik | 77252 [Gene Symbol: 9430038I01Rik] [Locus Tag: ] [Chromosome: 7] [Map Location: 7 | 6.273797642 | 5.99E-13 | 7.13E-12 |
| 8698 | ENSMUSG00000040710 | St8sia4 | 20452 [Gene Symbol: St8sia4] [Locus Tag: ] [Chromosome: 1] [Map Location: 1\|1 D] [De | 6.273588296 | 2.51E-13 | 3.16E-12 |
| 4687 | ENSMUSG00000020738 | Sumo2 | 170930 [Gene Symbol: Sumo2] [Locus Tag: ] [Chromosome: 11] [Map Location: 11\|11 E | 6.271860005 | 1.10E-10 | 9.14E-10 |
| 17873 | ENSMUSG00000030525 | Chrna7 | 11441 [Gene Symbol: Chrna7] [Locus Tag: ] [Chromosome: 7] [Map Location: 7 C\|7 34.4 | 6.256831245 | 4.74E-19 | 1.50E-17 |
| 7618 | ENSMUSG00000069184 | Zfp72 | 238722 [Gene Symbol: Zfp72] [Locus Tag: ] [Chromosome: 13] [Map Location: 13\|13 C1 | 6.256611188 | 2.51E-08 | 1.47E-07 |
| 10710 | ENSMUSG00000010608 | Rbm25 | 67039 [Gene Symbol: Rbm25] [Locus Tag: ] [Chromosome: 12] [Map Location: 12\|12 D | 6.252480948 | 2.53E-14 | 3.81E-13 |
| 9102 | ENSMUSG00000035840 | Lysmd3 | 80289 [Gene Symbol: Lysmd3] [Locus Tag: ] [Chromosome: 13] [Map Location: 13\|13 C | 6.250342179 | 2.51E-19 | 8.26E-18 |
| 15962 | ENSMUSG00000021621 | Zcchc9 | 69085 [Gene Symbol: Zcchc9] [Locus Tag: ] [Chromosome: 13] [Map Location: 13\|13 C3 | 6.239280473 | 2.13E-13 | 2.72E-12 |
| 4285 | ENSMUSG00000021047 | Nova1 | 664883 [Gene Symbol: Nova1] [Locus Tag: ] [Chromosome: 12] [Map Location: 12 B3\|1 | 6.223004546 | 2.51E-16 | 5.19E-15 |
| 15185 | ENSMUSG00000074829 | 2010315B03Rik | 630836 [Gene Symbol: 2010315B03Rik] [Locus Tag: ] [Chromosome: 9] [Map Location: | 6.22116526 | 3.77E-11 | 3.37E-10 |
| 17456 | ENSMUSG00000035033 | Tbr1 | 21375 [Gene Symbol: Tbr1] [Locus Tag: ] [Chromosome: 2] [Map Location: 2 C1.3\|2 35. | 6.22106775 | 1.14E-16 | 2.51E-15 |
| 761 | ENSMUSG00000059325 | Hopx | 74318 [Gene Symbol: Hopx] [Locus Tag: ] [Chromosome: 5] [Map Location: 5\|5 C3.3] [D | 6.220094568 | 3.52E-11 | 3.16E-10 |
| 17719 | ENSMUSG00000039086 | Ss18l1 | 269397 [Gene Symbol: Ss18l1] [Locus Tag: ] [Chromosome: 2] [Map Location: 2\|2 H4] [I | 6.208622627 | 9.06E-18 | 2.31E-16 |
| 7047 | ENSMUSG00000032051 | Fdx1 | 14148 [Gene Symbol: Fdx1] [Locus Tag: ] [Chromosome: 9] [Map Location: 9\|9 B] [Descr | 6.201443778 | 4.40E-10 | 3.33E-09 |
| 11266 | ENSMUSG00000039396 | Neil3 | 234258 [Gene Symbol: Neil3] [Locus Tag: ] [Chromosome: 8] [Map Location: 8\|8 B1.3] [ | 6.19424673 | 3.13E-11 | 2.82E-10 |
| 3165 | ENSMUSG00000016534 | Lamp2 | 16784 [Gene Symbol: Lamp2] [Locus Tag: ] [Chromosome: X] [Map Location: X A3.3\|X 2 | 6.194108937 | 1.16E-18 | 3.42E-17 |
| 7799 | ENSMUSG00000028578 | Caap1 | 67770 [Gene Symbol: Caap1] [Locus Tag: ] [Chromosome: 4] [Map Location: 4\|4 C5] [De | 6.191336199 | 2.66E-16 | 5.48E-15 |
| 11563 | ENSMUSG00000090266 | Mettl23 | 74319 [Gene Symbol: Mettl23] [Locus Tag: ] [Chromosome: 11] [Map Location: 11\|11 E | 6.191289147 | 7.03E-21 | 3.04E-19 |
| 8769 | ENSMUSG00000041459 | Tardbp | 230908 [Gene Symbol: Tardbp] [Locus Tag: ] [Chromosome: 4] [Map Location: 4\|4 E2] [I | 6.189357299 | 6.76E-21 | 2.93E-19 |
| 14486 | ENSMUSG00000050697 | Prkaa1 | 105787 [Gene Symbol: Prkaa1] [Locus Tag: ] [Chromosome: 15] [Map Location: 15\|15 A | 6.177589383 | 4.21E-19 | 1.34E-17 |

| | | | | | | |
|---|---|---|---|---|---|---|
| 11443 | ENSMUSG00000028955 | Vamp3 | 22319 [Gene Symbol: Vamp3] [Locus Tag: ] [Chromosome: 4] [Map Location: 4\|4 E1] [D | 6.175662497 | 9.16E-15 | 1.46E-13 |
| 15413 | ENSMUSG00000033184 | Tmed7 | 66676 [Gene Symbol: Tmed7] [Locus Tag: ] [Chromosome: 18] [Map Location: 18\|18 C] | 6.167668435 | 9.54E-21 | 4.00E-19 |
| 11932 | ENSMUSG00000032463 | Faim | 23873 [Gene Symbol: Faim] [Locus Tag: ] [Chromosome: 9] [Map Location: 9\|9 E3.3] [D | 6.162813154 | 2.80E-12 | 2.99E-11 |
| 3719 | ENSMUSG00000050875 | A730017C20Rik | 225583 [Gene Symbol: A730017C20Rik] [Locus Tag: ] [Chromosome: 18] [Map Locatior | 6.149297158 | 1.64E-18 | 4.74E-17 |
| 6184 | ENSMUSG00000004096 | Cwc15 | 66070 [Gene Symbol: Cwc15] [Locus Tag: ] [Chromosome: 9] [Map Location: 9\|9 A3] [D | 6.147660144 | 3.20E-15 | 5.45E-14 |
| 9423 | ENSMUSG00000046101 | Mcmdc2 | 240697 [Gene Symbol: Mcmdc2] [Locus Tag: ] [Chromosome: 1] [Map Location: 1\|1 A2 | 6.146083638 | 1.12E-05 | 4.21E-05 |
| 17613 | ENSMUSG00000003062 | Stard3nl | 76205 [Gene Symbol: Stard3nl] [Locus Tag: ] [Chromosome: 13] [Map Location: 13\|13 / | 6.143528617 | 2.36E-16 | 4.93E-15 |
| 13111 | ENSMUSG00000026239 | Pde6d | 18582 [Gene Symbol: Pde6d] [Locus Tag: ] [Chromosome: 1] [Map Location: 1\|1 D] [Des | 6.130677528 | 1.06E-12 | 1.21E-11 |
| 17948 | ENSMUSG00000041431 | Ccnb1 | 268697 [Gene Symbol: Ccnb1] [Locus Tag: ] [Chromosome: 13] [Map Location: 13 D1\|1 | 6.127175881 | 0.001796 | 0.004422 |
| 779 | ENSMUSG00000022507 | 1810013L24Rik | 69053 [Gene Symbol: 1810013L24Rik] [Locus Tag: ] [Chromosome: 16] [Map Location: | 6.124704632 | 8.54E-22 | 4.22E-20 |
| 2102 | ENSMUSG00000031683 | Lsm6 | 78651 [Gene Symbol: Lsm6] [Locus Tag: ] [Chromosome: 8] [Map Location: 8\|8 C1] [Des | 6.120698716 | 1.68E-17 | 4.11E-16 |
| 11727 | ENSMUSG00000090000 | Ier3ip1 | 66191 [Gene Symbol: Ier3ip1] [Locus Tag: ] [Chromosome: 18] [Map Location: 18\|18 E3 | 6.12048769 | 4.65E-17 | 1.07E-15 |
| 7134 | ENSMUSG00000096225 | Lhx8 | 16875 [Gene Symbol: Lhx8] [Locus Tag: ] [Chromosome: 3] [Map Location: 3\|3 H3-H4] [ | 6.109269211 | 6.52E-09 | 4.21E-08 |
| 7532 | ENSMUSG00000050914 | Ankrd37 | 654824 [Gene Symbol: Ankrd37] [Locus Tag: ] [Chromosome: 8] [Map Location: 8\|8 B1. | 6.102343652 | 0.000173 | 0.00052 |
| 15006 | ENSMUSG00000031885 | Cbfb | 12400 [Gene Symbol: Cbfb] [Locus Tag: ] [Chromosome: 8] [Map Location: 8 D3\|8 53.0 | 6.100834158 | 2.12E-15 | 3.73E-14 |
| 7893 | ENSMUSG00000002633 | Shh | 20423 [Gene Symbol: Shh] [Locus Tag: ] [Chromosome: 5] [Map Location: 5 B1\|5 14.39 | 6.098074937 | 1.19E-07 | 6.23E-07 |
| 13702 | ENSMUSG00000054204 | Fam150b | 100294583 [Gene Symbol: Fam150b] [Locus Tag: ] [Chromosome: 12] [Map Location: 1 | 6.092633956 | 5.06E-05 | 0.000168 |
| 10969 | ENSMUSG00000018800 | Abca5 | 217265 [Gene Symbol: Abca5] [Locus Tag: ] [Chromosome: 11] [Map Location: 11\|11 E | 6.088736597 | 4.07E-19 | 1.30E-17 |
| 11769 | ENSMUSG00000027287 | Snap23 | 20619 [Gene Symbol: Snap23] [Locus Tag: ] [Chromosome: 2] [Map Location: 2 60.37 cl | 6.0866655 | 5.24E-11 | 4.59E-10 |
| 7195 | ENSMUSG00000062397 | Zfp706 | 68036 [Gene Symbol: Zfp706] [Locus Tag: ] [Chromosome: 15] [Map Location: 15\|15 B3 | 6.077465153 | 4.67E-21 | 2.07E-19 |
| 4164 | ENSMUSG00000026718 | Stam | 20844 [Gene Symbol: Stam] [Locus Tag: ] [Chromosome: 2] [Map Location: 2\|2 A2-B] [D | 6.075391523 | 1.90E-20 | 7.52E-19 |
| 13515 | ENSMUSG00000043061 | Tmem18 | 211986 [Gene Symbol: Tmem18] [Locus Tag: ] [Chromosome: 12] [Map Location: 12\|12 | 6.073853527 | 5.06E-18 | 1.35E-16 |
| 572 | ENSMUSG00000030443 | Zfp583 | 213011 [Gene Symbol: Zfp583] [Locus Tag: ] [Chromosome: 7] [Map Location: 7\|7 A1] [ | 6.066497872 | 1.97E-12 | 2.16E-11 |
| 17265 | ENSMUSG00000021930 | Spryd7 | 66674 [Gene Symbol: Spryd7] [Locus Tag: ] [Chromosome: 14] [Map Location: 14\|14 D | 6.066259104 | 1.42E-15 | 2.59E-14 |
| 14551 | ENSMUSG00000020053 | Igf1 | 16000 [Gene Symbol: Igf1] [Locus Tag: ] [Chromosome: 10] [Map Location: 10 C1\|10 43 | 6.064687332 | 6.83E-11 | 5.88E-10 |
| 11676 | ENSMUSG00000024539 | Ptpn2 | 19255 [Gene Symbol: Ptpn2] [Locus Tag: ] [Chromosome: 18] [Map Location: 18\|18 E1 | 6.064590851 | 8.22E-14 | 1.13E-12 |
| 990 | ENSMUSG00000019863 | Qrsl1 | 76563 [Gene Symbol: Qrsl1] [Locus Tag: ] [Chromosome: 10] [Map Location: 10\|10 B2] | 6.053962831 | 4.10E-14 | 5.95E-13 |
| 17075 | ENSMUSG00000027018 | Hat1 | 107435 [Gene Symbol: Hat1] [Locus Tag: ] [Chromosome: 2] [Map Location: 2\|2 C2] [De | 6.050808785 | 2.70E-15 | 4.67E-14 |
| 10703 | ENSMUSG00000058704 | Memo1 | 76890 [Gene Symbol: Memo1] [Locus Tag: ] [Chromosome: 17] [Map Location: 17\|17 E | 6.050315194 | 1.41E-16 | 3.06E-15 |
| 5653 | ENSMUSG00000022528 | Hes1 | 15205 [Gene Symbol: Hes1] [Locus Tag: ] [Chromosome: 16] [Map Location: 16 B2\|16 2 | 6.049447918 | 2.84E-13 | 3.55E-12 |
| 6979 | ENSMUSG00000055373 | Fut9 | 14348 [Gene Symbol: Fut9] [Locus Tag: ] [Chromosome: 4] [Map Location: 4 A3\|4 10.5 | 6.047340778 | 6.55E-22 | 3.30E-20 |
| 4343 | ENSMUSG00000035597 | Prpf39 | 328110 [Gene Symbol: Prpf39] [Locus Tag: ] [Chromosome: 12] [Map Location: 12\|12 C | 6.043008547 | 1.39E-12 | 1.55E-11 |
| 740 | ENSMUSG00000071533 | Pcnp | 76302 [Gene Symbol: Pcnp] [Locus Tag: ] [Chromosome: 16] [Map Location: 16\|16 C1. | 6.026583825 | 9.45E-20 | 3.39E-18 |
| 16626 | ENSMUSG00000045414 | 1190002N15Rik | 68861 [Gene Symbol: 1190002N15Rik] [Locus Tag: ] [Chromosome: 9] [Map Location: | 6.022632972 | 4.07E-19 | 1.30E-17 |
| 10925 | ENSMUSG00000027900 | Dram2 | 67171 [Gene Symbol: Dram2] [Locus Tag: ] [Chromosome: 3] [Map Location: 3\|3 F3] [D | 6.017347696 | 7.12E-18 | 1.84E-16 |
| 1925 | ENSMUSG00000033981 | Gria2 | 14800 [Gene Symbol: Gria2] [Locus Tag: ] [Chromosome: 3] [Map Location: 3 E3\|3 35.5 | 6.010985691 | 2.04E-18 | 5.82E-17 |
| 11741 | ENSMUSG00000021748 | Pdhb | 68263 [Gene Symbol: Pdhb] [Locus Tag: ] [Chromosome: 14] [Map Location: 14\|14 A1] | 6.010562275 | 5.71E-18 | 1.50E-16 |
| 18447 | ENSMUSG00000039114 | Nrn1 | 68404 [Gene Symbol: Nrn1] [Locus Tag: ] [Chromosome: 13] [Map Location: 13\|13 A3. | 5.99808685 | 9.79E-13 | 1.13E-11 |
| 14682 | ENSMUSG00000026439 | Rbbp5 | 213464 [Gene Symbol: Rbbp5] [Locus Tag: ] [Chromosome: 1] [Map Location: 1\|1 E4] [D | 5.997233493 | 4.58E-18 | 1.23E-16 |
| 16819 | ENSMUSG00000029265 | Dr1 | 13486 [Gene Symbol: Dr1] [Locus Tag: ] [Chromosome: 5] [Map Location: 5 F\|5 52.82 c | 5.997126962 | 9.35E-16 | 1.76E-14 |
| 10277 | ENSMUSG00000031198 | Fundc2 | 67391 [Gene Symbol: Fundc2] [Locus Tag: ] [Chromosome: X] [Map Location: X\|X] [Desc | 5.996129266 | 2.52E-10 | 1.98E-09 |
| 9867 | ENSMUSG00000098923 | Tmem185b | 226351 [Gene Symbol: Tmem185b] [Locus Tag: ] [Chromosome: 1] [Map Location: 1\|1 | 5.99449612 | 2.89E-11 | 2.63E-10 |
| 3261 | ENSMUSG00000014547 | Wdfy2 | 268752 [Gene Symbol: Wdfy2] [Locus Tag: ] [Chromosome: 14] [Map Location: 14\|14 D | 5.994363097 | 1.64E-13 | 2.14E-12 |
| 6618 | ENSMUSG00000028896 | Rcc1 | 100088 [Gene Symbol: Rcc1] [Locus Tag: ] [Chromosome: 4] [Map Location: 4\|4 D2.3] [ | 5.988869531 | 1.15E-11 | 1.12E-10 |
| 6352 | ENSMUSG00000020385 | Clk4 | 12750 [Gene Symbol: Clk4] [Locus Tag: ] [Chromosome: 11] [Map Location: 11\|11 B1.3 | 5.988192337 | 1.89E-12 | 2.08E-11 |
| 7405 | ENSMUSG00000034158 | Lrrc58 | 320184 [Gene Symbol: Lrrc58] [Locus Tag: ] [Chromosome: 16] [Map Location: 16\|16 B | 5.981614763 | 1.06E-18 | 3.14E-17 |
| 17224 | ENSMUSG00000046516 | Cox17 | 12856 [Gene Symbol: Cox17] [Locus Tag: ] [Chromosome: 16] [Map Location: 16\|16 B3 | 5.970802266 | 1.87E-07 | 9.48E-07 |
| 15264 | ENSMUSG00000040367 | Lrrd1 | 242838 [Gene Symbol: Lrrd1] [Locus Tag: ] [Chromosome: 5] [Map Location: 5\|5 A1] [D | 5.969179526 | 5.98E-05 | 0.000196 |
| 14099 | ENSMUSG00000029687 | Ezh2 | 14056 [Gene Symbol: Ezh2] [Locus Tag: ] [Chromosome: 6] [Map Location: 6 22.92 cM\| | 5.965320147 | 1.23E-12 | 1.39E-11 |
| 4531 | ENSMUSG00000079283 | 2310009B15Rik | 69549 [Gene Symbol: 2310009B15Rik] [Locus Tag: ] [Chromosome: 1] [Map Location: 1 | 5.934175493 | 1.81E-05 | 6.60E-05 |
| 7380 | ENSMUSG00000022788 | Fgd4 | 224014 [Gene Symbol: Fgd4] [Locus Tag: ] [Chromosome: 16] [Map Location: 16\|16 A3 | 5.932401014 | 2.81E-13 | 3.52E-12 |
| 7777 | ENSMUSG00000022119 | Rbm26 | 74213 [Gene Symbol: Rbm26] [Locus Tag: ] [Chromosome: 14] [Map Location: 14\|14 E | 5.932157424 | 3.36E-14 | 4.94E-13 |
| 5012 | ENSMUSG00000028568 | Btf3l4 | 70533 [Gene Symbol: Btf3l4] [Locus Tag: ] [Chromosome: 4] [Map Location: 4\|4 C7] [De | 5.931815274 | 3.71E-15 | 6.25E-14 |
| 9498 | ENSMUSG00000038736 | Nudcd1 | 67429 [Gene Symbol: Nudcd1] [Locus Tag: ] [Chromosome: 15] [Map Location: 15\|15 B | 5.931197406 | 4.69E-16 | 9.26E-15 |
| 12016 | ENSMUSG00000022797 | Tfrc | 22042 [Gene Symbol: Tfrc] [Locus Tag: ] [Chromosome: 16] [Map Location: 16 B3\|16 2 | 5.923953801 | 4.67E-16 | 9.24E-15 |
| 6947 | ENSMUSG00000021597 | Slf1 | 105377 [Gene Symbol: Slf1] [Locus Tag: ] [Chromosome: 13] [Map Location: 13\|13 C1] | 5.904869888 | 1.07E-18 | 3.15E-17 |
| 16578 | ENSMUSG00000035946 | Gsx2 | 14843 [Gene Symbol: Gsx2] [Locus Tag: ] [Chromosome: 5] [Map Location: 5 C3.3\|5 39. | 5.902429726 | 5.56E-06 | 2.20E-05 |
| 578 | ENSMUSG00000025981 | Coq10b | 67876 [Gene Symbol: Coq10b] [Locus Tag: ] [Chromosome: 1] [Map Location: 1\|1 C1.2 | 5.900998044 | 7.20E-12 | 7.25E-11 |
| 4059 | ENSMUSG00000022130 | Tgds | 76355 [Gene Symbol: Tgds] [Locus Tag: ] [Chromosome: 14] [Map Location: 14\|14 E4] [ | 5.89902751 | 5.13E-11 | 4.50E-10 |
| 9688 | ENSMUSG00000042225 | Ammecr1 | 56068 [Gene Symbol: Ammecr1] [Locus Tag: ] [Chromosome: X] [Map Location: X\|X F2] | 5.897897753 | 1.78E-17 | 4.34E-16 |
| 12123 | ENSMUSG00000020265 | Sumo3 | 20610 [Gene Symbol: Sumo3] [Locus Tag: ] [Chromosome: 10] [Map Location: 10 C1\|1 | 5.892125001 | 4.84E-21 | 2.14E-19 |
| 14734 | ENSMUSG00000030615 | Tmem126a | 66271 [Gene Symbol: Tmem126a] [Locus Tag: ] [Chromosome: 7] [Map Location: 7\|7 E | 5.888942633 | 2.81E-14 | 4.18E-13 |
| 16520 | ENSMUSG00000045624 | Esf1 | 66580 [Gene Symbol: Esf1] [Locus Tag: ] [Chromosome: 2] [Map Location: 2\|2 G3] [Desc | 5.887976958 | 2.08E-13 | 2.66E-12 |
| 9698 | ENSMUSG00000027011 | Ube2e3 | 22193 [Gene Symbol: Ube2e3] [Locus Tag: ] [Chromosome: 2] [Map Location: 2\|2 D] [D | 5.887569886 | 8.68E-21 | 3.67E-19 |
| 18105 | ENSMUSG00000047854 | Stx19 | 68159 [Gene Symbol: Stx19] [Locus Tag: ] [Chromosome: 16] [Map Location: 16\|16 C1. | 5.880940203 | 3.13E-05 | 0.000109 |
| 2165 | ENSMUSG00000032905 | Atg12 | 67526 [Gene Symbol: Atg12] [Locus Tag: ] [Chromosome: 18] [Map Location: 18\|18 C] [ | 5.879682088 | 1.13E-17 | 2.82E-16 |
| 1985 | ENSMUSG00000003131 | Pafah1b2 | 18475 [Gene Symbol: Pafah1b2] [Locus Tag: ] [Chromosome: 9] [Map Location: 9\|9 A5. | 5.878284166 | 8.01E-21 | 3.42E-19 |
| 1990 | ENSMUSG00000021432 | Slc35b3 | 108652 [Gene Symbol: Slc35b3] [Locus Tag: ] [Chromosome: 13] [Map Location: 13\|13 | 5.871833076 | 3.65E-13 | 4.47E-12 |
| 1222 | ENSMUSG00000033953 | Ppp3r1 | 19058 [Gene Symbol: Ppp3r1] [Locus Tag: ] [Chromosome: 11] [Map Location: 11\|11 A | 5.864211401 | 3.64E-21 | 1.63E-19 |
| 2808 | ENSMUSG00000028228 | Cpne3 | 70568 [Gene Symbol: Cpne3] [Locus Tag: ] [Chromosome: 4] [Map Location: 4\|4 A3] [De | 5.857870007 | 1.34E-15 | 2.45E-14 |
| 14876 | ENSMUSG00000028132 | Tmem56 | 99887 [Gene Symbol: Tmem56] [Locus Tag: ] [Chromosome: 3] [Map Location: 3\|3 G3] | 5.856210393 | 7.19E-20 | 2.62E-18 |
| 14588 | ENSMUSG00000034723 | Tmx4 | 52837 [Gene Symbol: Tmx4] [Locus Tag: ] [Chromosome: 2] [Map Location: 2 65.66 cM | 5.855086308 | 6.41E-20 | 2.36E-18 |
| 14654 | ENSMUSG00000030643 | Rab30 | 75985 [Gene Symbol: Rab30] [Locus Tag: ] [Chromosome: 7] [Map Location: 7\|7 E1] [De | 5.85378967 | 1.87E-18 | 5.36E-17 |
| 11339 | ENSMUSG00000031176 | Dynlt3 | 67117 [Gene Symbol: Dynlt3] [Locus Tag: ] [Chromosome: X] [Map Location: X\|X A1.1] | 5.852937309 | 1.06E-11 | 1.04E-10 |
| 7572 | ENSMUSG00000014907 | Naf1 | 234344 [Gene Symbol: Naf1] [Locus Tag: ] [Chromosome: 8] [Map Location: 8\|8 B3.3] [ | 5.852606521 | 1.17E-13 | 1.56E-12 |
| 14379 | ENSMUSG00000026784 | Pdss1 | 56075 [Gene Symbol: Pdss1] [Locus Tag: ] [Chromosome: 2] [Map Location: 2\|2 A3] [De | 5.849460299 | 8.82E-15 | 1.42E-13 |
| 1951 | ENSMUSG00000027775 | Mfsd1 | 66868 [Gene Symbol: Mfsd1] [Locus Tag: ] [Chromosome: 3] [Map Location: 3\|3 E2] [De | 5.841291061 | 2.31E-15 | 4.04E-14 |
| 8192 | ENSMUSG00000068250 | Amn1 | 232566 [Gene Symbol: Amn1] [Locus Tag: ] [Chromosome: 6] [Map Location: 6\|6 G3] [D | 5.838681375 | 1.21E-15 | 2.22E-14 |
| 8436 | ENSMUSG00000054408 | Spcs3 | 76687 [Gene Symbol: Spcs3] [Locus Tag: ] [Chromosome: 8] [Map Location: 8\|8 B1.3] [D | 5.835508542 | 2.18E-18 | 6.17E-17 |
| 1052 | ENSMUSG00000049775 | Tmsb4x | 19241 [Gene Symbol: Tmsb4x] [Locus Tag: ] [Chromosome: X] [Map Location: X F5\|X 78 | 5.834871852 | 1.80E-14 | 2.76E-13 |
| 18211 | ENSMUSG00000024985 | Tcf7l2 | 21416 [Gene Symbol: Tcf7l2] [Locus Tag: ] [Chromosome: 19] [Map Location: 19 D2\|19 | 5.833628977 | 1.46E-11 | 1.39E-10 |
| 9985 | ENSMUSG00000029617 | Ccz1 | 231874 [Gene Symbol: Ccz1] [Locus Tag: ] [Chromosome: 5] [Map Location: 5\|5 G2] [De | 5.825104418 | 1.54E-19 | 5.27E-18 |
| 5949 | ENSMUSG00000024069 | Slc30a6 | 210148 [Gene Symbol: Slc30a6] [Locus Tag: ] [Chromosome: 17] [Map Location: 17\|17 | 5.822278576 | 1.76E-13 | 2.28E-12 |
| 7448 | ENSMUSG00000001403 | Ube2c | 68612 [Gene Symbol: Ube2c] [Locus Tag: ] [Chromosome: 2] [Map Location: 2 H3\|2 85. | 5.819439956 | 0.000139 | 0.000427 |

| | | | | | | |
|---|---|---|---|---|---|---|
| 5759 | ENSMUSG00000022359 | Wdyhv1 | 76773 [Gene Symbol: Wdyhv1] [Locus Tag: ] [Chromosome: 15] [Map Location: 15\|15 I | 5.81808073 | 3.76E-10 | 2.88E-09 |
| 12877 | ENSMUSG00000048249 | Crebrf | 77128 [Gene Symbol: Crebrf] [Locus Tag: ] [Chromosome: 17] [Map Location: 17\|17 B1 | 5.816524319 | 8.12E-17 | 1.83E-15 |
| 9121 | ENSMUSG00000019817 | Plagl1 | 22634 [Gene Symbol: Plagl1] [Locus Tag: ] [Chromosome: 10] [Map Location: 10 A2\|10 | 5.81036191 | 5.31E-14 | 7.56E-13 |
| 15910 | ENSMUSG00000020273 | Papolg | 216578 [Gene Symbol: Papolg] [Locus Tag: ] [Chromosome: 11] [Map Location: 11\|11 A | 5.804210993 | 2.06E-15 | 3.63E-14 |
| 9346 | ENSMUSG00000033882 | Rbm46 | 633285 [Gene Symbol: Rbm46] [Locus Tag: ] [Chromosome: 3] [Map Location: 3\|3 E3] [ | 5.803483482 | 0.000268 | 0.000777 |
| 10374 | ENSMUSG00000074500 | Zfp558 | 72230 [Gene Symbol: Zfp558] [Locus Tag: ] [Chromosome: 9] [Map Location: 9\|9 A3] [D | 5.791893581 | 2.43E-12 | 2.61E-11 |
| 1001 | ENSMUSG00000061778 | Mospd2 | 76763 [Gene Symbol: Mospd2] [Locus Tag: ] [Chromosome: X] [Map Location: X\|X F5] [I | 5.770088798 | 3.28E-16 | 6.64E-15 |
| 2941 | ENSMUSG00000031112 | Stk26 | 70415 [Gene Symbol: Stk26] [Locus Tag: ] [Chromosome: X] [Map Location: X\|X A3.3] [I | 5.768987017 | 5.00E-05 | 0.000167 |
| 2690 | ENSMUSG00000054099 | Slc25a40 | 319653 [Gene Symbol: Slc25a40] [Locus Tag: ] [Chromosome: 5] [Map Location: 5\|5 A1 | 5.765504075 | 4.74E-13 | 5.74E-12 |
| 9101 | ENSMUSG00000033502 | Cdc14a | 229776 [Gene Symbol: Cdc14a] [Locus Tag: ] [Chromosome: 3] [Map Location: 3\|3 G1] | 5.7652018 | 5.80E-14 | 8.22E-13 |
| 13464 | ENSMUSG00000021713 | Ppwd1 | 238831 [Gene Symbol: Ppwd1] [Locus Tag: ] [Chromosome: 13] [Map Location: 13\|13 I | 5.765030531 | 8.47E-12 | 8.41E-11 |
| 8726 | ENSMUSG00000022698 | Naa50 | 72117 [Gene Symbol: Naa50] [Locus Tag: ] [Chromosome: 16] [Map Location: 16\|16 B4 | 5.758695543 | 1.25E-19 | 4.39E-18 |
| 16258 | ENSMUSG00000022860 | Chodl | 246048 [Gene Symbol: Chodl] [Locus Tag: ] [Chromosome: 16] [Map Location: 16\|16 C | 5.75470233 | 0.000126 | 0.000388 |
| 5588 | ENSMUSG00000061130 | Ppm1b | 19043 [Gene Symbol: Ppm1b] [Locus Tag: ] [Chromosome: 17] [Map Location: 17 55.13 | 5.753621688 | 7.72E-20 | 2.79E-18 |
| 5425 | ENSMUSG00000028822 | Tmem50a | 71817 [Gene Symbol: Tmem50a] [Locus Tag: ] [Chromosome: 4] [Map Location: 4 D3\|4 | 5.753566466 | 9.62E-15 | 1.53E-13 |
| 12418 | ENSMUSG00000025898 | Cwf19l2 | 244672 [Gene Symbol: Cwf19l2] [Locus Tag: ] [Chromosome: 9] [Map Location: 9\|9 A1] | 5.746563725 | 3.33E-15 | 5.66E-14 |
| 6467 | ENSMUSG00000055963 | Triqk | 208820 [Gene Symbol: Triqk] [Locus Tag: ] [Chromosome: 4] [Map Location: 4 A1\|4] [De | 5.74334418 | 1.24E-12 | 1.40E-11 |
| 5046 | ENSMUSG00000039131 | Gipc2 | 54120 [Gene Symbol: Gipc2] [Locus Tag: ] [Chromosome: 3] [Map Location: 3\|3 H3] [De | 5.742974303 | 1.43E-08 | 8.74E-08 |
| 11400 | ENSMUSG00000051022 | Hs3st1 | 15476 [Gene Symbol: Hs3st1] [Locus Tag: ] [Chromosome: 5] [Map Location: 5 B3\|5 21. | 5.738263698 | 5.97E-19 | 1.83E-17 |
| 14675 | ENSMUSG00000038079 | Tmem237 | 381259 [Gene Symbol: Tmem237] [Locus Tag: ] [Chromosome: 1] [Map Location: 1\|1 C | 5.731650629 | 2.02E-15 | 3.57E-14 |
| 4855 | ENSMUSG00000047260 | Emc6 | 66048 [Gene Symbol: Emc6] [Locus Tag: ] [Chromosome: 11] [Map Location: 11\|11 B4] | 5.729504446 | 1.34E-16 | 2.93E-15 |
| 14502 | ENSMUSG00000020464 | Pnpt1 | 71701 [Gene Symbol: Pnpt1] [Locus Tag: ] [Chromosome: 11] [Map Location: 11\|11 A4 | 5.724634534 | 4.90E-11 | 4.32E-10 |
| 12980 | ENSMUSG00000027349 | Fam98b | 68215 [Gene Symbol: Fam98b] [Locus Tag: ] [Chromosome: 2] [Map Location: 2\|2 E5] [I | 5.723308314 | 6.50E-17 | 1.48E-15 |
| 15957 | ENSMUSG00000038215 | Cep44 | 382010 [Gene Symbol: Cep44] [Locus Tag: ] [Chromosome: 8] [Map Location: 8\|8 B2] [I | 5.718750927 | 6.71E-12 | 6.79E-11 |
| 2954 | ENSMUSG00000062929 | Cfl2 | 12632 [Gene Symbol: Cfl2] [Locus Tag: ] [Chromosome: 12] [Map Location: 12\|12 C1] [I | 5.709132294 | 2.53E-20 | 9.82E-19 |
| 12602 | ENSMUSG00000036390 | Gadd45a | 13197 [Gene Symbol: Gadd45a] [Locus Tag: ] [Chromosome: 6] [Map Location: 6\|6 C1] | 5.708883326 | 5.23E-08 | 2.90E-07 |
| 9732 | ENSMUSG00000028821 | Syf2 | 68592 [Gene Symbol: Syf2] [Locus Tag: ] [Chromosome: 4] [Map Location: 4 D3\|4 67.15 | 5.707526971 | 5.26E-14 | 7.50E-13 |
| 3321 | ENSMUSG00000037984 | Neurod6 | 11922 [Gene Symbol: Neurod6] [Locus Tag: ] [Chromosome: 6] [Map Location: 6 B3\|6 2 | 5.702142972 | 7.78E-15 | 1.26E-13 |
| 14188 | ENSMUSG00000040268 | Plekha1 | 101476 [Gene Symbol: Plekha1] [Locus Tag: ] [Chromosome: 7] [Map Location: 7\|7 F3] | 5.693354662 | 4.90E-20 | 1.83E-18 |
| 697 | ENSMUSG00000068798 | Rap1a | 109905 [Gene Symbol: Rap1a] [Locus Tag: ] [Chromosome: 3] [Map Location: 3 F2.2\|3 | 5.692575546 | 3.46E-14 | 5.07E-13 |
| 12349 | ENSMUSG00000027882 | Stxbp3 | 20912 [Gene Symbol: Stxbp3] [Locus Tag: ] [Chromosome: 3] [Map Location: 3\|3 F3] [D | 5.692111327 | 9.51E-10 | 6.87E-09 |
| 11069 | ENSMUSG00000019877 | Serinc1 | 56442 [Gene Symbol: Serinc1] [Locus Tag: ] [Chromosome: 10] [Map Location: 10\|10 B | 5.69205468 | 2.55E-17 | 6.09E-16 |
| 7578 | ENSMUSG00000053164 | Gpr21 | 338346 [Gene Symbol: Gpr21] [Locus Tag: ] [Chromosome: 2] [Map Location: 2\|2 B] [De | 5.688258907 | 7.76E-15 | 1.26E-13 |
| 16851 | ENSMUSG00000048355 | Arxes1 | 76219 [Gene Symbol: Arxes1] [Locus Tag: ] [Chromosome: X] [Map Location: X\|X F1] [De | 5.681186376 | 1.96E-08 | 1.17E-07 |
| 3374 | ENSMUSG00000064351 | COX1 | 17708 [Gene Symbol: COX1] [Locus Tag: ] [Chromosome: MT] [Map Location: ] [Descript | 5.679048657 | 2.73E-20 | 1.06E-18 |
| 1391 | ENSMUSG00000023961 | Enpp4 | 224794 [Gene Symbol: Enpp4] [Locus Tag: ] [Chromosome: 17] [Map Location: 17\|17 B | 5.677555253 | 4.81E-14 | 6.89E-13 |
| 4417 | ENSMUSG00000029571 | Tmem106b | 71900 [Gene Symbol: Tmem106b] [Locus Tag: ] [Chromosome: 6] [Map Location: 6\|6 A | 5.675064288 | 1.98E-19 | 6.67E-18 |
| 4703 | ENSMUSG00000030557 | Mef2a | 17258 [Gene Symbol: Mef2a] [Locus Tag: ] [Chromosome: 7] [Map Location: 7 C\|7 36.7 | 5.669025793 | 4.72E-16 | 9.29E-15 |
| 5450 | ENSMUSG00000024878 | Cbwd1 | 226043 [Gene Symbol: Cbwd1] [Locus Tag: ] [Chromosome: 19] [Map Location: 19\|19 I | 5.668200289 | 4.87E-13 | 5.88E-12 |
| 8802 | ENSMUSG00000068617 | Efcab1 | 66793 [Gene Symbol: Efcab1] [Locus Tag: ] [Chromosome: 16] [Map Location: 16\|16 B1 | 5.661941063 | 1.75E-08 | 1.05E-07 |
| 16745 | ENSMUSG00000019868 | Vta1 | 66201 [Gene Symbol: Vta1] [Locus Tag: ] [Chromosome: 10] [Map Location: 10\|10 A2] [ | 5.652368215 | 2.68E-17 | 6.39E-16 |
| 12833 | ENSMUSG00000032116 | Stt3a | 16430 [Gene Symbol: Stt3a] [Locus Tag: ] [Chromosome: 9] [Map Location: 9 A4\|9 20.6 | 5.650402212 | 1.18E-17 | 2.93E-16 |
| 848 | ENSMUSG00000019857 | Asf1a | 66403 [Gene Symbol: Asf1a] [Locus Tag: ] [Chromosome: 10] [Map Location: 10\|10 B3] | 5.648069915 | 1.70E-15 | 3.03E-14 |
| 16600 | ENSMUSG00000039782 | Cpeb2 | 231207 [Gene Symbol: Cpeb2] [Locus Tag: ] [Chromosome: 5] [Map Location: 5\|5 B] [De | 5.647826984 | 1.05E-13 | 1.42E-12 |
| 14369 | ENSMUSG00000027770 | Dhx36 | 72162 [Gene Symbol: Dhx36] [Locus Tag: ] [Chromosome: 3] [Map Location: 3\|3 E1] [De | 5.64686157 | 4.82E-19 | 1.51E-17 |
| 8532 | ENSMUSG00000015341 | Golga7 | 57437 [Gene Symbol: Golga7] [Locus Tag: ] [Chromosome: 8] [Map Location: 8\|8 A2] [D | 5.641990838 | 3.12E-16 | 6.36E-15 |
| 4605 | ENSMUSG00000025816 | Sec61a2 | 57743 [Gene Symbol: Sec61a2] [Locus Tag: ] [Chromosome: 2] [Map Location: 2\|2 A1] [ | 5.635736606 | 1.55E-12 | 1.72E-11 |
| 4440 | ENSMUSG00000029836 | Cbx3 | 12417 [Gene Symbol: Cbx3] [Locus Tag: ] [Chromosome: 6] [Map Location: 6 24.89 cM\| | 5.632859256 | 1.57E-12 | 1.74E-11 |
| 11255 | ENSMUSG00000027834 | Serpini1 | 20713 [Gene Symbol: Serpini1] [Locus Tag: ] [Chromosome: 3] [Map Location: 3\|3 E3] [ | 5.632737912 | 3.52E-18 | 9.63E-17 |
| 8664 | ENSMUSG00000017421 | Zfp207 | 22680 [Gene Symbol: Zfp207] [Locus Tag: ] [Chromosome: 11] [Map Location: 11\|11 B5 | 5.632161465 | 3.96E-14 | 5.76E-13 |
| 5754 | ENSMUSG00000000340 | Dbt | 13171 [Gene Symbol: Dbt] [Locus Tag: ] [Chromosome: 3] [Map Location: 3 G1\|3 50.37 | 5.625661079 | 8.07E-14 | 1.11E-12 |
| 14263 | ENSMUSG00000069631 | Strada | 72149 [Gene Symbol: Strada] [Locus Tag: ] [Chromosome: 11] [Map Location: 11\|11 E1 | 5.624397775 | 9.22E-19 | 2.75E-17 |
| 17021 | ENSMUSG00000034467 | Dynlrb2 | 75465 [Gene Symbol: Dynlrb2] [Locus Tag: ] [Chromosome: 8] [Map Location: 8\|8 E1] [I | 5.620761761 | 0.000513 | 0.001413 |
| 8049 | ENSMUSG00000028367 | Txn1 | 22166 [Gene Symbol: Txn1] [Locus Tag: ] [Chromosome: 4] [Map Location: 4 B3\|4 31.8 | 5.615755914 | 6.77E-16 | 1.31E-14 |
| 8568 | ENSMUSG00000005687 | Bcas2 | 68183 [Gene Symbol: Bcas2] [Locus Tag: ] [Chromosome: 3] [Map Location: 3\|3 F2.2] [D | 5.61199932 | 5.46E-12 | 5.59E-11 |
| 6385 | ENSMUSG00000002679 | Med6 | 69792 [Gene Symbol: Med6] [Locus Tag: ] [Chromosome: 12] [Map Location: 12\|12 D3 | 5.589283458 | 1.45E-12 | 1.62E-11 |
| 9783 | ENSMUSG00000025326 | Ube3a | 22215 [Gene Symbol: Ube3a] [Locus Tag: ] [Chromosome: 7] [Map Location: 7 C\|7 33.9 | 5.585754598 | 1.68E-20 | 6.73E-19 |
| 16061 | ENSMUSG00000020078 | Vps26a | 30930 [Gene Symbol: Vps26a] [Locus Tag: ] [Chromosome: 10] [Map Location: 10\|10 B | 5.585009194 | 4.08E-18 | 1.10E-16 |
| 2959 | ENSMUSG00000022020 | Naa16 | 66897 [Gene Symbol: Naa16] [Locus Tag: ] [Chromosome: 14] [Map Location: 14\|14 D3 | 5.581415374 | 3.67E-14 | 5.36E-13 |
| 7990 | ENSMUSG00000073016 | Uprt | 331487 [Gene Symbol: Uprt] [Locus Tag: ] [Chromosome: X] [Map Location: X\|X D] [Des | 5.57235808 | 6.12E-15 | 1.01E-13 |
| 8583 | ENSMUSG00000095464 | Cox17 | 12856 [Gene Symbol: Cox17] [Locus Tag: ] [Chromosome: 16] [Map Location: 16\|16 B3 | 5.56931342 | 1.59E-07 | 8.17E-07 |
| 2158 | ENSMUSG00000030188 | Magohb | 66441 [Gene Symbol: Magohb] [Locus Tag: ] [Chromosome: 6] [Map Location: 6\|6 C7] [ | 5.568831053 | 1.51E-09 | 1.06E-08 |
| 12649 | ENSMUSG00000047213 | Ythdf3 | 229096 [Gene Symbol: Ythdf3] [Locus Tag: ] [Chromosome: 3] [Map Location: 3\|3 A1] [ | 5.568435763 | 7.59E-20 | 2.75E-18 |
| 1378 | ENSMUSG00000020180 | Snrpd3 | 67332 [Gene Symbol: Snrpd3] [Locus Tag: ] [Chromosome: 10] [Map Location: 10\|10 C | 5.558376242 | 3.33E-18 | 9.14E-17 |
| 6233 | ENSMUSG00000023945 | Slc5a7 | 63993 [Gene Symbol: Slc5a7] [Locus Tag: ] [Chromosome: 17] [Map Location: 17\|17 D] | 5.555883736 | 1.45E-05 | 5.38E-05 |
| 6152 | ENSMUSG00000027012 | Dync1i2 | 13427 [Gene Symbol: Dync1i2] [Locus Tag: ] [Chromosome: 2] [Map Location: 2 C2\|2 4 | 5.545461753 | 4.18E-15 | 6.99E-14 |
| 4801 | ENSMUSG00000027763 | Mbnl1 | 56758 [Gene Symbol: Mbnl1] [Locus Tag: ] [Chromosome: 3] [Map Location: 3\|3 E1] [De | 5.544661675 | 1.30E-12 | 1.46E-11 |
| 11156 | ENSMUSG00000019920 | Lims1 | 110829 [Gene Symbol: Lims1] [Locus Tag: ] [Chromosome: 10] [Map Location: 10 B4\|1 | 5.543426009 | 1.82E-18 | 5.24E-17 |
| 14077 | ENSMUSG00000028864 | Hgf | 15234 [Gene Symbol: Hgf] [Locus Tag: ] [Chromosome: 5] [Map Location: 5 A2-A3\|5 7.0 | 5.538466395 | 2.48E-07 | 1.23E-06 |
| 2157 | ENSMUSG00000041777 | Cir1 | 66935 [Gene Symbol: Cir1] [Locus Tag: ] [Chromosome: 2] [Map Location: 2\|2 C3] [Desc | 5.538282377 | 1.76E-12 | 1.94E-11 |
| 2647 | ENSMUSG00000028221 | Tmem55a | 72519 [Gene Symbol: Tmem55a] [Locus Tag: ] [Chromosome: 4] [Map Location: 4\|4 A1- | 5.534343259 | 1.01E-18 | 2.99E-17 |
| 13173 | ENSMUSG00000035383 | Pmch | 110312 [Gene Symbol: Pmch] [Locus Tag: ] [Chromosome: 10] [Map Location: 10 43.7 | 5.527435193 | 0.007421 | 0.01592 |
| 9345 | ENSMUSG00000039735 | Fnbp1l | 214459 [Gene Symbol: Fnbp1l] [Locus Tag: ] [Chromosome: 3] [Map Location: 3\|3 G1] [ | 5.523665571 | 1.29E-19 | 4.50E-18 |
| 3978 | ENSMUSG00000068882 | Ssb | 20823 [Gene Symbol: Ssb] [Locus Tag: ] [Chromosome: 2] [Map Location: 2 C2\|2 40.95 | 5.522114792 | 2.46E-18 | 6.92E-17 |
| 17313 | ENSMUSG00000093536 | Gm16532 | 100042450 [Gene Symbol: Gm16532] [Locus Tag: ] [Chromosome: 7] [Map Location: 7 | 5.509602577 | 4.04E-10 | 3.08E-09 |
| 8089 | ENSMUSG00000027180 | Fbxo3 | 57443 [Gene Symbol: Fbxo3] [Locus Tag: ] [Chromosome: 2] [Map Location: 2\|2 E2] [De | 5.508328379 | 2.82E-18 | 7.85E-17 |
| 9188 | ENSMUSG00000031688 | Pou4f2 | 18997 [Gene Symbol: Pou4f2] [Locus Tag: ] [Chromosome: 8] [Map Location: 8\|8] [Desc | 5.507714347 | 3.24E-09 | 2.17E-08 |
| 2409 | ENSMUSG00000030031 | Kbtbd8 | 243574 [Gene Symbol: Kbtbd8] [Locus Tag: ] [Chromosome: 6] [Map Location: 6\|6 D2] | 5.507648291 | 3.09E-16 | 6.30E-15 |
| 3482 | ENSMUSG00000043969 | Emx2 | 13797 [Gene Symbol: Emx2] [Locus Tag: ] [Chromosome: 19] [Map Location: 19 D3\|19 | 5.499042386 | 9.88E-10 | 7.11E-09 |
| 16988 | ENSMUSG00000073664 | Nbeal1 | 269198 [Gene Symbol: Nbeal1] [Locus Tag: ] [Chromosome: 1] [Map Location: 1\|1 C2] [ | 5.498507452 | 1.30E-19 | 4.55E-18 |
| 10084 | ENSMUSG00000063015 | Ccni | 12453 [Gene Symbol: Ccni] [Locus Tag: ] [Chromosome: 5] [Map Location: 5\|5 E3.3-F1. | 5.498181635 | 6.14E-18 | 1.60E-16 |
| 189 | ENSMUSG00000067377 | Tspan6 | 56496 [Gene Symbol: Tspan6] [Locus Tag: ] [Chromosome: X] [Map Location: X\|X E3] [D | 5.497618341 | 1.15E-12 | 1.31E-11 |
| 11739 | ENSMUSG00000028179 | Cth | 107869 [Gene Symbol: Cth] [Locus Tag: ] [Chromosome: 3] [Map Location: 3\|3 H4] [Des | 5.496472683 | 0.003192 | 0.007436 |
| 4162 | ENSMUSG00000036934 | 4921524J17Rik | 66714 [Gene Symbol: 4921524J17Rik] [Locus Tag: ] [Chromosome: 8] [Map Location: 8 | 5.495752988 | 9.99E-15 | 1.58E-13 |

| | | | | | | |
|---|---|---|---|---|---|---|
| 4136 | ENSMUSG00000031568 | Rwdd4a | 192174 [Gene Symbol: Rwdd4a] [Locus Tag: ] [Chromosome: 8] [Map Location: 8\|8 B1. | 5.494784132 | 2.16E-13 | 2.75E-12 |
| 4893 | ENSMUSG00000038836 | Agbl3 | 76223 [Gene Symbol: Agbl3] [Locus Tag: ] [Chromosome: 6] [Map Location: 6\|6 B1] [De | 5.492892391 | 1.61E-11 | 1.53E-10 |
| 15540 | ENSMUSG00000023025 | Larp4 | 207214 [Gene Symbol: Larp4] [Locus Tag: ] [Chromosome: 15] [Map Location: 15\|15 F1 | 5.489175567 | 9.44E-19 | 2.81E-17 |
| 5855 | ENSMUSG00000022800 | Fyttd1 | 69823 [Gene Symbol: Fyttd1] [Locus Tag: ] [Chromosome: 16] [Map Location: 16\|16 B3 | 5.484026267 | 4.73E-18 | 1.27E-16 |
| 9804 | ENSMUSG00000062169 | Cnih4 | 98417 [Gene Symbol: Cnih4] [Locus Tag: ] [Chromosome: 1] [Map Location: 1\|1 H4] [De | 5.472045617 | 6.14E-16 | 1.19E-14 |
| 7542 | ENSMUSG00000031007 | Atp6ap2 | 70495 [Gene Symbol: Atp6ap2] [Locus Tag: ] [Chromosome: X] [Map Location: X\|X A1.1 | 5.458066319 | 1.35E-16 | 2.93E-15 |
| 3718 | ENSMUSG00000040204 | 2810417H13Rik | 68026 [Gene Symbol: 2810417H13Rik] [Locus Tag: ] [Chromosome: 9] [Map Location: 9 | 5.455558579 | 4.03E-07 | 1.94E-06 |
| 8866 | ENSMUSG00000051705 | Senp8 | 71599 [Gene Symbol: Senp8] [Locus Tag: ] [Chromosome: 9] [Map Location: 9\|9 C] [Des | 5.453417181 | 1.88E-13 | 2.42E-12 |
| 18575 | ENSMUSG00000041658 | Rragb | 245670 [Gene Symbol: Rragb] [Locus Tag: ] [Chromosome: X] [Map Location: X\|X F3] [D | 5.447296679 | 2.37E-13 | 3.00E-12 |
| 7959 | ENSMUSG00000042498 | D330045A20Rik | 102871 [Gene Symbol: D330045A20Rik] [Locus Tag: ] [Chromosome: X] [Map Location: | 5.441485786 | 7.92E-06 | 3.06E-05 |
| 16236 | ENSMUSG00000036422 | Pcdh8 | 18530 [Gene Symbol: Pcdh8] [Locus Tag: ] [Chromosome: 14] [Map Location: 14 D3\|14 | 5.418795816 | 9.75E-10 | 7.02E-09 |
| 10557 | ENSMUSG00000020309 | Chac2 | 68044 [Gene Symbol: Chac2] [Locus Tag: ] [Chromosome: 11] [Map Location: 11\|11 A4 | 5.417818058 | 5.31E-07 | 2.51E-06 |
| 1171 | ENSMUSG00000012422 | Tmem167 | 66074 [Gene Symbol: Tmem167] [Locus Tag: ] [Chromosome: 13] [Map Location: 13\|13 | 5.417587756 | 1.74E-18 | 5.01E-17 |
| 12795 | ENSMUSG00000044528 | Tram1l1 | 229801 [Gene Symbol: Tram1l1] [Locus Tag: ] [Chromosome: 3] [Map Location: 3\|3 G1 | 5.416659593 | 2.04E-16 | 4.31E-15 |
| 4792 | ENSMUSG00000010290 | AI597479 | 98404 [Gene Symbol: AI597479] [Locus Tag: ] [Chromosome: 1] [Map Location: 1\|1 B] [ | 5.416083401 | 1.77E-16 | 3.77E-15 |
| 9454 | ENSMUSG00000023089 | Ndufa5 | 68202 [Gene Symbol: Ndufa5] [Locus Tag: ] [Chromosome: 6] [Map Location: 6\|6 A3] [D | 5.414596324 | 3.40E-14 | 4.99E-13 |
| 7181 | ENSMUSG00000027341 | Tmem230 | 70612 [Gene Symbol: Tmem230] [Locus Tag: ] [Chromosome: 2] [Map Location: 2\|2 F2 | 5.413445487 | 1.64E-10 | 1.33E-09 |
| 12700 | ENSMUSG00000072889 | Nfxl1 | 100978 [Gene Symbol: Nfxl1] [Locus Tag: ] [Chromosome: 5] [Map Location: 5\|5 D] [De | 5.411827094 | 4.35E-15 | 7.24E-14 |
| 12851 | ENSMUSG00000027881 | Prpf38b | 66921 [Gene Symbol: Prpf38b] [Locus Tag: ] [Chromosome: 3] [Map Location: 3\|3 G1] [ | 5.408794989 | 8.26E-16 | 1.57E-14 |
| 8280 | ENSMUSG00000051224 | Tceanc | 245695 [Gene Symbol: Tceanc] [Locus Tag: ] [Chromosome: X] [Map Location: X\|X F5] [ | 5.405888378 | 9.01E-07 | 4.09E-06 |
| 17291 | ENSMUSG00000015189 | Casd1 | 213819 [Gene Symbol: Casd1] [Locus Tag: ] [Chromosome: 6] [Map Location: 6 A1\|6 1.8 | 5.401268983 | 2.71E-18 | 7.56E-17 |
| 16417 | ENSMUSG00000020794 | Ube2g1 | 67128 [Gene Symbol: Ube2g1] [Locus Tag: ] [Chromosome: 11] [Map Location: 11\|11 B | 5.399708995 | 3.87E-19 | 1.25E-17 |
| 12191 | ENSMUSG00000049357 | 4933408B17Rik | 271508 [Gene Symbol: 4933408B17Rik] [Locus Tag: ] [Chromosome: 18] [Map Location | 5.395076031 | 0.000335 | 0.000956 |
| 6517 | ENSMUSG00000025912 | Mybl1 | 17864 [Gene Symbol: Mybl1] [Locus Tag: ] [Chromosome: 1] [Map Location: 1 A2\|1 2.0 | 5.392387434 | 8.48E-12 | 8.42E-11 |
| 1304 | ENSMUSG00000039166 | Akap7 | 432442 [Gene Symbol: Akap7] [Locus Tag: ] [Chromosome: 10] [Map Location: 10\|10 A | 5.389774499 | 3.35E-05 | 0.000116 |
| 13052 | ENSMUSG00000000037 | Scml2 | 107815 [Gene Symbol: Scml2] [Locus Tag: ] [Chromosome: X] [Map Location: X\|X F4] [D | 5.386300411 | 5.13E-10 | 3.85E-09 |
| 1586 | ENSMUSG00000029229 | Chic2 | 74277 [Gene Symbol: Chic2] [Locus Tag: ] [Chromosome: 5] [Map Location: 5\|5 D] [Des | 5.385996577 | 2.88E-17 | 6.85E-16 |
| 7816 | ENSMUSG00000029177 | Cenpa | 12615 [Gene Symbol: Cenpa] [Locus Tag: ] [Chromosome: 5] [Map Location: 5 B1\|5 16. | 5.377495464 | 0.000214 | 0.000634 |
| 1252 | ENSMUSG00000044408 | Sptssa | 104725 [Gene Symbol: Sptssa] [Locus Tag: ] [Chromosome: 12] [Map Location: 12\|12 C | 5.376730196 | 8.76E-15 | 1.41E-13 |
| 1291 | ENSMUSG00000042032 | Mat2b | 108645 [Gene Symbol: Mat2b] [Locus Tag: ] [Chromosome: 11] [Map Location: 11\|11 A | 5.376013817 | 1.02E-15 | 1.90E-14 |
| 10426 | ENSMUSG00000027967 | Neurog2 | 11924 [Gene Symbol: Neurog2] [Locus Tag: ] [Chromosome: 3] [Map Location: 3\|3 H1] | 5.374023575 | 0.0187 | 0.03632 |
| 13598 | ENSMUSG00000034317 | Trim59 | 66949 [Gene Symbol: Trim59] [Locus Tag: ] [Chromosome: 3] [Map Location: 3\|3 E2] [D | 5.371324796 | 4.81E-10 | 3.63E-09 |
| 301 | ENSMUSG00000017144 | Rnd3 | 74194 [Gene Symbol: Rnd3] [Locus Tag: ] [Chromosome: 2] [Map Location: 2\|2 C1.1] [D | 5.366704219 | 2.50E-13 | 3.16E-12 |
| 2771 | ENSMUSG00000020974 | Pole2 | 18974 [Gene Symbol: Pole2] [Locus Tag: ] [Chromosome: 12] [Map Location: 12\|12 C3] | 5.363520371 | 1.68E-07 | 8.58E-07 |
| 10315 | ENSMUSG00000027620 | Rbm39 | 170791 [Gene Symbol: Rbm39] [Locus Tag: ] [Chromosome: 2] [Map Location: 2\|2 H1] | 5.359370165 | 2.28E-16 | 4.79E-15 |
| 17588 | ENSMUSG00000033918 | Parl | 381038 [Gene Symbol: Parl] [Locus Tag: ] [Chromosome: 16] [Map Location: 16 A3\|16 | 5.356645463 | 3.58E-12 | 3.75E-11 |
| 10485 | ENSMUSG00000027104 | LOC102641666; Atf2 | 102641666 [Gene Symbol: LOC102641666] [Locus Tag: ] [Chromosome: 4] [Map Locati | 5.354839911 | 1.17E-19 | 4.15E-18 |
| 11401 | ENSMUSG00000039530 | Tusc3 | 80286 [Gene Symbol: Tusc3] [Locus Tag: ] [Chromosome: 8] [Map Location: 8\|8 B1.2] [D | 5.350450658 | 1.03E-15 | 1.93E-14 |
| 13313 | ENSMUSG00000006373 | Pgrmc1 | 53328 [Gene Symbol: Pgrmc1] [Locus Tag: ] [Chromosome: X] [Map Location: X\|X A3.3] | 5.348843285 | 5.15E-18 | 1.37E-16 |
| 14275 | ENSMUSG00000073295 | Nudt10; Nudt11 | 102954 [Gene Symbol: Nudt10] [Locus Tag: ] [Chromosome: X] [Map Location: X\|X A1. | 5.348180632 | 1.29E-16 | 2.82E-15 |
| 5031 | ENSMUSG00000060904 | Arl1 | 104303 [Gene Symbol: Arl1] [Locus Tag: ] [Chromosome: 10] [Map Location: 10\|10 C1] | 5.340509496 | 3.21E-15 | 5.47E-14 |
| 709 | ENSMUSG00000075703 | Ept1 | 28042 [Gene Symbol: Ept1] [Locus Tag: ] [Chromosome: 5] [Map Location: 5 B1\|5 16.2 | 5.338419955 | 3.99E-18 | 1.09E-16 |
| 13389 | ENSMUSG00000091155 | Serpine3 | 319433 [Gene Symbol: Serpine3] [Locus Tag: ] [Chromosome: 14] [Map Location: 14\|14 | 5.336171643 | 3.89E-11 | 3.48E-10 |
| 7992 | ENSMUSG00000059482 | 2610301B20Rik | 67157 [Gene Symbol: 2610301B20Rik] [Locus Tag: ] [Chromosome: 4] [Map Location: 4 | 5.325451669 | 1.76E-13 | 2.28E-12 |
| 12698 | ENSMUSG00000071252 | 2210408I21Rik | 72371 [Gene Symbol: 2210408I21Rik] [Locus Tag: ] [Chromosome: 13] [Map Location: | 5.322440147 | 8.20E-19 | 2.46E-17 |
| 9861 | ENSMUSG00000022159 | Rab2b | 76338 [Gene Symbol: Rab2b] [Locus Tag: ] [Chromosome: 14] [Map Location: 14\|14 C2 | 5.317525327 | 1.48E-15 | 2.69E-14 |
| 12774 | ENSMUSG00000029500 | Pgam5 | 72542 [Gene Symbol: Pgam5] [Locus Tag: ] [Chromosome: 5] [Map Location: 5\|5 F] [Des | 5.315416328 | 7.84E-18 | 2.01E-16 |
| 15053 | ENSMUSG00000006412 | Pfdn2 | 18637 [Gene Symbol: Pfdn2] [Locus Tag: ] [Chromosome: 1] [Map Location: 1 H3\|1 79. | 5.310770697 | 2.86E-15 | 4.92E-14 |
| 2078 | ENSMUSG00000056679 | Gpr173 | 70771 [Gene Symbol: Gpr173] [Locus Tag: ] [Chromosome: X] [Map Location: X\|X F2] [D | 5.309030107 | 3.01E-16 | 6.16E-15 |
| 1681 | ENSMUSG00000032375 | Aph1b | 208117 [Gene Symbol: Aph1b] [Locus Tag: ] [Chromosome: 9] [Map Location: 9\|9 D] [D | 5.30759374 | 1.35E-12 | 1.51E-11 |
| 18320 | ENSMUSG00000034653 | Ythdc2 | 240255 [Gene Symbol: Ythdc2] [Locus Tag: ] [Chromosome: 18] [Map Location: 18\|18 | 5.306623344 | 5.66E-14 | 8.04E-13 |
| 6008 | ENSMUSG00000030342 | Cd9 | 12527 [Gene Symbol: Cd9] [Locus Tag: ] [Chromosome: 6] [Map Location: 6 F3\|6 59.32 | 5.295371846 | 1.44E-15 | 2.62E-14 |
| 9921 | ENSMUSG00000047897 | Ripply2 | 382089 [Gene Symbol: Ripply2] [Locus Tag: ] [Chromosome: 9] [Map Location: 9\|9 E3.1 | 5.290430379 | 6.72E-05 | 0.000218 |
| 6597 | ENSMUSG00000000078 | Klf6 | 23849 [Gene Symbol: Klf6] [Locus Tag: ] [Chromosome: 13] [Map Location: 13\|13 A1] [ | 5.280551834 | 1.24E-13 | 1.66E-12 |
| 7214 | ENSMUSG00000054115 | Skp2 | 27401 [Gene Symbol: Skp2] [Locus Tag: ] [Chromosome: 15] [Map Location: 15\|15 A2] | 5.280545899 | 1.76E-11 | 1.66E-10 |
| 6586 | ENSMUSG00000032661 | Oas3 | 246727 [Gene Symbol: Oas3] [Locus Tag: ] [Chromosome: 5] [Map Location: 5 F\|5 60.6 | 5.273397317 | 0.000219 | 0.000647 |
| 17883 | ENSMUSG00000027428 | Rbbp9 | 26450 [Gene Symbol: Rbbp9] [Locus Tag: ] [Chromosome: 2] [Map Location: 2\|2 G1-H1 | 5.273372971 | 2.20E-13 | 2.80E-12 |
| 16317 | ENSMUSG00000062328 | Rpl17; Gm10362 | 319195 [Gene Symbol: Rpl17] [Locus Tag: ] [Chromosome: 18] [Map Location: 18\|18 E | 5.27242596 | 0.000328 | 0.000939 |
| 10745 | ENSMUSG00000029463 | Fam216a | 68948 [Gene Symbol: Fam216a] [Locus Tag: ] [Chromosome: 5] [Map Location: 5\|5 F] [ | 5.265815617 | 1.51E-17 | 3.70E-16 |
| 17943 | ENSMUSG00000025451 | Paip1 | 218693 [Gene Symbol: Paip1] [Locus Tag: ] [Chromosome: 13] [Map Location: 13\|13 D | 5.265150752 | 2.50E-19 | 8.26E-18 |
| 963 | ENSMUSG00000027706 | Sec62 | 69276 [Gene Symbol: Sec62] [Locus Tag: ] [Chromosome: 3] [Map Location: 3\|3 A3] [De | 5.260015488 | 2.46E-18 | 6.92E-17 |
| 5043 | ENSMUSG00000033793 | Atp6v1h | 108664 [Gene Symbol: Atp6v1h] [Locus Tag: ] [Chromosome: 1] [Map Location: 1\|1 A1 | 5.2453193 | 8.01E-18 | 2.06E-16 |
| 15027 | ENSMUSG00000027778 | Ift80 | 68259 [Gene Symbol: Ift80] [Locus Tag: ] [Chromosome: 3] [Map Location: 3\|3 E2] [Des | 5.236434307 | 2.22E-14 | 3.35E-13 |
| 6592 | ENSMUSG00000043410 | Hfm1 | 330149 [Gene Symbol: Hfm1] [Locus Tag: ] [Chromosome: 5] [Map Location: 5\|5 E5] [D | 5.232724047 | 5.51E-10 | 4.12E-09 |
| 5027 | ENSMUSG00000031422 | Morf4l2 | 56397 [Gene Symbol: Morf4l2] [Locus Tag: ] [Chromosome: X] [Map Location: X\|X F1] [ | 5.22981822 | 6.03E-18 | 1.57E-16 |
| 4258 | ENSMUSG00000023068 | Nus1 | 52014 [Gene Symbol: Nus1] [Locus Tag: ] [Chromosome: 10] [Map Location: 10 B3\|10 2 | 5.229727156 | 3.86E-19 | 1.24E-17 |
| 6556 | ENSMUSG00000037625 | Cldn11 | 18417 [Gene Symbol: Cldn11] [Locus Tag: ] [Chromosome: 3] [Map Location: 3 A3\|3 15 | 5.222706059 | 4.85E-08 | 2.71E-07 |
| 13276 | ENSMUSG00000052525 | Spdya | 70891 [Gene Symbol: Spdya] [Locus Tag: ] [Chromosome: 17] [Map Location: 17\|17 E1. | 5.222166832 | 7.05E-08 | 3.83E-07 |
| 18061 | ENSMUSG00000027536 | Chmp4c | 66371 [Gene Symbol: Chmp4c] [Locus Tag: ] [Chromosome: 3] [Map Location: 3\|3 A1] [ | 5.219508925 | 9.31E-14 | 1.27E-12 |
| 9228 | ENSMUSG00000079419 | Ms4a6c | 73656 [Gene Symbol: Ms4a6c] [Locus Tag: ] [Chromosome: 19] [Map Location: 19\|19 A | 5.202048045 | 0.021294 | 0.040741 |
| 10458 | ENSMUSG00000017550 | Atad5 | 237877 [Gene Symbol: Atad5] [Locus Tag: ] [Chromosome: 11] [Map Location: 11\|11 B | 5.198802063 | 8.24E-12 | 8.20E-11 |
| 8471 | ENSMUSG00000063895 | Nupl1 | 71844 [Gene Symbol: Nupl1] [Locus Tag: ] [Chromosome: 14] [Map Location: 14\|14 D1 | 5.193524315 | 2.48E-18 | 6.97E-17 |
| 10018 | ENSMUSG00000026500 | Cox20 | 66359 [Gene Symbol: Cox20] [Locus Tag: ] [Chromosome: 1] [Map Location: 1\|1 H3] [D | 5.192853036 | 9.46E-11 | 7.94E-10 |
| 16589 | ENSMUSG00000027499 | Pkia | 18767 [Gene Symbol: Pkia] [Locus Tag: ] [Chromosome: 3] [Map Location: 3\|3 A1] [Desc | 5.191768847 | 4.52E-14 | 6.53E-13 |
| 8178 | ENSMUSG00000030069 | Prok2 | 50501 [Gene Symbol: Prok2] [Locus Tag: ] [Chromosome: 6] [Map Location: 6 D3\|6 46. | 5.191516847 | 0.00025 | 0.000731 |
| 13427 | ENSMUSG00000025630 | Hprt | 15452 [Gene Symbol: Hprt] [Locus Tag: ] [Chromosome: X] [Map Location: X 29.31 cM\| | 5.190219727 | 5.92E-14 | 8.36E-13 |
| 14032 | ENSMUSG00000020116 | Pno1 | 66249 [Gene Symbol: Pno1] [Locus Tag: ] [Chromosome: 11] [Map Location: 11\|11 A2] | 5.1881083 | 1.21E-10 | 1.00E-09 |
| 17367 | ENSMUSG00000009030 | Pdcl | 67466 [Gene Symbol: Pdcl] [Locus Tag: ] [Chromosome: 2] [Map Location: 2\|2 B] [Descr | 5.185955934 | 2.18E-13 | 2.78E-12 |
| 7007 | ENSMUSG00000079038 | D130040H23Rik | 211135 [Gene Symbol: D130040H23Rik] [Locus Tag: ] [Chromosome: 8] [Map Location | 5.179372613 | 2.76E-07 | 1.36E-06 |
| 14102 | ENSMUSG00000020650 | Bcap29 | 12033 [Gene Symbol: Bcap29] [Locus Tag: ] [Chromosome: 12] [Map Location: 12 A3\|1 | 5.178218122 | 2.11E-11 | 1.96E-10 |
| 4943 | ENSMUSG00000021519 | Mterf3 | 66410 [Gene Symbol: Mterf3] [Locus Tag: ] [Chromosome: 13] [Map Location: 13\|13 B | 5.17575195 | 8.17E-15 | 1.32E-13 |
| 6186 | ENSMUSG00000038235 | F11r | 16456 [Gene Symbol: F11r] [Locus Tag: ] [Chromosome: 1] [Map Location: 1 H2\|1 79.4 | 5.174240866 | 2.03E-07 | 1.02E-06 |
| 12502 | ENSMUSG00000037822 | Smim14 | 68552 [Gene Symbol: Smim14] [Locus Tag: ] [Chromosome: 5] [Map Location: 5\|5 C3.1 | 5.172741102 | 3.07E-18 | 8.46E-17 |
| 4533 | ENSMUSG00000032235 | Ice2 | 93697 [Gene Symbol: Ice2] [Locus Tag: ] [Chromosome: 9] [Map Location: 9\|9 C] [Descr | 5.168284101 | 7.00E-08 | 3.80E-07 |

| | | | | | | |
|---|---|---|---|---|---|---|
| 17292 | ENSMUSG00000026739 | Bmi1 | 12151 [Gene Symbol: Bmi1] [Locus Tag: ] [Chromosome: 2] [Map Location: 2 A3\|2 12.9 | 5.167692728 | 1.39E-12 | 1.55E-11 |
| 14797 | ENSMUSG00000021687 | Scamp1 | 107767 [Gene Symbol: Scamp1] [Locus Tag: ] [Chromosome: 13] [Map Location: 13\|13 | 5.167647884 | 3.05E-18 | 8.42E-17 |
| 4528 | ENSMUSG00000021936 | Mapk8 | 26419 [Gene Symbol: Mapk8] [Locus Tag: ] [Chromosome: 14] [Map Location: 14\|14 B] | 5.160756909 | 1.21E-18 | 3.54E-17 |
| 9697 | ENSMUSG00000069135 | Fgfr1op | 75296 [Gene Symbol: Fgfr1op] [Locus Tag: ] [Chromosome: 17] [Map Location: 17\|17 A | 5.160718727 | 1.89E-12 | 2.08E-11 |
| 7231 | ENSMUSG00000025925 | Terf1 | 21749 [Gene Symbol: Terf1] [Locus Tag: ] [Chromosome: 1] [Map Location: 1 4.88 cM\|1 | 5.160244262 | 2.20E-11 | 2.04E-10 |
| 8858 | ENSMUSG00000006010 | BC003331 | 226499 [Gene Symbol: BC003331] [Locus Tag: ] [Chromosome: 1] [Map Location: 1\|1 G | 5.1576933 | 2.15E-14 | 3.26E-13 |
| 4860 | ENSMUSG00000026655 | Fam107b | 66540 [Gene Symbol: Fam107b] [Locus Tag: ] [Chromosome: 2] [Map Location: 2\|2 A1] | 5.156724912 | 2.37E-12 | 2.55E-11 |
| 2129 | ENSMUSG00000025283 | Sat1 | 20229 [Gene Symbol: Sat1] [Locus Tag: ] [Chromosome: X] [Map Location: X 72.38 cM\| | 5.15633933 | 4.53E-05 | 0.000152 |
| 6760 | ENSMUSG00000060636 | Rpl35a | 57808 [Gene Symbol: Rpl35a] [Locus Tag: ] [Chromosome: 16] [Map Location: 16\|16 B | 5.152970126 | 6.92E-10 | 5.10E-09 |
| 16076 | ENSMUSG00000025903 | Lypla1 | 18777 [Gene Symbol: Lypla1] [Locus Tag: ] [Chromosome: 1] [Map Location: 1\|1 A1] [D | 5.151689288 | 1.00E-12 | 1.16E-11 |
| 10704 | ENSMUSG00000028476 | Reck | 53614 [Gene Symbol: Reck] [Locus Tag: ] [Chromosome: 4] [Map Location: 4\|4 B1] [Desc | 5.149383003 | 6.11E-14 | 8.62E-13 |
| 449 | ENSMUSG00000042541 | Shfm1 | 20422 [Gene Symbol: Shfm1] [Locus Tag: ] [Chromosome: 6] [Map Location: 6\|6 A2] [De | 5.149306917 | 1.25E-11 | 1.21E-10 |
| 2879 | ENSMUSG00000022299 | Slc25a32 | 69906 [Gene Symbol: Slc25a32] [Locus Tag: ] [Chromosome: 15] [Map Location: 15\|15 | 5.148276265 | 2.77E-12 | 2.96E-11 |
| 18592 | ENSMUSG00000002845 | Tmem39a | 67846 [Gene Symbol: Tmem39a] [Locus Tag: ] [Chromosome: 16] [Map Location: 16\|16 | 5.142027269 | 5.86E-13 | 6.98E-12 |
| 11369 | ENSMUSG00000006699 | Cdc42 | 12540 [Gene Symbol: Cdc42] [Locus Tag: ] [Chromosome: 4] [Map Location: 4 D3\|4 69. | 5.138438205 | 2.84E-18 | 7.91E-17 |
| 6299 | ENSMUSG00000040760 | Appl1 | 72993 [Gene Symbol: Appl1] [Locus Tag: ] [Chromosome: 14] [Map Location: 14\|14 B] [ | 5.134728551 | 5.22E-18 | 1.39E-16 |
| 16203 | ENSMUSG00000040929 | Rfx3 | 19726 [Gene Symbol: Rfx3] [Locus Tag: ] [Chromosome: 19] [Map Location: 19\|19 C1] [ | 5.134151767 | 7.22E-19 | 2.19E-17 |
| 6324 | ENSMUSG00000027115 | Kif18a | 228421 [Gene Symbol: Kif18a] [Locus Tag: ] [Chromosome: 2] [Map Location: 2\|2 E3] [D | 5.125738162 | 6.57E-10 | 4.85E-09 |
| 12253 | ENSMUSG00000036030 | Prtg | 235472 [Gene Symbol: Prtg] [Locus Tag: ] [Chromosome: 9] [Map Location: 9\|9 D] [Desc | 5.118407784 | 4.20E-10 | 3.19E-09 |
| 15239 | ENSMUSG00000005124 | Wisp1 | 22402 [Gene Symbol: Wisp1] [Locus Tag: ] [Chromosome: 15] [Map Location: 15 D2\|15 | 5.11757473 | 9.18E-06 | 3.51E-05 |
| 16562 | ENSMUSG00000044165 | Bcl2l15 | 229672 [Gene Symbol: Bcl2l15] [Locus Tag: ] [Chromosome: 3] [Map Location: 3\|3 F2.2 | 5.116771641 | 0.005472 | 0.012076 |
| 4919 | ENSMUSG00000094724 | Rnaset2b | 68195 [Gene Symbol: Rnaset2b] [Locus Tag: ] [Chromosome: 17] [Map Location: 17\|17 | 5.111498329 | 0.020413 | 0.039229 |
| 5310 | ENSMUSG00000025050 | Pcgf6 | 71041 [Gene Symbol: Pcgf6] [Locus Tag: ] [Chromosome: 19] [Map Location: 19\|19 C3] | 5.10428414 | 1.22E-11 | 1.18E-10 |
| 3332 | ENSMUSG00000028998 | Tomm7 | 66169 [Gene Symbol: Tomm7] [Locus Tag: ] [Chromosome: 5] [Map Location: 5\|5 A3] [D | 5.095131043 | 6.83E-11 | 5.88E-10 |
| 10338 | ENSMUSG00000030846 | Tial1 | 21843 [Gene Symbol: Tial1] [Locus Tag: ] [Chromosome: 7] [Map Location: 7\|7 F4] [Desc | 5.079661223 | 1.08E-12 | 1.24E-11 |
| 7125 | ENSMUSG00000027787 | Nmd3 | 97112 [Gene Symbol: Nmd3] [Locus Tag: ] [Chromosome: 3] [Map Location: 3\|3 E1] [De | 5.076323525 | 1.56E-15 | 2.81E-14 |
| 17123 | ENSMUSG00000012429 | Mplkip; Gm7102 | 66308 [Gene Symbol: Mplkip] [Locus Tag: ] [Chromosome: 13] [Map Location: 13\|13 A | 5.071624135 | 1.26E-10 | 1.04E-09 |
| 13233 | ENSMUSG00000027589 | Pcmtd2 | 245867 [Gene Symbol: Pcmtd2] [Locus Tag: ] [Chromosome: 2] [Map Location: 2\|2 H4] | 5.068099608 | 1.38E-12 | 1.55E-11 |
| 16369 | ENSMUSG00000036503 | Rnf13 | 24017 [Gene Symbol: Rnf13] [Locus Tag: ] [Chromosome: 3] [Map Location: 3\|3 D] [Des | 5.066128602 | 4.18E-16 | 8.34E-15 |
| 18023 | ENSMUSG00000031016 | Wee1 | 22390 [Gene Symbol: Wee1] [Locus Tag: ] [Chromosome: 7] [Map Location: 7\|7 F1] [Des | 5.065694209 | 2.14E-06 | 9.10E-06 |
| 5040 | ENSMUSG00000041483 | Zfp281 | 226442 [Gene Symbol: Zfp281] [Locus Tag: ] [Chromosome: 1] [Map Location: 1\|1 E4] [ | 5.064987208 | 4.01E-18 | 1.09E-16 |
| 6461 | ENSMUSG00000028842 | Ago3 | 214150 [Gene Symbol: Ago3] [Locus Tag: ] [Chromosome: 4] [Map Location: 4\|4 D2.2] [ | 5.062660051 | 1.61E-15 | 2.89E-14 |
| 7616 | ENSMUSG00000038206 | Fbxo8 | 50753 [Gene Symbol: Fbxo8] [Locus Tag: ] [Chromosome: 8] [Map Location: 8\|8 B3.2] [ | 5.061809145 | 4.71E-15 | 7.82E-14 |
| 9821 | ENSMUSG00000061353 | Cxcl12 | 20315 [Gene Symbol: Cxcl12] [Locus Tag: ] [Chromosome: 6] [Map Location: 6 54.81 cM | 5.060684481 | 2.90E-15 | 4.98E-14 |
| 1417 | ENSMUSG00000020907 | Rcvrn | 19674 [Gene Symbol: Rcvrn] [Locus Tag: ] [Chromosome: 11] [Map Location: 11 B3\|11 | 5.053503858 | 0.021101 | 0.040421 |
| 13879 | ENSMUSG00000024172 | St6gal2 | 240119 [Gene Symbol: St6gal2] [Locus Tag: ] [Chromosome: 17] [Map Location: 17\|17 | 5.048339195 | 1.48E-18 | 4.30E-17 |
| 6027 | ENSMUSG00000028553 | Angptl3 | 30924 [Gene Symbol: Angptl3] [Locus Tag: ] [Chromosome: 4] [Map Location: 4 C6\|4 4 | 5.041830556 | 5.00E-07 | 2.37E-06 |
| 13555 | ENSMUSG00000036977 | Anapc10 | 68999 [Gene Symbol: Anapc10] [Locus Tag: ] [Chromosome: 8] [Map Location: 8 37.79 | 5.041812082 | 4.10E-13 | 4.99E-12 |
| 9974 | ENSMUSG00000040370 | Lyrm5 | 67636 [Gene Symbol: Lyrm5] [Locus Tag: ] [Chromosome: 6] [Map Location: 6 G3\|6 77. | 5.037381994 | 8.13E-10 | 5.95E-09 |
| 3821 | ENSMUSG00000038822 | Hace1 | 209462 [Gene Symbol: Hace1] [Locus Tag: ] [Chromosome: 10] [Map Location: 10\|10 B | 5.037024558 | 1.35E-13 | 1.80E-12 |
| 6714 | ENSMUSG00000036275 | 9530068E07Rik | 213673 [Gene Symbol: 9530068E07Rik] [Locus Tag: ] [Chromosome: 11] [Map Location | 5.035971712 | 4.89E-13 | 5.90E-12 |
| 2970 | ENSMUSG00000031613 | Hpgd | 15446 [Gene Symbol: Hpgd] [Locus Tag: ] [Chromosome: 8] [Map Location: 8\|8 B3.2] [D | 5.03550875 | 1.82E-06 | 7.87E-06 |
| 9776 | ENSMUSG00000029263 | Pigg | 433931 [Gene Symbol: Pigg] [Locus Tag: ] [Chromosome: 5] [Map Location: 5\|5 F] [Desc | 5.03235462 | 1.72E-12 | 1.90E-11 |
| 12166 | ENSMUSG00000039652 | Cpeb3 | 208922 [Gene Symbol: Cpeb3] [Locus Tag: ] [Chromosome: 19] [Map Location: 19\|19 C | 5.030443924 | 9.26E-18 | 2.35E-16 |
| 13441 | ENSMUSG00000074884 | Serf2 | 378702 [Gene Symbol: Serf2] [Locus Tag: ] [Chromosome: 2] [Map Location: 2\|2 E5] [De | 5.030231122 | 8.74E-12 | 8.66E-11 |
| 521 | ENSMUSG00000032349 | Elovl5 | 68801 [Gene Symbol: Elovl5] [Locus Tag: ] [Chromosome: 9] [Map Location: 9\|9 E1] [De | 5.025187003 | 2.78E-16 | 5.71E-15 |
| 12952 | ENSMUSG00000004771 | Rab11a | 53869 [Gene Symbol: Rab11a] [Locus Tag: ] [Chromosome: 9] [Map Location: 9\|9 C] [De | 5.024359149 | 5.70E-18 | 1.49E-16 |
| 3553 | ENSMUSG00000041891 | Lman1 | 70361 [Gene Symbol: Lman1] [Locus Tag: ] [Chromosome: 18] [Map Location: 18\|18 E1 | 5.019129465 | 1.24E-14 | 1.93E-13 |
| 12399 | ENSMUSG00000022525 | Hrasls | 27281 [Gene Symbol: Hrasls] [Locus Tag: ] [Chromosome: 16] [Map Location: 16\|16 B2 | 5.01842041 | 9.24E-07 | 4.19E-06 |
| 3081 | ENSMUSG00000028172 | Tacr3 | 21338 [Gene Symbol: Tacr3] [Locus Tag: ] [Chromosome: 3] [Map Location: 3\|3 H2] [De | 5.015971289 | 3.52E-06 | 1.44E-05 |
| 72 | ENSMUSG00000079658 | Tceb1; LOC10264281 | 67923 [Gene Symbol: Tceb1] [Locus Tag: ] [Chromosome: 1] [Map Location: 1\|1 A3] [De | 5.015426659 | 2.54E-14 | 3.81E-13 |
| 2485 | ENSMUSG00000021774 | Ube2e1 | 22194 [Gene Symbol: Ube2e1] [Locus Tag: ] [Chromosome: 14] [Map Location: 14\|14 A | 5.011034419 | 1.34E-15 | 2.45E-14 |
| 7054 | ENSMUSG00000028016 | Ints12 | 71793 [Gene Symbol: Ints12] [Locus Tag: ] [Chromosome: 3] [Map Location: 3\|3 H2] [D | 5.010151678 | 6.84E-12 | 6.91E-11 |
| 8860 | ENSMUSG00000015619 | Gata3 | 14462 [Gene Symbol: Gata3] [Locus Tag: ] [Chromosome: 2] [Map Location: 2 A1\|2 6.6 | 5.009297101 | 1.02E-10 | 8.52E-10 |
| 6765 | ENSMUSG00000047844 | Bex4 | 406217 [Gene Symbol: Bex4] [Locus Tag: ] [Chromosome: X] [Map Location: XE3\|X] [De | 5.000224926 | 2.24E-11 | 2.07E-10 |
| 15282 | ENSMUSG00000055430 | Nap1l5 | 58243 [Gene Symbol: Nap1l5] [Locus Tag: ] [Chromosome: 6] [Map Location: 6\|6 C1] [D | 4.997151143 | 7.36E-11 | 6.30E-10 |
| 15510 | ENSMUSG00000024293 | Esco1 | 77805 [Gene Symbol: Esco1] [Locus Tag: ] [Chromosome: 18] [Map Location: 18\|18 A2] | 4.991091318 | 3.30E-10 | 2.55E-09 |
| 15603 | ENSMUSG00000072501 | Phf20l1 | 239510 [Gene Symbol: Phf20l1] [Locus Tag: ] [Chromosome: 15] [Map Location: 15\|15 | 4.987890521 | 1.06E-16 | 2.35E-15 |
| 10934 | ENSMUSG00000062901 | Klhl24 | 75785 [Gene Symbol: Klhl24] [Locus Tag: ] [Chromosome: 16] [Map Location: 16\|16 B1 | 4.987539956 | 4.85E-13 | 5.86E-12 |
| 9771 | ENSMUSG00000032839 | Trpc1 | 22063 [Gene Symbol: Trpc1] [Locus Tag: ] [Chromosome: 9] [Map Location: 9 50.2 cM\| | 4.986298586 | 1.85E-12 | 2.03E-11 |
| 10671 | ENSMUSG00000048379 | Socs4 | 67296 [Gene Symbol: Socs4] [Locus Tag: ] [Chromosome: 14] [Map Location: 14\|14 C1] | 4.986297972 | 2.69E-11 | 2.46E-10 |
| 9491 | ENSMUSG00000022228 | Zscan26 | 432731 [Gene Symbol: Zscan26] [Locus Tag: ] [Chromosome: 13] [Map Location: 13\|13 | 4.976567416 | 2.39E-15 | 4.17E-14 |
| 8477 | ENSMUSG00000030654 | Arl6ip1 | 54208 [Gene Symbol: Arl6ip1] [Locus Tag: ] [Chromosome: 7] [Map Location: 7\|7 F3] [D | 4.971124432 | 1.97E-10 | 1.57E-09 |
| 4848 | ENSMUSG00000062949 | Atp11c | 320940 [Gene Symbol: Atp11c] [Locus Tag: ] [Chromosome: X] [Map Location: X\|X A5] [ | 4.970846702 | 2.77E-17 | 6.60E-16 |
| 2877 | ENSMUSG00000027429 | Sec23b | 27054 [Gene Symbol: Sec23b] [Locus Tag: ] [Chromosome: 2] [Map Location: 2\|2 H1] [D | 4.967272652 | 1.02E-15 | 1.91E-14 |
| 1295 | ENSMUSG00000039191 | Rbpj | 19664 [Gene Symbol: Rbpj] [Locus Tag: ] [Chromosome: 5] [Map Location: 5 29.37 cM\| | 4.966967112 | 1.49E-15 | 2.69E-14 |
| 12458 | ENSMUSG00000040693 | Slco4c1 | 227394 [Gene Symbol: Slco4c1] [Locus Tag: ] [Chromosome: 1] [Map Location: 1\|1 D] [ | 4.964116732 | 9.57E-07 | 4.32E-06 |
| 16980 | ENSMUSG00000058748 | Zfp958 | 233987 [Gene Symbol: Zfp958] [Locus Tag: ] [Chromosome: 8] [Map Location: 8\|8 A1.1 | 4.963405503 | 1.78E-09 | 1.23E-08 |
| 11031 | ENSMUSG00000036513 | Commd2 | 52245 [Gene Symbol: Commd2] [Locus Tag: ] [Chromosome: 3] [Map Location: 3 D\|3 2 | 4.960139271 | 9.96E-13 | 1.15E-11 |
| 14167 | ENSMUSG00000060715 | 1700019A02Rik | 69397 [Gene Symbol: 1700019A02Rik] [Locus Tag: ] [Chromosome: 1] [Map Location: 1 | 4.955566686 | 1.87E-05 | 6.79E-05 |
| 17875 | ENSMUSG00000025889 | Snca | 20617 [Gene Symbol: Snca] [Locus Tag: ] [Chromosome: 6] [Map Location: 6 B3\|6 29.1 | 4.955357608 | 3.32E-06 | 1.36E-05 |
| 17418 | ENSMUSG00000021090 | Lrrc9 | 78257 [Gene Symbol: Lrrc9] [Locus Tag: ] [Chromosome: 12] [Map Location: 12\|12 C3] | 4.951981237 | 3.38E-07 | 1.64E-06 |
| 17674 | ENSMUSG00000053477 | Tcf4 | 21413 [Gene Symbol: Tcf4] [Locus Tag: ] [Chromosome: 18] [Map Location: 18\|18 E2] [ | 4.947875426 | 2.27E-18 | 6.43E-17 |
| 12726 | ENSMUSG00000032407 | U2surp | 67958 [Gene Symbol: U2surp] [Locus Tag: ] [Chromosome: 9] [Map Location: 9\|9 E4] [D | 4.947501752 | 6.26E-15 | 1.03E-13 |
| 14656 | ENSMUSG00000005763 | Cd247 | 12503 [Gene Symbol: Cd247] [Locus Tag: ] [Chromosome: 1] [Map Location: 1 73.14 cM | 4.945238555 | 2.13E-14 | 3.23E-13 |
| 15782 | ENSMUSG00000027166 | Dnajc24 | 99349 [Gene Symbol: Dnajc24] [Locus Tag: ] [Chromosome: 2] [Map Location: 2\|2 E3] [ | 4.937982959 | 6.32E-10 | 4.69E-09 |
| 13426 | ENSMUSG00000048482 | Bdnf | 12064 [Gene Symbol: Bdnf] [Locus Tag: ] [Chromosome: 2] [Map Location: 2 E3\|2 56.63 | 4.937622174 | 3.05E-12 | 3.23E-11 |
| 13191 | ENSMUSG00000055436 | Srsf11 | 69207 [Gene Symbol: Srsf11] [Locus Tag: ] [Chromosome: 3] [Map Location: 3\|3 H4] [De | 4.929597007 | 8.86E-18 | 2.26E-16 |
| 636 | ENSMUSG00000034457 | Eda2r | 245527 [Gene Symbol: Eda2r] [Locus Tag: ] [Chromosome: X] [Map Location: X\|X C3] [D | 4.929384293 | 0.001217 | 0.003116 |
| 3753 | ENSMUSG00000002731 | Prkra | 23992 [Gene Symbol: Prkra] [Locus Tag: ] [Chromosome: 2] [Map Location: 2\|2 C3] [Des | 4.926231384 | 3.39E-14 | 4.98E-13 |
| 131 | ENSMUSG00000034160 | Ogt | 108155 [Gene Symbol: Ogt] [Locus Tag: ] [Chromosome: X] [Map Location: X\|X D] [Desc | 4.926131905 | 4.58E-17 | 1.06E-15 |
| 6599 | ENSMUSG00000041718 | Alg13 | 67574 [Gene Symbol: Alg13] [Locus Tag: ] [Chromosome: X] [Map Location: X\|X F2] [De | 4.915125298 | 1.71E-14 | 2.63E-13 |
| 640 | ENSMUSG00000021025 | Nfkbia | 18035 [Gene Symbol: Nfkbia] [Locus Tag: ] [Chromosome: 12] [Map Location: 12\|12 C1 | 4.914384729 | 3.84E-15 | 6.46E-14 |
| 15636 | ENSMUSG00000051285 | Pcmtd1 | 319263 [Gene Symbol: Pcmtd1] [Locus Tag: ] [Chromosome: 1] [Map Location: 1\|1 A1] | 4.912970337 | 7.28E-12 | 7.32E-11 |

| | | | | | | |
|---|---|---|---|---|---|---|
| 8874 | ENSMUSG00000089996 | Tmsb15b2 | 100034363 [Gene Symbol: Tmsb15b2] [Locus Tag: ] [Chromosome: X] [Map Location: X | 4.910422512 | 0.013903 | 0.027925 |
| 15994 | ENSMUSG00000026675 | Hsd17b7 | 15490 [Gene Symbol: Hsd17b7] [Locus Tag: ] [Chromosome: 1] [Map Location: 1\|1 H3] | 4.906457146 | 2.47E-13 | 3.11E-12 |
| 13774 | ENSMUSG00000030275 | Etnk1 | 75320 [Gene Symbol: Etnk1] [Locus Tag: ] [Chromosome: 6] [Map Location: 6 G3\|6 75.4 | 4.902181822 | 3.39E-18 | 9.29E-17 |
| 16159 | ENSMUSG00000026342 | Slc35f5 | 74150 [Gene Symbol: Slc35f5] [Locus Tag: ] [Chromosome: 1] [Map Location: 1\|1 E3] [D | 4.900033545 | 4.30E-14 | 6.24E-13 |
| 18226 | ENSMUSG00000026565 | Pou2f1 | 18986 [Gene Symbol: Pou2f1] [Locus Tag: ] [Chromosome: 1] [Map Location: 1 H2.3\|1 | 4.899491382 | 1.25E-17 | 3.09E-16 |
| 6496 | ENSMUSG00000018068 | Ints2 | 70422 [Gene Symbol: Ints2] [Locus Tag: ] [Chromosome: 11] [Map Location: 11\|11 C] [D | 4.89934097 | 5.76E-15 | 9.52E-14 |
| 5550 | ENSMUSG00000031601 | Cnot7 | 18983 [Gene Symbol: Cnot7] [Locus Tag: ] [Chromosome: 8] [Map Location: 8\|8 A4] [De | 4.896970595 | 1.47E-17 | 3.62E-16 |
| 6081 | ENSMUSG00000020003 | Pex7 | 18634 [Gene Symbol: Pex7] [Locus Tag: ] [Chromosome: 10] [Map Location: 10\|10 A3] | 4.894042015 | 1.32E-14 | 2.05E-13 |
| 11348 | ENSMUSG00000026600 | Soat1 | 20652 [Gene Symbol: Soat1] [Locus Tag: ] [Chromosome: 1] [Map Location: 1 G3\|1 67.7 | 4.891339723 | 1.27E-15 | 2.32E-14 |
| 2896 | ENSMUSG00000078970 | Wdr92 | 103784 [Gene Symbol: Wdr92] [Locus Tag: ] [Chromosome: 11] [Map Location: 11\|11 A | 4.890357243 | 6.73E-18 | 1.74E-16 |
| 116 | ENSMUSG00000040167 | Ikzf5 | 67143 [Gene Symbol: Ikzf5] [Locus Tag: ] [Chromosome: 7] [Map Location: 7\|7 F3] [Desc | 4.889994842 | 4.59E-13 | 5.57E-12 |
| 11995 | ENSMUSG00000039219 | Arid4b | 94246 [Gene Symbol: Arid4b] [Locus Tag: ] [Chromosome: 13] [Map Location: 13 A1\|13 | 4.882878216 | 1.67E-17 | 4.09E-16 |
| 4585 | ENSMUSG00000057858 | Fam204a | 76539 [Gene Symbol: Fam204a] [Locus Tag: ] [Chromosome: 19] [Map Location: 19 D3 | 4.875771151 | 4.37E-14 | 6.34E-13 |
| 13002 | ENSMUSG00000056018 | Ccdc7b | 75453 [Gene Symbol: Ccdc7b] [Locus Tag: ] [Chromosome: 8] [Map Location: 8\|8 E2] [D | 4.873874563 | 3.83E-09 | 2.54E-08 |
| 16733 | ENSMUSG00000042363 | Lgalsl | 216551 [Gene Symbol: Lgalsl] [Locus Tag: ] [Chromosome: 11] [Map Location: 11 A3.1\| | 4.871470675 | 4.65E-17 | 1.07E-15 |
| 16360 | ENSMUSG00000042595 | Fam199x | 245622 [Gene Symbol: Fam199x] [Locus Tag: ] [Chromosome: X] [Map Location: X\|X F1 | 4.868107026 | 4.34E-15 | 7.24E-14 |
| 13573 | ENSMUSG00000050926 | Dcaf12l2 | 245403 [Gene Symbol: Dcaf12l2] [Locus Tag: ] [Chromosome: X] [Map Location: X\|X A4 | 4.866998619 | 0.003145 | 0.007339 |
| 5487 | ENSMUSG00000028128 | F3 | 14066 [Gene Symbol: F3] [Locus Tag: ] [Chromosome: 3] [Map Location: 3 G1\|3 52.94 c | 4.864214136 | 3.60E-08 | 2.06E-07 |
| 13958 | ENSMUSG00000026238 | Ptma | 19231 [Gene Symbol: Ptma] [Locus Tag: ] [Chromosome: 1] [Map Location: 1\|1 D] [Desc | 4.863962827 | 1.52E-07 | 7.86E-07 |
| 2179 | ENSMUSG00000040359 | Ufl1 | 67490 [Gene Symbol: Ufl1] [Locus Tag: ] [Chromosome: 4] [Map Location: 4\|4 A3] [Desc | 4.862388363 | 7.02E-12 | 7.09E-11 |
| 4866 | ENSMUSG00000053253 | Ndfip2 | 76273 [Gene Symbol: Ndfip2] [Locus Tag: ] [Chromosome: 14] [Map Location: 14\|14 E2 | 4.859217573 | 1.67E-16 | 3.58E-15 |
| 15853 | ENSMUSG00000022801 | Lrch3 | 70144 [Gene Symbol: Lrch3] [Locus Tag: ] [Chromosome: 16] [Map Location: 16\|16 B3] | 4.858618094 | 2.22E-15 | 3.89E-14 |
| 10082 | ENSMUSG00000020460 | Rps27a; LOC100042C | 78294 [Gene Symbol: Rps27a] [Locus Tag: ] [Chromosome: 11] [Map Location: 11\|11 A | 4.854781661 | 0.003829 | 0.008756 |
| 12067 | ENSMUSG00000029684 | Wasl | 73178 [Gene Symbol: Wasl] [Locus Tag: ] [Chromosome: 6] [Map Location: 6\|6 A3] [Des | 4.853364612 | 2.08E-17 | 5.04E-16 |
| 14814 | ENSMUSG00000037608 | Bclaf1 | 72567 [Gene Symbol: Bclaf1] [Locus Tag: ] [Chromosome: 10] [Map Location: 10\|10 A3 | 4.850829151 | 2.39E-12 | 2.57E-11 |
| 3120 | ENSMUSG00000052144 | Ppp4r2 | 232314 [Gene Symbol: Ppp4r2] [Locus Tag: ] [Chromosome: 6] [Map Location: 6\|6 D3] | 4.840248572 | 5.80E-16 | 1.13E-14 |
| 9746 | ENSMUSG00000027381 | Bcl2l11 | 12125 [Gene Symbol: Bcl2l11] [Locus Tag: ] [Chromosome: 2] [Map Location: 2\|2 F3-G1 | 4.839556856 | 1.62E-15 | 2.91E-14 |
| 9563 | ENSMUSG00000020189 | Osbpl8 | 237542 [Gene Symbol: Osbpl8] [Locus Tag: ] [Chromosome: 10] [Map Location: 10\|10 I | 4.83650992 | 7.43E-18 | 1.91E-16 |
| 4936 | ENSMUSG00000028188 | Spata1 | 70951 [Gene Symbol: Spata1] [Locus Tag: ] [Chromosome: 3] [Map Location: 3\|3 H2] [D | 4.82492386 | 6.50E-10 | 4.80E-09 |
| 8243 | ENSMUSG00000075502 | Kbtbd6 | 432879 [Gene Symbol: Kbtbd6] [Locus Tag: ] [Chromosome: 14] [Map Location: 14\|14 | 4.824573378 | 0.000171 | 0.000515 |
| 1573 | ENSMUSG00000026393 | Nek7 | 59125 [Gene Symbol: Nek7] [Locus Tag: ] [Chromosome: 1] [Map Location: 1 E4\|1 60.8 | 4.823840756 | 9.69E-14 | 1.32E-12 |
| 5549 | ENSMUSG00000022601 | Zbtb11 | 271377 [Gene Symbol: Zbtb11] [Locus Tag: ] [Chromosome: 16] [Map Location: 16\|16 | 4.820827627 | 8.60E-12 | 8.53E-11 |
| 4721 | ENSMUSG00000023755 | Rhebl1 | 69159 [Gene Symbol: Rhebl1] [Locus Tag: ] [Chromosome: 15] [Map Location: 15\|15 F2 | 4.817027374 | 1.74E-08 | 1.05E-07 |
| 8598 | ENSMUSG00000020436 | Gabrg2 | 14406 [Gene Symbol: Gabrg2] [Locus Tag: ] [Chromosome: 11] [Map Location: 11 A5\|1 | 4.813778093 | 1.86E-16 | 3.95E-15 |
| 14187 | ENSMUSG00000022339 | Ebag9 | 55960 [Gene Symbol: Ebag9] [Locus Tag: ] [Chromosome: 15] [Map Location: 15\|15 D1 | 4.813203309 | 1.18E-13 | 1.59E-12 |
| 8535 | ENSMUSG00000026360 | Rgs2 | 19735 [Gene Symbol: Rgs2] [Locus Tag: ] [Chromosome: 1] [Map Location: 1 F\|1 62.56 | 4.806482658 | 2.70E-09 | 1.83E-08 |
| 13551 | ENSMUSG00000063108 | Zfp26 | 22688 [Gene Symbol: Zfp26] [Locus Tag: ] [Chromosome: 9] [Map Location: 9 A3\|9 7.55 | 4.805150481 | 4.04E-12 | 4.19E-11 |
| 16568 | ENSMUSG00000027361 | Gabpb1 | 14391 [Gene Symbol: Gabpb1] [Locus Tag: ] [Chromosome: 2] [Map Location: 2 F1\|2 61 | 4.79812263 | 1.45E-16 | 3.14E-15 |
| 7890 | ENSMUSG00000071866 | Ppia | 268373 [Gene Symbol: Ppia] [Locus Tag: ] [Chromosome: 11] [Map Location: 11 A1\|11 | 4.791577491 | 2.98E-05 | 0.000104 |
| 9204 | ENSMUSG00000061244 | Exoc5 | 105504 [Gene Symbol: Exoc5] [Locus Tag: ] [Chromosome: 14] [Map Location: 14\|14 C | 4.785902496 | 5.78E-17 | 1.32E-15 |
| 5239 | ENSMUSG00000054737 | Zfp182 | 319535 [Gene Symbol: Zfp182] [Locus Tag: ] [Chromosome: X] [Map Location: X\|X A1.3 | 4.785742241 | 2.02E-09 | 1.39E-08 |
| 13918 | ENSMUSG00000029290 | Zfp326 | 54367 [Gene Symbol: Zfp326] [Locus Tag: ] [Chromosome: 5] [Map Location: 5\|5 E5] [D | 4.781735619 | 6.25E-15 | 1.03E-13 |
| 3943 | ENSMUSG00000051098 | Mblac2 | 72852 [Gene Symbol: Mblac2] [Locus Tag: ] [Chromosome: 13] [Map Location: 13\|13 C | 4.775531195 | 1.23E-12 | 1.39E-11 |
| 14662 | ENSMUSG00000031284 | Pak3 | 18481 [Gene Symbol: Pak3] [Locus Tag: ] [Chromosome: X] [Map Location: X\|X F2] [Des | 4.772302898 | 6.15E-18 | 1.60E-16 |
| 16090 | ENSMUSG00000034610 | Zcchc11 | 230594 [Gene Symbol: Zcchc11] [Locus Tag: ] [Chromosome: 4] [Map Location: 4\|4 C7] | 4.770907939 | 8.98E-15 | 1.44E-13 |
| 2170 | ENSMUSG00000039100 | March6 | 223455 [Gene Symbol: March6] [Locus Tag: ] [Chromosome: 15] [Map Location: 15\|15 | 4.761546512 | 6.29E-17 | 1.44E-15 |
| 13833 | ENSMUSG00000054679 | Srsf12 | 272009 [Gene Symbol: Srsf12] [Locus Tag: ] [Chromosome: 4] [Map Location: 4\|4 A5] [D | 4.757490907 | 2.84E-10 | 2.22E-09 |
| 15551 | ENSMUSG00000032030 | Cul5 | 75717 [Gene Symbol: Cul5] [Locus Tag: ] [Chromosome: 9] [Map Location: 9\|9 C] [Descr | 4.755464246 | 1.05E-16 | 2.32E-15 |
| 1418 | ENSMUSG00000032360 | Hcrtr2 | 387285 [Gene Symbol: Hcrtr2] [Locus Tag: ] [Chromosome: 9] [Map Location: 9\|9 D] [D | 4.755461909 | 5.77E-08 | 3.17E-07 |
| 18448 | ENSMUSG00000029279 | Brdt | 114642 [Gene Symbol: Brdt] [Locus Tag: ] [Chromosome: 5] [Map Location: 5\|5 E5] [Des | 4.752289557 | 2.85E-08 | 1.65E-07 |
| 17393 | ENSMUSG00000040322 | Slc25a24 | 229731 [Gene Symbol: Slc25a24] [Locus Tag: ] [Chromosome: 3] [Map Location: 3\|3 F3 | 4.751965678 | 4.38E-11 | 3.89E-10 |
| 14165 | ENSMUSG00000036990 | Otud4 | 73945 [Gene Symbol: Otud4] [Locus Tag: ] [Chromosome: 8] [Map Location: 8 37.74 cM | 4.748663443 | 1.40E-17 | 3.45E-16 |
| 10776 | ENSMUSG00000037369 | Kdm6a | 22289 [Gene Symbol: Kdm6a] [Locus Tag: ] [Chromosome: X] [Map Location: X A1.2-A1. | 4.744685145 | 1.64E-16 | 3.53E-15 |
| 4316 | ENSMUSG00000032468 | Armc8 | 74125 [Gene Symbol: Armc8] [Locus Tag: ] [Chromosome: 9] [Map Location: 9\|9 F1] [De | 4.742799259 | 8.57E-16 | 1.62E-14 |
| 10214 | ENSMUSG00000055489 | Ano5 | 233246 [Gene Symbol: Ano5] [Locus Tag: ] [Chromosome: 7] [Map Location: 7\|7 B5] [De | 4.742182149 | 6.94E-07 | 3.21E-06 |
| 7469 | ENSMUSG00000052748 | Swt1 | 66875 [Gene Symbol: Swt1] [Locus Tag: ] [Chromosome: 1] [Map Location: 1\|1 G2] [Des | 4.742033875 | 3.94E-11 | 3.52E-10 |
| 8060 | ENSMUSG00000021076 | Actr10 | 56444 [Gene Symbol: Actr10] [Locus Tag: ] [Chromosome: 12] [Map Location: 12\|12 C3 | 4.74017707 | 1.96E-15 | 3.48E-14 |
| 4868 | ENSMUSG00000030867 | Plk1 | 18817 [Gene Symbol: Plk1] [Locus Tag: ] [Chromosome: 7] [Map Location: 7 F3\|7 65.52 | 4.739410865 | 0.000239 | 0.000702 |
| 1590 | ENSMUSG00000040747 | Cd53 | 12508 [Gene Symbol: Cd53] [Locus Tag: ] [Chromosome: 3] [Map Location: 3 F2.3\|3 46 | 4.738409576 | 0.00771 | 0.016471 |
| 17970 | ENSMUSG00000058587 | Tmod3 | 50875 [Gene Symbol: Tmod3] [Locus Tag: ] [Chromosome: 9] [Map Location: 9 D\|9 42.3 | 4.737823834 | 9.49E-16 | 1.78E-14 |
| 6080 | ENSMUSG00000020994 | Pnn | 18949 [Gene Symbol: Pnn] [Locus Tag: ] [Chromosome: 12] [Map Location: 12 C1\|12 26 | 4.728359177 | 3.46E-13 | 4.26E-12 |
| 11793 | ENSMUSG00000025892 | Gria4 | 14802 [Gene Symbol: Gria4] [Locus Tag: ] [Chromosome: 9] [Map Location: 9 A1\|9 2.46 | 4.727071631 | 1.63E-13 | 2.13E-12 |
| 4630 | ENSMUSG00000033192 | Lpcat2 | 270084 [Gene Symbol: Lpcat2] [Locus Tag: ] [Chromosome: 8] [Map Location: 8\|8 C5] [ | 4.726131088 | 7.25E-08 | 3.93E-07 |
| 4541 | ENSMUSG00000021706 | Zfyve16 | 218441 [Gene Symbol: Zfyve16] [Locus Tag: ] [Chromosome: 13] [Map Location: 13\|13 | 4.726130356 | 6.81E-11 | 5.87E-10 |
| 9431 | ENSMUSG00000027699 | Ect2 | 13605 [Gene Symbol: Ect2] [Locus Tag: ] [Chromosome: 3] [Map Location: 3\|3 B] [Descr | 4.724753909 | 3.83E-06 | 1.56E-05 |
| 18389 | ENSMUSG00000039740 | Alg2 | 56737 [Gene Symbol: Alg2] [Locus Tag: ] [Chromosome: 4] [Map Location: 4\|4 B2] [Desc | 4.722388729 | 2.44E-16 | 5.06E-15 |
| 12880 | ENSMUSG00000019961 | Tmpo | 21917 [Gene Symbol: Tmpo] [Locus Tag: ] [Chromosome: 10] [Map Location: 10 C2\|10 | 4.720689759 | 5.86E-11 | 5.10E-10 |
| 17751 | ENSMUSG00000036097 | Fam178a | 226151 [Gene Symbol: Fam178a] [Locus Tag: ] [Chromosome: 19] [Map Location: 19\|1 | 4.718347439 | 6.85E-14 | 9.57E-13 |
| 16974 | ENSMUSG00000021703 | Serinc5 | 218442 [Gene Symbol: Serinc5] [Locus Tag: ] [Chromosome: 13] [Map Location: 13\|13 | 4.718031842 | 4.35E-13 | 5.29E-12 |
| 10344 | ENSMUSG00000033847 | Pla2g4c | 232889 [Gene Symbol: Pla2g4c] [Locus Tag: ] [Chromosome: 7] [Map Location: 7 A1\|7 | 4.716124959 | 0.003242 | 0.007542 |
| 10037 | ENSMUSG00000066595 | Mfsd7b | 226844 [Gene Symbol: Mfsd7b] [Locus Tag: ] [Chromosome: 1] [Map Location: 1\|1 H6] | 4.710154058 | 5.01E-13 | 6.03E-12 |
| 4839 | ENSMUSG00000062184 | Hs6st2 | 50786 [Gene Symbol: Hs6st2] [Locus Tag: ] [Chromosome: X] [Map Location: X\|X A3.3] | 4.707881597 | 3.83E-15 | 6.44E-14 |
| 17431 | ENSMUSG00000050029 | Rap2c | 72065 [Gene Symbol: Rap2c] [Locus Tag: ] [Chromosome: X] [Map Location: X\|X A5] [De | 4.705527755 | 1.47E-16 | 3.18E-15 |
| 16938 | ENSMUSG00000043015 | Tmem194b | 227094 [Gene Symbol: Tmem194b] [Locus Tag: ] [Chromosome: 1] [Map Location: 1\|1 | 4.703631429 | 4.06E-11 | 3.62E-10 |
| 7679 | ENSMUSG00000020653 | Klf11 | 194655 [Gene Symbol: Klf11] [Locus Tag: ] [Chromosome: 12] [Map Location: 12 A1.3\| | 4.702451479 | 4.83E-16 | 9.51E-15 |
| 867 | ENSMUSG00000075701 | Vimp | 109815 [Gene Symbol: Vimp] [Locus Tag: ] [Chromosome: 7] [Map Location: 7 C\|7 35.4 | 4.702442088 | 9.00E-13 | 1.05E-11 |
| 2293 | ENSMUSG00000041986 | Elmod1 | 270162 [Gene Symbol: Elmod1] [Locus Tag: ] [Chromosome: 9] [Map Location: 9\|9 A5.3 | 4.700218444 | 1.02E-16 | 2.25E-15 |
| 3467 | ENSMUSG00000022429 | Dmc1 | 13404 [Gene Symbol: Dmc1] [Locus Tag: ] [Chromosome: 15] [Map Location: 15 E1\|15 | 4.695074628 | 8.31E-05 | 0.000265 |
| 11062 | ENSMUSG00000043463 | Rab9b | 319642 [Gene Symbol: Rab9b] [Locus Tag: ] [Chromosome: X] [Map Location: X\|X F1] [D | 4.694381628 | 2.22E-11 | 2.06E-10 |
| 9240 | ENSMUSG00000024472 | Dcp2 | 70640 [Gene Symbol: Dcp2] [Locus Tag: ] [Chromosome: 18] [Map Location: 18\|18 B3] | 4.690936324 | 9.25E-17 | 2.07E-15 |
| 8987 | ENSMUSG00000059588 | Calcrl | 54598 [Gene Symbol: Calcrl] [Locus Tag: ] [Chromosome: 2] [Map Location: 2\|2 D] [Des | 4.688352761 | 2.71E-10 | 2.12E-09 |
| 1395 | ENSMUSG00000020493 | Prr11 | 270906 [Gene Symbol: Prr11] [Locus Tag: ] [Chromosome: 11] [Map Location: 11\|11 C] | 4.682223539 | 1.70E-06 | 7.39E-06 |
| 6411 | ENSMUSG00000035171 | 1110059E24Rik | 66206 [Gene Symbol: 1110059E24Rik] [Locus Tag: ] [Chromosome: 19] [Map Location: | 4.680534817 | 1.45E-11 | 1.38E-10 |
| 4771 | ENSMUSG00000055228 | Gm36298; Zfp935 | 102640165 [Gene Symbol: Gm36298] [Locus Tag: ] [Chromosome: 13] [Map Location: | 4.679626253 | 3.45E-08 | 1.98E-07 |

| | | | | | | |
|---|---|---|---|---|---|---|
| 11211 | ENSMUSG00000028214 | Gem | 14579 [Gene Symbol: Gem] [Locus Tag: ] [Chromosome: 4] [Map Location: 4 A1\|4 5.34 | 4.676619381 | 0.000242 | 0.000709 |
| 1384 | ENSMUSG00000063320 | 1190007I07Rik | 544717 [Gene Symbol: 1190007I07Rik] [Locus Tag: ] [Chromosome: 10] [Map Location | 4.673016886 | 1.74E-06 | 7.55E-06 |
| 7151 | ENSMUSG00000056211 | R3hdm1 | 226412 [Gene Symbol: R3hdm1] [Locus Tag: ] [Chromosome: 1] [Map Location: 1\|1 E4] | 4.672014462 | 1.34E-11 | 1.29E-10 |
| 17798 | ENSMUSG00000015757 | Ppil4 | 67418 [Gene Symbol: Ppil4] [Locus Tag: ] [Chromosome: 10] [Map Location: 10\|10 A1] | 4.671242189 | 5.26E-16 | 1.03E-14 |
| 5371 | ENSMUSG00000079427 | Mthfsl | 100039707 [Gene Symbol: Mthfsl] [Locus Tag: ] [Chromosome: 9] [Map Location: 9 E3.: | 4.669884899 | 2.21E-07 | 1.11E-06 |
| 17049 | ENSMUSG00000027286 | Lrrc57 | 66606 [Gene Symbol: Lrrc57] [Locus Tag: ] [Chromosome: 2] [Map Location: 2\|2 E5] [De | 4.666705498 | 3.37E-10 | 2.60E-09 |
| 9640 | ENSMUSG00000053317 | Sec61b | 66212 [Gene Symbol: Sec61b] [Locus Tag: ] [Chromosome: 4] [Map Location: 4\|4 B1] [D | 4.663212274 | 1.18E-05 | 4.42E-05 |
| 8946 | ENSMUSG00000005804 | Bloc1s6 | 18457 [Gene Symbol: Bloc1s6] [Locus Tag: ] [Chromosome: 2] [Map Location: 2 E5\|2 60 | 4.663128229 | 2.18E-15 | 3.82E-14 |
| 12050 | ENSMUSG00000021377 | Dek | 110052 [Gene Symbol: Dek] [Locus Tag: ] [Chromosome: 13] [Map Location: 13 24.5 cM | 4.661203713 | 5.02E-16 | 9.85E-15 |
| 7202 | ENSMUSG00000024425 | Ndfip1 | 65113 [Gene Symbol: Ndfip1] [Locus Tag: ] [Chromosome: 18] [Map Location: 18\|18 B3 | 4.660132856 | 2.50E-16 | 5.17E-15 |
| 10658 | ENSMUSG00000060445 | Sycp2 | 320558 [Gene Symbol: Sycp2] [Locus Tag: ] [Chromosome: 2] [Map Location: 2\|2 H4] [D | 4.658505134 | 6.31E-05 | 0.000206 |
| 12344 | ENSMUSG00000023027 | Atf1 | 11908 [Gene Symbol: Atf1] [Locus Tag: ] [Chromosome: 15] [Map Location: 15\|15 F3] [I | 4.65797022 | 3.66E-13 | 4.48E-12 |
| 9890 | ENSMUSG00000039531 | Zufsp | 72580 [Gene Symbol: Zufsp] [Locus Tag: ] [Chromosome: 10] [Map Location: 10\|10 B1] | 4.656239877 | 6.46E-10 | 4.78E-09 |
| 6960 | ENSMUSG00000039968 | Rsbn1l | 242860 [Gene Symbol: Rsbn1l] [Locus Tag: ] [Chromosome: 5] [Map Location: 5\|5 A3] [I | 4.650651079 | 2.39E-16 | 4.99E-15 |
| 7365 | ENSMUSG00000027981 | Rnpc3 | 67225 [Gene Symbol: Rnpc3] [Locus Tag: ] [Chromosome: 3] [Map Location: 3\|3 G1] [De | 4.642745179 | 5.90E-10 | 4.40E-09 |
| 6790 | ENSMUSG00000024259 | Slc25a46 | 67453 [Gene Symbol: Slc25a46] [Locus Tag: ] [Chromosome: 18] [Map Location: 18\|18 | 4.641532087 | 1.95E-16 | 4.14E-15 |
| 11237 | ENSMUSG00000031403 | Dkc1 | 245474 [Gene Symbol: Dkc1] [Locus Tag: ] [Chromosome: X] [Map Location: X\|X A7.3] [ | 4.639332269 | 8.89E-15 | 1.43E-13 |
| 4432 | ENSMUSG00000059839 | Zfp874b | 408067 [Gene Symbol: Zfp874b] [Locus Tag: ] [Chromosome: 13] [Map Location: 13\|13 | 4.639205337 | 1.13E-06 | 5.05E-06 |
| 4571 | ENSMUSG00000029916 | Agk | 69923 [Gene Symbol: Agk] [Locus Tag: ] [Chromosome: 6] [Map Location: 6\|6 B1] [Descr | 4.638602065 | 2.84E-09 | 1.92E-08 |
| 10278 | ENSMUSG00000018666 | Cbx1 | 12412 [Gene Symbol: Cbx1] [Locus Tag: ] [Chromosome: 11] [Map Location: 11 D\|11 60 | 4.638541244 | 3.40E-15 | 5.76E-14 |
| 17327 | ENSMUSG00000021760 | Gpx8 | 69590 [Gene Symbol: Gpx8] [Locus Tag: ] [Chromosome: 13] [Map Location: 13\|13 D2. | 4.637288319 | 1.81E-05 | 6.57E-05 |
| 5142 | ENSMUSG00000033031 | C330027C09Rik | 224171 [Gene Symbol: C330027C09Rik] [Locus Tag: ] [Chromosome: 16] [Map Locatior | 4.636703349 | 1.16E-08 | 7.19E-08 |
| 6504 | ENSMUSG00000015882 | Lcorl | 209707 [Gene Symbol: Lcorl] [Locus Tag: ] [Chromosome: 5] [Map Location: 5\|5 B3] [De | 4.63096836 | 1.40E-16 | 3.04E-15 |
| 11737 | ENSMUSG00000024943 | Smc5 | 226026 [Gene Symbol: Smc5] [Locus Tag: ] [Chromosome: 19] [Map Location: 19\|19 B] | 4.628617423 | 9.67E-11 | 8.11E-10 |
| 17893 | ENSMUSG00000055760 | Gemin6 | 67242 [Gene Symbol: Gemin6] [Locus Tag: ] [Chromosome: 17] [Map Location: 17\|17 E | 4.62636107 | 0.000396 | 0.001116 |
| 12750 | ENSMUSG00000030265 | Kras | 16653 [Gene Symbol: Kras] [Locus Tag: ] [Chromosome: 6] [Map Location: 6 77.37 cM\|6 | 4.621666223 | 4.57E-17 | 1.06E-15 |
| 4530 | ENSMUSG00000054162 | Spock3 | 72902 [Gene Symbol: Spock3] [Locus Tag: ] [Chromosome: 8] [Map Location: 8\|8 B3.1] | 4.621031486 | 3.91E-06 | 1.59E-05 |
| 7200 | ENSMUSG00000032925 | Itgbl1 | 223272 [Gene Symbol: Itgbl1] [Locus Tag: ] [Chromosome: 14] [Map Location: 14\|14 E5 | 4.618837597 | 1.63E-11 | 1.55E-10 |
| 479 | ENSMUSG00000055150 | Zfp78 | 330463 [Gene Symbol: Zfp78] [Locus Tag: ] [Chromosome: 7] [Map Location: 7 A1\|7 3.6 | 4.618819145 | 8.67E-05 | 0.000275 |
| 7035 | ENSMUSG00000020954 | Strn3 | 94186 [Gene Symbol: Strn3] [Locus Tag: ] [Chromosome: 12] [Map Location: 12\|12 C1] | 4.617418057 | 2.01E-12 | 2.19E-11 |
| 3171 | ENSMUSG00000067261 | Foxd3 | 15221 [Gene Symbol: Foxd3] [Locus Tag: ] [Chromosome: 4] [Map Location: 4 C6\|4 45.: | 4.616596979 | 0.001275 | 0.003246 |
| 12422 | ENSMUSG00000035967 | Ddx26b | 236790 [Gene Symbol: Ddx26b] [Locus Tag: ] [Chromosome: X] [Map Location: X\|X A5] | 4.61464507 | 3.26E-12 | 3.43E-11 |
| 2737 | ENSMUSG00000016493 | Cd46 | 17221 [Gene Symbol: Cd46] [Locus Tag: ] [Chromosome: 1] [Map Location: 1 H6\|1 98.4 | 4.611285328 | 4.25E-05 | 0.000143 |
| 11091 | ENSMUSG00000044681 | Cnpy1 | 269637 [Gene Symbol: Cnpy1] [Locus Tag: ] [Chromosome: 5] [Map Location: 5\|5 B1] [I | 4.610176297 | 0.003746 | 0.008582 |
| 18112 | ENSMUSG00000020132 | Rab21 | 216344 [Gene Symbol: Rab21] [Locus Tag: ] [Chromosome: 10] [Map Location: 10\|10 D | 4.608640043 | 1.88E-15 | 3.36E-14 |
| 3960 | ENSMUSG00000029128 | Rab28 | 100972 [Gene Symbol: Rab28] [Locus Tag: ] [Chromosome: 5] [Map Location: 5\|5 B2] [I | 4.606165908 | 5.59E-13 | 6.69E-12 |
| 13863 | ENSMUSG00000073062 | Zxdb | 668166 [Gene Symbol: Zxdb] [Locus Tag: ] [Chromosome: X] [Map Location: X\|X C3] [De | 4.605685908 | 1.03E-12 | 1.19E-11 |
| 18316 | ENSMUSG00000039987 | Phtf2 | 68770 [Gene Symbol: Phtf2] [Locus Tag: ] [Chromosome: 5] [Map Location: 5\|5 A3] [De: | 4.60551986 | 3.28E-16 | 6.64E-15 |
| 10035 | ENSMUSG00000022512 | Cldn1 | 12737 [Gene Symbol: Cldn1] [Locus Tag: ] [Chromosome: 16] [Map Location: 16\|16 B2 | 4.601657522 | 1.10E-06 | 4.91E-06 |
| 533 | ENSMUSG00000020580 | Rock2 | 19878 [Gene Symbol: Rock2] [Locus Tag: ] [Chromosome: 12] [Map Location: 12\|12 A3 | 4.600911957 | 5.25E-12 | 5.38E-11 |
| 3389 | ENSMUSG00000022704 | Qtrtd1 | 106248 [Gene Symbol: Qtrtd1] [Locus Tag: ] [Chromosome: 16] [Map Location: 16\|16 E | 4.600115561 | 9.72E-14 | 1.32E-12 |
| 10655 | ENSMUSG00000035725 | Prkx | 19108 [Gene Symbol: Prkx] [Locus Tag: ] [Chromosome: X] [Map Location: X\|X A7.3] [De | 4.598556417 | 2.08E-11 | 1.94E-10 |
| 224 | ENSMUSG00000043019 | Edem3 | 66967 [Gene Symbol: Edem3] [Locus Tag: ] [Chromosome: 1] [Map Location: 1\|1 G2] [D | 4.597810532 | 1.33E-14 | 2.07E-13 |
| 9049 | ENSMUSG00000034755 | Pcdh11x | 245578 [Gene Symbol: Pcdh11x] [Locus Tag: ] [Chromosome: X] [Map Location: X\|X E2 | 4.587649557 | 6.30E-13 | 7.49E-12 |
| 11868 | ENSMUSG00000037400 | Atp11b | 76295 [Gene Symbol: Atp11b] [Locus Tag: ] [Chromosome: 3] [Map Location: 3\|3 B] [De | 4.586873704 | 6.88E-16 | 1.33E-14 |
| 17592 | ENSMUSG00000058446 | Znrf2 | 387524 [Gene Symbol: Znrf2] [Locus Tag: ] [Chromosome: 6] [Map Location: 6 B3\|6 27. | 4.584921155 | 3.63E-13 | 4.46E-12 |
| 17821 | ENSMUSG00000024302 | Dtna | 13527 [Gene Symbol: Dtna] [Locus Tag: ] [Chromosome: 18] [Map Location: 18 A2\|18 1 | 4.583664076 | 1.89E-10 | 1.52E-09 |
| 13097 | ENSMUSG00000021413 | Prpf4b | 19134 [Gene Symbol: Prpf4b] [Locus Tag: ] [Chromosome: 13] [Map Location: 13\|13 A5 | 4.582272298 | 3.59E-12 | 3.76E-11 |
| 12451 | ENSMUSG00000005470 | Asf1b | 66929 [Gene Symbol: Asf1b] [Locus Tag: ] [Chromosome: 8] [Map Location: 8\|8 C3] [De: | 4.580104885 | 9.91E-06 | 3.76E-05 |
| 17643 | ENSMUSG00000014353 | Tmem87b | 72477 [Gene Symbol: Tmem87b] [Locus Tag: ] [Chromosome: 2] [Map Location: 2\|2 F3 | 4.576241245 | 1.83E-14 | 2.80E-13 |
| 5835 | ENSMUSG00000094441 | Zfp955a | 77652 [Gene Symbol: Zfp955a] [Locus Tag: ] [Chromosome: 17] [Map Location: 17\|17 E | 4.575718267 | 5.87E-09 | 3.81E-08 |
| 1922 | ENSMUSG00000062270 | Morf4l1b; Morf4l1 | 627352 [Gene Symbol: Morf4l1b] [Locus Tag: ] [Chromosome: 19] [Map Location: 19\|1 | 4.571434554 | 1.30E-12 | 1.46E-11 |
| 1047 | ENSMUSG00000055447 | Cd47 | 16423 [Gene Symbol: Cd47] [Locus Tag: ] [Chromosome: 16] [Map Location: 16\|16 B5] | 4.570936557 | 2.63E-16 | 5.43E-15 |
| 13634 | ENSMUSG00000026241 | Nppc | 18159 [Gene Symbol: Nppc] [Locus Tag: ] [Chromosome: 1] [Map Location: 1 D\|1 43.98 | 4.569194415 | 0.009707 | 0.020282 |
| 11508 | ENSMUSG00000047675 | Rps8 | 20116 [Gene Symbol: Rps8] [Locus Tag: ] [Chromosome: 4] [Map Location: 4\|4 D1] [Des | 4.569107345 | 3.47E-09 | 2.32E-08 |
| 1548 | ENSMUSG00000095253 | Zfp799 | 240064 [Gene Symbol: Zfp799] [Locus Tag: ] [Chromosome: 17] [Map Location: 17\|17 E | 4.568187987 | 1.23E-11 | 1.19E-10 |
| 13063 | ENSMUSG00000025742 | Prps2 | 110639 [Gene Symbol: Prps2] [Locus Tag: ] [Chromosome: X] [Map Location: X F2-F3\|X | 4.561633963 | 1.95E-10 | 1.56E-09 |
| 7620 | ENSMUSG00000016257 | Slmo2 | 66390 [Gene Symbol: Slmo2] [Locus Tag: ] [Chromosome: 2] [Map Location: 2\|2 H4] [De | 4.559369604 | 2.98E-15 | 5.11E-14 |
| 2813 | ENSMUSG00000054387 | Mdm4 | 17248 [Gene Symbol: Mdm4] [Locus Tag: ] [Chromosome: 1] [Map Location: 1 57.75 cM | 4.552301056 | 2.33E-16 | 4.88E-15 |
| 12014 | ENSMUSG00000035329 | Fbxo33 | 70611 [Gene Symbol: Fbxo33] [Locus Tag: ] [Chromosome: 12] [Map Location: 12\|12 C | 4.549565282 | 1.94E-14 | 2.96E-13 |
| 4550 | ENSMUSG00000075316 | Scn9a | 20274 [Gene Symbol: Scn9a] [Locus Tag: ] [Chromosome: 2] [Map Location: 2 C1.3\|2 39 | 4.548432712 | 1.12E-08 | 6.93E-08 |
| 5551 | ENSMUSG00000034607 | Pof1b | 69693 [Gene Symbol: Pof1b] [Locus Tag: ] [Chromosome: X] [Map Location: X\|X E1] [De | 4.547546508 | 0.000338 | 0.000963 |
| 15391 | ENSMUSG00000079436 | Kcnj13 | 100040591 [Gene Symbol: Kcnj13] [Locus Tag: ] [Chromosome: 1] [Map Location: 1 D\| | 4.546147145 | 0.000173 | 0.000521 |
| 1317 | ENSMUSG00000022762 | Ncam2 | 17968 [Gene Symbol: Ncam2] [Locus Tag: ] [Chromosome: 16] [Map Location: 16 C1-3\| | 4.544659608 | 7.32E-05 | 0.000236 |
| 17186 | ENSMUSG00000019865 | Nmbr | 18101 [Gene Symbol: Nmbr] [Locus Tag: ] [Chromosome: 10] [Map Location: 10\|10 A2] | 4.544514715 | 4.12E-06 | 1.66E-05 |
| 138 | ENSMUSG00000021135 | Slc10a1 | 20493 [Gene Symbol: Slc10a1] [Locus Tag: ] [Chromosome: 12] [Map Location: 12 D1\|1 | 4.543535461 | 0.006521 | 0.014147 |
| 3683 | ENSMUSG00000062762 | Ei24 | 13663 [Gene Symbol: Ei24] [Locus Tag: ] [Chromosome: 9] [Map Location: 9 A4\|9 20.68 | 4.540442961 | 2.40E-16 | 4.99E-15 |
| 1239 | ENSMUSG00000025037 | Maoa | 17161 [Gene Symbol: Maoa] [Locus Tag: ] [Chromosome: X] [Map Location: X A2\|X 11.7 | 4.537936103 | 9.23E-08 | 4.93E-07 |
| 4218 | ENSMUSG00000067942 | Zfp160 | 224585 [Gene Symbol: Zfp160] [Locus Tag: ] [Chromosome: 17] [Map Location: 17 A3.2 | 4.534785016 | 2.80E-14 | 4.17E-13 |
| 171 | ENSMUSG00000009633 | G0s2 | 14373 [Gene Symbol: G0s2] [Locus Tag: ] [Chromosome: 1] [Map Location: 1 H6\|1 97.6 | 4.529996379 | 0.000333 | 0.000951 |
| 10446 | ENSMUSG00000023845 | Lnpep | 240028 [Gene Symbol: Lnpep] [Locus Tag: ] [Chromosome: 17] [Map Location: 17\|17 A | 4.529541399 | 1.48E-16 | 3.21E-15 |
| 12018 | ENSMUSG00000003038 | Hmgn2 | 15331 [Gene Symbol: Hmgn2] [Locus Tag: ] [Chromosome: 4] [Map Location: 4 D3\|4 66 | 4.528455973 | 1.83E-07 | 9.33E-07 |
| 15250 | ENSMUSG00000024594 | Prrc1 | 73137 [Gene Symbol: Prrc1] [Locus Tag: ] [Chromosome: 18] [Map Location: 18\|18 D3] | 4.521673762 | 5.04E-15 | 8.34E-14 |
| 4311 | ENSMUSG00000028522 | Mier1 | 71148 [Gene Symbol: Mier1] [Locus Tag: ] [Chromosome: 4] [Map Location: 4\|4 C6] [De | 4.521362765 | 1.14E-12 | 1.30E-11 |
| 4642 | ENSMUSG00000059447 | Hadhb | 231086 [Gene Symbol: Hadhb] [Locus Tag: ] [Chromosome: 5] [Map Location: 5\|5 B1] [I | 4.520864302 | 1.29E-08 | 7.95E-08 |
| 17725 | ENSMUSG00000037214 | Thap1 | 73754 [Gene Symbol: Thap1] [Locus Tag: ] [Chromosome: 8] [Map Location: 8\|8 A2] [De | 4.518495855 | 0.000201 | 0.000597 |
| 14081 | ENSMUSG00000034321 | Exosc1 | 66583 [Gene Symbol: Exosc1] [Locus Tag: ] [Chromosome: 19] [Map Location: 19\|19 D: | 4.51804067 | 1.16E-11 | 1.13E-10 |
| 787 | ENSMUSG00000022992 | Kansl2 | 69612 [Gene Symbol: Kansl2] [Locus Tag: ] [Chromosome: 15] [Map Location: 15\|15 F2 | 4.51281798 | 3.41E-15 | 5.77E-14 |
| 913 | ENSMUSG00000021815 | Mss51 | 74843 [Gene Symbol: Mss51] [Locus Tag: ] [Chromosome: 14] [Map Location: 14\|14 B] | 4.512641634 | 1.54E-06 | 6.74E-06 |
| 9515 | ENSMUSG00000050312 | Nsun3 | 106338 [Gene Symbol: Nsun3] [Locus Tag: ] [Chromosome: 16] [Map Location: 16\|16 C | 4.507450321 | 3.16E-13 | 3.92E-12 |
| 13579 | ENSMUSG00000073792 | Alg6 | 320438 [Gene Symbol: Alg6] [Locus Tag: ] [Chromosome: 4] [Map Location: 4\|4 C6] [De | 4.49612947 | 2.92E-07 | 1.43E-06 |
| 18411 | ENSMUSG00000046351 | Zfp322a | 218100 [Gene Symbol: Zfp322a] [Locus Tag: ] [Chromosome: 13] [Map Location: 13\|13 | 4.49531525 | 5.54E-12 | 5.66E-11 |
| 10497 | ENSMUSG00000068854 | Hist2h2be | 319190 [Gene Symbol: Hist2h2be] [Locus Tag: ] [Chromosome: 3] [Map Location: 3\|3 F | 4.494244775 | 6.91E-06 | 2.70E-05 |
| 10871 | ENSMUSG00000034883 | Lrr1 | 69706 [Gene Symbol: Lrr1] [Locus Tag: ] [Chromosome: 12] [Map Location: 12\|12 C3] [I | 4.493061259 | 0.020756 | 0.039826 |

| | | | | | | |
|---|---|---|---|---|---|---|
| 117 | ENSMUSG00000026715 | Serpinc1 | 11905 [Gene Symbol: Serpinc1] [Locus Tag: ] [Chromosome: 1] [Map Location: 1 H2.1\|1 | 4.492017204 | 2.90E-10 | 2.27E-09 |
| 4593 | ENSMUSG00000024800 | Rpp30 | 54364 [Gene Symbol: Rpp30] [Locus Tag: ] [Chromosome: 19] [Map Location: 19\|19 C2 | 4.487262273 | 7.30E-10 | 5.37E-09 |
| 8 | ENSMUSG00000033972 | Zfp944 | 319615 [Gene Symbol: Zfp944] [Locus Tag: ] [Chromosome: 17] [Map Location: 17\|17 A | 4.487195887 | 3.51E-12 | 3.68E-11 |
| 16041 | ENSMUSG00000068457 | Uty | 22290 [Gene Symbol: Uty] [Locus Tag: ] [Chromosome: Y] [Map Location: Y\|Y A1] [Descr | 4.486574628 | 1.04E-07 | 5.50E-07 |
| 9138 | ENSMUSG00000026893 | Gca | 227960 [Gene Symbol: Gca] [Locus Tag: ] [Chromosome: 2] [Map Location: 2\|2 C1.3] [D | 4.472301274 | 4.21E-09 | 2.79E-08 |
| 6398 | ENSMUSG00000038784 | Cnot4 | 53621 [Gene Symbol: Cnot4] [Locus Tag: ] [Chromosome: 6] [Map Location: 6\|6 B1] [De | 4.471853173 | 5.45E-16 | 1.06E-14 |
| 12317 | ENSMUSG00000036197 | Gxylt1 | 223827 [Gene Symbol: Gxylt1] [Locus Tag: ] [Chromosome: 15] [Map Location: 15\|15 E | 4.469536523 | 4.47E-12 | 4.62E-11 |
| 1349 | ENSMUSG00000054237 | Fra10ac1 | 70567 [Gene Symbol: Fra10ac1] [Locus Tag: ] [Chromosome: 19] [Map Location: 19\|19 | 4.46761928 | 1.30E-12 | 1.46E-11 |
| 4683 | ENSMUSG00000030208 | Emp1 | 13730 [Gene Symbol: Emp1] [Locus Tag: ] [Chromosome: 6] [Map Location: 6 G1\|6 66.2 | 4.465677375 | 1.76E-06 | 7.63E-06 |
| 11842 | ENSMUSG00000054976 | Nyap2 | 241134 [Gene Symbol: Nyap2] [Locus Tag: ] [Chromosome: 1] [Map Location: 1\|1 C5] [D | 4.464709163 | 1.17E-16 | 2.57E-15 |
| 156 | ENSMUSG00000032251 | Irak1bp1 | 65099 [Gene Symbol: Irak1bp1] [Locus Tag: ] [Chromosome: 9] [Map Location: 9\|9 E2] [ | 4.464342071 | 1.66E-13 | 2.16E-12 |
| 9868 | ENSMUSG00000042305 | Tmem183a | 57439 [Gene Symbol: Tmem183a] [Locus Tag: ] [Chromosome: 1] [Map Location: 1\|1 E4 | 4.464093195 | 1.10E-15 | 2.05E-14 |
| 11825 | ENSMUSG00000062683 | Atp5g2 | 67942 [Gene Symbol: Atp5g2] [Locus Tag: ] [Chromosome: 15] [Map Location: 15\|15 F3 | 4.464039708 | 0.000406 | 0.001142 |
| 14296 | ENSMUSG00000022151 | Ttc33 | 67515 [Gene Symbol: Ttc33] [Locus Tag: ] [Chromosome: 15] [Map Location: 15\|15 A1] | 4.461116346 | 3.17E-14 | 4.68E-13 |
| 9057 | ENSMUSG00000058835 | Abi1 | 11308 [Gene Symbol: Abi1] [Locus Tag: ] [Chromosome: 2] [Map Location: 2 A3\|2 15.18 | 4.460480742 | 4.19E-16 | 8.35E-15 |
| 3920 | ENSMUSG00000038374 | Rbm8a | 60365 [Gene Symbol: Rbm8a] [Locus Tag: ] [Chromosome: 3] [Map Location: 3\|3 H3] [D | 4.458871711 | 3.53E-13 | 4.34E-12 |
| 12933 | ENSMUSG00000021215 | Net1 | 56349 [Gene Symbol: Net1] [Locus Tag: ] [Chromosome: 13] [Map Location: 13\|13 A1] [ | 4.456565343 | 1.41E-12 | 1.57E-11 |
| 17248 | ENSMUSG00000048495 | Tyw5 | 68736 [Gene Symbol: Tyw5] [Locus Tag: ] [Chromosome: 1] [Map Location: 1\|1 C1.3] [D | 4.451134128 | 1.50E-07 | 7.76E-07 |
| 2281 | ENSMUSG00000023861 | Mpc1 | 55951 [Gene Symbol: Mpc1] [Locus Tag: ] [Chromosome: 17] [Map Location: 17 A1\|17 | 4.450191296 | 1.54E-10 | 1.25E-09 |
| 6859 | ENSMUSG00000025453 | Nnt | 18115 [Gene Symbol: Nnt] [Locus Tag: ] [Chromosome: 13] [Map Location: 13 67.21 cM | 4.445605432 | 3.63E-13 | 4.46E-12 |
| 17655 | ENSMUSG00000027534 | Snx16 | 74718 [Gene Symbol: Snx16] [Locus Tag: ] [Chromosome: 3] [Map Location: 3\|3 A1] [De | 4.443467306 | 6.22E-10 | 4.62E-09 |
| 12432 | ENSMUSG00000006423 | C330007P06Rik | 77644 [Gene Symbol: C330007P06Rik] [Locus Tag: ] [Chromosome: X] [Map Location: X | 4.442967622 | 1.42E-11 | 1.36E-10 |
| 8497 | ENSMUSG00000024989 | Cep55 | 74107 [Gene Symbol: Cep55] [Locus Tag: ] [Chromosome: 19] [Map Location: 19\|19 C3 | 4.442427492 | 0.000841 | 0.002219 |
| 3870 | ENSMUSG00000056073 | Grik2 | 14806 [Gene Symbol: Grik2] [Locus Tag: ] [Chromosome: 10] [Map Location: 10 B3\|10 | 4.439430509 | 4.26E-12 | 4.41E-11 |
| 7221 | ENSMUSG00000060679 | Mrps9 | 69527 [Gene Symbol: Mrps9] [Locus Tag: ] [Chromosome: 1] [Map Location: 1\|1 B] [Des | 4.432668138 | 1.05E-12 | 1.21E-11 |
| 11106 | ENSMUSG00000021427 | Ssr1 | 107513 [Gene Symbol: Ssr1] [Locus Tag: ] [Chromosome: 13] [Map Location: 13\|13 A3. | 4.432168097 | 2.61E-14 | 3.90E-13 |
| 3502 | ENSMUSG00000032320 | Rcn2 | 26611 [Gene Symbol: Rcn2] [Locus Tag: ] [Chromosome: 9] [Map Location: 9\|9 C] [Desc | 4.427323647 | 2.20E-15 | 3.86E-14 |
| 16479 | ENSMUSG00000009013 | Dynll1 | 56455 [Gene Symbol: Dynll1] [Locus Tag: ] [Chromosome: 5] [Map Location: 5\|5 F] [Des | 4.425938808 | 1.15E-13 | 1.54E-12 |
| 9504 | ENSMUSG00000030231 | Plekha5 | 109135 [Gene Symbol: Plekha5] [Locus Tag: ] [Chromosome: 6] [Map Location: 6\|6 G2] | 4.422514485 | 3.23E-10 | 2.49E-09 |
| 5510 | ENSMUSG00000015002 | Efr3a | 76740 [Gene Symbol: Efr3a] [Locus Tag: ] [Chromosome: 15] [Map Location: 15\|15 D1] | 4.421826304 | 6.90E-16 | 1.33E-14 |
| 16312 | ENSMUSG00000074934 | Grem1 | 23892 [Gene Symbol: Grem1] [Locus Tag: ] [Chromosome: 2] [Map Location: 2 E4\|2 57. | 4.417869769 | 1.29E-07 | 6.74E-07 |
| 7463 | ENSMUSG00000026826 | Nr4a2 | 18227 [Gene Symbol: Nr4a2] [Locus Tag: ] [Chromosome: 2] [Map Location: 2 C1.1\|2 3 | 4.416075453 | 5.76E-12 | 5.87E-11 |
| 11241 | ENSMUSG00000051730 | Mettl5 | 75422 [Gene Symbol: Mettl5] [Locus Tag: ] [Chromosome: 2] [Map Location: 2\|2 C3] [D | 4.414440124 | 1.96E-06 | 8.41E-06 |
| 15426 | ENSMUSG00000028771 | Ptpn12 | 19248 [Gene Symbol: Ptpn12] [Locus Tag: ] [Chromosome: 5] [Map Location: 5\|5 A3-B] | 4.413328719 | 7.88E-14 | 1.09E-12 |
| 4591 | ENSMUSG00000008435 | Rdh13 | 108841 [Gene Symbol: Rdh13] [Locus Tag: ] [Chromosome: 7] [Map Location: 7\|7 A1] [ | 4.411659762 | 2.91E-12 | 3.09E-11 |
| 16039 | ENSMUSG00000038402 | Foxf2 | 14238 [Gene Symbol: Foxf2] [Locus Tag: ] [Chromosome: 13] [Map Location: 13\|13 A4] | 4.411223706 | 7.80E-06 | 3.02E-05 |
| 18232 | ENSMUSG00000058883 | Zfp708 | 432769 [Gene Symbol: Zfp708] [Locus Tag: ] [Chromosome: 13] [Map Location: 13\|13 | 4.410908273 | 3.00E-10 | 2.34E-09 |
| 3144 | ENSMUSG00000020594 | Pum2 | 80913 [Gene Symbol: Pum2] [Locus Tag: ] [Chromosome: 12] [Map Location: 12\|12 A1. | 4.410445296 | 9.65E-17 | 2.15E-15 |
| 14593 | ENSMUSG00000032018 | Sc5d | 235293 [Gene Symbol: Sc5d] [Locus Tag: ] [Chromosome: 9] [Map Location: 9\|9 B] [Des | 4.409922909 | 3.56E-14 | 5.20E-13 |
| 5794 | ENSMUSG00000034488 | Edil3 | 13612 [Gene Symbol: Edil3] [Locus Tag: ] [Chromosome: 13] [Map Location: 13\|13 C3] | 4.408908917 | 2.75E-15 | 4.74E-14 |
| 1370 | ENSMUSG00000027879 | Sec22b | 20333 [Gene Symbol: Sec22b] [Locus Tag: ] [Chromosome: 3] [Map Location: 3\|3 F2.2] | 4.40507147 | 1.58E-14 | 2.43E-13 |
| 5900 | ENSMUSG00000043991 | Pura | 19290 [Gene Symbol: Pura] [Locus Tag: ] [Chromosome: 18] [Map Location: 18\|18 B3] [ | 4.40436763 | 1.71E-10 | 1.38E-09 |
| 8449 | ENSMUSG00000027601 | Mtfr1 | 67472 [Gene Symbol: Mtfr1] [Locus Tag: ] [Chromosome: 3] [Map Location: 3\|3 A2] [De | 4.403823503 | 7.62E-10 | 5.59E-09 |
| 7957 | ENSMUSG00000020321 | Mdh1 | 17449 [Gene Symbol: Mdh1] [Locus Tag: ] [Chromosome: 11] [Map Location: 11 A3.1\|1 | 4.403256136 | 1.99E-14 | 3.03E-13 |
| 18595 | ENSMUSG00000061474 | Mrps36 | 66128 [Gene Symbol: Mrps36] [Locus Tag: ] [Chromosome: 13] [Map Location: 13\|13 D | 4.402990772 | 0.001204 | 0.003089 |
| 8952 | ENSMUSG00000002107 | Celf2 | 14007 [Gene Symbol: Celf2] [Locus Tag: ] [Chromosome: 2] [Map Location: 2\|2 A2-A3] [ | 4.40256304 | 1.51E-16 | 3.26E-15 |
| 2914 | ENSMUSG00000006818 | Sod2 | 20656 [Gene Symbol: Sod2] [Locus Tag: ] [Chromosome: 17] [Map Location: 17 A1\|17 8 | 4.402463081 | 2.57E-15 | 4.46E-14 |
| 1360 | ENSMUSG00000022003 | Slc25a30 | 67554 [Gene Symbol: Slc25a30] [Locus Tag: ] [Chromosome: 14] [Map Location: 14\|14 | 4.399422837 | 2.28E-12 | 2.46E-11 |
| 4971 | ENSMUSG00000022722 | Arl6 | 56297 [Gene Symbol: Arl6] [Locus Tag: ] [Chromosome: 16] [Map Location: 16\|16 C1.2 | 4.397429565 | 5.95E-10 | 4.43E-09 |
| 7829 | ENSMUSG00000022261 | Sdc2 | 15529 [Gene Symbol: Sdc2] [Locus Tag: ] [Chromosome: 15] [Map Location: 15 B3.1\|15 | 4.396077265 | 3.41E-13 | 4.21E-12 |
| 3129 | ENSMUSG00000036257 | Pnpla8 | 67452 [Gene Symbol: Pnpla8] [Locus Tag: ] [Chromosome: 12] [Map Location: 12\|12 B3 | 4.395277181 | 4.43E-16 | 8.78E-15 |
| 7304 | ENSMUSG00000035649 | Zcchc7 | 319885 [Gene Symbol: Zcchc7] [Locus Tag: ] [Chromosome: 4] [Map Location: 4 B1\|4 2 | 4.394675674 | 3.87E-10 | 2.96E-09 |
| 13400 | ENSMUSG00000025104 | Hdgfrp3 | 29877 [Gene Symbol: Hdgfrp3] [Locus Tag: ] [Chromosome: 7] [Map Location: 7\|7 D3] [ | 4.392191872 | 5.17E-16 | 1.01E-14 |
| 7220 | ENSMUSG00000026698 | Pigc | 67292 [Gene Symbol: Pigc] [Locus Tag: ] [Chromosome: 1] [Map Location: 1\|1 H2.1] [De | 4.391053258 | 9.90E-12 | 9.73E-11 |
| 5634 | ENSMUSG00000042523 | Dnal1 | 105000 [Gene Symbol: Dnal1] [Locus Tag: ] [Chromosome: 12] [Map Location: 12\|12 D | 4.389189089 | 2.01E-15 | 3.56E-14 |
| 2856 | ENSMUSG00000074212 | Dnajb14 | 70604 [Gene Symbol: Dnajb14] [Locus Tag: ] [Chromosome: 3] [Map Location: 3\|3 G3] [ | 4.388966203 | 3.45E-16 | 6.95E-15 |
| 10030 | ENSMUSG00000030019 | Fbxl14 | 101358 [Gene Symbol: Fbxl14] [Locus Tag: ] [Chromosome: 6] [Map Location: 6\|6 F1] [ | 4.386885412 | 1.09E-12 | 1.24E-11 |
| 15408 | ENSMUSG00000032067 | Pts | 19286 [Gene Symbol: Pts] [Locus Tag: ] [Chromosome: 9] [Map Location: 9\|9 A5.3] [Des | 4.384451102 | 2.20E-10 | 1.75E-09 |
| 5259 | ENSMUSG00000056531 | Ccdc18 | 73254 [Gene Symbol: Ccdc18] [Locus Tag: ] [Chromosome: 5] [Map Location: 5\|5 E5] [D | 4.383789832 | 2.87E-07 | 1.41E-06 |
| 16755 | ENSMUSG00000052917 | Senp7 | 66315 [Gene Symbol: Senp7] [Locus Tag: ] [Chromosome: 16] [Map Location: 16\|16 C1 | 4.379007118 | 1.17E-15 | 2.16E-14 |
| 13255 | ENSMUSG00000015120 | LOC102641751; Ube | 102641751 [Gene Symbol: LOC102641751] [Locus Tag: ] [Chromosome: 18] [Map Loca | 4.378929913 | 1.46E-13 | 1.93E-12 |
| 7385 | ENSMUSG00000019777 | Hdac2 | 15182 [Gene Symbol: Hdac2] [Locus Tag: ] [Chromosome: 10] [Map Location: 10 19.44 | 4.370382814 | 1.03E-15 | 1.93E-14 |
| 7311 | ENSMUSG00000028247 | Coq3 | 230027 [Gene Symbol: Coq3] [Locus Tag: ] [Chromosome: 4] [Map Location: 4 A3\|4 9.1 | 4.370177675 | 1.58E-09 | 1.11E-08 |
| 18414 | ENSMUSG00000022024 | Sugt1 | 67955 [Gene Symbol: Sugt1] [Locus Tag: ] [Chromosome: 14] [Map Location: 14\|14 D3] | 4.362481596 | 2.45E-13 | 3.10E-12 |
| 3757 | ENSMUSG00000041231 | Ublcp1 | 79560 [Gene Symbol: Ublcp1] [Locus Tag: ] [Chromosome: 11] [Map Location: 11\|11 B | 4.362275238 | 1.97E-07 | 9.96E-07 |
| 14643 | ENSMUSG00000032076 | Cadm1 | 54725 [Gene Symbol: Cadm1] [Locus Tag: ] [Chromosome: 9] [Map Location: 9\|9 B-C] [ | 4.362142845 | 3.35E-16 | 6.77E-15 |
| 18413 | ENSMUSG00000028618 | Tmem59 | 56374 [Gene Symbol: Tmem59] [Locus Tag: ] [Chromosome: 4] [Map Location: 4 C7\|4 5 | 4.361954943 | 1.28E-12 | 1.44E-11 |
| 778 | ENSMUSG00000037818 | Abhd18 | 269423 [Gene Symbol: Abhd18] [Locus Tag: ] [Chromosome: 3] [Map Location: 3\|3 C] [ | 4.361361505 | 3.12E-11 | 2.82E-10 |
| 805 | ENSMUSG00000026775 | Yme1l1 | 27377 [Gene Symbol: Yme1l1] [Locus Tag: ] [Chromosome: 2] [Map Location: 2\|2 A3] [D | 4.360653547 | 6.36E-15 | 1.04E-13 |
| 17772 | ENSMUSG00000022200 | Golph3 | 66629 [Gene Symbol: Golph3] [Locus Tag: ] [Chromosome: 15] [Map Location: 15\|15 A | 4.355746255 | 3.43E-15 | 5.80E-14 |
| 11320 | ENSMUSG00000049076 | Acap2 | 78618 [Gene Symbol: Acap2] [Locus Tag: ] [Chromosome: 16] [Map Location: 16\|16 B2 | 4.354695232 | 1.15E-15 | 2.14E-14 |
| 9548 | ENSMUSG00000024298 | Zfp871 | 208292 [Gene Symbol: Zfp871] [Locus Tag: ] [Chromosome: 17] [Map Location: 17\|17 | 4.354688819 | 5.07E-16 | 9.94E-15 |
| 658 | ENSMUSG00000051890 | Klhdc1 | 271005 [Gene Symbol: Klhdc1] [Locus Tag: ] [Chromosome: 12] [Map Location: 12\|12 | 4.353483622 | 2.81E-07 | 1.38E-06 |
| 3797 | ENSMUSG00000044934 | Zfp367 | 238673 [Gene Symbol: Zfp367] [Locus Tag: ] [Chromosome: 13] [Map Location: 13\|13 | 4.353219542 | 9.05E-08 | 4.84E-07 |
| 2686 | ENSMUSG00000068686 | Cd59b | 333883 [Gene Symbol: Cd59b] [Locus Tag: ] [Chromosome: 2] [Map Location: 2\|2 E2] [ | 4.350094884 | 0.000561 | 0.001532 |
| 3868 | ENSMUSG00000027427 | Polr3f | 70408 [Gene Symbol: Polr3f] [Locus Tag: ] [Chromosome: 2] [Map Location: 2\|2 H1] [De | 4.344158289 | 5.45E-10 | 4.08E-09 |
| 12189 | ENSMUSG00000056313 | 1810011O10Rik | 69068 [Gene Symbol: 1810011O10Rik] [Locus Tag: ] [Chromosome: 8] [Map Location: 8 | 4.34333161 | 7.43E-07 | 3.43E-06 |
| 275 | ENSMUSG00000034551 | Hdx | 245596 [Gene Symbol: Hdx] [Locus Tag: ] [Chromosome: X] [Map Location: X\|X E1] [Des | 4.343291585 | 1.78E-12 | 1.96E-11 |
| 10294 | ENSMUSG00000069769 | Msi2 | 76626 [Gene Symbol: Msi2] [Locus Tag: ] [Chromosome: 11] [Map Location: 11\|11 C] [D | 4.342180497 | 2.27E-16 | 4.76E-15 |
| 16714 | ENSMUSG00000063894 | Zkscan8 | 93681 [Gene Symbol: Zkscan8] [Locus Tag: ] [Chromosome: 13] [Map Location: 13\|13 A | 4.334166316 | 9.75E-10 | 7.02E-09 |
| 14961 | ENSMUSG00000073725 | Lmbrd1 | 68421 [Gene Symbol: Lmbrd1] [Locus Tag: ] [Chromosome: 1] [Map Location: 1\|1 A5] [ | 4.334044273 | 2.50E-14 | 3.75E-13 |
| 1828 | ENSMUSG00000031174 | Rpgr | 19893 [Gene Symbol: Rpgr] [Locus Tag: ] [Chromosome: X] [Map Location: X\|X A1.1] [D | 4.332793833 | 1.41E-10 | 1.16E-09 |
| 1433 | ENSMUSG00000025997 | Ikzf2 | 22779 [Gene Symbol: Ikzf2] [Locus Tag: ] [Chromosome: 1] [Map Location: 1\|1 C3] [Des | 4.33217163 | 4.72E-12 | 4.87E-11 |
| 3529 | ENSMUSG00000038628 | Polr3k | 67005 [Gene Symbol: Polr3k] [Locus Tag: ] [Chromosome: 2] [Map Location: 2\|2 H4] [D | 4.331717732 | 8.56E-13 | 1.00E-11 |

| | | | | | | |
|---|---|---|---|---|---|---|
| 5151 | ENSMUSG00000022462 | Slc38a2 | 67760 [Gene Symbol: Slc38a2] [Locus Tag: ] [Chromosome: 15] [Map Location: 15\|15 F | 4.325458935 | 8.15E-13 | 9.57E-12 |
| 8754 | ENSMUSG00000056854 | Pou3f4 | 18994 [Gene Symbol: Pou3f4] [Locus Tag: ] [Chromosome: X] [Map Location: X E1\|X 48 | 4.325224502 | 1.02E-09 | 7.35E-09 |
| 11385 | ENSMUSG00000035798 | Zdhhc17 | 320150 [Gene Symbol: Zdhhc17] [Locus Tag: ] [Chromosome: 10] [Map Location: 10\|10 | 4.320357455 | 7.93E-15 | 1.28E-13 |
| 8330 | ENSMUSG00000071708 | Gm14680; Sms | 671878 [Gene Symbol: Gm14680] [Locus Tag: ] [Chromosome: X] [Map Location: X A6\| | 4.320236205 | 1.10E-09 | 7.88E-09 |
| 11331 | ENSMUSG00000026333 | Gin1 | 252876 [Gene Symbol: Gin1] [Locus Tag: ] [Chromosome: 1] [Map Location: 1\|1 D] [Des | 4.319363322 | 5.39E-06 | 2.14E-05 |
| 16116 | ENSMUSG00000047242 | Taf9b | 407786 [Gene Symbol: Taf9b] [Locus Tag: ] [Chromosome: X] [Map Location: X\|X D] [De | 4.31352191 | 2.39E-07 | 1.19E-06 |
| 16124 | ENSMUSG00000055069 | Rab39 | 270160 [Gene Symbol: Rab39] [Locus Tag: ] [Chromosome: 9] [Map Location: 9\|9 A5.3] | 4.312726249 | 0.000505 | 0.001393 |
| 4991 | ENSMUSG00000040569 | Slc26a7 | 208890 [Gene Symbol: Slc26a7] [Locus Tag: ] [Chromosome: 4] [Map Location: 4\|4 A1] | 4.312639252 | 6.14E-09 | 3.97E-08 |
| 7303 | ENSMUSG00000066613 | Zfp932 | 69504 [Gene Symbol: Zfp932] [Locus Tag: ] [Chromosome: 5] [Map Location: 5\|5 F] [De | 4.307283897 | 3.34E-09 | 2.24E-08 |
| 9835 | ENSMUSG00000086277 | 4930558K02Rik | 75368 [Gene Symbol: 4930558K02Rik] [Locus Tag: ] [Chromosome: 1] [Map Location: 1 | 4.305947521 | 9.00E-12 | 8.89E-11 |
| 1779 | ENSMUSG00000032913 | Lrig2 | 269473 [Gene Symbol: Lrig2] [Locus Tag: ] [Chromosome: 3] [Map Location: 3\|3 F2.2] [ | 4.303925517 | 2.50E-11 | 2.30E-10 |
| 577 | ENSMUSG00000021816 | Ppp3cb | 19056 [Gene Symbol: Ppp3cb] [Locus Tag: ] [Chromosome: 14] [Map Location: 14\|14 B | 4.303782172 | 4.50E-16 | 8.92E-15 |
| 17480 | ENSMUSG00000027510 | Rbm38 | 56190 [Gene Symbol: Rbm38] [Locus Tag: ] [Chromosome: 2] [Map Location: 2\|2 H3] [D | 4.300228356 | 4.92E-08 | 2.74E-07 |
| 1155 | ENSMUSG00000042197 | Zfp451 | 98403 [Gene Symbol: Zfp451] [Locus Tag: ] [Chromosome: 1] [Map Location: 1 B\|1 12.8 | 4.29749194 | 3.88E-16 | 7.76E-15 |
| 18475 | ENSMUSG00000026112 | Coa5 | 76178 [Gene Symbol: Coa5] [Locus Tag: ] [Chromosome: 1] [Map Location: 1\|1 B] [Desc | 4.295254876 | 5.02E-11 | 4.42E-10 |
| 5242 | ENSMUSG00000025958 | Creb1 | 12912 [Gene Symbol: Creb1] [Locus Tag: ] [Chromosome: 1] [Map Location: 1 C2\|1 32.7 | 4.29514518 | 2.48E-11 | 2.28E-10 |
| 6293 | ENSMUSG00000020115 | Tbk1 | 56480 [Gene Symbol: Tbk1] [Locus Tag: ] [Chromosome: 10] [Map Location: 10\|10 D2] | 4.294501197 | 7.53E-12 | 7.54E-11 |
| 3492 | ENSMUSG00000071014 | Ndufb6 | 230075 [Gene Symbol: Ndufb6] [Locus Tag: ] [Chromosome: 4] [Map Location: 4\|4 A5] | 4.294421086 | 6.83E-08 | 3.72E-07 |
| 6262 | ENSMUSG00000046753 | Ccdc66 | 320234 [Gene Symbol: Ccdc66] [Locus Tag: ] [Chromosome: 14] [Map Location: 14\|14 | 4.291635053 | 3.50E-14 | 5.12E-13 |
| 6396 | ENSMUSG00000032525 | Nktr | 18087 [Gene Symbol: Nktr] [Locus Tag: ] [Chromosome: 9] [Map Location: 9 F4\|9 72.57 | 4.288802028 | 3.68E-09 | 2.45E-08 |
| 9051 | ENSMUSG00000025531 | Chm | 12662 [Gene Symbol: Chm] [Locus Tag: ] [Chromosome: X] [Map Location: X\|X E1] [Desc | 4.28665609 | 2.13E-10 | 1.69E-09 |
| 10031 | ENSMUSG00000046794 | Ppp1r3b | 244416 [Gene Symbol: Ppp1r3b] [Locus Tag: ] [Chromosome: 8] [Map Location: 8\|8 A4 | 4.285948786 | 0.001186 | 0.003046 |
| 8119 | ENSMUSG00000043467 | Zbtb37 | 240869 [Gene Symbol: Zbtb37] [Locus Tag: ] [Chromosome: 1] [Map Location: 1\|1 H2.1 | 4.28310383 | 4.00E-12 | 4.16E-11 |
| 17989 | ENSMUSG00000041380 | Htr2c | 15560 [Gene Symbol: Htr2c] [Locus Tag: ] [Chromosome: X] [Map Location: X 68.46 cM | 4.277307947 | 8.96E-14 | 1.23E-12 |
| 12023 | ENSMUSG00000021241 | Isca2 | 74316 [Gene Symbol: Isca2] [Locus Tag: ] [Chromosome: 12] [Map Location: 12\|12 D3] | 4.271849866 | 2.05E-09 | 1.41E-08 |
| 6472 | ENSMUSG00000020413 | Hus1 | 15574 [Gene Symbol: Hus1] [Locus Tag: ] [Chromosome: 11] [Map Location: 11 A1\|11 5 | 4.26947358 | 2.66E-10 | 2.09E-09 |
| 62 | ENSMUSG00000061665 | Cd2ap | 12488 [Gene Symbol: Cd2ap] [Locus Tag: ] [Chromosome: 17] [Map Location: 17\|17 B3 | 4.267240165 | 2.96E-10 | 2.30E-09 |
| 2911 | ENSMUSG00000022292 | Rrm2b | 382985 [Gene Symbol: Rrm2b] [Locus Tag: ] [Chromosome: 15] [Map Location: 15\|15 E | 4.266152832 | 2.60E-12 | 2.78E-11 |
| 3090 | ENSMUSG00000035125 | Gcfc2 | 330361 [Gene Symbol: Gcfc2] [Locus Tag: ] [Chromosome: 6] [Map Location: 6\|6 C3] [D | 4.262495362 | 9.69E-10 | 6.99E-09 |
| 12712 | ENSMUSG00000022323 | Hrsp12 | 15473 [Gene Symbol: Hrsp12] [Locus Tag: ] [Chromosome: 15] [Map Location: 15\|15 B3 | 4.260697361 | 0.005559 | 0.012258 |
| 12177 | ENSMUSG00000026248 | Mrpl44 | 69163 [Gene Symbol: Mrpl44] [Locus Tag: ] [Chromosome: 1] [Map Location: 1\|1 C4] [D | 4.259125172 | 2.94E-10 | 2.30E-09 |
| 8169 | ENSMUSG00000048280 | Zfp738 | 408068 [Gene Symbol: Zfp738] [Locus Tag: ] [Chromosome: 13] [Map Location: 13\|13 I | 4.254348466 | 1.52E-13 | 2.00E-12 |
| 8898 | ENSMUSG00000028549 | Itgb3bp | 67733 [Gene Symbol: Itgb3bp] [Locus Tag: ] [Chromosome: 4] [Map Location: 4\|4 C6] [ | 4.25303457 | 1.76E-11 | 1.66E-10 |
| 7791 | ENSMUSG00000061313 | Ddhd2 | 72108 [Gene Symbol: Ddhd2] [Locus Tag: ] [Chromosome: 8] [Map Location: 8\|8 A3] [D | 4.252599209 | 1.06E-12 | 1.22E-11 |
| 7998 | ENSMUSG00000030303 | Far2 | 330450 [Gene Symbol: Far2] [Locus Tag: ] [Chromosome: 6] [Map Location: 6\|6 G3] [De | 4.252381006 | 1.17E-14 | 1.84E-13 |
| 4696 | ENSMUSG00000030061 | Uba3 | 22200 [Gene Symbol: Uba3] [Locus Tag: ] [Chromosome: 6] [Map Location: 6\|6 D3] [Des | 4.251540176 | 4.87E-14 | 6.97E-13 |
| 6097 | ENSMUSG00000040649 | Rimklb | 108653 [Gene Symbol: Rimklb] [Locus Tag: ] [Chromosome: 6] [Map Location: 6\|6 F2] [ | 4.250455424 | 1.09E-10 | 9.09E-10 |
| 17260 | ENSMUSG00000032253 | Phip | 83946 [Gene Symbol: Phip] [Locus Tag: ] [Chromosome: 9] [Map Location: 9\|9 E3.1] [De | 4.249963423 | 1.53E-11 | 1.46E-10 |
| 3406 | ENSMUSG00000028879 | Stx12 | 100226 [Gene Symbol: Stx12] [Locus Tag: ] [Chromosome: 4] [Map Location: 4 D2.3\|4 6 | 4.249723674 | 2.02E-15 | 3.57E-14 |
| 6015 | ENSMUSG00000030870 | Ubfd1 | 28018 [Gene Symbol: Ubfd1] [Locus Tag: ] [Chromosome: 7] [Map Location: 7 F3\|7 65.3 | 4.249348565 | 2.81E-15 | 4.84E-14 |
| 9988 | ENSMUSG00000024290 | Rock1 | 19877 [Gene Symbol: Rock1] [Locus Tag: ] [Chromosome: 18] [Map Location: 18\|18 A2 | 4.246190073 | 8.59E-08 | 4.61E-07 |
| 7094 | ENSMUSG00000004609 | Cd33 | 12489 [Gene Symbol: Cd33] [Locus Tag: ] [Chromosome: 7] [Map Location: 7 B4\|7 28.2 | 4.244754592 | 0.001595 | 0.003975 |
| 18602 | ENSMUSG00000025016 | Tm9sf3 | 107358 [Gene Symbol: Tm9sf3] [Locus Tag: ] [Chromosome: 19] [Map Location: 19\|19 | 4.243291967 | 2.47E-15 | 4.29E-14 |
| 4636 | ENSMUSG00000026159 | Agfg1 | 15463 [Gene Symbol: Agfg1] [Locus Tag: ] [Chromosome: 1] [Map Location: 1\|1 C5] [De | 4.243131287 | 9.95E-16 | 1.87E-14 |
| 3043 | ENSMUSG00000029253 | Cenpc1 | 12617 [Gene Symbol: Cenpc1] [Locus Tag: ] [Chromosome: 5] [Map Location: 5\|5 E2-E5 | 4.241132243 | 1.60E-13 | 2.10E-12 |
| 8394 | ENSMUSG00000020914 | Top2a | 21973 [Gene Symbol: Top2a] [Locus Tag: ] [Chromosome: 11] [Map Location: 11 D\|11 6 | 4.239452441 | 5.26E-09 | 3.43E-08 |
| 6242 | ENSMUSG00000035851 | Ythdc1 | 231386 [Gene Symbol: Ythdc1] [Locus Tag: ] [Chromosome: 5] [Map Location: 5\|5 E1] [ | 4.236291689 | 3.93E-12 | 4.09E-11 |
| 7213 | ENSMUSG00000019992 | Mtfr2 | 71804 [Gene Symbol: Mtfr2] [Locus Tag: ] [Chromosome: 10] [Map Location: 10\|10 A3] | 4.231179009 | 0.000394 | 0.001111 |
| 11526 | ENSMUSG00000049858 | Suox | 211389 [Gene Symbol: Suox] [Locus Tag: ] [Chromosome: 10] [Map Location: 10\|10 D3 | 4.228263099 | 1.86E-08 | 1.11E-07 |
| 7350 | ENSMUSG00000025921 | Rdh10 | 98711 [Gene Symbol: Rdh10] [Locus Tag: ] [Chromosome: 1] [Map Location: 1 A3\|1 4.9 | 4.227626095 | 1.37E-12 | 1.53E-11 |
| 14080 | ENSMUSG00000069089 | Cdk7 | 12572 [Gene Symbol: Cdk7] [Locus Tag: ] [Chromosome: 13] [Map Location: 13 D1\|13 5 | 4.224475751 | 4.62E-14 | 6.65E-13 |
| 5129 | ENSMUSG00000040195 | Tmem194 | 210035 [Gene Symbol: Tmem194] [Locus Tag: ] [Chromosome: 10] [Map Location: 10\|1 | 4.224337962 | 2.06E-07 | 1.04E-06 |
| 18433 | ENSMUSG00000043631 | Ecm2 | 407800 [Gene Symbol: Ecm2] [Locus Tag: ] [Chromosome: 13] [Map Location: 13\|13 B1 | 4.218666364 | 0.000469 | 0.0013 |
| 13665 | ENSMUSG00000087370 | Tmem170b | 621976 [Gene Symbol: Tmem170b] [Locus Tag: ] [Chromosome: 13] [Map Location: 13 | 4.218520158 | 3.68E-16 | 7.41E-15 |
| 6270 | ENSMUSG00000034210 | Efcab14 | 230648 [Gene Symbol: Efcab14] [Locus Tag: ] [Chromosome: 4] [Map Location: 4\|4 D1] | 4.217755469 | 1.99E-12 | 2.18E-11 |
| 11010 | ENSMUSG00000031333 | Abcb7 | 11306 [Gene Symbol: Abcb7] [Locus Tag: ] [Chromosome: X] [Map Location: X 46.58 cM | 4.214781784 | 1.28E-13 | 1.71E-12 |
| 127 | ENSMUSG00000024486 | Hbegf | 15200 [Gene Symbol: Hbegf] [Locus Tag: ] [Chromosome: 18] [Map Location: 18 B2\|18 | 4.213729363 | 8.91E-12 | 8.81E-11 |
| 2332 | ENSMUSG00000014980 | Tsen15 | 66637 [Gene Symbol: Tsen15] [Locus Tag: ] [Chromosome: 1] [Map Location: 1\|1 G3] [D | 4.212746111 | 7.35E-05 | 0.000237 |
| 12566 | ENSMUSG00000064368 | ND6 | 17722 [Gene Symbol: ND6] [Locus Tag: ] [Chromosome: MT] [Map Location: ] [Descripti | 4.212486509 | 3.07E-14 | 4.55E-13 |
| 5988 | ENSMUSG00000025810 | Nrp1 | 18186 [Gene Symbol: Nrp1] [Locus Tag: ] [Chromosome: 8] [Map Location: 8 75.78 cM\| | 4.212130616 | 1.68E-11 | 1.59E-10 |
| 8586 | ENSMUSG00000031202 | Rab39b | 67790 [Gene Symbol: Rab39b] [Locus Tag: ] [Chromosome: X] [Map Location: X 38.26 c | 4.211681963 | 5.80E-12 | 5.91E-11 |
| 1301 | ENSMUSG00000044288 | Cnr1 | 12801 [Gene Symbol: Cnr1] [Locus Tag: ] [Chromosome: 4] [Map Location: 4 A5\|4 16.2 | 4.209288389 | 1.29E-15 | 2.37E-14 |
| 8608 | ENSMUSG00000000568 | Hnrnpd | 11991 [Gene Symbol: Hnrnpd] [Locus Tag: ] [Chromosome: 5] [Map Location: 5\|5 E3] [D | 4.208830334 | 1.10E-12 | 1.25E-11 |
| 13179 | ENSMUSG00000026004 | Kansl1l | 68691 [Gene Symbol: Kansl1l] [Locus Tag: ] [Chromosome: 1] [Map Location: 1\|1 C3] [D | 4.206926272 | 4.94E-12 | 5.07E-11 |
| 18368 | ENSMUSG00000028518 | Prkaa2 | 108079 [Gene Symbol: Prkaa2] [Locus Tag: ] [Chromosome: 4] [Map Location: 4\|4 C6] [ | 4.205441336 | 2.99E-15 | 5.11E-14 |
| 7942 | ENSMUSG00000026434 | Nucks1 | 98415 [Gene Symbol: Nucks1] [Locus Tag: ] [Chromosome: 1] [Map Location: 1\|1 E4] [D | 4.203913439 | 1.63E-15 | 2.93E-14 |
| 3241 | ENSMUSG00000021716 | Srek1ip1 | 67288 [Gene Symbol: Srek1ip1] [Locus Tag: ] [Chromosome: 13] [Map Location: 13\|13 | 4.198584509 | 4.66E-11 | 4.11E-10 |
| 2649 | ENSMUSG00000022663 | Atg3 | 67841 [Gene Symbol: Atg3] [Locus Tag: ] [Chromosome: 16] [Map Location: 16\|16 B5] [ | 4.196344818 | 1.85E-14 | 2.83E-13 |
| 17065 | ENSMUSG00000026319 | 2310035C23Rik | 227446 [Gene Symbol: 2310035C23Rik] [Locus Tag: ] [Chromosome: 1] [Map Location: | 4.196043004 | 1.11E-11 | 1.08E-10 |
| 16264 | ENSMUSG00000058900 | Rsl1 | 380855 [Gene Symbol: Rsl1] [Locus Tag: ] [Chromosome: 13] [Map Location: 13 B3\|13 | 4.194861265 | 0.000949 | 0.002482 |
| 172 | ENSMUSG00000031519 | Asb5 | 76294 [Gene Symbol: Asb5] [Locus Tag: ] [Chromosome: 8] [Map Location: 8\|8 B3.1] [D | 4.193343503 | 0.000482 | 0.001332 |
| 16098 | ENSMUSG00000021302 | Ggps1 | 14593 [Gene Symbol: Ggps1] [Locus Tag: ] [Chromosome: 13] [Map Location: 13 A1\|13 | 4.18577275 | 3.04E-13 | 3.77E-12 |
| 8379 | ENSMUSG00000025724 | Sec11a | 56529 [Gene Symbol: Sec11a] [Locus Tag: ] [Chromosome: 7] [Map Location: 7\|7 D3] [D | 4.185199028 | 1.08E-11 | 1.05E-10 |
| 13134 | ENSMUSG00000026361 | Cdc73 | 214498 [Gene Symbol: Cdc73] [Locus Tag: ] [Chromosome: 1] [Map Location: 1 F\|1 62.5 | 4.185066979 | 1.63E-15 | 2.92E-14 |
| 1476 | ENSMUSG00000057069 | Ero1lb | 67475 [Gene Symbol: Ero1lb] [Locus Tag: ] [Chromosome: 13] [Map Location: 13\|13 A1 | 4.18298166 | 1.29E-09 | 9.12E-09 |
| 4670 | ENSMUSG00000042821 | Snai1 | 20613 [Gene Symbol: Snai1] [Locus Tag: ] [Chromosome: 2] [Map Location: 2 H3\|2 87.3 | 4.181378425 | 0.000558 | 0.001527 |
| 17231 | ENSMUSG00000025785 | Exosc7 | 66446 [Gene Symbol: Exosc7] [Locus Tag: ] [Chromosome: 9] [Map Location: 9\|9 F4] [D | 4.181258981 | 3.52E-10 | 2.70E-09 |
| 18427 | ENSMUSG00000031210 | Gpr165 | 76206 [Gene Symbol: Gpr165] [Locus Tag: ] [Chromosome: X] [Map Location: X\|X C1] [D | 4.179621553 | 1.65E-05 | 6.04E-05 |
| 5972 | ENSMUSG00000021496 | Pcbd2 | 72562 [Gene Symbol: Pcbd2] [Locus Tag: ] [Chromosome: 13] [Map Location: 13\|13 B2 | 4.173541268 | 5.88E-06 | 2.32E-05 |
| 10959 | ENSMUSG00000027384 | Ndufaf5 | 69487 [Gene Symbol: Ndufaf5] [Locus Tag: ] [Chromosome: 2] [Map Location: 2\|2 G3] [ | 4.172995743 | 5.78E-11 | 5.03E-10 |
| 14057 | ENSMUSG00000027804 | Ppid | 67738 [Gene Symbol: Ppid] [Locus Tag: ] [Chromosome: 3] [Map Location: 3\|3 F1] [Des | 4.171775478 | 1.16E-11 | 1.13E-10 |
| 15252 | ENSMUSG00000074345 | Tnfaip8l3 | 244882 [Gene Symbol: Tnfaip8l3] [Locus Tag: ] [Chromosome: 9] [Map Location: 9\|9 A5 | 4.169926254 | 2.75E-06 | 1.15E-05 |
| 15933 | ENSMUSG00000035704 | Alg8 | 381903 [Gene Symbol: Alg8] [Locus Tag: ] [Chromosome: 7] [Map Location: 7\|7 E1] [De | 4.166229797 | 2.39E-06 | 1.01E-05 |
| 17105 | ENSMUSG00000029676 | Pot1a | 101185 [Gene Symbol: Pot1a] [Locus Tag: ] [Chromosome: 6] [Map Location: 6\|6 A3.1] | 4.16437178 | 6.88E-11 | 5.92E-10 |

| | | | | | | |
|---|---|---|---|---|---|---|
| 14759 | ENSMUSG00000014077 | Chp1 | 56398 [Gene Symbol: Chp1] [Locus Tag: ] [Chromosome: 2] [Map Location: 2\|2 E5] [Des | 4.163153317 | 1.67E-13 | 2.17E-12 |
| 8037 | ENSMUSG00000007655 | Cav1 | 12389 [Gene Symbol: Cav1] [Locus Tag: ] [Chromosome: 6] [Map Location: 6\|6 A2] [Des | 4.159485019 | 2.45E-11 | 2.26E-10 |
| 15930 | ENSMUSG00000034345 | Gtf2h5 | 66467 [Gene Symbol: Gtf2h5] [Locus Tag: ] [Chromosome: 17] [Map Location: 17 A1\|17 | 4.157764458 | 5.53E-14 | 7.85E-13 |
| 12763 | ENSMUSG00000035992 | Fnip1 | 216742 [Gene Symbol: Fnip1] [Locus Tag: ] [Chromosome: 11] [Map Location: 11\|11 B1 | 4.150801865 | 7.38E-15 | 1.20E-13 |
| 7976 | ENSMUSG00000037896 | Rcor1 | 217864 [Gene Symbol: Rcor1] [Locus Tag: ] [Chromosome: 12] [Map Location: 12 F1\|12 | 4.150349971 | 3.65E-10 | 2.80E-09 |
| 13907 | ENSMUSG00000043881 | Kbtbd7 | 211255 [Gene Symbol: Kbtbd7] [Locus Tag: ] [Chromosome: 14] [Map Location: 14\|14 | 4.148149297 | 3.39E-14 | 4.98E-13 |
| 1046 | ENSMUSG00000022124 | Fbxl3 | 50789 [Gene Symbol: Fbxl3] [Locus Tag: ] [Chromosome: 14] [Map Location: 14\|14 E2.1 | 4.146525423 | 1.24E-14 | 1.93E-13 |
| 3928 | ENSMUSG00000041935 | AW549877 | 106064 [Gene Symbol: AW549877] [Locus Tag: ] [Chromosome: 15] [Map Location: 15 | 4.137625571 | 1.05E-13 | 1.42E-12 |
| 8102 | ENSMUSG00000022858 | Tra2b | 20462 [Gene Symbol: Tra2b] [Locus Tag: ] [Chromosome: 16] [Map Location: 16 B1\|16 | 4.136642422 | 9.77E-14 | 1.33E-12 |
| 3203 | ENSMUSG00000033585 | Ndn | 17984 [Gene Symbol: Ndn] [Locus Tag: ] [Chromosome: 7] [Map Location: 7 34.36 cM\|7 | 4.134452728 | 4.44E-10 | 3.36E-09 |
| 165 | ENSMUSG00000023882 | Zfp54 | 22712 [Gene Symbol: Zfp54] [Locus Tag: ] [Chromosome: 17] [Map Location: 17 A3.2\|1 | 4.127314127 | 0.00013 | 0.000401 |
| 17142 | ENSMUSG00000016495 | Plgrkt | 67759 [Gene Symbol: Plgrkt] [Locus Tag: ] [Chromosome: 19] [Map Location: 19\|19 C1] | 4.126895856 | 9.17E-09 | 5.79E-08 |
| 4623 | ENSMUSG00000024986 | Hhex | 15242 [Gene Symbol: Hhex] [Locus Tag: ] [Chromosome: 19] [Map Location: 19 32.28 c | 4.125110943 | 2.57E-05 | 9.08E-05 |
| 2531 | ENSMUSG00000023075 | Akirin1 | 68050 [Gene Symbol: Akirin1] [Locus Tag: ] [Chromosome: 4] [Map Location: 4\|4 D1] [D | 4.120680254 | 1.70E-12 | 1.88E-11 |
| 10647 | ENSMUSG00000018287 | Spag7 | 216873 [Gene Symbol: Spag7] [Locus Tag: ] [Chromosome: 11] [Map Location: 11 B4\|11 | 4.111524124 | 1.46E-09 | 1.03E-08 |
| 15414 | ENSMUSG00000078861 | Zfp931 | 353208 [Gene Symbol: Zfp931] [Locus Tag: ] [Chromosome: 2] [Map Location: 2\|2 H4] [ | 4.109682713 | 0.00146 | 0.003671 |
| 10656 | ENSMUSG00000039789 | Zfp597 | 71063 [Gene Symbol: Zfp597] [Locus Tag: ] [Chromosome: 16] [Map Location: 16\|16 A1 | 4.109039802 | 1.56E-14 | 2.41E-13 |
| 4906 | ENSMUSG00000031381 | Piga | 18700 [Gene Symbol: Piga] [Locus Tag: ] [Chromosome: X] [Map Location: X 76.49 cM\|X | 4.106440764 | 3.53E-06 | 1.44E-05 |
| 4871 | ENSMUSG00000022629 | Kif21a | 16564 [Gene Symbol: Kif21a] [Locus Tag: ] [Chromosome: 15] [Map Location: 15 E3\|15 | 4.104675987 | 1.03E-11 | 1.01E-10 |
| 8023 | ENSMUSG00000026511 | Srp9 | 27058 [Gene Symbol: Srp9] [Locus Tag: ] [Chromosome: 1] [Map Location: 1\|1 H5] [Des | 4.101684063 | 1.22E-10 | 1.01E-09 |
| 8546 | ENSMUSG00000038132 | Rbm24 | 666794 [Gene Symbol: Rbm24] [Locus Tag: ] [Chromosome: 13] [Map Location: 13\|13 A | 4.101325818 | 1.45E-08 | 8.82E-08 |
| 12545 | ENSMUSG00000031843 | Mphosph6 | 68533 [Gene Symbol: Mphosph6] [Locus Tag: ] [Chromosome: 8] [Map Location: 8\|8 E1 | 4.10093714 | 1.07E-07 | 5.69E-07 |
| 16531 | ENSMUSG00000068130 | Zfp442 | 668923 [Gene Symbol: Zfp442] [Locus Tag: ] [Chromosome: 2] [Map Location: 2 G3\|2] [ | 4.099848529 | 9.33E-08 | 4.98E-07 |
| 8621 | ENSMUSG00000024726 | Carnmt1 | 67383 [Gene Symbol: Carnmt1] [Locus Tag: ] [Chromosome: 19] [Map Location: 19\|19 | 4.099663539 | 6.56E-07 | 3.05E-06 |
| 13525 | ENSMUSG00000073460 | Pnldc1 | 240023 [Gene Symbol: Pnldc1] [Locus Tag: ] [Chromosome: 17] [Map Location: 17\|17 A | 4.097776606 | 0.000301 | 0.000868 |
| 16070 | ENSMUSG00000034484 | Snx2 | 67804 [Gene Symbol: Snx2] [Locus Tag: ] [Chromosome: 18] [Map Location: 18\|18 D1] | 4.093770435 | 1.63E-09 | 1.14E-08 |
| 11347 | ENSMUSG00000036438 | Calm2 | 12314 [Gene Symbol: Calm2] [Locus Tag: ] [Chromosome: 17] [Map Location: 17\|17 E4 | 4.08892456 | 8.28E-10 | 6.05E-09 |
| 14813 | ENSMUSG00000052430 | Bmpr1b | 12167 [Gene Symbol: Bmpr1b] [Locus Tag: ] [Chromosome: 3] [Map Location: 3\|3 H1] [ | 4.086749847 | 2.36E-13 | 2.99E-12 |
| 14838 | ENSMUSG00000028609 | Magoh | 17149 [Gene Symbol: Magoh] [Locus Tag: ] [Chromosome: 4] [Map Location: 4 C7\|4 50. | 4.086415935 | 9.97E-12 | 9.80E-11 |
| 10676 | ENSMUSG00000035004 | Igsf6 | 80719 [Gene Symbol: Igsf6] [Locus Tag: ] [Chromosome: 7] [Map Location: 7\|7 F2-F3] [D | 4.084071574 | 0.000142 | 0.000435 |
| 13047 | ENSMUSG00000029917 | C130060K24Rik | 243407 [Gene Symbol: C130060K24Rik] [Locus Tag: ] [Chromosome: 6] [Map Location: | 4.08152971 | 0.000416 | 0.001167 |
| 4881 | ENSMUSG00000055723 | Rras2 | 66922 [Gene Symbol: Rras2] [Locus Tag: ] [Chromosome: 7] [Map Location: 7\|7 F2] [Des | 4.080044837 | 4.60E-14 | 6.64E-13 |
| 14121 | ENSMUSG00000027823 | Gmps | 229363 [Gene Symbol: Gmps] [Locus Tag: ] [Chromosome: 3] [Map Location: 3\|3 E1] [D | 4.07967772 | 2.34E-11 | 2.16E-10 |
| 7244 | ENSMUSG00000047909 | Ankrd16 | 320816 [Gene Symbol: Ankrd16] [Locus Tag: ] [Chromosome: 2] [Map Location: 2\|2 A1] | 4.077880522 | 7.03E-12 | 7.09E-11 |
| 10016 | ENSMUSG00000038623 | Tm6sf1 | 107769 [Gene Symbol: Tm6sf1] [Locus Tag: ] [Chromosome: 7] [Map Location: 7\|7 D1] | 4.074254922 | 6.90E-14 | 9.63E-13 |
| 1113 | ENSMUSG00000027171 | Prrg4 | 228413 [Gene Symbol: Prrg4] [Locus Tag: ] [Chromosome: 2] [Map Location: 2\|2 E2] [D | 4.074137192 | 0.000911 | 0.002389 |
| 16858 | ENSMUSG00000020300 | Cpeb4 | 67579 [Gene Symbol: Cpeb4] [Locus Tag: ] [Chromosome: 11] [Map Location: 11\|11 A4 | 4.074098579 | 7.39E-11 | 6.33E-10 |
| 12629 | ENSMUSG00000033386 | Frrs1 | 20321 [Gene Symbol: Frrs1] [Locus Tag: ] [Chromosome: 3] [Map Location: 3\|3 G1] [Des | 4.073076937 | 5.44E-07 | 2.57E-06 |
| 10372 | ENSMUSG00000035678 | Tnfsf9 | 21950 [Gene Symbol: Tnfsf9] [Locus Tag: ] [Chromosome: 17] [Map Location: 17\|17 D] | 4.072181815 | 7.46E-08 | 4.03E-07 |
| 238 | ENSMUSG00000024597 | Slc12a2 | 20496 [Gene Symbol: Slc12a2] [Locus Tag: ] [Chromosome: 18] [Map Location: 18 D3\|1 | 4.065363086 | 3.00E-14 | 4.46E-13 |
| 5156 | ENSMUSG00000033382 | Trappc8 | 75964 [Gene Symbol: Trappc8] [Locus Tag: ] [Chromosome: 18] [Map Location: 18\|18 A | 4.064077101 | 2.68E-11 | 2.45E-10 |
| 3987 | ENSMUSG00000070287 | Slc35g2 | 245020 [Gene Symbol: Slc35g2] [Locus Tag: ] [Chromosome: 9] [Map Location: 9\|9 E3.3 | 4.058767611 | 1.12E-05 | 4.22E-05 |
| 6241 | ENSMUSG00000048281 | Dleu7 | 239133 [Gene Symbol: Dleu7] [Locus Tag: ] [Chromosome: 14] [Map Location: 14\|14 D | 4.058576032 | 1.98E-08 | 1.18E-07 |
| 15190 | ENSMUSG00000044501 | Zfp758 | 224598 [Gene Symbol: Zfp758] [Locus Tag: ] [Chromosome: 17] [Map Location: 17\|17 A | 4.056935798 | 1.70E-07 | 8.70E-07 |
| 10918 | ENSMUSG00000027160 | Ccdc34 | 68201 [Gene Symbol: Ccdc34] [Locus Tag: ] [Chromosome: 2] [Map Location: 2\|2 E3] [D | 4.056866707 | 2.65E-13 | 3.33E-12 |
| 17706 | ENSMUSG00000026484 | Rnf2 | 19821 [Gene Symbol: Rnf2] [Locus Tag: ] [Chromosome: 1] [Map Location: 1\|1 G2] [Des | 4.055063658 | 4.74E-14 | 6.80E-13 |
| 5643 | ENSMUSG00000028273 | Pdlim5 | 56376 [Gene Symbol: Pdlim5] [Locus Tag: ] [Chromosome: 3] [Map Location: 3\|3 H3] [D | 4.054502405 | 5.72E-08 | 3.15E-07 |
| 12631 | ENSMUSG00000058013 | Sept11 | 52398 [Gene Symbol: Sept11] [Locus Tag: ] [Chromosome: 5] [Map Location: 5 E2\|5 47. | 4.051891069 | 1.07E-14 | 1.68E-13 |
| 10587 | ENSMUSG00000003992 | Ssbp2 | 66970 [Gene Symbol: Ssbp2] [Locus Tag: ] [Chromosome: 13] [Map Location: 13\|13 C3] | 4.050850378 | 4.77E-15 | 7.93E-14 |
| 4549 | ENSMUSG00000030660 | Pik3c2a | 18704 [Gene Symbol: Pik3c2a] [Locus Tag: ] [Chromosome: 7] [Map Location: 7 F1\|7 61 | 4.050798907 | 1.88E-13 | 2.42E-12 |
| 9639 | ENSMUSG00000024068 | Spast | 50850 [Gene Symbol: Spast] [Locus Tag: ] [Chromosome: 17] [Map Location: 17\|17 E3] | 4.047379013 | 3.37E-15 | 5.72E-14 |
| 7848 | ENSMUSG00000058258 | Idi1 | 319554 [Gene Symbol: Idi1] [Locus Tag: ] [Chromosome: 13] [Map Location: 13\|13 A1] | 4.047050921 | 0.002276 | 0.005476 |
| 4590 | ENSMUSG00000008226 | Scrn3 | 74616 [Gene Symbol: Scrn3] [Locus Tag: ] [Chromosome: 2] [Map Location: 2\|2 C3] [De | 4.046440762 | 4.00E-06 | 1.62E-05 |
| 12343 | ENSMUSG00000005078 | Jkamp | 104771 [Gene Symbol: Jkamp] [Locus Tag: ] [Chromosome: 12] [Map Location: 12\|12 C | 4.042057122 | 8.20E-13 | 9.61E-12 |
| 12006 | ENSMUSG00000060681 | Slc9a6 | 236794 [Gene Symbol: Slc9a6] [Locus Tag: ] [Chromosome: X] [Map Location: X\|X A5] [ | 4.040220661 | 4.59E-14 | 6.63E-13 |
| 11124 | ENSMUSG00000035847 | Ids | 15931 [Gene Symbol: Ids] [Locus Tag: ] [Chromosome: X] [Map Location: X A7.1\|X 36.0 | 4.039901076 | 1.26E-15 | 2.32E-14 |
| 14399 | ENSMUSG00000063415 | Cyp26b1 | 232174 [Gene Symbol: Cyp26b1] [Locus Tag: ] [Chromosome: 6] [Map Location: 6\|6 C3 | 4.039499354 | 8.99E-14 | 1.23E-12 |
| 10320 | ENSMUSG00000000686 | Abhd15 | 67477 [Gene Symbol: Abhd15] [Locus Tag: ] [Chromosome: 11] [Map Location: 11\|11 B | 4.039031579 | 6.71E-15 | 1.10E-13 |
| 10623 | ENSMUSG00000059834 | Sclt1 | 67161 [Gene Symbol: Sclt1] [Locus Tag: ] [Chromosome: 3] [Map Location: 3\|3 C] [Desc | 4.035796456 | 5.58E-09 | 3.63E-08 |
| 2971 | ENSMUSG00000066735 | Vkorc1l1 | 69568 [Gene Symbol: Vkorc1l1] [Locus Tag: ] [Chromosome: 5] [Map Location: 5\|5 G1.3 | 4.03198435 | 1.71E-14 | 2.63E-13 |
| 14478 | ENSMUSG00000001627 | Ifrd1 | 15982 [Gene Symbol: Ifrd1] [Locus Tag: ] [Chromosome: 12] [Map Location: 12 B1\|12 1 | 4.030295077 | 1.25E-13 | 1.67E-12 |
| 3138 | ENSMUSG00000066798 | Zbtb6 | 241322 [Gene Symbol: Zbtb6] [Locus Tag: ] [Chromosome: 2] [Map Location: 2\|2 B] [De | 4.026432041 | 1.02E-14 | 1.61E-13 |
| 11786 | ENSMUSG00000027276 | Jag1 | 16449 [Gene Symbol: Jag1] [Locus Tag: ] [Chromosome: 2] [Map Location: 2 F3\|2 67.73 | 4.025257849 | 4.88E-08 | 2.72E-07 |
| 7734 | ENSMUSG00000030077 | Chl1 | 12661 [Gene Symbol: Chl1] [Locus Tag: ] [Chromosome: 6] [Map Location: 6\|6 E1] [Desc | 4.023069851 | 1.80E-15 | 3.21E-14 |
| 37 | ENSMUSG00000029629 | Phf14 | 75725 [Gene Symbol: Phf14] [Locus Tag: ] [Chromosome: 6] [Map Location: 6\|6 A2] [De | 4.022683771 | 1.00E-11 | 9.82E-11 |
| 13154 | ENSMUSG00000047539 | Fbxo28 | 67948 [Gene Symbol: Fbxo28] [Locus Tag: ] [Chromosome: 1] [Map Location: 1 H5\|1 84 | 4.019890477 | 7.60E-11 | 6.50E-10 |
| 9521 | ENSMUSG00000025982 | Sf3b1 | 81898 [Gene Symbol: Sf3b1] [Locus Tag: ] [Chromosome: 1] [Map Location: 1 C1.2\|1 27 | 4.019096232 | 2.62E-08 | 1.53E-07 |
| 12534 | ENSMUSG00000038344 | Txlng | 353170 [Gene Symbol: Txlng] [Locus Tag: ] [Chromosome: X] [Map Location: X\|X F4] [De | 4.018953261 | 1.11E-07 | 5.88E-07 |
| 6932 | ENSMUSG00000026743 | Mllt10 | 17354 [Gene Symbol: Mllt10] [Locus Tag: ] [Chromosome: 2] [Map Location: 2\|2 A2] [D | 4.018142728 | 8.72E-15 | 1.40E-13 |
| 14804 | ENSMUSG00000074579 | Lekr1 | 624866 [Gene Symbol: Lekr1] [Locus Tag: ] [Chromosome: 3] [Map Location: 3\|3 E1] [D | 4.015787988 | 2.93E-07 | 1.44E-06 |
| 9365 | ENSMUSG00000020521 | Rnft1 | 76892 [Gene Symbol: Rnft1] [Locus Tag: ] [Chromosome: 11] [Map Location: 11\|11 C] [ | 4.01470994 | 4.59E-07 | 2.19E-06 |
| 9844 | ENSMUSG00000048100 | Taf13 | 99730 [Gene Symbol: Taf13] [Locus Tag: ] [Chromosome: 3] [Map Location: 3\|3 F3] [Des | 4.012075515 | 3.98E-09 | 2.64E-08 |
| 5577 | ENSMUSG00000030562 | Nox4 | 50490 [Gene Symbol: Nox4] [Locus Tag: ] [Chromosome: 7] [Map Location: 7\|7 D3] [Des | 4.009547832 | 9.67E-07 | 4.37E-06 |
| 7087 | ENSMUSG00000021890 | Eaf1 | 74427 [Gene Symbol: Eaf1] [Locus Tag: ] [Chromosome: 14] [Map Location: 14\|14 B] [D | 4.00885833 | 1.87E-12 | 2.06E-11 |
| 13386 | ENSMUSG00000033446 | Lpar6 | 67168 [Gene Symbol: Lpar6] [Locus Tag: ] [Chromosome: 14] [Map Location: 14\|14 D3] | 4.008231703 | 3.11E-05 | 0.000108 |
| 10400 | ENSMUSG00000020612 | Prkar1a | 19084 [Gene Symbol: Prkar1a] [Locus Tag: ] [Chromosome: 11] [Map Location: 11 E1\|1 | 4.005651317 | 3.13E-15 | 5.35E-14 |
| 17601 | ENSMUSG00000049047 | Armcx3 | 71703 [Gene Symbol: Armcx3] [Locus Tag: ] [Chromosome: X] [Map Location: X\|X E3] [D | 3.998919448 | 1.43E-08 | 8.71E-08 |
| 4382 | ENSMUSG00000074867 | Zfp808 | 630579 [Gene Symbol: Zfp808] [Locus Tag: ] [Chromosome: 13] [Map Location: 13 B3\|1 | 3.991538958 | 0.001338 | 0.003394 |
| 570 | ENSMUSG00000032336 | Nptn | 20320 [Gene Symbol: Nptn] [Locus Tag: ] [Chromosome: 9] [Map Location: 9\|9 B] [Desc | 3.991052509 | 2.87E-12 | 3.05E-11 |
| 11437 | ENSMUSG00000023235 | Ccl25 | 20300 [Gene Symbol: Ccl25] [Locus Tag: ] [Chromosome: 8] [Map Location: 8 A1.1\|8 2. | 3.99051169 | 6.13E-10 | 4.56E-09 |
| 11313 | ENSMUSG00000036291 | Ap5m1 | 74385 [Gene Symbol: Ap5m1] [Locus Tag: ] [Chromosome: 14] [Map Location: 14\|14 C1 | 3.988512593 | 3.10E-09 | 2.09E-08 |
| 3780 | ENSMUSG00000027531 | Impa1 | 55980 [Gene Symbol: Impa1] [Locus Tag: ] [Chromosome: 3] [Map Location: 3\|3 A1] [De | 3.987421978 | 1.18E-11 | 1.15E-10 |
| 7661 | ENSMUSG00000009734 | Pou6f2 | 218030 [Gene Symbol: Pou6f2] [Locus Tag: ] [Chromosome: 13] [Map Location: 13\|13 A | 3.986114568 | 2.00E-08 | 1.19E-07 |
| 10360 | ENSMUSG00000028127 | Abcd3 | 19299 [Gene Symbol: Abcd3] [Locus Tag: ] [Chromosome: 3] [Map Location: 3 52.94 cM | 3.985598907 | 1.86E-10 | 1.50E-09 |

| | | | | | | |
|---|---|---|---|---|---|---|
| 14444 | ENSMUSG00000024498 | Tcerg1 | 56070 [Gene Symbol: Tcerg1] [Locus Tag: ] [Chromosome: 18] [Map Location: 18\|18 B3 | 3.985145582 | 3.81E-09 | 2.53E-08 |
| 3058 | ENSMUSG00000028114 | Mettl14 | 210529 [Gene Symbol: Mettl14] [Locus Tag: ] [Chromosome: 3] [Map Location: 3\|3 G1] | 3.978371 | 8.52E-08 | 4.57E-07 |
| 17887 | ENSMUSG00000021379 | Id4 | 15904 [Gene Symbol: Id4] [Locus Tag: ] [Chromosome: 13] [Map Location: 13 24.71 cM | 3.971547853 | 6.63E-07 | 3.08E-06 |
| 14820 | ENSMUSG00000070336 | Fbxo47 | 72973 [Gene Symbol: Fbxo47] [Locus Tag: ] [Chromosome: 11] [Map Location: 11\|11 D | 3.970566608 | 0.025091 | 0.047209 |
| 7754 | ENSMUSG00000027206 | Cops2 | 12848 [Gene Symbol: Cops2] [Locus Tag: ] [Chromosome: 2] [Map Location: 2 F1\|2 61. | 3.968651692 | 4.75E-13 | 5.75E-12 |
| 11370 | ENSMUSG00000039178 | Tbc1d19 | 67249 [Gene Symbol: Tbc1d19] [Locus Tag: ] [Chromosome: 5] [Map Location: 5\|5 C1] | 3.968214729 | 1.30E-09 | 9.23E-09 |
| 13802 | ENSMUSG00000028034 | Fubp1 | 51886 [Gene Symbol: Fubp1] [Locus Tag: ] [Chromosome: 3] [Map Location: 3 H3\|3 77. | 3.967333035 | 1.75E-10 | 1.41E-09 |
| 2300 | ENSMUSG00000034848 | Ttc21b | 73668 [Gene Symbol: Ttc21b] [Locus Tag: ] [Chromosome: 2] [Map Location: 2\|2 C1.3] | 3.966500142 | 1.89E-08 | 1.13E-07 |
| 15240 | ENSMUSG00000050549 | 5730508B09Rik | 70617 [Gene Symbol: 5730508B09Rik] [Locus Tag: ] [Chromosome: 3] [Map Location: 3 | 3.964016215 | 3.36E-10 | 2.59E-09 |
| 766 | ENSMUSG00000029422 | Rsrc2 | 208606 [Gene Symbol: Rsrc2] [Locus Tag: ] [Chromosome: 5] [Map Location: 5\|5 F] [De | 3.963740464 | 4.97E-09 | 3.25E-08 |
| 649 | ENSMUSG00000005696 | Sh2d1a | 20400 [Gene Symbol: Sh2d1a] [Locus Tag: ] [Chromosome: X] [Map Location: X\|X A5] [D | 3.963234433 | 1.09E-08 | 6.79E-08 |
| 11885 | ENSMUSG00000073987 | Ggh | 14590 [Gene Symbol: Ggh] [Locus Tag: ] [Chromosome: 4] [Map Location: 4\|4 A3] [Desc | 3.963029205 | 1.05E-07 | 5.56E-07 |
| 524 | ENSMUSG00000040006 | Ginm1 | 215751 [Gene Symbol: Ginm1] [Locus Tag: ] [Chromosome: 10] [Map Location: 10\|10 A | 3.96238446 | 1.24E-09 | 8.80E-09 |
| 16334 | ENSMUSG00000045767 | B230219D22Rik | 78521 [Gene Symbol: B230219D22Rik] [Locus Tag: ] [Chromosome: 13] [Map Location: | 3.962052692 | 2.33E-14 | 3.51E-13 |
| 17122 | ENSMUSG00000032898 | Fbxo21 | 231670 [Gene Symbol: Fbxo21] [Locus Tag: ] [Chromosome: 5] [Map Location: 5\|5 F] [D | 3.960315276 | 5.86E-14 | 8.29E-13 |
| 2778 | ENSMUSG00000036902 | Neto2 | 74513 [Gene Symbol: Neto2] [Locus Tag: ] [Chromosome: 8] [Map Location: 8\|8 C4] [De | 3.959474097 | 1.37E-14 | 2.13E-13 |
| 5138 | ENSMUSG00000003161 | Sri | 109552 [Gene Symbol: Sri] [Locus Tag: ] [Chromosome: 5] [Map Location: 5 A1-h\|5 3.3 | 3.957978118 | 4.51E-08 | 2.54E-07 |
| 1561 | ENSMUSG00000019851 | Perp | 64058 [Gene Symbol: Perp] [Locus Tag: ] [Chromosome: 10] [Map Location: 10\|10 A2] [ | 3.957629132 | 1.18E-06 | 5.25E-06 |
| 16467 | ENSMUSG00000064289 | Tank | 21353 [Gene Symbol: Tank] [Locus Tag: ] [Chromosome: 2] [Map Location: 2\|2 C1.3] [D | 3.957129819 | 3.34E-08 | 1.92E-07 |
| 7642 | ENSMUSG00000019878 | Hsf2 | 15500 [Gene Symbol: Hsf2] [Locus Tag: ] [Chromosome: 10] [Map Location: 10\|10 B3-B | 3.95568111 | 3.79E-10 | 2.90E-09 |
| 10530 | ENSMUSG00000021694 | Ercc8 | 71991 [Gene Symbol: Ercc8] [Locus Tag: ] [Chromosome: 13] [Map Location: 13\|13 D] [ | 3.954005919 | 1.43E-11 | 1.36E-10 |
| 5579 | ENSMUSG00000021785 | Ngly1 | 59007 [Gene Symbol: Ngly1] [Locus Tag: ] [Chromosome: 14] [Map Location: 14 A2\|14 | 3.951543195 | 1.37E-11 | 1.31E-10 |
| 9931 | ENSMUSG00000027195 | Hsd17b12 | 56348 [Gene Symbol: Hsd17b12] [Locus Tag: ] [Chromosome: 2] [Map Location: 2\|2 E1 | 3.948929277 | 4.35E-11 | 3.87E-10 |
| 13591 | ENSMUSG00000055733 | Nap1l3 | 54561 [Gene Symbol: Nap1l3] [Locus Tag: ] [Chromosome: X] [Map Location: X\|X E3] [D | 3.948482849 | 1.49E-08 | 9.07E-08 |
| 15559 | ENSMUSG00000048076 | Arf1 | 11840 [Gene Symbol: Arf1] [Locus Tag: ] [Chromosome: 11] [Map Location: 11\|11 B2] [ | 3.946341604 | 4.66E-14 | 6.71E-13 |
| 2441 | ENSMUSG00000046314 | Stxbp6 | 217517 [Gene Symbol: Stxbp6] [Locus Tag: ] [Chromosome: 12] [Map Location: 12\|12 | 3.944411182 | 4.90E-09 | 3.21E-08 |
| 2110 | ENSMUSG00000022533 | Atp13a3 | 224088 [Gene Symbol: Atp13a3] [Locus Tag: ] [Chromosome: 16] [Map Location: 16\|16 | 3.942526642 | 7.96E-15 | 1.29E-13 |
| 7690 | ENSMUSG00000031538 | Plat | 18791 [Gene Symbol: Plat] [Locus Tag: ] [Chromosome: 8] [Map Location: 8 A2\|8 11.42 | 3.94240634 | 2.23E-09 | 1.53E-08 |
| 9293 | ENSMUSG00000068154 | Insm1 | 53626 [Gene Symbol: Insm1] [Locus Tag: ] [Chromosome: 2] [Map Location: 2\|2 G1] [De | 3.940837343 | 0.000189 | 0.000566 |
| 1107 | ENSMUSG00000024817 | Uhrf2 | 109113 [Gene Symbol: Uhrf2] [Locus Tag: ] [Chromosome: 19] [Map Location: 19\|19 C | 3.940346139 | 3.48E-11 | 3.13E-10 |
| 5231 | ENSMUSG00000028990 | Lzic | 69151 [Gene Symbol: Lzic] [Locus Tag: ] [Chromosome: 4] [Map Location: 4\|4 E2] [Desc | 3.939667542 | 2.02E-11 | 1.89E-10 |
| 18556 | ENSMUSG00000029068 | Ccnl2 | 56036 [Gene Symbol: Ccnl2] [Locus Tag: ] [Chromosome: 4] [Map Location: 4 E2\|4 87.4 | 3.939628768 | 1.68E-12 | 1.86E-11 |
| 18280 | ENSMUSG00000057098 | Ebf1 | 13591 [Gene Symbol: Ebf1] [Locus Tag: ] [Chromosome: 11] [Map Location: 11 B1.1\|11 | 3.938137813 | 3.22E-10 | 2.49E-09 |
| 16688 | ENSMUSG00000002428 | Hltf | 20585 [Gene Symbol: Hltf] [Locus Tag: ] [Chromosome: 3] [Map Location: 3 A2\|3 6.15 c | 3.936788012 | 8.72E-10 | 6.36E-09 |
| 12289 | ENSMUSG00000026977 | March7 | 57438 [Gene Symbol: March7] [Locus Tag: ] [Chromosome: 2] [Map Location: 2\|2 C3] [D | 3.935279468 | 3.24E-14 | 4.78E-13 |
| 7855 | ENSMUSG00000021635 | Rad17 | 19356 [Gene Symbol: Rad17] [Locus Tag: ] [Chromosome: 13] [Map Location: 13\|13 D1 | 3.929925324 | 5.94E-12 | 6.04E-11 |
| 2940 | ENSMUSG00000036093 | Arl5a | 75423 [Gene Symbol: Arl5a] [Locus Tag: ] [Chromosome: 2] [Map Location: 2\|2 C1.1] [D | 3.929671709 | 6.20E-12 | 6.30E-11 |
| 17647 | ENSMUSG00000026830 | Ermn | 77767 [Gene Symbol: Ermn] [Locus Tag: ] [Chromosome: 2] [Map Location: 2\|2 C2] [Des | 3.928982696 | 0.000751 | 0.002001 |
| 1560 | ENSMUSG00000052584 | Serp2 | 72661 [Gene Symbol: Serp2] [Locus Tag: ] [Chromosome: 14] [Map Location: 14\|14 D3 | 3.927537727 | 2.46E-09 | 1.67E-08 |
| 6475 | ENSMUSG00000040297 | Suco | 226551 [Gene Symbol: Suco] [Locus Tag: ] [Chromosome: 1] [Map Location: 1\|1 H2.1] [ | 3.926002388 | 1.27E-13 | 1.69E-12 |
| 18401 | ENSMUSG00000026276 | Sept2 | 18000 [Gene Symbol: Sept2] [Locus Tag: ] [Chromosome: 1] [Map Location: 1\|1 D] [Des | 3.925998055 | 1.17E-13 | 1.56E-12 |
| 4698 | ENSMUSG00000022656 | Pvrl3 | 58998 [Gene Symbol: Pvrl3] [Locus Tag: ] [Chromosome: 16] [Map Location: 16\|16 B5] | 3.925974721 | 1.20E-12 | 1.36E-11 |
| 9725 | ENSMUSG00000035674 | Ndufa3 | 66091 [Gene Symbol: Ndufa3] [Locus Tag: ] [Chromosome: 7] [Map Location: 7\|7 A1] [D | 3.924428122 | 2.90E-06 | 1.20E-05 |
| 11614 | ENSMUSG00000094500 | Smim18 | 72632 [Gene Symbol: Smim18] [Locus Tag: ] [Chromosome: 8] [Map Location: 8\|8] [Des | 3.924140377 | 4.80E-05 | 0.00016 |
| 16962 | ENSMUSG00000025198 | Erlin1 | 226144 [Gene Symbol: Erlin1] [Locus Tag: ] [Chromosome: 19] [Map Location: 19\|19 C | 3.923299311 | 2.09E-11 | 1.95E-10 |
| 7018 | ENSMUSG00000030452 | Nipa2 | 93790 [Gene Symbol: Nipa2] [Locus Tag: ] [Chromosome: 7] [Map Location: 7\|7 B5] [De | 3.92045369 | 1.39E-10 | 1.14E-09 |
| 17420 | ENSMUSG00000008475 | Arpc5 | 67771 [Gene Symbol: Arpc5] [Locus Tag: ] [Chromosome: 1] [Map Location: 1\|1 A3] [De | 3.920419447 | 7.52E-14 | 1.04E-12 |
| 18546 | ENSMUSG00000005413 | Hmox1 | 15368 [Gene Symbol: Hmox1] [Locus Tag: ] [Chromosome: 8] [Map Location: 8 35.59 c | 3.92002119 | 0.00537 | 0.011876 |
| 4126 | ENSMUSG00000047514 | Tspyl1 | 22110 [Gene Symbol: Tspyl1] [Locus Tag: ] [Chromosome: 10] [Map Location: 10\|10 B1 | 3.919297578 | 4.87E-13 | 5.88E-12 |
| 6055 | ENSMUSG00000042750 | Bex2 | 12069 [Gene Symbol: Bex2] [Locus Tag: ] [Chromosome: X] [Map Location: X F1\|X 57.3 | 3.91841914 | 6.48E-14 | 9.09E-13 |
| 11332 | ENSMUSG00000024122 | Pdpk1 | 18607 [Gene Symbol: Pdpk1] [Locus Tag: ] [Chromosome: 17] [Map Location: 17\|17 A3 | 3.918399814 | 4.16E-14 | 6.04E-13 |
| 15996 | ENSMUSG00000021003 | Galc | 14420 [Gene Symbol: Galc] [Locus Tag: ] [Chromosome: 12] [Map Location: 12 E\|12 49. | 3.917298376 | 3.02E-12 | 3.20E-11 |
| 6939 | ENSMUSG00000078546 | 2210404O09Rik | 70081 [Gene Symbol: 2210404O09Rik] [Locus Tag: ] [Chromosome: 17] [Map Location: | 3.91401437 | 4.76E-06 | 1.90E-05 |
| 18451 | ENSMUSG00000002015 | Bcap31 | 27061 [Gene Symbol: Bcap31] [Locus Tag: ] [Chromosome: X] [Map Location: X\|X A6] [D | 3.908777858 | 4.62E-09 | 3.04E-08 |
| 11781 | ENSMUSG00000046994 | Mars2 | 212679 [Gene Symbol: Mars2] [Locus Tag: ] [Chromosome: 1] [Map Location: 1\|1 C1.2] | 3.907798675 | 1.55E-13 | 2.04E-12 |
| 16423 | ENSMUSG00000020170 | Frs2 | 327826 [Gene Symbol: Frs2] [Locus Tag: ] [Chromosome: 10] [Map Location: 10\|10 D2] | 3.907525601 | 1.44E-11 | 1.37E-10 |
| 5802 | ENSMUSG00000009907 | Vps4b | 20479 [Gene Symbol: Vps4b] [Locus Tag: ] [Chromosome: 1] [Map Location: 1\|1 D] [Des | 3.904482487 | 8.42E-13 | 9.86E-12 |
| 6774 | ENSMUSG00000040928 | S100pbp | 74648 [Gene Symbol: S100pbp] [Locus Tag: ] [Chromosome: 4] [Map Location: 4\|4 D2. | 3.903303462 | 2.66E-13 | 3.34E-12 |
| 10003 | ENSMUSG00000015214 | Mtmr1 | 53332 [Gene Symbol: Mtmr1] [Locus Tag: ] [Chromosome: X] [Map Location: X\|X A7.2] | 3.903174173 | 9.01E-13 | 1.05E-11 |
| 6053 | ENSMUSG00000069132 | Nxph2 | 18232 [Gene Symbol: Nxph2] [Locus Tag: ] [Chromosome: 2] [Map Location: 2\|2 A3] [D | 3.902206397 | 9.32E-09 | 5.86E-08 |
| 5335 | ENSMUSG00000062373 | Tmem65 | 74868 [Gene Symbol: Tmem65] [Locus Tag: ] [Chromosome: 15] [Map Location: 15\|15 | 3.90139216 | 2.42E-14 | 3.64E-13 |
| 9672 | ENSMUSG00000039001 | Rps21 | 66481 [Gene Symbol: Rps21] [Locus Tag: ] [Chromosome: 2] [Map Location: 2\|2 H4] [De | 3.900995325 | 1.51E-06 | 6.62E-06 |
| 8146 | ENSMUSG00000058503 | Fam133b | 68152 [Gene Symbol: Fam133b] [Locus Tag: ] [Chromosome: 5] [Map Location: 5\|5 A2] | 3.900334055 | 2.04E-08 | 1.21E-07 |
| 1027 | ENSMUSG00000019872 | Smpdl3a | 57319 [Gene Symbol: Smpdl3a] [Locus Tag: ] [Chromosome: 10] [Map Location: 10\|10 | 3.893754575 | 2.84E-07 | 1.40E-06 |
| 15716 | ENSMUSG00000052155 | Acvr2a | 11480 [Gene Symbol: Acvr2a] [Locus Tag: ] [Chromosome: 2] [Map Location: 2 C1.1\|2 2 | 3.892184734 | 3.43E-13 | 4.22E-12 |
| 3628 | ENSMUSG00000038069 | Cdkn2aip | 70925 [Gene Symbol: Cdkn2aip] [Locus Tag: ] [Chromosome: 8] [Map Location: 8\|8 B1. | 3.890995073 | 1.82E-12 | 2.01E-11 |
| 648 | ENSMUSG00000036391 | Sec24a | 77371 [Gene Symbol: Sec24a] [Locus Tag: ] [Chromosome: 11] [Map Location: 11\|11 B | 3.890849223 | 1.51E-10 | 1.23E-09 |
| 16673 | ENSMUSG00000005873 | Reep5 | 13476 [Gene Symbol: Reep5] [Locus Tag: ] [Chromosome: 18] [Map Location: 18\|18 B1 | 3.887414569 | 2.56E-14 | 3.83E-13 |
| 585 | ENSMUSG00000030170 | Wnt5b | 22419 [Gene Symbol: Wnt5b] [Locus Tag: ] [Chromosome: 6] [Map Location: 6 F1\|6 56. | 3.886091104 | 9.06E-12 | 8.95E-11 |
| 11551 | ENSMUSG00000079450 | Cldn34c1 | 73061 [Gene Symbol: Cldn34c1] [Locus Tag: ] [Chromosome: X] [Map Location: X\|X E3] | 3.883659882 | 2.51E-05 | 8.88E-05 |
| 313 | ENSMUSG00000055639 | Dach1 | 13134 [Gene Symbol: Dach1] [Locus Tag: ] [Chromosome: 14] [Map Location: 14 48.25 | 3.883433701 | 7.87E-14 | 1.09E-12 |
| 15133 | ENSMUSG00000052446 | Zfp961 | 234413 [Gene Symbol: Zfp961] [Locus Tag: ] [Chromosome: 8] [Map Location: 8\|8 B3.3 | 3.883284332 | 1.21E-11 | 1.17E-10 |
| 986 | ENSMUSG00000042354 | Gnl3 | 30877 [Gene Symbol: Gnl3] [Locus Tag: ] [Chromosome: 14] [Map Location: 14 B\|14 19 | 3.879956847 | 5.87E-10 | 4.38E-09 |
| 11657 | ENSMUSG00000031604 | Msmo1 | 66234 [Gene Symbol: Msmo1] [Locus Tag: ] [Chromosome: 8] [Map Location: 8\|8 B3.1] | 3.878833787 | 3.19E-12 | 3.36E-11 |
| 13136 | ENSMUSG00000028266 | Lmo4 | 16911 [Gene Symbol: Lmo4] [Locus Tag: ] [Chromosome: 3] [Map Location: 3 H2\|3 68. | 3.878536194 | 1.43E-14 | 2.22E-13 |
| 13850 | ENSMUSG00000062078 | Qk | 19317 [Gene Symbol: Qk] [Locus Tag: ] [Chromosome: 17] [Map Location: 17 A1\|17 7.7 | 3.87718873 | 6.68E-15 | 1.09E-13 |
| 8476 | ENSMUSG00000031596 | Slc7a2 | 11988 [Gene Symbol: Slc7a2] [Locus Tag: ] [Chromosome: 8] [Map Location: 8 A4\|8 23. | 3.876228003 | 1.96E-10 | 1.57E-09 |
| 16354 | ENSMUSG00000031697 | Orc6 | 56452 [Gene Symbol: Orc6] [Locus Tag: ] [Chromosome: 8] [Map Location: 8\|8 C3] [Des | 3.874624527 | 1.61E-10 | 1.31E-09 |
| 12791 | ENSMUSG00000020669 | Sh3yl1 | 24057 [Gene Symbol: Sh3yl1] [Locus Tag: ] [Chromosome: 12] [Map Location: 12\|12 A | 3.873703717 | 1.89E-09 | 1.31E-08 |
| 17929 | ENSMUSG00000027342 | Pcna | 18538 [Gene Symbol: Pcna] [Locus Tag: ] [Chromosome: 2] [Map Location: 2 F2\|2 64.1 | 3.873700864 | 3.43E-08 | 1.97E-07 |
| 14342 | ENSMUSG00000019763 | Rmnd1 | 66084 [Gene Symbol: Rmnd1] [Locus Tag: ] [Chromosome: 10] [Map Location: 10\|10 A | 3.872965527 | 8.83E-07 | 4.02E-06 |
| 17872 | ENSMUSG00000037058 | Paip2 | 67869 [Gene Symbol: Paip2] [Locus Tag: ] [Chromosome: 18] [Map Location: 18\|18 B2] | 3.871078114 | 2.43E-12 | 2.61E-11 |
| 4859 | ENSMUSG00000009575 | Cbx5 | 12419 [Gene Symbol: Cbx5] [Locus Tag: ] [Chromosome: 15] [Map Location: 15\|15 F3] | 3.870410673 | 6.20E-11 | 5.39E-10 |
| 16396 | ENSMUSG00000014226 | Cacybp | 12301 [Gene Symbol: Cacybp] [Locus Tag: ] [Chromosome: 1] [Map Location: 1\|1 H1] [D | 3.870025058 | 2.64E-13 | 3.32E-12 |

| | | | | | | |
|---|---|---|---|---|---|---|
| 18156 | ENSMUSG00000033966 | Cdkl4 | 381113 [Gene Symbol: Cdkl4] [Locus Tag: ] [Chromosome: 17] [Map Location: 17\|17 E3 | 3.86795329 | 3.28E-07 | 1.60E-06 |
| 6595 | ENSMUSG00000028497 | Hacd4 | 66775 [Gene Symbol: Hacd4] [Locus Tag: ] [Chromosome: 4] [Map Location: 4\|4 C4] [De | 3.867926429 | 1.44E-11 | 1.37E-10 |
| 17448 | ENSMUSG00000051451 | Crebzf | 233490 [Gene Symbol: Crebzf] [Locus Tag: ] [Chromosome: 7] [Map Location: 7\|7 E1] [D | 3.866492557 | 1.44E-11 | 1.37E-10 |
| 13116 | ENSMUSG00000024583 | Txnl1 | 53382 [Gene Symbol: Txnl1] [Locus Tag: ] [Chromosome: 18] [Map Location: 18\|18 D3] | 3.862703612 | 3.21E-12 | 3.38E-11 |
| 8900 | ENSMUSG00000036357 | Gpr101 | 245424 [Gene Symbol: Gpr101] [Locus Tag: ] [Chromosome: X] [Map Location: X\|X A6] | 3.861789556 | 0.017459 | 0.034147 |
| 11787 | ENSMUSG00000028226 | Mmp16 | 17389 [Gene Symbol: Mmp16] [Locus Tag: ] [Chromosome: 4] [Map Location: 4 A3\|4 7. | 3.8584963 | 9.36E-13 | 1.08E-11 |
| 1535 | ENSMUSG00000091896 | Ube2d2a | 56550 [Gene Symbol: Ube2d2a] [Locus Tag: ] [Chromosome: 18] [Map Location: 18\|18 | 3.856762375 | 6.31E-13 | 7.49E-12 |
| 3155 | ENSMUSG00000070372 | Capza1 | 12340 [Gene Symbol: Capza1] [Locus Tag: ] [Chromosome: 3] [Map Location: 3 F2.2\|3 4 | 3.85496803 | 8.32E-10 | 6.07E-09 |
| 8773 | ENSMUSG00000079508 | Apoo | 68316 [Gene Symbol: Apoo] [Locus Tag: ] [Chromosome: X] [Map Location: X\|X] [Descri | 3.850975956 | 2.37E-06 | 1.00E-05 |
| 12635 | ENSMUSG00000071342 | Lsmem1 | 380755 [Gene Symbol: Lsmem1] [Locus Tag: ] [Chromosome: 12] [Map Location: 12\|12 | 3.847088029 | 9.02E-07 | 4.10E-06 |
| 17305 | ENSMUSG00000041828 | Abca8a | 217258 [Gene Symbol: Abca8a] [Locus Tag: ] [Chromosome: 11] [Map Location: 11\|11 | 3.847035769 | 2.40E-06 | 1.01E-05 |
| 647 | ENSMUSG00000040521 | Tsfm | 66399 [Gene Symbol: Tsfm] [Locus Tag: ] [Chromosome: 10] [Map Location: 10\|10 D3] | 3.8447215 | 2.12E-09 | 1.46E-08 |
| 2576 | ENSMUSG00000043131 | Mob1a | 232157 [Gene Symbol: Mob1a] [Locus Tag: ] [Chromosome: 6] [Map Location: 6\|6 C3] [ | 3.841710215 | 2.65E-10 | 2.08E-09 |
| 6224 | ENSMUSG00000040040 | Ift88 | 21821 [Gene Symbol: Ift88] [Locus Tag: ] [Chromosome: 14] [Map Location: 14 C3\|14 3 | 3.840854009 | 2.42E-07 | 1.21E-06 |
| 595 | ENSMUSG00000030861 | Acadsb | 66885 [Gene Symbol: Acadsb] [Locus Tag: ] [Chromosome: 7] [Map Location: 7\|7 F4] [D | 3.83879434 | 7.03E-10 | 5.18E-09 |
| 4286 | ENSMUSG00000089798 | 1700028K03Rik | 76421 [Gene Symbol: 1700028K03Rik] [Locus Tag: ] [Chromosome: 5] [Map Location: 5 | 3.831324946 | 0.001981 | 0.00483 |
| 16109 | ENSMUSG00000021638 | Ocln | 18260 [Gene Symbol: Ocln] [Locus Tag: ] [Chromosome: 13] [Map Location: 13 D1\|13 5 | 3.830088321 | 5.19E-07 | 2.45E-06 |
| 4840 | ENSMUSG00000035849 | Krt222 | 268481 [Gene Symbol: Krt222] [Locus Tag: ] [Chromosome: 11] [Map Location: 11\|11 D | 3.829445469 | 2.34E-06 | 9.86E-06 |
| 10079 | ENSMUSG00000016308 | Ube2a; Ube2b | 22209 [Gene Symbol: Ube2a] [Locus Tag: ] [Chromosome: X] [Map Location: X\|X A3.3] [ | 3.827066321 | 1.21E-08 | 7.46E-08 |
| 9534 | ENSMUSG00000029089 | Pacrgl | 66768 [Gene Symbol: Pacrgl] [Locus Tag: ] [Chromosome: 5] [Map Location: 5\|5 B3] [De | 3.824920222 | 1.85E-11 | 1.74E-10 |
| 1340 | ENSMUSG00000032374 | Plod2 | 26432 [Gene Symbol: Plod2] [Locus Tag: ] [Chromosome: 9] [Map Location: 9 E3.3\|9 48 | 3.824379778 | 1.53E-07 | 7.88E-07 |
| 6105 | ENSMUSG00000033487 | Fndc3a | 319448 [Gene Symbol: Fndc3a] [Locus Tag: ] [Chromosome: 14] [Map Location: 14 D2\| | 3.823028797 | 5.63E-13 | 6.74E-12 |
| 4114 | ENSMUSG00000001833 | Sep7 | 235072 [Gene Symbol: Sept7] [Locus Tag: ] [Chromosome: 9] [Map Location: 9\|9 A4] [D | 3.820770884 | 1.39E-12 | 1.55E-11 |
| 413 | ENSMUSG00000008333 | Snrpb2 | 20639 [Gene Symbol: Snrpb2] [Locus Tag: ] [Chromosome: 2] [Map Location: 2\|2 G1] [D | 3.819292535 | 1.32E-08 | 8.11E-08 |
| 6689 | ENSMUSG00000036087 | Slain2 | 75991 [Gene Symbol: Slain2] [Locus Tag: ] [Chromosome: 5] [Map Location: 5\|5 C3.2] [ | 3.818114841 | 1.98E-12 | 2.17E-11 |
| 5250 | ENSMUSG00000028233 | Tgs1 | 116940 [Gene Symbol: Tgs1] [Locus Tag: ] [Chromosome: 4] [Map Location: 4 A1\|4 1.99 | 3.817177624 | 1.58E-10 | 1.29E-09 |
| 2692 | ENSMUSG00000037847 | Nmrk1 | 225994 [Gene Symbol: Nmrk1] [Locus Tag: ] [Chromosome: 19] [Map Location: 19\|19 B | 3.816641336 | 1.26E-05 | 4.71E-05 |
| 5286 | ENSMUSG00000047022 | Mipol1 | 73490 [Gene Symbol: Mipol1] [Locus Tag: ] [Chromosome: 12] [Map Location: 12 24.7 | 3.815965258 | 5.81E-11 | 5.06E-10 |
| 1270 | ENSMUSG00000040118 | Cacna2d1 | 12293 [Gene Symbol: Cacna2d1] [Locus Tag: ] [Chromosome: 5] [Map Location: 5 A2\|5 | 3.809445479 | 2.98E-14 | 4.42E-13 |
| 5633 | ENSMUSG00000030521 | Mphosph10 | 67973 [Gene Symbol: Mphosph10] [Locus Tag: ] [Chromosome: 7] [Map Location: 7\|7 C | 3.808378478 | 1.65E-10 | 1.33E-09 |
| 954 | ENSMUSG00000008206 | Cers4 | 67260 [Gene Symbol: Cers4] [Locus Tag: ] [Chromosome: 8] [Map Location: 8\|8 A1.2] [D | 3.807345343 | 5.39E-11 | 4.71E-10 |
| 11228 | ENSMUSG00000025609 | Mkln1 | 27418 [Gene Symbol: Mkln1] [Locus Tag: ] [Chromosome: 6] [Map Location: 6\|6 B1-B2] | 3.801899221 | 7.51E-14 | 1.04E-12 |
| 17637 | ENSMUSG00000046785 | Epm2aip1 | 77781 [Gene Symbol: Epm2aip1] [Locus Tag: ] [Chromosome: 9] [Map Location: 9\|9 F3] | 3.800065735 | 1.40E-13 | 1.86E-12 |
| 16096 | ENSMUSG00000038172 | Ttc39b | 69863 [Gene Symbol: Ttc39b] [Locus Tag: ] [Chromosome: 4] [Map Location: 4\|4 C3] [D | 3.799463944 | 8.12E-11 | 6.91E-10 |
| 7646 | ENSMUSG00000079509 | Zfx | 22764 [Gene Symbol: Zfx] [Locus Tag: ] [Chromosome: X] [Map Location: X 41.48 cM\|X | 3.799077194 | 2.19E-12 | 2.37E-11 |
| 12962 | ENSMUSG00000068566 | Myadm | 50918 [Gene Symbol: Myadm] [Locus Tag: ] [Chromosome: 7] [Map Location: 7 A1\|7 1. | 3.798553602 | 4.40E-11 | 3.90E-10 |
| 11235 | ENSMUSG00000036381 | P2ry14 | 140795 [Gene Symbol: P2ry14] [Locus Tag: ] [Chromosome: 3] [Map Location: 3 D\|3 28 | 3.79846798 | 1.47E-08 | 8.93E-08 |
| 8147 | ENSMUSG00000029804 | Herc3 | 73998 [Gene Symbol: Herc3] [Locus Tag: ] [Chromosome: 6] [Map Location: 6\|6 B3] [De | 3.798037404 | 1.78E-09 | 1.24E-08 |
| 2888 | ENSMUSG00000026999 | Nup35 | 69482 [Gene Symbol: Nup35] [Locus Tag: ] [Chromosome: 2] [Map Location: 2\|2 C3] [D | 3.796909062 | 5.77E-07 | 2.71E-06 |
| 8343 | ENSMUSG00000036009 | Mettl25 | 216292 [Gene Symbol: Mettl25] [Locus Tag: ] [Chromosome: 10] [Map Location: 10\|10 | 3.793813082 | 1.89E-10 | 1.52E-09 |
| 9241 | ENSMUSG00000037411 | Serpine1 | 18787 [Gene Symbol: Serpine1] [Locus Tag: ] [Chromosome: 5] [Map Location: 5\|5 G2] | 3.793206784 | 0.014935 | 0.029771 |
| 17195 | ENSMUSG00000037152 | Ndufc1 | 66377 [Gene Symbol: Ndufc1] [Locus Tag: ] [Chromosome: 3] [Map Location: 3\|3 D] [De | 3.792280556 | 8.85E-05 | 0.00028 |
| 16817 | ENSMUSG00000030327 | Necap1 | 67602 [Gene Symbol: Necap1] [Locus Tag: ] [Chromosome: 6] [Map Location: 6\|6 F2] [D | 3.789781641 | 1.79E-13 | 2.32E-12 |
| 17840 | ENSMUSG00000005069 | Pex5 | 19305 [Gene Symbol: Pex5] [Locus Tag: ] [Chromosome: 6] [Map Location: 6 F2\|6 59.1 | 3.789592529 | 3.18E-13 | 3.94E-12 |
| 14222 | ENSMUSG00000021066 | Atl1 | 73991 [Gene Symbol: Atl1] [Locus Tag: ] [Chromosome: 12] [Map Location: 12\|12 C2] [ | 3.789371841 | 3.14E-12 | 3.31E-11 |
| 4969 | ENSMUSG00000051444 | Bbs12 | 241950 [Gene Symbol: Bbs12] [Locus Tag: ] [Chromosome: 3] [Map Location: 3\|3 B] [De | 3.786867726 | 5.85E-06 | 2.31E-05 |
| 10734 | ENSMUSG00000027303 | Ptpra | 19262 [Gene Symbol: Ptpra] [Locus Tag: ] [Chromosome: 2] [Map Location: 2 63.23 cM | 3.786363253 | 5.30E-14 | 7.56E-13 |
| 15177 | ENSMUSG00000022337 | Emc2 | 66736 [Gene Symbol: Emc2] [Locus Tag: ] [Chromosome: 15] [Map Location: 15\|15 B3. | 3.78610396 | 1.99E-07 | 1.00E-06 |
| 10996 | ENSMUSG00000031231 | Cox7b | 66142 [Gene Symbol: Cox7b] [Locus Tag: ] [Chromosome: X] [Map Location: X\|X C3] [De | 3.784227525 | 1.94E-07 | 9.80E-07 |
| 1496 | ENSMUSG00000033906 | Zdhhc15 | 108672 [Gene Symbol: Zdhhc15] [Locus Tag: ] [Chromosome: X] [Map Location: X\|X D] | 3.783550715 | 4.25E-08 | 2.40E-07 |
| 1692 | ENSMUSG00000024491 | Rbm27 | 225432 [Gene Symbol: Rbm27] [Locus Tag: ] [Chromosome: 18] [Map Location: 18\|18 | 3.783123798 | 1.05E-13 | 1.42E-12 |
| 11167 | ENSMUSG00000027961 | Lrrc39 | 109245 [Gene Symbol: Lrrc39] [Locus Tag: ] [Chromosome: 3] [Map Location: 3\|3 G2] [ | 3.783023053 | 2.82E-05 | 9.88E-05 |
| 9818 | ENSMUSG00000026904 | Slc4a10 | 94229 [Gene Symbol: Slc4a10] [Locus Tag: ] [Chromosome: 2] [Map Location: 2\|2 C3] [ | 3.781635182 | 2.39E-05 | 8.50E-05 |
| 15676 | ENSMUSG00000063972 | Nr6a1 | 14536 [Gene Symbol: Nr6a1] [Locus Tag: ] [Chromosome: 2] [Map Location: 2\|2 B] [Des | 3.781175724 | 2.00E-11 | 1.88E-10 |
| 15373 | ENSMUSG00000025993 | Slc40a1 | 53945 [Gene Symbol: Slc40a1] [Locus Tag: ] [Chromosome: 1] [Map Location: 1\|1 B] [D | 3.778015062 | 2.30E-09 | 1.57E-08 |
| 17477 | ENSMUSG00000027366 | Sppl2a | 66552 [Gene Symbol: Sppl2a] [Locus Tag: ] [Chromosome: 2] [Map Location: 2\|2 F3] [D | 3.77784978 | 6.95E-12 | 7.02E-11 |
| 12583 | ENSMUSG00000020620 | Abca8b | 27404 [Gene Symbol: Abca8b] [Locus Tag: ] [Chromosome: 11] [Map Location: 11 E1\|1 | 3.776992157 | 9.22E-05 | 0.000291 |
| 11204 | ENSMUSG00000049092 | Gpr137c | 70713 [Gene Symbol: Gpr137c] [Locus Tag: ] [Chromosome: 14] [Map Location: 14\|14 | 3.776958529 | 9.68E-12 | 9.53E-11 |
| 16818 | ENSMUSG00000020130 | Tbc1d15 | 66687 [Gene Symbol: Tbc1d15] [Locus Tag: ] [Chromosome: 10] [Map Location: 10\|10 | 3.775682707 | 5.16E-08 | 2.87E-07 |
| 4838 | ENSMUSG00000063179 | Pstk | 214580 [Gene Symbol: Pstk] [Locus Tag: ] [Chromosome: 7] [Map Location: 7\|7 F3] [Des | 3.775375075 | 1.67E-06 | 7.25E-06 |
| 11892 | ENSMUSG00000045573 | Penk | 18619 [Gene Symbol: Penk] [Locus Tag: ] [Chromosome: 4] [Map Location: 4 A1\|4 2.31 | 3.775061007 | 2.71E-11 | 2.48E-10 |
| 6405 | ENSMUSG00000019802 | Sec63 | 140740 [Gene Symbol: Sec63] [Locus Tag: ] [Chromosome: 10] [Map Location: 10 B2\|1 | 3.774820527 | 5.46E-13 | 6.54E-12 |
| 17084 | ENSMUSG00000057147 | Dph6 | 66632 [Gene Symbol: Dph6] [Locus Tag: ] [Chromosome: 2] [Map Location: 2\|2 E5] [Des | 3.772316356 | 1.44E-09 | 1.01E-08 |
| 4038 | ENSMUSG00000030346 | Rad51ap1 | 19362 [Gene Symbol: Rad51ap1] [Locus Tag: ] [Chromosome: 6] [Map Location: 6 F3\|6 | 3.770693041 | 0.000335 | 0.000956 |
| 3365 | ENSMUSG00000019809 | Pex3 | 56535 [Gene Symbol: Pex3] [Locus Tag: ] [Chromosome: 10] [Map Location: 10\|10 A] [D | 3.768777383 | 2.22E-10 | 1.76E-09 |
| 4570 | ENSMUSG00000025766 | D3Ertd751e | 73852 [Gene Symbol: D3Ertd751e] [Locus Tag: ] [Chromosome: 3] [Map Location: 3\|3 C | 3.766625689 | 2.89E-05 | 0.000101 |
| 10312 | ENSMUSG00000019929 | Dcn | 13179 [Gene Symbol: Dcn] [Locus Tag: ] [Chromosome: 10] [Map Location: 10 C3\|10 50 | 3.765008927 | 1.00E-06 | 4.51E-06 |
| 2090 | ENSMUSG00000039795 | Zfand1 | 66361 [Gene Symbol: Zfand1] [Locus Tag: ] [Chromosome: 3] [Map Location: 3\|3 A2] [D | 3.764462981 | 1.53E-05 | 5.65E-05 |
| 15708 | ENSMUSG00000025777 | Gdap1 | 14545 [Gene Symbol: Gdap1] [Locus Tag: ] [Chromosome: 1] [Map Location: 1\|1 A4] [D | 3.76296121 | 1.05E-13 | 1.42E-12 |
| 12217 | ENSMUSG00000075486 | Commd6 | 66200 [Gene Symbol: Commd6] [Locus Tag: ] [Chromosome: 14] [Map Location: 14\|14 | 3.762438933 | 5.41E-06 | 2.14E-05 |
| 17434 | ENSMUSG00000061458 | Nol10 | 217431 [Gene Symbol: Nol10] [Locus Tag: ] [Chromosome: 12] [Map Location: 12\|12 A | 3.760896397 | 4.22E-11 | 3.75E-10 |
| 16026 | ENSMUSG00000044098 | Rsbn1 | 229675 [Gene Symbol: Rsbn1] [Locus Tag: ] [Chromosome: 3] [Map Location: 3\|3 F2.2] | 3.759223201 | 9.49E-11 | 7.97E-10 |
| 4074 | ENSMUSG00000023074 | Mospd1 | 70380 [Gene Symbol: Mospd1] [Locus Tag: ] [Chromosome: X] [Map Location: X\|X A4] [ | 3.758798563 | 0.000153 | 0.000467 |
| 4385 | ENSMUSG00000066197 | Gpr139 | 209776 [Gene Symbol: Gpr139] [Locus Tag: ] [Chromosome: 7] [Map Location: 7\|7 F2] | 3.758681625 | 1.41E-07 | 7.31E-07 |
| 15034 | ENSMUSG00000019732 | Calr3 | 73316 [Gene Symbol: Calr3] [Locus Tag: ] [Chromosome: 8] [Map Location: 8\|8 C1] [Des | 3.75766451 | 0.002057 | 0.004994 |
| 14899 | ENSMUSG00000054453 | Sytl5 | 236643 [Gene Symbol: Sytl5] [Locus Tag: ] [Chromosome: X] [Map Location: X\|X A1.1] [ | 3.757642003 | 1.87E-06 | 8.06E-06 |
| 17787 | ENSMUSG00000072704 | Smim10l1 | 381820 [Gene Symbol: Smim10l1] [Locus Tag: ] [Chromosome: 6] [Map Location: 6\|6 G | 3.756675214 | 1.10E-08 | 6.84E-08 |
| 10025 | ENSMUSG00000025529 | Zfp711 | 245595 [Gene Symbol: Zfp711] [Locus Tag: ] [Chromosome: X] [Map Location: X\|X E1] [ | 3.756286627 | 3.03E-09 | 2.04E-08 |
| 12403 | ENSMUSG00000062209 | Erbb4 | 13869 [Gene Symbol: Erbb4] [Locus Tag: ] [Chromosome: 1] [Map Location: 1 C3\|1 33.8 | 3.753896465 | 4.62E-14 | 6.65E-13 |
| 13707 | ENSMUSG00000028719 | Cmpk1 | 66588 [Gene Symbol: Cmpk1] [Locus Tag: ] [Chromosome: 4] [Map Location: 4\|4 C7] [D | 3.751101912 | 2.71E-13 | 3.39E-12 |
| 15635 | ENSMUSG00000019461 | Plscr3 | 70310 [Gene Symbol: Plscr3] [Locus Tag: ] [Chromosome: 11] [Map Location: 11 B3\|11 | 3.750398154 | 6.80E-11 | 5.86E-10 |
| 11840 | ENSMUSG00000035696 | Rnf38 | 73469 [Gene Symbol: Rnf38] [Locus Tag: ] [Chromosome: 4] [Map Location: 4\|4 B1] [De | 3.75014389 | 9.22E-10 | 6.68E-09 |
| 16343 | ENSMUSG00000029641 | Rasl11a | 68895 [Gene Symbol: Rasl11a] [Locus Tag: ] [Chromosome: 5] [Map Location: 5\|5 G3] [ | 3.749566219 | 0.000267 | 0.000776 |
| 401 | ENSMUSG00000026102 | Inpp1 | 16329 [Gene Symbol: Inpp1] [Locus Tag: ] [Chromosome: 1] [Map Location: 1 C1.1\|1 26 | 3.748449559 | 1.09E-11 | 1.07E-10 |

| | | | | | | |
|---|---|---|---|---|---|---|
| 8981 | ENSMUSG00000047714 | Ppp1r2 | 66849 [Gene Symbol: Ppp1r2] [Locus Tag: ] [Chromosome: 16] [Map Location: 16 B2\|1 | 3.747266345 | 1.84E-12 | 2.03E-11 |
| 3439 | ENSMUSG00000067931 | Zfp948 | 381066 [Gene Symbol: Zfp948] [Locus Tag: ] [Chromosome: 17] [Map Location: 17\|17 / | 3.747125516 | 3.44E-10 | 2.64E-09 |
| 4762 | ENSMUSG00000033808 | Tmem87a | 211499 [Gene Symbol: Tmem87a] [Locus Tag: ] [Chromosome: 2] [Map Location: 2\|2 E5 | 3.746591543 | 2.54E-09 | 1.72E-08 |
| 5279 | ENSMUSG00000009905 | Kdsr | 70750 [Gene Symbol: Kdsr] [Locus Tag: ] [Chromosome: 1] [Map Location: 1\|1 E2.1] [De | 3.744889364 | 8.40E-11 | 7.12E-10 |
| 6538 | ENSMUSG00000032215 | Rsl24d1 | 225215 [Gene Symbol: Rsl24d1] [Locus Tag: ] [Chromosome: 9] [Map Location: 9\|9 D] [ | 3.739668223 | 3.67E-05 | 0.000126 |
| 11767 | ENSMUSG00000049612 | Omg | 18377 [Gene Symbol: Omg] [Locus Tag: ] [Chromosome: 11] [Map Location: 11 B5\|11 4 | 3.73869869 | 2.51E-06 | 1.05E-05 |
| 6470 | ENSMUSG00000046519 | Golph3l | 229593 [Gene Symbol: Golph3l] [Locus Tag: ] [Chromosome: 3] [Map Location: 3\|3 F2. | 3.735893314 | 1.46E-06 | 6.43E-06 |
| 14772 | ENSMUSG00000040473 | Cfap69 | 207686 [Gene Symbol: Cfap69] [Locus Tag: ] [Chromosome: 5] [Map Location: 5\|5 A1] [ | 3.735833352 | 2.18E-06 | 9.27E-06 |
| 17474 | ENSMUSG00000050953 | Gja1 | 14609 [Gene Symbol: Gja1] [Locus Tag: ] [Chromosome: 10] [Map Location: 10 B4\|10 2 | 3.734272047 | 7.39E-08 | 4.00E-07 |
| 1078 | ENSMUSG00000078919 | Dpm1 | 13480 [Gene Symbol: Dpm1] [Locus Tag: ] [Chromosome: 2] [Map Location: 2\|2 H3] [De | 3.732657698 | 2.90E-08 | 1.68E-07 |
| 14489 | ENSMUSG00000026659 | Dusp12 | 80915 [Gene Symbol: Dusp12] [Locus Tag: ] [Chromosome: 1] [Map Location: 1\|1 H3] [I | 3.732542045 | 5.74E-08 | 3.16E-07 |
| 2365 | ENSMUSG00000031591 | Asah1 | 11886 [Gene Symbol: Asah1] [Locus Tag: ] [Chromosome: 8] [Map Location: 8\|8 A4] [De | 3.731895911 | 3.66E-08 | 2.09E-07 |
| 14181 | ENSMUSG00000069662 | Marcks | 17118 [Gene Symbol: Marcks] [Locus Tag: ] [Chromosome: 10] [Map Location: 10 B1\|1( | 3.730077417 | 2.94E-13 | 3.65E-12 |
| 10663 | ENSMUSG00000036502 | Tmem255a | 245386 [Gene Symbol: Tmem255a] [Locus Tag: ] [Chromosome: X] [Map Location: X\|X / | 3.729443509 | 2.97E-08 | 1.72E-07 |
| 14184 | ENSMUSG00000047534 | Mis18bp1 | 217653 [Gene Symbol: Mis18bp1] [Locus Tag: ] [Chromosome: 12] [Map Location: 12\| | 3.729390529 | 2.45E-05 | 8.70E-05 |
| 13383 | ENSMUSG00000026707 | Nsun6 | 74455 [Gene Symbol: Nsun6] [Locus Tag: ] [Chromosome: 2] [Map Location: 2\|2 A2] [De | 3.728755438 | 9.53E-09 | 5.99E-08 |
| 968 | ENSMUSG00000022367 | Has2 | 15117 [Gene Symbol: Has2] [Locus Tag: ] [Chromosome: 15] [Map Location: 15 D1\|15 2 | 3.723016511 | 1.44E-06 | 6.36E-06 |
| 9593 | ENSMUSG00000047648 | Fbxo30 | 71865 [Gene Symbol: Fbxo30] [Locus Tag: ] [Chromosome: 10] [Map Location: 10\|10 A | 3.72296465 | 6.88E-10 | 5.07E-09 |
| 5317 | ENSMUSG00000028195 | Cyr61 | 16007 [Gene Symbol: Cyr61] [Locus Tag: ] [Chromosome: 3] [Map Location: 3 H2\|3 70. | 3.720462272 | 3.21E-05 | 0.000111 |
| 8757 | ENSMUSG00000022034 | Esco2 | 71988 [Gene Symbol: Esco2] [Locus Tag: ] [Chromosome: 14] [Map Location: 14\|14 D1] | 3.720265096 | 0.000617 | 0.001672 |
| 6964 | ENSMUSG00000060708 | Bloc1s4 | 117197 [Gene Symbol: Bloc1s4] [Locus Tag: ] [Chromosome: 5] [Map Location: 5\|5 B3] | 3.719352992 | 1.80E-08 | 1.08E-07 |
| 9463 | ENSMUSG00000042198 | Chchd7 | 66433 [Gene Symbol: Chchd7] [Locus Tag: ] [Chromosome: 4] [Map Location: 4\|4 A1] [I | 3.719305445 | 5.39E-06 | 2.14E-05 |
| 13871 | ENSMUSG00000037010 | Apln | 30878 [Gene Symbol: Apln] [Locus Tag: ] [Chromosome: X] [Map Location: X\|X A3.2] [De | 3.716813286 | 1.19E-07 | 6.26E-07 |
| 2335 | ENSMUSG00000035356 | Nfkbiz | 80859 [Gene Symbol: Nfkbiz] [Locus Tag: ] [Chromosome: 16] [Map Location: 16\|16 C1 | 3.715755778 | 9.77E-07 | 4.41E-06 |
| 307 | ENSMUSG00000035161 | Ints6 | 18130 [Gene Symbol: Ints6] [Locus Tag: ] [Chromosome: 14] [Map Location: 14\|14 C3] | 3.713776448 | 7.92E-12 | 7.90E-11 |
| 8128 | ENSMUSG00000022679 | Mpv17l | 93734 [Gene Symbol: Mpv17l] [Locus Tag: ] [Chromosome: 16] [Map Location: 16\|16 A | 3.712502994 | 2.89E-07 | 1.42E-06 |
| 3739 | ENSMUSG00000031513 | Leprotl1 | 68192 [Gene Symbol: Leprotl1] [Locus Tag: ] [Chromosome: 8] [Map Location: 8\|8 A4] [ | 3.711712047 | 6.57E-12 | 6.66E-11 |
| 2113 | ENSMUSG00000040452 | Cdh12 | 215654 [Gene Symbol: Cdh12] [Locus Tag: ] [Chromosome: 15] [Map Location: 15 A2\|1 | 3.710331833 | 2.18E-12 | 2.36E-11 |
| 10056 | ENSMUSG00000079104 | Prps1l3 | 328099 [Gene Symbol: Prps1l3] [Locus Tag: ] [Chromosome: 12] [Map Location: 12 C1\| | 3.709962572 | 1.77E-07 | 9.03E-07 |
| 1493 | ENSMUSG00000041301 | Cftr | 12638 [Gene Symbol: Cftr] [Locus Tag: ] [Chromosome: 6] [Map Location: 6 A3\|6 8.1 cN | 3.709828344 | 0.00014 | 0.000427 |
| 1460 | ENSMUSG00000045871 | Slitrk6 | 239250 [Gene Symbol: Slitrk6] [Locus Tag: ] [Chromosome: 14] [Map Location: 14\|14 E | 3.706766385 | 3.08E-05 | 0.000107 |
| 7310 | ENSMUSG00000052102 | Gnpda1 | 26384 [Gene Symbol: Gnpda1] [Locus Tag: ] [Chromosome: 18] [Map Location: 18\|18 E | 3.70670618 | 1.37E-07 | 7.12E-07 |
| 7876 | ENSMUSG00000035521 | Gnptg | 214505 [Gene Symbol: Gnptg] [Locus Tag: ] [Chromosome: 17] [Map Location: 17\|17 A | 3.702939797 | 2.79E-10 | 2.18E-09 |
| 6281 | ENSMUSG00000027598 | Itch | 16396 [Gene Symbol: Itch] [Locus Tag: ] [Chromosome: 2] [Map Location: 2 H1\|2 76.94 | 3.701314383 | 3.34E-12 | 3.52E-11 |
| 12084 | ENSMUSG00000020052 | Ascl1 | 17172 [Gene Symbol: Ascl1] [Locus Tag: ] [Chromosome: 10] [Map Location: 10\|10 C1] | 3.700901777 | 1.25E-08 | 7.70E-08 |
| 8391 | ENSMUSG00000027852 | Nras | 18176 [Gene Symbol: Nras] [Locus Tag: ] [Chromosome: 3] [Map Location: 3 F2.2\|3 45. | 3.699641941 | 3.32E-13 | 4.10E-12 |
| 13322 | ENSMUSG00000062931 | Zfp938 | 237411 [Gene Symbol: Zfp938] [Locus Tag: ] [Chromosome: 10] [Map Location: 10 C1\| | 3.698952706 | 5.47E-08 | 3.03E-07 |
| 6434 | ENSMUSG00000050323 | Ndufaf6 | 76947 [Gene Symbol: Ndufaf6] [Locus Tag: ] [Chromosome: 4] [Map Location: 4\|4 A1] [I | 3.697904014 | 1.48E-05 | 5.46E-05 |
| 10696 | ENSMUSG00000031342 | Gpm6b | 14758 [Gene Symbol: Gpm6b] [Locus Tag: ] [Chromosome: X] [Map Location: X\|X F5] [D | 3.696551095 | 1.61E-13 | 2.11E-12 |
| 2421 | ENSMUSG00000047044 | D030056L22Rik | 225995 [Gene Symbol: D030056L22Rik] [Locus Tag: ] [Chromosome: 19] [Map Locatior | 3.69646264 | 1.87E-07 | 9.50E-07 |
| 15887 | ENSMUSG00000032068 | Plet1 | 76509 [Gene Symbol: Plet1] [Locus Tag: ] [Chromosome: 9] [Map Location: 9\|9 B] [Desc | 3.695714409 | 0.022869 | 0.043411 |
| 4350 | ENSMUSG00000050010 | Shisa3 | 330096 [Gene Symbol: Shisa3] [Locus Tag: ] [Chromosome: 5] [Map Location: 5\|5 C3.1] | 3.69353631 | 3.14E-13 | 3.90E-12 |
| 3592 | ENSMUSG00000020063 | Sirt1 | 93759 [Gene Symbol: Sirt1] [Locus Tag: ] [Chromosome: 10] [Map Location: 10\|10 B4] | 3.690537279 | 2.91E-10 | 2.28E-09 |
| 6896 | ENSMUSG00000020659 | Cbll1 | 104836 [Gene Symbol: Cbll1] [Locus Tag: ] [Chromosome: 12] [Map Location: 12\|12 A3 | 3.689811259 | 9.18E-13 | 1.07E-11 |
| 5907 | ENSMUSG00000035623 | Rsf1 | 233532 [Gene Symbol: Rsf1] [Locus Tag: ] [Chromosome: 7] [Map Location: 7\|7 E2] [Des | 3.688389054 | 2.05E-12 | 2.23E-11 |
| 5638 | ENSMUSG00000020024 | Cep83 | 77048 [Gene Symbol: Cep83] [Locus Tag: ] [Chromosome: 10] [Map Location: 10\|10 C2 | 3.688303651 | 3.66E-09 | 2.44E-08 |
| 596 | ENSMUSG00000006273 | Atp6v1b2 | 11966 [Gene Symbol: Atp6v1b2] [Locus Tag: ] [Chromosome: 8] [Map Location: 8\|8 B3. | 3.688164238 | 1.01E-11 | 9.96E-11 |
| 16740 | ENSMUSG00000039031 | Arhgap18 | 73910 [Gene Symbol: Arhgap18] [Locus Tag: ] [Chromosome: 10] [Map Location: 10\|10 | 3.686939313 | 3.14E-10 | 2.43E-09 |
| 13013 | ENSMUSG00000018102 | Hist1h2bc | 68024 [Gene Symbol: Hist1h2bc] [Locus Tag: ] [Chromosome: 13] [Map Location: 13\|1 | 3.686573578 | 6.31E-05 | 0.000206 |
| 8798 | ENSMUSG00000029152 | Ociad1 | 68095 [Gene Symbol: Ociad1] [Locus Tag: ] [Chromosome: 5] [Map Location: 5\|5 C3.2] | 3.685196296 | 8.73E-13 | 1.02E-11 |
| 12488 | ENSMUSG00000022863 | Btg3 | 12228 [Gene Symbol: Btg3] [Locus Tag: ] [Chromosome: 16] [Map Location: 16\|16 C3.1 | 3.684705497 | 0.000438 | 0.001222 |
| 17627 | ENSMUSG00000029227 | Fip1l1 | 66899 [Gene Symbol: Fip1l1] [Locus Tag: ] [Chromosome: 5] [Map Location: 5\|5 D] [Des | 3.68319945 | 9.86E-10 | 7.10E-09 |
| 13596 | ENSMUSG00000021054 | Sgpp1 | 81535 [Gene Symbol: Sgpp1] [Locus Tag: ] [Chromosome: 12] [Map Location: 12\|12 C3 | 3.682433571 | 2.59E-12 | 2.77E-11 |
| 676 | ENSMUSG00000024529 | Lox | 16948 [Gene Symbol: Lox] [Locus Tag: ] [Chromosome: 18] [Map Location: 18 D1\|18 28 | 3.681714027 | 6.70E-05 | 0.000218 |
| 13074 | ENSMUSG00000031603 | Fgf20 | 80857 [Gene Symbol: Fgf20] [Locus Tag: ] [Chromosome: 8] [Map Location: 8\|8 B1.2] [D | 3.680884101 | 6.92E-05 | 0.000224 |
| 2261 | ENSMUSG00000039037 | St6galnac5 | 26938 [Gene Symbol: St6galnac5] [Locus Tag: ] [Chromosome: 3] [Map Location: 3\|3 H | 3.680235981 | 1.90E-13 | 2.45E-12 |
| 13135 | ENSMUSG00000018395 | Kif3a | 16568 [Gene Symbol: Kif3a] [Locus Tag: ] [Chromosome: 11] [Map Location: 11 A5-B1\| | 3.679717146 | 3.15E-09 | 2.12E-08 |
| 8986 | ENSMUSG00000044337 | Ackr3 | 12778 [Gene Symbol: Ackr3] [Locus Tag: ] [Chromosome: 1] [Map Location: 1 D\|1 45.28 | 3.67850455 | 0.000403 | 0.001135 |
| 2472 | ENSMUSG00000068749 | Psma5 | 26442 [Gene Symbol: Psma5] [Locus Tag: ] [Chromosome: 3] [Map Location: 3\|3 F3] [De | 3.675385132 | 1.05E-05 | 3.97E-05 |
| 3166 | ENSMUSG00000029328 | Hnrnpdl | 50926 [Gene Symbol: Hnrnpdl] [Locus Tag: ] [Chromosome: 5] [Map Location: 5 E4\|5 4 | 3.672195885 | 1.11E-08 | 6.91E-08 |
| 22 | ENSMUSG00000060373 | Hnrnpc | 15381 [Gene Symbol: Hnrnpc] [Locus Tag: ] [Chromosome: 14] [Map Location: 14 C2\|1 | 3.67137497 | 5.76E-11 | 5.02E-10 |
| 9406 | ENSMUSG00000021711 | Trappc13 | 66975 [Gene Symbol: Trappc13] [Locus Tag: ] [Chromosome: 13] [Map Location: 13\|13 | 3.671005893 | 3.06E-11 | 2.77E-10 |
| 12658 | ENSMUSG00000038462 | Uqcrfs1 | 66694 [Gene Symbol: Uqcrfs1] [Locus Tag: ] [Chromosome: 13] [Map Location: 13\|13 A | 3.669058707 | 3.35E-07 | 1.63E-06 |
| 5513 | ENSMUSG00000001767 | Crnkl1 | 66877 [Gene Symbol: Crnkl1] [Locus Tag: ] [Chromosome: 2] [Map Location: 2\|2 G1] [D | 3.662394518 | 1.92E-09 | 1.33E-08 |
| 14359 | ENSMUSG00000024521 | Pmaip1 | 58801 [Gene Symbol: Pmaip1] [Locus Tag: ] [Chromosome: 18] [Map Location: 18\|18 E | 3.660137405 | 0.004741 | 0.010612 |
| 13899 | ENSMUSG00000022285 | Ywhaz | 22631 [Gene Symbol: Ywhaz] [Locus Tag: ] [Chromosome: 15] [Map Location: 15\|15 B3 | 3.65938887 | 1.92E-13 | 2.46E-12 |
| 15851 | ENSMUSG00000062014 | Gmfb | 63985 [Gene Symbol: Gmfb] [Locus Tag: ] [Chromosome: 14] [Map Location: 14 C1\|14 | 3.659086159 | 1.27E-12 | 1.43E-11 |
| 1343 | ENSMUSG00000021364 | Elovl2 | 54326 [Gene Symbol: Elovl2] [Locus Tag: ] [Chromosome: 13] [Map Location: 13\|13 A3 | 3.655993181 | 4.07E-12 | 4.22E-11 |
| 16604 | ENSMUSG00000004896 | Rrnad1 | 229503 [Gene Symbol: Rrnad1] [Locus Tag: ] [Chromosome: 3] [Map Location: 3\|3 F1] [ | 3.656457693 | 1.18E-09 | 8.43E-09 |
| 18038 | ENSMUSG00000028191 | Bcl10 | 12042 [Gene Symbol: Bcl10] [Locus Tag: ] [Chromosome: 3] [Map Location: 3\|3 H3] [De | 3.654855735 | 1.07E-09 | 7.65E-09 |
| 10608 | ENSMUSG00000049764 | Zfp280b | 64453 [Gene Symbol: Zfp280b] [Locus Tag: ] [Chromosome: 10] [Map Location: 10\|10 ( | 3.654369475 | 1.89E-10 | 1.52E-09 |
| 17173 | ENSMUSG00000046168 | Kcnrg | 328424 [Gene Symbol: Kcnrg] [Locus Tag: ] [Chromosome: 14] [Map Location: 14\|14 D | 3.650796775 | 0.000115 | 0.000358 |
| 1515 | ENSMUSG00000034858 | Fam214a | 235493 [Gene Symbol: Fam214a] [Locus Tag: ] [Chromosome: 9] [Map Location: 9\|9 D] | 3.645210837 | 8.04E-13 | 9.45E-12 |
| 9494 | ENSMUSG00000037434 | Slc30a1 | 22782 [Gene Symbol: Slc30a1] [Locus Tag: ] [Chromosome: 1] [Map Location: 1 H6\|1 9 | 3.64206542 | 8.83E-09 | 5.58E-08 |
| 490 | ENSMUSG00000008200 | Fnbp4 | 55935 [Gene Symbol: Fnbp4] [Locus Tag: ] [Chromosome: 2] [Map Location: 2\|2 E1] [De | 3.636821001 | 3.46E-09 | 2.31E-08 |
| 11101 | ENSMUSG00000004631 | Sgce | 20392 [Gene Symbol: Sgce] [Locus Tag: ] [Chromosome: 6] [Map Location: 6 A1\|6 1.81 | 3.636350574 | 8.28E-11 | 7.03E-10 |
| 14439 | ENSMUSG00000043190 | Rfesd | 218341 [Gene Symbol: Rfesd] [Locus Tag: ] [Chromosome: 13] [Map Location: 13\|13 C1 | 3.63442507 | 1.65E-09 | 1.15E-08 |
| 18031 | ENSMUSG00000042607 | Asb4 | 65255 [Gene Symbol: Asb4] [Locus Tag: ] [Chromosome: 6] [Map Location: 6 A1\|6 1.99 | 3.63229143 | 0.000109 | 0.000339 |
| 5930 | ENSMUSG00000032883 | Acsl3 | 74205 [Gene Symbol: Acsl3] [Locus Tag: ] [Chromosome: 1] [Map Location: 1 C4\|1 40.8 | 3.628335139 | 2.86E-13 | 3.57E-12 |
| 9856 | ENSMUSG00000046567 | 4930430F08Rik | 68281 [Gene Symbol: 4930430F08Rik] [Locus Tag: ] [Chromosome: 10] [Map Location: | 3.626602312 | 1.75E-06 | 7.60E-06 |
| 14795 | ENSMUSG00000032413 | Rasa2 | 114713 [Gene Symbol: Rasa2] [Locus Tag: ] [Chromosome: 9] [Map Location: 9\|9 E4] [D | 3.626218748 | 1.90E-12 | 2.09E-11 |
| 15734 | ENSMUSG00000000184 | Ccnd2 | 12444 [Gene Symbol: Ccnd2] [Locus Tag: ] [Chromosome: 6] [Map Location: 6 F3\|6 61. | 3.625625682 | 5.12E-07 | 2.42E-06 |
| 8011 | ENSMUSG00000026409 | Pfkfb2 | 18640 [Gene Symbol: Pfkfb2] [Locus Tag: ] [Chromosome: 1] [Map Location: 1\|1 E4] [De | 3.624763431 | 2.64E-13 | 3.32E-12 |
| 13283 | ENSMUSG00000027599 | Armc1 | 74252 [Gene Symbol: Armc1] [Locus Tag: ] [Chromosome: 3] [Map Location: 3\|3 A3] [De | 3.621853436 | 1.85E-11 | 1.74E-10 |

| | | | | | | |
|---|---|---|---|---|---|---|
| 17676 | ENSMUSG00000040429 | Mterf1a | 545725 [Gene Symbol: Mterf1a] [Locus Tag: ] [Chromosome: 5] [Map Location: 5\|5 A1] | 3.621195842 | 0.005852 | 0.012837 |
| 13676 | ENSMUSG00000020629 | Adi1 | 104923 [Gene Symbol: Adi1] [Locus Tag: ] [Chromosome: 12] [Map Location: 12\|12 A2] | 3.62093054 | 2.90E-06 | 1.20E-05 |
| 384 | ENSMUSG00000096433 | Gm4944 | 240038 [Gene Symbol: Gm4944] [Locus Tag: ] [Chromosome: 17] [Map Location: 17\|17 | 3.620134521 | 3.85E-05 | 0.000131 |
| 15956 | ENSMUSG00000018362 | Kpna2 | 16647 [Gene Symbol: Kpna2] [Locus Tag: ] [Chromosome: 11] [Map Location: 11\|11 E1 | 3.61910075 | 0.000544 | 0.001489 |
| 11938 | ENSMUSG00000090641 | Zfp712 | 78251 [Gene Symbol: Zfp712] [Locus Tag: ] [Chromosome: 13] [Map Location: 13\|13 B3 | 3.616159454 | 4.29E-10 | 3.26E-09 |
| 15959 | ENSMUSG00000022139 | Mbnl2 | 105559 [Gene Symbol: Mbnl2] [Locus Tag: ] [Chromosome: 14] [Map Location: 14\|14 E | 3.614699926 | 4.11E-09 | 2.72E-08 |
| 8338 | ENSMUSG00000053774 | Ubxn7 | 224111 [Gene Symbol: Ubxn7] [Locus Tag: ] [Chromosome: 16] [Map Location: 16\|16 A | 3.613720377 | 6.88E-13 | 8.14E-12 |
| 8321 | ENSMUSG00000040749 | Siah1b | 20438 [Gene Symbol: Siah1b] [Locus Tag: ] [Chromosome: X] [Map Location: X F5\|X 76. | 3.611477287 | 1.85E-05 | 6.73E-05 |
| 4982 | ENSMUSG00000042104 | Uggt2 | 66435 [Gene Symbol: Uggt2] [Locus Tag: ] [Chromosome: 14] [Map Location: 14\|14 E4] | 3.610273541 | 3.79E-07 | 1.83E-06 |
| 7504 | ENSMUSG00000095362 | Gm14325 | 329575 [Gene Symbol: Gm14325] [Locus Tag: ] [Chromosome: 2] [Map Location: 2\|2 H | 3.608804434 | 0.000443 | 0.001237 |
| 15840 | ENSMUSG00000028004 | Npy2r | 18167 [Gene Symbol: Npy2r] [Locus Tag: ] [Chromosome: 3] [Map Location: 3 E3\|3 36.4 | 3.605731321 | 3.94E-06 | 1.60E-05 |
| 8214 | ENSMUSG00000073565 | Prr16 | 71373 [Gene Symbol: Prr16] [Locus Tag: ] [Chromosome: 18] [Map Location: 18\|18 D1] | 3.602608198 | 4.93E-06 | 1.96E-05 |
| 4306 | ENSMUSG00000031600 | Vps37a | 52348 [Gene Symbol: Vps37a] [Locus Tag: ] [Chromosome: 8] [Map Location: 8 23.89 cM | 3.600467299 | 7.00E-13 | 8.25E-12 |
| 8703 | ENSMUSG00000037894 | H2afz | 51788 [Gene Symbol: H2afz] [Locus Tag: ] [Chromosome: 3] [Map Location: 3\|3 G3] [De | 3.595724126 | 0.001898 | 0.004646 |
| 13989 | ENSMUSG00000059173 | Pde1a | 18573 [Gene Symbol: Pde1a] [Locus Tag: ] [Chromosome: 2] [Map Location: 2\|2 D] [Des | 3.591311637 | 7.46E-07 | 3.43E-06 |
| 2346 | ENSMUSG00000038446 | Cdc40 | 71713 [Gene Symbol: Cdc40] [Locus Tag: ] [Chromosome: 10] [Map Location: 10\|10 B1 | 3.590262726 | 2.33E-10 | 1.85E-09 |
| 11495 | ENSMUSG00000040818 | Dennd6a | 211922 [Gene Symbol: Dennd6a] [Locus Tag: ] [Chromosome: 14] [Map Location: 14\|14 | 3.588633669 | 1.06E-11 | 1.04E-10 |
| 7139 | ENSMUSG00000035062 | Zc4h2 | 245522 [Gene Symbol: Zc4h2] [Locus Tag: ] [Chromosome: X] [Map Location: X\|X C3] [D | 3.585288286 | 5.17E-10 | 3.88E-09 |
| 11368 | ENSMUSG00000018481 | Appbp2 | 66884 [Gene Symbol: Appbp2] [Locus Tag: ] [Chromosome: 11] [Map Location: 11\|11 E | 3.585209981 | 1.16E-12 | 1.32E-11 |
| 9006 | ENSMUSG00000044365 | Cxxc4 | 319478 [Gene Symbol: Cxxc4] [Locus Tag: ] [Chromosome: 3] [Map Location: 3\|3 G3] [D | 3.58439106 | 1.72E-09 | 1.20E-08 |
| 16528 | ENSMUSG00000056832 | Ttc26 | 264134 [Gene Symbol: Ttc26] [Locus Tag: ] [Chromosome: 6] [Map Location: 6\|6 B1] [D | 3.583965231 | 5.70E-07 | 2.68E-06 |
| 15023 | ENSMUSG00000024758 | Rtn3 | 20168 [Gene Symbol: Rtn3] [Locus Tag: ] [Chromosome: 19] [Map Location: 19\|19 A] [D | 3.583365416 | 5.21E-13 | 6.25E-12 |
| 6678 | ENSMUSG00000027282 | Mtch2 | 56428 [Gene Symbol: Mtch2] [Locus Tag: ] [Chromosome: 2] [Map Location: 2\|2 E1] [De | 3.580383231 | 3.39E-11 | 3.05E-10 |
| 8873 | ENSMUSG00000031684 | Slc10a7 | 76775 [Gene Symbol: Slc10a7] [Locus Tag: ] [Chromosome: 8] [Map Location: 8\|8 C3] [D | 3.578850056 | 2.17E-10 | 1.73E-09 |
| 12677 | ENSMUSG00000024474 | Ik | 24010 [Gene Symbol: Ik] [Locus Tag: ] [Chromosome: 18] [Map Location: 18\|18 B2] [De | 3.577816444 | 6.36E-11 | 5.52E-10 |
| 13601 | ENSMUSG00000036371 | Serbp1 | 66870 [Gene Symbol: Serbp1] [Locus Tag: ] [Chromosome: 6] [Map Location: 6\|6 C1] [D | 3.574782602 | 3.34E-13 | 4.13E-12 |
| 16883 | ENSMUSG00000024601 | Isoc1 | 66307 [Gene Symbol: Isoc1] [Locus Tag: ] [Chromosome: 18] [Map Location: 18\|18 D2] | 3.574104854 | 2.62E-11 | 2.41E-10 |
| 9012 | ENSMUSG00000073551 | Spink13 | 100038417 [Gene Symbol: Spink13] [Locus Tag: ] [Chromosome: 18] [Map Location: 18 | 3.573335547 | 7.35E-07 | 3.39E-06 |
| 12914 | ENSMUSG00000032400 | Zwilch | 68014 [Gene Symbol: Zwilch] [Locus Tag: ] [Chromosome: 9] [Map Location: 9\|9 D] [Des | 3.571426217 | 8.12E-06 | 3.13E-05 |
| 17320 | ENSMUSG00000042379 | Esm1 | 71690 [Gene Symbol: Esm1] [Locus Tag: ] [Chromosome: 13] [Map Location: 13\|13 D2. | 3.571359905 | 0.008211 | 0.017431 |
| 10549 | ENSMUSG00000029464 | Gpn3 | 68080 [Gene Symbol: Gpn3] [Locus Tag: ] [Chromosome: 5] [Map Location: 5 F\|5 62.33 | 3.570068913 | 1.09E-10 | 9.08E-10 |
| 18185 | ENSMUSG00000075232 | Amd1 | 11702 [Gene Symbol: Amd1] [Locus Tag: ] [Chromosome: 10] [Map Location: 10 21.85 c | 3.569742549 | 4.65E-08 | 2.61E-07 |
| 5749 | ENSMUSG00000019577 | Pdk4 | 27273 [Gene Symbol: Pdk4] [Locus Tag: ] [Chromosome: 6] [Map Location: 6 A1\|6 2.06 | 3.569089339 | 0.000125 | 0.000386 |
| 17554 | ENSMUSG00000022508 | Bcl6 | 12053 [Gene Symbol: Bcl6] [Locus Tag: ] [Chromosome: 16] [Map Location: 16 B1\|16 1 | 3.568617586 | 3.68E-08 | 2.10E-07 |
| 15589 | ENSMUSG00000025665 | Rps6ka6 | 67071 [Gene Symbol: Rps6ka6] [Locus Tag: ] [Chromosome: X] [Map Location: X\|X E1] [ | 3.565087167 | 1.04E-10 | 8.70E-10 |
| 6877 | ENSMUSG00000064138 | Fam172a | 68675 [Gene Symbol: Fam172a] [Locus Tag: ] [Chromosome: 13] [Map Location: 13\|13 | 3.564890328 | 3.24E-12 | 3.41E-11 |
| 2159 | ENSMUSG00000036873 | 2410004B18Rik | 66421 [Gene Symbol: 2410004B18Rik] [Locus Tag: ] [Chromosome: 3] [Map Location: 3 | 3.563379034 | 1.79E-10 | 1.45E-09 |
| 15807 | ENSMUSG00000025266 | Gnl3l | 237107 [Gene Symbol: Gnl3l] [Locus Tag: ] [Chromosome: X] [Map Location: X\|X F3] [D | 3.561772682 | 7.70E-12 | 7.69E-11 |
| 2003 | ENSMUSG00000020585 | Laptm4a | 17775 [Gene Symbol: Laptm4a] [Locus Tag: ] [Chromosome: 12] [Map Location: 12\|12 | 3.559934933 | 1.89E-11 | 1.77E-10 |
| 5116 | ENSMUSG00000061759 | Armt1 | 73419 [Gene Symbol: Armt1] [Locus Tag: ] [Chromosome: 10] [Map Location: 10\|10 A1 | 3.558637257 | 8.66E-06 | 3.32E-05 |
| 10616 | ENSMUSG00000063172 | Hspb11 | 72938 [Gene Symbol: Hspb11] [Locus Tag: ] [Chromosome: 4] [Map Location: 4\|4 C7] [D | 3.558274436 | 1.68E-07 | 8.58E-07 |
| 16373 | ENSMUSG00000017291 | Taok1 | 216965 [Gene Symbol: Taok1] [Locus Tag: ] [Chromosome: 11] [Map Location: 11\|11 B5 | 3.556879289 | 2.00E-12 | 2.18E-11 |
| 8063 | ENSMUSG00000001157 | Gmcl1 | 23885 [Gene Symbol: Gmcl1] [Locus Tag: ] [Chromosome: 6] [Map Location: 6 D1\|6 37. | 3.555784371 | 5.70E-12 | 5.82E-11 |
| 4403 | ENSMUSG00000045576 | St7l | 229681 [Gene Symbol: St7l] [Locus Tag: ] [Chromosome: 3] [Map Location: 3\|3 F2.2] [D | 3.555507014 | 2.93E-11 | 2.66E-10 |
| 17153 | ENSMUSG00000031516 | Dctn6 | 22428 [Gene Symbol: Dctn6] [Locus Tag: ] [Chromosome: 8] [Map Location: 8\|8 A4] [De | 3.554258767 | 2.21E-11 | 2.05E-10 |
| 10173 | ENSMUSG00000069206 | Zfp874a | 238692 [Gene Symbol: Zfp874a] [Locus Tag: ] [Chromosome: 13] [Map Location: 13\|13 | 3.554011598 | 5.68E-06 | 2.24E-05 |
| 15384 | ENSMUSG00000026014 | Raph1 | 77300 [Gene Symbol: Raph1] [Locus Tag: ] [Chromosome: 1] [Map Location: 1\|1 C2] [De | 3.552661163 | 8.21E-11 | 6.98E-10 |
| 5153 | ENSMUSG00000032422 | Snx14 | 244962 [Gene Symbol: Snx14] [Locus Tag: ] [Chromosome: 9] [Map Location: 9\|9 E3.1] | 3.552020636 | 6.01E-09 | 3.90E-08 |
| 1271 | ENSMUSG00000026679 | Enkur | 71233 [Gene Symbol: Enkur] [Locus Tag: ] [Chromosome: 2] [Map Location: 2\|2 A3] [De | 3.550825034 | 0.007585 | 0.016235 |
| 18558 | ENSMUSG00000064254 | Ethe1 | 66071 [Gene Symbol: Ethe1] [Locus Tag: ] [Chromosome: 7] [Map Location: 7\|7 A3] [De | 3.549291854 | 1.23E-07 | 6.47E-07 |
| 9888 | ENSMUSG00000029669 | Tspan12 | 269831 [Gene Symbol: Tspan12] [Locus Tag: ] [Chromosome: 6] [Map Location: 6\|6 A3. | 3.548577234 | 9.26E-09 | 5.83E-08 |
| 11930 | ENSMUSG00000039706 | Ldb2 | 16826 [Gene Symbol: Ldb2] [Locus Tag: ] [Chromosome: 5] [Map Location: 5\|5 B3] [Des | 3.548166198 | 2.31E-08 | 1.36E-07 |
| 5391 | ENSMUSG00000074656 | Eif2s2 | 67204 [Gene Symbol: Eif2s2] [Locus Tag: ] [Chromosome: 2] [Map Location: 2 H1\|2 76.8 | 3.548014038 | 1.44E-10 | 1.18E-09 |
| 12039 | ENSMUSG00000020561 | Twistnb | 28071 [Gene Symbol: Twistnb] [Locus Tag: ] [Chromosome: 12] [Map Location: 12\|12 A | 3.547994439 | 3.98E-08 | 2.26E-07 |
| 10765 | ENSMUSG00000042111 | Ccdc115 | 69668 [Gene Symbol: Ccdc115] [Locus Tag: ] [Chromosome: 1] [Map Location: 1\|1 B] [D | 3.547856139 | 4.63E-10 | 3.50E-09 |
| 13381 | ENSMUSG00000046352 | Gjb2 | 14619 [Gene Symbol: Gjb2] [Locus Tag: ] [Chromosome: 14] [Map Location: 14 30.1 cM | 3.54721385 | 1.63E-05 | 6.00E-05 |
| 16021 | ENSMUSG00000021697 | Depdc1b | 218581 [Gene Symbol: Depdc1b] [Locus Tag: ] [Chromosome: 13] [Map Location: 13\|13 | 3.545293788 | 0.019405 | 0.037502 |
| 4545 | ENSMUSG00000072294 | Klf12 | 16597 [Gene Symbol: Klf12] [Locus Tag: ] [Chromosome: 14] [Map Location: 14 E2.2\|14 | 3.544908331 | 3.38E-11 | 3.04E-10 |
| 822 | ENSMUSG00000025878 | Uimc1 | 20184 [Gene Symbol: Uimc1] [Locus Tag: ] [Chromosome: 13] [Map Location: 13\|13 B2 | 3.543802414 | 1.49E-08 | 9.03E-08 |
| 5277 | ENSMUSG00000075284 | Wipf1 | 215280 [Gene Symbol: Wipf1] [Locus Tag: ] [Chromosome: 2] [Map Location: 2 C3\|2 43 | 3.537873252 | 9.03E-10 | 6.55E-09 |
| 8806 | ENSMUSG00000017776 | Crk | 12928 [Gene Symbol: Crk] [Locus Tag: ] [Chromosome: 11] [Map Location: 11 45.92 cM | 3.536222825 | 4.52E-13 | 5.48E-12 |
| 15766 | ENSMUSG00000093661 | Eif4e3 | 66892 [Gene Symbol: Eif4e3] [Locus Tag: ] [Chromosome: 6] [Map Location: 6\|6 D3] [De | 3.531490818 | 1.30E-08 | 7.96E-08 |
| 15223 | ENSMUSG00000025862 | Stag2 | 20843 [Gene Symbol: Stag2] [Locus Tag: ] [Chromosome: X] [Map Location: X\|X A2] [De | 3.530808091 | 1.55E-12 | 1.72E-11 |
| 8747 | ENSMUSG00000032932 | Hspa13 | 110920 [Gene Symbol: Hspa13] [Locus Tag: ] [Chromosome: 16] [Map Location: 16 C3.2 | 3.530521276 | 2.31E-10 | 1.83E-09 |
| 7175 | ENSMUSG00000002332 | Dhrs1 | 52585 [Gene Symbol: Dhrs1] [Locus Tag: ] [Chromosome: 14] [Map Location: 14 C3\|14 | 3.530434382 | 1.91E-09 | 1.32E-08 |
| 14262 | ENSMUSG00000036499 | Eea1 | 216238 [Gene Symbol: Eea1] [Locus Tag: ] [Chromosome: 10] [Map Location: 10\|10 C3] | 3.529562343 | 2.25E-07 | 1.13E-06 |
| 18432 | ENSMUSG00000029203 | Ube2k | 53323 [Gene Symbol: Ube2k] [Locus Tag: ] [Chromosome: 5] [Map Location: 5 C3.1\|5 3 | 3.529395218 | 1.07E-12 | 1.22E-11 |
| 6835 | ENSMUSG00000050310 | Rictor | 78757 [Gene Symbol: Rictor] [Locus Tag: ] [Chromosome: 15] [Map Location: 15\|15 A1 | 3.528961645 | 6.82E-13 | 8.07E-12 |
| 11172 | ENSMUSG00000029186 | Pi4k2b | 67073 [Gene Symbol: Pi4k2b] [Locus Tag: ] [Chromosome: 5] [Map Location: 5\|5 C1] [D | 3.52799764 | 4.47E-06 | 1.79E-05 |
| 10914 | ENSMUSG00000027714 | Exosc9 | 50911 [Gene Symbol: Exosc9] [Locus Tag: ] [Chromosome: 3] [Map Location: 3\|3 B] [Des | 3.527211996 | 1.09E-09 | 7.81E-09 |
| 15875 | ENSMUSG00000019888 | Mgat4c | 67569 [Gene Symbol: Mgat4c] [Locus Tag: ] [Chromosome: 10] [Map Location: 10\|10 D | 3.525977731 | 0.000308 | 0.000886 |
| 17029 | ENSMUSG00000021112 | Mpp5 | 56217 [Gene Symbol: Mpp5] [Locus Tag: ] [Chromosome: 12] [Map Location: 12\|12 C3] | 3.525929577 | 8.29E-11 | 7.04E-10 |
| 6414 | ENSMUSG00000007989 | Fzd3 | 14365 [Gene Symbol: Fzd3] [Locus Tag: ] [Chromosome: 14] [Map Location: 14 D1\|14 3 | 3.52572208 | 5.26E-10 | 3.94E-09 |
| 3188 | ENSMUSG00000010067 | Rassf1 | 56289 [Gene Symbol: Rassf1] [Locus Tag: ] [Chromosome: 9] [Map Location: 9\|9 F1] [De | 3.525468882 | 2.43E-08 | 1.43E-07 |
| 2966 | ENSMUSG00000026411 | Tmem9 | 66241 [Gene Symbol: Tmem9] [Locus Tag: ] [Chromosome: 1] [Map Location: 1\|1 E4] [D | 3.523652699 | 1.50E-09 | 1.05E-08 |
| 11239 | ENSMUSG00000037474 | Dtl | 76843 [Gene Symbol: Dtl] [Locus Tag: ] [Chromosome: 1] [Map Location: 1\|1 H6] [Descr | 3.521718407 | 1.16E-05 | 4.35E-05 |
| 3247 | ENSMUSG00000027805 | Pfn2 | 18645 [Gene Symbol: Pfn2] [Locus Tag: ] [Chromosome: 3] [Map Location: 3 D\|3 28.15 | 3.521547808 | 7.66E-13 | 9.01E-12 |
| 7029 | ENSMUSG00000020102 | Slc16a7 | 20503 [Gene Symbol: Slc16a7] [Locus Tag: ] [Chromosome: 10] [Map Location: 10\|10 D | 3.52103032 | 3.67E-10 | 2.82E-09 |
| 16994 | ENSMUSG00000032324 | Tspan3 | 56434 [Gene Symbol: Tspan3] [Locus Tag: ] [Chromosome: 9] [Map Location: 9\|9 C] [De | 3.520428528 | 4.25E-12 | 4.40E-11 |
| 10222 | ENSMUSG00000061371 | Zfp873 | 408062 [Gene Symbol: Zfp873] [Locus Tag: ] [Chromosome: 10] [Map Location: 10\|10 C | 3.520290042 | 6.70E-05 | 0.000218 |
| 6740 | ENSMUSG00000037169 | Mycn | 18109 [Gene Symbol: Mycn] [Locus Tag: ] [Chromosome: 12] [Map Location: 12 6.14 cM | 3.519765406 | 5.12E-11 | 4.49E-10 |
| 16189 | ENSMUSG00000052033 | Pfdn4 | 109054 [Gene Symbol: Pfdn4] [Locus Tag: ] [Chromosome: 2] [Map Location: 2 H3\|2 91 | 3.51892851 | 1.37E-09 | 9.67E-09 |
| 13819 | ENSMUSG00000020516 | Rps6kb1 | 72508 [Gene Symbol: Rps6kb1] [Locus Tag: ] [Chromosome: 11] [Map Location: 11\|11 C | 3.517284985 | 9.32E-13 | 1.08E-11 |
| 15534 | ENSMUSG00000025370 | Cdh9 | 12565 [Gene Symbol: Cdh9] [Locus Tag: ] [Chromosome: 15] [Map Location: 15\|15 A2] | 3.515147373 | 0.004058 | 0.009227 |

| | | | | | | |
|---|---|---|---|---|---|---|
| 638 | ENSMUSG00000019230 | Lhx9 | 16876 [Gene Symbol: Lhx9] [Locus Tag: ] [Chromosome: 1] [Map Location: 1\|1 G-H2] [D | 3.514889429 | 3.46E-09 | 2.31E-08 |
| 17445 | ENSMUSG00000040550 | Otud6b | 72201 [Gene Symbol: Otud6b] [Locus Tag: ] [Chromosome: 4] [Map Location: 4\|4 A1] [D | 3.513382323 | 6.12E-12 | 6.23E-11 |
| 15312 | ENSMUSG00000036309 | Skp1a | 21402 [Gene Symbol: Skp1a] [Locus Tag: ] [Chromosome: 11] [Map Location: 11 B1.3\|1 | 3.513177077 | 4.12E-10 | 3.13E-09 |
| 15633 | ENSMUSG00000008307 | 1700109H08Rik | 77036 [Gene Symbol: 1700109H08Rik] [Locus Tag: ] [Chromosome: 5] [Map Location: 5 | 3.512588206 | 2.48E-06 | 1.04E-05 |
| 10185 | ENSMUSG00000020649 | Rrm2 | 20135 [Gene Symbol: Rrm2] [Locus Tag: ] [Chromosome: 12] [Map Location: 12 A1.3\|1 | 3.511740952 | 2.22E-05 | 7.94E-05 |
| 16737 | ENSMUSG00000056812 | St8sia3 | 20451 [Gene Symbol: St8sia3] [Locus Tag: ] [Chromosome: 18] [Map Location: 18\|18 E | 3.509972657 | 3.10E-12 | 3.28E-11 |
| 18353 | ENSMUSG00000014767 | Tbp | 21374 [Gene Symbol: Tbp] [Locus Tag: ] [Chromosome: 17] [Map Location: 17 A2\|17 8. | 3.509568881 | 1.06E-10 | 8.82E-10 |
| 9645 | ENSMUSG00000029126 | Nsg1 | 18196 [Gene Symbol: Nsg1] [Locus Tag: ] [Chromosome: 5] [Map Location: 5 B3\|5 20.2 | 3.505714078 | 1.63E-12 | 1.81E-11 |
| 15705 | ENSMUSG00000021171 | Esyt2 | 52635 [Gene Symbol: Esyt2] [Locus Tag: ] [Chromosome: 12] [Map Location: 12 F2\|12 6 | 3.501330957 | 6.78E-12 | 6.85E-11 |
| 3142 | ENSMUSG00000028381 | Ugcg | 22234 [Gene Symbol: Ugcg] [Locus Tag: ] [Chromosome: 4] [Map Location: 4 32.44 cM\| | 3.501047313 | 1.45E-12 | 1.62E-11 |
| 15416 | ENSMUSG00000032998 | Foxj3 | 230700 [Gene Symbol: Foxj3] [Locus Tag: ] [Chromosome: 4] [Map Location: 4\|4 D2.1] | 3.500771855 | 2.10E-12 | 2.28E-11 |
| 3130 | ENSMUSG00000029430 | LOC102641332; Ran | 102641332 [Gene Symbol: LOC102641332] [Locus Tag: ] [Chromosome: 9] [Map Locati | 3.499647074 | 4.64E-08 | 2.60E-07 |
| 9295 | ENSMUSG00000039470 | Zdhhc2 | 70546 [Gene Symbol: Zdhhc2] [Locus Tag: ] [Chromosome: 8] [Map Location: 8\|8 A4] [D | 3.493768668 | 6.51E-13 | 7.73E-12 |
| 2136 | ENSMUSG00000054619 | Mettl7a1 | 70152 [Gene Symbol: Mettl7a1] [Locus Tag: ] [Chromosome: 15] [Map Location: 15\|15 | 3.490577635 | 6.47E-05 | 0.000211 |
| 17611 | ENSMUSG00000039770 | Ypel5 | 383295 [Gene Symbol: Ypel5] [Locus Tag: ] [Chromosome: 17] [Map Location: 17\|17 E2 | 3.490477658 | 5.17E-11 | 4.53E-10 |
| 13159 | ENSMUSG00000033964 | Zbtb41 | 226470 [Gene Symbol: Zbtb41] [Locus Tag: ] [Chromosome: 1] [Map Location: 1\|1 F] [D | 3.488273112 | 3.03E-11 | 2.74E-10 |
| 14334 | ENSMUSG00000030629 | Zfand6 | 65098 [Gene Symbol: Zfand6] [Locus Tag: ] [Chromosome: 7] [Map Location: 7\|7 D3] [D | 3.487910422 | 2.67E-10 | 2.09E-09 |
| 16332 | ENSMUSG00000035365 | Parpbp | 75317 [Gene Symbol: Parpbp] [Locus Tag: ] [Chromosome: 10] [Map Location: 10\|10 C | 3.486559766 | 4.63E-06 | 1.85E-05 |
| 16024 | ENSMUSG00000006586 | Runx1t1 | 12395 [Gene Symbol: Runx1t1] [Locus Tag: ] [Chromosome: 4] [Map Location: 4 5.88 cl | 3.485784703 | 3.43E-13 | 4.22E-12 |
| 4145 | ENSMUSG00000021967 | Mrpl57 | 67840 [Gene Symbol: Mrpl57] [Locus Tag: ] [Chromosome: 14] [Map Location: 14\|14 C | 3.483946961 | 2.60E-07 | 1.29E-06 |
| 5051 | ENSMUSG00000029838 | Ptn | 19242 [Gene Symbol: Ptn] [Locus Tag: ] [Chromosome: 6] [Map Location: 6 B1\|6 15.48 | 3.477286768 | 1.38E-12 | 1.55E-11 |
| 3576 | ENSMUSG00000035069 | Oma1 | 67013 [Gene Symbol: Oma1] [Locus Tag: ] [Chromosome: 4] [Map Location: 4\|4 C6] [De | 3.476448198 | 1.37E-07 | 7.11E-07 |
| 3679 | ENSMUSG00000041747 | Utp15 | 105372 [Gene Symbol: Utp15] [Locus Tag: ] [Chromosome: 13] [Map Location: 13\|13 D | 3.476066402 | 1.47E-10 | 1.20E-09 |
| 7772 | ENSMUSG00000028389 | Zfp37 | 22696 [Gene Symbol: Zfp37] [Locus Tag: ] [Chromosome: 4] [Map Location: 4 B3\|4 33.0 | 3.475959682 | 4.02E-11 | 3.59E-10 |
| 18466 | ENSMUSG00000032288 | Imp3 | 102462 [Gene Symbol: Imp3] [Locus Tag: ] [Chromosome: 9] [Map Location: 9\|9 B] [Des | 3.47580377 | 1.95E-07 | 9.85E-07 |
| 6858 | ENSMUSG00000025592 | Dach2 | 93837 [Gene Symbol: Dach2] [Locus Tag: ] [Chromosome: X] [Map Location: X E1\|X 49. | 3.475749476 | 8.97E-07 | 4.08E-06 |
| 15036 | ENSMUSG00000005656 | Snx6 | 72183 [Gene Symbol: Snx6] [Locus Tag: ] [Chromosome: 12] [Map Location: 12\|12 C1] | 3.475334722 | 1.68E-10 | 1.36E-09 |
| 17858 | ENSMUSG00000022634 | Yaf2 | 67057 [Gene Symbol: Yaf2] [Locus Tag: ] [Chromosome: 15] [Map Location: 15\|15 F1] [ | 3.475034219 | 3.41E-10 | 2.63E-09 |
| 7419 | ENSMUSG00000036109 | Mbnl3 | 171170 [Gene Symbol: Mbnl3] [Locus Tag: ] [Chromosome: X] [Map Location: X\|X A5] [ | 3.473511838 | 0.000373 | 0.001056 |
| 5595 | ENSMUSG00000039428 | Tmem135 | 72759 [Gene Symbol: Tmem135] [Locus Tag: ] [Chromosome: 7] [Map Location: 7\|7 E1 | 3.471699279 | 2.49E-11 | 2.29E-10 |
| 12479 | ENSMUSG00000026312 | Cdh7 | 241201 [Gene Symbol: Cdh7] [Locus Tag: ] [Chromosome: 1] [Map Location: 1\|1 E2.1] [ | 3.471563883 | 1.12E-10 | 9.28E-10 |
| 6867 | ENSMUSG00000028759 | Hp1bp3 | 15441 [Gene Symbol: Hp1bp3] [Locus Tag: ] [Chromosome: 4] [Map Location: 4\|4 D3] [ | 3.466773613 | 2.01E-12 | 2.19E-11 |
| 3622 | ENSMUSG00000071302 | 2610044O15Rik8 | 72139 [Gene Symbol: 2610044O15Rik8] [Locus Tag: ] [Chromosome: 8] [Map Location: | 3.46591378 | 2.45E-09 | 1.67E-08 |
| 17478 | ENSMUSG00000032688 | Malt1 | 240354 [Gene Symbol: Malt1] [Locus Tag: ] [Chromosome: 18] [Map Location: 18\|18 E | 3.46477973 | 5.07E-05 | 0.000168 |
| 9187 | ENSMUSG00000031711 | Zfp330 | 30932 [Gene Symbol: Zfp330] [Locus Tag: ] [Chromosome: 8] [Map Location: 8\|8 C2] [D | 3.463538485 | 1.54E-07 | 7.93E-07 |
| 5372 | ENSMUSG00000021546 | Hnrnpk | 15387 [Gene Symbol: Hnrnpk] [Locus Tag: ] [Chromosome: 13] [Map Location: 13\|13 B | 3.463265159 | 4.81E-12 | 4.95E-11 |
| 11482 | ENSMUSG00000019808 | Adat2 | 66757 [Gene Symbol: Adat2] [Locus Tag: ] [Chromosome: 10] [Map Location: 10\|10 A2 | 3.462503023 | 8.78E-05 | 0.000278 |
| 9936 | ENSMUSG00000022008 | Gpalpp1 | 67467 [Gene Symbol: Gpalpp1] [Locus Tag: ] [Chromosome: 14] [Map Location: 14\|14 | 3.459635534 | 1.23E-10 | 1.02E-09 |
| 11674 | ENSMUSG00000017843 | Ppp2r5c | 26931 [Gene Symbol: Ppp2r5c] [Locus Tag: ] [Chromosome: 12] [Map Location: 12 F1\| | 3.45700998 | 2.92E-10 | 2.28E-09 |
| 4418 | ENSMUSG00000024505 | Dtwd2 | 68857 [Gene Symbol: Dtwd2] [Locus Tag: ] [Chromosome: 18] [Map Location: 18\|18 C] | 3.454943177 | 1.37E-06 | 6.05E-06 |
| 13984 | ENSMUSG00000025935 | Tram1 | 72265 [Gene Symbol: Tram1] [Locus Tag: ] [Chromosome: 1] [Map Location: 1\|1 A3] [De | 3.453394934 | 2.41E-10 | 1.91E-09 |
| 1069 | ENSMUSG00000021952 | Xpo4 | 57258 [Gene Symbol: Xpo4] [Locus Tag: ] [Chromosome: 14] [Map Location: 14\|14 C3] | 3.452572853 | 2.63E-10 | 2.07E-09 |
| 5491 | ENSMUSG00000027001 | Dusp19 | 68082 [Gene Symbol: Dusp19] [Locus Tag: ] [Chromosome: 2] [Map Location: 2\|2 C3] [ | 3.452168268 | 0.000191 | 0.000571 |
| 2063 | ENSMUSG00000063362 | Alg11 | 207958 [Gene Symbol: Alg11] [Locus Tag: ] [Chromosome: 8] [Map Location: 8\|8 A2] [D | 3.450974202 | 8.99E-11 | 7.58E-10 |
| 6032 | ENSMUSG00000029130 | Rnf32 | 56874 [Gene Symbol: Rnf32] [Locus Tag: ] [Chromosome: 5] [Map Location: 5 B1\|5 14. | 3.450075808 | 8.01E-10 | 5.86E-09 |
| 960 | ENSMUSG00000034532 | Fbxo16 | 50759 [Gene Symbol: Fbxo16] [Locus Tag: ] [Chromosome: 14] [Map Location: 14\|14 D | 3.449741853 | 2.00E-07 | 1.01E-06 |
| 6131 | ENSMUSG00000032002 | Dcun1d5 | 76863 [Gene Symbol: Dcun1d5] [Locus Tag: ] [Chromosome: 9] [Map Location: 9\|9 A1] | 3.448825376 | 1.65E-10 | 1.34E-09 |
| 8197 | ENSMUSG00000048661 | Lemd3 | 380664 [Gene Symbol: Lemd3] [Locus Tag: ] [Chromosome: 10] [Map Location: 10\|10 | 3.448183317 | 5.89E-12 | 6.00E-11 |
| 8085 | ENSMUSG00000020801 | Med31 | 67279 [Gene Symbol: Med31] [Locus Tag: ] [Chromosome: 11] [Map Location: 11\|11 B | 3.447855191 | 3.82E-07 | 1.84E-06 |
| 1589 | ENSMUSG00000020950 | Foxg1 | 15228 [Gene Symbol: Foxg1] [Locus Tag: ] [Chromosome: 12] [Map Location: 12 B3\|12 | 3.447747781 | 6.49E-10 | 4.80E-09 |
| 8816 | ENSMUSG00000026834 | Acvr1c | 269275 [Gene Symbol: Acvr1c] [Locus Tag: ] [Chromosome: 2] [Map Location: 2\|2 C1.1 | 3.4466658 | 4.84E-08 | 2.70E-07 |
| 12921 | ENSMUSG00000024059 | Clip4 | 78785 [Gene Symbol: Clip4] [Locus Tag: ] [Chromosome: 17] [Map Location: 17\|17 E2] | 3.44661352 | 7.58E-11 | 6.48E-10 |
| 11011 | ENSMUSG00000042599 | Kdm7a | 338523 [Gene Symbol: Kdm7a] [Locus Tag: ] [Chromosome: 6] [Map Location: 6\|6 B1] [ | 3.445404739 | 1.55E-08 | 9.38E-08 |
| 3645 | ENSMUSG00000025759 | Mfsd8 | 72175 [Gene Symbol: Mfsd8] [Locus Tag: ] [Chromosome: 3] [Map Location: 3\|3 C] [Des | 3.442542777 | 6.78E-08 | 3.69E-07 |
| 6400 | ENSMUSG00000041135 | Ripk2 | 192656 [Gene Symbol: Ripk2] [Locus Tag: ] [Chromosome: 4] [Map Location: 4 A2\|4 6. | 3.442432144 | 5.05E-07 | 2.39E-06 |
| 15322 | ENSMUSG00000025024 | Smndc1 | 76479 [Gene Symbol: Smndc1] [Locus Tag: ] [Chromosome: 19] [Map Location: 19\|19 | 3.441591216 | 9.04E-10 | 6.56E-09 |
| 15420 | ENSMUSG00000070527 | Mkrn3 | 22652 [Gene Symbol: Mkrn3] [Locus Tag: ] [Chromosome: 7] [Map Location: 7 C\|7 34.3 | 3.440346039 | 2.48E-10 | 1.96E-09 |
| 7763 | ENSMUSG00000078958 | Atp6ap1l | 435376 [Gene Symbol: Atp6ap1l] [Locus Tag: ] [Chromosome: 13] [Map Location: 13\|1 | 3.440311907 | 0.000961 | 0.002507 |
| 17969 | ENSMUSG00000020248 | Nfyb | 18045 [Gene Symbol: Nfyb] [Locus Tag: ] [Chromosome: 10] [Map Location: 10 40.25 cl | 3.43596053 | 5.70E-09 | 3.70E-08 |
| 9726 | ENSMUSG00000024477 | Pggt1b | 225467 [Gene Symbol: Pggt1b] [Locus Tag: ] [Chromosome: 18] [Map Location: 18\|18 | 3.43567889 | 8.74E-10 | 6.37E-09 |
| 16513 | ENSMUSG00000023921 | Mut | 17850 [Gene Symbol: Mut] [Locus Tag: ] [Chromosome: 17] [Map Location: 17 19.55 cM | 3.43450365 | 5.31E-08 | 2.94E-07 |
| 5223 | ENSMUSG00000042708 | Shcbp1l | 71836 [Gene Symbol: Shcbp1l] [Locus Tag: ] [Chromosome: 1] [Map Location: 1\|1 G3] [ | 3.433971134 | 0.001745 | 0.00431 |
| 11450 | ENSMUSG00000022056 | Adam7 | 11500 [Gene Symbol: Adam7] [Locus Tag: ] [Chromosome: 14] [Map Location: 14\|14 D | 3.433787835 | 0.002731 | 0.006466 |
| 11681 | ENSMUSG00000003541 | Ier3 | 15937 [Gene Symbol: Ier3] [Locus Tag: ] [Chromosome: 17] [Map Location: 17\|17 B1] [ | 3.431888238 | 0.01684 | 0.033076 |
| 14640 | ENSMUSG00000021139 | Synj2bp-cox16 | 105940408 [Gene Symbol: Synj2bp-cox16] [Locus Tag: ] [Chromosome: 12] [Map Locat | 3.424121136 | 3.96E-12 | 4.13E-11 |
| 4737 | ENSMUSG00000094870 | Zfp131 | 72465 [Gene Symbol: Zfp131] [Locus Tag: ] [Chromosome: 13] [Map Location: 13\|13] [ | 3.423588938 | 1.13E-10 | 9.39E-10 |
| 17577 | ENSMUSG00000071662 | Polr2g | 67710 [Gene Symbol: Polr2g] [Locus Tag: ] [Chromosome: 19] [Map Location: 19\|19 A] | 3.418654381 | 8.00E-08 | 4.31E-07 |
| 17667 | ENSMUSG00000022865 | Cxadr | 13052 [Gene Symbol: Cxadr] [Locus Tag: ] [Chromosome: 16] [Map Location: 16\|16 C] [ | 3.417936796 | 2.16E-12 | 2.34E-11 |
| 6303 | ENSMUSG00000042340 | Ctf1 | 13019 [Gene Symbol: Ctf1] [Locus Tag: ] [Chromosome: 7] [Map Location: 7\|7 F3] [Desc | 3.416376624 | 0.007994 | 0.017024 |
| 15214 | ENSMUSG00000052299 | Ltn1 | 78913 [Gene Symbol: Ltn1] [Locus Tag: ] [Chromosome: 16] [Map Location: 16\|16 C3.3 | 3.4154918 | 6.37E-11 | 5.52E-10 |
| 14996 | ENSMUSG00000043262 | Uevld | 54122 [Gene Symbol: Uevld] [Locus Tag: ] [Chromosome: 7] [Map Location: 7\|7 B4] [De | 3.415268448 | 2.21E-06 | 9.40E-06 |
| 926 | ENSMUSG00000018509 | Cenpv | 73139 [Gene Symbol: Cenpv] [Locus Tag: ] [Chromosome: 11] [Map Location: 11\|11 B2 | 3.414515387 | 5.23E-08 | 2.90E-07 |
| 15711 | ENSMUSG00000025899 | Alkbh8 | 67667 [Gene Symbol: Alkbh8] [Locus Tag: ] [Chromosome: 9] [Map Location: 9\|9 A1] [D | 3.414269155 | 1.77E-11 | 1.67E-10 |
| 10904 | ENSMUSG00000016496 | Cd274 | 60533 [Gene Symbol: Cd274] [Locus Tag: ] [Chromosome: 19] [Map Location: 19\|19 C2 | 3.41419386 | 4.04E-05 | 0.000137 |
| 4031 | ENSMUSG00000050447 | Lypd6 | 320343 [Gene Symbol: Lypd6] [Locus Tag: ] [Chromosome: 2] [Map Location: 2\|2 C1.1] | 3.413584264 | 7.90E-11 | 6.74E-10 |
| 6540 | ENSMUSG00000019916 | P4ha1 | 18451 [Gene Symbol: P4ha1] [Locus Tag: ] [Chromosome: 10] [Map Location: 10\|10 B4 | 3.412476273 | 1.60E-09 | 1.12E-08 |
| 15661 | ENSMUSG00000014773 | Dll1 | 13388 [Gene Symbol: Dll1] [Locus Tag: ] [Chromosome: 17] [Map Location: 17 A2\|17 8. | 3.412139077 | 1.64E-05 | 6.01E-05 |
| 9122 | ENSMUSG00000026616 | Cr2 | 12902 [Gene Symbol: Cr2] [Locus Tag: ] [Chromosome: 1] [Map Location: 1 H6\|1 98.44 | 3.411819099 | 6.37E-07 | 2.97E-06 |
| 13641 | ENSMUSG00000028557 | Rnf11 | 29864 [Gene Symbol: Rnf11] [Locus Tag: ] [Chromosome: 4] [Map Location: 4\|4 C7] [De | 3.410254564 | 8.41E-11 | 7.13E-10 |
| 6374 | ENSMUSG00000021576 | Pdcd6 | 18570 [Gene Symbol: Pdcd6] [Locus Tag: ] [Chromosome: 13] [Map Location: 13\|13 C1 | 3.409846541 | 1.04E-10 | 8.67E-10 |
| 15370 | ENSMUSG00000025340 | Rabgef1 | 56715 [Gene Symbol: Rabgef1] [Locus Tag: ] [Chromosome: 5] [Map Location: 5\|5 F] [De | 3.407592112 | 4.07E-08 | 2.31E-07 |
| 7484 | ENSMUSG00000038630 | Zkscan16 | 100041581 [Gene Symbol: Zkscan16] [Locus Tag: ] [Chromosome: 4] [Map Location: 4\| | 3.403171682 | 3.33E-06 | 1.37E-05 |
| 17383 | ENSMUSG00000037994 | Slc9b2 | 97086 [Gene Symbol: Slc9b2] [Locus Tag: ] [Chromosome: 3] [Map Location: 3\|3 G3] [D | 3.402630649 | 3.60E-05 | 0.000123 |
| 16234 | ENSMUSG00000032373 | Car12 | 76459 [Gene Symbol: Car12] [Locus Tag: ] [Chromosome: 9] [Map Location: 9\|9 C] [Des | 3.40222941 | 3.04E-06 | 1.26E-05 |

| | | | | | | |
|---|---|---|---|---|---|---|
| 4894 | ENSMUSG00000025436 | Xrcc6bp1 | 68876 [Gene Symbol: Xrcc6bp1] [Locus Tag: ] [Chromosome: 10] [Map Location: 10\|10 | 3.401988607 | 2.55E-07 | 1.27E-06 |
| 17361 | ENSMUSG00000020949 | Fkbp3 | 30795 [Gene Symbol: Fkbp3] [Locus Tag: ] [Chromosome: 12] [Map Location: 12\|12 C3 | 3.399413568 | 2.10E-10 | 1.67E-09 |
| 3271 | ENSMUSG00000021127 | Zfp36l1 | 12192 [Gene Symbol: Zfp36l1] [Locus Tag: ] [Chromosome: 12] [Map Location: 12\|12 C | 3.398836966 | 1.30E-10 | 1.07E-09 |
| 7871 | ENSMUSG00000022893 | Adamts1 | 11504 [Gene Symbol: Adamts1] [Locus Tag: ] [Chromosome: 16] [Map Location: 16 48.2 | 3.398160258 | 1.56E-08 | 9.42E-08 |
| 8091 | ENSMUSG00000026740 | Dnajc1 | 13418 [Gene Symbol: Dnajc1] [Locus Tag: ] [Chromosome: 2] [Map Location: 2\|2 A2] [D | 3.397634844 | 4.73E-12 | 4.88E-11 |
| 3319 | ENSMUSG00000034522 | Zfp395 | 380912 [Gene Symbol: Zfp395] [Locus Tag: ] [Chromosome: 14] [Map Location: 14\|14 D | 3.397526768 | 1.92E-10 | 1.54E-09 |
| 8744 | ENSMUSG00000029840 | Mtpn | 14489 [Gene Symbol: Mtpn] [Locus Tag: ] [Chromosome: 6] [Map Location: 6\|6 B1] [De | 3.39603678 | 2.36E-12 | 2.54E-11 |
| 3194 | ENSMUSG00000030549 | Rhcg | 56315 [Gene Symbol: Rhcg] [Locus Tag: ] [Chromosome: 7] [Map Location: 7 D3\|7 45.0 | 3.395816084 | 0.001071 | 0.00277 |
| 16248 | ENSMUSG00000073176 | Zfp449 | 78619 [Gene Symbol: Zfp449] [Locus Tag: ] [Chromosome: X] [Map Location: X\|X A4] [D | 3.3956858 | 6.11E-07 | 2.85E-06 |
| 2482 | ENSMUSG00000090112 | Shprh | 268281 [Gene Symbol: Shprh] [Locus Tag: ] [Chromosome: 10] [Map Location: 10\|10 A | 3.394962069 | 3.63E-11 | 3.26E-10 |
| 14910 | ENSMUSG00000031262 | Cenpi | 102920 [Gene Symbol: Cenpi] [Locus Tag: ] [Chromosome: X] [Map Location: X\|X E3] [D | 3.394356378 | 3.88E-05 | 0.000132 |
| 6902 | ENSMUSG00000070520 | Ndnl2 | 66647 [Gene Symbol: Ndnl2] [Locus Tag: ] [Chromosome: 7] [Map Location: 7\|7 C] [Des | 3.394203687 | 7.29E-09 | 4.65E-08 |
| 17550 | ENSMUSG00000034928 | Rnf44 | 105239 [Gene Symbol: Rnf44] [Locus Tag: ] [Chromosome: 13] [Map Location: 13\|13 B | 3.393838961 | 8.49E-10 | 6.19E-09 |
| 1498 | ENSMUSG00000029863 | Casp2 | 12366 [Gene Symbol: Casp2] [Locus Tag: ] [Chromosome: 6] [Map Location: 6 B2.1\|6 20 | 3.391545784 | 6.48E-08 | 3.54E-07 |
| 18550 | ENSMUSG00000031990 | Jam3 | 83964 [Gene Symbol: Jam3] [Locus Tag: ] [Chromosome: 9] [Map Location: 9\|9 A4] [Des | 3.389930473 | 1.06E-08 | 6.63E-08 |
| 4964 | ENSMUSG00000028496 | Mllt3 | 70122 [Gene Symbol: Mllt3] [Locus Tag: ] [Chromosome: 4] [Map Location: 4 C4\|4 41.0 | 3.389909646 | 6.48E-11 | 5.61E-10 |
| 276 | ENSMUSG00000026384 | Ptpn4 | 19258 [Gene Symbol: Ptpn4] [Locus Tag: ] [Chromosome: 1] [Map Location: 1 E2\|1 52.3 | 3.38814456 | 6.71E-09 | 4.31E-08 |
| 12168 | ENSMUSG00000040729 | Cep126 | 234915 [Gene Symbol: Cep126] [Locus Tag: ] [Chromosome: 9] [Map Location: 9\|9 A1] | 3.386753007 | 6.57E-07 | 3.05E-06 |
| 2916 | ENSMUSG00000019814 | Ltv1 | 353258 [Gene Symbol: Ltv1] [Locus Tag: ] [Chromosome: 10] [Map Location: 10\|10 A2] | 3.38501607 | 5.57E-08 | 3.07E-07 |
| 16062 | ENSMUSG00000026158 | Ogfrl1 | 70155 [Gene Symbol: Ogfrl1] [Locus Tag: ] [Chromosome: 1] [Map Location: 1\|1 A5] [De | 3.38398241 | 1.60E-09 | 1.12E-08 |
| 14657 | ENSMUSG00000035505 | Cox18 | 231430 [Gene Symbol: Cox18] [Locus Tag: ] [Chromosome: 5] [Map Location: 5\|5 E1] [D | 3.38218095 | 7.00E-07 | 3.24E-06 |
| 13402 | ENSMUSG00000090626 | Tex9 | 21778 [Gene Symbol: Tex9] [Locus Tag: ] [Chromosome: 9] [Map Location: 9\|9 D] [Desc | 3.379692732 | 3.95E-10 | 3.02E-09 |
| 4611 | ENSMUSG00000004980 | LOC102642938; Hnrr | 102642938 [Gene Symbol: LOC102642938] [Locus Tag: ] [Chromosome: Un] [Map Loca | 3.378166028 | 3.08E-12 | 3.26E-11 |
| 3067 | ENSMUSG00000015536 | Mocs2 | 17434 [Gene Symbol: Mocs2] [Locus Tag: ] [Chromosome: 13] [Map Location: 13\|13 D2 | 3.377402843 | 2.77E-08 | 1.61E-07 |
| 6672 | ENSMUSG00000005672 | Kit | 16590 [Gene Symbol: Kit] [Locus Tag: ] [Chromosome: 5] [Map Location: 5 C3.3\|5 39.5 | 3.376750399 | 4.86E-11 | 4.28E-10 |
| 14828 | ENSMUSG00000026721 | Rabgap1l | 29809 [Gene Symbol: Rabgap1l] [Locus Tag: ] [Chromosome: 1] [Map Location: 1\|1 H2. | 3.375305559 | 2.00E-12 | 2.18E-11 |
| 3351 | ENSMUSG00000024712 | Rfk | 54391 [Gene Symbol: Rfk] [Locus Tag: ] [Chromosome: 19] [Map Location: 19\|19 B] [De | 3.374608028 | 6.36E-10 | 4.71E-09 |
| 17942 | ENSMUSG00000039117 | Taf4a | 228980 [Gene Symbol: Taf4a] [Locus Tag: ] [Chromosome: 2] [Map Location: 2\|2 H4] [D | 3.374225594 | 3.16E-10 | 2.45E-09 |
| 14243 | ENSMUSG00000029563 | Foxp2 | 114142 [Gene Symbol: Foxp2] [Locus Tag: ] [Chromosome: 6] [Map Location: 6\|6 A2] [D | 3.373659029 | 3.01E-12 | 3.19E-11 |
| 14559 | ENSMUSG00000052812 | Atad2b | 320817 [Gene Symbol: Atad2b] [Locus Tag: ] [Chromosome: 12] [Map Location: 12\|12 A | 3.372129839 | 4.45E-09 | 2.93E-08 |
| 758 | ENSMUSG00000046169 | Adamts6 | 108154 [Gene Symbol: Adamts6] [Locus Tag: ] [Chromosome: 13] [Map Location: 13\|13 | 3.366477222 | 4.97E-08 | 2.77E-07 |
| 8256 | ENSMUSG00000034480 | Diaph2 | 54004 [Gene Symbol: Diaph2] [Locus Tag: ] [Chromosome: X] [Map Location: X\|X E3] [D | 3.366064159 | 3.51E-09 | 2.34E-08 |
| 18404 | ENSMUSG00000020577 | Tspan13 | 66109 [Gene Symbol: Tspan13] [Locus Tag: ] [Chromosome: 12] [Map Location: 12\|12 B | 3.365677458 | 5.86E-11 | 5.10E-10 |
| 9571 | ENSMUSG00000033486 | Catsper2 | 212670 [Gene Symbol: Catsper2] [Locus Tag: ] [Chromosome: 2] [Map Location: 2\|2 E5 | 3.365148028 | 0.000122 | 0.000376 |
| 11981 | ENSMUSG00000025505 | Tmem80 | 71448 [Gene Symbol: Tmem80] [Locus Tag: ] [Chromosome: 7] [Map Location: 7\|7 F5] [ | 3.364981559 | 1.56E-06 | 6.84E-06 |
| 17632 | ENSMUSG00000078578 | Ube2d3 | 66105 [Gene Symbol: Ube2d3] [Locus Tag: ] [Chromosome: 3] [Map Location: 3\|3 G3] [ | 3.363956829 | 5.72E-12 | 5.84E-11 |
| 8506 | ENSMUSG00000048047 | Zbtb33 | 56805 [Gene Symbol: Zbtb33] [Locus Tag: ] [Chromosome: X] [Map Location: X\|X A3.3] | 3.363096622 | 1.35E-10 | 1.11E-09 |
| 2571 | ENSMUSG00000051455 | Meioc | 268491 [Gene Symbol: Meioc] [Locus Tag: ] [Chromosome: 11] [Map Location: 11\|11 E | 3.356801867 | 0.011152 | 0.022971 |
| 623 | ENSMUSG00000028035 | Dnajb4 | 67035 [Gene Symbol: Dnajb4] [Locus Tag: ] [Chromosome: 3] [Map Location: 3\|3 H3] [D | 3.356384454 | 1.58E-07 | 8.13E-07 |
| 15622 | ENSMUSG00000041624 | Gucy1a2 | 234889 [Gene Symbol: Gucy1a2] [Locus Tag: ] [Chromosome: 9] [Map Location: 9\|9 A1 | 3.35422736 | 5.06E-12 | 5.19E-11 |
| 5899 | ENSMUSG00000035694 | Caps2 | 353025 [Gene Symbol: Caps2] [Locus Tag: ] [Chromosome: 10] [Map Location: 10\|10 D | 3.354065527 | 0.011394 | 0.023415 |
| 10293 | ENSMUSG00000024354 | Slc23a1 | 20522 [Gene Symbol: Slc23a1] [Locus Tag: ] [Chromosome: 18] [Map Location: 18 B2\|1 | 3.352651851 | 4.93E-10 | 3.72E-09 |
| 14135 | ENSMUSG00000052504 | Epha3 | 13837 [Gene Symbol: Epha3] [Locus Tag: ] [Chromosome: 16] [Map Location: 16\|16 C1 | 3.352273044 | 1.03E-06 | 4.61E-06 |
| 14851 | ENSMUSG00000024158 | Hagh | 14651 [Gene Symbol: Hagh] [Locus Tag: ] [Chromosome: 17] [Map Location: 17 A3.3\|17 | 3.352136387 | 2.87E-09 | 1.94E-08 |
| 15447 | ENSMUSG00000063663 | Brwd3 | 382236 [Gene Symbol: Brwd3] [Locus Tag: ] [Chromosome: X] [Map Location: X\|X D] [D | 3.351629662 | 2.76E-10 | 2.16E-09 |
| 18515 | ENSMUSG00000017485 | Top2b | 21974 [Gene Symbol: Top2b] [Locus Tag: ] [Chromosome: 14] [Map Location: 14\|14 A3 | 3.350507124 | 7.58E-11 | 6.48E-10 |
| 9285 | ENSMUSG00000014932 | Yes1 | 22612 [Gene Symbol: Yes1] [Locus Tag: ] [Chromosome: 5] [Map Location: 5 B1\|5 17.33 | 3.348951066 | 1.48E-10 | 1.21E-09 |
| 4208 | ENSMUSG00000028647 | Mycbp | 56309 [Gene Symbol: Mycbp] [Locus Tag: ] [Chromosome: 4] [Map Location: 4\|4 D2.2] | 3.347592583 | 1.27E-07 | 6.66E-07 |
| 11607 | ENSMUSG00000022827 | Rabl3 | 67657 [Gene Symbol: Rabl3] [Locus Tag: ] [Chromosome: 16] [Map Location: 16\|16 B3] | 3.346683417 | 4.24E-05 | 0.000143 |
| 2749 | ENSMUSG00000040177 | 2310057M21Rik | 68277 [Gene Symbol: 2310057M21Rik] [Locus Tag: ] [Chromosome: 7] [Map Location: | 3.346244495 | 2.55E-08 | 1.49E-07 |
| 12703 | ENSMUSG00000007908 | Hmgcll1 | 208982 [Gene Symbol: Hmgcll1] [Locus Tag: ] [Chromosome: 9] [Map Location: 9\|9 D] [ | 3.345956554 | 1.46E-06 | 6.43E-06 |
| 5194 | ENSMUSG00000025956 | Mettl21a | 67099 [Gene Symbol: Mettl21a] [Locus Tag: ] [Chromosome: 1] [Map Location: 1\|1 C2] | 3.344016604 | 2.76E-06 | 1.15E-05 |
| 8263 | ENSMUSG00000040254 | Sema3d | 108151 [Gene Symbol: Sema3d] [Locus Tag: ] [Chromosome: 5] [Map Location: 5\|5 A2] | 3.342783473 | 9.66E-06 | 3.68E-05 |
| 4194 | ENSMUSG00000034218 | Atm | 11920 [Gene Symbol: Atm] [Locus Tag: ] [Chromosome: 9] [Map Location: 9 29.12 cM\|9 | 3.341394366 | 1.96E-08 | 1.17E-07 |
| 6141 | ENSMUSG00000039478 | Micu3 | 78506 [Gene Symbol: Micu3] [Locus Tag: ] [Chromosome: 8] [Map Location: 8\|8 A4] [De | 3.33807154 | 6.51E-08 | 3.56E-07 |
| 4353 | ENSMUSG00000032368 | Zic1 | 22771 [Gene Symbol: Zic1] [Locus Tag: ] [Chromosome: 9] [Map Location: 9 E3.3\|9 48.2 | 3.337971057 | 6.67E-12 | 6.75E-11 |
| 6052 | ENSMUSG00000028412 | Slc44a1 | 100434 [Gene Symbol: Slc44a1] [Locus Tag: ] [Chromosome: 4] [Map Location: 4\|4 B2] | 3.334655095 | 2.79E-11 | 2.54E-10 |
| 3946 | ENSMUSG00000025162 | Csnk1d | 104318 [Gene Symbol: Csnk1d] [Locus Tag: ] [Chromosome: 11] [Map Location: 11\|11 | 3.333120079 | 4.36E-11 | 3.87E-10 |
| 12209 | ENSMUSG00000027630 | Tbl1xr1 | 81004 [Gene Symbol: Tbl1xr1] [Locus Tag: ] [Chromosome: 3] [Map Location: 3\|3 A3] [D | 3.332958691 | 6.31E-12 | 6.40E-11 |
| 12000 | ENSMUSG00000002617 | Zfp40 | 22700 [Gene Symbol: Zfp40] [Locus Tag: ] [Chromosome: 17] [Map Location: 17 A3.3\|1 | 3.332748738 | 1.23E-08 | 7.58E-08 |
| 17345 | ENSMUSG00000022765 | Snap29 | 67474 [Gene Symbol: Snap29] [Locus Tag: ] [Chromosome: 16] [Map Location: 16\|16 A | 3.332548986 | 2.95E-10 | 2.30E-09 |
| 16921 | ENSMUSG00000032405 | Pias1 | 56469 [Gene Symbol: Pias1] [Locus Tag: ] [Chromosome: 9] [Map Location: 9\|9 B] [Desc | 3.332100276 | 5.03E-11 | 4.42E-10 |
| 6174 | ENSMUSG00000005034 | Prkacb | 18749 [Gene Symbol: Prkacb] [Locus Tag: ] [Chromosome: 3] [Map Location: 3\|3 H3] [D | 3.331975602 | 2.52E-12 | 2.70E-11 |
| 17728 | ENSMUSG00000047757 | Fancb | 237211 [Gene Symbol: Fancb] [Locus Tag: ] [Chromosome: X] [Map Location: X\|X F5] [D | 3.331783071 | 0.000348 | 0.000991 |
| 12292 | ENSMUSG00000022365 | Derl1 | 67819 [Gene Symbol: Derl1] [Locus Tag: ] [Chromosome: 15] [Map Location: 15\|15 D2] | 3.330432826 | 5.55E-07 | 2.61E-06 |
| 4837 | ENSMUSG00000045106 | Ccdc73 | 211936 [Gene Symbol: Ccdc73] [Locus Tag: ] [Chromosome: 2] [Map Location: 2\|2 E2] [ | 3.330148511 | 5.31E-10 | 3.98E-09 |
| 971 | ENSMUSG00000100235 | Zfp708 | 432769 [Gene Symbol: Zfp708] [Locus Tag: ] [Chromosome: 13] [Map Location: 13\|13 | 3.328832301 | 3.67E-08 | 2.09E-07 |
| 10107 | ENSMUSG00000019988 | Nedd1 | 17997 [Gene Symbol: Nedd1] [Locus Tag: ] [Chromosome: 10] [Map Location: 10\|10 C2 | 3.326285878 | 1.50E-06 | 6.59E-06 |
| 15302 | ENSMUSG00000040044 | Orc3 | 50793 [Gene Symbol: Orc3] [Locus Tag: ] [Chromosome: 4] [Map Location: 4\|4 A3] [Des | 3.326180303 | 9.48E-12 | 9.35E-11 |
| 14884 | ENSMUSG00000074802 | Gas2l3 | 237436 [Gene Symbol: Gas2l3] [Locus Tag: ] [Chromosome: 10] [Map Location: 10\|10 C | 3.326030354 | 1.52E-08 | 9.23E-08 |
| 1480 | ENSMUSG00000025019 | Lcor | 212391 [Gene Symbol: Lcor] [Locus Tag: ] [Chromosome: 19] [Map Location: 19\|19 C3] | 3.325919606 | 9.20E-09 | 5.80E-08 |
| 5665 | ENSMUSG00000011884 | Gltp | 56356 [Gene Symbol: Gltp] [Locus Tag: ] [Chromosome: 5] [Map Location: 5\|5 F] [Descr | 3.323964603 | 1.42E-08 | 8.67E-08 |
| 17299 | ENSMUSG00000020427 | Igfbp3 | 16009 [Gene Symbol: Igfbp3] [Locus Tag: ] [Chromosome: 11] [Map Location: 11 A1\|11 | 3.323633322 | 1.20E-09 | 8.54E-09 |
| 4335 | ENSMUSG00000038816 | Ctnnal1 | 54366 [Gene Symbol: Ctnnal1] [Locus Tag: ] [Chromosome: 4] [Map Location: 4\|4 C1] [ | 3.322616543 | 9.18E-09 | 5.79E-08 |
| 2097 | ENSMUSG00000010554 | Mettl16 | 67493 [Gene Symbol: Mettl16] [Locus Tag: ] [Chromosome: 11] [Map Location: 11\|11 E | 3.322173 | 5.56E-08 | 3.07E-07 |
| 5932 | ENSMUSG00000030934 | Oat | 18242 [Gene Symbol: Oat] [Locus Tag: ] [Chromosome: 7] [Map Location: 7 F3\|7 76.3 c | 3.322157206 | 9.02E-08 | 4.82E-07 |
| 18287 | ENSMUSG00000032221 | Mns1 | 17427 [Gene Symbol: Mns1] [Locus Tag: ] [Chromosome: 9] [Map Location: 9\|9 D] [Desc | 3.320632035 | 1.81E-09 | 1.25E-08 |
| 17782 | ENSMUSG00000022450 | Ndufa6 | 67130 [Gene Symbol: Ndufa6] [Locus Tag: ] [Chromosome: 15] [Map Location: 15\|15 E | 3.317790267 | 1.76E-09 | 1.22E-08 |
| 18520 | ENSMUSG00000021540 | Smad5 | 17129 [Gene Symbol: Smad5] [Locus Tag: ] [Chromosome: 13] [Map Location: 13 B1\|13 | 3.317605712 | 6.88E-08 | 3.74E-07 |
| 6176 | ENSMUSG00000026849 | Tor1a | 30931 [Gene Symbol: Tor1a] [Locus Tag: ] [Chromosome: 2] [Map Location: 2\|2 B] [Desc | 3.316281779 | 4.48E-11 | 3.97E-10 |
| 16438 | ENSMUSG00000022360 | Atad2 | 70472 [Gene Symbol: Atad2] [Locus Tag: ] [Chromosome: 15] [Map Location: 15\|15 D2 | 3.31508318 | 3.41E-06 | 1.40E-05 |
| 11168 | ENSMUSG00000020034 | Tcp11l2 | 216198 [Gene Symbol: Tcp11l2] [Locus Tag: ] [Chromosome: 10] [Map Location: 10\|10 | 3.313591247 | 3.52E-06 | 1.44E-05 |
| 11622 | ENSMUSG00000045427 | Hnrnph2 | 56258 [Gene Symbol: Hnrnph2] [Locus Tag: ] [Chromosome: X] [Map Location: X E3\|X 5 | 3.313577876 | 1.28E-07 | 6.72E-07 |
| 14371 | ENSMUSG00000053819 | Camk2d | 108058 [Gene Symbol: Camk2d] [Locus Tag: ] [Chromosome: 3] [Map Location: 3\|3 G1- | 3.313147418 | 2.89E-07 | 1.42E-06 |

| | | | | | | |
|---|---|---|---|---|---|---|
| 11129 | ENSMUSG00000047789 | Slc38a9 | 268706 [Gene Symbol: Slc38a9] [Locus Tag: ] [Chromosome: 13] [Map Location: 13\|13 | 3.311942268 | 1.29E-08 | 7.95E-08 |
| 7842 | ENSMUSG00000049488 | Tmem67 | 329795 [Gene Symbol: Tmem67] [Locus Tag: ] [Chromosome: 4] [Map Location: 4\|4 A1 | 3.310342551 | 1.34E-07 | 6.98E-07 |
| 16899 | ENSMUSG00000079470 | Utp14b | 195434 [Gene Symbol: Utp14b] [Locus Tag: ] [Chromosome: 1] [Map Location: 1 C4\|1 4 | 3.309405422 | 7.33E-10 | 5.39E-09 |
| 18526 | ENSMUSG00000030279 | C2cd5 | 74741 [Gene Symbol: C2cd5] [Locus Tag: ] [Chromosome: 6] [Map Location: 6\|6 G2] [De | 3.309278759 | 1.09E-08 | 6.79E-08 |
| 8318 | ENSMUSG00000072955 | Tmsb15l | 399591 [Gene Symbol: Tmsb15l] [Locus Tag: ] [Chromosome: X] [Map Location: X\|X F1] | 3.309253604 | 0.001788 | 0.004404 |
| 2039 | ENSMUSG00000039456 | Morc3 | 338467 [Gene Symbol: Morc3] [Locus Tag: ] [Chromosome: 16] [Map Location: 16\|16 C | 3.309234431 | 7.12E-11 | 6.11E-10 |
| 2215 | ENSMUSG00000040584 | Abcb1a | 18671 [Gene Symbol: Abcb1a] [Locus Tag: ] [Chromosome: 5] [Map Location: 5 A1\|5 3.4 | 3.308176326 | 2.82E-05 | 9.88E-05 |
| 18502 | ENSMUSG00000021987 | Mtmr6 | 219135 [Gene Symbol: Mtmr6] [Locus Tag: ] [Chromosome: 14] [Map Location: 14\|14 D | 3.307772897 | 2.05E-11 | 1.92E-10 |
| 13586 | ENSMUSG00000031754 | Nudt21 | 68219 [Gene Symbol: Nudt21] [Locus Tag: ] [Chromosome: 8] [Map Location: 8\|8 C5] [D | 3.307318567 | 1.38E-06 | 6.09E-06 |
| 7355 | ENSMUSG00000017376 | Nlk | 18099 [Gene Symbol: Nlk] [Locus Tag: ] [Chromosome: 11] [Map Location: 11\|11 B5] [D | 3.304498381 | 7.26E-11 | 6.22E-10 |
| 9543 | ENSMUSG00000045991 | Onecut2 | 225631 [Gene Symbol: Onecut2] [Locus Tag: ] [Chromosome: 18] [Map Location: 18\|18 | 3.30376519 | 5.57E-08 | 3.08E-07 |
| 11189 | ENSMUSG00000028580 | Pum1 | 80912 [Gene Symbol: Pum1] [Locus Tag: ] [Chromosome: 4] [Map Location: 4\|4 D2.2-D | 3.303493036 | 1.13E-11 | 1.10E-10 |
| 16322 | ENSMUSG00000028882 | Ppp1r8 | 100336 [Gene Symbol: Ppp1r8] [Locus Tag: ] [Chromosome: 4] [Map Location: 4\|4 D2.3 | 3.303359794 | 1.09E-10 | 9.05E-10 |
| 18588 | ENSMUSG00000038594 | Cep85l | 100038725 [Gene Symbol: Cep85l] [Locus Tag: ] [Chromosome: 10] [Map Location: 10\| | 3.300579467 | 1.12E-10 | 9.31E-10 |
| 13279 | ENSMUSG00000029701 | Rbm28 | 68272 [Gene Symbol: Rbm28] [Locus Tag: ] [Chromosome: 6] [Map Location: 6\|6 A3.3] | 3.299637197 | 7.45E-07 | 3.43E-06 |
| 16822 | ENSMUSG00000079037 | Prnp | 19122 [Gene Symbol: Prnp] [Locus Tag: ] [Chromosome: 2] [Map Location: 2 F2\|2 64.07 | 3.299159193 | 6.09E-10 | 4.53E-09 |
| 16833 | ENSMUSG00000028277 | Ube2j1 | 56228 [Gene Symbol: Ube2j1] [Locus Tag: ] [Chromosome: 4] [Map Location: 4\|4 A5] [D | 3.298969164 | 1.56E-11 | 1.48E-10 |
| 16508 | ENSMUSG00000046058 | Eid2 | 386655 [Gene Symbol: Eid2] [Locus Tag: ] [Chromosome: 7] [Map Location: 7\|7 A3] [De | 3.298570032 | 2.11E-10 | 1.68E-09 |
| 3810 | ENSMUSG00000030309 | Caprin2 | 232560 [Gene Symbol: Caprin2] [Locus Tag: ] [Chromosome: 6] [Map Location: 6\|6 G3] | 3.298439506 | 6.39E-07 | 2.97E-06 |
| 17814 | ENSMUSG00000045519 | Zfp560 | 434377 [Gene Symbol: Zfp560] [Locus Tag: ] [Chromosome: 9] [Map Location: 9\|9 A3] [ | 3.297536563 | 3.34E-08 | 1.92E-07 |
| 147 | ENSMUSG00000020280 | Pus10 | 74467 [Gene Symbol: Pus10] [Locus Tag: ] [Chromosome: 11] [Map Location: 11\|11 A3 | 3.297238544 | 9.81E-09 | 6.14E-08 |
| 15911 | ENSMUSG00000021087 | Rtn1 | 104001 [Gene Symbol: Rtn1] [Locus Tag: ] [Chromosome: 12] [Map Location: 12\|12 C3 | 3.297034882 | 3.32E-08 | 1.91E-07 |
| 15225 | ENSMUSG00000025776 | Crispld1 | 83691 [Gene Symbol: Crispld1] [Locus Tag: ] [Chromosome: 1] [Map Location: 1\|1 A4] [ | 3.296765284 | 0.00014 | 0.00043 |
| 11792 | ENSMUSG00000030417 | Pdcd5; Gm3837 | 56330 [Gene Symbol: Pdcd5] [Locus Tag: ] [Chromosome: 7] [Map Location: 7\|7 B2] [De | 3.295908532 | 1.70E-05 | 6.23E-05 |
| 14 | ENSMUSG00000044573 | LOC102642088; Acp1 | 102642088 [Gene Symbol: LOC102642088] [Locus Tag: ] [Chromosome: 10] [Map Loca | 3.295346869 | 3.44E-06 | 1.41E-05 |
| 8465 | ENSMUSG00000025825 | Iscu | 66383 [Gene Symbol: Iscu] [Locus Tag: ] [Chromosome: 5] [Map Location: 5\|5 F] [Descri | 3.294082739 | 3.24E-06 | 1.33E-05 |
| 15762 | ENSMUSG00000037013 | Ss18 | 268996 [Gene Symbol: Ss18] [Locus Tag: ] [Chromosome: 18] [Map Location: 18\|18 B1] | 3.291914858 | 2.88E-09 | 1.95E-08 |
| 12245 | ENSMUSG00000067860 | Zic3 | 22773 [Gene Symbol: Zic3] [Locus Tag: ] [Chromosome: X] [Map Location: X A6\|X 32.56 | 3.290531758 | 3.42E-07 | 1.66E-06 |
| 9816 | ENSMUSG00000030538 | Cib1 | 23991 [Gene Symbol: Cib1] [Locus Tag: ] [Chromosome: 7] [Map Location: 7\|7 D3] [Des | 3.289706313 | 2.60E-05 | 9.17E-05 |
| 11182 | ENSMUSG00000074925 | Ptar1 | 72351 [Gene Symbol: Ptar1] [Locus Tag: ] [Chromosome: 19] [Map Location: 19\|19 B] [ | 3.287932084 | 1.99E-09 | 1.37E-08 |
| 3397 | ENSMUSG00000029787 | Avl9 | 78937 [Gene Symbol: Avl9] [Locus Tag: ] [Chromosome: 6] [Map Location: 6\|6 C1] [Desc | 3.287026471 | 1.09E-09 | 7.81E-09 |
| 11325 | ENSMUSG00000045690 | Wdr89 | 72338 [Gene Symbol: Wdr89] [Locus Tag: ] [Chromosome: 12] [Map Location: 12\|12 C3 | 3.286803228 | 8.48E-06 | 3.26E-05 |
| 312 | ENSMUSG00000027831 | Veph1 | 72789 [Gene Symbol: Veph1] [Locus Tag: ] [Chromosome: 3] [Map Location: 3\|3 E1] [De | 3.285588563 | 0.000625 | 0.001692 |
| 873 | ENSMUSG00000036437 | Npy1r | 18166 [Gene Symbol: Npy1r] [Locus Tag: ] [Chromosome: 8] [Map Location: 8 33.15 cM | 3.284699908 | 6.75E-08 | 3.68E-07 |
| 4961 | ENSMUSG00000091183 | Gm5141 | 380850 [Gene Symbol: Gm5141] [Locus Tag: ] [Chromosome: 13] [Map Location: 13 B3 | 3.283291552 | 0.000155 | 0.000473 |
| 12665 | ENSMUSG00000036198 | Arhgap36 | 75404 [Gene Symbol: Arhgap36] [Locus Tag: ] [Chromosome: X] [Map Location: X\|X A5] | 3.283235973 | 0.000345 | 0.000982 |
| 4747 | ENSMUSG00000027615 | Hps3 | 12807 [Gene Symbol: Hps3] [Locus Tag: ] [Chromosome: 3] [Map Location: 3 A2\|3 6.12 | 3.281773567 | 1.79E-08 | 1.07E-07 |
| 1327 | ENSMUSG00000047187 | Rab2a | 59021 [Gene Symbol: Rab2a] [Locus Tag: ] [Chromosome: 4] [Map Location: 4\|4 A1] [De | 3.278798326 | 6.27E-12 | 6.36E-11 |
| 9837 | ENSMUSG00000025933 | Tmem14a | 75712 [Gene Symbol: Tmem14a] [Locus Tag: ] [Chromosome: 1] [Map Location: 1\|1 A4] | 3.278240357 | 1.27E-05 | 4.74E-05 |
| 6111 | ENSMUSG00000029832 | Nfe2l3 | 18025 [Gene Symbol: Nfe2l3] [Locus Tag: ] [Chromosome: 6] [Map Location: 6 B3\|6 24. | 3.275620787 | 1.02E-08 | 6.39E-08 |
| 18 | ENSMUSG00000025240 | Sacm1l | 83493 [Gene Symbol: Sacm1l] [Locus Tag: ] [Chromosome: 9] [Map Location: 9 74.08 cM | 3.272968164 | 1.29E-08 | 7.95E-08 |
| 14411 | ENSMUSG00000001305 | Rrp15 | 67223 [Gene Symbol: Rrp15] [Locus Tag: ] [Chromosome: 1] [Map Location: 1\|1 H5] [De | 3.271590035 | 3.76E-07 | 1.82E-06 |
| 11631 | ENSMUSG00000035601 | Trmt10b | 69934 [Gene Symbol: Trmt10b] [Locus Tag: ] [Chromosome: 4] [Map Location: 4\|4 B2] [ | 3.268351716 | 5.79E-06 | 2.29E-05 |
| 4128 | ENSMUSG00000026781 | Acbd5 | 74159 [Gene Symbol: Acbd5] [Locus Tag: ] [Chromosome: 2] [Map Location: 2\|2 A3] [De | 3.268066062 | 8.17E-11 | 6.95E-10 |
| 1482 | ENSMUSG00000042473 | Tbc1d8b | 245638 [Gene Symbol: Tbc1d8b] [Locus Tag: ] [Chromosome: X] [Map Location: X\|X F1] | 3.267178155 | 8.34E-05 | 0.000266 |
| 7141 | ENSMUSG00000008822 | Acyp1 | 66204 [Gene Symbol: Acyp1] [Locus Tag: ] [Chromosome: 12] [Map Location: 12\|12 D2 | 3.265427509 | 7.96E-05 | 0.000255 |
| 12136 | ENSMUSG00000097328 | Tnfsf12 | 21944 [Gene Symbol: Tnfsf12] [Locus Tag: ] [Chromosome: 11] [Map Location: 11\|11 B4 | 3.264623703 | 4.09E-05 | 0.000139 |
| 9281 | ENSMUSG00000014313 | Cox6c | 12864 [Gene Symbol: Cox6c] [Locus Tag: ] [Chromosome: 15] [Map Location: 15\|15 B3 | 3.264083963 | 1.45E-06 | 6.40E-06 |
| 14581 | ENSMUSG00000024691 | Fam111a | 107373 [Gene Symbol: Fam111a] [Locus Tag: ] [Chromosome: 19] [Map Location: 19\|1 | 3.26304932 | 4.56E-05 | 0.000153 |
| 17091 | ENSMUSG00000058799 | Nap1l1 | 53605 [Gene Symbol: Nap1l1] [Locus Tag: ] [Chromosome: 10] [Map Location: 10 D1\|1 | 3.260354673 | 4.19E-11 | 3.72E-10 |
| 9696 | ENSMUSG00000038039 | Gcc2 | 70297 [Gene Symbol: Gcc2] [Locus Tag: ] [Chromosome: 10] [Map Location: 10\|10 B4] | 3.259624618 | 1.44E-07 | 7.45E-07 |
| 1115 | ENSMUSG00000031337 | Mtm1 | 17772 [Gene Symbol: Mtm1] [Locus Tag: ] [Chromosome: X] [Map Location: X A7.2\|X 3 | 3.255008055 | 0.000176 | 0.000528 |
| 16985 | ENSMUSG00000028289 | Epha7 | 13841 [Gene Symbol: Epha7] [Locus Tag: ] [Chromosome: 4] [Map Location: 4 A4\|4 12.4 | 3.253736337 | 6.22E-09 | 4.02E-08 |
| 17188 | ENSMUSG00000036403 | Cep135 | 381644 [Gene Symbol: Cep135] [Locus Tag: ] [Chromosome: 5] [Map Location: 5\|5 C3.3 | 3.252486016 | 9.98E-07 | 4.49E-06 |
| 8058 | ENSMUSG00000035437 | Rabgap1 | 227800 [Gene Symbol: Rabgap1] [Locus Tag: ] [Chromosome: 2] [Map Location: 2\|2 B] [ | 3.252234496 | 1.02E-11 | 9.97E-11 |
| 14613 | ENSMUSG00000020571 | Pdia6 | 71853 [Gene Symbol: Pdia6] [Locus Tag: ] [Chromosome: 12] [Map Location: 12\|12 A1. | 3.250702791 | 4.53E-07 | 2.16E-06 |
| 248 | ENSMUSG00000071665 | Foxr2 | 436240 [Gene Symbol: Foxr2] [Locus Tag: ] [Chromosome: X] [Map Location: X\|X F3] [D | 3.250328458 | 0.001986 | 0.004841 |
| 16534 | ENSMUSG00000089862 | Umad1 | 100036521 [Gene Symbol: Umad1] [Locus Tag: ] [Chromosome: 6] [Map Location: 6\|6 A | 3.249743312 | 1.43E-11 | 1.37E-10 |
| 17382 | ENSMUSG00000021692 | Dimt1 | 66254 [Gene Symbol: Dimt1] [Locus Tag: ] [Chromosome: 13] [Map Location: 13\|13 D1 | 3.249374443 | 8.95E-08 | 4.79E-07 |
| 14552 | ENSMUSG00000056267 | Cep70 | 68121 [Gene Symbol: Cep70] [Locus Tag: ] [Chromosome: 9] [Map Location: 9\|9 F1] [De | 3.246768479 | 3.62E-08 | 2.07E-07 |
| 8616 | ENSMUSG00000069539 | Scyl2 | 213326 [Gene Symbol: Scyl2] [Locus Tag: ] [Chromosome: 10] [Map Location: 10 C2\|10 | 3.246018395 | 6.17E-11 | 5.36E-10 |
| 15604 | ENSMUSG00000031644 | Nek1 | 18004 [Gene Symbol: Nek1] [Locus Tag: ] [Chromosome: 8] [Map Location: 8 B3.1\|8 30 | 3.24219515 | 3.56E-06 | 1.45E-05 |
| 11567 | ENSMUSG00000062376 | Borcs7 | 66439 [Gene Symbol: Borcs7] [Locus Tag: ] [Chromosome: 19] [Map Location: 19\|19 D1 | 3.241386849 | 8.53E-06 | 3.28E-05 |
| 10560 | ENSMUSG00000055240 | Zfp101 | 22643 [Gene Symbol: Zfp101] [Locus Tag: ] [Chromosome: 17] [Map Location: 17\|17 B1 | 3.237994203 | 4.11E-08 | 2.32E-07 |
| 14308 | ENSMUSG00000034109 | Golim4 | 73124 [Gene Symbol: Golim4] [Locus Tag: ] [Chromosome: 3] [Map Location: 3\|3 E3] [D | 3.237522022 | 1.53E-08 | 9.29E-08 |
| 6555 | ENSMUSG00000033629 | Hacd3 | 57874 [Gene Symbol: Hacd3] [Locus Tag: ] [Chromosome: 9] [Map Location: 9\|9 D] [Des | 3.236617008 | 1.02E-10 | 8.55E-10 |
| 17811 | ENSMUSG00000029173 | Sepsecs | 211006 [Gene Symbol: Sepsecs] [Locus Tag: ] [Chromosome: 5] [Map Location: 5 C1\|5 2 | 3.236575436 | 2.13E-08 | 1.27E-07 |
| 13053 | ENSMUSG00000028410 | Dnaja1 | 15502 [Gene Symbol: Dnaja1] [Locus Tag: ] [Chromosome: 4] [Map Location: 4\|4 A5] [D | 3.235505482 | 7.60E-10 | 5.58E-09 |
| 1939 | ENSMUSG00000043943 | Naalad2 | 72560 [Gene Symbol: Naalad2] [Locus Tag: ] [Chromosome: 9] [Map Location: 9 7.02 cM | 3.234271228 | 0.000847 | 0.002236 |
| 10358 | ENSMUSG00000029390 | Tmed2; Gm21540 | 56334 [Gene Symbol: Tmed2] [Locus Tag: ] [Chromosome: 5] [Map Location: 5\|5 F] [Des | 3.233547403 | 5.29E-05 | 0.000175 |
| 5858 | ENSMUSG00000041353 | Tmem29 | 382245 [Gene Symbol: Tmem29] [Locus Tag: ] [Chromosome: X] [Map Location: X\|X F3] | 3.233502286 | 6.43E-10 | 4.76E-09 |
| 15833 | ENSMUSG00000000876 | Pxmp4 | 59038 [Gene Symbol: Pxmp4] [Locus Tag: ] [Chromosome: 2] [Map Location: 2\|2 H2] [D | 3.232927881 | 1.34E-06 | 5.94E-06 |
| 8835 | ENSMUSG00000032262 | Elovl4 | 83603 [Gene Symbol: Elovl4] [Locus Tag: ] [Chromosome: 9] [Map Location: 9\|9 E2] [De | 3.232252655 | 2.30E-10 | 1.82E-09 |
| 14440 | ENSMUSG00000073468 | Sft2d1 | 106489 [Gene Symbol: Sft2d1] [Locus Tag: ] [Chromosome: 17] [Map Location: 17\|17 A | 3.230555794 | 4.22E-09 | 2.79E-08 |
| 15427 | ENSMUSG00000021877 | Arf4 | 11843 [Gene Symbol: Arf4] [Locus Tag: ] [Chromosome: 14] [Map Location: 14 A3\|14 1 | 3.230292975 | 6.97E-11 | 6.00E-10 |
| 11621 | ENSMUSG00000027122 | Arl14ep | 212772 [Gene Symbol: Arl14ep] [Locus Tag: ] [Chromosome: 2] [Map Location: 2\|2 E3] | 3.225322746 | 6.58E-09 | 4.24E-08 |
| 497 | ENSMUSG00000029782 | Tmem209 | 72649 [Gene Symbol: Tmem209] [Locus Tag: ] [Chromosome: 6] [Map Location: 6\|6 A3 | 3.222382397 | 3.97E-07 | 1.91E-06 |
| 12799 | ENSMUSG00000019907 | Ppp1r12a | 17931 [Gene Symbol: Ppp1r12a] [Locus Tag: ] [Chromosome: 10] [Map Location: 10 56 | 3.219561776 | 2.26E-08 | 1.34E-07 |
| 8553 | ENSMUSG00000042460 | C1galt1 | 94192 [Gene Symbol: C1galt1] [Locus Tag: ] [Chromosome: 6] [Map Location: 6\|6 A1] [D | 3.218790631 | 5.79E-08 | 3.18E-07 |
| 7915 | ENSMUSG00000021754 | Map3k1 | 26401 [Gene Symbol: Map3k1] [Locus Tag: ] [Chromosome: 13] [Map Location: 13 D2.2 | 3.218717522 | 6.83E-09 | 4.38E-08 |
| 3843 | ENSMUSG00000031609 | Sap30 | 60406 [Gene Symbol: Sap30] [Locus Tag: ] [Chromosome: 8] [Map Location: 8 B2\|8 29.8 | 3.218305068 | 2.34E-05 | 8.34E-05 |
| 11519 | ENSMUSG00000016458 | Wt1 | 22431 [Gene Symbol: Wt1] [Locus Tag: ] [Chromosome: 2] [Map Location: 2 55.06 cM\|2 | 3.216477873 | 0.025373 | 0.047703 |
| 2196 | ENSMUSG00000051037 | Zfp455 | 218311 [Gene Symbol: Zfp455] [Locus Tag: ] [Chromosome: 13] [Map Location: 13 B3\|1 | 3.215211444 | 5.56E-07 | 2.62E-06 |
| 8715 | ENSMUSG00000069045 | Ddx3y | 26900 [Gene Symbol: Ddx3y] [Locus Tag: ] [Chromosome: Y] [Map Location: Y\|Y A1] [De | 3.214850879 | 0.020369 | 0.039153 |

| | | | | | | |
|---|---|---|---|---|---|---|
| 6585 | ENSMUSG00000026586 | Prrx1 | 18933 [Gene Symbol: Prrx1] [Locus Tag: ] [Chromosome: 1] [Map Location: 1 H2.1\|1 70 | 3.214740794 | 1.03E-07 | 5.49E-07 |
| 7802 | ENSMUSG00000005268 | Prlr | 19116 [Gene Symbol: Prlr] [Locus Tag: ] [Chromosome: 15] [Map Location: 15 A1\|15 5. | 3.214494379 | 2.31E-05 | 8.24E-05 |
| 11075 | ENSMUSG00000043929 | Klhl15 | 236904 [Gene Symbol: Klhl15] [Locus Tag: ] [Chromosome: X] [Map Location: X\|X C3] [ | 3.214441485 | 5.54E-10 | 4.14E-09 |
| 16664 | ENSMUSG00000079065 | BC005561 | 100042165 [Gene Symbol: BC005561] [Locus Tag: ] [Chromosome: 5] [Map Location: 5 | 3.214123838 | 1.01E-08 | 6.29E-08 |
| 6007 | ENSMUSG00000025915 | Sgk3 | 170755 [Gene Symbol: Sgk3] [Locus Tag: ] [Chromosome: 1] [Map Location: 1 A2\|1 2.08 | 3.213476758 | 1.59E-05 | 5.84E-05 |
| 2735 | ENSMUSG00000027550 | Lrrcc1 | 71710 [Gene Symbol: Lrrcc1] [Locus Tag: ] [Chromosome: 3] [Map Location: 3\|3 A2] [De | 3.212261045 | 5.14E-08 | 2.86E-07 |
| 11756 | ENSMUSG00000043067 | Dpy19l1 | 244745 [Gene Symbol: Dpy19l1] [Locus Tag: ] [Chromosome: 9] [Map Location: 9\|9 A4] | 3.2122297 | 4.75E-07 | 2.26E-06 |
| 10274 | ENSMUSG00000058076 | Sdhc | 66052 [Gene Symbol: Sdhc] [Locus Tag: ] [Chromosome: 1] [Map Location: 1\|1 H3] [Des | 3.211578911 | 5.41E-08 | 3.00E-07 |
| 44 | ENSMUSG00000051674 | Dcun1d4 | 100737 [Gene Symbol: Dcun1d4] [Locus Tag: ] [Chromosome: 5] [Map Location: 5\|5 C3 | 3.211494632 | 5.73E-11 | 5.00E-10 |
| 3784 | ENSMUSG00000024642 | Tle4 | 21888 [Gene Symbol: Tle4] [Locus Tag: ] [Chromosome: 19] [Map Location: 19 A\|19 9.1 | 3.210489325 | 2.82E-11 | 2.56E-10 |
| 1022 | ENSMUSG00000028576 | Ift74 | 67694 [Gene Symbol: Ift74] [Locus Tag: ] [Chromosome: 4] [Map Location: 4\|4 C5] [Des | 3.210050601 | 2.58E-08 | 1.51E-07 |
| 1762 | ENSMUSG00000031616 | Ednra | 13617 [Gene Symbol: Ednra] [Locus Tag: ] [Chromosome: 8] [Map Location: 8\|8 C2] [De | 3.20962544 | 1.90E-05 | 6.88E-05 |
| 12104 | ENSMUSG00000044950 | Pwwp2a | 70802 [Gene Symbol: Pwwp2a] [Locus Tag: ] [Chromosome: 11] [Map Location: 11\|11 | 3.20895979 | 1.85E-10 | 1.49E-09 |
| 7745 | ENSMUSG00000026634 | Angel2 | 52477 [Gene Symbol: Angel2] [Locus Tag: ] [Chromosome: 1] [Map Location: 1 H6\|1 96 | 3.208372849 | 1.38E-10 | 1.13E-09 |
| 1457 | ENSMUSG00000027959 | Sass6 | 72776 [Gene Symbol: Sass6] [Locus Tag: ] [Chromosome: 3] [Map Location: 3\|3 G2] [De | 3.208162222 | 6.46E-07 | 3.00E-06 |
| 4938 | ENSMUSG00000055302 | Mrfap1 | 67568 [Gene Symbol: Mrfap1] [Locus Tag: ] [Chromosome: 5] [Map Location: 5\|5 B3] [D | 3.206691618 | 2.92E-10 | 2.28E-09 |
| 9705 | ENSMUSG00000049288 | Lix1l | 280411 [Gene Symbol: Lix1l] [Locus Tag: ] [Chromosome: 3] [Map Location: 3\|3 F2.1] [ | 3.206647555 | 1.33E-10 | 1.09E-09 |
| 7422 | ENSMUSG00000058254 | Tspan7 | 21912 [Gene Symbol: Tspan7] [Locus Tag: ] [Chromosome: X] [Map Location: X\|X A1.3- | 3.202131216 | 1.96E-10 | 1.57E-09 |
| 11265 | ENSMUSG00000020044 | Timp3 | 21859 [Gene Symbol: Timp3] [Locus Tag: ] [Chromosome: 10] [Map Location: 10 42.83 | 3.200565315 | 1.09E-10 | 9.08E-10 |
| 2068 | ENSMUSG00000035984 | Nme5 | 75533 [Gene Symbol: Nme5] [Locus Tag: ] [Chromosome: 18] [Map Location: 18\|18 B3] | 3.199709029 | 0.001603 | 0.003991 |
| 3209 | ENSMUSG00000044067 | Gpr22 | 73010 [Gene Symbol: Gpr22] [Locus Tag: ] [Chromosome: 12] [Map Location: 12\|12 A3 | 3.199560255 | 1.05E-09 | 7.52E-09 |
| 7360 | ENSMUSG00000036580 | Spg20 | 229285 [Gene Symbol: Spg20] [Locus Tag: ] [Chromosome: 3] [Map Location: 3\|3 C] [De | 3.199442326 | 1.63E-09 | 1.14E-08 |
| 3958 | ENSMUSG00000049001 | Ndnf | 68169 [Gene Symbol: Ndnf] [Locus Tag: ] [Chromosome: 6] [Map Location: 6\|6 C1] [Des | 3.198966529 | 9.48E-10 | 6.86E-09 |
| 6452 | ENSMUSG00000021282 | LOC102641436; Eif5 | 102641436 [Gene Symbol: LOC102641436] [Locus Tag: ] [Chromosome: 10] [Map Loca | 3.197375097 | 2.65E-11 | 2.43E-10 |
| 2083 | ENSMUSG00000035199 | Arl6ip5 | 65106 [Gene Symbol: Arl6ip5] [Locus Tag: ] [Chromosome: 6] [Map Location: 6\|6 D3] [ | 3.196753966 | 2.55E-09 | 1.73E-08 |
| 6166 | ENSMUSG00000038538 | Ubn2 | 320538 [Gene Symbol: Ubn2] [Locus Tag: ] [Chromosome: 6] [Map Location: 6\|6 B1] [D | 3.195763595 | 4.34E-09 | 2.86E-08 |
| 14156 | ENSMUSG00000023150 | Ivns1abp | 117198 [Gene Symbol: Ivns1abp] [Locus Tag: ] [Chromosome: 1] [Map Location: 1\|1 G2 | 3.19542114 | 2.25E-11 | 2.08E-10 |
| 3702 | ENSMUSG00000020526 | Znhit3 | 448850 [Gene Symbol: Znhit3] [Locus Tag: ] [Chromosome: 11] [Map Location: 11\|11 C | 3.191957005 | 1.56E-05 | 5.75E-05 |
| 4019 | ENSMUSG00000000787 | Ddx3x | 13205 [Gene Symbol: Ddx3x] [Locus Tag: ] [Chromosome: X] [Map Location: X A1.1\|X 8 | 3.191931823 | 2.23E-11 | 2.07E-10 |
| 12907 | ENSMUSG00000027751 | Supt20 | 56790 [Gene Symbol: Supt20] [Locus Tag: ] [Chromosome: 3] [Map Location: 3\|3 D] [De | 3.191551501 | 2.85E-08 | 1.65E-07 |
| 16337 | ENSMUSG00000098754 | Prn | 111368 [Gene Symbol: Prn] [Locus Tag: ] [Chromosome: 2] [Map Location: 2 F2\|2] [Des | 3.19031582 | 4.94E-10 | 3.72E-09 |
| 17636 | ENSMUSG00000035150 | Eif2s3x | 26905 [Gene Symbol: Eif2s3x] [Locus Tag: ] [Chromosome: X] [Map Location: X 41.52 cM | 3.188957385 | 0.000144 | 0.000439 |
| 13558 | ENSMUSG00000029178 | Klf3 | 16599 [Gene Symbol: Klf3] [Locus Tag: ] [Chromosome: 5] [Map Location: 5\|5 C3.1] [De | 3.188833742 | 8.36E-11 | 7.09E-10 |
| 10989 | ENSMUSG00000020059 | Sycp3 | 20962 [Gene Symbol: Sycp3] [Locus Tag: ] [Chromosome: 10] [Map Location: 10\|10 C] [ | 3.18661394 | 0.001582 | 0.003945 |
| 4637 | ENSMUSG00000033361 | Prrg3 | 208748 [Gene Symbol: Prrg3] [Locus Tag: ] [Chromosome: X] [Map Location: X\|X A7.3] | 3.185316856 | 6.33E-10 | 4.70E-09 |
| 12001 | ENSMUSG00000037617 | Spag1 | 26942 [Gene Symbol: Spag1] [Locus Tag: ] [Chromosome: 15] [Map Location: 15\|15 C] [ | 3.182788784 | 8.40E-06 | 3.24E-05 |
| 9716 | ENSMUSG00000026348 | Acmsd | 266645 [Gene Symbol: Acmsd] [Locus Tag: ] [Chromosome: 1] [Map Location: 1\|1 E3] [D | 3.178954838 | 0.003742 | 0.008576 |
| 8467 | ENSMUSG00000021972 | Hmbox1 | 219150 [Gene Symbol: Hmbox1] [Locus Tag: ] [Chromosome: 14] [Map Location: 14\|14 | 3.178715485 | 9.32E-11 | 7.83E-10 |
| 8207 | ENSMUSG00000026058 | Khdrbs2 | 170771 [Gene Symbol: Khdrbs2] [Locus Tag: ] [Chromosome: 1] [Map Location: 1\|1 B] [ | 3.177878283 | 9.11E-11 | 7.67E-10 |
| 17147 | ENSMUSG00000015217 | Hmgb3 | 15354 [Gene Symbol: Hmgb3] [Locus Tag: ] [Chromosome: X] [Map Location: X\|X A1] [D | 3.175476945 | 7.18E-09 | 4.59E-08 |
| 8650 | ENSMUSG00000003282 | Plag1 | 56711 [Gene Symbol: Plag1] [Locus Tag: ] [Chromosome: 4] [Map Location: 4\|4 A1] [De | 3.174935609 | 2.13E-05 | 7.66E-05 |
| 14362 | ENSMUSG00000001280 | Sp1 | 20683 [Gene Symbol: Sp1] [Locus Tag: ] [Chromosome: 15] [Map Location: 15\|15 F3] [D | 3.174043476 | 1.64E-10 | 1.33E-09 |
| 16590 | ENSMUSG00000024576 | Csnk1a1 | 93687 [Gene Symbol: Csnk1a1] [Locus Tag: ] [Chromosome: 18] [Map Location: 18\|18 | 3.17245097 | 2.89E-11 | 2.63E-10 |
| 4459 | ENSMUSG00000024287 | Thoc1 | 225160 [Gene Symbol: Thoc1] [Locus Tag: ] [Chromosome: 18] [Map Location: 18\|18 A | 3.170875176 | 5.72E-08 | 3.15E-07 |
| 9744 | ENSMUSG00000030322 | Mbd4 | 17193 [Gene Symbol: Mbd4] [Locus Tag: ] [Chromosome: 6] [Map Location: 6 E3\|6 53.7 | 3.169830561 | 4.84E-06 | 1.93E-05 |
| 6516 | ENSMUSG00000040410 | Fbxl4 | 269514 [Gene Symbol: Fbxl4] [Locus Tag: ] [Chromosome: 4] [Map Location: 4\|4 A3] [D | 3.168882271 | 1.77E-08 | 1.06E-07 |
| 3677 | ENSMUSG00000027306 | Nusap1 | 108907 [Gene Symbol: Nusap1] [Locus Tag: ] [Chromosome: 2] [Map Location: 2\|2 E5] [ | 3.168328679 | 3.67E-05 | 0.000126 |
| 14094 | ENSMUSG00000073481 | March2 | 67247 [Gene Symbol: Marc2] [Locus Tag: ] [Chromosome: 1] [Map Location: 1\|1 H4] [D | 3.164339234 | 4.39E-07 | 2.10E-06 |
| 15657 | ENSMUSG00000089774 | Slc5a3 | 53881 [Gene Symbol: Slc5a3] [Locus Tag: ] [Chromosome: 16] [Map Location: 16\|16 C4 | 3.161706016 | 2.43E-08 | 1.43E-07 |
| 7498 | ENSMUSG00000032135 | Mcam | 84004 [Gene Symbol: Mcam] [Locus Tag: ] [Chromosome: 9] [Map Location: 9\|9 A5.1] [ | 3.161094267 | 0.000156 | 0.000475 |
| 234 | ENSMUSG00000022018 | Rgcc | 66214 [Gene Symbol: Rgcc] [Locus Tag: ] [Chromosome: 14] [Map Location: 14\|14 D3] | 3.160571039 | 8.68E-05 | 0.000275 |
| 15361 | ENSMUSG00000017057 | Il13ra1 | 16164 [Gene Symbol: Il13ra1] [Locus Tag: ] [Chromosome: X] [Map Location: X A3.3\|X 2 | 3.157450025 | 1.86E-06 | 8.02E-06 |
| 15571 | ENSMUSG00000020152 | Actr2 | 66713 [Gene Symbol: Actr2] [Locus Tag: ] [Chromosome: 11] [Map Location: 11 A3.1\|1 | 3.155444744 | 9.74E-11 | 8.17E-10 |
| 1472 | ENSMUSG00000005506 | Celf1 | 13046 [Gene Symbol: Celf1] [Locus Tag: ] [Chromosome: 2] [Map Location: 2 E1\|2 50.4 | 3.155101463 | 6.72E-11 | 5.81E-10 |
| 11166 | ENSMUSG00000048058 | Ldlrad3 | 241576 [Gene Symbol: Ldlrad3] [Locus Tag: ] [Chromosome: 2] [Map Location: 2\|2 E2] | 3.152892971 | 6.89E-10 | 5.08E-09 |
| 1614 | ENSMUSG00000036282 | Naa30 | 70646 [Gene Symbol: Naa30] [Locus Tag: ] [Chromosome: 14] [Map Location: 14\|14 C1 | 3.149410753 | 8.79E-11 | 7.42E-10 |
| 7633 | ENSMUSG00000037262 | Kin | 16588 [Gene Symbol: Kin] [Locus Tag: ] [Chromosome: 2] [Map Location: 2\|2 A1] [Descr | 3.144807497 | 3.59E-06 | 1.46E-05 |
| 16388 | ENSMUSG00000035953 | Tmem55b | 219024 [Gene Symbol: Tmem55b] [Locus Tag: ] [Chromosome: 14] [Map Location: 14\| | 3.144097914 | 3.13E-10 | 2.43E-09 |
| 8683 | ENSMUSG00000029427 | Zcchc8 | 70650 [Gene Symbol: Zcchc8] [Locus Tag: ] [Chromosome: 5] [Map Location: 5\|5 F] [De | 3.143229109 | 1.66E-08 | 1.00E-07 |
| 708 | ENSMUSG00000034560 | A230046K03Rik | 319277 [Gene Symbol: A230046K03Rik] [Locus Tag: ] [Chromosome: 10] [Map Location | 3.143146222 | 8.79E-10 | 6.39E-09 |
| 4498 | ENSMUSG00000007653 | Gabrb2 | 14401 [Gene Symbol: Gabrb2] [Locus Tag: ] [Chromosome: 11] [Map Location: 11 A5\|1 | 3.142669649 | 5.51E-11 | 4.81E-10 |
| 1092 | ENSMUSG00000040843 | Tiprl | 226591 [Gene Symbol: Tiprl] [Locus Tag: ] [Chromosome: 1] [Map Location: 1\|1 H2.3] [ | 3.140761638 | 3.51E-10 | 2.69E-09 |
| 14125 | ENSMUSG00000058402 | Zfp420 | 233058 [Gene Symbol: Zfp420] [Locus Tag: ] [Chromosome: 7] [Map Location: 7\|7 B1] [ | 3.136880663 | 4.32E-05 | 0.000146 |
| 13983 | ENSMUSG00000058355 | Abce1 | 24015 [Gene Symbol: Abce1] [Locus Tag: ] [Chromosome: 8] [Map Location: 8\|8 C] [Des | 3.136699314 | 2.18E-10 | 1.73E-09 |
| 67 | ENSMUSG00000067424 | Zfp563 | 240068 [Gene Symbol: Zfp563] [Locus Tag: ] [Chromosome: 17] [Map Location: 17\|17 | 3.135991007 | 1.30E-08 | 7.98E-08 |
| 1236 | ENSMUSG00000020123 | Avpr1a | 54140 [Gene Symbol: Avpr1a] [Locus Tag: ] [Chromosome: 10] [Map Location: 10\|10 D | 3.133583795 | 0.000661 | 0.001779 |
| 14172 | ENSMUSG00000062797 | l7Rn6 | 67669 [Gene Symbol: l7Rn6] [Locus Tag: ] [Chromosome: 7] [Map Location: 7 50.4 cM\| | 3.133197885 | 4.19E-07 | 2.01E-06 |
| 14162 | ENSMUSG00000032387 | Rbpms2 | 71973 [Gene Symbol: Rbpms2] [Locus Tag: ] [Chromosome: 9] [Map Location: 9\|9 C] [D | 3.132689363 | 0.003493 | 0.008056 |
| 17488 | ENSMUSG00000011958 | Bnip2 | 12175 [Gene Symbol: Bnip2] [Locus Tag: ] [Chromosome: 9] [Map Location: 9\|9 D] [Des | 3.132207302 | 5.30E-09 | 3.46E-08 |
| 7069 | ENSMUSG00000041594 | Tmtc4 | 70551 [Gene Symbol: Tmtc4] [Locus Tag: ] [Chromosome: 14] [Map Location: 14\|14 E5 | 3.131798445 | 1.31E-09 | 9.24E-09 |
| 10414 | ENSMUSG00000024378 | Stard4 | 170459 [Gene Symbol: Stard4] [Locus Tag: ] [Chromosome: 18] [Map Location: 18 B1\|1 | 3.125533749 | 1.05E-09 | 7.51E-09 |
| 10178 | ENSMUSG00000000766 | Oprm1 | 18390 [Gene Symbol: Oprm1] [Locus Tag: ] [Chromosome: 10] [Map Location: 10 1.85 c | 3.123639538 | 1.17E-10 | 9.73E-10 |
| 11340 | ENSMUSG00000073733 | Rsg1 | 76166 [Gene Symbol: Rsg1] [Locus Tag: ] [Chromosome: 4] [Map Location: 4\|4 E1] [Desc | 3.122905498 | 2.57E-05 | 9.09E-05 |
| 16177 | ENSMUSG00000074578 | Zfos1 | 68949 [Gene Symbol: Zfos1] [Locus Tag: ] [Chromosome: 2] [Map Location: 2\|2 H3] [De | 3.122365971 | 2.38E-06 | 1.00E-05 |
| 8174 | ENSMUSG00000071072 | LOC102641464; Ptge | 102641464 [Gene Symbol: LOC102641464] [Locus Tag: ] [Chromosome: 6] [Map Locati | 3.121784554 | 9.20E-09 | 5.80E-08 |
| 8328 | ENSMUSG00000037905 | Bri3bp | 76809 [Gene Symbol: Bri3bp] [Locus Tag: ] [Chromosome: 5] [Map Location: 5\|5 F] [De | 3.121043283 | 4.17E-11 | 3.71E-10 |
| 11125 | ENSMUSG00000031520 | Vegfc | 22341 [Gene Symbol: Vegfc] [Locus Tag: ] [Chromosome: 8] [Map Location: 8\|8 B3] [De | 3.120421131 | 1.95E-06 | 8.38E-06 |
| 12680 | ENSMUSG00000028780 | Sema3c | 20348 [Gene Symbol: Sema3c] [Locus Tag: ] [Chromosome: 5] [Map Location: 5\|5 A3] [ | 3.118596556 | 4.81E-10 | 3.63E-09 |
| 6307 | ENSMUSG00000056536 | Pign | 27392 [Gene Symbol: Pign] [Locus Tag: ] [Chromosome: 1] [Map Location: 1\|1 D] [Descr | 3.118495992 | 2.03E-09 | 1.40E-08 |
| 17661 | ENSMUSG00000050705 | 2310061I04Rik | 69662 [Gene Symbol: 2310061I04Rik] [Locus Tag: ] [Chromosome: 17] [Map Location: | 3.117572216 | 3.08E-09 | 2.08E-08 |
| 4502 | ENSMUSG00000020671 | Rab10 | 19325 [Gene Symbol: Rab10] [Locus Tag: ] [Chromosome: 12] [Map Location: 12 A1.1\| | 3.116825521 | 6.56E-11 | 5.67E-10 |
| 15789 | ENSMUSG00000020648 | Dus4l | 71916 [Gene Symbol: Dus4l] [Locus Tag: ] [Chromosome: 12] [Map Location: 12\|12 A3] | 3.115746901 | 3.47E-05 | 0.00012 |
| 7724 | ENSMUSG00000031467 | Agpat5 | 52123 [Gene Symbol: Agpat5] [Locus Tag: ] [Chromosome: 8] [Map Location: 8 10.3 cM | 3.11426994 | 2.57E-10 | 2.02E-09 |
| 17150 | ENSMUSG00000040565 | Btaf1 | 107182 [Gene Symbol: Btaf1] [Locus Tag: ] [Chromosome: 19] [Map Location: 19\|19 C2 | 3.112903528 | 1.02E-06 | 4.57E-06 |

| | | | | | | |
|---|---|---|---|---|---|---|
| 5781 | ENSMUSG00000042410 | Agps | 228061 [Gene Symbol: Agps] [Locus Tag: ] [Chromosome: 2] [Map Location: 2\|2 C3] [De | 3.112771341 | 3.36E-10 | 2.59E-09 |
| 11686 | ENSMUSG00000087598 | Zfp111 | 56707 [Gene Symbol: Zfp111] [Locus Tag: ] [Chromosome: 7] [Map Location: 7\|7 A3] [D | 3.111619338 | 4.57E-07 | 2.18E-06 |
| 8370 | ENSMUSG00000073077 | Gm8787; Gm7173 | 667736 [Gene Symbol: Gm8787] [Locus Tag: ] [Chromosome: X] [Map Location: X\|X B] | 3.109061877 | 2.26E-06 | 9.57E-06 |
| 1325 | ENSMUSG00000050931 | Sgms2 | 74442 [Gene Symbol: Sgms2] [Locus Tag: ] [Chromosome: 3] [Map Location: 3\|3 G3] [D | 3.106620235 | 6.00E-06 | 2.36E-05 |
| 9929 | ENSMUSG00000031950 | Gabarapl2 | 93739 [Gene Symbol: Gabarapl2] [Locus Tag: ] [Chromosome: 8] [Map Location: 8\|8 E1 | 3.106405769 | 2.05E-05 | 7.39E-05 |
| 2469 | ENSMUSG00000033009 | Ogfod1 | 270086 [Gene Symbol: Ogfod1] [Locus Tag: ] [Chromosome: 8] [Map Location: 8\|8 C5] [ | 3.105632746 | 1.42E-09 | 1.00E-08 |
| 1163 | ENSMUSG00000001018 | Snapin | 20615 [Gene Symbol: Snapin] [Locus Tag: ] [Chromosome: 3] [Map Location: 3\|3 F1] [D | 3.104724809 | 1.60E-08 | 9.64E-08 |
| 9887 | ENSMUSG00000028098 | Rnf115 | 67845 [Gene Symbol: Rnf115] [Locus Tag: ] [Chromosome: 3] [Map Location: 3\|3 F2.1] | 3.104608806 | 1.15E-09 | 8.17E-09 |
| 11562 | ENSMUSG00000054894 | Atp5s | 68055 [Gene Symbol: Atp5s] [Locus Tag: ] [Chromosome: 12] [Map Location: 12\|12 C3] | 3.103340128 | 0.000162 | 0.00049 |
| 8044 | ENSMUSG00000021696 | Elovl7 | 74559 [Gene Symbol: Elovl7] [Locus Tag: ] [Chromosome: 13] [Map Location: 13\|13 D2 | 3.102687832 | 0.000473 | 0.001312 |
| 9662 | ENSMUSG00000041245 | Wnk3 | 279561 [Gene Symbol: Wnk3] [Locus Tag: ] [Chromosome: X] [Map Location: X\|X F3] [D | 3.102626197 | 1.03E-10 | 8.61E-10 |
| 14065 | ENSMUSG00000028274 | Rngtt | 24018 [Gene Symbol: Rngtt] [Locus Tag: ] [Chromosome: 4] [Map Location: 4\|4 A5] [De | 3.102518502 | 6.44E-10 | 4.77E-09 |
| 15149 | ENSMUSG00000040446 | Rprd1a | 225283 [Gene Symbol: Rprd1a] [Locus Tag: ] [Chromosome: 18] [Map Location: 18\|18 | 3.101874509 | 4.18E-10 | 3.18E-09 |
| 4471 | ENSMUSG00000026189 | Pecr | 111175 [Gene Symbol: Pecr] [Locus Tag: ] [Chromosome: 1] [Map Location: 1 C3\|1 36. | 3.101708433 | 8.97E-05 | 0.000283 |
| 11985 | ENSMUSG00000060002 | Chpt1 | 212862 [Gene Symbol: Chpt1] [Locus Tag: ] [Chromosome: 10] [Map Location: 10\|10 C | 3.099743924 | 7.76E-05 | 0.000249 |
| 5540 | ENSMUSG00000078784 | 1810022K09Rik | 69126 [Gene Symbol: 1810022K09Rik] [Locus Tag: ] [Chromosome: 3] [Map Location: 3 | 3.098907771 | 0.000116 | 0.00036 |
| 1740 | ENSMUSG00000024231 | Cul2 | 71745 [Gene Symbol: Cul2] [Locus Tag: ] [Chromosome: 18] [Map Location: 18\|18 A1] [ | 3.098072255 | 1.14E-09 | 8.14E-09 |
| 11471 | ENSMUSG00000025817 | Nudt5 | 53893 [Gene Symbol: Nudt5] [Locus Tag: ] [Chromosome: 2] [Map Location: 2\|2 A1] [De | 3.098007464 | 1.15E-07 | 6.04E-07 |
| 15260 | ENSMUSG00000054414 | Slc30a7 | 66500 [Gene Symbol: Slc30a7] [Locus Tag: ] [Chromosome: 3] [Map Location: 3\|3 G2] [ | 3.098006218 | 2.02E-08 | 1.20E-07 |
| 11623 | ENSMUSG00000032567 | Aste1 | 66595 [Gene Symbol: Aste1] [Locus Tag: ] [Chromosome: 9] [Map Location: 9\|9 F1] [Des | 3.097049959 | 8.06E-08 | 4.33E-07 |
| 16038 | ENSMUSG00000035509 | Fbxl21 | 213311 [Gene Symbol: Fbxl21] [Locus Tag: ] [Chromosome: 13] [Map Location: 13\|13 E | 3.096701762 | 0.000126 | 0.000388 |
| 2337 | ENSMUSG00000036781 | Rps27l | 67941 [Gene Symbol: Rps27l] [Locus Tag: ] [Chromosome: 9] [Map Location: 9\|9 D] [De | 3.095941726 | 0.002266 | 0.005453 |
| 17357 | ENSMUSG00000027679 | Dnajc19; Dnajc19-ps | 67713 [Gene Symbol: Dnajc19] [Locus Tag: ] [Chromosome: 3] [Map Location: ] [Descrip | 3.094817287 | 2.26E-10 | 1.80E-09 |
| 17321 | ENSMUSG00000015289 | Lage3 | 66192 [Gene Symbol: Lage3] [Locus Tag: ] [Chromosome: X] [Map Location: X\|X A6] [De | 3.091872781 | 0.00038 | 0.001075 |
| 10063 | ENSMUSG00000050074 | Spink8 | 78709 [Gene Symbol: Spink8] [Locus Tag: ] [Chromosome: 9] [Map Location: 9\|9 F2] [D | 3.090917084 | 0.005889 | 0.01291 |
| 16804 | ENSMUSG00000032265 | Fam46a | 212943 [Gene Symbol: Fam46a] [Locus Tag: ] [Chromosome: 9] [Map Location: 9\|9 E3. | 3.088892858 | 1.62E-05 | 5.95E-05 |
| 2554 | ENSMUSG00000061524 | Zic2 | 22772 [Gene Symbol: Zic2] [Locus Tag: ] [Chromosome: 14] [Map Location: 14 E5\|14 6 | 3.083422901 | 2.00E-10 | 1.60E-09 |
| 924 | ENSMUSG00000021555 | Naa35 | 78689 [Gene Symbol: Naa35] [Locus Tag: ] [Chromosome: 13] [Map Location: 13\|13 B3 | 3.08175522 | 9.64E-10 | 6.96E-09 |
| 206 | ENSMUSG00000032026 | Rexo2 | 104444 [Gene Symbol: Rexo2] [Locus Tag: ] [Chromosome: 9] [Map Location: 9\|9 A5.3] | 3.081669081 | 2.34E-05 | 8.35E-05 |
| 16829 | ENSMUSG00000047843 | Bri3 | 55950 [Gene Symbol: Bri3] [Locus Tag: ] [Chromosome: 5] [Map Location: 5\|5 G2] [Des | 3.081212321 | 1.80E-06 | 7.79E-06 |
| 9653 | ENSMUSG00000027997 | Casp6 | 12368 [Gene Symbol: Casp6] [Locus Tag: ] [Chromosome: 3] [Map Location: 3\|3 H1] [De | 3.080765343 | 0.000168 | 0.000508 |
| 15451 | ENSMUSG00000031352 | Hccs | 15159 [Gene Symbol: Hccs] [Locus Tag: ] [Chromosome: X] [Map Location: X\|X F5] [Des | 3.079574637 | 4.30E-07 | 2.06E-06 |
| 6831 | ENSMUSG00000026111 | Unc50 | 67387 [Gene Symbol: Unc50] [Locus Tag: ] [Chromosome: 1] [Map Location: 1\|1 B] [Des | 3.07944898 | 2.77E-08 | 1.61E-07 |
| 9838 | ENSMUSG00000037958 | Nsrp1 | 237859 [Gene Symbol: Nsrp1] [Locus Tag: ] [Chromosome: 11] [Map Location: 11\|11 B | 3.079066509 | 4.43E-09 | 2.92E-08 |
| 14284 | ENSMUSG00000033740 | St18 | 240690 [Gene Symbol: St18] [Locus Tag: ] [Chromosome: 1] [Map Location: 1\|1 A1] [De | 3.07827125 | 3.22E-07 | 1.57E-06 |
| 8078 | ENSMUSG00000022336 | Eif3e | 16341 [Gene Symbol: Eif3e] [Locus Tag: ] [Chromosome: 15] [Map Location: 15 B3.2\|15 | 3.077870829 | 1.55E-08 | 9.38E-08 |
| 7337 | ENSMUSG00000025961 | 4933402D24Rik | 74426 [Gene Symbol: 4933402D24Rik] [Locus Tag: ] [Chromosome: 1] [Map Location: | 3.074667909 | 0.001594 | 0.003972 |
| 1225 | ENSMUSG00000030663 | 1110004F10Rik | 56372 [Gene Symbol: 1110004F10Rik] [Locus Tag: ] [Chromosome: 7] [Map Location: 7 | 3.07465405 | 1.69E-09 | 1.18E-08 |
| 1655 | ENSMUSG00000026492 | Tfb2m | 15278 [Gene Symbol: Tfb2m] [Locus Tag: ] [Chromosome: 1] [Map Location: 1 83.56 cM | 3.073830694 | 1.76E-06 | 7.64E-06 |
| 13498 | ENSMUSG00000063273 | Naa15 | 74838 [Gene Symbol: Naa15] [Locus Tag: ] [Chromosome: 3] [Map Location: 3\|3 D] [Des | 3.072910882 | 9.86E-11 | 8.26E-10 |
| 15682 | ENSMUSG00000062822 | 4833420G17Rik | 67392 [Gene Symbol: 4833420G17Rik] [Locus Tag: ] [Chromosome: 13] [Map Location: | 3.07133456 | 4.63E-08 | 2.60E-07 |
| 11987 | ENSMUSG00000070737 | Gm12942 | 100039968 [Gene Symbol: Gm12942] [Locus Tag: ] [Chromosome: 4] [Map Location: 4 | 3.071152783 | 2.88E-05 | 0.000101 |
| 14282 | ENSMUSG00000089857 | Zfp882 | 382019 [Gene Symbol: Zfp882] [Locus Tag: ] [Chromosome: 8] [Map Location: 8 B3.3\|8 | 3.069472791 | 1.25E-06 | 5.57E-06 |
| 5090 | ENSMUSG00000048170 | Mcmbp | 210711 [Gene Symbol: Mcmbp] [Locus Tag: ] [Chromosome: 7] [Map Location: 7\|7 F3] | 3.069175878 | 1.36E-09 | 9.64E-09 |
| 1051 | ENSMUSG00000049411 | Tmem241 | 338363 [Gene Symbol: Tmem241] [Locus Tag: ] [Chromosome: 18] [Map Location: 18\| | 3.067335138 | 1.54E-07 | 7.96E-07 |
| 11045 | ENSMUSG00000043065 | Spice1 | 212514 [Gene Symbol: Spice1] [Locus Tag: ] [Chromosome: 16] [Map Location: 16 B4\|1 | 3.066702441 | 4.67E-05 | 0.000156 |
| 948 | ENSMUSG00000096856 | Gm10778 | 100233208 [Gene Symbol: Gm10778] [Locus Tag: ] [Chromosome: 10] [Map Location: | 3.066529587 | 0.015575 | 0.030898 |
| 7369 | ENSMUSG00000008682 | Rpl10; Gm5621 | 110954 [Gene Symbol: Rpl10] [Locus Tag: ] [Chromosome: X] [Map Location: X A7.3\|X | 3.064443026 | 0.018831 | 0.03654 |
| 11910 | ENSMUSG00000036315 | Znrd1 | 66136 [Gene Symbol: Znrd1] [Locus Tag: ] [Chromosome: 17] [Map Location: 17\|17 B3 | 3.063247355 | 8.10E-05 | 0.000258 |
| 11442 | ENSMUSG00000042246 | Tmc7 | 209760 [Gene Symbol: Tmc7] [Locus Tag: ] [Chromosome: 7] [Map Location: 7\|7 F2] [D | 3.061769081 | 2.87E-07 | 1.41E-06 |
| 12646 | ENSMUSG00000030002 | Dusp11 | 72102 [Gene Symbol: Dusp11] [Locus Tag: ] [Chromosome: 6] [Map Location: 6\|6 C3] [ | 3.061471035 | 3.15E-10 | 2.44E-09 |
| 1422 | ENSMUSG00000057894 | Zfp329 | 67230 [Gene Symbol: Zfp329] [Locus Tag: ] [Chromosome: 7] [Map Location: 7\|7 A2] [D | 3.061189452 | 3.98E-07 | 1.91E-06 |
| 18594 | ENSMUSG00000020544 | Cox11 | 69802 [Gene Symbol: Cox11] [Locus Tag: ] [Chromosome: 11] [Map Location: 11\|11 D] | 3.060806209 | 4.30E-05 | 0.000145 |
| 2809 | ENSMUSG00000027111 | Itga6 | 16403 [Gene Symbol: Itga6] [Locus Tag: ] [Chromosome: 2] [Map Location: 2 C2-C3\|2 4 | 3.059454587 | 2.30E-08 | 1.36E-07 |
| 14965 | ENSMUSG00000033352 | Map2k4 | 26398 [Gene Symbol: Map2k4] [Locus Tag: ] [Chromosome: 11] [Map Location: 11 B3\|1 | 3.058973277 | 6.44E-11 | 5.58E-10 |
| 2561 | ENSMUSG00000032034 | Kcnj5 | 16521 [Gene Symbol: Kcnj5] [Locus Tag: ] [Chromosome: 9] [Map Location: 9 A4\|9 17. | 3.057461256 | 2.35E-05 | 8.36E-05 |
| 14722 | ENSMUSG00000078566 | Bnip3 | 12176 [Gene Symbol: Bnip3] [Locus Tag: ] [Chromosome: 7] [Map Location: 7\|7 F5] [De | 3.056941582 | 2.04E-06 | 8.70E-06 |
| 7428 | ENSMUSG00000055761 | Nkain3 | 269513 [Gene Symbol: Nkain3] [Locus Tag: ] [Chromosome: 4] [Map Location: 4\|4 A3] | 3.056469049 | 2.96E-10 | 2.30E-09 |
| 3432 | ENSMUSG00000041028 | Ghitm | 66092 [Gene Symbol: Ghitm] [Locus Tag: ] [Chromosome: 14] [Map Location: 14 B\|14 | 3.056132848 | 6.33E-08 | 3.46E-07 |
| 4992 | ENSMUSG00000032965 | Ift57 | 73916 [Gene Symbol: Ift57] [Locus Tag: ] [Chromosome: 16] [Map Location: 16\|16 B5] | 3.056070084 | 9.52E-07 | 4.30E-06 |
| 3146 | ENSMUSG00000048234 | Rnf149 | 67702 [Gene Symbol: Rnf149] [Locus Tag: ] [Chromosome: 1] [Map Location: 1\|1 B] [De | 3.055941643 | 1.02E-08 | 6.40E-08 |
| 10717 | ENSMUSG00000024067 | Dpy30 | 66310 [Gene Symbol: Dpy30] [Locus Tag: ] [Chromosome: 17] [Map Location: 17\|17 E2 | 3.055340952 | 9.22E-07 | 4.18E-06 |
| 17784 | ENSMUSG00000035376 | Hacd2 | 70757 [Gene Symbol: Hacd2] [Locus Tag: ] [Chromosome: 16] [Map Location: 16\|16 B3 | 3.052667662 | 2.01E-07 | 1.01E-06 |
| 13687 | ENSMUSG00000071660 | Ttc9c | 70387 [Gene Symbol: Ttc9c] [Locus Tag: ] [Chromosome: 19] [Map Location: 19\|19 A] [ | 3.052093068 | 8.67E-07 | 3.96E-06 |
| 17359 | ENSMUSG00000018589 | Glra2 | 237213 [Gene Symbol: Glra2] [Locus Tag: ] [Chromosome: X] [Map Location: X F5\|X 76. | 3.051545026 | 2.19E-09 | 1.50E-08 |
| 13109 | ENSMUSG00000028145 | Them4 | 75778 [Gene Symbol: Them4] [Locus Tag: ] [Chromosome: 3] [Map Location: 3\|3 F2.1] [ | 3.050213563 | 4.31E-08 | 2.44E-07 |
| 1731 | ENSMUSG00000066232 | Ipo7 | 233726 [Gene Symbol: Ipo7] [Locus Tag: ] [Chromosome: 7] [Map Location: 7\|7 F1] [De | 3.049938798 | 4.63E-10 | 3.50E-09 |
| 5526 | ENSMUSG00000022449 | Adamts20 | 223838 [Gene Symbol: Adamts20] [Locus Tag: ] [Chromosome: 15] [Map Location: 15 E | 3.048844325 | 4.93E-07 | 2.34E-06 |
| 7686 | ENSMUSG00000038510 | Rpf2 | 67239 [Gene Symbol: Rpf2] [Locus Tag: ] [Chromosome: 10] [Map Location: 10\|10 B1] [ | 3.047982875 | 5.67E-07 | 2.66E-06 |
| 15430 | ENSMUSG00000031536 | Polb | 18970 [Gene Symbol: Polb] [Locus Tag: ] [Chromosome: 8] [Map Location: 8 A2\|8 11.4 | 3.044696504 | 1.40E-07 | 7.29E-07 |
| 11859 | ENSMUSG00000047259 | Mc4r | 17202 [Gene Symbol: Mc4r] [Locus Tag: ] [Chromosome: 18] [Map Location: 18\|18 E1] | 3.044050875 | 0.001881 | 0.00461 |
| 6046 | ENSMUSG00000078851 | Hist3h2a | 319162 [Gene Symbol: Hist3h2a] [Locus Tag: ] [Chromosome: 11] [Map Location: 11\|1 | 3.042774413 | 9.24E-09 | 5.82E-08 |
| 16593 | ENSMUSG00000025525 | Apool | 68117 [Gene Symbol: Apool] [Locus Tag: ] [Chromosome: X] [Map Location: X\|X E2] [De | 3.04070234 | 0.003694 | 0.008478 |
| 7490 | ENSMUSG00000025959 | Klf7 | 93691 [Gene Symbol: Klf7] [Locus Tag: ] [Chromosome: 1] [Map Location: 1\|1 C1-C3] [D | 3.04069903 | 3.82E-10 | 2.92E-09 |
| 17252 | ENSMUSG00000036461 | Elf1 | 13709 [Gene Symbol: Elf1] [Locus Tag: ] [Chromosome: 14] [Map Location: 14\|14 D3] [ | 3.038730272 | 3.50E-08 | 2.00E-07 |
| 6611 | ENSMUSG00000028413 | B4galt1 | 14595 [Gene Symbol: B4galt1] [Locus Tag: ] [Chromosome: 4] [Map Location: 4 A5\|4 2 | 3.038455329 | 5.82E-07 | 2.73E-06 |
| 4207 | ENSMUSG00000053007 | Creb5 | 231991 [Gene Symbol: Creb5] [Locus Tag: ] [Chromosome: 6] [Map Location: 6 B3\|6 25 | 3.038329376 | 7.01E-06 | 2.73E-05 |
| 10048 | ENSMUSG00000015478 | Rnf5 | 54197 [Gene Symbol: Rnf5] [Locus Tag: ] [Chromosome: 17] [Map Location: 17 B1\|17 1 | 3.03830122 | 8.44E-09 | 5.35E-08 |
| 7013 | ENSMUSG00000027340 | Slc23a2 | 54338 [Gene Symbol: Slc23a2] [Locus Tag: ] [Chromosome: 2] [Map Location: 2\|2 G2] [ | 3.036771172 | 8.97E-10 | 6.51E-09 |
| 3747 | ENSMUSG00000052534 | Pbx1 | 18514 [Gene Symbol: Pbx1] [Locus Tag: ] [Chromosome: 1] [Map Location: 1 H2.3\|1 75 | 3.03623767 | 1.26E-10 | 1.04E-09 |
| 16644 | ENSMUSG00000021363 | Mak | 17152 [Gene Symbol: Mak] [Locus Tag: ] [Chromosome: 13] [Map Location: 13 A3.3\|13 | 3.035060997 | 3.08E-06 | 1.27E-05 |
| 11684 | ENSMUSG00000040918 | Slc19a2 | 116914 [Gene Symbol: Slc19a2] [Locus Tag: ] [Chromosome: 1] [Map Location: 1 H2.2\| | 3.034870625 | 3.96E-06 | 1.61E-05 |
| 733 | ENSMUSG00000027479 | Mapre1 | 13589 [Gene Symbol: Mapre1] [Locus Tag: ] [Chromosome: 2] [Map Location: 2 H1\|2 7 | 3.032615444 | 2.92E-10 | 2.28E-09 |
| 18424 | ENSMUSG00000027765 | P2ry1 | 18441 [Gene Symbol: P2ry1] [Locus Tag: ] [Chromosome: 3] [Map Location: 3\|3 D] [Des | 3.031868093 | 1.60E-09 | 1.12E-08 |

| | | | | | | |
|---|---|---|---|---|---|---|
| 16383 | ENSMUSG00000032184 | Lysmd2 | 70082 [Gene Symbol: Lysmd2] [Locus Tag: ] [Chromosome: 9] [Map Location: 9\|9 D] [D | 3.03088783 | 1.25E-06 | 5.55E-06 |
| 10072 | ENSMUSG00000038379 | Ttk | 22137 [Gene Symbol: Ttk] [Locus Tag: ] [Chromosome: 9] [Map Location: 9\|9 E2] [Descr | 3.030804545 | 0.00023 | 0.000677 |
| 14893 | ENSMUSG00000015759 | Cnih1 | 12793 [Gene Symbol: Cnih1] [Locus Tag: ] [Chromosome: 14] [Map Location: 14\|14 C1 | 3.028472956 | 9.63E-08 | 5.13E-07 |
| 16170 | ENSMUSG00000027680 | Fxr1 | 14359 [Gene Symbol: Fxr1] [Locus Tag: ] [Chromosome: 3] [Map Location: 3\|3 B] [Descr | 3.027669947 | 3.03E-10 | 2.36E-09 |
| 2274 | ENSMUSG00000052137 | Rbm12b2 | 77604 [Gene Symbol: Rbm12b2] [Locus Tag: ] [Chromosome: 4] [Map Location: 4\|4 A1 | 3.027157994 | 1.79E-08 | 1.08E-07 |
| 2922 | ENSMUSG00000018595 | Glra4 | 14657 [Gene Symbol: Glra4] [Locus Tag: ] [Chromosome: X] [Map Location: X F1\|X 59.1 | 3.025779373 | 0.020575 | 0.03952 |
| 6629 | ENSMUSG00000026915 | Strbp | 20744 [Gene Symbol: Strbp] [Locus Tag: ] [Chromosome: 2] [Map Location: 2\|2 B] [Desc | 3.024315985 | 6.26E-11 | 5.43E-10 |
| 95 | ENSMUSG00000003949 | Hlf | 217082 [Gene Symbol: Hlf] [Locus Tag: ] [Chromosome: 11] [Map Location: 11 C-D\|11 5 | 3.024034839 | 0.000349 | 0.000992 |
| 15649 | ENSMUSG00000039680 | Mrps6 | 121022 [Gene Symbol: Mrps6] [Locus Tag: ] [Chromosome: 16] [Map Location: 16\|16 C | 3.023247406 | 4.79E-08 | 2.68E-07 |
| 441 | ENSMUSG00000078572 | 1810043H04Rik | 208501 [Gene Symbol: 1810043H04Rik] [Locus Tag: ] [Chromosome: 11] [Map Locatior | 3.02136767 | 0.006894 | 0.014872 |
| 18144 | ENSMUSG00000026779 | Mastl | 67121 [Gene Symbol: Mastl] [Locus Tag: ] [Chromosome: 2] [Map Location: 2\|2 A3] [De | 3.021334661 | 4.21E-05 | 0.000142 |
| 16867 | ENSMUSG00000022895 | Ets2 | 23872 [Gene Symbol: Ets2] [Locus Tag: ] [Chromosome: 16] [Map Location: 16 56.64 cN | 3.018959286 | 3.36E-08 | 1.93E-07 |
| 13268 | ENSMUSG00000024807 | Syvn1 | 74126 [Gene Symbol: Syvn1] [Locus Tag: ] [Chromosome: 19] [Map Location: 19\|19 A] [ | 3.018956749 | 1.26E-06 | 5.59E-06 |
| 7654 | ENSMUSG00000063659 | Zbtb18 | 30928 [Gene Symbol: Zbtb18] [Locus Tag: ] [Chromosome: 1] [Map Location: 1\|1 H3] [D | 3.017078992 | 9.89E-11 | 8.29E-10 |
| 1452 | ENSMUSG00000067365 | Tmem128 | 66309 [Gene Symbol: Tmem128] [Locus Tag: ] [Chromosome: 5] [Map Location: 5\|5 B3 | 3.01660328 | 1.73E-06 | 7.52E-06 |
| 241 | ENSMUSG00000036202 | Rif1 | 51869 [Gene Symbol: Rif1] [Locus Tag: ] [Chromosome: 2] [Map Location: 2 C1.1\|2 29.9 | 3.013104681 | 5.31E-09 | 3.46E-08 |
| 17414 | ENSMUSG00000039958 | Mettl20 | 320204 [Gene Symbol: Mettl20] [Locus Tag: ] [Chromosome: 6] [Map Location: 6\|6 G3] | 3.013016882 | 0.0001 | 0.000314 |
| 18532 | ENSMUSG00000047412 | Zbtb44 | 235132 [Gene Symbol: Zbtb44] [Locus Tag: ] [Chromosome: 9] [Map Location: 9\|9 A4] [ | 3.012233515 | 8.75E-11 | 7.39E-10 |
| 11000 | ENSMUSG00000034701 | Neurod1 | 18012 [Gene Symbol: Neurod1] [Locus Tag: ] [Chromosome: 2] [Map Location: 2 C3\|2 4 | 3.009225381 | 2.20E-07 | 1.10E-06 |
| 9759 | ENSMUSG00000053070 | 9230110C19Rik | 234912 [Gene Symbol: 9230110C19Rik] [Locus Tag: ] [Chromosome: 9] [Map Location: | 3.008992915 | 7.65E-06 | 2.97E-05 |
| 2844 | ENSMUSG00000020184 | Mdm2 | 17246 [Gene Symbol: Mdm2] [Locus Tag: ] [Chromosome: 10] [Map Location: 10 66.32 | 3.007789248 | 4.23E-10 | 3.22E-09 |
| 6648 | ENSMUSG00000039841 | Zfp800 | 627049 [Gene Symbol: Zfp800] [Locus Tag: ] [Chromosome: 6] [Map Location: 6\|6 A3.2 | 3.007291009 | 1.39E-09 | 9.83E-09 |
| 13609 | ENSMUSG00000027999 | Pla2g12a | 66350 [Gene Symbol: Pla2g12a] [Locus Tag: ] [Chromosome: 3] [Map Location: 3\|3 H1] | 3.006635409 | 8.50E-07 | 3.88E-06 |
| 18535 | ENSMUSG00000069208 | Zfp825 | 235956 [Gene Symbol: Zfp825] [Locus Tag: ] [Chromosome: 13] [Map Location: 13\|13 ( | 3.005657115 | 0.002287 | 0.005497 |
| 13450 | ENSMUSG00000073676 | Hspe1 | 15528 [Gene Symbol: Hspe1] [Locus Tag: ] [Chromosome: 1] [Map Location: 1\|1 C1] [De | 3.003417452 | 1.06E-06 | 4.74E-06 |
| 11772 | ENSMUSG00000034327 | Kctd9 | 105440 [Gene Symbol: Kctd9] [Locus Tag: ] [Chromosome: 14] [Map Location: 14\|14 D | 3.002801854 | 6.63E-06 | 2.59E-05 |
| 13454 | ENSMUSG00000069072 | Slc7a14 | 241919 [Gene Symbol: Slc7a14] [Locus Tag: ] [Chromosome: 3] [Map Location: 3\|3 A3] | 3.001231144 | 1.13E-07 | 5.94E-07 |
| 12430 | ENSMUSG00000053730 | Tmem39b | 230770 [Gene Symbol: Tmem39b] [Locus Tag: ] [Chromosome: 4] [Map Location: 4\|4 D | 3.001004699 | 4.04E-06 | 1.64E-05 |
| 7809 | ENSMUSG00000032252 | Glce | 93683 [Gene Symbol: Glce] [Locus Tag: ] [Chromosome: 9] [Map Location: 9\|9 C-D] [De | 3.000296567 | 9.72E-10 | 7.01E-09 |
| 4050 | ENSMUSG00000015880 | Ncapg | 54392 [Gene Symbol: Ncapg] [Locus Tag: ] [Chromosome: 5] [Map Location: 5\|5 B3] [De | 2.999318484 | 2.75E-08 | 1.61E-07 |
| 909 | ENSMUSG00000003452 | Bicd1 | 12121 [Gene Symbol: Bicd1] [Locus Tag: ] [Chromosome: 6] [Map Location: 6\|6 G3] [De | 2.997072514 | 8.92E-10 | 6.48E-09 |
| 13660 | ENSMUSG00000036769 | Wdr44 | 72404 [Gene Symbol: Wdr44] [Locus Tag: ] [Chromosome: X] [Map Location: X\|X A2] [D | 2.996826866 | 1.17E-08 | 7.25E-08 |
| 17831 | ENSMUSG00000030016 | Zfp638 | 18139 [Gene Symbol: Zfp638] [Locus Tag: ] [Chromosome: 6] [Map Location: 6\|6 C3] [D | 2.996401114 | 1.23E-10 | 1.02E-09 |
| 10356 | ENSMUSG00000075256 | Cerkl | 228094 [Gene Symbol: Cerkl] [Locus Tag: ] [Chromosome: 2] [Map Location: 2\|2 C3] [De | 2.995991459 | 1.32E-07 | 6.91E-07 |
| 9516 | ENSMUSG00000019873 | Reep3 | 28193 [Gene Symbol: Reep3] [Locus Tag: ] [Chromosome: 10] [Map Location: 10 B5.1\|1 | 2.994570695 | 1.65E-09 | 1.15E-08 |
| 3153 | ENSMUSG00000033282 | Rpgrip1l | 244585 [Gene Symbol: Rpgrip1l] [Locus Tag: ] [Chromosome: 8] [Map Location: 8 C5\|8 | 2.994079001 | 1.27E-06 | 5.64E-06 |
| 16133 | ENSMUSG00000034681 | Rnps1 | 19826 [Gene Symbol: Rnps1] [Locus Tag: ] [Chromosome: 17] [Map Location: 17\|17 A3 | 2.992578692 | 6.44E-10 | 4.77E-09 |
| 14283 | ENSMUSG00000004880 | Lbr | 98386 [Gene Symbol: Lbr] [Locus Tag: ] [Chromosome: 1] [Map Location: 1 H5\|1 84.89 | 2.992155722 | 2.82E-07 | 1.39E-06 |
| 13250 | ENSMUSG00000022369 | Mtbp | 105837 [Gene Symbol: Mtbp] [Locus Tag: ] [Chromosome: 15] [Map Location: 15\|15 D1 | 2.99171675 | 8.97E-06 | 3.43E-05 |
| 11217 | ENSMUSG00000028820 | Sfpq | 71514 [Gene Symbol: Sfpq] [Locus Tag: ] [Chromosome: 4] [Map Location: 4 61.21 cM\|4 | 2.991602392 | 1.20E-10 | 9.94E-10 |
| 756 | ENSMUSG00000031370 | Zrsr2 | 22184 [Gene Symbol: Zrsr2] [Locus Tag: ] [Chromosome: X] [Map Location: X F5\|X 75.9 | 2.990783932 | 4.11E-09 | 2.72E-08 |
| 14670 | ENSMUSG00000040242 | Fgfr1op2 | 67529 [Gene Symbol: Fgfr1op2] [Locus Tag: ] [Chromosome: 6] [Map Location: 6\|6 G3] | 2.990436087 | 4.22E-08 | 2.38E-07 |
| 2029 | ENSMUSG00000032727 | Mier3 | 218613 [Gene Symbol: Mier3] [Locus Tag: ] [Chromosome: 13] [Map Location: 13\|13 D | 2.990143554 | 1.88E-09 | 1.30E-08 |
| 14212 | ENSMUSG00000008859 | Rala | 56044 [Gene Symbol: Rala] [Locus Tag: ] [Chromosome: 13] [Map Location: 13 6.05 cM | 2.987950363 | 1.48E-10 | 1.21E-09 |
| 6530 | ENSMUSG00000022142 | Nup155 | 170762 [Gene Symbol: Nup155] [Locus Tag: ] [Chromosome: 15] [Map Location: 15\|15 | 2.985380774 | 3.20E-09 | 2.15E-08 |
| 14405 | ENSMUSG00000051223 | Bzw1 | 66882 [Gene Symbol: Bzw1] [Locus Tag: ] [Chromosome: 1] [Map Location: 1\|1 C2] [De | 2.98473279 | 3.69E-10 | 2.83E-09 |
| 1240 | ENSMUSG00000020268 | Lyrm7 | 75530 [Gene Symbol: Lyrm7] [Locus Tag: ] [Chromosome: 11] [Map Location: 11\|11 B1 | 2.984170971 | 6.46E-06 | 2.53E-05 |
| 6872 | ENSMUSG00000030872 | Gga2 | 74105 [Gene Symbol: Gga2] [Locus Tag: ] [Chromosome: 7] [Map Location: 7\|7 F3] [Des | 2.983249271 | 6.58E-09 | 4.24E-08 |
| 3583 | ENSMUSG00000040128 | Pnrc1 | 108767 [Gene Symbol: Pnrc1] [Locus Tag: ] [Chromosome: 4] [Map Location: 4\|4 A5] [D | 2.980907269 | 6.25E-08 | 3.43E-07 |
| 18309 | ENSMUSG00000028347 | Tmeff1 | 230157 [Gene Symbol: Tmeff1] [Locus Tag: ] [Chromosome: 4] [Map Location: 4\|4 B2] [ | 2.980546432 | 1.69E-10 | 1.37E-09 |
| 3995 | ENSMUSG00000049164 | Zfp518a | 72672 [Gene Symbol: Zfp518a] [Locus Tag: ] [Chromosome: 19] [Map Location: 19\|19 D | 2.980514566 | 2.51E-08 | 1.47E-07 |
| 5268 | ENSMUSG00000021500 | Ddx46 | 212880 [Gene Symbol: Ddx46] [Locus Tag: ] [Chromosome: 13] [Map Location: 13\|13 B | 2.979781232 | 7.32E-07 | 3.38E-06 |
| 4397 | ENSMUSG00000040374 | Pex2 | 19302 [Gene Symbol: Pex2] [Locus Tag: ] [Chromosome: 3] [Map Location: 3 A1\|3 1.96 | 2.979685406 | 3.12E-07 | 1.52E-06 |
| 10627 | ENSMUSG00000026502 | Desi2 | 78825 [Gene Symbol: Desi2] [Locus Tag: ] [Chromosome: 1] [Map Location: 1\|1 H3] [De | 2.977578972 | 7.10E-09 | 4.54E-08 |
| 701 | ENSMUSG00000026663 | Atf6 | 226641 [Gene Symbol: Atf6] [Locus Tag: ] [Chromosome: 1] [Map Location: 1 H3\|1 76.9 | 2.975018402 | 1.43E-09 | 1.01E-08 |
| 8875 | ENSMUSG00000027965 | Olfm3 | 229759 [Gene Symbol: Olfm3] [Locus Tag: ] [Chromosome: 3] [Map Location: 3\|3 F3] [D | 2.974841799 | 1.16E-05 | 4.36E-05 |
| 13712 | ENSMUSG00000032314 | Etfa | 110842 [Gene Symbol: Etfa] [Locus Tag: ] [Chromosome: 9] [Map Location: 9\|9 B] [Desc | 2.973803115 | 3.81E-05 | 0.00013 |
| 10013 | ENSMUSG00000036006 | Fam65b | 193385 [Gene Symbol: Fam65b] [Locus Tag: ] [Chromosome: 13] [Map Location: 13\|13 | 2.97291147 | 4.96E-09 | 3.25E-08 |
| 15538 | ENSMUSG00000091264 | Smim13 | 108934 [Gene Symbol: Smim13] [Locus Tag: ] [Chromosome: 13] [Map Location: 13A4\| | 2.972691996 | 2.14E-10 | 1.71E-09 |
| 4901 | ENSMUSG00000036572 | Upf3b | 68134 [Gene Symbol: Upf3b] [Locus Tag: ] [Chromosome: X] [Map Location: X\|X A2] [De | 2.970503789 | 2.85E-06 | 1.18E-05 |
| 5789 | ENSMUSG00000070934 | Rraga | 68441 [Gene Symbol: Rraga] [Locus Tag: ] [Chromosome: 4] [Map Location: 4\|4 C4] [De | 2.970006256 | 1.35E-08 | 8.24E-08 |
| 14702 | ENSMUSG00000020841 | Cpd | 12874 [Gene Symbol: Cpd] [Locus Tag: ] [Chromosome: 11] [Map Location: 11 B5\|11 46 | 2.969425334 | 3.63E-10 | 2.78E-09 |
| 4089 | ENSMUSG00000029486 | Mrpl1 | 94061 [Gene Symbol: Mrpl1] [Locus Tag: ] [Chromosome: 5] [Map Location: 5\|5 E3] [De | 2.968963762 | 2.21E-07 | 1.11E-06 |
| 9478 | ENSMUSG00000018427 | Ypel2 | 77864 [Gene Symbol: Ypel2] [Locus Tag: ] [Chromosome: 11] [Map Location: 11\|11 C] [ | 2.968609867 | 2.87E-06 | 1.20E-05 |
| 16575 | ENSMUSG00000029213 | Commd8 | 27784 [Gene Symbol: Commd8] [Locus Tag: ] [Chromosome: 5] [Map Location: 5 C3.2\| | 2.968421076 | 3.88E-08 | 2.20E-07 |
| 5531 | ENSMUSG00000029634 | Rnf6 | 74132 [Gene Symbol: Rnf6] [Locus Tag: ] [Chromosome: 5] [Map Location: 5\|5 G3] [Des | 2.968097148 | 2.37E-06 | 1.00E-05 |
| 8004 | ENSMUSG00000042408 | Zmym6 | 100177 [Gene Symbol: Zmym6] [Locus Tag: ] [Chromosome: 4] [Map Location: 4 D2.2\|4 | 2.967767432 | 2.49E-09 | 1.69E-08 |
| 6712 | ENSMUSG00000041695 | Kcnj2 | 16518 [Gene Symbol: Kcnj2] [Locus Tag: ] [Chromosome: 11] [Map Location: 11 E2\|11 | 2.967715405 | 9.28E-09 | 5.84E-08 |
| 14132 | ENSMUSG00000026020 | Nop58 | 55989 [Gene Symbol: Nop58] [Locus Tag: ] [Chromosome: 1] [Map Location: 1\|1 C1.3] | 2.967147068 | 1.46E-06 | 6.41E-06 |
| 12325 | ENSMUSG00000035227 | Spcs2 | 66624 [Gene Symbol: Spcs2] [Locus Tag: ] [Chromosome: 7] [Map Location: 7\|7 E2] [De | 2.96605271 | 4.45E-08 | 2.51E-07 |
| 1153 | ENSMUSG00000071267 | Zfp942 | 73233 [Gene Symbol: Zfp942] [Locus Tag: ] [Chromosome: 17] [Map Location: 17\|17 A3 | 2.965550536 | 2.10E-07 | 1.05E-06 |
| 14064 | ENSMUSG00000007827 | Ankrd26 | 232339 [Gene Symbol: Ankrd26] [Locus Tag: ] [Chromosome: 6] [Map Location: 6\|6 F1] | 2.964615989 | 3.36E-07 | 1.63E-06 |
| 11425 | ENSMUSG00000041012 | Cmtm8 | 70031 [Gene Symbol: Cmtm8] [Locus Tag: ] [Chromosome: 9] [Map Location: 9\|9 F3] [D | 2.962829658 | 0.004357 | 0.009835 |
| 2934 | ENSMUSG00000025395 | Prim1 | 19075 [Gene Symbol: Prim1] [Locus Tag: ] [Chromosome: 10] [Map Location: 10 76.39 | 2.962370771 | 1.77E-05 | 6.46E-05 |
| 13275 | ENSMUSG00000020803 | Txndc17 | 52700 [Gene Symbol: Txndc17] [Locus Tag: ] [Chromosome: 11] [Map Location: 11 B4\| | 2.961959111 | 2.63E-07 | 1.30E-06 |
| 2968 | ENSMUSG00000057982 | Zfp809 | 235047 [Gene Symbol: Zfp809] [Locus Tag: ] [Chromosome: 9] [Map Location: 9\|9 A3] [ | 2.9607559 | 3.17E-06 | 1.31E-05 |
| 8054 | ENSMUSG00000021929 | Kpna3 | 16648 [Gene Symbol: Kpna3] [Locus Tag: ] [Chromosome: 14] [Map Location: 14\|14 D1 | 2.959189263 | 6.25E-10 | 4.64E-09 |
| 13473 | ENSMUSG00000004591 | Pkn2 | 109333 [Gene Symbol: Pkn2] [Locus Tag: ] [Chromosome: 3] [Map Location: 3 H1\|3 66. | 2.958656781 | 2.88E-09 | 1.95E-08 |
| 13700 | ENSMUSG00000029012 | Orc5 | 26429 [Gene Symbol: Orc5] [Locus Tag: ] [Chromosome: 5] [Map Location: 5\|5 A3] [Des | 2.953691665 | 1.69E-06 | 7.37E-06 |
| 8265 | ENSMUSG00000025134 | Alyref | 21681 [Gene Symbol: Alyref] [Locus Tag: ] [Chromosome: 11] [Map Location: 11\|11 E2] | 2.95316126 | 1.20E-08 | 7.43E-08 |
| 6463 | ENSMUSG00000016481 | Cr1l | 12946 [Gene Symbol: Cr1l] [Locus Tag: ] [Chromosome: 1] [Map Location: 1 H6\|1 98.43 | 2.9508071 | 6.49E-08 | 3.54E-07 |
| 10903 | ENSMUSG00000027203 | Dut | 110074 [Gene Symbol: Dut] [Locus Tag: ] [Chromosome: 2] [Map Location: 2 F1\|2 61.3 | 2.949012118 | 8.28E-08 | 4.45E-07 |
| 17673 | ENSMUSG00000051232 | Tmem199 | 195040 [Gene Symbol: Tmem199] [Locus Tag: ] [Chromosome: 11] [Map Location: 11\| | 2.948038974 | 6.23E-06 | 2.44E-05 |
| 12866 | ENSMUSG00000040423 | Rc3h1 | 381305 [Gene Symbol: Rc3h1] [Locus Tag: ] [Chromosome: 1] [Map Location: 1\|1 H2.1 | 2.947975664 | 3.95E-10 | 3.01E-09 |

| | | | | | | |
|---|---|---|---|---|---|---|
| 1281 | ENSMUSG00000038323 | 1700066M21Rik | 73467 [Gene Symbol: 1700066M21Rik] [Locus Tag: ] [Chromosome: 1] [Map Location: | 2.946688009 | 4.16E-08 | 2.35E-07 |
| 4442 | ENSMUSG00000037685 | Atp8a1 | 11980 [Gene Symbol: Atp8a1] [Locus Tag: ] [Chromosome: 5] [Map Location: 5\|5 C3.1] | 2.946313267 | 3.28E-10 | 2.53E-09 |
| 4931 | ENSMUSG00000060798 | Intu | 380614 [Gene Symbol: Intu] [Locus Tag: ] [Chromosome: 3] [Map Location: 3\|3 B] [Desc | 2.945258346 | 1.26E-08 | 7.79E-08 |
| 7960 | ENSMUSG00000029276 | Glmn | 170823 [Gene Symbol: Glmn] [Locus Tag: ] [Chromosome: 5] [Map Location: 5\|5 E5] [De | 2.945195921 | 0.000121 | 0.000374 |
| 9441 | ENSMUSG00000027715 | Ccna2 | 12428 [Gene Symbol: Ccna2] [Locus Tag: ] [Chromosome: 3] [Map Location: 3 B\|3 17.6 | 2.943503837 | 0.0007 | 0.001876 |
| 8043 | ENSMUSG00000029376 | Mthfd2l | 665563 [Gene Symbol: Mthfd2l] [Locus Tag: ] [Chromosome: 5] [Map Location: 5\|5 E1] | 2.943297426 | 1.14E-07 | 6.01E-07 |
| 7386 | ENSMUSG00000022912 | Pros1 | 19128 [Gene Symbol: Pros1] [Locus Tag: ] [Chromosome: 16] [Map Location: 16\|16 C1. | 2.941331425 | 1.27E-05 | 4.74E-05 |
| 3339 | ENSMUSG00000027739 | Rab33b | 19338 [Gene Symbol: Rab33b] [Locus Tag: ] [Chromosome: 3] [Map Location: 3\|3 D] [D | 2.939138332 | 5.53E-09 | 3.60E-08 |
| 1916 | ENSMUSG00000038225 | Primpol | 408022 [Gene Symbol: Primpol] [Locus Tag: ] [Chromosome: 8] [Map Location: 8\|8 B1. | 2.93821981 | 5.78E-06 | 2.28E-05 |
| 12109 | ENSMUSG00000028525 | Pde4b | 18578 [Gene Symbol: Pde4b] [Locus Tag: ] [Chromosome: 4] [Map Location: 4 C6\|4 46. | 2.937229352 | 3.42E-07 | 1.66E-06 |
| 12340 | ENSMUSG00000009621 | Vav2 | 22325 [Gene Symbol: Vav2] [Locus Tag: ] [Chromosome: 2] [Map Location: 2 A3\|2 19.3 | 2.937093363 | 2.75E-10 | 2.15E-09 |
| 1805 | ENSMUSG00000055943 | Emc7 | 73024 [Gene Symbol: Emc7] [Locus Tag: ] [Chromosome: 2] [Map Location: 2\|2 E4] [De | 2.936667584 | 2.71E-08 | 1.58E-07 |
| 18397 | ENSMUSG00000068015 | Lrch1 | 380916 [Gene Symbol: Lrch1] [Locus Tag: ] [Chromosome: 14] [Map Location: 14\|14 D | 2.935717921 | 2.16E-09 | 1.48E-08 |
| 4222 | ENSMUSG00000024750 | Zfand5 | 22682 [Gene Symbol: Zfand5] [Locus Tag: ] [Chromosome: 19] [Map Location: 19 B\|19 | 2.93481939 | 8.15E-10 | 5.95E-09 |
| 4527 | ENSMUSG00000019789 | Hey2 | 15214 [Gene Symbol: Hey2] [Locus Tag: ] [Chromosome: 10] [Map Location: 10\|10 A4] | 2.932083046 | 0.00088 | 0.002315 |
| 10131 | ENSMUSG00000047343 | Mettl21c | 433294 [Gene Symbol: Mettl21c] [Locus Tag: ] [Chromosome: 1] [Map Location: 1\|1 C1 | 2.932070707 | 0.004988 | 0.011121 |
| 11389 | ENSMUSG00000030760 | Acer3 | 66190 [Gene Symbol: Acer3] [Locus Tag: ] [Chromosome: 7] [Map Location: 7\|7 E2] [De | 2.931741592 | 2.74E-05 | 9.63E-05 |
| 6277 | ENSMUSG00000020305 | Asb3 | 65257 [Gene Symbol: Asb3] [Locus Tag: ] [Chromosome: 11] [Map Location: 11 A4\|11 1 | 2.931695301 | 7.03E-08 | 3.82E-07 |
| 13420 | ENSMUSG00000027082 | Tfpi | 21788 [Gene Symbol: Tfpi] [Locus Tag: ] [Chromosome: 2] [Map Location: 2\|2 D] [Descri | 2.931540833 | 0.000223 | 0.000659 |
| 9224 | ENSMUSG00000022035 | Ccdc25 | 67179 [Gene Symbol: Ccdc25] [Locus Tag: ] [Chromosome: 14] [Map Location: 14\|14 D | 2.930718849 | 1.35E-07 | 7.03E-07 |
| 1102 | ENSMUSG00000029206 | Nsun7 | 70918 [Gene Symbol: Nsun7] [Locus Tag: ] [Chromosome: 5] [Map Location: 5\|5 D] [De | 2.930490524 | 0.00078 | 0.00207 |
| 12624 | ENSMUSG00000041058 | Wwp1 | 107568 [Gene Symbol: Wwp1] [Locus Tag: ] [Chromosome: 4] [Map Location: 4\|4 A3] [ | 2.929153211 | 1.46E-08 | 8.90E-08 |
| 1730 | ENSMUSG00000038642 | Ctss | 13040 [Gene Symbol: Ctss] [Locus Tag: ] [Chromosome: 3] [Map Location: 3 F2.1\|3 40. | 2.928373624 | 0.000158 | 0.000479 |
| 18430 | ENSMUSG00000024228 | Nudt12 | 67993 [Gene Symbol: Nudt12] [Locus Tag: ] [Chromosome: 17] [Map Location: 17\|17 E | 2.926520757 | 0.001563 | 0.003903 |
| 12383 | ENSMUSG00000031429 | Psmd10 | 53380 [Gene Symbol: Psmd10] [Locus Tag: ] [Chromosome: X] [Map Location: X\|X F1] [ | 2.924430309 | 0.000144 | 0.00044 |
| 7336 | ENSMUSG00000031099 | Smarca1 | 93761 [Gene Symbol: Smarca1] [Locus Tag: ] [Chromosome: X] [Map Location: X\|X A3.2 | 2.924308884 | 3.18E-06 | 1.31E-05 |
| 5088 | ENSMUSG00000033904 | Ccp110 | 101565 [Gene Symbol: Ccp110] [Locus Tag: ] [Chromosome: 7] [Map Location: 7\|7 F2] | 2.924236373 | 7.62E-09 | 4.85E-08 |
| 13003 | ENSMUSG00000020134 | Peli1 | 67245 [Gene Symbol: Peli1] [Locus Tag: ] [Chromosome: 11] [Map Location: 11 13.81 c | 2.92413478 | 5.33E-08 | 2.95E-07 |
| 9180 | ENSMUSG00000022105 | Rb1 | 19645 [Gene Symbol: Rb1] [Locus Tag: ] [Chromosome: 14] [Map Location: 14 38.73 cM | 2.921842173 | 3.47E-09 | 2.32E-08 |
| 18570 | ENSMUSG00000037652 | Phc3 | 241915 [Gene Symbol: Phc3] [Locus Tag: ] [Chromosome: 3] [Map Location: 3\|3 A3] [De | 2.920295617 | 5.64E-10 | 4.21E-09 |
| 7032 | ENSMUSG00000022257 | Laptm4b | 114128 [Gene Symbol: Laptm4b] [Locus Tag: ] [Chromosome: 15] [Map Location: 15\|1 | 2.917458216 | 3.31E-05 | 0.000115 |
| 7806 | ENSMUSG00000073702 | Rpl31 | 114641 [Gene Symbol: Rpl31] [Locus Tag: ] [Chromosome: 1] [Map Location: 1\|1 B] [De | 2.916499482 | 0.006703 | 0.014493 |
| 998 | ENSMUSG00000008429 | Herpud2 | 80517 [Gene Symbol: Herpud2] [Locus Tag: ] [Chromosome: 9] [Map Location: 9\|9 A4] | 2.914991139 | 1.01E-09 | 7.24E-09 |
| 12379 | ENSMUSG00000046230 | Vps13a | 271564 [Gene Symbol: Vps13a] [Locus Tag: ] [Chromosome: 19] [Map Location: 19\|19 | 2.911674727 | 3.12E-06 | 1.29E-05 |
| 15496 | ENSMUSG00000032035 | Ets1 | 23871 [Gene Symbol: Ets1] [Locus Tag: ] [Chromosome: 9] [Map Location: 9 A4\|9 17.97 | 2.907957981 | 1.73E-08 | 1.04E-07 |
| 14017 | ENSMUSG00000022668 | Gtpbp8 | 66067 [Gene Symbol: Gtpbp8] [Locus Tag: ] [Chromosome: 16] [Map Location: 16\|16 B | 2.907562591 | 1.87E-07 | 9.47E-07 |
| 3863 | ENSMUSG00000040621 | Gemin8 | 237221 [Gene Symbol: Gemin8] [Locus Tag: ] [Chromosome: X] [Map Location: X\|X F5] | 2.907352132 | 0.002188 | 0.00528 |
| 10011 | ENSMUSG00000030929 | Eri2 | 71151 [Gene Symbol: Eri2] [Locus Tag: ] [Chromosome: 7] [Map Location: 7\|7 F3] [Desc | 2.907160612 | 2.90E-05 | 0.000101 |
| 10109 | ENSMUSG00000068859 | Sp9 | 381373 [Gene Symbol: Sp9] [Locus Tag: ] [Chromosome: 2] [Map Location: 2\|2 C3] [Des | 2.90420123 | 5.18E-07 | 2.45E-06 |
| 9508 | ENSMUSG00000032370 | Lactb | 80907 [Gene Symbol: Lactb] [Locus Tag: ] [Chromosome: 9] [Map Location: 9\|9 D] [Desc | 2.904082396 | 3.88E-06 | 1.58E-05 |
| 7148 | ENSMUSG00000020124 | Usp15 | 14479 [Gene Symbol: Usp15] [Locus Tag: ] [Chromosome: 10] [Map Location: 10\|10 D3 | 2.903929406 | 3.38E-06 | 1.39E-05 |
| 16105 | ENSMUSG00000063275 | Hacd1 | 30963 [Gene Symbol: Hacd1] [Locus Tag: ] [Chromosome: 2] [Map Location: 2\|2 A2] [De | 2.903297277 | 6.09E-06 | 2.40E-05 |
| 13251 | ENSMUSG00000032258 | Lca5 | 75782 [Gene Symbol: Lca5] [Locus Tag: ] [Chromosome: 9] [Map Location: 9\|9 E2] [Desc | 2.901075913 | 4.42E-07 | 2.12E-06 |
| 8024 | ENSMUSG00000022311 | Csmd3 | 239420 [Gene Symbol: Csmd3] [Locus Tag: ] [Chromosome: 15] [Map Location: 15\|15 | 2.901014408 | 4.89E-06 | 1.95E-05 |
| 13594 | ENSMUSG00000024516 | Sec11c | 66286 [Gene Symbol: Sec11c] [Locus Tag: ] [Chromosome: 18] [Map Location: 18\|18 E1 | 2.899398266 | 5.12E-05 | 0.00017 |
| 5229 | ENSMUSG00000008035 | Mid1ip1 | 68041 [Gene Symbol: Mid1ip1] [Locus Tag: ] [Chromosome: X] [Map Location: X\|X A1.2 | 2.8971107 | 4.05E-09 | 2.69E-08 |
| 2205 | ENSMUSG00000010914 | Pdhx | 27402 [Gene Symbol: Pdhx] [Locus Tag: ] [Chromosome: 2] [Map Location: 2\|2 E2] [Des | 2.895547245 | 2.96E-08 | 1.71E-07 |
| 15064 | ENSMUSG00000045518 | Onecut3 | 246086 [Gene Symbol: Onecut3] [Locus Tag: ] [Chromosome: 10] [Map Location: 10\|1 | 2.895396829 | 0.000294 | 0.000848 |
| 7348 | ENSMUSG00000044629 | Cnrip1 | 380686 [Gene Symbol: Cnrip1] [Locus Tag: ] [Chromosome: 11] [Map Location: 11\|11 | 2.895053404 | 9.49E-10 | 6.87E-09 |
| 6245 | ENSMUSG00000032846 | Zswim6 | 67263 [Gene Symbol: Zswim6] [Locus Tag: ] [Chromosome: 13] [Map Location: 13\|13 D | 2.893856081 | 9.24E-10 | 6.69E-09 |
| 11904 | ENSMUSG00000079316 | Rab9 | 56382 [Gene Symbol: Rab9] [Locus Tag: ] [Chromosome: X] [Map Location: X\|X F5] [Des | 2.893682231 | 3.31E-06 | 1.36E-05 |
| 9424 | ENSMUSG00000028020 | Glrb | 14658 [Gene Symbol: Glrb] [Locus Tag: ] [Chromosome: 3] [Map Location: 3 35.71 cM\| | 2.893235617 | 7.16E-08 | 3.88E-07 |
| 13967 | ENSMUSG00000004317 | Clcn5 | 12728 [Gene Symbol: Clcn5] [Locus Tag: ] [Chromosome: X] [Map Location: X A1.1\|X 3. | 2.893214547 | 6.24E-09 | 4.03E-08 |
| 5989 | ENSMUSG00000054604 | Cggbp1 | 106143 [Gene Symbol: Cggbp1] [Locus Tag: ] [Chromosome: 16] [Map Location: 16\|16 | 2.89306683 | 1.65E-09 | 1.15E-08 |
| 8850 | ENSMUSG00000079418 | Atg4a | 666468 [Gene Symbol: Atg4a] [Locus Tag: ] [Chromosome: X] [Map Location: X\|X F1] [D | 2.890345824 | 0.000223 | 0.000659 |
| 12365 | ENSMUSG00000032412 | Atp1b3 | 11933 [Gene Symbol: Atp1b3] [Locus Tag: ] [Chromosome: 9] [Map Location: 9 E3.3\|9 | 2.886613021 | 5.00E-09 | 3.27E-08 |
| 1758 | ENSMUSG00000021460 | Auh | 11992 [Gene Symbol: Auh] [Locus Tag: ] [Chromosome: 13] [Map Location: 13\|13 B1] [ | 2.886256068 | 3.34E-08 | 1.92E-07 |
| 3305 | ENSMUSG00000061983 | Rps12 | 20042 [Gene Symbol: Rps12] [Locus Tag: ] [Chromosome: 10] [Map Location: 10\|10 A3 | 2.884354541 | 0.00328 | 0.007621 |
| 15872 | ENSMUSG00000019897 | Ccdc59 | 52713 [Gene Symbol: Ccdc59] [Locus Tag: ] [Chromosome: 10] [Map Location: 10 D1\|1 | 2.883351881 | 8.94E-05 | 0.000283 |
| 1696 | ENSMUSG00000021994 | Wnt5a | 22418 [Gene Symbol: Wnt5a] [Locus Tag: ] [Chromosome: 14] [Map Location: 14 A3\|1 | 2.882855736 | 9.61E-09 | 6.03E-08 |
| 9638 | ENSMUSG00000001998 | Ap4e1 | 108011 [Gene Symbol: Ap4e1] [Locus Tag: ] [Chromosome: 2] [Map Location: 2 F1\|2 61 | 2.881009844 | 2.81E-08 | 1.63E-07 |
| 17244 | ENSMUSG00000018377 | Vezf1 | 22344 [Gene Symbol: Vezf1] [Locus Tag: ] [Chromosome: 11] [Map Location: 11\|11 C] [ | 2.876631865 | 7.91E-09 | 5.03E-08 |
| 7644 | ENSMUSG00000035941 | Ibtk | 108837 [Gene Symbol: Ibtk] [Locus Tag: ] [Chromosome: 9] [Map Location: 9\|9 E3.1] [D | 2.875463022 | 5.50E-07 | 2.59E-06 |
| 1750 | ENSMUSG00000000001 | Gnai3 | 14679 [Gene Symbol: Gnai3] [Locus Tag: ] [Chromosome: 3] [Map Location: 3 F2.3\|3 4 | 2.874504854 | 6.78E-09 | 4.35E-08 |
| 2317 | ENSMUSG00000031715 | Smarca5 | 93762 [Gene Symbol: Smarca5] [Locus Tag: ] [Chromosome: 8] [Map Location: 8\|8 C2] | 2.873922168 | 7.13E-08 | 3.87E-07 |
| 9001 | ENSMUSG00000033429 | Mcee | 73724 [Gene Symbol: Mcee] [Locus Tag: ] [Chromosome: 7] [Map Location: 7\|7 C] [Desc | 2.871171631 | 0.000516 | 0.001421 |
| 4731 | ENSMUSG00000036779 | Papd5 | 214627 [Gene Symbol: Papd5] [Locus Tag: ] [Chromosome: 8] [Map Location: 8\|8 C3] [ | 2.870593644 | 1.86E-09 | 1.29E-08 |
| 11766 | ENSMUSG00000038151 | Prdm1 | 12142 [Gene Symbol: Prdm1] [Locus Tag: ] [Chromosome: 10] [Map Location: 10 B2\|1 | 2.870584783 | 0.000322 | 0.000924 |
| 14609 | ENSMUSG00000049658 | Bdp1 | 544971 [Gene Symbol: Bdp1] [Locus Tag: ] [Chromosome: 13] [Map Location: 13 D1\|1 | 2.867923995 | 3.40E-08 | 1.95E-07 |
| 2116 | ENSMUSG00000022370 | Mrpl13 | 68537 [Gene Symbol: Mrpl13] [Locus Tag: ] [Chromosome: 15] [Map Location: 15\|15 D | 2.866412607 | 1.23E-05 | 4.61E-05 |
| 9560 | ENSMUSG00000037580 | Gch1 | 14528 [Gene Symbol: Gch1] [Locus Tag: ] [Chromosome: 14] [Map Location: 14 24.6 cM | 2.865444462 | 0.009165 | 0.019249 |
| 7347 | ENSMUSG00000020105 | Lrig3 | 320398 [Gene Symbol: Lrig3] [Locus Tag: ] [Chromosome: 10] [Map Location: 10\|10 D3 | 2.864949652 | 9.13E-05 | 0.000288 |
| 6391 | ENSMUSG00000023004 | Tuba1b | 22143 [Gene Symbol: Tuba1b] [Locus Tag: ] [Chromosome: 15] [Map Location: 15\|15 F | 2.864929945 | 1.57E-06 | 6.87E-06 |
| 15213 | ENSMUSG00000012076 | Brms1l | 52592 [Gene Symbol: Brms1l] [Locus Tag: ] [Chromosome: 12] [Map Location: 12 C1\|1 | 2.864696125 | 1.51E-08 | 9.16E-08 |
| 6693 | ENSMUSG00000026761 | Orc4 | 26428 [Gene Symbol: Orc4] [Locus Tag: ] [Chromosome: 2] [Map Location: 2\|2 C1.1] [D | 2.864493744 | 1.70E-09 | 1.19E-08 |
| 6634 | ENSMUSG00000071855 | Ccdc112 | 240261 [Gene Symbol: Ccdc112] [Locus Tag: ] [Chromosome: 18] [Map Location: 18\|1 | 2.864053636 | 2.24E-05 | 8.03E-05 |
| 3371 | ENSMUSG00000021000 | Ctage5 | 217615 [Gene Symbol: Ctage5] [Locus Tag: ] [Chromosome: 12] [Map Location: 12 C1\|1 | 2.864016007 | 9.75E-09 | 6.11E-08 |
| 15830 | ENSMUSG00000060034 | Ctf2 | 244218 [Gene Symbol: Ctf2] [Locus Tag: ] [Chromosome: 7] [Map Location: 7\|7 F3] [De | 2.863948174 | 0.022518 | 0.042814 |
| 9609 | ENSMUSG00000022009 | Nufip1 | 27275 [Gene Symbol: Nufip1] [Locus Tag: ] [Chromosome: 14] [Map Location: 14\|14 D | 2.863840457 | 4.37E-06 | 1.76E-05 |
| 11790 | ENSMUSG00000001986 | Gria3 | 53623 [Gene Symbol: Gria3] [Locus Tag: ] [Chromosome: X] [Map Location: X A3.3-A4\| | 2.863170185 | 2.24E-09 | 1.53E-08 |
| 311 | ENSMUSG00000072066 | 6720489N17Rik | 211378 [Gene Symbol: 6720489N17Rik] [Locus Tag: ] [Chromosome: 13] [Map Locatio | 2.862024665 | 0.000737 | 0.001966 |
| 16419 | ENSMUSG00000067795 | 4930444P10Rik | 75799 [Gene Symbol: 4930444P10Rik] [Locus Tag: ] [Chromosome: 1] [Map Location: 1 | 2.861990614 | 0.001766 | 0.004354 |
| 16240 | ENSMUSG00000024174 | Pot1b | 72836 [Gene Symbol: Pot1b] [Locus Tag: ] [Chromosome: 17] [Map Location: 17\|17 C] | 2.861340047 | 2.03E-05 | 7.33E-05 |
| 9284 | ENSMUSG00000048285 | Frmd6 | 319710 [Gene Symbol: Frmd6] [Locus Tag: ] [Chromosome: 12] [Map Location: 12\|12 C | 2.86099665 | 5.46E-08 | 3.02E-07 |

| | | | | | | |
|---|---|---|---|---|---|---|
| 12513 | ENSMUSG00000024120 | Lrpprc | 72416 [Gene Symbol: Lrpprc] [Locus Tag: ] [Chromosome: 17] [Map Location: 17\|17 E4 | 2.860679341 | 2.32E-09 | 1.58E-08 |
| 17874 | ENSMUSG00000022864 | D16Ertd472e | 67102 [Gene Symbol: D16Ertd472e] [Locus Tag: ] [Chromosome: 16] [Map Location: 16 | 2.859399714 | 3.86E-06 | 1.57E-05 |
| 11645 | ENSMUSG00000041303 | Gtf3c3 | 98488 [Gene Symbol: Gtf3c3] [Locus Tag: ] [Chromosome: 1] [Map Location: 1\|1 C1.1] | 2.858900174 | 1.85E-07 | 9.42E-07 |
| 12732 | ENSMUSG00000032403 | 2300009A05Rik | 69478 [Gene Symbol: 2300009A05Rik] [Locus Tag: ] [Chromosome: 9] [Map Location: 9 | 2.858831712 | 3.75E-06 | 1.53E-05 |
| 164 | ENSMUSG00000021772 | Nkiras1 | 69721 [Gene Symbol: Nkiras1] [Locus Tag: ] [Chromosome: 14] [Map Location: 14\|14 A | 2.858392038 | 7.72E-08 | 4.16E-07 |
| 18608 | ENSMUSG00000066797 | Zfp648 | 100503355 [Gene Symbol: Zfp648] [Locus Tag: ] [Chromosome: 1] [Map Location: 1\|1 C | 2.857604107 | 0.002585 | 0.006151 |
| 987 | ENSMUSG00000024581 | Napg | 108123 [Gene Symbol: Napg] [Locus Tag: ] [Chromosome: 18] [Map Location: 18\|18 D3 | 2.85721807 | 8.69E-08 | 4.66E-07 |
| 12219 | ENSMUSG00000033460 | Armcx1 | 78248 [Gene Symbol: Armcx1] [Locus Tag: ] [Chromosome: X] [Map Location: X\|X E3] [D | 2.857161854 | 5.43E-08 | 3.00E-07 |
| 2731 | ENSMUSG00000064115 | Cadm2 | 239857 [Gene Symbol: Cadm2] [Locus Tag: ] [Chromosome: 16] [Map Location: 16\|16 C | 2.852254631 | 2.86E-05 | 0.0001 |
| 10266 | ENSMUSG00000018651 | Tada2a | 217031 [Gene Symbol: Tada2a] [Locus Tag: ] [Chromosome: 11] [Map Location: 11\|11 C | 2.852225806 | 4.50E-06 | 1.81E-05 |
| 5204 | ENSMUSG00000047344 | Lancl3 | 236285 [Gene Symbol: Lancl3] [Locus Tag: ] [Chromosome: X] [Map Location: X\|X A1.1 | 2.851850199 | 1.32E-07 | 6.88E-07 |
| 9900 | ENSMUSG00000039908 | Slc26a11 | 268512 [Gene Symbol: Slc26a11] [Locus Tag: ] [Chromosome: 11] [Map Location: 11\|1 | 2.850898165 | 0.012214 | 0.024883 |
| 14239 | ENSMUSG00000003948 | Mmd | 67468 [Gene Symbol: Mmd] [Locus Tag: ] [Chromosome: 11] [Map Location: 11\|11 C] [ | 2.850541845 | 6.34E-10 | 4.70E-09 |
| 17718 | ENSMUSG00000025986 | Slc39a10 | 227059 [Gene Symbol: Slc39a10] [Locus Tag: ] [Chromosome: 1] [Map Location: 1\|1 C1 | 2.84830144 | 3.37E-08 | 1.93E-07 |
| 9078 | ENSMUSG00000029228 | Lnx1 | 16924 [Gene Symbol: Lnx1] [Locus Tag: ] [Chromosome: 5] [Map Location: 5\|5 D] [Desc | 2.847949283 | 4.70E-06 | 1.88E-05 |
| 17791 | ENSMUSG00000070305 | Mpzl3 | 319742 [Gene Symbol: Mpzl3] [Locus Tag: ] [Chromosome: 9] [Map Location: 9 A5.2\|9 | 2.846589024 | 0.014981 | 0.029836 |
| 9644 | ENSMUSG00000022255 | Mtdh | 67154 [Gene Symbol: Mtdh] [Locus Tag: ] [Chromosome: 15] [Map Location: 15 13.98 c | 2.846550059 | 1.81E-09 | 1.25E-08 |
| 10387 | ENSMUSG00000028312 | Smc2 | 14211 [Gene Symbol: Smc2] [Locus Tag: ] [Chromosome: 4] [Map Location: 4 28.31 cM | 2.845912886 | 2.29E-07 | 1.14E-06 |
| 12760 | ENSMUSG00000044150 | A830080D01Rik | 382252 [Gene Symbol: A830080D01Rik] [Locus Tag: ] [Chromosome: X] [Map Location: | 2.845836543 | 3.83E-06 | 1.56E-05 |
| 12338 | ENSMUSG00000022793 | B4galt4 | 56375 [Gene Symbol: B4galt4] [Locus Tag: ] [Chromosome: 16] [Map Location: 16\|16 B | 2.845129935 | 2.06E-08 | 1.23E-07 |
| 9324 | ENSMUSG00000011960 | Ccnt1 | 12455 [Gene Symbol: Ccnt1] [Locus Tag: ] [Chromosome: 15] [Map Location: 15\|15 F2] | 2.845071525 | 2.25E-08 | 1.33E-07 |
| 6915 | ENSMUSG00000024528 | Srfbp1; LOC1026419 | 67222 [Gene Symbol: Srfbp1] [Locus Tag: ] [Chromosome: 18] [Map Location: 18\|18 D1 | 2.844078483 | 4.38E-05 | 0.000148 |
| 18601 | ENSMUSG00000017548 | Suz12 | 52615 [Gene Symbol: Suz12] [Locus Tag: ] [Chromosome: 11] [Map Location: 11 B5\|11 | 2.843749543 | 1.25E-08 | 7.69E-08 |
| 16247 | ENSMUSG00000030060 | Hmces | 232210 [Gene Symbol: Hmces] [Locus Tag: ] [Chromosome: 6] [Map Location: 6\|6 D1] [ | 2.840970445 | 1.88E-06 | 8.10E-06 |
| 11532 | ENSMUSG00000021103 | Mnat1 | 17420 [Gene Symbol: Mnat1] [Locus Tag: ] [Chromosome: 12] [Map Location: 12\|12 C3 | 2.839760322 | 7.03E-06 | 2.74E-05 |
| 18537 | ENSMUSG00000041907 | Gpr45 | 93690 [Gene Symbol: Gpr45] [Locus Tag: ] [Chromosome: 1] [Map Location: 1\|1 C1] [De | 2.839485345 | 6.77E-09 | 4.35E-08 |
| 4574 | ENSMUSG00000002342 | Tmem161a | 234371 [Gene Symbol: Tmem161a] [Locus Tag: ] [Chromosome: 8] [Map Location: 8\|8 | 2.838829763 | 4.86E-06 | 1.94E-05 |
| 2359 | ENSMUSG00000063358 | Mapk1 | 26413 [Gene Symbol: Mapk1] [Locus Tag: ] [Chromosome: 16] [Map Location: 16 A3\|16 | 2.838485885 | 8.97E-10 | 6.51E-09 |
| 8596 | ENSMUSG00000028343 | Erp44 | 76299 [Gene Symbol: Erp44] [Locus Tag: ] [Chromosome: 4] [Map Location: 4\|4 B1] [De | 2.836859738 | 7.51E-08 | 4.06E-07 |
| 2491 | ENSMUSG00000029672 | Fam3c | 27999 [Gene Symbol: Fam3c] [Locus Tag: ] [Chromosome: 6] [Map Location: 6 A3.1\|6 9 | 2.83682523 | 3.26E-09 | 2.19E-08 |
| 2906 | ENSMUSG00000025321 | Itgb8 | 320910 [Gene Symbol: Itgb8] [Locus Tag: ] [Chromosome: 12] [Map Location: 12\|12 F2 | 2.831259769 | 5.07E-09 | 3.32E-08 |
| 14635 | ENSMUSG00000027010 | Slc25a12 | 78830 [Gene Symbol: Slc25a12] [Locus Tag: ] [Chromosome: 2] [Map Location: 2\|2 C2] | 2.827325117 | 3.51E-08 | 2.01E-07 |
| 9401 | ENSMUSG00000022822 | Abcc5 | 27416 [Gene Symbol: Abcc5] [Locus Tag: ] [Chromosome: 16] [Map Location: 16 A3\|16 | 2.826964783 | 8.20E-08 | 4.41E-07 |
| 11276 | ENSMUSG00000076431 | Sox4 | 20677 [Gene Symbol: Sox4] [Locus Tag: ] [Chromosome: 13] [Map Location: 13 12.87 c | 2.825762522 | 0.001843 | 0.004526 |
| 9229 | ENSMUSG00000031278 | Acsl4 | 50790 [Gene Symbol: Acsl4] [Locus Tag: ] [Chromosome: X] [Map Location: X\|X F2] [Des | 2.825721718 | 1.04E-06 | 4.66E-06 |
| 7089 | ENSMUSG00000021728 | Emb | 13723 [Gene Symbol: Emb] [Locus Tag: ] [Chromosome: 13] [Map Location: 13\|13 D2.3 | 2.825487748 | 0.000881 | 0.002316 |
| 18116 | ENSMUSG00000024423 | Impact | 16210 [Gene Symbol: Impact] [Locus Tag: ] [Chromosome: 18] [Map Location: 18\|18 A | 2.824878927 | 4.17E-08 | 2.36E-07 |
| 556 | ENSMUSG00000028809 | Srrm1 | 51796 [Gene Symbol: Srrm1] [Locus Tag: ] [Chromosome: 4] [Map Location: 4\|4 D3] [De | 2.82300648 | 2.20E-05 | 7.90E-05 |
| 18449 | ENSMUSG00000061455 | Stx17 | 67727 [Gene Symbol: Stx17] [Locus Tag: ] [Chromosome: 4] [Map Location: 4\|4 B1] [De | 2.820782205 | 7.90E-08 | 4.26E-07 |
| 10012 | ENSMUSG00000020656 | Grhl1 | 195733 [Gene Symbol: Grhl1] [Locus Tag: ] [Chromosome: 12] [Map Location: 12\|12 A | 2.819668553 | 0.014016 | 0.028129 |
| 12367 | ENSMUSG00000022833 | Ccdc14 | 239839 [Gene Symbol: Ccdc14] [Locus Tag: ] [Chromosome: 16] [Map Location: 16\|16 | 2.819632065 | 7.33E-05 | 0.000236 |
| 4510 | ENSMUSG00000059423 | Zfp933 | 242747 [Gene Symbol: Zfp933] [Locus Tag: ] [Chromosome: 4] [Map Location: 4\|4 E2] [ | 2.819269186 | 3.91E-06 | 1.59E-05 |
| 16853 | ENSMUSG00000004356 | Utp20 | 70683 [Gene Symbol: Utp20] [Locus Tag: ] [Chromosome: 10] [Map Location: 10\|10 C1 | 2.818193834 | 2.66E-08 | 1.55E-07 |
| 80 | ENSMUSG00000031221 | Igbp1 | 18518 [Gene Symbol: Igbp1] [Locus Tag: ] [Chromosome: X] [Map Location: X\|X C3] [De | 2.81480719 | 1.53E-06 | 6.70E-06 |
| 2615 | ENSMUSG00000042390 | Gatad2b | 229542 [Gene Symbol: Gatad2b] [Locus Tag: ] [Chromosome: 3] [Map Location: 3\|3 F1] | 2.814686979 | 1.49E-09 | 1.05E-08 |
| 8039 | ENSMUSG00000045053 | Kcng3 | 225030 [Gene Symbol: Kcng3] [Locus Tag: ] [Chromosome: 17] [Map Location: 17\|17 E | 2.813376013 | 6.54E-05 | 0.000213 |
| 16757 | ENSMUSG00000027709 | Mccc1 | 72039 [Gene Symbol: Mccc1] [Locus Tag: ] [Chromosome: 3] [Map Location: 3\|3 B] [Des | 2.811978317 | 2.75E-06 | 1.15E-05 |
| 2128 | ENSMUSG00000019796 | Lrp11 | 237253 [Gene Symbol: Lrp11] [Locus Tag: ] [Chromosome: 10] [Map Location: 10\|10 A | 2.811887711 | 1.50E-09 | 1.05E-08 |
| 5854 | ENSMUSG00000066760 | Psg16 | 26436 [Gene Symbol: Psg16] [Locus Tag: ] [Chromosome: 7] [Map Location: 7\|7 A2] [De | 2.811347746 | 7.35E-07 | 3.39E-06 |
| 1697 | ENSMUSG00000037997 | Parp11 | 101187 [Gene Symbol: Parp11] [Locus Tag: ] [Chromosome: 6] [Map Location: 6\|6 F3] [ | 2.811013948 | 2.29E-06 | 9.69E-06 |
| 10433 | ENSMUSG00000053289 | Ddx10 | 77591 [Gene Symbol: Ddx10] [Locus Tag: ] [Chromosome: 9] [Map Location: 9\|9 A5.3] [ | 2.810830942 | 1.84E-06 | 7.94E-06 |
| 9337 | ENSMUSG00000050229 | Pigm | 67556 [Gene Symbol: Pigm] [Locus Tag: ] [Chromosome: 1] [Map Location: 1 H3\|1 79.7 | 2.810190613 | 3.11E-06 | 1.28E-05 |
| 9870 | ENSMUSG00000047368 | Abhd17b | 226016 [Gene Symbol: Abhd17b] [Locus Tag: ] [Chromosome: 19] [Map Location: 19\|1 | 2.80907929 | 2.74E-08 | 1.60E-07 |
| 8443 | ENSMUSG00000030232 | Aebp2 | 11569 [Gene Symbol: Aebp2] [Locus Tag: ] [Chromosome: 6] [Map Location: 6\|6 G1] [D | 2.808879268 | 5.02E-08 | 2.79E-07 |
| 6244 | ENSMUSG00000028528 | Dnajc6 | 72685 [Gene Symbol: Dnajc6] [Locus Tag: ] [Chromosome: 4] [Map Location: 4\|4 C6] [D | 2.808566902 | 8.50E-10 | 6.20E-09 |
| 10619 | ENSMUSG00000031148 | Gpkow | 209416 [Gene Symbol: Gpkow] [Locus Tag: ] [Chromosome: X] [Map Location: X A1.1\|X | 2.808543747 | 1.20E-07 | 6.32E-07 |
| 8514 | ENSMUSG00000020728 | Cep112 | 76380 [Gene Symbol: Cep112] [Locus Tag: ] [Chromosome: 11] [Map Location: 11\|11 E | 2.807121615 | 1.66E-08 | 1.00E-07 |
| 14068 | ENSMUSG00000036894 | Rap2b | 74012 [Gene Symbol: Rap2b] [Locus Tag: ] [Chromosome: 3] [Map Location: 3\|3 E1] [De | 2.80430512 | 1.91E-09 | 1.32E-08 |
| 13620 | ENSMUSG00000035310 | Lin54 | 231506 [Gene Symbol: Lin54] [Locus Tag: ] [Chromosome: 5] [Map Location: 5\|5 E4] [D | 2.80426004 | 3.85E-08 | 2.19E-07 |
| 15650 | ENSMUSG00000059182 | Skap2 | 54353 [Gene Symbol: Skap2] [Locus Tag: ] [Chromosome: 6] [Map Location: 6\|6 B3] [De | 2.80308726 | 0.000101 | 0.000317 |
| 6992 | ENSMUSG00000029094 | Afap1 | 70292 [Gene Symbol: Afap1] [Locus Tag: ] [Chromosome: 5] [Map Location: 5\|5 B1] [De | 2.802555734 | 1.13E-09 | 8.03E-09 |
| 14220 | ENSMUSG00000020623 | Map2k6 | 26399 [Gene Symbol: Map2k6] [Locus Tag: ] [Chromosome: 11] [Map Location: 11\|11 E | 2.802010153 | 2.46E-06 | 1.03E-05 |
| 7912 | ENSMUSG00000045671 | Spred2 | 114716 [Gene Symbol: Spred2] [Locus Tag: ] [Chromosome: 11] [Map Location: 11\|11 A | 2.799792805 | 4.06E-08 | 2.30E-07 |
| 3617 | ENSMUSG00000052852 | Reep1 | 52250 [Gene Symbol: Reep1] [Locus Tag: ] [Chromosome: 6] [Map Location: 6 C1\|6 32. | 2.7980583 | 3.11E-09 | 2.09E-08 |
| 16207 | ENSMUSG00000047496 | Rnf152 | 320311 [Gene Symbol: Rnf152] [Locus Tag: ] [Chromosome: 1] [Map Location: 1\|1 E2.1 | 2.795482857 | 1.36E-09 | 9.62E-09 |
| 1368 | ENSMUSG00000021786 | Oxsm | 71147 [Gene Symbol: Oxsm] [Locus Tag: ] [Chromosome: 14] [Map Location: 14\|14 A2] | 2.794395559 | 1.15E-05 | 4.32E-05 |
| 1345 | ENSMUSG00000030869 | Ndufab1 | 70316 [Gene Symbol: Ndufab1] [Locus Tag: ] [Chromosome: 7] [Map Location: 7\|7 F3] [ | 2.792876369 | 4.97E-05 | 0.000166 |
| 10069 | ENSMUSG00000031344 | Gabrq | 57249 [Gene Symbol: Gabrq] [Locus Tag: ] [Chromosome: X] [Map Location: X\|X A6] [De | 2.792396258 | 0.000971 | 0.00253 |
| 17828 | ENSMUSG00000031245 | Hmgn5 | 50887 [Gene Symbol: Hmgn5] [Locus Tag: ] [Chromosome: X] [Map Location: X\|X D] [De | 2.791702886 | 2.85E-05 | 9.99E-05 |
| 1628 | ENSMUSG00000028936 | LOC102642602; Rpl2 | 102642602 [Gene Symbol: LOC102642602] [Locus Tag: ] [Chromosome: 7] [Map Locati | 2.79071309 | 9.25E-09 | 5.83E-08 |
| 8778 | ENSMUSG00000026708 | Cenpl | 70454 [Gene Symbol: Cenpl] [Locus Tag: ] [Chromosome: 1] [Map Location: 1\|1 H1] [De | 2.788091088 | 0.002185 | 0.005275 |
| 7805 | ENSMUSG00000016477 | E2f3 | 13557 [Gene Symbol: E2f3] [Locus Tag: ] [Chromosome: 13] [Map Location: 13\|13 A3.3 | 2.786585626 | 1.04E-08 | 6.51E-08 |
| 15440 | ENSMUSG00000030226 | Lmo3 | 109593 [Gene Symbol: Lmo3] [Locus Tag: ] [Chromosome: 6] [Map Location: 6 G1\|6 69 | 2.786488204 | 1.42E-09 | 1.00E-08 |
| 12517 | ENSMUSG00000019173 | Rab5c | 19345 [Gene Symbol: Rab5c] [Locus Tag: ] [Chromosome: 11] [Map Location: 11 D\|11 6 | 2.785813963 | 5.55E-09 | 3.61E-08 |
| 4070 | ENSMUSG00000090258 | Churc1 | 211151 [Gene Symbol: Churc1] [Locus Tag: ] [Chromosome: 12] [Map Location: 12\|12 | 2.785404744 | 0.000903 | 0.002369 |
| 15316 | ENSMUSG00000079608 | Stard6 | 170461 [Gene Symbol: Stard6] [Locus Tag: ] [Chromosome: 18] [Map Location: 18 E2\|1 | 2.784564179 | 0.005932 | 0.01299 |
| 8259 | ENSMUSG00000005148 | Klf5 | 12224 [Gene Symbol: Klf5] [Locus Tag: ] [Chromosome: 14] [Map Location: 14\|14 E2.2] | 2.783606504 | 2.75E-05 | 9.66E-05 |
| 694 | ENSMUSG00000024754 | Tmem2 | 83921 [Gene Symbol: Tmem2] [Locus Tag: ] [Chromosome: 19] [Map Location: 19\|19 B | 2.782415631 | 1.57E-06 | 6.86E-06 |
| 7201 | ENSMUSG00000030788 | Rnf141 | 67150 [Gene Symbol: Rnf141] [Locus Tag: ] [Chromosome: 7] [Map Location: 7\|7 F2] [D | 2.782380678 | 9.27E-09 | 5.84E-08 |
| 2350 | ENSMUSG00000020015 | Cdk17 | 237459 [Gene Symbol: Cdk17] [Locus Tag: ] [Chromosome: 10] [Map Location: 10\|10 C | 2.782113445 | 2.11E-09 | 1.45E-08 |
| 16218 | ENSMUSG00000061136 | Prpf40a | 56194 [Gene Symbol: Prpf40a] [Locus Tag: ] [Chromosome: 2] [Map Location: 2\|2 C1] [ | 2.781623637 | 1.41E-06 | 6.22E-06 |
| 2784 | ENSMUSG00000027009 | Itga4 | 16401 [Gene Symbol: Itga4] [Locus Tag: ] [Chromosome: 2] [Map Location: 2 C3\|2 47.3 | 2.780067786 | 4.46E-06 | 1.79E-05 |
| 14723 | ENSMUSG00000003198 | Zfp959 | 224893 [Gene Symbol: Zfp959] [Locus Tag: ] [Chromosome: 17] [Map Location: 17\|17 | 2.779890495 | 0.000779 | 0.002069 |
| 2636 | ENSMUSG00000031156 | Slc35a2 | 22232 [Gene Symbol: Slc35a2] [Locus Tag: ] [Chromosome: X] [Map Location: X 3.56 cM | 2.778717691 | 3.63E-05 | 0.000125 |

| | | | | | | |
|---|---|---|---|---|---|---|
| 16522 | ENSMUSG00000025757 | Hspa4l | 18415 [Gene Symbol: Hspa4l] [Locus Tag: ] [Chromosome: 3] [Map Location: 3\|3 B] [De | 2.778178471 | 1.85E-07 | 9.42E-07 |
| 9157 | ENSMUSG00000072946 | Ptgr2 | 77219 [Gene Symbol: Ptgr2] [Locus Tag: ] [Chromosome: 12] [Map Location: 12\|12 D3] | 2.776548378 | 8.78E-06 | 3.36E-05 |
| 3393 | ENSMUSG00000095139 | Pou3f2 | 18992 [Gene Symbol: Pou3f2] [Locus Tag: ] [Chromosome: 4] [Map Location: 4 A3\|4 9. | 2.776356438 | 5.68E-07 | 2.67E-06 |
| 12520 | ENSMUSG00000026153 | Fam135a | 68187 [Gene Symbol: Fam135a] [Locus Tag: ] [Chromosome: 1] [Map Location: 1\|1 A5] | 2.77571829 | 2.76E-09 | 1.87E-08 |
| 18036 | ENSMUSG00000058174 | Gm5148 | 381438 [Gene Symbol: Gm5148] [Locus Tag: ] [Chromosome: 3] [Map Location: 3 B\|3] | 2.774523625 | 0.000436 | 0.001217 |
| 15868 | ENSMUSG00000021180 | Rps6ka5 | 73086 [Gene Symbol: Rps6ka5] [Locus Tag: ] [Chromosome: 12] [Map Location: 12\|12 E | 2.773475083 | 1.64E-08 | 9.90E-08 |
| 18027 | ENSMUSG00000055320 | Tead1 | 21676 [Gene Symbol: Tead1] [Locus Tag: ] [Chromosome: 7] [Map Location: 7\|7 F2] [De | 2.77336647 | 8.44E-08 | 4.53E-07 |
| 7282 | ENSMUSG00000047989 | Ino80c | 225280 [Gene Symbol: Ino80c] [Locus Tag: ] [Chromosome: 18] [Map Location: 18\|18 A | 2.773157708 | 5.23E-07 | 2.47E-06 |
| 6476 | ENSMUSG00000000827 | Tpd52l2 | 66314 [Gene Symbol: Tpd52l2] [Locus Tag: ] [Chromosome: 2] [Map Location: 2\|2 H4] [ | 2.77181634 | 2.01E-07 | 1.01E-06 |
| 11782 | ENSMUSG00000021248 | Tmed10 | 68581 [Gene Symbol: Tmed10] [Locus Tag: ] [Chromosome: 12] [Map Location: 12\|12 D | 2.769985866 | 5.45E-08 | 3.02E-07 |
| 368 | ENSMUSG00000034544 | Rsrc1 | 66880 [Gene Symbol: Rsrc1] [Locus Tag: ] [Chromosome: 3] [Map Location: 3\|3 E2] [De | 2.769982262 | 2.82E-08 | 1.64E-07 |
| 2340 | ENSMUSG00000020120 | Plek | 56193 [Gene Symbol: Plek] [Locus Tag: ] [Chromosome: 11] [Map Location: 11 A2\|11 9. | 2.769539042 | 3.07E-08 | 1.77E-07 |
| 6535 | ENSMUSG00000035270 | Impg2 | 224224 [Gene Symbol: Impg2] [Locus Tag: ] [Chromosome: 16] [Map Location: 16\|16 C | 2.769196589 | 0.003636 | 0.008362 |
| 6466 | ENSMUSG00000055371 | Stam2 | 56324 [Gene Symbol: Stam2] [Locus Tag: ] [Chromosome: 2] [Map Location: 2\|2 C1.1] [ | 2.769123243 | 1.87E-08 | 1.12E-07 |
| 3924 | ENSMUSG00000021395 | Spin1 | 20729 [Gene Symbol: Spin1] [Locus Tag: ] [Chromosome: 13] [Map Location: 13 A5\|13 | 2.768276515 | 2.06E-09 | 1.42E-08 |
| 9602 | ENSMUSG00000009291 | Pttg1ip | 108705 [Gene Symbol: Pttg1ip] [Locus Tag: ] [Chromosome: 10] [Map Location: 10\|10 | 2.765815407 | 1.93E-06 | 8.31E-06 |
| 14292 | ENSMUSG00000029245 | Epha5 | 13839 [Gene Symbol: Epha5] [Locus Tag: ] [Chromosome: 5] [Map Location: 5 E1\|5 43.( | 2.765297413 | 7.00E-08 | 3.80E-07 |
| 2632 | ENSMUSG00000020634 | Ubxn2a | 217379 [Gene Symbol: Ubxn2a] [Locus Tag: ] [Chromosome: 12] [Map Location: 12\|12 | 2.765197977 | 4.10E-08 | 2.32E-07 |
| 3732 | ENSMUSG00000042487 | Leo1 | 235497 [Gene Symbol: Leo1] [Locus Tag: ] [Chromosome: 9] [Map Location: 9\|9 D] [Des | 2.76419982 | 6.18E-08 | 3.39E-07 |
| 8713 | ENSMUSG00000035770 | Dync1li2 | 234663 [Gene Symbol: Dync1li2] [Locus Tag: ] [Chromosome: 8] [Map Location: 8 D3\|8 | 2.763599376 | 4.57E-09 | 3.01E-08 |
| 7022 | ENSMUSG00000026914 | Psmd14 | 59029 [Gene Symbol: Psmd14] [Locus Tag: ] [Chromosome: 2] [Map Location: 2\|2 C1.3 | 2.763506617 | 3.52E-08 | 2.01E-07 |
| 13281 | ENSMUSG00000025647 | Shisa5 | 66940 [Gene Symbol: Shisa5] [Locus Tag: ] [Chromosome: 9] [Map Location: 9\|9 F2] [De | 2.761707955 | 2.90E-06 | 1.21E-05 |
| 10450 | ENSMUSG00000022120 | Rnf219 | 72486 [Gene Symbol: Rnf219] [Locus Tag: ] [Chromosome: 14] [Map Location: 14\|14 E | 2.760227123 | 1.40E-08 | 8.57E-08 |
| 9899 | ENSMUSG00000025086 | Trub1 | 72133 [Gene Symbol: Trub1] [Locus Tag: ] [Chromosome: 19] [Map Location: 19\|19 D2 | 2.759745854 | 2.31E-07 | 1.15E-06 |
| 734 | ENSMUSG00000019913 | Sim1 | 20464 [Gene Symbol: Sim1] [Locus Tag: ] [Chromosome: 10] [Map Location: 10 B3\|10 2 | 2.75888854 | 0.020759 | 0.039828 |
| 5715 | ENSMUSG00000026643 | Nmt2 | 18108 [Gene Symbol: Nmt2] [Locus Tag: ] [Chromosome: 2] [Map Location: 2\|2 A1] [De | 2.758548535 | 5.64E-09 | 3.67E-08 |
| 2920 | ENSMUSG00000033610 | Pank1 | 75735 [Gene Symbol: Pank1] [Locus Tag: ] [Chromosome: 19] [Map Location: 19\|19 G1 | 2.757823845 | 1.65E-07 | 8.48E-07 |
| 3775 | ENSMUSG00000045410 | Akr1e1 | 56043 [Gene Symbol: Akr1e1] [Locus Tag: ] [Chromosome: 13] [Map Location: 13\|13 A | 2.757685027 | 0.00023 | 0.000677 |
| 133 | ENSMUSG00000021965 | Ska3 | 219114 [Gene Symbol: Ska3] [Locus Tag: ] [Chromosome: 14] [Map Location: 14\|14 C3 | 2.75727293 | 0.000166 | 0.000501 |
| 17208 | ENSMUSG00000034868 | Myl12b | 67938 [Gene Symbol: Myl12b] [Locus Tag: ] [Chromosome: 17] [Map Location: 17\|17 E | 2.756719332 | 1.59E-08 | 9.60E-08 |
| 17388 | ENSMUSG00000026019 | Wdr12 | 57750 [Gene Symbol: Wdr12] [Locus Tag: ] [Chromosome: 1] [Map Location: 1\|1 C1-C2 | 2.756364104 | 1.18E-06 | 5.26E-06 |
| 6106 | ENSMUSG00000033499 | Larp4b | 217980 [Gene Symbol: Larp4b] [Locus Tag: ] [Chromosome: 13] [Map Location: 13 A1\| | 2.753206694 | 1.61E-09 | 1.12E-08 |
| 6370 | ENSMUSG00000033623 | Pcgf3 | 69587 [Gene Symbol: Pcgf3] [Locus Tag: ] [Chromosome: 5] [Map Location: 5\|5 F] [Desc | 2.75069422 | 6.27E-09 | 4.05E-08 |
| 17501 | ENSMUSG00000015932 | Dstn | 56431 [Gene Symbol: Dstn] [Locus Tag: ] [Chromosome: 2] [Map Location: 2 70.89 cM\| | 2.750673038 | 1.05E-05 | 3.96E-05 |
| 11078 | ENSMUSG00000040648 | Ppip5k2 | 227399 [Gene Symbol: Ppip5k2] [Locus Tag: ] [Chromosome: 1] [Map Location: 1\|1 D] [ | 2.750315863 | 3.39E-09 | 2.27E-08 |
| 18474 | ENSMUSG00000020687 | Cdc27 | 217232 [Gene Symbol: Cdc27] [Locus Tag: ] [Chromosome: 11] [Map Location: 11 E1\|1 | 2.750011454 | 7.20E-09 | 4.60E-08 |
| 16772 | ENSMUSG00000053600 | Zfp472 | 224691 [Gene Symbol: Zfp472] [Locus Tag: ] [Chromosome: 17] [Map Location: 17\|17 I | 2.749407226 | 7.20E-05 | 0.000232 |
| 12433 | ENSMUSG00000025911 | Adhfe1 | 76187 [Gene Symbol: Adhfe1] [Locus Tag: ] [Chromosome: 1] [Map Location: 1\|1 A2] [D | 2.748033745 | 0.000496 | 0.001369 |
| 4554 | ENSMUSG00000028007 | Snx7 | 76561 [Gene Symbol: Snx7] [Locus Tag: ] [Chromosome: 3] [Map Location: 3\|3 G2] [Des | 2.74746516 | 0.000346 | 0.000985 |
| 13035 | ENSMUSG00000028300 | 3110043O21Rik | 73205 [Gene Symbol: 3110043O21Rik] [Locus Tag: ] [Chromosome: 4] [Map Location: 4 | 2.747405922 | 1.60E-08 | 9.68E-08 |
| 16319 | ENSMUSG00000027357 | Crls1 | 66586 [Gene Symbol: Crls1] [Locus Tag: ] [Chromosome: 2] [Map Location: 2\|2 F3] [Des | 2.746706963 | 1.72E-07 | 8.77E-07 |
| 6773 | ENSMUSG00000074968 | Ano3 | 228432 [Gene Symbol: Ano3] [Locus Tag: ] [Chromosome: 2] [Map Location: 2 E3\|2] [De | 2.744690639 | 0.001036 | 0.002686 |
| 9579 | ENSMUSG00000017561 | Crlf3 | 54394 [Gene Symbol: Crlf3] [Locus Tag: ] [Chromosome: 11] [Map Location: 11 B5\|11 4 | 2.742348116 | 5.27E-05 | 0.000174 |
| 15364 | ENSMUSG00000060121 | Gemin2 | 66603 [Gene Symbol: Gemin2] [Locus Tag: ] [Chromosome: 12] [Map Location: 12\|12 C | 2.742194786 | 0.001666 | 0.004132 |
| 9561 | ENSMUSG00000053012 | Krcc1 | 57896 [Gene Symbol: Krcc1] [Locus Tag: ] [Chromosome: 6] [Map Location: 6\|6 C1] [De | 2.742164986 | 2.55E-06 | 1.07E-05 |
| 4699 | ENSMUSG00000067578 | Cbln4 | 228942 [Gene Symbol: Cbln4] [Locus Tag: ] [Chromosome: 2] [Map Location: 2\|2 H3] [D | 2.742068564 | 9.49E-07 | 4.29E-06 |
| 11994 | ENSMUSG00000022400 | Rbx1; Gm9840 | 56438 [Gene Symbol: Rbx1] [Locus Tag: ] [Chromosome: 15] [Map Location: 15\|15 E1] | 2.741962697 | 1.96E-07 | 9.92E-07 |
| 10782 | ENSMUSG00000060534 | Dcc | 13176 [Gene Symbol: Dcc] [Locus Tag: ] [Chromosome: 18] [Map Location: 18 E2\|18 45 | 2.741594221 | 3.66E-09 | 2.44E-08 |
| 11717 | ENSMUSG00000039753 | Fbxl5 | 242960 [Gene Symbol: Fbxl5] [Locus Tag: ] [Chromosome: 5] [Map Location: 5\|5 B3] [D | 2.73990671 | 5.54E-09 | 3.61E-08 |
| 17937 | ENSMUSG00000078302 | Foxd1 | 15229 [Gene Symbol: Foxd1] [Locus Tag: ] [Chromosome: 13] [Map Location: 13 D1\|13 | 2.738124472 | 0.006028 | 0.013181 |
| 8495 | ENSMUSG00000038943 | Prc1 | 233406 [Gene Symbol: Prc1] [Locus Tag: ] [Chromosome: 7] [Map Location: 7 D3\|7 45.( | 2.737617535 | 1.07E-05 | 4.05E-05 |
| 634 | ENSMUSG00000021758 | Ddx4 | 13206 [Gene Symbol: Ddx4] [Locus Tag: ] [Chromosome: 13] [Map Location: 13 D2.2\|1 | 2.736842206 | 0.000968 | 0.002524 |
| 7008 | ENSMUSG00000027671 | Actl6a | 56456 [Gene Symbol: Actl6a] [Locus Tag: ] [Chromosome: 3] [Map Location: 3\|3 A3] [De | 2.736222976 | 4.75E-06 | 1.90E-05 |
| 17705 | ENSMUSG00000021147 | Wdr37 | 207615 [Gene Symbol: Wdr37] [Locus Tag: ] [Chromosome: 13] [Map Location: 13\|13 A | 2.735798783 | 2.95E-08 | 1.71E-07 |
| 9408 | ENSMUSG00000042256 | Ptchd4 | 627626 [Gene Symbol: Ptchd4] [Locus Tag: ] [Chromosome: 17] [Map Location: 17\|17 | 2.733845771 | 1.84E-06 | 7.94E-06 |
| 5132 | ENSMUSG00000020657 | Dnajc27 | 217378 [Gene Symbol: Dnajc27] [Locus Tag: ] [Chromosome: 12] [Map Location: 12\|12 | 2.733747104 | 5.55E-06 | 2.20E-05 |
| 13075 | ENSMUSG00000008976 | Gabpa | 14390 [Gene Symbol: Gabpa] [Locus Tag: ] [Chromosome: 16] [Map Location: 16 C3-4\| | 2.732901775 | 2.64E-08 | 1.55E-07 |
| 8246 | ENSMUSG00000021796 | Bmpr1a | 12166 [Gene Symbol: Bmpr1a] [Locus Tag: ] [Chromosome: 14] [Map Location: 14 B\|14 | 2.731778519 | 5.25E-09 | 3.43E-08 |
| 8047 | ENSMUSG00000020283 | Pex13 | 72129 [Gene Symbol: Pex13] [Locus Tag: ] [Chromosome: 11] [Map Location: 11\|11 A3 | 2.728700656 | 1.29E-06 | 5.71E-06 |
| 7064 | ENSMUSG00000020572 | Nampt | 59027 [Gene Symbol: Nampt] [Locus Tag: ] [Chromosome: 12] [Map Location: 12\|12 B1 | 2.727901382 | 3.31E-08 | 1.90E-07 |
| 16581 | ENSMUSG00000025907 | Rb1cc1 | 12421 [Gene Symbol: Rb1cc1] [Locus Tag: ] [Chromosome: 1] [Map Location: 1\|1 A2] [D | 2.727787116 | 4.31E-09 | 2.84E-08 |
| 12881 | ENSMUSG00000003746 | Man1a | 17155 [Gene Symbol: Man1a] [Locus Tag: ] [Chromosome: 10] [Map Location: 10\|10 B3 | 2.72775291 | 8.44E-08 | 4.53E-07 |
| 5958 | ENSMUSG00000073293 | Nudt10 | 102954 [Gene Symbol: Nudt10] [Locus Tag: ] [Chromosome: X] [Map Location: X\|X A1. | 2.726959372 | 3.56E-07 | 1.72E-06 |
| 11484 | ENSMUSG00000020326 | Ccng1 | 12450 [Gene Symbol: Ccng1] [Locus Tag: ] [Chromosome: 11] [Map Location: 11\|11 B1 | 2.726930217 | 3.11E-07 | 1.52E-06 |
| 17965 | ENSMUSG00000066880 | Zfp617 | 170938 [Gene Symbol: Zfp617] [Locus Tag: ] [Chromosome: 8] [Map Location: 8\|8 B3.3 | 2.726508445 | 3.53E-06 | 1.45E-05 |
| 15915 | ENSMUSG00000039047 | Pigk | 329777 [Gene Symbol: Pigk] [Locus Tag: ] [Chromosome: 3] [Map Location: 3\|3 H4] [De | 2.726064475 | 3.16E-09 | 2.12E-08 |
| 3605 | ENSMUSG00000040372 | Gpr63 | 81006 [Gene Symbol: Gpr63] [Locus Tag: ] [Chromosome: 4] [Map Location: 4\|4 A3] [De | 2.725594351 | 3.51E-07 | 1.70E-06 |
| 17783 | ENSMUSG00000026889 | Rbm18 | 67889 [Gene Symbol: Rbm18] [Locus Tag: ] [Chromosome: 2] [Map Location: 2\|2 B] [De | 2.72525123 | 3.06E-07 | 1.49E-06 |
| 68 | ENSMUSG00000025758 | Plk4 | 20873 [Gene Symbol: Plk4] [Locus Tag: ] [Chromosome: 3] [Map Location: 3\|3 B] [Descr | 2.725193644 | 2.93E-06 | 1.22E-05 |
| 18534 | ENSMUSG00000026098 | Pms1 | 227099 [Gene Symbol: Pms1] [Locus Tag: ] [Chromosome: 1] [Map Location: 1\|1 C1.1] | 2.724968189 | 2.52E-05 | 8.94E-05 |
| 6013 | ENSMUSG00000021669 | Col4a3bp | 68018 [Gene Symbol: Col4a3bp] [Locus Tag: ] [Chromosome: 13] [Map Location: 13\|13 | 2.723681326 | 9.18E-09 | 5.79E-08 |
| 5555 | ENSMUSG00000025154 | Arhgap19 | 71085 [Gene Symbol: Arhgap19] [Locus Tag: ] [Chromosome: 19] [Map Location: 19\|19 | 2.72335726 | 0.024934 | 0.04695 |
| 10211 | ENSMUSG00000001815 | Evx2 | 14029 [Gene Symbol: Evx2] [Locus Tag: ] [Chromosome: 2] [Map Location: 2 C3\|2 44.1 | 2.722697064 | 0.014784 | 0.029514 |
| 17171 | ENSMUSG00000028710 | Atpaf1 | 230649 [Gene Symbol: Atpaf1] [Locus Tag: ] [Chromosome: 4] [Map Location: 4\|4 D1] [ | 2.722321296 | 9.72E-06 | 3.70E-05 |
| 14546 | ENSMUSG00000028655 | Mfsd2a | 76574 [Gene Symbol: Mfsd2a] [Locus Tag: ] [Chromosome: 4] [Map Location: 4\|4 D1] [D | 2.721855907 | 0.000937 | 0.00245 |
| 14624 | ENSMUSG00000038619 | Ensa | 56205 [Gene Symbol: Ensa] [Locus Tag: ] [Chromosome: 3] [Map Location: 3\|3 F2.1] [De | 2.72184203 | 4.31E-08 | 2.44E-07 |
| 3200 | ENSMUSG00000063884 | Ptcd3 | 69956 [Gene Symbol: Ptcd3] [Locus Tag: ] [Chromosome: 6] [Map Location: 6\|6 C1] [De | 2.720336048 | 6.46E-08 | 3.53E-07 |
| 6254 | ENSMUSG00000021719 | Rgs7bp | 52882 [Gene Symbol: Rgs7bp] [Locus Tag: ] [Chromosome: 13] [Map Location: 13 D1\|1 | 2.720264779 | 2.62E-05 | 9.25E-05 |
| 15634 | ENSMUSG00000041915 | Ammecr1l | 225339 [Gene Symbol: Ammecr1l] [Locus Tag: ] [Chromosome: 18] [Map Location: 18\| | 2.718593249 | 2.60E-06 | 1.09E-05 |
| 15362 | ENSMUSG00000047904 | Sstr2 | 20606 [Gene Symbol: Sstr2] [Locus Tag: ] [Chromosome: 11] [Map Location: 11 E2\|11 7 | 2.718348078 | 0.003992 | 0.009091 |
| 2315 | ENSMUSG00000004018 | Fancl | 67030 [Gene Symbol: Fancl] [Locus Tag: ] [Chromosome: 11] [Map Location: 11\|11 A2- | 2.717625157 | 3.69E-05 | 0.000126 |
| 7117 | ENSMUSG00000027597 | Ahcy | 269378 [Gene Symbol: Ahcy] [Locus Tag: ] [Chromosome: 2] [Map Location: 2 H1\|2 76. | 2.716742259 | 0.013489 | 0.027174 |
| 13065 | ENSMUSG00000020386 | Sar1b | 66397 [Gene Symbol: Sar1b] [Locus Tag: ] [Chromosome: 11] [Map Location: 11\|11 B1. | 2.716639548 | 3.24E-05 | 0.000112 |
| 4139 | ENSMUSG00000024234 | Mtpap | 67440 [Gene Symbol: Mtpap] [Locus Tag: ] [Chromosome: 18] [Map Location: 18\|18 A1 | 2.716635933 | 1.14E-06 | 5.09E-06 |

| | | | | | | |
|---|---|---|---|---|---|---|
| 7822 | ENSMUSG00000025277 | Abhd6 | 66082 [Gene Symbol: Abhd6] [Locus Tag: ] [Chromosome: 14] [Map Location: 14\|14 A1 | 2.713191071 | 1.43E-05 | 5.28E-05 |
| 18071 | ENSMUSG00000022808 | Snx4 | 69150 [Gene Symbol: Snx4] [Locus Tag: ] [Chromosome: 16] [Map Location: 16\|16 B3] | 2.712810723 | 5.26E-08 | 2.92E-07 |
| 11277 | ENSMUSG00000091931 | AK010878 | 100233175 [Gene Symbol: AK010878] [Locus Tag: ] [Chromosome: 12] [Map Location: | 2.711857934 | 4.19E-06 | 1.69E-05 |
| 12339 | ENSMUSG00000027954 | Efna1 | 13636 [Gene Symbol: Efna1] [Locus Tag: ] [Chromosome: 3] [Map Location: 3 F1\|3 39.0 | 2.708365437 | 0.006219 | 0.013556 |
| 13373 | ENSMUSG00000045294 | Insig1 | 231070 [Gene Symbol: Insig1] [Locus Tag: ] [Chromosome: 5] [Map Location: 5\|5 B1] [D | 2.707572214 | 6.93E-08 | 3.77E-07 |
| 12186 | ENSMUSG00000004151 | Etv1 | 14009 [Gene Symbol: Etv1] [Locus Tag: ] [Chromosome: 12] [Map Location: 12 A3-B1\|1 | 2.706568227 | 5.93E-08 | 3.26E-07 |
| 13430 | ENSMUSG00000030924 | 2610020H08Rik | 434234 [Gene Symbol: 2610020H08Rik] [Locus Tag: ] [Chromosome: 7] [Map Location: | 2.702986283 | 7.78E-05 | 0.000249 |
| 7302 | ENSMUSG00000020462 | Cfap36 | 216618 [Gene Symbol: Cfap36] [Locus Tag: ] [Chromosome: 11] [Map Location: 11\|11 A | 2.702472582 | 1.15E-05 | 4.33E-05 |
| 6500 | ENSMUSG00000021131 | Erh | 13877 [Gene Symbol: Erh] [Locus Tag: ] [Chromosome: 12] [Map Location: 12\|12 D1] [D | 2.702124172 | 0.000273 | 0.00079 |
| 10883 | ENSMUSG00000036112 | Metap2 | 56307 [Gene Symbol: Metap2] [Locus Tag: ] [Chromosome: 10] [Map Location: 10\|10 C | 2.701349216 | 1.66E-07 | 8.53E-07 |
| 17690 | ENSMUSG00000046593 | Tmem215 | 320500 [Gene Symbol: Tmem215] [Locus Tag: ] [Chromosome: 4] [Map Location: 4\|4 A | 2.699496521 | 0.006093 | 0.013306 |
| 3550 | ENSMUSG00000045636 | Mtus1 | 102103 [Gene Symbol: Mtus1] [Locus Tag: ] [Chromosome: 8] [Map Location: 8\|8 A4] [D | 2.698179969 | 8.11E-07 | 3.71E-06 |
| 4113 | ENSMUSG00000042258 | Isl1 | 16392 [Gene Symbol: Isl1] [Locus Tag: ] [Chromosome: 13] [Map Location: 13\|13 D2.3] | 2.696990321 | 5.02E-05 | 0.000167 |
| 16796 | ENSMUSG00000058317 | Ube2e2 | 218793 [Gene Symbol: Ube2e2] [Locus Tag: ] [Chromosome: 14] [Map Location: 14\|14 | 2.696070882 | 1.60E-08 | 9.66E-08 |
| 7065 | ENSMUSG00000031347 | Cetn2 | 26370 [Gene Symbol: Cetn2] [Locus Tag: ] [Chromosome: X] [Map Location: X 37.29 cM | 2.695457368 | 7.92E-05 | 0.000253 |
| 7995 | ENSMUSG00000029211 | Gabra4 | 14397 [Gene Symbol: Gabra4] [Locus Tag: ] [Chromosome: 5] [Map Location: 5\|5 C3.2] | 2.695202981 | 5.55E-05 | 0.000183 |
| 17573 | ENSMUSG00000044018 | Mrpl50 | 28028 [Gene Symbol: Mrpl50] [Locus Tag: ] [Chromosome: 4] [Map Location: 4 B1\|4 26 | 2.69494677 | 2.28E-07 | 1.14E-06 |
| 18100 | ENSMUSG00000056537 | Rlim | 19820 [Gene Symbol: Rlim] [Locus Tag: ] [Chromosome: X] [Map Location: X 46.42 cM\| | 2.693933575 | 6.55E-08 | 3.58E-07 |
| 5891 | ENSMUSG00000005054 | Cstb | 13014 [Gene Symbol: Cstb] [Locus Tag: ] [Chromosome: 10] [Map Location: 10 C1\|10 3 | 2.69361708 | 0.001237 | 0.003162 |
| 308 | ENSMUSG00000031762 | Mt2 | 17750 [Gene Symbol: Mt2] [Locus Tag: ] [Chromosome: 8] [Map Location: 8 C5\|8 46.32 | 2.692451145 | 4.23E-05 | 0.000143 |
| 15212 | ENSMUSG00000027746 | Ufm1 | 67890 [Gene Symbol: Ufm1] [Locus Tag: ] [Chromosome: 3] [Map Location: 3\|3 C] [Desc | 2.692262145 | 2.57E-07 | 1.28E-06 |
| 14834 | ENSMUSG00000021548 | Ccnh | 66671 [Gene Symbol: Ccnh] [Locus Tag: ] [Chromosome: 13] [Map Location: 13\|13 C3] | 2.68675668 | 2.10E-08 | 1.25E-07 |
| 15624 | ENSMUSG00000028398 | Tmem261 | 66928 [Gene Symbol: Tmem261] [Locus Tag: ] [Chromosome: 4] [Map Location: 4\|4 C3 | 2.685974702 | 0.000249 | 0.000728 |
| 7707 | ENSMUSG00000023066 | Rttn | 246102 [Gene Symbol: Rttn] [Locus Tag: ] [Chromosome: 18] [Map Location: 18\|18 E4] | 2.684933067 | 1.61E-05 | 5.92E-05 |
| 8235 | ENSMUSG00000009828 | Ick | 56542 [Gene Symbol: Ick] [Locus Tag: ] [Chromosome: 9] [Map Location: 9\|9 E1] [Descri | 2.683802827 | 6.32E-09 | 4.08E-08 |
| 15347 | ENSMUSG00000062991 | Nrg1 | 211323 [Gene Symbol: Nrg1] [Locus Tag: ] [Chromosome: 8] [Map Location: 8\|8 A3] [De | 2.682644959 | 2.38E-06 | 1.00E-05 |
| 13719 | ENSMUSG00000000266 | Mid2 | 23947 [Gene Symbol: Mid2] [Locus Tag: ] [Chromosome: X] [Map Location: X F1\|X 61.3 | 2.682432676 | 1.22E-06 | 5.41E-06 |
| 5160 | ENSMUSG00000038187 | Btbd10 | 68815 [Gene Symbol: Btbd10] [Locus Tag: ] [Chromosome: 7] [Map Location: 7\|7 F2] [D | 2.681803281 | 2.61E-05 | 9.21E-05 |
| 10023 | ENSMUSG00000048728 | Zfp454 | 237758 [Gene Symbol: Zfp454] [Locus Tag: ] [Chromosome: 11] [Map Location: 11\|11 I | 2.681113831 | 0.000464 | 0.001288 |
| 17392 | ENSMUSG00000034761 | Map4k5 | 399510 [Gene Symbol: Map4k5] [Locus Tag: ] [Chromosome: 12] [Map Location: 12\|12 | 2.678767844 | 2.82E-08 | 1.64E-07 |
| 11575 | ENSMUSG00000021390 | Ogn | 18295 [Gene Symbol: Ogn] [Locus Tag: ] [Chromosome: 13] [Map Location: 13\|13 A5] [I | 2.678465094 | 0.004098 | 0.0093 |
| 13734 | ENSMUSG00000070883 | Ccdc173 | 75051 [Gene Symbol: Ccdc173] [Locus Tag: ] [Chromosome: 2] [Map Location: 2\|2 C2] | 2.675876796 | 0.011433 | 0.023483 |
| 12670 | ENSMUSG00000022781 | Pak2 | 224105 [Gene Symbol: Pak2] [Locus Tag: ] [Chromosome: 16] [Map Location: 16 B2\|16 | 2.674878069 | 3.24E-08 | 1.87E-07 |
| 13572 | ENSMUSG00000063297 | Luzp2 | 233271 [Gene Symbol: Luzp2] [Locus Tag: ] [Chromosome: 7] [Map Location: 7 B5\|7 33 | 2.674296296 | 5.20E-06 | 2.07E-05 |
| 16759 | ENSMUSG00000031917 | Nip7 | 66164 [Gene Symbol: Nip7] [Locus Tag: ] [Chromosome: 8] [Map Location: 8\|8 D2] [Des | 2.674122566 | 1.59E-06 | 6.95E-06 |
| 3354 | ENSMUSG00000049583 | Grm5 | 108071 [Gene Symbol: Grm5] [Locus Tag: ] [Chromosome: 7] [Map Location: 7\|7 E1] [D | 2.674041315 | 8.74E-07 | 3.98E-06 |
| 7503 | ENSMUSG00000001376 | Vps50 | 73288 [Gene Symbol: Vps50] [Locus Tag: ] [Chromosome: 6] [Map Location: 6\|6 A1] [De | 2.673874387 | 9.66E-08 | 5.15E-07 |
| 15508 | ENSMUSG00000046818 | Ddit4l | 73284 [Gene Symbol: Ddit4l] [Locus Tag: ] [Chromosome: 3] [Map Location: 3\|3 G3] [D | 2.67360005 | 0.000888 | 0.002333 |
| 17896 | ENSMUSG00000031303 | Map3k15 | 270672 [Gene Symbol: Map3k15] [Locus Tag: ] [Chromosome: X] [Map Location: X\|X F4 | 2.673083247 | 1.35E-07 | 7.04E-07 |
| 17300 | ENSMUSG00000071415 | LOC100862455; Rpl2 | 100862455 [Gene Symbol: LOC100862455] [Locus Tag: ] [Chromosome: 17] [Map Loca | 2.671444911 | 6.69E-05 | 0.000217 |
| 2386 | ENSMUSG00000047959 | Kcna3 | 16491 [Gene Symbol: Kcna3] [Locus Tag: ] [Chromosome: 3] [Map Location: 3 F2.3\|3 46 | 2.671056316 | 2.22E-06 | 9.42E-06 |
| 7481 | ENSMUSG00000020255 | D10Wsu102e | 28109 [Gene Symbol: D10Wsu102e] [Locus Tag: ] [Chromosome: 10] [Map Location: 10 | 2.669562848 | 2.52E-07 | 1.25E-06 |
| 16649 | ENSMUSG00000067916 | Gm13139 | 666532 [Gene Symbol: Gm13139] [Locus Tag: ] [Chromosome: 4] [Map Location: 4\|4 E | 2.669217738 | 0.017842 | 0.034806 |
| 9300 | ENSMUSG00000018189 | Uchl5 | 56207 [Gene Symbol: Uchl5] [Locus Tag: ] [Chromosome: 1] [Map Location: 1 F\|1 62.54 | 2.66784669 | 8.32E-07 | 3.81E-06 |
| 10546 | ENSMUSG00000020514 | Mrpl22 | 216767 [Gene Symbol: Mrpl22] [Locus Tag: ] [Chromosome: 11] [Map Location: 11\|11 | 2.667470633 | 0.000387 | 0.001093 |
| 10977 | ENSMUSG00000041762 | Gpr155 | 68526 [Gene Symbol: Gpr155] [Locus Tag: ] [Chromosome: 2] [Map Location: 2\|2 C3] [I | 2.666689906 | 4.17E-07 | 2.00E-06 |
| 16490 | ENSMUSG00000015749 | Anp32e | 66471 [Gene Symbol: Anp32e] [Locus Tag: ] [Chromosome: 3] [Map Location: 3\|3 F2.1] | 2.665920704 | 4.43E-08 | 2.50E-07 |
| 3280 | ENSMUSG00000006498 | Ptbp1 | 19205 [Gene Symbol: Ptbp1] [Locus Tag: ] [Chromosome: 10] [Map Location: 10 C1\|10 | 2.665652569 | 8.27E-08 | 4.45E-07 |
| 14636 | ENSMUSG00000055660 | Mettl4 | 76781 [Gene Symbol: Mettl4] [Locus Tag: ] [Chromosome: 17] [Map Location: 17\|17 E5 | 2.665108192 | 7.01E-05 | 0.000226 |
| 18600 | ENSMUSG00000001158 | Snrnp27 | 66618 [Gene Symbol: Snrnp27] [Locus Tag: ] [Chromosome: 6] [Map Location: 6\|6 D1] | 2.6650315 | 1.84E-05 | 6.69E-05 |
| 16440 | ENSMUSG00000024095 | Hnrnpll | 72692 [Gene Symbol: Hnrnpll] [Locus Tag: ] [Chromosome: 17] [Map Location: 17\|17 E | 2.664986642 | 3.75E-08 | 2.14E-07 |
| 9950 | ENSMUSG00000028552 | Eps15 | 13858 [Gene Symbol: Eps15] [Locus Tag: ] [Chromosome: 4] [Map Location: 4\|4 C7] [De | 2.66459233 | 4.88E-09 | 3.20E-08 |
| 6837 | ENSMUSG00000029670 | Ing3 | 71777 [Gene Symbol: Ing3] [Locus Tag: ] [Chromosome: 6] [Map Location: 6\|6 A3] [Desc | 2.66339069 | 3.39E-07 | 1.65E-06 |
| 736 | ENSMUSG00000031343 | Gabra3 | 14396 [Gene Symbol: Gabra3] [Locus Tag: ] [Chromosome: X] [Map Location: X A7.3\|X 3 | 2.662876553 | 5.62E-08 | 3.10E-07 |
| 13328 | ENSMUSG00000042396 | Rbm7 | 67010 [Gene Symbol: Rbm7] [Locus Tag: ] [Chromosome: 9] [Map Location: 9\|9 A5.3] [I | 2.662763643 | 6.09E-06 | 2.40E-05 |
| 3116 | ENSMUSG00000054942 | Fam73a | 215708 [Gene Symbol: Fam73a] [Locus Tag: ] [Chromosome: 3] [Map Location: 3\|3 H3] | 2.662548118 | 1.85E-08 | 1.11E-07 |
| 5997 | ENSMUSG00000043300 | B3galnt1 | 26879 [Gene Symbol: B3galnt1] [Locus Tag: ] [Chromosome: 3] [Map Location: 3\|3 E1] | 2.661900051 | 1.32E-07 | 6.91E-07 |
| 2344 | ENSMUSG00000039242 | B3galnt2 | 97884 [Gene Symbol: B3galnt2] [Locus Tag: ] [Chromosome: 13] [Map Location: 13\|13 | 2.661706251 | 4.31E-06 | 1.74E-05 |
| 16184 | ENSMUSG00000037643 | Prkci | 18759 [Gene Symbol: Prkci] [Locus Tag: ] [Chromosome: 3] [Map Location: 3 A3\|3 14.6 | 2.660257506 | 1.09E-08 | 6.76E-08 |
| 335 | ENSMUSG00000034211 | Mrps17 | 66258 [Gene Symbol: Mrps17] [Locus Tag: ] [Chromosome: 5] [Map Location: 5\|5 F] [De | 2.660051368 | 2.34E-05 | 8.34E-05 |
| 5898 | ENSMUSG00000079477 | Rab7 | 19349 [Gene Symbol: Rab7] [Locus Tag: ] [Chromosome: 6] [Map Location: 6 D1\|6 39.1 | 2.659810647 | 1.49E-08 | 9.05E-08 |
| 7741 | ENSMUSG00000048787 | Dcun1d3 | 233805 [Gene Symbol: Dcun1d3] [Locus Tag: ] [Chromosome: 7] [Map Location: 7\|7 F2 | 2.659247745 | 3.48E-07 | 1.69E-06 |
| 12431 | ENSMUSG00000026399 | Cd55 | 13136 [Gene Symbol: Cd55] [Locus Tag: ] [Chromosome: 1] [Map Location: 1 E4\|1 56.8 | 2.659182782 | 0.006522 | 0.014148 |
| 8660 | ENSMUSG00000028453 | Fancg | 60534 [Gene Symbol: Fancg] [Locus Tag: ] [Chromosome: 4] [Map Location: 4\|4 B1] [De | 2.658858057 | 0.001314 | 0.003339 |
| 5104 | ENSMUSG00000079036 | Alkbh1; Nrp | 211064 [Gene Symbol: Alkbh1] [Locus Tag: ] [Chromosome: 12] [Map Location: 12\|12 I | 2.6562188 | 9.34E-07 | 4.23E-06 |
| 5914 | ENSMUSG00000079614 | Seh1l | 72124 [Gene Symbol: Seh1l] [Locus Tag: ] [Chromosome: 18] [Map Location: 18\|18 E1] | 2.655881736 | 1.18E-07 | 6.20E-07 |
| 17251 | ENSMUSG00000030256 | Bhlhe41 | 79362 [Gene Symbol: Bhlhe41] [Locus Tag: ] [Chromosome: 6] [Map Location: 6 77.7 cM | 2.655385514 | 0.001438 | 0.003623 |
| 16377 | ENSMUSG00000004455 | Gm5601; Ppp1cc | 434233 [Gene Symbol: Gm5601] [Locus Tag: ] [Chromosome: 7] [Map Location: 7\|7 F2 | 2.654317252 | 5.04E-05 | 0.000168 |
| 2465 | ENSMUSG00000022812 | Gsk3b | 56637 [Gene Symbol: Gsk3b] [Locus Tag: ] [Chromosome: 16] [Map Location: 16\|16 B4 | 2.654310425 | 1.10E-08 | 6.86E-08 |
| 13024 | ENSMUSG00000036840 | Siah1a | 20437 [Gene Symbol: Siah1a] [Locus Tag: ] [Chromosome: 8] [Map Location: 8 C3\|8 42. | 2.653270354 | 4.01E-08 | 2.28E-07 |
| 15997 | ENSMUSG00000050711 | Scg2 | 20254 [Gene Symbol: Scg2] [Locus Tag: ] [Chromosome: 1] [Map Location: 1 C4\|1 40.8 | 2.652773561 | 9.11E-08 | 4.86E-07 |
| 216 | ENSMUSG00000020719 | Ddx5 | 13207 [Gene Symbol: Ddx5] [Locus Tag: ] [Chromosome: 11] [Map Location: 11 70.01 c | 2.651892236 | 1.55E-05 | 5.72E-05 |
| 12234 | ENSMUSG00000023963 | Cyp39a1 | 56050 [Gene Symbol: Cyp39a1] [Locus Tag: ] [Chromosome: 17] [Map Location: 17\|17 | 2.650798254 | 0.001085 | 0.002803 |
| 3888 | ENSMUSG00000067194 | Eif1ax | 66235 [Gene Symbol: Eif1ax] [Locus Tag: ] [Chromosome: X] [Map Location: X\|X F4] [De | 2.650340963 | 1.04E-08 | 6.47E-08 |
| 7675 | ENSMUSG00000036061 | Smug1 | 71726 [Gene Symbol: Smug1] [Locus Tag: ] [Chromosome: 15] [Map Location: 15\|15 F3 | 2.650198452 | 5.18E-06 | 2.06E-05 |
| 15567 | ENSMUSG00000061024 | Rrs1 | 59014 [Gene Symbol: Rrs1] [Locus Tag: ] [Chromosome: 1] [Map Location: 1 A2\|1 2.08 | 2.648983743 | 1.27E-05 | 4.74E-05 |
| 707 | ENSMUSG00000058589 | Anks1b | 77531 [Gene Symbol: Anks1b] [Locus Tag: ] [Chromosome: 10] [Map Location: 10\|10 C | 2.648414417 | 4.82E-08 | 2.69E-07 |
| 1523 | ENSMUSG00000002413 | Braf | 109880 [Gene Symbol: Braf] [Locus Tag: ] [Chromosome: 6] [Map Location: 6 B1\|6 18.4 | 2.648229929 | 6.65E-09 | 4.28E-08 |
| 4824 | ENSMUSG00000061315 | Naca | 17938 [Gene Symbol: Naca] [Locus Tag: ] [Chromosome: 10] [Map Location: 10\|10 D2-I | 2.648017068 | 9.90E-06 | 3.76E-05 |
| 4270 | ENSMUSG00000038160 | Atg5 | 11793 [Gene Symbol: Atg5] [Locus Tag: ] [Chromosome: 10] [Map Location: 10 B2\|10 2 | 2.647405957 | 2.46E-07 | 1.22E-06 |
| 1273 | ENSMUSG00000024077 | Strn | 268980 [Gene Symbol: Strn] [Locus Tag: ] [Chromosome: 17] [Map Location: 17\|17 E3] | 2.647060131 | 8.14E-09 | 5.17E-08 |
| 579 | ENSMUSG00000025937 | Lactb2 | 212442 [Gene Symbol: Lactb2] [Locus Tag: ] [Chromosome: 1] [Map Location: 1\|1 A3] [I | 2.647047344 | 0.002364 | 0.005667 |
| 14980 | ENSMUSG00000067367 | Lyar | 17089 [Gene Symbol: Lyar] [Locus Tag: ] [Chromosome: 5] [Map Location: 5 B3\|5 20.29 | 2.646868054 | 1.04E-05 | 3.95E-05 |
| 4454 | ENSMUSG00000024511 | Rab27b | 80718 [Gene Symbol: Rab27b] [Locus Tag: ] [Chromosome: 18] [Map Location: 18 E2\|1 | 2.646693021 | 0.000184 | 0.000552 |

| | | | | | | |
|---|---|---|---|---|---|---|
| 531 | ENSMUSG00000041084 | Ostc | 66357 [Gene Symbol: Ostc] [Locus Tag: ] [Chromosome: 3] [Map Location: 3\|3 H1] [Des | 2.646485587 | 1.85E-05 | 6.71E-05 |
| 16345 | ENSMUSG00000042505 | Sdhaf3 | 71238 [Gene Symbol: Sdhaf3] [Locus Tag: ] [Chromosome: 6] [Map Location: 6\|6 A1] [D | 2.644055745 | 0.000186 | 0.000556 |
| 2042 | ENSMUSG00000000171 | Sdhd | 66925 [Gene Symbol: Sdhd] [Locus Tag: ] [Chromosome: 9] [Map Location: 9\|9 B] [Desc | 2.642624552 | 8.67E-06 | 3.33E-05 |
| 7635 | ENSMUSG00000037325 | Bbs7 | 71492 [Gene Symbol: Bbs7] [Locus Tag: ] [Chromosome: 3] [Map Location: 3\|3 B] [Desc | 2.642167324 | 0.000379 | 0.001073 |
| 3826 | ENSMUSG00000049148 | Plcxd3 | 239318 [Gene Symbol: Plcxd3] [Locus Tag: ] [Chromosome: 15] [Map Location: 15\|15 A | 2.641540709 | 6.61E-07 | 3.07E-06 |
| 6612 | ENSMUSG00000034187 | Nsf | 18195 [Gene Symbol: Nsf] [Locus Tag: ] [Chromosome: 11] [Map Location: 11 E1\|11 67. | 2.640594009 | 0.000166 | 0.000501 |
| 12246 | ENSMUSG00000026096 | Osgepl1 | 72085 [Gene Symbol: Osgepl1] [Locus Tag: ] [Chromosome: 1] [Map Location: 1 C1.1\|1 | 2.640144421 | 7.71E-05 | 0.000247 |
| 14197 | ENSMUSG00000000416 | Cttnbp2 | 30785 [Gene Symbol: Cttnbp2] [Locus Tag: ] [Chromosome: 6] [Map Location: 6\|6 A2] [ | 2.64001462 | 4.95E-08 | 2.76E-07 |
| 84 | ENSMUSG00000022637 | Cblb | 208650 [Gene Symbol: Cblb] [Locus Tag: ] [Chromosome: 16] [Map Location: 16\|16 B5] | 2.639664016 | 3.00E-07 | 1.47E-06 |
| 12701 | ENSMUSG00000031105 | Slc25a14 | 20523 [Gene Symbol: Slc25a14] [Locus Tag: ] [Chromosome: X] [Map Location: X A4\|X | 2.639000335 | 3.09E-06 | 1.28E-05 |
| 16290 | ENSMUSG00000029063 | Nadk | 192185 [Gene Symbol: Nadk] [Locus Tag: ] [Chromosome: 4] [Map Location: 4\|4 E2] [De | 2.63769838 | 2.59E-08 | 1.52E-07 |
| 14116 | ENSMUSG00000040738 | Ints8 | 72656 [Gene Symbol: Ints8] [Locus Tag: ] [Chromosome: 4] [Map Location: 4\|4 A1] [Des | 2.636916813 | 2.37E-07 | 1.18E-06 |
| 12358 | ENSMUSG00000035173 | Ccdc186 | 213993 [Gene Symbol: Ccdc186] [Locus Tag: ] [Chromosome: 19] [Map Location: 19\|19 | 2.636280266 | 4.68E-07 | 2.23E-06 |
| 17532 | ENSMUSG00000032050 | Rdx | 19684 [Gene Symbol: Rdx] [Locus Tag: ] [Chromosome: 9] [Map Location: 9\|9 A5.3] [De | 2.636058997 | 2.36E-08 | 1.39E-07 |
| 7177 | ENSMUSG00000041649 | Klf8 | 245671 [Gene Symbol: Klf8] [Locus Tag: ] [Chromosome: X] [Map Location: X\|X F3] [Des | 2.635386258 | 6.55E-08 | 3.58E-07 |
| 14336 | ENSMUSG00000015671 | Psma2 | 19166 [Gene Symbol: Psma2] [Locus Tag: ] [Chromosome: 13] [Map Location: 13\|13 A1 | 2.634565067 | 1.17E-05 | 4.41E-05 |
| 1009 | ENSMUSG00000026228 | Htr2b | 15559 [Gene Symbol: Htr2b] [Locus Tag: ] [Chromosome: 1] [Map Location: 1\|1 C5] [De | 2.633910908 | 0.013994 | 0.028096 |
| 15274 | ENSMUSG00000028527 | Ak4 | 11639 [Gene Symbol: Ak4] [Locus Tag: ] [Chromosome: 4] [Map Location: 4 C6\|4 46.84 | 2.633065729 | 2.76E-05 | 9.69E-05 |
| 6408 | ENSMUSG00000021532 | Fastkd3 | 69577 [Gene Symbol: Fastkd3] [Locus Tag: ] [Chromosome: 13] [Map Location: 13\|13 B | 2.632668626 | 1.18E-05 | 4.44E-05 |
| 1449 | ENSMUSG00000038482 | Tfdp1 | 21781 [Gene Symbol: Tfdp1] [Locus Tag: ] [Chromosome: 8] [Map Location: 8\|8 A1-A3] | 2.631920788 | 6.98E-08 | 3.79E-07 |
| 17891 | ENSMUSG00000067928 | Zfp760 | 240034 [Gene Symbol: Zfp760] [Locus Tag: ] [Chromosome: 17] [Map Location: 17\|17 A | 2.631331247 | 1.57E-07 | 8.10E-07 |
| 10075 | ENSMUSG00000001847 | Rac1 | 19353 [Gene Symbol: Rac1] [Locus Tag: ] [Chromosome: 5] [Map Location: 5 G2\|5 82.2 | 2.631330651 | 2.35E-08 | 1.39E-07 |
| 9537 | ENSMUSG00000078652 | Psme3 | 19192 [Gene Symbol: Psme3] [Locus Tag: ] [Chromosome: 11] [Map Location: 11 D\|11 | 2.631011553 | 6.26E-08 | 3.43E-07 |
| 11617 | ENSMUSG00000013736 | Trnt1 | 70047 [Gene Symbol: Trnt1] [Locus Tag: ] [Chromosome: 6] [Map Location: 6\|6 E2] [Des | 2.630549541 | 6.57E-07 | 3.05E-06 |
| 15284 | ENSMUSG00000031605 | Klhl2 | 77113 [Gene Symbol: Klhl2] [Locus Tag: ] [Chromosome: 8] [Map Location: 8\|8 B3.1] [D | 2.630227683 | 4.83E-09 | 3.17E-08 |
| 3560 | ENSMUSG00000036104 | Rab3gap1 | 226407 [Gene Symbol: Rab3gap1] [Locus Tag: ] [Chromosome: 1] [Map Location: 1\|1 E | 2.628951333 | 1.37E-07 | 7.11E-07 |
| 9980 | ENSMUSG00000035681 | Kcnc2 | 268345 [Gene Symbol: Kcnc2] [Locus Tag: ] [Chromosome: 10] [Map Location: 10 D2\|1 | 2.628366022 | 2.05E-07 | 1.03E-06 |
| 8455 | ENSMUSG00000060510 | Zfp266 | 77519 [Gene Symbol: Zfp266] [Locus Tag: ] [Chromosome: 9] [Map Location: 9\|9 A3] [D | 2.62797796 | 1.22E-06 | 5.43E-06 |
| 15144 | ENSMUSG00000038010 | Ccdc138 | 76138 [Gene Symbol: Ccdc138] [Locus Tag: ] [Chromosome: 10] [Map Location: 10\|10 | 2.626986292 | 4.84E-06 | 1.93E-05 |
| 17634 | ENSMUSG00000028800 | Hdac1 | 433759 [Gene Symbol: Hdac1] [Locus Tag: ] [Chromosome: 4] [Map Location: 4 D2.2\|4 | 2.626974475 | 0.000278 | 0.000806 |
| 14669 | ENSMUSG00000021400 | Wrnip1 | 78903 [Gene Symbol: Wrnip1] [Locus Tag: ] [Chromosome: 13] [Map Location: 13\|13 A | 2.626966484 | 4.48E-08 | 2.53E-07 |
| 8093 | ENSMUSG00000025809 | Itgb1 | 16412 [Gene Symbol: Itgb1] [Locus Tag: ] [Chromosome: 8] [Map Location: 8\|8 E2] [Des | 2.626339189 | 3.77E-08 | 2.15E-07 |
| 4518 | ENSMUSG00000053347 | Zfp943 | 74670 [Gene Symbol: Zfp943] [Locus Tag: ] [Chromosome: 17] [Map Location: 17\|17 A | 2.625143644 | 2.52E-06 | 1.06E-05 |
| 16143 | ENSMUSG00000067279 | Ppp1r3c | 53412 [Gene Symbol: Ppp1r3c] [Locus Tag: ] [Chromosome: 19] [Map Location: 19\|19 | 2.624235494 | 3.80E-05 | 0.00013 |
| 18418 | ENSMUSG00000030092 | Cntn6 | 53870 [Gene Symbol: Cntn6] [Locus Tag: ] [Chromosome: 6] [Map Location: 6\|6 E2] [De | 2.623161951 | 0.019319 | 0.037351 |
| 7327 | ENSMUSG00000021224 | Numb | 18222 [Gene Symbol: Numb] [Locus Tag: ] [Chromosome: 12] [Map Location: 12\|12 D3 | 2.622287721 | 2.66E-07 | 1.32E-06 |
| 14619 | ENSMUSG00000084957 | Bbip1 | 100503572 [Gene Symbol: Bbip1] [Locus Tag: ] [Chromosome: 19] [Map Location: 19\|1 | 2.621020292 | 2.83E-07 | 1.39E-06 |
| 6157 | ENSMUSG00000059474 | Mbtd1 | 103537 [Gene Symbol: Mbtd1] [Locus Tag: ] [Chromosome: 11] [Map Location: 11\|11 D | 2.617358228 | 5.89E-07 | 2.76E-06 |
| 2842 | ENSMUSG00000008683 | Rps15a | 267019 [Gene Symbol: Rps15a] [Locus Tag: ] [Chromosome: 7] [Map Location: 7\|7 F2] [ | 2.616950763 | 2.83E-08 | 1.64E-07 |
| 6161 | ENSMUSG00000035133 | Arhgap5 | 11855 [Gene Symbol: Arhgap5] [Locus Tag: ] [Chromosome: 12] [Map Location: 12 C1\| | 2.616143039 | 3.22E-08 | 1.85E-07 |
| 16212 | ENSMUSG00000005362 | Crbn | 58799 [Gene Symbol: Crbn] [Locus Tag: ] [Chromosome: 6] [Map Location: 6\|6 E2] [Des | 2.61459773 | 2.67E-07 | 1.32E-06 |
| 17278 | ENSMUSG00000038570 | Saxo2 | 330577 [Gene Symbol: Saxo2] [Locus Tag: ] [Chromosome: 7] [Map Location: 7\|7 D3] [D | 2.614560333 | 0.005847 | 0.012828 |
| 13987 | ENSMUSG00000033628 | Pik3c3 | 225326 [Gene Symbol: Pik3c3] [Locus Tag: ] [Chromosome: 18] [Map Location: 18\|18 E | 2.613751592 | 6.13E-07 | 2.86E-06 |
| 11522 | ENSMUSG00000051354 | Samd3 | 268288 [Gene Symbol: Samd3] [Locus Tag: ] [Chromosome: 10] [Map Location: 10\|10 A | 2.61329691 | 0.019392 | 0.03748 |
| 2396 | ENSMUSG00000023147 | Wrb | 71446 [Gene Symbol: Wrb] [Locus Tag: ] [Chromosome: 16] [Map Location: 16\|16 C4] [ | 2.612976981 | 1.50E-07 | 7.75E-07 |
| 13655 | ENSMUSG00000020064 | Herc4 | 67345 [Gene Symbol: Herc4] [Locus Tag: ] [Chromosome: 10] [Map Location: 10\|10 B4] | 2.610342888 | 9.23E-06 | 3.52E-05 |
| 3641 | ENSMUSG00000020463 | Smek2 | 104570 [Gene Symbol: Smek2] [Locus Tag: ] [Chromosome: 11] [Map Location: 11\|11 A | 2.608967337 | 5.25E-08 | 2.91E-07 |
| 3205 | ENSMUSG00000075054 | Yae1d1 | 67008 [Gene Symbol: Yae1d1] [Locus Tag: ] [Chromosome: 13] [Map Location: 13\|13 A | 2.608616907 | 0.000177 | 0.000532 |
| 12115 | ENSMUSG00000057469 | E2f6 | 50496 [Gene Symbol: E2f6] [Locus Tag: ] [Chromosome: 12] [Map Location: 12 A1.1\|12 | 2.60820232 | 8.23E-06 | 3.17E-05 |
| 2687 | ENSMUSG00000037395 | Rcor3 | 214742 [Gene Symbol: Rcor3] [Locus Tag: ] [Chromosome: 1] [Map Location: 1\|1 H6] [D | 2.606307318 | 8.00E-08 | 4.31E-07 |
| 15879 | ENSMUSG00000074166 | AW146154 | 101835 [Gene Symbol: AW146154] [Locus Tag: ] [Chromosome: 7] [Map Location: 7\|7 | 2.606244706 | 0.001476 | 0.003707 |
| 1375 | ENSMUSG00000021411 | Pxdc1 | 66895 [Gene Symbol: Pxdc1] [Locus Tag: ] [Chromosome: 13] [Map Location: 13\|13 A4 | 2.605636329 | 6.89E-05 | 0.000223 |
| 700 | ENSMUSG00000024993 | Fam45a | 67894 [Gene Symbol: Fam45a] [Locus Tag: ] [Chromosome: 19] [Map Location: 19\|19 D | 2.605367535 | 5.33E-05 | 0.000176 |
| 10722 | ENSMUSG00000013878 | Rnf170 | 77733 [Gene Symbol: Rnf170] [Locus Tag: ] [Chromosome: 8] [Map Location: 8\|8] [Desc | 2.602592248 | 1.90E-05 | 6.88E-05 |
| 13025 | ENSMUSG00000030671 | Pde3b | 18576 [Gene Symbol: Pde3b] [Locus Tag: ] [Chromosome: 7] [Map Location: 7 F1\|7 59. | 2.601768776 | 6.04E-07 | 2.82E-06 |
| 17332 | ENSMUSG00000022325 | Pop1 | 67724 [Gene Symbol: Pop1] [Locus Tag: ] [Chromosome: 15] [Map Location: 15\|15 B3. | 2.601447746 | 1.18E-05 | 4.42E-05 |
| 656 | ENSMUSG00000090272 | Mndal | 100040462 [Gene Symbol: Mndal] [Locus Tag: ] [Chromosome: 1] [Map Location: 1 H3\| | 2.600424023 | 0.002351 | 0.005636 |
| 4476 | ENSMUSG00000037075 | Rnf139 | 75841 [Gene Symbol: Rnf139] [Locus Tag: ] [Chromosome: 15] [Map Location: 15\|15 D | 2.598547774 | 2.75E-08 | 1.61E-07 |
| 15969 | ENSMUSG00000047879 | Usp14 | 59025 [Gene Symbol: Usp14] [Locus Tag: ] [Chromosome: 18] [Map Location: 18 A1\|18 | 2.598329871 | 1.84E-07 | 9.34E-07 |
| 12794 | ENSMUSG00000033849 | B3galt2 | 26878 [Gene Symbol: B3galt2] [Locus Tag: ] [Chromosome: 1] [Map Location: 1\|1 F] [De | 2.597540471 | 3.44E-08 | 1.97E-07 |
| 2617 | ENSMUSG00000055184 | Fam72a | 108900 [Gene Symbol: Fam72a] [Locus Tag: ] [Chromosome: 1] [Map Location: 1\|1 E4] | 2.595283589 | 0.007478 | 0.016023 |
| 17866 | ENSMUSG00000032397 | Tipin | 66131 [Gene Symbol: Tipin] [Locus Tag: ] [Chromosome: 9] [Map Location: 9 34.6 cM\|9 | 2.595088545 | 6.86E-07 | 3.18E-06 |
| 10106 | ENSMUSG00000050796 | B3galt6 | 117592 [Gene Symbol: B3galt6] [Locus Tag: ] [Chromosome: 4] [Map Location: 4\|4 E2] | 2.594985828 | 6.57E-06 | 2.57E-05 |
| 16250 | ENSMUSG00000070345 | Hsf5 | 327992 [Gene Symbol: Hsf5] [Locus Tag: ] [Chromosome: 11] [Map Location: 11\|11 C] [ | 2.594712592 | 0.015086 | 0.030007 |
| 3331 | ENSMUSG00000079555 | Haus3 | 231123 [Gene Symbol: Haus3] [Locus Tag: ] [Chromosome: 5] [Map Location: 5 B2\|5 17 | 2.59283261 | 0.000103 | 0.000322 |
| 12660 | ENSMUSG00000022313 | Utp23 | 78581 [Gene Symbol: Utp23] [Locus Tag: ] [Chromosome: 15] [Map Location: 15\|15 D1 | 2.591513101 | 0.001099 | 0.002837 |
| 956 | ENSMUSG00000049327 | Setd8 | 67956 [Gene Symbol: Setd8] [Locus Tag: ] [Chromosome: 5] [Map Location: 5\|5 F] [Desc | 2.588207218 | 9.46E-07 | 4.28E-06 |
| 2057 | ENSMUSG00000038298 | Pdzk1 | 59020 [Gene Symbol: Pdzk1] [Locus Tag: ] [Chromosome: 3] [Map Location: 3\|3 F2.1] [D | 2.587957457 | 1.46E-06 | 6.43E-06 |
| 5886 | ENSMUSG00000022876 | Samsn1 | 67742 [Gene Symbol: Samsn1] [Locus Tag: ] [Chromosome: 16] [Map Location: 16\|16 C | 2.587143129 | 0.004511 | 0.010149 |
| 782 | ENSMUSG00000033063 | Cntnap3 | 238680 [Gene Symbol: Cntnap3] [Locus Tag: ] [Chromosome: 13] [Map Location: 13\|13 | 2.586541797 | 1.46E-05 | 5.40E-05 |
| 5447 | ENSMUSG00000031158 | Timm17b | 21855 [Gene Symbol: Timm17b] [Locus Tag: ] [Chromosome: X] [Map Location: X A1.1\| | 2.584815985 | 3.02E-05 | 0.000105 |
| 3255 | ENSMUSG00000024241 | Sos1 | 20662 [Gene Symbol: Sos1] [Locus Tag: ] [Chromosome: 17] [Map Location: 17 E3\|17 5 | 2.584472752 | 1.57E-08 | 9.50E-08 |
| 10094 | ENSMUSG00000062937 | Mtap | 66902 [Gene Symbol: Mtap] [Locus Tag: ] [Chromosome: 4] [Map Location: 4\|4 C4] [Des | 2.584118795 | 0.000331 | 0.000946 |
| 18567 | ENSMUSG00000015937 | H2afy | 26914 [Gene Symbol: H2afy] [Locus Tag: ] [Chromosome: 13] [Map Location: 13\|13 B2] | 2.584053763 | 2.23E-06 | 9.47E-06 |
| 13731 | ENSMUSG00000050199 | Lgr4 | 107515 [Gene Symbol: Lgr4] [Locus Tag: ] [Chromosome: 2] [Map Location: 2\|2 E3] [Des | 2.583222617 | 9.04E-07 | 4.11E-06 |
| 18066 | ENSMUSG00000029131 | Dnajb6 | 23950 [Gene Symbol: Dnajb6] [Locus Tag: ] [Chromosome: 5] [Map Location: 5\|5 B1] [D | 2.581520302 | 2.32E-06 | 9.80E-06 |
| 16307 | ENSMUSG00000017716 | Birc5 | 11799 [Gene Symbol: Birc5] [Locus Tag: ] [Chromosome: 11] [Map Location: 11\|11 E2] | 2.581025382 | 0.01302 | 0.026305 |
| 14821 | ENSMUSG00000004394 | Tmed4 | 103694 [Gene Symbol: Tmed4] [Locus Tag: ] [Chromosome: 11] [Map Location: 11\|11 A | 2.580648106 | 0.000149 | 0.000453 |
| 7756 | ENSMUSG00000044712 | Slc38a6 | 625098 [Gene Symbol: Slc38a6] [Locus Tag: ] [Chromosome: 12] [Map Location: 12\|12 | 2.580165454 | 8.86E-05 | 0.00028 |
| 13775 | ENSMUSG00000033105 | Lss | 16987 [Gene Symbol: Lss] [Locus Tag: ] [Chromosome: 10] [Map Location: 10 C1\|10 39. | 2.579727502 | 4.57E-08 | 2.57E-07 |
| 7009 | ENSMUSG00000024063 | Lbh | 77889 [Gene Symbol: Lbh] [Locus Tag: ] [Chromosome: 17] [Map Location: 17\|17 E2] [D | 2.579660077 | 1.13E-07 | 5.95E-07 |
| 14191 | ENSMUSG00000025645 | Ccdc51 | 66658 [Gene Symbol: Ccdc51] [Locus Tag: ] [Chromosome: 9] [Map Location: 9\|9 F2] [D | 2.579140896 | 7.65E-05 | 0.000246 |
| 11279 | ENSMUSG00000057455 | Rit2 | 19762 [Gene Symbol: Rit2] [Locus Tag: ] [Chromosome: 18] [Map Location: 18\|18 B1] [ | 2.579013414 | 0.002703 | 0.006402 |
| 14870 | ENSMUSG00000031781 | Ciapin1 | 109006 [Gene Symbol: Ciapin1] [Locus Tag: ] [Chromosome: 8] [Map Location: 8\|8 C5] | 2.576396104 | 7.58E-07 | 3.49E-06 |

| | | | | | | |
|---|---|---|---|---|---|---|
| 15865 | ENSMUSG00000035530 | Eif1 | 20918 [Gene Symbol: Eif1] [Locus Tag: ] [Chromosome: 11] [Map Location: 11\|11 D] [D | 2.575670702 | 2.55E-07 | 1.26E-06 |
| 13928 | ENSMUSG00000039701 | Usp53 | 99526 [Gene Symbol: Usp53] [Locus Tag: ] [Chromosome: 3] [Map Location: 3\|3 G3] [D | 2.575575197 | 8.05E-05 | 0.000257 |
| 5901 | ENSMUSG00000003617 | Cp | 12870 [Gene Symbol: Cp] [Locus Tag: ] [Chromosome: 3] [Map Location: 3\|3 D] [Descrip | 2.575337057 | 6.71E-05 | 0.000218 |
| 15411 | ENSMUSG00000032410 | Xrn1 | 24127 [Gene Symbol: Xrn1] [Locus Tag: ] [Chromosome: 9] [Map Location: 9\|9 E4] [Des | 2.57476229 | 5.83E-08 | 3.20E-07 |
| 14114 | ENSMUSG00000026374 | Tsn | 22099 [Gene Symbol: Tsn] [Locus Tag: ] [Chromosome: 1] [Map Location: 1\|1 E2.3] [Des | 2.574637369 | 7.31E-07 | 3.37E-06 |
| 6498 | ENSMUSG00000039985 | Fam60a; LOC102636 | 56306 [Gene Symbol: Fam60a] [Locus Tag: ] [Chromosome: 6] [Map Location: 6\|6 G3] [ | 2.574438065 | 0.003602 | 0.008288 |
| 2825 | ENSMUSG00000022201 | Zfr | 22763 [Gene Symbol: Zfr] [Locus Tag: ] [Chromosome: 15] [Map Location: 15\|15 A] [Des | 2.574311533 | 8.51E-06 | 3.27E-05 |
| 4283 | ENSMUSG00000027035 | Cers6 | 241447 [Gene Symbol: Cers6] [Locus Tag: ] [Chromosome: 2] [Map Location: 2\|2 C2] [D | 2.573522302 | 4.96E-07 | 2.36E-06 |
| 3769 | ENSMUSG00000079083 | Jrkl | 77532 [Gene Symbol: Jrkl] [Locus Tag: ] [Chromosome: 9] [Map Location: 9\|9] [Descript | 2.573049811 | 8.34E-06 | 3.21E-05 |
| 11116 | ENSMUSG00000036167 | Pphln1 | 223828 [Gene Symbol: Pphln1] [Locus Tag: ] [Chromosome: 15] [Map Location: 15\|15 | 2.572110644 | 5.07E-07 | 2.40E-06 |
| 12393 | ENSMUSG00000025995 | Wdr75 | 73674 [Gene Symbol: Wdr75] [Locus Tag: ] [Chromosome: 1] [Map Location: 1\|1 C1] [D | 2.572108889 | 1.69E-06 | 7.34E-06 |
| 8570 | ENSMUSG00000025893 | Kbtbd3 | 69149 [Gene Symbol: Kbtbd3] [Locus Tag: ] [Chromosome: 9] [Map Location: 9\|9 A1] [D | 2.571634082 | 0.000103 | 0.000322 |
| 1507 | ENSMUSG00000020021 | Fgd6 | 13998 [Gene Symbol: Fgd6] [Locus Tag: ] [Chromosome: 10] [Map Location: 10\|10 C3] | 2.570842328 | 7.90E-07 | 3.62E-06 |
| 12281 | ENSMUSG00000019710 | Mrpl24 | 67707 [Gene Symbol: Mrpl24] [Locus Tag: ] [Chromosome: 3] [Map Location: 3\|3 F1] [D | 2.570515224 | 8.54E-06 | 3.28E-05 |
| 7962 | ENSMUSG00000054836 | Elp6 | 72341 [Gene Symbol: Elp6] [Locus Tag: ] [Chromosome: 9] [Map Location: 9\|9 F2] [Desc | 2.569780208 | 2.03E-05 | 7.31E-05 |
| 17531 | ENSMUSG00000046688 | Tifa | 211550 [Gene Symbol: Tifa] [Locus Tag: ] [Chromosome: 3] [Map Location: 3\|3 G2] [Des | 2.569060703 | 4.20E-06 | 1.70E-05 |
| 7130 | ENSMUSG00000025597 | Klhl4 | 237010 [Gene Symbol: Klhl4] [Locus Tag: ] [Chromosome: X] [Map Location: X\|X E1] [De | 2.568297172 | 0.000626 | 0.001694 |
| 7888 | ENSMUSG00000038732 | Mboat1 | 218121 [Gene Symbol: Mboat1] [Locus Tag: ] [Chromosome: 13] [Map Location: 13\|13 | 2.567914863 | 0.000531 | 0.001457 |
| 58 | ENSMUSG00000037455 | Slc18b1 | 76306 [Gene Symbol: Slc18b1] [Locus Tag: ] [Chromosome: 10] [Map Location: 10\|10 A | 2.567701754 | 4.37E-06 | 1.76E-05 |
| 17177 | ENSMUSG00000016200 | Syt14 | 329324 [Gene Symbol: Syt14] [Locus Tag: ] [Chromosome: 1] [Map Location: 1\|1 H6] [D | 2.567619019 | 1.17E-06 | 5.23E-06 |
| 12163 | ENSMUSG00000031095 | Cul4b | 72584 [Gene Symbol: Cul4b] [Locus Tag: ] [Chromosome: X] [Map Location: X\|X A2] [De | 2.566733271 | 5.22E-07 | 2.47E-06 |
| 7269 | ENSMUSG00000049717 | Lig4 | 319583 [Gene Symbol: Lig4] [Locus Tag: ] [Chromosome: 8] [Map Location: 8\|8 A1.1] [D | 2.566517294 | 2.46E-07 | 1.22E-06 |
| 2976 | ENSMUSG00000059920 | 4930453N24Rik | 67609 [Gene Symbol: 4930453N24Rik] [Locus Tag: ] [Chromosome: 16] [Map Location: | 2.566153464 | 7.88E-07 | 3.61E-06 |
| 17582 | ENSMUSG00000032469 | Dbr1 | 83703 [Gene Symbol: Dbr1] [Locus Tag: ] [Chromosome: 9] [Map Location: 9\|9 F1] [Des | 2.565959705 | 1.64E-06 | 7.14E-06 |
| 1045 | ENSMUSG00000074238 | Ap1ar | 211556 [Gene Symbol: Ap1ar] [Locus Tag: ] [Chromosome: 3] [Map Location: 3\|3 G2] [D | 2.56564479 | 3.82E-06 | 1.55E-05 |
| 1004 | ENSMUSG00000027668 | Mfn1 | 67414 [Gene Symbol: Mfn1] [Locus Tag: ] [Chromosome: 3] [Map Location: 3 15.75 cM\| | 2.565491967 | 4.26E-05 | 0.000144 |
| 17376 | ENSMUSG00000073207 | Ccdc160 | 434778 [Gene Symbol: Ccdc160] [Locus Tag: ] [Chromosome: X] [Map Location: X\|X A5 | 2.565192925 | 0.01513 | 0.030089 |
| 12176 | ENSMUSG00000027884 | Clcc1 | 229725 [Gene Symbol: Clcc1] [Locus Tag: ] [Chromosome: 3] [Map Location: 3\|3 F3] [D | 2.564511619 | 2.00E-05 | 7.23E-05 |
| 3526 | ENSMUSG00000051034 | Zfp11 | 22648 [Gene Symbol: Zfp11] [Locus Tag: ] [Chromosome: 5] [Map Location: 5\|5 G1.3] [ | 2.562939722 | 1.02E-06 | 4.57E-06 |
| 3985 | ENSMUSG00000046844 | Vat1l | 270097 [Gene Symbol: Vat1l] [Locus Tag: ] [Chromosome: 8] [Map Location: 8\|8 E1] [De | 2.562528265 | 1.14E-05 | 4.30E-05 |
| 8617 | ENSMUSG00000024084 | Qpct | 70536 [Gene Symbol: Qpct] [Locus Tag: ] [Chromosome: 17] [Map Location: 17\|17 E3] [ | 2.561528835 | 0.000984 | 0.002562 |
| 3625 | ENSMUSG00000020523 | Fam114a2 | 67726 [Gene Symbol: Fam114a2] [Locus Tag: ] [Chromosome: 11] [Map Location: 11\|1 | 2.560374351 | 1.40E-06 | 6.19E-06 |
| 18425 | ENSMUSG00000037463 | Fbxo27 | 233040 [Gene Symbol: Fbxo27] [Locus Tag: ] [Chromosome: 7] [Map Location: 7\|7 A3] | 2.560109349 | 0.002537 | 0.00605 |
| 13061 | ENSMUSG00000079428 | Tceal7 | 100040972 [Gene Symbol: Tceal7] [Locus Tag: ] [Chromosome: X] [Map Location: X\|X F | 2.558026338 | 0.003854 | 0.008805 |
| 15055 | ENSMUSG00000048109 | Rbm15 | 229700 [Gene Symbol: Rbm15] [Locus Tag: ] [Chromosome: 3] [Map Location: 3\|3 F2.3 | 2.557190357 | 5.10E-07 | 2.42E-06 |
| 12510 | ENSMUSG00000031508 | Ankrd10 | 102334 [Gene Symbol: Ankrd10] [Locus Tag: ] [Chromosome: 8] [Map Location: 8\|8 A1. | 2.555758139 | 1.35E-07 | 7.04E-07 |
| 12777 | ENSMUSG00000008575 | Nfib | 18028 [Gene Symbol: Nfib] [Locus Tag: ] [Chromosome: 4] [Map Location: 4 38.4 cM\|4 | 2.555448781 | 0.000235 | 0.00069 |
| 10191 | ENSMUSG00000043140 | Tmem186 | 66690 [Gene Symbol: Tmem186] [Locus Tag: ] [Chromosome: 16] [Map Location: 16\|16 | 2.554168318 | 1.62E-05 | 5.96E-05 |
| 1668 | ENSMUSG00000029776 | Hibadh | 58875 [Gene Symbol: Hibadh] [Locus Tag: ] [Chromosome: 6] [Map Location: 6\|6 B3] [D | 2.553625754 | 5.10E-06 | 2.03E-05 |
| 11806 | ENSMUSG00000026196 | Bard1 | 12021 [Gene Symbol: Bard1] [Locus Tag: ] [Chromosome: 1] [Map Location: 1\|1 C3] [De | 2.553412842 | 0.001279 | 0.003255 |
| 16209 | ENSMUSG00000030720 | Cln3 | 12752 [Gene Symbol: Cln3] [Locus Tag: ] [Chromosome: 7] [Map Location: 7 F3\|7 69.16 | 2.550864389 | 0.001266 | 0.003227 |
| 2893 | ENSMUSG00000022770 | Dlg1 | 13383 [Gene Symbol: Dlg1] [Locus Tag: ] [Chromosome: 16] [Map Location: 16 B2\|16 2 | 2.549951906 | 4.42E-07 | 2.12E-06 |
| 14018 | ENSMUSG00000038119 | Cdon | 57810 [Gene Symbol: Cdon] [Locus Tag: ] [Chromosome: 9] [Map Location: 9\|9 A4] [Des | 2.549192682 | 0.010525 | 0.021802 |
| 4372 | ENSMUSG00000058298 | Mcm9 | 71567 [Gene Symbol: Mcm9] [Locus Tag: ] [Chromosome: 10] [Map Location: 10\|10 B3 | 2.548844323 | 1.22E-07 | 6.39E-07 |
| 18059 | ENSMUSG00000020745 | Pafah1b1 | 18472 [Gene Symbol: Pafah1b1] [Locus Tag: ] [Chromosome: 11] [Map Location: 11 45. | 2.546461307 | 1.31E-08 | 8.07E-08 |
| 4046 | ENSMUSG00000045259 | Klhdc9 | 68874 [Gene Symbol: Klhdc9] [Locus Tag: ] [Chromosome: 1] [Map Location: 1 H3\|1 79 | 2.545721475 | 0.000232 | 0.000684 |
| 8718 | ENSMUSG00000008892 | Vdac3 | 22335 [Gene Symbol: Vdac3] [Locus Tag: ] [Chromosome: 8] [Map Location: 8 A2\|8 11. | 2.54557145 | 0.001264 | 0.003224 |
| 5038 | ENSMUSG00000026970 | Rbms1 | 56878 [Gene Symbol: Rbms1] [Locus Tag: ] [Chromosome: 2] [Map Location: 2\|2 C3] [D | 2.545065573 | 0.000221 | 0.000651 |
| 12055 | ENSMUSG00000039672 | Kcne2 | 246133 [Gene Symbol: Kcne2] [Locus Tag: ] [Chromosome: 16] [Map Location: 16\|16 C | 2.542472871 | 0.023004 | 0.043624 |
| 6078 | ENSMUSG00000017686 | Rhot1 | 59040 [Gene Symbol: Rhot1] [Locus Tag: ] [Chromosome: 11] [Map Location: 11 B5\|11 | 2.542370313 | 1.33E-07 | 6.94E-07 |
| 13554 | ENSMUSG00000041308 | Sntb2 | 20650 [Gene Symbol: Sntb2] [Locus Tag: ] [Chromosome: 8] [Map Location: 8 D3\|8 53. | 2.541766904 | 3.83E-05 | 0.000131 |
| 6280 | ENSMUSG00000023951 | Vegfa | 22339 [Gene Symbol: Vegfa] [Locus Tag: ] [Chromosome: 17] [Map Location: 17 C\|17 2 | 2.541576684 | 0.000912 | 0.002391 |
| 10662 | ENSMUSG00000026005 | Rpe | 66646 [Gene Symbol: Rpe] [Locus Tag: ] [Chromosome: 1] [Map Location: 1\|1 C3] [Desc | 2.541427468 | 1.91E-06 | 8.23E-06 |
| 15818 | ENSMUSG00000031953 | Tmem170 | 66817 [Gene Symbol: Tmem170] [Locus Tag: ] [Chromosome: 8] [Map Location: 8 58.0 | 2.541394867 | 0.003561 | 0.008202 |
| 12861 | ENSMUSG00000039633 | Lonrf1 | 244421 [Gene Symbol: Lonrf1] [Locus Tag: ] [Chromosome: 8] [Map Location: 8\|8 A4] [ | 2.540768431 | 1.22E-07 | 6.42E-07 |
| 3400 | ENSMUSG00000048911 | Rnf24 | 51902 [Gene Symbol: Rnf24] [Locus Tag: ] [Chromosome: 2] [Map Location: 2 F1\|2 63. | 2.540017428 | 2.80E-06 | 1.17E-05 |
| 16461 | ENSMUSG00000023939 | Mrpl14 | 68463 [Gene Symbol: Mrpl14] [Locus Tag: ] [Chromosome: 17] [Map Location: 17\|17 B | 2.539215345 | 0.00766 | 0.016375 |
| 12463 | ENSMUSG00000040028 | Elavl1 | 15568 [Gene Symbol: Elavl1] [Locus Tag: ] [Chromosome: 8] [Map Location: 8 A1.1\|8 2. | 2.53885004 | 1.81E-06 | 7.83E-06 |
| 9814 | ENSMUSG00000053931 | Cnn3 | 71994 [Gene Symbol: Cnn3] [Locus Tag: ] [Chromosome: 3] [Map Location: 3\|3 G1] [De | 2.537588144 | 0.000643 | 0.001736 |
| 6481 | ENSMUSG00000064367 | ND5 | 17721 [Gene Symbol: ND5] [Locus Tag: ] [Chromosome: MT] [Map Location: ] [Descripti | 2.536999446 | 3.50E-08 | 2.00E-07 |
| 6090 | ENSMUSG00000098274 | LOC102641064; Rpl2 | 102641064 [Gene Symbol: LOC102641064] [Locus Tag: ] [Chromosome: 4] [Map Locati | 2.536495861 | 0.013992 | 0.028096 |
| 16093 | ENSMUSG00000025234 | Arih1 | 23806 [Gene Symbol: Arih1] [Locus Tag: ] [Chromosome: 9] [Map Location: 9\|9 C] [Desc | 2.53542429 | 4.63E-08 | 2.60E-07 |
| 8979 | ENSMUSG00000040855 | Reps2 | 194590 [Gene Symbol: Reps2] [Locus Tag: ] [Chromosome: X] [Map Location: X\|X F4] [D | 2.535326854 | 1.16E-06 | 5.19E-06 |
| 3487 | ENSMUSG00000039942 | Ptger4 | 19219 [Gene Symbol: Ptger4] [Locus Tag: ] [Chromosome: 15] [Map Location: 15 A1\|15 | 2.534893839 | 7.62E-08 | 4.11E-07 |
| 1504 | ENSMUSG00000056888 | Glipr1 | 73690 [Gene Symbol: Glipr1] [Locus Tag: ] [Chromosome: 10] [Map Location: 10\|10 D1 | 2.534738157 | 0.000878 | 0.002311 |
| 17642 | ENSMUSG00000015653 | Steap2 | 74051 [Gene Symbol: Steap2] [Locus Tag: ] [Chromosome: 5] [Map Location: 5\|5 A1] [D | 2.534587085 | 7.70E-06 | 2.99E-05 |
| 9296 | ENSMUSG00000034098 | Fstl5 | 213262 [Gene Symbol: Fstl5] [Locus Tag: ] [Chromosome: 3] [Map Location: 3\|3 E3] [De | 2.534479405 | 1.95E-07 | 9.85E-07 |
| 4041 | ENSMUSG00000036036 | Zfp57 | 22715 [Gene Symbol: Zfp57] [Locus Tag: ] [Chromosome: 17] [Map Location: 17\|17 B3] | 2.534356312 | 2.74E-06 | 1.14E-05 |
| 15246 | ENSMUSG00000046139 | Patl1 | 225929 [Gene Symbol: Patl1] [Locus Tag: ] [Chromosome: 19] [Map Location: 19\|19 A] | 2.53315928 | 1.67E-07 | 8.55E-07 |
| 8852 | ENSMUSG00000045679 | Pqlc3 | 217430 [Gene Symbol: Pqlc3] [Locus Tag: ] [Chromosome: 12] [Map Location: 12\|12 A | 2.532586616 | 0.004339 | 0.009806 |
| 7041 | ENSMUSG00000050071 | Bex1 | 19716 [Gene Symbol: Bex1] [Locus Tag: ] [Chromosome: X] [Map Location: X F1\|X 57.4 | 2.532400496 | 4.81E-07 | 2.29E-06 |
| 18557 | ENSMUSG00000036339 | Tmem260 | 218989 [Gene Symbol: Tmem260] [Locus Tag: ] [Chromosome: 14] [Map Location: 14\| | 2.532389737 | 7.68E-07 | 3.53E-06 |
| 5048 | ENSMUSG00000046010 | Zfp830 | 66983 [Gene Symbol: Zfp830] [Locus Tag: ] [Chromosome: 11] [Map Location: 11\|11 C] | 2.531983762 | 2.02E-05 | 7.28E-05 |
| 17272 | ENSMUSG00000078495 | Gm13157 | 100041677 [Gene Symbol: Gm13157] [Locus Tag: ] [Chromosome: 4] [Map Location: 4 | 2.531922606 | 0.002054 | 0.004989 |
| 15141 | ENSMUSG00000038299 | Wdr36 | 225348 [Gene Symbol: Wdr36] [Locus Tag: ] [Chromosome: 18] [Map Location: 18\|18 | 2.531017499 | 4.71E-06 | 1.89E-05 |
| 9438 | ENSMUSG00000068966 | Zbtb34 | 241311 [Gene Symbol: Zbtb34] [Locus Tag: ] [Chromosome: 2] [Map Location: 2\|2 B] [D | 2.529318378 | 5.82E-08 | 3.20E-07 |
| 386 | ENSMUSG00000041040 | Fam117b | 72750 [Gene Symbol: Fam117b] [Locus Tag: ] [Chromosome: 1] [Map Location: 1\|1 C2] | 2.527753127 | 3.88E-08 | 2.21E-07 |
| 36 | ENSMUSG00000106744 | Riok2 | 67045 [Gene Symbol: Riok2] [Locus Tag: ] [Chromosome: 17] [Map Location: 17\|17 A3. | 2.52628028 | 8.65E-06 | 3.32E-05 |
| 4631 | ENSMUSG00000026469 | Xpr1 | 19775 [Gene Symbol: Xpr1] [Locus Tag: ] [Chromosome: 1] [Map Location: 1 G3\|1 66.4 | 2.525477136 | 2.15E-06 | 9.16E-06 |
| 5274 | ENSMUSG00000046550 | Spin2c | 278240 [Gene Symbol: Spin2c] [Locus Tag: ] [Chromosome: X] [Map Location: X\|X F3] [ | 2.525267826 | 0.00078 | 0.00207 |
| 199 | ENSMUSG00000012443 | Kif11 | 16551 [Gene Symbol: Kif11] [Locus Tag: ] [Chromosome: 19] [Map Location: 19\|19 C2] | 2.524625659 | 0.00024 | 0.000705 |
| 14214 | ENSMUSG00000053870 | Fpgt | 75540 [Gene Symbol: Fpgt] [Locus Tag: ] [Chromosome: 3] [Map Location: 3\|3 H4] [Des | 2.523851926 | 2.13E-06 | 9.08E-06 |
| 3334 | ENSMUSG00000042195 | Slc35f2 | 72022 [Gene Symbol: Slc35f2] [Locus Tag: ] [Chromosome: 9] [Map Location: 9\|9 A5.3] | 2.523672717 | 0.000175 | 0.000526 |
| 15684 | ENSMUSG00000093904 | Tomm20 | 67952 [Gene Symbol: Tomm20] [Locus Tag: ] [Chromosome: 8] [Map Location: 8\|8 E2] | 2.520479092 | 1.50E-07 | 7.79E-07 |

| | | | | | | |
|---|---|---|---|---|---|---|
| 1819 | ENSMUSG00000004768 | Rab23 | 19335 [Gene Symbol: Rab23] [Locus Tag: ] [Chromosome: 1] [Map Location: 1 B\|1 12.8 | 2.520430013 | 1.08E-07 | 5.72E-07 |
| 2666 | ENSMUSG00000003099 | Ppp5c | 19060 [Gene Symbol: Ppp5c] [Locus Tag: ] [Chromosome: 7] [Map Location: 7 A2\|7 9.1 | 2.51981539 | 1.54E-07 | 7.93E-07 |
| 1564 | ENSMUSG00000035764 | Fbxo45 | 268882 [Gene Symbol: Fbxo45] [Locus Tag: ] [Chromosome: 16] [Map Location: 16\|16 | 2.51942036 | 3.84E-07 | 1.85E-06 |
| 2456 | ENSMUSG00000047446 | Arl4a | 11861 [Gene Symbol: Arl4a] [Locus Tag: ] [Chromosome: 12] [Map Location: 12 B1\|12 | 2.519315622 | 0.000175 | 0.000526 |
| 13866 | ENSMUSG00000058240 | Cryzl1 | 66609 [Gene Symbol: Cryzl1] [Locus Tag: ] [Chromosome: 16] [Map Location: 16\|16 C4 | 2.517619856 | 2.82E-06 | 1.18E-05 |
| 16422 | ENSMUSG00000041215 | Yeats2 | 208146 [Gene Symbol: Yeats2] [Locus Tag: ] [Chromosome: 16] [Map Location: 16\|16 A | 2.517114132 | 1.71E-07 | 8.73E-07 |
| 16370 | ENSMUSG00000028124 | Gclm | 14630 [Gene Symbol: Gclm] [Locus Tag: ] [Chromosome: 3] [Map Location: 3 52.94 cM\| | 2.51660204 | 1.39E-07 | 7.21E-07 |
| 11245 | ENSMUSG00000041841 | Rpl37rt; Rpl37 | 100502825 [Gene Symbol: Rpl37rt] [Locus Tag: ] [Chromosome: 5] [Map Location: 5\|5 | 2.516596301 | 0.003851 | 0.0088 |
| 9796 | ENSMUSG00000028565 | Nfia | 18027 [Gene Symbol: Nfia] [Locus Tag: ] [Chromosome: 4] [Map Location: 4 45.52 cM\|4 | 2.51585823 | 1.98E-06 | 8.47E-06 |
| 232 | ENSMUSG00000049086 | Bmyc | 107771 [Gene Symbol: Bmyc] [Locus Tag: ] [Chromosome: 2] [Map Location: 2\|2 A3] [D | 2.512641837 | 1.38E-06 | 6.10E-06 |
| 277 | ENSMUSG00000039233 | Tbce | 70430 [Gene Symbol: Tbce] [Locus Tag: ] [Chromosome: 13] [Map Location: 13 A1\|13 5 | 2.511926729 | 1.32E-06 | 5.85E-06 |
| 11941 | ENSMUSG00000041974 | Spidr | 224008 [Gene Symbol: Spidr] [Locus Tag: ] [Chromosome: 16] [Map Location: 16\|16 A2 | 2.51095804 | 4.66E-05 | 0.000156 |
| 6696 | ENSMUSG00000029179 | Zcchc4 | 78796 [Gene Symbol: Zcchc4] [Locus Tag: ] [Chromosome: 5] [Map Location: 5\|5 C1] [D | 2.510565984 | 2.32E-05 | 8.27E-05 |
| 11269 | ENSMUSG00000036817 | Sun1 | 77053 [Gene Symbol: Sun1] [Locus Tag: ] [Chromosome: 5] [Map Location: 5\|5 G1] [Des | 2.510221623 | 1.39E-07 | 7.20E-07 |
| 16253 | ENSMUSG00000045100 | Slc25a26 | 67582 [Gene Symbol: Slc25a26] [Locus Tag: ] [Chromosome: 6] [Map Location: 6\|6 D3] | 2.507874092 | 4.85E-05 | 0.000162 |
| 827 | ENSMUSG00000054693 | Adam10 | 11487 [Gene Symbol: Adam10] [Locus Tag: ] [Chromosome: 9] [Map Location: 9 D\|9 39 | 2.507423312 | 1.51E-07 | 7.82E-07 |
| 14100 | ENSMUSG00000020986 | Sec23a | 20334 [Gene Symbol: Sec23a] [Locus Tag: ] [Chromosome: 12] [Map Location: 12\|12 C1 | 2.507365401 | 4.48E-08 | 2.52E-07 |
| 1397 | ENSMUSG00000004661 | Arid3b | 56380 [Gene Symbol: Arid3b] [Locus Tag: ] [Chromosome: 9] [Map Location: 9\|9 C] [Des | 2.506697146 | 0.000235 | 0.000691 |
| 3879 | ENSMUSG00000039033 | Tasp1 | 75812 [Gene Symbol: Tasp1] [Locus Tag: ] [Chromosome: 2] [Map Location: 2\|2 F3] [De | 2.504847981 | 4.62E-07 | 2.20E-06 |
| 18300 | ENSMUSG00000075376 | Rc3h2 | 319817 [Gene Symbol: Rc3h2] [Locus Tag: ] [Chromosome: 2] [Map Location: 2\|2 B] [De | 2.502173946 | 2.98E-08 | 1.72E-07 |
| 15847 | ENSMUSG00000022635 | Zcrb1 | 67197 [Gene Symbol: Zcrb1] [Locus Tag: ] [Chromosome: 15] [Map Location: 15\|15 F1] | 2.501669669 | 4.11E-07 | 1.97E-06 |
| 14011 | ENSMUSG00000056296 | Synpr | 72003 [Gene Symbol: Synpr] [Locus Tag: ] [Chromosome: 14] [Map Location: 14\|14 A2] | 2.501359324 | 1.68E-05 | 6.17E-05 |
| 12524 | ENSMUSG00000031090 | Nadsyn1 | 78914 [Gene Symbol: Nadsyn1] [Locus Tag: ] [Chromosome: 7] [Map Location: 7\|7 F5] [ | 2.499722021 | 0.008032 | 0.017092 |
| 754 | ENSMUSG00000018995 | Nars2 | 244141 [Gene Symbol: Nars2] [Locus Tag: ] [Chromosome: 7] [Map Location: 7\|7 E1] [D | 2.498144435 | 3.86E-05 | 0.000132 |
| 1911 | ENSMUSG00000031479 | Vps36 | 70160 [Gene Symbol: Vps36] [Locus Tag: ] [Chromosome: 8] [Map Location: 8\|8 A3] [De | 2.497330711 | 2.81E-07 | 1.38E-06 |
| 515 | ENSMUSG00000015354 | Pcolce2 | 76477 [Gene Symbol: Pcolce2] [Locus Tag: ] [Chromosome: 9] [Map Location: 9\|9 E3.3] | 2.497117849 | 0.008572 | 0.018123 |
| 8964 | ENSMUSG00000064280 | Ccdc146 | 75172 [Gene Symbol: Ccdc146] [Locus Tag: ] [Chromosome: 5] [Map Location: 5\|5 A3] | 2.495915455 | 0.001029 | 0.00267 |
| 11996 | ENSMUSG00000022022 | Mtrf1 | 211253 [Gene Symbol: Mtrf1] [Locus Tag: ] [Chromosome: 14] [Map Location: 14\|14 D | 2.495857676 | 0.000688 | 0.001847 |
| 13914 | ENSMUSG00000066551 | Gm21596; Hmgb1 | 100862258 [Gene Symbol: Gm21596] [Locus Tag: ] [Chromosome: 2] [Map Location: 2 | 2.494430837 | 2.02E-06 | 8.62E-06 |
| 4012 | ENSMUSG00000018752 | Tnfsfm13 | 619441 [Gene Symbol: Tnfsfm13] [Locus Tag: ] [Chromosome: 11] [Map Location: 11\|1 | 2.49235689 | 0.000803 | 0.002127 |
| 972 | ENSMUSG00000028271 | Gtf2b | 229906 [Gene Symbol: Gtf2b] [Locus Tag: ] [Chromosome: 3] [Map Location: 3\|3 H1] [D | 2.492245445 | 2.15E-05 | 7.72E-05 |
| 7955 | ENSMUSG00000039159 | Ube2h | 22214 [Gene Symbol: Ube2h] [Locus Tag: ] [Chromosome: 6] [Map Location: 6 A3.3\|6 1 | 2.491818815 | 2.75E-07 | 1.35E-06 |
| 1126 | ENSMUSG00000024999 | Noc3l | 57753 [Gene Symbol: Noc3l] [Locus Tag: ] [Chromosome: 19] [Map Location: 19\|19 C3 | 2.491657712 | 2.34E-05 | 8.34E-05 |
| 5736 | ENSMUSG00000038718 | Pbx3 | 18516 [Gene Symbol: Pbx3] [Locus Tag: ] [Chromosome: 2] [Map Location: 2 B\|2 22.84 | 2.491612235 | 2.38E-07 | 1.19E-06 |
| 15271 | ENSMUSG00000092416 | Zfp141 | 434178 [Gene Symbol: Zfp141] [Locus Tag: ] [Chromosome: 7] [Map Location: 7\|7 B3] [ | 2.490417171 | 2.00E-06 | 8.57E-06 |
| 10228 | ENSMUSG00000048004 | Tmem196 | 217951 [Gene Symbol: Tmem196] [Locus Tag: ] [Chromosome: 12] [Map Location: 12\| | 2.490270551 | 0.000741 | 0.001976 |
| 17868 | ENSMUSG00000042590 | Ipo11 | 76582 [Gene Symbol: Ipo11] [Locus Tag: ] [Chromosome: 13] [Map Location: 13\|13 D2 | 2.488854144 | 2.25E-06 | 9.54E-06 |
| 2414 | ENSMUSG00000029649 | Pomp | 66537 [Gene Symbol: Pomp] [Locus Tag: ] [Chromosome: 5] [Map Location: 5\|5 G3] [De | 2.488572347 | 2.12E-06 | 9.04E-06 |
| 5130 | ENSMUSG00000039286 | Fndc3b | 72007 [Gene Symbol: Fndc3b] [Locus Tag: ] [Chromosome: 3] [Map Location: 3\|3 A3] [D | 2.488067813 | 1.13E-07 | 5.94E-07 |
| 6073 | ENSMUSG00000026839 | Upp2 | 76654 [Gene Symbol: Upp2] [Locus Tag: ] [Chromosome: 2] [Map Location: 2\|2 C3] [Des | 2.485469808 | 4.11E-05 | 0.000139 |
| 17633 | ENSMUSG00000036972 | Zic4 | 22774 [Gene Symbol: Zic4] [Locus Tag: ] [Chromosome: 9] [Map Location: 9 E3.3\|9 48. | 2.485345388 | 1.19E-07 | 6.26E-07 |
| 15606 | ENSMUSG00000020311 | Erlec1 | 66753 [Gene Symbol: Erlec1] [Locus Tag: ] [Chromosome: 11] [Map Location: 11\|11 A4 | 2.482397744 | 1.75E-06 | 7.59E-06 |
| 10877 | ENSMUSG00000022742 | Cpox | 12892 [Gene Symbol: Cpox] [Locus Tag: ] [Chromosome: 16] [Map Location: 16\|16 C1. | 2.481390516 | 0.0002 | 0.000596 |
| 14558 | ENSMUSG00000094936 | Rbm4 | 19653 [Gene Symbol: Rbm4] [Locus Tag: ] [Chromosome: 19] [Map Location: 19\|19 A] [ | 2.480894512 | 3.15E-06 | 1.30E-05 |
| 10188 | ENSMUSG00000056131 | Pgm3 | 109785 [Gene Symbol: Pgm3] [Locus Tag: ] [Chromosome: 9] [Map Location: 9 46.58 cM | 2.480802392 | 3.21E-05 | 0.000111 |
| 4341 | ENSMUSG00000051278 | Zgrf1 | 71643 [Gene Symbol: Zgrf1] [Locus Tag: ] [Chromosome: 3] [Map Location: 3\|3 G2] [Des | 2.480596998 | 0.0014 | 0.003538 |
| 4005 | ENSMUSG00000021051 | Ppp2r5e | 26932 [Gene Symbol: Ppp2r5e] [Locus Tag: ] [Chromosome: 12] [Map Location: 12\|12 | 2.479792471 | 7.89E-08 | 4.25E-07 |
| 18330 | ENSMUSG00000020640 | Itsn2 | 20403 [Gene Symbol: Itsn2] [Locus Tag: ] [Chromosome: 12] [Map Location: 12\|12 A1. | 2.479582557 | 0.000207 | 0.000615 |
| 2729 | ENSMUSG00000035126 | Wdr78 | 242584 [Gene Symbol: Wdr78] [Locus Tag: ] [Chromosome: 4] [Map Location: 4\|4 C6] [ | 2.479516314 | 0.00146 | 0.003671 |
| 4750 | ENSMUSG00000042105 | Inpp5f | 101490 [Gene Symbol: Inpp5f] [Locus Tag: ] [Chromosome: 7] [Map Location: 7\|7 F3] [D | 2.479440806 | 3.49E-08 | 2.00E-07 |
| 2863 | ENSMUSG00000038530 | Rgs4 | 19736 [Gene Symbol: Rgs4] [Locus Tag: ] [Chromosome: 1] [Map Location: 1 H3\|1 76.8 | 2.478971763 | 2.56E-07 | 1.27E-06 |
| 6199 | ENSMUSG00000033885 | Pxk | 218699 [Gene Symbol: Pxk] [Locus Tag: ] [Chromosome: 14] [Map Location: 14 A1\|14 4 | 2.478337838 | 5.17E-05 | 0.000171 |
| 16879 | ENSMUSG00000045257 | Morn2 | 378462 [Gene Symbol: Morn2] [Locus Tag: ] [Chromosome: 17] [Map Location: 17\|17 E | 2.477956239 | 0.013299 | 0.026825 |
| 666 | ENSMUSG00000029287 | Tgfbr3 | 21814 [Gene Symbol: Tgfbr3] [Locus Tag: ] [Chromosome: 5] [Map Location: 5\|5 E5] [De | 2.477894663 | 8.24E-06 | 3.18E-05 |
| 2203 | ENSMUSG00000031292 | Cdkl5 | 382253 [Gene Symbol: Cdkl5] [Locus Tag: ] [Chromosome: X] [Map Location: X F4\|X 73. | 2.477364529 | 6.49E-08 | 3.54E-07 |
| 6998 | ENSMUSG00000020069 | Hnrnph3 | 432467 [Gene Symbol: Hnrnph3] [Locus Tag: ] [Chromosome: 10] [Map Location: 10\|1 | 2.476776275 | 1.20E-05 | 4.51E-05 |
| 10378 | ENSMUSG00000024045 | Akap8 | 56399 [Gene Symbol: Akap8] [Locus Tag: ] [Chromosome: 17] [Map Location: 17\|17 B2 | 2.476330691 | 3.17E-06 | 1.31E-05 |
| 11306 | ENSMUSG00000056383 | AI987944 | 233168 [Gene Symbol: AI987944] [Locus Tag: ] [Chromosome: 7] [Map Location: 7\|7 B | 2.47585914 | 0.004242 | 0.009603 |
| 6949 | ENSMUSG00000041966 | Dcaf17 | 75763 [Gene Symbol: Dcaf17] [Locus Tag: ] [Chromosome: 2] [Map Location: 2\|2 C2] [D | 2.475669715 | 1.40E-06 | 6.19E-06 |
| 9417 | ENSMUSG00000052593 | Adam17 | 11491 [Gene Symbol: Adam17] [Locus Tag: ] [Chromosome: 12] [Map Location: 12 A1.3 | 2.475002283 | 3.35E-06 | 1.38E-05 |
| 11435 | ENSMUSG00000022136 | Dnajc3 | 100037258 [Gene Symbol: Dnajc3] [Locus Tag: ] [Chromosome: 14] [Map Location: 14\| | 2.474038783 | 6.07E-07 | 2.84E-06 |
| 7230 | ENSMUSG00000045948 | Mrps12 | 24030 [Gene Symbol: Mrps12] [Locus Tag: ] [Chromosome: 7] [Map Location: 7\|7 A3] [D | 2.474021775 | 0.000305 | 0.000877 |
| 4201 | ENSMUSG00000041219 | Arhgap11a | 228482 [Gene Symbol: Arhgap11a] [Locus Tag: ] [Chromosome: 2] [Map Location: 2\|2 E | 2.472848712 | 0.022367 | 0.04255 |
| 14615 | ENSMUSG00000027014 | Cwc22 | 80744 [Gene Symbol: Cwc22] [Locus Tag: ] [Chromosome: 2] [Map Location: 2\|2 C3] [D | 2.47283833 | 0.000168 | 0.000506 |
| 10073 | ENSMUSG00000024271 | Elp2 | 58523 [Gene Symbol: Elp2] [Locus Tag: ] [Chromosome: 18] [Map Location: 18\|18 A2] [ | 2.471437657 | 3.44E-07 | 1.67E-06 |
| 7739 | ENSMUSG00000021477 | Ctsl | 13039 [Gene Symbol: Ctsl] [Locus Tag: ] [Chromosome: 13] [Map Location: 13 B3\|13 33 | 2.470447005 | 5.10E-05 | 0.000169 |
| 7502 | ENSMUSG00000026109 | Tmeff2 | 56363 [Gene Symbol: Tmeff2] [Locus Tag: ] [Chromosome: 1] [Map Location: 1\|1 C1] [D | 2.468957378 | 3.20E-06 | 1.32E-05 |
| 14900 | ENSMUSG00000041540 | Sox5 | 20678 [Gene Symbol: Sox5] [Locus Tag: ] [Chromosome: 6] [Map Location: 6 G3\|6 76.1 | 2.466835534 | 9.32E-08 | 4.97E-07 |
| 5537 | ENSMUSG00000068823 | Csde1 | 229663 [Gene Symbol: Csde1] [Locus Tag: ] [Chromosome: 3] [Map Location: 3 F2.2\|3 4 | 2.466436342 | 7.72E-08 | 4.16E-07 |
| 4526 | ENSMUSG00000070866 | Zfp804a | 241514 [Gene Symbol: Zfp804a] [Locus Tag: ] [Chromosome: 2] [Map Location: 2\|2 D] [ | 2.464908913 | 7.67E-05 | 0.000246 |
| 9581 | ENSMUSG00000030104 | Edem1 | 192193 [Gene Symbol: Edem1] [Locus Tag: ] [Chromosome: 6] [Map Location: 6\|6 E2] [ | 2.46457806 | 1.25E-07 | 6.57E-07 |
| 4853 | ENSMUSG00000048701 | Ccdc6 | 76551 [Gene Symbol: Ccdc6] [Locus Tag: ] [Chromosome: 10] [Map Location: 10\|10 B5 | 2.464263037 | 9.67E-08 | 5.15E-07 |
| 12610 | ENSMUSG00000020160 | Meis1 | 17268 [Gene Symbol: Meis1] [Locus Tag: ] [Chromosome: 11] [Map Location: 11 A3.1\|1 | 2.464228629 | 2.04E-05 | 7.35E-05 |
| 15738 | ENSMUSG00000019828 | Grm1 | 14816 [Gene Symbol: Grm1] [Locus Tag: ] [Chromosome: 10] [Map Location: 10\|10 A2] | 2.464168904 | 3.56E-06 | 1.45E-05 |
| 2328 | ENSMUSG00000032479 | Map4 | 17758 [Gene Symbol: Map4] [Locus Tag: ] [Chromosome: 9] [Map Location: 9 F2\|9 59.8 | 2.463781823 | 3.68E-05 | 0.000126 |
| 10153 | ENSMUSG00000033377 | Palmd | 114301 [Gene Symbol: Palmd] [Locus Tag: ] [Chromosome: 3] [Map Location: 3 G1\|3 5 | 2.463682392 | 3.38E-05 | 0.000117 |
| 7026 | ENSMUSG00000027281 | Slx4ip | 74243 [Gene Symbol: Slx4ip] [Locus Tag: ] [Chromosome: 2] [Map Location: 2\|2 G1] [De | 2.463242838 | 2.07E-05 | 7.45E-05 |
| 1873 | ENSMUSG00000032698 | Lmo2 | 16909 [Gene Symbol: Lmo2] [Locus Tag: ] [Chromosome: 2] [Map Location: 2 E2\|2 54.4 | 2.463202701 | 3.43E-05 | 0.000118 |
| 135 | ENSMUSG00000060301 | 2610008E11Rik | 72128 [Gene Symbol: 2610008E11Rik] [Locus Tag: ] [Chromosome: 10] [Map Location: | 2.461863743 | 4.95E-06 | 1.97E-05 |
| 11590 | ENSMUSG00000031432 | Prps1 | 19139 [Gene Symbol: Prps1] [Locus Tag: ] [Chromosome: X] [Map Location: X 61.35 cM | 2.461545754 | 1.91E-05 | 6.91E-05 |
| 14625 | ENSMUSG00000015575 | Atp6v0e | 11974 [Gene Symbol: Atp6v0e] [Locus Tag: ] [Chromosome: 17] [Map Location: 17\|17 | 2.461487516 | 0.000516 | 0.001421 |
| 9962 | ENSMUSG00000050538 | B230217C12Rik | 68127 [Gene Symbol: B230217C12Rik] [Locus Tag: ] [Chromosome: 11] [Map Location: | 2.461404171 | 2.36E-05 | 8.41E-05 |
| 16188 | ENSMUSG00000052914 | Cyp2j6 | 13110 [Gene Symbol: Cyp2j6] [Locus Tag: ] [Chromosome: 4] [Map Location: 4 C5\|4 44 | 2.461336293 | 0.00139 | 0.003516 |
| 12155 | ENSMUSG00000021660 | Btf3 | 218490 [Gene Symbol: Btf3] [Locus Tag: ] [Chromosome: 13] [Map Location: 13\|13 D1] | 2.461029763 | 5.12E-05 | 0.00017 |

| | | | | | | |
|---|---|---|---|---|---|---|
| 15461 | ENSMUSG00000036777 | Anln | 68743 [Gene Symbol: Anln] [Locus Tag: ] [Chromosome: 9] [Map Location: 9\|9 A4] [Des | 2.460119431 | 4.33E-05 | 0.000146 |
| 9780 | ENSMUSG00000073514 | Dok6 | 623279 [Gene Symbol: Dok6] [Locus Tag: ] [Chromosome: 18] [Map Location: 18\|18 E4 | 2.459902301 | 2.14E-07 | 1.07E-06 |
| 15045 | ENSMUSG00000057396 | Zfp759 | 268670 [Gene Symbol: Zfp759] [Locus Tag: ] [Chromosome: 13] [Map Location: 13 B3\|1 | 2.458649248 | 0.00025 | 0.000731 |
| 6801 | ENSMUSG00000029534 | St7 | 64213 [Gene Symbol: St7] [Locus Tag: ] [Chromosome: 6] [Map Location: 6\|6 A2] [Descr | 2.458105633 | 4.36E-07 | 2.09E-06 |
| 6294 | ENSMUSG00000034343 | Ube2f | 67921 [Gene Symbol: Ube2f] [Locus Tag: ] [Chromosome: 1] [Map Location: 1\|1 D] [Des | 2.456599648 | 7.59E-07 | 3.49E-06 |
| 670 | ENSMUSG00000045648 | Vwc2l | 320460 [Gene Symbol: Vwc2l] [Locus Tag: ] [Chromosome: 1] [Map Location: 1\|1 C3] [D | 2.456498295 | 7.98E-06 | 3.09E-05 |
| 14419 | ENSMUSG00000054871 | Tmem158 | 72309 [Gene Symbol: Tmem158] [Locus Tag: ] [Chromosome: 9] [Map Location: 9\|9 F4] | 2.456237434 | 0.000175 | 0.000527 |
| 3659 | ENSMUSG00000035954 | Dock4 | 238130 [Gene Symbol: Dock4] [Locus Tag: ] [Chromosome: 12] [Map Location: 12 B1\|1 | 2.453934819 | 8.76E-08 | 4.69E-07 |
| 10574 | ENSMUSG00000002808 | Epdr1 | 105298 [Gene Symbol: Epdr1] [Locus Tag: ] [Chromosome: 13] [Map Location: 13\|13 A | 2.453193277 | 3.64E-05 | 0.000125 |
| 18375 | ENSMUSG00000019984 | Med23 | 70208 [Gene Symbol: Med23] [Locus Tag: ] [Chromosome: 10] [Map Location: 10 A4\|1( | 2.45317627 | 2.16E-06 | 9.17E-06 |
| 16544 | ENSMUSG00000093938 | Evi2b | 216984 [Gene Symbol: Evi2b] [Locus Tag: ] [Chromosome: 11] [Map Location: 11\|11 B5 | 2.449870027 | 0.020332 | 0.039089 |
| 11085 | ENSMUSG00000028207 | Asph | 65973 [Gene Symbol: Asph] [Locus Tag: ] [Chromosome: 4] [Map Location: 4\|4 A1] [Des | 2.449183929 | 1.32E-07 | 6.91E-07 |
| 1781 | ENSMUSG00000030101 | Sumf1 | 58911 [Gene Symbol: Sumf1] [Locus Tag: ] [Chromosome: 6] [Map Location: 6\|6 E1] [De | 2.447860795 | 0.000244 | 0.000715 |
| 11084 | ENSMUSG00000029790 | Cep41 | 83922 [Gene Symbol: Cep41] [Locus Tag: ] [Chromosome: 6] [Map Location: 6\|6 A3.3] [ | 2.447103687 | 0.000253 | 0.00074 |
| 8306 | ENSMUSG00000048450 | Msx1 | 17701 [Gene Symbol: Msx1] [Locus Tag: ] [Chromosome: 5] [Map Location: 5 B3\|5 20.2 | 2.446641391 | 0.00097 | 0.002529 |
| 9448 | ENSMUSG00000024766 | Lipo1 | 381236 [Gene Symbol: Lipo1] [Locus Tag: ] [Chromosome: 19] [Map Location: 19\|19 C | 2.446193253 | 0.00044 | 0.001229 |
| 7359 | ENSMUSG00000048027 | Rgmb | 68799 [Gene Symbol: Rgmb] [Locus Tag: ] [Chromosome: 17] [Map Location: 17\|17 A2] | 2.445173481 | 1.65E-07 | 8.46E-07 |
| 17311 | ENSMUSG00000060985 | Tdrd5 | 214575 [Gene Symbol: Tdrd5] [Locus Tag: ] [Chromosome: 1] [Map Location: 1\|1 G3] [D | 2.444658944 | 0.023249 | 0.044037 |
| 5928 | ENSMUSG00000041298 | Katnal1 | 231912 [Gene Symbol: Katnal1] [Locus Tag: ] [Chromosome: 5] [Map Location: 5\|5 G3] | 2.444246337 | 1.52E-07 | 7.84E-07 |
| 3949 | ENSMUSG00000074698 | Csnk2a1 | 12995 [Gene Symbol: Csnk2a1] [Locus Tag: ] [Chromosome: 2] [Map Location: 2\|2 G3] [ | 2.443204614 | 4.49E-07 | 2.15E-06 |
| 12147 | ENSMUSG00000038533 | Cbfa2t2 | 12396 [Gene Symbol: Cbfa2t2] [Locus Tag: ] [Chromosome: 2] [Map Location: 2\|2 C3.2 | 2.443078785 | 1.76E-07 | 9.00E-07 |
| 16533 | ENSMUSG00000030534 | Vps33b | 233405 [Gene Symbol: Vps33b] [Locus Tag: ] [Chromosome: 7] [Map Location: 7\|7 D3] | 2.4429816 | 1.85E-05 | 6.72E-05 |
| 6845 | ENSMUSG00000041124 | Msantd4 | 78100 [Gene Symbol: Msantd4] [Locus Tag: ] [Chromosome: 9] [Map Location: 9\|9 A1] | 2.442976962 | 1.19E-05 | 4.46E-05 |
| 13113 | ENSMUSG00000015342 | Xk | 22439 [Gene Symbol: Xk] [Locus Tag: ] [Chromosome: X] [Map Location: X A1.1\|X 4.22 | 2.442083512 | 6.09E-06 | 2.40E-05 |
| 9205 | ENSMUSG00000006678 | Pola1 | 18968 [Gene Symbol: Pola1] [Locus Tag: ] [Chromosome: X] [Map Location: X 41.06 cM | 2.438239731 | 3.57E-06 | 1.46E-05 |
| 16035 | ENSMUSG00000039449 | Prpf18 | 67229 [Gene Symbol: Prpf18] [Locus Tag: ] [Chromosome: 2] [Map Location: 2\|2 A1] [D | 2.438114439 | 1.73E-06 | 7.52E-06 |
| 13130 | ENSMUSG00000029552 | Tes | 21753 [Gene Symbol: Tes] [Locus Tag: ] [Chromosome: 6] [Map Location: 6 A2\|6 7.67 cl | 2.437998923 | 2.15E-06 | 9.15E-06 |
| 1362 | ENSMUSG00000022708 | Zbtb20 | 56490 [Gene Symbol: Zbtb20] [Locus Tag: ] [Chromosome: 16] [Map Location: 16 B4\|1 | 2.437907358 | 1.85E-06 | 8.00E-06 |
| 10791 | ENSMUSG00000059436 | Max | 17187 [Gene Symbol: Max] [Locus Tag: ] [Chromosome: 12] [Map Location: 12 33.78 cM | 2.436825932 | 5.82E-07 | 2.73E-06 |
| 2891 | ENSMUSG00000026994 | Galnt3 | 14425 [Gene Symbol: Galnt3] [Locus Tag: ] [Chromosome: 2] [Map Location: 2\|2 C3] [D | 2.435895472 | 0.016616 | 0.032715 |
| 7328 | ENSMUSG00000025868 | Higd2a | 67044 [Gene Symbol: Higd2a] [Locus Tag: ] [Chromosome: 13] [Map Location: 13\|13 B | 2.435658122 | 2.21E-05 | 7.90E-05 |
| 2500 | ENSMUSG00000028360 | Slc44a5 | 242259 [Gene Symbol: Slc44a5] [Locus Tag: ] [Chromosome: 3] [Map Location: 3\|3 H3- | 2.4355935 | 6.12E-07 | 2.86E-06 |
| 15545 | ENSMUSG00000037860 | Aim2 | 383619 [Gene Symbol: Aim2] [Locus Tag: ] [Chromosome: 1] [Map Location: 1\|1 H3] [D | 2.434464706 | 4.89E-05 | 0.000163 |
| 15113 | ENSMUSG00000006740 | Kif5b | 16573 [Gene Symbol: Kif5b] [Locus Tag: ] [Chromosome: 18] [Map Location: 18 A2-B1\| | 2.433924436 | 1.48E-07 | 7.68E-07 |
| 18056 | ENSMUSG00000029432 | Gbas | 14467 [Gene Symbol: Gbas] [Locus Tag: ] [Chromosome: 5] [Map Location: 5\|5 G1.3] [D | 2.433665833 | 2.37E-06 | 1.00E-05 |
| 16200 | ENSMUSG00000049038 | Mterf2 | 74238 [Gene Symbol: Mterf2] [Locus Tag: ] [Chromosome: 10] [Map Location: 10\|10 C1 | 2.433288359 | 5.00E-05 | 0.000167 |
| 15982 | ENSMUSG00000040652 | Oaz2 | 18247 [Gene Symbol: Oaz2] [Locus Tag: ] [Chromosome: 9] [Map Location: 9\|9 C] [Descr | 2.433211131 | 9.09E-06 | 3.47E-05 |
| 15549 | ENSMUSG00000035284 | Vps13c | 320528 [Gene Symbol: Vps13c] [Locus Tag: ] [Chromosome: 9] [Map Location: 9\|9 C] [D | 2.431899147 | 2.85E-07 | 1.40E-06 |
| 17348 | ENSMUSG00000047141 | Zfp654 | 72020 [Gene Symbol: Zfp654] [Locus Tag: ] [Chromosome: 16] [Map Location: 16\|16 C | 2.431780605 | 2.62E-06 | 1.10E-05 |
| 9587 | ENSMUSG00000043259 | Fam13c | 71721 [Gene Symbol: Fam13c] [Locus Tag: ] [Chromosome: 10] [Map Location: 10\|10 B | 2.431626192 | 3.99E-07 | 1.92E-06 |
| 435 | ENSMUSG00000026712 | Mrc1 | 17533 [Gene Symbol: Mrc1] [Locus Tag: ] [Chromosome: 2] [Map Location: 2 A2\|2 10.4 | 2.431402822 | 0.000347 | 0.000987 |
| 1005 | ENSMUSG00000060475 | Wtap | 60532 [Gene Symbol: Wtap] [Locus Tag: ] [Chromosome: 17] [Map Location: 17\|17 A1] | 2.430819199 | 2.25E-07 | 1.13E-06 |
| 9810 | ENSMUSG00000024487 | Yipf5 | 67180 [Gene Symbol: Yipf5] [Locus Tag: ] [Chromosome: 18] [Map Location: 18\|18 B3] | 2.430425571 | 2.80E-05 | 9.84E-05 |
| 4909 | ENSMUSG00000023021 | Cers5 | 71949 [Gene Symbol: Cers5] [Locus Tag: ] [Chromosome: 15] [Map Location: 15\|15 F1] | 2.428957479 | 6.92E-07 | 3.20E-06 |
| 1884 | ENSMUSG00000037296 | Lsm1 | 67207 [Gene Symbol: Lsm1] [Locus Tag: ] [Chromosome: 8] [Map Location: 8\|8 A3] [Des | 2.428264031 | 2.88E-06 | 1.20E-05 |
| 5478 | ENSMUSG00000004843 | Chmp2b | 68942 [Gene Symbol: Chmp2b] [Locus Tag: ] [Chromosome: 16] [Map Location: 16\|16 | 2.425999772 | 1.81E-06 | 7.82E-06 |
| 668 | ENSMUSG00000030498 | Gas2 | 14453 [Gene Symbol: Gas2] [Locus Tag: ] [Chromosome: 7] [Map Location: 7 B5\|7 32.8 | 2.425738862 | 1.76E-05 | 6.42E-05 |
| 557 | ENSMUSG00000042029 | Ncapg2 | 76044 [Gene Symbol: Ncapg2] [Locus Tag: ] [Chromosome: 12] [Map Location: 12\|12 F | 2.425364308 | 1.89E-05 | 6.86E-05 |
| 16923 | ENSMUSG00000037111 | Setd7 | 73251 [Gene Symbol: Setd7] [Locus Tag: ] [Chromosome: 3] [Map Location: 3\|3 C] [Desc | 2.424987617 | 1.04E-06 | 4.69E-06 |
| 15858 | ENSMUSG00000063200 | Nol7 | 70078 [Gene Symbol: Nol7] [Locus Tag: ] [Chromosome: 13] [Map Location: 13\|13 A4] | 2.423872831 | 1.55E-05 | 5.72E-05 |
| 361 | ENSMUSG00000066979 | Bub3 | 12237 [Gene Symbol: Bub3] [Locus Tag: ] [Chromosome: 7] [Map Location: 7 F3\|7 74.3 | 2.423796195 | 7.68E-05 | 0.000246 |
| 2411 | ENSMUSG00000025968 | Ndufs1 | 227197 [Gene Symbol: Ndufs1] [Locus Tag: ] [Chromosome: 1] [Map Location: 1\|1 C2] [ | 2.422025912 | 8.54E-07 | 3.90E-06 |
| 4977 | ENSMUSG00000043207 | Zmpste24 | 230709 [Gene Symbol: Zmpste24] [Locus Tag: ] [Chromosome: 4] [Map Location: 4\|4 D | 2.420791654 | 7.23E-05 | 0.000233 |
| 7179 | ENSMUSG00000031641 | Cbr4 | 234309 [Gene Symbol: Cbr4] [Locus Tag: ] [Chromosome: 8] [Map Location: 8\|8 B3.1] [ | 2.420554552 | 0.013802 | 0.027741 |
| 9003 | ENSMUSG00000061119 | Prcp | 72461 [Gene Symbol: Prcp] [Locus Tag: ] [Chromosome: 7] [Map Location: 7\|7 E2] [Desc | 2.419846686 | 9.30E-05 | 0.000293 |
| 8725 | ENSMUSG00000039270 | Megf9 | 230316 [Gene Symbol: Megf9] [Locus Tag: ] [Chromosome: 4] [Map Location: 4\|4 C2] [D | 2.419501419 | 1.52E-07 | 7.86E-07 |
| 7594 | ENSMUSG00000068134 | Zfp120 | 104348 [Gene Symbol: Zfp120] [Locus Tag: ] [Chromosome: 2] [Map Location: 2\|2 G3] [ | 2.419133413 | 3.88E-05 | 0.000132 |
| 9668 | ENSMUSG00000041390 | Mdfic | 16543 [Gene Symbol: Mdfic] [Locus Tag: ] [Chromosome: 6] [Map Location: 6\|6 A1] [De | 2.419026056 | 7.27E-05 | 0.000234 |
| 2917 | ENSMUSG00000047414 | Flrt2 | 399558 [Gene Symbol: Flrt2] [Locus Tag: ] [Chromosome: 12] [Map Location: 12\|12 E] [ | 2.418838318 | 3.76E-06 | 1.53E-05 |
| 1827 | ENSMUSG00000032419 | Tbx18 | 76365 [Gene Symbol: Tbx18] [Locus Tag: ] [Chromosome: 9] [Map Location: 9 47.06 cM | 2.418518436 | 0.001042 | 0.002701 |
| 5933 | ENSMUSG00000079466 | Prdm12 | 381359 [Gene Symbol: Prdm12] [Locus Tag: ] [Chromosome: 2] [Map Location: 2\|2 B] [ | 2.417158214 | 0.003531 | 0.008138 |
| 14206 | ENSMUSG00000026577 | Blzf1 | 66352 [Gene Symbol: Blzf1] [Locus Tag: ] [Chromosome: 1] [Map Location: 1\|1 H1] [Des | 2.413704974 | 0.000158 | 0.000478 |
| 8075 | ENSMUSG00000045078 | Rnf216 | 108086 [Gene Symbol: Rnf216] [Locus Tag: ] [Chromosome: 5] [Map Location: 5\|5 G2] | 2.412485357 | 7.20E-06 | 2.80E-05 |
| 12426 | ENSMUSG00000058897 | Col25a1 | 77018 [Gene Symbol: Col25a1] [Locus Tag: ] [Chromosome: 3] [Map Location: 3\|3 H1] [ | 2.410840762 | 2.18E-05 | 7.81E-05 |
| 7335 | ENSMUSG00000031458 | Coprs | 66423 [Gene Symbol: Coprs] [Locus Tag: ] [Chromosome: 8] [Map Location: 8\|8 A1.1] [ | 2.410271234 | 2.76E-05 | 9.69E-05 |
| 16689 | ENSMUSG00000019295 | Tmem129 | 68366 [Gene Symbol: Tmem129] [Locus Tag: ] [Chromosome: 5] [Map Location: 5\|5 B2 | 2.409887668 | 4.93E-06 | 1.96E-05 |
| 7252 | ENSMUSG00000023791 | Pigx | 72084 [Gene Symbol: Pigx] [Locus Tag: ] [Chromosome: 16] [Map Location: 16\|16 B2] [ | 2.409610835 | 8.31E-05 | 0.000265 |
| 13638 | ENSMUSG00000078348 | Sf3b5 | 66125 [Gene Symbol: Sf3b5] [Locus Tag: ] [Chromosome: 10] [Map Location: 10\|10 A1] | 2.409347398 | 0.000326 | 0.000934 |
| 18115 | ENSMUSG00000032290 | Ptpn9 | 56294 [Gene Symbol: Ptpn9] [Locus Tag: ] [Chromosome: 9] [Map Location: 9\|9 C] [Des | 2.409203463 | 1.78E-07 | 9.07E-07 |
| 327 | ENSMUSG00000060733 | Ipmk | 69718 [Gene Symbol: Ipmk] [Locus Tag: ] [Chromosome: 10] [Map Location: 10\|10 B5. | 2.409166854 | 1.79E-07 | 9.10E-07 |
| 10043 | ENSMUSG00000061544 | Zfp229 | 381067 [Gene Symbol: Zfp229] [Locus Tag: ] [Chromosome: 17] [Map Location: 17\|17 | 2.408635525 | 3.02E-05 | 0.000105 |
| 265 | ENSMUSG00000003847 | Nfat5 | 54446 [Gene Symbol: Nfat5] [Locus Tag: ] [Chromosome: 8] [Map Location: 8 53.93 cM | 2.407775467 | 1.40E-07 | 7.25E-07 |
| 17365 | ENSMUSG00000093930 | Hmgcs1 | 208715 [Gene Symbol: Hmgcs1] [Locus Tag: ] [Chromosome: 13] [Map Location: 13\|13 | 2.407700642 | 0.000538 | 0.001477 |
| 14627 | ENSMUSG00000030145 | Zfp248 | 72720 [Gene Symbol: Zfp248] [Locus Tag: ] [Chromosome: 6] [Map Location: 6\|6 F1] [D | 2.407094526 | 3.13E-06 | 1.29E-05 |
| 11140 | ENSMUSG00000041272 | Tox | 252838 [Gene Symbol: Tox] [Locus Tag: ] [Chromosome: 4] [Map Location: 4\|4 A1] [Des | 2.406091701 | 6.09E-07 | 2.85E-06 |
| 15492 | ENSMUSG00000030245 | Golt1b | 66964 [Gene Symbol: Golt1b] [Locus Tag: ] [Chromosome: 6] [Map Location: 6\|6 G1] [D | 2.405884758 | 6.79E-05 | 0.00022 |
| 2897 | ENSMUSG00000031478 | Nek3 | 23954 [Gene Symbol: Nek3] [Locus Tag: ] [Chromosome: 8] [Map Location: 8 A2\|8 11.0 | 2.405210594 | 0.004093 | 0.009291 |
| 15608 | ENSMUSG00000042745 | Id1 | 15901 [Gene Symbol: Id1] [Locus Tag: ] [Chromosome: 2] [Map Location: 2 H1\|2 75.41 | 2.405174864 | 0.000709 | 0.001897 |
| 5645 | ENSMUSG00000028907 | Utp11l | 67205 [Gene Symbol: Utp11l] [Locus Tag: ] [Chromosome: 4] [Map Location: 4\|4 D1] [D | 2.404939495 | 0.000185 | 0.000554 |
| 9304 | ENSMUSG00000028484 | Psip1 | 101739 [Gene Symbol: Psip1] [Locus Tag: ] [Chromosome: 4] [Map Location: 4\|4 C3] [D | 2.404154276 | 7.51E-08 | 4.06E-07 |
| 615 | ENSMUSG00000020541 | Tom1l1 | 71943 [Gene Symbol: Tom1l1] [Locus Tag: ] [Chromosome: 11] [Map Location: 11 D\|11 | 2.404124726 | 1.75E-05 | 6.40E-05 |
| 8206 | ENSMUSG00000026727 | Rsu1 | 20163 [Gene Symbol: Rsu1] [Locus Tag: ] [Chromosome: 2] [Map Location: 2\|2 A1] [Des | 2.404104606 | 7.96E-06 | 3.08E-05 |
| 5641 | ENSMUSG00000022070 | Bora | 77744 [Gene Symbol: Bora] [Locus Tag: ] [Chromosome: 14] [Map Location: 14\|14 E2.2 | 2.403769101 | 7.63E-05 | 0.000245 |
| 9851 | ENSMUSG00000027176 | Cstf3 | 228410 [Gene Symbol: Cstf3] [Locus Tag: ] [Chromosome: 2] [Map Location: 2 E2\|2 54. | 2.403676801 | 5.17E-07 | 2.45E-06 |

| | | | | | | |
|---|---|---|---|---|---|---|
| 16411 | ENSMUSG00000037531 | Mrpl47 | 74600 [Gene Symbol: Mrpl47] [Locus Tag: ] [Chromosome: 3] [Map Location: 3\|3 B] [De | 2.403100839 | 0.000784 | 0.00208 |
| 6718 | ENSMUSG00000060988 | Galnt13 | 271786 [Gene Symbol: Galnt13] [Locus Tag: ] [Chromosome: 2] [Map Location: 2\|2 C1. | 2.402802826 | 0.000273 | 0.00079 |
| 1606 | ENSMUSG00000050545 | Fam228b | 207921 [Gene Symbol: Fam228b] [Locus Tag: ] [Chromosome: 12] [Map Location: 12\|1 | 2.402288455 | 0.001134 | 0.002922 |
| 17270 | ENSMUSG00000003623 | Crot | 74114 [Gene Symbol: Crot] [Locus Tag: ] [Chromosome: 5] [Map Location: 5\|5 A1] [Desc | 2.402020399 | 0.000363 | 0.00103 |
| 7477 | ENSMUSG00000071550 | Cfap44 | 212517 [Gene Symbol: Cfap44] [Locus Tag: ] [Chromosome: 16] [Map Location: 16 B4\|1 | 2.401094612 | 0.001252 | 0.003197 |
| 16560 | ENSMUSG00000057440 | Mpp7 | 75739 [Gene Symbol: Mpp7] [Locus Tag: ] [Chromosome: 18] [Map Location: 18\|18 A1] | 2.398719666 | 2.19E-05 | 7.84E-05 |
| 4572 | ENSMUSG00000034826 | Nup54 | 269113 [Gene Symbol: Nup54] [Locus Tag: ] [Chromosome: 5] [Map Location: 5\|5 E3] [I | 2.398491758 | 0.000164 | 0.000497 |
| 2582 | ENSMUSG00000029234 | Tmem165 | 21982 [Gene Symbol: Tmem165] [Locus Tag: ] [Chromosome: 5] [Map Location: 5 C3.3\| | 2.398287854 | 1.97E-05 | 7.12E-05 |
| 2556 | ENSMUSG00000040302 | Rbm48 | 269623 [Gene Symbol: Rbm48] [Locus Tag: ] [Chromosome: 5] [Map Location: 5\|5 A1] [ | 2.398271894 | 0.001066 | 0.002759 |
| 6185 | ENSMUSG00000090546 | Cdr1 | 631990 [Gene Symbol: Cdr1] [Locus Tag: ] [Chromosome: X] [Map Location: X A6\|X 34.: | 2.397935857 | 3.94E-06 | 1.60E-05 |
| 9299 | ENSMUSG00000026632 | Tatdn3 | 68972 [Gene Symbol: Tatdn3] [Locus Tag: ] [Chromosome: 1] [Map Location: 1\|1 H6] [D | 2.397684321 | 0.003371 | 0.007805 |
| 8883 | ENSMUSG00000021668 | Polk | 27015 [Gene Symbol: Polk] [Locus Tag: ] [Chromosome: 13] [Map Location: 13\|13 D1] [ | 2.396382561 | 0.000301 | 0.000868 |
| 2296 | ENSMUSG00000053702 | Nebl | 74103 [Gene Symbol: Nebl] [Locus Tag: ] [Chromosome: 2] [Map Location: 2\|2 A2] [Des | 2.395554247 | 7.09E-05 | 0.000229 |
| 6380 | ENSMUSG00000002477 | Snrpd1 | 20641 [Gene Symbol: Snrpd1] [Locus Tag: ] [Chromosome: 18] [Map Location: 18\|18 A: | 2.395108215 | 1.99E-05 | 7.20E-05 |
| 9155 | ENSMUSG00000038569 | Rad9b | 231724 [Gene Symbol: Rad9b] [Locus Tag: ] [Chromosome: 5] [Map Location: 5\|5 F] [De | 2.394588172 | 1.79E-05 | 6.54E-05 |
| 5106 | ENSMUSG00000027133 | Nop10 | 66181 [Gene Symbol: Nop10] [Locus Tag: ] [Chromosome: 2] [Map Location: 2\|2 E4] [D | 2.394075409 | 0.001216 | 0.003116 |
| 5521 | ENSMUSG00000030225 | Dera | 232449 [Gene Symbol: Dera] [Locus Tag: ] [Chromosome: 6] [Map Location: 6\|6 G1] [De | 2.393737914 | 0.001884 | 0.004616 |
| 3875 | ENSMUSG00000028399 | Ptprd | 19266 [Gene Symbol: Ptprd] [Locus Tag: ] [Chromosome: 4] [Map Location: 4 C3\|4 36.9 | 2.393591967 | 1.56E-06 | 6.82E-06 |
| 3046 | ENSMUSG00000020680 | Taf15 | 70439 [Gene Symbol: Taf15] [Locus Tag: ] [Chromosome: 11] [Map Location: 11\|11 B5] | 2.391632846 | 0.000143 | 0.000436 |
| 17915 | ENSMUSG00000067369 | Trmt2b | 215201 [Gene Symbol: Trmt2b] [Locus Tag: ] [Chromosome: X] [Map Location: X\|X E3] [ | 2.39132839 | 0.002699 | 0.006394 |
| 991 | ENSMUSG00000049882 | Vcpkmt; LOC102641 | 207965 [Gene Symbol: Vcpkmt] [Locus Tag: ] [Chromosome: 12] [Map Location: 12\|12 | 2.390648275 | 0.000234 | 0.000688 |
| 6232 | ENSMUSG00000050473 | Slc35d3 | 76157 [Gene Symbol: Slc35d3] [Locus Tag: ] [Chromosome: 10] [Map Location: 10\|10 A | 2.390585764 | 0.003894 | 0.00889 |
| 17436 | ENSMUSG00000055945 | Prr18 | 320111 [Gene Symbol: Prr18] [Locus Tag: ] [Chromosome: 17] [Map Location: 17\|17 A: | 2.390508095 | 4.71E-06 | 1.88E-05 |
| 17884 | ENSMUSG00000032740 | Ccdc88a | 108686 [Gene Symbol: Ccdc88a] [Locus Tag: ] [Chromosome: 11] [Map Location: 11\|11 | 2.390482284 | 0.000163 | 0.000493 |
| 11090 | ENSMUSG00000066568 | Lsm14a | 67070 [Gene Symbol: Lsm14a] [Locus Tag: ] [Chromosome: 7] [Map Location: 7\|7 B1] [I | 2.390471368 | 6.63E-07 | 3.08E-06 |
| 16908 | ENSMUSG00000032218 | Ccnb2 | 12442 [Gene Symbol: Ccnb2] [Locus Tag: ] [Chromosome: 9] [Map Location: 9\|9 D] [Des | 2.388933792 | 0.014735 | 0.029432 |
| 10176 | ENSMUSG00000028173 | Wls | 68151 [Gene Symbol: Wls] [Locus Tag: ] [Chromosome: 3] [Map Location: 3\|3 H4] [Desc | 2.386320822 | 8.68E-05 | 0.000275 |
| 17184 | ENSMUSG00000034007 | Scaper | 244891 [Gene Symbol: Scaper] [Locus Tag: ] [Chromosome: 9] [Map Location: 9\|9 B] [D | 2.385870948 | 1.88E-07 | 9.55E-07 |
| 11493 | ENSMUSG00000027797 | Dclk1 | 13175 [Gene Symbol: Dclk1] [Locus Tag: ] [Chromosome: 3] [Map Location: 3\|3 D] [Des | 2.383758867 | 1.02E-07 | 5.44E-07 |
| 8588 | ENSMUSG00000062098 | Btbd3 | 228662 [Gene Symbol: Btbd3] [Locus Tag: ] [Chromosome: 2] [Map Location: 2\|2 F3] [D | 2.383675113 | 2.36E-07 | 1.18E-06 |
| 7325 | ENSMUSG00000041438 | Cirh1a | 21771 [Gene Symbol: Cirh1a] [Locus Tag: ] [Chromosome: 8] [Map Location: 8\|8 D3] [D | 2.382620111 | 3.06E-05 | 0.000107 |
| 9325 | ENSMUSG00000027942 | 4933434E20Rik; 170 | 99650 [Gene Symbol: 4933434E20Rik] [Locus Tag: ] [Chromosome: 3] [Map Location: 3 | 2.381766587 | 2.04E-05 | 7.35E-05 |
| 7443 | ENSMUSG00000037072 | Sep15 | 93684 [Gene Symbol: Sep15] [Locus Tag: ] [Chromosome: 3] [Map Location: 3\|3 H3] [De | 2.381497716 | 1.99E-06 | 8.54E-06 |
| 208 | ENSMUSG00000028218 | Fam92a | 68099 [Gene Symbol: Fam92a] [Locus Tag: ] [Chromosome: 4] [Map Location: 4\|4 A2] [ | 2.380407947 | 5.37E-07 | 2.53E-06 |
| 2111 | ENSMUSG00000034071 | Zfp551 | 619331 [Gene Symbol: Zfp551] [Locus Tag: ] [Chromosome: 7] [Map Location: 7\|7 A1] [ | 2.380077069 | 2.75E-05 | 9.66E-05 |
| 5572 | ENSMUSG00000050891 | Tatdn1 | 69694 [Gene Symbol: Tatdn1] [Locus Tag: ] [Chromosome: 15] [Map Location: 15\|15 D: | 2.379971199 | 2.29E-07 | 1.15E-06 |
| 1790 | ENSMUSG00000055200 | Sertad3 | 170742 [Gene Symbol: Sertad3] [Locus Tag: ] [Chromosome: 7] [Map Location: 7\|7 A3] | 2.379322655 | 0.005912 | 0.012951 |
| 12912 | ENSMUSG00000016833 | Mrps18c | 68735 [Gene Symbol: Mrps18c] [Locus Tag: ] [Chromosome: 5] [Map Location: 5\|5 E3] [ | 2.377900638 | 0.004077 | 0.009263 |
| 2633 | ENSMUSG00000062101 | Zfp119b | 240120 [Gene Symbol: Zfp119b] [Locus Tag: ] [Chromosome: 17] [Map Location: 17\|17 | 2.377303016 | 0.020804 | 0.039906 |
| 13238 | ENSMUSG00000000296 | Tpd52l1 | 21987 [Gene Symbol: Tpd52l1] [Locus Tag: ] [Chromosome: 10] [Map Location: 10\|10 A | 2.377013096 | 0.003383 | 0.00783 |
| 2460 | ENSMUSG00000034674 | Tdg; Tdg-ps | 21665 [Gene Symbol: Tdg] [Locus Tag: ] [Chromosome: 10] [Map Location: 10 C1\|10 39 | 2.375874291 | 4.88E-05 | 0.000163 |
| 10692 | ENSMUSG00000032570 | Atp2c1 | 235574 [Gene Symbol: Atp2c1] [Locus Tag: ] [Chromosome: 9] [Map Location: 9\|9 F1] [ | 2.374932601 | 1.78E-07 | 9.06E-07 |
| 9073 | ENSMUSG00000025578 | Cbx8 | 30951 [Gene Symbol: Cbx8] [Locus Tag: ] [Chromosome: 11] [Map Location: 11\|11 E2] [ | 2.374719793 | 3.54E-05 | 0.000122 |
| 4633 | ENSMUSG00000026890 | Lhx6 | 16874 [Gene Symbol: Lhx6] [Locus Tag: ] [Chromosome: 2] [Map Location: 2\|2 B] [Descr | 2.374567574 | 1.85E-06 | 7.99E-06 |
| 1633 | ENSMUSG00000020290 | Xpo1 | 103573 [Gene Symbol: Xpo1] [Locus Tag: ] [Chromosome: 11] [Map Location: 11\|11 A3 | 2.374485742 | 8.15E-07 | 3.73E-06 |
| 7242 | ENSMUSG00000029638 | Glcci1 | 170772 [Gene Symbol: Glcci1] [Locus Tag: ] [Chromosome: 6] [Map Location: 6\|6 A1] [I | 2.374163448 | 3.94E-07 | 1.90E-06 |
| 11945 | ENSMUSG00000022364 | Tbc1d31 | 210544 [Gene Symbol: Tbc1d31] [Locus Tag: ] [Chromosome: 15] [Map Location: 15\|1! | 2.373295365 | 0.001529 | 0.003824 |
| 16137 | ENSMUSG00000069670 | Nkain2 | 432450 [Gene Symbol: Nkain2] [Locus Tag: ] [Chromosome: 10] [Map Location: 10\|10 A | 2.37278951 | 2.21E-06 | 9.40E-06 |
| 12265 | ENSMUSG00000095115 | Itpripl2 | 319622 [Gene Symbol: Itpripl2] [Locus Tag: ] [Chromosome: 7] [Map Location: 7\|7 F1] | 2.372506988 | 3.57E-05 | 0.000123 |
| 5548 | ENSMUSG00000021848 | Otx2 | 18424 [Gene Symbol: Otx2] [Locus Tag: ] [Chromosome: 14] [Map Location: 14 C1\|14 2 | 2.371644121 | 0.003037 | 0.007112 |
| 1090 | ENSMUSG00000056216 | Cebpg | 12611 [Gene Symbol: Cebpg] [Locus Tag: ] [Chromosome: 7] [Map Location: 7\|7 B1] [De | 2.370727396 | 4.47E-05 | 0.00015 |
| 3877 | ENSMUSG00000038257 | Glra3 | 110304 [Gene Symbol: Glra3] [Locus Tag: ] [Chromosome: 8] [Map Location: 8 B2\|8 29. | 2.369258295 | 0.005076 | 0.011292 |
| 4410 | ENSMUSG00000007617 | Homer1 | 26556 [Gene Symbol: Homer1] [Locus Tag: ] [Chromosome: 13] [Map Location: 13\|13 C | 2.368248229 | 2.44E-06 | 1.03E-05 |
| 13964 | ENSMUSG00000031657 | Heatr3 | 234549 [Gene Symbol: Heatr3] [Locus Tag: ] [Chromosome: 8] [Map Location: 8\|8 C3] [ | 2.367995885 | 1.02E-05 | 3.88E-05 |
| 15679 | ENSMUSG00000031924 | Cyb5b | 66427 [Gene Symbol: Cyb5b] [Locus Tag: ] [Chromosome: 8] [Map Location: 8\|8 D2] [D | 2.367558793 | 3.88E-07 | 1.87E-06 |
| 4884 | ENSMUSG00000026678 | Rgs5 | 19737 [Gene Symbol: Rgs5] [Locus Tag: ] [Chromosome: 1] [Map Location: 1 76.84 cM\| | 2.366215663 | 9.34E-06 | 3.56E-05 |
| 11387 | ENSMUSG00000028223 | Decr1 | 67460 [Gene Symbol: Decr1] [Locus Tag: ] [Chromosome: 4] [Map Location: 4\|4 A2] [De | 2.366076256 | 0.002592 | 0.006162 |
| 13762 | ENSMUSG00000045589 | Frrs1l | 230235 [Gene Symbol: Frrs1l] [Locus Tag: ] [Chromosome: 4] [Map Location: 4\|4 B3] [D | 2.363978445 | 1.32E-07 | 6.89E-07 |
| 12116 | ENSMUSG00000025423 | Pias2 | 17344 [Gene Symbol: Pias2] [Locus Tag: ] [Chromosome: 18] [Map Location: 18\|18 E3] | 2.363543135 | 2.81E-07 | 1.38E-06 |
| 3303 | ENSMUSG00000051950 | B3glct | 381694 [Gene Symbol: B3glct] [Locus Tag: ] [Chromosome: 5] [Map Location: 5\|5 G3] [ | 2.363072094 | 2.17E-06 | 9.21E-06 |
| 15317 | ENSMUSG00000027273 | Snap25 | 20614 [Gene Symbol: Snap25] [Locus Tag: ] [Chromosome: 2] [Map Location: 2 F3\|2 67 | 2.362359945 | 1.72E-06 | 7.49E-06 |
| 8072 | ENSMUSG00000061666 | Gdpd1 | 66569 [Gene Symbol: Gdpd1] [Locus Tag: ] [Chromosome: 11] [Map Location: 11\|11 C] | 2.361814453 | 2.42E-06 | 1.02E-05 |
| 3854 | ENSMUSG00000031299 | Pdha1 | 18597 [Gene Symbol: Pdha1] [Locus Tag: ] [Chromosome: X] [Map Location: X 73.95 cN | 2.361739782 | 8.88E-07 | 4.04E-06 |
| 9554 | ENSMUSG00000046185 | Zfp84 | 74352 [Gene Symbol: Zfp84] [Locus Tag: ] [Chromosome: 7] [Map Location: 7 B1\|7 17.2 | 2.361359473 | 4.19E-05 | 0.000142 |
| 5473 | ENSMUSG00000042202 | Slc35e2 | 320541 [Gene Symbol: Slc35e2] [Locus Tag: ] [Chromosome: 4] [Map Location: 4\|4 E2] | 2.36099737 | 5.82E-07 | 2.73E-06 |
| 17967 | ENSMUSG00000035933 | Cog5 | 238123 [Gene Symbol: Cog5] [Locus Tag: ] [Chromosome: 12] [Map Location: 12\|12 A2 | 2.3597344 | 4.71E-07 | 2.24E-06 |
| 8741 | ENSMUSG00000021619 | Atg10 | 66795 [Gene Symbol: Atg10] [Locus Tag: ] [Chromosome: 13] [Map Location: 13\|13 C3] | 2.358527421 | 0.000165 | 0.000499 |
| 3857 | ENSMUSG00000029397 | Rchy1 | 68098 [Gene Symbol: Rchy1] [Locus Tag: ] [Chromosome: 5] [Map Location: 5\|5 E2] [De | 2.358085683 | 4.95E-06 | 1.97E-05 |
| 9211 | ENSMUSG00000034320 | Slc26a2 | 13521 [Gene Symbol: Slc26a2] [Locus Tag: ] [Chromosome: 18] [Map Location: 18\|18 D | 2.357936419 | 1.40E-06 | 6.18E-06 |
| 15106 | ENSMUSG00000034875 | Nudt19 | 110959 [Gene Symbol: Nudt19] [Locus Tag: ] [Chromosome: 7] [Map Location: 7 B2\|7 2 | 2.357804549 | 6.98E-06 | 2.72E-05 |
| 471 | ENSMUSG00000039717 | Ralyl | 76897 [Gene Symbol: Ralyl] [Locus Tag: ] [Chromosome: 3] [Map Location: 3\|3 A1] [Des | 2.350482081 | 9.83E-05 | 0.000309 |
| 12187 | ENSMUSG00000024479 | Mal2 | 105853 [Gene Symbol: Mal2] [Locus Tag: ] [Chromosome: 15] [Map Location: 15\|15 D1 | 2.350236363 | 0.000723 | 0.001932 |
| 3127 | ENSMUSG00000046178 | Nxph1 | 18231 [Gene Symbol: Nxph1] [Locus Tag: ] [Chromosome: 6] [Map Location: 6\|6 A1] [D | 2.349830123 | 2.99E-05 | 0.000104 |
| 15432 | ENSMUSG00000032179 | Bmp5 | 12160 [Gene Symbol: Bmp5] [Locus Tag: ] [Chromosome: 9] [Map Location: 9 D\|9 42.3 | 2.349387465 | 0.001 | 0.002598 |
| 10753 | ENSMUSG00000068742 | Cry2 | 12953 [Gene Symbol: Cry2] [Locus Tag: ] [Chromosome: 2] [Map Location: 2\|2 E] [Descr | 2.348959253 | 1.09E-06 | 4.90E-06 |
| 2495 | ENSMUSG00000041408 | Wapl | 218914 [Gene Symbol: Wapl] [Locus Tag: ] [Chromosome: 14] [Map Location: 14\|14 B] | 2.348905051 | 2.69E-07 | 1.33E-06 |
| 955 | ENSMUSG00000020530 | Ggnbp2 | 217039 [Gene Symbol: Ggnbp2] [Locus Tag: ] [Chromosome: 11] [Map Location: 11\|11 | 2.348163371 | 7.89E-05 | 0.000253 |
| 12075 | ENSMUSG00000020621 | Rdh14 | 105014 [Gene Symbol: Rdh14] [Locus Tag: ] [Chromosome: 12] [Map Location: 12\|12 A | 2.348126807 | 0.000317 | 0.000908 |
| 13648 | ENSMUSG00000044715 | Gskip | 66787 [Gene Symbol: Gskip] [Locus Tag: ] [Chromosome: 12] [Map Location: 12\|12 E] [I | 2.347766142 | 8.40E-05 | 0.000267 |
| 3930 | ENSMUSG00000072258 | Taf1a | 21339 [Gene Symbol: Taf1a] [Locus Tag: ] [Chromosome: 1] [Map Location: 1\|1] [Descri | 2.34684352 | 0.002847 | 0.006711 |
| 5297 | ENSMUSG00000051989 | Smim11 | 68936 [Gene Symbol: Smim11] [Locus Tag: ] [Chromosome: 16] [Map Location: 16\|16 C | 2.346401785 | 0.006889 | 0.014862 |
| 12535 | ENSMUSG00000024098 | Twsg1 | 65960 [Gene Symbol: Twsg1] [Locus Tag: ] [Chromosome: 17] [Map Location: 17 E1.1\|1 | 2.344549144 | 2.19E-06 | 9.29E-06 |
| 12319 | ENSMUSG00000074088 | Snrnp40 | 66585 [Gene Symbol: Snrnp40] [Locus Tag: ] [Chromosome: 4] [Map Location: 4\|4 D3] [ | 2.344334435 | 2.40E-05 | 8.53E-05 |
| 2566 | ENSMUSG00000034303 | Ccdc15 | 245902 [Gene Symbol: Ccdc15] [Locus Tag: ] [Chromosome: 9] [Map Location: 9\|9 A4] [ | 2.341178618 | 5.33E-05 | 0.000176 |

| | | | | | | |
|---|---|---|---|---|---|---|
| 12958 | ENSMUSG00000022452 | Smdt1 | 69029 [Gene Symbol: Smdt1] [Locus Tag: ] [Chromosome: 15] [Map Location: 15\|15 E2 | 2.340445202 | 0.000267 | 0.000776 |
| 11934 | ENSMUSG00000072770 | Acrbp | 54137 [Gene Symbol: Acrbp] [Locus Tag: ] [Chromosome: 6] [Map Location: 6\|6 F2] [De | 2.339661315 | 0.003778 | 0.008652 |
| 9305 | ENSMUSG00000044763 | Trmt10c | 52575 [Gene Symbol: Trmt10c] [Locus Tag: ] [Chromosome: 16] [Map Location: 16 C1.1 | 2.339609369 | 7.47E-05 | 0.00024 |
| 1314 | ENSMUSG00000029910 | Mad2l1 | 56150 [Gene Symbol: Mad2l1] [Locus Tag: ] [Chromosome: 6] [Map Location: 6 C1\|6 30 | 2.339405054 | 0.000544 | 0.001491 |
| 14272 | ENSMUSG00000000402 | Egfl6 | 54156 [Gene Symbol: Egfl6] [Locus Tag: ] [Chromosome: X] [Map Location: X F5\|X 77.6 | 2.339328111 | 0.020987 | 0.040216 |
| 6993 | ENSMUSG00000031617 | Tmem184c | 234463 [Gene Symbol: Tmem184c] [Locus Tag: ] [Chromosome: 8] [Map Location: 8\|8 | 2.339069137 | 7.56E-07 | 3.48E-06 |
| 6638 | ENSMUSG00000028292 | Rars2 | 109093 [Gene Symbol: Rars2] [Locus Tag: ] [Chromosome: 4] [Map Location: 4\|4 A5] [D | 2.338091698 | 2.67E-05 | 9.40E-05 |
| 4223 | ENSMUSG00000020883 | Fbxl20 | 72194 [Gene Symbol: Fbxl20] [Locus Tag: ] [Chromosome: 11] [Map Location: 11\|11 D] | 2.337844611 | 1.50E-06 | 6.59E-06 |
| 15433 | ENSMUSG00000057234 | Mettl15 | 76894 [Gene Symbol: Mettl15] [Locus Tag: ] [Chromosome: 2] [Map Location: 2\|2 E3] [ | 2.336745531 | 0.00072 | 0.001925 |
| 3030 | ENSMUSG00000027963 | Extl2 | 58193 [Gene Symbol: Extl2] [Locus Tag: ] [Chromosome: 3] [Map Location: 3\|3 G1] [De | 2.336574117 | 4.42E-06 | 1.78E-05 |
| 8249 | ENSMUSG00000037640 | Zfp60 | 22718 [Gene Symbol: Zfp60] [Locus Tag: ] [Chromosome: 7] [Map Location: 7 A3\|7 15.9 | 2.33652348 | 5.52E-05 | 0.000182 |
| 14147 | ENSMUSG00000053390 | Zfp952 | 240067 [Gene Symbol: Zfp952] [Locus Tag: ] [Chromosome: 17] [Map Location: 17\|17 | 2.335974279 | 0.000283 | 0.000818 |
| 4072 | ENSMUSG00000075415 | Fnbp1 | 14269 [Gene Symbol: Fnbp1] [Locus Tag: ] [Chromosome: 2] [Map Location: 2\|2 B] [De | 2.335635577 | 0.000451 | 0.001257 |
| 9683 | ENSMUSG00000029111 | Nelfa | 24116 [Gene Symbol: Nelfa] [Locus Tag: ] [Chromosome: 5] [Map Location: 5\|5 B2] [De | 2.333849917 | 8.49E-06 | 3.27E-05 |
| 16548 | ENSMUSG00000021243 | Fcf1 | 73736 [Gene Symbol: Fcf1] [Locus Tag: ] [Chromosome: 12] [Map Location: 12\|12 D3] [ | 2.333842261 | 0.011673 | 0.023927 |
| 9737 | ENSMUSG00000026730 | Pter | 19212 [Gene Symbol: Pter] [Locus Tag: ] [Chromosome: 2] [Map Location: 2\|2 A1] [Des | 2.33324857 | 9.38E-05 | 0.000295 |
| 4056 | ENSMUSG00000042596 | Tfap2d | 226896 [Gene Symbol: Tfap2d] [Locus Tag: ] [Chromosome: 1] [Map Location: 1\|1 A3] [ | 2.333010237 | 0.000314 | 0.0009 |
| 1379 | ENSMUSG00000013701 | Timm23 | 53600 [Gene Symbol: Timm23] [Locus Tag: ] [Chromosome: 14] [Map Location: 14\|14 | 2.332817731 | 0.002799 | 0.006608 |
| 16917 | ENSMUSG00000037315 | Jade3 | 382207 [Gene Symbol: Jade3] [Locus Tag: ] [Chromosome: X] [Map Location: X\|X A1.3] | 2.33171131 | 3.93E-06 | 1.59E-05 |
| 15590 | ENSMUSG00000024150 | Mcfd2 | 193813 [Gene Symbol: Mcfd2] [Locus Tag: ] [Chromosome: 17] [Map Location: 17\|17 E | 2.330860134 | 5.64E-05 | 0.000185 |
| 12745 | ENSMUSG00000027669 | Gnb4 | 14696 [Gene Symbol: Gnb4] [Locus Tag: ] [Chromosome: 3] [Map Location: 3\|3 B] [Desc | 2.330837662 | 1.21E-06 | 5.40E-06 |
| 17452 | ENSMUSG00000060739 | LOC102642514; Nsa2 | 102642514 [Gene Symbol: LOC102642514] [Locus Tag: ] [Chromosome: 9] [Map Locati | 2.330093344 | 0.003487 | 0.008044 |
| 6757 | ENSMUSG00000020590 | Snx13 | 217463 [Gene Symbol: Snx13] [Locus Tag: ] [Chromosome: 12] [Map Location: 12\|12 A | 2.32951928 | 6.30E-07 | 2.94E-06 |
| 14288 | ENSMUSG00000071281 | Zfp65 | 235907 [Gene Symbol: Zfp65] [Locus Tag: ] [Chromosome: 13] [Map Location: 13\|13 34 | 2.328998223 | 8.34E-05 | 0.000266 |
| 7685 | ENSMUSG00000024805 | Pcgf5 | 76073 [Gene Symbol: Pcgf5] [Locus Tag: ] [Chromosome: 19] [Map Location: 19\|19 C2] | 2.327634087 | 0.001007 | 0.002615 |
| 16537 | ENSMUSG00000062510 | Nsl1 | 381318 [Gene Symbol: Nsl1] [Locus Tag: ] [Chromosome: 1] [Map Location: 1\|1 H6] [De | 2.327050612 | 0.022817 | 0.04333 |
| 5640 | ENSMUSG00000020859 | Spag9 | 70834 [Gene Symbol: Spag9] [Locus Tag: ] [Chromosome: 11] [Map Location: 11\|11 C] [ | 2.32630331 | 3.42E-07 | 1.66E-06 |
| 3102 | ENSMUSG00000031584 | Gsr | 14782 [Gene Symbol: Gsr] [Locus Tag: ] [Chromosome: 8] [Map Location: 8 A4\|8 20.69 | 2.324816393 | 3.28E-06 | 1.35E-05 |
| 15837 | ENSMUSG00000028517 | Plpp3 | 67916 [Gene Symbol: Plpp3] [Locus Tag: ] [Chromosome: 4] [Map Location: 4 C6\|4 49. | 2.324210429 | 1.00E-05 | 3.80E-05 |
| 13856 | ENSMUSG00000030935 | Acsm3 | 20216 [Gene Symbol: Acsm3] [Locus Tag: ] [Chromosome: 7] [Map Location: 7\|7 F3] [De | 2.323726324 | 0.001211 | 0.003104 |
| 10348 | ENSMUSG00000030614 | Tmem126b | 68472 [Gene Symbol: Tmem126b] [Locus Tag: ] [Chromosome: 7] [Map Location: 7\|7 E | 2.323538392 | 0.000775 | 0.002058 |
| 15029 | ENSMUSG00000032872 | Cyb5r4 | 266690 [Gene Symbol: Cyb5r4] [Locus Tag: ] [Chromosome: 9] [Map Location: 9\|9 E3.1 | 2.322955269 | 3.62E-06 | 1.48E-05 |
| 14730 | ENSMUSG00000055296 | Tmem245 | 242474 [Gene Symbol: Tmem245] [Locus Tag: ] [Chromosome: 4] [Map Location: 4\|4 B | 2.321913297 | 1.03E-06 | 4.63E-06 |
| 4249 | ENSMUSG00000002733 | Plekha3 | 83435 [Gene Symbol: Plekha3] [Locus Tag: ] [Chromosome: 2] [Map Location: 2\|2 D] [D | 2.320659818 | 1.93E-05 | 6.99E-05 |
| 6655 | ENSMUSG00000040250 | Asun | 71177 [Gene Symbol: Asun] [Locus Tag: ] [Chromosome: 6] [Map Location: 6\|6 G3] [Des | 2.319989672 | 2.01E-05 | 7.26E-05 |
| 13685 | ENSMUSG00000046667 | Rbm12b2; Rbm12b1 | 77604 [Gene Symbol: Rbm12b2] [Locus Tag: ] [Chromosome: 4] [Map Location: 4\|4 A1 | 2.319966165 | 0.002108 | 0.005107 |
| 9096 | ENSMUSG00000054814 | Usp46 | 69727 [Gene Symbol: Usp46] [Locus Tag: ] [Chromosome: 5] [Map Location: 5\|5 C3.3] [ | 2.319695722 | 5.26E-07 | 2.48E-06 |
| 3296 | ENSMUSG00000023460 | Rab12 | 19328 [Gene Symbol: Rab12] [Locus Tag: ] [Chromosome: 17] [Map Location: 17\|17 E1 | 2.31868695 | 8.79E-06 | 3.37E-05 |
| 10053 | ENSMUSG00000047881 | Rell1 | 100532 [Gene Symbol: Rell1] [Locus Tag: ] [Chromosome: 5] [Map Location: 5\|5 C3.1] [ | 2.31809986 | 0.000121 | 0.000373 |
| 10586 | ENSMUSG00000074748 | Atxn7l3b | 382423 [Gene Symbol: Atxn7l3b] [Locus Tag: ] [Chromosome: 10] [Map Location: 10\|1 | 2.317419604 | 5.41E-07 | 2.55E-06 |
| 17836 | ENSMUSG00000023967 | Mrps18a | 68565 [Gene Symbol: Mrps18a] [Locus Tag: ] [Chromosome: 17] [Map Location: 17\|17 | 2.316761809 | 0.002433 | 0.005821 |
| 4051 | ENSMUSG00000051235 | Gen1 | 209334 [Gene Symbol: Gen1] [Locus Tag: ] [Chromosome: 12] [Map Location: 12\|12 A1 | 2.316082558 | 0.011491 | 0.02359 |
| 17005 | ENSMUSG00000037892 | Pcdh18 | 73173 [Gene Symbol: Pcdh18] [Locus Tag: ] [Chromosome: 3] [Map Location: 3\|3 C] [De | 2.315291827 | 5.15E-06 | 2.05E-05 |
| 16780 | ENSMUSG00000019880 | Rspo3 | 72780 [Gene Symbol: Rspo3] [Locus Tag: ] [Chromosome: 10] [Map Location: 10\|10 A4 | 2.312713508 | 0.007594 | 0.016248 |
| 1973 | ENSMUSG00000016921 | Srsf6 | 67996 [Gene Symbol: Srsf6] [Locus Tag: ] [Chromosome: 2] [Map Location: 2 H2\|2 83.8 | 2.312023189 | 2.11E-06 | 9.00E-06 |
| 11293 | ENSMUSG00000020715 | Ern1 | 78943 [Gene Symbol: Ern1] [Locus Tag: ] [Chromosome: 11] [Map Location: 11\|11 E1] [ | 2.309221718 | 4.95E-05 | 0.000165 |
| 15965 | ENSMUSG00000056153 | Socs6 | 54607 [Gene Symbol: Socs6] [Locus Tag: ] [Chromosome: 18] [Map Location: 18\|18 E4] | 2.309012774 | 1.58E-06 | 6.91E-06 |
| 1674 | ENSMUSG00000058331 | Zfp85 | 22746 [Gene Symbol: Zfp85] [Locus Tag: ] [Chromosome: 13] [Map Location: 13 B3\|13 | 2.308311571 | 0.000666 | 0.00179 |
| 13376 | ENSMUSG00000075334 | Rprm | 67874 [Gene Symbol: Rprm] [Locus Tag: ] [Chromosome: 2] [Map Location: 2\|2 C1] [De | 2.307714525 | 8.55E-07 | 3.90E-06 |
| 18014 | ENSMUSG00000020297 | Nsg2 | 18197 [Gene Symbol: Nsg2] [Locus Tag: ] [Chromosome: 11] [Map Location: 11\|11 A4] | 2.30730495 | 1.00E-05 | 3.81E-05 |
| 4979 | ENSMUSG00000059486 | Kbtbd2 | 210973 [Gene Symbol: Kbtbd2] [Locus Tag: ] [Chromosome: 6] [Map Location: 6\|6 B3] | 2.306993272 | 1.33E-06 | 5.91E-06 |
| 10283 | ENSMUSG00000027792 | Bche | 12038 [Gene Symbol: Bche] [Locus Tag: ] [Chromosome: 3] [Map Location: 3\|3 E3] [Des | 2.306799574 | 0.003822 | 0.008741 |
| 6783 | ENSMUSG00000051469 | Zfp24 | 59057 [Gene Symbol: Zfp24] [Locus Tag: ] [Chromosome: 18] [Map Location: 18\|18 B1] | 2.306777839 | 1.13E-05 | 4.26E-05 |
| 6903 | ENSMUSG00000026768 | Itga8 | 241226 [Gene Symbol: Itga8] [Locus Tag: ] [Chromosome: 2] [Map Location: 2 A1\|2 9.1 | 2.306627964 | 0.001085 | 0.002804 |
| 9334 | ENSMUSG00000047221 | Fam185a | 330050 [Gene Symbol: Fam185a] [Locus Tag: ] [Chromosome: 5] [Map Location: 5\|5 A3 | 2.306511428 | 8.63E-05 | 0.000274 |
| 3639 | ENSMUSG00000078307 | AI593442 | 330941 [Gene Symbol: AI593442] [Locus Tag: ] [Chromosome: 9] [Map Location: 9\|9 A | 2.305000619 | 8.29E-05 | 0.000264 |
| 12661 | ENSMUSG00000022971 | Ifnar2 | 15976 [Gene Symbol: Ifnar2] [Locus Tag: ] [Chromosome: 16] [Map Location: 16 C3.3\|1 | 2.304889243 | 0.000164 | 0.000496 |
| 749 | ENSMUSG00000000711 | Rab5b | 19344 [Gene Symbol: Rab5b] [Locus Tag: ] [Chromosome: 10] [Map Location: 10\|10 D3 | 2.30407879 | 2.83E-06 | 1.18E-05 |
| 17160 | ENSMUSG00000032409 | Atr | 245000 [Gene Symbol: Atr] [Locus Tag: ] [Chromosome: 9] [Map Location: 9\|9 E4] [Desc | 2.302962957 | 6.10E-05 | 0.000199 |
| 8701 | ENSMUSG00000033102 | Cdc14b | 218294 [Gene Symbol: Cdc14b] [Locus Tag: ] [Chromosome: 13] [Map Location: 13\|13 | 2.301452561 | 0.000674 | 0.00181 |
| 6928 | ENSMUSG00000030850 | Ate1 | 11907 [Gene Symbol: Ate1] [Locus Tag: ] [Chromosome: 7] [Map Location: 7\|7 F3] [Desc | 2.300897895 | 1.70E-06 | 7.39E-06 |
| 10544 | ENSMUSG00000063632 | Sox11 | 20666 [Gene Symbol: Sox11] [Locus Tag: ] [Chromosome: 12] [Map Location: 12\|12 A3 | 2.30054746 | 0.009688 | 0.020245 |
| 9475 | ENSMUSG00000042705 | Commd10 | 69456 [Gene Symbol: Commd10] [Locus Tag: ] [Chromosome: 18] [Map Location: 18\|1 | 2.300417758 | 4.55E-05 | 0.000153 |
| 16827 | ENSMUSG00000071037 | Camkmt | 73582 [Gene Symbol: Camkmt] [Locus Tag: ] [Chromosome: 17] [Map Location: 17\|17 | 2.298662023 | 5.03E-05 | 0.000167 |
| 18189 | ENSMUSG00000020074 | Ccar1 | 67500 [Gene Symbol: Ccar1] [Locus Tag: ] [Chromosome: 10] [Map Location: 10\|10 B4] | 2.29823585 | 3.45E-05 | 0.000119 |
| 13425 | ENSMUSG00000039246 | Lyplal1 | 226791 [Gene Symbol: Lyplal1] [Locus Tag: ] [Chromosome: 1] [Map Location: 1\|1 H5] | 2.29762176 | 0.00232 | 0.005569 |
| 10653 | ENSMUSG00000019303 | Psmc3ip | 19183 [Gene Symbol: Psmc3ip] [Locus Tag: ] [Chromosome: 11] [Map Location: 11\|11 | 2.297370818 | 0.004454 | 0.01003 |
| 4815 | ENSMUSG00000020397 | Med7 | 66213 [Gene Symbol: Med7] [Locus Tag: ] [Chromosome: 11] [Map Location: 11\|11 B1. | 2.29651468 | 0.000217 | 0.000641 |
| 2235 | ENSMUSG00000026594 | Ralgps2 | 78255 [Gene Symbol: Ralgps2] [Locus Tag: ] [Chromosome: 1] [Map Location: 1\|1 H1] [ | 2.296257375 | 5.83E-07 | 2.73E-06 |
| 2898 | ENSMUSG00000066043 | Phactr4 | 100169 [Gene Symbol: Phactr4] [Locus Tag: ] [Chromosome: 4] [Map Location: 4\|4 D2. | 2.29528206 | 9.06E-05 | 0.000286 |
| 8606 | ENSMUSG00000031647 | Mfap3l | 71306 [Gene Symbol: Mfap3l] [Locus Tag: ] [Chromosome: 8] [Map Location: 8\|8 B3.2] | 2.29452918 | 0.001091 | 0.002817 |
| 13241 | ENSMUSG00000042369 | Rbm45 | 241490 [Gene Symbol: Rbm45] [Locus Tag: ] [Chromosome: 2] [Map Location: 2\|2 C3] [ | 2.294501605 | 3.21E-05 | 0.000111 |
| 16599 | ENSMUSG00000036469 | March1 | 72925 [Gene Symbol: March1] [Locus Tag: ] [Chromosome: 8] [Map Location: 8\|8 B3.1- | 2.294336483 | 8.09E-05 | 0.000258 |
| 9614 | ENSMUSG00000030201 | Lrp6 | 16974 [Gene Symbol: Lrp6] [Locus Tag: ] [Chromosome: 6] [Map Location: 6 G1\|6 65.3 | 2.292923751 | 1.35E-05 | 5.03E-05 |
| 13055 | ENSMUSG00000024384 | Iws1 | 73473 [Gene Symbol: Iws1] [Locus Tag: ] [Chromosome: 18] [Map Location: 18\|18 B3] [ | 2.292914147 | 8.47E-06 | 3.26E-05 |
| 1492 | ENSMUSG00000072872 | Rybp | 56353 [Gene Symbol: Rybp] [Locus Tag: ] [Chromosome: 6] [Map Location: 6\|6 E1] [Des | 2.292906098 | 2.99E-06 | 1.24E-05 |
| 3384 | ENSMUSG00000034501 | Pcnxl4 | 67708 [Gene Symbol: Pcnxl4] [Locus Tag: ] [Chromosome: 12] [Map Location: 12\|12 C3 | 2.288648022 | 1.39E-06 | 6.14E-06 |
| 3598 | ENSMUSG00000029878 | Dbpht2 | 386753 [Gene Symbol: Dbpht2] [Locus Tag: ] [Chromosome: 12] [Map Location: 12\|12 | 2.287384282 | 0.000147 | 0.000448 |
| 12058 | ENSMUSG00000038393 | Txnip | 56338 [Gene Symbol: Txnip] [Locus Tag: ] [Chromosome: 3] [Map Location: 3 F2.2\|3 41 | 2.286444183 | 4.36E-06 | 1.76E-05 |
| 9924 | ENSMUSG00000048371 | Pdp2 | 382051 [Gene Symbol: Pdp2] [Locus Tag: ] [Chromosome: 8] [Map Location: 8\|8 D3] [D | 2.284927992 | 2.77E-05 | 9.73E-05 |
| 13308 | ENSMUSG00000022053 | Ebf2 | 13592 [Gene Symbol: Ebf2] [Locus Tag: ] [Chromosome: 14] [Map Location: 14\|14 D-E1 | 2.284179753 | 7.17E-05 | 0.000231 |
| 1437 | ENSMUSG00000027996 | Sfrp2 | 20319 [Gene Symbol: Sfrp2] [Locus Tag: ] [Chromosome: 3] [Map Location: 3 E3\|3 37.3 | 2.283101403 | 2.67E-05 | 9.40E-05 |
| 9385 | ENSMUSG00000023912 | Slc25a27 | 74011 [Gene Symbol: Slc25a27] [Locus Tag: ] [Chromosome: 17] [Map Location: 17\|17 | 2.282430942 | 2.72E-06 | 1.14E-05 |
| 8154 | ENSMUSG00000024277 | Mapre2 | 212307 [Gene Symbol: Mapre2] [Locus Tag: ] [Chromosome: 18] [Map Location: 18 A2\| | 2.282395432 | 2.56E-06 | 1.07E-05 |

| | | | | | | |
|---|---|---|---|---|---|---|
| 14978 | ENSMUSG00000001467 | Cyp51 | 13121 [Gene Symbol: Cyp51] [Locus Tag: ] [Chromosome: 5] [Map Location: 5 2.3 cM\|5 | 2.281814321 | 2.62E-06 | 1.10E-05 |
| 1287 | ENSMUSG00000054484 | Tmem62 | 96957 [Gene Symbol: Tmem62] [Locus Tag: ] [Chromosome: 2] [Map Location: 2\|2 E5] [ | 2.281302733 | 0.000152 | 0.000464 |
| 11650 | ENSMUSG00000018433 | Nol11 | 68979 [Gene Symbol: Nol11] [Locus Tag: ] [Chromosome: 11] [Map Location: 11\|11 E1 | 2.280858364 | 6.07E-05 | 0.000198 |
| 8135 | ENSMUSG00000025337 | Sbds | 66711 [Gene Symbol: Sbds] [Locus Tag: ] [Chromosome: 5] [Map Location: 5\|5 G1.3] [D | 2.280614243 | 0.000137 | 0.00042 |
| 98 | ENSMUSG00000060336 | Zfp937 | 245174 [Gene Symbol: Zfp937] [Locus Tag: ] [Chromosome: 2] [Map Location: 2\|2 G3] [ | 2.280517853 | 0.017546 | 0.034289 |
| 6371 | ENSMUSG00000044968 | Napepld | 242864 [Gene Symbol: Napepld] [Locus Tag: ] [Chromosome: 5] [Map Location: 5\|5 A3] | 2.279852814 | 0.006944 | 0.01497 |
| 13246 | ENSMUSG00000034462 | Pkd2 | 18764 [Gene Symbol: Pkd2] [Locus Tag: ] [Chromosome: 5] [Map Location: 5 E5\|5 50.6 | 2.279478809 | 1.95E-06 | 8.38E-06 |
| 1963 | ENSMUSG00000034042 | Gpbp1l1 | 77110 [Gene Symbol: Gpbp1l1] [Locus Tag: ] [Chromosome: 4] [Map Location: 4\|4 C7] | 2.279228945 | 2.02E-05 | 7.29E-05 |
| 16315 | ENSMUSG00000031939 | Taf1d | 75316 [Gene Symbol: Taf1d] [Locus Tag: ] [Chromosome: 9] [Map Location: 9\|9 A3] [De | 2.278183122 | 0.000465 | 0.001291 |
| 1673 | ENSMUSG00000032384 | Csnk1g1 | 214897 [Gene Symbol: Csnk1g1] [Locus Tag: ] [Chromosome: 9] [Map Location: 9\|9 C] [ | 2.276743926 | 3.24E-06 | 1.33E-05 |
| 14815 | ENSMUSG00000029426 | Scarb2 | 12492 [Gene Symbol: Scarb2] [Locus Tag: ] [Chromosome: 5] [Map Location: 5\|5 E3] [D | 2.274468119 | 0.000115 | 0.000356 |
| 2040 | ENSMUSG00000024556 | Me2 | 107029 [Gene Symbol: Me2] [Locus Tag: ] [Chromosome: 18] [Map Location: 18\|18 E2] | 2.272646657 | 1.35E-05 | 5.03E-05 |
| 16587 | ENSMUSG00000066442 | Mthfs | 107885 [Gene Symbol: Mthfs] [Locus Tag: ] [Chromosome: 9] [Map Location: 9\|9 E3.1] | 2.272621879 | 0.007476 | 0.016022 |
| 2309 | ENSMUSG00000040865 | Ino80d | 227195 [Gene Symbol: Ino80d] [Locus Tag: ] [Chromosome: 1] [Map Location: 1\|1 C2] [ | 2.270570377 | 6.13E-07 | 2.86E-06 |
| 13732 | ENSMUSG00000031628 | Casp3 | 12367 [Gene Symbol: Casp3] [Locus Tag: ] [Chromosome: 8] [Map Location: 8 B1.1\|8 26 | 2.269556724 | 6.83E-06 | 2.67E-05 |
| 12425 | ENSMUSG00000024083 | Pja2 | 224938 [Gene Symbol: Pja2] [Locus Tag: ] [Chromosome: 17] [Map Location: 17\|17 E1. | 2.26844042 | 8.79E-07 | 4.01E-06 |
| 13976 | ENSMUSG00000027200 | Sema6d | 214968 [Gene Symbol: Sema6d] [Locus Tag: ] [Chromosome: 2] [Map Location: 2\|2 F1] | 2.268365836 | 8.94E-07 | 4.07E-06 |
| 8248 | ENSMUSG00000034021 | Pds5b | 100710 [Gene Symbol: Pds5b] [Locus Tag: ] [Chromosome: 5] [Map Location: 5\|5 G3] [D | 2.267456347 | 2.12E-06 | 9.05E-06 |
| 13673 | ENSMUSG00000061613 | U2af1 | 108121 [Gene Symbol: U2af1] [Locus Tag: ] [Chromosome: 17] [Map Location: 17\|17 B | 2.265492661 | 0.00041 | 0.001151 |
| 9466 | ENSMUSG00000029775 | Klhdc10 | 76788 [Gene Symbol: Klhdc10] [Locus Tag: ] [Chromosome: 6] [Map Location: 6\|6 A3] [ | 2.264649536 | 9.41E-07 | 4.26E-06 |
| 10468 | ENSMUSG00000027274 | Mkks | 59030 [Gene Symbol: Mkks] [Locus Tag: ] [Chromosome: 2] [Map Location: 2\|2 F3] [Des | 2.264613017 | 6.35E-05 | 0.000207 |
| 1470 | ENSMUSG00000021468 | Sptlc1 | 268656 [Gene Symbol: Sptlc1] [Locus Tag: ] [Chromosome: 13] [Map Location: 13\|13 B | 2.263812263 | 0.000166 | 0.000503 |
| 11618 | ENSMUSG00000041132 | N4bp2l1 | 100637 [Gene Symbol: N4bp2l1] [Locus Tag: ] [Chromosome: 5] [Map Location: 5\|5 G3 | 2.263778716 | 0.001761 | 0.004344 |
| 18542 | ENSMUSG00000024270 | Slc39a6 | 106957 [Gene Symbol: Slc39a6] [Locus Tag: ] [Chromosome: 18] [Map Location: 18\|18 | 2.263145162 | 8.88E-05 | 0.000281 |
| 9111 | ENSMUSG00000025920 | Stau2 | 29819 [Gene Symbol: Stau2] [Locus Tag: ] [Chromosome: 1] [Map Location: 1\|1 A3] [De | 2.262637672 | 9.48E-07 | 4.29E-06 |
| 9164 | ENSMUSG00000004347 | Pde1c | 18575 [Gene Symbol: Pde1c] [Locus Tag: ] [Chromosome: 6] [Map Location: 6 B3\|6 27. | 2.26237637 | 2.44E-05 | 8.68E-05 |
| 11108 | ENSMUSG00000040289 | Hey1 | 15213 [Gene Symbol: Hey1] [Locus Tag: ] [Chromosome: 3] [Map Location: 3 A1\|3 2.15 | 2.261405541 | 0.000111 | 0.000346 |
| 15379 | ENSMUSG00000034518 | Hmgxb4 | 70823 [Gene Symbol: Hmgxb4] [Locus Tag: ] [Chromosome: 8] [Map Location: 8\|8 C1] [ | 2.260760155 | 2.72E-05 | 9.58E-05 |
| 13642 | ENSMUSG00000024146 | Cript | 56724 [Gene Symbol: Cript] [Locus Tag: ] [Chromosome: 17] [Map Location: 17\|17 E4] | 2.259415267 | 0.000189 | 0.000564 |
| 6623 | ENSMUSG00000022021 | Diaph3 | 56419 [Gene Symbol: Diaph3] [Locus Tag: ] [Chromosome: 14] [Map Location: 14\|14 D | 2.258000278 | 8.67E-05 | 0.000275 |
| 9707 | ENSMUSG00000024085 | Man2a1 | 17158 [Gene Symbol: Man2a1] [Locus Tag: ] [Chromosome: 17] [Map Location: 17\|17 E | 2.257964029 | 3.89E-06 | 1.58E-05 |
| 5562 | ENSMUSG00000032599 | Ip6k2 | 76500 [Gene Symbol: Ip6k2] [Locus Tag: ] [Chromosome: 9] [Map Location: 9\|9 F2] [De | 2.257599219 | 4.62E-06 | 1.85E-05 |
| 9458 | ENSMUSG00000003031 | Cdkn1b | 12576 [Gene Symbol: Cdkn1b] [Locus Tag: ] [Chromosome: 6] [Map Location: 6 G1\|6 65 | 2.257580417 | 5.84E-05 | 0.000192 |
| 3549 | ENSMUSG00000038346 | Zfp384 | 269800 [Gene Symbol: Zfp384] [Locus Tag: ] [Chromosome: 6] [Map Location: 6\|6 F2] [ | 2.257497876 | 2.43E-06 | 1.02E-05 |
| 11829 | ENSMUSG00000070695 | Cntnap5a | 636808 [Gene Symbol: Cntnap5a] [Locus Tag: ] [Chromosome: 1] [Map Location: 1\|1 E | 2.2570087 | 0.001375 | 0.003481 |
| 5680 | ENSMUSG00000022106 | Rcbtb2 | 105670 [Gene Symbol: Rcbtb2] [Locus Tag: ] [Chromosome: 14] [Map Location: 14\|14 | 2.256175775 | 5.86E-06 | 2.31E-05 |
| 3654 | ENSMUSG00000030207 | Fam234b | 74525 [Gene Symbol: Fam234b] [Locus Tag: ] [Chromosome: 6] [Map Location: 6\|6 G1] | 2.255722394 | 6.93E-06 | 2.70E-05 |
| 6991 | ENSMUSG00000043252 | Tmem64 | 100201 [Gene Symbol: Tmem64] [Locus Tag: ] [Chromosome: 4] [Map Location: 4\|4 A2 | 2.254694905 | 1.80E-05 | 6.57E-05 |
| 11883 | ENSMUSG00000091337 | Eid1 | 58521 [Gene Symbol: Eid1] [Locus Tag: ] [Chromosome: 2] [Map Location: 2\|2 F1] [Desc | 2.254323778 | 8.53E-07 | 3.90E-06 |
| 10771 | ENSMUSG00000060314 | Zfp941 | 407812 [Gene Symbol: Zfp941] [Locus Tag: ] [Chromosome: 7] [Map Location: 7\|7 F5] [ | 2.253820327 | 0.000119 | 0.000368 |
| 15617 | ENSMUSG00000031079 | Zfp300 | 245368 [Gene Symbol: Zfp300] [Locus Tag: ] [Chromosome: X] [Map Location: X\|X A2] [ | 2.253505512 | 0.001989 | 0.004847 |
| 14993 | ENSMUSG00000027562 | Car2 | 12349 [Gene Symbol: Car2] [Locus Tag: ] [Chromosome: 3] [Map Location: 3 A1\|3 3.23 | 2.252730198 | 0.014098 | 0.028281 |
| 9763 | ENSMUSG00000063568 | Jazf1 | 231986 [Gene Symbol: Jazf1] [Locus Tag: ] [Chromosome: 6] [Map Location: 6 B3\|6 25. | 2.25212334 | 8.73E-06 | 3.35E-05 |
| 8350 | ENSMUSG00000034538 | Zfp418 | 232854 [Gene Symbol: Zfp418] [Locus Tag: ] [Chromosome: 7] [Map Location: 7\|7 A1] [ | 2.251544341 | 0.00131 | 0.00333 |
| 1231 | ENSMUSG00000036904 | Fzd8 | 14370 [Gene Symbol: Fzd8] [Locus Tag: ] [Chromosome: 18] [Map Location: 18 A1\|18 4 | 2.251264733 | 0.000548 | 0.001499 |
| 18113 | ENSMUSG00000032186 | Tmod2 | 50876 [Gene Symbol: Tmod2] [Locus Tag: ] [Chromosome: 9] [Map Location: 9 D\|9 42. | 2.25121356 | 3.32E-05 | 0.000115 |
| 8788 | ENSMUSG00000029335 | Bmp3 | 110075 [Gene Symbol: Bmp3] [Locus Tag: ] [Chromosome: 5] [Map Location: 5 E3\|5 48 | 2.250761627 | 2.83E-05 | 9.92E-05 |
| 10907 | ENSMUSG00000023034 | Nr4a1 | 15370 [Gene Symbol: Nr4a1] [Locus Tag: ] [Chromosome: 15] [Map Location: 15\|15 F] [ | 2.250357308 | 0.013538 | 0.027263 |
| 444 | ENSMUSG00000053113 | Socs3 | 12702 [Gene Symbol: Socs3] [Locus Tag: ] [Chromosome: 11] [Map Location: 11\|11 E2] | 2.249999402 | 0.007433 | 0.015935 |
| 1506 | ENSMUSG00000031532 | Saraf | 67887 [Gene Symbol: Saraf] [Locus Tag: ] [Chromosome: 8] [Map Location: 8\|8 A4] [Des | 2.249544849 | 0.000968 | 0.002524 |
| 8359 | ENSMUSG00000026610 | Esrrg | 26381 [Gene Symbol: Esrrg] [Locus Tag: ] [Chromosome: 1] [Map Location: 1\|1 H6] [Des | 2.248388306 | 1.89E-05 | 6.86E-05 |
| 10938 | ENSMUSG00000028341 | Nr4a3 | 18124 [Gene Symbol: Nr4a3] [Locus Tag: ] [Chromosome: 4] [Map Location: 4\|4 B2] [De | 2.248249289 | 1.53E-06 | 6.70E-06 |
| 7540 | ENSMUSG00000029176 | Anapc4 | 52206 [Gene Symbol: Anapc4] [Locus Tag: ] [Chromosome: 5] [Map Location: 5 C1\|5 28 | 2.248060118 | 1.69E-05 | 6.18E-05 |
| 4370 | ENSMUSG00000034460 | Six4 | 20474 [Gene Symbol: Six4] [Locus Tag: ] [Chromosome: 12] [Map Location: 12\|12 C3] [ | 2.247814117 | 0.006038 | 0.0132 |
| 17402 | ENSMUSG00000025701 | Alox5 | 11689 [Gene Symbol: Alox5] [Locus Tag: ] [Chromosome: 6] [Map Location: 6 F1\|6 53.7 | 2.247626666 | 0.019056 | 0.036902 |
| 2080 | ENSMUSG00000033931 | Rbm34 | 52202 [Gene Symbol: Rbm34] [Locus Tag: ] [Chromosome: 8] [Map Location: 8 E2\|8 74 | 2.247578455 | 1.52E-05 | 5.60E-05 |
| 9891 | ENSMUSG00000025358 | Cdk2 | 12566 [Gene Symbol: Cdk2] [Locus Tag: ] [Chromosome: 10] [Map Location: 10\|10 D3] | 2.247103121 | 0.009911 | 0.020672 |
| 976 | ENSMUSG00000006599 | Gtf2h1 | 14884 [Gene Symbol: Gtf2h1] [Locus Tag: ] [Chromosome: 7] [Map Location: 7\|7 B4] [D | 2.246903777 | 3.18E-05 | 0.00011 |
| 9066 | ENSMUSG00000003662 | Ciao1 | 26371 [Gene Symbol: Ciao1] [Locus Tag: ] [Chromosome: 2] [Map Location: 2 61.86 cM | 2.245455935 | 4.89E-05 | 0.000163 |
| 14374 | ENSMUSG00000044667 | Plppr4 | 229791 [Gene Symbol: Plppr4] [Locus Tag: ] [Chromosome: 3] [Map Location: 3 G1\|3 5 | 2.245128694 | 0.000946 | 0.002474 |
| 2265 | ENSMUSG00000019906 | Lin7a | 108030 [Gene Symbol: Lin7a] [Locus Tag: ] [Chromosome: 10] [Map Location: 10\|10 D | 2.245080561 | 6.90E-05 | 0.000223 |
| 14606 | ENSMUSG00000028345 | Tex10 | 269536 [Gene Symbol: Tex10] [Locus Tag: ] [Chromosome: 4] [Map Location: 4\|4 B1] [D | 2.244891191 | 2.12E-05 | 7.62E-05 |
| 376 | ENSMUSG00000040265 | Dnm3 | 103967 [Gene Symbol: Dnm3] [Locus Tag: ] [Chromosome: 1] [Map Location: 1\|1 H2.1] | 2.244618123 | 5.95E-05 | 0.000195 |
| 3838 | ENSMUSG00000042050 | Wdr60 | 217935 [Gene Symbol: Wdr60] [Locus Tag: ] [Chromosome: 12] [Map Location: 12\|12 | 2.244506163 | 0.002221 | 0.005352 |
| 17850 | ENSMUSG00000027803 | Wwtr1 | 97064 [Gene Symbol: Wwtr1] [Locus Tag: ] [Chromosome: 3] [Map Location: 3\|3 D] [De | 2.24302034 | 4.28E-06 | 1.73E-05 |
| 3111 | ENSMUSG00000020097 | Sgpl1 | 20397 [Gene Symbol: Sgpl1] [Locus Tag: ] [Chromosome: 10] [Map Location: 10 B4\|10 | 2.242593578 | 1.02E-05 | 3.86E-05 |
| 10260 | ENSMUSG00000022677 | Fopnl | 66086 [Gene Symbol: Fopnl] [Locus Tag: ] [Chromosome: 16] [Map Location: 16\|16 B1] | 2.242307898 | 4.30E-05 | 0.000145 |
| 9834 | ENSMUSG00000056268 | Dennd1b | 329260 [Gene Symbol: Dennd1b] [Locus Tag: ] [Chromosome: 1] [Map Location: 1\|1 F] | 2.241805157 | 3.31E-06 | 1.36E-05 |
| 7030 | ENSMUSG00000029310 | Nudt9 | 74167 [Gene Symbol: Nudt9] [Locus Tag: ] [Chromosome: 5] [Map Location: 5\|5 E5] [De | 2.241690677 | 5.14E-05 | 0.000171 |
| 5083 | ENSMUSG00000039828 | Wdr70 | 545085 [Gene Symbol: Wdr70] [Locus Tag: ] [Chromosome: 15] [Map Location: 15\|15 | 2.240765062 | 8.82E-06 | 3.38E-05 |
| 9775 | ENSMUSG00000070871 | Ccnyl1 | 227210 [Gene Symbol: Ccnyl1] [Locus Tag: ] [Chromosome: 1] [Map Location: 1\|1 C2] [ | 2.240363657 | 1.76E-05 | 6.43E-05 |
| 10866 | ENSMUSG00000022789 | Dnm1l | 74006 [Gene Symbol: Dnm1l] [Locus Tag: ] [Chromosome: 16] [Map Location: 16\|16 A | 2.24025341 | 0.000486 | 0.001343 |
| 5583 | ENSMUSG00000033569 | Adgrb3 | 210933 [Gene Symbol: Adgrb3] [Locus Tag: ] [Chromosome: 1] [Map Location: 1\|1 A5] [ | 2.239428234 | 0.000201 | 0.000597 |
| 8412 | ENSMUSG00000037416 | Dmxl1 | 240283 [Gene Symbol: Dmxl1] [Locus Tag: ] [Chromosome: 18] [Map Location: 18\|18 D | 2.238082849 | 8.49E-07 | 3.88E-06 |
| 12797 | ENSMUSG00000021133 | Susd6 | 217684 [Gene Symbol: Susd6] [Locus Tag: ] [Chromosome: 12] [Map Location: 12\|12 D | 2.237968783 | 1.63E-05 | 5.97E-05 |
| 6319 | ENSMUSG00000027655 | Dhx35 | 71715 [Gene Symbol: Dhx35] [Locus Tag: ] [Chromosome: 2] [Map Location: 2\|2 H1] [D | 2.237797042 | 2.40E-05 | 8.53E-05 |
| 8923 | ENSMUSG00000027242 | Wdr76 | 241627 [Gene Symbol: Wdr76] [Locus Tag: ] [Chromosome: 2] [Map Location: 2\|2 F1] [ | 2.236965148 | 0.001673 | 0.004148 |
| 5323 | ENSMUSG00000010025 | Aldh3a2 | 11671 [Gene Symbol: Aldh3a2] [Locus Tag: ] [Chromosome: 11] [Map Location: 11 B2\| | 2.236882639 | 8.23E-05 | 0.000262 |
| 11019 | ENSMUSG00000042272 | Sestd1 | 228071 [Gene Symbol: Sestd1] [Locus Tag: ] [Chromosome: 2] [Map Location: 2\|2 C3] [ | 2.236856459 | 6.80E-07 | 3.15E-06 |
| 9056 | ENSMUSG00000046675 | Tmem251 | 320351 [Gene Symbol: Tmem251] [Locus Tag: ] [Chromosome: 12] [Map Location: 12\| | 2.235969225 | 0.000264 | 0.000767 |
| 4682 | ENSMUSG00000020925 | Ccdc43 | 52715 [Gene Symbol: Ccdc43] [Locus Tag: ] [Chromosome: 11] [Map Location: 11 E1\|1 | 2.235930077 | 0.000105 | 0.000328 |
| 2943 | ENSMUSG00000020185 | E2f7 | 52679 [Gene Symbol: E2f7] [Locus Tag: ] [Chromosome: 10] [Map Location: 10 D1\|10 5 | 2.235897853 | 0.001953 | 0.004767 |
| 13932 | ENSMUSG00000024781 | Lipa | 16889 [Gene Symbol: Lipa] [Locus Tag: ] [Chromosome: 19] [Map Location: 19\|19 C1] [ | 2.234512848 | 0.001393 | 0.003523 |
| 1647 | ENSMUSG00000041958 | Pigs | 276846 [Gene Symbol: Pigs] [Locus Tag: ] [Chromosome: 11] [Map Location: 11\|11 B5] | 2.234264957 | 4.37E-05 | 0.000147 |

| | | | | | | |
|---|---|---|---|---|---|---|
| 16791 | ENSMUSG00000044768 | D1Ertd622e | 52392 [Gene Symbol: D1Ertd622e] [Locus Tag: ] [Chromosome: 1] [Map Location: 1 D\| | 2.233643463 | 2.21E-06 | 9.36E-06 |
| 6878 | ENSMUSG00000006288 | Ttc5 | 219022 [Gene Symbol: Ttc5] [Locus Tag: ] [Chromosome: 14] [Map Location: 14\|14 C1] | 2.23326359 | 0.000186 | 0.000556 |
| 6024 | ENSMUSG00000031093 | Dock11 | 75974 [Gene Symbol: Dock11] [Locus Tag: ] [Chromosome: X] [Map Location: X\|X A3.2] | 2.230340307 | 2.07E-05 | 7.45E-05 |
| 2144 | ENSMUSG00000074916 | Chst14 | 72136 [Gene Symbol: Chst14] [Locus Tag: ] [Chromosome: 2] [Map Location: 2\|2 E5] [D | 2.230073672 | 0.01626 | 0.032086 |
| 17823 | ENSMUSG00000020952 | Scfd1 | 76983 [Gene Symbol: Scfd1] [Locus Tag: ] [Chromosome: 12] [Map Location: 12\|12 C1] | 2.229580384 | 0.000121 | 0.000374 |
| 488 | ENSMUSG00000034349 | Smc4 | 70099 [Gene Symbol: Smc4] [Locus Tag: ] [Chromosome: 3] [Map Location: 3\|3 E2] [Des | 2.22911987 | 0.001066 | 0.002759 |
| 14291 | ENSMUSG00000009079 | Ewsr1 | 14030 [Gene Symbol: Ewsr1] [Locus Tag: ] [Chromosome: 11] [Map Location: 11\|11 A1 | 2.229001087 | 9.84E-07 | 4.44E-06 |
| 16702 | ENSMUSG00000044176 | Spink10 | 328971 [Gene Symbol: Spink10] [Locus Tag: ] [Chromosome: 18] [Map Location: 18\|18 | 2.228727463 | 0.006319 | 0.013746 |
| 4641 | ENSMUSG00000027858 | Tspan2 | 70747 [Gene Symbol: Tspan2] [Locus Tag: ] [Chromosome: 3] [Map Location: 3\|3 F3] [D | 2.227556579 | 0.000623 | 0.001688 |
| 4608 | ENSMUSG00000038074 | Fkbp14 | 231997 [Gene Symbol: Fkbp14] [Locus Tag: ] [Chromosome: 6] [Map Location: 6\|6 B3] | 2.227338094 | 0.005792 | 0.012717 |
| 6499 | ENSMUSG00000033054 | Npat | 244879 [Gene Symbol: Npat] [Locus Tag: ] [Chromosome: 9] [Map Location: 9 A5.3\|9 2 | 2.226886832 | 3.47E-06 | 1.42E-05 |
| 13945 | ENSMUSG00000026475 | Rgs16 | 19734 [Gene Symbol: Rgs16] [Locus Tag: ] [Chromosome: 1] [Map Location: 1 G3\|1 65. | 2.226754031 | 0.000577 | 0.001574 |
| 3650 | ENSMUSG00000029462 | Vps29 | 56433 [Gene Symbol: Vps29] [Locus Tag: ] [Chromosome: 5] [Map Location: 5\|5 F] [Des | 2.225118397 | 9.26E-06 | 3.53E-05 |
| 18151 | ENSMUSG00000029648 | Flt1 | 14254 [Gene Symbol: Flt1] [Locus Tag: ] [Chromosome: 5] [Map Location: 5 87.01 cM\|5 | 2.225085262 | 5.58E-05 | 0.000184 |
| 4462 | ENSMUSG00000090698 | Apold1 | 381823 [Gene Symbol: Apold1] [Locus Tag: ] [Chromosome: 6] [Map Location: 6\|6 G1] | 2.224632639 | 0.002603 | 0.006186 |
| 641 | ENSMUSG00000059708 | Akap17b | 338351 [Gene Symbol: Akap17b] [Locus Tag: ] [Chromosome: X] [Map Location: X\|X A3 | 2.224174355 | 0.000415 | 0.001164 |
| 1993 | ENSMUSG00000031358 | Msl3 | 17692 [Gene Symbol: Msl3] [Locus Tag: ] [Chromosome: X] [Map Location: X\|X F5] [Des | 2.223672026 | 5.77E-05 | 0.000189 |
| 5780 | ENSMUSG00000052997 | Uba2 | 50995 [Gene Symbol: Uba2] [Locus Tag: ] [Chromosome: 7] [Map Location: 7\|7 B1] [Des | 2.222138716 | 9.97E-06 | 3.79E-05 |
| 4775 | ENSMUSG00000040653 | Ppp1r14c | 76142 [Gene Symbol: Ppp1r14c] [Locus Tag: ] [Chromosome: 10] [Map Location: 10\|10 | 2.220572778 | 2.26E-06 | 9.55E-06 |
| 7016 | ENSMUSG00000035759 | Bbs10 | 71769 [Gene Symbol: Bbs10] [Locus Tag: ] [Chromosome: 10] [Map Location: 10\|10 D1 | 2.220235349 | 0.006641 | 0.014372 |
| 3556 | ENSMUSG00000041765 | Ubac2 | 68889 [Gene Symbol: Ubac2] [Locus Tag: ] [Chromosome: 14] [Map Location: 14\|14 E5 | 2.219477344 | 4.05E-05 | 0.000137 |
| 11569 | ENSMUSG00000035158 | Mitf | 17342 [Gene Symbol: Mitf] [Locus Tag: ] [Chromosome: 6] [Map Location: 6 D3\|6 45.05 | 2.219055079 | 0.000215 | 0.000636 |
| 5427 | ENSMUSG00000025738 | Fbxl16 | 214931 [Gene Symbol: Fbxl16] [Locus Tag: ] [Chromosome: 17] [Map Location: 17\|17 | 2.217080554 | 0.000264 | 0.000768 |
| 650 | ENSMUSG00000053825 | Ppfia2 | 327814 [Gene Symbol: Ppfia2] [Locus Tag: ] [Chromosome: 10] [Map Location: 10\|10 D | 2.217046288 | 0.000114 | 0.000356 |
| 1195 | ENSMUSG00000029238 | Clock | 12753 [Gene Symbol: Clock] [Locus Tag: ] [Chromosome: 5] [Map Location: 5 C3.3\|5 40 | 2.216711661 | 3.19E-05 | 0.000111 |
| 14989 | ENSMUSG00000009535 | Rnmt | 67897 [Gene Symbol: Rnmt] [Locus Tag: ] [Chromosome: 18] [Map Location: 18\|18 E1] | 2.215644452 | 7.02E-06 | 2.74E-05 |
| 9815 | ENSMUSG00000056124 | B4galt6 | 56386 [Gene Symbol: B4galt6] [Locus Tag: ] [Chromosome: 18] [Map Location: 18\|18 A | 2.213165551 | 9.93E-07 | 4.48E-06 |
| 144 | ENSMUSG00000022390 | Zc3h7b | 20286 [Gene Symbol: Zc3h7b] [Locus Tag: ] [Chromosome: 15] [Map Location: 15\|15 E | 2.213114272 | 1.22E-06 | 5.43E-06 |
| 3440 | ENSMUSG00000053453 | Thoc7 | 66231 [Gene Symbol: Thoc7] [Locus Tag: ] [Chromosome: 14] [Map Location: 14\|14 A2 | 2.21306616 | 4.58E-05 | 0.000154 |
| 6580 | ENSMUSG00000046836 | Brox | 71678 [Gene Symbol: Brox] [Locus Tag: ] [Chromosome: 1] [Map Location: 1\|1] [Descrip | 2.212845464 | 0.00054 | 0.001481 |
| 7509 | ENSMUSG00000041881 | Ndufa7 | 66416 [Gene Symbol: Ndufa7] [Locus Tag: ] [Chromosome: 17] [Map Location: 17\|17 B | 2.212408941 | 0.002626 | 0.006237 |
| 15233 | ENSMUSG00000063236 | 1110038F14Rik | 117171 [Gene Symbol: 1110038F14Rik] [Locus Tag: ] [Chromosome: 15] [Map Location | 2.211501564 | 0.000384 | 0.001085 |
| 17274 | ENSMUSG00000021764 | Ndufs4 | 17993 [Gene Symbol: Ndufs4] [Locus Tag: ] [Chromosome: 13] [Map Location: 13\|13 D | 2.211396036 | 9.53E-05 | 0.0003 |
| 16889 | ENSMUSG00000040022 | Rab11fip2 | 74998 [Gene Symbol: Rab11fip2] [Locus Tag: ] [Chromosome: 19] [Map Location: 19\|1 | 2.210418218 | 7.34E-06 | 2.85E-05 |
| 1540 | ENSMUSG00000037664 | Cdkn1c | 12577 [Gene Symbol: Cdkn1c] [Locus Tag: ] [Chromosome: 7] [Map Location: 7 F5\|7 88 | 2.209853098 | 0.000712 | 0.001906 |
| 3519 | ENSMUSG00000009741 | Ubp1 | 22221 [Gene Symbol: Ubp1] [Locus Tag: ] [Chromosome: 9] [Map Location: 9\|9 F2] [Des | 2.209173505 | 1.41E-06 | 6.21E-06 |
| 4079 | ENSMUSG00000022305 | Lrp12 | 239393 [Gene Symbol: Lrp12] [Locus Tag: ] [Chromosome: 15] [Map Location: 15\|15 B | 2.207423469 | 2.92E-06 | 1.21E-05 |
| 4920 | ENSMUSG00000027365 | Trpm7 | 58800 [Gene Symbol: Trpm7] [Locus Tag: ] [Chromosome: 2] [Map Location: 2\|2 F2] [De | 2.206853565 | 8.77E-06 | 3.36E-05 |
| 7627 | ENSMUSG00000030500 | Slc17a6 | 140919 [Gene Symbol: Slc17a6] [Locus Tag: ] [Chromosome: 7] [Map Location: 7\|7 B5] | 2.206273316 | 3.70E-05 | 0.000127 |
| 10205 | ENSMUSG00000032386 | Trip4 | 56404 [Gene Symbol: Trip4] [Locus Tag: ] [Chromosome: 9] [Map Location: 9\|9 D] [Desc | 2.204851429 | 1.83E-05 | 6.66E-05 |
| 12261 | ENSMUSG00000062110 | Scfd2 | 212986 [Gene Symbol: Scfd2] [Locus Tag: ] [Chromosome: 5] [Map Location: 5\|5 C3.3] | 2.203778584 | 2.74E-05 | 9.63E-05 |
| 13401 | ENSMUSG00000031357 | Syap1 | 67043 [Gene Symbol: Syap1] [Locus Tag: ] [Chromosome: X] [Map Location: X\|X F4] [De | 2.20373594 | 0.000154 | 0.00047 |
| 12210 | ENSMUSG00000027820 | Mme | 17380 [Gene Symbol: Mme] [Locus Tag: ] [Chromosome: 3] [Map Location: 3 E1\|3 29.9 | 2.203330126 | 0.000249 | 0.000727 |
| 18093 | ENSMUSG00000037742 | Eef1a1 | 13627 [Gene Symbol: Eef1a1] [Locus Tag: ] [Chromosome: 9] [Map Location: 9\|9 E1] [De | 2.203289169 | 2.90E-05 | 0.000101 |
| 4953 | ENSMUSG00000047777 | Phf13 | 230936 [Gene Symbol: Phf13] [Locus Tag: ] [Chromosome: 4] [Map Location: 4\|4 E2] [D | 2.202986914 | 3.30E-05 | 0.000114 |
| 14036 | ENSMUSG00000041112 | Elmo1 | 140580 [Gene Symbol: Elmo1] [Locus Tag: ] [Chromosome: 13] [Map Location: 13\|13 A | 2.202448937 | 3.98E-06 | 1.61E-05 |
| 8416 | ENSMUSG00000064325 | Hhip | 15245 [Gene Symbol: Hhip] [Locus Tag: ] [Chromosome: 8] [Map Location: 8 37.95 cM\| | 2.202075831 | 0.005243 | 0.011628 |
| 11046 | ENSMUSG00000027519 | Rab22a | 19334 [Gene Symbol: Rab22a] [Locus Tag: ] [Chromosome: 2] [Map Location: 2 H4\|2 97 | 2.200462597 | 3.05E-06 | 1.26E-05 |
| 1242 | ENSMUSG00000021067 | Sav1 | 64010 [Gene Symbol: Sav1] [Locus Tag: ] [Chromosome: 12] [Map Location: 12\|12 C3] | 2.19984302 | 2.65E-06 | 1.11E-05 |
| 1483 | ENSMUSG00000015357 | Clpx | 270166 [Gene Symbol: Clpx] [Locus Tag: ] [Chromosome: 9] [Map Location: 9 C\|9 35.2 | 2.198577219 | 7.72E-05 | 0.000247 |
| 13680 | ENSMUSG00000014402 | Tsg101 | 22088 [Gene Symbol: Tsg101] [Locus Tag: ] [Chromosome: 7] [Map Location: 7\|7 B5] [D | 2.197696065 | 1.64E-05 | 6.02E-05 |
| 4598 | ENSMUSG00000059669 | Taf1b | 21340 [Gene Symbol: Taf1b] [Locus Tag: ] [Chromosome: 12] [Map Location: 12\|12 A3] | 2.196201391 | 0.001437 | 0.003622 |
| 11028 | ENSMUSG00000027091 | Zc3h15 | 69082 [Gene Symbol: Zc3h15] [Locus Tag: ] [Chromosome: 2] [Map Location: 2\|2 E1] [D | 2.195681334 | 1.33E-05 | 4.94E-05 |
| 6497 | ENSMUSG00000019838 | Slc16a10 | 72472 [Gene Symbol: Slc16a10] [Locus Tag: ] [Chromosome: 10] [Map Location: 10\|10 | 2.194601559 | 0.0021 | 0.00509 |
| 2765 | ENSMUSG00000040282 | BC052040 | 399568 [Gene Symbol: BC052040] [Locus Tag: ] [Chromosome: 2] [Map Location: 2\|2 E | 2.193792476 | 6.87E-05 | 0.000222 |
| 8176 | ENSMUSG00000061080 | Lsamp | 268890 [Gene Symbol: Lsamp] [Locus Tag: ] [Chromosome: 16] [Map Location: 16\|16 B | 2.193184121 | 3.66E-06 | 1.49E-05 |
| 4651 | ENSMUSG00000021693 | Kif2a | 16563 [Gene Symbol: Kif2a] [Locus Tag: ] [Chromosome: 13] [Map Location: 13\|13 F] [D | 2.19194172 | 1.41E-06 | 6.21E-06 |
| 12064 | ENSMUSG00000055421 | Pcdh9 | 211712 [Gene Symbol: Pcdh9] [Locus Tag: ] [Chromosome: 14] [Map Location: 14 E2.1 | 2.191877078 | 8.14E-07 | 3.73E-06 |
| 4657 | ENSMUSG00000042564 | Fam227a | 75729 [Gene Symbol: Fam227a] [Locus Tag: ] [Chromosome: 15] [Map Location: 15\|15 | 2.191653505 | 0.000995 | 0.002588 |
| 16658 | ENSMUSG00000102252 | Snrpn | 20646 [Gene Symbol: Snrpn] [Locus Tag: ] [Chromosome: 7] [Map Location: 7 C\|7 34.0 | 2.189021006 | 0.001137 | 0.002927 |
| 3068 | ENSMUSG00000037533 | Rapgef6 | 192786 [Gene Symbol: Rapgef6] [Locus Tag: ] [Chromosome: 11] [Map Location: 11\|11 | 2.188139372 | 3.04E-06 | 1.26E-05 |
| 1652 | ENSMUSG00000038582 | Pptc7 | 320717 [Gene Symbol: Pptc7] [Locus Tag: ] [Chromosome: 5] [Map Location: 5\|5 F] [De | 2.186899719 | 1.42E-05 | 5.27E-05 |
| 8396 | ENSMUSG00000027737 | Slc7a11 | 26570 [Gene Symbol: Slc7a11] [Locus Tag: ] [Chromosome: 3] [Map Location: 3 21.72 c | 2.186584026 | 0.001388 | 0.003512 |
| 7196 | ENSMUSG00000020000 | Moxd1 | 59012 [Gene Symbol: Moxd1] [Locus Tag: ] [Chromosome: 10] [Map Location: 10\|10 A | 2.186350121 | 0.010889 | 0.022476 |
| 14836 | ENSMUSG00000031631 | Cfap97 | 66756 [Gene Symbol: Cfap97] [Locus Tag: ] [Chromosome: 8] [Map Location: 8 26.16 c | 2.186325386 | 5.11E-05 | 0.00017 |
| 10514 | ENSMUSG00000031696 | Vps35 | 65114 [Gene Symbol: Vps35] [Locus Tag: ] [Chromosome: 8] [Map Location: 8 C3\|8 41. | 2.185988595 | 6.44E-06 | 2.52E-05 |
| 10780 | ENSMUSG00000024258 | Polr2d | 69241 [Gene Symbol: Polr2d] [Locus Tag: ] [Chromosome: 18] [Map Location: 18\|18 B | 2.185825938 | 8.65E-05 | 0.000275 |
| 13000 | ENSMUSG00000020978 | Klhdc2 | 69554 [Gene Symbol: Klhdc2] [Locus Tag: ] [Chromosome: 12] [Map Location: 12 28.77 | 2.183283436 | 4.30E-06 | 1.74E-05 |
| 8052 | ENSMUSG00000024776 | Stambpl1 | 76630 [Gene Symbol: Stambpl1] [Locus Tag: ] [Chromosome: 19] [Map Location: 19\|19 | 2.183210538 | 0.004776 | 0.010683 |
| 10390 | ENSMUSG00000042747 | Krtcap2 | 66059 [Gene Symbol: Krtcap2] [Locus Tag: ] [Chromosome: 3] [Map Location: 3\|3 F2] [ | 2.182743735 | 0.000491 | 0.001357 |
| 14988 | ENSMUSG00000027170 | Eif3m | 98221 [Gene Symbol: Eif3m] [Locus Tag: ] [Chromosome: 2] [Map Location: 2 E2\|2 55.0 | 2.182182398 | 0.000666 | 0.00179 |
| 15429 | ENSMUSG00000020056 | Ccdc53 | 67282 [Gene Symbol: Ccdc53] [Locus Tag: ] [Chromosome: 10] [Map Location: 10\|10 C | 2.182154268 | 0.000972 | 0.002533 |
| 12931 | ENSMUSG00000031529 | Tnks | 21951 [Gene Symbol: Tnks] [Locus Tag: ] [Chromosome: 8] [Map Location: 8 A4\|8 21.1 | 2.180183989 | 2.34E-06 | 9.87E-06 |
| 539 | ENSMUSG00000028222 | Calb1 | 12307 [Gene Symbol: Calb1] [Locus Tag: ] [Chromosome: 4] [Map Location: 4 A2\|4 6.6 | 2.179520053 | 0.005419 | 0.011972 |
| 13968 | ENSMUSG00000069114 | Zbtb10 | 229055 [Gene Symbol: Zbtb10] [Locus Tag: ] [Chromosome: 3] [Map Location: 3\|3 A1] | 2.17885766 | 2.50E-06 | 1.05E-05 |
| 5782 | ENSMUSG00000021963 | Sap18 | 20220 [Gene Symbol: Sap18] [Locus Tag: ] [Chromosome: 14] [Map Location: 14 C3\|14 | 2.178795868 | 0.001602 | 0.00399 |
| 7084 | ENSMUSG00000019256 | Ahr | 11622 [Gene Symbol: Ahr] [Locus Tag: ] [Chromosome: 12] [Map Location: 12 A3\|12 15 | 2.177351879 | 0.006583 | 0.014263 |
| 8444 | ENSMUSG00000005774 | Rfx5 | 53970 [Gene Symbol: Rfx5] [Locus Tag: ] [Chromosome: 3] [Map Location: 3 40.74 cM\| | 2.177023383 | 4.37E-05 | 0.000147 |
| 4181 | ENSMUSG00000041702 | Btbd7 | 238386 [Gene Symbol: Btbd7] [Locus Tag: ] [Chromosome: 12] [Map Location: 12\|12 E | 2.176800076 | 5.46E-05 | 0.00018 |
| 5188 | ENSMUSG00000028295 | Smim8 | 66291 [Gene Symbol: Smim8] [Locus Tag: ] [Chromosome: 4] [Map Location: 4\|4 A5] [D | 2.176204116 | 0.003959 | 0.009019 |
| 14269 | ENSMUSG00000029246 | Ppat | 231327 [Gene Symbol: Ppat] [Locus Tag: ] [Chromosome: 5] [Map Location: 5\|5 C3.3] [ | 2.176045022 | 9.49E-05 | 0.000299 |
| 8141 | ENSMUSG00000049516 | Spty2d1 | 101685 [Gene Symbol: Spty2d1] [Locus Tag: ] [Chromosome: 7] [Map Location: 7\|7 B4] | 2.175588842 | 8.56E-05 | 0.000272 |
| 2348 | ENSMUSG00000019810 | Fuca2 | 66848 [Gene Symbol: Fuca2] [Locus Tag: ] [Chromosome: 10] [Map Location: 10\|10 A2 | 2.174987651 | 0.000127 | 0.000392 |
| 6477 | ENSMUSG00000057506 | Bloc1s2 | 73689 [Gene Symbol: Bloc1s2] [Locus Tag: ] [Chromosome: 19] [Map Location: 19\|19 C | 2.17487209 | 0.018604 | 0.036151 |

| | | | | | | |
|---|---|---|---|---|---|---|
| 10193 | ENSMUSG00000039765 | Cc2d2a | 231214 [Gene Symbol: Cc2d2a] [Locus Tag: ] [Chromosome: 5] [Map Location: 5\|5 B3] | 2.172936674 | 4.00E-05 | 0.000136 |
| 12860 | ENSMUSG00000057715 | A830018L16Rik | 320492 [Gene Symbol: A830018L16Rik] [Locus Tag: ] [Chromosome: 1] [Map Location: | 2.172390795 | 0.018594 | 0.036139 |
| 10350 | ENSMUSG00000008730 | Hipk1 | 15257 [Gene Symbol: Hipk1] [Locus Tag: ] [Chromosome: 3] [Map Location: 3\|3 F3] [De | 2.171689512 | 4.39E-06 | 1.77E-05 |
| 11423 | ENSMUSG00000074994 | Qser1 | 99003 [Gene Symbol: Qser1] [Locus Tag: ] [Chromosome: 2] [Map Location: 2\|2 E2] [De | 2.170992031 | 8.81E-05 | 0.000279 |
| 4292 | ENSMUSG00000050505 | Pcdh20 | 219257 [Gene Symbol: Pcdh20] [Locus Tag: ] [Chromosome: 14] [Map Location: 14\|14 | 2.17079185 | 0.000177 | 0.000533 |
| 2012 | ENSMUSG00000033478 | Fam160b1 | 226252 [Gene Symbol: Fam160b1] [Locus Tag: ] [Chromosome: 19] [Map Location: 19\| | 2.170243456 | 8.83E-06 | 3.38E-05 |
| 2630 | ENSMUSG00000039043 | Arpin | 70420 [Gene Symbol: Arpin] [Locus Tag: ] [Chromosome: 7] [Map Location: 7\|7 D2] [De | 2.170090254 | 0.000896 | 0.002354 |
| 4515 | ENSMUSG00000032411 | Tfdp2 | 211586 [Gene Symbol: Tfdp2] [Locus Tag: ] [Chromosome: 9] [Map Location: 9\|9 E3.3] | 2.169768899 | 0.000209 | 0.00062 |
| 11060 | ENSMUSG00000047193 | Dync2h1 | 110350 [Gene Symbol: Dync2h1] [Locus Tag: ] [Chromosome: 9] [Map Location: 9 A1\|9 | 2.165597304 | 7.68E-06 | 2.98E-05 |
| 15062 | ENSMUSG00000028883 | Sema3a | 20346 [Gene Symbol: Sema3a] [Locus Tag: ] [Chromosome: 5] [Map Location: 5\|5 A1] [ | 2.165281967 | 2.39E-05 | 8.50E-05 |
| 698 | ENSMUSG00000036790 | Slitrk2 | 245450 [Gene Symbol: Slitrk2] [Locus Tag: ] [Chromosome: X] [Map Location: X\|X A5] [ | 2.164947544 | 7.50E-06 | 2.92E-05 |
| 2333 | ENSMUSG00000020528 | Prpsap2 | 212627 [Gene Symbol: Prpsap2] [Locus Tag: ] [Chromosome: 11] [Map Location: 11\|11 | 2.164808373 | 0.001238 | 0.003165 |
| 13553 | ENSMUSG00000056673 | Kdm5d | 20592 [Gene Symbol: Kdm5d] [Locus Tag: ] [Chromosome: Y] [Map Location: Y\|Y A1] [D | 2.16441825 | 0.004861 | 0.010861 |
| 16656 | ENSMUSG00000054079 | Utp18 | 217109 [Gene Symbol: Utp18] [Locus Tag: ] [Chromosome: 11] [Map Location: 11\|11 D | 2.164170722 | 4.22E-05 | 0.000142 |
| 17247 | ENSMUSG00000055027 | Smyd1 | 12180 [Gene Symbol: Smyd1] [Locus Tag: ] [Chromosome: 6] [Map Location: 6 C1\|6 32. | 2.163152229 | 0.000135 | 0.000415 |
| 2586 | ENSMUSG00000031634 | Ufsp2 | 192169 [Gene Symbol: Ufsp2] [Locus Tag: ] [Chromosome: 8] [Map Location: 8\|8 B1.1] | 2.163053721 | 0.00135 | 0.003424 |
| 5418 | ENSMUSG00000024248 | Cox7a2l | 20463 [Gene Symbol: Cox7a2l] [Locus Tag: ] [Chromosome: 17] [Map Location: 17\|17 E | 2.162347772 | 6.81E-05 | 0.000221 |
| 732 | ENSMUSG00000041205 | Map6d1 | 208158 [Gene Symbol: Map6d1] [Locus Tag: ] [Chromosome: 16] [Map Location: 16\|16 | 2.159167339 | 0.00053 | 0.001455 |
| 11286 | ENSMUSG00000070923 | Klhl9 | 242521 [Gene Symbol: Klhl9] [Locus Tag: ] [Chromosome: 4] [Map Location: 4\|4 C4] [D | 2.157060399 | 5.65E-05 | 0.000186 |
| 12852 | ENSMUSG00000106379 | Lhfpl3 | 269629 [Gene Symbol: Lhfpl3] [Locus Tag: ] [Chromosome: 5] [Map Location: 5\|5 A3] [ | 2.155896395 | 9.76E-05 | 0.000307 |
| 3921 | ENSMUSG00000029992 | Gfpt1 | 14583 [Gene Symbol: Gfpt1] [Locus Tag: ] [Chromosome: 6] [Map Location: 6 D1\|6 37.8 | 2.155566964 | 7.66E-06 | 2.97E-05 |
| 4558 | ENSMUSG00000029999 | Tgfa | 21802 [Gene Symbol: Tgfa] [Locus Tag: ] [Chromosome: 6] [Map Location: 6 D1\|6 37.62 | 2.155527925 | 3.49E-06 | 1.43E-05 |
| 718 | ENSMUSG00000020482 | Ccdc117 | 104479 [Gene Symbol: Ccdc117] [Locus Tag: ] [Chromosome: 11] [Map Location: 11\|1 | 2.154767896 | 0.001106 | 0.002855 |
| 1446 | ENSMUSG00000018196 | Glrx2 | 69367 [Gene Symbol: Glrx2] [Locus Tag: ] [Chromosome: 1] [Map Location: 1 F\|1 62.53 | 2.154539674 | 2.40E-05 | 8.53E-05 |
| 14710 | ENSMUSG00000062604 | Srpk2 | 20817 [Gene Symbol: Srpk2] [Locus Tag: ] [Chromosome: 5] [Map Location: 5 A3\|5 10.3 | 2.15438474 | 7.10E-06 | 2.77E-05 |
| 15256 | ENSMUSG00000027642 | Rpn2 | 20014 [Gene Symbol: Rpn2] [Locus Tag: ] [Chromosome: 2] [Map Location: 2 H1\|2 78.2 | 2.153705645 | 0.000313 | 0.000899 |
| 13980 | ENSMUSG00000003382 | Etv3 | 27049 [Gene Symbol: Etv3] [Locus Tag: ] [Chromosome: 3] [Map Location: 3\|3 F1] [Desc | 2.153057963 | 4.71E-05 | 0.000158 |
| 14587 | ENSMUSG00000002059 | Rab34 | 19376 [Gene Symbol: Rab34] [Locus Tag: ] [Chromosome: 11] [Map Location: 11 B5\|11 | 2.152994278 | 0.001291 | 0.003284 |
| 14768 | ENSMUSG00000069094 | Pde7a | 18583 [Gene Symbol: Pde7a] [Locus Tag: ] [Chromosome: 3] [Map Location: 3 A2\|3 5.3 | 2.152722093 | 1.98E-05 | 7.16E-05 |
| 14774 | ENSMUSG00000000826 | Dnajc5 | 13002 [Gene Symbol: Dnajc5] [Locus Tag: ] [Chromosome: 2] [Map Location: 2 H4\|2 10 | 2.151796485 | 4.00E-06 | 1.62E-05 |
| 6355 | ENSMUSG00000039887 | Alg14 | 66789 [Gene Symbol: Alg14] [Locus Tag: ] [Chromosome: 3] [Map Location: 3\|3 G1] [De | 2.151268724 | 0.001499 | 0.003758 |
| 12197 | ENSMUSG00000019971 | Cep290 | 216274 [Gene Symbol: Cep290] [Locus Tag: ] [Chromosome: 10] [Map Location: 10 D1\| | 2.151171769 | 0.007076 | 0.015229 |
| 18119 | ENSMUSG00000061393 | Acvr2b | 11481 [Gene Symbol: Acvr2b] [Locus Tag: ] [Chromosome: 9] [Map Location: 9\|9 F3] [D | 2.150879175 | 0.004303 | 0.009735 |
| 14761 | ENSMUSG00000064128 | Cenpj | 219103 [Gene Symbol: Cenpj] [Locus Tag: ] [Chromosome: 14] [Map Location: 14\|14 C | 2.149739648 | 8.75E-05 | 0.000277 |
| 10651 | ENSMUSG00000042208 | 0610010F05Rik | 71675 [Gene Symbol: 0610010F05Rik] [Locus Tag: ] [Chromosome: 11] [Map Location: | 2.149090366 | 3.36E-06 | 1.38E-05 |
| 18276 | ENSMUSG00000026341 | Actr3 | 74117 [Gene Symbol: Actr3] [Locus Tag: ] [Chromosome: 1] [Map Location: 1\|1 E3] [De | 2.148297878 | 1.40E-05 | 5.21E-05 |
| 4677 | ENSMUSG00000021974 | Fgf9 | 14180 [Gene Symbol: Fgf9] [Locus Tag: ] [Chromosome: 14] [Map Location: 14 30.51 c | 2.146946776 | 0.002222 | 0.005355 |
| 18266 | ENSMUSG00000031902 | Nfatc3 | 18021 [Gene Symbol: Nfatc3] [Locus Tag: ] [Chromosome: 8] [Map Location: 8 53.08 c | 2.146794843 | 7.52E-05 | 0.000242 |
| 735 | ENSMUSG00000071862 | Lrrtm2 | 107065 [Gene Symbol: Lrrtm2] [Locus Tag: ] [Chromosome: 18] [Map Location: 18\|18 | 2.146659733 | 1.90E-05 | 6.89E-05 |
| 3991 | ENSMUSG00000021592 | Arsk | 77041 [Gene Symbol: Arsk] [Locus Tag: ] [Chromosome: 13] [Map Location: 13\|13 C1] [ | 2.145744885 | 0.001751 | 0.004322 |
| 4656 | ENSMUSG00000025870 | Arl10 | 56795 [Gene Symbol: Arl10] [Locus Tag: ] [Chromosome: 13] [Map Location: 13\|13 B1] | 2.145133425 | 0.000122 | 0.000378 |
| 209 | ENSMUSG00000019883 | Echdc1 | 52665 [Gene Symbol: Echdc1] [Locus Tag: ] [Chromosome: 10] [Map Location: 10 A4\|1 | 2.143802354 | 2.74E-05 | 9.64E-05 |
| 6887 | ENSMUSG00000052331 | Ankrd44 | 329154 [Gene Symbol: Ankrd44] [Locus Tag: ] [Chromosome: 1] [Map Location: 1\|1 C1. | 2.143631662 | 5.00E-06 | 1.99E-05 |
| 7104 | ENSMUSG00000015243 | Abca1 | 11303 [Gene Symbol: Abca1] [Locus Tag: ] [Chromosome: 4] [Map Location: 4 28.57 c | 2.142321879 | 1.10E-05 | 4.15E-05 |
| 137 | ENSMUSG00000019804 | Snx3 | 54198 [Gene Symbol: Snx3] [Locus Tag: ] [Chromosome: 10] [Map Location: 10 B2\|10 2 | 2.141940736 | 7.84E-06 | 3.04E-05 |
| 15697 | ENSMUSG00000027344 | Fsip1 | 71313 [Gene Symbol: Fsip1] [Locus Tag: ] [Chromosome: 2] [Map Location: 2\|2 E5] [Des | 2.141867967 | 0.00876 | 0.018487 |
| 10683 | ENSMUSG00000010175 | Prox1 | 19130 [Gene Symbol: Prox1] [Locus Tag: ] [Chromosome: 1] [Map Location: 1 H6\|1 95. | 2.141189781 | 2.37E-05 | 8.45E-05 |
| 3923 | ENSMUSG00000060288 | Ppih | 66101 [Gene Symbol: Ppih] [Locus Tag: ] [Chromosome: 4] [Map Location: 4 55.34 cM\| | 2.140858469 | 0.013489 | 0.027174 |
| 17061 | ENSMUSG00000030691 | Fchsd2 | 207278 [Gene Symbol: Fchsd2] [Locus Tag: ] [Chromosome: 7] [Map Location: 7\|7 E3] [ | 2.139806273 | 2.30E-06 | 9.73E-06 |
| 193 | ENSMUSG00000028898 | Trnau1ap | 71787 [Gene Symbol: Trnau1ap] [Locus Tag: ] [Chromosome: 4] [Map Location: 4\|4 D3] | 2.139798249 | 0.000469 | 0.0013 |
| 13563 | ENSMUSG00000063334 | Krr1 | 52705 [Gene Symbol: Krr1] [Locus Tag: ] [Chromosome: 10] [Map Location: 10 D2\|10 5 | 2.139528733 | 0.000134 | 0.000411 |
| 2607 | ENSMUSG00000019856 | Fam184a | 75906 [Gene Symbol: Fam184a] [Locus Tag: ] [Chromosome: 10] [Map Location: 10\|10 | 2.139403206 | 0.005471 | 0.012075 |
| 17964 | ENSMUSG00000041238 | Rbbp8 | 225182 [Gene Symbol: Rbbp8] [Locus Tag: ] [Chromosome: 18] [Map Location: 18\|18 A | 2.138148446 | 1.90E-05 | 6.89E-05 |
| 817 | ENSMUSG00000033096 | Apmap | 71881 [Gene Symbol: Apmap] [Locus Tag: ] [Chromosome: 2] [Map Location: 2\|2 G3] [D | 2.1377309 | 0.000686 | 0.001842 |
| 17789 | ENSMUSG00000003153 | Slc2a3 | 20527 [Gene Symbol: Slc2a3] [Locus Tag: ] [Chromosome: 6] [Map Location: 6 F2\|6 57. | 2.137594074 | 2.99E-05 | 0.000104 |
| 1849 | ENSMUSG00000032212 | Sltm | 66660 [Gene Symbol: Sltm] [Locus Tag: ] [Chromosome: 9] [Map Location: 9\|9 D] [Desc | 2.135494266 | 0.000347 | 0.000988 |
| 6513 | ENSMUSG00000031137 | Fgf13 | 14168 [Gene Symbol: Fgf13] [Locus Tag: ] [Chromosome: X] [Map Location: X A6\|X 33.3 | 2.133933061 | 7.35E-05 | 0.000237 |
| 17926 | ENSMUSG00000038095 | Sbno1 | 243272 [Gene Symbol: Sbno1] [Locus Tag: ] [Chromosome: 5] [Map Location: 5\|5 F] [De | 2.133787745 | 7.77E-06 | 3.01E-05 |
| 7256 | ENSMUSG00000027806 | Tsc22d2 | 72033 [Gene Symbol: Tsc22d2] [Locus Tag: ] [Chromosome: 3] [Map Location: 3\|3 D] [D | 2.133705802 | 1.04E-05 | 3.93E-05 |
| 12582 | ENSMUSG00000026223 | Itm2c | 64294 [Gene Symbol: Itm2c] [Locus Tag: ] [Chromosome: 1] [Map Location: 1\|1 C5] [De | 2.131978299 | 0.002113 | 0.005116 |
| 1648 | ENSMUSG00000035109 | Shc4 | 271849 [Gene Symbol: Shc4] [Locus Tag: ] [Chromosome: 2] [Map Location: 2\|2 F1] [De | 2.131235361 | 3.74E-06 | 1.52E-05 |
| 11709 | ENSMUSG00000020910 | Adprm | 66358 [Gene Symbol: Adprm] [Locus Tag: ] [Chromosome: 11] [Map Location: 11\|11 B | 2.130214023 | 0.000851 | 0.002246 |
| 6200 | ENSMUSG00000028933 | Xrcc2 | 57434 [Gene Symbol: Xrcc2] [Locus Tag: ] [Chromosome: 5] [Map Location: 5\|5 A3] [De | 2.129443299 | 0.020157 | 0.038769 |
| 18395 | ENSMUSG00000019359 | Gdpd2 | 71584 [Gene Symbol: Gdpd2] [Locus Tag: ] [Chromosome: X] [Map Location: X\|X C2] [D | 2.127049015 | 0.006233 | 0.013581 |
| 8274 | ENSMUSG00000028019 | Pdgfc | 54635 [Gene Symbol: Pdgfc] [Locus Tag: ] [Chromosome: 3] [Map Location: 3\|3 E3] [De | 2.126196996 | 3.71E-05 | 0.000127 |
| 3596 | ENSMUSG00000032854 | Ugt8a | 22239 [Gene Symbol: Ugt8a] [Locus Tag: ] [Chromosome: 3] [Map Location: 3\|3 E3-F1] | 2.125744588 | 0.005234 | 0.011612 |
| 12825 | ENSMUSG00000067430 | Zfp763 | 73451 [Gene Symbol: Zfp763] [Locus Tag: ] [Chromosome: 17] [Map Location: 17\|17 B | 2.125201078 | 0.000393 | 0.00111 |
| 10398 | ENSMUSG00000031627 | Irf2 | 16363 [Gene Symbol: Irf2] [Locus Tag: ] [Chromosome: 8] [Map Location: 8\|8 B2] [Desc | 2.124884867 | 0.000299 | 0.000862 |
| 5256 | ENSMUSG00000022403 | St13 | 70356 [Gene Symbol: St13] [Locus Tag: ] [Chromosome: 15] [Map Location: 15\|15 E2] [ | 2.123987626 | 4.52E-06 | 1.81E-05 |
| 15570 | ENSMUSG00000053040 | Aph1c | 68318 [Gene Symbol: Aph1c] [Locus Tag: ] [Chromosome: 9] [Map Location: 9\|9 C] [Des | 2.123751336 | 0.005169 | 0.01148 |
| 14387 | ENSMUSG00000022914 | Brwd1 | 93871 [Gene Symbol: Brwd1] [Locus Tag: ] [Chromosome: 16] [Map Location: 16\|16 C4 | 2.123511738 | 3.10E-05 | 0.000108 |
| 5870 | ENSMUSG00000035572 | Dcaf10 | 242418 [Gene Symbol: Dcaf10] [Locus Tag: ] [Chromosome: 4] [Map Location: 4\|4 B1] [ | 2.123231094 | 1.30E-05 | 4.85E-05 |
| 10306 | ENSMUSG00000039542 | Ncam1 | 17967 [Gene Symbol: Ncam1] [Locus Tag: ] [Chromosome: 9] [Map Location: 9 A5.3\|9 2 | 2.123017127 | 3.37E-06 | 1.38E-05 |
| 3804 | ENSMUSG00000047379 | B4gat1 | 108902 [Gene Symbol: B4gat1] [Locus Tag: ] [Chromosome: 19] [Map Location: 19\|19 | 2.122870292 | 0.003104 | 0.007254 |
| 10498 | ENSMUSG00000000567 | Sox9 | 20682 [Gene Symbol: Sox9] [Locus Tag: ] [Chromosome: 11] [Map Location: 11 E2\|11 7 | 2.122469017 | 5.63E-05 | 0.000185 |
| 8792 | ENSMUSG00000037788 | Vopp1 | 232023 [Gene Symbol: Vopp1] [Locus Tag: ] [Chromosome: 6] [Map Location: 6\|6 B3] [ | 2.122022751 | 6.07E-06 | 2.39E-05 |
| 3103 | ENSMUSG00000078926 | Cdc20b | 238896 [Gene Symbol: Cdc20b] [Locus Tag: ] [Chromosome: 13] [Map Location: 13\|13 | 2.120212222 | 0.007803 | 0.016651 |
| 1538 | ENSMUSG00000037712 | Fermt2 | 218952 [Gene Symbol: Fermt2] [Locus Tag: ] [Chromosome: 14] [Map Location: 14\|14 | 2.118390403 | 3.03E-05 | 0.000105 |
| 3289 | ENSMUSG00000021478 | Drd1 | 13488 [Gene Symbol: Drd1] [Locus Tag: ] [Chromosome: 13] [Map Location: 13 B1\|13 2 | 2.117759417 | 5.03E-05 | 0.000167 |
| 14660 | ENSMUSG00000026031 | Cflar | 12633 [Gene Symbol: Cflar] [Locus Tag: ] [Chromosome: 1] [Map Location: 1 C1.3\|1 29. | 2.116465 | 1.39E-05 | 5.17E-05 |
| 3278 | ENSMUSG00000023026 | Dip2b | 239667 [Gene Symbol: Dip2b] [Locus Tag: ] [Chromosome: 15] [Map Location: 15\|15 F | 2.115772876 | 8.18E-06 | 3.16E-05 |
| 285 | ENSMUSG00000037486 | Asxl2 | 75302 [Gene Symbol: Asxl2] [Locus Tag: ] [Chromosome: 12] [Map Location: 12\|12 A1. | 2.115217832 | 1.04E-05 | 3.94E-05 |
| 8220 | ENSMUSG00000058152 | Chsy3 | 78923 [Gene Symbol: Chsy3] [Locus Tag: ] [Chromosome: 18] [Map Location: 18\|18 D3 | 2.11492899 | 0.005235 | 0.011613 |
| 5060 | ENSMUSG00000022636 | Alcam | 11658 [Gene Symbol: Alcam] [Locus Tag: ] [Chromosome: 16] [Map Location: 16\|16 B5 | 2.114824846 | 9.49E-06 | 3.62E-05 |

| | | | | | | |
|---|---|---|---|---|---|---|
| 2651 | ENSMUSG00000053119 | Chmp3 | 66700 [Gene Symbol: Chmp3] [Locus Tag: ] [Chromosome: 6] [Map Location: 6 32.17 c | 2.114526191 | 1.25E-05 | 4.66E-05 |
| 17595 | ENSMUSG00000020646 | Mboat2 | 67216 [Gene Symbol: Mboat2] [Locus Tag: ] [Chromosome: 12] [Map Location: 12\|12 A | 2.114409467 | 1.76E-05 | 6.42E-05 |
| 2198 | ENSMUSG00000015971 | Actr8 | 56249 [Gene Symbol: Actr8] [Locus Tag: ] [Chromosome: 14] [Map Location: 14\|14 B] [ | 2.113753718 | 8.72E-05 | 0.000276 |
| 672 | ENSMUSG00000046727 | Cystm1 | 66060 [Gene Symbol: Cystm1] [Locus Tag: ] [Chromosome: 18] [Map Location: 18\|18 B | 2.113716025 | 0.00671 | 0.014505 |
| 1786 | ENSMUSG00000050558 | Prokr2 | 246313 [Gene Symbol: Prokr2] [Locus Tag: ] [Chromosome: 2] [Map Location: 2\|2 F2] [ | 2.113482872 | 4.63E-05 | 0.000155 |
| 3413 | ENSMUSG00000072774 | Zfp951 | 626391 [Gene Symbol: Zfp951] [Locus Tag: ] [Chromosome: 5] [Map Location: 5\|5 E5] [ | 2.113273269 | 0.003842 | 0.008783 |
| 3516 | ENSMUSG00000026380 | Tfcp2l1 | 81879 [Gene Symbol: Tfcp2l1] [Locus Tag: ] [Chromosome: 1] [Map Location: 1\|1 E2] [D | 2.111952673 | 0.019887 | 0.038321 |
| 15287 | ENSMUSG00000016984 | Etaa1 | 68145 [Gene Symbol: Etaa1] [Locus Tag: ] [Chromosome: 11] [Map Location: 11\|11 A3. | 2.111011006 | 0.00017 | 0.000512 |
| 1854 | ENSMUSG00000022514 | Il1rap | 16180 [Gene Symbol: Il1rap] [Locus Tag: ] [Chromosome: 16] [Map Location: 16\|16 B2 | 2.110955523 | 0.000766 | 0.002039 |
| 2881 | ENSMUSG00000032803 | Cdv3 | 321022 [Gene Symbol: Cdv3] [Locus Tag: ] [Chromosome: 9] [Map Location: 9\|9 F1] [De | 2.110455078 | 3.35E-05 | 0.000116 |
| 6169 | ENSMUSG00000004285 | Atp6v1f | 66144 [Gene Symbol: Atp6v1f] [Locus Tag: ] [Chromosome: 6] [Map Location: 6\|6 A3] [ | 2.110452605 | 0.004734 | 0.010599 |
| 5570 | ENSMUSG00000026322 | Htr4 | 15562 [Gene Symbol: Htr4] [Locus Tag: ] [Chromosome: 18] [Map Location: 18\|18 D3] | 2.109789118 | 0.007276 | 0.015634 |
| 2222 | ENSMUSG00000022231 | Sema5a | 20356 [Gene Symbol: Sema5a] [Locus Tag: ] [Chromosome: 15] [Map Location: 15 B3.1 | 2.109512659 | 6.34E-05 | 0.000207 |
| 3668 | ENSMUSG00000027272 | Ubr1 | 22222 [Gene Symbol: Ubr1] [Locus Tag: ] [Chromosome: 2] [Map Location: 2 E5\|2 60.3 | 2.109358896 | 1.74E-05 | 6.37E-05 |
| 9339 | ENSMUSG00000017412 | Cacnb4 | 12298 [Gene Symbol: Cacnb4] [Locus Tag: ] [Chromosome: 2] [Map Location: 2 C1.1\|2 | 2.108820766 | 2.97E-05 | 0.000104 |
| 17615 | ENSMUSG00000031229 | Atrx | 22589 [Gene Symbol: Atrx] [Locus Tag: ] [Chromosome: X] [Map Location: X D\|X 47.26 | 2.108789612 | 0.000415 | 0.001164 |
| 11410 | ENSMUSG00000063550 | Nup98 | 269966 [Gene Symbol: Nup98] [Locus Tag: ] [Chromosome: 7] [Map Location: 7 E2\|7 5 | 2.107046516 | 1.51E-05 | 5.57E-05 |
| 17736 | ENSMUSG00000020823 | Sec14l1 | 74136 [Gene Symbol: Sec14l1] [Locus Tag: ] [Chromosome: 11] [Map Location: 11\|11 E | 2.106093115 | 7.62E-06 | 2.96E-05 |
| 7197 | ENSMUSG00000028126 | Pip5k1a | 18720 [Gene Symbol: Pip5k1a] [Locus Tag: ] [Chromosome: 3] [Map Location: 3\|3 F2.1 | 2.102840779 | 4.28E-05 | 0.000145 |
| 5459 | ENSMUSG00000038331 | Satb2 | 212712 [Gene Symbol: Satb2] [Locus Tag: ] [Chromosome: 1] [Map Location: 1\|1 C1.3] | 2.102380456 | 5.28E-06 | 2.10E-05 |
| 12841 | ENSMUSG00000031922 | Cep57 | 74360 [Gene Symbol: Cep57] [Locus Tag: ] [Chromosome: 9] [Map Location: 9\|9 A2] [De | 2.101612927 | 0.000246 | 0.00072 |
| 16855 | ENSMUSG00000026566 | Mpzl1 | 68481 [Gene Symbol: Mpzl1] [Locus Tag: ] [Chromosome: 1] [Map Location: 1\|1 H2.3] [ | 2.100971191 | 0.000127 | 0.000391 |
| 1904 | ENSMUSG00000066441 | Rdh11 | 17252 [Gene Symbol: Rdh11] [Locus Tag: ] [Chromosome: 12] [Map Location: 12 C3\|12 | 2.100354262 | 0.000282 | 0.000816 |
| 347 | ENSMUSG00000054659 | Pm20d2 | 242377 [Gene Symbol: Pm20d2] [Locus Tag: ] [Chromosome: 4] [Map Location: 4\|4 A5 | 2.099286357 | 0.008096 | 0.017221 |
| 15655 | ENSMUSG00000027522 | Stx16 | 228960 [Gene Symbol: Stx16] [Locus Tag: ] [Chromosome: 2] [Map Location: 2\|2 H4] [D | 2.099141373 | 1.80E-05 | 6.56E-05 |
| 1682 | ENSMUSG00000033760 | Rbm4b | 66704 [Gene Symbol: Rbm4b] [Locus Tag: ] [Chromosome: 19] [Map Location: 19\|19 A | 2.098949016 | 0.00031 | 0.00089 |
| 3021 | ENSMUSG00000053626 | Tll1 | 21892 [Gene Symbol: Tll1] [Locus Tag: ] [Chromosome: 8] [Map Location: 8 B3.1\|8 32.1 | 2.098540726 | 0.001003 | 0.002607 |
| 10437 | ENSMUSG00000062519 | Zfp398 | 272347 [Gene Symbol: Zfp398] [Locus Tag: ] [Chromosome: 6] [Map Location: 6\|6 B3] [ | 2.098449938 | 4.51E-05 | 0.000152 |
| 2193 | ENSMUSG00000019978 | Epb41l2 | 13822 [Gene Symbol: Epb41l2] [Locus Tag: ] [Chromosome: 10] [Map Location: 10 A4\| | 2.097897361 | 0.003641 | 0.008371 |
| 2685 | ENSMUSG00000019699 | Akt3 | 23797 [Gene Symbol: Akt3] [Locus Tag: ] [Chromosome: 1] [Map Location: 1\|1 H4-H6] [ | 2.097618138 | 3.87E-06 | 1.57E-05 |
| 7346 | ENSMUSG00000040105 | Plpp6 | 74411 [Gene Symbol: Plpp6] [Locus Tag: ] [Chromosome: 19] [Map Location: 19\|19 C1 | 2.097477161 | 3.62E-05 | 0.000124 |
| 424 | ENSMUSG00000063364 | 3300002I08Rik | 69277 [Gene Symbol: 3300002I08Rik] [Locus Tag: ] [Chromosome: 2] [Map Location: 2 | 2.097273444 | 0.000453 | 0.001261 |
| 13939 | ENSMUSG00000036292 | Gramd1c | 207798 [Gene Symbol: Gramd1c] [Locus Tag: ] [Chromosome: 16] [Map Location: 16\|1 | 2.096815099 | 0.026107 | 0.048905 |
| 4445 | ENSMUSG00000047804 | Akap10 | 56697 [Gene Symbol: Akap10] [Locus Tag: ] [Chromosome: 11] [Map Location: 11\|11 B | 2.096795439 | 1.72E-05 | 6.30E-05 |
| 18159 | ENSMUSG00000032340 | Neo1 | 18007 [Gene Symbol: Neo1] [Locus Tag: ] [Chromosome: 9] [Map Location: 9\|9 B] [Desc | 2.096066927 | 1.13E-05 | 4.26E-05 |
| 16397 | ENSMUSG00000022300 | Dcaf13 | 223499 [Gene Symbol: Dcaf13] [Locus Tag: ] [Chromosome: 15] [Map Location: 15\|15 | 2.09595657 | 0.000284 | 0.00082 |
| 2454 | ENSMUSG00000039988 | Ankrd13c | 433667 [Gene Symbol: Ankrd13c] [Locus Tag: ] [Chromosome: 3] [Map Location: 3\|3 H | 2.09591768 | 4.97E-05 | 0.000166 |
| 14782 | ENSMUSG00000075600 | Zc3h3 | 223642 [Gene Symbol: Zc3h3] [Locus Tag: ] [Chromosome: 15] [Map Location: 15\|15 D | 2.095367483 | 0.001374 | 0.003481 |
| 2605 | ENSMUSG00000038759 | Nup205 | 70699 [Gene Symbol: Nup205] [Locus Tag: ] [Chromosome: 6] [Map Location: 6\|6 B1] [ | 2.094818994 | 1.28E-05 | 4.79E-05 |
| 9882 | ENSMUSG00000021636 | Marveld2 | 218518 [Gene Symbol: Marveld2] [Locus Tag: ] [Chromosome: 13] [Map Location: 13\|1 | 2.094270632 | 0.025169 | 0.047347 |
| 1491 | ENSMUSG00000029156 | Sgcb | 24051 [Gene Symbol: Sgcb] [Locus Tag: ] [Chromosome: 5] [Map Location: 5\|5 C3.3] [D | 2.094052085 | 0.000101 | 0.000315 |
| 2437 | ENSMUSG00000040842 | Szrd1 | 213491 [Gene Symbol: Szrd1] [Locus Tag: ] [Chromosome: 4] [Map Location: 4 D3\|4 73 | 2.091584731 | 0.000111 | 0.000344 |
| 13351 | ENSMUSG00000031353 | Rbbp7 | 245688 [Gene Symbol: Rbbp7] [Locus Tag: ] [Chromosome: X] [Map Location: X\|X F4] [ | 2.089823973 | 3.26E-05 | 0.000113 |
| 741 | ENSMUSG00000020070 | Rufy2 | 70432 [Gene Symbol: Rufy2] [Locus Tag: ] [Chromosome: 10] [Map Location: 10 B4\|10 | 2.089154096 | 2.02E-05 | 7.29E-05 |
| 203 | ENSMUSG00000037071 | Scd1 | 20249 [Gene Symbol: Scd1] [Locus Tag: ] [Chromosome: 19] [Map Location: 19 C3\|19 3 | 2.086783536 | 6.54E-05 | 0.000213 |
| 16482 | ENSMUSG00000025938 | Slco5a1 | 240726 [Gene Symbol: Slco5a1] [Locus Tag: ] [Chromosome: 1] [Map Location: 1\|1 A3] | 2.085911716 | 0.013999 | 0.028103 |
| 13770 | ENSMUSG00000027660 | Skil | 20482 [Gene Symbol: Skil] [Locus Tag: ] [Chromosome: 3] [Map Location: 3 A3\|3 15.12 | 2.085418641 | 5.22E-06 | 2.07E-05 |
| 8071 | ENSMUSG00000022270 | Fam134b | 66270 [Gene Symbol: Fam134b] [Locus Tag: ] [Chromosome: 15] [Map Location: 15\|15 | 2.085068731 | 0.000159 | 0.000482 |
| 12259 | ENSMUSG00000056947 | Mab21l1 | 17116 [Gene Symbol: Mab21l1] [Locus Tag: ] [Chromosome: 3] [Map Location: 3\|3 E3- | 2.083657245 | 0.000574 | 0.001564 |
| 6563 | ENSMUSG00000041057 | Wdr43 | 72515 [Gene Symbol: Wdr43] [Locus Tag: ] [Chromosome: 17] [Map Location: 17\|17 E1 | 2.082851033 | 8.21E-05 | 0.000262 |
| 5244 | ENSMUSG00000078671 | Chd2 | 244059 [Gene Symbol: Chd2] [Locus Tag: ] [Chromosome: 7] [Map Location: 7\|7 D1] [D | 2.08257478 | 6.79E-06 | 2.65E-05 |
| 5669 | ENSMUSG00000020303 | Stc2 | 20856 [Gene Symbol: Stc2] [Locus Tag: ] [Chromosome: 11] [Map Location: 11\|11 A4] [ | 2.081697638 | 0.010146 | 0.021102 |
| 17108 | ENSMUSG00000024242 | Map4k3 | 225028 [Gene Symbol: Map4k3] [Locus Tag: ] [Chromosome: 17] [Map Location: 17\|17 | 2.081352925 | 8.77E-06 | 3.36E-05 |
| 3933 | ENSMUSG00000044949 | Ubtd2 | 327900 [Gene Symbol: Ubtd2] [Locus Tag: ] [Chromosome: 11] [Map Location: 11\|11 A | 2.081339497 | 3.11E-05 | 0.000108 |
| 13942 | ENSMUSG00000052407 | Ccdc171 | 320226 [Gene Symbol: Ccdc171] [Locus Tag: ] [Chromosome: 4] [Map Location: 4\|4 C3 | 2.080596943 | 0.0008 | 0.002118 |
| 8441 | ENSMUSG00000022297 | Fzd6 | 14368 [Gene Symbol: Fzd6] [Locus Tag: ] [Chromosome: 15] [Map Location: 15 B3.1\|15 | 2.080535458 | 0.001655 | 0.00411 |
| 7980 | ENSMUSG00000053411 | Cbx7 | 52609 [Gene Symbol: Cbx7] [Locus Tag: ] [Chromosome: 15] [Map Location: 15 E1\|15 3 | 2.080363231 | 0.003677 | 0.008445 |
| 7972 | ENSMUSG00000050714 | Zbtb26 | 320633 [Gene Symbol: Zbtb26] [Locus Tag: ] [Chromosome: 2] [Map Location: 2\|2 B] [D | 2.080353448 | 0.000173 | 0.000521 |
| AVE=Mean=Geomean | | | | 2.90420123 | | |
| 1583 | ENSMUSG00000022889 | Mrpl39 | 27393 [Gene Symbol: Mrpl39] [Locus Tag: ] [Chromosome: 16] [Map Location: 16\|16 C | 2.080025317 | 0.000311 | 0.000892 |
| 8630 | ENSMUSG00000074165 | Zfp788 | 67607 [Gene Symbol: Zfp788] [Locus Tag: ] [Chromosome: 7] [Map Location: 7 B3\|7] [D | 2.079560013 | 3.87E-05 | 0.000132 |
| 723 | ENSMUSG00000021759 | Plpp1 | 19012 [Gene Symbol: Plpp1] [Locus Tag: ] [Chromosome: 13] [Map Location: 13\|13 D2 | 2.079491866 | 0.000316 | 0.000905 |
| 17459 | ENSMUSG00000019082 | Slc25a22 | 68267 [Gene Symbol: Slc25a22] [Locus Tag: ] [Chromosome: 7] [Map Location: 7\|7 F5] | 2.079398834 | 3.52E-05 | 0.000121 |
| 11920 | ENSMUSG00000021549 | Rasa1 | 218397 [Gene Symbol: Rasa1] [Locus Tag: ] [Chromosome: 13] [Map Location: 13 C3\|1 | 2.079193314 | 7.82E-06 | 3.03E-05 |
| 10435 | ENSMUSG00000028093 | Acp6 | 66659 [Gene Symbol: Acp6] [Locus Tag: ] [Chromosome: 3] [Map Location: 3\|3 F2.1] [D | 2.078843102 | 0.017777 | 0.034704 |
| 8805 | ENSMUSG00000027673 | Ndufb5 | 66046 [Gene Symbol: Ndufb5] [Locus Tag: ] [Chromosome: 3] [Map Location: 3\|3 A3] [ | 2.078713065 | 0.000698 | 0.001873 |
| 6023 | ENSMUSG00000024188 | Luc7l | 66978 [Gene Symbol: Luc7l] [Locus Tag: ] [Chromosome: 17] [Map Location: 17 13.09 | 2.07725576 | 1.46E-05 | 5.40E-05 |
| 15494 | ENSMUSG00000020716 | Nf1 | 18015 [Gene Symbol: Nf1] [Locus Tag: ] [Chromosome: 11] [Map Location: 11 46.74 c | 2.077049755 | 5.94E-06 | 2.34E-05 |
| 8369 | ENSMUSG00000038481 | Cdk19 | 78334 [Gene Symbol: Cdk19] [Locus Tag: ] [Chromosome: 10] [Map Location: 10\|10 B1 | 2.076496808 | 1.07E-05 | 4.05E-05 |
| 13895 | ENSMUSG00000072969 | Armcx5 | 494468 [Gene Symbol: Armcx5] [Locus Tag: ] [Chromosome: X] [Map Location: X\|X F1] | 2.0764848 | 2.95E-05 | 0.000103 |
| 18032 | ENSMUSG00000028614 | Ndc1 | 72787 [Gene Symbol: Ndc1] [Locus Tag: ] [Chromosome: 4] [Map Location: 4\|4 C7] [De | 2.07423632 | 0.000143 | 0.000437 |
| 11860 | ENSMUSG00000031731 | Ap1g1 | 11765 [Gene Symbol: Ap1g1] [Locus Tag: ] [Chromosome: 8] [Map Location: 8 D3\|8 57. | 2.073519538 | 1.45E-05 | 5.36E-05 |
| 6685 | ENSMUSG00000040325 | Vprbp | 321006 [Gene Symbol: Vprbp] [Locus Tag: ] [Chromosome: 9] [Map Location: 9\|9 F1] [D | 2.073280082 | 5.35E-05 | 0.000177 |
| 17997 | ENSMUSG00000044674 | Fzd1 | 14362 [Gene Symbol: Fzd1] [Locus Tag: ] [Chromosome: 5] [Map Location: 5 A1\|5 2.61 | 2.071592288 | 0.018024 | 0.035128 |
| 1234 | ENSMUSG00000063760 | Rnf217 | 268291 [Gene Symbol: Rnf217] [Locus Tag: ] [Chromosome: 10] [Map Location: 10\|10 | 2.071393465 | 9.68E-06 | 3.68E-05 |
| 11560 | ENSMUSG00000005667 | Mthfd2 | 17768 [Gene Symbol: Mthfd2] [Locus Tag: ] [Chromosome: 6] [Map Location: 6 C3\|6 35 | 2.070917183 | 0.001427 | 0.003599 |
| 9546 | ENSMUSG00000047037 | Nipa1 | 233280 [Gene Symbol: Nipa1] [Locus Tag: ] [Chromosome: 7] [Map Location: 7\|7 C] [De | 2.070268779 | 1.91E-05 | 6.93E-05 |
| 4543 | ENSMUSG00000024048 | Myl12a | 67268 [Gene Symbol: Myl12a] [Locus Tag: ] [Chromosome: 17] [Map Location: 17\|17 E | 2.06876951 | 0.00406 | 0.009231 |
| 8793 | ENSMUSG00000028282 | Casp8ap2 | 26885 [Gene Symbol: Casp8ap2] [Locus Tag: ] [Chromosome: 4] [Map Location: 4 A5\|4 | 2.06859802 | 8.56E-06 | 3.29E-05 |
| 11275 | ENSMUSG00000047485 | Klhl34 | 245683 [Gene Symbol: Klhl34] [Locus Tag: ] [Chromosome: X] [Map Location: X\|X F4] [ | 2.06839906 | 0.000286 | 0.000826 |
| 10525 | ENSMUSG00000049122 | Frmd3 | 242506 [Gene Symbol: Frmd3] [Locus Tag: ] [Chromosome: 4] [Map Location: 4\|4 C3] [ | 2.068278578 | 2.26E-05 | 8.09E-05 |
| 8915 | ENSMUSG00000026426 | Arl8a | 68724 [Gene Symbol: Arl8a] [Locus Tag: ] [Chromosome: 1] [Map Location: 1\|1 F] [Desc | 2.06754993 | 4.81E-05 | 0.000161 |
| 2220 | ENSMUSG00000068747 | Sort1 | 20661 [Gene Symbol: Sort1] [Locus Tag: ] [Chromosome: 3] [Map Location: 3\|3 F3] [Des | 2.06749296 | 1.57E-05 | 5.77E-05 |
| 4035 | ENSMUSG00000052707 | Tnrc6a | 233833 [Gene Symbol: Tnrc6a] [Locus Tag: ] [Chromosome: 7] [Map Location: 7\|7 F3] [ | 2.06632414 | 0.000175 | 0.000526 |

| | | | | | | |
|---|---|---|---|---|---|---|
| 5336 | ENSMUSG00000022516 | Nudt16l1 | 66911 [Gene Symbol: Nudt16l1] [Locus Tag: ] [Chromosome: 16] [Map Location: 16 A1 | 2.065615142 | 0.004384 | 0.00989 |
| 8611 | ENSMUSG00000038007 | Acer2 | 230379 [Gene Symbol: Acer2] [Locus Tag: ] [Chromosome: 4] [Map Location: 4\|4 C4] [D | 2.065567244 | 0.010385 | 0.021541 |
| 8488 | ENSMUSG00000002996 | Hbp1 | 73389 [Gene Symbol: Hbp1] [Locus Tag: ] [Chromosome: 12] [Map Location: 12\|12 A3] | 2.065264623 | 8.45E-05 | 0.000269 |
| 6769 | ENSMUSG00000021598 | Med10 | 28077 [Gene Symbol: Med10] [Locus Tag: ] [Chromosome: 13] [Map Location: 13 C1\|13 | 2.065040408 | 0.001495 | 0.003749 |
| 4007 | ENSMUSG00000039285 | Azi2 | 27215 [Gene Symbol: Azi2] [Locus Tag: ] [Chromosome: 9] [Map Location: 9\|9 F3] [Desc | 2.064205042 | 2.28E-05 | 8.14E-05 |
| 8745 | ENSMUSG00000056708 | Ier5 | 15939 [Gene Symbol: Ier5] [Locus Tag: ] [Chromosome: 1] [Map Location: 1 G3\|1 66.45 | 2.063882313 | 0.004349 | 0.009823 |
| 14297 | ENSMUSG00000027777 | Schip1 | 30953 [Gene Symbol: Schip1] [Locus Tag: ] [Chromosome: 3] [Map Location: 3\|3 E2] [D | 2.063746409 | 8.63E-06 | 3.32E-05 |
| 9613 | ENSMUSG00000021264 | Yy1 | 22632 [Gene Symbol: Yy1] [Locus Tag: ] [Chromosome: 12] [Map Location: 12 F1\|12 59 | 2.062849436 | 2.75E-05 | 9.68E-05 |
| 16729 | ENSMUSG00000056069 | Fam105a | 223433 [Gene Symbol: Fam105a] [Locus Tag: ] [Chromosome: 15] [Map Location: 15\|1 | 2.062229621 | 0.005876 | 0.012886 |
| 13443 | ENSMUSG00000037286 | Stag1 | 20842 [Gene Symbol: Stag1] [Locus Tag: ] [Chromosome: 9] [Map Location: 9\|9 F1] [Des | 2.062054219 | 1.75E-05 | 6.38E-05 |
| 9989 | ENSMUSG00000034912 | Mdga2 | 320772 [Gene Symbol: Mdga2] [Locus Tag: ] [Chromosome: 12] [Map Location: 12 C2\|1 | 2.061438124 | 0.000224 | 0.000659 |
| 14124 | ENSMUSG00000031770 | Herpud1 | 64209 [Gene Symbol: Herpud1] [Locus Tag: ] [Chromosome: 8] [Map Location: 8\|8 C5] | 2.061182796 | 0.00048 | 0.001327 |
| 13123 | ENSMUSG00000028483 | Snapc3 | 77634 [Gene Symbol: Snapc3] [Locus Tag: ] [Chromosome: 4] [Map Location: 4\|4 C3] [D | 2.061055575 | 6.54E-06 | 2.56E-05 |
| 2447 | ENSMUSG00000024283 | Wac | 225131 [Gene Symbol: Wac] [Locus Tag: ] [Chromosome: 18] [Map Location: 18\|18 A1] | 2.060085854 | 9.58E-06 | 3.65E-05 |
| 15761 | ENSMUSG00000020364 | Zfp354a | 21408 [Gene Symbol: Zfp354a] [Locus Tag: ] [Chromosome: 11] [Map Location: 11 B1.3 | 2.059466461 | 0.003187 | 0.007428 |
| 4011 | ENSMUSG00000030960 | Mettl10 | 72096 [Gene Symbol: Mettl10] [Locus Tag: ] [Chromosome: 7] [Map Location: 7\|7 F4] [ | 2.059397916 | 0.001998 | 0.004868 |
| 17528 | ENSMUSG00000040396 | Abhd13 | 68904 [Gene Symbol: Abhd13] [Locus Tag: ] [Chromosome: 8] [Map Location: 8\|8 A2] [ | 2.058452779 | 4.16E-05 | 0.000141 |
| 12199 | ENSMUSG00000075000 | Nrbf2 | 641340 [Gene Symbol: Nrbf2] [Locus Tag: ] [Chromosome: 10] [Map Location: 10\|10 B | 2.05843988 | 0.000468 | 0.001298 |
| 2641 | ENSMUSG00000028920 | Fbxo42 | 213499 [Gene Symbol: Fbxo42] [Locus Tag: ] [Chromosome: 4] [Map Location: 4\|4 D3] | 2.058161796 | 1.81E-05 | 6.59E-05 |
| 10389 | ENSMUSG00000094483 | Purb | 19291 [Gene Symbol: Purb] [Locus Tag: ] [Chromosome: 11] [Map Location: 11 A1\|11 4 | 2.057391015 | 8.56E-06 | 3.29E-05 |
| 3752 | ENSMUSG00000049439 | Cyp20a1 | 77951 [Gene Symbol: Cyp20a1] [Locus Tag: ] [Chromosome: 1] [Map Location: 1\|1 C2] | 2.056326611 | 0.004557 | 0.01024 |
| 2653 | ENSMUSG00000049184 | Purg | 75029 [Gene Symbol: Purg] [Locus Tag: ] [Chromosome: 8] [Map Location: 8\|8 A4] [Des | 2.056291035 | 0.005459 | 0.012053 |
| 16693 | ENSMUSG00000020864 | Ankrd40 | 71452 [Gene Symbol: Ankrd40] [Locus Tag: ] [Chromosome: 11] [Map Location: 11\|11 | 2.055295275 | 0.000231 | 0.000678 |
| 16784 | ENSMUSG00000036052 | Dnajb5 | 56323 [Gene Symbol: Dnajb5] [Locus Tag: ] [Chromosome: 4] [Map Location: 4\|4 B1] [D | 2.055046563 | 9.60E-06 | 3.66E-05 |
| 15449 | ENSMUSG00000027326 | Casc5 | 76464 [Gene Symbol: Casc5] [Locus Tag: ] [Chromosome: 2] [Map Location: 2\|2 E5] [De | 2.053383756 | 0.007057 | 0.015189 |
| 13236 | ENSMUSG00000023980 | Taf8 | 63856 [Gene Symbol: Taf8] [Locus Tag: ] [Chromosome: 17] [Map Location: 17\|17 C] [D | 2.052995806 | 0.000138 | 0.000424 |
| 12901 | ENSMUSG00000031425 | Plp1 | 18823 [Gene Symbol: Plp1] [Locus Tag: ] [Chromosome: X] [Map Location: X 59.1 cM\|X | 2.051838327 | 0.001988 | 0.004846 |
| 17128 | ENSMUSG00000073139 | BC023829 | 236848 [Gene Symbol: BC023829] [Locus Tag: ] [Chromosome: X] [Map Location: X\|X A | 2.051418487 | 0.000866 | 0.002283 |
| 16278 | ENSMUSG00000025616 | Usp16 | 74112 [Gene Symbol: Usp16] [Locus Tag: ] [Chromosome: 16] [Map Location: 16\|16 C3 | 2.050797758 | 0.001651 | 0.004102 |
| 9294 | ENSMUSG00000039081 | Zfp503 | 218820 [Gene Symbol: Zfp503] [Locus Tag: ] [Chromosome: 14] [Map Location: 14 A3\| | 2.050795101 | 0.001484 | 0.003723 |
| 4030 | ENSMUSG00000074643 | Cpne1 | 266692 [Gene Symbol: Cpne1] [Locus Tag: ] [Chromosome: 2] [Map Location: 2\|2 H2] [ | 2.050301473 | 2.06E-05 | 7.42E-05 |
| 1983 | ENSMUSG00000041225 | Arhgap12 | 75415 [Gene Symbol: Arhgap12] [Locus Tag: ] [Chromosome: 18] [Map Location: 18\|18 | 2.050013909 | 0.000174 | 0.000523 |
| 4010 | ENSMUSG00000092094 | Zfp804b | 207618 [Gene Symbol: Zfp804b] [Locus Tag: ] [Chromosome: 5] [Map Location: 5\|5 A1] | 2.04975765 | 0.007801 | 0.016649 |
| 2237 | ENSMUSG00000035868 | 3110052M02Rik | 73229 [Gene Symbol: 3110052M02Rik] [Locus Tag: ] [Chromosome: 17] [Map Location | 2.049490876 | 0.018683 | 0.036294 |
| 14539 | ENSMUSG00000078676 | Casc3 | 192160 [Gene Symbol: Casc3] [Locus Tag: ] [Chromosome: 11] [Map Location: 11\|11 D | 2.048985832 | 3.38E-05 | 0.000117 |
| 8930 | ENSMUSG00000037572 | Wdhd1 | 218973 [Gene Symbol: Wdhd1] [Locus Tag: ] [Chromosome: 14] [Map Location: 14\|14 | 2.045622212 | 0.0059 | 0.01293 |
| 9674 | ENSMUSG00000022378 | Fam49b | 223601 [Gene Symbol: Fam49b] [Locus Tag: ] [Chromosome: 15] [Map Location: 15\|15 | 2.044899639 | 0.001683 | 0.004171 |
| 18094 | ENSMUSG00000091002 | Tcerg1l | 70571 [Gene Symbol: Tcerg1l] [Locus Tag: ] [Chromosome: 7] [Map Location: 7\|7 F4] [D | 2.044884664 | 0.005338 | 0.011814 |
| 4980 | ENSMUSG00000052837 | Junb | 16477 [Gene Symbol: Junb] [Locus Tag: ] [Chromosome: 8] [Map Location: 8 41.41 cM\| | 2.044876552 | 0.002901 | 0.006817 |
| 13319 | ENSMUSG00000026499 | Acbd3 | 170760 [Gene Symbol: Acbd3] [Locus Tag: ] [Chromosome: 1] [Map Location: 1 H4\|1 8 | 2.044837119 | 3.87E-05 | 0.000132 |
| 11333 | ENSMUSG00000016150 | Tenm1 | 23963 [Gene Symbol: Tenm1] [Locus Tag: ] [Chromosome: X] [Map Location: X\|X A2] [D | 2.044079864 | 0.000128 | 0.000393 |
| 33 | ENSMUSG00000021111 | Papola | 18789 [Gene Symbol: Papola] [Locus Tag: ] [Chromosome: 12] [Map Location: 12\|12 F1 | 2.043891877 | 1.56E-05 | 5.76E-05 |
| 4067 | ENSMUSG00000029169 | Dhx15 | 13204 [Gene Symbol: Dhx15] [Locus Tag: ] [Chromosome: 5] [Map Location: 5\|5 C1] [D | 2.043612504 | 4.02E-05 | 0.000137 |
| 1206 | ENSMUSG00000006057 | Atp5g1 | 11951 [Gene Symbol: Atp5g1] [Locus Tag: ] [Chromosome: 11] [Map Location: 11\|11 D | 2.043387386 | 0.006561 | 0.01422 |
| 8565 | ENSMUSG00000040782 | Rfwd2 | 26374 [Gene Symbol: Rfwd2] [Locus Tag: ] [Chromosome: 1] [Map Location: 1\|1 C3] [De | 2.043178586 | 3.44E-05 | 0.000118 |
| 363 | ENSMUSG00000062908 | Acadm | 11364 [Gene Symbol: Acadm] [Locus Tag: ] [Chromosome: 3] [Map Location: 3 H3\|3 78 | 2.042798877 | 0.002867 | 0.006744 |
| 15488 | ENSMUSG00000032902 | Slc16a1 | 20501 [Gene Symbol: Slc16a1] [Locus Tag: ] [Chromosome: 3] [Map Location: 3\|3 F2.2] | 2.042554442 | 0.000934 | 0.002444 |
| 282 | ENSMUSG00000052459 | Atp6v1a | 11964 [Gene Symbol: Atp6v1a] [Locus Tag: ] [Chromosome: 16] [Map Location: 16\|16 | 2.042344962 | 3.33E-05 | 0.000115 |
| 15550 | ENSMUSG00000054423 | Cadps | 27062 [Gene Symbol: Cadps] [Locus Tag: ] [Chromosome: 14] [Map Location: 14\|14 A2 | 2.041045059 | 0.000444 | 0.001237 |
| 7869 | ENSMUSG00000073968 | Trim68 | 101700 [Gene Symbol: Trim68] [Locus Tag: ] [Chromosome: 7] [Map Location: 7\|7 E3] [ | 2.040757314 | 0.018927 | 0.036695 |
| 13042 | ENSMUSG00000020193 | Zpbp | 53604 [Gene Symbol: Zpbp] [Locus Tag: ] [Chromosome: 11] [Map Location: 11\|11 A2] | 2.040016196 | 0.017773 | 0.034699 |
| 12967 | ENSMUSG00000028291 | Akirin2 | 433693 [Gene Symbol: Akirin2] [Locus Tag: ] [Chromosome: 4] [Map Location: 4\|4 A5] [ | 2.038644007 | 5.93E-05 | 0.000194 |
| 10041 | ENSMUSG00000042942 | Greb1l | 381157 [Gene Symbol: Greb1l] [Locus Tag: ] [Chromosome: 18] [Map Location: 18\|18 A | 2.037636704 | 0.002347 | 0.005627 |
| 2559 | ENSMUSG00000038301 | Snx10 | 71982 [Gene Symbol: Snx10] [Locus Tag: ] [Chromosome: 6] [Map Location: 6\|6 B3] [De | 2.035332263 | 0.000181 | 0.000542 |
| 1299 | ENSMUSG00000040321 | Zfp770 | 228491 [Gene Symbol: Zfp770] [Locus Tag: ] [Chromosome: 2] [Map Location: 2\|2 E4] [ | 2.035106346 | 1.78E-05 | 6.48E-05 |
| 1948 | ENSMUSG00000047490 | 4932411E22Rik | 214604 [Gene Symbol: 4932411E22Rik] [Locus Tag: ] [Chromosome: 11] [Map Location | 2.034978067 | 0.003511 | 0.008096 |
| 17419 | ENSMUSG00000046603 | Tcaim | 382117 [Gene Symbol: Tcaim] [Locus Tag: ] [Chromosome: 9] [Map Location: 9 F4\|9 73 | 2.034530702 | 0.003546 | 0.008168 |
| 3833 | ENSMUSG00000038280 | Ostm1 | 14628 [Gene Symbol: Ostm1] [Locus Tag: ] [Chromosome: 10] [Map Location: 10 B2\|10 | 2.033847244 | 0.001072 | 0.002773 |
| 9480 | ENSMUSG00000031285 | Dcx | 13193 [Gene Symbol: Dcx] [Locus Tag: ] [Chromosome: X] [Map Location: X\|X F2] [Desc | 2.03366291 | 1.37E-05 | 5.11E-05 |
| 17604 | ENSMUSG00000025969 | Nrp2 | 18187 [Gene Symbol: Nrp2] [Locus Tag: ] [Chromosome: 1] [Map Location: 1\|1 C2] [Des | 2.032592055 | 0.000203 | 0.000605 |
| 9841 | ENSMUSG00000045083 | Lingo2 | 242384 [Gene Symbol: Lingo2] [Locus Tag: ] [Chromosome: 4] [Map Location: 4\|4 A5] [ | 2.031813332 | 0.002078 | 0.00504 |
| 412 | ENSMUSG00000001017 | Chtop | 66511 [Gene Symbol: Chtop] [Locus Tag: ] [Chromosome: 3] [Map Location: 3\|3 F2] [De | 2.03171625 | 0.000305 | 0.000878 |
| 14548 | ENSMUSG00000021715 | Cwc27 | 67285 [Gene Symbol: Cwc27] [Locus Tag: ] [Chromosome: 13] [Map Location: 13\|13 D | 2.03171598 | 0.001436 | 0.003619 |
| 4561 | ENSMUSG00000044148 | 1810030O07Rik | 69155 [Gene Symbol: 1810030O07Rik] [Locus Tag: ] [Chromosome: X] [Map Location: | 2.031293958 | 0.000589 | 0.001602 |
| 6334 | ENSMUSG00000048720 | Tbc1d12 | 209478 [Gene Symbol: Tbc1d12] [Locus Tag: ] [Chromosome: 19] [Map Location: 19\|1 | 2.029818307 | 5.00E-05 | 0.000167 |
| 5887 | ENSMUSG00000064202 | 4430402I18Rik | 381218 [Gene Symbol: 4430402I18Rik] [Locus Tag: ] [Chromosome: 19] [Map Location | 2.029344705 | 4.17E-05 | 0.000141 |
| 7492 | ENSMUSG00000027810 | Eif2a | 229317 [Gene Symbol: Eif2a] [Locus Tag: ] [Chromosome: 3] [Map Location: 3 D\|3 28.5 | 2.029099177 | 0.000234 | 0.000688 |
| 12254 | ENSMUSG00000028556 | Dock7 | 67299 [Gene Symbol: Dock7] [Locus Tag: ] [Chromosome: 4] [Map Location: 4 C6\|4 45. | 2.028657261 | 1.36E-05 | 5.06E-05 |
| 7000 | ENSMUSG00000061904 | Slc25a3; LOC102641 | 18674 [Gene Symbol: Slc25a3] [Locus Tag: ] [Chromosome: 10] [Map Location: 10\|10 C | 2.028020675 | 0.000103 | 0.000323 |
| 16040 | ENSMUSG00000018648 | Dusp14 | 56405 [Gene Symbol: Dusp14] [Locus Tag: ] [Chromosome: 11] [Map Location: 11 C\|11 | 2.026734228 | 0.012636 | 0.025636 |
| 2924 | ENSMUSG00000044647 | Csrnp3 | 77771 [Gene Symbol: Csrnp3] [Locus Tag: ] [Chromosome: 2] [Map Location: 2\|2 C1.3] | 2.026477266 | 1.25E-05 | 4.67E-05 |
| 17876 | ENSMUSG00000038893 | Fam117a | 215512 [Gene Symbol: Fam117a] [Locus Tag: ] [Chromosome: 11] [Map Location: 11\|1 | 2.026188233 | 0.005575 | 0.01229 |
| 725 | ENSMUSG00000030942 | Thumpd1 | 233802 [Gene Symbol: Thumpd1] [Locus Tag: ] [Chromosome: 7] [Map Location: 7 F2\|7 | 2.025981712 | 0.00016 | 0.000485 |
| 15193 | ENSMUSG00000019982 | Myb | 17863 [Gene Symbol: Myb] [Locus Tag: ] [Chromosome: 10] [Map Location: 10 A3\|10 9 | 2.025346169 | 0.006088 | 0.013297 |
| 6423 | ENSMUSG00000005683 | Cs | 12974 [Gene Symbol: Cs] [Locus Tag: ] [Chromosome: 10] [Map Location: 10\|10 D3] [De | 2.025067465 | 3.53E-05 | 0.000121 |
| 7541 | ENSMUSG00000021470 | Ercc6l2 | 76251 [Gene Symbol: Ercc6l2] [Locus Tag: ] [Chromosome: 13] [Map Location: 13\|13 B | 2.024767621 | 3.26E-05 | 0.000113 |
| 4239 | ENSMUSG00000064127 | Med14 | 26896 [Gene Symbol: Med14] [Locus Tag: ] [Chromosome: X] [Map Location: X\|X A1.1] | 2.024312272 | 8.47E-05 | 0.000269 |
| 7127 | ENSMUSG00000030105 | Arl8b | 67166 [Gene Symbol: Arl8b] [Locus Tag: ] [Chromosome: 6] [Map Location: 6\|6 E2] [Des | 2.02403357 | 2.83E-05 | 9.92E-05 |
| 3811 | ENSMUSG00000051978 | Erich1 | 234086 [Gene Symbol: Erich1] [Locus Tag: ] [Chromosome: 8] [Map Location: 8\|8 A1.1] | 2.023452272 | 0.003738 | 0.008569 |
| 12963 | ENSMUSG00000036898 | Zfp157 | 72154 [Gene Symbol: Zfp157] [Locus Tag: ] [Chromosome: 5] [Map Location: 5\|5 G2] [D | 2.023449999 | 5.24E-05 | 0.000174 |
| 17766 | ENSMUSG00000024074 | Crim1 | 50766 [Gene Symbol: Crim1] [Locus Tag: ] [Chromosome: 17] [Map Location: 17\|17 E2 | 2.023273027 | 1.43E-05 | 5.31E-05 |
| 8103 | ENSMUSG00000020674 | Pxdn | 69675 [Gene Symbol: Pxdn] [Locus Tag: ] [Chromosome: 12] [Map Location: 12\|12 A2] | 2.022653751 | 4.06E-05 | 0.000138 |
| 18269 | ENSMUSG00000019894 | Slc6a15 | 103098 [Gene Symbol: Slc6a15] [Locus Tag: ] [Chromosome: 10] [Map Location: 10\|10 | 2.021889968 | 3.40E-05 | 0.000117 |
| 4408 | ENSMUSG00000029153 | Ociad2 | 433904 [Gene Symbol: Ociad2] [Locus Tag: ] [Chromosome: 5] [Map Location: 5\|5 D] [D | 2.021743877 | 0.004407 | 0.009938 |

| | | | | | | |
|---|---|---|---|---|---|---|
| 5016 | ENSMUSG00000052293 | Taf9 | 108143 [Gene Symbol: Taf9] [Locus Tag: ] [Chromosome: 13] [Map Location: 13\|13 D1] | 2.021494302 | 0.000236 | 0.000694 |
| 13699 | ENSMUSG00000025551 | Fgf14 | 14169 [Gene Symbol: Fgf14] [Locus Tag: ] [Chromosome: 14] [Map Location: 14 E5\|14 ( | 2.020354638 | 0.0037 | 0.008489 |
| 5215 | ENSMUSG00000069911 | Fam196b | 574403 [Gene Symbol: Fam196b] [Locus Tag: ] [Chromosome: 11] [Map Location: 11 A4 | 2.020233548 | 0.000126 | 0.000389 |
| 6168 | ENSMUSG00000011831 | Evi5 | 14020 [Gene Symbol: Evi5] [Locus Tag: ] [Chromosome: 5] [Map Location: 5 F\|5 52.23 c | 2.020211721 | 0.000977 | 0.002543 |
| 6337 | ENSMUSG00000033883 | D3Ertd254e | 241944 [Gene Symbol: D3Ertd254e] [Locus Tag: ] [Chromosome: 3] [Map Location: 3 B\| | 2.019175933 | 5.74E-05 | 0.000188 |
| 8953 | ENSMUSG00000102422 | Gm21949 | 100505386 [Gene Symbol: Gm21949] [Locus Tag: ] [Chromosome: 3] [Map Location: 3 | 2.018945745 | 1.43E-05 | 5.31E-05 |
| 14963 | ENSMUSG00000026806 | Ddx31 | 227674 [Gene Symbol: Ddx31] [Locus Tag: ] [Chromosome: 2] [Map Location: 2\|2 A3] [ | 2.018641528 | 0.001024 | 0.002658 |
| 9097 | ENSMUSG00000032745 | Gpbp1 | 73274 [Gene Symbol: Gpbp1] [Locus Tag: ] [Chromosome: 13] [Map Location: 13\|13 D2 | 2.018613727 | 3.82E-05 | 0.00013 |
| 11365 | ENSMUSG00000031931 | Ankrd49 | 56503 [Gene Symbol: Ankrd49] [Locus Tag: ] [Chromosome: 9] [Map Location: 9\|9 A3] [ | 2.018236433 | 0.002563 | 0.006106 |
| 8127 | ENSMUSG00000054855 | Rnd1 | 223881 [Gene Symbol: Rnd1] [Locus Tag: ] [Chromosome: 15] [Map Location: 15\|15 F1 | 2.018160688 | 0.000991 | 0.002579 |
| 13531 | ENSMUSG00000020515 | Cnot8 | 69125 [Gene Symbol: Cnot8] [Locus Tag: ] [Chromosome: 11] [Map Location: 11\|11 B1 | 2.017773815 | 6.09E-05 | 0.000199 |
| 1848 | ENSMUSG00000038668 | Lpar1 | 14745 [Gene Symbol: Lpar1] [Locus Tag: ] [Chromosome: 4] [Map Location: 4 B3\|4 32.2 | 2.017451116 | 0.000438 | 0.001223 |
| 3896 | ENSMUSG00000021613 | Hapln1 | 12950 [Gene Symbol: Hapln1] [Locus Tag: ] [Chromosome: 13] [Map Location: 13 C3\|1: | 2.017289231 | 0.016404 | 0.032346 |
| 13237 | ENSMUSG00000022849 | Hspbap1 | 66667 [Gene Symbol: Hspbap1] [Locus Tag: ] [Chromosome: 16] [Map Location: 16\|16 | 2.016416931 | 0.001653 | 0.004105 |
| 17307 | ENSMUSG00000024537 | Psmg2 | 107047 [Gene Symbol: Psmg2] [Locus Tag: ] [Chromosome: 18] [Map Location: 18\|18 E | 2.015815293 | 0.004108 | 0.009322 |
| 2710 | ENSMUSG00000034161 | Scx | 20289 [Gene Symbol: Scx] [Locus Tag: ] [Chromosome: 15] [Map Location: 15 D3\|15 35 | 2.015788631 | 0.016099 | 0.031799 |
| 7162 | ENSMUSG00000017707 | Serinc3 | 26943 [Gene Symbol: Serinc3] [Locus Tag: ] [Chromosome: 2] [Map Location: 2\|2 H3] [D | 2.01558073 | 0.000122 | 0.000378 |
| 12669 | ENSMUSG00000027108 | Ola1 | 67059 [Gene Symbol: Ola1] [Locus Tag: ] [Chromosome: 2] [Map Location: 2\|2 C3] [Des | 2.014962211 | 5.15E-05 | 0.000171 |
| 6659 | ENSMUSG00000021589 | Rhobtb3 | 73296 [Gene Symbol: Rhobtb3] [Locus Tag: ] [Chromosome: 13] [Map Location: 13\|13 | 2.014907693 | 0.000127 | 0.000392 |
| 2853 | ENSMUSG00000031010 | Usp9x | 22284 [Gene Symbol: Usp9x] [Locus Tag: ] [Chromosome: X] [Map Location: X A1.1\|X 7. | 2.014772291 | 1.67E-05 | 6.12E-05 |
| 15507 | ENSMUSG00000057367 | Birc2 | 11797 [Gene Symbol: Birc2] [Locus Tag: ] [Chromosome: 9] [Map Location: 9\|9 A1] [Des | 2.014256653 | 0.000374 | 0.001059 |
| 2017 | ENSMUSG00000024542 | Cep192 | 70799 [Gene Symbol: Cep192] [Locus Tag: ] [Chromosome: 18] [Map Location: 18\|18 E | 2.013632116 | 0.000412 | 0.001157 |
| 12573 | ENSMUSG00000102627 | Snurf | 84704 [Gene Symbol: Snurf] [Locus Tag: ] [Chromosome: 7] [Map Location: 7\|7 C] [Desc | 2.012912264 | 0.003624 | 0.008337 |
| 15609 | ENSMUSG00000038368 | Focad | 230393 [Gene Symbol: Focad] [Locus Tag: ] [Chromosome: 4] [Map Location: 4\|4 C4] [D | 2.012455099 | 6.18E-05 | 0.000202 |
| 6574 | ENSMUSG00000020275 | Rel | 19696 [Gene Symbol: Rel] [Locus Tag: ] [Chromosome: 11] [Map Location: 11 A3.2\|11 1 | 2.01206147 | 0.017006 | 0.033354 |
| 14611 | ENSMUSG00000044641 | Pard6b | 58220 [Gene Symbol: Pard6b] [Locus Tag: ] [Chromosome: 2] [Map Location: 2\|2 H3] [D | 2.011454915 | 0.000594 | 0.001614 |
| 15502 | ENSMUSG00000026774 | 4931423N10Rik | 70981 [Gene Symbol: 4931423N10Rik] [Locus Tag: ] [Chromosome: 2] [Map Location: 2 | 2.011220055 | 0.02148 | 0.041058 |
| 7306 | ENSMUSG00000042406 | Atf4 | 11911 [Gene Symbol: Atf4] [Locus Tag: ] [Chromosome: 15] [Map Location: 15 E1\|15 37 | 2.011060671 | 3.51E-05 | 0.000121 |
| 5542 | ENSMUSG00000023852 | Chd1 | 12648 [Gene Symbol: Chd1] [Locus Tag: ] [Chromosome: 17] [Map Location: 17 A2\|17 8 | 2.008200902 | 0.000573 | 0.001564 |
| 9368 | ENSMUSG00000031935 | Med17 | 234959 [Gene Symbol: Med17] [Locus Tag: ] [Chromosome: 9] [Map Location: 9\|9 A2] [ | 2.007122302 | 0.000656 | 0.001768 |
| 7489 | ENSMUSG00000032580 | Rbm5 | 83486 [Gene Symbol: Rbm5] [Locus Tag: ] [Chromosome: 9] [Map Location: 9\|9 F1] [De | 2.00692319 | 0.000343 | 0.000979 |
| 6101 | ENSMUSG00000046449 | C77370 | 245555 [Gene Symbol: C77370] [Locus Tag: ] [Chromosome: X] [Map Location: X\|X D] [ | 2.00651655 | 1.04E-05 | 3.94E-05 |
| 10455 | ENSMUSG00000020955 | Ap4s1 | 11782 [Gene Symbol: Ap4s1] [Locus Tag: ] [Chromosome: 12] [Map Location: 12\|12 C1 | 2.006463861 | 0.001816 | 0.004467 |
| 96 | ENSMUSG00000061288 | Taok3 | 330177 [Gene Symbol: Taok3] [Locus Tag: ] [Chromosome: 5] [Map Location: 5\|5 F] [De | 2.00514319 | 0.002488 | 0.005942 |
| 15100 | ENSMUSG00000027109 | Sp3 | 20687 [Gene Symbol: Sp3] [Locus Tag: ] [Chromosome: 2] [Map Location: 2\|2 C3] [Desc | 2.005004839 | 1.89E-05 | 6.86E-05 |
| 564 | ENSMUSG00000021852 | Slc35f4 | 75288 [Gene Symbol: Slc35f4] [Locus Tag: ] [Chromosome: 14] [Map Location: 14\|14 C | 2.00486445 | 0.023429 | 0.044342 |
| 15010 | ENSMUSG00000022828 | Gtf2e1 | 74197 [Gene Symbol: Gtf2e1] [Locus Tag: ] [Chromosome: 16] [Map Location: 16\|16 B3 | 2.004396941 | 0.000736 | 0.001965 |
| 8329 | ENSMUSG00000040018 | Cox15 | 226139 [Gene Symbol: Cox15] [Locus Tag: ] [Chromosome: 19] [Map Location: 19\|19 C | 2.004243672 | 0.000158 | 0.000481 |
| 5744 | ENSMUSG00000039630 | Hnrnpu | 51810 [Gene Symbol: Hnrnpu] [Locus Tag: ] [Chromosome: 1] [Map Location: 1\|1 H4] [ | 2.003606546 | 0.00054 | 0.001481 |
| 7803 | ENSMUSG00000020720 | Psmd12 | 66997 [Gene Symbol: Psmd12] [Locus Tag: ] [Chromosome: 11] [Map Location: 11\|11 E | 2.003471015 | 0.000277 | 0.000801 |
| 11699 | ENSMUSG00000013663 | Pten | 19211 [Gene Symbol: Pten] [Locus Tag: ] [Chromosome: 19] [Map Location: 19 C1\|19 2 | 2.003194064 | 1.32E-05 | 4.94E-05 |
| 4573 | ENSMUSG00000037376 | Trmt6 | 66926 [Gene Symbol: Trmt6] [Locus Tag: ] [Chromosome: 2] [Map Location: 2\|2 F2] [De | 2.003164273 | 0.000822 | 0.002173 |
| 1013 | ENSMUSG00000028524 | Sgip1 | 73094 [Gene Symbol: Sgip1] [Locus Tag: ] [Chromosome: 4] [Map Location: 4\|4 C6] [De | 2.003135477 | 0.000212 | 0.000628 |
| 10345 | ENSMUSG00000041426 | Hibch | 227095 [Gene Symbol: Hibch] [Locus Tag: ] [Chromosome: 1] [Map Location: 1\|1 C1.1] | 2.003122682 | 0.006358 | 0.01382 |
| 8767 | ENSMUSG00000021306 | Gpr137b | 83924 [Gene Symbol: Gpr137b] [Locus Tag: ] [Chromosome: 13] [Map Location: 13 A1\| | 2.002118998 | 0.002028 | 0.004932 |
| 3218 | ENSMUSG00000017831 | Rab5a | 271457 [Gene Symbol: Rab5a] [Locus Tag: ] [Chromosome: 17] [Map Location: 17 C\|17 | 2.002010166 | 8.47E-05 | 0.000269 |
| 17895 | ENSMUSG00000022314 | Rad21 | 19357 [Gene Symbol: Rad21] [Locus Tag: ] [Chromosome: 15] [Map Location: 15\|15 D3 | 2.001229179 | 3.40E-05 | 0.000117 |
| 13805 | ENSMUSG00000031703 | Itfg1 | 71927 [Gene Symbol: Itfg1] [Locus Tag: ] [Chromosome: 8] [Map Location: 8 41.61 cM\| | 2.000908285 | 4.91E-05 | 0.000164 |
| 3567 | ENSMUSG00000024145 | Pigf | 18701 [Gene Symbol: Pigf] [Locus Tag: ] [Chromosome: 17] [Map Location: 17 56.9 cM\| | 1.999850626 | 9.15E-05 | 0.000289 |
| 5983 | ENSMUSG00000037573 | Tob1 | 22057 [Gene Symbol: Tob1] [Locus Tag: ] [Chromosome: 11] [Map Location: 11\|11 C] [ | 1.999630008 | 3.11E-05 | 0.000108 |
| 1372 | ENSMUSG00000021614 | Vcan | 13003 [Gene Symbol: Vcan] [Locus Tag: ] [Chromosome: 13] [Map Location: 13 C3\|13 4 | 1.999455982 | 2.11E-05 | 7.58E-05 |
| 15699 | ENSMUSG00000024843 | Chka | 12660 [Gene Symbol: Chka] [Locus Tag: ] [Chromosome: 19] [Map Location: 19 A\|19 3. | 1.998745221 | 0.000109 | 0.00034 |
| 10370 | ENSMUSG00000024639 | Gnaq | 14682 [Gene Symbol: Gnaq] [Locus Tag: ] [Chromosome: 19] [Map Location: 19 A\|19 1: | 1.998289702 | 1.19E-05 | 4.46E-05 |
| 17449 | ENSMUSG00000037062 | Sh3glb1 | 54673 [Gene Symbol: Sh3glb1] [Locus Tag: ] [Chromosome: 3] [Map Location: 3\|3 H2] [ | 1.997993935 | 2.26E-05 | 8.09E-05 |
| 12949 | ENSMUSG00000021770 | Samd8 | 67630 [Gene Symbol: Samd8] [Locus Tag: ] [Chromosome: 14] [Map Location: 14\|14 B] | 1.997082663 | 2.90E-05 | 0.000101 |
| 15308 | ENSMUSG00000041684 | Bivm | 246229 [Gene Symbol: Bivm] [Locus Tag: ] [Chromosome: 1] [Map Location: 1\|1 C1.1] [ | 1.995378797 | 0.000147 | 0.000448 |
| 3055 | ENSMUSG00000021629 | Slc30a5 | 69048 [Gene Symbol: Slc30a5] [Locus Tag: ] [Chromosome: 13] [Map Location: 13\|13 D | 1.995109303 | 9.07E-05 | 0.000286 |
| 748 | ENSMUSG00000056870 | Gulp1 | 70676 [Gene Symbol: Gulp1] [Locus Tag: ] [Chromosome: 1] [Map Location: 1\|1 C1] [De | 1.995101753 | 0.001575 | 0.00393 |
| 15255 | ENSMUSG00000044921 | Rassf9 | 237504 [Gene Symbol: Rassf9] [Locus Tag: ] [Chromosome: 10] [Map Location: 10\|10 D | 1.994699053 | 0.024758 | 0.04665 |
| 13625 | ENSMUSG00000024997 | Prdx3 | 11757 [Gene Symbol: Prdx3] [Locus Tag: ] [Chromosome: 19] [Map Location: 19 D3\|19 | 1.993905321 | 0.006563 | 0.014223 |
| 1485 | ENSMUSG00000068184 | Ndufaf2 | 75597 [Gene Symbol: Ndufaf2] [Locus Tag: ] [Chromosome: 13] [Map Location: 13\|13 D | 1.993329135 | 0.005255 | 0.011648 |
| 4769 | ENSMUSG00000063626 | Unc5d | 210801 [Gene Symbol: Unc5d] [Locus Tag: ] [Chromosome: 8] [Map Location: 8\|8 A2-A: | 1.992647956 | 5.02E-05 | 0.000167 |
| 13584 | ENSMUSG00000014554 | Dguok | 27369 [Gene Symbol: Dguok] [Locus Tag: ] [Chromosome: 6] [Map Location: 6 C3\|6 35. | 1.991361785 | 0.003999 | 0.009106 |
| 5247 | ENSMUSG00000079562 | Maea | 59003 [Gene Symbol: Maea] [Locus Tag: ] [Chromosome: 5] [Map Location: 5\|5 B1] [De | 1.991223709 | 0.000117 | 0.000362 |
| 2260 | ENSMUSG00000038047 | Haus6 | 230376 [Gene Symbol: Haus6] [Locus Tag: ] [Chromosome: 4] [Map Location: 4 C4\|4 40 | 1.991063996 | 0.001333 | 0.003385 |
| 14357 | ENSMUSG00000039128 | Cdc123 | 98828 [Gene Symbol: Cdc123] [Locus Tag: ] [Chromosome: 2] [Map Location: 2\|2 A1] [ | 1.990712888 | 0.000175 | 0.000528 |
| 12150 | ENSMUSG00000050174 | Nudt6 | 229228 [Gene Symbol: Nudt6] [Locus Tag: ] [Chromosome: 3] [Map Location: 3\|3 B] [De | 1.989936455 | 0.009859 | 0.020567 |
| 18270 | ENSMUSG00000027962 | Vcam1 | 22329 [Gene Symbol: Vcam1] [Locus Tag: ] [Chromosome: 3] [Map Location: 3 G1\|3 50 | 1.988179087 | 0.00054 | 0.001481 |
| 10407 | ENSMUSG00000033565 | Rbfox2 | 93686 [Gene Symbol: Rbfox2] [Locus Tag: ] [Chromosome: 15] [Map Location: 15\|15 E1 | 1.987797871 | 0.000146 | 0.000445 |
| 11999 | ENSMUSG00000026127 | Imp4 | 27993 [Gene Symbol: Imp4] [Locus Tag: ] [Chromosome: 1] [Map Location: 1 B\|1 13.22 | 1.986946057 | 0.000263 | 0.000766 |
| 8978 | ENSMUSG00000025950 | Idh1 | 15926 [Gene Symbol: Idh1] [Locus Tag: ] [Chromosome: 1] [Map Location: 1 C2\|1 32.9: | 1.986770559 | 8.43E-05 | 0.000268 |
| 9318 | ENSMUSG00000024253 | Dync2li1 | 213575 [Gene Symbol: Dync2li1] [Locus Tag: ] [Chromosome: 17] [Map Location: 17\|1 | 1.985973926 | 0.004556 | 0.01024 |
| 5865 | ENSMUSG00000021466 | Ptch1 | 19206 [Gene Symbol: Ptch1] [Locus Tag: ] [Chromosome: 13] [Map Location: 13 B3\|13 | 1.98596113 | 4.00E-05 | 0.000136 |
| 16931 | ENSMUSG00000021116 | Eif2s1 | 13665 [Gene Symbol: Eif2s1] [Locus Tag: ] [Chromosome: 12] [Map Location: 12\|12 D1 | 1.985031952 | 0.001648 | 0.004094 |
| 17980 | ENSMUSG00000028444 | Cntfr | 12804 [Gene Symbol: Cntfr] [Locus Tag: ] [Chromosome: 4] [Map Location: 4 A5\|4 21.8 | 1.984888005 | 0.005488 | 0.012108 |
| 9802 | ENSMUSG00000029171 | Pgm1 | 66681 [Gene Symbol: Pgm1] [Locus Tag: ] [Chromosome: 5] [Map Location: 5 32.8 cM\| | 1.983877253 | 0.00097 | 0.002528 |
| 16608 | ENSMUSG00000047604 | Frat2 | 212398 [Gene Symbol: Frat2] [Locus Tag: ] [Chromosome: 19] [Map Location: 19 C3\|19 | 1.983698438 | 0.000585 | 0.001594 |
| 2694 | ENSMUSG00000026383 | Epb41l5 | 226352 [Gene Symbol: Epb41l5] [Locus Tag: ] [Chromosome: 1] [Map Location: 1\|1 E2. | 1.981852603 | 0.000205 | 0.000609 |
| 15158 | ENSMUSG00000060227 | Casc4 | 319996 [Gene Symbol: Casc4] [Locus Tag: ] [Chromosome: 2] [Map Location: 2\|2 E5] [D | 1.98183296 | 2.05E-05 | 7.38E-05 |
| 9354 | ENSMUSG00000018415 | Gid4 | 66771 [Gene Symbol: Gid4] [Locus Tag: ] [Chromosome: 11] [Map Location: 11\|11 B2] | 1.98084152 | 6.12E-05 | 0.0002 |
| 9265 | ENSMUSG00000028261 | Ndufaf4 | 68493 [Gene Symbol: Ndufaf4] [Locus Tag: ] [Chromosome: 4] [Map Location: 4\|4 A3] [ | 1.980374149 | 0.001323 | 0.003361 |
| 14069 | ENSMUSG00000011179 | Odc1 | 18263 [Gene Symbol: Odc1] [Locus Tag: ] [Chromosome: 12] [Map Location: 12 A1.1\|1: | 1.979583702 | 0.004335 | 0.009796 |
| 12832 | ENSMUSG00000022404 | Slc25a17 | 20524 [Gene Symbol: Slc25a17] [Locus Tag: ] [Chromosome: 15] [Map Location: 15\|15 | 1.978615978 | 0.001521 | 0.003804 |
| 17253 | ENSMUSG00000031309 | Rps6ka3 | 110651 [Gene Symbol: Rps6ka3] [Locus Tag: ] [Chromosome: X] [Map Location: X F4\|X | 1.978354638 | 3.31E-05 | 0.000115 |

| | | | | | | |
|---|---|---|---|---|---|---|
| 3117 | ENSMUSG00000049940 | Pgrmc2 | 70804 [Gene Symbol: Pgrmc2] [Locus Tag: ] [Chromosome: 3] [Map Location: 3\|3 C] [De | 1.97791693 | 0.00015 | 0.000457 |
| 6490 | ENSMUSG00000040151 | Hs2st1 | 23908 [Gene Symbol: Hs2st1] [Locus Tag: ] [Chromosome: 3] [Map Location: 3\|3 H2] [D | 1.976376043 | 6.70E-05 | 0.000217 |
| 15737 | ENSMUSG00000079184 | Mphosph8 | 75339 [Gene Symbol: Mphosph8] [Locus Tag: ] [Chromosome: 14] [Map Location: 14\|1 | 1.975996144 | 0.000619 | 0.001677 |
| 16167 | ENSMUSG00000073563 | Csnk1g3 | 70425 [Gene Symbol: Csnk1g3] [Locus Tag: ] [Chromosome: 18] [Map Location: 18\|18 | 1.975132308 | 3.54E-05 | 0.000122 |
| 8972 | ENSMUSG00000021830 | Txndc16 | 70561 [Gene Symbol: Txndc16] [Locus Tag: ] [Chromosome: 14] [Map Location: 14\|14 | 1.974462977 | 0.000101 | 0.000317 |
| 2514 | ENSMUSG00000051256 | Jagn1 | 67767 [Gene Symbol: Jagn1] [Locus Tag: ] [Chromosome: 6] [Map Location: 6\|6 E3] [De | 1.974070474 | 0.009474 | 0.019835 |
| 5214 | ENSMUSG00000010054 | Tusc2 | 80385 [Gene Symbol: Tusc2] [Locus Tag: ] [Chromosome: 9] [Map Location: 9\|9 F1] [De | 1.973377552 | 4.45E-05 | 0.00015 |
| 10263 | ENSMUSG00000040037 | Negr1 | 320840 [Gene Symbol: Negr1] [Locus Tag: ] [Chromosome: 3] [Map Location: 3\|3 H4] [D | 1.973083275 | 0.000106 | 0.00033 |
| 15938 | ENSMUSG00000018372 | Cep95 | 320162 [Gene Symbol: Cep95] [Locus Tag: ] [Chromosome: 11] [Map Location: 11\|11 E | 1.972608511 | 0.011255 | 0.02316 |
| 3851 | ENSMUSG00000015222 | Map2 | 17756 [Gene Symbol: Map2] [Locus Tag: ] [Chromosome: 1] [Map Location: 1 C3\|1 33.4 | 1.972386114 | 1.63E-05 | 5.99E-05 |
| 12373 | ENSMUSG00000074715 | Ccl28 | 56838 [Gene Symbol: Ccl28] [Locus Tag: ] [Chromosome: 13] [Map Location: 13\|13] [De | 1.971445277 | 0.003423 | 0.007911 |
| 18143 | ENSMUSG00000028234 | Rps20 | 67427 [Gene Symbol: Rps20] [Locus Tag: ] [Chromosome: 4] [Map Location: 4\|4 A1] [De | 1.970563174 | 0.000533 | 0.001463 |
| 834 | ENSMUSG00000034621 | Gpatch8 | 237943 [Gene Symbol: Gpatch8] [Locus Tag: ] [Chromosome: 11] [Map Location: 11\|11 | 1.970234374 | 0.000234 | 0.000688 |
| 11088 | ENSMUSG00000041975 | Mettl8 | 228019 [Gene Symbol: Mettl8] [Locus Tag: ] [Chromosome: 2] [Map Location: 2\|2 C2] [ | 1.969949314 | 0.003822 | 0.008741 |
| 1101 | ENSMUSG00000017858 | Ift52 | 245866 [Gene Symbol: Ift52] [Locus Tag: ] [Chromosome: 2] [Map Location: 2\|2 H2] [De | 1.969816379 | 0.000592 | 0.00161 |
| 5897 | ENSMUSG00000020721 | Helz | 78455 [Gene Symbol: Helz] [Locus Tag: ] [Chromosome: 11] [Map Location: 11\|11 E1] [ | 1.96929154 | 0.000177 | 0.000532 |
| 8217 | ENSMUSG00000024513 | Mbd2 | 17191 [Gene Symbol: Mbd2] [Locus Tag: ] [Chromosome: 18] [Map Location: 18 E2\|18 | 1.968761228 | 0.00112 | 0.002887 |
| 15582 | ENSMUSG00000060802 | B2m | 12010 [Gene Symbol: B2m] [Locus Tag: ] [Chromosome: 2] [Map Location: 2 60.55 cM\| | 1.968619032 | 0.010383 | 0.021539 |
| 10383 | ENSMUSG00000031812 | Map1lc3b | 67443 [Gene Symbol: Map1lc3b] [Locus Tag: ] [Chromosome: 8] [Map Location: 8\|8 E1 | 1.967937099 | 0.000122 | 0.000378 |
| 15372 | ENSMUSG00000058440 | Nrf1 | 18181 [Gene Symbol: Nrf1] [Locus Tag: ] [Chromosome: 6] [Map Location: 6 A3.3\|6 12. | 1.967263399 | 0.000193 | 0.000576 |
| 5219 | ENSMUSG00000032462 | Pik3cb | 74769 [Gene Symbol: Pik3cb] [Locus Tag: ] [Chromosome: 9] [Map Location: 9\|9 E4] [D | 1.966680687 | 7.38E-05 | 0.000238 |
| 7566 | ENSMUSG00000030213 | Atf7ip | 54343 [Gene Symbol: Atf7ip] [Locus Tag: ] [Chromosome: 6] [Map Location: 6\|6 G1] [De | 1.966677914 | 0.000262 | 0.000762 |
| 17095 | ENSMUSG00000048616 | Nog | 18121 [Gene Symbol: Nog] [Locus Tag: ] [Chromosome: 11] [Map Location: 11 C\|11 54. | 1.966281462 | 0.004206 | 0.009528 |
| 18383 | ENSMUSG00000020212 | Mdm1 | 17245 [Gene Symbol: Mdm1] [Locus Tag: ] [Chromosome: 10] [Map Location: 10 66.65 | 1.965181581 | 0.000406 | 0.001142 |
| 8990 | ENSMUSG00000021391 | Cenpp | 66336 [Gene Symbol: Cenpp] [Locus Tag: ] [Chromosome: 13] [Map Location: 13\|13 B1 | 1.964898458 | 0.000312 | 0.000895 |
| 18379 | ENSMUSG00000021541 | Trpc7 | 26946 [Gene Symbol: Trpc7] [Locus Tag: ] [Chromosome: 13] [Map Location: 13\|13 B2] | 1.964772122 | 0.000467 | 0.001295 |
| 16117 | ENSMUSG00000040029 | Ipo8 | 320727 [Gene Symbol: Ipo8] [Locus Tag: ] [Chromosome: 6] [Map Location: 6\|6 G3] [De | 1.963807747 | 0.000152 | 0.000462 |
| 5984 | ENSMUSG00000025917 | Cops5 | 26754 [Gene Symbol: Cops5] [Locus Tag: ] [Chromosome: 1] [Map Location: 1 A2\|1 2.2 | 1.963375533 | 0.001331 | 0.003381 |
| 11725 | ENSMUSG00000010392 | Gosr1 | 53334 [Gene Symbol: Gosr1] [Locus Tag: ] [Chromosome: 11] [Map Location: 11\|11 B5 | 1.962728075 | 0.000188 | 0.000562 |
| 3071 | ENSMUSG00000041817 | Fam169a | 320557 [Gene Symbol: Fam169a] [Locus Tag: ] [Chromosome: 13] [Map Location: 13\|1 | 1.962667543 | 0.001816 | 0.004468 |
| 15712 | ENSMUSG00000040570 | Rundc3b | 242819 [Gene Symbol: Rundc3b] [Locus Tag: ] [Chromosome: 5] [Map Location: 5\|5 A1 | 1.962564383 | 7.54E-05 | 0.000242 |
| 18086 | ENSMUSG00000069727 | Gm5595 | 434179 [Gene Symbol: Gm5595] [Locus Tag: ] [Chromosome: 7] [Map Location: 7 B3\|7 | 1.962186558 | 0.012151 | 0.024776 |
| 14431 | ENSMUSG00000030446 | Zfp273 | 212569 [Gene Symbol: Zfp273] [Locus Tag: ] [Chromosome: 13] [Map Location: 13\|13 | 1.961529066 | 0.003744 | 0.008578 |
| 7954 | ENSMUSG00000050830 | Vwc2 | 319922 [Gene Symbol: Vwc2] [Locus Tag: ] [Chromosome: 11] [Map Location: 11\|11 A1 | 1.961112079 | 0.007432 | 0.015935 |
| 16362 | ENSMUSG00000020894 | Vamp2 | 22318 [Gene Symbol: Vamp2] [Locus Tag: ] [Chromosome: 11] [Map Location: 11\|11 B | 1.961078935 | 2.13E-05 | 7.65E-05 |
| 4954 | ENSMUSG00000057236 | Rbbp4 | 19646 [Gene Symbol: Rbbp4] [Locus Tag: ] [Chromosome: 4] [Map Location: 4\|4 D2.3] | 1.961038838 | 0.000157 | 0.000476 |
| 3427 | ENSMUSG00000066037 | Hnrnpr | 74326 [Gene Symbol: Hnrnpr] [Locus Tag: ] [Chromosome: 4] [Map Location: 4\|4 D3] [D | 1.959535609 | 5.03E-05 | 0.000167 |
| 3507 | ENSMUSG00000040536 | Necab1 | 69352 [Gene Symbol: Necab1] [Locus Tag: ] [Chromosome: 4] [Map Location: 4\|4 A2] [D | 1.95838355 | 0.020725 | 0.039784 |
| 6569 | ENSMUSG00000036964 | Trim17 | 56631 [Gene Symbol: Trim17] [Locus Tag: ] [Chromosome: 11] [Map Location: 11\|11 B | 1.958055918 | 0.001244 | 0.003179 |
| 17212 | ENSMUSG00000079737 | 3110001I22Rik | 66598 [Gene Symbol: 3110001I22Rik] [Locus Tag: ] [Chromosome: 16] [Map Location: | 1.955455789 | 0.014225 | 0.028517 |
| 14874 | ENSMUSG00000038546 | Ranbp9 | 56705 [Gene Symbol: Ranbp9] [Locus Tag: ] [Chromosome: 13] [Map Location: 13\|13 A | 1.953553882 | 9.65E-05 | 0.000304 |
| 2316 | ENSMUSG00000062691 | Cebpzos | 68554 [Gene Symbol: Cebpzos] [Locus Tag: ] [Chromosome: 17] [Map Location: 17\|17 | 1.952462517 | 0.019358 | 0.037418 |
| 7851 | ENSMUSG00000003226 | Ranbp2 | 19386 [Gene Symbol: Ranbp2] [Locus Tag: ] [Chromosome: 10] [Map Location: 10 B4\|1 | 1.952349655 | 6.85E-05 | 0.000222 |
| 11301 | ENSMUSG00000047146 | Tet1 | 52463 [Gene Symbol: Tet1] [Locus Tag: ] [Chromosome: 10] [Map Location: 10 B4\|10 3 | 1.950938155 | 4.75E-05 | 0.000159 |
| 12996 | ENSMUSG00000099083 | Atf7 | 223922 [Gene Symbol: Atf7] [Locus Tag: ] [Chromosome: 15] [Map Location: 15\|15 F3] | 1.949612897 | 0.000115 | 0.000357 |
| 16839 | ENSMUSG00000001674 | Ddx18 | 66942 [Gene Symbol: Ddx18] [Locus Tag: ] [Chromosome: 1] [Map Location: 1\|1 E2] [De | 1.949480733 | 0.017213 | 0.033717 |
| 16210 | ENSMUSG00000031629 | Cenpu | 71876 [Gene Symbol: Cenpu] [Locus Tag: ] [Chromosome: 8] [Map Location: 8\|8 B2] [De | 1.949412077 | 0.019176 | 0.037121 |
| 10693 | ENSMUSG00000032507 | Fbxl2 | 72179 [Gene Symbol: Fbxl2] [Locus Tag: ] [Chromosome: 9] [Map Location: 9\|9 F3] [De | 1.948881701 | 4.60E-05 | 0.000154 |
| 15469 | ENSMUSG00000026987 | Baz2b | 407823 [Gene Symbol: Baz2b] [Locus Tag: ] [Chromosome: 2] [Map Location: 2 C1.1\|2 | 1.948294133 | 3.62E-05 | 0.000124 |
| 18441 | ENSMUSG00000037270 | 4932438A13Rik | 229227 [Gene Symbol: 4932438A13Rik] [Locus Tag: ] [Chromosome: 3] [Map Location: | 1.948231996 | 3.31E-05 | 0.000115 |
| 10670 | ENSMUSG00000025323 | Sp4 | 20688 [Gene Symbol: Sp4] [Locus Tag: ] [Chromosome: 12] [Map Location: 12 F2\|12 63 | 1.946993216 | 0.000265 | 0.000769 |
| 8419 | ENSMUSG00000028096 | Gpr89 | 67549 [Gene Symbol: Gpr89] [Locus Tag: ] [Chromosome: 3] [Map Location: 3\|3 F2] [De | 1.946406353 | 0.000325 | 0.000931 |
| 13872 | ENSMUSG00000030310 | Slc6a1 | 232333 [Gene Symbol: Slc6a1] [Locus Tag: ] [Chromosome: 6] [Map Location: 6\|6 E3] [ | 1.946129492 | 7.75E-05 | 0.000248 |
| 15573 | ENSMUSG00000022051 | Bnip3l | 12177 [Gene Symbol: Bnip3l] [Locus Tag: ] [Chromosome: 14] [Map Location: 14 D1\|14 | 1.945611715 | 0.000642 | 0.001733 |
| 5315 | ENSMUSG00000020368 | Canx | 12330 [Gene Symbol: Canx] [Locus Tag: ] [Chromosome: 11] [Map Location: 11 B1.3\|11 | 1.945328091 | 7.54E-05 | 0.000242 |
| 12385 | ENSMUSG00000054409 | Tmem74 | 239408 [Gene Symbol: Tmem74] [Locus Tag: ] [Chromosome: 15] [Map Location: 15\|1 | 1.945327681 | 0.011642 | 0.023868 |
| 12195 | ENSMUSG00000024099 | Ndufv2 | 72900 [Gene Symbol: Ndufv2] [Locus Tag: ] [Chromosome: 17] [Map Location: 17 35.26 | 1.944768669 | 0.00022 | 0.000651 |
| 3619 | ENSMUSG00000037119 | D15Ertd621e | 210998 [Gene Symbol: D15Ertd621e] [Locus Tag: ] [Chromosome: 15] [Map Location: 1 | 1.944747545 | 8.92E-05 | 0.000282 |
| 16718 | ENSMUSG00000021775 | Nr1d2 | 353187 [Gene Symbol: Nr1d2] [Locus Tag: ] [Chromosome: 14] [Map Location: 14\|14 A | 1.944644742 | 0.000696 | 0.001867 |
| 16732 | ENSMUSG00000029632 | Ndufa4 | 17992 [Gene Symbol: Ndufa4] [Locus Tag: ] [Chromosome: 6] [Map Location: 6\|6 A2] [D | 1.944225311 | 0.019902 | 0.038346 |
| 16999 | ENSMUSG00000055078 | Gabra5 | 110886 [Gene Symbol: Gabra5] [Locus Tag: ] [Chromosome: 7] [Map Location: 7 C\|7 33 | 1.943916152 | 0.005599 | 0.012333 |
| 15685 | ENSMUSG00000068040 | Tm9sf4 | 99237 [Gene Symbol: Tm9sf4] [Locus Tag: ] [Chromosome: 2] [Map Location: 2\|2 H1] [D | 1.943775592 | 0.000228 | 0.000671 |
| 6661 | ENSMUSG00000027184 | Caprin1 | 53872 [Gene Symbol: Caprin1] [Locus Tag: ] [Chromosome: 2] [Map Location: 2\|2 E2] [ | 1.943692469 | 7.69E-05 | 0.000247 |
| 11445 | ENSMUSG00000026478 | Lamc1 | 226519 [Gene Symbol: Lamc1] [Locus Tag: ] [Chromosome: 1] [Map Location: 1 G3\|1 6 | 1.943670442 | 6.67E-05 | 0.000217 |
| 4766 | ENSMUSG00000027238 | Frmd5 | 228564 [Gene Symbol: Frmd5] [Locus Tag: ] [Chromosome: 2] [Map Location: 2\|2 E5] [ | 1.943642313 | 5.13E-05 | 0.00017 |
| 18288 | ENSMUSG00000059586 | Nsmce2 | 68501 [Gene Symbol: Nsmce2] [Locus Tag: ] [Chromosome: 15] [Map Location: 15\|15 D | 1.941934456 | 0.000427 | 0.001196 |
| 11702 | ENSMUSG00000031592 | Pcm1 | 18536 [Gene Symbol: Pcm1] [Locus Tag: ] [Chromosome: 8] [Map Location: 8 A4\|8 23.8 | 1.941924923 | 0.001821 | 0.004476 |
| 1859 | ENSMUSG00000001156 | Mxd1 | 17119 [Gene Symbol: Mxd1] [Locus Tag: ] [Chromosome: 6] [Map Location: 6 D1\|6 37. | 1.939881177 | 0.001289 | 0.003279 |
| 14837 | ENSMUSG00000034401 | Spata6 | 67946 [Gene Symbol: Spata6] [Locus Tag: ] [Chromosome: 4] [Map Location: 4\|4 D1] [D | 1.939631686 | 0.004061 | 0.009232 |
| 7929 | ENSMUSG00000021611 | Tert | 21752 [Gene Symbol: Tert] [Locus Tag: ] [Chromosome: 13] [Map Location: 13 C1\|13 4 | 1.939159845 | 0.012632 | 0.025631 |
| 5748 | ENSMUSG00000051329 | Nup160 | 59015 [Gene Symbol: Nup160] [Locus Tag: ] [Chromosome: 2] [Map Location: 2\|2 E1] [ | 1.938510491 | 0.000405 | 0.001139 |
| 8761 | ENSMUSG00000048000 | Gigyf2 | 227331 [Gene Symbol: Gigyf2] [Locus Tag: ] [Chromosome: 1] [Map Location: 1\|1 D] [D | 1.937999718 | 0.000285 | 0.000824 |
| 16848 | ENSMUSG00000024014 | Pim1 | 18712 [Gene Symbol: Pim1] [Locus Tag: ] [Chromosome: 17] [Map Location: 17 A3.3\|1 | 1.937973713 | 0.005208 | 0.011558 |
| 9948 | ENSMUSG00000030465 | Psd3 | 234353 [Gene Symbol: Psd3] [Locus Tag: ] [Chromosome: 8] [Map Location: 8\|8 B3.3] [ | 1.937034007 | 0.001722 | 0.004258 |
| 6125 | ENSMUSG00000025880 | Smad7 | 17131 [Gene Symbol: Smad7] [Locus Tag: ] [Chromosome: 18] [Map Location: 18\|18 E2 | 1.936542645 | 4.90E-05 | 0.000163 |
| 309 | ENSMUSG00000031755 | Bbs2 | 67378 [Gene Symbol: Bbs2] [Locus Tag: ] [Chromosome: 8] [Map Location: 8\|8 C5] [Des | 1.93647862 | 0.003615 | 0.008316 |
| 10007 | ENSMUSG00000046573 | Lyrm4 | 380840 [Gene Symbol: Lyrm4] [Locus Tag: ] [Chromosome: 13] [Map Location: 13\|13 A | 1.936377234 | 0.001436 | 0.003619 |
| 16965 | ENSMUSG00000051650 | B3gnt2 | 53625 [Gene Symbol: B3gnt2] [Locus Tag: ] [Chromosome: 11] [Map Location: 11 A3.2\| | 1.935862845 | 0.007703 | 0.016459 |
| 86 | ENSMUSG00000036768 | Kif15 | 209737 [Gene Symbol: Kif15] [Locus Tag: ] [Chromosome: 9] [Map Location: 9\|9 F4] [De | 1.935342427 | 0.011905 | 0.024346 |
| 11962 | ENSMUSG00000037890 | Wdr19 | 213081 [Gene Symbol: Wdr19] [Locus Tag: ] [Chromosome: 5] [Map Location: 5\|5 C3.1 | 1.935168677 | 0.000445 | 0.00124 |
| 6906 | ENSMUSG00000037940 | Inpp4b | 234515 [Gene Symbol: Inpp4b] [Locus Tag: ] [Chromosome: 8] [Map Location: 8 C2\|8 3 | 1.935036668 | 0.016053 | 0.03173 |
| 2760 | ENSMUSG00000026495 | Efcab2 | 68226 [Gene Symbol: Efcab2] [Locus Tag: ] [Chromosome: 1] [Map Location: 1\|1 H3] [D | 1.935031162 | 0.008787 | 0.018538 |
| 5975 | ENSMUSG00000045973 | Slc25a51 | 230125 [Gene Symbol: Slc25a51] [Locus Tag: ] [Chromosome: 4] [Map Location: 4\|4 B1 | 1.934975128 | 5.00E-05 | 0.000167 |
| 507 | ENSMUSG00000038005 | 2700029M09Rik | 72612 [Gene Symbol: 2700029M09Rik] [Locus Tag: ] [Chromosome: 8] [Map Location: | 1.933897363 | 0.001169 | 0.003005 |

| | | | | | | |
|---|---|---|---|---|---|---|
| 2683 | ENSMUSG00000053470 | Kdm3a | 104263 [Gene Symbol: Kdm3a] [Locus Tag: ] [Chromosome: 6] [Map Location: 6\|6 C1] [ | 1.933379556 | 0.00072 | 0.001924 |
| 15513 | ENSMUSG00000031590 | Frg1 | 14300 [Gene Symbol: Frg1] [Locus Tag: ] [Chromosome: 8] [Map Location: 8 A4\|8 23.8 | 1.933028668 | 0.002055 | 0.00499 |
| 10054 | ENSMUSG00000070880 | Gad1 | 14415 [Gene Symbol: Gad1] [Locus Tag: ] [Chromosome: 2] [Map Location: 2 41.63 cM\| | 1.931743251 | 8.02E-05 | 0.000256 |
| 11909 | ENSMUSG00000036712 | Cyld | 74256 [Gene Symbol: Cyld] [Locus Tag: ] [Chromosome: 8] [Map Location: 8\|8 C3] [Desc | 1.931687151 | 0.000107 | 0.000333 |
| 13922 | ENSMUSG00000016520 | Lnx2 | 140887 [Gene Symbol: Lnx2] [Locus Tag: ] [Chromosome: 5] [Map Location: 5\|5 G2] [De | 1.931553417 | 0.00069 | 0.001852 |
| 14831 | ENSMUSG00000041488 | Stx3 | 20908 [Gene Symbol: Stx3] [Locus Tag: ] [Chromosome: 19] [Map Location: 19\|19 A] [D | 1.931254539 | 0.00182 | 0.004474 |
| 9396 | ENSMUSG00000027508 | Pag1 | 94212 [Gene Symbol: Pag1] [Locus Tag: ] [Chromosome: 3] [Map Location: 3\|3 A1] [Des | 1.931204745 | 0.000534 | 0.001465 |
| 15293 | ENSMUSG00000036111 | Lmo1 | 109594 [Gene Symbol: Lmo1] [Locus Tag: ] [Chromosome: 7] [Map Location: 7 E3\|7 57. | 1.930170301 | 0.000811 | 0.002147 |
| 1921 | ENSMUSG00000037287 | Tbcel | 272589 [Gene Symbol: Tbcel] [Locus Tag: ] [Chromosome: 9] [Map Location: 9\|9 A5.1] [ | 1.92928276 | 0.000138 | 0.000422 |
| 4305 | ENSMUSG00000040152 | Thbs1 | 21825 [Gene Symbol: Thbs1] [Locus Tag: ] [Chromosome: 2] [Map Location: 2 59.34 cM | 1.926700004 | 0.003035 | 0.007109 |
| 7071 | ENSMUSG00000053293 | Pom121 | 107939 [Gene Symbol: Pom121] [Locus Tag: ] [Chromosome: 5] [Map Location: 5\|5 G2 | 1.925623034 | 0.002327 | 0.005585 |
| 6747 | ENSMUSG00000018209 | Stk4 | 58231 [Gene Symbol: Stk4] [Locus Tag: ] [Chromosome: 2] [Map Location: 2\|2 H3] [Des | 1.924333388 | 0.000329 | 0.00094 |
| 3717 | ENSMUSG00000052632 | Asap2 | 211914 [Gene Symbol: Asap2] [Locus Tag: ] [Chromosome: 12] [Map Location: 12\|12 A | 1.924174501 | 5.61E-05 | 0.000184 |
| 3335 | ENSMUSG00000035578 | Iqcg | 69707 [Gene Symbol: Iqcg] [Locus Tag: ] [Chromosome: 16] [Map Location: 16\|16 B3] [ | 1.923372543 | 0.009416 | 0.019728 |
| 4617 | ENSMUSG00000057706 | Mex3b | 108797 [Gene Symbol: Mex3b] [Locus Tag: ] [Chromosome: 7] [Map Location: 7 D3\|7 4 | 1.923335787 | 0.000244 | 0.000714 |
| 15968 | ENSMUSG00000029754 | Dlx6 | 13396 [Gene Symbol: Dlx6] [Locus Tag: ] [Chromosome: 6] [Map Location: 6 A1\|6 2.83 | 1.922885224 | 0.002061 | 0.005004 |
| 8055 | ENSMUSG00000043668 | Tox3 | 244579 [Gene Symbol: Tox3] [Locus Tag: ] [Chromosome: 8] [Map Location: 8 C4\|8 43. | 1.922689031 | 0.000169 | 0.00051 |
| 8795 | ENSMUSG00000014592 | Camta1 | 100072 [Gene Symbol: Camta1] [Locus Tag: ] [Chromosome: 4] [Map Location: 4\|4 E2] | 1.922206786 | 0.001565 | 0.003906 |
| 191 | ENSMUSG00000030235 | Slco1c1 | 58807 [Gene Symbol: Slco1c1] [Locus Tag: ] [Chromosome: 6] [Map Location: 6\|6 G1] [ | 1.92191092 | 0.004947 | 0.011037 |
| 5783 | ENSMUSG00000078435 | AU041133 | 216177 [Gene Symbol: AU041133] [Locus Tag: ] [Chromosome: 10] [Map Location: 10\| | 1.921137676 | 0.018656 | 0.036248 |
| 550 | ENSMUSG00000031864 | Ints10 | 70885 [Gene Symbol: Ints10] [Locus Tag: ] [Chromosome: 8] [Map Location: 8\|8 C1] [De | 1.92085086 | 0.000283 | 0.000817 |
| 15464 | ENSMUSG00000020022 | Ndufa12 | 66414 [Gene Symbol: Ndufa12] [Locus Tag: ] [Chromosome: 10] [Map Location: 10\|10 | 1.920655704 | 0.009719 | 0.020299 |
| 14074 | ENSMUSG00000025078 | Nhlrc2 | 66866 [Gene Symbol: Nhlrc2] [Locus Tag: ] [Chromosome: 19] [Map Location: 19\|19 D | 1.920060649 | 0.002044 | 0.004966 |
| 13647 | ENSMUSG00000034893 | Cog3 | 338337 [Gene Symbol: Cog3] [Locus Tag: ] [Chromosome: 14] [Map Location: 14\|14 D3 | 1.919200345 | 6.65E-05 | 0.000216 |
| 3464 | ENSMUSG00000032475 | Nck1 | 17973 [Gene Symbol: Nck1] [Locus Tag: ] [Chromosome: 9] [Map Location: 9\|9 E3.3] [D | 1.919141865 | 0.000791 | 0.002097 |
| 12129 | ENSMUSG00000050069 | Grem2 | 23893 [Gene Symbol: Grem2] [Locus Tag: ] [Chromosome: 1] [Map Location: 1\|1 H3] [D | 1.918596369 | 0.010059 | 0.020946 |
| 16391 | ENSMUSG00000029125 | Stx18 | 71116 [Gene Symbol: Stx18] [Locus Tag: ] [Chromosome: 5] [Map Location: 5\|5 B3] [De | 1.918573033 | 0.000313 | 0.000897 |
| 11469 | ENSMUSG00000022419 | Deptor | 97998 [Gene Symbol: Deptor] [Locus Tag: ] [Chromosome: 15] [Map Location: 15 D1\|1 | 1.918499809 | 0.008344 | 0.017687 |
| 17756 | ENSMUSG00000022206 | Npr3 | 18162 [Gene Symbol: Npr3] [Locus Tag: ] [Chromosome: 15] [Map Location: 15 A1\|15 | 1.918349357 | 0.000259 | 0.000754 |
| 1909 | ENSMUSG00000045932 | Ifit2 | 15958 [Gene Symbol: Ifit2] [Locus Tag: ] [Chromosome: 19] [Map Location: 19\|19 C1] [ | 1.917599791 | 0.001426 | 0.003598 |
| 12380 | ENSMUSG00000043885 | Slc36a4 | 234967 [Gene Symbol: Slc36a4] [Locus Tag: ] [Chromosome: 9] [Map Location: 9\|9 A2] | 1.917258168 | 0.000122 | 0.000378 |
| 233 | ENSMUSG00000055900 | Tmem69 | 230657 [Gene Symbol: Tmem69] [Locus Tag: ] [Chromosome: 4] [Map Location: 4\|4 D1 | 1.915726125 | 0.007424 | 0.015923 |
| 5504 | ENSMUSG00000031256 | Cstf2 | 108062 [Gene Symbol: Cstf2] [Locus Tag: ] [Chromosome: X] [Map Location: X\|X E3] [De | 1.915349534 | 0.000146 | 0.000446 |
| 2178 | ENSMUSG00000036815 | Dpp10 | 269109 [Gene Symbol: Dpp10] [Locus Tag: ] [Chromosome: 1] [Map Location: 1\|1 E2.3 | 1.915278938 | 0.006314 | 0.013738 |
| 15192 | ENSMUSG00000021569 | Trip13 | 69716 [Gene Symbol: Trip13] [Locus Tag: ] [Chromosome: 13] [Map Location: 13 C1\|13 | 1.915177954 | 0.015242 | 0.030301 |
| 2156 | ENSMUSG00000049538 | Adamts16 | 271127 [Gene Symbol: Adamts16] [Locus Tag: ] [Chromosome: 13] [Map Location: 13 C | 1.91410568 | 0.013185 | 0.026622 |
| 645 | ENSMUSG00000025613 | Cct8 | 12469 [Gene Symbol: Cct8] [Locus Tag: ] [Chromosome: 16] [Map Location: 16\|16 C3.3 | 1.914077596 | 0.000129 | 0.000397 |
| 13449 | ENSMUSG00000032316 | Clk3 | 102414 [Gene Symbol: Clk3] [Locus Tag: ] [Chromosome: 9] [Map Location: 9\|9 C] [Desc | 1.913988848 | 0.001015 | 0.002636 |
| 15198 | ENSMUSG00000024935 | Slc1a1 | 20510 [Gene Symbol: Slc1a1] [Locus Tag: ] [Chromosome: 19] [Map Location: 19\|19 C1 | 1.9133189 | 0.000543 | 0.001487 |
| 12918 | ENSMUSG00000030619 | Eed | 13626 [Gene Symbol: Eed] [Locus Tag: ] [Chromosome: 7] [Map Location: 7 E1\|7] [Descr | 1.913136572 | 0.004082 | 0.009272 |
| 6359 | ENSMUSG00000037795 | N4bp2 | 333789 [Gene Symbol: N4bp2] [Locus Tag: ] [Chromosome: 5] [Map Location: 5\|5 C3.1 | 1.912298277 | 0.00013 | 0.000401 |
| 15067 | ENSMUSG00000034118 | Tpst1 | 22021 [Gene Symbol: Tpst1] [Locus Tag: ] [Chromosome: 5] [Map Location: 5\|5 F-G1] [D | 1.910708349 | 0.000247 | 0.000723 |
| 9076 | ENSMUSG00000014503 | Pkd2l2 | 53871 [Gene Symbol: Pkd2l2] [Locus Tag: ] [Chromosome: 18] [Map Location: 18\|18 C] | 1.910284961 | 7.25E-05 | 0.000234 |
| 17572 | ENSMUSG00000037211 | Spry1 | 24063 [Gene Symbol: Spry1] [Locus Tag: ] [Chromosome: 3] [Map Location: 3\|3 B] [Desc | 1.90971622 | 0.000147 | 0.000448 |
| 14764 | ENSMUSG00000027236 | Eif3j1 | 78655 [Gene Symbol: Eif3j1] [Locus Tag: ] [Chromosome: 2] [Map Location: 2\|2 E5] [De | 1.909349543 | 0.003435 | 0.007935 |
| 9105 | ENSMUSG00000056429 | Tgoln2; Tgoln1 | 22135 [Gene Symbol: Tgoln2] [Locus Tag: ] [Chromosome: ] [Map Location: ] [Descriptic | 1.909110314 | 0.000487 | 0.001346 |
| 14681 | ENSMUSG00000005907 | Pex1 | 71382 [Gene Symbol: Pex1] [Locus Tag: ] [Chromosome: 5] [Map Location: 5\|5 A2] [Des | 1.907269527 | 0.000222 | 0.000654 |
| 16941 | ENSMUSG00000039485 | Tspyl4 | 72480 [Gene Symbol: Tspyl4] [Locus Tag: ] [Chromosome: 10] [Map Location: 10 B1\|10 | 1.906560485 | 0.001222 | 0.003128 |
| 16160 | ENSMUSG00000025962 | Fastkd2 | 75619 [Gene Symbol: Fastkd2] [Locus Tag: ] [Chromosome: 1] [Map Location: 1\|1 C2] [D | 1.906240481 | 0.007284 | 0.01565 |
| 18158 | ENSMUSG00000063488 | Zkscan7 | 382118 [Gene Symbol: Zkscan7] [Locus Tag: ] [Chromosome: 9] [Map Location: 9\|9 F4] | 1.905504186 | 0.010438 | 0.021642 |
| 5289 | ENSMUSG00000035021 | Baz1a | 217578 [Gene Symbol: Baz1a] [Locus Tag: ] [Chromosome: 12] [Map Location: 12\|12 C] | 1.905285938 | 0.007362 | 0.015804 |
| 8035 | ENSMUSG00000070354 | Gm21975 | 101488212 [Gene Symbol: Gm21975] [Locus Tag: ] [Chromosome: 11] [Map Location: | 1.90409559 | 0.025054 | 0.047145 |
| 7520 | ENSMUSG00000036402 | Gng12 | 14701 [Gene Symbol: Gng12] [Locus Tag: ] [Chromosome: 6] [Map Location: 6 C1\|6 30. | 1.903555101 | 0.002377 | 0.005697 |
| 16966 | ENSMUSG00000028995 | Fam126a | 84652 [Gene Symbol: Fam126a] [Locus Tag: ] [Chromosome: 5] [Map Location: 5\|5 A3] | 1.902556104 | 0.000159 | 0.000483 |
| 12637 | ENSMUSG00000038122 | Tbc1d32 | 544696 [Gene Symbol: Tbc1d32] [Locus Tag: ] [Chromosome: 10] [Map Location: 10\|1 | 1.902073848 | 0.000507 | 0.001397 |
| 11297 | ENSMUSG00000039480 | Nt5dc1 | 319638 [Gene Symbol: Nt5dc1] [Locus Tag: ] [Chromosome: 10] [Map Location: 10\|10 | 1.901756051 | 0.001205 | 0.003091 |
| 4047 | ENSMUSG00000032261 | Sh3bgrl2 | 212531 [Gene Symbol: Sh3bgrl2] [Locus Tag: ] [Chromosome: 9] [Map Location: 9\|9 E2 | 1.90135255 | 0.006319 | 0.013746 |
| 12985 | ENSMUSG00000029681 | Bcl7b | 12054 [Gene Symbol: Bcl7b] [Locus Tag: ] [Chromosome: 5] [Map Location: 5\|5 G2] [De | 1.899870411 | 0.000598 | 0.001626 |
| 261 | ENSMUSG00000040269 | Mrps28 | 66230 [Gene Symbol: Mrps28] [Locus Tag: ] [Chromosome: 3] [Map Location: 3\|3 A1] [D | 1.899789243 | 0.00453 | 0.010187 |
| 2688 | ENSMUSG00000044807 | Zfp354c | 30944 [Gene Symbol: Zfp354c] [Locus Tag: ] [Chromosome: 11] [Map Location: 11 B1.3 | 1.897081896 | 5.38E-05 | 0.000177 |
| 55 | ENSMUSG00000006567 | Atp7b | 11979 [Gene Symbol: Atp7b] [Locus Tag: ] [Chromosome: 8] [Map Location: 8 A2\|8 10. | 1.895261166 | 0.003194 | 0.007442 |
| 7411 | ENSMUSG00000000804 | Usp32 | 237898 [Gene Symbol: Usp32] [Locus Tag: ] [Chromosome: 11] [Map Location: 11\|11 C | 1.895082851 | 0.000426 | 0.001192 |
| 11043 | ENSMUSG00000037514 | Pank2 | 74450 [Gene Symbol: Pank2] [Locus Tag: ] [Chromosome: 2] [Map Location: 2\|2 F1] [De | 1.894821031 | 0.000222 | 0.000656 |
| 18574 | ENSMUSG00000062981 | Mrpl42 | 67270 [Gene Symbol: Mrpl42] [Locus Tag: ] [Chromosome: 10] [Map Location: 10 C2\|1 | 1.894482092 | 0.003421 | 0.007908 |
| 16771 | ENSMUSG00000078941 | Ak6; Taf9 | 102216272 [Gene Symbol: Ak6] [Locus Tag: ] [Chromosome: 13] [Map Location: 13 D1] | 1.893796778 | 0.000569 | 0.001552 |
| 9123 | ENSMUSG00000042496 | Prdm10 | 382066 [Gene Symbol: Prdm10] [Locus Tag: ] [Chromosome: 9] [Map Location: 9\|9 A4] | 1.893475386 | 0.000345 | 0.000983 |
| 1886 | ENSMUSG00000038375 | Trp53inp2 | 68728 [Gene Symbol: Trp53inp2] [Locus Tag: ] [Chromosome: 2] [Map Location: 2\|2 H1 | 1.892888727 | 0.000105 | 0.000328 |
| 16233 | ENSMUSG00000035476 | Tab3 | 66724 [Gene Symbol: Tab3] [Locus Tag: ] [Chromosome: X] [Map Location: X\|X C1] [Des | 1.892412363 | 0.000102 | 0.00032 |
| 18522 | ENSMUSG00000054720 | Lrrc8c | 100604 [Gene Symbol: Lrrc8c] [Locus Tag: ] [Chromosome: 5] [Map Location: 5\|5 E5] [D | 1.892327641 | 0.000199 | 0.000593 |
| 15321 | ENSMUSG00000029474 | Rnf34 | 80751 [Gene Symbol: Rnf34] [Locus Tag: ] [Chromosome: 5] [Map Location: 5\|5 F] [Desc | 1.892300948 | 0.000352 | 0.001001 |
| 562 | ENSMUSG00000051166 | Eml5 | 319670 [Gene Symbol: Eml5] [Locus Tag: ] [Chromosome: 12] [Map Location: 12\|12 E] | 1.892266879 | 0.000894 | 0.00235 |
| 9460 | ENSMUSG00000022253 | Nadk2 | 68646 [Gene Symbol: Nadk2] [Locus Tag: ] [Chromosome: 15] [Map Location: 15\|15 A2 | 1.892216531 | 0.000356 | 0.00101 |
| 14258 | ENSMUSG00000041477 | Dcp1b | 319618 [Gene Symbol: Dcp1b] [Locus Tag: ] [Chromosome: 6] [Map Location: 6\|6 F1] [D | 1.892080223 | 0.000567 | 0.001549 |
| 12481 | ENSMUSG00000026767 | Fam188a | 66960 [Gene Symbol: Fam188a] [Locus Tag: ] [Chromosome: 2] [Map Location: 2\|2 A1] | 1.891754103 | 0.001744 | 0.004309 |
| 4252 | ENSMUSG00000029212 | Gabrb1 | 14400 [Gene Symbol: Gabrb1] [Locus Tag: ] [Chromosome: 5] [Map Location: 5 C3.2\|5 | 1.8914811 | 0.007296 | 0.01567 |
| 12030 | ENSMUSG00000051695 | Pcbp1 | 23983 [Gene Symbol: Pcbp1] [Locus Tag: ] [Chromosome: 6] [Map Location: 6\|6 D1] [D | 1.890394347 | 0.000204 | 0.000608 |
| 11192 | ENSMUSG00000041020 | Map7d2 | 78283 [Gene Symbol: Map7d2] [Locus Tag: ] [Chromosome: X] [Map Location: X\|X F4] [ | 1.888380901 | 0.011204 | 0.023072 |
| 2018 | ENSMUSG00000035266 | Helq | 191578 [Gene Symbol: Helq] [Locus Tag: ] [Chromosome: 5] [Map Location: 5\|5 E] [Desc | 1.888066622 | 0.000561 | 0.001534 |
| 14085 | ENSMUSG00000022160 | Mettl3 | 56335 [Gene Symbol: Mettl3] [Locus Tag: ] [Chromosome: 14] [Map Location: 14\|14 C | 1.888051949 | 0.002465 | 0.005892 |
| 4309 | ENSMUSG00000025932 | Eya1 | 14048 [Gene Symbol: Eya1] [Locus Tag: ] [Chromosome: 1] [Map Location: 1 A3\|1 4.31 | 1.887482352 | 0.000705 | 0.001889 |
| 12675 | ENSMUSG00000000794 | Kcnn3 | 140493 [Gene Symbol: Kcnn3] [Locus Tag: ] [Chromosome: 3] [Map Location: 3\|3 F2] [D | 1.886817962 | 0.001621 | 0.004036 |
| 17991 | ENSMUSG00000032558 | Nphp3 | 74025 [Gene Symbol: Nphp3] [Locus Tag: ] [Chromosome: 9] [Map Location: 9 F1\|9 56. | 1.886205042 | 0.000186 | 0.000557 |
| 5243 | ENSMUSG00000040234 | Tm7sf3 | 67623 [Gene Symbol: Tm7sf3] [Locus Tag: ] [Chromosome: 6] [Map Location: 6\|6 G3] [D | 1.886058026 | 0.00062 | 0.001679 |
| 5873 | ENSMUSG00000000346 | Dazap2 | 23994 [Gene Symbol: Dazap2] [Locus Tag: ] [Chromosome: 15] [Map Location: 15 F1\|1 | 1.885380847 | 0.001587 | 0.003956 |

| | | | | | | |
|---|---|---|---|---|---|---|
| 16080 | ENSMUSG00000018654 | Ikzf1 | 22778 [Gene Symbol: Ikzf1] [Locus Tag: ] [Chromosome: 11] [Map Location: 11 A1\|11 7 | 1.88533668 | 0.007113 | 0.015304 |
| 3530 | ENSMUSG00000021102 | Glrx5 | 73046 [Gene Symbol: Glrx5] [Locus Tag: ] [Chromosome: 12] [Map Location: 12\|12 E] [ | 1.884000881 | 0.001311 | 0.003332 |
| 16435 | ENSMUSG00000025262 | Fam120c | 207375 [Gene Symbol: Fam120c] [Locus Tag: ] [Chromosome: X] [Map Location: X\|X F2 | 1.882832783 | 0.000164 | 0.000495 |
| 15838 | ENSMUSG00000042688 | Mapk6 | 50772 [Gene Symbol: Mapk6] [Locus Tag: ] [Chromosome: 9] [Map Location: 9 D\|9 42.3 | 1.882269311 | 6.87E-05 | 0.000223 |
| 7519 | ENSMUSG00000044499 | Hs3st5 | 319415 [Gene Symbol: Hs3st5] [Locus Tag: ] [Chromosome: 10] [Map Location: 10\|10 E | 1.882148023 | 0.001921 | 0.004696 |
| 14830 | ENSMUSG00000038876 | Rnf146 | 68031 [Gene Symbol: Rnf146] [Locus Tag: ] [Chromosome: 10] [Map Location: 10\|10 A | 1.881997369 | 0.000158 | 0.000478 |
| 3459 | ENSMUSG00000054051 | Ercc6 | 319955 [Gene Symbol: Ercc6] [Locus Tag: ] [Chromosome: 14] [Map Location: 14\|14 B] | 1.880961699 | 0.00036 | 0.001021 |
| 16395 | ENSMUSG00000037316 | Bag4 | 67384 [Gene Symbol: Bag4] [Locus Tag: ] [Chromosome: 8] [Map Location: 8\|8 A3] [Des | 1.880645667 | 0.000237 | 0.000694 |
| 3817 | ENSMUSG00000054226 | Tprkb | 69786 [Gene Symbol: Tprkb] [Locus Tag: ] [Chromosome: 6] [Map Location: 6\|6 C3] [De | 1.880225946 | 0.001392 | 0.00352 |
| 6600 | ENSMUSG00000045751 | Mms22l | 212377 [Gene Symbol: Mms22l] [Locus Tag: ] [Chromosome: 4] [Map Location: 4\|4 A3] | 1.879770789 | 0.01373 | 0.027611 |
| 1500 | ENSMUSG00000029192 | LOC102641289; Tbc1 | 102641289 [Gene Symbol: LOC102641289] [Locus Tag: ] [Chromosome: 5] [Map Locati | 1.879670136 | 8.60E-05 | 0.000273 |
| 4648 | ENSMUSG00000029772 | Ahcyl2 | 74340 [Gene Symbol: Ahcyl2] [Locus Tag: ] [Chromosome: 6] [Map Location: 6\|6 A3.3] | 1.879531103 | 0.000992 | 0.002579 |
| 1835 | ENSMUSG00000019087 | Atp6ap1 | 54411 [Gene Symbol: Atp6ap1] [Locus Tag: ] [Chromosome: X] [Map Location: X A7.3\|X | 1.879322728 | 0.009177 | 0.019271 |
| 16882 | ENSMUSG00000019874 | Fabp7 | 12140 [Gene Symbol: Fabp7] [Locus Tag: ] [Chromosome: 10] [Map Location: 10\|10 B4 | 1.879305347 | 0.000335 | 0.000958 |
| 11921 | ENSMUSG00000061436 | Hipk2 | 15258 [Gene Symbol: Hipk2] [Locus Tag: ] [Chromosome: 6] [Map Location: 6\|6 B] [Des | 1.879051454 | 0.000788 | 0.002089 |
| 5353 | ENSMUSG00000022747 | St3gal6 | 54613 [Gene Symbol: St3gal6] [Locus Tag: ] [Chromosome: 16] [Map Location: 16\|16 C | 1.879025048 | 0.001429 | 0.003603 |
| 5331 | ENSMUSG00000034800 | Zfp661 | 72180 [Gene Symbol: Zfp661] [Locus Tag: ] [Chromosome: 2] [Map Location: 2\|2 F1] [D | 1.877369371 | 0.002647 | 0.006282 |
| 17524 | ENSMUSG00000057182 | Scn3a | 20269 [Gene Symbol: Scn3a] [Locus Tag: ] [Chromosome: 2] [Map Location: 2 C1.3\|2 38 | 1.877258954 | 9.26E-05 | 0.000292 |
| 6018 | ENSMUSG00000024647 | Cbln2 | 12405 [Gene Symbol: Cbln2] [Locus Tag: ] [Chromosome: 18] [Map Location: 18 E4\|18 | 1.876956856 | 0.000628 | 0.001698 |
| 2540 | ENSMUSG00000033364 | Usp37 | 319651 [Gene Symbol: Usp37] [Locus Tag: ] [Chromosome: 1] [Map Location: 1\|1 C3] [ | 1.876702803 | 0.000426 | 0.001192 |
| 10820 | ENSMUSG00000041126 | H2afv | 77605 [Gene Symbol: H2afv] [Locus Tag: ] [Chromosome: 11] [Map Location: 11\|11 A1] | 1.87668209 | 0.003136 | 0.007321 |
| 3566 | ENSMUSG00000026667 | Uhmk1 | 16589 [Gene Symbol: Uhmk1] [Locus Tag: ] [Chromosome: 1] [Map Location: 1\|1 H2] [D | 1.876645448 | 9.70E-05 | 0.000305 |
| 4958 | ENSMUSG00000031963 | Bmper | 73230 [Gene Symbol: Bmper] [Locus Tag: ] [Chromosome: 9] [Map Location: 9\|9 A4] [D | 1.875399012 | 0.002648 | 0.006282 |
| 18519 | ENSMUSG00000035805 | Mlc1 | 170790 [Gene Symbol: Mlc1] [Locus Tag: ] [Chromosome: 15] [Map Location: 15\|15 E3 | 1.874732252 | 0.002945 | 0.006916 |
| 8970 | ENSMUSG00000042508 | Dmtf1 | 23857 [Gene Symbol: Dmtf1] [Locus Tag: ] [Chromosome: 5] [Map Location: 5\|5 A2-A3] | 1.874695946 | 0.000478 | 0.001323 |
| 966 | ENSMUSG00000030045 | Mrpl19 | 56284 [Gene Symbol: Mrpl19] [Locus Tag: ] [Chromosome: 6] [Map Location: 6 C3\|6 35 | 1.872147783 | 0.003133 | 0.007314 |
| 16615 | ENSMUSG00000020564 | Atxn7l1 | 380753 [Gene Symbol: Atxn7l1] [Locus Tag: ] [Chromosome: 12] [Map Location: 12\|12 | 1.872075494 | 0.000535 | 0.001469 |
| 1532 | ENSMUSG00000021811 | Dnajc9 | 108671 [Gene Symbol: Dnajc9] [Locus Tag: ] [Chromosome: 14] [Map Location: 14\|14 | 1.871213459 | 0.001267 | 0.003228 |
| 6901 | ENSMUSG00000063052 | Lrrc40 | 67144 [Gene Symbol: Lrrc40] [Locus Tag: ] [Chromosome: 3] [Map Location: 3\|3 H4] [D | 1.871133757 | 0.0002 | 0.000596 |
| 6594 | ENSMUSG00000030751 | Psma1 | 26440 [Gene Symbol: Psma1] [Locus Tag: ] [Chromosome: 7] [Map Location: 7 F1\|7 59. | 1.868904454 | 0.001636 | 0.004067 |
| 9558 | ENSMUSG00000027175 | Tcp11l1 | 320554 [Gene Symbol: Tcp11l1] [Locus Tag: ] [Chromosome: 2] [Map Location: 2\|2 E2] | 1.867137698 | 9.70E-05 | 0.000305 |
| 3693 | ENSMUSG00000056234 | Ncoa4 | 27057 [Gene Symbol: Ncoa4] [Locus Tag: ] [Chromosome: 14] [Map Location: 14\|14 B] | 1.866873968 | 0.015733 | 0.031185 |
| 9574 | ENSMUSG00000002416 | Ndufb2 | 68198 [Gene Symbol: Ndufb2] [Locus Tag: ] [Chromosome: 6] [Map Location: 6\|6 B1] [D | 1.864482544 | 0.006733 | 0.014547 |
| 5434 | ENSMUSG00000020029 | Nudt4 | 71207 [Gene Symbol: Nudt4] [Locus Tag: ] [Chromosome: 10] [Map Location: 10\|10] [D | 1.863794094 | 8.68E-05 | 0.000275 |
| 8751 | ENSMUSG00000041324 | Inhba | 16323 [Gene Symbol: Inhba] [Locus Tag: ] [Chromosome: 13] [Map Location: 13 A1\|13 | 1.863717097 | 0.00207 | 0.005022 |
| 1389 | ENSMUSG00000052798 | Nup107 | 103468 [Gene Symbol: Nup107] [Locus Tag: ] [Chromosome: 10] [Map Location: 10\|10 | 1.86321877 | 0.002032 | 0.004941 |
| 5611 | ENSMUSG00000002997 | Prkar2b | 19088 [Gene Symbol: Prkar2b] [Locus Tag: ] [Chromosome: 12] [Map Location: 12\|12 E | 1.86305047 | 0.000138 | 0.000422 |
| 780 | ENSMUSG00000020790 | Ankfy1 | 11736 [Gene Symbol: Ankfy1] [Locus Tag: ] [Chromosome: 11] [Map Location: 11\|11 B4 | 1.862996998 | 0.0002 | 0.000597 |
| 157 | ENSMUSG00000029202 | Pds5a | 71521 [Gene Symbol: Pds5a] [Locus Tag: ] [Chromosome: 5] [Map Location: 5\|5 C3.3] [ | 1.861902127 | 0.000139 | 0.000427 |
| 13974 | ENSMUSG00000039693 | Msantd3 | 66665 [Gene Symbol: Msantd3] [Locus Tag: ] [Chromosome: 4] [Map Location: 4\|4 B2] | 1.861628897 | 0.000353 | 0.001005 |
| 4328 | ENSMUSG00000025927 | Tfap2b | 21419 [Gene Symbol: Tfap2b] [Locus Tag: ] [Chromosome: 1] [Map Location: 1\|1 A2-A4 | 1.86117393 | 0.001046 | 0.002712 |
| 411 | ENSMUSG00000022204 | Ngdn | 68966 [Gene Symbol: Ngdn] [Locus Tag: ] [Chromosome: 14] [Map Location: 14\|14 C1] | 1.859930217 | 0.020015 | 0.038534 |
| 2524 | ENSMUSG00000031758 | Cdyl2 | 75796 [Gene Symbol: Cdyl2] [Locus Tag: ] [Chromosome: 8] [Map Location: 8\|8 E1] [De | 1.859208584 | 0.000941 | 0.00246 |
| 2458 | ENSMUSG00000004069 | Dnaja3 | 83945 [Gene Symbol: Dnaja3] [Locus Tag: ] [Chromosome: 16] [Map Location: 16 A1\|1 | 1.859084277 | 0.000228 | 0.000671 |
| 15984 | ENSMUSG00000079659 | Tmem243 | 652925 [Gene Symbol: Tmem243] [Locus Tag: ] [Chromosome: 5] [Map Location: 5\|5 A | 1.859020345 | 0.000527 | 0.001448 |
| 15691 | ENSMUSG00000030551 | Nr2f2 | 11819 [Gene Symbol: Nr2f2] [Locus Tag: ] [Chromosome: 7] [Map Location: 7 D1\|7 38. | 1.858947669 | 0.000737 | 0.001966 |
| 12057 | ENSMUSG00000045210 | Vcpip1 | 70675 [Gene Symbol: Vcpip1] [Locus Tag: ] [Chromosome: 1] [Map Location: 1\|1 A2] [D | 1.858877632 | 0.000141 | 0.00043 |
| 7789 | ENSMUSG00000024151 | Msh2 | 17685 [Gene Symbol: Msh2] [Locus Tag: ] [Chromosome: 17] [Map Location: 17 E4\|17 | 1.858375852 | 0.001602 | 0.00399 |
| 16382 | ENSMUSG00000031608 | Galnt7 | 108150 [Gene Symbol: Galnt7] [Locus Tag: ] [Chromosome: 8] [Map Location: 8\|8 B3.2 | 1.858204969 | 0.001685 | 0.004175 |
| 17333 | ENSMUSG00000021149 | Gtpbp4 | 69237 [Gene Symbol: Gtpbp4] [Locus Tag: ] [Chromosome: 13] [Map Location: 13\|13 A | 1.857680232 | 0.001454 | 0.003658 |
| 15751 | ENSMUSG00000041440 | Gk5 | 235533 [Gene Symbol: Gk5] [Locus Tag: ] [Chromosome: 9] [Map Location: 9\|9 E3.3] [D | 1.857390783 | 0.004063 | 0.009234 |
| 7495 | ENSMUSG00000039634 | Zfp189 | 230162 [Gene Symbol: Zfp189] [Locus Tag: ] [Chromosome: 4] [Map Location: 4\|4 B1] [ | 1.856642195 | 0.003742 | 0.008576 |
| 3528 | ENSMUSG00000024780 | Cdc37l1 | 67072 [Gene Symbol: Cdc37l1] [Locus Tag: ] [Chromosome: 19] [Map Location: 19\|19 | 1.856004562 | 0.000426 | 0.001192 |
| 1336 | ENSMUSG00000036099 | Vezt | 215008 [Gene Symbol: Vezt] [Locus Tag: ] [Chromosome: 10] [Map Location: 10\|10 C2] | 1.855802123 | 0.000996 | 0.002589 |
| 17678 | ENSMUSG00000021731 | Mrps30 | 59054 [Gene Symbol: Mrps30] [Locus Tag: ] [Chromosome: 13] [Map Location: 13\|13 D | 1.855381932 | 0.002283 | 0.005488 |
| 4887 | ENSMUSG00000031585 | Gtf2e2 | 68153 [Gene Symbol: Gtf2e2] [Locus Tag: ] [Chromosome: 8] [Map Location: 8\|8 A4] [D | 1.854159404 | 0.002773 | 0.00655 |
| 14929 | ENSMUSG00000070544 | Top1 | 21969 [Gene Symbol: Top1] [Locus Tag: ] [Chromosome: 2] [Map Location: 2 80.96 cM\| | 1.852681689 | 0.000259 | 0.000756 |
| 2820 | ENSMUSG00000039983 | Ccdc32 | 269336 [Gene Symbol: Ccdc32] [Locus Tag: ] [Chromosome: 2] [Map Location: 2\|2 E5] | 1.852449381 | 0.001577 | 0.003933 |
| 8241 | ENSMUSG00000063887 | Nlgn1 | 192167 [Gene Symbol: Nlgn1] [Locus Tag: ] [Chromosome: 3] [Map Location: 3\|3 A3] [D | 1.852398708 | 0.00095 | 0.002482 |
| 8872 | ENSMUSG00000019861 | Gopc | 94221 [Gene Symbol: Gopc] [Locus Tag: ] [Chromosome: 10] [Map Location: 10\|10 B3] | 1.852352959 | 0.00501 | 0.011168 |
| 13995 | ENSMUSG00000033799 | Fam208b | 105203 [Gene Symbol: Fam208b] [Locus Tag: ] [Chromosome: 13] [Map Location: 13\|1 | 1.851500859 | 0.00052 | 0.001431 |
| 7427 | ENSMUSG00000036327 | Qsox2 | 227638 [Gene Symbol: Qsox2] [Locus Tag: ] [Chromosome: 2] [Map Location: 2\|2 A3] [D | 1.851284255 | 0.002566 | 0.00611 |
| 4970 | ENSMUSG00000020267 | Hint1 | 15254 [Gene Symbol: Hint1] [Locus Tag: ] [Chromosome: 11] [Map Location: 11\|11 B1. | 1.850815386 | 0.001107 | 0.002857 |
| 8818 | ENSMUSG00000051236 | Msrb3 | 320183 [Gene Symbol: Msrb3] [Locus Tag: ] [Chromosome: 10] [Map Location: 10\|10 D | 1.850626239 | 0.011209 | 0.023081 |
| 16216 | ENSMUSG00000024563 | Smad2 | 17126 [Gene Symbol: Smad2] [Locus Tag: ] [Chromosome: 18] [Map Location: 18 E3\|18 | 1.850279412 | 0.000307 | 0.000884 |
| 11160 | ENSMUSG00000057554 | Lgals8 | 56048 [Gene Symbol: Lgals8] [Locus Tag: ] [Chromosome: 13] [Map Location: 13 A1\|13 | 1.849677515 | 0.003416 | 0.007899 |
| 13990 | ENSMUSG00000014771 | Pdcd2 | 18567 [Gene Symbol: Pdcd2] [Locus Tag: ] [Chromosome: 17] [Map Location: 17 A2\|17 | 1.849674394 | 0.00207 | 0.005022 |
| 16495 | ENSMUSG00000049791 | Fzd4 | 14366 [Gene Symbol: Fzd4] [Locus Tag: ] [Chromosome: 7] [Map Location: 7 E1\|7 49.32 | 1.848704112 | 0.006336 | 0.013776 |
| 9479 | ENSMUSG00000011256 | Adam19 | 11492 [Gene Symbol: Adam19] [Locus Tag: ] [Chromosome: 11] [Map Location: 11 27.5 | 1.848191517 | 0.000354 | 0.001005 |
| 4422 | ENSMUSG00000069255 | Dusp22 | 105352 [Gene Symbol: Dusp22] [Locus Tag: ] [Chromosome: 13] [Map Location: 13\|13 | 1.847964028 | 0.001252 | 0.003196 |
| 8521 | ENSMUSG00000020459 | Mtif2 | 76784 [Gene Symbol: Mtif2] [Locus Tag: ] [Chromosome: 11] [Map Location: 11\|11 A4] | 1.847652978 | 0.001221 | 0.003126 |
| 17553 | ENSMUSG00000033854 | Kcnk10 | 72258 [Gene Symbol: Kcnk10] [Locus Tag: ] [Chromosome: 12] [Map Location: 12\|12 E | 1.847611642 | 0.00027 | 0.000783 |
| 8513 | ENSMUSG00000029201 | Ugdh | 22235 [Gene Symbol: Ugdh] [Locus Tag: ] [Chromosome: 5] [Map Location: 5 C3.1\|5 33 | 1.84632531 | 0.001394 | 0.003525 |
| 9020 | ENSMUSG00000021615 | Xrcc4 | 108138 [Gene Symbol: Xrcc4] [Locus Tag: ] [Chromosome: 13] [Map Location: 13\|13 C | 1.846202847 | 0.004073 | 0.009256 |
| 7884 | ENSMUSG00000042213 | Zfand4 | 67492 [Gene Symbol: Zfand4] [Locus Tag: ] [Chromosome: 6] [Map Location: 6\|6 F1] [D | 1.846124546 | 0.001293 | 0.00329 |
| 439 | ENSMUSG00000022899 | Slc15a2 | 57738 [Gene Symbol: Slc15a2] [Locus Tag: ] [Chromosome: 16] [Map Location: 16\|16 B | 1.844159953 | 0.001273 | 0.003243 |
| 10483 | ENSMUSG00000037730 | Mynn | 80732 [Gene Symbol: Mynn] [Locus Tag: ] [Chromosome: 3] [Map Location: 3\|3 A3] [De | 1.842877848 | 0.000324 | 0.000927 |
| 13649 | ENSMUSG00000019889 | Ptprk | 19272 [Gene Symbol: Ptprk] [Locus Tag: ] [Chromosome: 10] [Map Location: 10 A4\|10 | 1.842616385 | 0.001633 | 0.004062 |
| 18576 | ENSMUSG00000026872 | Zeb2 | 24136 [Gene Symbol: Zeb2] [Locus Tag: ] [Chromosome: 2] [Map Location: 2\|2 C1] [Des | 1.842597213 | 0.000454 | 0.001264 |
| 396 | ENSMUSG00000026674 | Ddr2 | 18214 [Gene Symbol: Ddr2] [Locus Tag: ] [Chromosome: 1] [Map Location: 1 76.84 cM\| | 1.842578664 | 0.003536 | 0.008147 |
| 6930 | ENSMUSG00000028005 | Gucy1b3 | 54195 [Gene Symbol: Gucy1b3] [Locus Tag: ] [Chromosome: 3] [Map Location: 3\|3 E3] | 1.841768206 | 0.001056 | 0.002736 |
| 5629 | ENSMUSG00000028333 | Anp32b | 67628 [Gene Symbol: Anp32b] [Locus Tag: ] [Chromosome: 4] [Map Location: 4\|4 B2] [ | 1.84146928 | 0.000774 | 0.002058 |
| 5547 | ENSMUSG00000031729 | Ist1 | 71955 [Gene Symbol: Ist1] [Locus Tag: ] [Chromosome: 8] [Map Location: 8\|8 D3] [Desc | 1.841221601 | 0.000558 | 0.001525 |
| 8142 | ENSMUSG00000026466 | Tor1aip1 | 208263 [Gene Symbol: Tor1aip1] [Locus Tag: ] [Chromosome: 1] [Map Location: 1\|1 G3 | 1.841013547 | 0.000455 | 0.001266 |

| | | | | | | |
|---|---|---|---|---|---|---|
| 15099 | ENSMUSG00000044952 | Kctd21 | 622320 [Gene Symbol: Kctd21] [Locus Tag: ] [Chromosome: 7] [Map Location: 7\|7 E1] [ | 1.838919812 | 0.024039 | 0.045424 |
| 13965 | ENSMUSG00000073633 | Fbxo36 | 66153 [Gene Symbol: Fbxo36] [Locus Tag: ] [Chromosome: 1] [Map Location: 1 C5\|1 43 | 1.838408771 | 0.010126 | 0.02107 |
| 10922 | ENSMUSG00000033147 | Slc22a15 | 242126 [Gene Symbol: Slc22a15] [Locus Tag: ] [Chromosome: 3] [Map Location: 3\|3 F2 | 1.837747587 | 0.001229 | 0.003145 |
| 18148 | ENSMUSG00000028136 | Snx27 | 76742 [Gene Symbol: Snx27] [Locus Tag: ] [Chromosome: 3] [Map Location: 3\|3 F2] [De | 1.836452013 | 0.000273 | 0.000792 |
| 7647 | ENSMUSG00000062627 | Mysm1 | 320713 [Gene Symbol: Mysm1] [Locus Tag: ] [Chromosome: 4] [Map Location: 4\|4 C5] [ | 1.836335839 | 0.000306 | 0.000879 |
| 15444 | ENSMUSG00000002083 | Bbc3 | 170770 [Gene Symbol: Bbc3] [Locus Tag: ] [Chromosome: 7] [Map Location: 7\|7 A2] [De | 1.835009511 | 0.013234 | 0.02671 |
| 355 | ENSMUSG00000028617 | Lrrc42 | 77809 [Gene Symbol: Lrrc42] [Locus Tag: ] [Chromosome: 4] [Map Location: 4\|4 C7] [De | 1.83489398 | 0.004572 | 0.010269 |
| 8663 | ENSMUSG00000026516 | Nvl | 67459 [Gene Symbol: Nvl] [Locus Tag: ] [Chromosome: 1] [Map Location: 1\|1 H4] [Descr | 1.834282276 | 0.000507 | 0.001399 |
| 16789 | ENSMUSG00000031904 | Slc7a6 | 330836 [Gene Symbol: Slc7a6] [Locus Tag: ] [Chromosome: 8] [Map Location: 8\|8 D3] [ | 1.833910961 | 0.001207 | 0.003096 |
| 3122 | ENSMUSG00000032966 | Fkbp1a | 14225 [Gene Symbol: Fkbp1a] [Locus Tag: ] [Chromosome: 2] [Map Location: 2\|2 G3] [D | 1.833562805 | 0.000184 | 0.00055 |
| 2708 | ENSMUSG00000005718 | Tfap4 | 83383 [Gene Symbol: Tfap4] [Locus Tag: ] [Chromosome: 16] [Map Location: 16 A1\|16 | 1.832987902 | 0.006275 | 0.013662 |
| 9323 | ENSMUSG00000063446 | Plppr1 | 272031 [Gene Symbol: Plppr1] [Locus Tag: ] [Chromosome: 4] [Map Location: 4\|4 B1] [ | 1.832980299 | 0.000387 | 0.001093 |
| 1035 | ENSMUSG00000036782 | Klhl13 | 67455 [Gene Symbol: Klhl13] [Locus Tag: ] [Chromosome: X] [Map Location: X\|X A2] [De | 1.83195299 | 0.000756 | 0.002014 |
| 5697 | ENSMUSG00000033014 | Trim33 | 94093 [Gene Symbol: Trim33] [Locus Tag: ] [Chromosome: 3] [Map Location: 3\|3 F2.2] | 1.830157327 | 0.000177 | 0.000532 |
| 9355 | ENSMUSG00000027233 | Patl2 | 67578 [Gene Symbol: Patl2] [Locus Tag: ] [Chromosome: 2] [Map Location: 2\|2 F1] [Des | 1.830131509 | 0.020054 | 0.038599 |
| 9130 | ENSMUSG00000079111 | Kdelr2 | 66913 [Gene Symbol: Kdelr2] [Locus Tag: ] [Chromosome: 5] [Map Location: 5\|5 G2] [D | 1.829877569 | 0.020945 | 0.040151 |
| 2674 | ENSMUSG00000038497 | Tmco3 | 234076 [Gene Symbol: Tmco3] [Locus Tag: ] [Chromosome: 8] [Map Location: 8\|8 A1.1 | 1.829770482 | 0.002181 | 0.005267 |
| 15114 | ENSMUSG00000027167 | Elp4 | 77766 [Gene Symbol: Elp4] [Locus Tag: ] [Chromosome: 2] [Map Location: 2 E3\|2 55.33 | 1.829399715 | 0.004804 | 0.010743 |
| 2126 | ENSMUSG00000025139 | Tollip | 54473 [Gene Symbol: Tollip] [Locus Tag: ] [Chromosome: 7] [Map Location: 7\|7 F5] [De | 1.828640445 | 0.000467 | 0.001296 |
| 15460 | ENSMUSG00000042851 | Zc3h6 | 78751 [Gene Symbol: Zc3h6] [Locus Tag: ] [Chromosome: 2] [Map Location: 2\|2 F1] [De | 1.828633098 | 0.000255 | 0.000743 |
| 7119 | ENSMUSG00000034801 | Sos2 | 20663 [Gene Symbol: Sos2] [Locus Tag: ] [Chromosome: 12] [Map Location: 12 28.93 cM | 1.828294594 | 0.000158 | 0.000479 |
| 6628 | ENSMUSG00000033454 | Zbtb1 | 268564 [Gene Symbol: Zbtb1] [Locus Tag: ] [Chromosome: 12] [Map Location: 12\|12 C3 | 1.828056767 | 0.000511 | 0.001408 |
| 11669 | ENSMUSG00000039318 | Rab3gap2 | 98732 [Gene Symbol: Rab3gap2] [Locus Tag: ] [Chromosome: 1] [Map Location: 1\|1 H5 | 1.827871041 | 0.000877 | 0.00231 |
| 7588 | ENSMUSG00000010505 | Myt1 | 17932 [Gene Symbol: Myt1] [Locus Tag: ] [Chromosome: 2] [Map Location: 2 H4\|2 103. | 1.827870318 | 0.003179 | 0.00741 |
| 4437 | ENSMUSG00000032366 | Tpm1 | 22003 [Gene Symbol: Tpm1] [Locus Tag: ] [Chromosome: 9] [Map Location: 9 C\|9 36.27 | 1.827087291 | 0.000643 | 0.001736 |
| 7660 | ENSMUSG00000043153 | Ispd | 75847 [Gene Symbol: Ispd] [Locus Tag: ] [Chromosome: 12] [Map Location: 12\|12 A3] [ | 1.82684142 | 0.003443 | 0.00795 |
| 13162 | ENSMUSG00000021905 | Dph3 | 105638 [Gene Symbol: Dph3] [Locus Tag: ] [Chromosome: 14] [Map Location: 14\|14 B] | 1.82673777 | 0.001321 | 0.003357 |
| 8434 | ENSMUSG00000070348 | Ccnd1 | 12443 [Gene Symbol: Ccnd1] [Locus Tag: ] [Chromosome: 7] [Map Location: 7 F5\|7 88.9 | 1.826670513 | 0.001641 | 0.004081 |
| 14190 | ENSMUSG00000046432 | Ngfrap1 | 12070 [Gene Symbol: Ngfrap1] [Locus Tag: ] [Chromosome: X] [Map Location: X F1\|X 57 | 1.82618494 | 0.000287 | 0.00083 |
| 525 | ENSMUSG00000040717 | Il17rd | 171463 [Gene Symbol: Il17rd] [Locus Tag: ] [Chromosome: 14] [Map Location: 14\|14 A | 1.825497275 | 0.002111 | 0.005113 |
| 3795 | ENSMUSG00000046798 | Cldn12 | 64945 [Gene Symbol: Cldn12] [Locus Tag: ] [Chromosome: 5] [Map Location: 5\|5 A1] [D | 1.825421852 | 0.003014 | 0.007064 |
| 2620 | ENSMUSG00000026836 | Acvr1 | 11477 [Gene Symbol: Acvr1] [Locus Tag: ] [Chromosome: 2] [Map Location: 2\|2 C1.1] [ | 1.825283262 | 0.000389 | 0.001098 |
| 15707 | ENSMUSG00000047583 | Tyw3 | 209584 [Gene Symbol: Tyw3] [Locus Tag: ] [Chromosome: 3] [Map Location: 3\|3 H4] [D | 1.824933733 | 0.000555 | 0.001517 |
| 9529 | ENSMUSG00000061559 | Wdr61 | 66317 [Gene Symbol: Wdr61] [Locus Tag: ] [Chromosome: 9] [Map Location: 9\|9 C] [De | 1.824817143 | 0.003912 | 0.008923 |
| 8156 | ENSMUSG00000038806 | Sde2 | 208768 [Gene Symbol: Sde2] [Locus Tag: ] [Chromosome: 1] [Map Location: 1\|1 H4] [De | 1.824107656 | 0.001692 | 0.00419 |
| 17849 | ENSMUSG00000020611 | Gna13 | 14674 [Gene Symbol: Gna13] [Locus Tag: ] [Chromosome: 11] [Map Location: 11 E1\|11 | 1.822427136 | 0.000438 | 0.001224 |
| 18525 | ENSMUSG00000031583 | Wrn; LOC105242468 | 22427 [Gene Symbol: Wrn] [Locus Tag: ] [Chromosome: 8] [Map Location: 8 20.3 cM\|8 | 1.822224294 | 0.00097 | 0.002529 |
| 4650 | ENSMUSG00000030254 | Rad18 | 58186 [Gene Symbol: Rad18] [Locus Tag: ] [Chromosome: 6] [Map Location: 6\|6 F] [Des | 1.82124086 | 0.004657 | 0.010438 |
| 16228 | ENSMUSG00000056459 | Zbtb25 | 109929 [Gene Symbol: Zbtb25] [Locus Tag: ] [Chromosome: 12] [Map Location: 12 D2\| | 1.82119461 | 0.002128 | 0.00515 |
| 5152 | ENSMUSG00000032504 | Pdcd6ip | 18571 [Gene Symbol: Pdcd6ip] [Locus Tag: ] [Chromosome: 9] [Map Location: 9\|9 F2] [ | 1.821008314 | 0.000342 | 0.000976 |
| 6072 | ENSMUSG00000005583 | Mef2c | 17260 [Gene Symbol: Mef2c] [Locus Tag: ] [Chromosome: 13] [Map Location: 13 C3\|13 | 1.820732122 | 0.014554 | 0.029111 |
| 3073 | ENSMUSG00000021986 | Amer2 | 72125 [Gene Symbol: Amer2] [Locus Tag: ] [Chromosome: 14] [Map Location: 14\|14 D1 | 1.820527214 | 0.000397 | 0.001119 |
| 7438 | ENSMUSG00000038240 | Pdss2 | 71365 [Gene Symbol: Pdss2] [Locus Tag: ] [Chromosome: 10] [Map Location: 10 B2\|10 | 1.81867795 | 0.001752 | 0.004324 |
| 15201 | ENSMUSG00000028899 | Taf12 | 66464 [Gene Symbol: Taf12] [Locus Tag: ] [Chromosome: 4] [Map Location: 4\|4 D2.3] [ | 1.818237709 | 0.00617 | 0.013462 |
| 13710 | ENSMUSG00000025577 | Cbx2 | 12416 [Gene Symbol: Cbx2] [Locus Tag: ] [Chromosome: 11] [Map Location: 11 E2\|11 8 | 1.818138232 | 0.005879 | 0.01289 |
| 17729 | ENSMUSG00000042699 | Dhx9 | 13211 [Gene Symbol: Dhx9] [Locus Tag: ] [Chromosome: 1] [Map Location: 1 G3\|1 65.3 | 1.817330614 | 0.000183 | 0.000548 |
| 4243 | ENSMUSG00000033701 | Acbd6 | 72482 [Gene Symbol: Acbd6] [Locus Tag: ] [Chromosome: 1] [Map Location: 1\|1 G3] [De | 1.816820653 | 0.001441 | 0.003628 |
| 14066 | ENSMUSG00000015804 | Med28 | 66999 [Gene Symbol: Med28] [Locus Tag: ] [Chromosome: 5] [Map Location: 5\|5 B3] [D | 1.816346304 | 0.00495 | 0.011043 |
| 7033 | ENSMUSG00000020075 | Ddx21 | 56200 [Gene Symbol: Ddx21] [Locus Tag: ] [Chromosome: 10] [Map Location: 10 32.43 | 1.816080003 | 0.000477 | 0.001321 |
| 16793 | ENSMUSG00000026788 | Zbtb43 | 71834 [Gene Symbol: Zbtb43] [Locus Tag: ] [Chromosome: 2] [Map Location: 2\|2 B] [De | 1.815935636 | 0.000769 | 0.002044 |
| 7558 | ENSMUSG00000027966 | Col11a1 | 12814 [Gene Symbol: Col11a1] [Locus Tag: ] [Chromosome: 3] [Map Location: 3 F3\|3 4 | 1.815646027 | 0.00332 | 0.007704 |
| 4121 | ENSMUSG00000057409 | Zfp53 | 24132 [Gene Symbol: Zfp53] [Locus Tag: ] [Chromosome: 17] [Map Location: 17 A3.2\|1 | 1.814410057 | 0.005293 | 0.011725 |
| 16768 | ENSMUSG00000022110 | Sucla2 | 20916 [Gene Symbol: Sucla2] [Locus Tag: ] [Chromosome: 14] [Map Location: 14\|14 D3 | 1.814199593 | 0.001213 | 0.003108 |
| 13706 | ENSMUSG00000020076 | Ddx50 | 94213 [Gene Symbol: Ddx50] [Locus Tag: ] [Chromosome: 10] [Map Location: 10\|10 B4 | 1.813976095 | 0.000877 | 0.00231 |
| 10966 | ENSMUSG00000074182 | Znhit6 | 229937 [Gene Symbol: Znhit6] [Locus Tag: ] [Chromosome: 3] [Map Location: 3\|3 H3] [ | 1.813689549 | 0.012597 | 0.025575 |
| 866 | ENSMUSG00000033949 | Trim36 | 28105 [Gene Symbol: Trim36] [Locus Tag: ] [Chromosome: 18] [Map Location: 18 C\|18 | 1.813341013 | 0.000788 | 0.002089 |
| 15736 | ENSMUSG00000037343 | Taf2 | 319944 [Gene Symbol: Taf2] [Locus Tag: ] [Chromosome: 15] [Map Location: 15\|15 D1] | 1.812915009 | 0.000775 | 0.002059 |
| 13050 | ENSMUSG00000038879 | Nipal2 | 223473 [Gene Symbol: Nipal2] [Locus Tag: ] [Chromosome: 15] [Map Location: 15\|15 B | 1.812885946 | 0.018439 | 0.035868 |
| 2507 | ENSMUSG00000026283 | Ing5 | 66262 [Gene Symbol: Ing5] [Locus Tag: ] [Chromosome: 1] [Map Location: 1\|1 D] [Descr | 1.811959676 | 0.001581 | 0.003942 |
| 18004 | ENSMUSG00000036501 | Fam13b | 225358 [Gene Symbol: Fam13b] [Locus Tag: ] [Chromosome: 18] [Map Location: 18\|18 | 1.811576537 | 0.001068 | 0.002763 |
| 17616 | ENSMUSG00000042063 | Zfp386 | 56220 [Gene Symbol: Zfp386] [Locus Tag: ] [Chromosome: 12] [Map Location: 12\|12 F2 | 1.811530487 | 0.001664 | 0.004128 |
| 9445 | ENSMUSG00000070632 | Btbd8 | 100503185 [Gene Symbol: Btbd8] [Locus Tag: ] [Chromosome: 5] [Map Location: 5\|5 E | 1.811241599 | 0.018966 | 0.036759 |
| 2704 | ENSMUSG00000020154 | Ptprb | 19263 [Gene Symbol: Ptprb] [Locus Tag: ] [Chromosome: 10] [Map Location: 10\|10 D2 | 1.811143798 | 0.001628 | 0.00405 |
| 6825 | ENSMUSG00000058351 | Smim4 | 66487 [Gene Symbol: Smim4] [Locus Tag: ] [Chromosome: 14] [Map Location: 14\|14 B] | 1.81089187 | 0.001122 | 0.002894 |
| 2866 | ENSMUSG00000022441 | Efcab6 | 77627 [Gene Symbol: Efcab6] [Locus Tag: ] [Chromosome: 15] [Map Location: 15\|15 E3 | 1.810864671 | 0.001182 | 0.003036 |
| 198 | ENSMUSG00000051154 | Commd3 | 12238 [Gene Symbol: Commd3] [Locus Tag: ] [Chromosome: 2] [Map Location: 2 A3\|2 | 1.810287331 | 0.002078 | 0.00504 |
| 7589 | ENSMUSG00000031938 | 4931406C07Rik | 70984 [Gene Symbol: 4931406C07Rik] [Locus Tag: ] [Chromosome: 9] [Map Location: 9 | 1.808829591 | 0.003346 | 0.007754 |
| 5771 | ENSMUSG00000044627 | Swi5 | 72931 [Gene Symbol: Swi5] [Locus Tag: ] [Chromosome: 2] [Map Location: 2\|2 B] [Descr | 1.808654432 | 0.013019 | 0.026305 |
| 18246 | ENSMUSG00000022048 | Dpysl2 | 12934 [Gene Symbol: Dpysl2] [Locus Tag: ] [Chromosome: 14] [Map Location: 14 D1\|14 | 1.808556645 | 0.000174 | 0.000523 |
| 17109 | ENSMUSG00000039934 | Gsap | 212167 [Gene Symbol: Gsap] [Locus Tag: ] [Chromosome: 5] [Map Location: 5\|5 A3] [De | 1.808127957 | 0.005203 | 0.011547 |
| 16053 | ENSMUSG00000037465 | Klf10 | 21847 [Gene Symbol: Klf10] [Locus Tag: ] [Chromosome: 15] [Map Location: 15\|15 B3. | 1.807483647 | 0.003143 | 0.007335 |
| 13571 | ENSMUSG00000020982 | Nemf | 66244 [Gene Symbol: Nemf] [Locus Tag: ] [Chromosome: 12] [Map Location: 12\|12 C3] | 1.807391919 | 0.011423 | 0.023472 |
| 17796 | ENSMUSG00000035164 | Zc3h12c | 244871 [Gene Symbol: Zc3h12c] [Locus Tag: ] [Chromosome: 9] [Map Location: 9\|9 A5. | 1.806824491 | 0.00063 | 0.001704 |
| 7406 | ENSMUSG00000021578 | Ccdc127 | 67433 [Gene Symbol: Ccdc127] [Locus Tag: ] [Chromosome: 13] [Map Location: 13\|13 | 1.806764463 | 0.001437 | 0.003622 |
| 8247 | ENSMUSG00000000168 | Dlat | 235339 [Gene Symbol: Dlat] [Locus Tag: ] [Chromosome: 9] [Map Location: 9\|9 A5.3] [D | 1.805630166 | 0.00119 | 0.003054 |
| 14084 | ENSMUSG00000020818 | Mfsd11 | 69900 [Gene Symbol: Mfsd11] [Locus Tag: ] [Chromosome: 11] [Map Location: 11\|11 E | 1.805492744 | 0.010781 | 0.022274 |
| 13143 | ENSMUSG00000029326 | Enoph1 | 67870 [Gene Symbol: Enoph1] [Locus Tag: ] [Chromosome: 5] [Map Location: 5\|5 E4] [D | 1.805448383 | 0.001373 | 0.003478 |
| 16979 | ENSMUSG00000092558 | Med20 | 56771 [Gene Symbol: Med20] [Locus Tag: ] [Chromosome: 17] [Map Location: 17\|17 C] | 1.804226767 | 0.006629 | 0.014352 |
| 15899 | ENSMUSG00000033417 | Cacul1 | 78832 [Gene Symbol: Cacul1] [Locus Tag: ] [Chromosome: 19] [Map Location: 19\|19 D3 | 1.80381837 | 0.000756 | 0.002014 |
| 11632 | ENSMUSG00000045092 | S1pr1 | 13609 [Gene Symbol: S1pr1] [Locus Tag: ] [Chromosome: 3] [Map Location: 3\|3 G1] [De | 1.803502402 | 0.001645 | 0.004088 |
| 13078 | ENSMUSG00000030329 | Pianp | 319352 [Gene Symbol: Pianp] [Locus Tag: ] [Chromosome: 6] [Map Location: 6\|6 F2] [D | 1.802887144 | 0.000416 | 0.001166 |
| 17547 | ENSMUSG00000027742 | Cog6 | 67542 [Gene Symbol: Cog6] [Locus Tag: ] [Chromosome: 3] [Map Location: 3\|3 D] [Desc | 1.802617957 | 0.000898 | 0.002357 |
| 12507 | ENSMUSG00000049630 | C1ql3 | 227580 [Gene Symbol: C1ql3] [Locus Tag: ] [Chromosome: 2] [Map Location: 2\|2 A1] [D | 1.802589432 | 0.006222 | 0.01356 |
| 7999 | ENSMUSG00000024764 | Naa40 | 70999 [Gene Symbol: Naa40] [Locus Tag: ] [Chromosome: 19] [Map Location: 19\|19 A] | 1.802487024 | 0.00286 | 0.006729 |

| | | | | | | |
|---|---|---|---|---|---|---|
| 13607 | ENSMUSG00000021385 | Ippk | 75678 [Gene Symbol: Ippk] [Locus Tag: ] [Chromosome: 13] [Map Location: 13\|13 A5] [ | 1.802231708 | 0.002182 | 0.005269 |
| 8862 | ENSMUSG00000034601 | 2700049A03Rik | 76967 [Gene Symbol: 2700049A03Rik] [Locus Tag: ] [Chromosome: 12] [Map Location: | 1.802080215 | 0.004299 | 0.009727 |
| 6192 | ENSMUSG00000063060 | Sox7 | 20680 [Gene Symbol: Sox7] [Locus Tag: ] [Chromosome: 14] [Map Location: 14\|14 C3] | 1.801703233 | 0.021908 | 0.041796 |
| 3700 | ENSMUSG00000020122 | Egfr | 13649 [Gene Symbol: Egfr] [Locus Tag: ] [Chromosome: 11] [Map Location: 11 9.41 cM\| | 1.80168467 | 0.00085 | 0.002242 |
| 2399 | ENSMUSG00000032437 | Stt3b | 68292 [Gene Symbol: Stt3b] [Locus Tag: ] [Chromosome: 9] [Map Location: 9\|9 F3] [Des | 1.801678719 | 0.001131 | 0.002914 |
| 16272 | ENSMUSG00000033430 | Terf2ip | 57321 [Gene Symbol: Terf2ip] [Locus Tag: ] [Chromosome: 8] [Map Location: 8\|8 E1] [D | 1.799065246 | 0.001135 | 0.002923 |
| 2303 | ENSMUSG00000063434 | Sorcs3 | 66673 [Gene Symbol: Sorcs3] [Locus Tag: ] [Chromosome: 19] [Map Location: 19\|19 D1 | 1.797520577 | 0.005705 | 0.012538 |
| 5560 | ENSMUSG00000004364 | Cul3 | 26554 [Gene Symbol: Cul3] [Locus Tag: ] [Chromosome: 1] [Map Location: 1\|1 C4] [Des | 1.796775261 | 0.000248 | 0.000723 |
| 9383 | ENSMUSG00000034292 | Traf3ip1 | 74019 [Gene Symbol: Traf3ip1] [Locus Tag: ] [Chromosome: 1] [Map Location: 1\|1 D] [D | 1.796522368 | 0.004091 | 0.009288 |
| 4904 | ENSMUSG00000021868 | Ppif | 105675 [Gene Symbol: Ppif] [Locus Tag: ] [Chromosome: 14] [Map Location: 14\|14 A3] | 1.79496072 | 0.005066 | 0.011276 |
| 3442 | ENSMUSG00000022965 | Ifngr2 | 15980 [Gene Symbol: Ifngr2] [Locus Tag: ] [Chromosome: 16] [Map Location: 16 C3.3\|1 | 1.794890654 | 0.000247 | 0.000721 |
| 11238 | ENSMUSG00000026245 | Farsb | 23874 [Gene Symbol: Farsb] [Locus Tag: ] [Chromosome: 1] [Map Location: 1\|1 C4] [Des | 1.794696919 | 0.00173 | 0.004278 |
| 5516 | ENSMUSG00000031925 | Maml2 | 270118 [Gene Symbol: Maml2] [Locus Tag: ] [Chromosome: 9] [Map Location: 9\|9 A1] [ | 1.793699306 | 0.000416 | 0.001166 |
| 9624 | ENSMUSG00000035107 | Dcbld2 | 73379 [Gene Symbol: Dcbld2] [Locus Tag: ] [Chromosome: 16] [Map Location: 16\|16 C | 1.791648769 | 0.000663 | 0.001784 |
| 10856 | ENSMUSG00000002741 | Ykt6 | 56418 [Gene Symbol: Ykt6] [Locus Tag: ] [Chromosome: 11] [Map Location: 11\|11 A1] [ | 1.791415113 | 0.00038 | 0.001076 |
| 18293 | ENSMUSG00000038774 | Ascc3 | 77987 [Gene Symbol: Ascc3] [Locus Tag: ] [Chromosome: 10] [Map Location: 10\|10 B3] | 1.790619037 | 0.000359 | 0.00102 |
| 13520 | ENSMUSG00000001441 | Npepps | 19155 [Gene Symbol: Npepps] [Locus Tag: ] [Chromosome: 11] [Map Location: 11 D\|11 | 1.790374847 | 0.000387 | 0.001093 |
| 15811 | ENSMUSG00000047786 | Lix1 | 66643 [Gene Symbol: Lix1] [Locus Tag: ] [Chromosome: 17] [Map Location: 17\|17 A3.1 | 1.790216227 | 0.006185 | 0.013488 |
| 8576 | ENSMUSG00000057914 | Cacnb2 | 12296 [Gene Symbol: Cacnb2] [Locus Tag: ] [Chromosome: 2] [Map Location: 2 A2\|2 10 | 1.78948971 | 0.020548 | 0.039475 |
| 16892 | ENSMUSG00000026664 | Phyh | 16922 [Gene Symbol: Phyh] [Locus Tag: ] [Chromosome: 2] [Map Location: 2\|2 A1] [Des | 1.789020985 | 0.018936 | 0.036709 |
| 17373 | ENSMUSG00000022321 | Cdh10 | 320873 [Gene Symbol: Cdh10] [Locus Tag: ] [Chromosome: 15] [Map Location: 15 A2\|1 | 1.787824313 | 0.002746 | 0.006498 |
| 17476 | ENSMUSG00000000276 | Dgke | 56077 [Gene Symbol: Dgke] [Locus Tag: ] [Chromosome: 11] [Map Location: 11\|11 C] [D | 1.786465552 | 0.000571 | 0.001559 |
| 14043 | ENSMUSG00000016087 | Fli1 | 14247 [Gene Symbol: Fli1] [Locus Tag: ] [Chromosome: 9] [Map Location: 9 A4\|9 17.74 | 1.786068559 | 0.00469 | 0.010509 |
| 15080 | ENSMUSG00000008398 | Elk3 | 13713 [Gene Symbol: Elk3] [Locus Tag: ] [Chromosome: 10] [Map Location: 10 48.04 cM | 1.785803055 | 0.002484 | 0.005935 |
| 9971 | ENSMUSG00000044548 | Dact1 | 59036 [Gene Symbol: Dact1] [Locus Tag: ] [Chromosome: 12] [Map Location: 12\|12 C3] | 1.785751897 | 0.000463 | 0.001285 |
| 15216 | ENSMUSG00000091803 | Cox16 | 66272 [Gene Symbol: Cox16] [Locus Tag: ] [Chromosome: 12] [Map Location: 12\|12 D2 | 1.785164346 | 0.000717 | 0.001918 |
| 16856 | ENSMUSG00000068732 | Tmem167b | 67495 [Gene Symbol: Tmem167b] [Locus Tag: ] [Chromosome: 3] [Map Location: 3\|3 F | 1.784239244 | 0.002618 | 0.006219 |
| 13445 | ENSMUSG00000022494 | Shisa9 | 72555 [Gene Symbol: Shisa9] [Locus Tag: ] [Chromosome: 16] [Map Location: 16\|16 B1 | 1.783888448 | 0.002433 | 0.005821 |
| 3113 | ENSMUSG00000021670 | Hmgcr | 15357 [Gene Symbol: Hmgcr] [Locus Tag: ] [Chromosome: 13] [Map Location: 13 D1\|13 | 1.783758632 | 0.000496 | 0.001369 |
| 14138 | ENSMUSG00000036745 | Ttll7 | 70892 [Gene Symbol: Ttll7] [Locus Tag: ] [Chromosome: 3] [Map Location: 3\|3 H2] [Des | 1.782612477 | 0.001085 | 0.002803 |
| 1987 | ENSMUSG00000021981 | Cab39l | 69008 [Gene Symbol: Cab39l] [Locus Tag: ] [Chromosome: 14] [Map Location: 14\|14 C | 1.782070527 | 0.005393 | 0.011924 |
| 14196 | ENSMUSG00000030750 | Nsmce1 | 67711 [Gene Symbol: Nsmce1] [Locus Tag: ] [Chromosome: 7] [Map Location: 7\|7 F3] [D | 1.781539744 | 0.014777 | 0.02951 |
| 18490 | ENSMUSG00000031533 | Mrps31 | 57312 [Gene Symbol: Mrps31] [Locus Tag: ] [Chromosome: 8] [Map Location: 8\|8 A3] [D | 1.78142427 | 0.02606 | 0.048841 |
| 669 | ENSMUSG00000066392 | Nrxn3 | 18191 [Gene Symbol: Nrxn3] [Locus Tag: ] [Chromosome: 12] [Map Location: 12D2\|12 | 1.780288504 | 0.000276 | 0.0008 |
| 17167 | ENSMUSG00000052551 | Adarb2 | 94191 [Gene Symbol: Adarb2] [Locus Tag: ] [Chromosome: 13] [Map Location: 13\|13 A | 1.779596347 | 0.000567 | 0.001549 |
| 18324 | ENSMUSG00000071723 | Gspt2 | 14853 [Gene Symbol: Gspt2] [Locus Tag: ] [Chromosome: X] [Map Location: X\|X B1] [De | 1.778562314 | 0.01675 | 0.032937 |
| 8624 | ENSMUSG00000044149 | Nkrf | 77286 [Gene Symbol: Nkrf] [Locus Tag: ] [Chromosome: X] [Map Location: X\|X A3.3] [De | 1.778082215 | 0.002684 | 0.006361 |
| 15747 | ENSMUSG00000021115 | Vrk1 | 22367 [Gene Symbol: Vrk1] [Locus Tag: ] [Chromosome: 12] [Map Location: 12\|12 F1] [ | 1.777375737 | 0.003482 | 0.008034 |
| 1890 | ENSMUSG00000006386 | Tek | 21687 [Gene Symbol: Tek] [Locus Tag: ] [Chromosome: 4] [Map Location: 4 C5\|4 43.34 | 1.776870346 | 0.015472 | 0.030719 |
| 14923 | ENSMUSG00000038412 | Higd1a | 56295 [Gene Symbol: Higd1a] [Locus Tag: ] [Chromosome: 9] [Map Location: 9\|9 F4] [D | 1.776801909 | 0.026453 | 0.049508 |
| 15693 | ENSMUSG00000029004 | Kmt2e | 69188 [Gene Symbol: Kmt2e] [Locus Tag: ] [Chromosome: 5] [Map Location: 5\|5 A3] [D | 1.776723324 | 0.00045 | 0.001254 |
| 8428 | ENSMUSG00000025060 | Slk | 20874 [Gene Symbol: Slk] [Locus Tag: ] [Chromosome: 19] [Map Location: 19\|19 D2] [D | 1.776108938 | 0.000477 | 0.00132 |
| 5734 | ENSMUSG00000067879 | 3110035E14Rik | 76982 [Gene Symbol: 3110035E14Rik] [Locus Tag: ] [Chromosome: 1] [Map Location: 1 | 1.775554753 | 0.000731 | 0.001952 |
| 336 | ENSMUSG00000089762 | Ier5l | 72500 [Gene Symbol: Ier5l] [Locus Tag: ] [Chromosome: 2] [Map Location: 2\|2 B] [Descr | 1.775450579 | 0.003858 | 0.008814 |
| 5105 | ENSMUSG00000022064 | Pibf1 | 52023 [Gene Symbol: Pibf1] [Locus Tag: ] [Chromosome: 14] [Map Location: 14 E2.2\|14 | 1.775329839 | 0.004028 | 0.009168 |
| 2951 | ENSMUSG00000037259 | Dzank1 | 241688 [Gene Symbol: Dzank1] [Locus Tag: ] [Chromosome: 2] [Map Location: 2\|2 G1] | 1.773404395 | 0.001378 | 0.003488 |
| 15479 | ENSMUSG00000029735 | Tpk1 | 29807 [Gene Symbol: Tpk1] [Locus Tag: ] [Chromosome: 6] [Map Location: 6\|6 B2] [Des | 1.773018765 | 0.010433 | 0.021636 |
| 2681 | ENSMUSG00000026766 | Mmadhc | 109129 [Gene Symbol: Mmadhc] [Locus Tag: ] [Chromosome: 2] [Map Location: 2\|2 C1 | 1.772646885 | 0.003912 | 0.008923 |
| 2494 | ENSMUSG00000021552 | Gkap1 | 56278 [Gene Symbol: Gkap1] [Locus Tag: ] [Chromosome: 13] [Map Location: 13 B1\|13 | 1.772465098 | 0.021324 | 0.040791 |
| 12323 | ENSMUSG00000020525 | Ppm1d | 53892 [Gene Symbol: Ppm1d] [Locus Tag: ] [Chromosome: 11] [Map Location: 11 C\|11 | 1.771976171 | 0.004103 | 0.009311 |
| 18554 | ENSMUSG00000026575 | Nme7 | 171567 [Gene Symbol: Nme7] [Locus Tag: ] [Chromosome: 1] [Map Location: 1\|1 H2.2] | 1.771899098 | 0.001161 | 0.002985 |
| 5964 | ENSMUSG00000002578 | Ikzf4 | 22781 [Gene Symbol: Ikzf4] [Locus Tag: ] [Chromosome: 10] [Map Location: 10\|10 D3] | 1.771128468 | 0.0005 | 0.001379 |
| 13393 | ENSMUSG00000029469 | Ift81 | 12589 [Gene Symbol: Ift81] [Locus Tag: ] [Chromosome: 5] [Map Location: 5 F\|5 62.44 | 1.770874167 | 0.003437 | 0.007939 |
| 4720 | ENSMUSG00000022122 | Ednrb | 13618 [Gene Symbol: Ednrb] [Locus Tag: ] [Chromosome: 14] [Map Location: 14 E2.3\|1 | 1.770640268 | 0.002238 | 0.005389 |
| 7357 | ENSMUSG00000057363 | Uxs1 | 67883 [Gene Symbol: Uxs1] [Locus Tag: ] [Chromosome: 1] [Map Location: 1\|1 C1.1] [D | 1.770494657 | 0.002158 | 0.005217 |
| 1233 | ENSMUSG00000044043 | Pcdhb14 | 93885 [Gene Symbol: Pcdhb14] [Locus Tag: ] [Chromosome: 18] [Map Location: 18\|18 | 1.770308839 | 0.001913 | 0.004679 |
| 10355 | ENSMUSG00000005540 | Fcer2a | 14128 [Gene Symbol: Fcer2a] [Locus Tag: ] [Chromosome: 8] [Map Location: 8 1.92 cM\| | 1.770140142 | 0.010349 | 0.021473 |
| 1394 | ENSMUSG00000025173 | Wdr45b | 66840 [Gene Symbol: Wdr45b] [Locus Tag: ] [Chromosome: 11] [Map Location: 11 E2\|1 | 1.769671898 | 0.00094 | 0.002459 |
| 14286 | ENSMUSG00000035367 | Rmi1 | 74386 [Gene Symbol: Rmi1] [Locus Tag: ] [Chromosome: 13] [Map Location: 13\|13 B1] | 1.769630085 | 0.020383 | 0.039174 |
| 4525 | ENSMUSG00000022131 | Gpr180 | 58245 [Gene Symbol: Gpr180] [Locus Tag: ] [Chromosome: 14] [Map Location: 14\|14 E | 1.768884295 | 0.002277 | 0.005476 |
| 2562 | ENSMUSG00000036192 | Rorb | 225998 [Gene Symbol: Rorb] [Locus Tag: ] [Chromosome: 19] [Map Location: 19\|19 B] | 1.76883904 | 0.000358 | 0.001017 |
| 16171 | ENSMUSG00000006134 | Crkl | 12929 [Gene Symbol: Crkl] [Locus Tag: ] [Chromosome: 16] [Map Location: 16\|16 A3] [ | 1.76854466 | 0.011285 | 0.023217 |
| 3441 | ENSMUSG00000001300 | Efnb2 | 13642 [Gene Symbol: Efnb2] [Locus Tag: ] [Chromosome: 8] [Map Location: 8 A1.1\|8 3. | 1.768254673 | 0.003297 | 0.007655 |
| 8470 | ENSMUSG00000030641 | Ddias | 74041 [Gene Symbol: Ddias] [Locus Tag: ] [Chromosome: 7] [Map Location: 7\|7 E2] [De | 1.76764043 | 0.016056 | 0.031733 |
| 7689 | ENSMUSG00000026646 | Suv39h2 | 64707 [Gene Symbol: Suv39h2] [Locus Tag: ] [Chromosome: 2] [Map Location: 2 A\|2 1. | 1.766692591 | 0.002515 | 0.006 |
| 6347 | ENSMUSG00000028967 | Errfi1 | 74155 [Gene Symbol: Errfi1] [Locus Tag: ] [Chromosome: 4] [Map Location: 4\|4 E1] [De | 1.76667596 | 0.002257 | 0.005433 |
| 2025 | ENSMUSG00000039210 | Gpatch2 | 67769 [Gene Symbol: Gpatch2] [Locus Tag: ] [Chromosome: 1] [Map Location: 1\|1 H5] | 1.765835828 | 0.001528 | 0.003822 |
| 16631 | ENSMUSG00000070972 | Dnajc25 | 72429 [Gene Symbol: Dnajc25] [Locus Tag: ] [Chromosome: 4] [Map Location: 4\|4 B3] | 1.765773085 | 0.016784 | 0.033002 |
| 7762 | ENSMUSG00000026377 | Nifk | 67949 [Gene Symbol: Nifk] [Locus Tag: ] [Chromosome: 1] [Map Location: 1\|1 E2.3] [De | 1.765541247 | 0.01661 | 0.032707 |
| 14524 | ENSMUSG00000025949 | Pikfyve | 18711 [Gene Symbol: Pikfyve] [Locus Tag: ] [Chromosome: 1] [Map Location: 1\|1 C2] [D | 1.765469529 | 0.000659 | 0.001775 |
| 18073 | ENSMUSG00000016946 | Kctd5 | 69259 [Gene Symbol: Kctd5] [Locus Tag: ] [Chromosome: 17] [Map Location: 17\|17 A3. | 1.764456734 | 0.000565 | 0.001544 |
| 10121 | ENSMUSG00000070705 | Eid2b | 434156 [Gene Symbol: Eid2b] [Locus Tag: ] [Chromosome: 7] [Map Location: 7\|7 A3] [D | 1.764061605 | 0.001476 | 0.003707 |
| 10174 | ENSMUSG00000041220 | Elovl6 | 170439 [Gene Symbol: Elovl6] [Locus Tag: ] [Chromosome: 3] [Map Location: 3\|3 G3] [ | 1.763240448 | 0.000767 | 0.002041 |
| 17394 | ENSMUSG00000022019 | Tdrd3 | 219249 [Gene Symbol: Tdrd3] [Locus Tag: ] [Chromosome: 14] [Map Location: 14\|14 E | 1.762883079 | 0.000614 | 0.001665 |
| 9943 | ENSMUSG00000007815 | Rhoa | 11848 [Gene Symbol: Rhoa] [Locus Tag: ] [Chromosome: 9] [Map Location: 9\|9 F2] [Des | 1.762318449 | 0.002088 | 0.00506 |
| 1337 | ENSMUSG00000021932 | Rnaseh2b | 67153 [Gene Symbol: Rnaseh2b] [Locus Tag: ] [Chromosome: 14] [Map Location: 14\|14 | 1.762233479 | 0.013002 | 0.026284 |
| 7240 | ENSMUSG00000026179 | Pnkd | 56695 [Gene Symbol: Pnkd] [Locus Tag: ] [Chromosome: 1] [Map Location: 1\|1 C3] [Des | 1.761110909 | 0.000965 | 0.002518 |
| 11534 | ENSMUSG00000028975 | Pex14 | 56273 [Gene Symbol: Pex14] [Locus Tag: ] [Chromosome: 4] [Map Location: 4\|4 E2] [De | 1.760529277 | 0.00285 | 0.006715 |
| 5059 | ENSMUSG00000049090 | Zadh2 | 225791 [Gene Symbol: Zadh2] [Locus Tag: ] [Chromosome: 18] [Map Location: 18\|18 E | 1.760507108 | 0.0026 | 0.00618 |
| 11647 | ENSMUSG00000051855 | Mest | 17294 [Gene Symbol: Mest] [Locus Tag: ] [Chromosome: 6] [Map Location: 6 12.53 cM\| | 1.760204286 | 0.001207 | 0.003096 |
| 17870 | ENSMUSG00000035401 | Emsy | 233545 [Gene Symbol: Emsy] [Locus Tag: ] [Chromosome: 7] [Map Location: 7\|7 E2] [De | 1.76019428 | 0.0012 | 0.003079 |
| 17670 | ENSMUSG00000026623 | Lpgat1 | 226856 [Gene Symbol: Lpgat1] [Locus Tag: ] [Chromosome: 1] [Map Location: 1\|1 H6] [ | 1.759947777 | 0.000447 | 0.001246 |
| 15117 | ENSMUSG00000028370 | Pappa | 18491 [Gene Symbol: Pappa] [Locus Tag: ] [Chromosome: 4] [Map Location: 4 C1\|4 34. | 1.759861894 | 0.011038 | 0.022753 |
| 11978 | ENSMUSG00000033676 | Gabrb3 | 14402 [Gene Symbol: Gabrb3] [Locus Tag: ] [Chromosome: 7] [Map Location: 7 C\|7 33. | 1.759633573 | 0.000335 | 0.000957 |

| | | | | | | |
|---|---|---|---|---|---|---|
| 8728 | ENSMUSG00000040124 | Gorab | 98376 [Gene Symbol: Gorab] [Locus Tag: ] [Chromosome: 1] [Map Location: 1\|1 H2.1] [ | 1.758725321 | 0.015432 | 0.030646 |
| 8938 | ENSMUSG00000030447 | Cyfip1 | 20430 [Gene Symbol: Cyfip1] [Locus Tag: ] [Chromosome: 7] [Map Location: 7\|7 B5] [De | 1.758553481 | 0.001664 | 0.004128 |
| 8021 | ENSMUSG00000050240 | Hic2 | 58180 [Gene Symbol: Hic2] [Locus Tag: ] [Chromosome: 16] [Map Location: 16\|16 B1] [ | 1.758101347 | 0.000857 | 0.00226 |
| 2145 | ENSMUSG00000019143 | Hars2 | 70791 [Gene Symbol: Hars2] [Locus Tag: ] [Chromosome: 18] [Map Location: 18\|18 B2] | 1.756825776 | 0.006337 | 0.013776 |
| 8438 | ENSMUSG00000019960 | Dusp6 | 67603 [Gene Symbol: Dusp6] [Locus Tag: ] [Chromosome: 10] [Map Location: 10\|10 C3 | 1.756728719 | 0.001471 | 0.003697 |
| 8522 | ENSMUSG00000031134 | Rbmx | 19655 [Gene Symbol: Rbmx] [Locus Tag: ] [Chromosome: X] [Map Location: X\|X A5] [De | 1.754618075 | 0.001304 | 0.003316 |
| 17223 | ENSMUSG00000027209 | Fam227b | 75823 [Gene Symbol: Fam227b] [Locus Tag: ] [Chromosome: 2] [Map Location: 2\|2 F1] | 1.75438006 | 0.011225 | 0.023106 |
| 14604 | ENSMUSG00000021738 | Atxn7 | 246103 [Gene Symbol: Atxn7] [Locus Tag: ] [Chromosome: 14] [Map Location: 14\|14 A | 1.754270972 | 0.001172 | 0.003012 |
| 13812 | ENSMUSG00000024043 | Arhgap28 | 268970 [Gene Symbol: Arhgap28] [Locus Tag: ] [Chromosome: 17] [Map Location: 17\|1 | 1.753317544 | 0.001979 | 0.004825 |
| 6130 | ENSMUSG00000035614 | Fam179b | 328108 [Gene Symbol: Fam179b] [Locus Tag: ] [Chromosome: 12] [Map Location: 12\|1 | 1.752824345 | 0.000716 | 0.001915 |
| 17119 | ENSMUSG00000030878 | Cdr2 | 12585 [Gene Symbol: Cdr2] [Locus Tag: ] [Chromosome: 7] [Map Location: 7 F2\|7 65.0 | 1.751805252 | 0.0023 | 0.005527 |
| 17316 | ENSMUSG00000041261 | Car8 | 12319 [Gene Symbol: Car8] [Locus Tag: ] [Chromosome: 4] [Map Location: 4 A1\|4 3.53 | 1.751380016 | 0.019447 | 0.037574 |
| 12559 | ENSMUSG00000038608 | Dock10 | 210293 [Gene Symbol: Dock10] [Locus Tag: ] [Chromosome: 1] [Map Location: 1\|1 C4] | 1.751270903 | 0.010003 | 0.020846 |
| 8346 | ENSMUSG00000020705 | Ddx42 | 72047 [Gene Symbol: Ddx42] [Locus Tag: ] [Chromosome: 11] [Map Location: 11\|11 E1 | 1.750574971 | 0.003634 | 0.008359 |
| 9459 | ENSMUSG00000025591 | Tma16 | 66282 [Gene Symbol: Tma16] [Locus Tag: ] [Chromosome: 8] [Map Location: 8\|8 B3.3] | 1.750494103 | 0.009823 | 0.020498 |
| 13285 | ENSMUSG00000057561 | Eif1a | 13664 [Gene Symbol: Eif1a] [Locus Tag: ] [Chromosome: 18] [Map Location: 18\|18 C] [D | 1.749784898 | 0.001801 | 0.004433 |
| 1454 | ENSMUSG00000032531 | Amotl2 | 56332 [Gene Symbol: Amotl2] [Locus Tag: ] [Chromosome: 9] [Map Location: 9\|9 F1] [D | 1.749255141 | 0.003732 | 0.008557 |
| 16800 | ENSMUSG00000050321 | Neto1 | 246317 [Gene Symbol: Neto1] [Locus Tag: ] [Chromosome: 18] [Map Location: 18\|18 E | 1.748140027 | 0.005656 | 0.012444 |
| 17403 | ENSMUSG00000035778 | Ggta1 | 14594 [Gene Symbol: Ggta1] [Locus Tag: ] [Chromosome: 2] [Map Location: 2 B\|2 23.6 | 1.747905747 | 0.014782 | 0.029513 |
| 2501 | ENSMUSG00000021022 | Ppp2r3c | 59032 [Gene Symbol: Ppp2r3c] [Locus Tag: ] [Chromosome: 12] [Map Location: 12\|12 | 1.747672061 | 0.00868 | 0.018329 |
| 8841 | ENSMUSG00000037363 | Letm2 | 270035 [Gene Symbol: Letm2] [Locus Tag: ] [Chromosome: 8] [Map Location: 8\|8 A2] [ | 1.747526421 | 0.022296 | 0.042444 |
| 6827 | ENSMUSG00000021245 | Mlh3 | 217716 [Gene Symbol: Mlh3] [Locus Tag: ] [Chromosome: 12] [Map Location: 12\|12 D | 1.746904009 | 0.002493 | 0.005953 |
| 11176 | ENSMUSG00000021572 | Cep72 | 74470 [Gene Symbol: Cep72] [Locus Tag: ] [Chromosome: 13] [Map Location: 13\|13 C1 | 1.746462894 | 0.021707 | 0.041455 |
| 3534 | ENSMUSG00000005225 | Plekha8 | 231999 [Gene Symbol: Plekha8] [Locus Tag: ] [Chromosome: 6] [Map Location: 6\|6 B3] | 1.745383366 | 0.006151 | 0.013423 |
| 4661 | ENSMUSG00000045817 | Zfp36l2 | 12193 [Gene Symbol: Zfp36l2] [Locus Tag: ] [Chromosome: 17] [Map Location: 17\|17 E | 1.744976556 | 0.001865 | 0.004574 |
| 2163 | ENSMUSG00000037885 | Stk35 | 67333 [Gene Symbol: Stk35] [Locus Tag: ] [Chromosome: 2] [Map Location: 2\|2 F1] [De | 1.744655911 | 0.001592 | 0.003968 |
| 10329 | ENSMUSG00000019505 | Ubb | 22187 [Gene Symbol: Ubb] [Locus Tag: ] [Chromosome: 11] [Map Location: 11 B2\|11 3 | 1.743700216 | 0.018179 | 0.035404 |
| 14324 | ENSMUSG00000027827 | Kcnab1 | 16497 [Gene Symbol: Kcnab1] [Locus Tag: ] [Chromosome: 3] [Map Location: 3 E1\|3 30 | 1.74277861 | 0.001449 | 0.003648 |
| 16439 | ENSMUSG00000024970 | AI846148 | 68229 [Gene Symbol: AI846148] [Locus Tag: ] [Chromosome: 19] [Map Location: 19\|1 | 1.742706419 | 0.00221 | 0.005329 |
| 11847 | ENSMUSG00000025290 | LOC102642752; Rps2 | 102642752 [Gene Symbol: LOC102642752] [Locus Tag: ] [Chromosome: 16] [Map Loca | 1.741818619 | 0.006909 | 0.0149 |
| 17697 | ENSMUSG00000028383 | Hsdl2 | 72479 [Gene Symbol: Hsdl2] [Locus Tag: ] [Chromosome: 4] [Map Location: 4\|4 C1] [De | 1.741695431 | 0.009589 | 0.020064 |
| 7413 | ENSMUSG00000038048 | Cntnap5c | 620292 [Gene Symbol: Cntnap5c] [Locus Tag: ] [Chromosome: 17] [Map Location: 17\|1 | 1.741606503 | 0.002851 | 0.006715 |
| 6883 | ENSMUSG00000024087 | Cyp1b1 | 13078 [Gene Symbol: Cyp1b1] [Locus Tag: ] [Chromosome: 17] [Map Location: 17\|17 E | 1.741051086 | 0.025781 | 0.048381 |
| 11003 | ENSMUSG00000031673 | Cdh11 | 12552 [Gene Symbol: Cdh11] [Locus Tag: ] [Chromosome: 8] [Map Location: 8 D2\|8 50. | 1.740949805 | 0.000687 | 0.001846 |
| 21 | ENSMUSG00000022685 | Parn | 74108 [Gene Symbol: Parn] [Locus Tag: ] [Chromosome: 16] [Map Location: 16\|16 B1] [ | 1.740511586 | 0.004451 | 0.010027 |
| 14290 | ENSMUSG00000043441 | Gpr149 | 229357 [Gene Symbol: Gpr149] [Locus Tag: ] [Chromosome: 3] [Map Location: 3\|3 E1] | 1.740505098 | 0.005576 | 0.012291 |
| 13305 | ENSMUSG00000063600 | Egfem1 | 75740 [Gene Symbol: Egfem1] [Locus Tag: ] [Chromosome: 3] [Map Location: 3\|3 A3] [D | 1.738831009 | 0.009801 | 0.020455 |
| 73 | ENSMUSG00000034329 | Brip1 | 237911 [Gene Symbol: Brip1] [Locus Tag: ] [Chromosome: 11] [Map Location: 11\|11 C] | 1.738594962 | 0.026192 | 0.049048 |
| 678 | ENSMUSG00000085925 | Rtl1 | 353326 [Gene Symbol: Rtl1] [Locus Tag: ] [Chromosome: 12] [Map Location: 12\|12 F1] | 1.738543638 | 0.009354 | 0.019613 |
| 10846 | ENSMUSG00000022816 | Fstl1 | 14314 [Gene Symbol: Fstl1] [Locus Tag: ] [Chromosome: 16] [Map Location: 16 B3\|16 2 | 1.738246604 | 0.002769 | 0.006542 |
| 3029 | ENSMUSG00000040594 | Ranbp17 | 66011 [Gene Symbol: Ranbp17] [Locus Tag: ] [Chromosome: 11] [Map Location: 11\|11 | 1.737373322 | 0.003643 | 0.008375 |
| 18173 | ENSMUSG00000019970 | Sgk1 | 20393 [Gene Symbol: Sgk1] [Locus Tag: ] [Chromosome: 10] [Map Location: 10\|10 A3] [ | 1.737028211 | 0.001155 | 0.002971 |
| 6302 | ENSMUSG00000033721 | Vav3 | 57257 [Gene Symbol: Vav3] [Locus Tag: ] [Chromosome: 3] [Map Location: 3\|3 G1] [Des | 1.736677466 | 0.002502 | 0.005973 |
| 606 | ENSMUSG00000022246 | Rai14 | 75646 [Gene Symbol: Rai14] [Locus Tag: ] [Chromosome: 15] [Map Location: 15\|15 A2] | 1.736098062 | 0.005899 | 0.012928 |
| 15825 | ENSMUSG00000032009 | Sesn3 | 75747 [Gene Symbol: Sesn3] [Locus Tag: ] [Chromosome: 9] [Map Location: 9\|9 A3] [De | 1.735390101 | 0.001249 | 0.003189 |
| 5690 | ENSMUSG00000036208 | Nepro | 212547 [Gene Symbol: Nepro] [Locus Tag: ] [Chromosome: 16] [Map Location: 16\|16 B | 1.734690124 | 0.022145 | 0.042192 |
| 2706 | ENSMUSG00000023284 | Zfp605 | 675812 [Gene Symbol: Zfp605] [Locus Tag: ] [Chromosome: 5] [Map Location: 5\|5 F] [D | 1.734135202 | 0.000913 | 0.002393 |
| 16967 | ENSMUSG00000091476 | Gm16432 | 545391 [Gene Symbol: Gm16432] [Locus Tag: ] [Chromosome: 1] [Map Location: 1\|1 H | 1.734030084 | 0.003307 | 0.007675 |
| 621 | ENSMUSG00000051920 | Rspo2 | 239405 [Gene Symbol: Rspo2] [Locus Tag: ] [Chromosome: 15] [Map Location: 15 B3.1 | 1.733116912 | 0.006945 | 0.01497 |
| 17318 | ENSMUSG00000039831 | Arhgap29 | 214137 [Gene Symbol: Arhgap29] [Locus Tag: ] [Chromosome: 3] [Map Location: 3\|3 G | 1.732241274 | 0.0006 | 0.001629 |
| 10991 | ENSMUSG00000041716 | Gm20604 | 100859931 [Gene Symbol: Gm20604] [Locus Tag: ] [Chromosome: 12] [Map Location: | 1.730727617 | 0.001095 | 0.002826 |
| 9817 | ENSMUSG00000044167 | Foxo1 | 56458 [Gene Symbol: Foxo1] [Locus Tag: ] [Chromosome: 3] [Map Location: 3 C\|3 23.1 | 1.730451181 | 0.000775 | 0.002059 |
| 10835 | ENSMUSG00000044835 | Ankrd45 | 73844 [Gene Symbol: Ankrd45] [Locus Tag: ] [Chromosome: 1] [Map Location: 1\|1 H1] | 1.730350313 | 0.003215 | 0.007482 |
| 6457 | ENSMUSG00000030057 | Cnbp | 12785 [Gene Symbol: Cnbp] [Locus Tag: ] [Chromosome: 6] [Map Location: 6\|6 D1-D2] | 1.730157139 | 0.00137 | 0.003471 |
| 8225 | ENSMUSG00000038612 | Mcl1 | 17210 [Gene Symbol: Mcl1] [Locus Tag: ] [Chromosome: 3] [Map Location: 3 F2.1\|3 40. | 1.730147367 | 0.001269 | 0.003234 |
| 8070 | ENSMUSG00000004936 | Map2k1 | 26395 [Gene Symbol: Map2k1] [Locus Tag: ] [Chromosome: 9] [Map Location: 9 C\|9 34. | 1.729837224 | 0.005297 | 0.01173 |
| 15951 | ENSMUSG00000026640 | Plxna2 | 18845 [Gene Symbol: Plxna2] [Locus Tag: ] [Chromosome: 1] [Map Location: 1\|1 H6] [D | 1.729728188 | 0.017427 | 0.034093 |
| 10261 | ENSMUSG00000027971 | Ndst4 | 64580 [Gene Symbol: Ndst4] [Locus Tag: ] [Chromosome: 3] [Map Location: 3\|3 H1] [De | 1.729324335 | 0.002911 | 0.006839 |
| 1626 | ENSMUSG00000031772 | Cntnap4 | 170571 [Gene Symbol: Cntnap4] [Locus Tag: ] [Chromosome: 8] [Map Location: 8\|8 E1] | 1.728695368 | 0.001819 | 0.004472 |
| 16298 | ENSMUSG00000038658 | Ric1 | 226089 [Gene Symbol: Ric1] [Locus Tag: ] [Chromosome: 19] [Map Location: 19\|19 C1] | 1.728634734 | 0.001185 | 0.003044 |
| 7783 | ENSMUSG00000028832 | Stmn1 | 16765 [Gene Symbol: Stmn1] [Locus Tag: ] [Chromosome: 4] [Map Location: 4 D3\|4 66. | 1.728420095 | 0.001256 | 0.003205 |
| 7261 | ENSMUSG00000046191 | Pcdhb20 | 93891 [Gene Symbol: Pcdhb20] [Locus Tag: ] [Chromosome: 18] [Map Location: 18\|18 | 1.728237655 | 0.00207 | 0.005022 |
| 11731 | ENSMUSG00000052581 | Lrrtm4 | 243499 [Gene Symbol: Lrrtm4] [Locus Tag: ] [Chromosome: 6] [Map Location: 6\|6 C3] [ | 1.727881356 | 0.019301 | 0.037328 |
| 6085 | ENSMUSG00000032652 | Crebl2 | 232430 [Gene Symbol: Crebl2] [Locus Tag: ] [Chromosome: 6] [Map Location: 6\|6 G1] [ | 1.727704098 | 0.011157 | 0.022978 |
| 8381 | ENSMUSG00000020363 | Gfpt2 | 14584 [Gene Symbol: Gfpt2] [Locus Tag: ] [Chromosome: 11] [Map Location: 11 B1.2\|1 | 1.726233326 | 0.007616 | 0.016291 |
| 17903 | ENSMUSG00000032216 | Nedd4 | 17999 [Gene Symbol: Nedd4] [Locus Tag: ] [Chromosome: 9] [Map Location: 9 D\|9 40.0 | 1.726102366 | 0.00052 | 0.001431 |
| 3941 | ENSMUSG00000031555 | Adam9 | 11502 [Gene Symbol: Adam9] [Locus Tag: ] [Chromosome: 8] [Map Location: 8 A2\|8 13 | 1.725961728 | 0.001144 | 0.002946 |
| 3137 | ENSMUSG00000014769 | Psmb1 | 19170 [Gene Symbol: Psmb1] [Locus Tag: ] [Chromosome: 17] [Map Location: 17 A2\|17 | 1.725425396 | 0.003296 | 0.007653 |
| 16870 | ENSMUSG00000089824 | Rbm12 | 75710 [Gene Symbol: Rbm12] [Locus Tag: ] [Chromosome: 2] [Map Location: 2\|2 H2] [D | 1.724675701 | 0.001478 | 0.003709 |
| 13509 | ENSMUSG00000039357 | Fut11 | 73068 [Gene Symbol: Fut11] [Locus Tag: ] [Chromosome: 14] [Map Location: 14\|14 B] [ | 1.723864108 | 0.004652 | 0.010433 |
| 16617 | ENSMUSG00000067567 | Hdac8 | 70315 [Gene Symbol: Hdac8] [Locus Tag: ] [Chromosome: X] [Map Location: X\|X C3] [De | 1.723216494 | 0.005319 | 0.011776 |
| 10474 | ENSMUSG00000038024 | Dennd4c | 329877 [Gene Symbol: Dennd4c] [Locus Tag: ] [Chromosome: 4] [Map Location: 4\|4 C4 | 1.722359817 | 0.001743 | 0.004304 |
| 18166 | ENSMUSG00000040537 | Adam22 | 11496 [Gene Symbol: Adam22] [Locus Tag: ] [Chromosome: 5] [Map Location: 5 A1\|5 3 | 1.722350182 | 0.000708 | 0.001896 |
| 10116 | ENSMUSG00000026648 | Dclre1c | 227525 [Gene Symbol: Dclre1c] [Locus Tag: ] [Chromosome: 2] [Map Location: 2\|2 A1] | 1.722115344 | 0.002147 | 0.005192 |
| 16802 | ENSMUSG00000021327 | Zkscan3 | 72739 [Gene Symbol: Zkscan3] [Locus Tag: ] [Chromosome: 13] [Map Location: 13\|13 A | 1.722062085 | 0.002757 | 0.006521 |
| 13910 | ENSMUSG00000038525 | Armc10 | 67211 [Gene Symbol: Armc10] [Locus Tag: ] [Chromosome: 5] [Map Location: 5\|5 A3] [ | 1.722003602 | 0.005812 | 0.012755 |
| 10720 | ENSMUSG00000096916 | Zfp850 | 100043772 [Gene Symbol: Zfp850] [Locus Tag: ] [Chromosome: 7] [Map Location: 7 A3 | 1.721965943 | 0.01231 | 0.025061 |
| 15852 | ENSMUSG00000027774 | Gfm1 | 28030 [Gene Symbol: Gfm1] [Locus Tag: ] [Chromosome: 3] [Map Location: 3 E1\|3 30.9 | 1.721160785 | 0.004355 | 0.009834 |
| 3709 | ENSMUSG00000034110 | Kctd7 | 212919 [Gene Symbol: Kctd7] [Locus Tag: ] [Chromosome: 5] [Map Location: 5\|5 G1.3] | 1.719928628 | 0.006931 | 0.014946 |
| 3733 | ENSMUSG00000056999 | Ide | 15925 [Gene Symbol: Ide] [Locus Tag: ] [Chromosome: 19] [Map Location: 19 C2\|19 32 | 1.719915077 | 0.012044 | 0.024599 |
| 1595 | ENSMUSG00000015747 | Vps45 | 22365 [Gene Symbol: Vps45] [Locus Tag: ] [Chromosome: 3] [Map Location: 3 F2.1\|3 4 | 1.719398716 | 0.002847 | 0.00671 |
| 8508 | ENSMUSG00000045102 | Poln | 272158 [Gene Symbol: Poln] [Locus Tag: ] [Chromosome: 5] [Map Location: 5\|5 C1] [De | 1.717931343 | 0.015825 | 0.031353 |
| 4780 | ENSMUSG00000000278 | Scpep1 | 74617 [Gene Symbol: Scpep1] [Locus Tag: ] [Chromosome: 11] [Map Location: 11\|11 C | 1.717867883 | 0.022345 | 0.042521 |
| 10218 | ENSMUSG00000000861 | Bcl11a | 14025 [Gene Symbol: Bcl11a] [Locus Tag: ] [Chromosome: 11] [Map Location: 11\|11 A | 1.716564143 | 0.003178 | 0.007408 |

| | | | | | | |
|---|---|---|---|---|---|---|
| 3587 | ENSMUSG00000018697 | Aatf | 56321 [Gene Symbol: Aatf] [Locus Tag: ] [Chromosome: 11] [Map Location: 11\|11 B5] [ | 1.716266642 | 0.003455 | 0.007976 |
| 12280 | ENSMUSG00000053907 | Mat2a | 232087 [Gene Symbol: Mat2a] [Locus Tag: ] [Chromosome: 6] [Map Location: 6\|6 C1] [D | 1.714879058 | 0.001695 | 0.004196 |
| 16429 | ENSMUSG00000046441 | Cmtr2 | 234728 [Gene Symbol: Cmtr2] [Locus Tag: ] [Chromosome: 8] [Map Location: 8\|8 E1] [D | 1.714336132 | 0.02305 | 0.043705 |
| 10668 | ENSMUSG00000034024 | Cct2 | 12461 [Gene Symbol: Cct2] [Locus Tag: ] [Chromosome: 10] [Map Location: 10\|10 D2] [ | 1.714010355 | 0.001279 | 0.003256 |
| 11738 | ENSMUSG00000045062 | Pcdhb7 | 93878 [Gene Symbol: Pcdhb7] [Locus Tag: ] [Chromosome: 18] [Map Location: 18\|18 B | 1.713515511 | 0.01508 | 0.03 |
| 7837 | ENSMUSG00000000561 | Wdr77 | 70465 [Gene Symbol: Wdr77] [Locus Tag: ] [Chromosome: 3] [Map Location: 3\|3 F2.2] [ | 1.713512605 | 0.008509 | 0.018003 |
| 17877 | ENSMUSG00000053062 | Jam2 | 67374 [Gene Symbol: Jam2] [Locus Tag: ] [Chromosome: 16] [Map Location: 16\|16 C3. | 1.71116908 | 0.008685 | 0.018336 |
| 11187 | ENSMUSG00000044071 | Fam19a2 | 268354 [Gene Symbol: Fam19a2] [Locus Tag: ] [Chromosome: 10] [Map Location: 10\|1 | 1.711083054 | 0.000579 | 0.001578 |
| 10961 | ENSMUSG00000036478 | Btg1 | 12226 [Gene Symbol: Btg1] [Locus Tag: ] [Chromosome: 10] [Map Location: 10\|10 C3] [ | 1.711062994 | 0.00306 | 0.00716 |
| 17760 | ENSMUSG00000053846 | Lipg | 16891 [Gene Symbol: Lipg] [Locus Tag: ] [Chromosome: 18] [Map Location: 18\|18 E2] [ | 1.711024527 | 0.01257 | 0.025536 |
| 1 | ENSMUSG00000036323 | Srp72 | 66661 [Gene Symbol: Srp72] [Locus Tag: ] [Chromosome: 5] [Map Location: 5\|5 C3.3] [ | 1.710943841 | 0.001923 | 0.004699 |
| 7623 | ENSMUSG00000024668 | Sdhaf2 | 66072 [Gene Symbol: Sdhaf2] [Locus Tag: ] [Chromosome: 19] [Map Location: 19\|19 A] | 1.710851283 | 0.005083 | 0.011306 |
| 12935 | ENSMUSG00000022307 | Oxr1 | 170719 [Gene Symbol: Oxr1] [Locus Tag: ] [Chromosome: 15] [Map Location: 15\|15 D1 | 1.710327001 | 0.002628 | 0.006239 |
| 9608 | ENSMUSG00000019892 | Lrriq1 | 74978 [Gene Symbol: Lrriq1] [Locus Tag: ] [Chromosome: 10] [Map Location: 10\|10 D1 | 1.710102886 | 0.019155 | 0.037083 |
| 15402 | ENSMUSG00000037022 | Mmaa | 109136 [Gene Symbol: Mmaa] [Locus Tag: ] [Chromosome: 8] [Map Location: 8\|8 C2] [D | 1.709490936 | 0.005172 | 0.011486 |
| 7898 | ENSMUSG00000044221 | Grsf1 | 231413 [Gene Symbol: Grsf1] [Locus Tag: ] [Chromosome: 5] [Map Location: 5 E1\|5 43. | 1.708867349 | 0.002032 | 0.004942 |
| 9527 | ENSMUSG00000058291 | Zfp68 | 24135 [Gene Symbol: Zfp68] [Locus Tag: ] [Chromosome: 5] [Map Location: 5 G2\|5 77. | 1.708728396 | 0.003381 | 0.007827 |
| 10480 | ENSMUSG00000036510 | Cdh8 | 12564 [Gene Symbol: Cdh8] [Locus Tag: ] [Chromosome: 8] [Map Location: 8 D1\|8 49.4 | 1.707727005 | 0.000813 | 0.002152 |
| 3002 | ENSMUSG00000021824 | Ap3m1 | 55946 [Gene Symbol: Ap3m1] [Locus Tag: ] [Chromosome: 14] [Map Location: 14 A3\|1 | 1.707697336 | 0.003398 | 0.007859 |
| 3207 | ENSMUSG00000019794 | Katna1 | 23924 [Gene Symbol: Katna1] [Locus Tag: ] [Chromosome: 10] [Map Location: 10\|10 A | 1.706657109 | 0.026621 | 0.049797 |
| 5175 | ENSMUSG00000044308 | Ubr3 | 68795 [Gene Symbol: Ubr3] [Locus Tag: ] [Chromosome: 2] [Map Location: 2\|2 C2] [Des | 1.706562722 | 0.005281 | 0.0117 |
| 11308 | ENSMUSG00000059327 | Eda | 13607 [Gene Symbol: Eda] [Locus Tag: ] [Chromosome: X] [Map Location: X C3\|X 43.59 | 1.705827775 | 0.003286 | 0.007631 |
| 12813 | ENSMUSG00000024427 | Spry4 | 24066 [Gene Symbol: Spry4] [Locus Tag: ] [Chromosome: 18] [Map Location: 18 B3\|18 | 1.705749779 | 0.001972 | 0.00481 |
| 17146 | ENSMUSG00000058793 | Cds2 | 110911 [Gene Symbol: Cds2] [Locus Tag: ] [Chromosome: 2] [Map Location: 2 F2\|2 64. | 1.705048633 | 0.00307 | 0.007181 |
| 15478 | ENSMUSG00000026281 | Dtymk | 21915 [Gene Symbol: Dtymk] [Locus Tag: ] [Chromosome: 1] [Map Location: 1 D\|1 47.3 | 1.704459437 | 0.013598 | 0.027367 |
| 17427 | ENSMUSG00000026614 | Slc30a10 | 226781 [Gene Symbol: Slc30a10] [Locus Tag: ] [Chromosome: 1] [Map Location: 1\|1 H5 | 1.704423323 | 0.005899 | 0.012928 |
| 16276 | ENSMUSG00000031012 | Cask | 12361 [Gene Symbol: Cask] [Locus Tag: ] [Chromosome: X] [Map Location: X A1.1\|X 8.4 | 1.703153076 | 0.000578 | 0.001575 |
| 6325 | ENSMUSG00000025437 | Usp33 | 170822 [Gene Symbol: Usp33] [Locus Tag: ] [Chromosome: 3] [Map Location: 3\|3 H3] [ | 1.70308815 | 0.001703 | 0.004216 |
| 13151 | ENSMUSG00000052915 | Msl1 | 74026 [Gene Symbol: Msl1] [Locus Tag: ] [Chromosome: 11] [Map Location: 11\|11 D] [ | 1.70300594 | 0.000651 | 0.001755 |
| 14158 | ENSMUSG00000033632 | AW554918 | 225289 [Gene Symbol: AW554918] [Locus Tag: ] [Chromosome: 18] [Map Location: 18 | 1.702604054 | 0.001477 | 0.003708 |
| 18329 | ENSMUSG00000010721 | Lmbr1 | 56873 [Gene Symbol: Lmbr1] [Locus Tag: ] [Chromosome: 5] [Map Location: 5 B1\|5 14. | 1.700898157 | 0.002303 | 0.005532 |
| 3715 | ENSMUSG00000029722 | Agfg2 | 231801 [Gene Symbol: Agfg2] [Locus Tag: ] [Chromosome: 5] [Map Location: 5\|5 G2] [D | 1.700774863 | 0.017022 | 0.03338 |
| 11432 | ENSMUSG00000054477 | Kcnn2 | 140492 [Gene Symbol: Kcnn2] [Locus Tag: ] [Chromosome: 18] [Map Location: 18\|18 C | 1.700061264 | 0.001488 | 0.003732 |
| 4297 | ENSMUSG00000030314 | Atg7 | 74244 [Gene Symbol: Atg7] [Locus Tag: ] [Chromosome: 6] [Map Location: 6\|6 E3] [Desc | 1.700003312 | 0.024648 | 0.046474 |
| 13653 | ENSMUSG00000023307 | March5 | 69104 [Gene Symbol: March5] [Locus Tag: ] [Chromosome: 19] [Map Location: 19\|19 C | 1.69844916 | 0.002542 | 0.00606 |
| 16015 | ENSMUSG00000021918 | Nek4 | 23955 [Gene Symbol: Nek4] [Locus Tag: ] [Chromosome: 14] [Map Location: 14\|14 B] [ | 1.697879343 | 0.00419 | 0.009495 |
| 6057 | ENSMUSG00000073591 | Pcdhb22 | 93893 [Gene Symbol: Pcdhb22] [Locus Tag: ] [Chromosome: 18] [Map Location: 18\|18 | 1.697438662 | 0.001414 | 0.003572 |
| 4267 | ENSMUSG00000027374 | Mrps5 | 77721 [Gene Symbol: Mrps5] [Locus Tag: ] [Chromosome: 2] [Map Location: 2\|2 F3] [De | 1.695939578 | 0.006512 | 0.014132 |
| 8480 | ENSMUSG00000002297 | Dbf4 | 27214 [Gene Symbol: Dbf4] [Locus Tag: ] [Chromosome: 5] [Map Location: 5\|5 A2] [Des | 1.695606813 | 0.009617 | 0.020114 |
| 4520 | ENSMUSG00000052516 | Robo2 | 268902 [Gene Symbol: Robo2] [Locus Tag: ] [Chromosome: 16] [Map Location: 16\|16 C | 1.695535148 | 0.003283 | 0.007627 |
| 12830 | ENSMUSG00000025421 | Hdhd2 | 76987 [Gene Symbol: Hdhd2] [Locus Tag: ] [Chromosome: 18] [Map Location: 18\|18 E3 | 1.695519797 | 0.001644 | 0.004086 |
| 18459 | ENSMUSG00000050244 | Heatr1 | 217995 [Gene Symbol: Heatr1] [Locus Tag: ] [Chromosome: 13] [Map Location: 13 A1\|1 | 1.695125496 | 0.011781 | 0.02413 |
| 12928 | ENSMUSG00000021497 | Txndc15 | 69672 [Gene Symbol: Txndc15] [Locus Tag: ] [Chromosome: 13] [Map Location: 13\|13 | 1.695037938 | 0.010537 | 0.021824 |
| 11392 | ENSMUSG00000003070 | Efna2 | 13637 [Gene Symbol: Efna2] [Locus Tag: ] [Chromosome: 10] [Map Location: 10 C1\|10 | 1.694589311 | 0.002858 | 0.006727 |
| 5553 | ENSMUSG00000028653 | Trit1 | 66966 [Gene Symbol: Trit1] [Locus Tag: ] [Chromosome: 4] [Map Location: 4\|4 D2.2] [D | 1.694469197 | 0.012437 | 0.025289 |
| 14748 | ENSMUSG00000039168 | Dap | 223453 [Gene Symbol: Dap] [Locus Tag: ] [Chromosome: 15] [Map Location: 15\|15 B2] | 1.692877716 | 0.024778 | 0.046682 |
| 1756 | ENSMUSG00000034342 | Cbl | 12402 [Gene Symbol: Cbl] [Locus Tag: ] [Chromosome: 9] [Map Location: 9 A5.2\|9 24.7 | 1.692861523 | 0.001727 | 0.00427 |
| 18487 | ENSMUSG00000056770 | Setd3 | 52690 [Gene Symbol: Setd3] [Locus Tag: ] [Chromosome: 12] [Map Location: 12 F1\|12 | 1.692524049 | 0.001572 | 0.003921 |
| 7629 | ENSMUSG00000056820 | Tsnax | 53424 [Gene Symbol: Tsnax] [Locus Tag: ] [Chromosome: 8] [Map Location: 8 E2\|8 73.1 | 1.691240913 | 0.003396 | 0.007857 |
| 15665 | ENSMUSG00000051396 | Hspa14 | 50497 [Gene Symbol: Hspa14] [Locus Tag: ] [Chromosome: 2] [Map Location: 2\|2 A1] [D | 1.690076804 | 0.003598 | 0.008281 |
| 14264 | ENSMUSG00000062960 | Kdr | 16542 [Gene Symbol: Kdr] [Locus Tag: ] [Chromosome: 5] [Map Location: 5 C3.3\|5 40.2 | 1.689501554 | 0.017828 | 0.034786 |
| 17309 | ENSMUSG00000046442 | Ppm1e | 320472 [Gene Symbol: Ppm1e] [Locus Tag: ] [Chromosome: 11] [Map Location: 11\|11 | 1.689315507 | 0.000953 | 0.002491 |
| 2076 | ENSMUSG00000034973 | Dopey1 | 320615 [Gene Symbol: Dopey1] [Locus Tag: ] [Chromosome: 9] [Map Location: 9 E3.1\|9 | 1.689046663 | 0.001339 | 0.003396 |
| 10388 | ENSMUSG00000032217 | Rnf111 | 93836 [Gene Symbol: Rnf111] [Locus Tag: ] [Chromosome: 9] [Map Location: 9 D\|9 39. | 1.688815989 | 0.002373 | 0.005689 |
| 15831 | ENSMUSG00000024604 | Rbm22 | 66810 [Gene Symbol: Rbm22] [Locus Tag: ] [Chromosome: 18] [Map Location: 18\|18 D | 1.688097432 | 0.00337 | 0.007803 |
| 9223 | ENSMUSG00000022340 | Sybu | 319613 [Gene Symbol: Sybu] [Locus Tag: ] [Chromosome: 15] [Map Location: 15\|15 B3 | 1.688043196 | 0.00118 | 0.003031 |
| 11436 | ENSMUSG00000026504 | Sdccag8 | 76816 [Gene Symbol: Sdccag8] [Locus Tag: ] [Chromosome: 1] [Map Location: 1\|1 H3] [ | 1.687341622 | 0.001545 | 0.003859 |
| 17259 | ENSMUSG00000021666 | Gfm2 | 320806 [Gene Symbol: Gfm2] [Locus Tag: ] [Chromosome: 13] [Map Location: 13\|13 D | 1.68592851 | 0.010072 | 0.020966 |
| 90 | ENSMUSG00000020495 | Smg8 | 74133 [Gene Symbol: Smg8] [Locus Tag: ] [Chromosome: 11] [Map Location: 11\|11 C] [ | 1.685810878 | 0.019773 | 0.038137 |
| 14127 | ENSMUSG00000054469 | Lclat1 | 225010 [Gene Symbol: Lclat1] [Locus Tag: ] [Chromosome: 17] [Map Location: 17\|17 E | 1.685176543 | 0.003363 | 0.007789 |
| 4371 | ENSMUSG00000056158 | Car10 | 72605 [Gene Symbol: Car10] [Locus Tag: ] [Chromosome: 11] [Map Location: 11\|11 C] [ | 1.684942044 | 0.016442 | 0.032411 |
| 2811 | ENSMUSG00000046110 | Serinc4 | 574418 [Gene Symbol: Serinc4] [Locus Tag: ] [Chromosome: 2] [Map Location: 2\|2 E5] [ | 1.683938627 | 0.010539 | 0.021826 |
| 5673 | ENSMUSG00000062580 | Timm17a | 21854 [Gene Symbol: Timm17a] [Locus Tag: ] [Chromosome: 1] [Map Location: 1\|1 F] [ | 1.683932584 | 0.004382 | 0.009889 |
| 15083 | ENSMUSG00000030306 | Tmtc1 | 387314 [Gene Symbol: Tmtc1] [Locus Tag: ] [Chromosome: 6] [Map Location: 6\|6 G3] [ | 1.683853806 | 0.001203 | 0.003086 |
| 6812 | ENSMUSG00000049421 | Zfp260 | 26466 [Gene Symbol: Zfp260] [Locus Tag: ] [Chromosome: 7] [Map Location: 7 B1\|7 17 | 1.682720246 | 0.007155 | 0.015387 |
| 3602 | ENSMUSG00000023106 | Denr | 68184 [Gene Symbol: Denr] [Locus Tag: ] [Chromosome: 5] [Map Location: 5\|5 F] [Descr | 1.68261143 | 0.003781 | 0.008655 |
| 882 | ENSMUSG00000024588 | Fech | 14151 [Gene Symbol: Fech] [Locus Tag: ] [Chromosome: 18] [Map Location: 18 E1\|18 3 | 1.682354878 | 0.003236 | 0.007529 |
| 8833 | ENSMUSG00000022023 | Wbp4 | 22380 [Gene Symbol: Wbp4] [Locus Tag: ] [Chromosome: 14] [Map Location: 14\|14 D3 | 1.682183254 | 0.004692 | 0.010512 |
| 14532 | ENSMUSG00000049580 | Tsku | 244152 [Gene Symbol: Tsku] [Locus Tag: ] [Chromosome: 7] [Map Location: 7\|7 E2] [De | 1.681885709 | 0.021262 | 0.040689 |
| 5909 | ENSMUSG00000036983 | Tfb1m | 224481 [Gene Symbol: Tfb1m] [Locus Tag: ] [Chromosome: 17] [Map Location: 17\|17 A | 1.681492215 | 0.015053 | 0.029959 |
| 18199 | ENSMUSG00000022295 | Atp6v1c1 | 66335 [Gene Symbol: Atp6v1c1] [Locus Tag: ] [Chromosome: 15] [Map Location: 15\|15 | 1.680821398 | 0.00391 | 0.008921 |
| 1098 | ENSMUSG00000041734 | Kirrel | 170643 [Gene Symbol: Kirrel] [Locus Tag: ] [Chromosome: 3] [Map Location: 3\|3 F1] [D | 1.680424327 | 0.014912 | 0.029732 |
| 13010 | ENSMUSG00000022802 | Lmln | 239833 [Gene Symbol: Lmln] [Locus Tag: ] [Chromosome: 16] [Map Location: 16\|16 B3 | 1.679871377 | 0.004074 | 0.009256 |
| 10047 | ENSMUSG00000036368 | Rmdn2 | 381110 [Gene Symbol: Rmdn2] [Locus Tag: ] [Chromosome: 17] [Map Location: 17\|17 | 1.679701374 | 0.024474 | 0.046194 |
| 17853 | ENSMUSG00000026018 | Ica1l | 70375 [Gene Symbol: Ica1l] [Locus Tag: ] [Chromosome: 1] [Map Location: 1\|1 C2] [Des | 1.679415808 | 0.00096 | 0.002506 |
| 11289 | ENSMUSG00000021962 | Dcp1a | 75901 [Gene Symbol: Dcp1a] [Locus Tag: ] [Chromosome: 14] [Map Location: 14 B\|14 1 | 1.679339184 | 0.004584 | 0.010294 |
| 2321 | ENSMUSG00000041763 | Tpp2 | 22019 [Gene Symbol: Tpp2] [Locus Tag: ] [Chromosome: 1] [Map Location: 1 C1.1\|1 23 | 1.679062571 | 0.002679 | 0.00635 |
| 8081 | ENSMUSG00000031986 | Sprtn | 244666 [Gene Symbol: Sprtn] [Locus Tag: ] [Chromosome: 8] [Map Location: 8\|8 E2] [D | 1.677610985 | 0.010765 | 0.022247 |
| 4214 | ENSMUSG00000035277 | Arx | 11878 [Gene Symbol: Arx] [Locus Tag: ] [Chromosome: X] [Map Location: X C3\|X 41.05 | 1.67608034 | 0.022216 | 0.042309 |
| 9116 | ENSMUSG00000059208 | Hnrnpm | 76936 [Gene Symbol: Hnrnpm] [Locus Tag: ] [Chromosome: 17] [Map Location: 17\|17 | 1.675865627 | 0.002312 | 0.005554 |
| 10302 | ENSMUSG00000042743 | Sgtb | 218544 [Gene Symbol: Sgtb] [Locus Tag: ] [Chromosome: 13] [Map Location: 13\|13 D1 | 1.675748757 | 0.005347 | 0.01183 |
| 15648 | ENSMUSG00000022883 | Robo1 | 19876 [Gene Symbol: Robo1] [Locus Tag: ] [Chromosome: 16] [Map Location: 16\|16 C3 | 1.675669332 | 0.007297 | 0.015671 |
| 2537 | ENSMUSG00000026705 | Klhl20 | 226541 [Gene Symbol: Klhl20] [Locus Tag: ] [Chromosome: 1] [Map Location: 1\|1 H2.1 | 1.673772739 | 0.00347 | 0.008008 |
| 13709 | ENSMUSG00000029401 | Rilpl2 | 80291 [Gene Symbol: Rilpl2] [Locus Tag: ] [Chromosome: 5] [Map Location: 5\|5 F] [Desc | 1.67256961 | 0.021777 | 0.041576 |

| | | | | | | |
|---|---|---|---|---|---|---|
| 10518 | ENSMUSG00000057614 | Gnai1 | 14677 [Gene Symbol: Gnai1] [Locus Tag: ] [Chromosome: 5] [Map Location: 5 A3\|5 8.1( | 1.671992017 | 0.004136 | 0.009381 |
| 13261 | ENSMUSG00000000708 | Kat2b | 18519 [Gene Symbol: Kat2b] [Locus Tag: ] [Chromosome: 17] [Map Location: 17 C\|17 2 | 1.671513062 | 0.007936 | 0.016915 |
| 5913 | ENSMUSG00000074637 | Sox2 | 20674 [Gene Symbol: Sox2] [Locus Tag: ] [Chromosome: 3] [Map Location: 3 16.93 cM\| | 1.671236045 | 0.011998 | 0.024506 |
| 9064 | ENSMUSG00000000884 | Gnb1l | 13972 [Gene Symbol: Gnb1l] [Locus Tag: ] [Chromosome: 16] [Map Location: 16 A3\|16 | 1.671116997 | 0.003691 | 0.008473 |
| 12371 | ENSMUSG00000057134 | Ado | 211488 [Gene Symbol: Ado] [Locus Tag: ] [Chromosome: 10] [Map Location: 10\|10 B5.: | 1.668815434 | 0.001858 | 0.004559 |
| 2792 | ENSMUSG00000031360 | Ctps2 | 55936 [Gene Symbol: Ctps2] [Locus Tag: ] [Chromosome: X] [Map Location: X\|X F5] [De: | 1.668761515 | 0.003665 | 0.008419 |
| 11573 | ENSMUSG00000077450 | Rab11b | 19326 [Gene Symbol: Rab11b] [Locus Tag: ] [Chromosome: 17] [Map Location: 17 B1\|1 | 1.668488507 | 0.003109 | 0.007264 |
| 760 | ENSMUSG00000024165 | Hn1l | 52009 [Gene Symbol: Hn1l] [Locus Tag: ] [Chromosome: 17] [Map Location: 17 A3.3\|17 | 1.667315074 | 0.007463 | 0.015998 |
| 17841 | ENSMUSG00000030421 | Uri1 | 19777 [Gene Symbol: Uri1] [Locus Tag: ] [Chromosome: 7] [Map Location: 7\|7 B2] [Desc | 1.667043693 | 0.001665 | 0.00413 |
| 8031 | ENSMUSG00000020205 | Phlda1 | 21664 [Gene Symbol: Phlda1] [Locus Tag: ] [Chromosome: 10] [Map Location: 10\|10 D: | 1.665872925 | 0.012686 | 0.025726 |
| 6701 | ENSMUSG00000053192 | Mllt11 | 56772 [Gene Symbol: Mllt11] [Locus Tag: ] [Chromosome: 3] [Map Location: 3\|3 F2.1] [ | 1.665385107 | 0.001502 | 0.003765 |
| 8407 | ENSMUSG00000047888 | Tnrc6b | 213988 [Gene Symbol: Tnrc6b] [Locus Tag: ] [Chromosome: 15] [Map Location: 15\|15 I | 1.665231909 | 0.001457 | 0.003666 |
| 17110 | ENSMUSG00000031821 | Gins2 | 272551 [Gene Symbol: Gins2] [Locus Tag: ] [Chromosome: 8] [Map Location: 8\|8 E1] [D | 1.66508001 | 0.021212 | 0.040609 |
| 659 | ENSMUSG00000048216 | Gpr85 | 64450 [Gene Symbol: Gpr85] [Locus Tag: ] [Chromosome: 6] [Map Location: 6\|6 A1] [De | 1.664955372 | 0.001499 | 0.003758 |
| 15730 | ENSMUSG00000039316 | Rftn1 | 76438 [Gene Symbol: Rftn1] [Locus Tag: ] [Chromosome: 17] [Map Location: 17\|17 C] [ | 1.664816986 | 0.010313 | 0.021405 |
| 3234 | ENSMUSG00000027210 | Meis2 | 17536 [Gene Symbol: Meis2] [Locus Tag: ] [Chromosome: 2] [Map Location: 2 E4-E5\|2 5 | 1.661867532 | 0.002649 | 0.006283 |
| 11585 | ENSMUSG00000056019 | Zfp709 | 236193 [Gene Symbol: Zfp709] [Locus Tag: ] [Chromosome: 8] [Map Location: 8\|8 B3.3 | 1.660564251 | 0.008093 | 0.017217 |
| 12691 | ENSMUSG00000027312 | Atrn | 11990 [Gene Symbol: Atrn] [Locus Tag: ] [Chromosome: 2] [Map Location: 2 F1\|2 63.26 | 1.660477728 | 0.001596 | 0.003977 |
| 16169 | ENSMUSG00000025128 | Bhlhe22 | 59058 [Gene Symbol: Bhlhe22] [Locus Tag: ] [Chromosome: 3] [Map Location: 3\|3 A2] [ | 1.66039651 | 0.003429 | 0.007923 |
| 14822 | ENSMUSG00000041328 | Pcf11 | 74737 [Gene Symbol: Pcf11] [Locus Tag: ] [Chromosome: 7] [Map Location: 7\|7 E2] [De: | 1.659888124 | 0.001538 | 0.003844 |
| 56 | ENSMUSG00000021373 | Cap2 | 67252 [Gene Symbol: Cap2] [Locus Tag: ] [Chromosome: 13] [Map Location: 13\|13 A5] | 1.658759475 | 0.010785 | 0.02228 |
| 18312 | ENSMUSG00000010476 | Ebf3 | 13593 [Gene Symbol: Ebf3] [Locus Tag: ] [Chromosome: 7] [Map Location: 7\|7 F3-F4] [D | 1.657977334 | 0.002801 | 0.006609 |
| 7657 | ENSMUSG00000044550 | Tceal3 | 594844 [Gene Symbol: Tceal3] [Locus Tag: ] [Chromosome: X] [Map Location: X\|X F1] [D | 1.657929769 | 0.016749 | 0.032937 |
| 1825 | ENSMUSG00000089682 | Bcl2l2 | 12050 [Gene Symbol: Bcl2l2] [Locus Tag: ] [Chromosome: 14] [Map Location: 14 C3\|14 | 1.657715522 | 0.005855 | 0.012843 |
| 767 | ENSMUSG00000021036 | Sptlc2 | 20773 [Gene Symbol: Sptlc2] [Locus Tag: ] [Chromosome: 12] [Map Location: 12\|12 E]\| | 1.657508471 | 0.002852 | 0.006717 |
| 1686 | ENSMUSG00000028039 | Efna3 | 13638 [Gene Symbol: Efna3] [Locus Tag: ] [Chromosome: 3] [Map Location: 3 F1\|3 39.0 | 1.657071219 | 0.002838 | 0.006691 |
| 8780 | ENSMUSG00000053799 | Exoc6 | 107371 [Gene Symbol: Exoc6] [Locus Tag: ] [Chromosome: 19] [Map Location: 19 C2\|1! | 1.656692809 | 0.002772 | 0.006547 |
| 15442 | ENSMUSG00000049672 | Zbtb14 | 22666 [Gene Symbol: Zbtb14] [Locus Tag: ] [Chromosome: 17] [Map Location: 17 E1.3\| | 1.653657429 | 0.003513 | 0.008098 |
| 3646 | ENSMUSG00000048271 | Rbm33 | 381626 [Gene Symbol: Rbm33] [Locus Tag: ] [Chromosome: 5] [Map Location: 5\|5 B1] [ | 1.653458906 | 0.002313 | 0.005555 |
| 3961 | ENSMUSG00000018412 | Kansl1 | 76719 [Gene Symbol: Kansl1] [Locus Tag: ] [Chromosome: 11] [Map Location: 11\|11 E1 | 1.650879342 | 0.001153 | 0.002967 |
| 1995 | ENSMUSG00000033306 | Lpp | 210126 [Gene Symbol: Lpp] [Locus Tag: ] [Chromosome: 16] [Map Location: 16\|16 B1] | 1.64973582 | 0.002881 | 0.006773 |
| 9917 | ENSMUSG00000036299 | BC031181; LOC1026 | 407819 [Gene Symbol: BC031181] [Locus Tag: ] [Chromosome: 18] [Map Location: 18\| | 1.649186468 | 0.018731 | 0.036368 |
| 5876 | ENSMUSG00000064363 | ND4 | 17719 [Gene Symbol: ND4] [Locus Tag: ] [Chromosome: MT] [Map Location: ] [Descripti | 1.648640226 | 0.002411 | 0.005775 |
| 1638 | ENSMUSG00000057265 | Bbof1 | 72873 [Gene Symbol: Bbof1] [Locus Tag: ] [Chromosome: 12] [Map Location: 12\|12 D1 | 1.647782437 | 0.01867 | 0.036273 |
| 14484 | ENSMUSG00000001366 | Fbxo9 | 71538 [Gene Symbol: Fbxo9] [Locus Tag: ] [Chromosome: 9] [Map Location: 9\|9 E1] [De | 1.647668111 | 0.009717 | 0.020299 |
| 9179 | ENSMUSG00000020440 | Arf5 | 11844 [Gene Symbol: Arf5] [Locus Tag: ] [Chromosome: 6] [Map Location: 6\|6 A3.3] [De | 1.646380298 | 0.006617 | 0.014333 |
| 10412 | ENSMUSG00000056476 | Med12l | 329650 [Gene Symbol: Med12l] [Locus Tag: ] [Chromosome: 3] [Map Location: 3\|3 D] [ | 1.646135722 | 0.001735 | 0.004287 |
| 10197 | ENSMUSG00000035580 | Kcnh8 | 211468 [Gene Symbol: Kcnh8] [Locus Tag: ] [Chromosome: 17] [Map Location: 17\|17 C | 1.645917557 | 0.011049 | 0.022772 |
| 5823 | ENSMUSG00000021365 | Nedd9 | 18003 [Gene Symbol: Nedd9] [Locus Tag: ] [Chromosome: 13] [Map Location: 13\|13 A3 | 1.645302377 | 0.010477 | 0.021711 |
| 6524 | ENSMUSG00000021610 | Clptm1l | 218335 [Gene Symbol: Clptm1l] [Locus Tag: ] [Chromosome: 13] [Map Location: 13\|13 | 1.645277256 | 0.007977 | 0.016989 |
| 4028 | ENSMUSG00000037876 | Jmjd1c | 108829 [Gene Symbol: Jmjd1c] [Locus Tag: ] [Chromosome: 10] [Map Location: 10\|10 I | 1.645021567 | 0.001755 | 0.004329 |
| 8491 | ENSMUSG00000039840 | Epg5 | 100502841 [Gene Symbol: Epg5] [Locus Tag: ] [Chromosome: 18] [Map Location: 18\|1 | 1.644391187 | 0.003316 | 0.007696 |
| 5683 | ENSMUSG00000021645 | Smn1 | 20595 [Gene Symbol: Smn1] [Locus Tag: ] [Chromosome: 13] [Map Location: 13 52.99 | 1.644174904 | 0.01443 | 0.028887 |
| 10436 | ENSMUSG00000041945 | Mfsd9 | 211798 [Gene Symbol: Mfsd9] [Locus Tag: ] [Chromosome: 1] [Map Location: 1\|1 B] [De | 1.643866312 | 0.021881 | 0.041748 |
| 115 | ENSMUSG00000040848 | Sft2d2 | 108735 [Gene Symbol: Sft2d2] [Locus Tag: ] [Chromosome: 1] [Map Location: 1\|1 H2.3 | 1.643733685 | 0.011682 | 0.02394 |
| 16703 | ENSMUSG00000024109 | Nrxn1 | 18189 [Gene Symbol: Nrxn1] [Locus Tag: ] [Chromosome: 17] [Map Location: 17\|17 E5 | 1.643487457 | 0.002146 | 0.005189 |
| 3955 | ENSMUSG00000033933 | Vhl | 22346 [Gene Symbol: Vhl] [Locus Tag: ] [Chromosome: 6] [Map Location: 6 E3\|6 52.81 | 1.643475653 | 0.009347 | 0.019602 |
| 7794 | ENSMUSG00000033392 | Clasp2 | 76499 [Gene Symbol: Clasp2] [Locus Tag: ] [Chromosome: 9] [Map Location: 9\|9 F2] [De | 1.643007179 | 0.001339 | 0.003397 |
| 17483 | ENSMUSG00000027977 | Ndst3 | 83398 [Gene Symbol: Ndst3] [Locus Tag: ] [Chromosome: 3] [Map Location: 3\|3 G3] [De | 1.642646329 | 0.017267 | 0.033812 |
| 1170 | ENSMUSG00000005089 | Slc1a2 | 20511 [Gene Symbol: Slc1a2] [Locus Tag: ] [Chromosome: 2] [Map Location: 2 E2\|2 54. | 1.641707758 | 0.001275 | 0.003246 |
| 4210 | ENSMUSG00000050565 | Tor1aip2 | 240832 [Gene Symbol: Tor1aip2] [Locus Tag: ] [Chromosome: 1] [Map Location: 1\|1 G3 | 1.640441121 | 0.010205 | 0.021215 |
| 11321 | ENSMUSG00000042348 | Arl15 | 218639 [Gene Symbol: Arl15] [Locus Tag: ] [Chromosome: 13] [Map Location: 13\|13 D2 | 1.639054776 | 0.002864 | 0.006738 |
| 3897 | ENSMUSG00000028436 | Dcaf12 | 68970 [Gene Symbol: Dcaf12] [Locus Tag: ] [Chromosome: 4] [Map Location: 4\|4 B1] [D | 1.638965144 | 0.009968 | 0.020779 |
| 12010 | ENSMUSG00000027259 | Adal | 75894 [Gene Symbol: Adal] [Locus Tag: ] [Chromosome: 2] [Map Location: 2\|2 F1] [Desc | 1.638692589 | 0.013428 | 0.02706 |
| 7610 | ENSMUSG00000036218 | Pdzrn4 | 239618 [Gene Symbol: Pdzrn4] [Locus Tag: ] [Chromosome: 15] [Map Location: 15\|15 I | 1.638561114 | 0.004293 | 0.009715 |
| 3720 | ENSMUSG00000068205 | Macrod2 | 72899 [Gene Symbol: Macrod2] [Locus Tag: ] [Chromosome: 2] [Map Location: 2\|2 F3]\| | 1.637887955 | 0.001372 | 0.003476 |
| 13264 | ENSMUSG00000027845 | Dclre1b | 140917 [Gene Symbol: Dclre1b] [Locus Tag: ] [Chromosome: 3] [Map Location: 3\|3 F2.2 | 1.637867485 | 0.014658 | 0.029294 |
| 14462 | ENSMUSG00000028608 | 0610037L13Rik | 74098 [Gene Symbol: 0610037L13Rik] [Locus Tag: ] [Chromosome: 4] [Map Location: 4 | 1.636626255 | 0.019654 | 0.037931 |
| 6523 | ENSMUSG00000036766 | Dner | 227325 [Gene Symbol: Dner] [Locus Tag: ] [Chromosome: 1] [Map Location: 1\|1 C5] [De | 1.636116827 | 0.003337 | 0.007739 |
| 11483 | ENSMUSG00000059923 | Grb2 | 14784 [Gene Symbol: Grb2] [Locus Tag: ] [Chromosome: 11] [Map Location: 11 E2\|11 8 | 1.635869235 | 0.002425 | 0.005805 |
| 16020 | ENSMUSG00000058656 | Samd12 | 320679 [Gene Symbol: Samd12] [Locus Tag: ] [Chromosome: 15] [Map Location: 15\|15 | 1.635795095 | 0.007617 | 0.016291 |
| 6250 | ENSMUSG00000020143 | Dock2 | 94176 [Gene Symbol: Dock2] [Locus Tag: ] [Chromosome: 11] [Map Location: 11\|11 A5 | 1.635783308 | 0.005128 | 0.011397 |
| 12753 | ENSMUSG00000038121 | Fam210a | 108654 [Gene Symbol: Fam210a] [Locus Tag: ] [Chromosome: 18] [Map Location: 18\|1 | 1.635569903 | 0.003845 | 0.008788 |
| 11948 | ENSMUSG00000044147 | Arf6 | 11845 [Gene Symbol: Arf6] [Locus Tag: ] [Chromosome: 12] [Map Location: 12\|12 C2] [ | 1.63535694 | 0.00323 | 0.007516 |
| 8781 | ENSMUSG00000002129 | Sf3a1 | 67465 [Gene Symbol: Sf3a1] [Locus Tag: ] [Chromosome: 11] [Map Location: 11\|11 A1] | 1.634596568 | 0.002227 | 0.005365 |
| 10852 | ENSMUSG00000034263 | Vwa9 | 69882 [Gene Symbol: Vwa9] [Locus Tag: ] [Chromosome: 9] [Map Location: 9\|9 C] [Desc | 1.63451628 | 0.012371 | 0.025173 |
| 101 | ENSMUSG00000041741 | Pde3a | 54611 [Gene Symbol: Pde3a] [Locus Tag: ] [Chromosome: 6] [Map Location: 6\|6 G1] [De | 1.633419873 | 0.007996 | 0.017025 |
| 2942 | ENSMUSG00000053684 | BC048403 | 270802 [Gene Symbol: BC048403] [Locus Tag: ] [Chromosome: 10] [Map Location: 10\| | 1.633007511 | 0.014427 | 0.028884 |
| 15417 | ENSMUSG00000031227 | Magee1 | 107528 [Gene Symbol: Magee1] [Locus Tag: ] [Chromosome: X] [Map Location: X\|X C3] | 1.630343366 | 0.002022 | 0.00492 |
| 5662 | ENSMUSG00000057335 | Cep170 | 545389 [Gene Symbol: Cep170] [Locus Tag: ] [Chromosome: 1] [Map Location: 1\|1 H4] | 1.629468728 | 0.00168 | 0.004165 |
| 11687 | ENSMUSG00000026491 | Ahctf1 | 226747 [Gene Symbol: Ahctf1] [Locus Tag: ] [Chromosome: 1] [Map Location: 1 H4\|1 8: | 1.629100687 | 0.005398 | 0.011935 |
| 7435 | ENSMUSG00000034780 | B3galt1 | 26877 [Gene Symbol: B3galt1] [Locus Tag: ] [Chromosome: 2] [Map Location: 2\|2 C3] [D | 1.62874359 | 0.002499 | 0.005966 |
| 10376 | ENSMUSG00000027351 | Spred1 | 114715 [Gene Symbol: Spred1] [Locus Tag: ] [Chromosome: 2] [Map Location: 2\|2 E5] [ | 1.626890525 | 0.00175 | 0.004321 |
| 7693 | ENSMUSG00000027438 | Napb | 17957 [Gene Symbol: Napb] [Locus Tag: ] [Chromosome: 2] [Map Location: 2\|2 G3] [De | 1.62618749 | 0.002459 | 0.005879 |
| 13175 | ENSMUSG00000033900 | Map9 | 213582 [Gene Symbol: Map9] [Locus Tag: ] [Chromosome: 3] [Map Location: 3\|3 E3] [D | 1.625817062 | 0.001419 | 0.003583 |
| 9916 | ENSMUSG00000042203 | Tbc1d22b | 381085 [Gene Symbol: Tbc1d22b] [Locus Tag: ] [Chromosome: 17] [Map Location: 17\|: | 1.625659792 | 0.00186 | 0.004563 |
| 18214 | ENSMUSG00000022194 | Pabpn1 | 54196 [Gene Symbol: Pabpn1] [Locus Tag: ] [Chromosome: 14] [Map Location: 14 C3\|1 | 1.625524483 | 0.002898 | 0.006812 |
| 10103 | ENSMUSG00000024382 | Ercc3 | 13872 [Gene Symbol: Ercc3] [Locus Tag: ] [Chromosome: 18] [Map Location: 18\|18 B3] | 1.624876904 | 0.005054 | 0.011251 |
| 18304 | ENSMUSG00000029465 | Arpc3 | 56378 [Gene Symbol: Arpc3] [Locus Tag: ] [Chromosome: 5] [Map Location: 5\|5 F] [Desc | 1.624134081 | 0.007713 | 0.016475 |
| 4380 | ENSMUSG00000047747 | Rnf150 | 330812 [Gene Symbol: Rnf150] [Locus Tag: ] [Chromosome: 8] [Map Location: 8\|8 C2] [ | 1.623329237 | 0.003946 | 0.008994 |
| 4681 | ENSMUSG00000095432 | Zfp748 | 212276 [Gene Symbol: Zfp748] [Locus Tag: ] [Chromosome: 13] [Map Location: 13\|13 I | 1.62324949 | 0.026097 | 0.048891 |
| 2901 | ENSMUSG00000021945 | Zmym2 | 76007 [Gene Symbol: Zmym2] [Locus Tag: ] [Chromosome: 14] [Map Location: 14\|14 C | 1.622753118 | 0.002544 | 0.006064 |
| 6767 | ENSMUSG00000032382 | Snx1 | 56440 [Gene Symbol: Snx1] [Locus Tag: ] [Chromosome: 9] [Map Location: 9\|9 D] [Desc | 1.622635714 | 0.014252 | 0.028566 |
| 12378 | ENSMUSG00000075324 | Fign | 60344 [Gene Symbol: Fign] [Locus Tag: ] [Chromosome: 2] [Map Location: 2 C1.3\|2 37.: | 1.621777929 | 0.010879 | 0.022457 |

| | | | | | | |
|---|---|---|---|---|---|---|
| 626 | ENSMUSG00000030386 | Zfp606 | 67370 [Gene Symbol: Zfp606] [Locus Tag: ] [Chromosome: 7] [Map Location: 7\|7 A2] [D | 1.621342957 | 0.005459 | 0.012053 |
| 9496 | ENSMUSG00000096472 | Cdkn2d | 12581 [Gene Symbol: Cdkn2d] [Locus Tag: ] [Chromosome: 9] [Map Location: 9\|9 A3] [D | 1.621064276 | 0.011879 | 0.024299 |
| 6120 | ENSMUSG00000002948 | Map2k7 | 26400 [Gene Symbol: Map2k7] [Locus Tag: ] [Chromosome: 8] [Map Location: 8\|8 A1.1 | 1.620789158 | 0.007365 | 0.015808 |
| 11539 | ENSMUSG00000041921 | Metap1d | 66559 [Gene Symbol: Metap1d] [Locus Tag: ] [Chromosome: 2] [Map Location: 2\|2 C2] | 1.620531807 | 0.024555 | 0.046321 |
| 14433 | ENSMUSG00000043313 | Pcdhb19 | 93890 [Gene Symbol: Pcdhb19] [Locus Tag: ] [Chromosome: 18] [Map Location: 18\|18 | 1.619526283 | 0.004975 | 0.011094 |
| 3195 | ENSMUSG00000044060 | A830010M20Rik | 231570 [Gene Symbol: A830010M20Rik] [Locus Tag: ] [Chromosome: 5] [Map Location | 1.617636705 | 0.011082 | 0.022833 |
| 1685 | ENSMUSG00000041992 | Rapgef5 | 217944 [Gene Symbol: Rapgef5] [Locus Tag: ] [Chromosome: 12] [Map Location: 12\|12 | 1.617241996 | 0.002029 | 0.004934 |
| 6386 | ENSMUSG00000036435 | Exoc1 | 69940 [Gene Symbol: Exoc1] [Locus Tag: ] [Chromosome: 5] [Map Location: 5\|5 E1] [De | 1.616081041 | 0.017196 | 0.033686 |
| 429 | ENSMUSG00000020458 | Rtn4 | 68585 [Gene Symbol: Rtn4] [Locus Tag: ] [Chromosome: 11] [Map Location: 11\|11 A3.3 | 1.615666236 | 0.00225 | 0.005417 |
| 16907 | ENSMUSG00000022995 | Enah | 13800 [Gene Symbol: Enah] [Locus Tag: ] [Chromosome: 1] [Map Location: 1 H5\|1 84.9 | 1.615185868 | 0.003665 | 0.008419 |
| 15558 | ENSMUSG00000075478 | Slitrk1 | 76965 [Gene Symbol: Slitrk1] [Locus Tag: ] [Chromosome: 14] [Map Location: 14\|14 E3 | 1.613797108 | 0.002939 | 0.006905 |
| 13756 | ENSMUSG00000030726 | Pold3 | 67967 [Gene Symbol: Pold3] [Locus Tag: ] [Chromosome: 7] [Map Location: 7\|7 F1] [De | 1.612809938 | 0.009609 | 0.020103 |
| 111 | ENSMUSG00000041961 | Znrf3 | 407821 [Gene Symbol: Znrf3] [Locus Tag: ] [Chromosome: 11] [Map Location: 11\|11 A1 | 1.611205758 | 0.004765 | 0.010662 |
| 106 | ENSMUSG00000060548 | Tnfrsf19 | 29820 [Gene Symbol: Tnfrsf19] [Locus Tag: ] [Chromosome: 14] [Map Location: 14\|14 D | 1.61093822 | 0.00445 | 0.010027 |
| 5354 | ENSMUSG00000024143 | Rhoq | 104215 [Gene Symbol: Rhoq] [Locus Tag: ] [Chromosome: 17] [Map Location: 17 E4\|17 | 1.610754638 | 0.004411 | 0.009944 |
| 4765 | ENSMUSG00000027854 | Sike1 | 66641 [Gene Symbol: Sike1] [Locus Tag: ] [Chromosome: 3] [Map Location: 3\|3 F3] [Des | 1.61035299 | 0.005308 | 0.011753 |
| 343 | ENSMUSG00000069895 | Atxn1l | 52335 [Gene Symbol: Atxn1l] [Locus Tag: ] [Chromosome: 8] [Map Location: 8 D3\|8 57. | 1.60985392 | 0.003321 | 0.007705 |
| 4355 | ENSMUSG00000038679 | Trps1 | 83925 [Gene Symbol: Trps1] [Locus Tag: ] [Chromosome: 15] [Map Location: 15 C\|15 1 | 1.608813475 | 0.004248 | 0.009616 |
| 4386 | ENSMUSG00000055065 | Ddx17 | 67040 [Gene Symbol: Ddx17] [Locus Tag: ] [Chromosome: 15] [Map Location: 15\|15 E1 | 1.608802627 | 0.006537 | 0.014173 |
| 5201 | ENSMUSG00000036298 | Slc2a13 | 239606 [Gene Symbol: Slc2a13] [Locus Tag: ] [Chromosome: 15] [Map Location: 15\|15 | 1.608489353 | 0.012682 | 0.02572 |
| 10602 | ENSMUSG00000092060 | Bend4 | 666938 [Gene Symbol: Bend4] [Locus Tag: ] [Chromosome: 5] [Map Location: 5 C3.1\|5] | 1.607455557 | 0.006629 | 0.014352 |
| 4856 | ENSMUSG00000023960 | Enpp5 | 83965 [Gene Symbol: Enpp5] [Locus Tag: ] [Chromosome: 17] [Map Location: 17 20.3 c | 1.607132975 | 0.006948 | 0.014976 |
| 3388 | ENSMUSG00000028550 | Atg4c | 242557 [Gene Symbol: Atg4c] [Locus Tag: ] [Chromosome: 4] [Map Location: 4\|4 C6] [D | 1.607069567 | 0.0168 | 0.033012 |
| 16577 | ENSMUSG00000046387 | Pcdhb17 | 93888 [Gene Symbol: Pcdhb17] [Locus Tag: ] [Chromosome: 18] [Map Location: 18\|18 | 1.606668419 | 0.009276 | 0.019458 |
| 8733 | ENSMUSG00000039578 | Ccser1 | 232035 [Gene Symbol: Ccser1] [Locus Tag: ] [Chromosome: 6] [Map Location: 6\|6 B3] [ | 1.605083428 | 0.00243 | 0.005815 |
| 5545 | ENSMUSG00000020088 | Sar1a | 20224 [Gene Symbol: Sar1a] [Locus Tag: ] [Chromosome: 10] [Map Location: 10\|10 B4] | 1.604736198 | 0.007914 | 0.016877 |
| 12174 | ENSMUSG00000035024 | Ncapd3 | 78658 [Gene Symbol: Ncapd3] [Locus Tag: ] [Chromosome: 9] [Map Location: 9\|9 A4] [D | 1.60448769 | 0.008858 | 0.018681 |
| 2067 | ENSMUSG00000030189 | Ybx3 | 56449 [Gene Symbol: Ybx3] [Locus Tag: ] [Chromosome: 6] [Map Location: 6\|6 F3] [Des | 1.604016067 | 0.005444 | 0.012024 |
| 4058 | ENSMUSG00000094114 | Gm7120 | 633640 [Gene Symbol: Gm7120] [Locus Tag: ] [Chromosome: 13] [Map Location: 13\|1 | 1.602684573 | 0.026205 | 0.049068 |
| 762 | ENSMUSG00000024131 | Slc3a1 | 20532 [Gene Symbol: Slc3a1] [Locus Tag: ] [Chromosome: 17] [Map Location: 17 E4\|17 | 1.602051897 | 0.003673 | 0.008437 |
| 7551 | ENSMUSG00000052031 | Tagap1 | 380608 [Gene Symbol: Tagap1] [Locus Tag: ] [Chromosome: 17] [Map Location: 17\|17 / | 1.601676779 | 0.015612 | 0.030969 |
| 6596 | ENSMUSG00000015127 | Unkl | 74154 [Gene Symbol: Unkl] [Locus Tag: ] [Chromosome: 17] [Map Location: 17\|17 A3.3 | 1.601656015 | 0.007505 | 0.016077 |
| 10250 | ENSMUSG00000055022 | Cntn1 | 12805 [Gene Symbol: Cntn1] [Locus Tag: ] [Chromosome: 15] [Map Location: 15 46.39 | 1.601573532 | 0.006114 | 0.013346 |
| 10594 | ENSMUSG00000033106 | Slc7a6os | 66432 [Gene Symbol: Slc7a6os] [Locus Tag: ] [Chromosome: 8] [Map Location: 8\|8 D3] | 1.601042666 | 0.014722 | 0.02941 |
| 14942 | ENSMUSG00000019990 | Pde7b | 29863 [Gene Symbol: Pde7b] [Locus Tag: ] [Chromosome: 10] [Map Location: 10\|10 A3 | 1.600048145 | 0.005408 | 0.011949 |
| 9715 | ENSMUSG00000021843 | Ktn1 | 16709 [Gene Symbol: Ktn1] [Locus Tag: ] [Chromosome: 14] [Map Location: 14\|14 C2] | 1.598966623 | 0.012331 | 0.025099 |
| 11341 | ENSMUSG00000018171 | Vmp1 | 75909 [Gene Symbol: Vmp1] [Locus Tag: ] [Chromosome: 11] [Map Location: 11\|11 C] [ | 1.59841713 | 0.019286 | 0.037302 |
| 16565 | ENSMUSG00000045180 | Shroom2 | 110380 [Gene Symbol: Shroom2] [Locus Tag: ] [Chromosome: X] [Map Location: X F2-F3 | 1.597991396 | 0.002564 | 0.006107 |
| 8098 | ENSMUSG00000038602 | Slc35f1 | 215085 [Gene Symbol: Slc35f1] [Locus Tag: ] [Chromosome: 10] [Map Location: 10\|10 | 1.597022517 | 0.004557 | 0.01024 |
| 4833 | ENSMUSG00000038949 | Cnst | 226744 [Gene Symbol: Cnst] [Locus Tag: ] [Chromosome: 1] [Map Location: 1\|1 H4] [De | 1.595769625 | 0.01197 | 0.024454 |
| 13610 | ENSMUSG00000031365 | Zfp275 | 27081 [Gene Symbol: Zfp275] [Locus Tag: ] [Chromosome: X] [Map Location: X A7.3\|X 3 | 1.592061663 | 0.009381 | 0.019662 |
| 16221 | ENSMUSG00000051934 | Spats2 | 72572 [Gene Symbol: Spats2] [Locus Tag: ] [Chromosome: 15] [Map Location: 15\|15 F1 | 1.590825101 | 0.004252 | 0.009625 |
| 13950 | ENSMUSG00000028790 | Khdrbs1 | 20218 [Gene Symbol: Khdrbs1] [Locus Tag: ] [Chromosome: 4] [Map Location: 4\|4 D2.2 | 1.588333617 | 0.004531 | 0.010189 |
| 14192 | ENSMUSG00000060181 | Slc35e3 | 215436 [Gene Symbol: Slc35e3] [Locus Tag: ] [Chromosome: 10] [Map Location: 10\|10 | 1.587560024 | 0.02334 | 0.044198 |
| 14356 | ENSMUSG00000022811 | Zfp148 | 22661 [Gene Symbol: Zfp148] [Locus Tag: ] [Chromosome: 16] [Map Location: 16 B3\|16 | 1.58723511 | 0.003093 | 0.007231 |
| 1591 | ENSMUSG00000041895 | Wipi1 | 52639 [Gene Symbol: Wipi1] [Locus Tag: ] [Chromosome: 11] [Map Location: 11 E1\|11 | 1.586344413 | 0.019624 | 0.037876 |
| 5985 | ENSMUSG00000038965 | Gm10705; Ube2l3 | 100042355 [Gene Symbol: Gm10705] [Locus Tag: ] [Chromosome: 3] [Map Location: 3 | 1.586065719 | 0.01216 | 0.024792 |
| 13675 | ENSMUSG00000035456 | Prdm8 | 77630 [Gene Symbol: Prdm8] [Locus Tag: ] [Chromosome: 5] [Map Location: 5\|5 E3] [D | 1.585354699 | 0.004829 | 0.010795 |
| 5318 | ENSMUSG00000021177 | Tdp1 | 104884 [Gene Symbol: Tdp1] [Locus Tag: ] [Chromosome: 12] [Map Location: 12\|12 E] | 1.585023592 | 0.01552 | 0.030804 |
| 13020 | ENSMUSG00000037824 | Tspan14 | 52588 [Gene Symbol: Tspan14] [Locus Tag: ] [Chromosome: 14] [Map Location: 14 B\|14 | 1.584521115 | 0.009007 | 0.01895 |
| 9394 | ENSMUSG00000024598 | Fbn2; LOC10264229 | 14119 [Gene Symbol: Fbn2] [Locus Tag: ] [Chromosome: 18] [Map Location: 18 32.15 c | 1.583570994 | 0.010077 | 0.020973 |
| 11038 | ENSMUSG00000031889 | D230025D16Rik | 234678 [Gene Symbol: D230025D16Rik] [Locus Tag: ] [Chromosome: 8] [Map Location | 1.582159967 | 0.018381 | 0.035775 |
| 3092 | ENSMUSG00000076432 | Ywhaq; Cdk5rap2 | 22630 [Gene Symbol: Ywhaq] [Locus Tag: ] [Chromosome: 12] [Map Location: 12\|12 A1 | 1.581750504 | 0.011822 | 0.024202 |
| 16355 | ENSMUSG00000035245 | Eogt | 101351 [Gene Symbol: Eogt] [Locus Tag: ] [Chromosome: 6] [Map Location: 6\|6 D3] [De | 1.581748726 | 0.016455 | 0.032433 |
| 1708 | ENSMUSG00000057181 | 5730455P16Rik | 70591 [Gene Symbol: 5730455P16Rik] [Locus Tag: ] [Chromosome: 11] [Map Location: | 1.581580601 | 0.008112 | 0.017242 |
| 14525 | ENSMUSG00000030876 | Mettl9 | 59052 [Gene Symbol: Mettl9] [Locus Tag: ] [Chromosome: 7] [Map Location: 7\|7 F2] [D | 1.580989806 | 0.012994 | 0.026274 |
| 472 | ENSMUSG00000075033 | Nxpe3 | 385658 [Gene Symbol: Nxpe3] [Locus Tag: ] [Chromosome: 16] [Map Location: 16\|16 C | 1.58027894 | 0.014966 | 0.02982 |
| 13613 | ENSMUSG00000037341 | Slc9a7 | 236727 [Gene Symbol: Slc9a7] [Locus Tag: ] [Chromosome: X] [Map Location: X\|X A1.3 | 1.580181925 | 0.025706 | 0.04825 |
| 12603 | ENSMUSG00000029250 | Polr2b | 231329 [Gene Symbol: Polr2b] [Locus Tag: ] [Chromosome: 5] [Map Location: 5\|5 C3.3 | 1.580172741 | 0.008017 | 0.017066 |
| 1224 | ENSMUSG00000041313 | Slc7a1 | 11987 [Gene Symbol: Slc7a1] [Locus Tag: ] [Chromosome: 5] [Map Location: 5 G3\|5 88. | 1.579711413 | 0.007261 | 0.015602 |
| 5482 | ENSMUSG00000027523 | Gnas | 14683 [Gene Symbol: Gnas] [Locus Tag: ] [Chromosome: 2] [Map Location: 2 97.89 cM\| | 1.579688306 | 0.002449 | 0.005858 |
| 6061 | ENSMUSG00000028414 | Fktn | 246179 [Gene Symbol: Fktn] [Locus Tag: ] [Chromosome: 4] [Map Location: 4 B2\|4 28.7 | 1.579427545 | 0.008638 | 0.018252 |
| 3025 | ENSMUSG00000037747 | Phyhipl | 70911 [Gene Symbol: Phyhipl] [Locus Tag: ] [Chromosome: 10] [Map Location: 10\|10 B | 1.579275022 | 0.011426 | 0.023474 |
| 12548 | ENSMUSG00000048920 | Fkrp | 243853 [Gene Symbol: Fkrp] [Locus Tag: ] [Chromosome: 7] [Map Location: 7\|7 A2] [De | 1.578126437 | 0.004823 | 0.010781 |
| 8958 | ENSMUSG00000023915 | Tnfrsf21 | 94185 [Gene Symbol: Tnfrsf21] [Locus Tag: ] [Chromosome: 17] [Map Location: 17\|17 C | 1.576297385 | 0.004963 | 0.011071 |
| 5237 | ENSMUSG00000034903 | Cobll1 | 319876 [Gene Symbol: Cobll1] [Locus Tag: ] [Chromosome: 2] [Map Location: 2\|2 C3] [ | 1.576240981 | 0.016354 | 0.032258 |
| 5264 | ENSMUSG00000036053 | Fmnl2 | 71409 [Gene Symbol: Fmnl2] [Locus Tag: ] [Chromosome: 2] [Map Location: 2\|2 C1.1] [ | 1.576001115 | 0.004518 | 0.010163 |
| 13483 | ENSMUSG00000043702 | Pde12 | 211948 [Gene Symbol: Pde12] [Locus Tag: ] [Chromosome: 14] [Map Location: 14\|14 A | 1.575678247 | 0.019108 | 0.037 |
| 6838 | ENSMUSG00000051910 | Sox6 | 20679 [Gene Symbol: Sox6] [Locus Tag: ] [Chromosome: 7] [Map Location: 7 F1\|7 61.2 | 1.575672897 | 0.014635 | 0.029252 |
| 16392 | ENSMUSG00000055239 | Kcmf1 | 74287 [Gene Symbol: Kcmf1] [Locus Tag: ] [Chromosome: 6] [Map Location: 6 32.3 cM\| | 1.57566451 | 0.004272 | 0.009668 |
| 2367 | ENSMUSG00000036641 | Ccdc148 | 227933 [Gene Symbol: Ccdc148] [Locus Tag: ] [Chromosome: 2] [Map Location: 2\|2 C1 | 1.575376708 | 0.022891 | 0.043449 |
| 16648 | ENSMUSG00000025255 | Zfhx4 | 80892 [Gene Symbol: Zfhx4] [Locus Tag: ] [Chromosome: 3] [Map Location: 3\|3 A1] [De | 1.574895529 | 0.005913 | 0.012953 |
| 4264 | ENSMUSG00000035828 | Pim3 | 223775 [Gene Symbol: Pim3] [Locus Tag: ] [Chromosome: 15] [Map Location: 15\|15 E3 | 1.574691881 | 0.017998 | 0.035084 |
| 18208 | ENSMUSG00000049907 | Rasl11b | 68939 [Gene Symbol: Rasl11b] [Locus Tag: ] [Chromosome: 5] [Map Location: 5\|5 D] [D | 1.574619752 | 0.023786 | 0.044986 |
| 983 | ENSMUSG00000049300 | Prmt6 | 99890 [Gene Symbol: Prmt6] [Locus Tag: ] [Chromosome: 3] [Map Location: 3\|3 F3] [De | 1.573728458 | 0.018835 | 0.036543 |
| 14649 | ENSMUSG00000054920 | Klhl5 | 71778 [Gene Symbol: Klhl5] [Locus Tag: ] [Chromosome: 5] [Map Location: 5\|5 C3.1] [D | 1.573273652 | 0.003373 | 0.007808 |
| 13618 | ENSMUSG00000018425 | Dhx40 | 67487 [Gene Symbol: Dhx40] [Locus Tag: ] [Chromosome: 11] [Map Location: 11\|11 C] | 1.571713063 | 0.024012 | 0.045378 |
| 17165 | ENSMUSG00000068959 | Zfp619 | 70227 [Gene Symbol: Zfp619] [Locus Tag: ] [Chromosome: 7] [Map Location: 7\|7 B3] [D | 1.571702876 | 0.014755 | 0.029469 |
| 1784 | ENSMUSG00000074785 | Plxnc1 | 54712 [Gene Symbol: Plxnc1] [Locus Tag: ] [Chromosome: 10] [Map Location: 10\|10 C | 1.570809091 | 0.014662 | 0.029298 |
| 4612 | ENSMUSG00000001419 | Mef2d | 17261 [Gene Symbol: Mef2d] [Locus Tag: ] [Chromosome: 3] [Map Location: 3 F1\|3 38. | 1.570427017 | 0.004331 | 0.00979 |
| 15134 | ENSMUSG00000022641 | Bbx | 70508 [Gene Symbol: Bbx] [Locus Tag: ] [Chromosome: 16] [Map Location: 16\|16 B5] [ | 1.570264912 | 0.01869 | 0.036304 |
| 10809 | ENSMUSG00000070565 | Rasal2 | 226525 [Gene Symbol: Rasal2] [Locus Tag: ] [Chromosome: 1] [Map Location: 1\|1 H1] [ | 1.570022359 | 0.006263 | 0.013637 |
| 18400 | ENSMUSG00000009733 | Tfcp2 | 21422 [Gene Symbol: Tfcp2] [Locus Tag: ] [Chromosome: 15] [Map Location: 15 F1\|15 | 1.569405769 | 0.012048 | 0.0246 |
| 15641 | ENSMUSG00000045613 | Chrm2 | 243764 [Gene Symbol: Chrm2] [Locus Tag: ] [Chromosome: 6] [Map Location: 6\|6 B1] [ | 1.568792607 | 0.024989 | 0.047038 |

| | | | | | | |
|---|---|---|---|---|---|---|
| 3994 | ENSMUSG00000048874 | Phf3 | 213109 [Gene Symbol: Phf3] [Locus Tag: ] [Chromosome: 1] [Map Location: 1\|1 A5] [De | 1.568140363 | 0.003458 | 0.007983 |
| 15906 | ENSMUSG00000020739 | Nup85 | 445007 [Gene Symbol: Nup85] [Locus Tag: ] [Chromosome: 11] [Map Location: 11\|11 E | 1.567924629 | 0.017938 | 0.034971 |
| 9905 | ENSMUSG00000040447 | Spns2 | 216892 [Gene Symbol: Spns2] [Locus Tag: ] [Chromosome: 11] [Map Location: 11\|11 B | 1.567816422 | 0.008647 | 0.018267 |
| 2975 | ENSMUSG00000035864 | Syt1 | 20979 [Gene Symbol: Syt1] [Locus Tag: ] [Chromosome: 10] [Map Location: 10 D1\|10 5 | 1.567750099 | 0.004853 | 0.010844 |
| 11349 | ENSMUSG00000030505 | Prmt3 | 71974 [Gene Symbol: Prmt3] [Locus Tag: ] [Chromosome: 7] [Map Location: 7\|7 B5] [De | 1.564448156 | 0.014781 | 0.029513 |
| 2459 | ENSMUSG00000068036 | Mllt4 | 17356 [Gene Symbol: Mllt4] [Locus Tag: ] [Chromosome: 17] [Map Location: 17\|17 A2] | 1.56403516 | 0.004029 | 0.009168 |
| 4273 | ENSMUSG00000062590 | Armc9 | 78795 [Gene Symbol: Armc9] [Locus Tag: ] [Chromosome: 1] [Map Location: 1\|1 C5] [D | 1.562460989 | 0.011472 | 0.023553 |
| 2377 | ENSMUSG00000025439 | Clns1a | 12729 [Gene Symbol: Clns1a] [Locus Tag: ] [Chromosome: 7] [Map Location: 7 53.57 cM | 1.562436132 | 0.024067 | 0.045467 |
| 17328 | ENSMUSG00000030313 | Dennd5b | 320560 [Gene Symbol: Dennd5b] [Locus Tag: ] [Chromosome: 6] [Map Location: 6\|6 G3 | 1.561578041 | 0.005456 | 0.012049 |
| 9399 | ENSMUSG00000022757 | Tfg | 21787 [Gene Symbol: Tfg] [Locus Tag: ] [Chromosome: 16] [Map Location: 16\|16 C1.1] | 1.560188145 | 0.011696 | 0.023966 |
| 7683 | ENSMUSG00000000532 | Acvr1b | 11479 [Gene Symbol: Acvr1b] [Locus Tag: ] [Chromosome: 15] [Map Location: 15 F2\|15 | 1.55981979 | 0.005096 | 0.011329 |
| 6469 | ENSMUSG00000048347 | Pcdhb18 | 93889 [Gene Symbol: Pcdhb18] [Locus Tag: ] [Chromosome: 18] [Map Location: 18\|18 | 1.557483485 | 0.014583 | 0.029163 |
| 4609 | ENSMUSG00000026510 | Trp53bp2 | 209456 [Gene Symbol: Trp53bp2] [Locus Tag: ] [Chromosome: 1] [Map Location: 1 H5\| | 1.556860905 | 0.008942 | 0.01883 |
| 16489 | ENSMUSG00000024240 | Epc1 | 13831 [Gene Symbol: Epc1] [Locus Tag: ] [Chromosome: 18] [Map Location: 18\|18 A1] | 1.556849093 | 0.00577 | 0.012673 |
| 11233 | ENSMUSG00000026021 | Sumo1 | 22218 [Gene Symbol: Sumo1] [Locus Tag: ] [Chromosome: 1] [Map Location: 1\|1 C2-C3 | 1.556526336 | 0.020564 | 0.039502 |
| 14747 | ENSMUSG00000031776 | Arl2bp | 107566 [Gene Symbol: Arl2bp] [Locus Tag: ] [Chromosome: 8] [Map Location: 8\|8 C5] | 1.556407444 | 0.005511 | 0.012156 |
| 12330 | ENSMUSG00000035517 | Tdrd7 | 100121 [Gene Symbol: Tdrd7] [Locus Tag: ] [Chromosome: 4] [Map Location: 4\|4 B1] [D | 1.556000821 | 0.008777 | 0.018521 |
| 15369 | ENSMUSG00000031189 | Aff2 | 14266 [Gene Symbol: Aff2] [Locus Tag: ] [Chromosome: X] [Map Location: X\|X A5] [Desc | 1.553161688 | 0.008571 | 0.018123 |
| 13556 | ENSMUSG00000029048 | Rer1 | 67830 [Gene Symbol: Rer1] [Locus Tag: ] [Chromosome: 4] [Map Location: 4\|4 E2] [Des | 1.552273464 | 0.016731 | 0.03291 |
| 3149 | ENSMUSG00000058056 | Palld | 72333 [Gene Symbol: Palld] [Locus Tag: ] [Chromosome: 8] [Map Location: 8\|8 B3.3] [D | 1.551824795 | 0.022589 | 0.04294 |
| 5082 | ENSMUSG00000026037 | Orc2 | 18393 [Gene Symbol: Orc2] [Locus Tag: ] [Chromosome: 1] [Map Location: 1\|1 C1.3] [D | 1.550394575 | 0.017758 | 0.034674 |
| 14628 | ENSMUSG00000026353 | Ubxn4 | 67812 [Gene Symbol: Ubxn4] [Locus Tag: ] [Chromosome: 1] [Map Location: 1\|1 E4] [De | 1.548821553 | 0.008951 | 0.018848 |
| 17889 | ENSMUSG00000036353 | P2ry12 | 70839 [Gene Symbol: P2ry12] [Locus Tag: ] [Chromosome: 3] [Map Location: 3\|3 D] [De | 1.547987 | 0.012208 | 0.024873 |
| 4 | ENSMUSG00000035189 | Ano4 | 320091 [Gene Symbol: Ano4] [Locus Tag: ] [Chromosome: 10] [Map Location: 10\|10 C1 | 1.547428431 | 0.01263 | 0.02563 |
| 1980 | ENSMUSG00000090213 | Tmem189 | 407243 [Gene Symbol: Tmem189] [Locus Tag: ] [Chromosome: 2] [Map Location: 2\|2 H | 1.547130713 | 0.013819 | 0.02777 |
| 18206 | ENSMUSG00000020257 | Wdr82 | 77305 [Gene Symbol: Wdr82] [Locus Tag: ] [Chromosome: 9] [Map Location: 9\|9 F1] [D | 1.546972844 | 0.007238 | 0.015559 |
| 1885 | ENSMUSG00000027993 | Trim2 | 80890 [Gene Symbol: Trim2] [Locus Tag: ] [Chromosome: 3] [Map Location: 3\|3 F1] [De | 1.543886658 | 0.006043 | 0.01321 |
| 14385 | ENSMUSG00000026270 | Capn10 | 23830 [Gene Symbol: Capn10] [Locus Tag: ] [Chromosome: 1] [Map Location: 1\|1 D] [D | 1.54228314 | 0.011712 | 0.023993 |
| 9327 | ENSMUSG00000059742 | Kcnh7 | 170738 [Gene Symbol: Kcnh7] [Locus Tag: ] [Chromosome: 2] [Map Location: 2\|2 C1.3] | 1.542147753 | 0.010066 | 0.020956 |
| 13242 | ENSMUSG00000021893 | Capn7 | 12339 [Gene Symbol: Capn7] [Locus Tag: ] [Chromosome: 14] [Map Location: 14\|14 B] | 1.541826062 | 0.020725 | 0.039784 |
| 17744 | ENSMUSG00000053641 | Dennd4a | 102442 [Gene Symbol: Dennd4a] [Locus Tag: ] [Chromosome: 9] [Map Location: 9\|9 C] | 1.54134379 | 0.006943 | 0.01497 |
| 10122 | ENSMUSG00000020354 | Sgcd | 24052 [Gene Symbol: Sgcd] [Locus Tag: ] [Chromosome: 11] [Map Location: 11\|11 B1.2 | 1.540775348 | 0.006707 | 0.014499 |
| 89 | ENSMUSG00000032350 | Gclc | 14629 [Gene Symbol: Gclc] [Locus Tag: ] [Chromosome: 9] [Map Location: 9 43.36 cM\| | 1.540373572 | 0.009677 | 0.020225 |
| 5428 | ENSMUSG00000055923 | Aasdh | 231326 [Gene Symbol: Aasdh] [Locus Tag: ] [Chromosome: 5] [Map Location: 5\|5 C3.3] | 1.54032685 | 0.019 | 0.03681 |
| 5664 | ENSMUSG00000000197 | Nalcn | 338370 [Gene Symbol: Nalcn] [Locus Tag: ] [Chromosome: 14] [Map Location: 14\|14 E | 1.540106806 | 0.01002 | 0.020875 |
| 17396 | ENSMUSG00000034602 | Mon2 | 67074 [Gene Symbol: Mon2] [Locus Tag: ] [Chromosome: 10] [Map Location: 10\|10 D2 | 1.540034785 | 0.009295 | 0.019495 |
| 1034 | ENSMUSG00000046709 | Mapk10 | 26414 [Gene Symbol: Mapk10] [Locus Tag: ] [Chromosome: 5] [Map Location: 5\|5 E5] [ | 1.539751296 | 0.006503 | 0.014117 |
| 7595 | ENSMUSG00000041911 | Dlx1 | 13390 [Gene Symbol: Dlx1] [Locus Tag: ] [Chromosome: 2] [Map Location: 2 C2\|2 42.6 | 1.539363848 | 0.018384 | 0.035777 |
| 2103 | ENSMUSG00000031774 | Fam192a | 102122 [Gene Symbol: Fam192a] [Locus Tag: ] [Chromosome: 8] [Map Location: 8\|8 C5 | 1.538367899 | 0.019569 | 0.037789 |
| 4732 | ENSMUSG00000022451 | Gm4887; Twf1 | 233637 [Gene Symbol: Gm4887] [Locus Tag: ] [Chromosome: 7] [Map Location: 7\|7 E3 | 1.537194517 | 0.016194 | 0.031974 |
| 1912 | ENSMUSG00000032939 | Nup93 | 71805 [Gene Symbol: Nup93] [Locus Tag: ] [Chromosome: 8] [Map Location: 8\|8 C5] [D | 1.537107798 | 0.023423 | 0.044335 |
| 12142 | ENSMUSG00000025795 | Rassf3 | 192678 [Gene Symbol: Rassf3] [Locus Tag: ] [Chromosome: 10] [Map Location: 10\|10 D | 1.535618034 | 0.025959 | 0.048681 |
| 4179 | ENSMUSG00000048833 | Slc39a9 | 328133 [Gene Symbol: Slc39a9] [Locus Tag: ] [Chromosome: 12] [Map Location: 12\|12 | 1.535492178 | 0.017294 | 0.033858 |
| 11803 | ENSMUSG00000059921 | Unc5c | 22253 [Gene Symbol: Unc5c] [Locus Tag: ] [Chromosome: 3] [Map Location: 3 H1\|3 65. | 1.53462766 | 0.013678 | 0.027515 |
| 6546 | ENSMUSG00000032178 | Ilf3 | 16201 [Gene Symbol: Ilf3] [Locus Tag: ] [Chromosome: 9] [Map Location: 9 7.78 cM\|9 A | 1.533273335 | 0.008867 | 0.018696 |
| 13019 | ENSMUSG00000038702 | Dsel | 319901 [Gene Symbol: Dsel] [Locus Tag: ] [Chromosome: 1] [Map Location: 1\|1 E2.2] [D | 1.532725491 | 0.015863 | 0.031411 |
| 8732 | ENSMUSG00000020333 | Acsl6 | 216739 [Gene Symbol: Acsl6] [Locus Tag: ] [Chromosome: 11] [Map Location: 11 B1.3\| | 1.532515504 | 0.01013 | 0.021076 |
| 16813 | ENSMUSG00000021010 | Npas3 | 27386 [Gene Symbol: Npas3] [Locus Tag: ] [Chromosome: 12] [Map Location: 12\|12 C1 | 1.531437077 | 0.006279 | 0.013669 |
| 2288 | ENSMUSG00000050730 | Arhgap42 | 71544 [Gene Symbol: Arhgap42] [Locus Tag: ] [Chromosome: 9] [Map Location: 9\|9 A1] | 1.530801194 | 0.015023 | 0.029908 |
| 5148 | ENSMUSG00000050890 | Pdik1l | 230809 [Gene Symbol: Pdik1l] [Locus Tag: ] [Chromosome: 4] [Map Location: 4\|4 D3] [ | 1.530300465 | 0.015969 | 0.031591 |
| 14637 | ENSMUSG00000047261 | Gap43 | 14432 [Gene Symbol: Gap43] [Locus Tag: ] [Chromosome: 16] [Map Location: 16 B4\|16 | 1.530052841 | 0.010362 | 0.021498 |
| 3190 | ENSMUSG00000046546 | Fam43a | 224093 [Gene Symbol: Fam43a] [Locus Tag: ] [Chromosome: 16] [Map Location: 16\|16 | 1.528103797 | 0.019591 | 0.03782 |
| 11923 | ENSMUSG00000034620 | Tmem5 | 216395 [Gene Symbol: Tmem5] [Locus Tag: ] [Chromosome: 10] [Map Location: 10\|10 | 1.527492911 | 0.02673 | 0.049966 |
| 11801 | ENSMUSG00000028639 | Ybx1 | 22608 [Gene Symbol: Ybx1] [Locus Tag: ] [Chromosome: 4] [Map Location: 4\|4 D1] [Des | 1.526867712 | 0.025484 | 0.047891 |
| 746 | ENSMUSG00000004319 | Clcn3 | 12725 [Gene Symbol: Clcn3] [Locus Tag: ] [Chromosome: 8] [Map Location: 8 B3.1\|8 30 | 1.523473193 | 0.008806 | 0.018576 |
| 12472 | ENSMUSG00000038467 | Chmp4b | 75608 [Gene Symbol: Chmp4b] [Locus Tag: ] [Chromosome: 2] [Map Location: 2\|2 H2] | 1.521678203 | 0.012416 | 0.025256 |
| 14646 | ENSMUSG00000042757 | Tmem108 | 81907 [Gene Symbol: Tmem108] [Locus Tag: ] [Chromosome: 9] [Map Location: 9\|9 F1 | 1.520651184 | 0.012775 | 0.025885 |
| 10341 | ENSMUSG00000024082 | Ndufaf7 | 73694 [Gene Symbol: Ndufaf7] [Locus Tag: ] [Chromosome: 17] [Map Location: 17\|17 E | 1.520454874 | 0.024645 | 0.046474 |
| 3498 | ENSMUSG00000024360 | Etf1 | 225363 [Gene Symbol: Etf1] [Locus Tag: ] [Chromosome: 18] [Map Location: 18\|18 B1] | 1.520273248 | 0.016218 | 0.032016 |
| 16271 | ENSMUSG00000040746 | Rnf167 | 70510 [Gene Symbol: Rnf167] [Locus Tag: ] [Chromosome: 11] [Map Location: 11\|11 B | 1.520236408 | 0.02324 | 0.044025 |
| 9520 | ENSMUSG00000042444 | Fam63b | 235461 [Gene Symbol: Fam63b] [Locus Tag: ] [Chromosome: 9] [Map Location: 9\|9 D] [ | 1.519461266 | 0.007676 | 0.016406 |
| 15210 | ENSMUSG00000021114 | Atp6v1d | 73834 [Gene Symbol: Atp6v1d] [Locus Tag: ] [Chromosome: 12] [Map Location: 12 C3\| | 1.519005799 | 0.018364 | 0.035745 |
| 1241 | ENSMUSG00000037353 | Letmd1 | 68614 [Gene Symbol: Letmd1] [Locus Tag: ] [Chromosome: 15] [Map Location: 15\|15 F | 1.518668727 | 0.021319 | 0.040784 |
| 11161 | ENSMUSG00000024713 | Pcsk5 | 18552 [Gene Symbol: Pcsk5] [Locus Tag: ] [Chromosome: 19] [Map Location: 19 B\|19 1 | 1.517524643 | 0.017161 | 0.033625 |
| 11460 | ENSMUSG00000036698 | Ago2 | 239528 [Gene Symbol: Ago2] [Locus Tag: ] [Chromosome: 15] [Map Location: 15\|15 D3 | 1.517390597 | 0.01706 | 0.033445 |
| 2160 | ENSMUSG00000031799 | Tpm4 | 326618 [Gene Symbol: Tpm4] [Locus Tag: ] [Chromosome: 8] [Map Location: 8\|8 B3.3] | 1.517188884 | 0.022328 | 0.042492 |
| 12316 | ENSMUSG00000068917 | Clk2 | 12748 [Gene Symbol: Clk2] [Locus Tag: ] [Chromosome: 3] [Map Location: 3\|3 F1] [Desc | 1.516586652 | 0.021139 | 0.040482 |
| 11770 | ENSMUSG00000046062 | Ppp1r15b | 108954 [Gene Symbol: Ppp1r15b] [Locus Tag: ] [Chromosome: 1] [Map Location: 1\|1 E | 1.516080846 | 0.011797 | 0.024158 |
| 5399 | ENSMUSG00000041229 | Phf8 | 320595 [Gene Symbol: Phf8] [Locus Tag: ] [Chromosome: X] [Map Location: X\|X F3] [De | 1.513771497 | 0.019119 | 0.037018 |
| 8218 | ENSMUSG00000067336 | Bmpr2 | 12168 [Gene Symbol: Bmpr2] [Locus Tag: ] [Chromosome: 1] [Map Location: 1\|1 C1.3] | 1.512325186 | 0.009618 | 0.020114 |
| 5474 | ENSMUSG00000026764 | Kif5c | 16574 [Gene Symbol: Kif5c] [Locus Tag: ] [Chromosome: 2] [Map Location: 2 C\|2 28.68 | 1.512114134 | 0.00807 | 0.017172 |
| 9712 | ENSMUSG00000049800 | Sertad2 | 58172 [Gene Symbol: Sertad2] [Locus Tag: ] [Chromosome: 11] [Map Location: 11\|11 A | 1.512013496 | 0.019286 | 0.037302 |
| 13523 | ENSMUSG00000032115 | Hyou1 | 12282 [Gene Symbol: Hyou1] [Locus Tag: ] [Chromosome: 9] [Map Location: 9\|9 B] [Des | 1.511687049 | 0.019699 | 0.03801 |
| 1895 | ENSMUSG00000020393 | Kremen1 | 84035 [Gene Symbol: Kremen1] [Locus Tag: ] [Chromosome: 11] [Map Location: 11\|11 | 1.511393218 | 0.022779 | 0.043262 |
| 16583 | ENSMUSG00000001016 | Ilf2 | 67781 [Gene Symbol: Ilf2] [Locus Tag: ] [Chromosome: 3] [Map Location: 3\|3 F2] [Descr | 1.511390106 | 0.022283 | 0.042423 |
| 16524 | ENSMUSG00000039585 | Myo9a | 270163 [Gene Symbol: Myo9a] [Locus Tag: ] [Chromosome: 9] [Map Location: 9 B\|9 32. | 1.51036322 | 0.007403 | 0.015885 |
| 3379 | ENSMUSG00000052656 | Rnf103 | 22644 [Gene Symbol: Rnf103] [Locus Tag: ] [Chromosome: 6] [Map Location: 6 C1\|6 32 | 1.510260033 | 0.018541 | 0.036044 |
| 9319 | ENSMUSG00000048732 | Klhl11 | 217194 [Gene Symbol: Klhl11] [Locus Tag: ] [Chromosome: 11] [Map Location: 11\|11 D | 1.509303694 | 0.014607 | 0.029198 |
| 7619 | ENSMUSG00000030022 | Adamts9 | 101401 [Gene Symbol: Adamts9] [Locus Tag: ] [Chromosome: 6] [Map Location: 6\|6 D1 | 1.508430698 | 0.026075 | 0.048864 |
| 13423 | ENSMUSG00000024400 | Wdr33 | 74320 [Gene Symbol: Wdr33] [Locus Tag: ] [Chromosome: 18] [Map Location: 18\|18 B | 1.507935036 | 0.012605 | 0.025586 |
| 17220 | ENSMUSG00000024501 | Dpysl3 | 22240 [Gene Symbol: Dpysl3] [Locus Tag: ] [Chromosome: 18] [Map Location: 18\|18 B | 1.507705758 | 0.008102 | 0.017227 |
| 16779 | ENSMUSG00000028329 | Xpa | 22590 [Gene Symbol: Xpa] [Locus Tag: ] [Chromosome: 4] [Map Location: 4 24.49 cM\|4 | 1.507596884 | 0.020752 | 0.039826 |
| 16476 | ENSMUSG00000052713 | Zfp608 | 269023 [Gene Symbol: Zfp608] [Locus Tag: ] [Chromosome: 18] [Map Location: 18\|18 | 1.507164568 | 0.013314 | 0.026852 |
| 17812 | ENSMUSG00000034636 | Zyg11b | 414872 [Gene Symbol: Zyg11b] [Locus Tag: ] [Chromosome: 4] [Map Location: 4\|4 C7] | 1.505796214 | 0.009761 | 0.020379 |

| | | | | | | |
|---|---|---|---|---|---|---|
| 1055 | ENSMUSG00000041444 | Arhgap32 | 330914 [Gene Symbol: Arhgap32] [Locus Tag: ] [Chromosome: 9] [Map Location: 9\|9 A4 | 1.505478548 | 0.009403 | 0.019703 |
| 16140 | ENSMUSG00000020142 | Slc1a4 | 55963 [Gene Symbol: Slc1a4] [Locus Tag: ] [Chromosome: 11] [Map Location: 11 A3.1\| | 1.504742606 | 0.017394 | 0.034036 |
| 18095 | ENSMUSG00000022964 | Tmem50b | 77975 [Gene Symbol: Tmem50b] [Locus Tag: ] [Chromosome: 16] [Map Location: 16\|16 | 1.503098844 | 0.025587 | 0.048051 |
| 12892 | ENSMUSG00000000127 | Fer | 14158 [Gene Symbol: Fer] [Locus Tag: ] [Chromosome: 17] [Map Location: 17\|17 E1.1]\| | 1.501347122 | 0.021837 | 0.041673 |
| 2682 | ENSMUSG00000027221 | Chst1 | 76969 [Gene Symbol: Chst1] [Locus Tag: ] [Chromosome: 2] [Map Location: 2\|2 D-E1] [D | 1.501133953 | 0.009969 | 0.02078 |
| 5094 | ENSMUSG00000041846 | Smek1 | 68734 [Gene Symbol: Smek1] [Locus Tag: ] [Chromosome: 12] [Map Location: 12\|12 E] | 1.500831368 | 0.022704 | 0.043132 |
| 7212 | ENSMUSG00000010517 | Faf1 | 14084 [Gene Symbol: Faf1] [Locus Tag: ] [Chromosome: 4] [Map Location: 4 C6\|4 51.33 | 1.499666163 | 0.01603 | 0.031701 |
| 2819 | ENSMUSG00000020128 | Vps54 | 245944 [Gene Symbol: Vps54] [Locus Tag: ] [Chromosome: 11] [Map Location: 11 A3\|1 | 1.498131457 | 0.009074 | 0.019075 |
| 16304 | ENSMUSG00000031976 | Urb2 | 382038 [Gene Symbol: Urb2] [Locus Tag: ] [Chromosome: 8] [Map Location: 8\|8 E2] [De | 1.495275105 | 0.022974 | 0.04357 |
| 14112 | ENSMUSG00000039375 | Wdr17 | 244484 [Gene Symbol: Wdr17] [Locus Tag: ] [Chromosome: 8] [Map Location: 8\|8 B3.1 | 1.495234924 | 0.019305 | 0.03733 |
| 1858 | ENSMUSG00000020271 | Fbxw11 | 103583 [Gene Symbol: Fbxw11] [Locus Tag: ] [Chromosome: 11] [Map Location: 11\|11 | 1.495152519 | 0.009428 | 0.019751 |
| 7852 | ENSMUSG00000038252 | Ncapd2 | 68298 [Gene Symbol: Ncapd2] [Locus Tag: ] [Chromosome: 6] [Map Location: 6\|6 F2] [D | 1.49484952 | 0.025371 | 0.047703 |
| 11261 | ENSMUSG00000032481 | Smarcc1 | 20588 [Gene Symbol: Smarcc1] [Locus Tag: ] [Chromosome: 9] [Map Location: 9\|9 F2] [ | 1.494776727 | 0.013592 | 0.027361 |
| 5512 | ENSMUSG00000020598 | Nrcam | 319504 [Gene Symbol: Nrcam] [Locus Tag: ] [Chromosome: 12] [Map Location: 12 B3\|1 | 1.494334678 | 0.010553 | 0.02185 |
| 13149 | ENSMUSG00000032279 | Idh3a | 67834 [Gene Symbol: Idh3a] [Locus Tag: ] [Chromosome: 9] [Map Location: 9\|9 A5.3] [D | 1.493683025 | 0.024556 | 0.046321 |
| 15810 | ENSMUSG00000045962 | Wnk1 | 232341 [Gene Symbol: Wnk1] [Locus Tag: ] [Chromosome: 6] [Map Location: 6\|6 F1] [D | 1.493018254 | 0.015965 | 0.031588 |
| 10336 | ENSMUSG00000036615 | Rfxap | 170767 [Gene Symbol: Rfxap] [Locus Tag: ] [Chromosome: 3] [Map Location: 3\|3 D] [De | 1.492639985 | 0.022653 | 0.043045 |
| 11811 | ENSMUSG00000040389 | Wdr47 | 99512 [Gene Symbol: Wdr47] [Locus Tag: ] [Chromosome: 3] [Map Location: 3\|3 F3] [D | 1.490628505 | 0.010275 | 0.021341 |
| 15800 | ENSMUSG00000039607 | Rbms3 | 207181 [Gene Symbol: Rbms3] [Locus Tag: ] [Chromosome: 9] [Map Location: 9\|9 F3] [I | 1.490628308 | 0.011578 | 0.023746 |
| 6443 | ENSMUSG00000029763 | Exoc4 | 20336 [Gene Symbol: Exoc4] [Locus Tag: ] [Chromosome: 6] [Map Location: 6 A3.3\|6 14 | 1.490390751 | 0.015192 | 0.030206 |
| 9393 | ENSMUSG00000021027 | Ralgapa1 | 56784 [Gene Symbol: Ralgapa1] [Locus Tag: ] [Chromosome: 12] [Map Location: 12\|12 | 1.488528634 | 0.010136 | 0.021082 |
| 3674 | ENSMUSG00000049354 | Dcaf7 | 71833 [Gene Symbol: Dcaf7] [Locus Tag: ] [Chromosome: 11] [Map Location: 11\|11 E1] | 1.486924656 | 0.017418 | 0.034079 |
| 11338 | ENSMUSG00000037253 | Mex3c | 240396 [Gene Symbol: Mex3c] [Locus Tag: ] [Chromosome: 18] [Map Location: 18\|18 E | 1.486607539 | 0.016015 | 0.031677 |
| 14848 | ENSMUSG00000041362 | Shtn1 | 71653 [Gene Symbol: Shtn1] [Locus Tag: ] [Chromosome: 19] [Map Location: 19\|19 D3 | 1.484615371 | 0.009588 | 0.020064 |
| 9991 | ENSMUSG00000038665 | Dgki | 320127 [Gene Symbol: Dgki] [Locus Tag: ] [Chromosome: 6] [Map Location: 6\|6 B1] [De | 1.48373779 | 0.012147 | 0.024769 |
| 15961 | ENSMUSG00000027520 | Zdbf2 | 73884 [Gene Symbol: Zdbf2] [Locus Tag: ] [Chromosome: 1] [Map Location: 1\|1 C2] [De | 1.48371302 | 0.014845 | 0.029616 |
| 4189 | ENSMUSG00000023927 | Satb1 | 20230 [Gene Symbol: Satb1] [Locus Tag: ] [Chromosome: 17] [Map Location: 17\|17 C] [ | 1.483417977 | 0.009732 | 0.020322 |
| 6292 | ENSMUSG00000025612 | Bach1 | 12013 [Gene Symbol: Bach1] [Locus Tag: ] [Chromosome: 16] [Map Location: 16\|16 C3 | 1.483406638 | 0.015889 | 0.031456 |
| 10067 | ENSMUSG00000038485 | Socs7 | 192157 [Gene Symbol: Socs7] [Locus Tag: ] [Chromosome: 11] [Map Location: 11\|11 D] | 1.482621361 | 0.012205 | 0.024869 |
| 17754 | ENSMUSG00000031558 | Slit2 | 20563 [Gene Symbol: Slit2] [Locus Tag: ] [Chromosome: 5] [Map Location: 5\|5 B3] [Des | 1.481444915 | 0.019249 | 0.037243 |
| 6879 | ENSMUSG00000005312 | Ubqln1 | 56085 [Gene Symbol: Ubqln1] [Locus Tag: ] [Chromosome: 13] [Map Location: 13 B1\|1 | 1.480809931 | 0.019776 | 0.038139 |
| 11882 | ENSMUSG00000028944 | Prkag2 | 108099 [Gene Symbol: Prkag2] [Locus Tag: ] [Chromosome: 5] [Map Location: 5\|5 A3] [ | 1.478376955 | 0.01381 | 0.027754 |
| 2807 | ENSMUSG00000040021 | Lats1 | 16798 [Gene Symbol: Lats1] [Locus Tag: ] [Chromosome: 10] [Map Location: 10\|10 A1] | 1.478045524 | 0.01214 | 0.024761 |
| 1965 | ENSMUSG00000040720 | 1110037F02Rik | 66185 [Gene Symbol: 1110037F02Rik] [Locus Tag: ] [Chromosome: 4] [Map Location: 4 | 1.477143007 | 0.016922 | 0.033213 |
| 8301 | ENSMUSG00000057522 | Spop | 20747 [Gene Symbol: Spop] [Locus Tag: ] [Chromosome: 11] [Map Location: 11 D\|11 59 | 1.476162636 | 0.025911 | 0.048597 |
| 7872 | ENSMUSG00000029821 | Dfna5 | 54722 [Gene Symbol: Dfna5] [Locus Tag: ] [Chromosome: 6] [Map Location: 6\|6 B2.3] [I | 1.47582173 | 0.013238 | 0.026712 |
| 9210 | ENSMUSG00000024054 | Smchd1 | 74355 [Gene Symbol: Smchd1] [Locus Tag: ] [Chromosome: 17] [Map Location: 17\|17 E | 1.474518077 | 0.016996 | 0.033336 |
| 7986 | ENSMUSG00000042323 | Pbrm1 | 66923 [Gene Symbol: Pbrm1] [Locus Tag: ] [Chromosome: 14] [Map Location: 14\|14 B] | 1.472766596 | 0.022479 | 0.042753 |
| 11545 | ENSMUSG00000039163 | Cmc1 | 67899 [Gene Symbol: Cmc1] [Locus Tag: ] [Chromosome: 9] [Map Location: 9\|9 F3] [De | 1.472281879 | 0.025198 | 0.047397 |
| 7522 | ENSMUSG00000045763 | Basp1 | 70350 [Gene Symbol: Basp1] [Locus Tag: ] [Chromosome: 15] [Map Location: 15\|15 B1] | 1.471512007 | 0.012626 | 0.025626 |
| 7467 | ENSMUSG00000035653 | Lrfn5 | 238205 [Gene Symbol: Lrfn5] [Locus Tag: ] [Chromosome: 12] [Map Location: 12\|12 C1 | 1.471415768 | 0.011136 | 0.02294 |
| 9489 | ENSMUSG00000045071 | E130308A19Rik | 230259 [Gene Symbol: E130308A19Rik] [Locus Tag: ] [Chromosome: 4] [Map Location: | 1.470060557 | 0.018805 | 0.036503 |
| 13692 | ENSMUSG00000015755 | Tab2 | 68652 [Gene Symbol: Tab2] [Locus Tag: ] [Chromosome: 10] [Map Location: 10\|10 A1] [ | 1.469433095 | 0.012226 | 0.024904 |
| 2187 | ENSMUSG00000019977 | Hbs1l | 56422 [Gene Symbol: Hbs1l] [Locus Tag: ] [Chromosome: 10] [Map Location: 10\|10 A3] | 1.468749898 | 0.022914 | 0.043482 |
| 1425 | ENSMUSG00000026782 | Abi2 | 329165 [Gene Symbol: Abi2] [Locus Tag: ] [Chromosome: 1] [Map Location: 1\|1 C2] [De | 1.467945667 | 0.013411 | 0.027031 |
| 10645 | ENSMUSG00000042659 | Arrdc4 | 66412 [Gene Symbol: Arrdc4] [Locus Tag: ] [Chromosome: 7] [Map Location: 7\|7 D1] [D | 1.466627025 | 0.01665 | 0.032772 |
| 13973 | ENSMUSG00000055254 | Ntrk2 | 18212 [Gene Symbol: Ntrk2] [Locus Tag: ] [Chromosome: 13] [Map Location: 13 B1-B2\| | 1.465558579 | 0.010842 | 0.022386 |
| 6652 | ENSMUSG00000024867 | Pip5k1b | 18719 [Gene Symbol: Pip5k1b] [Locus Tag: ] [Chromosome: 19] [Map Location: 19\|19 ( | 1.463909201 | 0.024898 | 0.046881 |
| 7274 | ENSMUSG00000054640 | Slc8a1 | 20541 [Gene Symbol: Slc8a1] [Locus Tag: ] [Chromosome: 17] [Map Location: 17 E3\|17 | 1.462014544 | 0.01459 | 0.029172 |
| 9225 | ENSMUSG00000051344 | Plekhm3 | 241075 [Gene Symbol: Plekhm3] [Locus Tag: ] [Chromosome: 1] [Map Location: 1\|1 C2 | 1.461740682 | 0.019833 | 0.03823 |
| 6137 | ENSMUSG00000027006 | Dnajc10 | 66861 [Gene Symbol: Dnajc10] [Locus Tag: ] [Chromosome: 2] [Map Location: 2\|2 D] [D | 1.461479312 | 0.026565 | 0.049702 |
| 14209 | ENSMUSG00000040850 | Psme4 | 103554 [Gene Symbol: Psme4] [Locus Tag: ] [Chromosome: 11] [Map Location: 11\|11 A | 1.459658953 | 0.020355 | 0.039129 |
| 7340 | ENSMUSG00000039458 | Mtmr12 | 268783 [Gene Symbol: Mtmr12] [Locus Tag: ] [Chromosome: 15] [Map Location: 15\|15 | 1.458180899 | 0.022839 | 0.043358 |
| 14620 | ENSMUSG00000008658 | Rbfox1 | 268859 [Gene Symbol: Rbfox1] [Locus Tag: ] [Chromosome: 16] [Map Location: 16 A1\| | 1.456851613 | 0.013422 | 0.027051 |
| 14754 | ENSMUSG00000034647 | Ankrd12 | 106585 [Gene Symbol: Ankrd12] [Locus Tag: ] [Chromosome: 17] [Map Location: 17\|17 | 1.45529082 | 0.011913 | 0.024355 |
| 1028 | ENSMUSG00000049690 | Nckap5 | 210356 [Gene Symbol: Nckap5] [Locus Tag: ] [Chromosome: 1] [Map Location: 1\|1 E3] [ | 1.454562412 | 0.026097 | 0.048891 |
| 7387 | ENSMUSG00000024294 | Mib1 | 225164 [Gene Symbol: Mib1] [Locus Tag: ] [Chromosome: 18] [Map Location: 18\|18 A1 | 1.45256629 | 0.016296 | 0.032154 |
| 3239 | ENSMUSG00000025964 | Adam23 | 23792 [Gene Symbol: Adam23] [Locus Tag: ] [Chromosome: 1] [Map Location: 1 C2\|1 3 | 1.451868797 | 0.015267 | 0.03034 |
| 9530 | ENSMUSG00000069495 | Epc2 | 227867 [Gene Symbol: Epc2] [Locus Tag: ] [Chromosome: 2] [Map Location: 2 C1.1\|2 28 | 1.44660808 | 0.015421 | 0.030629 |
| 17927 | ENSMUSG00000033342 | Plppr5 | 75769 [Gene Symbol: Plppr5] [Locus Tag: ] [Chromosome: 3] [Map Location: 3\|3 G2] [D | 1.446459468 | 0.024898 | 0.046881 |
| 3523 | ENSMUSG00000014496 | Ankrd28 | 105522 [Gene Symbol: Ankrd28] [Locus Tag: ] [Chromosome: 14] [Map Location: 14\|14 | 1.44582248 | 0.019805 | 0.038191 |
| 11004 | ENSMUSG00000012640 | Zfp715 | 69930 [Gene Symbol: Zfp715] [Locus Tag: ] [Chromosome: 7] [Map Location: 7\|7 B4] [D | 1.444409488 | 0.02555 | 0.048006 |
| 17258 | ENSMUSG00000041417 | Pik3r1 | 18708 [Gene Symbol: Pik3r1] [Locus Tag: ] [Chromosome: 13] [Map Location: 13 D1\|13 | 1.443981789 | 0.014561 | 0.029121 |
| 11853 | ENSMUSG00000036737 | Oxsr1 | 108737 [Gene Symbol: Oxsr1] [Locus Tag: ] [Chromosome: 9] [Map Location: 9\|9 F3] [D | 1.440274848 | 0.024693 | 0.046546 |
| 3761 | ENSMUSG00000057649 | Brd9 | 105246 [Gene Symbol: Brd9] [Locus Tag: ] [Chromosome: 13] [Map Location: 13\|13 C1 | 1.439967071 | 0.021403 | 0.040933 |
| 10 | ENSMUSG00000030806 | Stx1b | 56216 [Gene Symbol: Stx1b] [Locus Tag: ] [Chromosome: 7] [Map Location: 7\|7 F3] [Des | 1.439417275 | 0.018451 | 0.035889 |
| 2133 | ENSMUSG00000029467 | Atp2a2 | 11938 [Gene Symbol: Atp2a2] [Locus Tag: ] [Chromosome: 5] [Map Location: 5 62.38 cM | 1.437973146 | 0.020008 | 0.038533 |
| 8908 | ENSMUSG00000068748 | Ptprz1 | 19283 [Gene Symbol: Ptprz1] [Locus Tag: ] [Chromosome: 6] [Map Location: 6\|6 A3.1] [ | 1.432670572 | 0.023699 | 0.044826 |
| 13530 | ENSMUSG00000092083 | Kcnb2 | 98741 [Gene Symbol: Kcnb2] [Locus Tag: ] [Chromosome: 1] [Map Location: 1 A3\|1 4.5 | 1.431313401 | 0.018405 | 0.035813 |
| 13588 | ENSMUSG00000021745 | Ptprg | 19270 [Gene Symbol: Ptprg] [Locus Tag: ] [Chromosome: 14] [Map Location: 14 6.33 cM | 1.430380144 | 0.024255 | 0.045794 |
| 16981 | ENSMUSG00000045098 | Suv420h1 | 225888 [Gene Symbol: Suv420h1] [Locus Tag: ] [Chromosome: 19] [Map Location: 19\| | 1.429762758 | 0.026209 | 0.04907 |
| 15326 | ENSMUSG00000028284 | Map3k7 | 26409 [Gene Symbol: Map3k7] [Locus Tag: ] [Chromosome: 4] [Map Location: 4\|4 A5] [ | 1.428588259 | 0.026035 | 0.048804 |
| 7155 | ENSMUSG00000055435 | Maf | 17132 [Gene Symbol: Maf] [Locus Tag: ] [Chromosome: 8] [Map Location: 8 E1\|8 62.61 | 1.426745242 | 0.021425 | 0.040971 |
| 17036 | ENSMUSG00000032560 | Dnajc13 | 235567 [Gene Symbol: Dnajc13] [Locus Tag: ] [Chromosome: 9] [Map Location: 9\|9 F1] | 1.424498544 | 0.026685 | 0.049897 |
| 17337 | ENSMUSG00000029454 | Mapkapk5 | 17165 [Gene Symbol: Mapkapk5] [Locus Tag: ] [Chromosome: 5] [Map Location: 5\|5 F] [ | 1.424146107 | 0.025554 | 0.048008 |
| 12742 | ENSMUSG00000035051 | Dhx57 | 106794 [Gene Symbol: Dhx57] [Locus Tag: ] [Chromosome: 17] [Map Location: 17\|17 E | 1.42229658 | 0.024652 | 0.046478 |
| 15786 | ENSMUSG00000025909 | Sntg1 | 71096 [Gene Symbol: Sntg1] [Locus Tag: ] [Chromosome: 1] [Map Location: 1\|1 A1-A2] | 1.422287364 | 0.024999 | 0.047051 |
| 1416 | ENSMUSG00000027304 | Rtf1 | 76246 [Gene Symbol: Rtf1] [Locus Tag: ] [Chromosome: 2] [Map Location: 2\|2 E5] [Desc | 1.419322298 | 0.026169 | 0.049011 |

**Supplemental Table S3: Overlapping datasets of KHSRP targets**

**527 common elements in "KHSRP RIP target" and "Upregulated in CTX":**

| | | | |
|---|---|---|---|
| Rabggtb | | | |
| Rnf138 | **444 common elements in "KHSRP RIP target", "Up in CTX" and "ARE+targets":** | | |
| Tmem68 | Rnf138 | | |
| Cxcl10 | Tmem68 | **NS morphology (58 genes)** | |
| Serf1 | Cxcl10 | **Gene symbol** | **Fold change** |
| Ormdl1 | Ormdl1 | Arhgap5 | 1.3696048 |
| Tvp23b | Tvp23b | Arl6 | 1.2880108 |
| Lnp | Lnp | Atrx | 1.3186031 |
| Fam151b | Fam151b | Bmpr1b | 1.8156981 |
| Eif5a2 | Eif5a2 | Cadps | 1.2885912 |
| Manea | Manea | Calb1 | 1.3776299 |
| Ccdc126 | Ccdc126 | Cd47 | 1.6362207 |
| Chp2 | Dck | Chm | 1.62005 |
| Dck | Pank3 | Cpeb2 | 1.3237228 |
| Pank3 | Zc2hc1a | Dnm3 | 1.3638535 |
| Zc2hc1a | Cav2 | Epha7 | 1.5449396 |
| Cav2 | Katnbl1 | Ercc8 | 1.4539671 |
| Zfp97 | Yipf6 | Esco2 | 1.5194742 |
| Katnbl1 | Papd4 | Fubp1 | 2.2033444 |
| Yipf6 | Derl2 | Gata3 | 4.2416436 |
| Papd4 | Eny2 | Gdap1 | 1.3758444 |
| Derl2 | Fsbp | Gnas | 1.485868 |
| Eny2 | Rap1b | Gpm6b | 1.3679772 |
| Fsbp | Gpm6a | Grik2 | 1.5242547 |
| Rap1b | Tmem33 | Ids | 1.3076414 |
| Gpm6a | Ccdc82 | Intu | 1.4184539 |
| Tmem33 | Cks2 | Itgb8 | 1.7785179 |
| Ccdc82 | Tubgcp5 | Kif15 | 2.3346397 |
| Cks2 | Vstm2a | Lamp2 | 1.5246622 |
| Tubgcp5 | Mtx2 | Lypla1 | 1.370059 |
| Vstm2a | Mob1b | Mapk1 | 1.3214001 |
| Mtx2 | Dpy19l4 | Marcks | 1.4660375 |
| Mob1b | Cnot6 | Mbnl1 | 1.6097215 |
| Ccdc90b | Hnrnpa3 | Ncam2 | 1.8516387 |
| Tmem30a | Zc3h8 | Ndufs4 | 1.8307418 |
| Dpy19l4 | Fbxo48 | Nfia | 1.4630476 |
| Klhl7 | Tmem123 | Nrn1 | 1.2754371 |
| Scoc | Aasdhppt | Ogt | 1.3811442 |
| Cnot6 | Yipf4 | Oma1 | 1.2558534 |
| Ccdc58 | Gtf2a2 | Pcdh9 | 1.3678143 |
| Gmnn | Spopl | Pdpk1 | 1.404662 |
| Gm527 | Yy2 | Pex13 | 1.2707991 |
| Hnrnpa3 | Dzip3 | Pex2 | 1.3114293 |
| Zc3h8 | Syncrip | Pex5 | 1.8517856 |
| Fbxo48 | Tdp2 | Phyh | 1.3136758 |

| | | | |
|---|---|---|---|
| Tmem123 | Sdcbp | Prok2 | 1.5952196 |
| Aasdhppt | Degs1 | Rac1 | 1.3804698 |
| Yipf4 | Tcea1 | Rb1 | 1.3391191 |
| Tm9sf2 | Srsf1 | Reep1 | 1.7508531 |
| Masp2 | Atl2 | Rhot1 | 1.5056095 |
| Gtf2a2 | Fam76b | Robo1 | 2.2469979 |
| Spopl | Arpp19 | Rtn4 | 1.2599812 |
| Yy2 | Vamp4 | Shh | 1.4188314 |
| Dzip3 | Chmp5 | Slc1a2 | 1.3337701 |
| Syncrip | Ccne2 | Slc4a10 | 1.3262585 |
| Tdp2 | Elavl2 | Smarcc1 | 1.2665257 |
| Sdcbp | Tmem168 | Snap25 | 1.4057893 |
| Degs1 | Cept1 | Sox4 | 1.2695117 |
| Tcea1 | Gm10767 | St8sia4 | 1.4939186 |
| Srsf1 | Ssbp1 | Syncrip | 1.7566578 |
| Atl2 | Fam175a | Tor1a | 1.4646115 |
| Fam76b | Slitrk4 | Vezt | 1.3223154 |
| Arpp19 | Ttc8 | Wnt5b | 1.7126268 |
| Vamp4 | Ergic2 | | |
| Chmp5 | Tmem70 | | |
| Ccne2 | Ppm1k | | |
| Elavl2 | Nufip2 | | |
| Tmem168 | Ppp1cb | | |
| Spc25 | Trove2 | | |
| Cept1 | Cep97 | | |
| Trmt11 | Slc16a4 | | |
| Gm10767 | Tmco1 | | |
| Serpinb9 | Crem | | |
| Ssbp1 | St8sia4 | | |
| Fam175a | Nova1 | | |
| Slitrk4 | Lamp2 | | |
| Ttc8 | Caap1 | | |
| Nae1 | Ccnb1 | | |
| Olfr543 | Cbfb | | |
| Fbxo5 | Shh | | |
| Ergic2 | Spryd7 | | |
| Cdca3 | Nrn1 | | |
| Tmem70 | Rbbp5 | | |
| Ppm1k | Tfrc | | |
| Nufip2 | Ammecr1 | | |
| Ppp1cb | Stx19 | | |
| Trove2 | Slc35b3 | | |
| Cep97 | Dynlt3 | | |
| Slc16a4 | Mospd2 | | |
| Tmco1 | Hs3st1 | | |
| Crem | Gpr21 | | |
| St8sia4 | Tmem106b | | |
| Nova1 | Stt3a | | |

| | |
|---|---|
| Neil3 | Cpeb2 |
| Lamp2 | Dhx36 |
| Caap1 | Serpini1 |
| Ccnb1 | Txn1 |
| Cbfb | Ythdf3 |
| Shh | Dync1i2 |
| Spryd7 | Mbnl1 |
| Memo1 | Tmem55a |
| Nrn1 | Fbxo3 |
| Rbbp5 | Kbtbd8 |
| Tfrc | Tmem230 |
| Ammecr1 | Tceanc |
| Stx19 | Mybl1 |
| Slc35b3 | Arl1 |
| Dynlt3 | Skp2 |
| Mospd2 | Morf4l2 |
| Hs3st1 | Prok2 |
| Gpr21 | Pdcl |
| Tmem106b | Lypla1 |
| Vta1 | Slc25a32 |
| Stt3a | Hace1 |
| Cpeb2 | Hpgd |
| Dhx36 | Serf2 |
| Serpini1 | Gata3 |
| Txn1 | Atp11c |
| Ythdf3 | Slco4c1 |
| Dync1i2 | U2surp |
| Mbnl1 | Ogt |
| Tmem55a | Pcmtd1 |
| Fbxo3 | Hsd17b7 |
| Kbtbd8 | Pou2f1 |
| Tmem230 | Cnot7 |
| Tceanc | Fam199x |
| Mybl1 | Ndfip2 |
| Arl1 | Osbpl8 |
| Skp2 | Gabrg2 |
| Morf4l2 | Kdm6a |
| Prok2 | Armc8 |
| Pdcl | Ano5 |
| Lypla1 | Gria4 |
| Slc25a32 | Tmpo |
| Kif18a | Slc25a46 |
| Hace1 | Agk |
| Hpgd | Lcorl |
| Serf2 | Spock3 |
| Gata3 | Ddx26b |
| Trpc1 | Tmem87b |
| Atp11c | Cd47 |

| | |
|---|---|
| Slco4c1 | Prps2 |
| U2surp | Fbxo33 |
| Ogt | Ncam2 |
| Pcmtd1 | Zfp160 |
| Hsd17b7 | Hist2h2be |
| Pou2f1 | Grik2 |
| Cnot7 | Rcn2 |
| Pex7 | Zfp708 |
| Fam199x | Edil3 |
| Ndfip2 | Arl6 |
| Osbpl8 | Sdc2 |
| Gabrg2 | Hdgfrp3 |
| Kdm6a | Senp7 |
| Armc8 | Rpgr |
| Ano5 | Zfp451 |
| Gria4 | Coa5 |
| Tmpo | Chm |
| Slc25a46 | Phip |
| Agk | Cdc73 |
| Lcorl | Ero1lb |
| Spock3 | Gpr165 |
| Ddx26b | Kbtbd7 |
| Tmem87b | Fbxl3 |
| Cd47 | Bmpr1b |
| Prps2 | Tm6sf1 |
| Fbxo33 | Ids |
| Ncam2 | Phf14 |
| Zfp160 | Rnft1 |
| Hist2h2be | Impa1 |
| Grik2 | Abcd3 |
| Rcn2 | Cops2 |
| Zfp708 | Fubp1 |
| Edil3 | Rsrc2 |
| Arl6 | Neto2 |
| Sdc2 | Tank |
| Hdgfrp3 | Ercc8 |
| Senp7 | Nap1l3 |
| Rpgr | Arl5a |
| Gin1 | Suco |
| Zfp451 | Pdpk1 |
| Coa5 | Tmem65 |
| Chm | Rmnd1 |
| Uba3 | Paip2 |
| Phip | Crebzf |
| Cdc73 | Ube2d2a |
| Ero1lb | Lsmem1 |
| Gpr165 | Acadsb |
| Pcbd2 | Krt222 |

| | |
|---|---|
| Kbtbd7 | Plod2 |
| Fbxl3 | Pex5 |
| Bmpr1b | Cox7b |
| Tm6sf1 | Slc4a10 |
| Ids | Abca8b |
| Phf14 | Dph6 |
| Rnft1 | Rad51ap1 |
| Zfp808 | Pex3 |
| Impa1 | Gdap1 |
| Abcd3 | Mospd1 |
| Cops2 | Marcks |
| Fubp1 | Mis18bp1 |
| Ttc21b | Esco2 |
| Rsrc2 | Gpm6b |
| Neto2 | Fip1l1 |
| Tank | Fnbp4 |
| Ercc8 | Rtn3 |
| Ngly1 | Fam172a |
| Nap1l3 | Eif2s2 |
| 7-Mar | Gjb2 |
| Arl5a | Depdc1b |
| Suco | Stag2 |
| Pdpk1 | Pi4k2b |
| Bcap31 | Zdhhc2 |
| Tmem65 | Oma1 |
| Smpdl3a | Mbnl3 |
| Wnt5b | Tmem135 |
| Rmnd1 | Tram1 |
| Paip2 | Xpo4 |
| Crebzf | Mkrn3 |
| Ube2d2a | Cxadr |
| Lsmem1 | Xrcc6bp1 |
| Ift88 | Mtpn |
| Acadsb | Ptpn4 |
| Krt222 | Rfk |
| Plod2 | Gucy1a2 |
| Pex5 | Mycbp |
| Cox7b | Rabl3 |
| Slc4a10 | Micu3 |
| Abca8b | Slc44a1 |
| Dph6 | Zfp40 |
| Rad51ap1 | Tor1a |
| Pex3 | Atad2 |
| Gdap1 | Camk2d |
| Mospd1 | C2cd5 |
| Marcks | Ube2j1 |
| Mis18bp1 | Tmem14a |
| Esco2 | Epha7 |

| | |
|---|---|
| Prps1l3 | Cep70 |
| Gpm6b | Tmem209 |
| Fip1l1 | Ppp1r12a |
| Fnbp4 | C1galt1 |
| Rfesd | Lrrcc1 |
| Rtn3 | Eif2s3x |
| Fam172a | Hmbox1 |
| Eif2s2 | Mbd4 |
| Gjb2 | Nusap1 |
| Depdc1b | Celf1 |
| Stag2 | Ldlrad3 |
| Pi4k2b | Tiprl |
| Zdhhc2 | Agps |
| Oma1 | Sgms2 |
| Mbnl3 | Atp5s |
| Tmem135 | Elovl7 |
| Tram1 | Ghitm |
| Xpo4 | Strbp |
| Mkrn3 | Pla2g12a |
| Cxadr | Kctd9 |
| Xrcc6bp1 | Wdr44 |
| Mtpn | Nup155 |
| Ptpn4 | Zfp518a |
| Rfk | Rraga |
| Gucy1a2 | Ypel2 |
| Mycbp | Zmym6 |
| Rabl3 | Kcnj2 |
| Micu3 | Kpna3 |
| Slc44a1 | Intu |
| Zfp40 | Ccna2 |
| Tor1a | Asb3 |
| Atad2 | Psmd10 |
| Camk2d | Rb1 |
| C2cd5 | Lca5 |
| Rbm28 | Glrb |
| Ube2j1 | Atp1b3 |
| Tmem14a | Slc39a10 |
| Acyp1 | Mtdh |
| Epha7 | Suz12 |
| Cep70 | Mapk1 |
| Tmem209 | Itgb8 |
| Ppp1r12a | Abcc5 |
| C1galt1 | Sox4 |
| Lrrcc1 | Grhl1 |
| Eif2s3x | Abhd17b |
| Hmbox1 | Aebp2 |
| Mbd4 | Reep1 |
| Nusap1 | Cdk17 |

| | |
|---|---|
| Celf1 | Hspa4l |
| Ldlrad3 | Fam135a |
| Tiprl | Stam2 |
| Vegfc | Snx7 |
| Agps | Ano3 |
| Sgms2 | Prc1 |
| Atp5s | Pex13 |
| Elovl7 | Rgs7bp |
| Ghitm | Gabra4 |
| Ipo7 | Gpr155 |
| Fxr1 | Anp32e |
| Strbp | Hnrnpll |
| Pla2g12a | Seh1l |
| Kctd9 | Osgepl1 |
| Wdr44 | Zfp760 |
| Nup155 | Rac1 |
| Zfp518a | Kcnc2 |
| Pex2 | Zfp943 |
| Rraga | Rps15a |
| Ypel2 | Fam45a |
| Zmym6 | Usp14 |
| Kcnj2 | Birc5 |
| Zfp809 | Tmem186 |
| Kpna3 | Bard1 |
| Intu | Mcm9 |
| Ccna2 | Rhot1 |
| Asb3 | Nfia |
| Psmd10 | Nars2 |
| Rb1 | Tmem196 |
| Lca5 | Pgm3 |
| Glrb | Rgs4 |
| Atp1b3 | Dcaf17 |
| Slc39a10 | Lmo2 |
| Mtdh | Med23 |
| Suz12 | Asph |
| Mapk1 | Setd7 |
| Itgb8 | Zfp120 |
| Abcc5 | Tbx18 |
| Sox4 | Zfp248 |
| Grhl1 | Tox |
| Abhd17b | Cstf3 |
| Aebp2 | Mpp7 |
| Reep1 | Wls |
| Ndufab1 | Atp2c1 |
| Cdk17 | Snap25 |
| Hspa4l | Mal2 |
| Fam135a | Mad2l1 |
| Stam2 | Extl2 |

| | |
|---|---|
| Snx7 | Atxn7l3b |
| Ano3 | Ern1 |
| Prc1 | Socs6 |
| Pex13 | Ate1 |
| Rgs7bp | Commd10 |
| Gabra4 | Lyplal1 |
| Nrg1 | Pdp2 |
| Gpr155 | Stau2 |
| Anp32e | Tmem64 |
| Mettl4 | Jazf1 |
| Hnrnpll | Dnm3 |
| Seh1l | Dmxl1 |
| Osgepl1 | Sestd1 |
| Zfp760 | Ndufs4 |
| Rac1 | Trip4 |
| Kcnc2 | Scfd2 |
| Zfp943 | Pcdh9 |
| Rps15a | Fam227a |
| Arhgap5 | Stambpl1 |
| Fam45a | Calb1 |
| Usp14 | Cc2d2a |
| Birc5 | Brwd1 |
| Kbtbd3 | Drd1 |
| Ccdc160 | Cflar |
| Tmem186 | Alcam |
| Bard1 | Atrx |
| Mcm9 | Mpzl1 |
| Rhot1 | Sgcb |
| Nfia | Mrpl39 |
| Nars2 | Frmd3 |
| Mtrf1 | Azi2 |
| Tmem196 | Wac |
| Pgm3 | Zfp354a |
| Wdr78 | Casc5 |
| Rgs4 | Rfwd2 |
| Dcaf17 | Cadps |
| Lmo2 | Zfp770 |
| Med23 | Arl8b |
| Asph | Serinc3 |
| Cep41 | Ola1 |
| Lipo1 | Pard6b |
| Setd7 | Rad21 |
| Zfp120 | Gulp1 |
| Tbx18 | Casc4 |
| Zfp248 | Hs2st1 |
| Tox | Tet1 |
| Nek3 | Nsmce2 |
| Cstf3 | Kif15 |

| | |
|---|---|
| Mpp7 | Wdr19 |
| Wls | Grem2 |
| Atp2c1 | Sh3bgrl2 |
| Snap25 | Dcp1b |
| Mal2 | Fabp7 |
| Mad2l1 | St3gal6 |
| Extl2 | Scn3a |
| Atxn7l3b | Vcpip1 |
| Gen1 | Gk5 |
| Ern1 | Vezt |
| Socs6 | Dusp22 |
| Ate1 | Gucy1b3 |
| Commd10 | Zc3h6 |
| Lyplal1 | Zbtb1 |
| Pdp2 | Tpm1 |
| Stau2 | Nipal2 |
| Tmem64 | Phyh |
| Jazf1 | Vrk1 |
| Dnm3 | Higd1a |
| Dnm1l | Uxs1 |
| Dmxl1 | Pcdhb14 |
| Sestd1 | Car8 |
| Ndufs4 | Hsdl2 |
| Trip4 | Ankrd45 |
| Scfd2 | Pcdhb20 |
| Pcdh9 | Gfpt2 |
| Fam227a | Dclre1c |
| Stambpl1 | Jam2 |
| Ccdc53 | Oxr1 |
| Calb1 | Ubr3 |
| Cc2d2a | Usp33 |
| Fsip1 | Gfm2 |
| Adprm | Tpp2 |
| Gdpd2 | Sgtb |
| Brwd1 | Robo1 |
| Drd1 | Gnai1 |
| Cflar | Ado |
| Alcam | Rab11b |
| Atrx | Uri1 |
| Mpzl1 | Pcf11 |
| Sgcb | Slc1a2 |
| Mrpl39 | Adal |
| Ndufb5 | B3galt1 |
| Frmd3 | Tnfrsf19 |
| Azi2 | Slc2a13 |
| Wac | Pcdhb17 |
| Zfp354a | Ccser1 |
| Casc5 | Slc7a6os |

| | |
|---|---|
| Rfwd2 | Fmnl2 |
| Acadm | Kcmf1 |
| Cadps | Ccdc148 |
| Zfp770 | Epc1 |
| Ercc6l2 | Kcnh7 |
| Arl8b | Arhgap42 |
| Zfp157 | Rnf103 |
| Gpbp1 | Wdr33 |
| Serinc3 | Zyg11b |
| Ola1 | Smek1 |
| Pard6b | Wdr17 |
| Hibch | Smarcc1 |
| Rab5a | Rbms3 |
| Rad21 | Ubqln1 |
| Gulp1 | Smchd1 |
| Casc4 | Dnajc10 |
| Hs2st1 | Atp2a2 |
| Ranbp2 | |
| Tet1 | |
| Nsmce2 | |
| Lyrm4 | |
| Kif15 | |
| Wdr19 | |
| Grem2 | |
| Clk3 | |
| Sh3bgrl2 | |
| Dcp1b | |
| Fabp7 | |
| St3gal6 | |
| Scn3a | |
| Vcpip1 | |
| Gk5 | |
| Vezt | |
| Dusp22 | |
| Gucy1b3 | |
| Fbxo36 | |
| Zc3h6 | |
| Zbtb1 | |
| Tpm1 | |
| Nipal2 | |
| Ascc3 | |
| Phyh | |
| Vrk1 | |
| Higd1a | |
| Uxs1 | |
| Pcdhb14 | |
| Car8 | |
| Hsdl2 | |

Ankrd45
Pcdhb20
Gfpt2
Dclre1c
Jam2
Oxr1
Ubr3
Usp33
Nek4
Gfm2
Tpp2
Sgtb
Robo1
Gnai1
Ado
Rab11b
Uri1
Pcf11
Slc1a2
Adal
Magee1
B3galt1
Rtn4
Tnfrsf19
Slc2a13
Pcdhb17
Ccser1
Slc7a6os
Tdp1
Gnas
Fmnl2
Kcmf1
Ccdc148
Epc1
Arl2bp
Kcnh7
Arhgap42
Ybx1
Rnf103
Wdr33
Zyg11b
Smek1
Wdr17
Smarcc1
Rbms3
Ubqln1
Smchd1
Dnajc10

Psme4
Atp2a2

**Supplemental Table S4: KSRP-RIP-Seq 3'UTR AU-rich analyses as described in Bolognani et al, 2010, https://doi.org/10.1093/nar/gkp863**

| | | | % nucleotides in 3' UTRs | | | | Number of motifs in 3' UTRs | | | | | | | | | | | | | | |
|---|---|---|---|---|---|---|---|---|---|---|---|---|---|---|---|---|---|---|---|---|---|
| Gene ID | Gene Symbol | Transcript ID | %A | %T | %C | %G | ARED | OL AUUUA | Type I | Type II | Type IIA | Type IIB | Type IIC | Type IID | Type IIE | All ARE types | GRE | Motif 1 | Motif 2 | Motif 3 | ARE+Motifs |
| ENSMUSG00000042208 | 0610010F05Rik | ENSMUST00000043356 | 28.7892377 | 34 | 17.2197309 | 20.3139013 | 2 | 0 | 0 | 0 | 0 | 0 | 0 | 0 | 0 | 0 | 0 | 0 | 0 | 0 | 2 |
| ENSMUSG00000028608 | 0610037L13Rik | ENSMUST00000135454 | 28.0788177 | 25 | 22.2906404 | 24.4663383 | 0 | 0 | 0 | 0 | 0 | 0 | 0 | 0 | 0 | 0 | 0 | 1 | 0 | 0 | 1 |
| ENSMUSG00000040720 | 1110037F02Rik | ENSMUST00000059914 | 30.7614213 | 33 | 19.2893401 | 16.8527919 | 0 | 1 | 0 | 0 | 0 | 0 | 0 | 0 | 0 | 0 | 0 | 0 | 0 | 0 | 1 |
| ENSMUSG00000032551 | 1110059G10Rik | ENSMUST00000035154 | 28.8057041 | 29 | 21.14082 | 20.6417112 | 1 | 1 | 0 | 1 | 0 | 0 | 0 | 0 | 1 | 2 | 0 | 0 | 1 | 1 | 6 |
| ENSMUSG00000045414 | 1190002N15Rik | ENSMUST00000113028 | 30.9669523 | 34 | 17.3398613 | 17.50306 | 0 | 0 | 0 | 1 | 0 | 0 | 0 | 0 | 1 | 2 | 0 | 0 | 1 | 0 | 3 |
| ENSMUSG00000063320 | 1190007I07Rik | ENSMUST00000079648 | 31.7343173 | 32 | 18.0811808 | 17.7121771 | 0 | 0 | 1 | 1 | 0 | 0 | 0 | 0 | 1 | 3 | 0 | 0 | 0 | 0 | 3 |
| ENSMUSG00000038323 | 1700066M21Rik | ENSMUST00000042734 | 31.3609467 | 34 | 17.0521786 | 17.859064 | 0 | 0 | 0 | 2 | 0 | 0 | 0 | 0 | 2 | 4 | 0 | 0 | 1 | 1 | 6 |
| ENSMUSG00000056313 | 1810011O10Rik | ENSMUST00000052622 | 27.9369628 | 31 | 20.2005731 | 20.6303725 | 1 | 0 | 0 | 2 | 0 | 0 | 0 | 0 | 2 | 4 | 0 | 0 | 3 | 3 | 11 |
| ENSMUSG00000022507 | 1810013L24Rik | ENSMUST00000023150 | 26.8589298 | 35 | 18.4503127 | 19.9096595 | 0 | 0 | 0 | 0 | 0 | 0 | 0 | 0 | 0 | 0 | 1 | 1 | 10 | 0 | 12 |
| ENSMUSG00000044148 | 1810030O07Rik | ENSMUST00000060108 | 28.464885 | 28 | 22.5605966 | 21.0689869 | 0 | 0 | 0 | 0 | 0 | 0 | 0 | 0 | 0 | 0 | 0 | 1 | 0 | 0 | 1 |
| ENSMUSG00000074829 | 2010315B03Rik | ENSMUST00000177714 | 30.6655434 | 34 | 16.6807077 | 18.9553496 | 0 | 0 | 0 | 0 | 0 | 0 | 0 | 0 | 0 | 0 | 1 | 0 | 4 | 0 | 5 |
| ENSMUSG00000032403 | 2300009A05Rik | ENSMUST00000168665 | 30.6930693 | 35 | 14.8514851 | 19.8019802 | 0 | 0 | 1 | 0 | 0 | 0 | 0 | 0 | 0 | 1 | 0 | 0 | 0 | 0 | 1 |
| ENSMUSG00000026319 | 2310035C23Rik | ENSMUST00000185692 | 28.678376 | 32 | 18.7158077 | 20.2994529 | 1 | 0 | 0 | 0 | 0 | 0 | 0 | 0 | 0 | 0 | 0 | 0 | 1 | 0 | 2 |
| ENSMUSG00000040177 | 2310057M21Rik | ENSMUST00000059438 | 24.6478873 | 36 | 17.1361502 | 22.6525822 | 0 | 0 | 2 | 0 | 0 | 0 | 0 | 0 | 0 | 2 | 0 | 0 | 1 | 0 | 3 |
| ENSMUSG00000050705 | 2310061I04Rik | ENSMUST00000154670 | 22.7272727 | 23 | 28.1818182 | 25.7575758 | 0 | 0 | 0 | 0 | 0 | 0 | 0 | 0 | 0 | 0 | 0 | 3 | 0 | 0 | 3 |
| ENSMUSG00000036873 | 2410004B18Rik | ENSMUST00000134575 | 27.8947368 | 27 | 20.3759398 | 24.8120301 | 0 | 0 | 1 | 0 | 0 | 0 | 0 | 0 | 0 | 1 | 0 | 0 | 0 | 2 | 3 |
| ENSMUSG00000030924 | 2610020H08Rik | ENSMUST00000133758 | 29.6432964 | 27 | 21.1562116 | 21.8942189 | 0 | 0 | 0 | 0 | 0 | 0 | 0 | 0 | 0 | 0 | 0 | 1 | 0 | 0 | 1 |
| ENSMUSG00000028060 | 2810403A07Rik | ENSMUST00000199684 | 25.9655654 | 33 | 19.032108 | 21.8706375 | 1 | 0 | 1 | 1 | 0 | 0 | 0 | 0 | 1 | 3 | 0 | 0 | 1 | 0 | 5 |
| ENSMUSG00000040204 | 2810417H13Rik | ENSMUST00000045802 | 28.2621768 | 34 | 18.0998196 | 20.0240529 | 0 | 0 | 0 | 0 | 0 | 0 | 0 | 0 | 0 | 0 | 0 | 0 | 5 | 0 | 5 |
| ENSMUSG00000067879 | 3110035E14Rik | ENSMUST00000088666 | 29.9510458 | 30 | 19.9376947 | 20.2047174 | 1 | 2 | 0 | 1 | 0 | 0 | 0 | 0 | 1 | 2 | 0 | 0 | 0 | 0 | 5 |
| ENSMUSG00000028300 | 3110043O21Rik | ENSMUST00000084724 | 25.1054852 | 34 | 19.6729958 | 21.6772152 | 1 | 0 | 0 | 1 | 0 | 0 | 0 | 0 | 1 | 2 | 0 | 0 | 4 | 0 | 7 |
| ENSMUSG00000033111 | 3830406C13Rik | ENSMUST00000112669 | 30.4143126 | 30 | 19.4915254 | 19.9152542 | 0 | 0 | 0 | 0 | 0 | 0 | 0 | 0 | 0 | 0 | 0 | 1 | 3 | 4 | 8 |
| ENSMUSG00000062822 | 4833420G17Rik | ENSMUST00000026519 | 30.500000 | ### | 13.000000 | 19.500000 | 0 | 0 | 0 | 0 | 0 | 0 | 0 | 0 | 0 | 0 | 1 | 0 | 0 | 0 | 1 |
| ENSMUSG00000036934 | 4921524J17Rik | ENSMUST00000047749 | 27.9368213 | 36 | 15.4985192 | 20.1382034 | 1 | 0 | 0 | 2 | 0 | 0 | 0 | 0 | 2 | 4 | 0 | 0 | 0 | 0 | 5 |
| ENSMUSG00000046567 | 4930430F08Rik | ENSMUST00000054471 | 34.5201238 | 36 | 15.247678 | 14.1640867 | 0 | 0 | 1 | 0 | 0 | 0 | 0 | 0 | 0 | 1 | 0 | 0 | 0 | 0 | 1 |
| ENSMUSG00000021098 | 4930447C04Rik | ENSMUST00000136075 | 31.2300829 | 35 | 16.3161249 | 17.654557 | 0 | 1 | 0 | 1 | 0 | 0 | 0 | 0 | 1 | 2 | 1 | 0 | 0 | 0 | 4 |
| ENSMUSG00000059920 | 4930453N24Rik | ENSMUST00000076991 | 29.734259 | 32 | 18.1709361 | 19.6313143 | 2 | 1 | 0 | 2 | 0 | 0 | 0 | 0 | 2 | 4 | 0 | 0 | 9 | 4 | 20 |
| ENSMUSG00000027811 | 4930579G24Rik | ENSMUST00000029388 | 25.6510417 | 34 | 19.7265625 | 20.8984375 | 0 | 0 | 0 | 1 | 0 | 0 | 0 | 0 | 1 | 2 | 0 | 1 | 0 | 1 | 4 |
| ENSMUSG00000049357 | 4933408B17Rik | ENSMUST00000129566 | 26.1316872 | 27 | 20.1646091 | 27.1604938 | 0 | 0 | 0 | 0 | 0 | 0 | 0 | 0 | 0 | 0 | 0 | 4 | 0 | 0 | 4 |
| ENSMUSG00000027942 | 4933434E20Rik; 1700094D03Rik | ENSMUST00000029551 | 26.1330409 | 30 | 24.378655 | 19.8099415 | 0 | 0 | 0 | 0 | 0 | 0 | 0 | 0 | 0 | 0 | 0 | 0 | 0 | 3 | 3 |
| ENSMUSG00000057181 | 5730455P16Rik | ENSMUST00000179332 | 32.1385542 | 27 | 19.3373494 | 21.2951807 | 0 | 0 | 0 | 2 | 0 | 0 | 0 | 0 | 2 | 4 | 0 | 3 | 0 | 0 | 7 |
| ENSMUSG00000050549 | 5730508B09Rik | ENSMUST00000199273 | 30.9369476 | 34 | 16.2639953 | 18.385386 | 1 | 0 | 0 | 1 | 0 | 0 | 0 | 0 | 1 | 2 | 0 | 0 | 0 | 0 | 3 |
| ENSMUSG00000047466 | 8030462N17Rik | ENSMUST00000074653 | 25.6189451 | 33 | 21.31324 | 19.6986006 | 1 | 0 | 0 | 1 | 0 | 0 | 0 | 0 | 1 | 2 | 0 | 0 | 0 | 0 | 3 |
| ENSMUSG00000040139 | 9430038I01Rik | ENSMUST00000120340 | 25.9604191 | 30 | 20.8381839 | 23.1199069 | 0 | 0 | 0 | 0 | 0 | 0 | 0 | 0 | 0 | 0 | 0 | 1 | 0 | 1 | 2 |
| ENSMUSG00000036275 | 9530068E07Rik | ENSMUST00000109057 | 24.5762712 | 31 | 21.4285714 | 23.2445521 | 0 | 0 | 0 | 1 | 0 | 0 | 0 | 0 | 1 | 2 | 0 | 0 | 0 | 1 | 3 |
| ENSMUSG00000034560 | A230046K03Rik | ENSMUST00000038388 | 31.017481 | 32 | 16.9430749 | 19.6772748 | 0 | 0 | 0 | 1 | 0 | 0 | 0 | 0 | 1 | 2 | 1 | 0 | 0 | 0 | 3 |
| ENSMUSG00000044060 | A830010M20Rik | ENSMUST00000112671 | 30.436055 | 33 | 17.7641206 | 18.7884109 | 1 | 0 | 1 | 3 | 0 | 0 | 0 | 0 | 3 | 7 | 2 | 1 | 1 | 0 | 12 |
| ENSMUSG00000057715 | A830018L16Rik | ENSMUST00000135014 | 31.4923397 | 33 | 17.7794591 | 17.7416304 | 1 | 0 | 0 | 1 | 0 | 0 | 0 | 0 | 1 | 2 | 0 | 0 | 1 | 1 | 5 |
| ENSMUSG00000044150 | A830080D01Rik | ENSMUST00000112464 | 28.9036545 | 39 | 16.3898117 | 16.1683278 | 0 | 0 | 0 | 0 | 0 | 0 | 0 | 0 | 0 | 0 | 1 | 1 | 0 | 0 | 2 |
| ENSMUSG00000055923 | Aasdh | ENSMUST00000123682 | 25.0555732 | 26 | 24.1346459 | 24.6744998 | 0 | 0 | 0 | 0 | 0 | 0 | 0 | 0 | 0 | 0 | 0 | 1 | 0 | 0 | 1 |
| ENSMUSG00000025894 | Aasdhppt | ENSMUST00000051589 | 30.9776833 | 35 | 16.3655685 | 17.3219979 | 1 | 0 | 0 | 0 | 0 | 0 | 0 | 0 | 0 | 0 | 0 | 0 | 3 | 2 | 6 |
| ENSMUSG00000018697 | Aatf | ENSMUST00000018841 | 29.7029703 | 28 | 29.7029703 | 12.8712871 | 0 | 0 | 0 | 0 | 0 | 0 | 0 | 0 | 0 | 0 | 0 | 0 | 0 | 1 | 1 |
| ENSMUSG00000015243 | Abca1 | ENSMUST00000030010 | 30.2915082 | 32 | 18.9797212 | 18.7896071 | 0 | 0 | 0 | 0 | 0 | 0 | 0 | 0 | 0 | 0 | 0 | 1 | 0 | 1 | 2 |
| ENSMUSG00000018800 | Abca5 | ENSMUST00000043961 | 29.0291262 | 36 | 16.6666667 | 18.7378641 | 0 | 0 | 0 | 1 | 0 | 0 | 0 | 0 | 1 | 2 | 0 | 2 | 0 | 2 | 6 |
| ENSMUSG00000041828 | Abca8a | ENSMUST00000100287 | 25.9259259 | 39 | 18.6666667 | 16.8888889 | 0 | 0 | 0 | 0 | 0 | 0 | 0 | 0 | 0 | 0 | 0 | 1 | 0 | 0 | 1 |
| ENSMUSG00000020620 | Abca8b | ENSMUST00000020948 | 33.1034483 | 31 | 15.9051724 | 19.9137931 | 0 | 0 | 0 | 0 | 0 | 0 | 0 | 0 | 0 | 0 | 0 | 0 | 3 | 1 | 4 |
| ENSMUSG00000040584 | Abcb1a | ENSMUST00000047753 | 35.8974359 | 29 | 15.4832347 | 19.6252465 | 0 | 0 | 0 | 1 | 0 | 0 | 0 | 0 | 1 | 2 | 0 | 0 | 1 | 0 | 3 |
| ENSMUSG00000031333 | Abcb7 | ENSMUST00000033695 | 30.8033077 | 36 | 15.9480213 | 17.4542233 | 1 | 1 | 0 | 0 | 0 | 0 | 0 | 0 | 0 | 0 | 0 | 1 | 3 | 3 | 9 |
| ENSMUSG00000022822 | Abcc5 | ENSMUST00000079158 | 20.5995388 | 31 | 25.5956956 | 22.6748655 | 0 | 0 | 1 | 0 | 0 | 0 | 0 | 0 | 0 | 1 | 0 | 1 | 0 | 0 | 2 |
| ENSMUSG00000028127 | Abcd3 | ENSMUST00000197662 | 29.6610169 | 29 | 18.6886708 | 22.4353256 | 0 | 0 | 0 | 0 | 0 | 0 | 0 | 0 | 0 | 0 | 0 | 0 | 1 | 3 | 4 |
| ENSMUSG00000033157 | Abhd10 | ENSMUST00000066983 | 27.8306878 | 35 | 19.047619 | 18.4656085 | 0 | 1 | 1 | 0 | 0 | 0 | 0 | 0 | 0 | 1 | 0 | 3 | 1 | 0 | 6 |
| ENSMUSG00000040396 | Abhd13 | ENSMUST00000048216 | 28.9269467 | 33 | 17.8485416 | 20.5512443 | 0 | 0 | 0 | 0 | 0 | 0 | 0 | 0 | 0 | 0 | 3 | 0 | 2 | 0 | 5 |
| ENSMUSG00000037818 | Abhd18 | ENSMUST00000204496 | 31.5130527 | 30 | 19.4192861 | 19.2061801 | 0 | 0 | 0 | 1 | 0 | 0 | 0 | 0 | 1 | 2 | 0 | 1 | 0 | 0 | 3 |
| ENSMUSG00000026782 | Abi2 | ENSMUST00000052332 | 27.7398067 | 33 | 19.4202215 | 19.6794721 | 1 | 1 | 0 | 1 | 0 | 0 | 0 | 0 | 1 | 2 | 0 | 1 | 9 | 2 | 16 |
| ENSMUSG00000049076 | Acap2 | ENSMUST00000058033 | 26.5191965 | 33 | 19.6796339 | 20.518688 | 0 | 0 | 0 | 0 | 0 | 0 | 0 | 0 | 0 | 0 | 0 | 0 | 1 | 3 | 4 |
| ENSMUSG00000026781 | Acbd5 | ENSMUST00000028121 | 32.814852 | 34 | 16.2067235 | 16.9091821 | 0 | 0 | 0 | 1 | 0 | 0 | 0 | 0 | 1 | 2 | 0 | 0 | 0 | 1 | 3 |
| ENSMUSG00000033701 | Acbd6 | ENSMUST00000035560 | 27.939437 | 30 | 21.1970665 | 20.6056305 | 0 | 0 | 0 | 0 | 0 | 0 | 0 | 0 | 0 | 0 | 0 | 30 | 3 | 1 | 34 |
| ENSMUSG00000038007 | Acer2 | ENSMUST00000045224 | 23.9757208 | 30 | 22.5493171 | 23.7632777 | 0 | 0 | 0 | 0 | 0 | 0 | 0 | 0 | 0 | 0 | 0 | 2 | 0 | 0 | 2 |
| ENSMUSG00000030760 | Acer3 | ENSMUST00000033020 | 29.7090352 | 30 | 21.0662328 | 19.5348392 | 2 | 1 | 0 | 3 | 0 | 0 | 0 | 1 | 2 | 6 | 11 | 3 | 15 | 1 | 39 |
| ENSMUSG00000032883 | Acsl3 | ENSMUST00000035779 | 28.1626506 | 34 | 15.6626506 | 22.6656627 | 0 | 0 | 0 | 0 | 0 | 0 | 0 | 0 | 0 | 0 | 0 | 0 | 0 | 2 | 2 |
| ENSMUSG00000031278 | Acsl4 | ENSMUST00000033634 | 31.0730253 | 35 | 15.3502235 | 18.2563338 | 0 | 0 | 0 | 1 | 0 | 0 | 0 | 0 | 1 | 2 | 0 | 0 | 8 | 1 | 11 |
| ENSMUSG00000020333 | Acsl6 | ENSMUST00000093106 | 30.8513627 | 30 | 20.3146951 | 18.5164372 | 0 | 0 | 0 | 0 | 0 | 0 | 0 | 0 | 0 | 0 | 0 | 0 | 0 | 1 | 1 |
| ENSMUSG00000030935 | Acsm3 | ENSMUST00000106529 | 33.5078534 | 32 | 18.6736475 | 16.1431065 | 1 | 0 | 1 | 0 | 0 | 0 | 0 | 0 | 0 | 1 | 0 | 0 | 0 | 0 | 2 |
| ENSMUSG00000021076 | Actr10 | ENSMUST00000021479 | 28.0898876 | 32 | 19.17603 | 20.3745318 | 2 | 1 | 0 | 3 | 0 | 0 | 0 | 1 | 2 | 6 | 0 | 0 | 8 | 2 | 19 |
| ENSMUSG00000020152 | Actr2 | ENSMUST00000000137 | 28.0503145 | 35 | 16.8553459 | 20.1677149 | 0 | 0 | 0 | 1 | 0 | 0 | 0 | 0 | 1 | 2 | 0 | 0 | 5 | 1 | 8 |
| ENSMUSG00000026836 | Acvr1 | ENSMUST00000112599 | 27.0588235 | 31 | 21.9004525 | 20.4524887 | 0 | 0 | 0 | 0 | 0 | 0 | 0 | 0 | 0 | 0 | 0 | 0 | 0 | 2 | 2 |
| ENSMUSG00000000532 | Acvr1b | ENSMUST00000000544 | 21.0152645 | 27 | 25.5591054 | 26.7660632 | 0 | 0 | 0 | 0 | 0 | 0 | 0 | 0 | 0 | 0 | 2 | 0 | 3 | 0 | 5 |
| ENSMUSG00000026834 | Acvr1c | ENSMUST00000028178 | 30.5157401 | 33 | 17.4815807 | 19.3703952 | 3 | 1 | 0 | 3 | 0 | 0 | 0 | 1 | 2 | 6 | 0 | 2 | 15 | 0 | 27 |
| ENSMUSG00000052155 | Acvr2a | ENSMUST00000063886 | 32.0022982 | 34 | 17.1215168 | 17.2938811 | 0 | 0 | 0 | 0 | 0 | 0 | 0 | 0 | 0 | 0 | 0 | 0 | 0 | 3 | 3 |
| ENSMUSG00000027259 | Adal | ENSMUST00000028702 | 27.0962048 | 31 | 21.5357458 | 20.6531333 | 0 | 0 | 0 | 0 | 0 | 0 | 0 | 0 | 0 | 0 | 0 | 0 | 1 | 0 | 1 |
| ENSMUSG00000054693 | Adam10 | ENSMUST00000067880 | 29.2271663 | 35 | 16.9086651 | 19.1569087 | 0 | 0 | 0 | 0 | 0 | 0 | 0 | 0 | 0 | 0 | 1 | 0 | 2 | 0 | 3 |
| ENSMUSG00000052593 | Adam17 | ENSMUST00000127974 | 28.6946841 | 27 | 20.4564431 | 23.7962705 | 0 | 0 | 0 | 0 | 0 | 0 | 0 | 0 | 0 | 0 | 1 | 0 | 7 | 0 | 8 |
| ENSMUSG00000011256 | Adam19 | ENSMUST00000011400 | 23.5563179 | 27 | 25.4145226 | 23.9565466 | 0 | 0 | 0 | 0 | 0 | 0 | 0 | 0 | 0 | 0 | 0 | 1 | 0 | 0 | 1 |
| ENSMUSG00000040537 | Adam22 | ENSMUST00000050166 | 26.8234923 | 31 | 20.6662457 | 21.8187559 | 0 | 0 | 0 | 1 | 0 | 0 | 0 | 0 | 1 | 2 | 0 | 2 | 20 | 2 | 26 |
| ENSMUSG00000025964 | Adam23 | ENSMUST00000087374 | 26.7679558 | 32 | 21.9060773 | 19.0607735 | 0 | 0 | 0 | 0 | 0 | 0 | 0 | 0 | 0 | 0 | 1 | 2 | 8 | 0 | 11 |
| ENSMUSG00000031555 | Adam9 | ENSMUST00000084035 | 31.3935681 | 35 | 16.0030628 | 17.7641654 | 0 | 0 | 0 | 0 | 0 | 0 | 0 | 0 | 0 | 0 | 0 | 0 | 3 | 0 | 3 |
| ENSMUSG00000022893 | Adamts1 | ENSMUST00000023610 | 32.0203304 | 29 | 15.9466328 | 22.8716645 | 1 | 0 | 0 | 0 | 0 | 0 | 0 | 0 | 0 | 0 | 0 | 0 | 0 | 0 | 1 |
| ENSMUSG00000022449 | Adamts20 | ENSMUST00000155907 | 29.5202952 | 34 | 17.5071751 | 18.7781878 | 1 | 0 | 0 | 0 | 0 | 0 | 0 | 0 | 0 | 0 | 0 | 0 | 1 | 3 | 5 |
| ENSMUSG00000030022 | Adamts9 | ENSMUST00000113438 | 29.2618163 | 32 | 19.118428 | 19.1715348 | 2 | 0 | 1 | 0 | 0 | 0 | 0 | 0 | 0 | 1 | 0 | 0 | 0 | 2 | 5 |
| ENSMUSG00000052551 | Adarb2 | ENSMUST00000064473 | 31.128876 | 28 | 20.494186 | 19.9854651 | 0 | 0 | 0 | 0 | 0 | 0 | 0 | 0 | 0 | 0 | 0 | 0 | 1 | 1 | 2 |
| ENSMUSG00000024256 | Adcyap1 | ENSMUST00000064775 | 28.2032401 | 34 | 17.9675994 | 20.1767305 | 0 | 0 | 0 | 0 | 0 | 0 | 0 | 0 | 0 | 0 | 0 | 1 | 0 | 0 | 1 |
| ENSMUSG00000033569 | Adgrb3 | ENSMUST00000126626 | 29.4797688 | 37 | 16.6184971 | 17.1965318 | 0 | 0 | 1 | 1 | 0 | 0 | 0 | 0 | 1 | 3 | 0 | 0 | 0 | 0 | 3 |
| ENSMUSG00000020629 | Adi1 | ENSMUST00000020957 | 25.3211009 | 31 | 22.5688073 | 20.6422018 | 0 | 0 | 0 | 0 | 0 | 0 | 0 | 0 | 0 | 0 | 0 | 1 | 0 | 0 | 1 |
| ENSMUSG00000057134 | Ado | ENSMUST00000075686 | 26.7831567 | 31 | 20.8822687 | 21.311945 | 0 | 0 | 0 | 0 | 0 | 0 | 0 | 0 | 0 | 0 | 4 | 1 | 22 | 0 | 27 |
| ENSMUSG00000045730 | Adrb2 | ENSMUST00000053640 | 21.9440353 | 43 | 14.5802651 | 20.6185567 | 0 | 0 | 0 | 0 | 0 | 0 | 0 | 0 | 0 | 0 | 3 | 2 | 14 | 0 | 19 |
| ENSMUSG00000030232 | Aebp2 | ENSMUST00000032359 | 29.1160809 | 33 | 17.0394036 | 21.0223642 | 0 | 1 | 0 | 1 | 0 | 0 | 0 | 0 | 1 | 2 | 0 | 4 | 0 | 4 | 11 |
| ENSMUSG00000029094 | Afap1 | ENSMUST00000064571 | 22.8680324 | 29 | 25.2024774 | 22.6536446 | 1 | 1 | 0 | 2 | 0 | 0 | 0 | 0 | 2 | 4 | 3 | 1 | 9 | 1 | 20 |
| ENSMUSG00000031189 | Aff2 | ENSMUST00000033532 | 24.2718447 | 38 | 18.4466019 | 18.9320388 | 0 | 0 | 0 | 0 | 0 | 0 | 0 | 0 | 0 | 0 | 1 | 0 | 1 | 0 | 2 |

| | | | | | | | | | | | | | | | | | | | | | |
|---|---|---|---|---|---|---|---|---|---|---|---|---|---|---|---|---|---|---|---|---|---|
| ENSMUSG00000038836 | Agbl3 | ENSMUST00000148834 | 33.2081142 | 26 | 21.387428 | 19.6343601 | 1 | 0 | 0 | 1 | 0 | 0 | 0 | 0 | 1 | 2 | 0 | 1 | 0 | 0 | 4 |
| ENSMUSG00000026159 | Agfg1 | ENSMUST00000189220 | 27.4493795 | 34 | 17.6028739 | 20.6237753 | 0 | 0 | 0 | 1 | 0 | 0 | 0 | 0 | 1 | 2 | 0 | 0 | 2 | 2 | 6 |
| ENSMUSG00000029722 | Agfg2 | ENSMUST00000031736 | 25.6508423 | 26 | 23.81317 | 24.4257274 | 0 | 0 | 0 | 1 | 0 | 0 | 0 | 0 | 1 | 2 | 0 | 0 | 9 | 0 | 11 |
| ENSMUSG00000029916 | Agk | ENSMUST00000200715 | 26.996587 | 32 | 19.6587031 | 21.5017065 | 0 | 0 | 0 | 0 | 0 | 0 | 0 | 0 | 0 | 0 | 0 | 0 | 1 | 0 | 1 |
| ENSMUSG00000036698 | Ago2 | ENSMUST00000044113 | 23.2194505 | 30 | 24.0057845 | 23.1561822 | 3 | 2 | 0 | 4 | 0 | 0 | 0 | 1 | 3 | 8 | 0 | 8 | 8 | 1 | 30 |
| ENSMUSG00000028842 | Ago3 | ENSMUST00000069097 | 28.5983362 | 31 | 20.0753414 | 20.2166065 | 2 | 1 | 0 | 4 | 0 | 0 | 1 | 2 | 1 | 8 | 1 | 2 | 14 | 5 | 33 |
| ENSMUSG00000031467 | Agpat5 | ENSMUST00000149565 | 26.6546074 | 32 | 20.5469249 | 20.9619068 | 1 | 0 | 0 | 1 | 0 | 0 | 0 | 0 | 1 | 2 | 3 | 23 | 10 | 6 | 45 |
| ENSMUSG00000042410 | Agps | ENSMUST00000047232 | 27.7746373 | 33 | 18.5227059 | 20.4258526 | 1 | 0 | 0 | 0 | 0 | 0 | 0 | 0 | 0 | 0 | 2 | 2 | 5 | 2 | 12 |
| ENSMUSG00000029772 | Ahcyl2 | ENSMUST00000115242 | 24.9387255 | 31 | 22.0894608 | 21.9669118 | 1 | 1 | 2 | 2 | 0 | 0 | 0 | 1 | 1 | 6 | 1 | 1 | 3 | 0 | 13 |
| ENSMUSG00000019256 | Ahr | ENSMUST00000116436 | 29.7605893 | 33 | 18.3793738 | 18.4162063 | 1 | 0 | 0 | 1 | 0 | 0 | 0 | 0 | 1 | 2 | 2 | 0 | 0 | 0 | 5 |
| ENSMUSG00000078307 | AI593442 | ENSMUST00000098768 | 28.6701209 | 33 | 19.5164076 | 18.6312608 | 0 | 0 | 0 | 1 | 0 | 0 | 0 | 0 | 1 | 2 | 2 | 0 | 23 | 0 | 27 |
| ENSMUSG00000010290 | AI597479 | ENSMUST00000010434 | 25.6561922 | 34 | 18.9279113 | 21.5526802 | 0 | 0 | 0 | 0 | 0 | 0 | 0 | 0 | 0 | 0 | 0 | 2 | 0 | 1 | 3 |
| ENSMUSG00000024970 | AI846148 | ENSMUST00000025924 | 23.2339956 | 27 | 23.6754967 | 25.6622517 | 0 | 0 | 0 | 0 | 0 | 0 | 0 | 0 | 0 | 0 | 0 | 1 | 0 | 2 | 3 |
| ENSMUSG00000056383 | AI987944 | ENSMUST00000071804 | 31.9943622 | 29 | 18.1818182 | 20.6483439 | 1 | 0 | 0 | 0 | 0 | 0 | 0 | 0 | 0 | 0 | 0 | 1 | 0 | 0 | 2 |
| ENSMUSG00000037860 | Aim2 | ENSMUST00000147604 | 32.2896282 | 27 | 16.8297456 | 23.9726027 | 0 | 0 | 0 | 1 | 0 | 0 | 0 | 0 | 1 | 2 | 0 | 0 | 0 | 0 | 2 |
| ENSMUSG00000091931 | AK010878 | ENSMUST00000173969 | 32.875 | 30 | 16.875 | 20 | 0 | 0 | 0 | 0 | 0 | 0 | 0 | 0 | 0 | 0 | 1 | 0 | 1 | 0 | 2 |
| ENSMUSG00000028527 | Ak4 | ENSMUST00000102780 | 23.6177885 | 30 | 21.6646635 | 24.8197115 | 0 | 0 | 0 | 0 | 0 | 0 | 0 | 0 | 0 | 0 | 0 | 0 | 4 | 0 | 4 |
| ENSMUSG00000047804 | Akap10 | ENSMUST00000102650 | 25.8045641 | 31 | 21.4160328 | 21.7671153 | 0 | 0 | 0 | 1 | 0 | 0 | 0 | 0 | 1 | 2 | 0 | 2 | 0 | 0 | 4 |
| ENSMUSG00000059708 | Akap17b | ENSMUST00000051906 | 28.958255 | 32 | 18.8792779 | 20.6468597 | 0 | 0 | 0 | 0 | 0 | 0 | 0 | 0 | 0 | 0 | 1 | 2 | 2 | 0 | 5 |
| ENSMUSG00000039166 | Akap7 | ENSMUST00000100012 | 28.6442406 | 33 | 18.6544343 | 19.6738022 | 0 | 0 | 0 | 0 | 0 | 0 | 0 | 0 | 0 | 0 | 1 | 0 | 0 | 1 | 2 |
| ENSMUSG00000024045 | Akap8 | ENSMUST00000002699 | 26.2492094 | 34 | 18.5958254 | 20.9993675 | 0 | 0 | 0 | 0 | 0 | 0 | 0 | 0 | 0 | 0 | 1 | 0 | 4 | 0 | 5 |
| ENSMUSG00000023075 | Akirin1 | ENSMUST00000102636 | 25.8823529 | 32 | 19.5206972 | 22.4836601 | 1 | 0 | 0 | 0 | 0 | 0 | 0 | 0 | 0 | 0 | 1 | 1 | 0 | 0 | 3 |
| ENSMUSG00000028291 | Akirin2 | ENSMUST00000084299 | 25.7686676 | 36 | 17.2767204 | 20.4978038 | 1 | 0 | 0 | 0 | 0 | 0 | 0 | 0 | 0 | 0 | 0 | 0 | 0 | 0 | 1 |
| ENSMUSG00000019699 | Akt3 | ENSMUST00000111160 | 26.161896 | 30 | 22.4068944 | 21.4835334 | 2 | 1 | 0 | 1 | 0 | 0 | 0 | 0 | 1 | 2 | 0 | 1 | 1 | 2 | 9 |
| ENSMUSG00000022636 | Alcam | ENSMUST00000023312 | 31.1028037 | 34 | 16.5607477 | 18.317757 | 0 | 0 | 0 | 0 | 0 | 0 | 0 | 0 | 0 | 0 | 0 | 13 | 5 | 0 | 18 |
| ENSMUSG00000010025 | Aldh3a2 | ENSMUST00000074127 | 26.7673301 | 27 | 23.8846946 | 21.9629375 | 0 | 0 | 0 | 0 | 0 | 0 | 0 | 0 | 0 | 0 | 1 | 0 | 0 | 0 | 1 |
| ENSMUSG00000075470 | Alg10b | ENSMUST00000100309 | 26.4367816 | 32 | 19.4482759 | 22.3908046 | 0 | 0 | 0 | 0 | 0 | 0 | 0 | 0 | 0 | 0 | 0 | 0 | 1 | 0 | 1 |
| ENSMUSG00000063362 | Alg11 | ENSMUST00000072572 | 28.6043165 | 31 | 21.352518 | 19.1366906 | 0 | 0 | 1 | 0 | 0 | 0 | 0 | 0 | 0 | 1 | 0 | 1 | 10 | 1 | 13 |
| ENSMUSG00000041718 | Alg13 | ENSMUST00000154827 | 29.0602933 | 28 | 21.9445953 | 20.8582292 | 0 | 0 | 0 | 1 | 0 | 0 | 0 | 0 | 1 | 2 | 0 | 0 | 0 | 0 | 2 |
| ENSMUSG00000039887 | Alg14 | ENSMUST00000039442 | 30.8103976 | 30 | 16.2844037 | 22.8593272 | 0 | 0 | 0 | 0 | 0 | 0 | 0 | 0 | 0 | 0 | 4 | 0 | 10 | 0 | 14 |
| ENSMUSG00000039740 | Alg2 | ENSMUST00000044148 | 27.6940904 | 30 | 21.6685979 | 21.0312862 | 1 | 0 | 0 | 0 | 0 | 0 | 0 | 0 | 0 | 0 | 0 | 0 | 2 | 0 | 3 |
| ENSMUSG00000073792 | Alg6 | ENSMUST00000097961 | 31.6091954 | 33 | 16.7385057 | 18.3908046 | 0 | 0 | 0 | 0 | 0 | 0 | 0 | 0 | 0 | 0 | 6 | 0 | 6 | 3 | 15 |
| ENSMUSG00000035704 | Alg8 | ENSMUST00000098300 | 28.1690141 | 31 | 18.3098592 | 22.7699531 | 1 | 1 | 0 | 0 | 0 | 0 | 0 | 0 | 0 | 0 | 0 | 0 | 0 | 0 | 2 |
| ENSMUSG00000079036 | Alkbh1; Nrp | ENSMUST00000162961 | 26.5344311 | 31 | 19.8353293 | 22.3053892 | 0 | 0 | 0 | 1 | 0 | 0 | 0 | 0 | 1 | 2 | 0 | 0 | 1 | 0 | 3 |
| ENSMUSG00000025701 | Alox5 | ENSMUST00000026795 | 23.9875389 | 27 | 27.1028037 | 22.2741433 | 0 | 0 | 1 | 0 | 0 | 0 | 0 | 0 | 0 | 1 | 0 | 0 | 0 | 0 | 1 |
| ENSMUSG00000025134 | Alyref | ENSMUST00000026125 | 23.682284 | 33 | 18.3382138 | 24.5973646 | 0 | 0 | 0 | 0 | 0 | 0 | 0 | 0 | 0 | 0 | 0 | 1 | 8 | 2 | 11 |
| ENSMUSG00000021986 | Amer2 | ENSMUST00000022561 | 28.99729 | 27 | 24.3902439 | 20.0542005 | 0 | 0 | 0 | 0 | 0 | 0 | 0 | 0 | 0 | 0 | 0 | 1 | 0 | 0 | 1 |
| ENSMUSG00000042225 | Ammecr1 | ENSMUST00000041317 | 30.1344883 | 33 | 17.7114201 | 19.0107135 | 0 | 0 | 0 | 0 | 0 | 0 | 0 | 0 | 0 | 0 | 0 | 3 | 5 | 0 | 8 |
| ENSMUSG00000041915 | Ammecr1l | ENSMUST00000115808 | 25.482509 | 30 | 21.2002413 | 23.4016888 | 0 | 0 | 0 | 0 | 0 | 0 | 0 | 0 | 0 | 0 | 0 | 0 | 11 | 1 | 12 |
| ENSMUSG00000068250 | Amn1 | ENSMUST00000111535 | 26.5139116 | 39 | 17.1849427 | 17.3486088 | 1 | 1 | 3 | 2 | 0 | 0 | 0 | 1 | 1 | 7 | 3 | 0 | 5 | 0 | 17 |
| ENSMUSG00000032531 | Amotl2 | ENSMUST00000142011 | 23.9736442 | 27 | 24.3791181 | 24.4298023 | 0 | 0 | 0 | 0 | 0 | 0 | 0 | 0 | 0 | 0 | 1 | 0 | 3 | 0 | 4 |
| ENSMUSG00000036977 | Anapc10 | ENSMUST00000048147 | 29.6433534 | 31 | 19.7776748 | 19.3608152 | 0 | 0 | 0 | 0 | 0 | 0 | 0 | 0 | 0 | 0 | 0 | 0 | 0 | 1 | 1 |
| ENSMUSG00000020790 | Ankfy1 | ENSMUST00000127610 | 25.1934652 | 27 | 24.2132416 | 23.422184 | 0 | 0 | 0 | 0 | 0 | 0 | 0 | 0 | 0 | 0 | 1 | 1 | 1 | 0 | 3 |
| ENSMUSG00000021661 | Ankra2 | ENSMUST00000123924 | 28.3365571 | 34 | 17.794971 | 20.0193424 | 0 | 0 | 0 | 0 | 0 | 0 | 0 | 0 | 0 | 0 | 0 | 1 | 0 | 0 | 1 |
| ENSMUSG00000034647 | Ankrd12 | ENSMUST00000038116 | 32.3651452 | 32 | 16.1825726 | 19.4643531 | 1 | 0 | 0 | 1 | 0 | 0 | 0 | 0 | 1 | 2 | 1 | 1 | 2 | 0 | 7 |
| ENSMUSG00000039988 | Ankrd13c | ENSMUST00000199727 | 30.0251256 | 31 | 19.0536013 | 19.7654941 | 1 | 1 | 0 | 2 | 0 | 0 | 0 | 1 | 1 | 4 | 0 | 4 | 5 | 1 | 16 |
| ENSMUSG00000020864 | Ankrd40 | ENSMUST00000107818 | 25.5368814 | 28 | 23.2026144 | 23.2492997 | 0 | 0 | 0 | 0 | 0 | 0 | 0 | 0 | 0 | 0 | 0 | 3 | 0 | 0 | 3 |
| ENSMUSG00000052331 | Ankrd44 | ENSMUST00000179030 | 26.2512768 | 34 | 20.1225741 | 19.7139939 | 1 | 0 | 0 | 0 | 0 | 0 | 0 | 0 | 0 | 0 | 0 | 10 | 1 | 0 | 12 |
| ENSMUSG00000044835 | Ankrd45 | ENSMUST00000111608 | 25.5160142 | 31 | 23.0960854 | 20.1067616 | 0 | 0 | 0 | 0 | 0 | 0 | 0 | 0 | 0 | 0 | 0 | 4 | 0 | 0 | 4 |
| ENSMUSG00000058589 | Anks1b | ENSMUST00000182356 | 26.1862917 | 26 | 25.4353731 | 22.1281355 | 0 | 0 | 0 | 1 | 0 | 0 | 0 | 0 | 1 | 2 | 1 | 4 | 3 | 0 | 10 |
| ENSMUSG00000036777 | Anln | ENSMUST00000040912 | 29.1938998 | 33 | 16.9389978 | 20.4793028 | 1 | 0 | 1 | 0 | 0 | 0 | 0 | 0 | 0 | 1 | 0 | 0 | 0 | 0 | 2 |
| ENSMUSG00000055489 | Ano5 | ENSMUST00000043944 | 29.4819079 | 31 | 20.2097039 | 18.9967105 | 0 | 0 | 0 | 0 | 0 | 0 | 0 | 0 | 0 | 0 | 1 | 5 | 9 | 1 | 16 |
| ENSMUSG00000028333 | Anp32b | ENSMUST00000102926 | 27.0377734 | 32 | 21.6699801 | 18.8866799 | 0 | 0 | 0 | 0 | 0 | 0 | 0 | 0 | 0 | 0 | 0 | 0 | 0 | 1 | 1 |
| ENSMUSG00000015749 | Anp32e | ENSMUST00000165307 | 24.3925234 | 34 | 18.6915888 | 22.4766355 | 0 | 0 | 0 | 0 | 0 | 0 | 0 | 0 | 0 | 0 | 0 | 5 | 4 | 0 | 9 |
| ENSMUSG00000074238 | Ap1ar | ENSMUST00000200409 | 27.0215633 | 34 | 19.2048518 | 20.0134771 | 0 | 0 | 0 | 0 | 0 | 0 | 0 | 0 | 0 | 0 | 0 | 0 | 0 | 2 | 2 |
| ENSMUSG00000031731 | Ap1g1 | ENSMUST00000093157 | 25.2930905 | 32 | 20.3791469 | 22.4494887 | 0 | 0 | 0 | 0 | 0 | 0 | 0 | 0 | 0 | 0 | 1 | 3 | 4 | 0 | 8 |
| ENSMUSG00000054702 | Ap1s3 | ENSMUST00000162342 | 26.4949804 | 36 | 20.2095155 | 17.6342209 | 1 | 1 | 0 | 1 | 0 | 0 | 0 | 0 | 1 | 2 | 0 | 40 | 6 | 1 | 51 |
| ENSMUSG00000021824 | Ap3m1 | ENSMUST00000154460 | 30.0716915 | 29 | 20.6549118 | 20.1898857 | 0 | 0 | 0 | 0 | 0 | 0 | 0 | 0 | 0 | 0 | 2 | 0 | 2 | 0 | 4 |
| ENSMUSG00000024480 | Ap3s1 | ENSMUST00000025357 | 28.0898876 | 38 | 16.6042447 | 17.1036205 | 0 | 0 | 0 | 0 | 0 | 0 | 0 | 0 | 0 | 0 | 0 | 0 | 3 | 1 | 4 |
| ENSMUSG00000036291 | Ap5m1 | ENSMUST00000037473 | 28.5816619 | 35 | 19.4842407 | 17.2636103 | 1 | 0 | 1 | 0 | 0 | 0 | 0 | 0 | 0 | 1 | 0 | 0 | 2 | 0 | 4 |
| ENSMUSG00000032375 | Aph1b | ENSMUST00000168309 | 27.7550091 | 30 | 20.3324226 | 21.4253188 | 0 | 0 | 0 | 0 | 0 | 0 | 0 | 0 | 0 | 0 | 1 | 2 | 0 | 0 | 3 |
| ENSMUSG00000053040 | Aph1c | ENSMUST00000169282 | 27.4883721 | 30 | 21.6744186 | 21.1860465 | 0 | 0 | 0 | 0 | 0 | 0 | 0 | 0 | 0 | 0 | 1 | 4 | 0 | 0 | 5 |
| ENSMUSG00000033096 | Apmap | ENSMUST00000046399 | 24.5828699 | 31 | 20.1334816 | 24.0266963 | 0 | 0 | 0 | 0 | 0 | 0 | 0 | 0 | 0 | 0 | 0 | 1 | 0 | 0 | 1 |
| ENSMUSG00000090698 | Apold1 | ENSMUST00000167323 | 27.5587415 | 31 | 19.7530864 | 21.9832736 | 2 | 1 | 0 | 2 | 0 | 0 | 0 | 1 | 1 | 4 | 1 | 2 | 11 | 0 | 21 |
| ENSMUSG00000025525 | Apool | ENSMUST00000026599 | 35.0951374 | 33 | 12.6849894 | 19.6617336 | 0 | 0 | 0 | 0 | 0 | 0 | 0 | 0 | 0 | 0 | 0 | 0 | 0 | 1 | 1 |
| ENSMUSG00000018481 | Appbp2 | ENSMUST00000018625 | 27.8284672 | 32 | 19.5939781 | 20.4607664 | 0 | 0 | 0 | 1 | 0 | 0 | 0 | 0 | 1 | 2 | 1 | 2 | 6 | 3 | 14 |
| ENSMUSG00000040760 | Appl1 | ENSMUST00000036570 | 26.8565401 | 35 | 18.3333333 | 20.0843882 | 0 | 0 | 0 | 0 | 0 | 0 | 0 | 0 | 0 | 0 | 1 | 3 | 2 | 0 | 6 |
| ENSMUSG00000048076 | Arf1 | ENSMUST00000163300 | 22.8163993 | 29 | 23.885918 | 24.2424242 | 0 | 0 | 0 | 0 | 0 | 0 | 0 | 0 | 0 | 0 | 0 | 1 | 0 | 0 | 1 |
| ENSMUSG00000020440 | Arf5 | ENSMUST00000020717 | 12.8378378 | 31 | 26.8018018 | 29.2792793 | 0 | 0 | 0 | 0 | 0 | 0 | 0 | 0 | 0 | 0 | 0 | 0 | 4 | 0 | 4 |
| ENSMUSG00000044147 | Arf6 | ENSMUST00000050063 | 30.1730625 | 32 | 17.0805117 | 20.4288939 | 0 | 0 | 0 | 0 | 0 | 0 | 0 | 0 | 0 | 0 | 0 | 0 | 8 | 2 | 10 |
| ENSMUSG00000074513 | Arfip1 | ENSMUST00000098990 | 27.6932441 | 36 | 15.0334753 | 20.8764455 | 0 | 0 | 0 | 1 | 0 | 0 | 0 | 0 | 1 | 2 | 0 | 0 | 0 | 1 | 3 |
| ENSMUSG00000040459 | Arglu1 | ENSMUST00000048545 | 23.7037037 | 38 | 17.1851852 | 20.7407407 | 0 | 0 | 0 | 0 | 0 | 0 | 0 | 0 | 0 | 0 | 0 | 0 | 12 | 1 | 13 |
| ENSMUSG00000041219 | Arhgap11a | ENSMUST00000110948 | 26.0231924 | 28 | 17.8035471 | 28.0013643 | 1 | 1 | 0 | 0 | 0 | 0 | 0 | 0 | 0 | 0 | 0 | 0 | 0 | 0 | 2 |
| ENSMUSG00000041225 | Arhgap12 | ENSMUST00000182066 | 27.5246595 | 36 | 16.9093471 | 19.8684829 | 0 | 0 | 0 | 0 | 0 | 0 | 0 | 0 | 0 | 0 | 0 | 0 | 2 | 0 | 2 |
| ENSMUSG00000039031 | Arhgap18 | ENSMUST00000039557 | 31.056701 | 32 | 17.9123711 | 19.2654639 | 0 | 0 | 0 | 0 | 0 | 0 | 0 | 0 | 0 | 0 | 0 | 2 | 0 | 0 | 2 |
| ENSMUSG00000025154 | Arhgap19 | ENSMUST00000026150 | 22.1789883 | 32 | 20.5669817 | 25.6253474 | 2 | 1 | 0 | 1 | 0 | 0 | 0 | 0 | 1 | 2 | 3 | 0 | 0 | 0 | 8 |
| ENSMUSG00000024043 | Arhgap28 | ENSMUST00000024840 | 24.7187293 | 29 | 23.2627399 | 23.1634679 | 0 | 0 | 0 | 0 | 0 | 0 | 0 | 0 | 0 | 0 | 3 | 11 | 0 | 2 | 16 |
| ENSMUSG00000039831 | Arhgap29 | ENSMUST00000037958 | 26.3779528 | 37 | 16.2542182 | 20.303712 | 0 | 1 | 0 | 0 | 0 | 0 | 0 | 0 | 0 | 0 | 0 | 0 | 3 | 0 | 4 |
| ENSMUSG00000041444 | Arhgap32 | ENSMUST00000174641 | 27.5280899 | 30 | 20.0465881 | 22.4445053 | 0 | 0 | 0 | 0 | 0 | 0 | 0 | 0 | 0 | 0 | 1 | 10 | 1 | 2 | 14 |
| ENSMUSG00000036198 | Arhgap36 | ENSMUST00000114904 | 28.7398005 | 28 | 23.118767 | 20.5802357 | 0 | 0 | 0 | 0 | 0 | 0 | 0 | 0 | 0 | 0 | 0 | 2 | 0 | 0 | 2 |
| ENSMUSG00000050730 | Arhgap42 | ENSMUST00000093893 | 29.8347613 | 33 | 18.0538556 | 19.2778458 | 3 | 1 | 0 | 3 | 0 | 0 | 0 | 1 | 2 | 6 | 0 | 0 | 2 | 0 | 12 |
| ENSMUSG00000004661 | Arid3b | ENSMUST00000171444 | 22.3611111 | 26 | 27.8240741 | 23.8425926 | 0 | 0 | 0 | 0 | 0 | 0 | 0 | 0 | 0 | 0 | 0 | 1 | 0 | 0 | 1 |
| ENSMUSG00000039219 | Arid4b | ENSMUST00000149579 | 27.3275862 | 30 | 19.2241379 | 23.8362069 | 2 | 1 | 0 | 1 | 0 | 0 | 0 | 1 | 0 | 2 | 0 | 1 | 0 | 0 | 6 |
| ENSMUSG00000025234 | Arih1 | ENSMUST00000171975 | 27.5218427 | 33 | 19.7974583 | 19.3208896 | 1 | 0 | 0 | 0 | 0 | 0 | 0 | 0 | 0 | 0 | 0 | 1 | 5 | 3 | 10 |
| ENSMUSG00000060904 | Arl1 | ENSMUST00000116234 | 30.9317963 | 32 | 17.4831892 | 20.0768492 | 0 | 0 | 0 | 0 | 0 | 0 | 0 | 0 | 0 | 0 | 0 | 1 | 0 | 0 | 1 |
| ENSMUSG00000022911 | Arl13b | ENSMUST00000089289 | 30.816641 | 34 | 17.4627632 | 17.5141243 | 0 | 0 | 0 | 1 | 0 | 0 | 0 | 0 | 1 | 2 | 0 | 0 | 0 | 0 | 2 |
| ENSMUSG00000027122 | Arl14ep | ENSMUST00000028536 | 29.750983 | 32 | 17.8243775 | 20.1834862 | 0 | 0 | 0 | 0 | 0 | 0 | 0 | 0 | 0 | 0 | 0 | 0 | 11 | 0 | 11 |
| ENSMUSG00000042348 | Arl15 | ENSMUST00000091201 | 27.4088292 | 32 | 20.268714 | 20.4990403 | 0 | 0 | 0 | 1 | 0 | 0 | 0 | 0 | 1 | 2 | 0 | 0 | 2 | 1 | 5 |
| ENSMUSG00000047446 | Arl4a | ENSMUST00000136441 | 26.0169492 | 30 | 23.7288136 | 20.4237288 | 0 | 0 | 0 | 0 | 0 | 0 | 0 | 0 | 0 | 0 | 0 | 40 | 3 | 0 | 43 |
| ENSMUSG00000036093 | Arl5a | ENSMUST00000036541 | 30.1601624 | 32 | 18.3622829 | 19.1518159 | 0 | 0 | 0 | 1 | 0 | 0 | 0 | 0 | 1 | 2 | 0 | 0 | 6 | 2 | 10 |
| ENSMUSG00000017418 | Arl5b | ENSMUST00000069870 | 27.2756827 | 35 | 18.0429129 | 19.8472042 | 3 | 2 | 0 | 3 | 0 | 0 | 0 | 0 | 3 | 6 | 3 | 1 | 4 | 2 | 21 |
| ENSMUSG00000030654 | Arl6ip1 | ENSMUST00000032888 | 23.5546039 | 33 | 20.6995004 | 22.5553176 | 0 | 0 | 0 | 0 | 0 | 0 | 0 | 0 | 0 | 0 | 2 | 1 | 1 | 0 | 4 |

| | | | | | | | | | | | | | | | | | | | | | |
|---|---|---|---|---|---|---|---|---|---|---|---|---|---|---|---|---|---|---|---|---|---|
| ENSMUSG00000026960 | Arl6ip6 | ENSMUST00000028336 | 29.8034934 | 39 | 15.0655022 | 16.430131 | 2 | 0 | 0 | 1 | 0 | 0 | 0 | 0 | 1 | 2 | 0 | 1 | 0 | 0 | 5 |
| ENSMUSG00000026426 | Arl8a | ENSMUST00000027684 | 20.741483 | 30 | 28.3567134 | 20.741483 | 0 | 0 | 0 | 1 | 0 | 0 | 0 | 0 | 1 | 2 | 4 | 1 | 6 | 0 | 13 |
| ENSMUSG00000030105 | Arl8b | ENSMUST00000032196 | 26.7042405 | 32 | 20.5582394 | 21.0144928 | 0 | 0 | 0 | 0 | 0 | 0 | 0 | 0 | 0 | 0 | 0 | 1 | 3 | 0 | 4 |
| ENSMUSG00000027599 | Armc1 | ENSMUST00000029125 | 27.7946768 | 33 | 18.8212928 | 20.3422053 | 0 | 1 | 0 | 1 | 0 | 0 | 0 | 0 | 1 | 2 | 0 | 0 | 0 | 0 | 3 |
| ENSMUSG00000038525 | Armc10 | ENSMUST00000095495 | 27.7806518 | 30 | 20.1241593 | 22.2452147 | 0 | 0 | 0 | 0 | 0 | 0 | 0 | 0 | 0 | 0 | 0 | 0 | 1 | 2 | 3 |
| ENSMUSG00000032468 | Armc8 | ENSMUST00000035043 | 27.4030354 | 29 | 19.6037099 | 24.0303541 | 0 | 0 | 0 | 0 | 0 | 0 | 0 | 0 | 0 | 0 | 0 | 2 | 2 | 0 | 4 |
| ENSMUSG00000062590 | Armc9 | ENSMUST00000027434 | 24.980784 | 26 | 25.7109915 | 23.443505 | 0 | 0 | 0 | 0 | 0 | 0 | 0 | 0 | 0 | 0 | 0 | 2 | 2 | 0 | 4 |
| ENSMUSG00000061759 | Armt1 | ENSMUST00000118544 | 23.7614463 | 31 | 20.5212491 | 24.395398 | 0 | 0 | 0 | 0 | 0 | 0 | 0 | 0 | 0 | 0 | 2 | 0 | 12 | 0 | 14 |
| ENSMUSG00000008475 | Arpc5 | ENSMUST00000077755 | 28.5490196 | 31 | 20.2352941 | 19.7647059 | 0 | 0 | 0 | 0 | 0 | 0 | 0 | 0 | 0 | 0 | 0 | 0 | 2 | 1 | 3 |
| ENSMUSG00000007656 | Arpp19 | ENSMUST00000007800 | 28.3067275 | 34 | 19.3842645 | 18.4720639 | 0 | 0 | 0 | 1 | 0 | 0 | 0 | 0 | 1 | 2 | 1 | 0 | 0 | 0 | 3 |
| ENSMUSG00000074794 | Arrdc3 | ENSMUST00000159690 | 28.7043665 | 33 | 17.9312813 | 20.5082319 | 0 | 1 | 0 | 0 | 0 | 0 | 0 | 0 | 0 | 0 | 2 | 0 | 4 | 0 | 7 |
| ENSMUSG00000042659 | Arrdc4 | ENSMUST00000048068 | 26.9736842 | 30 | 19.8601974 | 23.4786184 | 0 | 0 | 0 | 0 | 0 | 0 | 0 | 0 | 0 | 0 | 0 | 2 | 1 | 0 | 3 |
| ENSMUSG00000021592 | Arsk | ENSMUST00000120573 | 29.9345692 | 36 | 18.5387132 | 15.2671756 | 0 | 0 | 0 | 0 | 0 | 0 | 0 | 0 | 0 | 0 | 0 | 0 | 2 | 0 | 2 |
| ENSMUSG00000035277 | Arx | ENSMUST00000046565 | 28.3018868 | 28 | 24.1090147 | 19.8113208 | 1 | 0 | 0 | 0 | 0 | 0 | 0 | 0 | 0 | 0 | 0 | 1 | 0 | 0 | 2 |
| ENSMUSG00000048040 | Arxes2 | ENSMUST00000058119 | 27.9859485 | 37 | 16.5105386 | 18.735363 | 0 | 0 | 0 | 0 | 0 | 0 | 0 | 0 | 0 | 0 | 0 | 0 | 2 | 0 | 2 |
| ENSMUSG00000031591 | Asah1 | ENSMUST00000034000 | 28.0200143 | 32 | 19.8713367 | 20.586133 | 0 | 0 | 0 | 0 | 0 | 0 | 0 | 0 | 0 | 0 | 1 | 0 | 1 | 1 | 3 |
| ENSMUSG00000052632 | Asap2 | ENSMUST00000064595 | 25.7250946 | 31 | 22.9928541 | 19.9663724 | 0 | 0 | 0 | 0 | 0 | 0 | 0 | 0 | 0 | 0 | 0 | 30 | 0 | 0 | 30 |
| ENSMUSG00000020305 | Asb3 | ENSMUST00000137306 | 29.1216703 | 33 | 18.3945284 | 19.150468 | 0 | 0 | 0 | 0 | 0 | 0 | 0 | 0 | 0 | 0 | 0 | 0 | 0 | 2 | 2 |
| ENSMUSG00000042607 | Asb4 | ENSMUST00000043294 | 29.5696203 | 32 | 16.9113924 | 21.6202532 | 0 | 0 | 0 | 0 | 0 | 0 | 0 | 0 | 0 | 0 | 0 | 2 | 2 | 0 | 4 |
| ENSMUSG00000031519 | Asb5 | ENSMUST00000033918 | 32.6315789 | 40 | 11.7894737 | 15.3684211 | 0 | 0 | 0 | 0 | 0 | 0 | 0 | 0 | 0 | 0 | 0 | 0 | 0 | 1 | 1 |
| ENSMUSG00000020052 | Ascl1 | ENSMUST00000020243 | 30.2555647 | 27 | 23.9901072 | 18.6314922 | 0 | 0 | 0 | 0 | 0 | 0 | 0 | 0 | 0 | 0 | 0 | 1 | 0 | 0 | 1 |
| ENSMUSG00000028207 | Asph | ENSMUST00000078139 | 28.5104364 | 32 | 18.9990512 | 20.683112 | 1 | 0 | 0 | 0 | 0 | 0 | 0 | 0 | 0 | 0 | 0 | 0 | 2 | 2 | 5 |
| ENSMUSG00000040250 | Asun | ENSMUST00000032427 | 32.4840764 | 37 | 13.3757962 | 17.1974522 | 0 | 0 | 0 | 0 | 0 | 0 | 0 | 0 | 0 | 0 | 0 | 0 | 0 | 1 | 1 |
| ENSMUSG00000037486 | Asxl2 | ENSMUST00000092003 | 29.041764 | 31 | 19.967483 | 20.1808759 | 0 | 0 | 1 | 1 | 0 | 0 | 0 | 0 | 1 | 3 | 4 | 0 | 47 | 5 | 59 |
| ENSMUSG00000013662 | Atad1 | ENSMUST00000070210 | 28.2569899 | 36 | 15.8239143 | 19.5716835 | 0 | 0 | 0 | 1 | 0 | 0 | 0 | 0 | 1 | 2 | 0 | 1 | 0 | 0 | 3 |
| ENSMUSG00000022360 | Atad2 | ENSMUST00000038194 | 29.0540541 | 37 | 15.2702703 | 18.2432432 | 0 | 0 | 0 | 0 | 0 | 0 | 0 | 0 | 0 | 0 | 0 | 0 | 2 | 0 | 2 |
| ENSMUSG00000030850 | Ate1 | ENSMUST00000035458 | 25.560686 | 31 | 20.3496042 | 23.0540897 | 0 | 0 | 1 | 0 | 0 | 0 | 0 | 0 | 0 | 1 | 0 | 0 | 1 | 0 | 2 |
| ENSMUSG00000023027 | Atf1 | ENSMUST00000023769 | 27.8599222 | 37 | 13.229572 | 22.2568093 | 0 | 0 | 0 | 0 | 0 | 0 | 0 | 0 | 0 | 0 | 0 | 0 | 3 | 0 | 3 |
| ENSMUSG00000026663 | Atf6 | ENSMUST00000027974 | 27.7041306 | 28 | 22.1709894 | 22.4207493 | 0 | 0 | 0 | 1 | 0 | 0 | 0 | 0 | 1 | 2 | 0 | 2 | 0 | 0 | 4 |
| ENSMUSG00000099083 | Atf7 | ENSMUST00000184616 | 22.435094 | 32 | 25.4610564 | 20.1253357 | 0 | 0 | 0 | 0 | 0 | 0 | 0 | 0 | 0 | 0 | 1 | 20 | 5 | 0 | 26 |
| ENSMUSG00000030213 | Atf7ip | ENSMUST00000032335 | 25.069287 | 31 | 19.8790627 | 23.7087428 | 0 | 0 | 0 | 0 | 0 | 0 | 0 | 0 | 0 | 0 | 3 | 0 | 2 | 1 | 6 |
| ENSMUSG00000021619 | Atg10 | ENSMUST00000022119 | 32.7491785 | 31 | 16.6484118 | 19.2771084 | 0 | 0 | 0 | 0 | 0 | 0 | 0 | 0 | 0 | 0 | 0 | 0 | 0 | 4 | 4 |
| ENSMUSG00000032905 | Atg12 | ENSMUST00000035648 | 27.0536153 | 30 | 19.724545 | 22.8726021 | 0 | 0 | 0 | 0 | 0 | 0 | 0 | 0 | 0 | 0 | 0 | 0 | 6 | 0 | 6 |
| ENSMUSG00000022663 | Atg3 | ENSMUST00000023343 | 26.5997491 | 37 | 18.5696361 | 17.6913425 | 0 | 0 | 0 | 0 | 0 | 0 | 0 | 0 | 0 | 0 | 0 | 0 | 3 | 0 | 3 |
| ENSMUSG00000028550 | Atg4c | ENSMUST00000030279 | 26.1651676 | 36 | 21.3409648 | 16.6802944 | 1 | 0 | 0 | 0 | 0 | 0 | 0 | 0 | 0 | 0 | 0 | 33 | 0 | 0 | 34 |
| ENSMUSG00000038160 | Atg5 | ENSMUST00000039286 | 27.8450363 | 32 | 19.37046 | 20.7425343 | 0 | 0 | 0 | 0 | 0 | 0 | 0 | 0 | 0 | 0 | 0 | 0 | 2 | 1 | 3 |
| ENSMUSG00000030314 | Atg7 | ENSMUST00000032457 | 26.7756128 | 21 | 29.0383407 | 22.9415462 | 0 | 0 | 0 | 0 | 0 | 0 | 0 | 0 | 0 | 0 | 0 | 2 | 0 | 0 | 2 |
| ENSMUSG00000059811 | Atl2 | ENSMUST00000068282 | 27.3344652 | 36 | 16.581777 | 20.147142 | 0 | 0 | 0 | 0 | 0 | 0 | 0 | 0 | 0 | 0 | 1 | 2 | 0 | 0 | 3 |
| ENSMUSG00000024759 | Atl3 | ENSMUST00000025668 | 26.2685914 | 34 | 19.3350831 | 20.8005249 | 2 | 0 | 0 | 1 | 0 | 0 | 0 | 0 | 1 | 2 | 0 | 4 | 0 | 1 | 9 |
| ENSMUSG00000037400 | Atp11b | ENSMUST00000198599 | 27.8797997 | 35 | 18.6143573 | 18.5308848 | 1 | 0 | 0 | 1 | 0 | 0 | 0 | 0 | 1 | 2 | 0 | 1 | 0 | 0 | 4 |
| ENSMUSG00000062949 | Atp11c | ENSMUST00000154051 | 32.3095333 | 35 | 15.5963303 | 17.351416 | 0 | 0 | 0 | 0 | 0 | 0 | 0 | 0 | 0 | 0 | 0 | 1 | 0 | 3 | 4 |
| ENSMUSG00000022533 | Atp13a3 | ENSMUST00000100013 | 29.4117647 | 36 | 17.1898811 | 17.5861018 | 2 | 2 | 0 | 1 | 0 | 0 | 0 | 0 | 1 | 2 | 0 | 5 | 1 | 0 | 12 |
| ENSMUSG00000032412 | Atp1b3 | ENSMUST00000034983 | 27.2727273 | 32 | 17.9516686 | 23.2451093 | 0 | 0 | 0 | 0 | 0 | 0 | 0 | 0 | 0 | 0 | 0 | 0 | 1 | 0 | 1 |
| ENSMUSG00000029467 | Atp2a2 | ENSMUST00000179939 | 28.195122 | 33 | 19.1707317 | 19.3658537 | 0 | 1 | 0 | 2 | 0 | 0 | 0 | 1 | 1 | 4 | 0 | 0 | 1 | 0 | 6 |
| ENSMUSG00000032570 | Atp2c1 | ENSMUST00000112558 | 29.4724771 | 36 | 17.0489297 | 17.5458716 | 0 | 1 | 0 | 1 | 0 | 0 | 0 | 0 | 1 | 2 | 0 | 0 | 6 | 1 | 10 |
| ENSMUSG00000006057 | Atp5g1 | ENSMUST00000090541 | 26.1759172 | 28 | 22.4129821 | 23.3301976 | 0 | 0 | 0 | 1 | 0 | 0 | 0 | 0 | 1 | 2 | 1 | 1 | 2 | 2 | 8 |
| ENSMUSG00000054894 | Atp5s | ENSMUST00000021372 | 33.8181818 | 31 | 18.9090909 | 16 | 0 | 0 | 0 | 0 | 0 | 0 | 0 | 0 | 0 | 0 | 1 | 0 | 0 | 1 | 2 |
| ENSMUSG00000015575 | Atp6v0e | ENSMUST00000167352 | 24.3707551 | 32 | 22.0135837 | 21.6939672 | 4 | 1 | 0 | 2 | 0 | 0 | 0 | 0 | 2 | 4 | 2 | 1 | 5 | 1 | 18 |
| ENSMUSG00000052459 | Atp6v1a | ENSMUST00000114666 | 29.1790307 | 33 | 18.5459941 | 19.4362018 | 0 | 0 | 0 | 0 | 0 | 0 | 0 | 0 | 0 | 0 | 1 | 0 | 1 | 0 | 2 |
| ENSMUSG00000021114 | Atp6v1d | ENSMUST00000021536 | 25.2083333 | 34 | 21.875 | 18.75 | 0 | 0 | 1 | 0 | 0 | 0 | 0 | 0 | 0 | 1 | 0 | 0 | 0 | 0 | 1 |
| ENSMUSG00000004285 | Atp6v1f | ENSMUST00000004396 | 18.0327869 | 32 | 28.6885246 | 21.7213115 | 0 | 0 | 0 | 0 | 0 | 0 | 0 | 0 | 0 | 0 | 0 | 1 | 0 | 0 | 1 |
| ENSMUSG00000033793 | Atp6v1h | ENSMUST00000044369 | 29.3956044 | 40 | 14.2857143 | 16.2087912 | 0 | 0 | 1 | 0 | 0 | 0 | 0 | 0 | 0 | 1 | 1 | 0 | 0 | 0 | 2 |
| ENSMUSG00000006567 | Atp7b | ENSMUST00000006742 | 23.5798499 | 27 | 25.1339764 | 24.2229368 | 1 | 0 | 0 | 1 | 0 | 0 | 0 | 0 | 1 | 2 | 0 | 0 | 9 | 0 | 12 |
| ENSMUSG00000037685 | Atp8a1 | ENSMUST00000037380 | 26.7100247 | 33 | 20.0044853 | 20.4081633 | 2 | 0 | 0 | 0 | 0 | 0 | 0 | 0 | 0 | 0 | 2 | 0 | 11 | 2 | 17 |
| ENSMUSG00000028710 | Atpaf1 | ENSMUST00000184179 | 27.6480174 | 31 | 21.890277 | 19.934818 | 1 | 0 | 0 | 0 | 0 | 0 | 0 | 0 | 0 | 0 | 0 | 0 | 0 | 0 | 1 |
| ENSMUSG00000027312 | Atrn | ENSMUST00000028781 | 27.2313297 | 27 | 23.5655738 | 22.4499089 | 0 | 0 | 0 | 1 | 0 | 0 | 0 | 0 | 1 | 2 | 0 | 5 | 4 | 1 | 12 |
| ENSMUSG00000069895 | Atxn1l | ENSMUST00000093162 | 23.6018772 | 30 | 22.2330856 | 24.520923 | 0 | 0 | 0 | 0 | 0 | 0 | 0 | 0 | 0 | 0 | 0 | 2 | 0 | 0 | 2 |
| ENSMUSG00000021189 | Atxn3 | ENSMUST00000021606 | 27.8887304 | 33 | 17.831669 | 20.851165 | 0 | 0 | 0 | 1 | 0 | 0 | 0 | 0 | 1 | 2 | 0 | 1 | 4 | 6 | 13 |
| ENSMUSG00000021738 | Atxn7 | ENSMUST00000022257 | 28.8267875 | 32 | 16.9687815 | 22.4320242 | 2 | 1 | 0 | 1 | 0 | 0 | 0 | 0 | 1 | 2 | 1 | 0 | 2 | 0 | 8 |
| ENSMUSG00000020564 | Atxn7l1 | ENSMUST00000125192 | 25.6084272 | 30 | 21.6491101 | 23.0657465 | 0 | 0 | 0 | 0 | 0 | 0 | 0 | 0 | 0 | 0 | 0 | 2 | 7 | 1 | 10 |
| ENSMUSG00000074748 | Atxn7l3b | ENSMUST00000099276 | 26.042014 | 31 | 20.9736579 | 22.2074025 | 0 | 0 | 0 | 0 | 0 | 0 | 0 | 0 | 0 | 0 | 0 | 1 | 1 | 0 | 2 |
| ENSMUSG00000021460 | Auh | ENSMUST00000110031 | 23.1788079 | 29 | 24.3193525 | 23.8410596 | 0 | 0 | 0 | 0 | 0 | 0 | 0 | 0 | 0 | 0 | 2 | 0 | 15 | 0 | 17 |
| ENSMUSG00000029787 | Avl9 | ENSMUST00000031805 | 29.1491713 | 31 | 19.1160221 | 20.2651934 | 0 | 0 | 0 | 0 | 0 | 0 | 0 | 0 | 0 | 0 | 0 | 0 | 0 | 2 | 2 |
| ENSMUSG00000074166 | AW146154 | ENSMUST00000098509 | 29.858156 | 29 | 19.2907801 | 21.5602837 | 0 | 0 | 0 | 0 | 0 | 0 | 0 | 0 | 0 | 0 | 0 | 1 | 0 | 0 | 1 |
| ENSMUSG00000041935 | AW549877 | ENSMUST00000046633 | 27.3317308 | 36 | 20.2163462 | 16.7067308 | 1 | 0 | 0 | 0 | 0 | 0 | 0 | 0 | 0 | 0 | 0 | 0 | 1 | 0 | 2 |
| ENSMUSG00000033632 | AW554918 | ENSMUST00000097643 | 27.2 | 30 | 22.2 | 20.8857143 | 0 | 0 | 0 | 0 | 0 | 0 | 0 | 0 | 0 | 0 | 0 | 3 | 0 | 0 | 3 |
| ENSMUSG00000039285 | Azi2 | ENSMUST00000044454 | 32.9830706 | 33 | 15.6450671 | 18.5055458 | 0 | 0 | 0 | 0 | 0 | 0 | 0 | 0 | 0 | 0 | 0 | 0 | 1 | 2 | 3 |
| ENSMUSG00000050538 | B230217C12Rik | ENSMUST00000054783 | 29.3888166 | 31 | 21.976593 | 18.0754226 | 0 | 0 | 0 | 0 | 0 | 0 | 0 | 0 | 0 | 0 | 0 | 1 | 0 | 0 | 1 |
| ENSMUSG00000045767 | B230219D22Rik | ENSMUST00000057844 | 28.9084781 | 33 | 18.3821623 | 19.9118486 | 1 | 0 | 0 | 1 | 0 | 0 | 0 | 0 | 1 | 2 | 0 | 1 | 0 | 1 | 5 |
| ENSMUSG00000039242 | B3galnt2 | ENSMUST00000099747 | 30.5651106 | 30 | 18.9189189 | 20.8845209 | 0 | 0 | 2 | 1 | 0 | 0 | 0 | 0 | 1 | 4 | 0 | 0 | 0 | 0 | 4 |
| ENSMUSG00000034780 | B3galt1 | ENSMUST00000112346 | 30.5894849 | 31 | 18.7732342 | 19.2777483 | 0 | 0 | 0 | 0 | 0 | 0 | 0 | 0 | 0 | 0 | 0 | 1 | 6 | 4 | 11 |
| ENSMUSG00000033849 | B3galt2 | ENSMUST00000038252 | 36.0072042 | 31 | 17.1100028 | 16.1679135 | 2 | 0 | 0 | 2 | 0 | 0 | 0 | 0 | 2 | 4 | 2 | 1 | 2 | 2 | 13 |
| ENSMUSG00000050796 | B3galt6 | ENSMUST00000052185 | 26.4611137 | 30 | 21.2609296 | 22.5034514 | 0 | 0 | 0 | 0 | 0 | 0 | 0 | 0 | 0 | 0 | 4 | 0 | 1 | 0 | 5 |
| ENSMUSG00000051950 | B3glct | ENSMUST00000100404 | 27.931841 | 30 | 21.4500501 | 20.6481791 | 0 | 0 | 0 | 0 | 0 | 0 | 0 | 0 | 0 | 0 | 0 | 4 | 0 | 0 | 4 |
| ENSMUSG00000051650 | B3gnt2 | ENSMUST00000062844 | 25.8426966 | 35 | 17.8972713 | 21.187801 | 1 | 0 | 0 | 0 | 0 | 0 | 0 | 0 | 0 | 0 | 0 | 0 | 1 | 0 | 2 |
| ENSMUSG00000028413 | B4galt1 | ENSMUST00000030121 | 21.6415021 | 31 | 23.6933798 | 23.3449477 | 0 | 0 | 0 | 0 | 0 | 0 | 0 | 0 | 0 | 0 | 1 | 5 | 14 | 1 | 21 |
| ENSMUSG00000056124 | B4galt6 | ENSMUST00000070080 | 28.8823213 | 32 | 19.4250403 | 19.4787749 | 1 | 0 | 0 | 1 | 0 | 0 | 0 | 0 | 1 | 2 | 1 | 2 | 5 | 2 | 13 |
| ENSMUSG00000042742 | B630005N14Rik | ENSMUST00000203078 | 29.1288081 | 35 | 17.3703902 | 18.0117584 | 1 | 0 | 0 | 0 | 0 | 0 | 0 | 0 | 0 | 0 | 3 | 1 | 11 | 1 | 17 |
| ENSMUSG00000025612 | Bach1 | ENSMUST00000026703 | 27.2517986 | 31 | 18.9064748 | 22.8201439 | 1 | 0 | 0 | 1 | 0 | 0 | 0 | 0 | 1 | 2 | 0 | 0 | 0 | 2 | 5 |
| ENSMUSG00000037316 | Bag4 | ENSMUST00000038498 | 28.9427053 | 33 | 17.5723568 | 20.4370939 | 0 | 0 | 0 | 1 | 0 | 0 | 0 | 0 | 1 | 2 | 0 | 0 | 1 | 1 | 4 |
| ENSMUSG00000026196 | Bard1 | ENSMUST00000027393 | 23.2627399 | 35 | 22.2700199 | 19.4242224 | 1 | 1 | 0 | 3 | 0 | 0 | 0 | 1 | 2 | 6 | 0 | 16 | 14 | 0 | 38 |
| ENSMUSG00000045763 | Basp1 | ENSMUST00000058845 | 31.1440678 | 29 | 21.5042373 | 18.75 | 0 | 0 | 0 | 0 | 0 | 0 | 0 | 0 | 0 | 0 | 0 | 4 | 0 | 0 | 4 |
| ENSMUSG00000035021 | Baz1a | ENSMUST00000038926 | 28.7993921 | 33 | 17.5531915 | 20.5927052 | 2 | 1 | 0 | 0 | 0 | 0 | 0 | 0 | 0 | 0 | 0 | 0 | 0 | 0 | 3 |
| ENSMUSG00000026987 | Baz2b | ENSMUST00000112550 | 30.5598756 | 34 | 17.807154 | 17.8849145 | 0 | 0 | 0 | 1 | 0 | 0 | 0 | 0 | 1 | 2 | 0 | 0 | 0 | 0 | 2 |
| ENSMUSG00000002083 | Bbc3 | ENSMUST00000002152 | 18.9495366 | 20 | 28.2183316 | 32.6467559 | 0 | 0 | 0 | 0 | 0 | 0 | 0 | 0 | 0 | 0 | 0 | 1 | 0 | 0 | 1 |
| ENSMUSG00000084957 | Bbip1 | ENSMUST00000135402 | 33.5131894 | 29 | 19.3645084 | 17.8057554 | 0 | 0 | 0 | 0 | 0 | 0 | 0 | 0 | 0 | 0 | 0 | 0 | 1 | 1 | 2 |
| ENSMUSG00000057265 | Bbof1 | ENSMUST00000081828 | 29.9578059 | 26 | 19.4092827 | 24.6835443 | 0 | 0 | 0 | 0 | 0 | 0 | 0 | 0 | 0 | 0 | 0 | 0 | 1 | 0 | 1 |
| ENSMUSG00000035759 | Bbs10 | ENSMUST00000040454 | 27.1604938 | 30 | 22.7513228 | 20.4585538 | 0 | 0 | 0 | 0 | 0 | 0 | 0 | 0 | 0 | 0 | 0 | 1 | 0 | 0 | 1 |
| ENSMUSG00000037325 | Bbs7 | ENSMUST00000108155 | 27.4698795 | 36 | 16.8674699 | 19.7590361 | 0 | 0 | 0 | 0 | 0 | 0 | 0 | 0 | 0 | 0 | 2 | 0 | 3 | 0 | 5 |
| ENSMUSG00000022641 | Bbx | ENSMUST00000138166 | 27.7515311 | 32 | 19.1776028 | 20.6824147 | 1 | 0 | 0 | 1 | 0 | 0 | 0 | 0 | 1 | 2 | 0 | 0 | 10 | 2 | 15 |
| ENSMUSG00000073139 | BC023829 | ENSMUST00000101506 | 26.023166 | 27 | 25.1737452 | 22.2393822 | 0 | 0 | 0 | 0 | 0 | 0 | 0 | 0 | 0 | 0 | 0 | 1 | 1 | 0 | 2 |
| ENSMUSG00000046096 | BC030336 | ENSMUST00000060175 | 27.9274982 | 34 | 18.1015979 | 20.2718817 | 1 | 0 | 0 | 1 | 0 | 0 | 0 | 0 | 1 | 2 | 2 | 1 | 7 | 0 | 13 |

| | | | | | | | | | | | | | | | | | | | | | |
|---|---|---|---|---|---|---|---|---|---|---|---|---|---|---|---|---|---|---|---|---|---|
| ENSMUSG00000036299 | BC031181; LOC102641497 | ENSMUST00000040284 | 23.8983051 | 30 | 22.0338983 | 23.7288136 | 0 | 0 | 0 | 1 | 0 | 0 | 0 | 0 | 1 | 2 | 0 | 1 | 0 | 0 | 3 |
| ENSMUSG00000053684 | BC048403 | ENSMUST00000136432 | 26.0375213 | 32 | 22.2853894 | 19.9545196 | 1 | 0 | 1 | 0 | 0 | 0 | 0 | 0 | 0 | 1 | 0 | 29 | 1 | 0 | 32 |
| ENSMUSG00000040282 | BC052040 | ENSMUST00000110918 | 26.5400115 | 30 | 23.3160622 | 20.3223949 | 0 | 0 | 0 | 0 | 0 | 0 | 0 | 0 | 0 | 0 | 0 | 21 | 2 | 1 | 24 |
| ENSMUSG00000005687 | Bcas2 | ENSMUST00000135017 | 31.1910883 | 28 | 17.1379606 | 23.479006 | 0 | 0 | 0 | 0 | 0 | 0 | 0 | 0 | 0 | 0 | 0 | 0 | 0 | 1 | 1 |
| ENSMUSG00000027792 | Bche | ENSMUST00000029367 | 32.5626673 | 33 | 18.4473108 | 16.4760282 | 1 | 0 | 0 | 1 | 0 | 0 | 0 | 0 | 1 | 2 | 0 | 0 | 0 | 0 | 3 |
| ENSMUSG00000028191 | Bcl10 | ENSMUST00000029842 | 25.4285714 | 35 | 18.5714286 | 21.1428571 | 0 | 0 | 0 | 1 | 0 | 0 | 0 | 0 | 1 | 2 | 0 | 0 | 0 | 0 | 2 |
| ENSMUSG00000000861 | Bcl11a | ENSMUST00000109514 | 30.6772908 | 34 | 16.5491879 | 18.8476862 | 0 | 0 | 0 | 0 | 0 | 0 | 0 | 0 | 0 | 0 | 0 | 2 | 2 | 2 | 6 |
| ENSMUSG00000027381 | Bcl2l11 | ENSMUST00000110341 | 22.6410621 | 31 | 23.2100522 | 22.9966809 | 1 | 0 | 0 | 0 | 0 | 0 | 0 | 0 | 0 | 0 | 0 | 2 | 1 | 0 | 4 |
| ENSMUSG00000044165 | Bcl2l15 | ENSMUST00000106822 | 30.5286672 | 30 | 17.4236783 | 22.1891288 | 0 | 0 | 1 | 2 | 0 | 0 | 0 | 0 | 2 | 5 | 1 | 0 | 3 | 1 | 10 |
| ENSMUSG00000089682 | Bcl2l2 | ENSMUST00000133397 | 22.8202707 | 26 | 22.3796034 | 29.1784703 | 0 | 0 | 0 | 0 | 0 | 0 | 0 | 0 | 0 | 0 | 0 | 3 | 0 | 0 | 3 |
| ENSMUSG00000029681 | Bcl7b | ENSMUST00000031692 | 22.3783784 | 26 | 26.3783784 | 25.7297297 | 0 | 0 | 0 | 0 | 0 | 0 | 0 | 0 | 0 | 0 | 0 | 0 | 0 | 1 | 1 |
| ENSMUSG00000037608 | Bclaf1 | ENSMUST00000092678 | 32.6104746 | 35 | 16.1620295 | 16.6530278 | 0 | 0 | 0 | 1 | 0 | 0 | 0 | 0 | 1 | 2 | 0 | 0 | 4 | 1 | 7 |
| ENSMUSG00000048482 | Bdnf | ENSMUST00000053317 | 31.558669 | 32 | 16.7775832 | 19.3345009 | 1 | 0 | 0 | 0 | 0 | 0 | 0 | 0 | 0 | 0 | 2 | 2 | 8 | 2 | 15 |
| ENSMUSG00000049658 | Bdp1 | ENSMUST00000099262 | 30.24 | 31 | 18.1485714 | 20.4342857 | 1 | 0 | 0 | 2 | 0 | 0 | 0 | 0 | 2 | 4 | 3 | 0 | 7 | 2 | 17 |
| ENSMUSG00000092060 | Bend4 | ENSMUST00000201705 | 24.6890074 | 30 | 21.4140645 | 23.9908606 | 0 | 0 | 0 | 0 | 0 | 0 | 0 | 0 | 0 | 0 | 6 | 0 | 17 | 0 | 23 |
| ENSMUSG00000030256 | Bhlhe41 | ENSMUST00000032386 | 26.2529833 | 32 | 20.0477327 | 22.0438273 | 0 | 0 | 0 | 0 | 0 | 0 | 0 | 0 | 0 | 0 | 0 | 2 | 27 | 0 | 29 |
| ENSMUSG00000003452 | Bicd1 | ENSMUST00000086829 | 32.9311886 | 35 | 15.5495979 | 16.4432529 | 4 | 2 | 0 | 6 | 0 | 0 | 0 | 1 | 5 | 12 | 2 | 2 | 3 | 3 | 28 |
| ENSMUSG00000017716 | Birc5 | ENSMUST00000093906 | 23.2288828 | 30 | 23.2629428 | 23.2629428 | 3 | 1 | 1 | 6 | 0 | 0 | 1 | 2 | 3 | 13 | 1 | 0 | 9 | 0 | 27 |
| ENSMUSG00000057506 | Bloc1s2 | ENSMUST00000079033 | 24.0384615 | 31 | 20.6730769 | 24.2788462 | 0 | 0 | 0 | 0 | 0 | 0 | 0 | 0 | 0 | 0 | 0 | 0 | 3 | 0 | 3 |
| ENSMUSG00000005804 | Bloc1s6 | ENSMUST00000005954 | 26.3620387 | 31 | 21.6871705 | 21.2302285 | 0 | 0 | 0 | 0 | 0 | 0 | 0 | 0 | 0 | 0 | 0 | 19 | 0 | 0 | 19 |
| ENSMUSG00000026577 | Blzf1 | ENSMUST00000027866 | 33.3882935 | 29 | 17.1887881 | 20.7337181 | 1 | 0 | 0 | 2 | 0 | 0 | 0 | 0 | 2 | 4 | 0 | 0 | 0 | 0 | 5 |
| ENSMUSG00000026739 | Bmi1 | ENSMUST00000028071 | 30.726257 | 37 | 15.0837989 | 17.2718808 | 0 | 0 | 0 | 1 | 0 | 0 | 0 | 0 | 1 | 2 | 0 | 0 | 0 | 0 | 2 |
| ENSMUSG00000029335 | Bmp3 | ENSMUST00000200388 | 30.2382003 | 31 | 20.3352448 | 18.2179091 | 0 | 1 | 0 | 1 | 0 | 0 | 0 | 1 | 0 | 2 | 0 | 0 | 7 | 1 | 11 |
| ENSMUSG00000032179 | Bmp5 | ENSMUST00000012281 | 28.9930556 | 35 | 17.2453704 | 19.1550926 | 1 | 1 | 2 | 3 | 0 | 0 | 0 | 1 | 2 | 8 | 1 | 0 | 5 | 0 | 16 |
| ENSMUSG00000031963 | Bmper | ENSMUST00000071982 | 29.4871795 | 34 | 16.4957265 | 20.5128205 | 0 | 0 | 0 | 1 | 0 | 0 | 0 | 0 | 1 | 2 | 0 | 0 | 0 | 0 | 2 |
| ENSMUSG00000021796 | Bmpr1a | ENSMUST00000049005 | 29.6587927 | 33 | 18.7401575 | 18.5301837 | 0 | 0 | 0 | 0 | 0 | 0 | 0 | 0 | 0 | 0 | 0 | 0 | 5 | 0 | 5 |
| ENSMUSG00000052430 | Bmpr1b | ENSMUST00000029948 | 29.1463064 | 32 | 18.4248347 | 20.3219316 | 3 | 2 | 0 | 3 | 0 | 0 | 0 | 0 | 3 | 6 | 0 | 1 | 0 | 2 | 14 |
| ENSMUSG00000067336 | Bmpr2 | ENSMUST00000087435 | 29.5110798 | 35 | 18.3726111 | 16.9656466 | 3 | 3 | 0 | 6 | 0 | 0 | 0 | 1 | 5 | 12 | 2 | 0 | 4 | 1 | 25 |
| ENSMUSG00000049086 | Bmyc | ENSMUST00000061483 | 30.7420495 | 31 | 22.614841 | 15.1943463 | 0 | 0 | 0 | 0 | 0 | 0 | 0 | 0 | 0 | 0 | 0 | 1 | 0 | 0 | 1 |
| ENSMUSG00000078566 | Bnip3 | ENSMUST00000106112 | 29.3790547 | 35 | 17.7015755 | 18.0722892 | 0 | 0 | 1 | 0 | 0 | 0 | 0 | 0 | 0 | 1 | 0 | 0 | 0 | 0 | 1 |
| ENSMUSG00000022051 | Bnip3l | ENSMUST00000111115 | 29.1601866 | 32 | 19.2068429 | 19.5567652 | 0 | 0 | 0 | 0 | 0 | 0 | 0 | 0 | 0 | 0 | 0 | 0 | 2 | 0 | 2 |
| ENSMUSG00000022070 | Bora | ENSMUST00000022656 | 32.9297821 | 29 | 16.2227603 | 21.7917676 | 0 | 0 | 0 | 0 | 0 | 0 | 0 | 0 | 0 | 0 | 0 | 1 | 1 | 1 | 3 |
| ENSMUSG00000062376 | Borcs7 | ENSMUST00000074912 | 29.5154185 | 29 | 23.0726872 | 18.5022026 | 0 | 0 | 0 | 1 | 0 | 0 | 0 | 0 | 1 | 2 | 0 | 42 | 0 | 0 | 44 |
| ENSMUSG00000002413 | Braf | ENSMUST00000002487 | 28.8873338 | 32 | 19.650105 | 19.9020294 | 2 | 1 | 0 | 1 | 0 | 0 | 0 | 1 | 0 | 2 | 0 | 0 | 13 | 4 | 22 |
| ENSMUSG00000031201 | Brcc3 | ENSMUST00000033544 | 35.8360754 | 32 | 15.7044571 | 16.5121149 | 0 | 0 | 0 | 1 | 0 | 0 | 0 | 0 | 1 | 2 | 2 | 0 | 1 | 1 | 6 |
| ENSMUSG00000029279 | Brdt | ENSMUST00000031215 | 29.7650131 | 31 | 18.2767624 | 21.2793734 | 0 | 0 | 0 | 0 | 0 | 0 | 0 | 0 | 0 | 0 | 1 | 0 | 1 | 0 | 2 |
| ENSMUSG00000037905 | Bri3bp | ENSMUST00000049040 | 24.6572904 | 30 | 22.4675093 | 23.2508456 | 0 | 0 | 0 | 0 | 0 | 0 | 0 | 0 | 0 | 0 | 6 | 4 | 13 | 4 | 27 |
| ENSMUSG00000034329 | Brip1 | ENSMUST00000044423 | 29.5654816 | 33 | 19.021574 | 18.8696445 | 0 | 0 | 0 | 0 | 0 | 0 | 0 | 0 | 0 | 0 | 0 | 1 | 0 | 0 | 1 |
| ENSMUSG00000046836 | Brox | ENSMUST00000194327 | 28.583993 | 27 | 20.8443272 | 23.0870712 | 2 | 1 | 1 | 0 | 0 | 0 | 0 | 0 | 0 | 1 | 0 | 0 | 0 | 0 | 4 |
| ENSMUSG00000031130 | Brs3 | ENSMUST00000033464 | 29.3361884 | 33 | 17.4875089 | 19.9143469 | 0 | 0 | 0 | 0 | 0 | 0 | 0 | 0 | 0 | 0 | 0 | 0 | 2 | 0 | 2 |
| ENSMUSG00000022914 | Brwd1 | ENSMUST00000113829 | 25.3255655 | 35 | 17.9232351 | 21.7957505 | 1 | 0 | 1 | 1 | 0 | 0 | 0 | 0 | 1 | 3 | 0 | 0 | 2 | 0 | 6 |
| ENSMUSG00000063663 | Brwd3 | ENSMUST00000150434 | 30.3 | 37 | 16.45 | 16.6333333 | 1 | 0 | 0 | 1 | 0 | 0 | 0 | 0 | 1 | 2 | 0 | 2 | 2 | 2 | 9 |
| ENSMUSG00000038187 | Btbd10 | ENSMUST00000135510 | 28.8830139 | 31 | 18.2419035 | 22.2075347 | 1 | 1 | 0 | 1 | 0 | 0 | 0 | 0 | 1 | 2 | 0 | 0 | 0 | 0 | 4 |
| ENSMUSG00000062098 | Btbd3 | ENSMUST00000091556 | 27.728513 | 32 | 19.8499318 | 20.2933151 | 0 | 0 | 0 | 0 | 0 | 0 | 0 | 0 | 0 | 0 | 0 | 1 | 4 | 2 | 7 |
| ENSMUSG00000028568 | Btf3l4 | ENSMUST00000102742 | 26.5618861 | 36 | 19.1748527 | 18.6247544 | 0 | 0 | 0 | 0 | 0 | 0 | 0 | 0 | 0 | 0 | 0 | 0 | 1 | 0 | 1 |
| ENSMUSG00000036478 | Btg1 | ENSMUST00000038377 | 26.4748201 | 34 | 18.705036 | 21.1510791 | 0 | 0 | 0 | 1 | 0 | 0 | 0 | 0 | 1 | 2 | 3 | 0 | 21 | 1 | 27 |
| ENSMUSG00000020423 | Btg2 | ENSMUST00000020692 | 22.9649841 | 29 | 23.2833106 | 24.2837653 | 0 | 0 | 1 | 0 | 0 | 0 | 0 | 0 | 0 | 1 | 0 | 4 | 7 | 4 | 16 |
| ENSMUSG00000066979 | Bub3 | ENSMUST00000084502 | 20.990099 | 33 | 21.6831683 | 24.0594059 | 0 | 0 | 0 | 0 | 0 | 0 | 0 | 0 | 0 | 0 | 1 | 0 | 1 | 0 | 2 |
| ENSMUSG00000051223 | Bzw1 | ENSMUST00000050552 | 29.1648448 | 34 | 17.9711412 | 18.3646699 | 0 | 0 | 0 | 0 | 0 | 0 | 0 | 0 | 0 | 0 | 0 | 1 | 0 | 4 | 5 |
| ENSMUSG00000029917 | C130060K24Rik | ENSMUST00000170608 | 31.5511163 | 30 | 19.2714454 | 18.8601645 | 0 | 0 | 0 | 0 | 0 | 0 | 0 | 0 | 0 | 0 | 0 | 0 | 0 | 1 | 1 |
| ENSMUSG00000000581 | C1d | ENSMUST00000000594 | 29.7460701 | 36 | 14.5908908 | 19.5485691 | 0 | 0 | 0 | 1 | 0 | 0 | 0 | 0 | 1 | 2 | 0 | 0 | 1 | 0 | 3 |
| ENSMUSG00000042460 | C1galt1 | ENSMUST00000040159 | 30.2934537 | 34 | 17.4266366 | 17.8555305 | 0 | 0 | 1 | 1 | 0 | 0 | 0 | 0 | 1 | 3 | 0 | 0 | 3 | 0 | 6 |
| ENSMUSG00000049630 | C1ql3 | ENSMUST00000061545 | 26.7326733 | 36 | 17.8217822 | 19.4306931 | 0 | 0 | 0 | 0 | 0 | 0 | 0 | 0 | 0 | 0 | 0 | 0 | 0 | 2 | 2 |
| ENSMUSG00000030279 | C2cd5 | ENSMUST00000205119 | 27.1805815 | 26 | 22.4859963 | 24.3264871 | 0 | 0 | 0 | 0 | 0 | 0 | 0 | 0 | 0 | 0 | 1 | 0 | 0 | 1 | 2 |
| ENSMUSG00000006423 | C330007P06Rik | ENSMUST00000115248 | 28.079781 | 34 | 18.8502151 | 19.1239734 | 0 | 0 | 0 | 0 | 0 | 0 | 0 | 0 | 0 | 0 | 0 | 15 | 2 | 2 | 19 |
| ENSMUSG00000033031 | C330027C09Rik | ENSMUST00000048374 | 31.2068966 | 40 | 13.7931034 | 15.1724138 | 1 | 0 | 1 | 0 | 0 | 0 | 0 | 0 | 0 | 1 | 0 | 0 | 2 | 0 | 4 |
| ENSMUSG00000046449 | C77370 | ENSMUST00000118314 | 30.7572452 | 33 | 17.4665005 | 19.0401994 | 0 | 0 | 0 | 0 | 0 | 0 | 0 | 0 | 0 | 0 | 0 | 0 | 1 | 0 | 1 |
| ENSMUSG00000028578 | Caap1 | ENSMUST00000030313 | 32.8389831 | 39 | 12.3940678 | 15.7838983 | 0 | 0 | 0 | 0 | 0 | 0 | 0 | 0 | 0 | 0 | 0 | 0 | 0 | 1 | 1 |
| ENSMUSG00000021981 | Cab39l | ENSMUST00000022553 | 26.7548907 | 30 | 21.3463751 | 22.2094361 | 0 | 0 | 0 | 1 | 0 | 0 | 0 | 0 | 1 | 2 | 0 | 0 | 0 | 2 | 4 |
| ENSMUSG00000040118 | Cacna2d1 | ENSMUST00000167946 | 30.6750065 | 34 | 17.3573104 | 18.0349231 | 0 | 0 | 0 | 1 | 0 | 0 | 0 | 0 | 1 | 2 | 0 | 0 | 5 | 0 | 7 |
| ENSMUSG00000057914 | Cacnb2 | ENSMUST00000114723 | 28.4934498 | 31 | 20.6331878 | 19.4323144 | 0 | 0 | 0 | 0 | 0 | 0 | 0 | 0 | 0 | 0 | 0 | 5 | 1 | 0 | 6 |
| ENSMUSG00000017412 | Cacnb4 | ENSMUST00000178799 | 31.3340898 | 33 | 17.9121414 | 17.8959313 | 1 | 0 | 0 | 0 | 0 | 0 | 0 | 0 | 0 | 0 | 0 | 3 | 2 | 0 | 6 |
| ENSMUSG00000033417 | Cacul1 | ENSMUST00000081790 | 26.89037 | 33 | 19.2139738 | 21.2364974 | 0 | 0 | 0 | 0 | 0 | 0 | 0 | 0 | 0 | 0 | 2 | 2 | 8 | 2 | 14 |
| ENSMUSG00000014226 | Cacybp | ENSMUST00000014370 | 36 | 30 | 15.0833333 | 19.4166667 | 0 | 0 | 0 | 0 | 0 | 0 | 0 | 0 | 0 | 0 | 1 | 0 | 0 | 0 | 1 |
| ENSMUSG00000032076 | Cadm1 | ENSMUST00000114548 | 28.3806344 | 33 | 18.196995 | 20.2337229 | 0 | 1 | 0 | 1 | 0 | 0 | 0 | 0 | 1 | 2 | 0 | 1 | 10 | 1 | 15 |
| ENSMUSG00000064115 | Cadm2 | ENSMUST00000114292 | 33.6548488 | 34 | 15.8107404 | 16.6058394 | 0 | 0 | 0 | 2 | 0 | 0 | 0 | 0 | 2 | 4 | 0 | 1 | 3 | 1 | 9 |
| ENSMUSG00000028222 | Calb1 | ENSMUST00000029876 | 34.6561648 | 32 | 17.1766131 | 16.5707361 | 0 | 0 | 0 | 0 | 0 | 0 | 0 | 0 | 0 | 0 | 0 | 1 | 0 | 0 | 1 |
| ENSMUSG00000059588 | Calcrl | ENSMUST00000074262 | 32.3903819 | 34 | 16.1598303 | 17.291372 | 1 | 1 | 0 | 3 | 0 | 0 | 0 | 1 | 2 | 6 | 0 | 15 | 1 | 1 | 25 |
| ENSMUSG00000036438 | Calm2 | ENSMUST00000040440 | 27.1544715 | 37 | 18.5365854 | 17.0731707 | 0 | 0 | 0 | 0 | 0 | 0 | 0 | 0 | 0 | 0 | 1 | 0 | 0 | 0 | 1 |
| ENSMUSG00000019732 | Calr3 | ENSMUST00000019876 | 28.4738041 | 31 | 21.8678815 | 18.9066059 | 0 | 0 | 0 | 0 | 0 | 0 | 0 | 0 | 0 | 0 | 0 | 0 | 2 | 2 | 4 |
| ENSMUSG00000053819 | Camk2d | ENSMUST00000199300 | 27.3182957 | 34 | 19.3817878 | 19.9640769 | 0 | 0 | 0 | 1 | 0 | 0 | 0 | 0 | 1 | 2 | 1 | 7 | 7 | 1 | 18 |
| ENSMUSG00000014592 | Camta1 | ENSMUST00000097774 | 31.8677987 | 30 | 19.1024864 | 18.9812007 | 0 | 0 | 0 | 1 | 0 | 0 | 0 | 0 | 1 | 2 | 0 | 1 | 1 | 2 | 6 |
| ENSMUSG00000020368 | Canx | ENSMUST00000179865 | 26.7557087 | 36 | 18.0525636 | 19.2158552 | 0 | 0 | 0 | 0 | 0 | 0 | 0 | 0 | 0 | 0 | 3 | 3 | 11 | 1 | 18 |
| ENSMUSG00000027184 | Caprin1 | ENSMUST00000028607 | 29.0046656 | 35 | 16.8999482 | 19.5438051 | 0 | 0 | 0 | 0 | 0 | 0 | 0 | 0 | 0 | 0 | 0 | 2 | 4 | 0 | 6 |
| ENSMUSG00000070372 | Capza1 | ENSMUST00000094028 | 26.122449 | 33 | 20.7709751 | 20.0907029 | 0 | 0 | 0 | 0 | 0 | 0 | 0 | 0 | 0 | 0 | 0 | 1 | 0 | 0 | 1 |
| ENSMUSG00000015733 | Capza2 | ENSMUST00000015877 | 28.236825 | 37 | 16.9160703 | 18.2823682 | 0 | 0 | 0 | 0 | 0 | 0 | 0 | 0 | 0 | 0 | 0 | 10 | 1 | 3 | 14 |
| ENSMUSG00000056158 | Car10 | ENSMUST00000107863 | 30.8291991 | 28 | 25.6555634 | 15.5917789 | 0 | 0 | 0 | 0 | 0 | 0 | 0 | 0 | 0 | 0 | 0 | 4 | 0 | 0 | 4 |
| ENSMUSG00000032373 | Car12 | ENSMUST00000071889 | 25.4019293 | 28 | 24.0353698 | 22.7893891 | 0 | 0 | 0 | 0 | 0 | 0 | 0 | 0 | 0 | 0 | 0 | 6 | 0 | 0 | 6 |
| ENSMUSG00000041261 | Car8 | ENSMUST00000066674 | 31.1418685 | 30 | 17.8892734 | 21.0034602 | 0 | 0 | 0 | 0 | 0 | 0 | 0 | 0 | 0 | 0 | 1 | 0 | 0 | 1 | 2 |
| ENSMUSG00000026017 | Carf | ENSMUST00000187978 | 31.5676669 | 36 | 15.3704838 | 17.3913043 | 0 | 0 | 0 | 1 | 0 | 0 | 0 | 0 | 1 | 2 | 0 | 1 | 11 | 3 | 17 |
| ENSMUSG00000078676 | Casc3 | ENSMUST00000017384 | 21.7418546 | 30 | 24.122807 | 24.0601504 | 0 | 0 | 0 | 0 | 0 | 0 | 0 | 0 | 0 | 0 | 2 | 1 | 0 | 1 | 4 |
| ENSMUSG00000060227 | Casc4 | ENSMUST00000078752 | 32.2230539 | 32 | 18.1137725 | 17.5523952 | 1 | 0 | 0 | 1 | 0 | 0 | 0 | 0 | 1 | 2 | 0 | 0 | 2 | 2 | 7 |
| ENSMUSG00000027326 | Casc5 | ENSMUST00000028802 | 29.9157303 | 33 | 18.1882022 | 19.241573 | 0 | 0 | 1 | 0 | 0 | 0 | 0 | 0 | 0 | 1 | 0 | 0 | 1 | 0 | 2 |
| ENSMUSG00000015189 | Casd1 | ENSMUST00000015333 | 28.8601455 | 35 | 17.6232821 | 18.9975748 | 1 | 1 | 0 | 0 | 0 | 0 | 0 | 0 | 0 | 0 | 0 | 0 | 0 | 0 | 2 |
| ENSMUSG00000031012 | Cask | ENSMUST00000115436 | 26.6745237 | 30 | 21.8621096 | 21.9013946 | 0 | 0 | 0 | 0 | 0 | 0 | 0 | 0 | 0 | 0 | 0 | 2 | 31 | 0 | 33 |
| ENSMUSG00000029863 | Casp2 | ENSMUST00000031895 | 20.9256524 | 36 | 22.5504677 | 20.4825209 | 0 | 0 | 0 | 0 | 0 | 0 | 0 | 0 | 0 | 0 | 0 | 0 | 1 | 0 | 1 |
| ENSMUSG00000031628 | Casp3 | ENSMUST00000093517 | 28.6738351 | 32 | 17.7419355 | 21.6845878 | 0 | 0 | 0 | 1 | 0 | 0 | 0 | 0 | 1 | 2 | 0 | 0 | 0 | 0 | 2 |
| ENSMUSG00000028282 | Casp8ap2 | ENSMUST00000029950 | 29.1935484 | 40 | 13.8709677 | 16.9354839 | 0 | 0 | 0 | 1 | 0 | 0 | 0 | 0 | 1 | 2 | 0 | 0 | 0 | 0 | 2 |
| ENSMUSG00000033486 | Catsper2 | ENSMUST00000154604 | 29.7281324 | 25 | 22.3404255 | 22.4586288 | 0 | 0 | 0 | 0 | 0 | 0 | 0 | 0 | 0 | 0 | 1 | 0 | 0 | 0 | 1 |
| ENSMUSG00000007655 | Cav1 | ENSMUST00000115453 | 28.028028 | 33 | 20.0700701 | 18.7687688 | 0 | 0 | 0 | 0 | 0 | 0 | 0 | 0 | 0 | 0 | 2 | 2 | 1 | 0 | 5 |
| ENSMUSG00000038533 | Cbfa2t2 | ENSMUST00000045270 | 24.8802682 | 29 | 22.8448276 | 22.7969349 | 0 | 0 | 0 | 0 | 0 | 0 | 0 | 0 | 0 | 0 | 0 | 2 | 20 | 0 | 22 |
| ENSMUSG00000031885 | Cbfb | ENSMUST00000109392 | 29.0510949 | 37 | 16.5450122 | 17.8588808 | 1 | 0 | 0 | 0 | 0 | 0 | 0 | 0 | 0 | 0 | 1 | 0 | 1 | 2 | 5 |

| | | | | | | | | | | | | | | | | | | | | |
|---|---|---|---|---|---|---|---|---|---|---|---|---|---|---|---|---|---|---|---|---|---|
| ENSMUSG00000034342 | Cbl | ENSMUST00000206720 | 23.5801157 | 32 | 22.7417641 | 21.9034124 | 1 | 0 | 0 | 0 | 0 | 0 | 0 | 0 | 0 | 0 | 1 | 51 | 3 | 1 | 57 |
| ENSMUSG00000022637 | Cblb | ENSMUST00000114471 | 25.9016393 | 33 | 20.6557377 | 20.1788376 | 0 | 0 | 0 | 1 | 0 | 0 | 0 | 0 | 1 | 2 | 0 | 0 | 4 | 1 | 7 |
| ENSMUSG00000024647 | Cbln2 | ENSMUST00000068423 | 27.3089701 | 30 | 20.9302326 | 21.3289037 | 0 | 0 | 0 | 0 | 0 | 0 | 0 | 0 | 0 | 0 | 0 | 0 | 3 | 1 | 4 |
| ENSMUSG00000067578 | Cbln4 | ENSMUST00000087950 | 27.7924362 | 37 | 18.82146 | 16.7985928 | 0 | 0 | 0 | 0 | 0 | 0 | 0 | 0 | 0 | 0 | 0 | 0 | 2 | 0 | 2 |
| ENSMUSG00000024878 | Cbwd1 | ENSMUST00000025815 | 32.0531758 | 34 | 17.1344165 | 16.9867061 | 1 | 0 | 0 | 0 | 0 | 0 | 0 | 0 | 0 | 0 | 0 | 0 | 0 | 0 | 1 |
| ENSMUSG00000018666 | Cbx1 | ENSMUST00000134585 | 28.6049238 | 30 | 18.6987104 | 22.5087925 | 0 | 0 | 0 | 0 | 0 | 0 | 0 | 0 | 0 | 0 | 0 | 4 | 2 | 0 | 6 |
| ENSMUSG00000029836 | Cbx3 | ENSMUST00000114445 | 28.8749017 | 35 | 14.6341463 | 21.3217939 | 1 | 0 | 1 | 1 | 0 | 0 | 0 | 0 | 1 | 3 | 0 | 0 | 17 | 0 | 21 |
| ENSMUSG00000009575 | Cbx5 | ENSMUST00000118152 | 25.1821662 | 32 | 20.6866741 | 21.6499938 | 0 | 0 | 0 | 0 | 0 | 0 | 0 | 0 | 0 | 0 | 2 | 4 | 16 | 3 | 25 |
| ENSMUSG00000053411 | Cbx7 | ENSMUST00000089293 | 20.9246712 | 25 | 27.5408529 | 26.9031487 | 1 | 1 | 0 | 0 | 0 | 0 | 0 | 0 | 0 | 0 | 0 | 1 | 0 | 2 | 5 |
| ENSMUSG00000025578 | Cbx8 | ENSMUST00000026663 | 20.1939292 | 27 | 24.7470489 | 27.8667791 | 0 | 0 | 0 | 0 | 0 | 0 | 0 | 0 | 0 | 0 | 0 | 2 | 5 | 0 | 7 |
| ENSMUSG00000071855 | Ccdc112 | ENSMUST00000072835 | 29.5927602 | 38 | 14.3891403 | 17.5565611 | 0 | 0 | 0 | 0 | 0 | 0 | 0 | 0 | 0 | 0 | 0 | 0 | 2 | 1 | 3 |
| ENSMUSG00000020482 | Ccdc117 | ENSMUST00000020776 | 25.5292653 | 31 | 19.2611042 | 23.7442922 | 0 | 0 | 0 | 0 | 0 | 0 | 0 | 0 | 0 | 0 | 0 | 1 | 0 | 0 | 1 |
| ENSMUSG00000021578 | Ccdc127 | ENSMUST00000022063 | 25.9635357 | 29 | 22.2247865 | 22.7325179 | 0 | 0 | 0 | 0 | 0 | 0 | 0 | 0 | 0 | 0 | 5 | 4 | 10 | 2 | 21 |
| ENSMUSG00000022833 | Ccdc14 | ENSMUST00000023532 | 25.952381 | 36 | 17.6984127 | 20.5555556 | 1 | 0 | 0 | 1 | 0 | 0 | 0 | 0 | 1 | 2 | 1 | 0 | 2 | 0 | 6 |
| ENSMUSG00000036641 | Ccdc148 | ENSMUST00000077687 | 30.8023694 | 29 | 18.6322025 | 21.1093161 | 0 | 0 | 0 | 0 | 0 | 0 | 0 | 0 | 0 | 0 | 0 | 0 | 6 | 0 | 6 |
| ENSMUSG00000050050 | Ccdc158 | ENSMUST00000151180 | 28.4615385 | 31 | 19.1346154 | 21.3942308 | 0 | 0 | 0 | 0 | 0 | 0 | 0 | 0 | 0 | 0 | 6 | 5 | 10 | 0 | 21 |
| ENSMUSG00000035173 | Ccdc186 | ENSMUST00000118592 | 26.479035 | 32 | 19.8449167 | 21.7690982 | 0 | 1 | 0 | 0 | 0 | 0 | 0 | 0 | 0 | 0 | 0 | 0 | 1 | 0 | 2 |
| ENSMUSG00000059554 | Ccdc28a | ENSMUST00000052648 | 29.1970803 | 29 | 19.8407432 | 21.6323822 | 0 | 0 | 0 | 0 | 0 | 0 | 0 | 0 | 0 | 0 | 0 | 0 | 0 | 3 | 3 |
| ENSMUSG00000039983 | Ccdc32 | ENSMUST00000036470 | 27.1852446 | 25 | 21.8925421 | 25.9823577 | 0 | 0 | 0 | 0 | 0 | 0 | 0 | 0 | 0 | 0 | 0 | 1 | 0 | 0 | 1 |
| ENSMUSG00000027160 | Ccdc34 | ENSMUST00000127630 | 33.5705813 | 30 | 16.5480427 | 19.7508897 | 1 | 1 | 0 | 1 | 0 | 0 | 0 | 0 | 1 | 2 | 0 | 0 | 0 | 0 | 4 |
| ENSMUSG00000027676 | Ccdc39 | ENSMUST00000029222 | 34.1605839 | 34 | 15.3284672 | 16.2043796 | 0 | 0 | 0 | 0 | 0 | 0 | 0 | 0 | 0 | 0 | 2 | 0 | 5 | 2 | 9 |
| ENSMUSG00000020925 | Ccdc43 | ENSMUST00000092569 | 28.2193469 | 33 | 18.1146026 | 20.4559458 | 0 | 0 | 0 | 0 | 0 | 0 | 0 | 0 | 0 | 0 | 0 | 0 | 1 | 2 | 3 |
| ENSMUSG00000078622 | Ccdc47 | ENSMUST00000002043 | 25.5102041 | 34 | 18.3390023 | 22.5340136 | 0 | 0 | 0 | 2 | 0 | 0 | 0 | 0 | 2 | 4 | 0 | 0 | 7 | 0 | 11 |
| ENSMUSG00000038127 | Ccdc50 | ENSMUST00000100026 | 28.5330197 | 33 | 18.5010923 | 19.8454041 | 1 | 0 | 0 | 1 | 0 | 0 | 0 | 0 | 1 | 2 | 2 | 27 | 6 | 1 | 39 |
| ENSMUSG00000048701 | Ccdc6 | ENSMUST00000147545 | 24.4263131 | 30 | 22.845487 | 22.9984702 | 0 | 0 | 0 | 0 | 0 | 0 | 0 | 0 | 0 | 0 | 4 | 1 | 12 | 0 | 17 |
| ENSMUSG00000045106 | Ccdc73 | ENSMUST00000111114 | 26.9439421 | 37 | 21.880651 | 13.7432188 | 0 | 0 | 0 | 0 | 0 | 0 | 0 | 0 | 0 | 0 | 1 | 0 | 0 | 0 | 1 |
| ENSMUSG00000032740 | Ccdc88a | ENSMUST00000040182 | 33.386277 | 34 | 16.13723 | 16.296061 | 0 | 1 | 1 | 0 | 0 | 0 | 0 | 0 | 0 | 1 | 0 | 2 | 0 | 0 | 4 |
| ENSMUSG00000074715 | Ccl28 | ENSMUST00000099241 | 28.892944 | 32 | 20.4683698 | 18.8564477 | 0 | 0 | 0 | 0 | 0 | 0 | 0 | 0 | 0 | 0 | 0 | 18 | 3 | 0 | 21 |
| ENSMUSG00000041431 | Ccnb1 | ENSMUST00000072119 | 25.6809339 | 37 | 19.844358 | 17.1206226 | 1 | 0 | 1 | 0 | 0 | 0 | 0 | 0 | 0 | 1 | 0 | 0 | 0 | 0 | 2 |
| ENSMUSG00000032218 | Ccnb2 | ENSMUST00000034742 | 30 | 37 | 19.1304348 | 13.9130435 | 0 | 0 | 0 | 0 | 0 | 0 | 0 | 0 | 0 | 0 | 1 | 0 | 0 | 0 | 1 |
| ENSMUSG00000028252 | Ccnc | ENSMUST00000065928 | 32.5448613 | 35 | 15.3344209 | 16.8026101 | 0 | 0 | 0 | 0 | 0 | 0 | 0 | 0 | 0 | 0 | 0 | 0 | 4 | 0 | 4 |
| ENSMUSG00000070348 | Ccnd1 | ENSMUST00000093962 | 24.813154 | 28 | 21.6367713 | 25.6726457 | 0 | 0 | 0 | 0 | 0 | 0 | 0 | 0 | 0 | 0 | 0 | 0 | 4 | 0 | 4 |
| ENSMUSG00000000184 | Ccnd2 | ENSMUST00000000188 | 25.4591146 | 29 | 21.4768993 | 23.5646627 | 0 | 0 | 0 | 0 | 0 | 0 | 0 | 0 | 0 | 0 | 7 | 3 | 24 | 1 | 35 |
| ENSMUSG00000028212 | Ccne2 | ENSMUST00000029866 | 35.0981559 | 35 | 14.3961927 | 15.2885187 | 1 | 1 | 0 | 4 | 0 | 0 | 1 | 2 | 1 | 8 | 0 | 0 | 0 | 0 | 10 |
| ENSMUSG00000020326 | Ccng1 | ENSMUST00000020576 | 33.5060449 | 30 | 18.2642487 | 18.1778929 | 0 | 0 | 0 | 0 | 0 | 0 | 0 | 0 | 0 | 0 | 0 | 0 | 0 | 1 | 1 |
| ENSMUSG00000029385 | Ccng2 | ENSMUST00000031331 | 27.9578606 | 36 | 15.802269 | 19.9351702 | 0 | 0 | 0 | 0 | 0 | 0 | 0 | 0 | 0 | 0 | 0 | 1 | 0 | 0 | 1 |
| ENSMUSG00000063015 | Ccni | ENSMUST00000058550 | 28.7356322 | 35 | 18.2950192 | 18.2950192 | 0 | 0 | 0 | 0 | 0 | 0 | 0 | 0 | 0 | 0 | 0 | 1 | 0 | 0 | 1 |
| ENSMUSG00000029068 | Ccnl2 | ENSMUST00000126346 | 26.5179678 | 24 | 23.7298637 | 25.8364312 | 0 | 0 | 0 | 0 | 0 | 0 | 0 | 0 | 0 | 0 | 0 | 0 | 1 | 0 | 1 |
| ENSMUSG00000011960 | Ccnt1 | ENSMUST00000169707 | 27.0676692 | 33 | 17.0942061 | 22.9102167 | 0 | 0 | 0 | 1 | 0 | 0 | 0 | 0 | 1 | 2 | 1 | 0 | 8 | 0 | 11 |
| ENSMUSG00000026349 | Ccnt2 | ENSMUST00000027587 | 27.4988894 | 33 | 17.8365171 | 21.5459796 | 0 | 0 | 1 | 1 | 0 | 0 | 0 | 0 | 1 | 3 | 1 | 1 | 3 | 1 | 9 |
| ENSMUSG00000033904 | Ccp110 | ENSMUST00000106557 | 29.0492958 | 34 | 16.6079812 | 20.5399061 | 0 | 0 | 0 | 0 | 0 | 0 | 0 | 0 | 0 | 0 | 1 | 0 | 2 | 0 | 3 |
| ENSMUSG00000029530 | Ccr9 | ENSMUST00000180093 | 31.6844488 | 27 | 21.4354995 | 20.2392499 | 0 | 0 | 0 | 0 | 0 | 0 | 0 | 0 | 0 | 0 | 1 | 0 | 4 | 1 | 6 |
| ENSMUSG00000039578 | Ccser1 | ENSMUST00000126214 | 29.9461124 | 36 | 16.0892995 | 18.3987683 | 1 | 0 | 0 | 0 | 0 | 0 | 0 | 0 | 0 | 0 | 1 | 0 | 9 | 0 | 11 |
| ENSMUSG00000034024 | Cct2 | ENSMUST00000047672 | 28.6290323 | 36 | 18.9516129 | 16.5322581 | 0 | 1 | 1 | 0 | 0 | 0 | 0 | 0 | 0 | 1 | 0 | 0 | 0 | 0 | 2 |
| ENSMUSG00000019818 | Cd164 | ENSMUST00000019962 | 28.6361577 | 34 | 18.3053919 | 18.9850476 | 1 | 0 | 0 | 1 | 0 | 0 | 0 | 0 | 1 | 2 | 0 | 0 | 0 | 0 | 3 |
| ENSMUSG00000022661 | Cd200 | ENSMUST00000163230 | 27.1903323 | 33 | 19.939577 | 20.2416918 | 0 | 0 | 0 | 0 | 0 | 0 | 0 | 0 | 0 | 0 | 0 | 13 | 2 | 0 | 15 |
| ENSMUSG00000047139 | Cd24a | ENSMUST00000058714 | 29.1249165 | 31 | 19.6392786 | 20.1068804 | 0 | 0 | 0 | 0 | 0 | 0 | 0 | 0 | 0 | 0 | 0 | 0 | 1 | 2 | 3 |
| ENSMUSG00000016496 | Cd274 | ENSMUST00000016640 | 25.3447633 | 32 | 23.2202758 | 19.455833 | 2 | 0 | 1 | 3 | 0 | 0 | 0 | 0 | 3 | 7 | 0 | 21 | 0 | 1 | 31 |
| ENSMUSG00000061665 | Cd2ap | ENSMUST00000024709 | 33.0174711 | 32 | 17.5007403 | 17.6488007 | 0 | 1 | 0 | 0 | 0 | 0 | 0 | 0 | 0 | 0 | 0 | 1 | 0 | 0 | 2 |
| ENSMUSG00000004609 | Cd33 | ENSMUST00000205503 | 26.582006 | 35 | 20.9642703 | 17.6711149 | 1 | 0 | 0 | 0 | 0 | 0 | 0 | 0 | 0 | 0 | 0 | 5 | 2 | 1 | 9 |
| ENSMUSG00000016493 | Cd46 | ENSMUST00000162650 | 31.8621758 | 30 | 20.596206 | 17.9829655 | 1 | 0 | 0 | 1 | 0 | 0 | 0 | 0 | 1 | 2 | 0 | 0 | 3 | 2 | 8 |
| ENSMUSG00000055447 | Cd47 | ENSMUST00000084838 | 25.5504476 | 34 | 22.0421002 | 18.3885797 | 1 | 0 | 0 | 1 | 0 | 0 | 0 | 0 | 1 | 2 | 1 | 44 | 14 | 2 | 64 |
| ENSMUSG00000040747 | Cd53 | ENSMUST00000038845 | 29.6715742 | 35 | 20.8380521 | 14.8357871 | 0 | 0 | 0 | 0 | 0 | 0 | 0 | 0 | 0 | 0 | 0 | 0 | 7 | 2 | 9 |
| ENSMUSG00000026399 | Cd55 | ENSMUST00000027650 | 35.8754864 | 30 | 15.4085603 | 19.0661479 | 1 | 1 | 0 | 2 | 0 | 0 | 0 | 1 | 1 | 4 | 0 | 0 | 2 | 0 | 8 |
| ENSMUSG00000068686 | Cd59b | ENSMUST00000090429 | 24.9190939 | 29 | 23.3009709 | 22.3300971 | 0 | 0 | 0 | 0 | 0 | 0 | 0 | 0 | 0 | 0 | 0 | 4 | 0 | 0 | 4 |
| ENSMUSG00000030342 | Cd9 | ENSMUST00000032492 | 30.1265823 | 38 | 13.9240506 | 17.4683544 | 0 | 0 | 0 | 0 | 0 | 0 | 0 | 0 | 0 | 0 | 0 | 0 | 1 | 0 | 1 |
| ENSMUSG00000033502 | Cdc14a | ENSMUST00000090464 | 30.1505174 | 34 | 15.6632173 | 19.943556 | 1 | 0 | 0 | 0 | 0 | 0 | 0 | 0 | 0 | 0 | 0 | 1 | 1 | 0 | 3 |
| ENSMUSG00000033102 | Cdc14b | ENSMUST00000039318 | 28.5851549 | 30 | 21.0665386 | 20.8263272 | 0 | 0 | 1 | 0 | 0 | 0 | 0 | 0 | 0 | 0 | 0 | 0 | 0 | 3 | 4 |
| ENSMUSG00000020687 | Cdc27 | ENSMUST00000093923 | 28.636523 | 34 | 17.9810726 | 19.5583596 | 0 | 0 | 0 | 0 | 0 | 0 | 0 | 0 | 0 | 0 | 0 | 1 | 0 | 1 | 2 |
| ENSMUSG00000024780 | Cdc37l1 | ENSMUST00000050148 | 27.3720137 | 33 | 18.3617747 | 21.5699659 | 0 | 0 | 0 | 0 | 0 | 0 | 0 | 0 | 0 | 0 | 0 | 0 | 2 | 0 | 2 |
| ENSMUSG00000038446 | Cdc40 | ENSMUST00000044166 | 28.9117836 | 31 | 21.6355441 | 18.6091436 | 0 | 0 | 2 | 0 | 0 | 0 | 0 | 0 | 0 | 2 | 0 | 0 | 0 | 0 | 2 |
| ENSMUSG00000006699 | Cdc42 | ENSMUST00000051477 | 25.94172 | 34 | 20.4690832 | 19.9715707 | 0 | 0 | 0 | 0 | 0 | 0 | 0 | 0 | 0 | 0 | 1 | 0 | 5 | 2 | 8 |
| ENSMUSG00000029283 | Cdc7 | ENSMUST00000031221 | 26.5552461 | 34 | 15.2274838 | 24.1411328 | 0 | 0 | 1 | 0 | 0 | 0 | 0 | 0 | 0 | 1 | 0 | 1 | 0 | 0 | 2 |
| ENSMUSG00000026361 | Cdc73 | ENSMUST00000018337 | 30.6552824 | 35 | 15.6568254 | 18.7840757 | 2 | 1 | 0 | 3 | 0 | 0 | 0 | 0 | 3 | 6 | 0 | 4 | 4 | 0 | 17 |
| ENSMUSG00000040452 | Cdh12 | ENSMUST00000075132 | 33.9121887 | 34 | 17.3984273 | 15.1376147 | 0 | 0 | 0 | 1 | 0 | 0 | 0 | 0 | 1 | 2 | 0 | 0 | 2 | 0 | 4 |
| ENSMUSG00000026312 | Cdh7 | ENSMUST00000146282 | 32.9654849 | 35 | 15.003522 | 17.1166941 | 3 | 1 | 0 | 5 | 0 | 0 | 0 | 1 | 4 | 10 | 0 | 0 | 0 | 0 | 14 |
| ENSMUSG00000036510 | Cdh8 | ENSMUST00000145601 | 30.7397959 | 31 | 17.8996599 | 20.1105442 | 1 | 0 | 0 | 1 | 0 | 0 | 0 | 0 | 1 | 2 | 0 | 0 | 0 | 0 | 3 |
| ENSMUSG00000025370 | Cdh9 | ENSMUST00000026432 | 33.5680751 | 36 | 13.4976526 | 16.6666667 | 0 | 1 | 1 | 0 | 0 | 0 | 0 | 0 | 0 | 1 | 0 | 0 | 0 | 0 | 2 |
| ENSMUSG00000019942 | Cdk1 | ENSMUST00000020099 | 24.1545894 | 30 | 21.900161 | 23.6178207 | 0 | 0 | 0 | 0 | 0 | 0 | 0 | 0 | 0 | 0 | 1 | 0 | 0 | 0 | 1 |
| ENSMUSG00000020015 | Cdk17 | ENSMUST00000069965 | 25.9755387 | 37 | 16.4239953 | 20.3261503 | 1 | 0 | 0 | 0 | 0 | 0 | 0 | 0 | 0 | 0 | 6 | 0 | 5 | 5 | 17 |
| ENSMUSG00000038481 | Cdk19 | ENSMUST00000044672 | 26.3511831 | 32 | 20.373599 | 21.7185554 | 0 | 0 | 1 | 0 | 0 | 0 | 0 | 0 | 0 | 1 | 0 | 0 | 0 | 2 | 3 |
| ENSMUSG00000025358 | Cdk2 | ENSMUST00000026416 | 22.9648672 | 33 | 21.7652099 | 22.3650386 | 1 | 0 | 0 | 1 | 0 | 0 | 0 | 0 | 1 | 2 | 0 | 1 | 4 | 0 | 8 |
| ENSMUSG00000069089 | Cdk7 | ENSMUST00000091299 | 26.8175477 | 30 | 21.4075337 | 21.6555573 | 0 | 0 | 0 | 1 | 0 | 0 | 0 | 0 | 1 | 2 | 7 | 0 | 10 | 0 | 19 |
| ENSMUSG00000031292 | Cdkl5 | ENSMUST00000198931 | 26.7322436 | 33 | 20.0057862 | 20.3095617 | 2 | 1 | 0 | 2 | 0 | 0 | 0 | 0 | 2 | 4 | 0 | 2 | 1 | 0 | 10 |
| ENSMUSG00000003031 | Cdkn1b | ENSMUST00000067327 | 29.5434719 | 33 | 18.1157313 | 19.6277988 | 0 | 0 | 0 | 1 | 0 | 0 | 0 | 0 | 1 | 2 | 0 | 1 | 15 | 4 | 22 |
| ENSMUSG00000037664 | Cdkn1c | ENSMUST00000037287 | 30.8714919 | 26 | 20.5317578 | 22.1565731 | 0 | 0 | 2 | 0 | 0 | 0 | 0 | 0 | 0 | 2 | 0 | 0 | 0 | 0 | 2 |
| ENSMUSG00000038069 | Cdkn2aip | ENSMUST00000038738 | 31.7489336 | 39 | 12.6142596 | 16.7580743 | 0 | 0 | 0 | 0 | 0 | 0 | 0 | 0 | 0 | 0 | 0 | 0 | 14 | 0 | 14 |
| ENSMUSG00000038119 | Cdon | ENSMUST00000119129 | 25.495338 | 31 | 20.8041958 | 22.7564103 | 1 | 0 | 1 | 1 | 0 | 0 | 0 | 0 | 1 | 3 | 0 | 3 | 3 | 0 | 10 |
| ENSMUSG00000030878 | Cdr2 | ENSMUST00000033169 | 28.4280936 | 28 | 22.7424749 | 20.6243032 | 0 | 0 | 0 | 0 | 0 | 0 | 0 | 0 | 0 | 0 | 0 | 1 | 0 | 0 | 1 |
| ENSMUSG00000058793 | Cds2 | ENSMUST00000089461 | 21.2564671 | 28 | 23.7989653 | 27.0509978 | 0 | 0 | 0 | 0 | 0 | 0 | 0 | 0 | 0 | 0 | 0 | 3 | 2 | 0 | 5 |
| ENSMUSG00000032803 | Cdv3 | ENSMUST00000116517 | 27.766323 | 31 | 18.9003436 | 22.3024055 | 0 | 0 | 0 | 0 | 0 | 0 | 0 | 0 | 0 | 0 | 0 | 0 | 16 | 1 | 17 |
| ENSMUSG00000031758 | Cdyl2 | ENSMUST00000109102 | 24.4808743 | 31 | 21.7954723 | 22.8571429 | 2 | 1 | 0 | 4 | 0 | 0 | 0 | 1 | 3 | 8 | 0 | 23 | 14 | 2 | 50 |
| ENSMUSG00000056216 | Cebpg | ENSMUST00000130491 | 26.8046441 | 33 | 17.3649672 | 23.2710752 | 0 | 0 | 0 | 0 | 0 | 0 | 0 | 0 | 0 | 0 | 2 | 0 | 3 | 1 | 6 |
| ENSMUSG00000005506 | Celf1 | ENSMUST00000068726 | 24.1356817 | 31 | 21.4285714 | 23.1898239 | 0 | 0 | 0 | 1 | 0 | 0 | 0 | 0 | 1 | 2 | 1 | 5 | 6 | 0 | 14 |
| ENSMUSG00000002107 | Celf2 | ENSMUST00000170438 | 27.2998239 | 32 | 21.2708305 | 19.076006 | 0 | 0 | 0 | 1 | 0 | 0 | 0 | 0 | 1 | 2 | 0 | 6 | 2 | 1 | 11 |
| ENSMUSG00000029177 | Cenpa | ENSMUST00000031073 | 22.9828851 | 28 | 23.594132 | 25.5501222 | 0 | 0 | 0 | 1 | 0 | 0 | 0 | 0 | 1 | 2 | 0 | 0 | 0 | 0 | 2 |
| ENSMUSG00000029253 | Cenpc1 | ENSMUST00000031170 | 32.6180258 | 46 | 9.01287554 | 12.4463519 | 0 | 0 | 0 | 0 | 0 | 0 | 0 | 0 | 0 | 0 | 0 | 0 | 1 | 0 | 1 |
| ENSMUSG00000064128 | Cenpj | ENSMUST00000065302 | 32.278481 | 42 | 10.1265823 | 15.1898734 | 0 | 0 | 1 | 1 | 0 | 0 | 0 | 0 | 1 | 3 | 0 | 0 | 0 | 0 | 3 |
| ENSMUSG00000026708 | Cenpl | ENSMUST00000195571 | 28.7333522 | 32 | 18.8722018 | 20.034004 | 1 | 0 | 0 | 0 | 0 | 0 | 0 | 0 | 0 | 0 | 0 | 2 | 9 | 1 | 13 |
| ENSMUSG00000031629 | Cenpu | ENSMUST00000135432 | 31.3070977 | 25 | 20.7308503 | 23.1201687 | 0 | 0 | 0 | 0 | 0 | 0 | 0 | 0 | 0 | 0 | 0 | 0 | 0 | 1 | 1 |
| ENSMUSG00000075266 | Cenpw | ENSMUST00000099985 | 32.4642557 | 37 | 14.7182506 | 15.7275021 | 1 | 0 | 0 | 0 | 0 | 0 | 0 | 0 | 0 | 0 | 0 | 0 | 0 | 0 | 1 |
| ENSMUSG00000036403 | Cep135 | ENSMUST00000121979 | 30.0650054 | 32 | 19.1224269 | 18.4723727 | 0 | 0 | 0 | 0 | 0 | 0 | 0 | 0 | 0 | 0 | 0 | 0 | 0 | 2 | 2 |

| | | | | | | | | | | | | | | | | | | | | | |
|---|---|---|---|---|---|---|---|---|---|---|---|---|---|---|---|---|---|---|---|---|---|
| ENSMUSG00000057335 | Cep170 | ENSMUST00000195433 | 31.3604535 | 25 | 21.6738913 | 21.9073024 | 2 | 1 | 0 | 3 | 0 | 0 | 0 | 1 | 2 | 6 | 1 | 1 | 2 | 0 | 13 |
| ENSMUSG00000019971 | Cep290 | ENSMUST00000164751 | 36.4963504 | 38 | 12.9562044 | 12.5912409 | 1 | 0 | 0 | 1 | 0 | 0 | 0 | 0 | 1 | 2 | 0 | 0 | 0 | 2 | 5 |
| ENSMUSG00000038215 | Cep44 | ENSMUST00000040330 | 30.952381 | 46 | 10.7142857 | 11.9047619 | 0 | 0 | 1 | 0 | 0 | 0 | 0 | 0 | 0 | 1 | 0 | 0 | 0 | 0 | 1 |
| ENSMUSG00000024989 | Cep55 | ENSMUST00000096096 | 26.8608414 | 38 | 16.828479 | 18.3387271 | 0 | 0 | 0 | 1 | 0 | 0 | 0 | 0 | 1 | 2 | 0 | 0 | 3 | 1 | 6 |
| ENSMUSG00000056267 | Cep70 | ENSMUST00000190715 | 26.7389341 | 31 | 20.3252033 | 21.5898826 | 0 | 0 | 0 | 0 | 0 | 0 | 0 | 0 | 0 | 0 | 0 | 1 | 0 | 0 | 1 |
| ENSMUSG00000021572 | Cep72 | ENSMUST00000036456 | 25.1798561 | 35 | 20.2038369 | 20.0239808 | 0 | 0 | 0 | 1 | 0 | 0 | 0 | 0 | 1 | 2 | 1 | 0 | 0 | 0 | 3 |
| ENSMUSG00000073542 | Cep76 | ENSMUST00000097542 | 31.4099075 | 32 | 16.4942842 | 20.0870985 | 5 | 1 | 0 | 3 | 0 | 0 | 0 | 1 | 2 | 6 | 0 | 0 | 0 | 0 | 12 |
| ENSMUSG00000022604 | Cep97 | ENSMUST00000023270 | 27.3106061 | 34 | 17.7083333 | 21.25 | 1 | 0 | 0 | 1 | 0 | 0 | 0 | 0 | 1 | 2 | 6 | 3 | 6 | 0 | 18 |
| ENSMUSG00000040774 | Cept1 | ENSMUST00000068301 | 29.1575092 | 35 | 15.970696 | 20.3663004 | 0 | 0 | 0 | 1 | 0 | 0 | 0 | 0 | 1 | 2 | 0 | 0 | 0 | 1 | 3 |
| ENSMUSG00000008206 | Cers4 | ENSMUST00000008350 | 28.5822785 | 30 | 18.4810127 | 22.4810127 | 0 | 0 | 0 | 0 | 0 | 0 | 0 | 0 | 0 | 0 | 9 | 13 | 8 | 0 | 30 |
| ENSMUSG00000023021 | Cers5 | ENSMUST00000109035 | 25.9816754 | 30 | 21.9240838 | 22.447644 | 0 | 0 | 0 | 0 | 0 | 0 | 0 | 0 | 0 | 0 | 0 | 0 | 1 | 0 | 1 |
| ENSMUSG00000027035 | Cers6 | ENSMUST00000028426 | 27.0745429 | 30 | 21.5014065 | 21.7475387 | 1 | 0 | 1 | 1 | 0 | 0 | 0 | 0 | 1 | 3 | 2 | 2 | 10 | 0 | 18 |
| ENSMUSG00000031347 | Cetn2 | ENSMUST00000114551 | 25.8928571 | 36 | 18.8988095 | 19.047619 | 0 | 0 | 0 | 0 | 0 | 0 | 0 | 0 | 0 | 0 | 0 | 1 | 0 | 0 | 1 |
| ENSMUSG00000020462 | Cfap36 | ENSMUST00000020754 | 39.4886364 | 26 | 17.0454545 | 17.6136364 | 0 | 0 | 0 | 1 | 0 | 0 | 0 | 0 | 1 | 2 | 0 | 0 | 0 | 0 | 2 |
| ENSMUSG00000031631 | Cfap97 | ENSMUST00000034048 | 28.8394719 | 31 | 19.9791522 | 20.3961084 | 0 | 0 | 0 | 1 | 0 | 0 | 0 | 0 | 1 | 2 | 0 | 0 | 1 | 0 | 3 |
| ENSMUSG00000062929 | Cfl2 | ENSMUST00000078124 | 29.0294627 | 37 | 16.1611785 | 17.9809359 | 0 | 0 | 0 | 0 | 0 | 0 | 0 | 0 | 0 | 0 | 1 | 0 | 6 | 0 | 7 |
| ENSMUSG00000026031 | Cflar | ENSMUST00000114313 | 26.2998791 | 31 | 19.2462717 | 23.9016526 | 2 | 1 | 0 | 5 | 0 | 0 | 1 | 1 | 3 | 10 | 7 | 28 | 13 | 0 | 61 |
| ENSMUSG00000041301 | Cftr | ENSMUST00000140407 | 30.4491413 | 29 | 19.1985909 | 21.7084985 | 1 | 0 | 0 | 1 | 0 | 0 | 0 | 0 | 1 | 2 | 1 | 0 | 0 | 0 | 4 |
| ENSMUSG00000054604 | Cggbp1 | ENSMUST00000067744 | 29.0585835 | 35 | 16.6132323 | 18.944914 | 1 | 0 | 0 | 2 | 0 | 0 | 0 | 0 | 2 | 4 | 0 | 1 | 1 | 2 | 9 |
| ENSMUSG00000023852 | Chd1 | ENSMUST00000024627 | 29.6458242 | 36 | 17.2715348 | 17.5776126 | 0 | 0 | 0 | 0 | 0 | 0 | 0 | 0 | 0 | 0 | 0 | 0 | 3 | 6 | 9 |
| ENSMUSG00000078671 | Chd2 | ENSMUST00000199809 | 26.5989628 | 24 | 22.4503025 | 27.1607606 | 0 | 0 | 0 | 1 | 0 | 0 | 0 | 0 | 1 | 2 | 0 | 1 | 0 | 0 | 3 |
| ENSMUSG00000032113 | Chek1 | ENSMUST00000174105 | 30.5210918 | 30 | 18.6600496 | 20.4962779 | 0 | 0 | 1 | 1 | 0 | 0 | 0 | 0 | 1 | 3 | 0 | 0 | 0 | 0 | 3 |
| ENSMUSG00000031327 | Chic1 | ENSMUST00000116547 | 30.4770586 | 34 | 17.5782437 | 18.3834701 | 0 | 0 | 0 | 0 | 0 | 0 | 0 | 0 | 0 | 0 | 0 | 0 | 1 | 0 | 1 |
| ENSMUSG00000029229 | Chic2 | ENSMUST00000075452 | 28.6074499 | 31 | 19.9083095 | 20.034384 | 0 | 0 | 0 | 0 | 0 | 0 | 0 | 0 | 0 | 0 | 6 | 2 | 13 | 1 | 22 |
| ENSMUSG00000024843 | Chka | ENSMUST00000128694 | 26.5251989 | 27 | 22.1750663 | 24.2970822 | 0 | 0 | 0 | 0 | 0 | 0 | 0 | 0 | 0 | 0 | 0 | 2 | 0 | 0 | 2 |
| ENSMUSG00000030077 | Chl1 | ENSMUST00000066905 | 29.89468 | 35 | 17.0132325 | 18.2824737 | 0 | 0 | 0 | 1 | 0 | 0 | 0 | 0 | 1 | 2 | 4 | 28 | 7 | 1 | 42 |
| ENSMUSG00000025531 | Chm | ENSMUST00000026607 | 33.532094 | 33 | 17.1168011 | 16.5906699 | 0 | 0 | 0 | 0 | 0 | 0 | 0 | 0 | 0 | 0 | 1 | 0 | 1 | 0 | 2 |
| ENSMUSG00000004843 | Chmp2b | ENSMUST00000004965 | 30.4865938 | 33 | 17.1797418 | 19.0665343 | 0 | 0 | 0 | 0 | 0 | 0 | 0 | 0 | 0 | 0 | 0 | 0 | 2 | 0 | 2 |
| ENSMUSG00000053119 | Chmp3 | ENSMUST00000059462 | 23.2232792 | 32 | 21.8242865 | 22.7756016 | 0 | 0 | 0 | 0 | 0 | 0 | 0 | 0 | 0 | 0 | 0 | 0 | 1 | 0 | 1 |
| ENSMUSG00000027536 | Chmp4c | ENSMUST00000029049 | 27.6472782 | 28 | 19.6308725 | 24.3847875 | 0 | 0 | 0 | 0 | 0 | 0 | 0 | 0 | 0 | 0 | 6 | 0 | 4 | 0 | 10 |
| ENSMUSG00000028419 | Chmp5 | ENSMUST00000030128 | 28.4194529 | 38 | 15.3495441 | 18.6930091 | 0 | 1 | 0 | 0 | 0 | 0 | 0 | 0 | 0 | 0 | 0 | 0 | 2 | 0 | 3 |
| ENSMUSG00000022860 | Chodl | ENSMUST00000023568 | 31.1145511 | 34 | 16.1764706 | 18.5758514 | 0 | 0 | 0 | 0 | 0 | 0 | 0 | 0 | 0 | 0 | 0 | 1 | 1 | 0 | 2 |
| ENSMUSG00000001774 | Chordc1 | ENSMUST00000001825 | 28.3595922 | 37 | 15.6626506 | 19.0917516 | 0 | 0 | 0 | 0 | 0 | 0 | 0 | 0 | 0 | 0 | 0 | 0 | 3 | 0 | 3 |
| ENSMUSG00000014077 | Chp1 | ENSMUST00000119172 | 23.2658337 | 33 | 22.447221 | 21.1977596 | 0 | 0 | 0 | 0 | 0 | 0 | 0 | 0 | 0 | 0 | 0 | 1 | 0 | 0 | 1 |
| ENSMUSG00000060002 | Chpt1 | ENSMUST00000117440 | 37.3029772 | 30 | 15.7034442 | 16.5207239 | 0 | 0 | 0 | 0 | 0 | 0 | 0 | 0 | 0 | 0 | 0 | 0 | 0 | 3 | 3 |
| ENSMUSG00000045613 | Chrm2 | ENSMUST00000172278 | 31.8611987 | 31 | 18.7171399 | 18.3491062 | 0 | 0 | 0 | 0 | 0 | 0 | 0 | 0 | 0 | 0 | 1 | 1 | 2 | 0 | 4 |
| ENSMUSG00000027221 | Chst1 | ENSMUST00000065797 | 20.4819277 | 29 | 26.746988 | 23.9759036 | 0 | 0 | 0 | 0 | 0 | 0 | 0 | 0 | 0 | 0 | 0 | 3 | 0 | 0 | 3 |
| ENSMUSG00000074916 | Chst14 | ENSMUST00000099546 | 23.6612702 | 29 | 24.5330012 | 22.6650062 | 0 | 1 | 0 | 1 | 0 | 0 | 0 | 0 | 1 | 2 | 0 | 0 | 0 | 0 | 3 |
| ENSMUSG00000001017 | Chtop | ENSMUST00000001043 | 25.6871036 | 34 | 18.9217759 | 21.564482 | 0 | 0 | 0 | 0 | 0 | 0 | 0 | 0 | 0 | 0 | 0 | 0 | 1 | 0 | 1 |
| ENSMUSG00000025199 | Chuk | ENSMUST00000119591 | 28.8834951 | 36 | 15.9385113 | 19.3365696 | 0 | 0 | 0 | 0 | 0 | 0 | 0 | 0 | 0 | 0 | 1 | 0 | 1 | 0 | 2 |
| ENSMUSG00000003662 | Ciao1 | ENSMUST00000003759 | 31.4685315 | 31 | 18.2356105 | 19.6342119 | 0 | 0 | 0 | 1 | 0 | 0 | 0 | 0 | 1 | 2 | 0 | 0 | 0 | 4 | 6 |
| ENSMUSG00000031781 | Ciapin1 | ENSMUST00000162538 | 22.5068618 | 27 | 23.9707228 | 26.3799939 | 0 | 0 | 0 | 0 | 0 | 0 | 0 | 0 | 0 | 0 | 0 | 4 | 0 | 0 | 4 |
| ENSMUSG00000041777 | Cir1 | ENSMUST00000058615 | 29.8007968 | 33 | 15.8565737 | 21.8326693 | 0 | 0 | 0 | 0 | 0 | 0 | 0 | 0 | 0 | 0 | 0 | 0 | 2 | 3 | 5 |
| ENSMUSG00000041438 | Cirh1a | ENSMUST00000047629 | 21.4285714 | 46 | 16.0714286 | 16.9642857 | 0 | 0 | 0 | 0 | 0 | 0 | 0 | 0 | 0 | 0 | 4 | 0 | 5 | 0 | 9 |
| ENSMUSG00000037725 | Ckap2 | ENSMUST00000046916 | 29.1882556 | 36 | 16.5803109 | 18.4801382 | 0 | 0 | 0 | 1 | 0 | 0 | 0 | 0 | 1 | 2 | 0 | 0 | 1 | 0 | 3 |
| ENSMUSG00000033392 | Clasp2 | ENSMUST00000111838 | 26.1138614 | 36 | 18.5643564 | 19.6782178 | 0 | 0 | 0 | 0 | 0 | 0 | 0 | 0 | 0 | 0 | 0 | 0 | 15 | 2 | 17 |
| ENSMUSG00000004319 | Clcn3 | ENSMUST00000093490 | 27.5222552 | 34 | 18.694362 | 20.1409496 | 1 | 0 | 0 | 2 | 0 | 0 | 0 | 0 | 2 | 4 | 0 | 1 | 3 | 2 | 11 |
| ENSMUSG00000004317 | Clcn5 | ENSMUST00000004428 | 27.1498564 | 30 | 21.2366954 | 21.5070113 | 0 | 0 | 0 | 1 | 0 | 0 | 0 | 0 | 1 | 2 | 0 | 6 | 4 | 0 | 12 |
| ENSMUSG00000022512 | Cldn1 | ENSMUST00000023154 | 28.2971619 | 34 | 19.4490818 | 18.4056761 | 1 | 0 | 0 | 1 | 0 | 0 | 0 | 0 | 1 | 2 | 0 | 0 | 0 | 0 | 3 |
| ENSMUSG00000079450 | Cldn34c1 | ENSMUST00000113348 | 31.3338926 | 32 | 17.9949664 | 18.3724832 | 0 | 0 | 0 | 1 | 0 | 0 | 0 | 0 | 1 | 2 | 2 | 30 | 3 | 0 | 37 |
| ENSMUSG00000022744 | Cldnd1 | ENSMUST00000023426 | 29.0268456 | 35 | 19.3791946 | 16.3590604 | 2 | 1 | 0 | 1 | 0 | 0 | 0 | 0 | 1 | 2 | 1 | 1 | 1 | 1 | 9 |
| ENSMUSG00000043832 | Clec4a3 | ENSMUST00000117173 | 27.5482094 | 34 | 21.7630854 | 16.5289256 | 0 | 0 | 0 | 0 | 0 | 0 | 0 | 0 | 0 | 0 | 0 | 8 | 0 | 0 | 8 |
| ENSMUSG00000024059 | Clip4 | ENSMUST00000024854 | 29.6892342 | 32 | 18.0355161 | 19.9778024 | 0 | 0 | 0 | 0 | 0 | 0 | 0 | 0 | 0 | 0 | 0 | 0 | 0 | 4 | 4 |
| ENSMUSG00000068917 | Clk2 | ENSMUST00000128318 | 25.9286776 | 24 | 26.1515602 | 23.9970282 | 0 | 0 | 0 | 0 | 0 | 0 | 0 | 0 | 0 | 0 | 0 | 1 | 0 | 0 | 1 |
| ENSMUSG00000030720 | Cln3 | ENSMUST00000084589 | 27.8301887 | 31 | 20.3773585 | 20.4716981 | 0 | 0 | 0 | 0 | 0 | 0 | 0 | 0 | 0 | 0 | 0 | 1 | 0 | 0 | 1 |
| ENSMUSG00000029238 | Clock | ENSMUST00000202651 | 29.1172596 | 32 | 18.6795491 | 20.1581028 | 1 | 1 | 0 | 2 | 0 | 0 | 0 | 0 | 2 | 4 | 1 | 0 | 5 | 1 | 13 |
| ENSMUSG00000039163 | Cmc1 | ENSMUST00000044220 | 28.8984263 | 32 | 17.8826896 | 21.316166 | 1 | 0 | 0 | 0 | 0 | 0 | 0 | 0 | 0 | 0 | 0 | 0 | 0 | 0 | 1 |
| ENSMUSG00000028719 | Cmpk1 | ENSMUST00000030491 | 27.2918419 | 37 | 16.9049622 | 18.9655172 | 2 | 1 | 0 | 2 | 0 | 0 | 0 | 0 | 2 | 4 | 2 | 0 | 6 | 3 | 18 |
| ENSMUSG00000041012 | Cmtm8 | ENSMUST00000047013 | 35.9621451 | 29 | 17.0347003 | 17.6656151 | 0 | 0 | 1 | 0 | 0 | 0 | 0 | 0 | 0 | 1 | 0 | 0 | 0 | 0 | 1 |
| ENSMUSG00000046441 | Cmtr2 | ENSMUST00000056972 | 27.5155833 | 37 | 16.4737311 | 19.0560997 | 0 | 1 | 2 | 0 | 0 | 0 | 0 | 0 | 0 | 2 | 0 | 0 | 0 | 0 | 3 |
| ENSMUSG00000030057 | Cnbp | ENSMUST00000032138 | 30.5477528 | 30 | 16.2921348 | 22.752809 | 0 | 0 | 0 | 0 | 0 | 0 | 0 | 0 | 0 | 0 | 0 | 0 | 1 | 0 | 1 |
| ENSMUSG00000036810 | Cnep1r1 | ENSMUST00000121097 | 25.8817146 | 30 | 21.5409658 | 22.9517092 | 0 | 0 | 0 | 0 | 0 | 0 | 0 | 0 | 0 | 0 | 1 | 0 | 0 | 1 | 2 |
| ENSMUSG00000062169 | Cnih4 | ENSMUST00000134115 | 27.2433597 | 28 | 21.3567839 | 23.3668342 | 1 | 0 | 0 | 0 | 0 | 0 | 0 | 0 | 0 | 0 | 0 | 0 | 0 | 0 | 1 |
| ENSMUSG00000020362 | Cnot6 | ENSMUST00000145353 | 28.6720867 | 36 | 16.3143631 | 18.7804878 | 1 | 0 | 0 | 1 | 0 | 0 | 0 | 0 | 1 | 2 | 2 | 0 | 3 | 0 | 8 |
| ENSMUSG00000034724 | Cnot6l | ENSMUST00000155901 | 28.8739594 | 35 | 17.0877757 | 18.7965532 | 2 | 0 | 0 | 1 | 0 | 0 | 0 | 0 | 1 | 2 | 1 | 3 | 9 | 0 | 17 |
| ENSMUSG00000020515 | Cnot8 | ENSMUST00000020822 | 23.9562624 | 30 | 22.8628231 | 23.2604374 | 1 | 0 | 0 | 0 | 0 | 0 | 0 | 0 | 0 | 0 | 0 | 0 | 0 | 0 | 1 |
| ENSMUSG00000044681 | Cnpy1 | ENSMUST00000141601 | 29.1085271 | 31 | 18.7984496 | 21.5116279 | 2 | 0 | 0 | 0 | 0 | 0 | 0 | 0 | 0 | 0 | 1 | 0 | 0 | 1 | 4 |
| ENSMUSG00000044288 | Cnr1 | ENSMUST00000057188 | 29.2676502 | 32 | 18.5985248 | 19.6786091 | 0 | 0 | 0 | 0 | 0 | 0 | 0 | 0 | 0 | 0 | 1 | 0 | 8 | 0 | 9 |
| ENSMUSG00000038949 | Cnst | ENSMUST00000040706 | 25.769569 | 29 | 21.5919085 | 23.175022 | 1 | 1 | 0 | 0 | 0 | 0 | 0 | 0 | 0 | 0 | 0 | 0 | 0 | 1 | 3 |
| ENSMUSG00000028444 | Cntfr | ENSMUST00000102962 | 18.8073394 | 24 | 35.3211009 | 22.324159 | 0 | 0 | 0 | 0 | 0 | 0 | 0 | 0 | 0 | 0 | 0 | 2 | 0 | 0 | 2 |
| ENSMUSG00000055022 | Cntn1 | ENSMUST00000169825 | 30.4878049 | 34 | 17.1152229 | 17.9983179 | 0 | 0 | 0 | 0 | 0 | 0 | 0 | 0 | 0 | 0 | 1 | 0 | 5 | 0 | 6 |
| ENSMUSG00000070695 | Cntnap5a | ENSMUST00000043725 | 30.3053091 | 32 | 18.6945405 | 18.7998195 | 0 | 0 | 0 | 0 | 0 | 0 | 0 | 0 | 0 | 0 | 1 | 2 | 3 | 1 | 7 |
| ENSMUSG00000034903 | Cobll1 | ENSMUST00000112429 | 33.1272085 | 36 | 15.1060071 | 16.254417 | 0 | 0 | 0 | 1 | 0 | 0 | 0 | 0 | 1 | 2 | 0 | 0 | 0 | 0 | 2 |
| ENSMUSG00000034893 | Cog3 | ENSMUST00000049168 | 25.2380952 | 28 | 20.952381 | 26.1904762 | 0 | 0 | 0 | 1 | 0 | 0 | 0 | 0 | 1 | 2 | 1 | 0 | 0 | 0 | 3 |
| ENSMUSG00000035933 | Cog5 | ENSMUST00000036862 | 33.744856 | 32 | 17.6954733 | 16.255144 | 0 | 0 | 0 | 0 | 0 | 0 | 0 | 0 | 0 | 0 | 0 | 0 | 1 | 0 | 1 |
| ENSMUSG00000027966 | Col11a1 | ENSMUST00000092155 | 33.6858805 | 35 | 16.3934426 | 15.2829191 | 0 | 0 | 0 | 1 | 0 | 0 | 0 | 0 | 1 | 2 | 0 | 0 | 0 | 0 | 2 |
| ENSMUSG00000058897 | Col25a1 | ENSMUST00000106353 | 26.7622461 | 35 | 18.6857826 | 19.4026284 | 0 | 1 | 0 | 0 | 0 | 0 | 0 | 0 | 0 | 0 | 1 | 0 | 1 | 0 | 3 |
| ENSMUSG00000021669 | Col4a3bp | ENSMUST00000077672 | 30.7943772 | 32 | 16.9990193 | 20.0392285 | 0 | 0 | 0 | 0 | 0 | 0 | 0 | 0 | 0 | 0 | 0 | 0 | 0 | 1 | 1 |
| ENSMUSG00000042705 | Commd10 | ENSMUST00000049388 | 26.5658747 | 32 | 22.2462203 | 19.3304536 | 0 | 0 | 0 | 0 | 0 | 0 | 0 | 0 | 0 | 0 | 0 | 1 | 0 | 0 | 1 |
| ENSMUSG00000036513 | Commd2 | ENSMUST00000160959 | 27.2961373 | 33 | 21.1158798 | 18.4120172 | 2 | 0 | 0 | 0 | 0 | 0 | 0 | 0 | 0 | 0 | 2 | 2 | 3 | 0 | 9 |
| ENSMUSG00000075486 | Commd6 | ENSMUST00000100339 | 34.2013889 | 30 | 17.5347222 | 18.4027778 | 1 | 0 | 0 | 0 | 0 | 0 | 0 | 0 | 0 | 0 | 0 | 0 | 0 | 0 | 1 |
| ENSMUSG00000029213 | Commd8 | ENSMUST00000013693 | 25.3790176 | 33 | 20.6488781 | 20.6488781 | 3 | 1 | 0 | 2 | 0 | 0 | 0 | 1 | 1 | 4 | 0 | 0 | 9 | 0 | 17 |
| ENSMUSG00000027206 | Cops2 | ENSMUST00000110463 | 32.186342 | 33 | 15.8814187 | 19.0047644 | 0 | 0 | 0 | 0 | 0 | 0 | 0 | 0 | 0 | 0 | 0 | 0 | 0 | 2 | 2 |
| ENSMUSG00000025981 | Coq10b | ENSMUST00000027125 | 26.807564 | 38 | 16.1290323 | 19.0211346 | 0 | 0 | 0 | 0 | 0 | 0 | 0 | 0 | 0 | 0 | 0 | 0 | 3 | 3 | 6 |
| ENSMUSG00000028247 | Coq3 | ENSMUST00000029909 | 28.5620486 | 32 | 19.3696651 | 20.4858831 | 1 | 0 | 0 | 0 | 0 | 0 | 0 | 0 | 0 | 0 | 0 | 4 | 0 | 1 | 6 |
| ENSMUSG00000020544 | Cox11 | ENSMUST00000020851 | 30.6068602 | 34 | 16.6226913 | 19.2612137 | 0 | 0 | 0 | 0 | 0 | 0 | 0 | 0 | 0 | 0 | 0 | 1 | 0 | 0 | 1 |
| ENSMUSG00000040018 | Cox15 | ENSMUST00000045562 | 27.6608018 | 28 | 21.9498125 | 22.5266801 | 0 | 0 | 0 | 1 | 0 | 0 | 0 | 0 | 1 | 2 | 6 | 1 | 3 | 1 | 13 |
| ENSMUSG00000091803 | Cox16 | ENSMUST00000002757 | 30.1110385 | 35 | 19.0071848 | 15.610712 | 0 | 0 | 0 | 0 | 0 | 0 | 0 | 0 | 0 | 0 | 1 | 0 | 3 | 2 | 6 |
| ENSMUSG00000046516 | Cox17 | ENSMUST00000120495 | 26.3527347 | 29 | 21.8192454 | 23.3109096 | 0 | 0 | 0 | 1 | 0 | 0 | 0 | 0 | 1 | 2 | 0 | 3 | 16 | 0 | 21 |
| ENSMUSG00000024248 | Cox7a2l | ENSMUST00000025095 | 23.6842105 | 28 | 19.5906433 | 28.3625731 | 0 | 0 | 0 | 0 | 0 | 0 | 0 | 0 | 0 | 0 | 0 | 0 | 1 | 0 | 1 |
| ENSMUSG00000031231 | Cox7b | ENSMUST00000033582 | 30.1435407 | 32 | 18.5007974 | 19.2982456 | 0 | 1 | 0 | 0 | 0 | 0 | 0 | 0 | 0 | 0 | 0 | 0 | 0 | 0 | 1 |
| ENSMUSG00000020841 | Cpd | ENSMUST00000021201 | 26.3829787 | 33 | 20.5585106 | 20.0265957 | 0 | 0 | 1 | 0 | 0 | 0 | 0 | 0 | 0 | 1 | 0 | 1 | 1 | 1 | 4 |

| | | | | | | | | | | | | | | | | | | | | | |
|---|---|---|---|---|---|---|---|---|---|---|---|---|---|---|---|---|---|---|---|---|---|
| ENSMUSG00000039782 | Cpeb2 | ENSMUST00000114065 | 28.798306 | 38 | 15.5108523 | 17.8930651 | 1 | 0 | 0 | 0 | 0 | 0 | 0 | 0 | 0 | 0 | 2 | 2 | 2 | 1 | 8 |
| ENSMUSG00000039652 | Cpeb3 | ENSMUST00000079754 | 27.4306528 | 33 | 18.5766321 | 20.8181563 | 0 | 0 | 0 | 0 | 0 | 0 | 0 | 0 | 0 | 0 | 1 | 1 | 1 | 1 | 4 |
| ENSMUSG00000020300 | Cpeb4 | ENSMUST00000109412 | 29.985373 | 36 | 16.6260361 | 17.7230619 | 0 | 1 | 0 | 0 | 0 | 0 | 0 | 0 | 0 | 0 | 0 | 2 | 2 | 2 | 7 |
| ENSMUSG00000074643 | Cpne1 | ENSMUST00000109607 | 25.6563246 | 34 | 20.1073986 | 20.2267303 | 0 | 0 | 0 | 0 | 0 | 0 | 0 | 0 | 0 | 0 | 0 | 1 | 0 | 0 | 1 |
| ENSMUSG00000028228 | Cpne3 | ENSMUST00000029885 | 28.2697201 | 35 | 17.9643766 | 18.9312977 | 1 | 1 | 0 | 1 | 0 | 0 | 0 | 0 | 1 | 2 | 2 | 1 | 4 | 2 | 13 |
| ENSMUSG00000022742 | Cpox | ENSMUST00000060077 | 27.8457772 | 35 | 17.9314565 | 19.0942472 | 0 | 0 | 0 | 2 | 0 | 0 | 0 | 0 | 2 | 4 | 0 | 0 | 0 | 7 | 11 |
| ENSMUSG00000055531 | Cpsf6 | ENSMUST00000069168 | 29.741559 | 37 | 14.7769904 | 18.5285536 | 0 | 0 | 0 | 1 | 0 | 0 | 0 | 0 | 1 | 2 | 0 | 0 | 1 | 5 | 8 |
| ENSMUSG00000026616 | Cr2 | ENSMUST00000195120 | 31.6033058 | 30 | 16.0330579 | 21.9834711 | 0 | 0 | 0 | 1 | 0 | 0 | 0 | 0 | 1 | 2 | 0 | 0 | 0 | 0 | 2 |
| ENSMUSG00000025958 | Creb1 | ENSMUST00000087366 | 29.3723849 | 33 | 17.4198047 | 20.237099 | 2 | 0 | 0 | 1 | 0 | 0 | 0 | 0 | 1 | 2 | 0 | 0 | 4 | 6 | 14 |
| ENSMUSG00000053007 | Creb5 | ENSMUST00000203528 | 28.4623947 | 31 | 21.2275401 | 19.6056607 | 0 | 0 | 0 | 1 | 0 | 0 | 0 | 0 | 1 | 2 | 2 | 1 | 3 | 2 | 10 |
| ENSMUSG00000032652 | Crebl2 | ENSMUST00000046303 | 23.8510567 | 31 | 21.1338477 | 24.119423 | 0 | 0 | 0 | 0 | 0 | 0 | 0 | 0 | 0 | 0 | 1 | 0 | 7 | 2 | 10 |
| ENSMUSG00000048249 | Crebrf | ENSMUST00000062519 | 28.5768936 | 35 | 17.8079572 | 18.7260903 | 0 | 0 | 0 | 0 | 0 | 0 | 0 | 0 | 0 | 0 | 3 | 1 | 3 | 2 | 9 |
| ENSMUSG00000051451 | Crebzf | ENSMUST00000107206 | 30.1158301 | 36 | 15.2785438 | 18.9464975 | 1 | 0 | 0 | 0 | 0 | 0 | 0 | 0 | 0 | 0 | 2 | 0 | 2 | 2 | 7 |
| ENSMUSG00000063889 | Crem | ENSMUST00000025069 | 28.422345 | 33 | 17.3112339 | 20.9330878 | 0 | 1 | 0 | 0 | 0 | 0 | 0 | 0 | 0 | 0 | 0 | 0 | 0 | 0 | 1 |
| ENSMUSG00000024074 | Crim1 | ENSMUST00000112498 | 30.1694915 | 32 | 16.3559322 | 21.1016949 | 0 | 0 | 0 | 1 | 0 | 0 | 0 | 0 | 1 | 2 | 0 | 0 | 4 | 1 | 7 |
| ENSMUSG00000024146 | Cript | ENSMUST00000024959 | 28.7760417 | 31 | 21.3541667 | 18.4895833 | 0 | 1 | 0 | 0 | 0 | 0 | 0 | 0 | 0 | 0 | 0 | 0 | 0 | 0 | 1 |
| ENSMUSG00000025776 | Crispld1 | ENSMUST00000095075 | 32.1973094 | 34 | 14.2152466 | 19.7757848 | 1 | 1 | 0 | 0 | 0 | 0 | 0 | 0 | 0 | 0 | 0 | 1 | 0 | 0 | 3 |
| ENSMUSG00000006134 | Crkl | ENSMUST00000006293 | 25.1219512 | 29 | 23.6178862 | 22.6829268 | 0 | 0 | 0 | 0 | 0 | 0 | 0 | 0 | 0 | 0 | 0 | 1 | 3 | 0 | 4 |
| ENSMUSG00000017561 | Crlf3 | ENSMUST00000103233 | 26.9461078 | 31 | 18.8195038 | 22.8400342 | 1 | 0 | 0 | 0 | 0 | 0 | 0 | 0 | 0 | 0 | 2 | 0 | 2 | 1 | 6 |
| ENSMUSG00000027357 | Crls1 | ENSMUST00000124834 | 32.6825127 | 31 | 17.3174873 | 19.0152801 | 0 | 0 | 0 | 0 | 0 | 0 | 0 | 0 | 0 | 0 | 0 | 0 | 11 | 1 | 12 |
| ENSMUSG00000001767 | Crnkl1 | ENSMUST00000001818 | 26.7724868 | 36 | 18.4126984 | 18.6243386 | 0 | 0 | 0 | 0 | 0 | 0 | 0 | 0 | 0 | 0 | 0 | 0 | 0 | 1 | 1 |
| ENSMUSG00000068742 | Cry2 | ENSMUST00000090559 | 21.405604 | 26 | 25.9990813 | 26.6881029 | 0 | 0 | 0 | 0 | 0 | 0 | 0 | 0 | 0 | 0 | 1 | 1 | 0 | 0 | 2 |
| ENSMUSG00000005683 | Cs | ENSMUST00000005826 | 25.0185598 | 30 | 24.2761693 | 21.0096511 | 0 | 0 | 0 | 0 | 0 | 0 | 0 | 0 | 0 | 0 | 0 | 0 | 1 | 0 | 1 |
| ENSMUSG00000068823 | Csde1 | ENSMUST00000198174 | 30.0845572 | 30 | 18.3800623 | 21.9403649 | 0 | 0 | 0 | 1 | 0 | 0 | 0 | 0 | 1 | 2 | 0 | 0 | 4 | 0 | 6 |
| ENSMUSG00000024576 | Csnk1a1 | ENSMUST00000165123 | 27.1689498 | 35 | 17.6179604 | 20.0913242 | 0 | 0 | 0 | 0 | 0 | 0 | 0 | 0 | 0 | 0 | 0 | 3 | 2 | 2 | 7 |
| ENSMUSG00000025162 | Csnk1d | ENSMUST00000070575 | 20.9323583 | 27 | 24.0402194 | 27.8793419 | 0 | 0 | 0 | 0 | 0 | 0 | 0 | 0 | 0 | 0 | 0 | 0 | 1 | 0 | 1 |
| ENSMUSG00000032384 | Csnk1g1 | ENSMUST00000034949 | 26.0215453 | 29 | 22.8826152 | 22.2511144 | 0 | 0 | 0 | 0 | 0 | 0 | 0 | 0 | 0 | 0 | 0 | 6 | 7 | 0 | 13 |
| ENSMUSG00000073563 | Csnk1g3 | ENSMUST00000069597 | 27.7987421 | 34 | 18.951782 | 19.706499 | 0 | 0 | 0 | 0 | 0 | 0 | 0 | 0 | 0 | 0 | 0 | 0 | 1 | 0 | 1 |
| ENSMUSG00000074698 | Csnk2a1 | ENSMUST00000099224 | 24.9277457 | 33 | 20.0144509 | 22.1459538 | 0 | 0 | 0 | 0 | 0 | 0 | 0 | 0 | 0 | 0 | 2 | 2 | 15 | 5 | 24 |
| ENSMUSG00000056763 | Cspp1 | ENSMUST00000122156 | 31.5500686 | 37 | 15.6378601 | 16.0493827 | 0 | 0 | 1 | 0 | 0 | 0 | 0 | 0 | 0 | 1 | 0 | 0 | 0 | 0 | 1 |
| ENSMUSG00000044647 | Csrnp3 | ENSMUST00000122912 | 28.4641558 | 32 | 19.7465681 | 19.5823067 | 1 | 0 | 1 | 1 | 0 | 0 | 0 | 0 | 1 | 3 | 0 | 0 | 0 | 1 | 5 |
| ENSMUSG00000031256 | Cstf2 | ENSMUST00000033609 | 31.6866075 | 31 | 18.5642609 | 18.3712852 | 0 | 0 | 0 | 0 | 0 | 0 | 0 | 0 | 0 | 0 | 0 | 1 | 0 | 0 | 1 |
| ENSMUSG00000027176 | Cstf3 | ENSMUST00000028599 | 28.4313725 | 35 | 17.4509804 | 18.8235294 | 0 | 0 | 0 | 0 | 0 | 0 | 0 | 0 | 0 | 0 | 1 | 0 | 1 | 1 | 3 |
| ENSMUSG00000021000 | Ctage5 | ENSMUST00000176752 | 32.5263569 | 25 | 21.007419 | 21.5150332 | 0 | 0 | 0 | 0 | 0 | 0 | 0 | 0 | 0 | 0 | 0 | 3 | 0 | 0 | 3 |
| ENSMUSG00000033411 | Ctdspl2 | ENSMUST00000036647 | 29.1307233 | 36 | 16.0583942 | 18.4472462 | 1 | 1 | 0 | 1 | 0 | 0 | 0 | 0 | 1 | 2 | 0 | 1 | 1 | 2 | 8 |
| ENSMUSG00000042340 | Ctf1 | ENSMUST00000047393 | 18.7239945 | 37 | 25.3814147 | 18.5852982 | 0 | 0 | 0 | 0 | 0 | 0 | 0 | 0 | 0 | 0 | 0 | 1 | 0 | 0 | 1 |
| ENSMUSG00000060034 | Ctf2 | ENSMUST00000076091 | 23.9819005 | 31 | 22.9864253 | 21.9909502 | 1 | 1 | 0 | 0 | 0 | 0 | 0 | 0 | 0 | 0 | 0 | 0 | 0 | 2 | 4 |
| ENSMUSG00000028179 | Cth | ENSMUST00000118539 | 29.9474606 | 38 | 14.8861646 | 16.8126095 | 0 | 0 | 0 | 0 | 0 | 0 | 0 | 0 | 0 | 0 | 0 | 0 | 0 | 2 | 2 |
| ENSMUSG00000031360 | Ctps2 | ENSMUST00000033727 | 27.5574113 | 32 | 21.5031315 | 18.8587335 | 0 | 0 | 0 | 0 | 0 | 0 | 0 | 0 | 0 | 0 | 4 | 0 | 4 | 0 | 8 |
| ENSMUSG00000021477 | Ctsl | ENSMUST00000021933 | 28.280543 | 35 | 16.9683258 | 20.2488688 | 1 | 0 | 0 | 1 | 0 | 0 | 0 | 0 | 1 | 2 | 0 | 1 | 0 | 0 | 4 |
| ENSMUSG00000004364 | Cul3 | ENSMUST00000163119 | 28.9744738 | 37 | 16.2113748 | 17.4205105 | 1 | 0 | 0 | 1 | 0 | 0 | 0 | 0 | 1 | 2 | 0 | 0 | 0 | 0 | 3 |
| ENSMUSG00000032030 | Cul5 | ENSMUST00000120122 | 29.2826221 | 33 | 17.563389 | 20.5318491 | 3 | 0 | 0 | 3 | 0 | 0 | 0 | 0 | 3 | 6 | 3 | 3 | 10 | 1 | 26 |
| ENSMUSG00000004096 | Cwc15 | ENSMUST00000004200 | 27.4193548 | 42 | 16.1290323 | 14.9193548 | 0 | 0 | 0 | 0 | 0 | 0 | 0 | 0 | 0 | 0 | 0 | 1 | 1 | 0 | 2 |
| ENSMUSG00000027014 | Cwc22 | ENSMUST00000065889 | 32.7683616 | 41 | 11.299435 | 14.6892655 | 0 | 0 | 0 | 0 | 0 | 0 | 0 | 0 | 0 | 0 | 0 | 0 | 0 | 1 | 1 |
| ENSMUSG00000025898 | Cwf19l2 | ENSMUST00000027027 | 29.8589342 | 38 | 14.968652 | 17.3981191 | 0 | 0 | 0 | 0 | 0 | 0 | 0 | 0 | 0 | 0 | 1 | 0 | 0 | 0 | 1 |
| ENSMUSG00000022865 | Cxadr | ENSMUST00000023572 | 29 | 33 | 17.3023256 | 20.3488372 | 1 | 1 | 0 | 0 | 0 | 0 | 0 | 0 | 0 | 0 | 0 | 1 | 0 | 1 | 4 |
| ENSMUSG00000034855 | Cxcl10 | ENSMUST00000118006 | 24.5983936 | 26 | 27.1084337 | 21.8875502 | 0 | 0 | 0 | 0 | 0 | 0 | 0 | 0 | 0 | 0 | 0 | 1 | 0 | 0 | 1 |
| ENSMUSG00000061353 | Cxcl12 | ENSMUST00000073043 | 22.6676947 | 27 | 25.7517348 | 24.7108712 | 0 | 0 | 0 | 0 | 0 | 0 | 0 | 0 | 0 | 0 | 5 | 2 | 5 | 0 | 12 |
| ENSMUSG00000045382 | Cxcr4 | ENSMUST00000052172 | 25.621118 | 40 | 15.0621118 | 18.9440994 | 0 | 0 | 0 | 0 | 0 | 0 | 0 | 0 | 0 | 0 | 5 | 0 | 11 | 0 | 16 |
| ENSMUSG00000044365 | Cxxc4 | ENSMUST00000181904 | 30.2411417 | 33 | 17.4458661 | 19.3651575 | 0 | 0 | 0 | 0 | 0 | 0 | 0 | 0 | 0 | 0 | 4 | 1 | 5 | 1 | 11 |
| ENSMUSG00000031924 | Cyb5b | ENSMUST00000034400 | 25.7233035 | 30 | 22.0936349 | 22.3566544 | 0 | 0 | 0 | 0 | 0 | 0 | 0 | 0 | 0 | 0 | 0 | 0 | 0 | 2 | 2 |
| ENSMUSG00000030447 | Cyfip1 | ENSMUST00000032629 | 33.8114754 | 28 | 18.5655738 | 19.6721311 | 0 | 0 | 0 | 0 | 0 | 0 | 0 | 0 | 0 | 0 | 0 | 1 | 0 | 0 | 1 |
| ENSMUSG00000036712 | Cyld | ENSMUST00000043526 | 25.5271695 | 32 | 18.6942417 | 24.0673155 | 1 | 0 | 0 | 0 | 0 | 0 | 0 | 0 | 0 | 0 | 0 | 0 | 1 | 3 | 5 |
| ENSMUSG00000024087 | Cyp1b1 | ENSMUST00000024894 | 30.3821656 | 31 | 17.9936306 | 20.5732484 | 0 | 0 | 0 | 0 | 0 | 0 | 0 | 0 | 0 | 0 | 0 | 0 | 1 | 1 | 2 |
| ENSMUSG00000049439 | Cyp20a1 | ENSMUST00000148443 | 26.7491927 | 30 | 20.452099 | 22.4434876 | 0 | 0 | 0 | 0 | 0 | 0 | 0 | 0 | 0 | 0 | 0 | 0 | 6 | 0 | 6 |
| ENSMUSG00000063415 | Cyp26b1 | ENSMUST00000205228 | 22.3273747 | 27 | 25.3066947 | 25.4118472 | 0 | 0 | 0 | 0 | 0 | 0 | 0 | 0 | 0 | 0 | 0 | 3 | 1 | 0 | 4 |
| ENSMUSG00000052914 | Cyp2j6 | ENSMUST00000030303 | 30.1746956 | 31 | 18.9518264 | 19.8517734 | 0 | 0 | 0 | 0 | 0 | 0 | 0 | 0 | 0 | 0 | 0 | 1 | 0 | 0 | 1 |
| ENSMUSG00000001467 | Cyp51 | ENSMUST00000001507 | 27.2222222 | 32 | 17.4747475 | 23.5858586 | 0 | 0 | 0 | 0 | 0 | 0 | 0 | 0 | 0 | 0 | 1 | 0 | 0 | 0 | 1 |
| ENSMUSG00000028195 | Cyr61 | ENSMUST00000029846 | 26.8115942 | 29 | 17.826087 | 26.5217391 | 0 | 1 | 0 | 1 | 0 | 0 | 0 | 0 | 1 | 2 | 0 | 0 | 0 | 0 | 3 |
| ENSMUSG00000046727 | Cystm1 | ENSMUST00000050584 | 25.729443 | 36 | 21.4854111 | 16.7108753 | 0 | 0 | 0 | 1 | 0 | 0 | 0 | 0 | 1 | 2 | 0 | 0 | 0 | 0 | 2 |
| ENSMUSG00000047044 | D030056L22Rik | ENSMUST00000062753 | 27.1347249 | 35 | 17.3624288 | 20.683112 | 0 | 0 | 0 | 0 | 0 | 0 | 0 | 0 | 0 | 0 | 0 | 0 | 1 | 0 | 1 |
| ENSMUSG00000020255 | D10Wsu102e | ENSMUST00000020488 | 25.5750086 | 28 | 22.0391349 | 24.3391692 | 0 | 0 | 0 | 0 | 0 | 0 | 0 | 0 | 0 | 0 | 0 | 1 | 0 | 0 | 1 |
| ENSMUSG00000037119 | D15Ertd621e | ENSMUST00000037270 | 27.3265946 | 33 | 18.5430464 | 20.6692227 | 0 | 0 | 0 | 0 | 0 | 0 | 0 | 0 | 0 | 0 | 2 | 0 | 0 | 0 | 2 |
| ENSMUSG00000044768 | D1Ertd622e | ENSMUST00000053033 | 28.0452826 | 31 | 19.262888 | 21.8722238 | 6 | 3 | 0 | 6 | 0 | 0 | 0 | 1 | 5 | 12 | 5 | 52 | 22 | 6 | 106 |
| ENSMUSG00000031889 | D230025D16Rik | ENSMUST00000124113 | 22.7470141 | 28 | 25.5157438 | 23.3984799 | 0 | 0 | 0 | 0 | 0 | 0 | 0 | 0 | 0 | 0 | 0 | 1 | 0 | 0 | 1 |
| ENSMUSG00000033883 | D3Ertd254e | ENSMUST00000197653 | 29.8156235 | 30 | 20.3299369 | 20.2086366 | 0 | 0 | 0 | 0 | 0 | 0 | 0 | 0 | 0 | 0 | 3 | 1 | 3 | 1 | 8 |
| ENSMUSG00000025766 | D3Ertd751e | ENSMUST00000143841 | 34.3089431 | 28 | 17.9132791 | 19.7289973 | 1 | 0 | 0 | 1 | 0 | 0 | 0 | 0 | 1 | 2 | 0 | 0 | 0 | 0 | 3 |
| ENSMUSG00000096039 | D830030K20Rik | ENSMUST00000112797 | 22.5419664 | 35 | 23.2613909 | 19.0647482 | 0 | 0 | 0 | 0 | 0 | 0 | 0 | 0 | 0 | 0 | 0 | 1 | 0 | 0 | 1 |
| ENSMUSG00000055639 | Dach1 | ENSMUST00000069334 | 30.7438622 | 36 | 15.5734701 | 17.955295 | 0 | 0 | 0 | 0 | 0 | 0 | 0 | 0 | 0 | 0 | 2 | 1 | 2 | 0 | 5 |
| ENSMUSG00000025592 | Dach2 | ENSMUST00000113382 | 32.585335 | 35 | 16.2768647 | 15.8343869 | 0 | 0 | 0 | 1 | 0 | 0 | 0 | 0 | 1 | 2 | 0 | 0 | 0 | 3 | 5 |
| ENSMUSG00000044548 | Dact1 | ENSMUST00000061273 | 27.0403957 | 35 | 17.6422094 | 20.4451772 | 0 | 0 | 0 | 0 | 0 | 0 | 0 | 0 | 0 | 0 | 0 | 1 | 0 | 0 | 1 |
| ENSMUSG00000000346 | Dazap2 | ENSMUST00000000356 | 25.5060729 | 32 | 22.5910931 | 19.9190283 | 0 | 0 | 0 | 0 | 0 | 0 | 0 | 0 | 0 | 0 | 0 | 0 | 1 | 0 | 1 |
| ENSMUSG00000029878 | Dbpht2 | ENSMUST00000072100 | 28.1469898 | 28 | 21.5793589 | 22.595778 | 0 | 0 | 0 | 0 | 0 | 0 | 0 | 0 | 0 | 0 | 0 | 0 | 0 | 2 | 2 |
| ENSMUSG00000035572 | Dcaf10 | ENSMUST00000155551 | 28.1831448 | 31 | 19.098869 | 21.5432324 | 1 | 1 | 0 | 1 | 0 | 0 | 0 | 0 | 1 | 2 | 0 | 0 | 3 | 0 | 7 |
| ENSMUSG00000028436 | Dcaf12 | ENSMUST00000030145 | 21.9817768 | 33 | 21.7539863 | 23.0068337 | 0 | 0 | 0 | 0 | 0 | 0 | 0 | 0 | 0 | 0 | 1 | 1 | 1 | 0 | 3 |
| ENSMUSG00000050926 | Dcaf12l2 | ENSMUST00000115057 | 32.4584427 | 36 | 12.6859143 | 19.3350831 | 0 | 0 | 0 | 0 | 0 | 0 | 0 | 0 | 0 | 0 | 0 | 2 | 2 | 1 | 5 |
| ENSMUSG00000041966 | Dcaf17 | ENSMUST00000154704 | 29.1617473 | 30 | 19.5985832 | 20.7792208 | 0 | 0 | 0 | 0 | 0 | 0 | 0 | 0 | 0 | 0 | 2 | 4 | 2 | 0 | 8 |
| ENSMUSG00000049354 | Dcaf7 | ENSMUST00000058438 | 21.0785411 | 30 | 23.2042963 | 25.3524278 | 0 | 0 | 0 | 0 | 0 | 0 | 0 | 0 | 0 | 0 | 1 | 2 | 0 | 0 | 3 |
| ENSMUSG00000035107 | Dcbld2 | ENSMUST00000046663 | 26.333164 | 31 | 21.4829863 | 20.9751143 | 0 | 0 | 0 | 1 | 0 | 0 | 0 | 0 | 1 | 2 | 0 | 0 | 4 | 0 | 6 |
| ENSMUSG00000029366 | Dck | ENSMUST00000031311 | 25.5528256 | 36 | 18.5257985 | 19.5085995 | 0 | 0 | 0 | 0 | 0 | 0 | 0 | 0 | 0 | 0 | 1 | 2 | 1 | 0 | 4 |
| ENSMUSG00000027797 | Dclk1 | ENSMUST00000070418 | 27.3680338 | 30 | 21.4456502 | 21.1697627 | 2 | 1 | 0 | 0 | 0 | 0 | 0 | 0 | 0 | 0 | 3 | 11 | 5 | 1 | 23 |
| ENSMUSG00000027845 | Dclre1b | ENSMUST00000029435 | 28.6500212 | 30 | 19.6783749 | 22.0482438 | 0 | 0 | 0 | 0 | 0 | 0 | 0 | 0 | 0 | 0 | 0 | 0 | 5 | 0 | 5 |
| ENSMUSG00000026648 | Dclre1c | ENSMUST00000061852 | 32.6700155 | 31 | 18.74034 | 17.6777434 | 3 | 1 | 0 | 6 | 0 | 0 | 0 | 0 | 6 | 12 | 0 | 1 | 1 | 0 | 18 |
| ENSMUSG00000021962 | Dcp1a | ENSMUST00000022535 | 24.32 | 30 | 22.0533333 | 24.08 | 0 | 0 | 0 | 0 | 0 | 0 | 0 | 0 | 0 | 0 | 0 | 1 | 1 | 0 | 2 |
| ENSMUSG00000041477 | Dcp1b | ENSMUST00000112777 | 26.2369792 | 28 | 20.8984375 | 25 | 0 | 0 | 0 | 0 | 0 | 0 | 0 | 0 | 0 | 0 | 0 | 0 | 0 | 4 | 4 |
| ENSMUSG00000024472 | Dcp2 | ENSMUST00000025350 | 26.5745856 | 32 | 17.9143646 | 23.660221 | 1 | 1 | 0 | 0 | 0 | 0 | 0 | 0 | 0 | 0 | 4 | 0 | 3 | 0 | 9 |
| ENSMUSG00000027708 | Dcun1d1 | ENSMUST00000108182 | 29.1322902 | 35 | 16.4438122 | 19.3172119 | 0 | 0 | 0 | 1 | 0 | 0 | 0 | 0 | 1 | 2 | 2 | 2 | 3 | 2 | 11 |
| ENSMUSG00000048787 | Dcun1d3 | ENSMUST00000059851 | 26.0770975 | 33 | 19.569161 | 21.2244898 | 1 | 0 | 0 | 0 | 0 | 0 | 0 | 0 | 0 | 0 | 0 | 1 | 5 | 1 | 8 |
| ENSMUSG00000051674 | Dcun1d4 | ENSMUST00000087181 | 25.812681 | 32 | 21.4032829 | 21.2745414 | 0 | 0 | 0 | 0 | 0 | 0 | 0 | 0 | 0 | 0 | 2 | 0 | 3 | 2 | 7 |
| ENSMUSG00000031285 | Dcx | ENSMUST00000033642 | 27.7727332 | 33 | 20.1452847 | 19.0556492 | 0 | 0 | 0 | 0 | 0 | 0 | 0 | 0 | 0 | 0 | 0 | 41 | 0 | 0 | 41 |
| ENSMUSG00000061313 | Ddhd2 | ENSMUST00000033975 | 23.0880929 | 30 | 22.5556631 | 24.1045499 | 0 | 0 | 0 | 0 | 0 | 0 | 0 | 0 | 0 | 0 | 0 | 0 | 10 | 0 | 10 |
| ENSMUSG00000046818 | Ddit4l | ENSMUST00000053855 | 25.3063399 | 32 | 23.2818327 | 19.3926478 | 0 | 0 | 0 | 0 | 0 | 0 | 0 | 0 | 0 | 0 | 0 | 26 | 2 | 0 | 28 |

| | | | | | | | | | | | | | | | | | | | | | |
|---|---|---|---|---|---|---|---|---|---|---|---|---|---|---|---|---|---|---|---|---|---|
| ENSMUSG00000026674 | Ddr2 | ENSMUST00000194690 | 28.5147801 | 30 | 19.1059841 | 22.3864456 | 0 | 0 | 0 | 0 | 0 | 0 | 0 | 0 | 0 | 0 | 1 | 33 | 4 | 1 | 39 |
| ENSMUSG00000053289 | Ddx10 | ENSMUST00000065630 | 32.1711569 | 36 | 15.5309033 | 15.8478605 | 0 | 0 | 0 | 1 | 0 | 0 | 0 | 0 | 1 | 2 | 1 | 0 | 3 | 0 | 6 |
| ENSMUSG00000055065 | Ddx17 | ENSMUST00000054014 | 23.8611714 | 32 | 21.2581345 | 23.1670282 | 0 | 0 | 0 | 0 | 0 | 0 | 0 | 0 | 0 | 0 | 0 | 1 | 0 | 0 | 1 |
| ENSMUSG00000001674 | Ddx18 | ENSMUST00000001724 | 26.7080745 | 38 | 24.2236025 | 11.1801242 | 0 | 0 | 0 | 0 | 0 | 0 | 0 | 0 | 0 | 0 | 0 | 0 | 0 | 1 | 1 |
| ENSMUSG00000020075 | Ddx21 | ENSMUST00000045866 | 26.1179829 | 32 | 19.6003806 | 22.26451 | 0 | 0 | 0 | 0 | 0 | 0 | 0 | 0 | 0 | 0 | 1 | 0 | 3 | 0 | 4 |
| ENSMUSG00000035967 | Ddx26b | ENSMUST00000101553 | 27.8291815 | 33 | 21.4234875 | 18.0782918 | 0 | 0 | 0 | 0 | 0 | 0 | 0 | 0 | 0 | 0 | 0 | 1 | 0 | 0 | 1 |
| ENSMUSG00000026806 | Ddx31 | ENSMUST00000113853 | 24.2014742 | 29 | 23.0958231 | 23.8329238 | 0 | 0 | 0 | 0 | 0 | 0 | 0 | 0 | 0 | 0 | 0 | 1 | 1 | 0 | 2 |
| ENSMUSG00000000787 | Ddx3x | ENSMUST00000000804 | 30.546875 | 34 | 16.328125 | 19.4921875 | 2 | 0 | 0 | 1 | 0 | 0 | 0 | 0 | 1 | 2 | 0 | 0 | 0 | 0 | 4 |
| ENSMUSG00000021758 | Ddx4 | ENSMUST00000075748 | 32.0261438 | 37 | 14.379085 | 16.6666667 | 0 | 0 | 0 | 0 | 0 | 0 | 0 | 0 | 0 | 0 | 0 | 0 | 0 | 2 | 2 |
| ENSMUSG00000020705 | Ddx42 | ENSMUST00000021046 | 26.618705 | 32 | 17.4717369 | 23.5354573 | 0 | 0 | 0 | 0 | 0 | 0 | 0 | 0 | 0 | 0 | 0 | 0 | 4 | 1 | 5 |
| ENSMUSG00000020719 | Ddx5 | ENSMUST00000133426 | 28.3956244 | 31 | 18.6417502 | 22.3336372 | 0 | 0 | 0 | 0 | 0 | 0 | 0 | 0 | 0 | 0 | 0 | 0 | 4 | 0 | 4 |
| ENSMUSG00000020076 | Ddx50 | ENSMUST00000020270 | 30.6976744 | 40 | 14.4186047 | 15.3488372 | 0 | 0 | 0 | 0 | 0 | 0 | 0 | 0 | 0 | 0 | 0 | 0 | 0 | 1 | 1 |
| ENSMUSG00000028223 | Decr1 | ENSMUST00000029877 | 26.4066852 | 33 | 19.4428969 | 21.3927577 | 0 | 0 | 0 | 0 | 0 | 0 | 0 | 0 | 0 | 0 | 0 | 3 | 0 | 0 | 3 |
| ENSMUSG00000038633 | Degs1 | ENSMUST00000035295 | 25.6410256 | 32 | 20.1025641 | 22.1538462 | 0 | 0 | 0 | 0 | 0 | 0 | 0 | 0 | 0 | 0 | 0 | 1 | 0 | 0 | 1 |
| ENSMUSG00000056268 | Dennd1b | ENSMUST00000200429 | 28.9774483 | 30 | 20.0494285 | 20.8423437 | 1 | 0 | 1 | 3 | 0 | 0 | 0 | 0 | 3 | 7 | 0 | 0 | 9 | 2 | 19 |
| ENSMUSG00000053641 | Dennd4a | ENSMUST00000038890 | 27.8771384 | 33 | 19.5178849 | 19.7122862 | 0 | 0 | 0 | 1 | 0 | 0 | 0 | 0 | 1 | 2 | 0 | 0 | 0 | 0 | 2 |
| ENSMUSG00000038024 | Dennd4c | ENSMUST00000082026 | 31.2315852 | 34 | 16.9711255 | 17.9728933 | 0 | 0 | 0 | 0 | 0 | 0 | 0 | 0 | 0 | 0 | 0 | 0 | 0 | 1 | 1 |
| ENSMUSG00000030313 | Dennd5b | ENSMUST00000111557 | 30.0621118 | 31 | 19.0771961 | 20.0709849 | 2 | 2 | 0 | 2 | 0 | 0 | 0 | 0 | 2 | 4 | 0 | 8 | 13 | 2 | 31 |
| ENSMUSG00000040818 | Dennd6a | ENSMUST00000037585 | 28.6584086 | 32 | 18.3707747 | 21.0371615 | 1 | 0 | 0 | 0 | 0 | 0 | 0 | 0 | 0 | 0 | 1 | 0 | 12 | 1 | 15 |
| ENSMUSG00000023106 | Denr | ENSMUST00000023869 | 28.5528877 | 32 | 17.6508761 | 21.6093446 | 0 | 0 | 0 | 0 | 0 | 0 | 0 | 0 | 0 | 0 | 0 | 0 | 5 | 1 | 6 |
| ENSMUSG00000028175 | Depdc1a | ENSMUST00000120272 | 31.8627451 | 38 | 15.3186275 | 14.9509804 | 0 | 0 | 1 | 0 | 0 | 0 | 0 | 0 | 0 | 1 | 0 | 0 | 0 | 0 | 1 |
| ENSMUSG00000021697 | Depdc1b | ENSMUST00000051594 | 21.6308041 | 34 | 18.4597961 | 25.8210646 | 0 | 0 | 0 | 0 | 0 | 0 | 0 | 0 | 0 | 0 | 4 | 0 | 4 | 0 | 8 |
| ENSMUSG00000022419 | Deptor | ENSMUST00000096433 | 27.5224215 | 30 | 21.7068386 | 20.3054933 | 0 | 0 | 0 | 1 | 0 | 0 | 0 | 0 | 1 | 2 | 9 | 24 | 10 | 5 | 50 |
| ENSMUSG00000022365 | Derl1 | ENSMUST00000022993 | 22.7272727 | 30 | 21.5798764 | 25.2427184 | 0 | 0 | 0 | 0 | 0 | 0 | 0 | 0 | 0 | 0 | 0 | 1 | 0 | 0 | 1 |
| ENSMUSG00000018442 | Derl2 | ENSMUST00000143850 | 26.0159778 | 33 | 20.4584925 | 20.7016325 | 1 | 0 | 0 | 1 | 0 | 0 | 0 | 0 | 1 | 2 | 0 | 0 | 1 | 0 | 4 |
| ENSMUSG00000026502 | Desi2 | ENSMUST00000027783 | 26.720286 | 31 | 22.9286353 | 19.5582791 | 0 | 0 | 0 | 0 | 0 | 0 | 0 | 0 | 0 | 0 | 6 | 0 | 15 | 1 | 22 |
| ENSMUSG00000029821 | Dfna5 | ENSMUST00000101405 | 32.1769574 | 31 | 18.8415022 | 17.8548695 | 0 | 0 | 0 | 0 | 0 | 0 | 0 | 0 | 0 | 0 | 0 | 0 | 1 | 1 | 2 |
| ENSMUSG00000000276 | Dgke | ENSMUST00000107894 | 26.3281109 | 31 | 20.6374932 | 22.2042139 | 6 | 2 | 0 | 3 | 0 | 0 | 0 | 0 | 3 | 6 | 5 | 0 | 10 | 3 | 32 |
| ENSMUSG00000038665 | Dgki | ENSMUST00000150300 | 25.2953813 | 24 | 25.5639098 | 25.1342642 | 0 | 0 | 0 | 0 | 0 | 0 | 0 | 0 | 0 | 0 | 0 | 2 | 0 | 0 | 2 |
| ENSMUSG00000014554 | Dguok | ENSMUST00000145455 | 23.1343284 | 27 | 25.4409769 | 24.2876526 | 0 | 0 | 0 | 0 | 0 | 0 | 0 | 0 | 0 | 0 | 0 | 2 | 1 | 0 | 3 |
| ENSMUSG00000025815 | Dhtkd1 | ENSMUST00000095147 | 25.6074046 | 34 | 21.2495179 | 18.8584651 | 0 | 0 | 0 | 1 | 0 | 0 | 0 | 0 | 1 | 2 | 3 | 4 | 3 | 0 | 12 |
| ENSMUSG00000027655 | Dhx35 | ENSMUST00000029186 | 19.3577982 | 27 | 27.0642202 | 26.3302752 | 0 | 0 | 0 | 0 | 0 | 0 | 0 | 0 | 0 | 0 | 2 | 0 | 0 | 0 | 2 |
| ENSMUSG00000027770 | Dhx36 | ENSMUST00000029336 | 27.5009808 | 36 | 17.8893684 | 18.4386034 | 1 | 0 | 0 | 0 | 0 | 0 | 0 | 0 | 0 | 0 | 0 | 0 | 1 | 0 | 2 |
| ENSMUSG00000018425 | Dhx40 | ENSMUST00000018569 | 25.9404389 | 35 | 22.4137931 | 16.6144201 | 0 | 0 | 0 | 0 | 0 | 0 | 0 | 0 | 0 | 0 | 0 | 48 | 0 | 0 | 48 |
| ENSMUSG00000042699 | Dhx9 | ENSMUST00000190544 | 29.2110874 | 30 | 18.4968017 | 21.9616205 | 0 | 0 | 0 | 0 | 0 | 0 | 0 | 0 | 0 | 0 | 0 | 0 | 0 | 1 | 1 |
| ENSMUSG00000034480 | Diaph2 | ENSMUST00000113320 | 29.2304552 | 33 | 21.0042866 | 16.8605838 | 0 | 0 | 0 | 0 | 0 | 0 | 0 | 0 | 0 | 0 | 2 | 14 | 2 | 0 | 18 |
| ENSMUSG00000021692 | Dimt1 | ENSMUST00000022203 | 33.3988212 | 35 | 17.3542895 | 14.4073346 | 1 | 0 | 1 | 2 | 0 | 0 | 0 | 0 | 2 | 5 | 0 | 0 | 0 | 0 | 6 |
| ENSMUSG00000023026 | Dip2b | ENSMUST00000100203 | 24.3630967 | 30 | 23.4974029 | 22.0380905 | 2 | 1 | 0 | 1 | 0 | 0 | 0 | 0 | 1 | 2 | 1 | 1 | 4 | 2 | 13 |
| ENSMUSG00000031403 | Dkc1 | ENSMUST00000033776 | 33.0708661 | 30 | 17.519685 | 19.7834646 | 0 | 0 | 0 | 0 | 0 | 0 | 0 | 0 | 0 | 0 | 0 | 0 | 2 | 4 | 6 |
| ENSMUSG00000000168 | Dlat | ENSMUST00000034567 | 31.0825893 | 31 | 17.96875 | 19.921875 | 1 | 1 | 1 | 1 | 0 | 0 | 0 | 0 | 1 | 3 | 0 | 0 | 0 | 0 | 5 |
| ENSMUSG00000022770 | Dlg1 | ENSMUST00000132176 | 29.2712551 | 30 | 19.0283401 | 21.659919 | 0 | 0 | 0 | 1 | 0 | 0 | 0 | 0 | 1 | 2 | 0 | 0 | 1 | 0 | 3 |
| ENSMUSG00000014773 | Dll1 | ENSMUST00000014917 | 28.5502959 | 35 | 15.8284024 | 21.0059172 | 0 | 0 | 0 | 1 | 0 | 0 | 0 | 0 | 1 | 2 | 0 | 0 | 0 | 0 | 2 |
| ENSMUSG00000029755 | Dlx5 | ENSMUST00000142635 | 26.8188302 | 26 | 26.2482168 | 20.6847361 | 0 | 0 | 0 | 0 | 0 | 0 | 0 | 0 | 0 | 0 | 0 | 0 | 5 | 0 | 5 |
| ENSMUSG00000029754 | Dlx6 | ENSMUST00000160937 | 31.0344828 | 27 | 22.0219436 | 19.9059561 | 0 | 0 | 0 | 0 | 0 | 0 | 0 | 0 | 0 | 0 | 0 | 25 | 7 | 1 | 33 |
| ENSMUSG00000022429 | Dmc1 | ENSMUST00000023065 | 31.9842053 | 32 | 16.5844028 | 19.7433366 | 1 | 0 | 0 | 0 | 0 | 0 | 0 | 0 | 0 | 0 | 0 | 1 | 1 | 0 | 3 |
| ENSMUSG00000042508 | Dmtf1 | ENSMUST00000183448 | 29.9793956 | 28 | 20.467033 | 21.6346154 | 0 | 0 | 0 | 0 | 0 | 0 | 0 | 0 | 0 | 0 | 0 | 0 | 2 | 1 | 3 |
| ENSMUSG00000037416 | Dmxl1 | ENSMUST00000041772 | 34.2327698 | 36 | 14.7269181 | 15.2145644 | 1 | 0 | 1 | 2 | 0 | 0 | 0 | 0 | 2 | 5 | 0 | 1 | 1 | 1 | 9 |
| ENSMUSG00000028410 | Dnaja1 | ENSMUST00000164233 | 30.0261718 | 31 | 19.4860814 | 19.8429693 | 0 | 0 | 0 | 1 | 0 | 0 | 0 | 0 | 1 | 2 | 0 | 2 | 0 | 0 | 4 |
| ENSMUSG00000004069 | Dnaja3 | ENSMUST00000060067 | 25.0657318 | 28 | 23.5758107 | 23.225241 | 0 | 0 | 0 | 0 | 0 | 0 | 0 | 0 | 0 | 0 | 0 | 0 | 4 | 0 | 4 |
| ENSMUSG00000074212 | Dnajb14 | ENSMUST00000090178 | 27.2873393 | 33 | 19.7700297 | 20.3264095 | 2 | 0 | 1 | 2 | 0 | 0 | 0 | 0 | 2 | 5 | 6 | 1 | 6 | 0 | 20 |
| ENSMUSG00000036052 | Dnajb5 | ENSMUST00000037872 | 25.6136606 | 23 | 26.9477054 | 24.3863394 | 0 | 0 | 0 | 0 | 0 | 0 | 0 | 0 | 0 | 0 | 0 | 2 | 1 | 0 | 3 |
| ENSMUSG00000029131 | Dnajb6 | ENSMUST00000008733 | 23.0769231 | 30 | 20.1465201 | 26.8864469 | 0 | 0 | 1 | 0 | 0 | 0 | 0 | 0 | 0 | 1 | 0 | 0 | 1 | 1 | 3 |
| ENSMUSG00000014905 | Dnajb9 | ENSMUST00000015049 | 28.3822138 | 38 | 16.7455061 | 16.461684 | 0 | 0 | 0 | 0 | 0 | 0 | 0 | 0 | 0 | 0 | 2 | 0 | 0 | 2 | 4 |
| ENSMUSG00000026740 | Dnajc1 | ENSMUST00000166495 | 27.1379704 | 26 | 24.6009122 | 22.4914481 | 0 | 0 | 0 | 0 | 0 | 0 | 0 | 0 | 0 | 0 | 0 | 2 | 1 | 0 | 3 |
| ENSMUSG00000027006 | Dnajc10 | ENSMUST00000028392 | 25.1491901 | 34 | 20.801364 | 20.5456095 | 1 | 0 | 0 | 0 | 0 | 0 | 0 | 0 | 0 | 0 | 0 | 0 | 2 | 0 | 3 |
| ENSMUSG00000032560 | Dnajc13 | ENSMUST00000035170 | 29.5652174 | 30 | 21.4492754 | 19.1304348 | 0 | 0 | 0 | 0 | 0 | 0 | 0 | 0 | 0 | 0 | 0 | 1 | 0 | 0 | 1 |
| ENSMUSG00000070972 | Dnajc25 | ENSMUST00000095070 | 26.0553633 | 32 | 22.0415225 | 19.550173 | 0 | 0 | 0 | 0 | 0 | 0 | 0 | 0 | 0 | 0 | 2 | 19 | 3 | 0 | 24 |
| ENSMUSG00000020657 | Dnajc27 | ENSMUST00000020986 | 22.8113105 | 29 | 24.4154432 | 24.252311 | 4 | 2 | 4 | 6 | 0 | 1 | 2 | 2 | 2 | 17 | 0 | 1 | 0 | 0 | 24 |
| ENSMUSG00000022136 | Dnajc3 | ENSMUST00000022734 | 28.0272882 | 32 | 18.3627061 | 21.8590108 | 0 | 0 | 0 | 0 | 0 | 0 | 0 | 0 | 0 | 0 | 0 | 0 | 1 | 2 | 3 |
| ENSMUSG00000000826 | Dnajc5 | ENSMUST00000072334 | 23.8144867 | 28 | 23.3454924 | 24.3529616 | 0 | 0 | 0 | 0 | 0 | 0 | 0 | 0 | 0 | 0 | 0 | 1 | 1 | 0 | 2 |
| ENSMUSG00000028528 | Dnajc6 | ENSMUST00000038207 | 28.032345 | 34 | 18.2839173 | 19.901168 | 1 | 0 | 0 | 1 | 0 | 0 | 0 | 0 | 1 | 2 | 4 | 0 | 8 | 1 | 16 |
| ENSMUSG00000042523 | Dnal1 | ENSMUST00000123491 | 26.4103065 | 29 | 21.4913137 | 22.7601015 | 0 | 1 | 1 | 1 | 0 | 0 | 0 | 1 | 0 | 3 | 1 | 40 | 2 | 0 | 47 |
| ENSMUSG00000036766 | Dner | ENSMUST00000049126 | 30.0540541 | 36 | 14.2702703 | 19.2432432 | 0 | 1 | 0 | 1 | 0 | 0 | 0 | 0 | 1 | 2 | 0 | 0 | 0 | 0 | 3 |
| ENSMUSG00000040265 | Dnm3 | ENSMUST00000086074 | 29.0466102 | 31 | 20.7838983 | 19.4491525 | 0 | 0 | 0 | 0 | 0 | 0 | 0 | 0 | 0 | 0 | 0 | 1 | 0 | 0 | 1 |
| ENSMUSG00000020143 | Dock2 | ENSMUST00000143540 | 25.4283756 | 27 | 25.8396162 | 21.5215901 | 0 | 0 | 0 | 0 | 0 | 0 | 0 | 0 | 0 | 0 | 0 | 1 | 0 | 0 | 1 |
| ENSMUSG00000035954 | Dock4 | ENSMUST00000037488 | 30.3130148 | 35 | 15.6507414 | 19.2202087 | 1 | 0 | 0 | 0 | 0 | 0 | 0 | 0 | 0 | 0 | 1 | 0 | 2 | 0 | 4 |
| ENSMUSG00000028556 | Dock7 | ENSMUST00000124466 | 29.6733569 | 27 | 21.2121212 | 22.3534042 | 2 | 1 | 0 | 2 | 0 | 0 | 0 | 0 | 2 | 4 | 0 | 1 | 0 | 0 | 8 |
| ENSMUSG00000034973 | Dopey1 | ENSMUST00000185919 | 29.3696669 | 29 | 21.8276906 | 20.2587393 | 0 | 0 | 0 | 1 | 0 | 0 | 0 | 0 | 1 | 2 | 0 | 0 | 0 | 0 | 2 |
| ENSMUSG00000057147 | Dph6 | ENSMUST00000028640 | 27.1305286 | 34 | 22.3840345 | 16.7745415 | 1 | 0 | 0 | 1 | 0 | 0 | 0 | 0 | 1 | 2 | 0 | 4 | 1 | 1 | 9 |
| ENSMUSG00000036815 | Dpp10 | ENSMUST00000112606 | 29.8560025 | 31 | 19.6274847 | 19.2361872 | 3 | 3 | 0 | 1 | 0 | 0 | 0 | 0 | 1 | 2 | 4 | 1 | 12 | 1 | 26 |
| ENSMUSG00000043067 | Dpy19l1 | ENSMUST00000115277 | 27.8201525 | 33 | 18.5869129 | 20.9152951 | 0 | 0 | 0 | 0 | 0 | 0 | 0 | 0 | 0 | 0 | 0 | 1 | 3 | 3 | 7 |
| ENSMUSG00000045205 | Dpy19l4 | ENSMUST00000084892 | 29.9350213 | 34 | 17.8355366 | 18.5077302 | 0 | 0 | 0 | 0 | 0 | 0 | 0 | 0 | 0 | 0 | 0 | 2 | 1 | 2 | 5 |
| ENSMUSG00000024067 | Dpy30 | ENSMUST00000112571 | 28.4172662 | 35 | 20.5035971 | 16.1870504 | 0 | 0 | 0 | 0 | 0 | 0 | 0 | 0 | 0 | 0 | 0 | 0 | 0 | 1 | 1 |
| ENSMUSG00000022048 | Dpysl2 | ENSMUST00000022629 | 22.5715437 | 34 | 21.4429665 | 21.8057235 | 0 | 0 | 0 | 0 | 0 | 0 | 0 | 0 | 0 | 0 | 1 | 0 | 38 | 6 | 45 |
| ENSMUSG00000024501 | Dpysl3 | ENSMUST00000121805 | 25.022368 | 29 | 22.3978527 | 23.471518 | 0 | 0 | 0 | 0 | 0 | 0 | 0 | 0 | 0 | 0 | 0 | 1 | 6 | 0 | 7 |
| ENSMUSG00000029265 | Dr1 | ENSMUST00000031190 | 30.3062869 | 38 | 12.9500269 | 18.323482 | 0 | 0 | 0 | 0 | 0 | 0 | 0 | 0 | 0 | 0 | 0 | 0 | 2 | 0 | 2 |
| ENSMUSG00000027900 | Dram2 | ENSMUST00000067630 | 36.2811791 | 33 | 13.5147392 | 17.6870748 | 0 | 0 | 0 | 0 | 0 | 0 | 0 | 0 | 0 | 0 | 7 | 0 | 6 | 0 | 13 |
| ENSMUSG00000021478 | Drd1 | ENSMUST00000021932 | 29.1541692 | 32 | 18.9562088 | 19.6160768 | 8 | 4 | 0 | 21 | 4 | 5 | 6 | 7 | 8 | 51 | 0 | 0 | 0 | 0 | 63 |
| ENSMUSG00000015932 | Dstn | ENSMUST00000103172 | 29.9632353 | 30 | 19.3933824 | 20.4963235 | 0 | 0 | 0 | 0 | 0 | 0 | 0 | 0 | 0 | 0 | 0 | 0 | 1 | 0 | 1 |
| ENSMUSG00000037474 | Dtl | ENSMUST00000193977 | 28.6726128 | 33 | 20.2256034 | 18.4679958 | 0 | 0 | 0 | 0 | 0 | 0 | 0 | 0 | 0 | 0 | 0 | 2 | 3 | 0 | 5 |
| ENSMUSG00000024302 | Dtna | ENSMUST00000115832 | 26.8243071 | 29 | 21.9425375 | 21.9425375 | 0 | 0 | 0 | 0 | 0 | 0 | 0 | 0 | 0 | 0 | 0 | 0 | 2 | 0 | 2 |
| ENSMUSG00000024505 | Dtwd2 | ENSMUST00000163590 | 27.826087 | 36 | 16.201373 | 20.4576659 | 0 | 0 | 0 | 0 | 0 | 0 | 0 | 0 | 0 | 0 | 2 | 0 | 17 | 4 | 23 |
| ENSMUSG00000026281 | Dtymk | ENSMUST00000112890 | 23.7380628 | 33 | 20.7366985 | 22.6466576 | 0 | 0 | 0 | 0 | 0 | 0 | 0 | 0 | 0 | 0 | 1 | 0 | 7 | 0 | 8 |
| ENSMUSG00000030002 | Dusp11 | ENSMUST00000032071 | 24.7892074 | 32 | 22.8780214 | 20.0487165 | 4 | 2 | 0 | 10 | 0 | 1 | 2 | 3 | 5 | 21 | 0 | 2 | 1 | 5 | 35 |
| ENSMUSG00000026659 | Dusp12 | ENSMUST00000170420 | 27.0192995 | 32 | 18.9421015 | 21.729807 | 0 | 0 | 0 | 0 | 0 | 0 | 0 | 0 | 0 | 0 | 0 | 1 | 10 | 3 | 14 |
| ENSMUSG00000018648 | Dusp14 | ENSMUST00000018792 | 22.9166667 | 32 | 22.3958333 | 22.5694444 | 0 | 0 | 0 | 0 | 0 | 0 | 0 | 0 | 0 | 0 | 0 | 3 | 0 | 0 | 3 |
| ENSMUSG00000027001 | Dusp19 | ENSMUST00000028384 | 29.8412698 | 37 | 18.7301587 | 14.2857143 | 0 | 0 | 0 | 0 | 0 | 0 | 0 | 0 | 0 | 0 | 0 | 1 | 0 | 0 | 1 |
| ENSMUSG00000019960 | Dusp6 | ENSMUST00000020118 | 26.8292683 | 35 | 17.2413793 | 21.1942809 | 0 | 0 | 0 | 0 | 0 | 0 | 0 | 0 | 0 | 0 | 1 | 0 | 5 | 2 | 8 |
| ENSMUSG00000027203 | Dut | ENSMUST00000051605 | 26.9456067 | 29 | 21.7573222 | 22.0083682 | 0 | 0 | 0 | 1 | 0 | 0 | 0 | 0 | 1 | 2 | 0 | 0 | 0 | 0 | 2 |
| ENSMUSG00000035770 | Dync1li2 | ENSMUST00000041769 | 25.4814305 | 32 | 17.9848693 | 25 | 0 | 0 | 0 | 0 | 0 | 0 | 0 | 0 | 0 | 0 | 0 | 0 | 4 | 1 | 5 |
| ENSMUSG00000009013 | Dynll1 | ENSMUST00000009157 | 22.887768 | 30 | 22.3833544 | 24.4640605 | 1 | 0 | 1 | 0 | 0 | 0 | 0 | 0 | 0 | 1 | 1 | 0 | 1 | 0 | 4 |
| ENSMUSG00000031176 | Dynlt3 | ENSMUST00000033519 | 32.7325581 | 32 | 16.5116279 | 19.244186 | 0 | 1 | 0 | 1 | 0 | 0 | 0 | 0 | 1 | 2 | 1 | 0 | 2 | 0 | 6 |

| | | | | | | | | | | | | | | | | | | | | | |
|---|---|---|---|---|---|---|---|---|---|---|---|---|---|---|---|---|---|---|---|---|---|
| ENSMUSG00000037259 | Dzank1 | ENSMUST00000081982 | 27.2956263 | 31 | 19.349668 | 22.5097321 | 1 | 0 | 0 | 0 | 0 | 0 | 0 | 0 | 0 | 0 | 2 | 2 | 1 | 1 | 7 |
| ENSMUSG00000064061 | Dzip3 | ENSMUST00000121869 | 31.25 | 36 | 14.6022727 | 17.7840909 | 0 | 0 | 0 | 0 | 0 | 0 | 0 | 0 | 0 | 0 | 0 | 0 | 1 | 0 | 1 |
| ENSMUSG00000045071 | E130308A19Rik | ENSMUST00000070150 | 27.5833333 | 28 | 22 | 22.0714286 | 0 | 0 | 0 | 0 | 0 | 0 | 0 | 0 | 0 | 0 | 1 | 12 | 9 | 0 | 22 |
| ENSMUSG00000016477 | E2f3 | ENSMUST00000102948 | 24.3088942 | 30 | 21.9951923 | 24.1286058 | 0 | 0 | 0 | 0 | 0 | 0 | 0 | 0 | 0 | 0 | 0 | 3 | 1 | 0 | 4 |
| ENSMUSG00000020185 | E2f7 | ENSMUST00000073781 | 24.7054203 | 31 | 21.9952867 | 21.8774548 | 1 | 0 | 1 | 1 | 0 | 0 | 0 | 0 | 1 | 3 | 0 | 1 | 1 | 0 | 6 |
| ENSMUSG00000021890 | Eaf1 | ENSMUST00000022446 | 25.8870719 | 30 | 21.5365628 | 22.369639 | 1 | 0 | 0 | 2 | 0 | 0 | 0 | 0 | 2 | 4 | 0 | 9 | 0 | 0 | 14 |
| ENSMUSG00000022838 | Eaf2 | ENSMUST00000075946 | 28.7148594 | 32 | 21.184739 | 18.1726908 | 0 | 0 | 0 | 0 | 0 | 0 | 0 | 0 | 0 | 0 | 0 | 1 | 0 | 0 | 1 |
| ENSMUSG00000057098 | Ebf1 | ENSMUST00000081265 | 31.1396949 | 31 | 18.6060425 | 19.1743943 | 1 | 0 | 1 | 1 | 0 | 0 | 0 | 0 | 1 | 3 | 1 | 2 | 1 | 3 | 11 |
| ENSMUSG00000022053 | Ebf2 | ENSMUST00000176161 | 28.7195994 | 32 | 20.4220315 | 18.7052933 | 0 | 0 | 0 | 1 | 0 | 0 | 0 | 0 | 1 | 2 | 3 | 6 | 4 | 2 | 17 |
| ENSMUSG00000010476 | Ebf3 | ENSMUST00000033378 | 28.0948851 | 32 | 20.4225352 | 19.273536 | 0 | 0 | 0 | 0 | 0 | 0 | 0 | 0 | 0 | 0 | 0 | 1 | 0 | 0 | 1 |
| ENSMUSG00000019883 | Echdc1 | ENSMUST00000020034 | 33.4205021 | 31 | 19.0899582 | 16.5794979 | 0 | 0 | 0 | 0 | 0 | 0 | 0 | 0 | 0 | 0 | 0 | 0 | 1 | 0 | 1 |
| ENSMUSG00000043631 | Ecm2 | ENSMUST00000051504 | 29.6759941 | 33 | 19.9558174 | 17.5257732 | 0 | 0 | 0 | 0 | 0 | 0 | 0 | 0 | 0 | 0 | 2 | 1 | 2 | 0 | 5 |
| ENSMUSG00000027699 | Ect2 | ENSMUST00000184113 | 30.6122449 | 30 | 19.2506853 | 20.2254036 | 0 | 0 | 0 | 0 | 0 | 0 | 0 | 0 | 0 | 0 | 0 | 1 | 0 | 1 | 2 |
| ENSMUSG00000034457 | Eda2r | ENSMUST00000037353 | 28.150798 | 33 | 18.5745735 | 19.9779857 | 0 | 0 | 0 | 0 | 0 | 0 | 0 | 0 | 0 | 0 | 0 | 2 | 1 | 0 | 3 |
| ENSMUSG00000030104 | Edem1 | ENSMUST00000204804 | 24.8967416 | 33 | 20.192749 | 21.9596145 | 0 | 0 | 0 | 0 | 0 | 0 | 0 | 0 | 0 | 0 | 1 | 10 | 9 | 0 | 20 |
| ENSMUSG00000043019 | Edem3 | ENSMUST00000059498 | 28.7597364 | 33 | 17.7950869 | 20.3714799 | 1 | 0 | 0 | 1 | 0 | 0 | 0 | 0 | 1 | 2 | 0 | 1 | 2 | 0 | 6 |
| ENSMUSG00000034488 | Edil3 | ENSMUST00000043111 | 32.2068763 | 37 | 15.5157214 | 15.4275639 | 1 | 0 | 0 | 1 | 0 | 0 | 0 | 0 | 1 | 2 | 0 | 0 | 1 | 0 | 4 |
| ENSMUSG00000031616 | Ednra | ENSMUST00000034029 | 26.8894602 | 31 | 22.159383 | 20.3084833 | 0 | 0 | 0 | 0 | 0 | 0 | 0 | 0 | 0 | 0 | 0 | 1 | 0 | 0 | 1 |
| ENSMUSG00000022122 | Ednrb | ENSMUST00000022718 | 31.1757476 | 30 | 20.4834084 | 18.4760344 | 0 | 0 | 0 | 0 | 0 | 0 | 0 | 0 | 0 | 0 | 0 | 0 | 7 | 2 | 9 |
| ENSMUSG00000036499 | Eea1 | ENSMUST00000053484 | 29.8430899 | 34 | 17.7429089 | 18.3464092 | 0 | 0 | 0 | 1 | 0 | 0 | 0 | 0 | 1 | 2 | 0 | 2 | 0 | 0 | 4 |
| ENSMUSG00000030619 | Eed | ENSMUST00000107234 | 28.113879 | 41 | 13.1672598 | 17.7935943 | 1 | 0 | 0 | 1 | 0 | 0 | 0 | 0 | 1 | 2 | 0 | 0 | 0 | 0 | 3 |
| ENSMUSG00000001707 | Eef1e1 | ENSMUST00000001757 | 31.1440678 | 39 | 12.2881356 | 17.5847458 | 1 | 0 | 0 | 1 | 0 | 0 | 0 | 0 | 1 | 2 | 1 | 0 | 0 | 0 | 4 |
| ENSMUSG00000034210 | Efcab14 | ENSMUST00000074425 | 24.6233522 | 35 | 21.4218456 | 18.9736347 | 0 | 0 | 0 | 0 | 0 | 0 | 0 | 0 | 0 | 0 | 0 | 3 | 7 | 0 | 10 |
| ENSMUSG00000026495 | Efcab2 | ENSMUST00000027775 | 32.2204214 | 27 | 16.2074554 | 25.0567261 | 1 | 1 | 0 | 1 | 0 | 0 | 0 | 0 | 1 | 2 | 0 | 0 | 0 | 0 | 4 |
| ENSMUSG00000003070 | Efna2 | ENSMUST00000003154 | 24.5081967 | 25 | 26.7213115 | 23.3606557 | 0 | 0 | 1 | 1 | 0 | 0 | 0 | 0 | 1 | 3 | 0 | 0 | 0 | 0 | 3 |
| ENSMUSG00000028039 | Efna3 | ENSMUST00000029673 | 22.8218466 | 25 | 27.0481144 | 25.1625488 | 0 | 0 | 0 | 0 | 0 | 0 | 0 | 0 | 0 | 0 | 0 | 1 | 0 | 0 | 1 |
| ENSMUSG00000001300 | Efnb2 | ENSMUST00000001319 | 26.6962587 | 28 | 21.4965124 | 24.2866202 | 2 | 1 | 0 | 2 | 0 | 0 | 0 | 1 | 1 | 4 | 0 | 2 | 4 | 1 | 14 |
| ENSMUSG00000063600 | Egfem1 | ENSMUST00000119598 | 29.4855709 | 33 | 19.9498118 | 17.1894605 | 0 | 1 | 2 | 0 | 0 | 0 | 0 | 0 | 0 | 2 | 2 | 0 | 0 | 0 | 5 |
| ENSMUSG00000020122 | Egfr | ENSMUST00000020329 | 30.0015865 | 30 | 21.0851975 | 18.6895129 | 1 | 0 | 0 | 0 | 0 | 0 | 0 | 0 | 0 | 0 | 1 | 0 | 6 | 0 | 8 |
| ENSMUSG00000062762 | Ei24 | ENSMUST00000115086 | 22.0464135 | 32 | 21.5189873 | 24.4725738 | 0 | 0 | 0 | 0 | 0 | 0 | 0 | 0 | 0 | 0 | 0 | 1 | 0 | 0 | 1 |
| ENSMUSG00000091337 | Eid1 | ENSMUST00000164756 | 31.956912 | 32 | 16.7863555 | 19.4793537 | 0 | 0 | 0 | 0 | 0 | 0 | 0 | 0 | 0 | 0 | 0 | 9 | 0 | 0 | 9 |
| ENSMUSG00000070705 | Eid2b | ENSMUST00000094651 | 25.4622293 | 30 | 18.8061278 | 25.4094031 | 0 | 0 | 0 | 0 | 0 | 0 | 0 | 0 | 0 | 0 | 3 | 1 | 5 | 1 | 10 |
| ENSMUSG00000035530 | Eif1 | ENSMUST00000049385 | 25.7142857 | 27 | 23.1292517 | 24.2176871 | 0 | 0 | 0 | 0 | 0 | 0 | 0 | 0 | 0 | 0 | 0 | 1 | 0 | 0 | 1 |
| ENSMUSG00000057561 | Eif1a | ENSMUST00000078079 | 29.9378288 | 33 | 18.1252989 | 19.0817791 | 0 | 0 | 0 | 0 | 0 | 0 | 0 | 0 | 0 | 0 | 0 | 0 | 5 | 2 | 7 |
| ENSMUSG00000067194 | Eif1ax | ENSMUST00000087143 | 28.4163898 | 33 | 17.7408638 | 20.3543743 | 0 | 0 | 0 | 0 | 0 | 0 | 0 | 0 | 0 | 0 | 1 | 0 | 3 | 0 | 4 |
| ENSMUSG00000027810 | Eif2a | ENSMUST00000154219 | 32.7468785 | 27 | 19.1827469 | 20.6583428 | 0 | 0 | 0 | 0 | 0 | 0 | 0 | 0 | 0 | 0 | 0 | 0 | 1 | 1 | 2 |
| ENSMUSG00000021116 | Eif2s1 | ENSMUST00000071230 | 30.3724928 | 32 | 18.8156638 | 19.1499522 | 0 | 0 | 1 | 0 | 0 | 0 | 0 | 0 | 0 | 1 | 0 | 0 | 0 | 0 | 1 |
| ENSMUSG00000074656 | Eif2s2 | ENSMUST00000099173 | 26.9201521 | 32 | 19.6197719 | 20.9885932 | 0 | 0 | 0 | 0 | 0 | 0 | 0 | 0 | 0 | 0 | 0 | 0 | 1 | 0 | 1 |
| ENSMUSG00000035150 | Eif2s3x | ENSMUST00000050328 | 26.2081784 | 31 | 21.1431227 | 21.8401487 | 0 | 0 | 1 | 0 | 0 | 0 | 0 | 0 | 0 | 1 | 5 | 1 | 7 | 0 | 14 |
| ENSMUSG00000027236 | Eif3j1 | ENSMUST00000028668 | 29.0666667 | 29 | 20.2222222 | 21.2888889 | 0 | 0 | 0 | 0 | 0 | 0 | 0 | 0 | 0 | 0 | 0 | 0 | 0 | 2 | 2 |
| ENSMUSG00000022884 | Eif4a2 | ENSMUST00000147117 | 29.9752781 | 31 | 16.7490729 | 22.6205192 | 0 | 1 | 0 | 0 | 0 | 0 | 0 | 0 | 0 | 0 | 0 | 0 | 0 | 0 | 1 |
| ENSMUSG00000028156 | Eif4e | ENSMUST00000029803 | 27.9818702 | 32 | 18.6545802 | 20.8730916 | 0 | 0 | 0 | 0 | 0 | 0 | 0 | 0 | 0 | 0 | 1 | 0 | 4 | 0 | 5 |
| ENSMUSG00000050192 | Eif5a2 | ENSMUST00000060500 | 29.0138921 | 34 | 18.0368936 | 18.5606923 | 0 | 0 | 0 | 1 | 0 | 0 | 0 | 0 | 1 | 2 | 4 | 0 | 7 | 2 | 15 |
| ENSMUSG00000040028 | Elavl1 | ENSMUST00000098950 | 25.7368203 | 33 | 20.6517227 | 20.1951017 | 8 | 3 | 0 | 14 | 0 | 1 | 3 | 5 | 6 | 29 | 4 | 2 | 7 | 0 | 53 |
| ENSMUSG00000008489 | Elavl2 | ENSMUST00000008633 | 30.2785628 | 36 | 14.4529673 | 18.974566 | 0 | 0 | 0 | 1 | 0 | 0 | 0 | 0 | 1 | 2 | 0 | 0 | 1 | 0 | 3 |
| ENSMUSG00000028546 | Elavl4 | ENSMUST00000106598 | 33.8314176 | 34 | 14.137931 | 17.8544061 | 1 | 1 | 0 | 1 | 0 | 0 | 0 | 1 | 0 | 2 | 0 | 0 | 0 | 0 | 4 |
| ENSMUSG00000036461 | Elf1 | ENSMUST00000040131 | 32.2977726 | 33 | 15.7092614 | 18.8745604 | 0 | 0 | 0 | 1 | 0 | 0 | 0 | 0 | 1 | 2 | 0 | 0 | 0 | 0 | 2 |
| ENSMUSG00000008398 | Elk3 | ENSMUST00000008542 | 28.3513097 | 29 | 19.9922958 | 23.0739599 | 0 | 0 | 0 | 0 | 0 | 0 | 0 | 0 | 0 | 0 | 0 | 2 | 0 | 0 | 2 |
| ENSMUSG00000041112 | Elmo1 | ENSMUST00000072519 | 22.5386493 | 31 | 28.4784378 | 17.7379984 | 0 | 0 | 0 | 0 | 0 | 0 | 0 | 0 | 0 | 0 | 0 | 16 | 2 | 0 | 18 |
| ENSMUSG00000035151 | Elmod2 | ENSMUST00000177594 | 27.9315961 | 34 | 17.6438654 | 20.5211726 | 0 | 0 | 0 | 0 | 0 | 0 | 0 | 0 | 0 | 0 | 5 | 0 | 5 | 0 | 10 |
| ENSMUSG00000021364 | Elovl2 | ENSMUST00000021793 | 27.6527331 | 30 | 22.2936763 | 20.2572347 | 0 | 0 | 0 | 0 | 0 | 0 | 0 | 0 | 0 | 0 | 0 | 0 | 0 | 1 | 1 |
| ENSMUSG00000032262 | Elovl4 | ENSMUST00000034796 | 29.7077922 | 33 | 18.0194805 | 18.8311688 | 0 | 0 | 1 | 0 | 0 | 0 | 0 | 0 | 0 | 1 | 0 | 0 | 4 | 2 | 7 |
| ENSMUSG00000032349 | Elovl5 | ENSMUST00000034904 | 27.6446523 | 33 | 18.8778492 | 20.8649912 | 0 | 0 | 0 | 0 | 0 | 0 | 0 | 0 | 0 | 0 | 0 | 0 | 0 | 1 | 1 |
| ENSMUSG00000041220 | Elovl6 | ENSMUST00000071402 | 29.97003 | 27 | 21.0989011 | 21.7182817 | 2 | 1 | 0 | 2 | 0 | 0 | 0 | 1 | 1 | 4 | 0 | 2 | 5 | 1 | 15 |
| ENSMUSG00000021696 | Elovl7 | ENSMUST00000022207 | 29.0461121 | 31 | 19.0325497 | 20.9990958 | 0 | 0 | 0 | 0 | 0 | 0 | 0 | 0 | 0 | 0 | 3 | 0 | 5 | 0 | 8 |
| ENSMUSG00000027167 | Elp4 | ENSMUST00000122965 | 31.7087493 | 33 | 17.2049325 | 18.144451 | 0 | 0 | 0 | 0 | 0 | 0 | 0 | 0 | 0 | 0 | 0 | 0 | 4 | 4 | 8 |
| ENSMUSG00000055943 | Emc7 | ENSMUST00000069747 | 30.0656194 | 28 | 19.5466295 | 22.1714058 | 0 | 0 | 0 | 0 | 0 | 0 | 0 | 0 | 0 | 0 | 0 | 2 | 1 | 0 | 3 |
| ENSMUSG00000054690 | Emcn | ENSMUST00000122064 | 30.509515 | 31 | 19.7053407 | 19.2142419 | 1 | 0 | 0 | 0 | 0 | 0 | 0 | 0 | 0 | 0 | 0 | 0 | 0 | 0 | 1 |
| ENSMUSG00000051166 | Eml5 | ENSMUST00000065716 | 32.1943406 | 31 | 19.9412707 | 16.9514148 | 1 | 0 | 0 | 2 | 0 | 0 | 0 | 0 | 2 | 4 | 0 | 0 | 0 | 0 | 5 |
| ENSMUSG00000035401 | Emsy | ENSMUST00000205276 | 28.6799887 | 31 | 18.8291585 | 21.840698 | 1 | 1 | 1 | 0 | 0 | 0 | 0 | 0 | 0 | 1 | 1 | 1 | 1 | 0 | 6 |
| ENSMUSG00000043969 | Emx2 | ENSMUST00000174353 | 31.7518248 | 26 | 18.540146 | 23.9416058 | 0 | 0 | 0 | 0 | 0 | 0 | 0 | 0 | 0 | 0 | 0 | 0 | 1 | 0 | 1 |
| ENSMUSG00000022995 | Enah | ENSMUST00000193703 | 25.9814268 | 32 | 19.723512 | 22.2351203 | 0 | 1 | 1 | 0 | 0 | 0 | 0 | 0 | 0 | 1 | 6 | 2 | 19 | 1 | 30 |
| ENSMUSG00000029326 | Enoph1 | ENSMUST00000169390 | 27.7575758 | 29 | 21.5757576 | 21.8181818 | 0 | 0 | 0 | 0 | 0 | 0 | 0 | 0 | 0 | 0 | 0 | 0 | 0 | 3 | 3 |
| ENSMUSG00000023961 | Enpp4 | ENSMUST00000024757 | 28.1530782 | 32 | 18.1364393 | 21.4975042 | 0 | 0 | 0 | 0 | 0 | 0 | 0 | 0 | 0 | 0 | 1 | 10 | 0 | 0 | 11 |
| ENSMUSG00000023960 | Enpp5 | ENSMUST00000126032 | 26.7922794 | 29 | 22.6332721 | 21.6911765 | 0 | 0 | 0 | 0 | 0 | 0 | 0 | 0 | 0 | 0 | 0 | 2 | 0 | 0 | 2 |
| ENSMUSG00000038619 | Ensa | ENSMUST00000037983 | 25.0663521 | 29 | 20.5544087 | 25.597169 | 0 | 0 | 0 | 0 | 0 | 0 | 0 | 0 | 0 | 0 | 3 | 2 | 7 | 0 | 12 |
| ENSMUSG00000035245 | Eogt | ENSMUST00000054344 | 28.2416503 | 31 | 19.9901768 | 20.7760314 | 0 | 0 | 0 | 0 | 0 | 0 | 0 | 0 | 0 | 0 | 1 | 0 | 1 | 0 | 2 |
| ENSMUSG00000019978 | Epb41l2 | ENSMUST00000053748 | 27.369281 | 33 | 21.3235294 | 18.1372549 | 0 | 0 | 0 | 0 | 0 | 0 | 0 | 0 | 0 | 0 | 0 | 1 | 0 | 0 | 1 |
| ENSMUSG00000026383 | Epb41l5 | ENSMUST00000191046 | 30.0369866 | 32 | 17.6562196 | 20.2258127 | 0 | 0 | 0 | 0 | 0 | 0 | 0 | 0 | 0 | 0 | 0 | 0 | 2 | 2 | 4 |
| ENSMUSG00000069495 | Epc2 | ENSMUST00000092123 | 30.8717949 | 34 | 15.5384615 | 19.8461538 | 1 | 0 | 1 | 1 | 0 | 0 | 0 | 0 | 1 | 3 | 1 | 0 | 0 | 0 | 5 |
| ENSMUSG00000002808 | Epdr1 | ENSMUST00000002885 | 27.1656272 | 31 | 20.6514207 | 21.0672211 | 0 | 0 | 0 | 0 | 0 | 0 | 0 | 0 | 0 | 0 | 1 | 0 | 0 | 0 | 1 |
| ENSMUSG00000039840 | Epg5 | ENSMUST00000044622 | 25.4166667 | 28 | 22.5520833 | 23.6979167 | 0 | 0 | 0 | 1 | 0 | 0 | 0 | 0 | 1 | 2 | 0 | 0 | 0 | 0 | 2 |
| ENSMUSG00000052504 | Epha3 | ENSMUST00000064405 | 32.3949094 | 34 | 16.2360201 | 17.7400694 | 1 | 0 | 0 | 0 | 0 | 0 | 0 | 0 | 0 | 0 | 0 | 0 | 1 | 0 | 2 |
| ENSMUSG00000029245 | Epha5 | ENSMUST00000053733 | 32.3288951 | 34 | 15.6271899 | 18.2667601 | 1 | 0 | 1 | 0 | 0 | 0 | 0 | 0 | 0 | 1 | 0 | 0 | 0 | 0 | 2 |
| ENSMUSG00000028289 | Epha7 | ENSMUST00000029964 | 31.7505721 | 32 | 18.2208238 | 17.9633867 | 1 | 0 | 0 | 1 | 0 | 0 | 0 | 0 | 1 | 2 | 0 | 0 | 0 | 0 | 3 |
| ENSMUSG00000046785 | Epm2aip1 | ENSMUST00000060711 | 27.6731928 | 34 | 19.6347892 | 18.4676205 | 0 | 0 | 0 | 0 | 0 | 0 | 0 | 0 | 0 | 0 | 0 | 2 | 7 | 0 | 9 |
| ENSMUSG00000028552 | Eps15 | ENSMUST00000102729 | 30.1020408 | 35 | 16.9642857 | 18.4098639 | 0 | 0 | 0 | 0 | 0 | 0 | 0 | 0 | 0 | 0 | 0 | 0 | 7 | 0 | 7 |
| ENSMUSG00000075703 | Ept1 | ENSMUST00000145167 | 25.7921887 | 32 | 19.1414886 | 22.9734709 | 1 | 0 | 0 | 1 | 0 | 0 | 0 | 0 | 1 | 2 | 2 | 2 | 5 | 0 | 12 |
| ENSMUSG00000019936 | Epyc | ENSMUST00000105285 | 37.5844595 | 32 | 14.9493243 | 15.4560811 | 0 | 0 | 0 | 1 | 0 | 0 | 0 | 0 | 1 | 2 | 0 | 0 | 0 | 0 | 2 |
| ENSMUSG00000062209 | Erbb4 | ENSMUST00000119142 | 31.2412476 | 33 | 17.6830045 | 18.255888 | 2 | 2 | 0 | 4 | 0 | 0 | 0 | 0 | 4 | 8 | 2 | 5 | 1 | 1 | 21 |
| ENSMUSG00000054051 | Ercc6 | ENSMUST00000066807 | 25.3923334 | 34 | 20.8386931 | 19.6809879 | 1 | 0 | 0 | 0 | 0 | 0 | 0 | 0 | 0 | 0 | 4 | 0 | 6 | 0 | 11 |
| ENSMUSG00000021694 | Ercc8 | ENSMUST00000123657 | 25.7182576 | 27 | 23.5866543 | 23.5866543 | 0 | 0 | 0 | 0 | 0 | 0 | 0 | 0 | 0 | 0 | 1 | 0 | 0 | 0 | 1 |
| ENSMUSG00000030304 | Ergic2 | ENSMUST00000136008 | 31.0238645 | 31 | 17.0515781 | 21.2856043 | 0 | 0 | 0 | 0 | 0 | 0 | 0 | 0 | 0 | 0 | 0 | 0 | 1 | 1 | 2 |
| ENSMUSG00000030929 | Eri2 | ENSMUST00000033224 | 29.7776442 | 30 | 20.5528846 | 19.5612981 | 0 | 0 | 0 | 1 | 0 | 0 | 0 | 0 | 1 | 2 | 0 | 0 | 1 | 0 | 3 |
| ENSMUSG00000051978 | Erich1 | ENSMUST00000110813 | 33.2038835 | 30 | 17.2815534 | 20 | 0 | 0 | 1 | 0 | 0 | 0 | 0 | 0 | 0 | 1 | 0 | 0 | 0 | 0 | 1 |
| ENSMUSG00000020311 | Erlec1 | ENSMUST00000073192 | 32.7018943 | 33 | 16.550349 | 17.6470588 | 1 | 0 | 0 | 0 | 0 | 0 | 0 | 0 | 0 | 0 | 0 | 0 | 3 | 2 | 6 |
| ENSMUSG00000025198 | Erlin1 | ENSMUST00000071698 | 26.8280383 | 30 | 21.2808875 | 21.9868886 | 0 | 0 | 0 | 0 | 0 | 0 | 0 | 0 | 0 | 0 | 0 | 1 | 0 | 2 | 3 |
| ENSMUSG00000020715 | Ern1 | ENSMUST00000001059 | 23.4069887 | 30 | 22.4563207 | 24.5375128 | 2 | 1 | 0 | 3 | 0 | 0 | 0 | 1 | 2 | 6 | 0 | 2 | 0 | 0 | 11 |
| ENSMUSG00000021831 | Ero1l | ENSMUST00000022378 | 27.6226158 | 34 | 19.7547684 | 18.3242507 | 2 | 0 | 0 | 0 | 0 | 0 | 0 | 0 | 0 | 0 | 0 | 1 | 1 | 1 | 5 |
| ENSMUSG00000057069 | Ero1lb | ENSMUST00000071973 | 25.6536567 | 34 | 19.7423266 | 20.6896552 | 0 | 1 | 0 | 0 | 0 | 0 | 0 | 0 | 0 | 0 | 1 | 0 | 1 | 2 | 5 |
| ENSMUSG00000024293 | Esco1 | ENSMUST00000097670 | 31.0810811 | 35 | 14.527027 | 19.0315315 | 0 | 0 | 0 | 0 | 0 | 0 | 0 | 0 | 0 | 0 | 0 | 0 | 2 | 1 | 3 |
| ENSMUSG00000022034 | Esco2 | ENSMUST00000022613 | 34.0517241 | 36 | 14.2241379 | 16.0560345 | 0 | 0 | 0 | 0 | 0 | 0 | 0 | 0 | 0 | 0 | 0 | 0 | 4 | 0 | 4 |

| | | | | | | | | | | | | | | | | | | | | | |
|---|---|---|---|---|---|---|---|---|---|---|---|---|---|---|---|---|---|---|---|---|---|
| ENSMUSG00000045624 | Esf1 | ENSMUST00000046030 | 32.6474623 | 33 | 15.9122085 | 18.2441701 | 1 | 0 | 1 | 1 | 0 | 0 | 0 | 0 | 1 | 3 | 0 | 0 | 0 | 0 | 4 |
| ENSMUSG00000042379 | Esm1 | ENSMUST00000038144 | 30.2650957 | 33 | 14.359352 | 22.3122239 | 0 | 0 | 0 | 1 | 0 | 0 | 0 | 0 | 1 | 2 | 0 | 0 | 0 | 0 | 2 |
| ENSMUSG00000026610 | Esrrg | ENSMUST00000027906 | 28.6438868 | 32 | 20.5018687 | 18.5264282 | 1 | 0 | 0 | 0 | 0 | 0 | 0 | 0 | 0 | 0 | 4 | 4 | 12 | 0 | 21 |
| ENSMUSG00000021171 | Esyt2 | ENSMUST00000100986 | 29.6527778 | 33 | 17.5 | 19.8263889 | 0 | 0 | 0 | 0 | 0 | 0 | 0 | 0 | 0 | 0 | 3 | 0 | 2 | 0 | 5 |
| ENSMUSG00000016984 | Etaa1 | ENSMUST00000076661 | 30.6122449 | 36 | 16.9387755 | 16.5986395 | 0 | 0 | 0 | 1 | 0 | 0 | 0 | 0 | 1 | 2 | 0 | 2 | 0 | 0 | 4 |
| ENSMUSG00000024360 | Etf1 | ENSMUST00000025218 | 26.4185202 | 30 | 20.154335 | 23.9219246 | 0 | 0 | 0 | 0 | 0 | 0 | 0 | 0 | 0 | 0 | 1 | 1 | 0 | 0 | 2 |
| ENSMUSG00000030275 | Etnk1 | ENSMUST00000032413 | 26.3261297 | 33 | 18.5265226 | 21.9253438 | 1 | 1 | 0 | 1 | 0 | 0 | 0 | 0 | 1 | 2 | 1 | 3 | 0 | 0 | 8 |
| ENSMUSG00000032035 | Ets1 | ENSMUST00000034534 | 26.6273932 | 30 | 19.4992636 | 23.65243 | 0 | 0 | 0 | 1 | 0 | 0 | 0 | 0 | 1 | 2 | 0 | 1 | 11 | 6 | 20 |
| ENSMUSG00000022895 | Ets2 | ENSMUST00000023612 | 24.2028228 | 30 | 22.9482488 | 22.7914271 | 0 | 0 | 0 | 0 | 0 | 0 | 0 | 0 | 0 | 0 | 1 | 1 | 2 | 1 | 5 |
| ENSMUSG00000004151 | Etv1 | ENSMUST00000095767 | 29.8855274 | 33 | 19.255928 | 18.3156173 | 0 | 0 | 0 | 0 | 0 | 0 | 0 | 0 | 0 | 0 | 0 | 1 | 2 | 1 | 4 |
| ENSMUSG00000003382 | Etv3 | ENSMUST00000119109 | 25.3995434 | 29 | 20.7477169 | 25.0856164 | 0 | 0 | 0 | 1 | 0 | 0 | 0 | 0 | 1 | 2 | 0 | 6 | 2 | 0 | 10 |
| ENSMUSG00000011831 | Evi5 | ENSMUST00000112642 | 29.0361086 | 33 | 18.5019397 | 19.7254551 | 0 | 0 | 0 | 0 | 0 | 0 | 0 | 0 | 0 | 0 | 0 | 1 | 4 | 0 | 5 |
| ENSMUSG00000001815 | Evx2 | ENSMUST00000001867 | 26.7047619 | 30 | 20.9904762 | 21.8666667 | 1 | 0 | 0 | 1 | 0 | 0 | 0 | 0 | 1 | 2 | 0 | 2 | 0 | 0 | 5 |
| ENSMUSG00000029763 | Exoc4 | ENSMUST00000052266 | 28.9934354 | 26 | 23.3588621 | 22.0459519 | 0 | 0 | 0 | 0 | 0 | 0 | 0 | 0 | 0 | 0 | 1 | 0 | 1 | 0 | 2 |
| ENSMUSG00000061244 | Exoc5 | ENSMUST00000162175 | 32.2580645 | 34 | 16.0297767 | 17.8163772 | 0 | 0 | 0 | 1 | 0 | 0 | 0 | 0 | 1 | 2 | 0 | 0 | 1 | 0 | 3 |
| ENSMUSG00000034321 | Exosc1 | ENSMUST00000075280 | 26.4471058 | 32 | 23.4530938 | 18.2634731 | 0 | 0 | 0 | 0 | 0 | 0 | 0 | 0 | 0 | 0 | 0 | 16 | 1 | 0 | 17 |
| ENSMUSG00000027963 | Extl2 | ENSMUST00000029575 | 26.8597762 | 33 | 19.3548387 | 20.5398288 | 0 | 0 | 0 | 0 | 0 | 0 | 0 | 0 | 0 | 0 | 0 | 0 | 4 | 1 | 5 |
| ENSMUSG00000025932 | Eya1 | ENSMUST00000027066 | 29.3577982 | 32 | 18.5524975 | 19.775739 | 0 | 0 | 0 | 0 | 0 | 0 | 0 | 0 | 0 | 0 | 0 | 1 | 0 | 0 | 1 |
| ENSMUSG00000038235 | F11r | ENSMUST00000043839 | 22.9953612 | 29 | 21.471173 | 26.1762757 | 0 | 0 | 0 | 1 | 0 | 0 | 0 | 0 | 1 | 2 | 0 | 2 | 0 | 0 | 4 |
| ENSMUSG00000028128 | F3 | ENSMUST00000029771 | 29.1718171 | 32 | 17.1817058 | 21.2608158 | 0 | 1 | 0 | 2 | 0 | 0 | 0 | 1 | 1 | 4 | 0 | 0 | 0 | 0 | 5 |
| ENSMUSG00000010517 | Faf1 | ENSMUST00000102724 | 24.1578441 | 33 | 22.1847931 | 20.7410972 | 0 | 0 | 0 | 0 | 0 | 0 | 0 | 0 | 0 | 0 | 11 | 1 | 10 | 0 | 22 |
| ENSMUSG00000038646 | Fam103a1 | ENSMUST00000042166 | 29.5819936 | 37 | 14.0407288 | 19.6141479 | 0 | 0 | 0 | 1 | 0 | 0 | 0 | 0 | 1 | 2 | 1 | 0 | 1 | 0 | 4 |
| ENSMUSG00000056069 | Fam105a | ENSMUST00000100739 | 29.5566502 | 30 | 18.3360701 | 22.0580186 | 0 | 1 | 0 | 1 | 0 | 0 | 0 | 0 | 1 | 2 | 0 | 0 | 0 | 0 | 3 |
| ENSMUSG00000026655 | Fam107b | ENSMUST00000027965 | 26.4950495 | 32 | 20.2772277 | 21.3069307 | 0 | 0 | 0 | 0 | 0 | 0 | 0 | 0 | 0 | 0 | 0 | 1 | 0 | 0 | 1 |
| ENSMUSG00000038893 | Fam117a | ENSMUST00000037502 | 23.5487404 | 28 | 24.8630887 | 23.5487404 | 0 | 0 | 0 | 0 | 0 | 0 | 0 | 0 | 0 | 0 | 2 | 1 | 2 | 0 | 5 |
| ENSMUSG00000041040 | Fam117b | ENSMUST00000036540 | 26.0845537 | 34 | 18.4857695 | 21.3318596 | 0 | 0 | 0 | 0 | 0 | 0 | 0 | 0 | 0 | 0 | 1 | 0 | 2 | 0 | 3 |
| ENSMUSG00000025262 | Fam120c | ENSMUST00000073364 | 30.0662879 | 31 | 18.4185606 | 20.217803 | 1 | 0 | 1 | 0 | 0 | 0 | 0 | 0 | 0 | 1 | 0 | 1 | 2 | 0 | 5 |
| ENSMUSG00000036022 | Fam122b | ENSMUST00000071023 | 28.1581838 | 36 | 18.1618455 | 17.9055291 | 0 | 0 | 0 | 0 | 0 | 0 | 0 | 0 | 0 | 0 | 0 | 0 | 1 | 0 | 1 |
| ENSMUSG00000028995 | Fam126a | ENSMUST00000030849 | 27.3812614 | 30 | 19.6210745 | 22.7614846 | 0 | 1 | 0 | 0 | 0 | 0 | 0 | 0 | 0 | 0 | 0 | 0 | 0 | 0 | 1 |
| ENSMUSG00000038174 | Fam126b | ENSMUST00000161600 | 31.3776999 | 34 | 16.2872154 | 18.2720374 | 0 | 0 | 0 | 0 | 0 | 0 | 0 | 0 | 0 | 0 | 1 | 1 | 2 | 3 | 7 |
| ENSMUSG00000058503 | Fam133b | ENSMUST00000197082 | 27.8111588 | 33 | 18.7124464 | 20.4291845 | 0 | 0 | 0 | 0 | 0 | 0 | 0 | 0 | 0 | 0 | 1 | 0 | 2 | 0 | 3 |
| ENSMUSG00000022270 | Fam134b | ENSMUST00000022881 | 30.591103 | 33 | 17.794028 | 18.4034126 | 1 | 1 | 0 | 0 | 0 | 0 | 0 | 0 | 0 | 0 | 0 | 0 | 4 | 0 | 6 |
| ENSMUSG00000026153 | Fam135a | ENSMUST00000188712 | 29.5552677 | 31 | 19.4300518 | 19.8618307 | 0 | 0 | 0 | 0 | 0 | 0 | 0 | 0 | 0 | 0 | 1 | 1 | 0 | 0 | 2 |
| ENSMUSG00000036501 | Fam13b | ENSMUST00000040506 | 23.7931034 | 32 | 23.1034483 | 21.0344828 | 0 | 0 | 0 | 0 | 0 | 0 | 0 | 0 | 0 | 0 | 1 | 0 | 0 | 0 | 1 |
| ENSMUSG00000041817 | Fam169a | ENSMUST00000042517 | 30.4873137 | 36 | 15.505437 | 17.841321 | 1 | 1 | 0 | 2 | 0 | 0 | 0 | 0 | 2 | 4 | 0 | 0 | 6 | 0 | 12 |
| ENSMUSG00000064138 | Fam172a | ENSMUST00000091459 | 27.1777003 | 31 | 21.3704994 | 20.7510647 | 0 | 0 | 0 | 0 | 0 | 0 | 0 | 0 | 0 | 0 | 2 | 1 | 1 | 0 | 4 |
| ENSMUSG00000051185 | Fam174a | ENSMUST00000186780 | 27.4647887 | 33 | 18.7546331 | 20.8302446 | 0 | 0 | 0 | 0 | 0 | 0 | 0 | 0 | 0 | 0 | 2 | 0 | 3 | 1 | 6 |
| ENSMUSG00000036097 | Fam178a | ENSMUST00000096053 | 26.0012516 | 33 | 18.3041302 | 22.6846058 | 0 | 0 | 0 | 1 | 0 | 0 | 0 | 0 | 1 | 2 | 3 | 1 | 8 | 1 | 15 |
| ENSMUSG00000031774 | Fam192a | ENSMUST00000034226 | 23.4870317 | 28 | 24.7838617 | 23.4870317 | 0 | 0 | 0 | 0 | 0 | 0 | 0 | 0 | 0 | 0 | 0 | 2 | 0 | 0 | 2 |
| ENSMUSG00000042595 | Fam199x | ENSMUST00000047852 | 30.0598278 | 35 | 15.2196118 | 19.5097038 | 0 | 0 | 0 | 1 | 0 | 0 | 0 | 0 | 1 | 2 | 2 | 2 | 8 | 5 | 19 |
| ENSMUSG00000044071 | Fam19a2 | ENSMUST00000050756 | 29.2267366 | 34 | 17.9554391 | 18.6544343 | 0 | 0 | 0 | 0 | 0 | 0 | 0 | 0 | 0 | 0 | 0 | 0 | 7 | 0 | 7 |
| ENSMUSG00000057858 | Fam204a | ENSMUST00000065286 | 31.1764706 | 38 | 13.0588235 | 17.5294118 | 1 | 1 | 0 | 0 | 0 | 0 | 0 | 0 | 0 | 0 | 0 | 0 | 1 | 0 | 3 |
| ENSMUSG00000040651 | Fam208a | ENSMUST00000022450 | 30.4728546 | 36 | 15.4115587 | 18.5288967 | 1 | 0 | 0 | 0 | 0 | 0 | 0 | 0 | 0 | 0 | 5 | 1 | 12 | 3 | 22 |
| ENSMUSG00000033799 | Fam208b | ENSMUST00000096069 | 33.2802548 | 39 | 12.8980892 | 15.2866242 | 0 | 0 | 0 | 0 | 0 | 0 | 0 | 0 | 0 | 0 | 0 | 0 | 6 | 2 | 8 |
| ENSMUSG00000038121 | Fam210a | ENSMUST00000042852 | 27.0862635 | 30 | 20.2883263 | 22.2105016 | 1 | 1 | 0 | 0 | 0 | 0 | 0 | 0 | 0 | 0 | 1 | 0 | 3 | 0 | 6 |
| ENSMUSG00000034858 | Fam214a | ENSMUST00000081746 | 28.375 | 34 | 19.25 | 18.625 | 1 | 0 | 1 | 1 | 0 | 0 | 0 | 0 | 1 | 3 | 0 | 0 | 0 | 0 | 4 |
| ENSMUSG00000047115 | Fam221a | ENSMUST00000060561 | 24.2442563 | 30 | 22.1886336 | 23.8210399 | 2 | 1 | 2 | 1 | 0 | 0 | 0 | 0 | 1 | 4 | 0 | 4 | 1 | 0 | 12 |
| ENSMUSG00000042564 | Fam227a | ENSMUST00000109648 | 27.9918173 | 29 | 22.1616093 | 20.5932492 | 1 | 1 | 0 | 0 | 0 | 0 | 0 | 0 | 0 | 0 | 3 | 2 | 2 | 0 | 9 |
| ENSMUSG00000050545 | Fam228b | ENSMUST00000053458 | 23.7645752 | 37 | 17.1016102 | 21.8767351 | 2 | 1 | 0 | 3 | 0 | 0 | 0 | 1 | 2 | 6 | 2 | 0 | 1 | 0 | 12 |
| ENSMUSG00000030207 | Fam234b | ENSMUST00000111915 | 20.9034544 | 30 | 24.3135518 | 24.8892826 | 0 | 0 | 0 | 1 | 0 | 0 | 0 | 0 | 1 | 2 | 0 | 1 | 0 | 0 | 3 |
| ENSMUSG00000029672 | Fam3c | ENSMUST00000165576 | 26.6977709 | 35 | 17.3665111 | 21.0471747 | 0 | 0 | 0 | 0 | 0 | 0 | 0 | 0 | 0 | 0 | 3 | 0 | 23 | 1 | 27 |
| ENSMUSG00000046546 | Fam43a | ENSMUST00000059078 | 25.135428 | 29 | 20.8017335 | 25.460455 | 0 | 0 | 0 | 0 | 0 | 0 | 0 | 0 | 0 | 0 | 0 | 1 | 2 | 0 | 3 |
| ENSMUSG00000024993 | Fam45a | ENSMUST00000025957 | 31.1020408 | 33 | 19.4285714 | 16 | 0 | 0 | 0 | 0 | 0 | 0 | 0 | 0 | 0 | 0 | 0 | 0 | 0 | 1 | 1 |
| ENSMUSG00000032265 | Fam46a | ENSMUST00000034802 | 31.159781 | 30 | 18.914883 | 19.5868591 | 0 | 0 | 0 | 1 | 0 | 0 | 0 | 0 | 1 | 2 | 1 | 1 | 5 | 0 | 9 |
| ENSMUSG00000022378 | Fam49b | ENSMUST00000063838 | 28.3445812 | 32 | 18.6581977 | 20.5240175 | 1 | 0 | 0 | 2 | 0 | 0 | 0 | 0 | 2 | 4 | 0 | 0 | 0 | 1 | 6 |
| ENSMUSG00000039985 | Fam60a; LOC102636734 | ENSMUST00000203164 | 27.1450075 | 32 | 19.2172604 | 21.4249875 | 1 | 1 | 0 | 3 | 0 | 0 | 1 | 1 | 1 | 6 | 0 | 1 | 10 | 2 | 21 |
| ENSMUSG00000055184 | Fam72a | ENSMUST00000068613 | 29.7985154 | 34 | 20.8907741 | 14.8462354 | 0 | 0 | 1 | 0 | 0 | 0 | 0 | 0 | 0 | 1 | 0 | 0 | 0 | 0 | 1 |
| ENSMUSG00000054942 | Fam73a | ENSMUST00000073089 | 29.7918162 | 32 | 16.9059584 | 21.4285714 | 1 | 1 | 0 | 1 | 0 | 0 | 0 | 0 | 1 | 2 | 0 | 0 | 0 | 0 | 4 |
| ENSMUSG00000037808 | Fam76b | ENSMUST00000156680 | 30.5108967 | 35 | 15.7913541 | 18.5423365 | 1 | 0 | 0 | 0 | 0 | 0 | 0 | 0 | 0 | 0 | 0 | 0 | 0 | 0 | 1 |
| ENSMUSG00000069237 | Fam8a1 | ENSMUST00000099547 | 30.0416667 | 32 | 17.9583333 | 19.5416667 | 0 | 0 | 0 | 1 | 0 | 0 | 0 | 0 | 1 | 2 | 0 | 0 | 0 | 0 | 2 |
| ENSMUSG00000032381 | Fam96a | ENSMUST00000034945 | 25.5278311 | 36 | 19.3857965 | 19.3857965 | 0 | 0 | 0 | 0 | 0 | 0 | 0 | 0 | 0 | 0 | 0 | 0 | 1 | 0 | 1 |
| ENSMUSG00000028453 | Fancg | ENSMUST00000030165 | 19.3447738 | 38 | 23.7129485 | 18.7207488 | 0 | 0 | 0 | 0 | 0 | 0 | 0 | 0 | 0 | 0 | 0 | 2 | 1 | 0 | 3 |
| ENSMUSG00000030759 | Far1 | ENSMUST00000033018 | 31.2538414 | 36 | 14.2593731 | 18.8383528 | 0 | 1 | 0 | 0 | 0 | 0 | 0 | 0 | 0 | 0 | 3 | 1 | 4 | 0 | 9 |
| ENSMUSG00000030303 | Far2 | ENSMUST00000111607 | 31.9063927 | 30 | 20.4908676 | 17.7511416 | 0 | 0 | 0 | 0 | 0 | 0 | 0 | 0 | 0 | 0 | 0 | 0 | 0 | 1 | 1 |
| ENSMUSG00000026245 | Farsb | ENSMUST00000189529 | 28.6343612 | 27 | 22.5928257 | 21.7747011 | 1 | 0 | 0 | 0 | 0 | 0 | 0 | 0 | 0 | 0 | 0 | 0 | 0 | 0 | 1 |
| ENSMUSG00000025962 | Fastkd2 | ENSMUST00000027103 | 26.5659707 | 31 | 18.8804976 | 23.411817 | 0 | 0 | 0 | 0 | 0 | 0 | 0 | 0 | 0 | 0 | 0 | 1 | 0 | 0 | 1 |
| ENSMUSG00000024598 | Fbn2; LOC102642295 | ENSMUST00000025497 | 30.6422018 | 31 | 18.5321101 | 19.3272171 | 1 | 0 | 0 | 0 | 0 | 0 | 0 | 0 | 0 | 0 | 3 | 1 | 7 | 2 | 14 |
| ENSMUSG00000030019 | Fbxl14 | ENSMUST00000032094 | 22.7040816 | 33 | 21.747449 | 22.3852041 | 0 | 0 | 0 | 0 | 0 | 0 | 0 | 0 | 0 | 0 | 5 | 0 | 7 | 0 | 12 |
| ENSMUSG00000025738 | Fbxl16 | ENSMUST00000045692 | 22.1835075 | 21 | 31.5911731 | 25.261324 | 0 | 0 | 0 | 0 | 0 | 0 | 0 | 0 | 0 | 0 | 0 | 1 | 0 | 0 | 1 |
| ENSMUSG00000020883 | Fbxl20 | ENSMUST00000103143 | 23.7668799 | 27 | 23.1556503 | 25.8706468 | 1 | 1 | 0 | 0 | 0 | 0 | 0 | 0 | 0 | 0 | 0 | 16 | 0 | 0 | 18 |
| ENSMUSG00000035509 | Fbxl21 | ENSMUST00000124981 | 27.899108 | 29 | 21.9317133 | 20.8858813 | 1 | 0 | 0 | 0 | 0 | 0 | 0 | 0 | 0 | 0 | 0 | 7 | 1 | 2 | 11 |
| ENSMUSG00000022124 | Fbxl3 | ENSMUST00000022720 | 33.0289193 | 34 | 14.6879756 | 18.7214612 | 0 | 0 | 0 | 1 | 0 | 0 | 0 | 0 | 1 | 2 | 0 | 0 | 2 | 0 | 4 |
| ENSMUSG00000040410 | Fbxl4 | ENSMUST00000185029 | 28.379773 | 32 | 19.8658411 | 20.123839 | 0 | 0 | 0 | 0 | 0 | 0 | 0 | 0 | 0 | 0 | 1 | 0 | 1 | 0 | 2 |
| ENSMUSG00000039753 | Fbxl5 | ENSMUST00000087465 | 25.4799302 | 36 | 17.7574171 | 20.5061082 | 1 | 0 | 0 | 0 | 0 | 0 | 0 | 0 | 0 | 0 | 6 | 1 | 7 | 0 | 15 |
| ENSMUSG00000032898 | Fbxo21 | ENSMUST00000202447 | 21.8562874 | 29 | 26.1477046 | 23.1037924 | 0 | 0 | 0 | 1 | 0 | 0 | 0 | 0 | 1 | 2 | 0 | 1 | 0 | 1 | 4 |
| ENSMUSG00000037463 | Fbxo27 | ENSMUST00000039998 | 21.7573222 | 29 | 25.7322176 | 23.2217573 | 0 | 0 | 1 | 0 | 0 | 0 | 0 | 0 | 0 | 1 | 0 | 0 | 0 | 0 | 1 |
| ENSMUSG00000047539 | Fbxo28 | ENSMUST00000051431 | 26.4449723 | 30 | 21.7207707 | 21.4040644 | 0 | 0 | 0 | 0 | 0 | 0 | 0 | 0 | 0 | 0 | 1 | 1 | 3 | 3 | 8 |
| ENSMUSG00000027180 | Fbxo3 | ENSMUST00000028603 | 29.5721078 | 31 | 19.207607 | 20.570523 | 0 | 0 | 0 | 0 | 0 | 0 | 0 | 0 | 0 | 0 | 1 | 1 | 2 | 0 | 4 |
| ENSMUSG00000047648 | Fbxo30 | ENSMUST00000129456 | 32.3708207 | 35 | 15.0075988 | 17.1352584 | 0 | 0 | 0 | 0 | 0 | 0 | 0 | 0 | 0 | 0 | 0 | 0 | 4 | 0 | 4 |
| ENSMUSG00000035329 | Fbxo33 | ENSMUST00000043204 | 31.8311466 | 33 | 16.2578437 | 19.3382772 | 0 | 0 | 0 | 0 | 0 | 0 | 0 | 0 | 0 | 0 | 1 | 0 | 1 | 0 | 2 |
| ENSMUSG00000028920 | Fbxo42 | ENSMUST00000030757 | 22.0697541 | 28 | 26.1006289 | 23.4705546 | 0 | 0 | 0 | 0 | 0 | 0 | 0 | 0 | 0 | 0 | 1 | 2 | 1 | 0 | 4 |
| ENSMUSG00000035764 | Fbxo45 | ENSMUST00000042732 | 28.1050151 | 31 | 17.8054527 | 23.2581622 | 0 | 0 | 1 | 1 | 0 | 0 | 0 | 0 | 1 | 3 | 2 | 0 | 3 | 0 | 8 |
| ENSMUSG00000044966 | Fbxo48 | ENSMUST00000061327 | 35.6979405 | 39 | 9.83981693 | 15.3318078 | 0 | 0 | 0 | 1 | 0 | 0 | 0 | 0 | 1 | 2 | 0 | 0 | 0 | 0 | 2 |
| ENSMUSG00000001366 | Fbxo9 | ENSMUST00000085311 | 30.3534304 | 32 | 16.6320166 | 20.7900208 | 0 | 0 | 1 | 0 | 0 | 0 | 0 | 0 | 0 | 1 | 0 | 0 | 0 | 2 | 3 |
| ENSMUSG00000020271 | Fbxw11 | ENSMUST00000076383 | 22.7607096 | 30 | 23.4530506 | 23.3232367 | 0 | 0 | 0 | 0 | 0 | 0 | 0 | 0 | 0 | 0 | 3 | 0 | 1 | 0 | 4 |
| ENSMUSG00000005540 | Fcer2a | ENSMUST00000005678 | 23.6719478 | 24 | 31.0344828 | 21.248835 | 0 | 0 | 0 | 0 | 0 | 0 | 0 | 0 | 0 | 0 | 0 | 2 | 0 | 0 | 2 |
| ENSMUSG00000041685 | Fcho2 | ENSMUST00000099277 | 33.4879798 | 36 | 15.0147617 | 15.7317588 | 1 | 0 | 0 | 2 | 0 | 0 | 0 | 0 | 2 | 4 | 0 | 0 | 1 | 0 | 6 |
| ENSMUSG00000024588 | Fech | ENSMUST00000025484 | 27.9843444 | 32 | 21.4611872 | 18.7214612 | 0 | 0 | 0 | 0 | 0 | 0 | 0 | 0 | 0 | 0 | 0 | 1 | 0 | 0 | 1 |
| ENSMUSG00000037712 | Fermt2 | ENSMUST00000045905 | 29.6786389 | 34 | 18.9981096 | 17.5803403 | 0 | 0 | 0 | 0 | 0 | 0 | 0 | 0 | 0 | 0 | 0 | 0 | 0 | 2 | 2 |
| ENSMUSG00000022788 | Fgd4 | ENSMUST00000161861 | 32.2529224 | 35 | 16.135317 | 17.0917464 | 2 | 1 | 0 | 3 | 0 | 0 | 0 | 0 | 3 | 6 | 0 | 3 | 2 | 1 | 15 |
| ENSMUSG00000020021 | Fgd6 | ENSMUST00000020208 | 25.5492958 | 32 | 23.7746479 | 18.8732394 | 0 | 0 | 1 | 0 | 0 | 0 | 0 | 0 | 0 | 1 | 0 | 2 | 1 | 2 | 6 |
| ENSMUSG00000031137 | Fgf13 | ENSMUST00000033473 | 27.8818444 | 34 | 21.2536023 | 16.4985591 | 0 | 0 | 0 | 0 | 0 | 0 | 0 | 0 | 0 | 0 | 0 | 49 | 0 | 0 | 49 |

| | | | | | | | | | | | | | | | | | | | | | |
|---|---|---|---|---|---|---|---|---|---|---|---|---|---|---|---|---|---|---|---|---|---|
| ENSMUSG00000025551 | Fgf14 | ENSMUST00000095529 | 28.2492825 | 35 | 19.0651907 | 17.9581796 | 2 | 1 | 0 | 1 | 0 | 0 | 0 | 0 | 1 | 2 | 0 | 0 | 5 | 0 | 10 |
| ENSMUSG00000031603 | Fgf20 | ENSMUST00000034014 | 35.1851852 | 32 | 15.4882155 | 17.6767677 | 3 | 1 | 1 | 4 | 0 | 0 | 0 | 1 | 3 | 9 | 0 | 0 | 0 | 0 | 13 |
| ENSMUSG00000021974 | Fgf9 | ENSMUST00000165526 | 28.1261327 | 33 | 19.1736136 | 19.4273287 | 4 | 2 | 2 | 6 | 0 | 0 | 0 | 2 | 4 | 14 | 1 | 0 | 6 | 0 | 27 |
| ENSMUSG00000047632 | Fgfbp3 | ENSMUST00000057337 | 32.4945295 | 30 | 15.4266958 | 21.7724289 | 0 | 0 | 0 | 1 | 0 | 0 | 0 | 0 | 1 | 2 | 0 | 0 | 1 | 1 | 4 |
| ENSMUSG00000069135 | Fgfr1op | ENSMUST00000097419 | 27.7501132 | 32 | 18.9225894 | 21.4124038 | 0 | 0 | 0 | 0 | 0 | 0 | 0 | 0 | 0 | 0 | 0 | 0 | 4 | 1 | 5 |
| ENSMUSG00000039899 | Fgl2 | ENSMUST00000035799 | 31.938691 | 35 | 16.7357084 | 16.8185584 | 1 | 0 | 0 | 0 | 0 | 0 | 0 | 0 | 0 | 0 | 0 | 0 | 0 | 0 | 1 |
| ENSMUSG00000075324 | Fign | ENSMUST00000131615 | 29.377977 | 33 | 19.3611656 | 18.3804987 | 0 | 0 | 0 | 1 | 0 | 0 | 0 | 0 | 1 | 2 | 3 | 45 | 7 | 1 | 58 |
| ENSMUSG00000029227 | Fip1l1 | ENSMUST00000113536 | 30.4193319 | 32 | 17.803838 | 19.7938877 | 0 | 0 | 0 | 0 | 0 | 0 | 0 | 0 | 0 | 0 | 0 | 0 | 0 | 1 | 1 |
| ENSMUSG00000038074 | Fkbp14 | ENSMUST00000155047 | 26.0471204 | 32 | 20.8987784 | 21.1169284 | 0 | 0 | 0 | 1 | 0 | 0 | 0 | 0 | 1 | 2 | 0 | 6 | 0 | 0 | 8 |
| ENSMUSG00000032966 | Fkbp1a | ENSMUST00000044011 | 22.4231465 | 34 | 22.4231465 | 21.6094033 | 0 | 0 | 0 | 0 | 0 | 0 | 0 | 0 | 0 | 0 | 0 | 1 | 0 | 0 | 1 |
| ENSMUSG00000048920 | Fkrp | ENSMUST00000061390 | 24.1493776 | 28 | 24.1493776 | 23.9834025 | 0 | 0 | 0 | 0 | 0 | 0 | 0 | 0 | 0 | 0 | 0 | 2 | 0 | 0 | 2 |
| ENSMUSG00000028414 | Fktn | ENSMUST00000128667 | 29.0359254 | 29 | 21.0095498 | 20.6684857 | 2 | 2 | 0 | 1 | 0 | 0 | 0 | 0 | 1 | 2 | 0 | 1 | 0 | 0 | 7 |
| ENSMUSG00000016087 | Fli1 | ENSMUST00000016231 | 30.2798982 | 36 | 15.9669211 | 17.6844784 | 0 | 0 | 0 | 0 | 0 | 0 | 0 | 0 | 0 | 0 | 4 | 0 | 16 | 3 | 23 |
| ENSMUSG00000047414 | Flrt2 | ENSMUST00000057324 | 31.4529127 | 32 | 17.9829611 | 18.8809579 | 0 | 0 | 0 | 1 | 0 | 0 | 0 | 0 | 1 | 2 | 0 | 6 | 0 | 1 | 9 |
| ENSMUSG00000029648 | Flt1 | ENSMUST00000110529 | 28.7333165 | 29 | 19.4157643 | 22.4880383 | 0 | 0 | 0 | 0 | 0 | 0 | 0 | 0 | 0 | 0 | 0 | 1 | 3 | 1 | 5 |
| ENSMUSG00000036053 | Fmnl2 | ENSMUST00000049483 | 29.5180723 | 35 | 19.2771084 | 16.4802065 | 0 | 0 | 0 | 0 | 0 | 0 | 0 | 0 | 0 | 0 | 0 | 2 | 0 | 0 | 2 |
| ENSMUSG00000075415 | Fnbp1 | ENSMUST00000073879 | 22.6337449 | 28 | 26.2492651 | 23.3686067 | 0 | 0 | 0 | 0 | 0 | 0 | 0 | 0 | 0 | 0 | 0 | 11 | 14 | 1 | 26 |
| ENSMUSG00000039735 | Fnbp1l | ENSMUST00000162409 | 26.5976331 | 33 | 18.6390533 | 22.1005917 | 0 | 0 | 0 | 0 | 0 | 0 | 0 | 0 | 0 | 0 | 0 | 1 | 0 | 0 | 1 |
| ENSMUSG00000008200 | Fnbp4 | ENSMUST00000013759 | 28.0291971 | 34 | 18.2481752 | 19.270073 | 0 | 0 | 0 | 0 | 0 | 0 | 0 | 0 | 0 | 0 | 0 | 1 | 0 | 0 | 1 |
| ENSMUSG00000033487 | Fndc3a | ENSMUST00000089017 | 29.716344 | 34 | 17.9198559 | 18.5952274 | 0 | 0 | 0 | 1 | 0 | 0 | 0 | 0 | 1 | 2 | 0 | 4 | 0 | 0 | 6 |
| ENSMUSG00000039286 | Fndc3b | ENSMUST00000046157 | 27.1013103 | 34 | 19.5270054 | 19.3991691 | 0 | 0 | 0 | 0 | 0 | 0 | 0 | 0 | 0 | 0 | 1 | 0 | 1 | 2 | 4 |
| ENSMUSG00000035992 | Fnip1 | ENSMUST00000046835 | 31.9292967 | 36 | 14.5543437 | 17.074088 | 0 | 0 | 0 | 1 | 0 | 0 | 0 | 0 | 1 | 2 | 1 | 0 | 1 | 3 | 7 |
| ENSMUSG00000038368 | Focad | ENSMUST00000107147 | 30.529595 | 29 | 18.6915888 | 22.1183801 | 0 | 0 | 0 | 0 | 0 | 0 | 0 | 0 | 0 | 0 | 0 | 1 | 0 | 0 | 1 |
| ENSMUSG00000078302 | Foxd1 | ENSMUST00000105098 | 27.9102384 | 31 | 20.1963534 | 21.3183731 | 0 | 0 | 0 | 0 | 0 | 0 | 0 | 0 | 0 | 0 | 0 | 1 | 0 | 0 | 1 |
| ENSMUSG00000055210 | Foxd2 | ENSMUST00000068654 | 22.5259189 | 28 | 24.5051838 | 24.8821866 | 0 | 0 | 0 | 0 | 0 | 0 | 0 | 0 | 0 | 0 | 0 | 2 | 0 | 2 | 4 |
| ENSMUSG00000038402 | Foxf2 | ENSMUST00000042054 | 26.6490765 | 30 | 21.6358839 | 22.0316623 | 0 | 0 | 0 | 0 | 0 | 0 | 0 | 0 | 0 | 0 | 0 | 0 | 1 | 1 | 2 |
| ENSMUSG00000020950 | Foxg1 | ENSMUST00000021333 | 30.8602151 | 35 | 15.2688172 | 19.1397849 | 0 | 0 | 1 | 0 | 0 | 0 | 0 | 0 | 0 | 1 | 0 | 0 | 0 | 0 | 1 |
| ENSMUSG00000032998 | Foxj3 | ENSMUST00000044564 | 26.1510129 | 31 | 17.9005525 | 24.5672192 | 0 | 0 | 0 | 1 | 0 | 0 | 0 | 0 | 1 | 2 | 0 | 0 | 0 | 0 | 2 |
| ENSMUSG00000034998 | Foxn2 | ENSMUST00000112238 | 28.7496535 | 35 | 17.0779041 | 19.3512614 | 2 | 0 | 0 | 1 | 0 | 0 | 0 | 0 | 1 | 2 | 0 | 1 | 11 | 0 | 16 |
| ENSMUSG00000044167 | Foxo1 | ENSMUST00000053764 | 26.5865385 | 30 | 21.3942308 | 22.0512821 | 0 | 0 | 0 | 0 | 0 | 0 | 0 | 0 | 0 | 0 | 0 | 2 | 5 | 0 | 7 |
| ENSMUSG00000029563 | Foxp2 | ENSMUST00000115477 | 32.0778564 | 32 | 17.6188069 | 18.0738119 | 1 | 0 | 1 | 0 | 0 | 0 | 0 | 0 | 0 | 1 | 3 | 1 | 4 | 0 | 10 |
| ENSMUSG00000053870 | Fpgt | ENSMUST00000066568 | 32.9210836 | 33 | 15.4888104 | 18.138987 | 1 | 0 | 0 | 1 | 0 | 0 | 0 | 0 | 1 | 2 | 1 | 0 | 1 | 0 | 5 |
| ENSMUSG00000031590 | Frg1 | ENSMUST00000033999 | 10.5960265 | 52 | 23.1788079 | 13.9072848 | 0 | 0 | 0 | 0 | 0 | 0 | 0 | 0 | 0 | 0 | 0 | 19 | 0 | 0 | 19 |
| ENSMUSG00000049122 | Frmd3 | ENSMUST00000084474 | 35.6083086 | 32 | 12.9574679 | 19.6834817 | 1 | 0 | 0 | 1 | 0 | 0 | 0 | 0 | 1 | 2 | 0 | 0 | 2 | 0 | 5 |
| ENSMUSG00000027238 | Frmd5 | ENSMUST00000110593 | 29.5454545 | 26 | 23.4710744 | 20.9504132 | 0 | 0 | 1 | 1 | 0 | 0 | 0 | 0 | 1 | 3 | 0 | 1 | 0 | 0 | 4 |
| ENSMUSG00000048285 | Frmd6 | ENSMUST00000057859 | 25.5070203 | 31 | 20.124805 | 23.400936 | 0 | 0 | 0 | 0 | 0 | 0 | 0 | 0 | 0 | 0 | 1 | 1 | 0 | 0 | 2 |
| ENSMUSG00000033386 | Frrs1 | ENSMUST00000195905 | 27.1331738 | 30 | 21.1124402 | 21.8102073 | 0 | 0 | 0 | 0 | 0 | 0 | 0 | 0 | 0 | 0 | 0 | 57 | 5 | 4 | 66 |
| ENSMUSG00000045589 | Frrs1l | ENSMUST00000053681 | 28.8907997 | 31 | 19.6732588 | 20.2923474 | 2 | 1 | 0 | 0 | 0 | 0 | 0 | 0 | 0 | 0 | 0 | 12 | 1 | 0 | 16 |
| ENSMUSG00000020170 | Frs2 | ENSMUST00000020381 | 24.1315789 | 33 | 22.0789474 | 21.2631579 | 0 | 1 | 0 | 0 | 0 | 0 | 0 | 0 | 0 | 0 | 1 | 15 | 0 | 0 | 17 |
| ENSMUSG00000094595 | Fsbp | ENSMUST00000180239 | 29.0646208 | 33 | 16.5535556 | 20.9206256 | 1 | 1 | 0 | 2 | 0 | 0 | 0 | 0 | 2 | 4 | 0 | 2 | 0 | 1 | 9 |
| ENSMUSG00000054752 | Fsd1l | ENSMUST00000132151 | 29.473144 | 36 | 16.9175505 | 17.9267875 | 0 | 0 | 0 | 0 | 0 | 0 | 0 | 0 | 0 | 0 | 3 | 0 | 8 | 1 | 12 |
| ENSMUSG00000021765 | Fst | ENSMUST00000022287 | 31.2661499 | 34 | 16.2790698 | 18.2601206 | 0 | 1 | 1 | 0 | 0 | 0 | 0 | 0 | 0 | 1 | 0 | 0 | 0 | 0 | 2 |
| ENSMUSG00000022816 | Fstl1 | ENSMUST00000114763 | 26.8547141 | 29 | 21.0200927 | 23.2612056 | 0 | 0 | 0 | 0 | 0 | 0 | 0 | 0 | 0 | 0 | 0 | 0 | 6 | 0 | 6 |
| ENSMUSG00000034098 | Fstl5 | ENSMUST00000038364 | 31.7046688 | 34 | 16.7752443 | 17.5895765 | 0 | 0 | 0 | 1 | 0 | 0 | 0 | 0 | 1 | 2 | 0 | 0 | 0 | 1 | 3 |
| ENSMUSG00000028034 | Fubp1 | ENSMUST00000166984 | 29.6405155 | 37 | 14.4698169 | 18.5394529 | 0 | 0 | 0 | 1 | 0 | 0 | 0 | 0 | 1 | 2 | 0 | 1 | 2 | 0 | 5 |
| ENSMUSG00000026843 | Fubp3 | ENSMUST00000055244 | 24.5848956 | 35 | 18.8002142 | 21.5318693 | 0 | 0 | 0 | 0 | 0 | 0 | 0 | 0 | 0 | 0 | 0 | 2 | 7 | 0 | 9 |
| ENSMUSG00000019810 | Fuca2 | ENSMUST00000121465 | 29.9426934 | 28 | 21.4422159 | 20.9169054 | 1 | 0 | 0 | 1 | 0 | 0 | 0 | 0 | 1 | 2 | 0 | 2 | 2 | 2 | 9 |
| ENSMUSG00000025040 | Fundc1 | ENSMUST00000026016 | 28.2481752 | 38 | 17.1532847 | 17.080292 | 0 | 0 | 1 | 0 | 0 | 0 | 0 | 0 | 0 | 1 | 0 | 1 | 1 | 0 | 3 |
| ENSMUSG00000031198 | Fundc2 | ENSMUST00000033541 | 30.453668 | 36 | 17.7123552 | 15.8783784 | 1 | 0 | 0 | 0 | 0 | 0 | 0 | 0 | 0 | 0 | 1 | 0 | 0 | 2 | 4 |
| ENSMUSG00000039357 | Fut11 | ENSMUST00000048016 | 26.2642741 | 33 | 19.6302338 | 21.0440457 | 0 | 0 | 0 | 0 | 0 | 0 | 0 | 0 | 0 | 0 | 3 | 1 | 4 | 0 | 8 |
| ENSMUSG00000055373 | Fut9 | ENSMUST00000084770 | 34.3428516 | 33 | 16.0080761 | 16.4887992 | 1 | 1 | 0 | 0 | 0 | 0 | 0 | 0 | 0 | 0 | 1 | 0 | 4 | 2 | 9 |
| ENSMUSG00000022800 | Fyttd1 | ENSMUST00000171325 | 29.7629763 | 35 | 16.3216322 | 19.1719172 | 0 | 0 | 0 | 0 | 0 | 0 | 0 | 0 | 0 | 0 | 0 | 0 | 7 | 1 | 8 |
| ENSMUSG00000044674 | Fzd1 | ENSMUST00000054294 | 25.3723111 | 30 | 22.6696084 | 21.6767788 | 0 | 0 | 1 | 0 | 0 | 0 | 0 | 0 | 0 | 1 | 0 | 0 | 0 | 2 | 3 |
| ENSMUSG00000022297 | Fzd6 | ENSMUST00000022906 | 26.4093137 | 34 | 19.3627451 | 20.2205882 | 0 | 0 | 0 | 0 | 0 | 0 | 0 | 0 | 0 | 0 | 6 | 1 | 6 | 0 | 13 |
| ENSMUSG00000036904 | Fzd8 | ENSMUST00000041080 | 27.7777778 | 29 | 21.5686275 | 21.5686275 | 0 | 0 | 1 | 0 | 0 | 0 | 0 | 0 | 0 | 1 | 0 | 0 | 0 | 0 | 1 |
| ENSMUSG00000035293 | G2e3 | ENSMUST00000054308 | 29.4559099 | 36 | 16.5103189 | 18.2223265 | 1 | 0 | 0 | 0 | 0 | 0 | 0 | 0 | 0 | 0 | 9 | 0 | 12 | 2 | 24 |
| ENSMUSG00000008976 | Gabpa | ENSMUST00000009120 | 30.4785894 | 33 | 16.9080605 | 19.2065491 | 0 | 0 | 0 | 2 | 0 | 0 | 0 | 0 | 2 | 4 | 1 | 1 | 0 | 2 | 8 |
| ENSMUSG00000027361 | Gabpb1 | ENSMUST00000110425 | 26.6499283 | 31 | 21.1262554 | 21.6284075 | 0 | 0 | 0 | 0 | 0 | 0 | 0 | 0 | 0 | 0 | 0 | 1 | 1 | 0 | 2 |
| ENSMUSG00000010803 | Gabra1 | ENSMUST00000020707 | 32.992126 | 34 | 16.1811024 | 16.9685039 | 0 | 0 | 0 | 1 | 0 | 0 | 0 | 0 | 1 | 2 | 0 | 0 | 0 | 1 | 3 |
| ENSMUSG00000000560 | Gabra2 | ENSMUST00000197284 | 33.745541 | 30 | 17.6932224 | 18.1450654 | 1 | 1 | 0 | 2 | 0 | 0 | 0 | 0 | 2 | 4 | 0 | 1 | 3 | 0 | 10 |
| ENSMUSG00000029212 | Gabrb1 | ENSMUST00000031122 | 30.2420674 | 30 | 21.229964 | 18.7602224 | 2 | 0 | 0 | 4 | 0 | 0 | 0 | 0 | 4 | 8 | 0 | 0 | 2 | 0 | 12 |
| ENSMUSG00000007653 | Gabrb2 | ENSMUST00000007797 | 28.1703322 | 32 | 18.0627047 | 21.853065 | 0 | 0 | 0 | 0 | 0 | 0 | 0 | 0 | 0 | 0 | 1 | 0 | 1 | 0 | 2 |
| ENSMUSG00000033676 | Gabrb3 | ENSMUST00000039697 | 30.126144 | 31 | 20.8013851 | 18.253772 | 0 | 0 | 0 | 0 | 0 | 0 | 0 | 0 | 0 | 0 | 0 | 2 | 0 | 0 | 2 |
| ENSMUSG00000001260 | Gabrg1 | ENSMUST00000031119 | 34.6554622 | 35 | 14.3529412 | 16.1008403 | 0 | 0 | 0 | 2 | 0 | 0 | 0 | 0 | 2 | 4 | 1 | 0 | 1 | 0 | 6 |
| ENSMUSG00000020436 | Gabrg2 | ENSMUST00000070725 | 32.3237104 | 34 | 16.3748225 | 17.3686701 | 0 | 0 | 0 | 0 | 0 | 0 | 0 | 0 | 0 | 0 | 1 | 1 | 1 | 0 | 3 |
| ENSMUSG00000031344 | Gabrq | ENSMUST00000114553 | 28.409459 | 30 | 21.218011 | 20.3757694 | 0 | 0 | 0 | 0 | 0 | 0 | 0 | 0 | 0 | 0 | 1 | 0 | 4 | 0 | 5 |
| ENSMUSG00000070880 | Gad1 | ENSMUST00000094934 | 29.0640394 | 32 | 19.2939245 | 19.7865353 | 0 | 0 | 0 | 0 | 0 | 0 | 0 | 0 | 0 | 0 | 0 | 1 | 0 | 0 | 1 |
| ENSMUSG00000026787 | Gad2 | ENSMUST00000028123 | 31.2358277 | 30 | 20.4931973 | 18.5941043 | 1 | 0 | 0 | 2 | 0 | 0 | 0 | 0 | 2 | 4 | 0 | 1 | 0 | 1 | 7 |
| ENSMUSG00000036390 | Gadd45a | ENSMUST00000043098 | 35.1687389 | 30 | 13.8543517 | 20.7815275 | 0 | 0 | 0 | 0 | 0 | 0 | 0 | 0 | 0 | 0 | 0 | 0 | 1 | 0 | 1 |
| ENSMUSG00000021003 | Galc | ENSMUST00000021390 | 25.5956017 | 31 | 21.5027489 | 21.4416616 | 0 | 0 | 0 | 0 | 0 | 0 | 0 | 0 | 0 | 0 | 0 | 6 | 1 | 0 | 7 |
| ENSMUSG00000060988 | Galnt13 | ENSMUST00000112635 | 32.9281975 | 34 | 17.0905011 | 15.6881077 | 1 | 1 | 0 | 0 | 0 | 0 | 0 | 0 | 0 | 0 | 3 | 2 | 10 | 0 | 17 |
| ENSMUSG00000026994 | Galnt3 | ENSMUST00000028378 | 33.6182336 | 34 | 16.0968661 | 16.2393162 | 0 | 0 | 1 | 0 | 0 | 0 | 0 | 0 | 0 | 1 | 0 | 0 | 0 | 0 | 1 |
| ENSMUSG00000031608 | Galnt7 | ENSMUST00000034021 | 30.5336833 | 33 | 17.3665792 | 19.0288714 | 0 | 0 | 0 | 1 | 0 | 0 | 0 | 0 | 1 | 2 | 0 | 1 | 3 | 0 | 6 |
| ENSMUSG00000047261 | Gap43 | ENSMUST00000102817 | 27.427598 | 32 | 21.2947189 | 18.9097104 | 0 | 0 | 0 | 0 | 0 | 0 | 0 | 0 | 0 | 0 | 0 | 2 | 0 | 0 | 2 |
| ENSMUSG00000074802 | Gas2l3 | ENSMUST00000099374 | 29.6803653 | 33 | 18.5844749 | 18.9726027 | 3 | 2 | 0 | 2 | 0 | 0 | 0 | 0 | 2 | 4 | 0 | 0 | 10 | 2 | 21 |
| ENSMUSG00000015619 | Gata3 | ENSMUST00000102976 | 31.0371517 | 32 | 16.0216718 | 20.8978328 | 1 | 1 | 0 | 1 | 0 | 0 | 0 | 0 | 1 | 2 | 1 | 1 | 12 | 0 | 18 |
| ENSMUSG00000042390 | Gatad2b | ENSMUST00000199754 | 26.4569843 | 30 | 21.9611471 | 21.5911193 | 0 | 0 | 0 | 0 | 0 | 0 | 0 | 0 | 0 | 0 | 0 | 3 | 10 | 2 | 15 |
| ENSMUSG00000029432 | Gbas | ENSMUST00000086046 | 28.4628378 | 30 | 21.7905405 | 20.1858108 | 0 | 0 | 0 | 0 | 0 | 0 | 0 | 0 | 0 | 0 | 0 | 0 | 6 | 2 | 8 |
| ENSMUSG00000026893 | Gca | ENSMUST00000028257 | 29.3767368 | 36 | 17.1496626 | 17.348154 | 1 | 0 | 0 | 0 | 0 | 0 | 0 | 0 | 0 | 0 | 1 | 0 | 2 | 4 | 8 |
| ENSMUSG00000035125 | Gcfc2 | ENSMUST00000152996 | 29.5581136 | 29 | 19.494987 | 22.4285184 | 1 | 0 | 0 | 0 | 0 | 0 | 0 | 0 | 0 | 0 | 0 | 0 | 0 | 0 | 1 |
| ENSMUSG00000037580 | Gch1 | ENSMUST00000089959 | 23.4366789 | 33 | 20.5990541 | 23.3315817 | 1 | 0 | 2 | 1 | 0 | 0 | 0 | 0 | 1 | 4 | 0 | 0 | 1 | 0 | 6 |
| ENSMUSG00000028124 | Gclm | ENSMUST00000029769 | 30.9803922 | 32 | 18.0622837 | 19.1464821 | 0 | 0 | 0 | 0 | 0 | 0 | 0 | 0 | 0 | 0 | 1 | 4 | 1 | 2 | 8 |
| ENSMUSG00000025777 | Gdap1 | ENSMUST00000026879 | 31.0282404 | 31 | 16.871832 | 21.0716872 | 0 | 0 | 0 | 0 | 0 | 0 | 0 | 0 | 0 | 0 | 0 | 3 | 0 | 0 | 3 |
| ENSMUSG00000061666 | Gdpd1 | ENSMUST00000020804 | 27.4119449 | 31 | 18.8361409 | 22.8943338 | 0 | 0 | 0 | 0 | 0 | 0 | 0 | 0 | 0 | 0 | 0 | 0 | 2 | 2 | 4 |
| ENSMUSG00000028214 | Gem | ENSMUST00000029868 | 29.3293293 | 35 | 17.017017 | 18.3183183 | 1 | 0 | 0 | 1 | 0 | 0 | 0 | 0 | 1 | 2 | 0 | 1 | 0 | 0 | 4 |
| ENSMUSG00000060121 | Gemin2 | ENSMUST00000021379 | 27.0893372 | 31 | 17.8674352 | 23.7752161 | 1 | 0 | 0 | 0 | 0 | 0 | 0 | 0 | 0 | 0 | 0 | 0 | 0 | 0 | 1 |
| ENSMUSG00000055760 | Gemin6 | ENSMUST00000069486 | 31.25 | 34 | 12.7604167 | 22.3958333 | 0 | 0 | 0 | 1 | 0 | 0 | 0 | 0 | 1 | 2 | 0 | 0 | 0 | 0 | 2 |
| ENSMUSG00000027774 | Gfm1 | ENSMUST00000077271 | 27.4568195 | 31 | 19.35676 | 22.4538416 | 0 | 0 | 0 | 0 | 0 | 0 | 0 | 0 | 0 | 0 | 0 | 0 | 1 | 1 | 2 |
| ENSMUSG00000021666 | Gfm2 | ENSMUST00000161825 | 28.3115672 | 25 | 23.6940299 | 22.6679104 | 0 | 0 | 0 | 0 | 0 | 0 | 0 | 0 | 0 | 0 | 0 | 0 | 0 | 1 | 1 |
| ENSMUSG00000029992 | Gfpt1 | ENSMUST00000113658 | 25.8479822 | 32 | 19.8564001 | 22.2084674 | 1 | 1 | 1 | 1 | 0 | 0 | 0 | 0 | 1 | 3 | 0 | 1 | 2 | 1 | 9 |
| ENSMUSG00000020363 | Gfpt2 | ENSMUST00000020629 | 22.6415094 | 31 | 26.7688679 | 20.0471698 | 0 | 0 | 0 | 1 | 0 | 0 | 0 | 0 | 1 | 2 | 0 | 0 | 0 | 0 | 2 |
| ENSMUSG00000030872 | Gga2 | ENSMUST00000033160 | 24.5200404 | 28 | 22.8022903 | 24.5200404 | 0 | 0 | 0 | 0 | 0 | 0 | 0 | 0 | 0 | 0 | 2 | 0 | 1 | 0 | 3 |

| | | | | | | | | | | | | | | | | | | | | | |
|---|---|---|---|---|---|---|---|---|---|---|---|---|---|---|---|---|---|---|---|---|---|
| ENSMUSG00000073987 | Ggh | ENSMUST00000098242 | 26.2092238 | 38 | 17.0978628 | 18.56018 | 0 | 0 | 0 | 0 | 0 | 0 | 0 | 0 | 0 | 0 | 0 | 0 | 2 | 0 | 2 |
| ENSMUSG00000021302 | Ggps1 | ENSMUST00000170957 | 30.1204819 | 31 | 20.3212851 | 18.3935743 | 2 | 1 | 0 | 1 | 0 | 0 | 0 | 0 | 1 | 2 | 1 | 0 | 5 | 4 | 15 |
| ENSMUSG00000035778 | Ggta1 | ENSMUST00000079424 | 28.6010363 | 28 | 20.4663212 | 22.746114 | 1 | 0 | 0 | 1 | 0 | 0 | 0 | 0 | 1 | 2 | 0 | 0 | 0 | 2 | 5 |
| ENSMUSG00000041028 | Ghitm | ENSMUST00000042564 | 27.5029446 | 33 | 18.7279152 | 20.9658422 | 0 | 0 | 0 | 0 | 0 | 0 | 0 | 0 | 0 | 0 | 0 | 0 | 3 | 0 | 3 |
| ENSMUSG00000018415 | Gid4 | ENSMUST00000070681 | 26.244204 | 28 | 22.7511592 | 23.2457496 | 0 | 0 | 0 | 0 | 0 | 0 | 0 | 0 | 0 | 0 | 0 | 0 | 10 | 0 | 10 |
| ENSMUSG00000048000 | Gigyf2 | ENSMUST00000027475 | 24.954573 | 30 | 22.7740763 | 22.1078134 | 0 | 0 | 0 | 0 | 0 | 0 | 0 | 0 | 0 | 0 | 0 | 2 | 0 | 0 | 2 |
| ENSMUSG00000040006 | Ginm1 | ENSMUST00000039763 | 34.5283019 | 30 | 19.4339623 | 16.2264151 | 1 | 0 | 0 | 0 | 0 | 0 | 0 | 0 | 0 | 0 | 0 | 0 | 0 | 0 | 1 |
| ENSMUSG00000031821 | Gins2 | ENSMUST00000034278 | 24.4677367 | 24 | 23.7143793 | 28.0707501 | 0 | 0 | 0 | 0 | 0 | 0 | 0 | 0 | 0 | 0 | 0 | 2 | 1 | 2 | 5 |
| ENSMUSG00000050953 | Gja1 | ENSMUST00000068581 | 28.5382831 | 33 | 17.2853828 | 20.7076566 | 0 | 0 | 0 | 0 | 0 | 0 | 0 | 0 | 0 | 0 | 0 | 2 | 6 | 0 | 8 |
| ENSMUSG00000046352 | Gjb2 | ENSMUST00000055698 | 28.6589517 | 28 | 22.2600408 | 21.1708645 | 0 | 0 | 0 | 0 | 0 | 0 | 0 | 0 | 0 | 0 | 0 | 4 | 0 | 1 | 5 |
| ENSMUSG00000025059 | Gk | ENSMUST00000026039 | 30.7064108 | 35 | 17.9665169 | 16.0065333 | 0 | 0 | 0 | 0 | 0 | 0 | 0 | 0 | 0 | 0 | 0 | 0 | 0 | 1 | 1 |
| ENSMUSG00000041440 | Gk5 | ENSMUST00000122383 | 25.4596888 | 33 | 20.8274399 | 20.7567185 | 0 | 0 | 0 | 0 | 0 | 0 | 0 | 0 | 0 | 0 | 2 | 0 | 6 | 0 | 8 |
| ENSMUSG00000029638 | Glcci1 | ENSMUST00000064285 | 27.7467412 | 32 | 19.4832402 | 20.5772812 | 0 | 0 | 0 | 0 | 0 | 0 | 0 | 0 | 0 | 0 | 2 | 1 | 2 | 3 | 8 |
| ENSMUSG00000032252 | Glce | ENSMUST00000034785 | 26.2034021 | 32 | 19.9420919 | 22.2222222 | 0 | 0 | 2 | 0 | 0 | 0 | 0 | 0 | 0 | 2 | 0 | 0 | 2 | 0 | 4 |
| ENSMUSG00000028020 | Glrb | ENSMUST00000135043 | 30.0560748 | 32 | 18.8411215 | 19.364486 | 1 | 0 | 0 | 0 | 0 | 0 | 0 | 0 | 0 | 0 | 0 | 1 | 0 | 0 | 2 |
| ENSMUSG00000018196 | Glrx2 | ENSMUST00000129653 | 26.4734936 | 32 | 20.7770826 | 21.0734277 | 0 | 0 | 0 | 0 | 0 | 0 | 0 | 0 | 0 | 0 | 0 | 1 | 6 | 1 | 8 |
| ENSMUSG00000026103 | Gls | ENSMUST00000114510 | 31.0948081 | 38 | 13.6851016 | 16.8171558 | 1 | 1 | 0 | 3 | 0 | 0 | 0 | 0 | 3 | 6 | 2 | 0 | 8 | 4 | 22 |
| ENSMUSG00000011884 | Gltp | ENSMUST00000012028 | 23.1912785 | 28 | 28.444004 | 20.0198216 | 0 | 0 | 0 | 0 | 0 | 0 | 0 | 0 | 0 | 0 | 0 | 14 | 0 | 0 | 14 |
| ENSMUSG00000038965 | Gm10705; Ube2l3 | ENSMUST00000115699 | 24.6808511 | 28 | 25.8156028 | 21.7021277 | 0 | 0 | 0 | 0 | 0 | 0 | 0 | 0 | 0 | 0 | 0 | 3 | 0 | 0 | 3 |
| ENSMUSG00000074826 | Gm10767 | ENSMUST00000168767 | 29.7959184 | 36 | 14.7959184 | 19.5918367 | 0 | 0 | 0 | 0 | 0 | 0 | 0 | 0 | 0 | 0 | 1 | 0 | 1 | 0 | 2 |
| ENSMUSG00000096856 | Gm10778 | ENSMUST00000189672 | 30.8945687 | 30 | 18.6261981 | 20.4153355 | 2 | 1 | 0 | 1 | 0 | 0 | 0 | 0 | 1 | 2 | 0 | 0 | 1 | 0 | 6 |
| ENSMUSG00000029918 | Gm12540; Mrps33 | ENSMUST00000031978 | 30.3937008 | 33 | 17.3228346 | 19.3700787 | 0 | 0 | 0 | 0 | 0 | 0 | 0 | 0 | 0 | 0 | 0 | 1 | 0 | 0 | 1 |
| ENSMUSG00000071708 | Gm14680; Sms | ENSMUST00000112529 | 29.5354952 | 32 | 17.572305 | 20.9465381 | 0 | 0 | 0 | 1 | 0 | 0 | 0 | 0 | 1 | 2 | 0 | 2 | 0 | 0 | 4 |
| ENSMUSG00000093536 | Gm16532 | ENSMUST00000160218 | 27.1550129 | 34 | 19.6078431 | 19.4598594 | 3 | 2 | 0 | 6 | 0 | 0 | 0 | 2 | 4 | 12 | 1 | 0 | 2 | 0 | 20 |
| ENSMUSG00000066551 | Gm21596; Hmgb1 | ENSMUST00000085546 | 27.9115479 | 35 | 17.1007371 | 19.5085995 | 0 | 0 | 0 | 0 | 0 | 0 | 0 | 0 | 0 | 0 | 0 | 0 | 5 | 0 | 5 |
| ENSMUSG00000102422 | Gm21949 | ENSMUST00000182006 | 27.1381579 | 38 | 17.4342105 | 17.7631579 | 0 | 0 | 0 | 0 | 0 | 0 | 0 | 0 | 0 | 0 | 0 | 0 | 8 | 0 | 8 |
| ENSMUSG00000070354 | Gm21975 | ENSMUST00000093983 | 30.9599308 | 28 | 23.5514558 | 17.0942635 | 1 | 1 | 0 | 0 | 0 | 0 | 0 | 0 | 0 | 0 | 0 | 1 | 0 | 0 | 3 |
| ENSMUSG00000022451 | Gm4887; Twf1 | ENSMUST00000023087 | 28.2504013 | 31 | 19.3686463 | 21.1342964 | 0 | 0 | 0 | 1 | 0 | 0 | 0 | 0 | 1 | 2 | 0 | 0 | 14 | 0 | 16 |
| ENSMUSG00000058174 | Gm5148 | ENSMUST00000079755 | 28.6746988 | 34 | 20.2409639 | 16.626506 | 0 | 0 | 0 | 0 | 0 | 0 | 0 | 0 | 0 | 0 | 5 | 0 | 4 | 0 | 9 |
| ENSMUSG00000074754 | Gm561 | ENSMUST00000136628 | 33.6363636 | 37 | 13.6363636 | 15.4545455 | 0 | 0 | 3 | 2 | 0 | 0 | 0 | 0 | 2 | 7 | 0 | 0 | 0 | 0 | 7 |
| ENSMUSG00000001157 | Gmcl1 | ENSMUST00000001185 | 26.9264069 | 38 | 16.5367965 | 18.7878788 | 1 | 0 | 0 | 0 | 0 | 0 | 0 | 0 | 0 | 0 | 0 | 0 | 2 | 0 | 3 |
| ENSMUSG00000062014 | Gmfb | ENSMUST00000079314 | 28.698307 | 34 | 17.5964474 | 20.1498751 | 1 | 0 | 0 | 0 | 0 | 0 | 0 | 0 | 0 | 0 | 0 | 0 | 0 | 0 | 1 |
| ENSMUSG00000027823 | Gmps | ENSMUST00000029405 | 29.2557742 | 34 | 17.673225 | 19.3156544 | 0 | 0 | 0 | 0 | 0 | 0 | 0 | 0 | 0 | 0 | 0 | 1 | 19 | 1 | 21 |
| ENSMUSG00000020611 | Gna13 | ENSMUST00000020930 | 24.73889 | 31 | 21.8308417 | 22.0765923 | 0 | 0 | 0 | 0 | 0 | 0 | 0 | 0 | 0 | 0 | 10 | 0 | 13 | 3 | 26 |
| ENSMUSG00000057614 | Gnai1 | ENSMUST00000074694 | 29.0870488 | 36 | 16.1358811 | 18.9490446 | 1 | 2 | 0 | 1 | 0 | 0 | 0 | 0 | 1 | 2 | 0 | 0 | 0 | 0 | 5 |
| ENSMUSG00000000001 | Gnai3 | ENSMUST00000000001 | 26.7996109 | 35 | 17.8988327 | 20.7684825 | 0 | 0 | 0 | 0 | 0 | 0 | 0 | 0 | 0 | 0 | 1 | 1 | 0 | 0 | 2 |
| ENSMUSG00000024639 | Gnaq | ENSMUST00000025541 | 27.2057016 | 31 | 21.110838 | 20.8405014 | 1 | 0 | 1 | 1 | 0 | 0 | 0 | 0 | 1 | 3 | 0 | 27 | 4 | 0 | 35 |
| ENSMUSG00000027669 | Gnb4 | ENSMUST00000155737 | 28.4252297 | 29 | 20.9482038 | 21.7627402 | 0 | 0 | 0 | 0 | 0 | 0 | 0 | 0 | 0 | 0 | 0 | 0 | 0 | 1 | 1 |
| ENSMUSG00000036402 | Gng12 | ENSMUST00000043148 | 28.396703 | 31 | 21.4304706 | 19.5426748 | 0 | 0 | 0 | 0 | 0 | 0 | 0 | 0 | 0 | 0 | 0 | 0 | 0 | 1 | 1 |
| ENSMUSG00000043004 | Gng2 | ENSMUST00000160013 | 28.3559113 | 31 | 20.5665025 | 20.320197 | 0 | 0 | 0 | 0 | 0 | 0 | 0 | 0 | 0 | 0 | 1 | 0 | 3 | 0 | 4 |
| ENSMUSG00000025266 | Gnl3l | ENSMUST00000026297 | 26.560918 | 31 | 21.363483 | 20.8909889 | 3 | 1 | 0 | 8 | 0 | 0 | 1 | 2 | 5 | 16 | 0 | 0 | 9 | 0 | 29 |
| ENSMUSG00000052102 | Gnpda1 | ENSMUST00000063814 | 21.0646388 | 32 | 25.3231939 | 22.0532319 | 0 | 0 | 0 | 1 | 0 | 0 | 0 | 0 | 1 | 2 | 0 | 1 | 0 | 0 | 3 |
| ENSMUSG00000035521 | Gnptg | ENSMUST00000115154 | 32.6878389 | 26 | 23.5476375 | 18.2029435 | 0 | 0 | 0 | 0 | 0 | 0 | 0 | 0 | 0 | 0 | 0 | 2 | 0 | 0 | 2 |
| ENSMUSG00000015341 | Golga7 | ENSMUST00000051094 | 24.6522411 | 33 | 21.7156105 | 20.4018547 | 0 | 0 | 0 | 0 | 0 | 0 | 0 | 0 | 0 | 0 | 0 | 0 | 1 | 0 | 1 |
| ENSMUSG00000034109 | Golim4 | ENSMUST00000038563 | 28.4696301 | 34 | 19.6131659 | 17.61113 | 0 | 0 | 0 | 0 | 0 | 0 | 0 | 0 | 0 | 0 | 0 | 0 | 1 | 0 | 1 |
| ENSMUSG00000022200 | Golph3 | ENSMUST00000059680 | 25.8655804 | 38 | 19.7556008 | 16.700611 | 0 | 0 | 0 | 0 | 0 | 0 | 0 | 0 | 0 | 0 | 0 | 0 | 2 | 0 | 2 |
| ENSMUSG00000046519 | Golph3l | ENSMUST00000060323 | 27.8153153 | 31 | 22.1283784 | 18.6936937 | 0 | 0 | 0 | 0 | 0 | 0 | 0 | 0 | 0 | 0 | 0 | 1 | 0 | 0 | 1 |
| ENSMUSG00000030245 | Golt1b | ENSMUST00000032372 | 25.8234519 | 32 | 21.3438735 | 21.0364515 | 1 | 1 | 0 | 1 | 0 | 0 | 0 | 0 | 1 | 2 | 0 | 0 | 13 | 3 | 20 |
| ENSMUSG00000019861 | Gopc | ENSMUST00000020008 | 29.4553538 | 34 | 16.8210448 | 20.1556132 | 0 | 0 | 0 | 0 | 0 | 0 | 0 | 0 | 0 | 0 | 0 | 0 | 0 | 2 | 2 |
| ENSMUSG00000010392 | Gosr1 | ENSMUST00000010536 | 23.6954662 | 30 | 23.0681494 | 22.897063 | 1 | 1 | 0 | 3 | 0 | 0 | 0 | 1 | 2 | 6 | 0 | 0 | 2 | 1 | 11 |
| ENSMUSG00000022008 | Gpalpp1 | ENSMUST00000022585 | 27.6536313 | 31 | 20.1350093 | 21.5549348 | 2 | 2 | 0 | 0 | 0 | 0 | 0 | 0 | 0 | 0 | 1 | 0 | 2 | 0 | 7 |
| ENSMUSG00000039210 | Gpatch2 | ENSMUST00000110943 | 27.3744803 | 29 | 22.3536936 | 21.3943076 | 0 | 0 | 1 | 0 | 0 | 0 | 0 | 0 | 0 | 1 | 0 | 3 | 3 | 1 | 8 |
| ENSMUSG00000034621 | Gpatch8 | ENSMUST00000143842 | 24.3195786 | 32 | 21.7295874 | 21.9490781 | 0 | 0 | 0 | 1 | 0 | 0 | 0 | 0 | 1 | 2 | 0 | 2 | 4 | 0 | 8 |
| ENSMUSG00000034042 | Gpbp1l1 | ENSMUST00000030460 | 28.030303 | 38 | 14.8268398 | 19.1558442 | 0 | 0 | 0 | 0 | 0 | 0 | 0 | 0 | 0 | 0 | 0 | 0 | 14 | 0 | 14 |
| ENSMUSG00000027346 | Gpcpd1 | ENSMUST00000149854 | 30.719587 | 31 | 18.2962246 | 19.8773798 | 0 | 1 | 0 | 0 | 0 | 0 | 0 | 0 | 0 | 0 | 0 | 0 | 0 | 0 | 1 |
| ENSMUSG00000031148 | Gpkow | ENSMUST00000049896 | 29.4992563 | 30 | 20.0793257 | 20.2776401 | 0 | 0 | 0 | 0 | 0 | 0 | 0 | 0 | 0 | 0 | 0 | 0 | 0 | 1 | 1 |
| ENSMUSG00000031342 | Gpm6b | ENSMUST00000112235 | 30.7648564 | 33 | 17.8845607 | 18.1973273 | 0 | 0 | 0 | 0 | 0 | 0 | 0 | 0 | 0 | 0 | 0 | 0 | 1 | 0 | 1 |
| ENSMUSG00000029464 | Gpn3 | ENSMUST00000127608 | 26.9273743 | 27 | 21.6759777 | 24.2458101 | 0 | 0 | 0 | 0 | 0 | 0 | 0 | 0 | 0 | 0 | 1 | 1 | 0 | 0 | 2 |
| ENSMUSG00000036357 | Gpr101 | ENSMUST00000057645 | 31.3807531 | 30 | 18.6610879 | 20.027894 | 1 | 0 | 0 | 1 | 0 | 0 | 0 | 0 | 1 | 2 | 1 | 6 | 1 | 2 | 13 |
| ENSMUSG00000021306 | Gpr137b | ENSMUST00000021738 | 25.7548845 | 29 | 23.2089994 | 22.0248668 | 0 | 0 | 0 | 0 | 0 | 0 | 0 | 0 | 0 | 0 | 0 | 1 | 0 | 0 | 1 |
| ENSMUSG00000049092 | Gpr137c | ENSMUST00000146150 | 32 | 31 | 18.3478261 | 19.1304348 | 0 | 0 | 0 | 0 | 0 | 0 | 0 | 0 | 0 | 0 | 0 | 0 | 2 | 1 | 3 |
| ENSMUSG00000043441 | Gpr149 | ENSMUST00000058535 | 31.8711446 | 33 | 17.3406443 | 17.6148047 | 0 | 0 | 0 | 0 | 0 | 0 | 0 | 0 | 0 | 0 | 0 | 0 | 0 | 2 | 2 |
| ENSMUSG00000041762 | Gpr155 | ENSMUST00000076463 | 30.9072271 | 32 | 18.7596105 | 18.2470528 | 1 | 0 | 0 | 1 | 0 | 0 | 0 | 0 | 1 | 2 | 0 | 0 | 1 | 0 | 4 |
| ENSMUSG00000056679 | Gpr173 | ENSMUST00000070316 | 26.5965167 | 29 | 23.1857765 | 21.6618287 | 2 | 1 | 0 | 0 | 0 | 0 | 0 | 0 | 0 | 0 | 0 | 17 | 0 | 0 | 20 |
| ENSMUSG00000073008 | Gpr174 | ENSMUST00000178838 | 33.1025802 | 34 | 16.0478288 | 16.9288861 | 0 | 0 | 0 | 1 | 0 | 0 | 0 | 0 | 1 | 2 | 1 | 0 | 0 | 0 | 3 |
| ENSMUSG00000022131 | Gpr180 | ENSMUST00000022728 | 28.1481481 | 32 | 16.2289562 | 23.6363636 | 0 | 0 | 0 | 0 | 0 | 0 | 0 | 0 | 0 | 0 | 4 | 0 | 0 | 0 | 4 |
| ENSMUSG00000044067 | Gpr22 | ENSMUST00000057783 | 37.5609756 | 33 | 15.2845528 | 13.8211382 | 2 | 0 | 0 | 2 | 0 | 0 | 0 | 0 | 2 | 4 | 0 | 0 | 0 | 0 | 6 |
| ENSMUSG00000041907 | Gpr45 | ENSMUST00000179766 | 25.3277154 | 27 | 24.1573034 | 23.6423221 | 0 | 0 | 0 | 0 | 0 | 0 | 0 | 0 | 0 | 0 | 0 | 4 | 0 | 0 | 4 |
| ENSMUSG00000040372 | Gpr63 | ENSMUST00000038920 | 24.6312684 | 38 | 13.8643068 | 23.5988201 | 0 | 0 | 0 | 1 | 0 | 0 | 0 | 0 | 1 | 2 | 0 | 1 | 2 | 0 | 5 |
| ENSMUSG00000048216 | Gpr85 | ENSMUST00000060442 | 30.6076519 | 33 | 16.204051 | 20.5551388 | 0 | 0 | 0 | 0 | 0 | 0 | 0 | 0 | 0 | 0 | 0 | 1 | 0 | 0 | 1 |
| ENSMUSG00000028096 | Gpr89 | ENSMUST00000029738 | 26.9069842 | 25 | 24.5574509 | 23.4953331 | 0 | 0 | 0 | 0 | 0 | 0 | 0 | 0 | 0 | 0 | 0 | 2 | 0 | 1 | 3 |
| ENSMUSG00000036292 | Gramd1c | ENSMUST00000179565 | 29.5667447 | 32 | 17.3887588 | 20.9601874 | 1 | 0 | 0 | 1 | 0 | 0 | 0 | 0 | 1 | 2 | 0 | 0 | 1 | 0 | 4 |
| ENSMUSG00000059923 | Grb2 | ENSMUST00000021090 | 22.5058005 | 29 | 24.9419954 | 23.7819026 | 0 | 0 | 0 | 0 | 0 | 0 | 0 | 0 | 0 | 0 | 4 | 2 | 4 | 3 | 13 |
| ENSMUSG00000042942 | Greb1l | ENSMUST00000048977 | 30.0693594 | 32 | 18.7270502 | 19.5430437 | 0 | 0 | 0 | 0 | 0 | 0 | 0 | 0 | 0 | 0 | 0 | 1 | 0 | 0 | 1 |
| ENSMUSG00000074934 | Grem1 | ENSMUST00000099575 | 31.0535506 | 27 | 18.975553 | 22.7881257 | 0 | 0 | 0 | 0 | 0 | 0 | 0 | 0 | 0 | 0 | 0 | 0 | 0 | 2 | 2 |
| ENSMUSG00000050069 | Grem2 | ENSMUST00000055294 | 28.5284281 | 27 | 21.4715719 | 22.5752508 | 1 | 0 | 1 | 1 | 0 | 0 | 0 | 0 | 1 | 3 | 0 | 1 | 0 | 0 | 5 |
| ENSMUSG00000033981 | Gria2 | ENSMUST00000075316 | 32.2647221 | 35 | 16.758393 | 16.1942763 | 3 | 0 | 0 | 3 | 0 | 0 | 0 | 0 | 3 | 6 | 0 | 1 | 1 | 1 | 12 |
| ENSMUSG00000001986 | Gria3 | ENSMUST00000139965 | 30.8584687 | 29 | 19.8762568 | 20.0309358 | 0 | 0 | 0 | 0 | 0 | 0 | 0 | 0 | 0 | 0 | 0 | 0 | 2 | 0 | 2 |
| ENSMUSG00000025892 | Gria4 | ENSMUST00000027020 | 30.2707501 | 30 | 20.5503773 | 19.3519751 | 0 | 0 | 0 | 1 | 0 | 0 | 0 | 0 | 1 | 2 | 0 | 2 | 0 | 0 | 4 |
| ENSMUSG00000056073 | Grik2 | ENSMUST00000105487 | 32.9490291 | 30 | 17.9004854 | 18.8713592 | 0 | 0 | 0 | 0 | 0 | 0 | 0 | 0 | 0 | 0 | 0 | 1 | 3 | 3 | 7 |
| ENSMUSG00000049583 | Grm5 | ENSMUST00000125009 | 32.4170381 | 33 | 17.0381377 | 17.5581971 | 0 | 0 | 0 | 0 | 0 | 0 | 0 | 0 | 0 | 0 | 3 | 1 | 4 | 1 | 9 |
| ENSMUSG00000039934 | Gsap | ENSMUST00000198014 | 25.8146067 | 29 | 22.4438202 | 23.1460674 | 0 | 0 | 0 | 0 | 0 | 0 | 0 | 0 | 0 | 0 | 0 | 0 | 2 | 0 | 2 |
| ENSMUSG00000022812 | Gsk3b | ENSMUST00000023507 | 26.3281958 | 32 | 20.0543971 | 21.7044424 | 2 | 1 | 0 | 2 | 0 | 0 | 0 | 1 | 1 | 4 | 0 | 7 | 25 | 5 | 44 |
| ENSMUSG00000031584 | Gsr | ENSMUST00000033992 | 23.156682 | 39 | 17.5115207 | 20.6221198 | 0 | 0 | 1 | 1 | 0 | 0 | 0 | 0 | 1 | 3 | 0 | 0 | 9 | 0 | 12 |
| ENSMUSG00000035946 | Gsx2 | ENSMUST00000160104 | 25.2427184 | 24 | 25.2427184 | 25.0809061 | 0 | 0 | 0 | 0 | 0 | 0 | 0 | 0 | 0 | 0 | 0 | 1 | 2 | 0 | 3 |
| ENSMUSG00000020962 | Gtf2a1 | ENSMUST00000021345 | 26.5673981 | 34 | 18.2601881 | 21.2121212 | 1 | 0 | 0 | 1 | 0 | 0 | 0 | 0 | 1 | 2 | 0 | 0 | 10 | 2 | 15 |
| ENSMUSG00000033543 | Gtf2a2 | ENSMUST00000140265 | 30.9701493 | 34 | 15.858209 | 18.8432836 | 1 | 0 | 0 | 0 | 0 | 0 | 0 | 0 | 0 | 0 | 0 | 0 | 0 | 0 | 1 |
| ENSMUSG00000022828 | Gtf2e1 | ENSMUST00000023525 | 27.1419637 | 32 | 19.6372733 | 21.0131332 | 0 | 0 | 0 | 0 | 0 | 0 | 0 | 0 | 0 | 0 | 0 | 0 | 0 | 2 | 2 |
| ENSMUSG00000031585 | Gtf2e2 | ENSMUST00000171010 | 26.6556291 | 33 | 16.5562914 | 23.3443709 | 0 | 0 | 0 | 1 | 0 | 0 | 0 | 0 | 1 | 2 | 1 | 0 | 2 | 0 | 5 |
| ENSMUSG00000041303 | Gtf3c3 | ENSMUST00000041638 | 28.891258 | 34 | 17.5906183 | 19.45629 | 2 | 0 | 0 | 0 | 0 | 0 | 0 | 0 | 0 | 0 | 1 | 0 | 17 | 0 | 20 |
| ENSMUSG00000040464 | Gtpbp10 | ENSMUST00000115441 | 30.5895439 | 34 | 16.5183537 | 19.1323693 | 0 | 0 | 0 | 1 | 0 | 0 | 0 | 0 | 1 | 2 | 0 | 0 | 0 | 0 | 2 |
| ENSMUSG00000021149 | Gtpbp4 | ENSMUST00000021574 | 30.0666032 | 32 | 16.1750714 | 21.5984776 | 0 | 0 | 0 | 0 | 0 | 0 | 0 | 0 | 0 | 0 | 0 | 0 | 1 | 1 | 2 |

| | | | | | | | | | | | | | | | | | | | | | |
|---|---|---|---|---|---|---|---|---|---|---|---|---|---|---|---|---|---|---|---|---|---|
| ENSMUSG00000022668 | Gtpbp8 | ENSMUST00000162512 | 29.1979455 | 27 | 21.9280917 | 21.8490715 | 0 | 0 | 1 | 0 | 0 | 0 | 0 | 0 | 0 | 1 | 0 | 0 | 1 | 0 | 2 |
| ENSMUSG00000041624 | Gucy1a2 | ENSMUST00000115733 | 32.9532497 | 34 | 17.1037628 | 16.381604 | 0 | 0 | 0 | 1 | 0 | 0 | 0 | 0 | 1 | 2 | 1 | 0 | 0 | 0 | 3 |
| ENSMUSG00000028005 | Gucy1b3 | ENSMUST00000029635 | 28.102482 | 36 | 16.4931946 | 19.5356285 | 0 | 0 | 0 | 1 | 0 | 0 | 0 | 0 | 1 | 2 | 0 | 0 | 0 | 0 | 2 |
| ENSMUSG00000029208 | Guf1 | ENSMUST00000154728 | 30.2926829 | 29 | 18.2926829 | 22.3902439 | 0 | 0 | 0 | 0 | 0 | 0 | 0 | 0 | 0 | 0 | 1 | 1 | 0 | 0 | 2 |
| ENSMUSG00000056870 | Gulp1 | ENSMUST00000159555 | 32.7409163 | 35 | 15.9952607 | 16.2322275 | 0 | 0 | 0 | 1 | 0 | 0 | 0 | 0 | 1 | 2 | 0 | 0 | 0 | 0 | 2 |
| ENSMUSG00000036197 | Gxylt1 | ENSMUST00000049484 | 27.9748707 | 29 | 18.5513673 | 24.6489283 | 0 | 0 | 0 | 0 | 0 | 0 | 0 | 0 | 0 | 0 | 3 | 0 | 1 | 0 | 4 |
| ENSMUSG00000041126 | H2afv | ENSMUST00000109737 | 28.1085814 | 31 | 19.614711 | 21.5411559 | 1 | 0 | 0 | 1 | 0 | 0 | 0 | 0 | 1 | 2 | 0 | 0 | 0 | 0 | 3 |
| ENSMUSG00000015937 | H2afy | ENSMUST00000016081 | 26.8092105 | 31 | 22.3684211 | 19.9013158 | 0 | 0 | 0 | 0 | 0 | 0 | 0 | 0 | 0 | 0 | 0 | 0 | 3 | 0 | 3 |
| ENSMUSG00000035376 | Hacd2 | ENSMUST00000061156 | 27.0698344 | 29 | 22.4262059 | 21.0223182 | 0 | 0 | 0 | 0 | 0 | 0 | 0 | 0 | 0 | 0 | 0 | 0 | 1 | 0 | 1 |
| ENSMUSG00000033629 | Hacd3 | ENSMUST00000036615 | 25.8174387 | 32 | 20.7765668 | 21.5940054 | 0 | 0 | 0 | 0 | 0 | 0 | 0 | 0 | 0 | 0 | 0 | 1 | 3 | 1 | 5 |
| ENSMUSG00000038822 | Hace1 | ENSMUST00000037044 | 29.9134734 | 39 | 14.5859085 | 16.5636588 | 1 | 0 | 0 | 1 | 0 | 0 | 0 | 0 | 1 | 2 | 0 | 0 | 0 | 0 | 3 |
| ENSMUSG00000021613 | Hapln1 | ENSMUST00000022108 | 34.1569373 | 33 | 16.5354331 | 16.3182188 | 0 | 0 | 0 | 1 | 0 | 0 | 0 | 0 | 1 | 2 | 3 | 0 | 8 | 0 | 13 |
| ENSMUSG00000019143 | Hars2 | ENSMUST00000152954 | 29.65806 | 28 | 19.1905094 | 22.9588276 | 0 | 0 | 0 | 0 | 0 | 0 | 0 | 0 | 0 | 0 | 0 | 1 | 0 | 0 | 1 |
| ENSMUSG00000022367 | Has2 | ENSMUST00000050544 | 31.25 | 32 | 18.4547244 | 18.1594488 | 1 | 0 | 0 | 0 | 0 | 0 | 0 | 0 | 0 | 0 | 0 | 2 | 0 | 0 | 3 |
| ENSMUSG00000027285 | Haus2 | ENSMUST00000124187 | 29.087852 | 31 | 19.209752 | 21.185372 | 0 | 0 | 0 | 0 | 0 | 0 | 0 | 0 | 0 | 0 | 0 | 1 | 0 | 0 | 1 |
| ENSMUSG00000038047 | Haus6 | ENSMUST00000070607 | 28.3236994 | 32 | 17.8869621 | 21.3551702 | 2 | 0 | 0 | 2 | 0 | 0 | 0 | 0 | 2 | 4 | 0 | 1 | 3 | 0 | 10 |
| ENSMUSG00000024486 | Hbegf | ENSMUST00000025363 | 25.65337 | 30 | 20.9078404 | 23.4525447 | 2 | 0 | 0 | 1 | 0 | 0 | 0 | 0 | 1 | 2 | 3 | 0 | 3 | 0 | 10 |
| ENSMUSG00000002996 | Hbp1 | ENSMUST00000167458 | 28.5714286 | 35 | 17.5289575 | 18.996139 | 0 | 0 | 0 | 0 | 0 | 0 | 0 | 0 | 0 | 0 | 0 | 0 | 1 | 0 | 1 |
| ENSMUSG00000019977 | Hbs1l | ENSMUST00000092674 | 29.8969072 | 41 | 12.371134 | 16.9366716 | 0 | 0 | 0 | 0 | 0 | 0 | 0 | 0 | 0 | 0 | 0 | 0 | 0 | 2 | 2 |
| ENSMUSG00000031352 | Hccs | ENSMUST00000154638 | 31.1939763 | 33 | 17.3180351 | 18.2144138 | 0 | 0 | 0 | 0 | 0 | 0 | 0 | 0 | 0 | 0 | 0 | 0 | 10 | 0 | 10 |
| ENSMUSG00000032360 | Hcrtr2 | ENSMUST00000063140 | 33.8177015 | 34 | 14.8392779 | 17.7014531 | 1 | 1 | 0 | 4 | 0 | 0 | 0 | 1 | 3 | 8 | 0 | 0 | 0 | 0 | 10 |
| ENSMUSG00000019777 | Hdac2 | ENSMUST00000105510 | 25.9637715 | 36 | 18.8109614 | 19.4612169 | 0 | 0 | 0 | 0 | 0 | 0 | 0 | 0 | 0 | 0 | 1 | 0 | 2 | 0 | 3 |
| ENSMUSG00000025104 | Hdgfrp3 | ENSMUST00000107305 | 31.075202 | 32 | 17.6299979 | 19.49451 | 1 | 0 | 0 | 0 | 0 | 0 | 0 | 0 | 0 | 0 | 7 | 1 | 4 | 2 | 15 |
| ENSMUSG00000025421 | Hdhd2 | ENSMUST00000148955 | 26.4625229 | 27 | 21.7550274 | 24.9085923 | 0 | 0 | 0 | 1 | 0 | 0 | 0 | 0 | 1 | 2 | 0 | 0 | 0 | 0 | 2 |
| ENSMUSG00000034551 | Hdx | ENSMUST00000113422 | 28.8613457 | 34 | 19.8174884 | 16.9980932 | 0 | 0 | 0 | 0 | 0 | 0 | 0 | 0 | 0 | 0 | 4 | 3 | 5 | 2 | 14 |
| ENSMUSG00000031657 | Heatr3 | ENSMUST00000034079 | 30.3059273 | 29 | 19.4072658 | 21.2237094 | 0 | 0 | 0 | 0 | 0 | 0 | 0 | 0 | 0 | 0 | 0 | 0 | 0 | 2 | 2 |
| ENSMUSG00000041180 | Hectd2 | ENSMUST00000139215 | 30.9575234 | 33 | 17.5905928 | 18.3585313 | 2 | 0 | 0 | 1 | 0 | 0 | 0 | 0 | 1 | 2 | 0 | 0 | 5 | 0 | 9 |
| ENSMUSG00000025001 | Hells | ENSMUST00000025965 | 27.7915633 | 37 | 16.9354839 | 18.3933002 | 2 | 2 | 0 | 5 | 0 | 0 | 0 | 2 | 3 | 10 | 1 | 0 | 1 | 0 | 16 |
| ENSMUSG00000020721 | Helz | ENSMUST00000106746 | 25.4397073 | 32 | 20.9933868 | 21.5280709 | 2 | 0 | 0 | 2 | 0 | 0 | 0 | 0 | 2 | 4 | 10 | 9 | 17 | 3 | 45 |
| ENSMUSG00000029804 | Herc3 | ENSMUST00000041401 | 24.5091401 | 37 | 19.7698037 | 18.7542316 | 2 | 1 | 0 | 0 | 0 | 0 | 0 | 0 | 0 | 0 | 0 | 0 | 3 | 2 | 8 |
| ENSMUSG00000008429 | Herpud2 | ENSMUST00000008573 | 33.7535014 | 35 | 15.5462185 | 15.9663866 | 0 | 0 | 0 | 0 | 0 | 0 | 0 | 0 | 0 | 0 | 1 | 0 | 0 | 0 | 1 |
| ENSMUSG00000040289 | Hey1 | ENSMUST00000042412 | 25.4177661 | 34 | 18.2058047 | 22.4274406 | 0 | 0 | 0 | 0 | 0 | 0 | 0 | 0 | 0 | 0 | 0 | 2 | 0 | 1 | 3 |
| ENSMUSG00000019789 | Hey2 | ENSMUST00000019924 | 31.372549 | 32 | 17.7923021 | 18.9542484 | 0 | 0 | 0 | 0 | 0 | 0 | 0 | 0 | 0 | 0 | 0 | 0 | 2 | 0 | 2 |
| ENSMUSG00000028864 | Hgf | ENSMUST00000199581 | 33.8222606 | 37 | 14.2364107 | 14.9266609 | 1 | 0 | 0 | 2 | 0 | 0 | 0 | 0 | 2 | 4 | 0 | 0 | 0 | 0 | 5 |
| ENSMUSG00000024986 | Hhex | ENSMUST00000025944 | 30.3455724 | 32 | 16.0907127 | 21.1663067 | 0 | 0 | 0 | 0 | 0 | 0 | 0 | 0 | 0 | 0 | 2 | 0 | 3 | 0 | 5 |
| ENSMUSG00000064325 | Hhip | ENSMUST00000079038 | 27.665996 | 34 | 19.6177062 | 18.360161 | 0 | 0 | 0 | 0 | 0 | 0 | 0 | 0 | 0 | 0 | 0 | 0 | 3 | 0 | 3 |
| ENSMUSG00000089911 | Hiat1 | ENSMUST00000029570 | 31.0810811 | 35 | 17.2072072 | 16.3063063 | 0 | 0 | 1 | 1 | 0 | 0 | 0 | 0 | 1 | 3 | 0 | 1 | 0 | 2 | 6 |
| ENSMUSG00000038212 | Hiatl1 | ENSMUST00000155487 | 29.7948718 | 36 | 16.9230769 | 17.3333333 | 0 | 0 | 0 | 0 | 0 | 0 | 0 | 0 | 0 | 0 | 1 | 0 | 0 | 1 | 2 |
| ENSMUSG00000050240 | Hic2 | ENSMUST00000090190 | 22.618489 | 26 | 25.8329423 | 25.786016 | 0 | 0 | 0 | 0 | 0 | 0 | 0 | 0 | 0 | 0 | 0 | 0 | 0 | 1 | 1 |
| ENSMUSG00000038412 | Higd1a | ENSMUST00000060251 | 28.2122905 | 31 | 16.6666667 | 24.5810056 | 0 | 0 | 0 | 0 | 0 | 0 | 0 | 0 | 0 | 0 | 0 | 0 | 1 | 0 | 1 |
| ENSMUSG00000025868 | Higd2a | ENSMUST00000026986 | 23.880597 | 29 | 26.4925373 | 20.1492537 | 0 | 0 | 0 | 0 | 0 | 0 | 0 | 0 | 0 | 0 | 0 | 1 | 0 | 0 | 1 |
| ENSMUSG00000019791 | Hint3 | ENSMUST00000161074 | 23.6 | 43 | 18 | 15.2 | 0 | 0 | 0 | 0 | 0 | 0 | 0 | 0 | 0 | 0 | 1 | 0 | 1 | 1 | 3 |
| ENSMUSG00000008730 | Hipk1 | ENSMUST00000118317 | 25.5988386 | 33 | 18.8240987 | 22.3566417 | 0 | 0 | 0 | 0 | 0 | 0 | 0 | 0 | 0 | 0 | 3 | 4 | 5 | 2 | 14 |
| ENSMUSG00000061436 | Hipk2 | ENSMUST00000161779 | 24.7263017 | 26 | 24.6728972 | 24.753004 | 1 | 0 | 0 | 2 | 0 | 0 | 0 | 0 | 2 | 4 | 0 | 3 | 0 | 2 | 10 |
| ENSMUSG00000068854 | Hist2h2be | ENSMUST00000090781 | 22.4452555 | 34 | 21.4872263 | 22.0346715 | 0 | 0 | 0 | 0 | 0 | 0 | 0 | 0 | 0 | 0 | 0 | 0 | 2 | 0 | 2 |
| ENSMUSG00000078851 | Hist3h2a | ENSMUST00000108817 | 27 | 30 | 18.4705882 | 24.0588235 | 0 | 1 | 0 | 1 | 0 | 0 | 0 | 0 | 1 | 2 | 0 | 0 | 0 | 0 | 3 |
| ENSMUSG00000003949 | Hlf | ENSMUST00000004051 | 27.4212645 | 31 | 21.0040256 | 20.4357092 | 0 | 0 | 0 | 0 | 0 | 0 | 0 | 0 | 0 | 0 | 4 | 2 | 0 | 0 | 6 |
| ENSMUSG00000054717 | Hmgb2 | ENSMUST00000067925 | 30.1816239 | 36 | 15.2777778 | 18.4294872 | 2 | 1 | 0 | 4 | 0 | 0 | 0 | 1 | 3 | 8 | 0 | 0 | 1 | 1 | 13 |
| ENSMUSG00000007908 | Hmgcll1 | ENSMUST00000008052 | 30.1810865 | 32 | 18.5110664 | 18.9134809 | 1 | 0 | 0 | 1 | 0 | 0 | 0 | 0 | 1 | 2 | 1 | 5 | 1 | 0 | 10 |
| ENSMUSG00000021670 | Hmgcr | ENSMUST00000022176 | 27.1091811 | 33 | 17.5558313 | 22.0223325 | 0 | 0 | 1 | 0 | 0 | 0 | 0 | 0 | 0 | 1 | 0 | 0 | 5 | 5 | 11 |
| ENSMUSG00000093930 | Hmgcs1 | ENSMUST00000179869 | 28.6069652 | 32 | 15.4228856 | 23.6318408 | 0 | 0 | 0 | 0 | 0 | 0 | 0 | 0 | 0 | 0 | 1 | 0 | 1 | 0 | 2 |
| ENSMUSG00000003038 | Hmgn2 | ENSMUST00000102553 | 27.25 | 34 | 18.75 | 20.375 | 0 | 0 | 0 | 0 | 0 | 0 | 0 | 0 | 0 | 0 | 0 | 0 | 0 | 1 | 1 |
| ENSMUSG00000066456 | Hmgn3 | ENSMUST00000185315 | 24.7544204 | 33 | 20.1375246 | 22.1021611 | 0 | 0 | 0 | 0 | 0 | 0 | 0 | 0 | 0 | 0 | 0 | 3 | 0 | 0 | 3 |
| ENSMUSG00000034518 | Hmgxb4 | ENSMUST00000145919 | 29.220415 | 25 | 21.9573752 | 23.3875491 | 0 | 0 | 0 | 1 | 0 | 0 | 0 | 0 | 1 | 2 | 0 | 0 | 4 | 1 | 7 |
| ENSMUSG00000005413 | Hmox1 | ENSMUST00000005548 | 22.0103986 | 32 | 24.7833622 | 21.6637782 | 0 | 0 | 0 | 0 | 0 | 0 | 0 | 0 | 0 | 0 | 0 | 0 | 3 | 0 | 3 |
| ENSMUSG00000024165 | Hn1l | ENSMUST00000024981 | 24.0685225 | 31 | 22.8265525 | 22.3554604 | 0 | 0 | 0 | 0 | 0 | 0 | 0 | 0 | 0 | 0 | 0 | 2 | 6 | 0 | 8 |
| ENSMUSG00000046434 | Hnrnpa1; Gm5803 | ENSMUST00000036004 | 30.3921569 | 32 | 16.3865546 | 21.2885154 | 0 | 1 | 0 | 0 | 0 | 0 | 0 | 0 | 0 | 0 | 0 | 0 | 5 | 2 | 8 |
| ENSMUSG00000060373 | Hnrnpc | ENSMUST00000111610 | 26.2941176 | 34 | 19.8823529 | 19.3529412 | 0 | 0 | 0 | 0 | 0 | 0 | 0 | 0 | 0 | 0 | 1 | 0 | 1 | 1 | 3 |
| ENSMUSG00000000568 | Hnrnpd | ENSMUST00000172361 | 27.1504881 | 34 | 17.3328421 | 21.0904402 | 0 | 0 | 0 | 1 | 0 | 0 | 0 | 0 | 1 | 2 | 2 | 1 | 6 | 4 | 15 |
| ENSMUSG00000007850 | Hnrnph1 | ENSMUST00000109142 | 28.856383 | 36 | 14.0957447 | 21.5425532 | 0 | 0 | 1 | 0 | 0 | 0 | 0 | 0 | 0 | 1 | 0 | 0 | 0 | 0 | 1 |
| ENSMUSG00000045427 | Hnrnph2 | ENSMUST00000113203 | 31.6313823 | 36 | 14.6948941 | 17.3100872 | 0 | 0 | 0 | 0 | 0 | 0 | 0 | 0 | 0 | 0 | 0 | 0 | 1 | 5 | 6 |
| ENSMUSG00000021546 | Hnrnpk | ENSMUST00000116403 | 28.6702537 | 36 | 14.3735588 | 20.5995388 | 0 | 0 | 0 | 0 | 0 | 0 | 0 | 0 | 0 | 0 | 1 | 0 | 1 | 2 | 4 |
| ENSMUSG00000024095 | Hnrnpll | ENSMUST00000184297 | 29.7816016 | 33 | 18.1336863 | 19.3249504 | 0 | 0 | 0 | 0 | 0 | 0 | 0 | 0 | 0 | 0 | 1 | 0 | 0 | 0 | 1 |
| ENSMUSG00000059208 | Hnrnpm | ENSMUST00000114385 | 23.943662 | 34 | 16.0965795 | 25.9557344 | 0 | 0 | 0 | 0 | 0 | 0 | 0 | 0 | 0 | 0 | 0 | 1 | 0 | 0 | 1 |
| ENSMUSG00000066037 | Hnrnpr | ENSMUST00000148843 | 28.6986972 | 31 | 18.6415366 | 21.5562078 | 1 | 0 | 0 | 1 | 0 | 0 | 0 | 0 | 1 | 2 | 3 | 3 | 32 | 4 | 45 |
| ENSMUSG00000039630 | Hnrnpu | ENSMUST00000037748 | 31.0915985 | 34 | 15.5457992 | 19.2576332 | 0 | 0 | 0 | 0 | 0 | 0 | 0 | 0 | 0 | 0 | 0 | 1 | 2 | 1 | 4 |
| ENSMUSG00000007617 | Homer1 | ENSMUST00000102752 | 30.7426995 | 36 | 15.7737411 | 17.7052196 | 1 | 0 | 1 | 1 | 0 | 0 | 0 | 0 | 1 | 3 | 0 | 1 | 0 | 3 | 8 |
| ENSMUSG00000028572 | Hook1 | ENSMUST00000030306 | 29.0637191 | 35 | 15.7022107 | 20.3836151 | 0 | 0 | 0 | 0 | 0 | 0 | 0 | 0 | 0 | 0 | 1 | 0 | 0 | 0 | 1 |
| ENSMUSG00000037234 | Hook3 | ENSMUST00000037182 | 29.3151208 | 33 | 17.9386761 | 19.5338619 | 2 | 2 | 0 | 2 | 0 | 0 | 0 | 0 | 2 | 4 | 8 | 3 | 9 | 0 | 28 |
| ENSMUSG00000059325 | Hopx | ENSMUST00000081964 | 29.6937417 | 33 | 17.8428762 | 19.7070573 | 1 | 0 | 0 | 1 | 0 | 0 | 0 | 0 | 1 | 2 | 0 | 1 | 1 | 0 | 5 |
| ENSMUSG00000028759 | Hp1bp3 | ENSMUST00000105825 | 25.7900102 | 31 | 20.8630649 | 22.7319062 | 0 | 0 | 0 | 0 | 0 | 0 | 0 | 0 | 0 | 0 | 1 | 1 | 0 | 0 | 2 |
| ENSMUSG00000031613 | Hpgd | ENSMUST00000034026 | 32.5802616 | 32 | 17.4791914 | 17.5980975 | 0 | 0 | 0 | 0 | 0 | 0 | 0 | 0 | 0 | 0 | 1 | 0 | 0 | 0 | 1 |
| ENSMUSG00000027615 | Hps3 | ENSMUST00000012580 | 27.2727273 | 37 | 18.1818182 | 17.1368861 | 2 | 1 | 0 | 3 | 0 | 0 | 0 | 1 | 2 | 6 | 1 | 1 | 1 | 0 | 12 |
| ENSMUSG00000022525 | Hrasls | ENSMUST00000089824 | 27.9930344 | 32 | 20.1131911 | 20.3744014 | 0 | 0 | 0 | 1 | 0 | 0 | 0 | 0 | 1 | 2 | 0 | 11 | 0 | 0 | 13 |
| ENSMUSG00000040151 | Hs2st1 | ENSMUST00000043325 | 25.7005437 | 32 | 21.0790464 | 21.371811 | 0 | 0 | 0 | 0 | 0 | 0 | 0 | 0 | 0 | 0 | 0 | 6 | 1 | 0 | 7 |
| ENSMUSG00000051022 | Hs3st1 | ENSMUST00000117944 | 27.8037383 | 41 | 14.4859813 | 16.588785 | 0 | 0 | 0 | 0 | 0 | 0 | 0 | 0 | 0 | 0 | 0 | 0 | 1 | 0 | 1 |
| ENSMUSG00000044499 | Hs3st5 | ENSMUST00000168572 | 31.6027088 | 34 | 16.3656885 | 17.6072235 | 0 | 0 | 0 | 0 | 0 | 0 | 0 | 0 | 0 | 0 | 3 | 0 | 5 | 0 | 8 |
| ENSMUSG00000026675 | Hsd17b7 | ENSMUST00000027989 | 24.0423099 | 35 | 20.9262436 | 20.3259005 | 5 | 1 | 0 | 0 | 0 | 0 | 0 | 0 | 0 | 0 | 0 | 1 | 3 | 0 | 10 |
| ENSMUSG00000028383 | Hsdl2 | ENSMUST00000030078 | 28.8747346 | 35 | 19.4267516 | 16.5605096 | 0 | 0 | 0 | 0 | 0 | 0 | 0 | 0 | 0 | 0 | 0 | 0 | 3 | 1 | 4 |
| ENSMUSG00000019878 | Hsf2 | ENSMUST00000079833 | 26.3101604 | 39 | 16.1497326 | 19.0374332 | 1 | 0 | 0 | 0 | 0 | 0 | 0 | 0 | 0 | 0 | 0 | 0 | 17 | 2 | 20 |
| ENSMUSG00000070345 | Hsf5 | ENSMUST00000093956 | 27.4022989 | 30 | 19.862069 | 22.4827586 | 1 | 0 | 0 | 2 | 0 | 0 | 0 | 0 | 2 | 4 | 0 | 0 | 5 | 1 | 11 |
| ENSMUSG00000032932 | Hspa13 | ENSMUST00000046283 | 30.9147047 | 34 | 16.2485527 | 18.8344269 | 0 | 0 | 0 | 0 | 0 | 0 | 0 | 0 | 0 | 0 | 2 | 0 | 4 | 2 | 8 |
| ENSMUSG00000051396 | Hspa14 | ENSMUST00000124331 | 25.932372 | 26 | 22.3023371 | 25.4599702 | 0 | 0 | 0 | 0 | 0 | 0 | 0 | 0 | 0 | 0 | 0 | 2 | 0 | 0 | 2 |
| ENSMUSG00000025757 | Hspa4l | ENSMUST00000204702 | 32.8120346 | 31 | 17.932678 | 18.5135538 | 2 | 1 | 0 | 0 | 0 | 0 | 0 | 0 | 0 | 0 | 0 | 0 | 2 | 3 | 8 |
| ENSMUSG00000063172 | Hspb11 | ENSMUST00000106749 | 38 | 40 | 13 | 9 | 1 | 0 | 0 | 0 | 0 | 0 | 0 | 0 | 0 | 0 | 0 | 0 | 0 | 0 | 1 |
| ENSMUSG00000022849 | Hspbap1 | ENSMUST00000023555 | 24.7703412 | 32 | 23.3595801 | 20.3412073 | 3 | 1 | 0 | 1 | 0 | 0 | 0 | 0 | 1 | 2 | 1 | 3 | 1 | 0 | 11 |
| ENSMUSG00000041380 | Htr2c | ENSMUST00000156697 | 29.6113689 | 31 | 19.6055684 | 19.5185615 | 0 | 0 | 0 | 1 | 0 | 0 | 0 | 0 | 1 | 2 | 1 | 0 | 1 | 0 | 4 |
| ENSMUSG00000026322 | Htr4 | ENSMUST00000027560 | 26.9066504 | 32 | 20.6528371 | 20.8968883 | 2 | 1 | 0 | 3 | 0 | 0 | 0 | 1 | 2 | 6 | 0 | 0 | 1 | 1 | 11 |
| ENSMUSG00000020413 | Hus1 | ENSMUST00000020683 | 25.8551037 | 31 | 22.2461621 | 21.3035281 | 0 | 0 | 0 | 1 | 0 | 0 | 0 | 0 | 1 | 2 | 0 | 0 | 1 | 0 | 3 |
| ENSMUSG00000032115 | Hyou1 | ENSMUST00000162560 | 25.8316008 | 21 | 24.8960499 | 28.7422037 | 0 | 0 | 0 | 0 | 0 | 0 | 0 | 0 | 0 | 0 | 0 | 1 | 0 | 0 | 1 |
| ENSMUSG00000027245 | Hypk | ENSMUST00000126764 | 26.8691589 | 37 | 18.2242991 | 17.5233645 | 0 | 0 | 0 | 0 | 0 | 0 | 0 | 0 | 0 | 0 | 0 | 0 | 1 | 0 | 1 |
| ENSMUSG00000009828 | Ick | ENSMUST00000044551 | 27.773949 | 29 | 21.6402481 | 21.2267402 | 1 | 0 | 0 | 1 | 0 | 0 | 0 | 0 | 1 | 2 | 1 | 0 | 0 | 4 | 8 |

| | | | | | | | | | | | | | | | | | | | | | |
|---|---|---|---|---|---|---|---|---|---|---|---|---|---|---|---|---|---|---|---|---|---|
| ENSMUSG00000042745 | Id1 | ENSMUST00000109824 | 23.0905861 | 29 | 23.0905861 | 25.044405 | 0 | 0 | 0 | 0 | 0 | 0 | 0 | 0 | 0 | 0 | 0 | 0 | 0 | 1 | 1 |
| ENSMUSG00000020644 | Id2 | ENSMUST00000020974 | 31.2899106 | 35 | 15.8365262 | 17.752235 | 1 | 0 | 0 | 1 | 0 | 0 | 0 | 0 | 1 | 2 | 0 | 0 | 0 | 0 | 3 |
| ENSMUSG00000021379 | Id4 | ENSMUST00000021810 | 27.8331822 | 35 | 15.4125113 | 22.1214869 | 0 | 0 | 0 | 0 | 0 | 0 | 0 | 0 | 0 | 0 | 0 | 0 | 2 | 1 | 3 |
| ENSMUSG00000056999 | Ide | ENSMUST00000131070 | 27.7638819 | 31 | 21.9609805 | 19.5597799 | 0 | 0 | 0 | 0 | 0 | 0 | 0 | 0 | 0 | 0 | 0 | 0 | 0 | 1 | 1 |
| ENSMUSG00000025950 | Idh1 | ENSMUST00000097709 | 28.8574794 | 35 | 16.8433451 | 19.1990577 | 0 | 0 | 0 | 0 | 0 | 0 | 0 | 0 | 0 | 0 | 0 | 0 | 4 | 0 | 4 |
| ENSMUSG00000032279 | Idh3a | ENSMUST00000167866 | 27.0833333 | 30 | 23.5339506 | 19.058642 | 0 | 0 | 0 | 0 | 0 | 0 | 0 | 0 | 0 | 0 | 0 | 1 | 0 | 0 | 1 |
| ENSMUSG00000058258 | Idi1 | ENSMUST00000169314 | 29.5704846 | 33 | 17.1255507 | 20.0440529 | 0 | 0 | 0 | 0 | 0 | 0 | 0 | 0 | 0 | 0 | 0 | 17 | 1 | 0 | 18 |
| ENSMUSG00000035847 | Ids | ENSMUST00000101509 | 28.7037037 | 36 | 18.614304 | 16.3154534 | 0 | 0 | 0 | 1 | 0 | 0 | 0 | 0 | 1 | 2 | 0 | 4 | 3 | 1 | 10 |
| ENSMUSG00000003541 | Ier3 | ENSMUST00000003635 | 23.4910277 | 27 | 20.228385 | 29.0375204 | 2 | 0 | 2 | 0 | 0 | 0 | 0 | 0 | 0 | 2 | 0 | 0 | 0 | 0 | 4 |
| ENSMUSG00000090000 | Ier3ip1 | ENSMUST00000026487 | 28.2545805 | 35 | 15.8148505 | 21.0221794 | 0 | 0 | 0 | 0 | 0 | 0 | 0 | 0 | 0 | 0 | 0 | 0 | 3 | 1 | 4 |
| ENSMUSG00000056708 | Ier5 | ENSMUST00000055322 | 28.0914606 | 29 | 18.8520765 | 24.4050397 | 0 | 0 | 0 | 0 | 0 | 0 | 0 | 0 | 0 | 0 | 1 | 0 | 0 | 0 | 1 |
| ENSMUSG00000089762 | Ier5l | ENSMUST00000065134 | 21.2121212 | 24 | 26.8740032 | 27.7511962 | 0 | 0 | 0 | 1 | 0 | 0 | 0 | 0 | 1 | 2 | 0 | 0 | 0 | 0 | 2 |
| ENSMUSG00000022965 | Ifngr2 | ENSMUST00000023687 | 25.323475 | 25 | 24.9075786 | 24.3068392 | 0 | 0 | 0 | 0 | 0 | 0 | 0 | 0 | 0 | 0 | 0 | 0 | 5 | 2 | 7 |
| ENSMUSG00000001627 | Ifrd1 | ENSMUST00000001672 | 25.2367688 | 34 | 18.1615599 | 22.1727019 | 0 | 0 | 0 | 0 | 0 | 0 | 0 | 0 | 0 | 0 | 0 | 1 | 0 | 0 | 1 |
| ENSMUSG00000028576 | Ift74 | ENSMUST00000107104 | 31.3513514 | 39 | 14.0540541 | 15.4054054 | 0 | 0 | 0 | 1 | 0 | 0 | 0 | 0 | 1 | 2 | 0 | 0 | 0 | 0 | 2 |
| ENSMUSG00000027778 | Ift80 | ENSMUST00000107812 | 29.6437659 | 34 | 17.3664122 | 19.3384224 | 0 | 0 | 0 | 0 | 0 | 0 | 0 | 0 | 0 | 0 | 4 | 0 | 3 | 2 | 9 |
| ENSMUSG00000029469 | Ift81 | ENSMUST00000031426 | 26.3157895 | 34 | 18.3598531 | 21.1750306 | 0 | 0 | 0 | 0 | 0 | 0 | 0 | 0 | 0 | 0 | 0 | 1 | 0 | 0 | 1 |
| ENSMUSG00000031221 | Igbp1 | ENSMUST00000033570 | 25.952381 | 30 | 19.7619048 | 24.2857143 | 0 | 0 | 0 | 0 | 0 | 0 | 0 | 0 | 0 | 0 | 0 | 0 | 1 | 0 | 1 |
| ENSMUSG00000020053 | Igf1 | ENSMUST00000095360 | 30.9606848 | 32 | 18.7222575 | 18.0088776 | 0 | 0 | 0 | 0 | 0 | 0 | 0 | 0 | 0 | 0 | 0 | 1 | 10 | 0 | 11 |
| ENSMUSG00000020427 | Igfbp3 | ENSMUST00000020702 | 27.804878 | 29 | 20.9059233 | 22.71777 | 0 | 0 | 0 | 0 | 0 | 0 | 0 | 0 | 0 | 0 | 0 | 1 | 0 | 0 | 1 |
| ENSMUSG00000019975 | Ikbip | ENSMUST00000020149 | 29.2156863 | 33 | 19.1176471 | 18.8235294 | 0 | 0 | 0 | 0 | 0 | 0 | 0 | 0 | 0 | 0 | 0 | 1 | 9 | 2 | 12 |
| ENSMUSG00000018654 | Ikzf1 | ENSMUST00000076700 | 27.8026906 | 30 | 20.8669656 | 20.896861 | 0 | 0 | 0 | 0 | 0 | 0 | 0 | 0 | 0 | 0 | 1 | 0 | 1 | 1 | 3 |
| ENSMUSG00000025997 | Ikzf2 | ENSMUST00000027146 | 28.1837437 | 34 | 18.0963728 | 19.8040773 | 0 | 0 | 0 | 1 | 0 | 0 | 0 | 0 | 1 | 2 | 0 | 5 | 18 | 1 | 26 |
| ENSMUSG00000002578 | Ikzf4 | ENSMUST00000133342 | 23.8689548 | 28 | 27.2698908 | 20.4368175 | 0 | 0 | 0 | 0 | 0 | 0 | 0 | 0 | 0 | 0 | 0 | 9 | 0 | 0 | 9 |
| ENSMUSG00000040167 | Ikzf5 | ENSMUST00000046306 | 30.8207705 | 34 | 16.9849246 | 18.4589615 | 0 | 0 | 0 | 1 | 0 | 0 | 0 | 0 | 1 | 2 | 0 | 4 | 0 | 0 | 6 |
| ENSMUSG00000018341 | Il12rb2 | ENSMUST00000117441 | 27.0056497 | 33 | 27.9096045 | 11.8644068 | 0 | 0 | 0 | 0 | 0 | 0 | 0 | 0 | 0 | 0 | 0 | 26 | 0 | 2 | 28 |
| ENSMUSG00000017057 | Il13ra1 | ENSMUST00000033418 | 28.4317892 | 33 | 19.3370166 | 19.2095198 | 0 | 0 | 0 | 0 | 0 | 0 | 0 | 0 | 0 | 0 | 0 | 2 | 0 | 0 | 2 |
| ENSMUSG00000040717 | Il17rd | ENSMUST00000035336 | 23.9830508 | 28 | 22.779661 | 25.1864407 | 0 | 0 | 0 | 0 | 0 | 0 | 0 | 0 | 0 | 0 | 0 | 6 | 1 | 0 | 7 |
| ENSMUSG00000022514 | Il1rap | ENSMUST00000173136 | 32.3901713 | 32 | 16.4184661 | 18.8384214 | 1 | 0 | 0 | 0 | 0 | 0 | 0 | 0 | 0 | 0 | 0 | 0 | 0 | 0 | 1 |
| ENSMUSG00000000869 | Il4 | ENSMUST00000000889 | 38.4615385 | 44 | 8.65384615 | 8.65384615 | 1 | 0 | 0 | 2 | 0 | 0 | 0 | 0 | 2 | 4 | 0 | 0 | 0 | 0 | 5 |
| ENSMUSG00000001016 | Ilf2 | ENSMUST00000001042 | 30.6578947 | 31 | 19.4736842 | 19.0789474 | 0 | 0 | 0 | 0 | 0 | 0 | 0 | 0 | 0 | 0 | 0 | 0 | 3 | 2 | 5 |
| ENSMUSG00000042670 | Immp1l | ENSMUST00000037499 | 38.9380531 | 38 | 11.9469027 | 11.0619469 | 1 | 0 | 1 | 0 | 0 | 0 | 0 | 0 | 0 | 1 | 0 | 0 | 0 | 0 | 2 |
| ENSMUSG00000026127 | Imp4 | ENSMUST00000027303 | 23.8845682 | 30 | 23.1887024 | 22.5133033 | 0 | 0 | 0 | 0 | 0 | 0 | 0 | 0 | 0 | 0 | 1 | 4 | 8 | 1 | 14 |
| ENSMUSG00000027531 | Impa1 | ENSMUST00000065938 | 25.8513932 | 31 | 20.495356 | 22.4767802 | 1 | 0 | 0 | 1 | 0 | 0 | 0 | 0 | 1 | 2 | 4 | 1 | 4 | 2 | 14 |
| ENSMUSG00000024423 | Impact | ENSMUST00000025290 | 27.4058577 | 34 | 18.8702929 | 20 | 1 | 0 | 0 | 1 | 0 | 0 | 0 | 0 | 1 | 2 | 0 | 0 | 12 | 0 | 15 |
| ENSMUSG00000066324 | Impad1 | ENSMUST00000084949 | 29.5210728 | 36 | 16.0153257 | 18.7931034 | 1 | 1 | 0 | 0 | 0 | 0 | 0 | 0 | 0 | 0 | 3 | 0 | 9 | 2 | 16 |
| ENSMUSG00000035270 | Impg2 | ENSMUST00000069936 | 31.7139001 | 31 | 17.1727395 | 19.7705803 | 1 | 0 | 0 | 1 | 0 | 0 | 0 | 0 | 1 | 2 | 0 | 1 | 0 | 0 | 4 |
| ENSMUSG00000029670 | Ing3 | ENSMUST00000031680 | 31.7657118 | 32 | 16.8020522 | 19.8375374 | 0 | 0 | 0 | 0 | 0 | 0 | 0 | 0 | 0 | 0 | 1 | 0 | 0 | 0 | 1 |
| ENSMUSG00000026283 | Ing5 | ENSMUST00000188402 | 24.4990893 | 28 | 22.0400729 | 25.273224 | 3 | 1 | 0 | 5 | 0 | 0 | 1 | 2 | 2 | 10 | 0 | 1 | 7 | 0 | 22 |
| ENSMUSG00000047989 | Ino80c | ENSMUST00000141489 | 28.700565 | 28 | 21.4124294 | 21.7514124 | 0 | 0 | 0 | 0 | 0 | 0 | 0 | 0 | 0 | 0 | 0 | 1 | 0 | 0 | 1 |
| ENSMUSG00000040865 | Ino80d | ENSMUST00000097718 | 26.3406011 | 33 | 19.4755451 | 21.1844431 | 1 | 3 | 0 | 1 | 0 | 0 | 0 | 0 | 1 | 2 | 8 | 2 | 13 | 3 | 32 |
| ENSMUSG00000026102 | Inpp1 | ENSMUST00000027271 | 30.6767272 | 32 | 18.5097854 | 18.7691582 | 1 | 1 | 0 | 3 | 0 | 0 | 0 | 1 | 2 | 6 | 0 | 4 | 0 | 0 | 12 |
| ENSMUSG00000042105 | Inpp5f | ENSMUST00000043138 | 30.5454545 | 34 | 17.6363636 | 17.6363636 | 0 | 0 | 1 | 1 | 0 | 0 | 0 | 0 | 1 | 3 | 0 | 3 | 0 | 0 | 6 |
| ENSMUSG00000045294 | Insig1 | ENSMUST00000059155 | 26.2214984 | 31 | 19.5439739 | 23.5613464 | 1 | 0 | 1 | 1 | 0 | 0 | 0 | 0 | 1 | 3 | 0 | 0 | 14 | 0 | 18 |
| ENSMUSG00000003721 | Insig2 | ENSMUST00000003818 | 27.6883451 | 31 | 18.3515776 | 22.5370251 | 0 | 0 | 0 | 0 | 0 | 0 | 0 | 0 | 0 | 0 | 0 | 1 | 0 | 0 | 1 |
| ENSMUSG00000068154 | Insm1 | ENSMUST00000089257 | 21.8671152 | 35 | 22.7922624 | 20.0168209 | 2 | 1 | 0 | 2 | 0 | 0 | 0 | 1 | 1 | 4 | 0 | 30 | 0 | 0 | 37 |
| ENSMUSG00000018068 | Ints2 | ENSMUST00000018212 | 33.3162481 | 33 | 15.3767299 | 18.195797 | 1 | 0 | 0 | 1 | 0 | 0 | 0 | 0 | 1 | 2 | 0 | 0 | 0 | 0 | 3 |
| ENSMUSG00000060798 | Intu | ENSMUST00000091186 | 29.1823172 | 31 | 19.9653279 | 19.8497544 | 0 | 0 | 1 | 0 | 0 | 0 | 0 | 0 | 0 | 1 | 0 | 1 | 0 | 0 | 2 |
| ENSMUSG00000060733 | Ipmk | ENSMUST00000079252 | 26.4322597 | 31 | 20.6048685 | 22.3998033 | 0 | 0 | 0 | 0 | 0 | 0 | 0 | 0 | 0 | 0 | 0 | 1 | 0 | 0 | 1 |
| ENSMUSG00000042590 | Ipo11 | ENSMUST00000190801 | 30.1719028 | 26 | 21.2803794 | 22.3473622 | 0 | 0 | 1 | 0 | 0 | 0 | 0 | 0 | 0 | 1 | 0 | 0 | 0 | 0 | 1 |
| ENSMUSG00000040029 | Ipo8 | ENSMUST00000048418 | 25.2491694 | 33 | 19.9335548 | 22.021832 | 0 | 0 | 0 | 0 | 0 | 0 | 0 | 0 | 0 | 0 | 0 | 0 | 1 | 1 | 2 |
| ENSMUSG00000021385 | Ippk | ENSMUST00000021817 | 23.9642567 | 31 | 19.8212835 | 24.9390739 | 0 | 0 | 0 | 0 | 0 | 0 | 0 | 0 | 0 | 0 | 0 | 1 | 2 | 2 | 5 |
| ENSMUSG00000022837 | Iqcb1 | ENSMUST00000114819 | 28.8613303 | 31 | 20.1803833 | 19.729425 | 0 | 0 | 0 | 0 | 0 | 0 | 0 | 0 | 0 | 0 | 0 | 0 | 1 | 2 | 3 |
| ENSMUSG00000032251 | Irak1bp1 | ENSMUST00000113245 | 31.3483146 | 30 | 21.7977528 | 16.9662921 | 0 | 0 | 0 | 0 | 0 | 0 | 0 | 0 | 0 | 0 | 1 | 1 | 0 | 1 | 3 |
| ENSMUSG00000032293 | Ireb2 | ENSMUST00000034843 | 30.8986753 | 35 | 16.0042965 | 18.5105621 | 0 | 0 | 0 | 0 | 0 | 0 | 0 | 0 | 0 | 0 | 1 | 0 | 2 | 1 | 4 |
| ENSMUSG00000031627 | Irf2 | ENSMUST00000034041 | 25.6472492 | 31 | 26.618123 | 16.828479 | 0 | 0 | 0 | 0 | 0 | 0 | 0 | 0 | 0 | 0 | 1 | 0 | 0 | 0 | 1 |
| ENSMUSG00000042258 | Isl1 | ENSMUST00000036060 | 30.3586322 | 34 | 16.9307756 | 18.6822352 | 1 | 0 | 0 | 0 | 0 | 0 | 0 | 0 | 0 | 0 | 0 | 0 | 6 | 1 | 8 |
| ENSMUSG00000024601 | Isoc1 | ENSMUST00000025503 | 27.8481013 | 33 | 19.5901145 | 19.6503918 | 0 | 0 | 0 | 0 | 0 | 0 | 0 | 0 | 0 | 0 | 0 | 2 | 0 | 0 | 2 |
| ENSMUSG00000031729 | Ist1 | ENSMUST00000034164 | 20.9591474 | 33 | 22.6465364 | 23.6234458 | 0 | 0 | 0 | 0 | 0 | 0 | 0 | 0 | 0 | 0 | 0 | 1 | 0 | 0 | 1 |
| ENSMUSG00000027598 | Itch | ENSMUST00000029126 | 27.7521976 | 32 | 21.2641272 | 18.6688991 | 0 | 0 | 0 | 0 | 0 | 0 | 0 | 0 | 0 | 0 | 0 | 1 | 0 | 0 | 1 |
| ENSMUSG00000031703 | Itfg1 | ENSMUST00000034140 | 30.0533944 | 34 | 16.8573608 | 18.993135 | 0 | 0 | 0 | 0 | 0 | 0 | 0 | 0 | 0 | 0 | 0 | 0 | 1 | 0 | 1 |
| ENSMUSG00000027009 | Itga4 | ENSMUST00000099972 | 29.9631223 | 31 | 21.3583282 | 17.3017824 | 2 | 1 | 0 | 2 | 0 | 0 | 0 | 0 | 2 | 4 | 0 | 5 | 0 | 0 | 12 |
| ENSMUSG00000027111 | Itga6 | ENSMUST00000155249 | 22.1374046 | 36 | 21.2977099 | 20.9160305 | 1 | 1 | 1 | 1 | 0 | 0 | 0 | 0 | 1 | 3 | 0 | 2 | 0 | 0 | 7 |
| ENSMUSG00000026768 | Itga8 | ENSMUST00000172791 | 30.7993475 | 28 | 20.3262643 | 20.5220228 | 2 | 1 | 0 | 1 | 0 | 0 | 0 | 0 | 1 | 2 | 0 | 0 | 1 | 0 | 6 |
| ENSMUSG00000025321 | Itgb8 | ENSMUST00000026360 | 31.7958783 | 34 | 17.4484789 | 16.6437684 | 0 | 0 | 0 | 0 | 0 | 0 | 0 | 0 | 0 | 0 | 0 | 1 | 0 | 0 | 1 |
| ENSMUSG00000032925 | Itgbl1 | ENSMUST00000142161 | 31.3502935 | 31 | 15.9295499 | 21.8786693 | 1 | 0 | 0 | 0 | 0 | 0 | 0 | 0 | 0 | 0 | 0 | 0 | 1 | 0 | 2 |
| ENSMUSG00000026223 | Itm2c | ENSMUST00000027425 | 18.611379 | 26 | 26.8081003 | 28.4474446 | 0 | 0 | 0 | 0 | 0 | 0 | 0 | 0 | 0 | 0 | 0 | 1 | 0 | 0 | 1 |
| ENSMUSG00000095115 | Itpripl2 | ENSMUST00000178344 | 27.0539779 | 30 | 19.4884023 | 23.1953176 | 1 | 0 | 0 | 0 | 0 | 0 | 0 | 0 | 0 | 0 | 4 | 1 | 2 | 3 | 11 |
| ENSMUSG00000020640 | Itsn2 | ENSMUST00000062580 | 29.8804781 | 30 | 18.7250996 | 21.1155378 | 1 | 0 | 0 | 0 | 0 | 0 | 0 | 0 | 0 | 0 | 0 | 0 | 0 | 0 | 1 |
| ENSMUSG00000023150 | Ivns1abp | ENSMUST00000186745 | 27.505618 | 29 | 19.505618 | 23.8651685 | 0 | 0 | 0 | 0 | 0 | 0 | 0 | 0 | 0 | 0 | 0 | 0 | 2 | 0 | 2 |
| ENSMUSG00000024384 | Iws1 | ENSMUST00000025243 | 25.8297872 | 35 | 18.7234043 | 20.0425532 | 0 | 0 | 0 | 0 | 0 | 0 | 0 | 0 | 0 | 0 | 8 | 0 | 12 | 0 | 20 |
| ENSMUSG00000027276 | Jag1 | ENSMUST00000028735 | 26.3392857 | 37 | 15.880102 | 21.1096939 | 4 | 0 | 2 | 3 | 0 | 0 | 0 | 0 | 3 | 8 | 0 | 1 | 15 | 0 | 28 |
| ENSMUSG00000051256 | Jagn1 | ENSMUST00000204254 | 22.5118483 | 29 | 25.3554502 | 22.985782 | 0 | 0 | 0 | 0 | 0 | 0 | 0 | 0 | 0 | 0 | 0 | 1 | 0 | 0 | 1 |
| ENSMUSG00000053062 | Jam2 | ENSMUST00000114195 | 28.7552268 | 29 | 23.0942425 | 19.491798 | 0 | 0 | 0 | 0 | 0 | 0 | 0 | 0 | 0 | 0 | 1 | 0 | 0 | 0 | 1 |
| ENSMUSG00000031990 | Jam3 | ENSMUST00000034472 | 26.1977574 | 30 | 21.1009174 | 22.9357798 | 0 | 0 | 0 | 0 | 0 | 0 | 0 | 0 | 0 | 0 | 0 | 2 | 0 | 0 | 2 |
| ENSMUSG00000037876 | Jmjd1c | ENSMUST00000051446 | 33.5868188 | 34 | 14.5754119 | 17.3637516 | 1 | 1 | 1 | 2 | 0 | 0 | 0 | 1 | 1 | 5 | 0 | 0 | 0 | 1 | 8 |
| ENSMUSG00000079083 | Jrkl | ENSMUST00000110582 | 33.0239753 | 40 | 10.9048724 | 15.6999227 | 2 | 0 | 0 | 1 | 0 | 0 | 0 | 0 | 1 | 2 | 0 | 0 | 5 | 2 | 11 |
| ENSMUSG00000052837 | Junb | ENSMUST00000064922 | 20.5761317 | 26 | 26.7489712 | 26.5432099 | 0 | 0 | 0 | 0 | 0 | 0 | 0 | 0 | 0 | 0 | 1 | 1 | 2 | 0 | 4 |
| ENSMUSG00000018412 | Kansl1 | ENSMUST00000106977 | 25.2144082 | 32 | 22.4128073 | 20.2401372 | 0 | 0 | 0 | 1 | 0 | 0 | 0 | 0 | 1 | 2 | 0 | 2 | 2 | 0 | 6 |
| ENSMUSG00000022992 | Kansl2 | ENSMUST00000023727 | 24.4592346 | 33 | 20.1331115 | 22.4625624 | 0 | 0 | 0 | 0 | 0 | 0 | 0 | 0 | 0 | 0 | 0 | 0 | 1 | 1 | 2 |
| ENSMUSG00000000708 | Kat2b | ENSMUST00000000724 | 29.1036717 | 35 | 16.7386609 | 18.7365011 | 0 | 0 | 0 | 0 | 0 | 0 | 0 | 0 | 0 | 0 | 1 | 0 | 3 | 1 | 5 |
| ENSMUSG00000041298 | Katnal1 | ENSMUST00000047257 | 27.3166447 | 28 | 22.5735617 | 21.8928415 | 3 | 2 | 0 | 3 | 0 | 0 | 0 | 1 | 2 | 6 | 0 | 2 | 1 | 2 | 16 |
| ENSMUSG00000027132 | Katnbl1 | ENSMUST00000028552 | 30.4109589 | 36 | 16.6027397 | 16.7671233 | 0 | 0 | 0 | 0 | 0 | 0 | 0 | 0 | 0 | 0 | 1 | 0 | 3 | 3 | 7 |
| ENSMUSG00000059486 | Kbtbd2 | ENSMUST00000114323 | 26.2186115 | 34 | 18.6853767 | 20.8271787 | 0 | 0 | 0 | 1 | 0 | 0 | 0 | 0 | 1 | 2 | 2 | 0 | 6 | 0 | 10 |
| ENSMUSG00000075502 | Kbtbd6 | ENSMUST00000100359 | 29.9230769 | 36 | 16.1538462 | 17.5384615 | 1 | 0 | 0 | 3 | 0 | 0 | 0 | 0 | 3 | 6 | 1 | 0 | 0 | 0 | 8 |
| ENSMUSG00000043881 | Kbtbd7 | ENSMUST00000061222 | 31.0683952 | 37 | 14.3495753 | 17.2999553 | 1 | 1 | 0 | 1 | 0 | 0 | 0 | 0 | 1 | 2 | 0 | 0 | 0 | 0 | 4 |
| ENSMUSG00000030031 | Kbtbd8 | ENSMUST00000032107 | 29.6842105 | 34 | 16.1403509 | 19.7192982 | 1 | 1 | 0 | 0 | 0 | 0 | 0 | 0 | 0 | 0 | 0 | 1 | 1 | 0 | 4 |
| ENSMUSG00000055239 | Kcmf1 | ENSMUST00000068697 | 25.8689459 | 32 | 21.4814815 | 20.4558405 | 0 | 0 | 0 | 1 | 0 | 0 | 0 | 0 | 1 | 2 | 0 | 1 | 0 | 1 | 4 |
| ENSMUSG00000027827 | Kcnab1 | ENSMUST00000049230 | 31.496063 | 32 | 19.3821926 | 17.0199879 | 0 | 0 | 0 | 0 | 0 | 0 | 0 | 0 | 0 | 0 | 0 | 0 | 1 | 0 | 1 |
| ENSMUSG00000092083 | Kcnb2 | ENSMUST00000170146 | 31.7465556 | 31 | 18.7278277 | 18.0678162 | 0 | 0 | 0 | 0 | 0 | 0 | 0 | 0 | 0 | 0 | 2 | 4 | 1 | 1 | 8 |
| ENSMUSG00000035681 | Kcnc2 | ENSMUST00000092175 | 27.6590693 | 34 | 19.6581197 | 18.9458689 | 7 | 2 | 0 | 12 | 0 | 0 | 1 | 3 | 8 | 24 | 2 | 0 | 7 | 5 | 47 |
| ENSMUSG00000039672 | Kcne2 | ENSMUST00000047383 | 29.3766012 | 28 | 19.7267293 | 22.6302306 | 0 | 0 | 1 | 0 | 0 | 0 | 0 | 0 | 0 | 1 | 0 | 0 | 0 | 0 | 1 |

| | | | | | | | | | | | | | | | | | | | | | |
|---|---|---|---|---|---|---|---|---|---|---|---|---|---|---|---|---|---|---|---|---|---|
| ENSMUSG00000045053 | Kcng3 | ENSMUST00000051482 | 26.6741699 | 31 | 21.5531795 | 20.8778841 | 1 | 0 | 0 | 0 | 0 | 0 | 0 | 0 | 0 | 0 | 0 | 1 | 0 | 0 | 2 |
| ENSMUSG00000059742 | Kcnh7 | ENSMUST00000075052 | 33.015107 | 30 | 18.1913554 | 18.7368863 | 2 | 0 | 1 | 2 | 0 | 0 | 0 | 0 | 2 | 5 | 0 | 0 | 0 | 4 | 11 |
| ENSMUSG00000041695 | Kcnj2 | ENSMUST00000042970 | 27.166089 | 34 | 18.5017329 | 20.1279659 | 0 | 0 | 0 | 0 | 0 | 0 | 0 | 0 | 0 | 0 | 3 | 19 | 8 | 1 | 31 |
| ENSMUSG00000033854 | Kcnk10 | ENSMUST00000110113 | 24.8462485 | 28 | 21.8942189 | 25.3382534 | 0 | 0 | 0 | 0 | 0 | 0 | 0 | 0 | 0 | 0 | 0 | 1 | 2 | 0 | 3 |
| ENSMUSG00000098557 | Kctd12 | ENSMUST00000184744 | 27.9942576 | 33 | 19.0319934 | 20.1599672 | 0 | 0 | 0 | 0 | 0 | 0 | 0 | 0 | 0 | 0 | 3 | 20 | 3 | 0 | 26 |
| ENSMUSG00000044952 | Kctd21 | ENSMUST00000054107 | 25.1303936 | 25 | 26.2209578 | 23.4708393 | 0 | 0 | 0 | 0 | 0 | 0 | 0 | 0 | 0 | 0 | 0 | 3 | 0 | 0 | 3 |
| ENSMUSG00000016946 | Kctd5 | ENSMUST00000017090 | 20.9399167 | 30 | 23.6168947 | 24.9851279 | 0 | 0 | 0 | 0 | 0 | 0 | 0 | 0 | 0 | 0 | 0 | 1 | 0 | 0 | 1 |
| ENSMUSG00000034110 | Kctd7 | ENSMUST00000040616 | 23.7261644 | 28 | 23.2572679 | 24.7889966 | 0 | 0 | 0 | 0 | 0 | 0 | 0 | 0 | 0 | 0 | 0 | 0 | 4 | 4 | 8 |
| ENSMUSG00000034327 | Kctd9 | ENSMUST00000156700 | 28.3169067 | 30 | 19.2570129 | 22.6686884 | 1 | 0 | 0 | 0 | 0 | 0 | 0 | 0 | 0 | 0 | 0 | 0 | 1 | 1 | 3 |
| ENSMUSG00000079111 | Kdelr2 | ENSMUST00000110731 | 25.2345216 | 32 | 20.8255159 | 21.4821764 | 0 | 0 | 0 | 0 | 0 | 0 | 0 | 0 | 0 | 0 | 1 | 0 | 12 | 1 | 14 |
| ENSMUSG00000028397 | Kdm4c | ENSMUST00000030102 | 30.3592121 | 33 | 17.7288528 | 18.8876014 | 0 | 0 | 1 | 1 | 0 | 0 | 0 | 0 | 1 | 3 | 0 | 0 | 0 | 0 | 3 |
| ENSMUSG00000037369 | Kdm6a | ENSMUST00000044484 | 32.8649493 | 38 | 14.3637783 | 14.9882904 | 0 | 0 | 0 | 0 | 0 | 0 | 0 | 0 | 0 | 0 | 0 | 0 | 0 | 1 | 1 |
| ENSMUSG00000042599 | Kdm7a | ENSMUST00000002305 | 29.5992714 | 34 | 17.0461445 | 19.3685489 | 1 | 1 | 1 | 1 | 0 | 0 | 0 | 0 | 1 | 3 | 0 | 1 | 0 | 6 | 12 |
| ENSMUSG00000009905 | Kdsr | ENSMUST00000010049 | 26.6036462 | 34 | 18.8836372 | 20.4141346 | 2 | 1 | 0 | 3 | 0 | 0 | 0 | 1 | 2 | 6 | 1 | 2 | 18 | 2 | 32 |
| ENSMUSG00000028790 | Khdrbs1 | ENSMUST00000066257 | 31.938326 | 32 | 17.3568282 | 18.7665198 | 0 | 0 | 0 | 0 | 0 | 0 | 0 | 0 | 0 | 0 | 0 | 1 | 0 | 2 | 3 |
| ENSMUSG00000026058 | Khdrbs2 | ENSMUST00000188257 | 29.7834733 | 34 | 18.3163942 | 17.4768007 | 3 | 0 | 0 | 1 | 0 | 0 | 0 | 0 | 1 | 2 | 1 | 0 | 0 | 1 | 7 |
| ENSMUSG00000007670 | Khsrp | ENSMUST00000007814 | 18.8379205 | 30 | 28.1957187 | 22.5076453 | 0 | 0 | 0 | 0 | 0 | 0 | 0 | 0 | 0 | 0 | 0 | | 9 | 0 | 9 |
| ENSMUSG00000012443 | Kif11 | ENSMUST00000012587 | 28.1143635 | 33 | 20.1497617 | 18.7882914 | 0 | 0 | 0 | 0 | 0 | 0 | 0 | 0 | 0 | 0 | 0 | 1 | 0 | 0 | 1 |
| ENSMUSG00000036768 | Kif15 | ENSMUST00000040717 | 22.5752508 | 38 | 20.0668896 | 19.5652174 | 1 | 0 | 0 | 0 | 0 | 0 | 0 | 0 | 0 | 0 | 0 | 0 | 4 | 0 | 5 |
| ENSMUSG00000022629 | Kif21a | ENSMUST00000067205 | 29.2163543 | 38 | 14.5655877 | 18.0579216 | 0 | 0 | 0 | 0 | 0 | 0 | 0 | 0 | 0 | 0 | 1 | 0 | 6 | 0 | 7 |
| ENSMUSG00000021693 | Kif2a | ENSMUST00000022204 | 31.9758193 | 33 | 15.9720013 | 19.0264079 | 0 | 1 | 0 | 0 | 0 | 0 | 0 | 0 | 0 | 0 | 0 | 0 | 5 | 1 | 7 |
| ENSMUSG00000006740 | Kif5b | ENSMUST00000025083 | 31.3876652 | 34 | 15.8223201 | 18.3920705 | 2 | 1 | 0 | 4 | 0 | 0 | 0 | 1 | 3 | 8 | 0 | 0 | 1 | 1 | 13 |
| ENSMUSG00000026764 | Kif5c | ENSMUST00000028102 | 25.7939796 | 32 | 19.4973764 | 22.5075946 | 1 | 0 | 0 | 0 | 0 | 0 | 0 | 0 | 0 | 0 | 4 | 0 | 5 | 0 | 10 |
| ENSMUSG00000041734 | Kirrel | ENSMUST00000159976 | 23.8628873 | 27 | 25.3570644 | 24.1924852 | 1 | 0 | 0 | 1 | 0 | 0 | 0 | 0 | 1 | 2 | 0 | 1 | 1 | 4 | 9 |
| ENSMUSG00000005672 | Kit | ENSMUST00000005815 | 24.7588424 | 30 | 22.3242995 | 22.6458429 | 0 | 0 | 0 | 0 | 0 | 0 | 0 | 0 | 0 | 0 | 0 | 1 | 0 | 0 | 1 |
| ENSMUSG00000019966 | Kitl | ENSMUST00000105283 | 31.3628159 | 32 | 17.2608303 | 19.4494585 | 0 | 1 | 0 | 0 | 0 | 0 | 0 | 0 | 0 | 0 | 1 | 0 | 0 | 0 | 2 |
| ENSMUSG00000037465 | Klf10 | ENSMUST00000074043 | 28.2841823 | 33 | 17.2252011 | 21.4477212 | 0 | 0 | 0 | 0 | 0 | 0 | 0 | 0 | 0 | 0 | 1 | 0 | 4 | 2 | 7 |
| ENSMUSG00000020653 | Klf11 | ENSMUST00000020982 | 25.0108178 | 34 | 18.5201212 | 22.3279965 | 1 | 0 | 0 | 1 | 0 | 0 | 0 | 0 | 1 | 2 | 0 | 5 | 0 | 0 | 8 |
| ENSMUSG00000029178 | Klf3 | ENSMUST00000165536 | 27.5714667 | 33 | 19.5030724 | 20.4114347 | 0 | 0 | 0 | 0 | 0 | 0 | 0 | 0 | 0 | 0 | 2 | 10 | 10 | 3 | 25 |
| ENSMUSG00000000078 | Klf6 | ENSMUST00000000080 | 29.5176849 | 32 | 18.0385852 | 20.6752412 | 0 | 0 | 0 | 0 | 0 | 0 | 0 | 0 | 0 | 0 | 0 | 1 | 5 | 1 | 7 |
| ENSMUSG00000025959 | Klf7 | ENSMUST00000114086 | 26.9869135 | 30 | 22.7736993 | 20.4117459 | 2 | 0 | 0 | 2 | 0 | 0 | 0 | 0 | 2 | 4 | 1 | 7 | 1 | 2 | 17 |
| ENSMUSG00000051890 | Klhdc1 | ENSMUST00000063445 | 27.9512567 | 32 | 19.7258187 | 20.4112719 | 0 | 0 | 0 | 0 | 0 | 0 | 0 | 0 | 0 | 0 | 1 | 0 | 0 | 0 | 1 |
| ENSMUSG00000029775 | Klhdc10 | ENSMUST00000068259 | 24.0017845 | 30 | 22.1503458 | 23.4441222 | 0 | 0 | 0 | 0 | 0 | 0 | 0 | 0 | 0 | 0 | 0 | 2 | 0 | 0 | 2 |
| ENSMUSG00000020978 | Klhdc2 | ENSMUST00000021362 | 23.8442822 | 35 | 19.7080292 | 21.4111922 | 1 | 1 | 0 | 0 | 0 | 0 | 0 | 0 | 0 | 0 | 0 | 0 | 1 | 1 | 4 |
| ENSMUSG00000043929 | Klhl15 | ENSMUST00000113915 | 28.7626203 | 38 | 15.9427096 | 17.3984503 | 0 | 0 | 0 | 1 | 0 | 0 | 0 | 0 | 1 | 2 | 0 | 0 | 23 | 2 | 27 |
| ENSMUSG00000031605 | Klhl2 | ENSMUST00000034017 | 29.0858726 | 37 | 15.5124654 | 18.6518929 | 0 | 0 | 0 | 0 | 0 | 0 | 0 | 0 | 0 | 0 | 0 | 0 | 0 | 1 | 1 |
| ENSMUSG00000026705 | Klhl20 | ENSMUST00000111611 | 26.9005848 | 33 | 21.3937622 | 18.5672515 | 0 | 0 | 2 | 0 | 0 | 0 | 0 | 0 | 0 | 2 | 0 | 2 | 2 | 1 | 7 |
| ENSMUSG00000062901 | Klhl24 | ENSMUST00000023509 | 30.0344976 | 31 | 18.1759379 | 20.5260888 | 0 | 1 | 0 | 0 | 0 | 0 | 0 | 0 | 0 | 0 | 0 | 0 | 9 | 0 | 10 |
| ENSMUSG00000020948 | Klhl28 | ENSMUST00000021331 | 27.5247525 | 37 | 14.0594059 | 21.0891089 | 0 | 0 | 0 | 0 | 0 | 0 | 0 | 0 | 0 | 0 | 0 | 0 | 1 | 0 | 1 |
| ENSMUSG00000054920 | Klhl5 | ENSMUST00000203538 | 29.8097252 | 31 | 19.5912615 | 20.0845666 | 0 | 0 | 0 | 0 | 0 | 0 | 0 | 0 | 0 | 0 | 0 | 0 | 1 | 0 | 1 |
| ENSMUSG00000070923 | Klhl9 | ENSMUST00000094993 | 29.994615 | 37 | 15.3473344 | 17.6628971 | 0 | 0 | 0 | 0 | 0 | 0 | 0 | 0 | 0 | 0 | 0 | 0 | 8 | 2 | 10 |
| ENSMUSG00000018362 | Kpna2 | ENSMUST00000145331 | 27.0411496 | 29 | 22.1423906 | 22.2077074 | 0 | 0 | 0 | 0 | 0 | 0 | 0 | 0 | 0 | 0 | 0 | 1 | 0 | 0 | 1 |
| ENSMUSG00000021929 | Kpna3 | ENSMUST00000022496 | 29.9795501 | 34 | 17.6278119 | 18.6503067 | 0 | 0 | 0 | 0 | 0 | 0 | 0 | 0 | 0 | 0 | 0 | 0 | 1 | 2 | 3 |
| ENSMUSG00000027782 | Kpna4 | ENSMUST00000029353 | 28.4582133 | 33 | 19.1354467 | 19.1066282 | 2 | 0 | 0 | 0 | 0 | 0 | 0 | 0 | 0 | 0 | 0 | 0 | 1 | 1 | 4 |
| ENSMUSG00000030265 | Kras | ENSMUST00000032399 | 30.0844206 | 34 | 17.7794832 | 17.9585572 | 0 | 0 | 0 | 1 | 0 | 0 | 0 | 0 | 1 | 2 | 5 | 1 | 7 | 0 | 15 |
| ENSMUSG00000053012 | Krcc1 | ENSMUST00000168700 | 28.2415631 | 40 | 14.3872114 | 16.8738899 | 0 | 1 | 0 | 1 | 0 | 0 | 0 | 0 | 1 | 2 | 0 | 0 | 0 | 0 | 3 |
| ENSMUSG00000020393 | Kremen1 | ENSMUST00000020662 | 23.0520435 | 23 | 26.4039988 | 27.462511 | 0 | 0 | 0 | 0 | 0 | 0 | 0 | 0 | 0 | 0 | 0 | 1 | 0 | 0 | 1 |
| ENSMUSG00000000600 | Krit1 | ENSMUST00000171023 | 26.0636715 | 37 | 18.8634335 | 18.1791134 | 0 | 0 | 0 | 0 | 0 | 0 | 0 | 0 | 0 | 0 | 0 | 1 | 1 | 2 | 4 |
| ENSMUSG00000063334 | Krr1 | ENSMUST00000163048 | 30.4985337 | 29 | 19.034924 | 21.5942415 | 0 | 0 | 0 | 0 | 0 | 0 | 0 | 0 | 0 | 0 | 0 | 1 | 0 | 0 | 1 |
| ENSMUSG00000035849 | Krt222 | ENSMUST00000103132 | 29.8887859 | 31 | 18.9063948 | 20.1112141 | 0 | 0 | 0 | 0 | 0 | 0 | 0 | 0 | 0 | 0 | 0 | 5 | 4 | 2 | 11 |
| ENSMUSG00000062797 | l7Rn6 | ENSMUST00000153470 | 28.0196629 | 35 | 17.5561798 | 19.1713483 | 0 | 0 | 0 | 0 | 0 | 0 | 0 | 0 | 0 | 0 | 0 | 1 | 13 | 0 | 14 |
| ENSMUSG00000025937 | Lactb2 | ENSMUST00000027071 | 32.7102804 | 29 | 19.22968 | 19.2013594 | 0 | 0 | 0 | 0 | 0 | 0 | 0 | 0 | 0 | 0 | 0 | 0 | 6 | 0 | 6 |
| ENSMUSG00000015289 | Lage3 | ENSMUST00000114153 | 26.4115433 | 31 | 21.0790464 | 21.2672522 | 3 | 2 | 0 | 1 | 0 | 0 | 0 | 0 | 1 | 2 | 2 | 0 | 6 | 1 | 16 |
| ENSMUSG00000026478 | Lamc1 | ENSMUST00000027752 | 23.2849863 | 30 | 25.4410035 | 21.4033712 | 0 | 0 | 0 | 1 | 0 | 0 | 0 | 0 | 1 | 2 | 1 | 1 | 2 | 2 | 8 |
| ENSMUSG00000016534 | Lamp2 | ENSMUST00000061755 | 28.6621315 | 36 | 17.2789116 | 17.6870748 | 0 | 0 | 0 | 0 | 0 | 0 | 0 | 0 | 0 | 0 | 0 | 0 | 2 | 0 | 2 |
| ENSMUSG00000091512 | Lamtor3 | ENSMUST00000168345 | 27.114094 | 31 | 21.0738255 | 20.8053691 | 0 | 0 | 1 | 0 | 0 | 0 | 0 | 0 | 0 | 1 | 0 | 0 | 0 | 0 | 1 |
| ENSMUSG00000047344 | Lancl3 | ENSMUST00000069763 | 30.875 | 31 | 18.4166667 | 20.1666667 | 0 | 0 | 0 | 0 | 0 | 0 | 0 | 0 | 0 | 0 | 0 | 0 | 0 | 2 | 2 |
| ENSMUSG00000023025 | Larp4 | ENSMUST00000057632 | 30.6679435 | 36 | 14.9868327 | 18.5779267 | 1 | 0 | 0 | 1 | 0 | 0 | 0 | 0 | 1 | 2 | 2 | 0 | 1 | 1 | 7 |
| ENSMUSG00000033499 | Larp4b | ENSMUST00000091829 | 28.7261328 | 30 | 18.6377885 | 22.5420348 | 0 | 0 | 0 | 0 | 0 | 0 | 0 | 0 | 0 | 0 | 0 | 0 | 4 | 3 | 7 |
| ENSMUSG00000040021 | Lats1 | ENSMUST00000165952 | 29.4953596 | 34 | 17.2563805 | 19.1705336 | 0 | 0 | 0 | 0 | 0 | 0 | 0 | 0 | 0 | 0 | 0 | 3 | 10 | 1 | 14 |
| ENSMUSG00000024063 | Lbh | ENSMUST00000024857 | 20.1761409 | 29 | 25.5404323 | 25.060048 | 0 | 0 | 0 | 0 | 0 | 0 | 0 | 0 | 0 | 0 | 0 | 1 | 15 | 0 | 16 |
| ENSMUSG00000004880 | Lbr | ENSMUST00000005003 | 29.0216678 | 38 | 15.1674327 | 17.9908076 | 0 | 0 | 0 | 0 | 0 | 0 | 0 | 0 | 0 | 0 | 0 | 0 | 3 | 1 | 4 |
| ENSMUSG00000054469 | Lclat1 | ENSMUST00000067545 | 27.3824651 | 28 | 23.6340534 | 20.8386277 | 0 | 0 | 0 | 1 | 0 | 0 | 0 | 0 | 1 | 2 | 0 | 1 | 0 | 0 | 3 |
| ENSMUSG00000025019 | Lcor | ENSMUST00000067795 | 27.7478085 | 34 | 17.9028995 | 19.9595415 | 0 | 0 | 0 | 0 | 0 | 0 | 0 | 0 | 0 | 0 | 0 | 9 | 1 | 0 | 10 |
| ENSMUSG00000015882 | Lcorl | ENSMUST00000190036 | 34.4583333 | 32 | 17.75 | 16.2916667 | 2 | 1 | 0 | 1 | 0 | 0 | 0 | 0 | 1 | 2 | 0 | 0 | 0 | 0 | 5 |
| ENSMUSG00000039706 | Ldb2 | ENSMUST00000199534 | 28.9640592 | 30 | 20.4369274 | 20.5073996 | 0 | 0 | 0 | 0 | 0 | 0 | 0 | 0 | 0 | 0 | 0 | 0 | 2 | 0 | 2 |
| ENSMUSG00000048058 | Ldlrad3 | ENSMUST00000058790 | 26.1060329 | 32 | 21.9744059 | 20.3656307 | 0 | 0 | 0 | 0 | 0 | 0 | 0 | 0 | 0 | 0 | 3 | 0 | 6 | 1 | 10 |
| ENSMUSG00000074579 | Lekr1 | ENSMUST00000099075 | 27.6574307 | 39 | 16.2720403 | 16.675063 | 1 | 1 | 0 | 1 | 0 | 0 | 0 | 0 | 1 | 2 | 0 | 0 | 4 | 1 | 9 |
| ENSMUSG00000042487 | Leo1 | ENSMUST00000048937 | 34.3283582 | 40 | 8.95522388 | 16.4179104 | 1 | 0 | 1 | 0 | 0 | 0 | 0 | 0 | 0 | 1 | 0 | 0 | 0 | 0 | 2 |
| ENSMUSG00000031513 | Leprotl1 | ENSMUST00000033910 | 25.8139535 | 32 | 20.7906977 | 21.7209302 | 0 | 0 | 0 | 0 | 0 | 0 | 0 | 0 | 0 | 0 | 0 | 1 | 2 | 0 | 3 |
| ENSMUSG00000037353 | Letmd1 | ENSMUST00000037001 | 21.4082504 | 28 | 26.6714083 | 23.7553343 | 0 | 0 | 0 | 0 | 0 | 0 | 0 | 0 | 0 | 0 | 0 | 0 | 0 | 2 | 2 |
| ENSMUSG00000057554 | Lgals8 | ENSMUST00000124888 | 28.8111888 | 29 | 21.1188811 | 20.6993007 | 0 | 0 | 0 | 0 | 0 | 0 | 0 | 0 | 0 | 0 | 0 | 0 | 0 | 4 | 4 |
| ENSMUSG00000042363 | Lgalsl | ENSMUST00000047028 | 29.7925608 | 31 | 18.8125894 | 20.4577969 | 2 | 1 | 0 | 3 | 0 | 0 | 0 | 1 | 2 | 6 | 0 | 0 | 5 | 2 | 16 |
| ENSMUSG00000050199 | Lgr4 | ENSMUST00000046548 | 29.9421965 | 35 | 16.3583815 | 18.7861272 | 0 | 1 | 0 | 0 | 0 | 0 | 0 | 0 | 0 | 0 | 0 | 0 | 12 | 2 | 15 |
| ENSMUSG00000106379 | Lhfpl3 | ENSMUST00000197992 | 34.0453357 | 31 | 19.0496949 | 15.9546643 | 0 | 0 | 0 | 0 | 0 | 0 | 0 | 0 | 0 | 0 | 0 | 0 | 1 | 0 | 1 |
| ENSMUSG00000026890 | Lhx6 | ENSMUST00000112961 | 25.8801142 | 23 | 27.4024738 | 23.2635585 | 0 | 1 | 0 | 0 | 0 | 0 | 0 | 0 | 0 | 0 | 0 | 2 | 0 | 1 | 4 |
| ENSMUSG00000019230 | Lhx9 | ENSMUST00000019374 | 31.5455594 | 32 | 17.3298731 | 19.4348328 | 0 | 0 | 0 | 0 | 0 | 0 | 0 | 0 | 0 | 0 | 0 | 0 | 5 | 2 | 7 |
| ENSMUSG00000019920 | Lims1 | ENSMUST00000020078 | 31.2520013 | 33 | 18.699968 | 17.3230868 | 0 | 0 | 0 | 0 | 0 | 0 | 0 | 0 | 0 | 0 | 0 | 0 | 0 | 1 | 1 |
| ENSMUSG00000063804 | Lin28b | ENSMUST00000079390 | 28.9313836 | 33 | 17.7502812 | 20.8098988 | 0 | 0 | 0 | 0 | 0 | 0 | 0 | 0 | 0 | 0 | 1 | 0 | 10 | 0 | 11 |
| ENSMUSG00000085793 | Lin52; Gm7020 | ENSMUST00000137170 | 25.7829978 | 31 | 20.8612975 | 21.8680089 | 0 | 0 | 0 | 0 | 0 | 0 | 0 | 0 | 0 | 0 | 0 | 2 | 2 | 0 | 4 |
| ENSMUSG00000035310 | Lin54 | ENSMUST00000144030 | 31.2098009 | 29 | 19.3568147 | 20.6125574 | 0 | 0 | 0 | 0 | 0 | 0 | 0 | 0 | 0 | 0 | 0 | 0 | 0 | 2 | 2 |
| ENSMUSG00000019906 | Lin7a | ENSMUST00000020057 | 32.1047527 | 30 | 20.3491756 | 17.5557711 | 2 | 1 | 0 | 2 | 0 | 0 | 1 | 0 | 1 | 4 | 0 | 0 | 2 | 1 | 10 |
| ENSMUSG00000027162 | Lin7c | ENSMUST00000028583 | 28.2343143 | 36 | 16.2687396 | 19.1282621 | 0 | 0 | 0 | 0 | 0 | 0 | 0 | 0 | 0 | 0 | 0 | 0 | 0 | 2 | 2 |
| ENSMUSG00000058729 | Lin9 | ENSMUST00000193892 | 30.6314797 | 30 | 18.8972667 | 20.5466541 | 1 | 1 | 1 | 0 | 0 | 0 | 0 | 0 | 0 | 1 | 0 | 0 | 4 | 0 | 7 |
| ENSMUSG00000045083 | Lingo2 | ENSMUST00000164772 | 32.3626737 | 29 | 19.1925877 | 19.655857 | 0 | 0 | 1 | 0 | 0 | 0 | 0 | 0 | 0 | 1 | 0 | 0 | 0 | 0 | 1 |
| ENSMUSG00000053846 | Lipg | ENSMUST00000066532 | 23.4586466 | 28 | 23.6090226 | 24.9122807 | 0 | 0 | 0 | 0 | 0 | 0 | 0 | 0 | 0 | 0 | 0 | 2 | 1 | 2 | 5 |
| ENSMUSG00000047786 | Lix1 | ENSMUST00000115576 | 30.8425276 | 32 | 19.3580742 | 18.1544634 | 0 | 0 | 0 | 0 | 0 | 0 | 0 | 0 | 0 | 0 | 0 | 0 | 4 | 3 | 7 |
| ENSMUSG00000049288 | Lix1l | ENSMUST00000062058 | 24.3267108 | 32 | 21.3686534 | 22.4724062 | 0 | 0 | 0 | 0 | 0 | 0 | 0 | 0 | 0 | 0 | 1 | 2 | 1 | 0 | 4 |
| ENSMUSG00000041891 | Lman1 | ENSMUST00000048260 | 26.5254665 | 31 | 22.1886031 | 20.373172 | 0 | 0 | 0 | 0 | 0 | 0 | 0 | 0 | 0 | 0 | 0 | 0 | 2 | 0 | 2 |
| ENSMUSG00000010721 | Lmbr1 | ENSMUST00000055195 | 27.6908924 | 30 | 19.6565471 | 22.232444 | 0 | 0 | 0 | 1 | 0 | 0 | 0 | 0 | 1 | 2 | 0 | 0 | 0 | 1 | 3 |
| ENSMUSG00000073725 | Lmbrd1 | ENSMUST00000095062 | 29.0603086 | 30 | 20.5890603 | 20.1963534 | 0 | 0 | 0 | 1 | 0 | 0 | 0 | 0 | 1 | 2 | 1 | 0 | 0 | 0 | 3 |
| ENSMUSG00000039704 | Lmbrd2 | ENSMUST00000090380 | 30.6491885 | 36 | 16.4794007 | 16.7290886 | 1 | 0 | 0 | 1 | 0 | 0 | 0 | 0 | 1 | 2 | 0 | 0 | 1 | 1 | 5 |

| | | | | | | | | | | | | | | | | | | | | | |
|---|---|---|---|---|---|---|---|---|---|---|---|---|---|---|---|---|---|---|---|---|---|
| ENSMUSG00000022802 | Lmln | ENSMUST00000023497 | 27.4981586 | 27 | 21.5811441 | 24.2818561 | 2 | 1 | 0 | 3 | 0 | 0 | 0 | 1 | 2 | 6 | 0 | 0 | 0 | 0 | 9 |
| ENSMUSG00000030226 | Lmo3 | ENSMUST00000162772 | 27.5132275 | 32 | 20.5026455 | 20.1719577 | 0 | 0 | 0 | 0 | 0 | 0 | 0 | 0 | 0 | 0 | 0 | 0 | 1 | 0 | 1 |
| ENSMUSG00000028266 | Lmo4 | ENSMUST00000121796 | 32.2033898 | 29 | 18.1225554 | 21.1212516 | 0 | 0 | 1 | 0 | 0 | 0 | 0 | 0 | 0 | 1 | 0 | 0 | 0 | 0 | 1 |
| ENSMUSG00000009207 | Lnp | ENSMUST00000111993 | 33.2332332 | 35 | 13.2632633 | 18.2682683 | 1 | 1 | 0 | 1 | 0 | 0 | 0 | 0 | 1 | 2 | 0 | 0 | 4 | 0 | 8 |
| ENSMUSG00000023845 | Lnpep | ENSMUST00000041047 | 27.0977295 | 37 | 16.2882527 | 19.3978282 | 0 | 0 | 0 | 0 | 0 | 0 | 0 | 0 | 0 | 0 | 9 | 0 | 8 | 0 | 17 |
| ENSMUSG00000016520 | Lnx2 | ENSMUST00000016664 | 24.9338041 | 29 | 24.4924978 | 21.8446602 | 0 | 0 | 1 | 0 | 0 | 0 | 0 | 0 | 0 | 1 | 0 | 0 | 1 | 1 | 3 |
| ENSMUSG00000072980 | LOC100862454; LOC100862416; | ENSMUST00000123818 | 29.5977011 | 35 | 17.816092 | 17.816092 | 1 | 0 | 1 | 1 | 0 | 0 | 0 | 0 | 1 | 3 | 0 | 0 | 0 | 0 | 4 |
| ENSMUSG00000029430 | LOC102641332; Ran | ENSMUST00000031383 | 25.7142857 | 33 | 18.2059801 | 23.0564784 | 0 | 0 | 0 | 0 | 0 | 0 | 0 | 0 | 0 | 0 | 0 | 1 | 3 | 0 | 4 |
| ENSMUSG00000021282 | LOC102641436; Eif5 | ENSMUST00000166123 | 29.3208431 | 34 | 16.4871194 | 19.9063232 | 0 | 0 | 1 | 0 | 0 | 0 | 0 | 0 | 0 | 1 | 0 | 0 | 0 | 0 | 1 |
| ENSMUSG00000071072 | LOC102641464; Ptges3 | ENSMUST00000052798 | 26.7284992 | 37 | 17.0320405 | 19.7301855 | 0 | 0 | 0 | 0 | 0 | 0 | 0 | 0 | 0 | 0 | 0 | 0 | 16 | 2 | 18 |
| ENSMUSG00000027104 | LOC102641666; Atf2 | ENSMUST00000124737 | 28.4009547 | 36 | 16.5473349 | 19.4908512 | 0 | 0 | 0 | 0 | 0 | 0 | 0 | 0 | 0 | 0 | 0 | 0 | 4 | 0 | 4 |
| ENSMUSG00000015120 | LOC102641751; Ube2i | ENSMUST00000172868 | 27.2342676 | 26 | 19.7250132 | 27.2342676 | 0 | 0 | 0 | 0 | 0 | 0 | 0 | 0 | 0 | 0 | 0 | 0 | 1 | 0 | 1 |
| ENSMUSG00000044573 | LOC102642088; Acp1 | ENSMUST00000074038 | 29.0980392 | 30 | 20 | 21.2941176 | 0 | 0 | 0 | 0 | 0 | 0 | 0 | 0 | 0 | 0 | 0 | 0 | 1 | 0 | 1 |
| ENSMUSG00000060739 | LOC102642514; Nsa2 | ENSMUST00000073456 | 35.005767 | 29 | 17.9930796 | 17.9354095 | 0 | 0 | 0 | 0 | 0 | 0 | 0 | 0 | 0 | 0 | 0 | 0 | 0 | 3 | 3 |
| ENSMUSG00000071180 | LOC102642895; LOC102642197; | ENSMUST00000095458 | 32.6027397 | 32 | 16.0958904 | 19.3835616 | 0 | 0 | 0 | 0 | 0 | 0 | 0 | 0 | 0 | 0 | 0 | 0 | 1 | 0 | 1 |
| ENSMUSG00000026064 | LOC102643131; Ptp4a1 | ENSMUST00000027232 | 29.8898072 | 35 | 17.0454545 | 18.5261708 | 0 | 0 | 0 | 0 | 0 | 0 | 0 | 0 | 0 | 0 | 0 | 0 | 2 | 1 | 3 |
| ENSMUSG00000039633 | Lonrf1 | ENSMUST00000065297 | 23.1327049 | 34 | 17.4764322 | 25.235678 | 1 | 0 | 0 | 0 | 0 | 0 | 0 | 0 | 0 | 0 | 0 | 0 | 1 | 0 | 2 |
| ENSMUSG00000038668 | Lpar1 | ENSMUST00000055018 | 30.7507218 | 30 | 17.4205967 | 21.9923003 | 0 | 0 | 0 | 0 | 0 | 0 | 0 | 0 | 0 | 0 | 0 | 0 | 7 | 2 | 9 |
| ENSMUSG00000049929 | Lpar4 | ENSMUST00000060576 | 37.7329193 | 35 | 12.0729814 | 15.6055901 | 1 | 0 | 0 | 2 | 0 | 0 | 0 | 0 | 2 | 4 | 0 | 0 | 1 | 2 | 8 |
| ENSMUSG00000033446 | Lpar6 | ENSMUST00000044405 | 33.6430719 | 33 | 17.2998087 | 16.2612736 | 0 | 0 | 0 | 0 | 0 | 0 | 0 | 0 | 0 | 0 | 4 | 1 | 5 | 2 | 12 |
| ENSMUSG00000026623 | Lpgat1 | ENSMUST00000110855 | 28.7711938 | 30 | 19.2798462 | 21.8143681 | 1 | 0 | 0 | 1 | 0 | 0 | 0 | 0 | 1 | 2 | 1 | 0 | 1 | 2 | 7 |
| ENSMUSG00000033306 | Lpp | ENSMUST00000038053 | 28.2610336 | 31 | 21.4937759 | 19.4190871 | 1 | 0 | 0 | 1 | 0 | 0 | 0 | 0 | 1 | 2 | 0 | 4 | 10 | 2 | 19 |
| ENSMUSG00000028003 | Lrat | ENSMUST00000029632 | 30.5188894 | 30 | 18.2976787 | 20.9376422 | 1 | 0 | 0 | 0 | 0 | 0 | 0 | 0 | 0 | 0 | 0 | 0 | 5 | 0 | 6 |
| ENSMUSG00000068015 | Lrch1 | ENSMUST00000088970 | 28.3170835 | 30 | 19.3726155 | 22.8062738 | 1 | 0 | 0 | 1 | 0 | 0 | 0 | 0 | 1 | 2 | 0 | 0 | 1 | 0 | 4 |
| ENSMUSG00000031290 | Lrch2 | ENSMUST00000152838 | 33.2300542 | 38 | 13.8652208 | 15.3756778 | 0 | 0 | 0 | 2 | 0 | 0 | 0 | 0 | 2 | 4 | 0 | 0 | 1 | 1 | 6 |
| ENSMUSG00000022801 | Lrch3 | ENSMUST00000023491 | 27.2410516 | 27 | 23.3449477 | 22.1095977 | 0 | 0 | 0 | 0 | 0 | 0 | 0 | 0 | 0 | 0 | 1 | 2 | 0 | 0 | 3 |
| ENSMUSG00000032913 | Lrig2 | ENSMUST00000046316 | 24.6235139 | 33 | 20.6340819 | 21.8229855 | 0 | 0 | 0 | 0 | 0 | 0 | 0 | 0 | 0 | 0 | 0 | 3 | 8 | 1 | 12 |
| ENSMUSG00000019796 | Lrp11 | ENSMUST00000130590 | 27.3224044 | 30 | 20.6557377 | 22.1857923 | 0 | 0 | 0 | 0 | 0 | 0 | 0 | 0 | 0 | 0 | 0 | 1 | 0 | 0 | 1 |
| ENSMUSG00000022305 | Lrp12 | ENSMUST00000022916 | 26.4164306 | 37 | 16.9971671 | 19.7592068 | 0 | 0 | 0 | 0 | 0 | 0 | 0 | 0 | 0 | 0 | 1 | 0 | 2 | 1 | 4 |
| ENSMUSG00000030201 | Lrp6 | ENSMUST00000032322 | 27.8886311 | 32 | 18.3526682 | 22.25058 | 0 | 1 | 1 | 0 | 0 | 0 | 0 | 0 | 0 | 1 | 4 | 0 | 9 | 0 | 15 |
| ENSMUSG00000034883 | Lrr1 | ENSMUST00000110621 | 35.2631579 | 40 | 7.89473684 | 16.8421053 | 0 | 0 | 0 | 1 | 0 | 0 | 0 | 0 | 1 | 2 | 0 | 0 | 0 | 0 | 2 |
| ENSMUSG00000027961 | Lrrc39 | ENSMUST00000029573 | 27.7447269 | 32 | 22.5527312 | 17.7934018 | 3 | 1 | 1 | 5 | 0 | 0 | 0 | 1 | 4 | 11 | 1 | 0 | 0 | 0 | 16 |
| ENSMUSG00000063052 | Lrrc40 | ENSMUST00000150525 | 29.1538799 | 30 | 19.9035511 | 20.7803595 | 0 | 0 | 0 | 0 | 0 | 0 | 0 | 0 | 0 | 0 | 0 | 0 | 0 | 1 | 1 |
| ENSMUSG00000027286 | Lrrc57 | ENSMUST00000102496 | 29.3156282 | 33 | 19.2032686 | 18.6925434 | 0 | 0 | 0 | 0 | 0 | 0 | 0 | 0 | 0 | 0 | 0 | 0 | 1 | 0 | 1 |
| ENSMUSG00000034158 | Lrrc58 | ENSMUST00000078717 | 27.5689562 | 32 | 19.4429421 | 20.8085452 | 0 | 0 | 0 | 1 | 0 | 0 | 0 | 0 | 1 | 2 | 1 | 0 | 10 | 1 | 14 |
| ENSMUSG00000054720 | Lrrc8c | ENSMUST00000067924 | 26.9665568 | 32 | 22.0913801 | 19.2416392 | 0 | 0 | 0 | 0 | 0 | 0 | 0 | 0 | 0 | 0 | 3 | 29 | 10 | 0 | 42 |
| ENSMUSG00000027550 | Lrrcc1 | ENSMUST00000169079 | 29.4766966 | 34 | 17.2935405 | 19.5421096 | 0 | 0 | 0 | 0 | 0 | 0 | 0 | 0 | 0 | 0 | 1 | 0 | 0 | 0 | 1 |
| ENSMUSG00000071862 | Lrrtm2 | ENSMUST00000091636 | 33.9658444 | 31 | 18.2163188 | 16.644077 | 0 | 0 | 0 | 0 | 0 | 0 | 0 | 0 | 0 | 0 | 0 | 1 | 2 | 0 | 3 |
| ENSMUSG00000052581 | Lrrtm4 | ENSMUST00000136421 | 31.4983595 | 36 | 14.7283996 | 17.8636529 | 1 | 0 | 0 | 0 | 0 | 0 | 0 | 0 | 0 | 0 | 6 | 1 | 21 | 0 | 29 |
| ENSMUSG00000061080 | Lsamp | ENSMUST00000099761 | 29.0036305 | 29 | 21.0434315 | 20.6803819 | 0 | 0 | 0 | 1 | 0 | 0 | 0 | 0 | 1 | 2 | 1 | 1 | 1 | 3 | 8 |
| ENSMUSG00000037296 | Lsm1 | ENSMUST00000038421 | 26.584976 | 31 | 22.1630261 | 19.9786894 | 0 | 0 | 0 | 0 | 0 | 0 | 0 | 0 | 0 | 0 | 0 | 2 | 1 | 2 | 5 |
| ENSMUSG00000066568 | Lsm14a | ENSMUST00000085585 | 28.5289747 | 34 | 16.2704309 | 20.8023774 | 0 | 0 | 0 | 0 | 0 | 0 | 0 | 0 | 0 | 0 | 0 | 0 | 1 | 0 | 1 |
| ENSMUSG00000031683 | Lsm6 | ENSMUST00000130325 | 25.3735804 | 30 | 21.3687986 | 23.6401674 | 0 | 0 | 0 | 0 | 0 | 0 | 0 | 0 | 0 | 0 | 3 | 2 | 4 | 0 | 9 |
| ENSMUSG00000071342 | Lsmem1 | ENSMUST00000095760 | 25.5617978 | 29 | 20.505618 | 25.2808989 | 0 | 0 | 0 | 1 | 0 | 0 | 0 | 0 | 1 | 2 | 0 | 0 | 0 | 0 | 2 |
| ENSMUSG00000033105 | Lss | ENSMUST00000048678 | 23.3098299 | 28 | 24.2223144 | 24.927416 | 0 | 0 | 0 | 1 | 0 | 0 | 0 | 0 | 1 | 2 | 0 | 1 | 0 | 0 | 3 |
| ENSMUSG00000052299 | Ltn1 | ENSMUST00000039449 | 30.8631212 | 34 | 16.5213601 | 18.657367 | 2 | 1 | 0 | 2 | 0 | 0 | 0 | 0 | 2 | 4 | 1 | 0 | 0 | 0 | 8 |
| ENSMUSG00000024188 | Luc7l | ENSMUST00000114976 | 24.3980738 | 30 | 21.1610487 | 24.023542 | 0 | 0 | 0 | 0 | 0 | 0 | 0 | 0 | 0 | 0 | 5 | 0 | 8 | 0 | 13 |
| ENSMUSG00000029823 | Luc7l2 | ENSMUST00000161538 | 29.1147994 | 35 | 15.3181189 | 20.4011065 | 0 | 0 | 0 | 1 | 0 | 0 | 0 | 0 | 1 | 2 | 1 | 0 | 1 | 0 | 4 |
| ENSMUSG00000020863 | Luc7l3 | ENSMUST00000166312 | 27.5824176 | 33 | 18.021978 | 21.7307692 | 0 | 0 | 0 | 0 | 0 | 0 | 0 | 0 | 0 | 0 | 2 | 1 | 0 | 0 | 3 |
| ENSMUSG00000063297 | Luzp2 | ENSMUST00000082373 | 32.1113953 | 34 | 16.8513782 | 16.9650469 | 0 | 0 | 0 | 0 | 0 | 0 | 0 | 0 | 0 | 0 | 2 | 0 | 2 | 2 | 6 |
| ENSMUSG00000050447 | Lypd6 | ENSMUST00000112712 | 29.0334302 | 31 | 19.6947674 | 20.494186 | 1 | 0 | 0 | 0 | 0 | 0 | 0 | 0 | 0 | 0 | 0 | 1 | 1 | 1 | 4 |
| ENSMUSG00000025903 | Lypla1 | ENSMUST00000027036 | 32.8496808 | 35 | 15.960534 | 16.2507255 | 0 | 0 | 0 | 2 | 0 | 0 | 0 | 0 | 2 | 4 | 0 | 0 | 0 | 0 | 4 |
| ENSMUSG00000020268 | Lyrm7 | ENSMUST00000141703 | 28.8584475 | 29 | 21.2328767 | 20.6849315 | 0 | 0 | 0 | 0 | 0 | 0 | 0 | 0 | 0 | 0 | 1 | 13 | 6 | 0 | 20 |
| ENSMUSG00000025245 | Lztfl1 | ENSMUST00000166097 | 33.0810811 | 31 | 17.972973 | 17.8918919 | 0 | 0 | 0 | 0 | 0 | 0 | 0 | 0 | 0 | 0 | 0 | 0 | 1 | 0 | 1 |
| ENSMUSG00000056947 | Mab21l1 | ENSMUST00000075422 | 33.512931 | 32 | 17.3491379 | 16.8103448 | 1 | 0 | 0 | 1 | 0 | 0 | 0 | 0 | 1 | 2 | 0 | 0 | 0 | 0 | 3 |
| ENSMUSG00000068205 | Macrod2 | ENSMUST00000110064 | 29.8245614 | 30 | 19.9043062 | 20.6060606 | 0 | 0 | 0 | 0 | 0 | 0 | 0 | 0 | 0 | 0 | 0 | 2 | 1 | 0 | 3 |
| ENSMUSG00000029910 | Mad2l1 | ENSMUST00000116605 | 30.2411056 | 30 | 18.9356072 | 20.4351661 | 0 | 0 | 1 | 0 | 0 | 0 | 0 | 0 | 0 | 1 | 0 | 1 | 2 | 1 | 5 |
| ENSMUSG00000055435 | Maf | ENSMUST00000109104 | 30.699722 | 31 | 19.2307692 | 18.7905468 | 0 | 0 | 0 | 0 | 0 | 0 | 0 | 0 | 0 | 0 | 2 | 0 | 6 | 3 | 11 |
| ENSMUSG00000028609 | Magoh | ENSMUST00000030348 | 29.7297297 | 43 | 9.45945946 | 18.2432432 | 0 | 0 | 0 | 0 | 0 | 0 | 0 | 0 | 0 | 0 | 0 | 0 | 1 | 0 | 1 |
| ENSMUSG00000031232 | Magt1 | ENSMUST00000151689 | 30.8284024 | 31 | 18.1952663 | 20.2662722 | 0 | 0 | 0 | 0 | 0 | 0 | 0 | 0 | 0 | 0 | 0 | 0 | 1 | 3 | 4 |
| ENSMUSG00000024479 | Mal2 | ENSMUST00000025356 | 30.7984791 | 33 | 18.4410646 | 17.9657795 | 2 | 0 | 0 | 1 | 0 | 0 | 0 | 0 | 1 | 2 | 0 | 0 | 2 | 0 | 6 |
| ENSMUSG00000031925 | Maml2 | ENSMUST00000177755 | 29.8395357 | 31 | 19.3581427 | 19.3240014 | 1 | 0 | 0 | 0 | 0 | 0 | 0 | 0 | 0 | 0 | 0 | 3 | 12 | 1 | 17 |
| ENSMUSG00000003746 | Man1a | ENSMUST00000003843 | 30.4643262 | 35 | 19.4790487 | 15.3265383 | 0 | 0 | 0 | 0 | 0 | 0 | 0 | 0 | 0 | 0 | 0 | 2 | 0 | 0 | 2 |
| ENSMUSG00000024085 | Man2a1 | ENSMUST00000086723 | 29.4720252 | 34 | 18.0851064 | 18.9125296 | 0 | 0 | 0 | 0 | 0 | 0 | 0 | 0 | 0 | 0 | 1 | 2 | 4 | 1 | 8 |
| ENSMUSG00000040520 | Manea | ENSMUST00000041374 | 32.782194 | 33 | 17.2655008 | 16.7567568 | 0 | 0 | 0 | 1 | 0 | 0 | 0 | 0 | 1 | 2 | 0 | 0 | 0 | 1 | 3 |
| ENSMUSG00000025037 | Maoa | ENSMUST00000026013 | 30.1268932 | 34 | 16.7826443 | 19.0749079 | 0 | 0 | 0 | 0 | 0 | 0 | 0 | 0 | 0 | 0 | 0 | 0 | 1 | 0 | 1 |
| ENSMUSG00000015222 | Map2 | ENSMUST00000077355 | 31.51566 | 32 | 17.2259508 | 18.8758389 | 1 | 0 | 0 | 1 | 0 | 0 | 0 | 0 | 1 | 2 | 0 | 1 | 3 | 3 | 10 |
| ENSMUSG00000004936 | Map2k1 | ENSMUST00000005066 | 25.7923497 | 34 | 19.6721311 | 20.5464481 | 1 | 0 | 0 | 0 | 0 | 0 | 0 | 0 | 0 | 0 | 0 | 0 | 0 | 0 | 1 |
| ENSMUSG00000033352 | Map2k4 | ENSMUST00000046963 | 26.6748366 | 29 | 22.7941176 | 21.119281 | 0 | 0 | 0 | 0 | 0 | 0 | 0 | 0 | 0 | 0 | 0 | 1 | 0 | 0 | 1 |
| ENSMUSG00000020623 | Map2k6 | ENSMUST00000100260 | 24.8366013 | 32 | 22.3311547 | 21.1328976 | 0 | 0 | 0 | 0 | 0 | 0 | 0 | 0 | 0 | 0 | 0 | 0 | 1 | 0 | 1 |
| ENSMUSG00000002948 | Map2k7 | ENSMUST00000062686 | 20.212766 | 25 | 27.9497099 | 27.1760155 | 0 | 0 | 0 | 0 | 0 | 0 | 0 | 0 | 0 | 0 | 0 | 6 | 0 | 0 | 6 |
| ENSMUSG00000021754 | Map3k1 | ENSMUST00000109267 | 28.236246 | 34 | 18.3252427 | 19.0938511 | 1 | 0 | 0 | 1 | 0 | 0 | 0 | 0 | 1 | 2 | 0 | 2 | 26 | 0 | 31 |
| ENSMUSG00000031303 | Map3k15 | ENSMUST00000033665 | 33.5227273 | 30 | 17.0454545 | 19.6022727 | 0 | 0 | 0 | 1 | 0 | 0 | 0 | 0 | 1 | 2 | 0 | 0 | 0 | 0 | 2 |
| ENSMUSG00000024383 | Map3k2 | ENSMUST00000096575 | 29.2514578 | 34 | 16.8868261 | 19.7072474 | 0 | 0 | 0 | 0 | 0 | 0 | 0 | 0 | 0 | 0 | 3 | 2 | 2 | 4 | 11 |
| ENSMUSG00000028284 | Map3k7 | ENSMUST00000108184 | 28.113879 | 30 | 20.8185053 | 20.9456024 | 1 | 0 | 0 | 0 | 0 | 0 | 0 | 0 | 0 | 0 | 0 | 1 | 0 | 0 | 2 |
| ENSMUSG00000032479 | Map4 | ENSMUST00000169851 | 27.19357 | 35 | 18.8546551 | 19.0890824 | 2 | 1 | 0 | 3 | 0 | 0 | 0 | 0 | 3 | 6 | 8 | 1 | 10 | 1 | 29 |
| ENSMUSG00000024242 | Map4k3 | ENSMUST00000025089 | 29.7155689 | 35 | 15.494012 | 19.3862275 | 0 | 0 | 1 | 0 | 0 | 0 | 0 | 0 | 0 | 1 | 0 | 0 | 0 | 0 | 1 |
| ENSMUSG00000034761 | Map4k5 | ENSMUST00000110567 | 30.9775429 | 34 | 15.1254954 | 20.2774108 | 0 | 0 | 0 | 0 | 0 | 0 | 0 | 0 | 0 | 0 | 0 | 0 | 2 | 0 | 2 |
| ENSMUSG00000041205 | Map6d1 | ENSMUST00000040880 | 25.4524887 | 27 | 25.9426848 | 21.7571644 | 2 | 1 | 2 | 4 | 0 | 0 | 0 | 1 | 3 | 10 | 0 | 0 | 0 | 0 | 13 |
| ENSMUSG00000041020 | Map7d2 | ENSMUST00000112471 | 26.8557131 | 34 | 19.0158465 | 19.6830692 | 0 | 0 | 0 | 0 | 0 | 0 | 0 | 0 | 0 | 0 | 0 | 2 | 1 | 0 | 3 |
| ENSMUSG00000033900 | Map9 | ENSMUST00000091014 | 28.1234445 | 31 | 19.2882031 | 21.2294674 | 6 | 3 | 0 | 7 | 0 | 0 | 1 | 3 | 3 | 14 | 0 | 3 | 0 | 0 | 26 |
| ENSMUSG00000063358 | Mapk1 | ENSMUST00000069107 | 25.8141382 | 32 | 20.9954991 | 20.8895949 | 0 | 0 | 0 | 0 | 0 | 0 | 0 | 0 | 0 | 0 | 0 | 2 | 1 | 1 | 4 |
| ENSMUSG00000046709 | Mapk10 | ENSMUST00000170792 | 27.5149031 | 28 | 22.6900149 | 21.7958271 | 0 | 0 | 0 | 0 | 0 | 0 | 0 | 0 | 0 | 0 | 0 | 5 | 1 | 2 | 8 |
| ENSMUSG00000021840 | Mapk1ip1l | ENSMUST00000164235 | 26.1042052 | 32 | 19.0690294 | 22.6924094 | 0 | 0 | 2 | 0 | 0 | 0 | 0 | 0 | 0 | 2 | 0 | 1 | 13 | 0 | 16 |
| ENSMUSG00000042688 | Mapk6 | ENSMUST00000168937 | 28.8695652 | 35 | 19.0434783 | 17.0434783 | 1 | 0 | 0 | 0 | 0 | 0 | 0 | 0 | 0 | 0 | 0 | 1 | 0 | 2 | 4 |
| ENSMUSG00000021936 | Mapk8 | ENSMUST00000111945 | 27.7097506 | 34 | 18.7755102 | 19.5918367 | 0 | 0 | 0 | 1 | 0 | 0 | 0 | 0 | 1 | 2 | 1 | 0 | 2 | 0 | 5 |
| ENSMUSG00000027479 | Mapre1 | ENSMUST00000028981 | 24.8902821 | 33 | 20.3605016 | 21.8808777 | 1 | 0 | 0 | 1 | 0 | 0 | 0 | 0 | 1 | 2 | 0 | 3 | 4 | 1 | 11 |
| ENSMUSG00000024277 | Mapre2 | ENSMUST00000170802 | 24.778157 | 29 | 25.3924915 | 20.4095563 | 0 | 0 | 0 | 0 | 0 | 0 | 0 | 0 | 0 | 0 | 0 | 3 | 1 | 0 | 4 |
| ENSMUSG00000036469 | March1 | ENSMUST00000110255 | 32.8934664 | 31 | 17.3801094 | 18.925008 | 0 | 0 | 0 | 1 | 0 | 0 | 0 | 0 | 1 | 2 | 0 | 0 | 5 | 1 | 8 |
| ENSMUSG00000023307 | March5 | ENSMUST00000024078 | 28.2780411 | 32 | 18.7993681 | 20.8530806 | 0 | 0 | 0 | 0 | 0 | 0 | 0 | 0 | 0 | 0 | 0 | 0 | 1 | 0 | 1 |
| ENSMUSG00000039100 | March6 | ENSMUST00000090227 | 27.3860182 | 33 | 18.8449848 | 21.0334347 | 0 | 0 | 0 | 2 | 0 | 0 | 0 | 0 | 2 | 4 | 0 | 0 | 0 | 0 | 4 |
| ENSMUSG00000026977 | March7 | ENSMUST00000102748 | 34.0245776 | 36 | 13.4408602 | 16.2058372 | 1 | 0 | 0 | 0 | 0 | 0 | 0 | 0 | 0 | 0 | 0 | 0 | 0 | 1 | 2 |

| | | | | | | | | | | | | | | | | | | | | |
|---|---|---|---|---|---|---|---|---|---|---|---|---|---|---|---|---|---|---|---|---|
| ENSMUSG00000069662 | Marcks | ENSMUST00000092584 | 29.4548116 | 36 | 15.9897548 | 18.3315038 | 0 | 0 | 0 | 0 | 0 | 0 | 0 | 0 | 0 | 0 | 1 | 8 | 8 | 3 | 20 |
| ENSMUSG00000046994 | Mars2 | ENSMUST00000061334 | 31.1510032 | 34 | 15.8817318 | 19.4086589 | 1 | 0 | 0 | 2 | 0 | 0 | 0 | 0 | 2 | 4 | 2 | 4 | 3 | 1 | 15 |
| ENSMUSG00000021636 | Marveld2 | ENSMUST00000022137 | 25.7507508 | 28 | 22.5975976 | 23.4234234 | 0 | 0 | 0 | 0 | 0 | 0 | 0 | 0 | 0 | 0 | 0 | 0 | 3 | 0 | 3 |
| ENSMUSG00000026779 | Mastl | ENSMUST00000028119 | 31.3538353 | 31 | 17.3559562 | 19.7949806 | 0 | 0 | 1 | 1 | 0 | 0 | 0 | 0 | 1 | 3 | 2 | 0 | 0 | 0 | 5 |
| ENSMUSG00000053907 | Mat2a | ENSMUST00000205335 | 28.6960514 | 31 | 19.651056 | 20.7529844 | 0 | 0 | 1 | 0 | 0 | 0 | 0 | 0 | 0 | 1 | 0 | 0 | 0 | 0 | 1 |
| ENSMUSG00000042032 | Mat2b | ENSMUST00000040167 | 26.6197183 | 38 | 15.0704225 | 20.2816901 | 0 | 0 | 0 | 0 | 0 | 0 | 0 | 0 | 0 | 0 | 0 | 2 | 3 | 0 | 5 |
| ENSMUSG00000059436 | Max | ENSMUST00000082136 | 22.8637413 | 31 | 24.0184758 | 22.0939184 | 0 | 0 | 0 | 0 | 0 | 0 | 0 | 0 | 0 | 0 | 0 | 1 | 0 | 0 | 1 |
| ENSMUSG00000047508 | Mbd3l2 | ENSMUST00000051008 | 28.0321285 | 36 | 16.626506 | 19.7590361 | 0 | 0 | 0 | 1 | 0 | 0 | 0 | 0 | 1 | 2 | 0 | 0 | 2 | 0 | 4 |
| ENSMUSG00000030322 | Mbd4 | ENSMUST00000032469 | 30.0894855 | 30 | 20.4697987 | 19.4071588 | 0 | 0 | 0 | 0 | 0 | 0 | 0 | 0 | 0 | 0 | 0 | 0 | 0 | 1 | 1 |
| ENSMUSG00000021028 | Mbip | ENSMUST00000021416 | 32.1336761 | 37 | 14.3958869 | 16.4524422 | 0 | 0 | 1 | 0 | 0 | 0 | 0 | 0 | 0 | 1 | 0 | 0 | 0 | 0 | 1 |
| ENSMUSG00000051098 | Mblac2 | ENSMUST00000057598 | 33.0488221 | 34 | 16.1829976 | 16.7975418 | 1 | 0 | 0 | 2 | 0 | 0 | 0 | 0 | 2 | 4 | 0 | 0 | 0 | 0 | 5 |
| ENSMUSG00000027763 | Mbnl1 | ENSMUST00000194069 | 32.9307806 | 34 | 15.7879234 | 17.3195876 | 1 | 0 | 0 | 1 | 0 | 0 | 0 | 0 | 1 | 2 | 0 | 0 | 0 | 0 | 3 |
| ENSMUSG00000022139 | Mbnl2 | ENSMUST00000088419 | 32.7147016 | 34 | 16.8850073 | 15.9388646 | 1 | 1 | 0 | 3 | 0 | 0 | 0 | 1 | 2 | 6 | 1 | 0 | 0 | 2 | 11 |
| ENSMUSG00000036109 | Mbnl3 | ENSMUST00000114875 | 30.0160514 | 33 | 17.1214553 | 19.6896736 | 0 | 0 | 0 | 0 | 0 | 0 | 0 | 0 | 0 | 0 | 0 | 1 | 0 | 0 | 1 |
| ENSMUSG00000038732 | Mboat1 | ENSMUST00000047311 | 29.0064103 | 29 | 20.1121795 | 22.3557692 | 0 | 0 | 0 | 1 | 0 | 0 | 0 | 0 | 1 | 2 | 0 | 0 | 1 | 0 | 3 |
| ENSMUSG00000020646 | Mboat2 | ENSMUST00000078902 | 29.3602103 | 30 | 17.4408414 | 23.4881683 | 0 | 0 | 0 | 0 | 0 | 0 | 0 | 0 | 0 | 0 | 0 | 0 | 0 | 1 | 1 |
| ENSMUSG00000059474 | Mbtd1 | ENSMUST00000107854 | 30.4980595 | 33 | 17.2380336 | 19.6959897 | 0 | 0 | 0 | 0 | 0 | 0 | 0 | 0 | 0 | 0 | 2 | 0 | 2 | 0 | 4 |
| ENSMUSG00000046873 | Mbtps2 | ENSMUST00000112522 | 27.7954257 | 33 | 18.1385006 | 21.1245235 | 0 | 0 | 0 | 0 | 0 | 0 | 0 | 0 | 0 | 0 | 1 | 0 | 5 | 0 | 6 |
| ENSMUSG00000047259 | Mc4r | ENSMUST00000057942 | 30.1431801 | 31 | 18.3119819 | 20.6480784 | 0 | 0 | 0 | 0 | 0 | 0 | 0 | 0 | 0 | 0 | 1 | 0 | 0 | 0 | 1 |
| ENSMUSG00000032135 | Mcam | ENSMUST00000098852 | 22.6332971 | 29 | 23.5038085 | 24.8095756 | 0 | 0 | 0 | 0 | 0 | 0 | 0 | 0 | 0 | 0 | 0 | 0 | 2 | 0 | 2 |
| ENSMUSG00000027709 | Mccc1 | ENSMUST00000199113 | 26.5373699 | 26 | 22.2800378 | 25.4493851 | 0 | 0 | 0 | 0 | 0 | 0 | 0 | 0 | 0 | 0 | 0 | 2 | 3 | 1 | 6 |
| ENSMUSG00000058298 | Mcm9 | ENSMUST00000075540 | 23.6180905 | 34 | 19.8994975 | 22.5125628 | 0 | 0 | 0 | 0 | 0 | 0 | 0 | 0 | 0 | 0 | 0 | 0 | 1 | 0 | 1 |
| ENSMUSG00000048170 | Mcmbp | ENSMUST00000057557 | 28.5571643 | 32 | 20.4692961 | 18.7219171 | 0 | 0 | 0 | 0 | 0 | 0 | 0 | 0 | 0 | 0 | 0 | 0 | 18 | 0 | 18 |
| ENSMUSG00000046101 | Mcmdc2 | ENSMUST00000052843 | 25.1038576 | 32 | 22.1364985 | 20.7121662 | 0 | 0 | 0 | 1 | 0 | 0 | 0 | 0 | 1 | 2 | 4 | 0 | 3 | 0 | 9 |
| ENSMUSG00000041390 | Mdfic | ENSMUST00000125326 | 29.3530499 | 33 | 19.297597 | 18.7430684 | 1 | 1 | 0 | 0 | 0 | 0 | 0 | 0 | 0 | 0 | 0 | 0 | 0 | 1 | 3 |
| ENSMUSG00000034912 | Mdga2 | ENSMUST00000101379 | 30.7980517 | 26 | 20.3446984 | 23.0798052 | 0 | 0 | 0 | 0 | 0 | 0 | 0 | 0 | 0 | 0 | 1 | 0 | 0 | 0 | 1 |
| ENSMUSG00000020212 | Mdm1 | ENSMUST00000020437 | 29.9760192 | 36 | 15.4676259 | 18.5851319 | 0 | 0 | 1 | 0 | 0 | 0 | 0 | 0 | 0 | 1 | 0 | 0 | 0 | 0 | 1 |
| ENSMUSG00000020184 | Mdm2 | ENSMUST00000155285 | 29.7407913 | 27 | 20.1000455 | 23.1014097 | 1 | 1 | 0 | 1 | 0 | 0 | 0 | 0 | 1 | 2 | 0 | 0 | 1 | 0 | 5 |
| ENSMUSG00000054387 | Mdm4 | ENSMUST00000188090 | 25.4047093 | 32 | 20.9529065 | 21.7807211 | 0 | 0 | 0 | 0 | 0 | 0 | 0 | 0 | 0 | 0 | 2 | 2 | 4 | 0 | 8 |
| ENSMUSG00000021598 | Med10 | ENSMUST00000022089 | 23.7037037 | 30 | 24.1975309 | 22.4691358 | 0 | 0 | 0 | 0 | 0 | 0 | 0 | 0 | 0 | 0 | 0 | 1 | 0 | 0 | 1 |
| ENSMUSG00000056476 | Med12l | ENSMUST00000199659 | 28.0701754 | 34 | 17.7218408 | 20.4169845 | 1 | 1 | 1 | 0 | 0 | 0 | 0 | 0 | 0 | 1 | 0 | 2 | 5 | 0 | 10 |
| ENSMUSG00000064127 | Med14 | ENSMUST00000096495 | 28.9271163 | 38 | 16.0253508 | 16.8854685 | 0 | 0 | 0 | 0 | 0 | 0 | 0 | 0 | 0 | 0 | 0 | 0 | 4 | 0 | 4 |
| ENSMUSG00000031935 | Med17 | ENSMUST00000034411 | 27.36 | 29 | 19.7866667 | 23.68 | 0 | 0 | 0 | 0 | 0 | 0 | 0 | 0 | 0 | 0 | 0 | 0 | 3 | 3 | 6 |
| ENSMUSG00000019984 | Med23 | ENSMUST00000177232 | 25.5250404 | 28 | 24.1518578 | 22.4286484 | 0 | 0 | 0 | 0 | 0 | 0 | 0 | 0 | 0 | 0 | 3 | 2 | 3 | 0 | 8 |
| ENSMUSG00000015804 | Med28 | ENSMUST00000156481 | 24.936322 | 31 | 22.4910851 | 21.3958227 | 1 | 0 | 0 | 0 | 0 | 0 | 0 | 0 | 0 | 0 | 2 | 1 | 1 | 0 | 5 |
| ENSMUSG00000038622 | Med30 | ENSMUST00000037115 | 31.2267658 | 41 | 14.1263941 | 13.3828996 | 0 | 0 | 1 | 0 | 0 | 0 | 0 | 0 | 0 | 1 | 0 | 0 | 0 | 1 | 2 |
| ENSMUSG00000022109 | Med4 | ENSMUST00000022705 | 28.3924843 | 24 | 20.0417537 | 27.1398747 | 0 | 0 | 0 | 0 | 0 | 0 | 0 | 0 | 0 | 0 | 1 | 0 | 0 | 0 | 1 |
| ENSMUSG00000020397 | Med7 | ENSMUST00000109232 | 31.4583333 | 34 | 18.2638889 | 16.5277778 | 0 | 0 | 0 | 0 | 0 | 0 | 0 | 0 | 0 | 0 | 0 | 1 | 0 | 0 | 1 |
| ENSMUSG00000030557 | Mef2a | ENSMUST00000156690 | 30.6072874 | 32 | 18.1916329 | 19.3252362 | 1 | 0 | 0 | 1 | 0 | 0 | 0 | 0 | 1 | 2 | 0 | 1 | 0 | 0 | 4 |
| ENSMUSG00000005583 | Mef2c | ENSMUST00000197146 | 29.8494044 | 33 | 17.8242302 | 19.2178018 | 0 | 0 | 0 | 0 | 0 | 0 | 0 | 0 | 0 | 0 | 5 | 0 | 6 | 0 | 11 |
| ENSMUSG00000001419 | Mef2d | ENSMUST00000001455 | 21.5800174 | 27 | 26.1690386 | 24.9491722 | 0 | 0 | 0 | 0 | 0 | 0 | 0 | 0 | 0 | 0 | 1 | 2 | 1 | 0 | 4 |
| ENSMUSG00000039270 | Megf9 | ENSMUST00000107359 | 30.2909393 | 32 | 19.201995 | 18.8694929 | 0 | 0 | 0 | 0 | 0 | 0 | 0 | 0 | 0 | 0 | 0 | 0 | 1 | 1 | 2 |
| ENSMUSG00000051455 | Meioc | ENSMUST00000100378 | 33.8451268 | 35 | 13.6849132 | 17.0894526 | 0 | 0 | 0 | 2 | 0 | 0 | 0 | 0 | 2 | 4 | 0 | 0 | 0 | 2 | 6 |
| ENSMUSG00000020160 | Meis1 | ENSMUST00000144988 | 28.3101811 | 31 | 21.4524354 | 19.0786003 | 0 | 0 | 0 | 0 | 0 | 0 | 0 | 0 | 0 | 0 | 2 | 30 | 7 | 1 | 40 |
| ENSMUSG00000005813 | Metap1 | ENSMUST00000197531 | 25.8610086 | 37 | 16.8819188 | 19.8339483 | 1 | 0 | 0 | 0 | 0 | 0 | 0 | 0 | 0 | 0 | 1 | 1 | 0 | 1 | 4 |
| ENSMUSG00000036112 | Metap2 | ENSMUST00000047910 | 28.4515904 | 33 | 18.6997553 | 19.6434813 | 0 | 0 | 0 | 0 | 0 | 0 | 0 | 0 | 0 | 0 | 0 | 0 | 13 | 2 | 15 |
| ENSMUSG00000028114 | Mettl14 | ENSMUST00000090371 | 26.2618419 | 33 | 19.7546203 | 20.6709116 | 0 | 0 | 0 | 0 | 0 | 0 | 0 | 0 | 0 | 0 | 0 | 1 | 2 | 0 | 3 |
| ENSMUSG00000010554 | Mettl16 | ENSMUST00000141755 | 25.9868116 | 30 | 20.5442556 | 23.2005201 | 3 | 1 | 0 | 3 | 0 | 0 | 0 | 1 | 2 | 6 | 6 | 4 | 13 | 3 | 36 |
| ENSMUSG00000025956 | Mettl21a | ENSMUST00000114079 | 24.5461721 | 34 | 19.1791634 | 22.1783741 | 0 | 0 | 0 | 0 | 0 | 0 | 0 | 0 | 0 | 0 | 0 | 0 | 3 | 0 | 3 |
| ENSMUSG00000022160 | Mettl3 | ENSMUST00000122962 | 26.835781 | 26 | 22.9639519 | 24.2990654 | 0 | 0 | 0 | 0 | 0 | 0 | 0 | 0 | 0 | 0 | 0 | 1 | 0 | 0 | 1 |
| ENSMUSG00000051730 | Mettl5 | ENSMUST00000142127 | 40.1098901 | 28 | 16.4835165 | 15.3846154 | 0 | 0 | 0 | 0 | 0 | 0 | 0 | 0 | 0 | 0 | 0 | 0 | 0 | 2 | 2 |
| ENSMUSG00000054619 | Mettl7a1 | ENSMUST00000067752 | 28.8908766 | 25 | 21.2880143 | 24.686941 | 0 | 0 | 0 | 0 | 0 | 0 | 0 | 0 | 0 | 0 | 0 | 2 | 0 | 0 | 2 |
| ENSMUSG00000041975 | Mettl8 | ENSMUST00000112179 | 25.2929014 | 29 | 22.26051 | 23.2942798 | 0 | 0 | 0 | 0 | 0 | 0 | 0 | 0 | 0 | 0 | 0 | 1 | 2 | 0 | 3 |
| ENSMUSG00000037253 | Mex3c | ENSMUST00000091852 | 30.3541315 | 36 | 15.9640247 | 18.1000562 | 0 | 0 | 0 | 0 | 0 | 0 | 0 | 0 | 0 | 0 | 0 | 0 | 2 | 0 | 2 |
| ENSMUSG00000031647 | Mfap3l | ENSMUST00000160719 | 30.9593632 | 31 | 18.9987432 | 18.8730624 | 1 | 0 | 0 | 1 | 0 | 0 | 0 | 0 | 1 | 2 | 0 | 2 | 2 | 1 | 8 |
| ENSMUSG00000026150 | Mff | ENSMUST00000162003 | 27.7286136 | 33 | 19.0265487 | 20.3539823 | 0 | 0 | 0 | 0 | 0 | 0 | 0 | 0 | 0 | 0 | 0 | 0 | 1 | 1 | 2 |
| ENSMUSG00000027668 | Mfn1 | ENSMUST00000091257 | 26.2631077 | 31 | 19.399428 | 23.7845567 | 0 | 0 | 0 | 0 | 0 | 0 | 0 | 0 | 0 | 0 | 0 | 2 | 0 | 0 | 2 |
| ENSMUSG00000027775 | Mfsd1 | ENSMUST00000029344 | 27.4204029 | 32 | 20.14295 | 20.9226771 | 1 | 0 | 0 | 1 | 0 | 0 | 0 | 0 | 1 | 2 | 0 | 0 | 1 | 0 | 4 |
| ENSMUSG00000066595 | Mfsd7b | ENSMUST00000192666 | 28.0683761 | 33 | 18.8376068 | 20.2393162 | 1 | 0 | 0 | 1 | 0 | 0 | 0 | 0 | 1 | 2 | 0 | 0 | 3 | 0 | 6 |
| ENSMUSG00000019888 | Mgat4c | ENSMUST00000020039 | 29.7720798 | 39 | 15.0522317 | 16.6191833 | 1 | 1 | 0 | 0 | 0 | 0 | 0 | 0 | 0 | 0 | 0 | 8 | 1 | 0 | 11 |
| ENSMUSG00000024294 | Mib1 | ENSMUST00000052838 | 28.6533395 | 35 | 17.0308384 | 19.6497753 | 0 | 0 | 0 | 0 | 0 | 0 | 0 | 0 | 0 | 0 | 0 | 4 | 11 | 5 | 20 |
| ENSMUSG00000039478 | Micu3 | ENSMUST00000136835 | 32.4117647 | 33 | 15.3529412 | 19 | 2 | 2 | 0 | 2 | 0 | 0 | 0 | 2 | 0 | 4 | 2 | 0 | 15 | 0 | 25 |
| ENSMUSG00000008035 | Mid1ip1 | ENSMUST00000008179 | 20.7236842 | 29 | 23.1359649 | 27.0833333 | 0 | 0 | 0 | 0 | 0 | 0 | 0 | 0 | 0 | 0 | 0 | 1 | 0 | 0 | 1 |
| ENSMUSG00000000266 | Mid2 | ENSMUST00000112990 | 30.2556077 | 33 | 18.7793427 | 18.2316119 | 0 | 0 | 0 | 0 | 0 | 0 | 0 | 0 | 0 | 0 | 0 | 2 | 0 | 0 | 2 |
| ENSMUSG00000028522 | Mier1 | ENSMUST00000106858 | 28.8378288 | 36 | 17.9487179 | 17.6157176 | 0 | 0 | 0 | 0 | 0 | 0 | 0 | 0 | 0 | 0 | 2 | 1 | 10 | 6 | 19 |
| ENSMUSG00000032727 | Mier3 | ENSMUST00000137268 | 28.8040075 | 32 | 18.7643498 | 20.4550198 | 0 | 0 | 1 | 1 | 0 | 0 | 0 | 0 | 1 | 3 | 0 | 0 | 11 | 4 | 18 |
| ENSMUSG00000022978 | Mis18a | ENSMUST00000099554 | 27.9863481 | 39 | 15.1877133 | 17.5767918 | 0 | 0 | 0 | 0 | 0 | 0 | 0 | 0 | 0 | 0 | 1 | 0 | 0 | 0 | 1 |
| ENSMUSG00000035158 | Mitf | ENSMUST00000043637 | 29.2690611 | 33 | 17.3913043 | 20.2268431 | 1 | 0 | 0 | 0 | 0 | 0 | 0 | 0 | 0 | 0 | 1 | 1 | 3 | 3 | 9 |
| ENSMUSG00000025609 | Mkln1 | ENSMUST00000026699 | 28.0088252 | 32 | 19.9338114 | 20.551572 | 1 | 0 | 0 | 0 | 0 | 0 | 0 | 0 | 0 | 0 | 4 | 7 | 2 | 1 | 15 |
| ENSMUSG00000070527 | Mkrn3 | ENSMUST00000094340 | 27.1498771 | 38 | 15.4791155 | 19.041769 | 1 | 0 | 0 | 1 | 0 | 0 | 0 | 0 | 1 | 2 | 1 | 0 | 0 | 0 | 4 |
| ENSMUSG00000026743 | Mllt10 | ENSMUST00000114671 | 29.251269 | 32 | 18.3375635 | 20.177665 | 1 | 0 | 0 | 1 | 0 | 0 | 0 | 0 | 1 | 2 | 0 | 0 | 0 | 0 | 3 |
| ENSMUSG00000053192 | Mllt11 | ENSMUST00000065482 | 28.952381 | 31 | 20.8571429 | 18.952381 | 4 | 2 | 3 | 9 | 1 | 2 | 3 | 3 | 3 | 24 | 1 | 0 | 1 | 0 | 32 |
| ENSMUSG00000028496 | Mllt3 | ENSMUST00000078090 | 30.0488998 | 33 | 18.3374083 | 19.0953545 | 0 | 0 | 0 | 0 | 0 | 0 | 0 | 0 | 0 | 0 | 3 | 1 | 3 | 2 | 9 |
| ENSMUSG00000068036 | Mllt4 | ENSMUST00000139666 | 26.7382175 | 32 | 19.6920205 | 21.3719085 | 0 | 0 | 0 | 0 | 0 | 0 | 0 | 0 | 0 | 0 | 0 | 0 | 1 | 0 | 1 |
| ENSMUSG00000037022 | Mmaa | ENSMUST00000048718 | 27.1276596 | 30 | 21.0771277 | 21.6755319 | 0 | 0 | 0 | 0 | 0 | 0 | 0 | 0 | 0 | 0 | 0 | 0 | 0 | 1 | 1 |
| ENSMUSG00000026766 | Mmadhc | ENSMUST00000102769 | 24.4648318 | 36 | 18.0428135 | 21.4067278 | 0 | 0 | 1 | 0 | 0 | 0 | 0 | 0 | 0 | 1 | 1 | 0 | 0 | 0 | 2 |
| ENSMUSG00000003948 | Mmd | ENSMUST00000004050 | 26.5932793 | 35 | 18.3082271 | 20.0463499 | 0 | 0 | 0 | 0 | 0 | 0 | 0 | 0 | 0 | 0 | 0 | 0 | 0 | 2 | 2 |
| ENSMUSG00000027820 | Mme | ENSMUST00000194134 | 32.4324324 | 31 | 18.041418 | 18.0940681 | 0 | 0 | 0 | 0 | 0 | 0 | 0 | 0 | 0 | 0 | 0 | 0 | 0 | 1 | 1 |
| ENSMUSG00000061273 | Mmgt1 | ENSMUST00000059899 | 28.6894273 | 32 | 19.9889868 | 18.9702643 | 1 | 1 | 0 | 0 | 0 | 0 | 0 | 0 | 0 | 0 | 1 | 0 | 1 | 0 | 4 |
| ENSMUSG00000028226 | Mmp16 | ENSMUST00000029881 | 30.4035568 | 32 | 17.9206566 | 20.1436389 | 0 | 0 | 0 | 1 | 0 | 0 | 0 | 0 | 1 | 2 | 1 | 0 | 0 | 2 | 5 |
| ENSMUSG00000045751 | Mms22l | ENSMUST00000131282 | 26.2611276 | 31 | 20.6231454 | 21.6617211 | 0 | 0 | 0 | 1 | 0 | 0 | 0 | 0 | 1 | 2 | 0 | 1 | 0 | 0 | 3 |
| ENSMUSG00000021103 | Mnat1 | ENSMUST00000187549 | 29.6414625 | 35 | 16.2229322 | 19.4888179 | 1 | 1 | 2 | 1 | 0 | 0 | 0 | 0 | 1 | 4 | 1 | 0 | 0 | 1 | 8 |
| ENSMUSG00000032221 | Mns1 | ENSMUST00000034746 | 26.8447137 | 33 | 20.2092511 | 19.9063877 | 1 | 0 | 0 | 1 | 0 | 0 | 0 | 0 | 1 | 2 | 0 | 0 | 0 | 1 | 4 |
| ENSMUSG00000043131 | Mob1a | ENSMUST00000055261 | 26.9372694 | 32 | 20.0965087 | 21.3454442 | 1 | 0 | 0 | 0 | 0 | 0 | 0 | 0 | 0 | 0 | 0 | 2 | 1 | 0 | 4 |
| ENSMUSG00000006262 | Mob1b | ENSMUST00000113229 | 29.7837972 | 34 | 16.9234592 | 19.5079523 | 0 | 0 | 0 | 0 | 0 | 0 | 0 | 0 | 0 | 0 | 3 | 3 | 6 | 0 | 12 |
| ENSMUSG00000025979 | Mob4 | ENSMUST00000162364 | 31.2470139 | 33 | 16.5312948 | 19.1113235 | 0 | 1 | 0 | 2 | 0 | 0 | 0 | 0 | 2 | 4 | 1 | 0 | 1 | 0 | 7 |
| ENSMUSG00000015536 | Mocs2 | ENSMUST00000184245 | 30.2325581 | 30 | 19.1684285 | 20.7657975 | 1 | 0 | 0 | 2 | 0 | 0 | 0 | 0 | 2 | 4 | 0 | 0 | 0 | 0 | 5 |
| ENSMUSG00000034602 | Mon2 | ENSMUST00000037557 | 28.3458022 | 31 | 20.2826268 | 20.8090884 | 1 | 1 | 1 | 1 | 0 | 0 | 0 | 0 | 1 | 3 | 0 | 0 | 0 | 0 | 5 |
| ENSMUSG00000062270 | Morf4l1b; Morf4l1 | ENSMUST00000085248 | 23.8897397 | 39 | 18.5298622 | 18.9892802 | 0 | 0 | 0 | 0 | 0 | 0 | 0 | 0 | 0 | 0 | 0 | 2 | 2 | 2 | 6 |
| ENSMUSG00000031422 | Morf4l2 | ENSMUST00000033797 | 26.7563528 | 37 | 17.4887892 | 18.3856502 | 0 | 0 | 0 | 0 | 0 | 0 | 0 | 0 | 0 | 0 | 0 | 0 | 1 | 0 | 1 |
| ENSMUSG00000031843 | Mphosph6 | ENSMUST00000034303 | 23.1053604 | 36 | 19.7781885 | 21.4417745 | 2 | 0 | 2 | 2 | 0 | 0 | 0 | 0 | 2 | 6 | 0 | 0 | 0 | 0 | 8 |
| ENSMUSG00000012429 | Mplkip; Gm7102 | ENSMUST00000049744 | 31.4105671 | 33 | 17.7411537 | 18.0804653 | 1 | 0 | 0 | 1 | 0 | 0 | 0 | 0 | 1 | 2 | 0 | 0 | 0 | 0 | 3 |

| | | | | | | | | | | | | | | | | | | | | | |
|---|---|---|---|---|---|---|---|---|---|---|---|---|---|---|---|---|---|---|---|---|---|
| ENSMUSG00000021112 | Mpp5 | ENSMUST00000082024 | 30.7442314 | 32 | 17.3220669 | 19.8245044 | 0 | 0 | 0 | 0 | 0 | 0 | 0 | 0 | 0 | 0 | 0 | 1 | 0 | 0 | 1 |
| ENSMUSG00000057440 | Mpp7 | ENSMUST00000115869 | 31.2354312 | 33 | 17.6157176 | 18.4482184 | 0 | 0 | 0 | 1 | 0 | 0 | 0 | 0 | 1 | 2 | 1 | 0 | 0 | 1 | 4 |
| ENSMUSG00000022679 | Mpv17l | ENSMUST00000128757 | 29.9554566 | 31 | 18.9680772 | 20.0074239 | 1 | 1 | 0 | 1 | 0 | 0 | 0 | 0 | 1 | 2 | 1 | 0 | 0 | 0 | 5 |
| ENSMUSG00000026712 | Mrc1 | ENSMUST00000028045 | 33.3333333 | 34 | 15.4195011 | 16.7800454 | 0 | 0 | 0 | 0 | 0 | 0 | 0 | 0 | 0 | 0 | 0 | 0 | 0 | 2 | 2 |
| ENSMUSG00000055302 | Mrfap1 | ENSMUST00000068795 | 25.1145738 | 32 | 18.6984418 | 23.9230064 | 0 | 0 | 0 | 0 | 0 | 0 | 0 | 0 | 0 | 0 | 2 | 0 | 4 | 0 | 6 |
| ENSMUSG00000029486 | Mrpl1 | ENSMUST00000117766 | 25.3974895 | 34 | 20.41841 | 20.41841 | 1 | 0 | 0 | 1 | 0 | 0 | 0 | 0 | 1 | 2 | 0 | 1 | 6 | 0 | 10 |
| ENSMUSG00000030045 | Mrpl19 | ENSMUST00000032124 | 26.6503667 | 27 | 22.3227384 | 23.7163814 | 0 | 0 | 0 | 0 | 0 | 0 | 0 | 0 | 0 | 0 | 0 | 0 | 2 | 2 | 4 |
| ENSMUSG00000019710 | Mrpl24 | ENSMUST00000019854 | 25.1533742 | 30 | 21.7791411 | 23.3128834 | 0 | 0 | 0 | 0 | 0 | 0 | 0 | 0 | 0 | 0 | 0 | 1 | 0 | 0 | 1 |
| ENSMUSG00000052962 | Mrpl35 | ENSMUST00000065103 | 25.0483559 | 32 | 20.696325 | 21.9213411 | 0 | 0 | 0 | 0 | 0 | 0 | 0 | 0 | 0 | 0 | 5 | 1 | 13 | 0 | 19 |
| ENSMUSG00000022889 | Mrpl39 | ENSMUST00000116584 | 26.2532982 | 34 | 18.8654354 | 20.9762533 | 0 | 0 | 0 | 0 | 0 | 0 | 0 | 0 | 0 | 0 | 1 | 1 | 0 | 0 | 2 |
| ENSMUSG00000037531 | Mrpl47 | ENSMUST00000043966 | 31.2699135 | 26 | 20.8921256 | 21.5293582 | 1 | 0 | 0 | 0 | 0 | 0 | 0 | 0 | 0 | 0 | 0 | 1 | 0 | 0 | 2 |
| ENSMUSG00000021967 | Mrpl57 | ENSMUST00000022538 | 25.8854167 | 35 | 19.0625 | 20.3645833 | 0 | 0 | 0 | 1 | 0 | 0 | 0 | 0 | 1 | 2 | 3 | 0 | 7 | 2 | 14 |
| ENSMUSG00000058267 | Mrps14 | ENSMUST00000135680 | 30.1953125 | 32 | 18.6328125 | 18.828125 | 0 | 0 | 0 | 0 | 0 | 0 | 0 | 0 | 0 | 0 | 0 | 0 | 0 | 2 | 2 |
| ENSMUSG00000034211 | Mrps17 | ENSMUST00000119576 | 28.1072484 | 29 | 20.6316746 | 22.4267212 | 0 | 0 | 0 | 0 | 0 | 0 | 0 | 0 | 0 | 0 | 1 | 0 | 2 | 2 | 5 |
| ENSMUSG00000040269 | Mrps28 | ENSMUST00000042148 | 40.000000 | ### | 14.000000 | 14.000000 | 0 | 0 | 0 | 1 | 0 | 0 | 0 | 0 | 1 | 2 | 0 | 0 | 0 | 0 | 2 |
| ENSMUSG00000027374 | Mrps5 | ENSMUST00000028852 | 22.6142473 | 35 | 20.6653226 | 21.9758065 | 0 | 0 | 0 | 0 | 0 | 0 | 0 | 0 | 0 | 0 | 8 | 0 | 7 | 3 | 18 |
| ENSMUSG00000039693 | Msantd3 | ENSMUST00000107704 | 22.3723724 | 34 | 19.5195195 | 24.3243243 | 0 | 0 | 0 | 0 | 0 | 0 | 0 | 0 | 0 | 0 | 1 | 0 | 4 | 2 | 7 |
| ENSMUSG00000069769 | Msi2 | ENSMUST00000107909 | 27.8531073 | 33 | 19.3973635 | 19.7551789 | 1 | 0 | 0 | 2 | 0 | 0 | 0 | 0 | 2 | 4 | 0 | 3 | 21 | 11 | 40 |
| ENSMUSG00000052915 | Msl1 | ENSMUST00000037915 | 23.6505974 | 31 | 19.6950968 | 25.1751133 | 0 | 0 | 0 | 0 | 0 | 0 | 0 | 0 | 0 | 0 | 0 | 1 | 1 | 0 | 2 |
| ENSMUSG00000031604 | Msmo1 | ENSMUST00000034015 | 27.390791 | 33 | 19.5985832 | 20.0708383 | 0 | 0 | 0 | 0 | 0 | 0 | 0 | 0 | 0 | 0 | 1 | 0 | 0 | 0 | 1 |
| ENSMUSG00000051236 | Msrb3 | ENSMUST00000092143 | 28.7685061 | 30 | 20.2557201 | 21.3997308 | 0 | 0 | 0 | 0 | 0 | 0 | 0 | 0 | 0 | 0 | 0 | 18 | 4 | 0 | 22 |
| ENSMUSG00000062937 | Mtap | ENSMUST00000058030 | 29.2063492 | 30 | 20.952381 | 20.2539683 | 0 | 0 | 0 | 0 | 0 | 0 | 0 | 0 | 0 | 0 | 0 | 0 | 1 | 0 | 1 |
| ENSMUSG00000022369 | Mtbp | ENSMUST00000170046 | 23.9469579 | 28 | 24.648986 | 23.0889236 | 0 | 0 | 0 | 0 | 0 | 0 | 0 | 0 | 0 | 0 | 0 | 2 | 0 | 0 | 2 |
| ENSMUSG00000022255 | Mtdh | ENSMUST00000022865 | 32.0812808 | 35 | 14.9630542 | 17.6724138 | 1 | 1 | 0 | 0 | 0 | 0 | 0 | 0 | 0 | 0 | 0 | 0 | 0 | 0 | 2 |
| ENSMUSG00000040429 | Mterf1a | ENSMUST00000044746 | 30.1369863 | 40 | 8.90410959 | 21.2328767 | 0 | 0 | 0 | 1 | 0 | 0 | 0 | 0 | 1 | 2 | 0 | 0 | 1 | 1 | 4 |
| ENSMUSG00000053178 | Mterf1b | ENSMUST00000177258 | 28.9655172 | 41 | 8.27586207 | 21.3793103 | 0 | 0 | 0 | 0 | 0 | 0 | 0 | 0 | 0 | 0 | 0 | 0 | 1 | 1 | 2 |
| ENSMUSG00000029267 | Mtf2 | ENSMUST00000143412 | 31.9535221 | 30 | 16.1583152 | 21.4233842 | 0 | 0 | 0 | 0 | 0 | 0 | 0 | 0 | 0 | 0 | 0 | 0 | 6 | 0 | 6 |
| ENSMUSG00000027601 | Mtfr1 | ENSMUST00000130645 | 29.2811222 | 33 | 17.8258328 | 20.2220923 | 0 | 0 | 0 | 0 | 0 | 0 | 0 | 0 | 0 | 0 | 0 | 1 | 0 | 0 | 1 |
| ENSMUSG00000005667 | Mthfd2 | ENSMUST00000203847 | 28.7804878 | 22 | 22.4390244 | 26.3414634 | 0 | 0 | 1 | 0 | 0 | 0 | 0 | 0 | 0 | 1 | 0 | 0 | 0 | 0 | 1 |
| ENSMUSG00000029376 | Mthfd2l | ENSMUST00000071652 | 32.5181159 | 33 | 18.2971014 | 16.3949275 | 3 | 1 | 1 | 2 | 0 | 0 | 0 | 1 | 1 | 5 | 1 | 0 | 0 | 0 | 10 |
| ENSMUSG00000031337 | Mtm1 | ENSMUST00000025391 | 32.1187584 | 36 | 14.9122807 | 17.2739541 | 0 | 0 | 0 | 1 | 0 | 0 | 0 | 0 | 1 | 2 | 0 | 0 | 1 | 0 | 3 |
| ENSMUSG00000015214 | Mtmr1 | ENSMUST00000015358 | 27.0149858 | 32 | 21.6686918 | 19.3195626 | 0 | 1 | 0 | 0 | 0 | 0 | 0 | 0 | 0 | 0 | 0 | 0 | 0 | 0 | 1 |
| ENSMUSG00000039458 | Mtmr12 | ENSMUST00000174418 | 25.4756871 | 28 | 24.127907 | 22.832981 | 0 | 0 | 0 | 0 | 0 | 0 | 0 | 0 | 0 | 0 | 0 | 1 | 0 | 0 | 1 |
| ENSMUSG00000021987 | Mtmr6 | ENSMUST00000022563 | 28.1773932 | 33 | 17.1444024 | 21.4710654 | 0 | 0 | 0 | 0 | 0 | 0 | 0 | 0 | 0 | 0 | 0 | 0 | 2 | 0 | 2 |
| ENSMUSG00000024234 | Mtpap | ENSMUST00000025077 | 26.0718424 | 35 | 22.0162225 | 16.6859791 | 0 | 0 | 0 | 0 | 0 | 0 | 0 | 0 | 0 | 0 | 0 | 1 | 0 | 0 | 1 |
| ENSMUSG00000029840 | Mtpn | ENSMUST00000031866 | 29.8947368 | 33 | 18.4360902 | 18.4360902 | 0 | 0 | 1 | 2 | 0 | 0 | 0 | 0 | 2 | 5 | 0 | 1 | 0 | 0 | 6 |
| ENSMUSG00000045636 | Mtus1 | ENSMUST00000059115 | 26.9752594 | 30 | 21.7478053 | 21.2689545 | 0 | 0 | 0 | 0 | 0 | 0 | 0 | 0 | 0 | 0 | 0 | 0 | 2 | 1 | 3 |
| ENSMUSG00000027099 | Mtx2 | ENSMUST00000028511 | 26.5712826 | 33 | 19.9795606 | 20.1328564 | 0 | 0 | 0 | 0 | 0 | 0 | 0 | 0 | 0 | 0 | 1 | 4 | 3 | 0 | 8 |
| ENSMUSG00000021704 | Mtx3 | ENSMUST00000076169 | 28.0172414 | 31 | 17.8973013 | 23.0697151 | 0 | 0 | 1 | 2 | 0 | 0 | 0 | 0 | 2 | 5 | 1 | 0 | 3 | 0 | 9 |
| ENSMUSG00000023921 | Mut | ENSMUST00000169611 | 33.87471 | 34 | 15.3132251 | 17.1693735 | 1 | 0 | 0 | 0 | 0 | 0 | 0 | 0 | 0 | 0 | 0 | 0 | 1 | 0 | 2 |
| ENSMUSG00000001156 | Mxd1 | ENSMUST00000001184 | 23.1392008 | 29 | 24.2622095 | 23.9749282 | 0 | 0 | 0 | 0 | 0 | 0 | 0 | 0 | 0 | 0 | 1 | 2 | 4 | 1 | 8 |
| ENSMUSG00000068566 | Myadm | ENSMUST00000203566 | 24.7460087 | 29 | 24.4920174 | 21.371553 | 1 | 0 | 0 | 0 | 0 | 0 | 0 | 0 | 0 | 0 | 0 | 6 | 1 | 4 | 12 |
| ENSMUSG00000019982 | Myb | ENSMUST00000188495 | 30.1535974 | 37 | 14.3088116 | 18.7550525 | 0 | 0 | 0 | 0 | 0 | 0 | 0 | 0 | 0 | 0 | 0 | 0 | 0 | 4 | 4 |
| ENSMUSG00000025912 | Mybl1 | ENSMUST00000088658 | 32.8473066 | 37 | 14.2162819 | 16.0793844 | 0 | 0 | 0 | 2 | 0 | 0 | 0 | 0 | 2 | 4 | 0 | 3 | 14 | 1 | 22 |
| ENSMUSG00000028647 | Mycbp | ENSMUST00000030400 | 28.0035336 | 34 | 18.3745583 | 19.5229682 | 1 | 0 | 1 | 1 | 0 | 0 | 0 | 0 | 1 | 3 | 0 | 1 | 0 | 0 | 5 |
| ENSMUSG00000037169 | Mycn | ENSMUST00000043396 | 26.4835165 | 35 | 17.8021978 | 20.6593407 | 0 | 0 | 0 | 0 | 0 | 0 | 0 | 0 | 0 | 0 | 0 | 0 | 4 | 2 | 6 |
| ENSMUSG00000027201 | Myef2 | ENSMUST00000142718 | 30.3618711 | 38 | 15.9752868 | 15.6222418 | 1 | 1 | 0 | 1 | 0 | 0 | 0 | 0 | 1 | 2 | 0 | 0 | 15 | 1 | 20 |
| ENSMUSG00000024048 | Myl12a | ENSMUST00000148960 | 23.4309623 | 35 | 22.1757322 | 19.3514644 | 1 | 0 | 0 | 0 | 0 | 0 | 0 | 0 | 0 | 0 | 1 | 0 | 0 | 0 | 2 |
| ENSMUSG00000034868 | Myl12b | ENSMUST00000038446 | 27.4752475 | 32 | 21.5346535 | 19.3069307 | 0 | 0 | 0 | 0 | 0 | 0 | 0 | 0 | 0 | 0 | 0 | 1 | 1 | 2 | 4 |
| ENSMUSG00000037730 | Mynn | ENSMUST00000195396 | 29.6263345 | 31 | 19.5284698 | 19.8398577 | 0 | 0 | 0 | 0 | 0 | 0 | 0 | 0 | 0 | 0 | 0 | 1 | 1 | 1 | 3 |
| ENSMUSG00000039585 | Myo9a | ENSMUST00000136740 | 28.0573664 | 32 | 19.6349413 | 19.8956975 | 0 | 0 | 0 | 1 | 0 | 0 | 0 | 0 | 1 | 2 | 0 | 0 | 1 | 0 | 3 |
| ENSMUSG00000062627 | Mysm1 | ENSMUST00000075872 | 29.1979695 | 32 | 19.2893401 | 19.4517766 | 0 | 0 | 0 | 0 | 0 | 0 | 0 | 0 | 0 | 0 | 0 | 0 | 11 | 0 | 11 |
| ENSMUSG00000010505 | Myt1 | ENSMUST00000081125 | 25.1568739 | 34 | 18.4826013 | 22.2475756 | 3 | 1 | 0 | 4 | 0 | 0 | 0 | 1 | 3 | 8 | 0 | 0 | 4 | 2 | 18 |
| ENSMUSG00000033186 | Mzt1 | ENSMUST00000042662 | 30.1644399 | 35 | 15.9815005 | 19.270298 | 0 | 0 | 2 | 0 | 0 | 0 | 0 | 0 | 0 | 2 | 0 | 0 | 3 | 0 | 5 |
| ENSMUSG00000037795 | N4bp2 | ENSMUST00000201489 | 29.3811452 | 34 | 17.4089069 | 19.1150954 | 0 | 0 | 0 | 1 | 0 | 0 | 0 | 0 | 1 | 2 | 3 | 0 | 2 | 0 | 7 |
| ENSMUSG00000041132 | N4bp2l1 | ENSMUST00000016279 | 28.5714286 | 30 | 18.4713376 | 22.7479527 | 0 | 0 | 1 | 0 | 0 | 0 | 0 | 0 | 0 | 1 | 0 | 0 | 0 | 0 | 1 |
| ENSMUSG00000029655 | N4bp2l2 | ENSMUST00000118316 | 28.5774293 | 37 | 16.2559507 | 18.1461775 | 1 | 1 | 0 | 0 | 0 | 0 | 0 | 0 | 0 | 0 | 4 | 0 | 10 | 6 | 22 |
| ENSMUSG00000063273 | Naa15 | ENSMUST00000193266 | 29.9163744 | 31 | 17.7232263 | 20.9063933 | 1 | 0 | 0 | 0 | 0 | 0 | 0 | 0 | 0 | 0 | 1 | 0 | 0 | 0 | 2 |
| ENSMUSG00000022020 | Naa16 | ENSMUST00000022597 | 25.99837 | 37 | 16.9519152 | 20.2933985 | 0 | 0 | 0 | 0 | 0 | 0 | 0 | 0 | 0 | 0 | 3 | 0 | 8 | 0 | 11 |
| ENSMUSG00000002728 | Naa20 | ENSMUST00000134759 | 31.6478287 | 28 | 18.6198691 | 21.8322427 | 1 | 0 | 0 | 2 | 0 | 0 | 0 | 0 | 2 | 4 | 0 | 0 | 0 | 0 | 5 |
| ENSMUSG00000036282 | Naa30 | ENSMUST00000037362 | 28.5885167 | 37 | 15.4007177 | 18.9892344 | 0 | 1 | 0 | 1 | 0 | 0 | 0 | 0 | 1 | 2 | 2 | 0 | 4 | 0 | 9 |
| ENSMUSG00000024764 | Naa40 | ENSMUST00000025675 | 20.8150745 | 29 | 26.5118317 | 23.9263804 | 0 | 0 | 0 | 0 | 0 | 0 | 0 | 0 | 0 | 0 | 0 | 0 | 1 | 0 | 1 |
| ENSMUSG00000022698 | Naa50 | ENSMUST00000161326 | 31.0520666 | 35 | 14.949007 | 19.1894793 | 0 | 0 | 0 | 0 | 0 | 0 | 0 | 0 | 0 | 0 | 0 | 1 | 3 | 0 | 4 |
| ENSMUSG00000029413 | Naaa | ENSMUST00000113102 | 27.7389277 | 28 | 21.7560218 | 22.1445221 | 0 | 0 | 0 | 0 | 0 | 0 | 0 | 0 | 0 | 0 | 1 | 0 | 0 | 0 | 1 |
| ENSMUSG00000026107 | Nabp1 | ENSMUST00000189542 | 30.1442041 | 32 | 17.9700499 | 20.160843 | 1 | 0 | 1 | 0 | 0 | 0 | 0 | 0 | 0 | 1 | 0 | 1 | 2 | 3 | 8 |
| ENSMUSG00000061315 | Naca | ENSMUST00000073868 | 23.8461538 | 36 | 20.7692308 | 19.2307692 | 0 | 0 | 0 | 0 | 0 | 0 | 0 | 0 | 0 | 0 | 0 | 1 | 0 | 0 | 1 |
| ENSMUSG00000029063 | Nadk | ENSMUST00000105612 | 23.197903 | 29 | 25.4259502 | 22.0838794 | 0 | 0 | 0 | 0 | 0 | 0 | 0 | 0 | 0 | 0 | 0 | 1 | 0 | 0 | 1 |
| ENSMUSG00000022253 | Nadk2 | ENSMUST00000100789 | 30.3755869 | 36 | 15.4460094 | 17.8403756 | 0 | 0 | 0 | 0 | 0 | 0 | 0 | 0 | 0 | 0 | 0 | 0 | 3 | 2 | 5 |
| ENSMUSG00000014907 | Naf1 | ENSMUST00000118009 | 32.5047801 | 36 | 15.2963671 | 16.6347992 | 0 | 1 | 0 | 1 | 0 | 0 | 0 | 0 | 1 | 2 | 0 | 0 | 1 | 0 | 4 |
| ENSMUSG00000000197 | Nalcn | ENSMUST00000000201 | 31.3032089 | 34 | 16.3719712 | 18.4675835 | 0 | 0 | 0 | 0 | 0 | 0 | 0 | 0 | 0 | 0 | 0 | 0 | 0 | 2 | 2 |
| ENSMUSG00000058799 | Nap1l1 | ENSMUST00000171797 | 27.7697146 | 34 | 16.7392356 | 21.4320271 | 0 | 0 | 0 | 0 | 0 | 0 | 0 | 0 | 0 | 0 | 1 | 1 | 0 | 0 | 2 |
| ENSMUSG00000055733 | Nap1l3 | ENSMUST00000079490 | 31.3414634 | 39 | 13.7804878 | 16.3414634 | 0 | 0 | 0 | 1 | 0 | 0 | 0 | 0 | 1 | 2 | 0 | 0 | 0 | 0 | 2 |
| ENSMUSG00000027438 | Napb | ENSMUST00000136513 | 28.0603118 | 29 | 20.546895 | 22.3613596 | 0 | 0 | 0 | 0 | 0 | 0 | 0 | 0 | 0 | 0 | 1 | 0 | 1 | 1 | 3 |
| ENSMUSG00000044968 | Napepld | ENSMUST00000115217 | 29.0502793 | 30 | 21.1431027 | 19.681994 | 1 | 0 | 0 | 0 | 0 | 0 | 0 | 0 | 0 | 0 | 0 | 1 | 1 | 2 | 5 |
| ENSMUSG00000024581 | Napg | ENSMUST00000025474 | 27.9518072 | 31 | 19.8393574 | 21.0040161 | 1 | 0 | 0 | 0 | 0 | 0 | 0 | 0 | 0 | 0 | 0 | 0 | 0 | 0 | 1 |
| ENSMUSG00000018995 | Nars2 | ENSMUST00000044466 | 28.0195599 | 33 | 18.5330073 | 20.4889976 | 0 | 0 | 0 | 1 | 0 | 0 | 0 | 0 | 1 | 2 | 1 | 0 | 1 | 0 | 4 |
| ENSMUSG00000028693 | Nasp | ENSMUST00000030457 | 25.4518072 | 35 | 16.1144578 | 23.1927711 | 0 | 0 | 0 | 0 | 0 | 0 | 0 | 0 | 0 | 0 | 0 | 0 | 0 | 1 | 1 |
| ENSMUSG00000073664 | Nbeal1 | ENSMUST00000160834 | 30.6161396 | 36 | 15.6761178 | 17.4345692 | 1 | 0 | 2 | 6 | 0 | 0 | 0 | 0 | 6 | 14 | 5 | 4 | 3 | 0 | 27 |
| ENSMUSG00000039542 | Ncam1 | ENSMUST00000193547 | 27.4569096 | 29 | 22.1953432 | 21.5905655 | 0 | 0 | 0 | 0 | 0 | 0 | 0 | 0 | 0 | 0 | 2 | 5 | 16 | 1 | 24 |
| ENSMUSG00000022762 | Ncam2 | ENSMUST00000037785 | 34.5505618 | 34 | 17.4157303 | 13.5634029 | 0 | 0 | 0 | 1 | 0 | 0 | 0 | 0 | 1 | 2 | 0 | 0 | 2 | 0 | 4 |
| ENSMUSG00000035024 | Ncapd3 | ENSMUST00000086198 | 29.1902072 | 34 | 17.5141243 | 18.8323917 | 0 | 0 | 0 | 1 | 0 | 0 | 0 | 0 | 1 | 2 | 0 | 0 | 0 | 0 | 2 |
| ENSMUSG00000015880 | Ncapg | ENSMUST00000117396 | 31.8493151 | 35 | 17.2945205 | 15.4109589 | 1 | 1 | 0 | 1 | 0 | 0 | 0 | 0 | 1 | 2 | 0 | 0 | 0 | 0 | 4 |
| ENSMUSG00000032475 | Nck1 | ENSMUST00000116522 | 30.4962312 | 33 | 18.1532663 | 18.3417085 | 0 | 0 | 0 | 0 | 0 | 0 | 0 | 0 | 0 | 0 | 0 | 0 | 2 | 2 | 4 |
| ENSMUSG00000049690 | Nckap5 | ENSMUST00000094609 | 28.5087719 | 33 | 16.5935673 | 22.0760234 | 0 | 0 | 0 | 0 | 0 | 0 | 0 | 0 | 0 | 0 | 0 | 1 | 0 | 1 | 2 |
| ENSMUSG00000028614 | Ndc1 | ENSMUST00000149366 | 26.1507937 | 31 | 20.1984127 | 22.3809524 | 0 | 0 | 0 | 0 | 0 | 0 | 0 | 0 | 0 | 0 | 0 | 0 | 0 | 1 | 1 |
| ENSMUSG00000053253 | Ndfip2 | ENSMUST00000181969 | 31.7314488 | 35 | 15.8303887 | 17.5971731 | 2 | 1 | 0 | 1 | 0 | 0 | 0 | 0 | 1 | 2 | 0 | 0 | 0 | 0 | 5 |
| ENSMUSG00000049001 | Ndnf | ENSMUST00000054351 | 30.262657 | 31 | 19.4899124 | 19.6548661 | 1 | 0 | 0 | 1 | 0 | 0 | 0 | 0 | 1 | 2 | 0 | 13 | 0 | 0 | 16 |
| ENSMUSG00000070520 | Ndnl2 | ENSMUST00000094331 | 24.726477 | 34 | 20.6484981 | 21.0463497 | 0 | 0 | 0 | 0 | 0 | 0 | 0 | 0 | 0 | 0 | 4 | 3 | 3 | 0 | 10 |
| ENSMUSG00000027977 | Ndst3 | ENSMUST00000137404 | 31.6939891 | 26 | 22.1311475 | 20.5525197 | 0 | 0 | 0 | 0 | 0 | 0 | 0 | 0 | 0 | 0 | 0 | 1 | 0 | 0 | 1 |
| ENSMUSG00000027971 | Ndst4 | ENSMUST00000173932 | 31.9205619 | 34 | 19.0845241 | 15.4516832 | 0 | 1 | 0 | 1 | 0 | 0 | 0 | 0 | 1 | 2 | 0 | 2 | 1 | 0 | 6 |
| ENSMUSG00000029632 | Ndufa4 | ENSMUST00000204978 | 32.0987654 | 35 | 13.5802469 | 19.3415638 | 0 | 0 | 0 | 0 | 0 | 0 | 0 | 0 | 0 | 0 | 0 | 0 | 2 | 0 | 2 |
| ENSMUSG00000028261 | Ndufaf4 | ENSMUST00000029925 | 31.526768 | 30 | 17.9114342 | 20.1916722 | 0 | 0 | 0 | 0 | 0 | 0 | 0 | 0 | 0 | 0 | 0 | 1 | 0 | 0 | 1 |

| | | | | | | | | | | | | | | | | | | | | | |
|---|---|---|---|---|---|---|---|---|---|---|---|---|---|---|---|---|---|---|---|---|---|
| ENSMUSG00000002416 | Ndufb2 | ENSMUST00000135671 | 27.9411765 | 30 | 19.7906281 | 22.6321037 | 0 | 0 | 0 | 0 | 0 | 0 | 0 | 0 | 0 | 0 | 0 | 0 | 6 | 4 | 10 |
| ENSMUSG00000026032 | Ndufb3 | ENSMUST00000027193 | 35.8490566 | 35 | 13.2075472 | 16.3522013 | 0 | 0 | 0 | 0 | 0 | 0 | 0 | 0 | 0 | 0 | 1 | 0 | 0 | 0 | 1 |
| ENSMUSG00000037152 | Ndufc1 | ENSMUST00000038108 | 29.5833333 | 30 | 18.8541667 | 21.7708333 | 0 | 0 | 0 | 0 | 0 | 0 | 0 | 0 | 0 | 0 | 0 | 0 | 2 | 0 | 2 |
| ENSMUSG00000024099 | Ndufv2 | ENSMUST00000143987 | 30.472103 | 36 | 18.0257511 | 15.3075823 | 2 | 1 | 0 | 2 | 0 | 0 | 0 | 1 | 1 | 4 | 0 | 0 | 0 | 0 | 7 |
| ENSMUSG00000053702 | Nebl | ENSMUST00000028080 | 29.9059561 | 33 | 17.9310345 | 19.5820272 | 0 | 0 | 0 | 1 | 0 | 0 | 0 | 0 | 1 | 2 | 1 | 2 | 0 | 0 | 5 |
| ENSMUSG00000040536 | Necab1 | ENSMUST00000041606 | 32.8947368 | 31 | 18.6951754 | 17.1326754 | 1 | 0 | 0 | 0 | 0 | 0 | 0 | 0 | 0 | 0 | 0 | 0 | 0 | 0 | 1 |
| ENSMUSG00000019988 | Nedd1 | ENSMUST00000020163 | 29.6193772 | 37 | 16.1937716 | 16.7474048 | 0 | 1 | 0 | 1 | 0 | 0 | 0 | 0 | 1 | 2 | 1 | 0 | 0 | 0 | 4 |
| ENSMUSG00000032216 | Nedd4 | ENSMUST00000034740 | 28.0487805 | 30 | 21.6463415 | 20.7698171 | 0 | 0 | 0 | 0 | 0 | 0 | 0 | 0 | 0 | 0 | 0 | 2 | 0 | 0 | 2 |
| ENSMUSG00000021365 | Nedd9 | ENSMUST00000021794 | 29.3982808 | 32 | 17.9942693 | 20.3438395 | 0 | 0 | 0 | 0 | 0 | 0 | 0 | 0 | 0 | 0 | 0 | 0 | 0 | 4 | 4 |
| ENSMUSG00000026393 | Nek7 | ENSMUST00000186017 | 28.6787879 | 33 | 18.1575758 | 19.7090909 | 0 | 0 | 0 | 0 | 0 | 0 | 0 | 0 | 0 | 0 | 0 | 0 | 5 | 0 | 5 |
| ENSMUSG00000029111 | Nelfa | ENSMUST00000030993 | 27.3176761 | 31 | 19.0358467 | 22.6205192 | 0 | 0 | 0 | 0 | 0 | 0 | 0 | 0 | 0 | 0 | 0 | 2 | 0 | 0 | 2 |
| ENSMUSG00000021215 | Net1 | ENSMUST00000091853 | 27.1760797 | 32 | 16.744186 | 23.5880399 | 0 | 0 | 1 | 0 | 0 | 0 | 0 | 0 | 0 | 1 | 2 | 2 | 0 | 0 | 5 |
| ENSMUSG00000050321 | Neto1 | ENSMUST00000058829 | 30.3607214 | 35 | 18.1362725 | 16.5330661 | 0 | 0 | 0 | 0 | 0 | 0 | 0 | 0 | 0 | 0 | 0 | 4 | 0 | 0 | 4 |
| ENSMUSG00000036902 | Neto2 | ENSMUST00000109686 | 31.1803445 | 35 | 17.7051672 | 16.2613982 | 1 | 0 | 0 | 2 | 0 | 0 | 0 | 0 | 2 | 4 | 0 | 0 | 1 | 2 | 8 |
| ENSMUSG00000037984 | Neurod6 | ENSMUST00000044767 | 29.8159509 | 37 | 16.6871166 | 16.9325153 | 0 | 0 | 0 | 0 | 0 | 0 | 0 | 0 | 0 | 0 | 0 | 18 | 0 | 0 | 18 |
| ENSMUSG00000027967 | Neurog2 | ENSMUST00000029587 | 26.1026103 | 30 | 21.4221422 | 22.1422142 | 1 | 0 | 0 | 0 | 0 | 0 | 0 | 0 | 0 | 0 | 0 | 4 | 1 | 0 | 6 |
| ENSMUSG00000020716 | Nf1 | ENSMUST00000071325 | 25.8706468 | 31 | 22.2947761 | 20.8022388 | 1 | 0 | 0 | 0 | 0 | 0 | 0 | 0 | 0 | 0 | 0 | 2 | 11 | 3 | 17 |
| ENSMUSG00000003847 | Nfat5 | ENSMUST00000133026 | 27.4497537 | 31 | 22.3117228 | 19.558927 | 0 | 1 | 0 | 3 | 0 | 0 | 0 | 0 | 3 | 6 | 1 | 77 | 3 | 0 | 88 |
| ENSMUSG00000031902 | Nfatc3 | ENSMUST00000109308 | 27.2945892 | 31 | 20.2004008 | 21.9639279 | 0 | 0 | 0 | 1 | 0 | 0 | 0 | 0 | 1 | 2 | 1 | 1 | 0 | 0 | 4 |
| ENSMUSG00000028565 | Nfia | ENSMUST00000092532 | 27.8177772 | 33 | 20.9058777 | 18.7197277 | 0 | 0 | 0 | 0 | 0 | 0 | 0 | 0 | 0 | 0 | 2 | 14 | 45 | 1 | 62 |
| ENSMUSG00000008575 | Nfib | ENSMUST00000107245 | 30.0558828 | 31 | 18.6527715 | 20.5104969 | 1 | 1 | 0 | 2 | 0 | 0 | 0 | 1 | 1 | 4 | 6 | 2 | 16 | 7 | 37 |
| ENSMUSG00000021025 | Nfkbia | ENSMUST00000021413 | 30.7240705 | 32 | 15.0684932 | 22.7005871 | 1 | 0 | 0 | 0 | 0 | 0 | 0 | 0 | 0 | 0 | 0 | 0 | 0 | 0 | 1 |
| ENSMUSG00000035356 | Nfkbiz | ENSMUST00000036273 | 29.3726937 | 35 | 15.498155 | 20.4428044 | 0 | 0 | 0 | 0 | 0 | 0 | 0 | 0 | 0 | 0 | 0 | 0 | 0 | 1 | 1 |
| ENSMUSG00000072889 | Nfxl1 | ENSMUST00000135318 | 28.5076847 | 28 | 20.2776401 | 23.5498265 | 0 | 0 | 0 | 0 | 0 | 0 | 0 | 0 | 0 | 0 | 0 | 0 | 1 | 0 | 1 |
| ENSMUSG00000020248 | Nfyb | ENSMUST00000130911 | 25.9626604 | 36 | 16.1610268 | 21.5869312 | 1 | 1 | 0 | 0 | 0 | 0 | 0 | 0 | 0 | 0 | 0 | 0 | 1 | 0 | 3 |
| ENSMUSG00000048540 | Nhlh2 | ENSMUST00000066187 | 30.6869601 | 29 | 19.2513369 | 21.3081037 | 1 | 1 | 0 | 0 | 0 | 0 | 0 | 0 | 0 | 0 | 0 | 1 | 0 | 0 | 3 |
| ENSMUSG00000025078 | Nhlrc2 | ENSMUST00000071423 | 29.1604198 | 36 | 16.4167916 | 18.7406297 | 0 | 0 | 0 | 1 | 0 | 0 | 0 | 0 | 1 | 2 | 0 | 0 | 0 | 0 | 2 |
| ENSMUSG00000063480 | Nhp2l1; LOC100862468 | ENSMUST00000166578 | 22.974607 | 36 | 19.2261185 | 22.0072551 | 0 | 0 | 0 | 0 | 0 | 0 | 0 | 0 | 0 | 0 | 1 | 0 | 1 | 0 | 2 |
| ENSMUSG00000031917 | Nip7 | ENSMUST00000034392 | 25.5967078 | 31 | 20.3703704 | 23.0864198 | 0 | 0 | 0 | 0 | 0 | 0 | 0 | 0 | 0 | 0 | 4 | 0 | 13 | 1 | 18 |
| ENSMUSG00000030452 | Nipa2 | ENSMUST00000032635 | 29.0362185 | 36 | 17.9864948 | 17.0042971 | 0 | 0 | 0 | 0 | 0 | 0 | 0 | 0 | 0 | 0 | 0 | 1 | 1 | 2 | 4 |
| ENSMUSG00000038879 | Nipal2 | ENSMUST00000040791 | 28.7969925 | 31 | 21.1654135 | 18.8345865 | 0 | 0 | 1 | 0 | 0 | 0 | 0 | 0 | 0 | 1 | 1 | 1 | 0 | 0 | 3 |
| ENSMUSG00000015247 | Nipsnap3b | ENSMUST00000015391 | 28.7142857 | 38 | 18 | 15.4285714 | 0 | 0 | 0 | 0 | 0 | 0 | 0 | 0 | 0 | 0 | 1 | 0 | 1 | 0 | 2 |
| ENSMUSG00000055761 | Nkain3 | ENSMUST00000119374 | 30.3528787 | 30 | 19.7930486 | 19.501194 | 0 | 0 | 0 | 0 | 0 | 0 | 0 | 0 | 0 | 0 | 0 | 1 | 7 | 1 | 9 |
| ENSMUSG00000016409 | Nkap | ENSMUST00000016553 | 29.5577131 | 33 | 17.9791442 | 19.3455592 | 0 | 1 | 0 | 1 | 0 | 0 | 0 | 0 | 1 | 2 | 0 | 0 | 3 | 0 | 6 |
| ENSMUSG00000021772 | Nkiras1 | ENSMUST00000132374 | 27.7911005 | 32 | 20.7833733 | 19.4244604 | 0 | 0 | 0 | 0 | 0 | 0 | 0 | 0 | 0 | 0 | 2 | 0 | 1 | 2 | 5 |
| ENSMUSG00000044149 | Nkrf | ENSMUST00000057093 | 31.1330049 | 33 | 17.3399015 | 18.1280788 | 0 | 0 | 0 | 0 | 0 | 0 | 0 | 0 | 0 | 0 | 2 | 0 | 0 | 0 | 2 |
| ENSMUSG00000032525 | Nktr | ENSMUST00000035112 | 25.4082039 | 31 | 21.6248507 | 22.3018718 | 0 | 0 | 0 | 0 | 0 | 0 | 0 | 0 | 0 | 0 | 1 | 0 | 2 | 2 | 5 |
| ENSMUSG00000063887 | Nlgn1 | ENSMUST00000193603 | 33.1806547 | 34 | 16.2737438 | 16.9473259 | 0 | 0 | 0 | 0 | 0 | 0 | 0 | 0 | 0 | 0 | 0 | 0 | 21 | 2 | 23 |
| ENSMUSG00000017376 | Nlk | ENSMUST00000142739 | 31.317741 | 32 | 16.1437536 | 20.2509983 | 0 | 0 | 0 | 0 | 0 | 0 | 0 | 0 | 0 | 0 | 0 | 0 | 3 | 1 | 4 |
| ENSMUSG00000027787 | Nmd3 | ENSMUST00000029358 | 26.9575693 | 35 | 18.5115781 | 19.2745282 | 1 | 0 | 0 | 2 | 0 | 0 | 0 | 0 | 2 | 4 | 2 | 24 | 19 | 1 | 51 |
| ENSMUSG00000037847 | Nmrk1 | ENSMUST00000159572 | 29.7581493 | 28 | 21.3459516 | 20.8201893 | 0 | 0 | 0 | 0 | 0 | 0 | 0 | 0 | 0 | 0 | 0 | 0 | 1 | 0 | 1 |
| ENSMUSG00000026643 | Nmt2 | ENSMUST00000081932 | 29.3969849 | 31 | 18.8083274 | 20.3876525 | 0 | 0 | 0 | 0 | 0 | 0 | 0 | 0 | 0 | 0 | 2 | 4 | 15 | 2 | 23 |
| ENSMUSG00000024999 | Noc3l | ENSMUST00000025963 | 26.6760762 | 32 | 20.8892025 | 20.3952011 | 1 | 1 | 0 | 0 | 0 | 0 | 0 | 0 | 0 | 0 | 0 | 2 | 4 | 0 | 8 |
| ENSMUSG00000048616 | Nog | ENSMUST00000061728 | 30.994152 | 30 | 19.005848 | 19.5906433 | 0 | 0 | 0 | 0 | 0 | 0 | 0 | 0 | 0 | 0 | 0 | 0 | 0 | 4 | 4 |
| ENSMUSG00000061458 | Nol10 | ENSMUST00000046011 | 21.2790698 | 29 | 25.5813953 | 24.3023256 | 0 | 0 | 0 | 0 | 0 | 0 | 0 | 0 | 0 | 0 | 1 | 7 | 4 | 0 | 12 |
| ENSMUSG00000026020 | Nop58 | ENSMUST00000191142 | 26.8419008 | 33 | 18.8657258 | 21.6755147 | 3 | 2 | 0 | 3 | 0 | 0 | 0 | 1 | 2 | 6 | 2 | 0 | 11 | 3 | 27 |
| ENSMUSG00000021047 | Nova1 | ENSMUST00000021438 | 29.7688921 | 35 | 17.4431401 | 17.6999266 | 1 | 1 | 0 | 1 | 0 | 0 | 0 | 0 | 1 | 2 | 0 | 0 | 10 | 4 | 18 |
| ENSMUSG00000030562 | Nox4 | ENSMUST00000126887 | 30.7942132 | 31 | 20.0767641 | 18.039563 | 0 | 0 | 2 | 0 | 0 | 0 | 0 | 0 | 0 | 2 | 0 | 0 | 0 | 0 | 2 |
| ENSMUSG00000021010 | Npas3 | ENSMUST00000101432 | 27.7007417 | 35 | 18.4779104 | 19.0906159 | 0 | 0 | 0 | 0 | 0 | 0 | 0 | 0 | 0 | 0 | 4 | 3 | 12 | 3 | 22 |
| ENSMUSG00000001441 | Npepps | ENSMUST00000001480 | 29.009434 | 28 | 22.327044 | 20.8333333 | 0 | 0 | 0 | 0 | 0 | 0 | 0 | 0 | 0 | 0 | 0 | 2 | 0 | 0 | 2 |
| ENSMUSG00000026241 | Nppc | ENSMUST00000027449 | 31.8181818 | 34 | 16.8560606 | 17.6136364 | 1 | 0 | 0 | 0 | 0 | 0 | 0 | 0 | 0 | 0 | 0 | 0 | 1 | 1 | 3 |
| ENSMUSG00000022206 | Npr3 | ENSMUST00000066529 | 29.5587957 | 30 | 18.4624124 | 21.6436281 | 0 | 0 | 0 | 1 | 0 | 0 | 0 | 0 | 1 | 2 | 0 | 1 | 1 | 1 | 5 |
| ENSMUSG00000032336 | Nptn | ENSMUST00000176250 | 27.3264781 | 34 | 19.4344473 | 19.1002571 | 1 | 0 | 0 | 2 | 0 | 0 | 0 | 0 | 2 | 4 | 0 | 5 | 6 | 0 | 16 |
| ENSMUSG00000036437 | Npy1r | ENSMUST00000039303 | 27.9503106 | 30 | 19.8757764 | 22.1118012 | 1 | 0 | 0 | 0 | 0 | 0 | 0 | 0 | 0 | 0 | 0 | 0 | 0 | 0 | 1 |
| ENSMUSG00000028004 | Npy2r | ENSMUST00000029633 | 32.868937 | 32 | 16.3570691 | 18.9886481 | 1 | 0 | 0 | 0 | 0 | 0 | 0 | 0 | 0 | 0 | 0 | 0 | 0 | 0 | 1 |
| ENSMUSG00000021775 | Nr1d2 | ENSMUST00000090543 | 30.1406309 | 34 | 15.2033447 | 20.2204485 | 0 | 0 | 1 | 0 | 0 | 0 | 0 | 0 | 0 | 1 | 0 | 1 | 1 | 0 | 3 |
| ENSMUSG00000005897 | Nr2c1 | ENSMUST00000105290 | 28.655914 | 34 | 20.3763441 | 17.0967742 | 0 | 0 | 0 | 0 | 0 | 0 | 0 | 0 | 0 | 0 | 0 | 8 | 0 | 0 | 8 |
| ENSMUSG00000030551 | Nr2f2 | ENSMUST00000032768 | 30.4684618 | 33 | 15.4186647 | 20.7303578 | 0 | 0 | 0 | 0 | 0 | 0 | 0 | 0 | 0 | 0 | 0 | 2 | 12 | 5 | 19 |
| ENSMUSG00000023034 | Nr4a1 | ENSMUST00000023779 | 24.6296296 | 28 | 24.0740741 | 22.962963 | 0 | 0 | 0 | 1 | 0 | 0 | 0 | 0 | 1 | 2 | 0 | 0 | 0 | 0 | 2 |
| ENSMUSG00000026826 | Nr4a2 | ENSMUST00000183542 | 32.3133415 | 31 | 18.7270502 | 18.1762546 | 0 | 0 | 0 | 0 | 0 | 0 | 0 | 0 | 0 | 0 | 0 | 0 | 2 | 0 | 2 |
| ENSMUSG00000028341 | Nr4a3 | ENSMUST00000153369 | 29.4345719 | 32 | 18.19063 | 19.9676898 | 0 | 0 | 0 | 1 | 0 | 0 | 0 | 0 | 1 | 2 | 0 | 0 | 6 | 6 | 14 |
| ENSMUSG00000063972 | Nr6a1 | ENSMUST00000112877 | 25.2411936 | 29 | 23.7155037 | 21.7410814 | 1 | 0 | 0 | 0 | 0 | 0 | 0 | 0 | 0 | 0 | 2 | 3 | 1 | 0 | 7 |
| ENSMUSG00000027852 | Nras | ENSMUST00000029445 | 27.3946874 | 32 | 19.9892675 | 20.740542 | 0 | 0 | 0 | 0 | 0 | 0 | 0 | 0 | 0 | 0 | 1 | 1 | 13 | 0 | 15 |
| ENSMUSG00000075000 | Nrbf2 | ENSMUST00000077839 | 31.1688312 | 31 | 16.8831169 | 21.4285714 | 0 | 0 | 0 | 1 | 0 | 0 | 0 | 0 | 1 | 2 | 1 | 0 | 0 | 0 | 3 |
| ENSMUSG00000020598 | Nrcam | ENSMUST00000020939 | 27.9840442 | 30 | 21.7244554 | 20.5277693 | 0 | 0 | 0 | 0 | 0 | 0 | 0 | 0 | 0 | 0 | 1 | 1 | 1 | 0 | 3 |
| ENSMUSG00000042834 | Nrep | ENSMUST00000051087 | 24.4837758 | 34 | 19.3510324 | 22.300885 | 0 | 0 | 0 | 0 | 0 | 0 | 0 | 0 | 0 | 0 | 0 | 1 | 0 | 0 | 1 |
| ENSMUSG00000058440 | Nrf1 | ENSMUST00000115212 | 25.1544076 | 29 | 21.111735 | 25.0982594 | 0 | 0 | 0 | 0 | 0 | 0 | 0 | 0 | 0 | 0 | 0 | 0 | 3 | 2 | 5 |
| ENSMUSG00000039114 | Nrn1 | ENSMUST00000037623 | 29.6111665 | 31 | 19.441675 | 20.2392822 | 0 | 0 | 0 | 0 | 0 | 0 | 0 | 0 | 0 | 0 | 0 | 1 | 0 | 0 | 1 |
| ENSMUSG00000025810 | Nrp1 | ENSMUST00000026917 | 30.5891239 | 30 | 19.826284 | 19.2975831 | 0 | 0 | 0 | 0 | 0 | 0 | 0 | 0 | 0 | 0 | 1 | 2 | 0 | 0 | 3 |
| ENSMUSG00000025969 | Nrp2 | ENSMUST00000114155 | 26.2556391 | 30 | 23.6390977 | 20.3609023 | 0 | 0 | 0 | 0 | 0 | 0 | 0 | 0 | 0 | 0 | 1 | 10 | 2 | 0 | 13 |
| ENSMUSG00000024109 | Nrxn1 | ENSMUST00000072671 | 34.9970981 | 31 | 17.2373767 | 16.2797446 | 1 | 0 | 0 | 1 | 0 | 0 | 0 | 0 | 1 | 2 | 0 | 1 | 3 | 0 | 7 |
| ENSMUSG00000066392 | Nrxn3 | ENSMUST00000163134 | 32.3564356 | 31 | 17.5445545 | 18.6138614 | 1 | 0 | 0 | 0 | 0 | 0 | 0 | 0 | 0 | 0 | 3 | 3 | 4 | 3 | 14 |
| ENSMUSG00000034187 | Nsf | ENSMUST00000103075 | 25.206044 | 28 | 24.9313187 | 21.6346154 | 0 | 0 | 1 | 0 | 0 | 0 | 0 | 0 | 0 | 1 | 0 | 0 | 0 | 0 | 1 |
| ENSMUSG00000029126 | Nsg1 | ENSMUST00000031009 | 24.0540541 | 29 | 20.7432432 | 26.3513514 | 0 | 0 | 0 | 0 | 0 | 0 | 0 | 0 | 0 | 0 | 0 | 0 | 3 | 0 | 3 |
| ENSMUSG00000020297 | Nsg2 | ENSMUST00000020537 | 24.4260204 | 26 | 21.2372449 | 27.9336735 | 0 | 0 | 0 | 0 | 0 | 0 | 0 | 0 | 0 | 0 | 0 | 1 | 0 | 0 | 1 |
| ENSMUSG00000062510 | Nsl1 | ENSMUST00000078259 | 24.2236025 | 27 | 22.0372671 | 26.5093168 | 0 | 0 | 0 | 0 | 0 | 0 | 0 | 0 | 0 | 0 | 0 | 0 | 6 | 0 | 6 |
| ENSMUSG00000040331 | Nsmce4a | ENSMUST00000161076 | 30.4439746 | 27 | 20.9302326 | 21.987315 | 0 | 0 | 0 | 0 | 0 | 0 | 0 | 0 | 0 | 0 | 0 | 1 | 0 | 0 | 1 |
| ENSMUSG00000037958 | Nsrp1 | ENSMUST00000102494 | 27.3480663 | 34 | 16.5745856 | 21.6850829 | 2 | 1 | 0 | 1 | 0 | 0 | 0 | 0 | 1 | 2 | 1 | 0 | 6 | 2 | 14 |
| ENSMUSG00000050312 | Nsun3 | ENSMUST00000143314 | 28.986403 | 32 | 19.4684796 | 19.6229913 | 2 | 1 | 0 | 1 | 0 | 0 | 0 | 0 | 1 | 2 | 0 | 0 | 4 | 0 | 9 |
| ENSMUSG00000026707 | Nsun6 | ENSMUST00000114713 | 30.808656 | 33 | 16.856492 | 19.8177677 | 3 | 3 | 0 | 5 | 0 | 1 | 1 | 2 | 2 | 11 | 1 | 0 | 0 | 0 | 18 |
| ENSMUSG00000055254 | Ntrk2 | ENSMUST00000079828 | 27.0785117 | 28 | 21.5061421 | 23.1084209 | 10 | 5 | 0 | 17 | 2 | 3 | 5 | 4 | 8 | 39 | 2 | 4 | 0 | 0 | 60 |
| ENSMUSG00000026434 | Nucks1 | ENSMUST00000062264 | 25.2785924 | 36 | 18.2209189 | 20.2737048 | 0 | 0 | 0 | 0 | 0 | 0 | 0 | 0 | 0 | 0 | 5 | 3 | 24 | 2 | 34 |
| ENSMUSG00000038736 | Nudcd1 | ENSMUST00000038719 | 31.7052271 | 34 | 16.3667524 | 18.251928 | 0 | 1 | 0 | 0 | 0 | 0 | 0 | 0 | 0 | 0 | 0 | 0 | 0 | 2 | 3 |
| ENSMUSG00000020328 | Nudcd2 | ENSMUST00000020578 | 33.7064677 | 33 | 13.9303483 | 19.7761194 | 0 | 0 | 1 | 0 | 0 | 0 | 0 | 0 | 0 | 1 | 0 | 1 | 0 | 0 | 2 |
| ENSMUSG00000073293 | Nudt10 | ENSMUST00000103006 | 30.8181818 | 34 | 16.2727273 | 18.6363636 | 0 | 0 | 0 | 0 | 0 | 0 | 0 | 0 | 0 | 0 | 0 | 0 | 1 | 0 | 1 |
| ENSMUSG00000024228 | Nudt12 | ENSMUST00000025065 | 30.7452204 | 33 | 19.1962544 | 17.0503316 | 2 | 1 | 0 | 1 | 0 | 0 | 0 | 0 | 1 | 2 | 0 | 22 | 2 | 0 | 29 |
| ENSMUSG00000022516 | Nudt16l1 | ENSMUST00000050881 | 18.9054726 | 29 | 26.3681592 | 26.0364842 | 0 | 0 | 0 | 0 | 0 | 0 | 0 | 0 | 0 | 0 | 0 | 1 | 0 | 0 | 1 |
| ENSMUSG00000025817 | Nudt5 | ENSMUST00000026927 | 29.3751691 | 30 | 20.5572085 | 20.4490127 | 0 | 0 | 0 | 1 | 0 | 0 | 0 | 0 | 1 | 2 | 0 | 1 | 12 | 0 | 15 |
| ENSMUSG00000022009 | Nufip1 | ENSMUST00000022586 | 27.7755349 | 30 | 19.5397658 | 22.2446508 | 7 | 4 | 0 | 20 | 4 | 5 | 6 | 7 | 7 | 49 | 0 | 1 | 5 | 2 | 68 |
| ENSMUSG00000037857 | Nufip2 | ENSMUST00000100802 | 27.7777778 | 37 | 16.0013307 | 18.762475 | 0 | 0 | 0 | 0 | 0 | 0 | 0 | 0 | 0 | 0 | 3 | 1 | 16 | 5 | 25 |
| ENSMUSG00000052798 | Nup107 | ENSMUST00000167943 | 22.4576271 | 46 | 11.8644068 | 19.4915254 | 0 | 0 | 0 | 0 | 0 | 0 | 0 | 0 | 0 | 0 | 4 | 0 | 3 | 0 | 7 |
| ENSMUSG00000022142 | Nup155 | ENSMUST00000163765 | 26.9947444 | 36 | 19.0635452 | 17.7735308 | 1 | 0 | 0 | 0 | 0 | 0 | 0 | 0 | 0 | 0 | 0 | 6 | 6 | 0 | 13 |

| | | | | | | | | | | | | | | | | | | | | | |
|---|---|---|---|---|---|---|---|---|---|---|---|---|---|---|---|---|---|---|---|---|---|
| ENSMUSG00000051329 | Nup160 | ENSMUST00000057481 | 30.7038835 | 35 | 18.3252427 | 16.1407767 | 0 | 0 | 0 | 0 | 0 | 0 | 0 | 0 | 0 | 0 | 1 | 0 | 0 | 0 | 1 |
| ENSMUSG00000034826 | Nup54 | ENSMUST00000038514 | 29.4029851 | 34 | 18.358209 | 18.0597015 | 0 | 0 | 0 | 0 | 0 | 0 | 0 | 0 | 0 | 0 | 0 | 0 | 3 | 0 | 3 |
| ENSMUSG00000032939 | Nup93 | ENSMUST00000109547 | 18.8679245 | 44 | 11.8598383 | 25.3369272 | 0 | 0 | 0 | 0 | 0 | 0 | 0 | 0 | 0 | 0 | 0 | 0 | 2 | 0 | 2 |
| ENSMUSG00000023068 | Nus1 | ENSMUST00000023830 | 30.5815603 | 33 | 16.4822695 | 20.141844 | 0 | 0 | 0 | 1 | 0 | 0 | 0 | 0 | 1 | 2 | 0 | 0 | 0 | 1 | 3 |
| ENSMUSG00000027306 | Nusap1 | ENSMUST00000028771 | 29.4456763 | 31 | 20.7538803 | 18.8026608 | 0 | 0 | 0 | 0 | 0 | 0 | 0 | 0 | 0 | 0 | 0 | 0 | 8 | 0 | 8 |
| ENSMUSG00000026516 | Nvl | ENSMUST00000027797 | 25.2415638 | 29 | 23.9928646 | 22.1495466 | 0 | 0 | 0 | 0 | 0 | 0 | 0 | 0 | 0 | 0 | 0 | 2 | 3 | 2 | 7 |
| ENSMUSG00000075033 | Nxpe3 | ENSMUST00000099705 | 29.1735733 | 32 | 19.2517168 | 19.8910727 | 2 | 1 | 2 | 4 | 0 | 0 | 0 | 1 | 3 | 10 | 0 | 1 | 1 | 0 | 15 |
| ENSMUSG00000069132 | Nxph2 | ENSMUST00000102945 | 31.6129032 | 36 | 14.7096774 | 17.6774194 | 0 | 0 | 0 | 1 | 0 | 0 | 0 | 0 | 1 | 2 | 0 | 0 | 0 | 0 | 2 |
| ENSMUSG00000042271 | Nxt2 | ENSMUST00000042329 | 31.9034853 | 35 | 17.1581769 | 15.8713137 | 0 | 0 | 0 | 1 | 0 | 0 | 0 | 0 | 1 | 2 | 0 | 0 | 0 | 0 | 2 |
| ENSMUSG00000054976 | Nyap2 | ENSMUST00000123285 | 33.3067888 | 34 | 15.3892096 | 17.3999602 | 1 | 0 | 0 | 2 | 0 | 0 | 0 | 0 | 2 | 4 | 0 | 0 | 1 | 0 | 6 |
| ENSMUSG00000032661 | Oas3 | ENSMUST00000044833 | 20.0631912 | 35 | 21.2480253 | 24.1706161 | 0 | 0 | 1 | 0 | 0 | 0 | 0 | 0 | 0 | 1 | 3 | 0 | 0 | 0 | 4 |
| ENSMUSG00000030934 | Oat | ENSMUST00000084500 | 27.9136691 | 34 | 16.6906475 | 21.5827338 | 1 | 0 | 0 | 0 | 0 | 0 | 0 | 0 | 0 | 0 | 0 | 0 | 0 | 0 | 1 |
| ENSMUSG00000029152 | Ociad1 | ENSMUST00000166823 | 26.7052023 | 33 | 18.8439306 | 21.734104 | 0 | 0 | 0 | 1 | 0 | 0 | 0 | 0 | 1 | 2 | 1 | 1 | 0 | 0 | 4 |
| ENSMUSG00000021638 | Ocln | ENSMUST00000069756 | 27.5249241 | 36 | 17.3818812 | 19.4191591 | 0 | 0 | 0 | 1 | 0 | 0 | 0 | 0 | 1 | 2 | 0 | 1 | 7 | 1 | 11 |
| ENSMUSG00000033009 | Ogfod1 | ENSMUST00000093301 | 25.1884948 | 29 | 23.1778833 | 22.8427813 | 0 | 0 | 0 | 0 | 0 | 0 | 0 | 0 | 0 | 0 | 0 | 1 | 4 | 0 | 5 |
| ENSMUSG00000026158 | Ogfrl1 | ENSMUST00000027343 | 30.7368421 | 34 | 16.9022556 | 18.0451128 | 0 | 0 | 0 | 0 | 0 | 0 | 0 | 0 | 0 | 0 | 0 | 0 | 1 | 1 | 2 |
| ENSMUSG00000021390 | Ogn | ENSMUST00000021822 | 34.4759394 | 31 | 19.5121951 | 14.8978247 | 0 | 0 | 0 | 0 | 0 | 0 | 0 | 0 | 0 | 0 | 0 | 0 | 1 | 0 | 1 |
| ENSMUSG00000034160 | Ogt | ENSMUST00000044475 | 27.2637308 | 35 | 17.714003 | 20.2375062 | 0 | 0 | 0 | 0 | 0 | 0 | 0 | 0 | 0 | 0 | 0 | 0 | 2 | 1 | 3 |
| ENSMUSG00000027108 | Ola1 | ENSMUST00000028517 | 29.8053528 | 33 | 16.0583942 | 21.4111922 | 0 | 0 | 0 | 0 | 0 | 0 | 0 | 0 | 0 | 0 | 0 | 0 | 1 | 0 | 1 |
| ENSMUSG00000027965 | Olfm3 | ENSMUST00000081752 | 34.0314136 | 36 | 15.4973822 | 14.7643979 | 0 | 0 | 0 | 0 | 0 | 0 | 0 | 0 | 0 | 0 | 1 | 0 | 3 | 0 | 4 |
| ENSMUSG00000043013 | Onecut1 | ENSMUST00000056006 | 34.0815392 | 26 | 20.6596427 | 19.0563445 | 0 | 0 | 0 | 1 | 0 | 0 | 0 | 0 | 1 | 2 | 1 | 0 | 0 | 0 | 3 |
| ENSMUSG00000045991 | Onecut2 | ENSMUST00000175965 | 30.2606055 | 31 | 19.9881926 | 18.88336 | 5 | 1 | 0 | 6 | 0 | 0 | 0 | 0 | 6 | 12 | 3 | 1 | 5 | 3 | 30 |
| ENSMUSG00000045518 | Onecut3 | ENSMUST00000051773 | 26.8690096 | 21 | 30.543131 | 21.5015974 | 0 | 0 | 0 | 0 | 0 | 0 | 0 | 0 | 0 | 0 | 6 | 3 | 7 | 0 | 16 |
| ENSMUSG00000026037 | Orc2 | ENSMUST00000027198 | 27.0760642 | 30 | 20.6559665 | 22.1214236 | 0 | 0 | 1 | 0 | 0 | 0 | 0 | 0 | 0 | 1 | 0 | 0 | 0 | 0 | 1 |
| ENSMUSG00000026761 | Orc4 | ENSMUST00000028098 | 33.5910321 | 31 | 17.3946656 | 17.6265945 | 0 | 0 | 0 | 1 | 0 | 0 | 0 | 0 | 1 | 2 | 0 | 0 | 0 | 0 | 2 |
| ENSMUSG00000031697 | Orc6 | ENSMUST00000034132 | 27.1126761 | 33 | 19.0140845 | 21.3028169 | 0 | 0 | 0 | 0 | 0 | 0 | 0 | 0 | 0 | 0 | 0 | 0 | 1 | 0 | 1 |
| ENSMUSG00000026097 | Ormdl1 | ENSMUST00000027266 | 29.4729027 | 33 | 16.8522643 | 20.7126949 | 0 | 0 | 0 | 0 | 0 | 0 | 0 | 0 | 0 | 0 | 0 | 1 | 0 | 0 | 1 |
| ENSMUSG00000020189 | Osbpl8 | ENSMUST00000095310 | 30.3504847 | 36 | 15.6599553 | 18.0213771 | 1 | 0 | 0 | 2 | 0 | 0 | 0 | 0 | 2 | 4 | 0 | 0 | 9 | 1 | 15 |
| ENSMUSG00000038280 | Ostm1 | ENSMUST00000040718 | 29.6199896 | 32 | 17.7511713 | 20.5622072 | 0 | 0 | 0 | 0 | 0 | 0 | 0 | 0 | 0 | 0 | 0 | 0 | 1 | 1 | 2 |
| ENSMUSG00000036990 | Otud4 | ENSMUST00000173078 | 26.6969535 | 37 | 16.6755746 | 20.1229289 | 1 | 0 | 0 | 0 | 0 | 0 | 0 | 0 | 0 | 0 | 3 | 0 | 7 | 0 | 11 |
| ENSMUSG00000022307 | Oxr1 | ENSMUST00000022918 | 30.8262108 | 34 | 16.7521368 | 18.9173789 | 0 | 0 | 0 | 0 | 0 | 0 | 0 | 0 | 0 | 0 | 0 | 0 | 3 | 0 | 3 |
| ENSMUSG00000021786 | Oxsm | ENSMUST00000022311 | 29.1749503 | 32 | 19.0357853 | 20.27833 | 1 | 1 | 0 | 1 | 0 | 0 | 0 | 0 | 1 | 2 | 0 | 2 | 0 | 0 | 6 |
| ENSMUSG00000036737 | Oxsr1 | ENSMUST00000040853 | 25.7195572 | 29 | 21.9557196 | 23.3210332 | 0 | 0 | 0 | 0 | 0 | 0 | 0 | 0 | 0 | 0 | 0 | 2 | 2 | 0 | 4 |
| ENSMUSG00000027765 | P2ry1 | ENSMUST00000029331 | 28.38923 | 36 | 14.9267832 | 20.8313651 | 0 | 0 | 0 | 0 | 0 | 0 | 0 | 0 | 0 | 0 | 5 | 0 | 9 | 0 | 14 |
| ENSMUSG00000036353 | P2ry12 | ENSMUST00000196583 | 38.8473853 | 32 | 16.0085379 | 13.3404482 | 0 | 0 | 0 | 0 | 0 | 0 | 0 | 0 | 0 | 0 | 0 | 1 | 0 | 0 | 1 |
| ENSMUSG00000019916 | P4ha1 | ENSMUST00000009789 | 29.3721973 | 28 | 20.4035874 | 22.2869955 | 0 | 0 | 0 | 0 | 0 | 0 | 0 | 0 | 0 | 0 | 1 | 0 | 0 | 3 | 4 |
| ENSMUSG00000090919 | Pabpc4l | ENSMUST00000166505 | 30.220626 | 38 | 16.8804515 | 15.2129297 | 1 | 1 | 0 | 1 | 0 | 0 | 0 | 0 | 1 | 2 | 1 | 0 | 14 | 0 | 19 |
| ENSMUSG00000034732 | Pabpc5 | ENSMUST00000040961 | 31.9417476 | 41 | 14.5631068 | 12.7184466 | 2 | 1 | 0 | 3 | 0 | 0 | 0 | 1 | 2 | 6 | 0 | 0 | 1 | 0 | 10 |
| ENSMUSG00000022194 | Pabpn1 | ENSMUST00000116476 | 21.9679634 | 29 | 23.1121281 | 26.0869565 | 0 | 0 | 0 | 0 | 0 | 0 | 0 | 0 | 0 | 0 | 0 | 3 | 0 | 0 | 3 |
| ENSMUSG00000029089 | Pacrgl | ENSMUST00000200566 | 26.7672008 | 32 | 19.3213949 | 21.6776626 | 0 | 0 | 0 | 0 | 0 | 0 | 0 | 0 | 0 | 0 | 2 | 0 | 0 | 0 | 2 |
| ENSMUSG00000020745 | Pafah1b1 | ENSMUST00000021091 | 28.2526882 | 33 | 17.9301075 | 20.4032258 | 0 | 0 | 0 | 0 | 0 | 0 | 0 | 0 | 0 | 0 | 2 | 0 | 1 | 0 | 3 |
| ENSMUSG00000003131 | Pafah1b2 | ENSMUST00000172450 | 24.5780278 | 35 | 19.5439739 | 20.9357418 | 0 | 0 | 0 | 0 | 0 | 0 | 0 | 0 | 0 | 0 | 2 | 3 | 4 | 0 | 9 |
| ENSMUSG00000027508 | Pag1 | ENSMUST00000161949 | 26.2881679 | 30 | 22.4077608 | 21.2945293 | 2 | 0 | 0 | 0 | 0 | 0 | 0 | 0 | 0 | 0 | 0 | 31 | 11 | 0 | 44 |
| ENSMUSG00000025451 | Paip1 | ENSMUST00000026520 | 29.5514512 | 41 | 14.6877748 | 15.1275286 | 0 | 0 | 0 | 0 | 0 | 0 | 0 | 0 | 0 | 0 | 2 | 0 | 22 | 2 | 26 |
| ENSMUSG00000037058 | Paip2 | ENSMUST00000041314 | 30.7196563 | 32 | 17.83029 | 19.8711063 | 0 | 0 | 0 | 2 | 0 | 0 | 0 | 0 | 2 | 4 | 0 | 0 | 0 | 0 | 4 |
| ENSMUSG00000022781 | Pak2 | ENSMUST00000023467 | 25.8097686 | 33 | 20.3856041 | 20.5912596 | 1 | 0 | 0 | 0 | 0 | 0 | 0 | 0 | 0 | 0 | 1 | 2 | 6 | 2 | 12 |
| ENSMUSG00000031284 | Pak3 | ENSMUST00000172330 | 30.4388133 | 33 | 17.7224969 | 19.236712 | 1 | 1 | 0 | 2 | 0 | 0 | 0 | 0 | 2 | 4 | 2 | 0 | 2 | 1 | 11 |
| ENSMUSG00000033610 | Pank1 | ENSMUST00000036584 | 27.3361227 | 34 | 19.2468619 | 19.525802 | 0 | 0 | 0 | 0 | 0 | 0 | 0 | 0 | 0 | 0 | 0 | 3 | 0 | 0 | 3 |
| ENSMUSG00000037514 | Pank2 | ENSMUST00000184932 | 26.031746 | 28 | 21.8412698 | 23.6825397 | 1 | 0 | 0 | 1 | 0 | 0 | 0 | 0 | 1 | 2 | 0 | 0 | 2 | 1 | 6 |
| ENSMUSG00000018846 | Pank3 | ENSMUST00000018990 | 28.8313026 | 35 | 17.8136262 | 18.5000858 | 1 | 0 | 0 | 0 | 0 | 0 | 0 | 0 | 0 | 0 | 4 | 3 | 6 | 0 | 14 |
| ENSMUSG00000042167 | Papd4 | ENSMUST00000048702 | 32.460733 | 43 | 14.1361257 | 9.94764398 | 0 | 0 | 0 | 0 | 0 | 0 | 0 | 0 | 0 | 0 | 0 | 0 | 0 | 2 | 2 |
| ENSMUSG00000036779 | Papd5 | ENSMUST00000118952 | 27.2578668 | 34 | 17.6951369 | 20.9644463 | 0 | 0 | 0 | 1 | 0 | 0 | 0 | 0 | 1 | 2 | 0 | 1 | 0 | 2 | 5 |
| ENSMUSG00000021111 | Papola | ENSMUST00000169938 | 28.9552239 | 30 | 21.5739484 | 19.8100407 | 0 | 0 | 0 | 0 | 0 | 0 | 0 | 0 | 0 | 0 | 0 | 0 | 1 | 0 | 1 |
| ENSMUSG00000020273 | Papolg | ENSMUST00000020513 | 27.6676829 | 37 | 17.3018293 | 18.2926829 | 0 | 0 | 0 | 0 | 0 | 0 | 0 | 0 | 0 | 0 | 0 | 0 | 9 | 0 | 9 |
| ENSMUSG00000028370 | Pappa | ENSMUST00000084501 | 29.9273859 | 31 | 19.5193638 | 19.813278 | 0 | 0 | 0 | 1 | 0 | 0 | 0 | 0 | 1 | 2 | 0 | 14 | 0 | 0 | 16 |
| ENSMUSG00000044641 | Pard6b | ENSMUST00000052125 | 21.6256899 | 31 | 22.7295534 | 24.8369293 | 0 | 0 | 0 | 0 | 0 | 0 | 0 | 0 | 0 | 0 | 2 | 4 | 0 | 0 | 6 |
| ENSMUSG00000022685 | Parn | ENSMUST00000058884 | 20.6546275 | 30 | 23.5891648 | 25.282167 | 0 | 0 | 0 | 0 | 0 | 0 | 0 | 0 | 0 | 0 | 0 | 1 | 0 | 0 | 1 |
| ENSMUSG00000037997 | Parp11 | ENSMUST00000039680 | 23.2779097 | 30 | 21.6943785 | 24.6239113 | 0 | 0 | 0 | 0 | 0 | 0 | 0 | 0 | 0 | 0 | 7 | 2 | 5 | 0 | 14 |
| ENSMUSG00000035365 | Parpbp | ENSMUST00000048518 | 29.1637386 | 32 | 19.7470134 | 18.9739986 | 0 | 1 | 0 | 0 | 0 | 0 | 0 | 0 | 0 | 0 | 2 | 0 | 3 | 0 | 6 |
| ENSMUSG00000046139 | Patl1 | ENSMUST00000061618 | 24.5986239 | 35 | 23.5665138 | 17.2018349 | 1 | 0 | 0 | 0 | 0 | 0 | 0 | 0 | 0 | 0 | 0 | 1 | 0 | 0 | 2 |
| ENSMUSG00000031226 | Pbdc1 | ENSMUST00000033577 | 29.2990419 | 33 | 17.1457388 | 20.3227433 | 1 | 0 | 0 | 1 | 0 | 0 | 0 | 0 | 1 | 2 | 0 | 0 | 0 | 0 | 3 |
| ENSMUSG00000042323 | Pbrm1 | ENSMUST00000112095 | 31.3638053 | 33 | 16.5366035 | 18.8034188 | 3 | 1 | 0 | 2 | 0 | 0 | 0 | 1 | 1 | 4 | 0 | 1 | 0 | 1 | 10 |
| ENSMUSG00000052534 | Pbx1 | ENSMUST00000176540 | 26.8978805 | 31 | 21.2524085 | 20.7514451 | 0 | 0 | 0 | 1 | 0 | 0 | 0 | 0 | 1 | 2 | 0 | 7 | 3 | 2 | 14 |
| ENSMUSG00000038718 | Pbx3 | ENSMUST00000113132 | 26.7307692 | 30 | 22.6282051 | 20.3205128 | 0 | 0 | 0 | 0 | 0 | 0 | 0 | 0 | 0 | 0 | 0 | 2 | 0 | 0 | 2 |
| ENSMUSG00000034755 | Pcdh11x | ENSMUST00000113358 | 33.4269006 | 36 | 13.9181287 | 16.374269 | 2 | 0 | 1 | 3 | 0 | 0 | 0 | 0 | 3 | 7 | 0 | 0 | 1 | 0 | 10 |
| ENSMUSG00000050505 | Pcdh20 | ENSMUST00000061628 | 32.1981424 | 35 | 15.5683326 | 16.9394073 | 0 | 0 | 1 | 0 | 0 | 0 | 0 | 0 | 0 | 1 | 0 | 1 | 4 | 2 | 8 |
| ENSMUSG00000036422 | Pcdh8 | ENSMUST00000039568 | 29.6979866 | 39 | 14.261745 | 17.4496644 | 0 | 0 | 1 | 0 | 0 | 0 | 0 | 0 | 0 | 1 | 0 | 0 | 0 | 0 | 1 |
| ENSMUSG00000055421 | Pcdh9 | ENSMUST00000195826 | 32.7307326 | 35 | 15.2235966 | 17.1741199 | 0 | 0 | 1 | 0 | 0 | 0 | 0 | 0 | 0 | 1 | 0 | 1 | 0 | 0 | 2 |
| ENSMUSG00000044043 | Pcdhb14 | ENSMUST00000052387 | 28.8703613 | 37 | 17.5085826 | 16.7892758 | 0 | 0 | 0 | 1 | 0 | 0 | 0 | 0 | 1 | 2 | 2 | 28 | 9 | 2 | 43 |
| ENSMUSG00000048347 | Pcdhb18 | ENSMUST00000055949 | 26.5550239 | 40 | 18.6204147 | 14.6331738 | 1 | 0 | 1 | 1 | 0 | 0 | 0 | 0 | 1 | 3 | 1 | 0 | 2 | 0 | 7 |
| ENSMUSG00000043313 | Pcdhb19 | ENSMUST00000059571 | 31.4948566 | 33 | 17.4874152 | 18.3409937 | 1 | 0 | 0 | 1 | 0 | 0 | 0 | 0 | 1 | 2 | 2 | 0 | 0 | 1 | 6 |
| ENSMUSG00000046191 | Pcdhb20 | ENSMUST00000052179 | 30 | 37 | 15.9 | 17 | 0 | 0 | 0 | 1 | 0 | 0 | 0 | 0 | 1 | 2 | 0 | 2 | 2 | 0 | 6 |
| ENSMUSG00000073591 | Pcdhb22 | ENSMUST00000192409 | 27.4488102 | 31 | 17.3491976 | 23.7686774 | 0 | 0 | 0 | 0 | 0 | 0 | 0 | 0 | 0 | 0 | 2 | 1 | 1 | 1 | 5 |
| ENSMUSG00000045062 | Pcdhb7 | ENSMUST00000053037 | 29.5555556 | 35 | 17.2222222 | 18.6666667 | 0 | 0 | 0 | 0 | 0 | 0 | 0 | 0 | 0 | 0 | 2 | 0 | 1 | 0 | 3 |
| ENSMUSG00000041328 | Pcf11 | ENSMUST00000119954 | 35.0588235 | 36 | 12.5882353 | 16.5882353 | 0 | 0 | 0 | 0 | 0 | 0 | 0 | 0 | 0 | 0 | 0 | 0 | 0 | 1 | 1 |
| ENSMUSG00000033623 | Pcgf3 | ENSMUST00000046975 | 26.9780889 | 31 | 21.5003043 | 20.4503956 | 2 | 0 | 1 | 0 | 0 | 0 | 0 | 0 | 0 | 1 | 0 | 1 | 3 | 0 | 7 |
| ENSMUSG00000025050 | Pcgf6 | ENSMUST00000026032 | 29.2079208 | 31 | 20.990099 | 19.1089109 | 1 | 0 | 0 | 1 | 0 | 0 | 0 | 0 | 1 | 2 | 0 | 0 | 0 | 0 | 3 |
| ENSMUSG00000031592 | Pcm1 | ENSMUST00000045218 | 31.3282002 | 36 | 16.8912416 | 16.26564 | 0 | 0 | 0 | 0 | 0 | 0 | 0 | 0 | 0 | 0 | 0 | 1 | 1 | 0 | 2 |
| ENSMUSG00000051285 | Pcmtd1 | ENSMUST00000061280 | 31.0464495 | 34 | 15.9636946 | 19.3539776 | 0 | 0 | 0 | 0 | 0 | 0 | 0 | 0 | 0 | 0 | 0 | 1 | 0 | 0 | 1 |
| ENSMUSG00000027589 | Pcmtd2 | ENSMUST00000108754 | 29.7738693 | 34 | 16.4991625 | 19.7654941 | 0 | 1 | 0 | 0 | 0 | 0 | 0 | 0 | 0 | 0 | 0 | 0 | 1 | 0 | 2 |
| ENSMUSG00000071533 | Pcnp | ENSMUST00000125040 | 29.4088176 | 32 | 18.0360721 | 20.6913828 | 0 | 0 | 0 | 0 | 0 | 0 | 0 | 0 | 0 | 0 | 0 | 0 | 10 | 4 | 14 |
| ENSMUSG00000034501 | Pcnxl4 | ENSMUST00000044352 | 29.4832827 | 35 | 15.3495441 | 20.212766 | 1 | 0 | 0 | 0 | 0 | 0 | 0 | 0 | 0 | 0 | 0 | 0 | 0 | 0 | 1 |
| ENSMUSG00000024713 | Pcsk5 | ENSMUST00000050715 | 28.5714286 | 32 | 21.1350294 | 18.4344423 | 0 | 1 | 0 | 1 | 0 | 0 | 0 | 0 | 1 | 2 | 1 | 5 | 4 | 1 | 14 |
| ENSMUSG00000027835 | Pdcd10 | ENSMUST00000161137 | 33.7864078 | 36 | 13.3980583 | 16.407767 | 0 | 0 | 0 | 0 | 0 | 0 | 0 | 0 | 0 | 0 | 1 | 0 | 0 | 0 | 1 |
| ENSMUSG00000021576 | Pdcd6 | ENSMUST00000022060 | 30.2774427 | 29 | 18.8178528 | 22.0747889 | 0 | 0 | 0 | 0 | 0 | 0 | 0 | 0 | 0 | 0 | 0 | 0 | 2 | 0 | 2 |
| ENSMUSG00000032504 | Pdcd6ip | ENSMUST00000035086 | 26.3091878 | 31 | 21.1351521 | 21.6368768 | 0 | 0 | 0 | 1 | 0 | 0 | 0 | 0 | 1 | 2 | 0 | 0 | 3 | 0 | 5 |
| ENSMUSG00000009030 | Pdcl | ENSMUST00000009174 | 28.1730222 | 37 | 16.1639158 | 18.554354 | 0 | 0 | 1 | 0 | 0 | 0 | 0 | 0 | 0 | 1 | 0 | 0 | 2 | 1 | 4 |
| ENSMUSG00000026078 | Pdcl3 | ENSMUST00000027247 | 31.0994764 | 34 | 15.8115183 | 19.5811518 | 0 | 0 | 0 | 1 | 0 | 0 | 0 | 0 | 1 | 2 | 0 | 0 | 0 | 0 | 2 |
| ENSMUSG00000043702 | Pde12 | ENSMUST00000052932 | 31.0041315 | 33 | 18.2863302 | 17.7474403 | 0 | 0 | 0 | 2 | 0 | 0 | 0 | 0 | 2 | 4 | 0 | 0 | 0 | 1 | 5 |
| ENSMUSG00000059173 | Pde1a | ENSMUST00000102655 | 29.5029821 | 32 | 18.6481113 | 20.1988072 | 0 | 0 | 0 | 0 | 0 | 0 | 0 | 0 | 0 | 0 | 1 | 0 | 0 | 0 | 1 |
| ENSMUSG00000004347 | Pde1c | ENSMUST00000166102 | 29.1692692 | 31 | 19.4253592 | 20.8619613 | 0 | 0 | 0 | 1 | 0 | 0 | 0 | 0 | 1 | 2 | 0 | 4 | 2 | 0 | 8 |
| ENSMUSG00000041741 | Pde3a | ENSMUST00000043259 | 30.0526008 | 33 | 18.012858 | 19.0882525 | 2 | 0 | 0 | 2 | 0 | 0 | 0 | 0 | 2 | 4 | 1 | 0 | 4 | 0 | 11 |

| | | | | | | | | | | | | | | | | | | | | | |
|---|---|---|---|---|---|---|---|---|---|---|---|---|---|---|---|---|---|---|---|---|---|
| ENSMUSG00000030671 | Pde3b | ENSMUST00000032909 | 31.968032 | 35 | 16.983017 | 16.3170163 | 2 | 0 | 0 | 0 | 0 | 0 | 0 | 0 | 0 | 0 | 0 | 0 | 1 | 0 | 3 |
| ENSMUSG00000028525 | Pde4b | ENSMUST00000106908 | 26.6145833 | 31 | 22.5 | 19.53125 | 0 | 0 | 0 | 1 | 0 | 0 | 0 | 0 | 1 | 2 | 0 | 2 | 0 | 0 | 4 |
| ENSMUSG00000069094 | Pde7a | ENSMUST00000099195 | 27.6501767 | 32 | 19.6113074 | 21.0909894 | 0 | 2 | 0 | 1 | 0 | 0 | 0 | 0 | 1 | 2 | 0 | 0 | 2 | 0 | 6 |
| ENSMUSG00000019990 | Pde7b | ENSMUST00000169016 | 26.4597191 | 26 | 25.2771619 | 22.0990392 | 1 | 0 | 0 | 0 | 0 | 0 | 0 | 0 | 0 | 0 | 0 | 0 | 0 | 0 | 1 |
| ENSMUSG00000028019 | Pdgfc | ENSMUST00000129285 | 29.0037831 | 29 | 19.840269 | 22.1941992 | 0 | 0 | 0 | 0 | 0 | 0 | 0 | 0 | 0 | 0 | 0 | 1 | 0 | 1 | 2 |
| ENSMUSG00000010914 | Pdhx | ENSMUST00000111183 | 31.3124108 | 31 | 16.5477889 | 21.0413695 | 0 | 0 | 0 | 1 | 0 | 0 | 0 | 0 | 1 | 2 | 0 | 1 | 0 | 0 | 3 |
| ENSMUSG00000050890 | Pdik1l | ENSMUST00000061234 | 29.8458803 | 31 | 18.3758151 | 20.5394191 | 2 | 0 | 0 | 1 | 0 | 0 | 0 | 0 | 1 | 2 | 0 | 1 | 4 | 3 | 12 |
| ENSMUSG00000019577 | Pdk4 | ENSMUST00000019721 | 26.9780744 | 32 | 18.6367969 | 22.0209724 | 0 | 0 | 1 | 0 | 0 | 0 | 0 | 0 | 0 | 1 | 0 | 23 | 0 | 0 | 24 |
| ENSMUSG00000028273 | Pdlim5 | ENSMUST00000029941 | 26.763754 | 35 | 17.7993528 | 20.776699 | 0 | 0 | 0 | 0 | 0 | 0 | 0 | 0 | 0 | 0 | 1 | 0 | 2 | 0 | 3 |
| ENSMUSG00000048371 | Pdp2 | ENSMUST00000059588 | 23.1799651 | 30 | 21.665696 | 24.6942341 | 0 | 0 | 0 | 0 | 0 | 0 | 0 | 0 | 0 | 0 | 0 | 0 | 16 | 0 | 16 |
| ENSMUSG00000024122 | Pdpk1 | ENSMUST00000102927 | 24.4855967 | 31 | 21.9042275 | 22.7459783 | 0 | 0 | 1 | 0 | 0 | 0 | 0 | 0 | 0 | 1 | 0 | 2 | 5 | 2 | 10 |
| ENSMUSG00000029202 | Pds5a | ENSMUST00000201948 | 28.3288651 | 32 | 18.9008043 | 20.4647006 | 0 | 0 | 0 | 0 | 0 | 0 | 0 | 0 | 0 | 0 | 1 | 2 | 7 | 0 | 10 |
| ENSMUSG00000034021 | Pds5b | ENSMUST00000016569 | 29.6973962 | 35 | 16.3617171 | 18.6136524 | 0 | 0 | 0 | 0 | 0 | 0 | 0 | 0 | 0 | 0 | 0 | 0 | 0 | 2 | 2 |
| ENSMUSG00000038240 | Pdss2 | ENSMUST00000160823 | 28.1976744 | 28 | 21.4389535 | 22.3110465 | 0 | 0 | 0 | 0 | 0 | 0 | 0 | 0 | 0 | 0 | 0 | 24 | 0 | 1 | 25 |
| ENSMUSG00000036218 | Pdzrn4 | ENSMUST00000035399 | 32.630273 | 36 | 16.3771712 | 15.3846154 | 0 | 0 | 0 | 0 | 0 | 0 | 0 | 0 | 0 | 0 | 1 | 0 | 0 | 0 | 1 |
| ENSMUSG00000020134 | Peli1 | ENSMUST00000101477 | 31 | 35 | 16.1666667 | 18.1111111 | 0 | 0 | 0 | 0 | 0 | 0 | 0 | 0 | 0 | 0 | 1 | 1 | 0 | 0 | 2 |
| ENSMUSG00000019851 | Perp | ENSMUST00000019998 | 29.3233083 | 31 | 17.0426065 | 22.3057644 | 0 | 0 | 0 | 0 | 0 | 0 | 0 | 0 | 0 | 0 | 0 | 0 | 1 | 2 | 3 |
| ENSMUSG00000005907 | Pex1 | ENSMUST00000143132 | 24.4803695 | 27 | 27.2517321 | 21.0161663 | 0 | 0 | 0 | 0 | 0 | 0 | 0 | 0 | 0 | 0 | 0 | 1 | 0 | 0 | 1 |
| ENSMUSG00000020283 | Pex13 | ENSMUST00000020523 | 30.8922858 | 35 | 15.6416637 | 18.556701 | 0 | 0 | 0 | 0 | 0 | 0 | 0 | 0 | 0 | 0 | 7 | 0 | 10 | 0 | 17 |
| ENSMUSG00000019809 | Pex3 | ENSMUST00000019945 | 31.2093628 | 34 | 17.6853056 | 16.7750325 | 0 | 0 | 0 | 1 | 0 | 0 | 0 | 0 | 1 | 2 | 0 | 0 | 0 | 0 | 2 |
| ENSMUSG00000005069 | Pex5 | ENSMUST00000112532 | 21.8560278 | 28 | 23.1569818 | 27.0598439 | 0 | 0 | 0 | 0 | 0 | 0 | 0 | 0 | 0 | 0 | 0 | 1 | 2 | 2 | 5 |
| ENSMUSG00000006412 | Pfdn2 | ENSMUST00000135941 | 25.4071661 | 31 | 22.2312704 | 21.4983713 | 0 | 0 | 0 | 0 | 0 | 0 | 0 | 0 | 0 | 0 | 0 | 0 | 3 | 0 | 3 |
| ENSMUSG00000026409 | Pfkfb2 | ENSMUST00000066863 | 23.7826394 | 33 | 19.3366267 | 23.8884968 | 0 | 0 | 0 | 0 | 0 | 0 | 0 | 0 | 0 | 0 | 1 | 1 | 0 | 0 | 2 |
| ENSMUSG00000027805 | Pfn2 | ENSMUST00000066882 | 25.3403933 | 35 | 19.9697428 | 20.0453858 | 0 | 0 | 0 | 0 | 0 | 0 | 0 | 0 | 0 | 0 | 0 | 1 | 1 | 0 | 2 |
| ENSMUSG00000029500 | Pgam5 | ENSMUST00000112505 | 25.5834054 | 32 | 19.9654278 | 22.3854797 | 0 | 0 | 1 | 0 | 0 | 0 | 0 | 0 | 0 | 1 | 1 | 2 | 1 | 0 | 5 |
| ENSMUSG00000073678 | Pgap1 | ENSMUST00000097739 | 28.3584025 | 34 | 17.2588174 | 20.5783195 | 1 | 1 | 0 | 2 | 0 | 0 | 0 | 0 | 2 | 4 | 0 | 0 | 3 | 2 | 11 |
| ENSMUSG00000024477 | Pggt1b | ENSMUST00000025354 | 32.3997454 | 33 | 15.5315086 | 18.6505411 | 0 | 0 | 0 | 0 | 0 | 0 | 0 | 0 | 0 | 0 | 0 | 0 | 1 | 0 | 1 |
| ENSMUSG00000056131 | Pgm3 | ENSMUST00000070064 | 33.9381003 | 30 | 17.6093917 | 18.5699039 | 0 | 0 | 0 | 1 | 0 | 0 | 0 | 0 | 1 | 2 | 0 | 0 | 0 | 0 | 2 |
| ENSMUSG00000006373 | Pgrmc1 | ENSMUST00000073339 | 32.3232323 | 31 | 17.5084175 | 18.9393939 | 0 | 0 | 0 | 0 | 0 | 0 | 0 | 0 | 0 | 0 | 1 | 0 | 0 | 0 | 1 |
| ENSMUSG00000049940 | Pgrmc2 | ENSMUST00000058578 | 26.3363755 | 29 | 21.8166015 | 22.9030856 | 0 | 0 | 0 | 0 | 0 | 0 | 0 | 0 | 0 | 0 | 0 | 3 | 1 | 0 | 4 |
| ENSMUSG00000066043 | Phactr4 | ENSMUST00000102568 | 23.1884058 | 29 | 26.342711 | 21.0997442 | 0 | 0 | 0 | 0 | 0 | 0 | 0 | 0 | 0 | 0 | 3 | 2 | 3 | 0 | 8 |
| ENSMUSG00000037652 | Phc3 | ENSMUST00000168645 | 27.6710609 | 32 | 18.9328311 | 21.6195857 | 1 | 1 | 0 | 3 | 0 | 0 | 0 | 0 | 3 | 6 | 1 | 1 | 0 | 2 | 12 |
| ENSMUSG00000047777 | Phf13 | ENSMUST00000055688 | 22.0833333 | 29 | 22.03125 | 26.7708333 | 0 | 0 | 0 | 0 | 0 | 0 | 0 | 0 | 0 | 0 | 0 | 0 | 0 | 1 | 1 |
| ENSMUSG00000029629 | Phf14 | ENSMUST00000090632 | 32.8527291 | 33 | 15.1390319 | 18.9495366 | 0 | 0 | 0 | 0 | 0 | 0 | 0 | 0 | 0 | 0 | 1 | 0 | 3 | 0 | 4 |
| ENSMUSG00000072501 | Phf20l1 | ENSMUST00000048188 | 32.6093514 | 33 | 16.1085973 | 17.918552 | 1 | 0 | 0 | 0 | 0 | 0 | 0 | 0 | 0 | 0 | 1 | 0 | 0 | 2 | 4 |
| ENSMUSG00000048874 | Phf3 | ENSMUST00000191329 | 31.4636957 | 21 | 23.127161 | 24.8943527 | 0 | 0 | 0 | 0 | 0 | 0 | 0 | 0 | 0 | 0 | 1 | 0 | 1 | 0 | 2 |
| ENSMUSG00000025626 | Phf6 | ENSMUST00000078944 | 28.4068466 | 35 | 18.0710994 | 18.4331797 | 0 | 0 | 0 | 0 | 0 | 0 | 0 | 0 | 0 | 0 | 0 | 0 | 1 | 0 | 1 |
| ENSMUSG00000041229 | Phf8 | ENSMUST00000046950 | 23.6430921 | 32 | 24.3832237 | 20.2302632 | 0 | 0 | 0 | 0 | 0 | 0 | 0 | 0 | 0 | 0 | 1 | 1 | 5 | 0 | 7 |
| ENSMUSG00000032253 | Phip | ENSMUST00000034787 | 32.8331688 | 35 | 14.5113524 | 17.828233 | 0 | 0 | 0 | 0 | 0 | 0 | 0 | 0 | 0 | 0 | 0 | 0 | 2 | 0 | 2 |
| ENSMUSG00000025537 | Phkg1 | ENSMUST00000026617 | 29.2156863 | 26 | 20.1960784 | 24.4117647 | 0 | 0 | 0 | 0 | 0 | 0 | 0 | 0 | 0 | 0 | 0 | 0 | 1 | 2 | 3 |
| ENSMUSG00000039987 | Phtf2 | ENSMUST00000118174 | 28.5953177 | 34 | 16.3043478 | 20.777592 | 2 | 1 | 1 | 1 | 0 | 0 | 0 | 0 | 1 | 3 | 1 | 0 | 5 | 0 | 12 |
| ENSMUSG00000026664 | Phyh | ENSMUST00000027975 | 35.1351351 | 29 | 16.2162162 | 20 | 0 | 0 | 0 | 0 | 0 | 0 | 0 | 0 | 0 | 0 | 0 | 1 | 0 | 0 | 1 |
| ENSMUSG00000067780 | Pi15 | ENSMUST00000088476 | 30.5076227 | 36 | 16.3511476 | 17.6076395 | 2 | 1 | 0 | 2 | 0 | 0 | 0 | 0 | 2 | 4 | 2 | 1 | 8 | 2 | 20 |
| ENSMUSG00000029186 | Pi4k2b | ENSMUST00000031081 | 32.0075758 | 33 | 15.9090909 | 18.8762626 | 0 | 0 | 0 | 1 | 0 | 0 | 0 | 0 | 1 | 2 | 0 | 0 | 2 | 0 | 4 |
| ENSMUSG00000030329 | Pianp | ENSMUST00000162000 | 16.8282548 | 28 | 32.2022161 | 22.9916898 | 0 | 0 | 0 | 0 | 0 | 0 | 0 | 0 | 0 | 0 | 0 | 6 | 0 | 0 | 6 |
| ENSMUSG00000032405 | Pias1 | ENSMUST00000098651 | 27.9217345 | 32 | 19.3019566 | 21.0999471 | 0 | 0 | 0 | 0 | 0 | 0 | 0 | 0 | 0 | 0 | 1 | 0 | 0 | 2 | 3 |
| ENSMUSG00000025423 | Pias2 | ENSMUST00000114777 | 27.5223061 | 36 | 18.2910089 | 18.2566918 | 1 | 0 | 0 | 0 | 0 | 0 | 0 | 0 | 0 | 0 | 0 | 0 | 0 | 2 | 3 |
| ENSMUSG00000039361 | Picalm | ENSMUST00000049537 | 29.7297297 | 35 | 16.6486486 | 19.0810811 | 1 | 1 | 0 | 1 | 0 | 0 | 0 | 0 | 1 | 2 | 0 | 0 | 12 | 2 | 18 |
| ENSMUSG00000031381 | Piga | ENSMUST00000151911 | 26.9527321 | 32 | 18.5048609 | 22.3935635 | 0 | 0 | 0 | 0 | 0 | 0 | 0 | 0 | 0 | 0 | 0 | 15 | 1 | 0 | 16 |
| ENSMUSG00000039047 | Pigk | ENSMUST00000159899 | 27.0347183 | 25 | 22.3392146 | 25.8110415 | 0 | 0 | 0 | 0 | 0 | 0 | 0 | 0 | 0 | 0 | 2 | 0 | 2 | 0 | 4 |
| ENSMUSG00000050229 | Pigm | ENSMUST00000052455 | 27.7859357 | 33 | 19.236417 | 20.1664219 | 1 | 0 | 0 | 1 | 0 | 0 | 0 | 0 | 1 | 2 | 5 | 1 | 14 | 2 | 25 |
| ENSMUSG00000056536 | Pign | ENSMUST00000070699 | 28.7548855 | 32 | 18.9838079 | 19.9329983 | 1 | 0 | 0 | 0 | 0 | 0 | 0 | 0 | 0 | 0 | 0 | 1 | 0 | 0 | 2 |
| ENSMUSG00000045140 | Pigw | ENSMUST00000067058 | 30.5185185 | 33 | 13.6296296 | 22.8148148 | 0 | 0 | 0 | 1 | 0 | 0 | 0 | 0 | 1 | 2 | 0 | 0 | 0 | 0 | 2 |
| ENSMUSG00000030660 | Pik3c2a | ENSMUST00000170430 | 33.1051873 | 33 | 16.462536 | 17.0389049 | 0 | 0 | 0 | 0 | 0 | 0 | 0 | 0 | 0 | 0 | 0 | 0 | 1 | 0 | 1 |
| ENSMUSG00000033628 | Pik3c3 | ENSMUST00000131405 | 28.5171103 | 24 | 22.0532319 | 25.4119138 | 0 | 0 | 0 | 0 | 0 | 0 | 0 | 0 | 0 | 0 | 0 | 1 | 0 | 0 | 1 |
| ENSMUSG00000032462 | Pik3cb | ENSMUST00000035037 | 25.1716247 | 30 | 22.6544622 | 21.935273 | 0 | 0 | 0 | 1 | 0 | 0 | 0 | 0 | 1 | 2 | 0 | 1 | 0 | 0 | 3 |
| ENSMUSG00000041417 | Pik3r1 | ENSMUST00000055518 | 25.9439426 | 32 | 19.3884642 | 22.2608293 | 0 | 0 | 0 | 0 | 0 | 0 | 0 | 0 | 0 | 0 | 0 | 19 | 11 | 2 | 32 |
| ENSMUSG00000025949 | Pikfyve | ENSMUST00000081154 | 26.7166295 | 33 | 18.594288 | 21.9161434 | 0 | 0 | 0 | 0 | 0 | 0 | 0 | 0 | 0 | 0 | 2 | 8 | 11 | 1 | 22 |
| ENSMUSG00000024014 | Pim1 | ENSMUST00000024811 | 21.1143695 | 31 | 23.8269795 | 24.4134897 | 4 | 2 | 0 | 6 | 0 | 1 | 2 | 2 | 2 | 13 | 0 | 1 | 12 | 1 | 33 |
| ENSMUSG00000028126 | Pip5k1a | ENSMUST00000107236 | 24.1468126 | 31 | 22.2150676 | 23.0521571 | 0 | 0 | 0 | 0 | 0 | 0 | 0 | 0 | 0 | 0 | 0 | 1 | 0 | 0 | 1 |
| ENSMUSG00000024867 | Pip5k1b | ENSMUST00000025800 | 29.2875989 | 30 | 21.2401055 | 19.2612137 | 1 | 1 | 0 | 1 | 0 | 0 | 0 | 0 | 1 | 2 | 0 | 0 | 0 | 0 | 4 |
| ENSMUSG00000024083 | Pja2 | ENSMUST00000172818 | 28.6129568 | 35 | 17.358804 | 19.0199336 | 0 | 0 | 0 | 1 | 0 | 0 | 0 | 0 | 1 | 2 | 2 | 0 | 3 | 0 | 7 |
| ENSMUSG00000034462 | Pkd2 | ENSMUST00000086831 | 28.531206 | 33 | 17.6912248 | 20.6945096 | 1 | 0 | 1 | 1 | 0 | 0 | 0 | 0 | 1 | 3 | 1 | 0 | 0 | 0 | 5 |
| ENSMUSG00000027499 | Pkia | ENSMUST00000028999 | 29.8534799 | 32 | 18.6813187 | 19.2918193 | 0 | 1 | 0 | 1 | 0 | 0 | 0 | 0 | 1 | 2 | 0 | 0 | 0 | 0 | 3 |
| ENSMUSG00000019876 | Pkib | ENSMUST00000075992 | 30.1156677 | 32 | 17.6235542 | 20.0630915 | 1 | 0 | 0 | 2 | 0 | 0 | 0 | 0 | 2 | 4 | 0 | 1 | 1 | 2 | 9 |
| ENSMUSG00000004591 | Pkn2 | ENSMUST00000043812 | 32.4853902 | 32 | 17.1880371 | 18.0818151 | 1 | 0 | 0 | 1 | 0 | 0 | 0 | 0 | 1 | 2 | 0 | 0 | 1 | 0 | 4 |
| ENSMUSG00000027999 | Pla2g12a | ENSMUST00000029629 | 26.5822785 | 31 | 20.8285386 | 21.9792865 | 1 | 0 | 0 | 1 | 0 | 0 | 0 | 0 | 1 | 2 | 0 | 0 | 0 | 0 | 3 |
| ENSMUSG00000033847 | Pla2g4c | ENSMUST00000043612 | 27.1209085 | 35 | 21.0420842 | 17.2344689 | 0 | 0 | 0 | 0 | 0 | 0 | 0 | 0 | 0 | 0 | 0 | 1 | 1 | 0 | 2 |
| ENSMUSG00000003282 | Plag1 | ENSMUST00000003369 | 29.8069785 | 33 | 17.5204157 | 19.5619896 | 0 | 0 | 0 | 0 | 0 | 0 | 0 | 0 | 0 | 0 | 0 | 0 | 2 | 0 | 2 |
| ENSMUSG00000019817 | Plagl1 | ENSMUST00000121646 | 25.5212355 | 39 | 19.7297297 | 16.0617761 | 0 | 0 | 0 | 0 | 0 | 0 | 0 | 0 | 0 | 0 | 1 | 0 | 0 | 0 | 1 |
| ENSMUSG00000020120 | Plek | ENSMUST00000102881 | 29.4497283 | 31 | 19.4972826 | 19.8029891 | 0 | 0 | 0 | 0 | 0 | 0 | 0 | 0 | 0 | 0 | 2 | 0 | 2 | 0 | 4 |
| ENSMUSG00000040268 | Plekha1 | ENSMUST00000120441 | 25.0792393 | 33 | 20.0871632 | 21.9096672 | 0 | 0 | 0 | 0 | 0 | 0 | 0 | 0 | 0 | 0 | 0 | 0 | 1 | 1 | 2 |
| ENSMUSG00000002733 | Plekha3 | ENSMUST00000111920 | 29.2609351 | 36 | 18.0241327 | 17.1945701 | 2 | 1 | 0 | 3 | 0 | 0 | 0 | 1 | 2 | 6 | 0 | 0 | 1 | 5 | 15 |
| ENSMUSG00000030231 | Plekha5 | ENSMUST00000087622 | 23.8659347 | 29 | 23.4325339 | 24.1548685 | 0 | 0 | 0 | 1 | 0 | 0 | 0 | 0 | 1 | 2 | 0 | 1 | 0 | 1 | 4 |
| ENSMUSG00000005225 | Plekha8 | ENSMUST00000119706 | 26.4100453 | 30 | 21.0168794 | 22.5195554 | 3 | 1 | 0 | 1 | 0 | 0 | 0 | 0 | 1 | 2 | 0 | 1 | 3 | 0 | 10 |
| ENSMUSG00000051344 | Plekhm3 | ENSMUST00000097713 | 24.8985801 | 28 | 24.5943205 | 22.3123732 | 2 | 1 | 2 | 2 | 0 | 0 | 0 | 1 | 1 | 6 | 5 | 6 | 7 | 2 | 29 |
| ENSMUSG00000030867 | Plk1 | ENSMUST00000033154 | 15.5477032 | 34 | 25.0883392 | 25.795053 | 0 | 0 | 0 | 0 | 0 | 0 | 0 | 0 | 0 | 0 | 0 | 1 | 0 | 0 | 1 |
| ENSMUSG00000025758 | Plk4 | ENSMUST00000203895 | 31.5710536 | 28 | 20.0224972 | 20.5099363 | 0 | 0 | 0 | 1 | 0 | 0 | 0 | 0 | 1 | 2 | 1 | 0 | 2 | 0 | 5 |
| ENSMUSG00000032374 | Plod2 | ENSMUST00000070522 | 30.8013356 | 33 | 17.1118531 | 19.115192 | 0 | 0 | 1 | 1 | 0 | 0 | 0 | 0 | 1 | 3 | 0 | 1 | 0 | 1 | 5 |
| ENSMUSG00000031425 | Plp1 | ENSMUST00000033800 | 32.3578134 | 30 | 18.332413 | 19.7128658 | 1 | 0 | 0 | 0 | 0 | 0 | 0 | 0 | 0 | 0 | 0 | 5 | 3 | 3 | 12 |
| ENSMUSG00000028517 | Plpp3 | ENSMUST00000064139 | 29.9081515 | 27 | 21.2399541 | 21.641791 | 0 | 0 | 0 | 0 | 0 | 0 | 0 | 0 | 0 | 0 | 0 | 0 | 4 | 0 | 4 |
| ENSMUSG00000040105 | Plpp6 | ENSMUST00000045674 | 29.8136646 | 29 | 20.8074534 | 20.0828157 | 0 | 0 | 1 | 0 | 0 | 0 | 0 | 0 | 0 | 1 | 0 | 0 | 0 | 0 | 1 |
| ENSMUSG00000063446 | Plppr1 | ENSMUST00000076670 | 28.4773344 | 33 | 18.3262301 | 19.759783 | 1 | 0 | 0 | 0 | 0 | 0 | 0 | 0 | 0 | 0 | 0 | 0 | 4 | 1 | 6 |
| ENSMUSG00000044667 | Plppr4 | ENSMUST00000061071 | 32.417781 | 32 | 17.3111673 | 18.5399349 | 0 | 0 | 0 | 0 | 0 | 0 | 0 | 0 | 0 | 0 | 0 | 0 | 3 | 0 | 3 |
| ENSMUSG00000033342 | Plppr5 | ENSMUST00000039564 | 33.3683106 | 32 | 18.0832459 | 16.7541098 | 1 | 0 | 0 | 0 | 0 | 0 | 0 | 0 | 0 | 0 | 0 | 2 | 0 | 0 | 3 |
| ENSMUSG00000019461 | Plscr3 | ENSMUST00000108633 | 22.3763571 | 25 | 26.8998794 | 25.6332931 | 0 | 0 | 0 | 0 | 0 | 0 | 0 | 0 | 0 | 0 | 0 | 1 | 0 | 0 | 1 |
| ENSMUSG00000026640 | Plxna2 | ENSMUST00000027952 | 29.6516568 | 27 | 22.4723874 | 21.2616822 | 0 | 0 | 1 | 0 | 0 | 0 | 0 | 0 | 0 | 1 | 0 | 0 | 1 | 0 | 2 |
| ENSMUSG00000074785 | Plxnc1 | ENSMUST00000099337 | 26.8534483 | 32 | 20.9482759 | 20.1293103 | 0 | 0 | 0 | 1 | 0 | 0 | 0 | 0 | 1 | 2 | 0 | 0 | 1 | 2 | 5 |
| ENSMUSG00000054659 | Pm20d2 | ENSMUST00000098181 | 22.9166667 | 39 | 15.625 | 22.9166667 | 0 | 0 | 0 | 0 | 0 | 0 | 0 | 0 | 0 | 0 | 1 | 0 | 1 | 0 | 2 |
| ENSMUSG00000024521 | Pmaip1 | ENSMUST00000025399 | 28.7092329 | 33 | 16.16904 | 22.1405604 | 1 | 0 | 0 | 1 | 0 | 0 | 0 | 0 | 1 | 2 | 0 | 0 | 8 | 0 | 11 |
| ENSMUSG00000028248 | Pnisr | ENSMUST00000098238 | 30.094919 | 38 | 14.9078727 | 17.1412619 | 2 | 1 | 0 | 1 | 0 | 0 | 0 | 0 | 1 | 2 | 0 | 6 | 0 | 1 | 12 |
| ENSMUSG00000026179 | Pnkd | ENSMUST00000087226 | 20.6466251 | 24 | 32.387975 | 22.802042 | 0 | 0 | 0 | 0 | 0 | 0 | 0 | 0 | 0 | 0 | 0 | 1 | 0 | 0 | 1 |

| | | | | | | | | | | | | | | | | | | | | | |
|---|---|---|---|---|---|---|---|---|---|---|---|---|---|---|---|---|---|---|---|---|---|
| ENSMUSG00000020464 | Pnpt1 | ENSMUST00000020756 | 32.4637681 | 41 | 14.7826087 | 11.3043478 | 1 | 0 | 0 | 1 | 0 | 0 | 0 | 0 | 1 | 2 | 0 | 1 | 0 | 0 | 4 |
| ENSMUSG00000028675 | Pnrc2 | ENSMUST00000030436 | 32.3905109 | 33 | 17.3357664 | 17.0620438 | 0 | 0 | 0 | 0 | 0 | 0 | 0 | 0 | 0 | 0 | 0 | 0 | 0 | 4 | 4 |
| ENSMUSG00000030726 | Pold3 | ENSMUST00000032969 | 24.6593121 | 29 | 22.8423102 | 23.880597 | 0 | 0 | 0 | 0 | 0 | 0 | 0 | 0 | 0 | 0 | 0 | 1 | 0 | 0 | 1 |
| ENSMUSG00000030042 | Pole4 | ENSMUST00000095786 | 26.096823 | 37 | 18.4568835 | 18.9107413 | 1 | 0 | 0 | 0 | 0 | 0 | 0 | 0 | 0 | 0 | 0 | 1 | 0 | 0 | 2 |
| ENSMUSG00000021668 | Polk | ENSMUST00000022172 | 29.7771776 | 34 | 18.6360567 | 17.9608373 | 1 | 0 | 1 | 0 | 0 | 0 | 0 | 0 | 0 | 1 | 0 | 0 | 0 | 0 | 2 |
| ENSMUSG00000045102 | Poln | ENSMUST00000042954 | 27.2727273 | 28 | 26.4069264 | 18.6147186 | 0 | 0 | 0 | 0 | 0 | 0 | 0 | 0 | 0 | 0 | 1 | 0 | 0 | 0 | 1 |
| ENSMUSG00000029250 | Polr2b | ENSMUST00000031167 | 29.0322581 | 43 | 11.6129032 | 16.1290323 | 0 | 0 | 0 | 0 | 0 | 0 | 0 | 0 | 0 | 0 | 0 | 0 | 2 | 0 | 2 |
| ENSMUSG00000027427 | Polr3f | ENSMUST00000028914 | 28.8868724 | 31 | 19.5402299 | 20.508167 | 2 | 2 | 0 | 0 | 0 | 0 | 0 | 0 | 0 | 0 | 1 | 0 | 1 | 0 | 6 |
| ENSMUSG00000038628 | Polr3k | ENSMUST00000039551 | 30.1720688 | 31 | 16.9267707 | 21.4085634 | 1 | 0 | 0 | 1 | 0 | 0 | 0 | 0 | 1 | 2 | 0 | 0 | 0 | 0 | 3 |
| ENSMUSG00000053293 | Pom121 | ENSMUST00000111171 | 21.4876033 | 29 | 24.9483471 | 25.0516529 | 1 | 0 | 2 | 1 | 0 | 0 | 0 | 0 | 1 | 4 | 0 | 0 | 0 | 0 | 5 |
| ENSMUSG00000029649 | Pomp | ENSMUST00000201120 | 20.4690214 | 31 | 25.0434279 | 23.8563984 | 0 | 0 | 0 | 0 | 0 | 0 | 0 | 0 | 0 | 0 | 0 | 4 | 0 | 0 | 4 |
| ENSMUSG00000029676 | Pot1a | ENSMUST00000115327 | 27.688172 | 37 | 19.6236559 | 15.6810036 | 1 | 0 | 0 | 0 | 0 | 0 | 0 | 0 | 0 | 0 | 0 | 0 | 0 | 0 | 1 |
| ENSMUSG00000024174 | Pot1b | ENSMUST00000086876 | 29.6162201 | 24 | 19.8406951 | 26.4301231 | 0 | 0 | 1 | 0 | 0 | 0 | 0 | 0 | 0 | 1 | 0 | 0 | 0 | 0 | 1 |
| ENSMUSG00000026565 | Pou2f1 | ENSMUST00000111427 | 26.8707483 | 32 | 19.68065 | 21.6742252 | 0 | 1 | 0 | 1 | 0 | 0 | 0 | 1 | 0 | 2 | 6 | 1 | 12 | 3 | 25 |
| ENSMUSG00000095139 | Pou3f2 | ENSMUST00000178174 | 30.9915296 | 33 | 17.1150972 | 19.0832088 | 1 | 0 | 0 | 0 | 0 | 0 | 0 | 0 | 0 | 0 | 5 | 2 | 17 | 5 | 30 |
| ENSMUSG00000056854 | Pou3f4 | ENSMUST00000078229 | 25.4677127 | 27 | 23.5968618 | 23.958962 | 0 | 0 | 0 | 0 | 0 | 0 | 0 | 0 | 0 | 0 | 0 | 0 | 0 | 2 | 2 |
| ENSMUSG00000031688 | Pou4f2 | ENSMUST00000034115 | 27.2148234 | 35 | 19.5136074 | 18.4134337 | 0 | 0 | 0 | 0 | 0 | 0 | 0 | 0 | 0 | 0 | 1 | 0 | 0 | 0 | 1 |
| ENSMUSG00000024497 | Pou4f3 | ENSMUST00000025374 | 24.1323648 | 29 | 23.0024213 | 24.2130751 | 0 | 0 | 1 | 0 | 0 | 0 | 0 | 0 | 0 | 1 | 0 | 2 | 0 | 0 | 3 |
| ENSMUSG00000009734 | Pou6f2 | ENSMUST00000139064 | 32.7751785 | 28 | 20.4629402 | 18.5914799 | 1 | 0 | 0 | 1 | 0 | 0 | 0 | 0 | 1 | 2 | 0 | 2 | 4 | 0 | 9 |
| ENSMUSG00000029246 | Ppat | ENSMUST00000155272 | 28.8198404 | 31 | 18.3295325 | 22.2063854 | 1 | 0 | 0 | 1 | 0 | 0 | 0 | 0 | 1 | 2 | 0 | 0 | 3 | 0 | 6 |
| ENSMUSG00000053825 | Ppfia2 | ENSMUST00000029404 | 30.9083263 | 32 | 19.1337258 | 17.9983179 | 0 | 0 | 0 | 0 | 0 | 0 | 0 | 0 | 0 | 0 | 0 | 2 | 9 | 1 | 12 |
| ENSMUSG00000036167 | Pphln1 | ENSMUST00000068457 | 27.5774135 | 33 | 20.4735883 | 19.2714026 | 0 | 0 | 0 | 0 | 0 | 0 | 0 | 0 | 0 | 0 | 0 | 0 | 8 | 1 | 9 |
| ENSMUSG00000042133 | Ppig | ENSMUST00000040915 | 29.0633609 | 38 | 16.0330579 | 16.6942149 | 0 | 0 | 0 | 0 | 0 | 0 | 0 | 0 | 0 | 0 | 1 | 2 | 5 | 4 | 12 |
| ENSMUSG00000015757 | Ppil4 | ENSMUST00000143214 | 27.1317829 | 32 | 20.7848837 | 20.4457364 | 0 | 0 | 0 | 0 | 0 | 0 | 0 | 0 | 0 | 0 | 0 | 1 | 2 | 0 | 3 |
| ENSMUSG00000040648 | Ppip5k2 | ENSMUST00000042509 | 31.7365269 | 35 | 15.1497006 | 17.8443114 | 0 | 0 | 0 | 2 | 0 | 0 | 0 | 0 | 2 | 4 | 0 | 0 | 4 | 0 | 8 |
| ENSMUSG00000061130 | Ppm1b | ENSMUST00000112304 | 27.5431291 | 36 | 16.5972635 | 19.6311719 | 1 | 0 | 0 | 0 | 0 | 0 | 0 | 0 | 0 | 0 | 0 | 0 | 0 | 0 | 1 |
| ENSMUSG00000020525 | Ppm1d | ENSMUST00000020835 | 30.0114548 | 36 | 16.838488 | 17.5257732 | 0 | 0 | 0 | 0 | 0 | 0 | 0 | 0 | 0 | 0 | 0 | 1 | 0 | 0 | 1 |
| ENSMUSG00000046442 | Ppm1e | ENSMUST00000055438 | 30.3472572 | 31 | 19.4765979 | 19.6527428 | 1 | 0 | 0 | 1 | 0 | 0 | 0 | 0 | 1 | 2 | 0 | 1 | 0 | 0 | 4 |
| ENSMUSG00000037826 | Ppm1k | ENSMUST00000042766 | 28.9611006 | 34 | 16.5796964 | 20.8965844 | 0 | 0 | 0 | 1 | 0 | 0 | 0 | 0 | 1 | 2 | 8 | 1 | 18 | 3 | 32 |
| ENSMUSG00000014956 | Ppp1cb | ENSMUST00000015100 | 29.5856663 | 34 | 17.2004479 | 19.6864502 | 0 | 0 | 0 | 0 | 0 | 0 | 0 | 0 | 0 | 0 | 5 | 2 | 15 | 2 | 24 |
| ENSMUSG00000019907 | Ppp1r12a | ENSMUST00000070663 | 31.7010309 | 24 | 21.3917526 | 23.1958763 | 0 | 0 | 0 | 0 | 0 | 0 | 0 | 0 | 0 | 0 | 0 | 0 | 1 | 0 | 1 |
| ENSMUSG00000040653 | Ppp1r14c | ENSMUST00000043374 | 27.7777778 | 33 | 19.2460317 | 20.0396825 | 1 | 1 | 0 | 0 | 0 | 0 | 0 | 0 | 0 | 0 | 0 | 7 | 1 | 2 | 12 |
| ENSMUSG00000046062 | Ppp1r15b | ENSMUST00000052529 | 25.3453039 | 33 | 19.198895 | 22.8936464 | 0 | 0 | 0 | 0 | 0 | 0 | 0 | 0 | 0 | 0 | 1 | 0 | 4 | 2 | 7 |
| ENSMUSG00000047714 | Ppp1r2 | ENSMUST00000060188 | 27.968254 | 34 | 17.8730159 | 20.1904762 | 0 | 0 | 0 | 0 | 0 | 0 | 0 | 0 | 0 | 0 | 1 | 3 | 0 | 0 | 4 |
| ENSMUSG00000046794 | Ppp1r3b | ENSMUST00000070481 | 24.1030359 | 29 | 21.8337933 | 24.5936829 | 0 | 0 | 1 | 0 | 0 | 0 | 0 | 0 | 0 | 1 | 0 | 2 | 1 | 0 | 4 |
| ENSMUSG00000021022 | Ppp2r3c | ENSMUST00000021410 | 25.4047323 | 35 | 18.6799502 | 21.1706102 | 0 | 0 | 0 | 0 | 0 | 0 | 0 | 0 | 0 | 0 | 0 | 0 | 5 | 1 | 6 |
| ENSMUSG00000021051 | Ppp2r5e | ENSMUST00000021447 | 28.3645355 | 30 | 19.7103497 | 21.9357118 | 0 | 0 | 0 | 0 | 0 | 0 | 0 | 0 | 0 | 0 | 0 | 0 | 0 | 1 | 1 |
| ENSMUSG00000021816 | Ppp3cb | ENSMUST00000159027 | 30.8962264 | 34 | 16.8396226 | 18.2075472 | 0 | 0 | 0 | 2 | 0 | 0 | 0 | 0 | 2 | 4 | 0 | 1 | 1 | 0 | 6 |
| ENSMUSG00000033953 | Ppp3r1 | ENSMUST00000102880 | 28.1778637 | 35 | 17.4963751 | 19.1880135 | 2 | 0 | 1 | 3 | 0 | 0 | 0 | 0 | 3 | 7 | 0 | 0 | 0 | 0 | 9 |
| ENSMUSG00000052144 | Ppp4r2 | ENSMUST00000063854 | 29.5413937 | 34 | 16.1703395 | 20.2799285 | 0 | 1 | 0 | 0 | 0 | 0 | 0 | 0 | 0 | 0 | 0 | 0 | 3 | 1 | 5 |
| ENSMUSG00000003099 | Ppp5c | ENSMUST00000003183 | 22.3880597 | 21 | 26.2260128 | 30.2771855 | 0 | 0 | 1 | 0 | 0 | 0 | 0 | 0 | 0 | 1 | 0 | 0 | 0 | 0 | 1 |
| ENSMUSG00000026753 | Ppp6c | ENSMUST00000028087 | 27.184466 | 37 | 17.9444258 | 17.4422497 | 0 | 0 | 0 | 0 | 0 | 0 | 0 | 0 | 0 | 0 | 0 | 0 | 1 | 3 | 4 |
| ENSMUSG00000038582 | Pptc7 | ENSMUST00000053426 | 26.5336073 | 30 | 20.8687995 | 22.835339 | 0 | 0 | 0 | 0 | 0 | 0 | 0 | 0 | 0 | 0 | 0 | 0 | 1 | 0 | 1 |
| ENSMUSG00000021713 | Ppwd1 | ENSMUST00000022226 | 32.1148825 | 41 | 14.0992167 | 13.0548303 | 0 | 0 | 0 | 0 | 0 | 0 | 0 | 0 | 0 | 0 | 0 | 0 | 12 | 3 | 15 |
| ENSMUSG00000045679 | Pqlc3 | ENSMUST00000156360 | 23.5182551 | 27 | 26.742532 | 22.2854433 | 0 | 0 | 0 | 0 | 0 | 0 | 0 | 0 | 0 | 0 | 0 | 4 | 0 | 0 | 4 |
| ENSMUSG00000061119 | Prcp | ENSMUST00000076052 | 23.3859397 | 34 | 22.8837877 | 19.7274032 | 0 | 0 | 0 | 0 | 0 | 0 | 0 | 0 | 0 | 0 | 0 | 1 | 10 | 0 | 11 |
| ENSMUSG00000038151 | Prdm1 | ENSMUST00000039174 | 28.1927711 | 33 | 18.5943775 | 19.9196787 | 0 | 0 | 0 | 0 | 0 | 0 | 0 | 0 | 0 | 0 | 3 | 3 | 3 | 0 | 9 |
| ENSMUSG00000042496 | Prdm10 | ENSMUST00000117389 | 26.6666667 | 35 | 17.4814815 | 21.3333333 | 0 | 1 | 1 | 1 | 0 | 0 | 0 | 0 | 1 | 3 | 0 | 1 | 0 | 0 | 5 |
| ENSMUSG00000035456 | Prdm8 | ENSMUST00000112959 | 29.3742018 | 29 | 22.0945083 | 19.6679438 | 0 | 0 | 0 | 0 | 0 | 0 | 0 | 0 | 0 | 0 | 0 | 10 | 2 | 0 | 12 |
| ENSMUSG00000024997 | Prdx3 | ENSMUST00000025961 | 26.9918699 | 34 | 20.1626016 | 18.8617886 | 0 | 0 | 0 | 0 | 0 | 0 | 0 | 0 | 0 | 0 | 0 | 0 | 2 | 2 | 4 |
| ENSMUSG00000050697 | Prkaa1 | ENSMUST00000051186 | 28.9518032 | 35 | 17.7283451 | 18.1327941 | 0 | 0 | 0 | 0 | 0 | 0 | 0 | 0 | 0 | 0 | 1 | 0 | 0 | 0 | 1 |
| ENSMUSG00000028518 | Prkaa2 | ENSMUST00000030243 | 29.7684143 | 32 | 18.5911869 | 19.1862335 | 1 | 0 | 0 | 0 | 0 | 0 | 0 | 0 | 0 | 0 | 0 | 1 | 1 | 1 | 4 |
| ENSMUSG00000005034 | Prkacb | ENSMUST00000102515 | 26.5096218 | 32 | 20.9024552 | 20.5706702 | 0 | 0 | 0 | 0 | 0 | 0 | 0 | 0 | 0 | 0 | 0 | 2 | 0 | 0 | 2 |
| ENSMUSG00000028944 | Prkag2 | ENSMUST00000030784 | 26.2589928 | 30 | 20.8633094 | 22.9316547 | 1 | 0 | 1 | 0 | 0 | 0 | 0 | 0 | 0 | 1 | 0 | 0 | 2 | 0 | 4 |
| ENSMUSG00000020612 | Prkar1a | ENSMUST00000049527 | 24.6835443 | 34 | 19.2307692 | 21.7137293 | 0 | 0 | 0 | 2 | 0 | 0 | 0 | 0 | 2 | 4 | 2 | 3 | 0 | 0 | 9 |
| ENSMUSG00000002997 | Prkar2b | ENSMUST00000036497 | 30.0575013 | 34 | 17.2503921 | 18.4004182 | 0 | 0 | 0 | 1 | 0 | 0 | 0 | 0 | 1 | 2 | 0 | 0 | 0 | 0 | 2 |
| ENSMUSG00000037643 | Prkci | ENSMUST00000108249 | 27.7404922 | 33 | 18.9038031 | 20.8426547 | 1 | 1 | 1 | 0 | 0 | 0 | 0 | 0 | 0 | 1 | 1 | 1 | 0 | 0 | 5 |
| ENSMUSG00000002731 | Prkra | ENSMUST00000002808 | 24.7563353 | 35 | 17.9337232 | 22.2222222 | 0 | 0 | 0 | 0 | 0 | 0 | 0 | 0 | 0 | 0 | 0 | 1 | 0 | 0 | 1 |
| ENSMUSG00000035725 | Prkx | ENSMUST00000036333 | 29.7923323 | 31 | 20.686901 | 18.5702875 | 0 | 0 | 0 | 1 | 0 | 0 | 0 | 0 | 1 | 2 | 5 | 0 | 5 | 0 | 12 |
| ENSMUSG00000005268 | Prlr | ENSMUST00000124470 | 31.2322397 | 32 | 19.5298373 | 17.4761044 | 0 | 1 | 0 | 0 | 0 | 0 | 0 | 0 | 0 | 0 | 0 | 3 | 1 | 1 | 6 |
| ENSMUSG00000049300 | Prmt6 | ENSMUST00000168412 | 28.9709763 | 28 | 21.7414248 | 21.1081794 | 0 | 1 | 0 | 0 | 0 | 0 | 0 | 0 | 0 | 0 | 0 | 1 | 0 | 0 | 2 |
| ENSMUSG00000098754 | Prn | ENSMUST00000136783 | 25.9029126 | 29 | 23.7669903 | 21.5533981 | 0 | 0 | 0 | 0 | 0 | 0 | 0 | 0 | 0 | 0 | 2 | 0 | 5 | 0 | 7 |
| ENSMUSG00000030069 | Prok2 | ENSMUST00000101120 | 25.142315 | 35 | 21.9165085 | 18.3111954 | 1 | 0 | 0 | 0 | 0 | 0 | 0 | 0 | 0 | 0 | 0 | 0 | 9 | 0 | 10 |
| ENSMUSG00000050558 | Prokr2 | ENSMUST00000110157 | 26.4327109 | 27 | 23.2775274 | 23.4063104 | 0 | 0 | 0 | 0 | 0 | 0 | 0 | 0 | 0 | 0 | 0 | 9 | 0 | 0 | 9 |
| ENSMUSG00000022912 | Pros1 | ENSMUST00000023629 | 28.7944493 | 35 | 15.6981787 | 20.2081526 | 0 | 0 | 0 | 0 | 0 | 0 | 0 | 0 | 0 | 0 | 1 | 0 | 7 | 1 | 9 |
| ENSMUSG00000010175 | Prox1 | ENSMUST00000175916 | 28.7367406 | 31 | 20.2314368 | 20.1542912 | 0 | 0 | 0 | 0 | 0 | 0 | 0 | 0 | 0 | 0 | 0 | 5 | 0 | 0 | 5 |
| ENSMUSG00000039449 | Prpf18 | ENSMUST00000035721 | 25.6281407 | 31 | 19.8911223 | 23.1155779 | 0 | 0 | 0 | 0 | 0 | 0 | 0 | 0 | 0 | 0 | 5 | 2 | 2 | 1 | 10 |
| ENSMUSG00000027881 | Prpf38b | ENSMUST00000129273 | 32.178415 | 25 | 16.8458781 | 26.4436479 | 0 | 0 | 0 | 0 | 0 | 0 | 0 | 0 | 0 | 0 | 2 | 2 | 1 | 1 | 6 |
| ENSMUSG00000035597 | Prpf39 | ENSMUST00000129956 | 32.9578976 | 31 | 16.8946098 | 19.4690265 | 0 | 0 | 0 | 0 | 0 | 0 | 0 | 0 | 0 | 0 | 0 | 0 | 8 | 0 | 8 |
| ENSMUSG00000061136 | Prpf40a | ENSMUST00000076313 | 30.1296379 | 36 | 16.5847117 | 17.5011176 | 0 | 0 | 0 | 0 | 0 | 0 | 0 | 0 | 0 | 0 | 0 | 1 | 2 | 0 | 3 |
| ENSMUSG00000021413 | Prpf4b | ENSMUST00000077853 | 28.2591725 | 37 | 15.5347385 | 19.5940671 | 0 | 0 | 0 | 0 | 0 | 0 | 0 | 0 | 0 | 0 | 0 | 0 | 0 | 1 | 1 |
| ENSMUSG00000025742 | Prps2 | ENSMUST00000026839 | 29.2569659 | 33 | 18.5371517 | 19.620743 | 0 | 0 | 0 | 0 | 0 | 0 | 0 | 0 | 0 | 0 | 0 | 1 | 1 | 0 | 2 |
| ENSMUSG00000020528 | Prpsap2 | ENSMUST00000168115 | 28.9473684 | 33 | 19.4331984 | 18.6234818 | 0 | 0 | 0 | 0 | 0 | 0 | 0 | 0 | 0 | 0 | 1 | 1 | 1 | 1 | 4 |
| ENSMUSG00000020493 | Prr11 | ENSMUST00000051395 | 29.6898638 | 31 | 21.2556732 | 18.419062 | 0 | 0 | 0 | 0 | 0 | 0 | 0 | 0 | 0 | 0 | 0 | 2 | 0 | 0 | 2 |
| ENSMUSG00000073565 | Prr16 | ENSMUST00000116639 | 31.8965517 | 34 | 16.69279 | 17.3981191 | 0 | 0 | 0 | 0 | 0 | 0 | 0 | 0 | 0 | 0 | 1 | 0 | 1 | 0 | 2 |
| ENSMUSG00000055945 | Prr18 | ENSMUST00000069742 | 23.3103448 | 27 | 25.6551724 | 24.045977 | 0 | 0 | 0 | 0 | 0 | 0 | 0 | 0 | 0 | 0 | 0 | 13 | 0 | 0 | 13 |
| ENSMUSG00000024594 | Prrc1 | ENSMUST00000025490 | 22.4012158 | 33 | 21.5197568 | 22.9483283 | 1 | 0 | 0 | 1 | 0 | 0 | 0 | 0 | 1 | 2 | 0 | 1 | 2 | 1 | 7 |
| ENSMUSG00000047996 | Prrg1 | ENSMUST00000114025 | 32.2055953 | 38 | 14.5413142 | 15.0292778 | 0 | 0 | 0 | 1 | 0 | 0 | 0 | 0 | 1 | 2 | 1 | 0 | 4 | 0 | 7 |
| ENSMUSG00000033361 | Prrg3 | ENSMUST00000048790 | 24.2896069 | 30 | 22.5184897 | 23.4527053 | 0 | 0 | 0 | 0 | 0 | 0 | 0 | 0 | 0 | 0 | 0 | 5 | 1 | 0 | 6 |
| ENSMUSG00000027171 | Prrg4 | ENSMUST00000028593 | 29.4341676 | 29 | 19.640914 | 21.4907508 | 0 | 0 | 0 | 0 | 0 | 0 | 0 | 0 | 0 | 0 | 2 | 0 | 8 | 0 | 10 |
| ENSMUSG00000026586 | Prrx1 | ENSMUST00000174397 | 29.6258434 | 30 | 19.7096708 | 20.9159681 | 0 | 0 | 0 | 0 | 0 | 0 | 0 | 0 | 0 | 0 | 1 | 25 | 1 | 1 | 28 |
| ENSMUSG00000036030 | Prtg | ENSMUST00000055535 | 28.4225296 | 31 | 19.1621567 | 20.9661255 | 0 | 1 | 0 | 0 | 0 | 0 | 0 | 0 | 0 | 0 | 2 | 1 | 12 | 5 | 21 |
| ENSMUSG00000030465 | Psd3 | ENSMUST00000038959 | 28.5551331 | 32 | 18.9480355 | 20.7858048 | 0 | 0 | 0 | 1 | 0 | 0 | 0 | 0 | 1 | 2 | 1 | 0 | 0 | 0 | 3 |
| ENSMUSG00000066760 | Psg16 | ENSMUST00000071399 | 30.3128371 | 26 | 16.4688961 | 26.7889248 | 1 | 0 | 0 | 1 | 0 | 0 | 0 | 0 | 1 | 2 | 3 | 0 | 4 | 0 | 10 |
| ENSMUSG00000015671 | Psma2 | ENSMUST00000170836 | 30.5825243 | 42 | 13.1067961 | 14.5631068 | 0 | 0 | 0 | 0 | 0 | 0 | 0 | 0 | 0 | 0 | 0 | 0 | 1 | 0 | 1 |
| ENSMUSG00000060073 | Psma3 | ENSMUST00000160027 | 29.2520492 | 32 | 19.3647541 | 19.2110656 | 1 | 0 | 0 | 0 | 0 | 0 | 0 | 0 | 0 | 0 | 0 | 0 | 8 | 0 | 9 |
| ENSMUSG00000019303 | Psmc3ip | ENSMUST00000019447 | 30.5025997 | 24 | 20.797227 | 24.4367418 | 1 | 0 | 0 | 0 | 0 | 0 | 0 | 0 | 0 | 0 | 0 | 0 | 0 | 0 | 1 |
| ENSMUSG00000031429 | Psmd10 | ENSMUST00000112978 | 29.8969072 | 32 | 17.067583 | 21.4203895 | 0 | 0 | 1 | 0 | 0 | 0 | 0 | 0 | 0 | 1 | 0 | 0 | 0 | 0 | 1 |
| ENSMUSG00000078652 | Psme3 | ENSMUST00000019470 | 20.6606607 | 31 | 25.8858859 | 22.5225225 | 0 | 0 | 0 | 0 | 0 | 0 | 0 | 0 | 0 | 0 | 1 | 3 | 1 | 0 | 5 |
| ENSMUSG00000021938 | Pspc1 | ENSMUST00000163924 | 28.9655172 | 37 | 16.1685824 | 18.0842912 | 0 | 0 | 0 | 0 | 0 | 0 | 0 | 0 | 0 | 0 | 2 | 0 | 3 | 0 | 5 |
| ENSMUSG00000074925 | Ptar1 | ENSMUST00000099560 | 27.5720165 | 37 | 18.5185185 | 16.872428 | 1 | 0 | 1 | 1 | 0 | 0 | 0 | 0 | 1 | 3 | 0 | 0 | 2 | 0 | 6 |
| ENSMUSG00000006498 | Ptbp1 | ENSMUST00000165704 | 23.5609103 | 26 | 25.6358768 | 24.3641232 | 0 | 0 | 0 | 0 | 0 | 0 | 0 | 0 | 0 | 0 | 0 | 0 | 0 | 1 | 1 |

| | | | | | | | | | | | | | | | | | | | | | |
|---|---|---|---|---|---|---|---|---|---|---|---|---|---|---|---|---|---|---|---|---|---|
| ENSMUSG00000028134 | Ptbp2 | ENSMUST00000029780 | 27.7427491 | 38 | 14.6910467 | 19.1677175 | 0 | 0 | 0 | 0 | 0 | 0 | 0 | 0 | 0 | 0 | 0 | 2 | 4 | 0 | 6 |
| ENSMUSG00000028382 | Ptbp3 | ENSMUST00000030076 | 29.0727817 | 36 | 16.3908275 | 18.4047856 | 0 | 0 | 0 | 0 | 0 | 0 | 0 | 0 | 0 | 0 | 0 | 1 | 5 | 0 | 6 |
| ENSMUSG00000021466 | Ptch1 | ENSMUST00000194663 | 23.8593327 | 28 | 24.1118124 | 24.1298467 | 1 | 1 | 0 | 2 | 0 | 0 | 0 | 1 | 1 | 4 | 0 | 3 | 1 | 0 | 10 |
| ENSMUSG00000042256 | Ptchd4 | ENSMUST00000048691 | 31.5734449 | 32 | 17.903816 | 18.9231573 | 0 | 0 | 0 | 1 | 0 | 0 | 0 | 0 | 1 | 2 | 5 | 0 | 4 | 0 | 11 |
| ENSMUSG00000013663 | Pten | ENSMUST00000013807 | 28.7834312 | 35 | 16.8297456 | 19.4390085 | 0 | 0 | 0 | 1 | 0 | 0 | 0 | 0 | 1 | 2 | 1 | 0 | 13 | 2 | 18 |
| ENSMUSG00000039942 | Ptger4 | ENSMUST00000047379 | 31.7923763 | 31 | 17.5993512 | 19.5458232 | 1 | 1 | 1 | 0 | 0 | 0 | 0 | 0 | 0 | 1 | 0 | 0 | 0 | 0 | 3 |
| ENSMUSG00000072946 | Ptgr2 | ENSMUST00000123614 | 28.1473337 | 33 | 18.0318857 | 21.2204508 | 0 | 0 | 0 | 1 | 0 | 0 | 0 | 0 | 1 | 2 | 1 | 0 | 0 | 1 | 4 |
| ENSMUSG00000026238 | Ptma | ENSMUST00000045897 | 31.5714286 | 31 | 20.4285714 | 16.8571429 | 0 | 0 | 0 | 0 | 0 | 0 | 0 | 0 | 0 | 0 | 0 | 0 | 2 | 2 | 4 |
| ENSMUSG00000024539 | Ptpn2 | ENSMUST00000122412 | 33.9456282 | 31 | 16.8993387 | 18.2218957 | 0 | 0 | 0 | 1 | 0 | 0 | 0 | 0 | 1 | 2 | 0 | 0 | 0 | 0 | 2 |
| ENSMUSG00000026384 | Ptpn4 | ENSMUST00000064091 | 30.9342749 | 35 | 16.5806627 | 17.7756654 | 0 | 0 | 0 | 1 | 0 | 0 | 0 | 0 | 1 | 2 | 0 | 0 | 0 | 0 | 2 |
| ENSMUSG00000027303 | Ptpra | ENSMUST00000028769 | 27.7777778 | 30 | 23.297491 | 19.3548387 | 0 | 0 | 0 | 0 | 0 | 0 | 0 | 0 | 0 | 0 | 0 | 2 | 0 | 0 | 2 |
| ENSMUSG00000020154 | Ptprb | ENSMUST00000092167 | 29.6839131 | 27 | 20.9606216 | 21.9671552 | 0 | 0 | 0 | 1 | 0 | 0 | 0 | 0 | 1 | 2 | 1 | 3 | 1 | 2 | 9 |
| ENSMUSG00000028399 | Ptprd | ENSMUST00000107289 | 30.9946237 | 33 | 18.0645161 | 17.6075269 | 0 | 0 | 0 | 0 | 0 | 0 | 0 | 0 | 0 | 0 | 2 | 1 | 7 | 6 | 16 |
| ENSMUSG00000021745 | Ptprg | ENSMUST00000022264 | 30.7341651 | 33 | 17.6823417 | 18.6900192 | 1 | 0 | 0 | 0 | 0 | 0 | 0 | 0 | 0 | 0 | 0 | 1 | 0 | 2 | 4 |
| ENSMUSG00000068748 | Ptprz1 | ENSMUST00000202579 | 35.2880125 | 34 | 14.5303581 | 16.6061235 | 0 | 0 | 0 | 0 | 0 | 0 | 0 | 0 | 0 | 0 | 1 | 0 | 0 | 0 | 1 |
| ENSMUSG00000032067 | Pts | ENSMUST00000034570 | 29.2436975 | 31 | 18.8235294 | 20.6722689 | 0 | 0 | 0 | 1 | 0 | 0 | 0 | 0 | 1 | 2 | 0 | 0 | 0 | 0 | 2 |
| ENSMUSG00000027832 | Ptx3 | ENSMUST00000029421 | 31 | 35 | 13.1666667 | 20.8333333 | 0 | 0 | 0 | 0 | 0 | 0 | 0 | 0 | 0 | 0 | 0 | 0 | 2 | 0 | 2 |
| ENSMUSG00000020594 | Pum2 | ENSMUST00000165293 | 29.0859667 | 30 | 20.1325178 | 20.795107 | 0 | 0 | 0 | 0 | 0 | 0 | 0 | 0 | 0 | 0 | 1 | 4 | 0 | 1 | 6 |
| ENSMUSG00000043991 | Pura | ENSMUST00000051301 | 30.4390025 | 35 | 15.9325071 | 19.0383754 | 1 | 0 | 0 | 1 | 0 | 0 | 0 | 0 | 1 | 2 | 2 | 3 | 6 | 0 | 14 |
| ENSMUSG00000094483 | Purb | ENSMUST00000179343 | 30.0697388 | 32 | 17.762888 | 20.1695611 | 0 | 0 | 0 | 0 | 0 | 0 | 0 | 0 | 0 | 0 | 2 | 1 | 1 | 5 | 9 |
| ENSMUSG00000049184 | Purg | ENSMUST00000078058 | 28.2527881 | 32 | 19.0830235 | 21.0656753 | 0 | 0 | 0 | 0 | 0 | 0 | 0 | 0 | 0 | 0 | 0 | 1 | 0 | 0 | 1 |
| ENSMUSG00000022656 | Pvrl3 | ENSMUST00000023335 | 26.9147497 | 29 | 22.9364005 | 21.4343708 | 0 | 0 | 0 | 0 | 0 | 0 | 0 | 0 | 0 | 0 | 2 | 1 | 1 | 0 | 4 |
| ENSMUSG00000044950 | Pwwp2a | ENSMUST00000061070 | 31.1425683 | 33 | 15.065723 | 20.5257836 | 0 | 0 | 0 | 0 | 0 | 0 | 0 | 0 | 0 | 0 | 1 | 0 | 0 | 0 | 1 |
| ENSMUSG00000021411 | Pxdc1 | ENSMUST00000125037 | 24.2088608 | 27 | 24.6835443 | 23.6550633 | 0 | 0 | 0 | 0 | 0 | 0 | 0 | 0 | 0 | 0 | 0 | 1 | 0 | 0 | 1 |
| ENSMUSG00000020674 | Pxdn | ENSMUST00000122328 | 25.9578544 | 28 | 22.2222222 | 23.5632184 | 1 | 1 | 1 | 1 | 0 | 0 | 0 | 0 | 1 | 3 | 0 | 0 | 0 | 0 | 5 |
| ENSMUSG00000033885 | Pxk | ENSMUST00000112689 | 25.4621849 | 28 | 25.9663866 | 20.9243697 | 0 | 0 | 0 | 0 | 0 | 0 | 0 | 0 | 0 | 0 | 0 | 0 | 0 | 1 | 1 |
| ENSMUSG00000000876 | Pxmp4 | ENSMUST00000000896 | 25.6397875 | 26 | 24.8672139 | 23.2737808 | 0 | 0 | 0 | 0 | 0 | 0 | 0 | 0 | 0 | 0 | 0 | 0 | 7 | 0 | 7 |
| ENSMUSG00000062078 | Qk | ENSMUST00000042296 | 30.3884235 | 36 | 16.7364813 | 16.9459254 | 0 | 0 | 0 | 1 | 0 | 0 | 0 | 0 | 1 | 2 | 0 | 25 | 16 | 1 | 44 |
| ENSMUSG00000024084 | Qpct | ENSMUST00000040789 | 34.3157895 | 38 | 12.2105263 | 15.3684211 | 1 | 1 | 0 | 0 | 0 | 0 | 0 | 0 | 0 | 0 | 0 | 0 | 0 | 0 | 2 |
| ENSMUSG00000074994 | Qser1 | ENSMUST00000117237 | 28.3923706 | 33 | 19.8365123 | 19.2370572 | 0 | 1 | 0 | 2 | 0 | 0 | 0 | 1 | 1 | 4 | 2 | 0 | 1 | 0 | 8 |
| ENSMUSG00000036327 | Qsox2 | ENSMUST00000091263 | 24.5274725 | 29 | 22.6373626 | 23.8241758 | 1 | 0 | 0 | 0 | 0 | 0 | 0 | 0 | 0 | 0 | 0 | 0 | 1 | 0 | 2 |
| ENSMUSG00000022704 | Qtrtd1 | ENSMUST00000023387 | 24.4137102 | 35 | 19.1822008 | 21.4070956 | 0 | 0 | 0 | 0 | 0 | 0 | 0 | 0 | 0 | 0 | 2 | 0 | 3 | 0 | 5 |
| ENSMUSG00000056211 | R3hdm1 | ENSMUST00000187900 | 30.3172127 | 33 | 17.9927197 | 18.7727509 | 1 | 1 | 1 | 0 | 0 | 0 | 0 | 0 | 0 | 1 | 3 | 1 | 3 | 1 | 11 |
| ENSMUSG00000020671 | Rab10 | ENSMUST00000021001 | 27.8319898 | 34 | 20.3224438 | 17.5222741 | 0 | 0 | 0 | 0 | 0 | 0 | 0 | 0 | 0 | 0 | 0 | 3 | 0 | 0 | 3 |
| ENSMUSG00000004771 | Rab11a | ENSMUST00000172298 | 27.2903226 | 38 | 16.1935484 | 18.5806452 | 0 | 0 | 0 | 0 | 0 | 0 | 0 | 0 | 0 | 0 | 0 | 0 | 2 | 0 | 2 |
| ENSMUSG00000077450 | Rab11b | ENSMUST00000057373 | 26.1188272 | 27 | 23.7654321 | 22.8587963 | 0 | 0 | 0 | 1 | 0 | 0 | 0 | 0 | 1 | 2 | 0 | 4 | 6 | 0 | 12 |
| ENSMUSG00000023460 | Rab12 | ENSMUST00000070538 | 27.9286927 | 30 | 19.9490662 | 21.6468591 | 0 | 0 | 1 | 0 | 0 | 0 | 0 | 0 | 0 | 1 | 0 | 0 | 0 | 0 | 1 |
| ENSMUSG00000026878 | Rab14 | ENSMUST00000113025 | 27.6547416 | 32 | 18.9381034 | 21.0675752 | 0 | 0 | 0 | 0 | 0 | 0 | 0 | 0 | 0 | 0 | 1 | 0 | 2 | 0 | 3 |
| ENSMUSG00000073639 | Rab18 | ENSMUST00000097680 | 26.4593909 | 38 | 16.3071066 | 19.035533 | 0 | 0 | 0 | 0 | 0 | 0 | 0 | 0 | 0 | 0 | 0 | 0 | 1 | 0 | 1 |
| ENSMUSG00000020149 | Rab1a | ENSMUST00000020358 | 27.2998508 | 33 | 19.6419692 | 19.6419692 | 1 | 0 | 0 | 0 | 0 | 0 | 0 | 0 | 0 | 0 | 0 | 3 | 0 | 0 | 4 |
| ENSMUSG00000020132 | Rab21 | ENSMUST00000020343 | 26.2926293 | 38 | 19.5819582 | 15.7315732 | 0 | 0 | 0 | 0 | 0 | 0 | 0 | 0 | 0 | 0 | 0 | 1 | 2 | 0 | 3 |
| ENSMUSG00000027519 | Rab22a | ENSMUST00000142820 | 25.8009858 | 28 | 23.1515712 | 22.9359211 | 0 | 0 | 0 | 0 | 0 | 0 | 0 | 0 | 0 | 0 | 0 | 6 | 4 | 2 | 12 |
| ENSMUSG00000004768 | Rab23 | ENSMUST00000138024 | 26.3422375 | 29 | 22.1370008 | 22.7982015 | 0 | 0 | 0 | 0 | 0 | 0 | 0 | 0 | 0 | 0 | 0 | 5 | 1 | 1 | 7 |
| ENSMUSG00000024511 | Rab27b | ENSMUST00000121693 | 31.6328215 | 32 | 17.7150404 | 18.7386495 | 0 | 0 | 0 | 1 | 0 | 0 | 0 | 0 | 1 | 2 | 0 | 1 | 2 | 0 | 5 |
| ENSMUSG00000029128 | Rab28 | ENSMUST00000201422 | 25.4484305 | 33 | 20.2914798 | 21.1883408 | 1 | 1 | 0 | 3 | 0 | 0 | 0 | 1 | 2 | 6 | 0 | 1 | 0 | 0 | 9 |
| ENSMUSG00000022159 | Rab2b | ENSMUST00000022765 | 24.7565351 | 31 | 24.551512 | 19.2721681 | 1 | 0 | 1 | 1 | 0 | 0 | 0 | 0 | 1 | 3 | 0 | 0 | 0 | 0 | 4 |
| ENSMUSG00000030643 | Rab30 | ENSMUST00000032879 | 30.3056027 | 29 | 18.8455008 | 21.901528 | 0 | 0 | 0 | 0 | 0 | 0 | 0 | 0 | 0 | 0 | 0 | 0 | 0 | 2 | 2 |
| ENSMUSG00000027739 | Rab33b | ENSMUST00000054387 | 28.2019704 | 33 | 18.4729064 | 20.7307061 | 0 | 0 | 0 | 0 | 0 | 0 | 0 | 0 | 0 | 0 | 0 | 0 | 1 | 0 | 1 |
| ENSMUSG00000055069 | Rab39 | ENSMUST00000068449 | 31.5454545 | 31 | 18.7727273 | 18.5909091 | 0 | 0 | 1 | 0 | 0 | 0 | 0 | 0 | 0 | 1 | 0 | 0 | 0 | 1 | 2 |
| ENSMUSG00000031202 | Rab39b | ENSMUST00000033545 | 34.2800789 | 31 | 17.0414201 | 18.1459566 | 0 | 0 | 1 | 1 | 0 | 0 | 0 | 0 | 1 | 3 | 0 | 0 | 0 | 0 | 3 |
| ENSMUSG00000021700 | Rab3c | ENSMUST00000167824 | 29.1688516 | 30 | 19.2711064 | 21.7226009 | 0 | 0 | 0 | 1 | 0 | 0 | 0 | 0 | 1 | 2 | 1 | 2 | 2 | 0 | 7 |
| ENSMUSG00000000711 | Rab5b | ENSMUST00000000727 | 21.9512195 | 30 | 26.0590501 | 22.0795892 | 1 | 0 | 1 | 0 | 0 | 0 | 0 | 0 | 0 | 1 | 0 | 13 | 0 | 0 | 15 |
| ENSMUSG00000019173 | Rab5c | ENSMUST00000107364 | 21.5799615 | 27 | 27.8420039 | 23.5067437 | 0 | 0 | 0 | 0 | 0 | 0 | 0 | 0 | 0 | 0 | 0 | 1 | 0 | 0 | 1 |
| ENSMUSG00000079477 | Rab7 | ENSMUST00000113600 | 26.3355907 | 29 | 24.5297216 | 20.6170053 | 0 | 0 | 0 | 0 | 0 | 0 | 0 | 0 | 0 | 0 | 0 | 0 | 1 | 0 | 1 |
| ENSMUSG00000036943 | Rab8b | ENSMUST00000041139 | 26.6617247 | 33 | 19.3229553 | 20.657277 | 0 | 0 | 0 | 0 | 0 | 0 | 0 | 0 | 0 | 0 | 3 | 1 | 7 | 4 | 15 |
| ENSMUSG00000043463 | Rab9b | ENSMUST00000058814 | 30.9285468 | 32 | 18.7090093 | 18.4328616 | 2 | 1 | 0 | 2 | 0 | 0 | 0 | 0 | 2 | 4 | 0 | 0 | 3 | 4 | 14 |
| ENSMUSG00000026721 | Rabgap1l | ENSMUST00000028052 | 31.1979373 | 33 | 18.6037287 | 16.8385561 | 0 | 0 | 0 | 0 | 0 | 0 | 0 | 0 | 0 | 0 | 1 | 0 | 3 | 0 | 4 |
| ENSMUSG00000025340 | Rabgef1 | ENSMUST00000119797 | 25.3153153 | 32 | 19.2792793 | 23.5135135 | 0 | 0 | 0 | 0 | 0 | 0 | 0 | 0 | 0 | 0 | 0 | 0 | 5 | 0 | 5 |
| ENSMUSG00000022827 | Rabl3 | ENSMUST00000130028 | 25.2090032 | 31 | 21.5434084 | 22.1221865 | 0 | 0 | 0 | 0 | 0 | 0 | 0 | 0 | 0 | 0 | 1 | 1 | 7 | 0 | 9 |
| ENSMUSG00000001847 | Rac1 | ENSMUST00000080537 | 24.6034122 | 29 | 23.5258905 | 22.5381622 | 2 | 1 | 0 | 5 | 0 | 0 | 1 | 2 | 2 | 10 | 0 | 0 | 1 | 0 | 14 |
| ENSMUSG00000021635 | Rad17 | ENSMUST00000022136 | 30.480167 | 35 | 14.1962422 | 20.6680585 | 1 | 1 | 1 | 0 | 0 | 0 | 0 | 0 | 0 | 1 | 0 | 0 | 0 | 0 | 3 |
| ENSMUSG00000030254 | Rad18 | ENSMUST00000077088 | 28.0628272 | 33 | 20.2094241 | 19.0575916 | 0 | 0 | 1 | 0 | 0 | 0 | 0 | 0 | 0 | 1 | 0 | 0 | 0 | 0 | 1 |
| ENSMUSG00000030346 | Rad51ap1 | ENSMUST00000112220 | 24.0174672 | 33 | 22.0960699 | 20.9606987 | 0 | 0 | 0 | 0 | 0 | 0 | 0 | 0 | 0 | 0 | 0 | 1 | 0 | 0 | 1 |
| ENSMUSG00000007646 | Rad51c | ENSMUST00000067692 | 24.2977016 | 32 | 20.1021525 | 23.8599051 | 0 | 0 | 0 | 0 | 0 | 0 | 0 | 0 | 0 | 0 | 0 | 1 | 6 | 0 | 7 |
| ENSMUSG00000078773 | Rad54b | ENSMUST00000095145 | 29.8315164 | 33 | 16.5014866 | 20.7631318 | 1 | 1 | 0 | 2 | 0 | 0 | 0 | 0 | 2 | 4 | 0 | 1 | 0 | 1 | 8 |
| ENSMUSG00000038569 | Rad9b | ENSMUST00000117263 | 29.13085 | 32 | 21.6809933 | 17.669532 | 0 | 0 | 0 | 0 | 0 | 0 | 0 | 0 | 0 | 0 | 0 | 0 | 12 | 0 | 12 |
| ENSMUSG00000021027 | Ralgapa1 | ENSMUST00000110687 | 27.5988287 | 31 | 21.1566618 | 20.3513909 | 0 | 1 | 1 | 0 | 0 | 0 | 0 | 0 | 0 | 1 | 0 | 0 | 0 | 0 | 2 |
| ENSMUSG00000026594 | Ralgps2 | ENSMUST00000172057 | 27.5310124 | 30 | 20.0480192 | 22.5690276 | 0 | 0 | 0 | 0 | 0 | 0 | 0 | 0 | 0 | 0 | 1 | 1 | 9 | 2 | 13 |
| ENSMUSG00000039717 | Ralyl | ENSMUST00000108372 | 30.8441558 | 34 | 17.288961 | 18.1818182 | 0 | 0 | 0 | 1 | 0 | 0 | 0 | 0 | 1 | 2 | 0 | 0 | 0 | 0 | 2 |
| ENSMUSG00000040594 | Ranbp17 | ENSMUST00000037522 | 23.8274707 | 36 | 21.1474037 | 19.0117253 | 8 | 4 | 0 | 18 | 0 | 2 | 4 | 6 | 8 | 38 | 1 | 0 | 4 | 0 | 55 |
| ENSMUSG00000068798 | Rap1a | ENSMUST00000090678 | 30.8048639 | 33 | 17.0237406 | 18.9924725 | 0 | 0 | 0 | 1 | 0 | 0 | 0 | 0 | 1 | 2 | 0 | 0 | 0 | 0 | 2 |
| ENSMUSG00000036894 | Rap2b | ENSMUST00000049064 | 26.8249468 | 32 | 19.1708009 | 22.2182849 | 0 | 0 | 0 | 0 | 0 | 0 | 0 | 0 | 0 | 0 | 0 | 0 | 14 | 0 | 14 |
| ENSMUSG00000050029 | Rap2c | ENSMUST00000053593 | 30.6996171 | 36 | 16.637661 | 16.8813087 | 0 | 0 | 0 | 0 | 0 | 0 | 0 | 0 | 0 | 0 | 0 | 0 | 1 | 0 | 1 |
| ENSMUSG00000026014 | Raph1 | ENSMUST00000140485 | 30.0932091 | 32 | 18.7250333 | 18.924767 | 0 | 0 | 1 | 0 | 0 | 0 | 0 | 0 | 0 | 1 | 2 | 2 | 1 | 5 | 11 |
| ENSMUSG00000021549 | Rasa1 | ENSMUST00000109552 | 31.4954683 | 36 | 16.5407855 | 15.7854985 | 0 | 1 | 0 | 0 | 0 | 0 | 0 | 0 | 0 | 0 | 0 | 0 | 0 | 2 | 3 |
| ENSMUSG00000032413 | Rasa2 | ENSMUST00000034984 | 28.0445151 | 33 | 20.1589825 | 19.2686804 | 0 | 0 | 0 | 1 | 0 | 0 | 0 | 0 | 1 | 2 | 0 | 1 | 7 | 0 | 10 |
| ENSMUSG00000070565 | Rasal2 | ENSMUST00000078308 | 27.8350515 | 31 | 20.0217037 | 20.90794 | 2 | 0 | 0 | 1 | 0 | 0 | 0 | 0 | 1 | 2 | 0 | 1 | 7 | 1 | 13 |
| ENSMUSG00000029641 | Rasl11a | ENSMUST00000031646 | 26.2948207 | 42 | 16.7330677 | 15.1394422 | 0 | 0 | 0 | 0 | 0 | 0 | 0 | 0 | 0 | 0 | 1 | 0 | 6 | 0 | 7 |
| ENSMUSG00000049907 | Rasl11b | ENSMUST00000051937 | 23.8359202 | 28 | 18.8470067 | 28.8248337 | 1 | 0 | 0 | 1 | 0 | 0 | 0 | 0 | 1 | 2 | 0 | 2 | 0 | 0 | 5 |
| ENSMUSG00000025795 | Rassf3 | ENSMUST00000026902 | 24.4006187 | 28 | 22.58314 | 25.4447022 | 0 | 0 | 0 | 0 | 0 | 0 | 0 | 0 | 0 | 0 | 0 | 1 | 1 | 0 | 2 |
| ENSMUSG00000022105 | Rb1 | ENSMUST00000022701 | 28.6132241 | 35 | 19.1925102 | 17.4956115 | 0 | 0 | 0 | 1 | 0 | 0 | 0 | 0 | 1 | 2 | 1 | 1 | 1 | 0 | 5 |
| ENSMUSG00000025907 | Rb1cc1 | ENSMUST00000027040 | 32.5747998 | 34 | 15.339233 | 17.7833965 | 1 | 0 | 0 | 2 | 0 | 0 | 0 | 0 | 2 | 4 | 4 | 0 | 4 | 3 | 16 |
| ENSMUSG00000057236 | Rbbp4 | ENSMUST00000102598 | 26.9385027 | 32 | 18.7834225 | 22.2927807 | 0 | 0 | 0 | 0 | 0 | 0 | 0 | 0 | 0 | 0 | 1 | 0 | 2 | 1 | 4 |
| ENSMUSG00000026439 | Rbbp5 | ENSMUST00000027700 | 22.7102303 | 32 | 22.6031066 | 22.6031066 | 0 | 0 | 0 | 0 | 0 | 0 | 0 | 0 | 0 | 0 | 0 | 1 | 0 | 0 | 1 |
| ENSMUSG00000031353 | Rbbp7 | ENSMUST00000033720 | 31.2130178 | 33 | 16.4201183 | 19.2307692 | 0 | 0 | 0 | 0 | 0 | 0 | 0 | 0 | 0 | 0 | 0 | 0 | 1 | 0 | 1 |
| ENSMUSG00000041238 | Rbbp8 | ENSMUST00000047322 | 26.9230769 | 41 | 14.3724696 | 17.611336 | 0 | 0 | 1 | 0 | 0 | 0 | 0 | 0 | 0 | 1 | 0 | 0 | 0 | 0 | 1 |
| ENSMUSG00000008658 | Rbfox1 | ENSMUST00000056416 | 32.3330833 | 30 | 18.492123 | 19.5048762 | 0 | 0 | 0 | 0 | 0 | 0 | 0 | 0 | 0 | 0 | 0 | 1 | 0 | 0 | 1 |
| ENSMUSG00000033565 | Rbfox2 | ENSMUST00000111581 | 22.5681686 | 29 | 23.0129569 | 25.3722684 | 0 | 0 | 0 | 0 | 0 | 0 | 0 | 0 | 0 | 0 | 0 | 11 | 8 | 0 | 19 |
| ENSMUSG00000027641 | Rbl1 | ENSMUST00000029170 | 28.2703038 | 36 | 17.3589585 | 18.1649101 | 0 | 0 | 0 | 0 | 0 | 0 | 0 | 0 | 0 | 0 | 0 | 0 | 0 | 2 | 2 |
| ENSMUSG00000089824 | Rbm12 | ENSMUST00000059647 | 28.6174575 | 36 | 15.6121851 | 19.7129467 | 1 | 0 | 0 | 0 | 0 | 0 | 0 | 0 | 0 | 0 | 0 | 0 | 2 | 2 | 5 |
| ENSMUSG00000052137 | Rbm12b2 | ENSMUST00000063839 | 29.3624161 | 40 | 13.9261745 | 16.442953 | 0 | 0 | 0 | 2 | 0 | 0 | 0 | 0 | 2 | 4 | 0 | 0 | 6 | 1 | 11 |
| ENSMUSG00000048109 | Rbm15 | ENSMUST00000061772 | 25.9001161 | 33 | 17.6538908 | 22.9965157 | 1 | 0 | 0 | 0 | 0 | 0 | 0 | 0 | 0 | 0 | 0 | 0 | 0 | 0 | 1 |

| | | | | | | | | | | | | | | | | | | | | | |
|---|---|---|---|---|---|---|---|---|---|---|---|---|---|---|---|---|---|---|---|---|---|
| ENSMUSG00000038132 | Rbm24 | ENSMUST00000037923 | 26.3959391 | 33 | 19.0073322 | 21.3761985 | 2 | 1 | 0 | 1 | 0 | 0 | 0 | 0 | 1 | 2 | 0 | 1 | 3 | 2 | 11 |
| ENSMUSG00000022119 | Rbm26 | ENSMUST00000172122 | 30.1550643 | 37 | 15.7703728 | 16.8921148 | 0 | 0 | 0 | 0 | 0 | 0 | 0 | 0 | 0 | 0 | 2 | 1 | 3 | 1 | 7 |
| ENSMUSG00000024491 | Rbm27 | ENSMUST00000091920 | 29.7685034 | 34 | 17.1503097 | 19.2044343 | 0 | 0 | 0 | 0 | 0 | 0 | 0 | 0 | 0 | 0 | 0 | 1 | 8 | 0 | 9 |
| ENSMUSG00000048271 | Rbm33 | ENSMUST00000059644 | 23.5639338 | 31 | 22.0167171 | 23.7417749 | 0 | 0 | 0 | 0 | 0 | 0 | 0 | 0 | 0 | 0 | 0 | 2 | 0 | 0 | 2 |
| ENSMUSG00000033931 | Rbm34 | ENSMUST00000045994 | 28.6893204 | 31 | 19.3203883 | 20.6796117 | 0 | 0 | 0 | 0 | 0 | 0 | 0 | 0 | 0 | 0 | 2 | 0 | 3 | 0 | 5 |
| ENSMUSG00000031433 | Rbm41 | ENSMUST00000113011 | 35.786802 | 29 | 16.751269 | 18.1472081 | 2 | 1 | 0 | 3 | 0 | 0 | 0 | 1 | 2 | 6 | 0 | 0 | 0 | 0 | 9 |
| ENSMUSG00000042369 | Rbm45 | ENSMUST00000046389 | 29.1925466 | 37 | 16.7701863 | 16.7701863 | 0 | 0 | 0 | 0 | 0 | 0 | 0 | 0 | 0 | 0 | 0 | 0 | 18 | 0 | 18 |
| ENSMUSG00000033882 | Rbm46 | ENSMUST00000182818 | 31.5789474 | 38 | 12.8482972 | 17.6470588 | 0 | 0 | 0 | 0 | 0 | 0 | 0 | 0 | 0 | 0 | 1 | 0 | 0 | 0 | 1 |
| ENSMUSG00000032580 | Rbm5 | ENSMUST00000182371 | 28.9782245 | 23 | 23.241206 | 24.2881072 | 0 | 0 | 0 | 0 | 0 | 0 | 0 | 0 | 0 | 0 | 0 | 1 | 0 | 0 | 1 |
| ENSMUSG00000038374 | Rbm8a | ENSMUST00000198027 | 25.6872465 | 32 | 18.2514646 | 23.9747634 | 2 | 1 | 1 | 1 | 0 | 0 | 0 | 0 | 1 | 3 | 0 | 3 | 3 | 3 | 15 |
| ENSMUSG00000026970 | Rbms1 | ENSMUST00000028347 | 29.6746514 | 33 | 17.0539864 | 19.9141938 | 0 | 0 | 0 | 1 | 0 | 0 | 0 | 0 | 1 | 2 | 0 | 13 | 3 | 0 | 18 |
| ENSMUSG00000039607 | Rbms3 | ENSMUST00000111773 | 29.8319328 | 32 | 19.3781513 | 19.1260504 | 1 | 0 | 0 | 1 | 0 | 0 | 0 | 0 | 1 | 2 | 0 | 4 | 4 | 3 | 14 |
| ENSMUSG00000031134 | Rbmx | ENSMUST00000098470 | 28.8442211 | 30 | 21.1055276 | 20.3015075 | 0 | 0 | 0 | 0 | 0 | 0 | 0 | 0 | 0 | 0 | 2 | 2 | 5 | 0 | 9 |
| ENSMUSG00000039191 | Rbpj | ENSMUST00000037618 | 25.0133618 | 34 | 20.8177445 | 20.3099947 | 0 | 0 | 0 | 0 | 0 | 0 | 0 | 0 | 0 | 0 | 7 | 0 | 10 | 0 | 17 |
| ENSMUSG00000032387 | Rbpms2 | ENSMUST00000169003 | 22.7686703 | 29 | 25.4098361 | 22.7686703 | 0 | 0 | 0 | 0 | 0 | 0 | 0 | 0 | 0 | 0 | 0 | 1 | 1 | 0 | 2 |
| ENSMUSG00000022400 | Rbx1; Gm9840 | ENSMUST00000023036 | 28.0094414 | 28 | 20.0629426 | 24.2328875 | 1 | 0 | 0 | 0 | 0 | 0 | 0 | 0 | 0 | 0 | 0 | 0 | 0 | 0 | 1 |
| ENSMUSG00000040423 | Rc3h1 | ENSMUST00000161609 | 28.1715638 | 35 | 17.7969642 | 19.3566356 | 0 | 0 | 0 | 2 | 0 | 0 | 0 | 0 | 2 | 4 | 0 | 2 | 13 | 6 | 25 |
| ENSMUSG00000075376 | Rc3h2 | ENSMUST00000112934 | 28.8126055 | 37 | 16.3946727 | 17.6514725 | 1 | 1 | 0 | 2 | 0 | 0 | 0 | 0 | 2 | 4 | 1 | 0 | 8 | 4 | 19 |
| ENSMUSG00000022106 | Rcbtb2 | ENSMUST00000169513 | 29.9567634 | 33 | 17.3563928 | 19.5182211 | 0 | 0 | 0 | 0 | 0 | 0 | 0 | 0 | 0 | 0 | 0 | 1 | 1 | 0 | 2 |
| ENSMUSG00000029397 | Rchy1 | ENSMUST00000031345 | 24.8798077 | 36 | 20.6730769 | 18.6298077 | 0 | 0 | 0 | 0 | 0 | 0 | 0 | 0 | 0 | 0 | 0 | 21 | 0 | 0 | 21 |
| ENSMUSG00000032320 | Rcn2 | ENSMUST00000147842 | 29.7704178 | 31 | 18.3289424 | 21.1893113 | 0 | 0 | 0 | 0 | 0 | 0 | 0 | 0 | 0 | 0 | 0 | 0 | 5 | 0 | 5 |
| ENSMUSG00000037896 | Rcor1 | ENSMUST00000084968 | 25.1881586 | 30 | 22.4034119 | 22.5790266 | 1 | 0 | 0 | 1 | 0 | 0 | 0 | 0 | 1 | 2 | 1 | 4 | 1 | 0 | 9 |
| ENSMUSG00000037395 | Rcor3 | ENSMUST00000192158 | 28.3535636 | 30 | 21.5375039 | 20.3548086 | 1 | 1 | 1 | 1 | 0 | 0 | 0 | 0 | 1 | 3 | 0 | 0 | 7 | 0 | 12 |
| ENSMUSG00000020907 | Rcvrn | ENSMUST00000021290 | 21.4705882 | 25 | 32.9411765 | 20.8823529 | 0 | 0 | 0 | 0 | 0 | 0 | 0 | 0 | 0 | 0 | 0 | 2 | 0 | 0 | 2 |
| ENSMUSG00000025921 | Rdh10 | ENSMUST00000027053 | 23.3059991 | 36 | 19.2921882 | 21.3206733 | 0 | 0 | 0 | 0 | 0 | 0 | 0 | 0 | 0 | 0 | 1 | 0 | 2 | 0 | 3 |
| ENSMUSG00000066441 | Rdh11 | ENSMUST00000161204 | 29.9225664 | 28 | 20.1327434 | 21.9579646 | 0 | 0 | 0 | 0 | 0 | 0 | 0 | 0 | 0 | 0 | 0 | 1 | 0 | 0 | 1 |
| ENSMUSG00000008435 | Rdh13 | ENSMUST00000008579 | 24.8545743 | 29 | 21.8931782 | 24.5901639 | 0 | 0 | 0 | 0 | 0 | 0 | 0 | 0 | 0 | 0 | 0 | 1 | 12 | 1 | 14 |
| ENSMUSG00000020621 | Rdh14 | ENSMUST00000020947 | 32.9238329 | 32 | 14.4963145 | 20.1474201 | 0 | 0 | 1 | 0 | 0 | 0 | 0 | 0 | 0 | 1 | 0 | 0 | 0 | 0 | 1 |
| ENSMUSG00000032050 | Rdx | ENSMUST00000000590 | 26.3507529 | 34 | 18.6448184 | 20.5491585 | 0 | 0 | 0 | 0 | 0 | 0 | 0 | 0 | 0 | 0 | 1 | 0 | 1 | 0 | 2 |
| ENSMUSG00000028476 | Reck | ENSMUST00000030198 | 27.9666898 | 31 | 20.9576683 | 19.8473282 | 0 | 0 | 0 | 1 | 0 | 0 | 0 | 0 | 1 | 2 | 0 | 0 | 0 | 0 | 2 |
| ENSMUSG00000052852 | Reep1 | ENSMUST00000121469 | 25.2664146 | 29 | 22.2756961 | 23.3413544 | 0 | 0 | 0 | 0 | 0 | 0 | 0 | 0 | 0 | 0 | 0 | 0 | 0 | 2 | 2 |
| ENSMUSG00000019873 | Reep3 | ENSMUST00000020023 | 26.9323937 | 32 | 21.8013653 | 18.7844087 | 0 | 0 | 0 | 0 | 0 | 0 | 0 | 0 | 0 | 0 | 2 | 2 | 4 | 1 | 9 |
| ENSMUSG00000020275 | Rel | ENSMUST00000102864 | 26.5411591 | 33 | 21.1517165 | 19.5090439 | 1 | 0 | 0 | 0 | 0 | 0 | 0 | 0 | 0 | 0 | 0 | 4 | 16 | 0 | 21 |
| ENSMUSG00000047881 | Rell1 | ENSMUST00000087327 | 28.3138918 | 27 | 20.7140332 | 23.8600212 | 0 | 0 | 0 | 0 | 0 | 0 | 0 | 0 | 0 | 0 | 3 | 1 | 3 | 2 | 9 |
| ENSMUSG00000040855 | Reps2 | ENSMUST00000101102 | 29.7906602 | 33 | 18.1248882 | 19.3952407 | 1 | 2 | 0 | 2 | 0 | 0 | 0 | 1 | 1 | 4 | 4 | 1 | 4 | 0 | 16 |
| ENSMUSG00000032026 | Rexo2 | ENSMUST00000034524 | 23.1023102 | 31 | 29.7029703 | 15.8415842 | 0 | 0 | 0 | 0 | 0 | 0 | 0 | 0 | 0 | 0 | 0 | 2 | 0 | 0 | 2 |
| ENSMUSG00000024712 | Rfk | ENSMUST00000025617 | 26.5750286 | 32 | 20.5612829 | 20.6185567 | 0 | 0 | 0 | 0 | 0 | 0 | 0 | 0 | 0 | 0 | 0 | 0 | 3 | 4 | 7 |
| ENSMUSG00000040782 | Rfwd2 | ENSMUST00000076894 | 31.834404 | 31 | 18.9507495 | 18.2369736 | 0 | 0 | 0 | 0 | 0 | 0 | 0 | 0 | 0 | 0 | 0 | 0 | 3 | 2 | 5 |
| ENSMUSG00000040929 | Rfx3 | ENSMUST00000165566 | 29.5069795 | 34 | 17.8942679 | 18.8595189 | 0 | 2 | 0 | 3 | 0 | 0 | 0 | 0 | 3 | 6 | 0 | 17 | 6 | 4 | 35 |
| ENSMUSG00000022018 | Rgcc | ENSMUST00000022595 | 28.0898876 | 31 | 19.3820225 | 21.3483146 | 0 | 0 | 0 | 1 | 0 | 0 | 0 | 0 | 1 | 2 | 0 | 0 | 0 | 0 | 2 |
| ENSMUSG00000048027 | Rgmb | ENSMUST00000170578 | 23.7668161 | 38 | 19.8430493 | 18.161435 | 0 | 0 | 0 | 0 | 0 | 0 | 0 | 0 | 0 | 0 | 1 | 0 | 4 | 0 | 5 |
| ENSMUSG00000026475 | Rgs16 | ENSMUST00000027748 | 23.0626162 | 24 | 24.3025418 | 28.2703038 | 2 | 1 | 0 | 2 | 0 | 0 | 0 | 1 | 1 | 4 | 0 | 0 | 0 | 0 | 7 |
| ENSMUSG00000019775 | Rgs17 | ENSMUST00000064225 | 31.6460905 | 33 | 17.6954733 | 17.5720165 | 0 | 0 | 0 | 1 | 0 | 0 | 0 | 0 | 1 | 2 | 4 | 0 | 5 | 1 | 12 |
| ENSMUSG00000026360 | Rgs2 | ENSMUST00000127206 | 28.9069172 | 29 | 21.7762596 | 20.5380017 | 1 | 0 | 0 | 1 | 0 | 0 | 0 | 0 | 1 | 2 | 0 | 0 | 0 | 0 | 3 |
| ENSMUSG00000026678 | Rgs5 | ENSMUST00000027997 | 30.4923697 | 30 | 19.5508206 | 20.1842787 | 0 | 0 | 0 | 1 | 0 | 0 | 0 | 0 | 1 | 2 | 0 | 3 | 2 | 0 | 7 |
| ENSMUSG00000021719 | Rgs7bp | ENSMUST00000063551 | 31.3047487 | 29 | 20.6546796 | 18.6030429 | 0 | 0 | 1 | 0 | 0 | 0 | 0 | 0 | 0 | 1 | 0 | 0 | 0 | 0 | 1 |
| ENSMUSG00000028945 | Rheb | ENSMUST00000030787 | 23.7812128 | 35 | 22.7110583 | 18.3115339 | 0 | 0 | 0 | 0 | 0 | 0 | 0 | 0 | 0 | 0 | 0 | 0 | 3 | 0 | 3 |
| ENSMUSG00000007815 | Rhoa | ENSMUST00000007959 | 25.0673854 | 36 | 21.3836478 | 17.9694519 | 0 | 0 | 0 | 0 | 0 | 0 | 0 | 0 | 0 | 0 | 0 | 1 | 7 | 2 | 10 |
| ENSMUSG00000021589 | Rhobtb3 | ENSMUST00000022078 | 30.5802777 | 34 | 16.2335351 | 19.508722 | 0 | 0 | 0 | 0 | 0 | 0 | 0 | 0 | 0 | 0 | 0 | 0 | 6 | 0 | 6 |
| ENSMUSG00000017686 | Rhot1 | ENSMUST00000055056 | 25.5371094 | 34 | 19.140625 | 21.3867188 | 0 | 0 | 0 | 0 | 0 | 0 | 0 | 0 | 0 | 0 | 0 | 1 | 1 | 0 | 2 |
| ENSMUSG00000038658 | Ric1 | ENSMUST00000043610 | 29.4776119 | 34 | 17.238806 | 18.8432836 | 0 | 0 | 0 | 0 | 0 | 0 | 0 | 0 | 0 | 0 | 1 | 0 | 18 | 1 | 20 |
| ENSMUSG00000050310 | Rictor | ENSMUST00000061656 | 29.1206151 | 32 | 18.5007208 | 20.06247 | 0 | 0 | 0 | 2 | 0 | 0 | 0 | 0 | 2 | 4 | 1 | 0 | 7 | 0 | 12 |
| ENSMUSG00000036202 | Rif1 | ENSMUST00000069794 | 32.7966102 | 38 | 13.1355932 | 15.7627119 | 1 | 0 | 0 | 3 | 0 | 0 | 0 | 0 | 3 | 6 | 0 | 0 | 0 | 0 | 7 |
| ENSMUSG00000040649 | Rimklb | ENSMUST00000068242 | 29.2635659 | 35 | 17.2803618 | 18.9276486 | 0 | 1 | 0 | 1 | 0 | 0 | 0 | 0 | 1 | 2 | 7 | 1 | 8 | 0 | 19 |
| ENSMUSG00000041135 | Ripk2 | ENSMUST00000183871 | 26.4139217 | 29 | 22.8713487 | 22.0633934 | 0 | 0 | 1 | 0 | 0 | 0 | 0 | 0 | 0 | 1 | 0 | 0 | 0 | 0 | 1 |
| ENSMUSG00000057455 | Rit2 | ENSMUST00000082070 | 30.8809747 | 32 | 18.6972821 | 17.9475164 | 0 | 0 | 0 | 0 | 0 | 0 | 0 | 0 | 0 | 0 | 1 | 6 | 0 | 0 | 7 |
| ENSMUSG00000056537 | Rlim | ENSMUST00000070705 | 27.9466967 | 34 | 17.4174174 | 20.7957958 | 0 | 0 | 0 | 0 | 0 | 0 | 0 | 0 | 0 | 0 | 0 | 1 | 10 | 1 | 12 |
| ENSMUSG00000039097 | Rln1 | ENSMUST00000044143 | 21.2389381 | 39 | 22.1238938 | 17.699115 | 0 | 0 | 1 | 0 | 0 | 0 | 0 | 0 | 0 | 1 | 0 | 0 | 0 | 0 | 1 |
| ENSMUSG00000035367 | Rmi1 | ENSMUST00000042450 | 27.5 | 32 | 19.3382353 | 21.6176471 | 0 | 0 | 0 | 1 | 0 | 0 | 0 | 0 | 1 | 2 | 1 | 0 | 3 | 0 | 6 |
| ENSMUSG00000019763 | Rmnd1 | ENSMUST00000042251 | 28.3140283 | 28 | 23.4234234 | 20.7207207 | 0 | 0 | 0 | 0 | 0 | 0 | 0 | 0 | 0 | 0 | 0 | 0 | 6 | 0 | 6 |
| ENSMUSG00000020630 | Rnaseh1 | ENSMUST00000020959 | 27.9296875 | 28 | 21.09375 | 22.65625 | 0 | 0 | 0 | 0 | 0 | 0 | 0 | 0 | 0 | 0 | 0 | 0 | 0 | 1 | 1 |
| ENSMUSG00000094724 | Rnaset2b | ENSMUST00000179728 | 26.4426126 | 30 | 22.8281547 | 21.1794547 | 0 | 0 | 0 | 0 | 0 | 0 | 0 | 0 | 0 | 0 | 0 | 3 | 0 | 0 | 3 |
| ENSMUSG00000054855 | Rnd1 | ENSMUST00000003451 | 21.9109195 | 27 | 22.4137931 | 28.591954 | 0 | 0 | 0 | 0 | 0 | 0 | 0 | 0 | 0 | 0 | 0 | 0 | 8 | 0 | 8 |
| ENSMUSG00000017144 | Rnd3 | ENSMUST00000154545 | 29.8774741 | 28 | 19.4156456 | 22.667295 | 0 | 0 | 0 | 1 | 0 | 0 | 0 | 0 | 1 | 2 | 0 | 0 | 1 | 0 | 3 |
| ENSMUSG00000052656 | Rnf103 | ENSMUST00000114178 | 29.7169811 | 31 | 17.5943396 | 21.2735849 | 1 | 1 | 0 | 1 | 0 | 0 | 0 | 0 | 1 | 2 | 0 | 0 | 0 | 0 | 4 |
| ENSMUSG00000028557 | Rnf11 | ENSMUST00000030284 | 28.5759594 | 35 | 16.5556613 | 20.0444022 | 0 | 0 | 0 | 0 | 0 | 0 | 0 | 0 | 0 | 0 | 1 | 0 | 0 | 0 | 1 |
| ENSMUSG00000032217 | Rnf111 | ENSMUST00000034739 | 28.3151826 | 38 | 16.8481742 | 17.2325432 | 0 | 0 | 1 | 0 | 0 | 0 | 0 | 0 | 0 | 1 | 0 | 1 | 7 | 1 | 10 |
| ENSMUSG00000036503 | Rnf13 | ENSMUST00000197205 | 30.117291 | 32 | 18.5773742 | 18.9557321 | 0 | 0 | 0 | 1 | 0 | 0 | 0 | 0 | 1 | 2 | 0 | 0 | 0 | 1 | 3 |
| ENSMUSG00000024317 | Rnf138 | ENSMUST00000052396 | 31.5611406 | 37 | 13.8714355 | 17.3513775 | 0 | 0 | 0 | 0 | 0 | 0 | 0 | 0 | 0 | 0 | 0 | 0 | 0 | 1 | 1 |
| ENSMUSG00000037075 | Rnf139 | ENSMUST00000036904 | 32.358156 | 30 | 16.9769504 | 20.3457447 | 0 | 0 | 0 | 1 | 0 | 0 | 0 | 0 | 1 | 2 | 0 | 0 | 1 | 1 | 4 |
| ENSMUSG00000030788 | Rnf141 | ENSMUST00000177236 | 28.1476807 | 32 | 19.5329757 | 20.6058694 | 1 | 0 | 0 | 1 | 0 | 0 | 0 | 0 | 1 | 2 | 0 | 0 | 0 | 0 | 3 |
| ENSMUSG00000038876 | Rnf146 | ENSMUST00000037548 | 30.1786393 | 32 | 18.6240973 | 18.9661726 | 1 | 2 | 0 | 2 | 0 | 0 | 0 | 1 | 1 | 4 | 0 | 0 | 0 | 1 | 8 |
| ENSMUSG00000047747 | Rnf150 | ENSMUST00000078525 | 27.5884411 | 28 | 22.223662 | 22.3014125 | 0 | 0 | 0 | 0 | 0 | 0 | 0 | 0 | 0 | 0 | 0 | 18 | 3 | 0 | 21 |
| ENSMUSG00000047496 | Rnf152 | ENSMUST00000058688 | 27.8439425 | 32 | 19.137577 | 21.1635866 | 1 | 0 | 0 | 1 | 0 | 0 | 0 | 0 | 1 | 2 | 1 | 46 | 1 | 2 | 53 |
| ENSMUSG00000044164 | Rnf182 | ENSMUST00000059986 | 24.831612 | 31 | 20.2514594 | 23.8886394 | 0 | 0 | 0 | 0 | 0 | 0 | 0 | 0 | 0 | 0 | 0 | 0 | 1 | 0 | 1 |
| ENSMUSG00000045078 | Rnf216 | ENSMUST00000053498 | 21.9817768 | 26 | 26.594533 | 25.7972665 | 0 | 0 | 0 | 0 | 0 | 0 | 0 | 0 | 0 | 0 | 0 | 1 | 0 | 0 | 1 |
| ENSMUSG00000063760 | Rnf217 | ENSMUST00000081989 | 30.6603275 | 34 | 17.7284733 | 17.3375594 | 1 | 0 | 0 | 0 | 0 | 0 | 0 | 0 | 0 | 0 | 1 | 2 | 2 | 4 | 10 |
| ENSMUSG00000022120 | Rnf219 | ENSMUST00000022716 | 23.4387672 | 36 | 16.6261152 | 23.6820762 | 0 | 0 | 0 | 0 | 0 | 0 | 0 | 0 | 0 | 0 | 2 | 1 | 0 | 0 | 3 |
| ENSMUSG00000048911 | Rnf24 | ENSMUST00000059372 | 25.2138341 | 29 | 24.8233544 | 21.3648196 | 0 | 0 | 0 | 0 | 0 | 0 | 0 | 0 | 0 | 0 | 0 | 0 | 0 | 2 | 2 |
| ENSMUSG00000029130 | Rnf32 | ENSMUST00000160383 | 27.6150628 | 26 | 22.3152022 | 23.5704324 | 0 | 0 | 0 | 0 | 0 | 0 | 0 | 0 | 0 | 0 | 1 | 0 | 0 | 0 | 1 |
| ENSMUSG00000029474 | Rnf34 | ENSMUST00000031434 | 29.7650131 | 30 | 19.7780679 | 20.3002611 | 1 | 0 | 0 | 1 | 0 | 0 | 0 | 0 | 1 | 2 | 0 | 0 | 4 | 2 | 9 |
| ENSMUSG00000035696 | Rnf38 | ENSMUST00000098098 | 24.7044917 | 35 | 20.1832151 | 20.2423168 | 0 | 0 | 0 | 0 | 0 | 0 | 0 | 0 | 0 | 0 | 2 | 1 | 7 | 2 | 12 |
| ENSMUSG00000034928 | Rnf44 | ENSMUST00000177950 | 20.8085106 | 28 | 26.6808511 | 24.8510638 | 0 | 0 | 0 | 0 | 0 | 0 | 0 | 0 | 0 | 0 | 3 | 0 | 2 | 0 | 5 |
| ENSMUSG00000015478 | Rnf5 | ENSMUST00000015622 | 23.0125523 | 30 | 26.3598326 | 20.292887 | 0 | 0 | 0 | 0 | 0 | 0 | 0 | 0 | 0 | 0 | 0 | 2 | 0 | 0 | 2 |
| ENSMUSG00000029634 | Rnf6 | ENSMUST00000161859 | 32.5420376 | 34 | 13.5509397 | 19.5845697 | 0 | 0 | 0 | 1 | 0 | 0 | 0 | 0 | 1 | 2 | 0 | 0 | 0 | 3 | 5 |
| ENSMUSG00000020521 | Rnft1 | ENSMUST00000020827 | 31.9218241 | 31 | 17.8175896 | 19.218241 | 0 | 0 | 0 | 1 | 0 | 0 | 0 | 0 | 1 | 2 | 0 | 0 | 0 | 1 | 3 |
| ENSMUSG00000028274 | Rngtt | ENSMUST00000108153 | 26.4798915 | 33 | 18.7980117 | 21.3737009 | 1 | 0 | 0 | 0 | 0 | 0 | 0 | 0 | 0 | 0 | 1 | 0 | 0 | 0 | 2 |
| ENSMUSG00000009535 | Rnmt | ENSMUST00000009679 | 27.3949099 | 30 | 20.1029454 | 22.3906205 | 0 | 0 | 0 | 0 | 0 | 0 | 0 | 0 | 0 | 0 | 1 | 0 | 1 | 0 | 2 |
| ENSMUSG00000027981 | Rnpc3 | ENSMUST00000153853 | 31.9647202 | 38 | 14.1727494 | 16.3017032 | 1 | 0 | 0 | 1 | 0 | 0 | 0 | 0 | 1 | 2 | 0 | 0 | 0 | 1 | 4 |
| ENSMUSG00000022883 | Robo1 | ENSMUST00000023600 | 30.813618 | 35 | 17.541835 | 17.0225043 | 1 | 0 | 0 | 1 | 0 | 0 | 0 | 0 | 1 | 2 | 0 | 0 | 3 | 0 | 6 |
| ENSMUSG00000052516 | Robo2 | ENSMUST00000117785 | 32.4715909 | 32 | 17.2727273 | 17.9545455 | 0 | 0 | 1 | 0 | 0 | 0 | 0 | 0 | 0 | 1 | 0 | 0 | 0 | 1 | 2 |
| ENSMUSG00000024290 | Rock1 | ENSMUST00000067947 | 31.9975713 | 37 | 13.8433515 | 17.5470553 | 0 | 0 | 0 | 1 | 0 | 0 | 0 | 0 | 1 | 2 | 0 | 0 | 10 | 0 | 12 |

| | | | | | | | | | | | | | | | | | | | | | |
|---|---|---|---|---|---|---|---|---|---|---|---|---|---|---|---|---|---|---|---|---|---|
| ENSMUSG00000020580 | Rock2 | ENSMUST00000020904 | 29.3696669 | 33 | 17.9466006 | 20.0110102 | 0 | 0 | 0 | 1 | 0 | 0 | 0 | 0 | 1 | 2 | 0 | 5 | 0 | 1 | 8 |
| ENSMUSG00000036192 | Rorb | ENSMUST00000040153 | 29.8831615 | 31 | 20.0274914 | 19.4776632 | 0 | 0 | 0 | 0 | 0 | 0 | 0 | 0 | 0 | 0 | 7 | 12 | 46 | 0 | 65 |
| ENSMUSG00000060090 | Rp2 | ENSMUST00000033372 | 33.4188582 | 36 | 14.5285439 | 16.3887107 | 0 | 0 | 0 | 0 | 0 | 0 | 0 | 0 | 0 | 0 | 1 | 0 | 0 | 0 | 1 |
| ENSMUSG00000028884 | Rpa2 | ENSMUST00000102561 | 21.4123007 | 29 | 20.9567198 | 28.8154897 | 0 | 0 | 0 | 0 | 0 | 0 | 0 | 0 | 0 | 0 | 0 | 1 | 0 | 0 | 1 |
| ENSMUSG00000012483 | Rpa3 | ENSMUST00000012627 | 34.0611354 | 40 | 13.5371179 | 12.6637555 | 1 | 0 | 1 | 1 | 0 | 0 | 0 | 0 | 1 | 3 | 0 | 0 | 0 | 0 | 4 |
| ENSMUSG00000026005 | Rpe | ENSMUST00000142920 | 32.5657895 | 31 | 16.8174342 | 19.6134868 | 0 | 0 | 0 | 0 | 0 | 0 | 0 | 0 | 0 | 0 | 0 | 0 | 0 | 1 | 1 |
| ENSMUSG00000031174 | Rpgr | ENSMUST00000115533 | 33.9055794 | 31 | 17.8111588 | 16.9527897 | 2 | 1 | 2 | 1 | 0 | 0 | 0 | 0 | 1 | 4 | 0 | 0 | 0 | 0 | 7 |
| ENSMUSG00000033282 | Rpgrip1l | ENSMUST00000139113 | 32.2610084 | 27 | 19.7809719 | 21.1955282 | 0 | 0 | 0 | 0 | 0 | 0 | 0 | 0 | 0 | 0 | 1 | 0 | 1 | 0 | 2 |
| ENSMUSG00000053604 | Rpia | ENSMUST00000066134 | 24.7002398 | 33 | 18.705036 | 23.8609113 | 0 | 0 | 0 | 0 | 0 | 0 | 0 | 0 | 0 | 0 | 0 | 0 | 0 | 1 | 1 |
| ENSMUSG00000047215 | Rpl9 | ENSMUST00000127874 | 25.0127877 | 31 | 16.9820972 | 27.1099744 | 0 | 0 | 0 | 0 | 0 | 0 | 0 | 0 | 0 | 0 | 2 | 0 | 1 | 0 | 3 |
| ENSMUSG00000040446 | Rprd1a | ENSMUST00000046206 | 27.1118012 | 32 | 19.5652174 | 21.7080745 | 0 | 0 | 0 | 0 | 0 | 0 | 0 | 0 | 0 | 0 | 1 | 0 | 1 | 1 | 3 |
| ENSMUSG00000008683 | Rps15a | ENSMUST00000131374 | 24.1962775 | 31 | 21.4043993 | 23.7732657 | 0 | 0 | 0 | 0 | 0 | 0 | 0 | 0 | 0 | 0 | 0 | 2 | 1 | 1 | 4 |
| ENSMUSG00000028234 | Rps20 | ENSMUST00000138502 | 28.4180613 | 33 | 18.7559204 | 20.1452479 | 0 | 0 | 0 | 0 | 0 | 0 | 0 | 0 | 0 | 0 | 0 | 0 | 2 | 1 | 3 |
| ENSMUSG00000031309 | Rps6ka3 | ENSMUST00000033671 | 29.9222526 | 32 | 18.9955873 | 18.7644463 | 0 | 0 | 0 | 0 | 0 | 0 | 0 | 0 | 0 | 0 | 0 | 1 | 1 | 0 | 2 |
| ENSMUSG00000021180 | Rps6ka5 | ENSMUST00000043599 | 28.6914766 | 35 | 17.4069628 | 18.4273709 | 0 | 0 | 0 | 0 | 0 | 0 | 0 | 0 | 0 | 0 | 0 | 0 | 1 | 1 | 2 |
| ENSMUSG00000070934 | Rraga | ENSMUST00000091064 | 28 | 39 | 13.6470588 | 19.7647059 | 0 | 0 | 0 | 0 | 0 | 0 | 0 | 0 | 0 | 0 | 0 | 0 | 3 | 0 | 3 |
| ENSMUSG00000055723 | Rras2 | ENSMUST00000069449 | 27.0485859 | 34 | 18.6366933 | 19.9419869 | 0 | 0 | 0 | 0 | 0 | 0 | 0 | 0 | 0 | 0 | 0 | 0 | 0 | 1 | 1 |
| ENSMUSG00000020649 | Rrm2 | ENSMUST00000020980 | 28.5557987 | 33 | 19.0371991 | 19.1466083 | 0 | 0 | 0 | 0 | 0 | 0 | 0 | 0 | 0 | 0 | 0 | 0 | 5 | 1 | 6 |
| ENSMUSG00000022292 | Rrm2b | ENSMUST00000022901 | 31.1873671 | 32 | 17.6738536 | 18.8278166 | 1 | 1 | 0 | 2 | 0 | 0 | 0 | 0 | 2 | 4 | 0 | 10 | 0 | 0 | 16 |
| ENSMUSG00000004896 | Rrnad1 | ENSMUST00000160648 | 25.7025103 | 25 | 25.3653054 | 24.3536905 | 0 | 0 | 0 | 1 | 0 | 0 | 0 | 0 | 1 | 2 | 0 | 2 | 0 | 0 | 4 |
| ENSMUSG00000044098 | Rsbn1 | ENSMUST00000068879 | 30.1689861 | 36 | 16.1530815 | 17.5695825 | 0 | 0 | 0 | 0 | 0 | 0 | 0 | 0 | 0 | 0 | 0 | 1 | 3 | 0 | 4 |
| ENSMUSG00000039968 | Rsbn1l | ENSMUST00000036489 | 29.6685264 | 35 | 15.4906465 | 19.4945848 | 1 | 1 | 0 | 3 | 0 | 0 | 0 | 0 | 3 | 6 | 1 | 0 | 2 | 1 | 12 |
| ENSMUSG00000035623 | Rsf1 | ENSMUST00000107153 | 28.9795317 | 33 | 18.1563834 | 19.5700191 | 0 | 0 | 0 | 1 | 0 | 0 | 0 | 0 | 1 | 2 | 1 | 0 | 3 | 2 | 8 |
| ENSMUSG00000032215 | Rsl24d1 | ENSMUST00000034738 | 21.686747 | 35 | 21.4859438 | 22.0883534 | 0 | 0 | 0 | 0 | 0 | 0 | 0 | 0 | 0 | 0 | 0 | 6 | 9 | 3 | 18 |
| ENSMUSG00000051920 | Rspo2 | ENSMUST00000063492 | 26.981352 | 33 | 18.7645688 | 21.3286713 | 0 | 0 | 0 | 0 | 0 | 0 | 0 | 0 | 0 | 0 | 0 | 0 | 1 | 0 | 1 |
| ENSMUSG00000034544 | Rsrc1 | ENSMUST00000161726 | 28.8046087 | 31 | 20.4992799 | 20.1152184 | 0 | 0 | 0 | 0 | 0 | 0 | 0 | 0 | 0 | 0 | 0 | 0 | 1 | 0 | 1 |
| ENSMUSG00000029422 | Rsrc2 | ENSMUST00000111515 | 29.2929293 | 35 | 16.8876263 | 18.844697 | 0 | 0 | 0 | 0 | 0 | 0 | 0 | 0 | 0 | 0 | 0 | 0 | 5 | 2 | 7 |
| ENSMUSG00000037266 | Rsrp1 | ENSMUST00000078084 | 29.3269231 | 33 | 15.7051282 | 21.9551282 | 0 | 0 | 0 | 0 | 0 | 0 | 0 | 0 | 0 | 0 | 0 | 0 | 0 | 1 | 1 |
| ENSMUSG00000026727 | Rsu1 | ENSMUST00000114791 | 28.2163743 | 36 | 22.8070175 | 13.3040936 | 0 | 0 | 0 | 0 | 0 | 0 | 0 | 0 | 0 | 0 | 0 | 1 | 0 | 0 | 1 |
| ENSMUSG00000027304 | Rtf1 | ENSMUST00000028767 | 24.4038156 | 30 | 22.4165342 | 23.5691574 | 0 | 0 | 0 | 0 | 0 | 0 | 0 | 0 | 0 | 0 | 0 | 2 | 1 | 0 | 3 |
| ENSMUSG00000024758 | Rtn3 | ENSMUST00000025667 | 26.5940902 | 30 | 21.87662 | 21.7729393 | 0 | 0 | 0 | 0 | 0 | 0 | 0 | 0 | 0 | 0 | 0 | 0 | 1 | 1 | 2 |
| ENSMUSG00000040570 | Rundc3b | ENSMUST00000047485 | 27.8672986 | 33 | 18.7677725 | 20.4739336 | 0 | 0 | 0 | 0 | 0 | 0 | 0 | 0 | 0 | 0 | 0 | 1 | 0 | 4 | 5 |
| ENSMUSG00000006586 | Runx1t1 | ENSMUST00000006761 | 30.595428 | 32 | 18.9261031 | 18.4476342 | 0 | 0 | 0 | 0 | 0 | 0 | 0 | 0 | 0 | 0 | 2 | 1 | 13 | 1 | 17 |
| ENSMUSG00000028133 | Rwdd3 | ENSMUST00000106466 | 33.5854766 | 34 | 14.8260212 | 17.8517398 | 1 | 0 | 0 | 1 | 0 | 0 | 0 | 0 | 1 | 2 | 0 | 0 | 1 | 0 | 4 |
| ENSMUSG00000031568 | Rwdd4a | ENSMUST00000033973 | 27.7883699 | 33 | 17.2545281 | 22.0209724 | 1 | 0 | 0 | 1 | 0 | 0 | 0 | 0 | 1 | 2 | 1 | 0 | 4 | 2 | 10 |
| ENSMUSG00000072872 | Rybp | ENSMUST00000101118 | 27.4944568 | 33 | 19.0687361 | 20.6485588 | 1 | 0 | 0 | 0 | 0 | 0 | 0 | 0 | 0 | 0 | 0 | 0 | 0 | 0 | 1 |
| ENSMUSG00000040928 | S100pbp | ENSMUST00000072431 | 23.2597623 | 34 | 21.0186757 | 21.8336163 | 0 | 0 | 0 | 0 | 0 | 0 | 0 | 0 | 0 | 0 | 1 | 2 | 6 | 0 | 9 |
| ENSMUSG00000045092 | S1pr1 | ENSMUST00000055676 | 27.4596182 | 31 | 19.8237885 | 21.5859031 | 0 | 0 | 0 | 0 | 0 | 0 | 0 | 0 | 0 | 0 | 0 | 3 | 0 | 0 | 3 |
| ENSMUSG00000051354 | Samd3 | ENSMUST00000164660 | 26.7985612 | 35 | 20.5635492 | 17.6858513 | 0 | 0 | 0 | 1 | 0 | 0 | 0 | 0 | 1 | 2 | 0 | 2 | 0 | 0 | 4 |
| ENSMUSG00000021770 | Samd8 | ENSMUST00000119430 | 27.6805252 | 32 | 19.055434 | 21.7177243 | 0 | 0 | 0 | 0 | 0 | 0 | 0 | 0 | 0 | 0 | 7 | 1 | 29 | 2 | 39 |
| ENSMUSG00000022876 | Samsn1 | ENSMUST00000114239 | 32.4574961 | 37 | 13.9103555 | 16.5378671 | 0 | 0 | 0 | 0 | 0 | 0 | 0 | 0 | 0 | 0 | 1 | 0 | 0 | 0 | 1 |
| ENSMUSG00000021963 | Sap18 | ENSMUST00000128764 | 28.4286699 | 29 | 19.9172984 | 22.7084769 | 1 | 1 | 0 | 0 | 0 | 0 | 0 | 0 | 0 | 0 | 0 | 2 | 2 | 0 | 6 |
| ENSMUSG00000031609 | Sap30 | ENSMUST00000034022 | 26.6968326 | 43 | 13.1221719 | 16.7420814 | 0 | 0 | 0 | 0 | 0 | 0 | 0 | 0 | 0 | 0 | 0 | 0 | 3 | 1 | 4 |
| ENSMUSG00000027959 | Sass6 | ENSMUST00000029571 | 32.2269012 | 27 | 20.4359673 | 20.113946 | 0 | 0 | 0 | 0 | 0 | 0 | 0 | 0 | 0 | 0 | 0 | 1 | 11 | 0 | 12 |
| ENSMUSG00000025283 | Sat1 | ENSMUST00000152463 | 28.7741935 | 32 | 16.6451613 | 22.8387097 | 0 | 0 | 0 | 0 | 0 | 0 | 0 | 0 | 0 | 0 | 0 | 0 | 0 | 1 | 1 |
| ENSMUSG00000023927 | Satb1 | ENSMUST00000144331 | 29.5338184 | 32 | 17.9466451 | 20.6413366 | 0 | 0 | 0 | 0 | 0 | 0 | 0 | 0 | 0 | 0 | 0 | 1 | 0 | 3 | 4 |
| ENSMUSG00000038331 | Satb2 | ENSMUST00000042857 | 26.2312634 | 34 | 21.0920771 | 18.5938615 | 0 | 0 | 0 | 0 | 0 | 0 | 0 | 0 | 0 | 0 | 1 | 13 | 12 | 1 | 27 |
| ENSMUSG00000038570 | Saxo2 | ENSMUST00000056728 | 35.2393617 | 33 | 15.2925532 | 16.6223404 | 0 | 0 | 0 | 0 | 0 | 0 | 0 | 0 | 0 | 0 | 1 | 0 | 0 | 0 | 1 |
| ENSMUSG00000038095 | Sbno1 | ENSMUST00000065263 | 24.3693153 | 32 | 21.1429552 | 22.7732967 | 1 | 1 | 0 | 0 | 0 | 0 | 0 | 0 | 0 | 0 | 1 | 2 | 13 | 1 | 19 |
| ENSMUSG00000032018 | Sc5d | ENSMUST00000052725 | 28.2403433 | 32 | 18.9699571 | 21.2017167 | 0 | 0 | 0 | 0 | 0 | 0 | 0 | 0 | 0 | 0 | 0 | 0 | 0 | 1 | 1 |
| ENSMUSG00000035236 | Scai | ENSMUST00000038874 | 30.1273306 | 36 | 16.4620282 | 17.7808095 | 0 | 0 | 0 | 0 | 0 | 0 | 0 | 0 | 0 | 0 | 1 | 1 | 10 | 8 | 20 |
| ENSMUSG00000021687 | Scamp1 | ENSMUST00000022197 | 27.2963927 | 35 | 16.4373373 | 21.4949795 | 0 | 0 | 0 | 0 | 0 | 0 | 0 | 0 | 0 | 0 | 3 | 1 | 1 | 0 | 5 |
| ENSMUSG00000029426 | Scarb2 | ENSMUST00000031377 | 25.9549795 | 31 | 19.9181446 | 23.0218281 | 0 | 0 | 0 | 0 | 0 | 0 | 0 | 0 | 0 | 0 | 1 | 0 | 2 | 1 | 4 |
| ENSMUSG00000037071 | Scd1 | ENSMUST00000041331 | 25.922722 | 31 | 21.1649366 | 22.3760092 | 0 | 0 | 0 | 0 | 0 | 0 | 0 | 0 | 0 | 0 | 1 | 3 | 1 | 0 | 5 |
| ENSMUSG00000062110 | Scfd2 | ENSMUST00000151474 | 29.4247788 | 33 | 18.7610619 | 18.3628319 | 0 | 0 | 0 | 0 | 0 | 0 | 0 | 0 | 0 | 0 | 0 | 2 | 1 | 0 | 3 |
| ENSMUSG00000050711 | Scg2 | ENSMUST00000049972 | 27.3858921 | 40 | 14.1078838 | 18.6721992 | 0 | 0 | 0 | 0 | 0 | 0 | 0 | 0 | 0 | 0 | 0 | 1 | 0 | 0 | 1 |
| ENSMUSG00000027777 | Schip1 | ENSMUST00000182532 | 26.6452648 | 38 | 17.9775281 | 17.6565008 | 0 | 0 | 0 | 0 | 0 | 0 | 0 | 0 | 0 | 0 | 0 | 0 | 8 | 0 | 8 |
| ENSMUSG00000057182 | Scn3a | ENSMUST00000066432 | 32.18959 | 34 | 17.7377145 | 15.6731608 | 0 | 0 | 0 | 1 | 0 | 0 | 0 | 0 | 1 | 2 | 6 | 0 | 9 | 2 | 19 |
| ENSMUSG00000000278 | Scpep1 | ENSMUST00000000287 | 29.2834891 | 26 | 24.2990654 | 20.8722741 | 0 | 0 | 0 | 0 | 0 | 0 | 0 | 0 | 0 | 0 | 0 | 1 | 0 | 0 | 1 |
| ENSMUSG00000008226 | Scrn3 | ENSMUST00000090811 | 27.2070212 | 34 | 18.5338152 | 19.9793495 | 1 | 0 | 0 | 0 | 0 | 0 | 0 | 0 | 0 | 0 | 0 | 2 | 0 | 0 | 3 |
| ENSMUSG00000022261 | Sdc2 | ENSMUST00000022871 | 30 | 33 | 17.716895 | 19.0410959 | 0 | 1 | 0 | 0 | 0 | 0 | 0 | 0 | 0 | 0 | 0 | 0 | 1 | 0 | 2 |
| ENSMUSG00000028249 | Sdcbp | ENSMUST00000029912 | 27.2093023 | 35 | 18.6046512 | 18.7209302 | 0 | 0 | 0 | 0 | 0 | 0 | 0 | 0 | 0 | 0 | 0 | 1 | 0 | 0 | 1 |
| ENSMUSG00000038806 | Sde2 | ENSMUST00000038091 | 23.4615385 | 38 | 19.1758242 | 19.8351648 | 0 | 0 | 0 | 0 | 0 | 0 | 0 | 0 | 0 | 0 | 0 | 0 | 2 | 0 | 2 |
| ENSMUSG00000042505 | Sdhaf3 | ENSMUST00000040826 | 32.2794493 | 29 | 19.836818 | 18.8169301 | 0 | 0 | 1 | 0 | 0 | 0 | 0 | 0 | 0 | 1 | 0 | 4 | 0 | 0 | 5 |
| ENSMUSG00000026154 | Sdhaf4 | ENSMUST00000027338 | 27.2435897 | 36 | 18.9102564 | 17.9487179 | 0 | 1 | 0 | 0 | 0 | 0 | 0 | 0 | 0 | 0 | 0 | 0 | 0 | 0 | 1 |
| ENSMUSG00000058076 | Sdhc | ENSMUST00000111336 | 24.8482549 | 27 | 22.875569 | 24.8482549 | 3 | 1 | 0 | 3 | 0 | 0 | 0 | 1 | 2 | 6 | 0 | 1 | 0 | 1 | 12 |
| ENSMUSG00000000171 | Sdhd | ENSMUST00000000175 | 27.1753681 | 30 | 19.0093708 | 23.6947791 | 0 | 0 | 1 | 0 | 0 | 0 | 0 | 0 | 0 | 1 | 0 | 1 | 0 | 0 | 2 |
| ENSMUSG00000020823 | Sec14l1 | ENSMUST00000090433 | 20.2370872 | 25 | 28.5351397 | 25.8679086 | 0 | 0 | 0 | 0 | 0 | 0 | 0 | 0 | 0 | 0 | 0 | 5 | 0 | 0 | 5 |
| ENSMUSG00000027879 | Sec22b | ENSMUST00000029476 | 25.9222333 | 34 | 20.0897308 | 19.5912263 | 0 | 0 | 0 | 1 | 0 | 0 | 0 | 0 | 1 | 2 | 3 | 1 | 20 | 0 | 26 |
| ENSMUSG00000020986 | Sec23a | ENSMUST00000021375 | 28.1288724 | 38 | 15.6133829 | 18.0916976 | 0 | 0 | 0 | 0 | 0 | 0 | 0 | 0 | 0 | 0 | 0 | 0 | 6 | 0 | 6 |
| ENSMUSG00000036391 | Sec24a | ENSMUST00000109097 | 29.5494237 | 37 | 16.3115613 | 17.3244848 | 0 | 0 | 0 | 0 | 0 | 0 | 0 | 0 | 0 | 0 | 0 | 1 | 8 | 3 | 12 |
| ENSMUSG00000025816 | Sec61a2 | ENSMUST00000102981 | 25.7075472 | 33 | 22.6415094 | 18.8679245 | 0 | 0 | 0 | 0 | 0 | 0 | 0 | 0 | 0 | 0 | 0 | 1 | 0 | 0 | 1 |
| ENSMUSG00000027706 | Sec62 | ENSMUST00000029256 | 29.1684806 | 35 | 16.8480627 | 19.06835 | 1 | 0 | 0 | 0 | 0 | 0 | 0 | 0 | 0 | 0 | 0 | 0 | 2 | 0 | 3 |
| ENSMUSG00000019802 | Sec63 | ENSMUST00000019937 | 26.2931034 | 33 | 19.9425287 | 21.2068966 | 0 | 0 | 0 | 2 | 0 | 0 | 0 | 0 | 2 | 4 | 3 | 0 | 3 | 0 | 10 |
| ENSMUSG00000079614 | Seh1l | ENSMUST00000025421 | 26.446281 | 36 | 19.0517616 | 18.1818182 | 0 | 0 | 0 | 0 | 0 | 0 | 0 | 0 | 0 | 0 | 0 | 3 | 1 | 1 | 5 |
| ENSMUSG00000042682 | Selk | ENSMUST00000112268 | 27.0588235 | 37 | 17.3109244 | 18.487395 | 0 | 0 | 0 | 0 | 0 | 0 | 0 | 0 | 0 | 0 | 0 | 0 | 4 | 0 | 4 |
| ENSMUSG00000075700 | Selt | ENSMUST00000107924 | 30.4294976 | 33 | 16.0858995 | 20.2593193 | 0 | 0 | 0 | 0 | 0 | 0 | 0 | 0 | 0 | 0 | 0 | 0 | 0 | 3 | 3 |
| ENSMUSG00000028883 | Sema3a | ENSMUST00000030714 | 33.4908845 | 31 | 17.6569885 | 17.4544227 | 0 | 0 | 0 | 0 | 0 | 0 | 0 | 0 | 0 | 0 | 1 | 0 | 0 | 0 | 1 |
| ENSMUSG00000028780 | Sema3c | ENSMUST00000030568 | 33.3996815 | 33 | 15.7245223 | 17.4761146 | 1 | 0 | 1 | 1 | 0 | 0 | 0 | 0 | 1 | 3 | 0 | 0 | 1 | 0 | 5 |
| ENSMUSG00000040254 | Sema3d | ENSMUST00000030868 | 33.3790455 | 34 | 16.1272628 | 16.7580911 | 0 | 0 | 0 | 0 | 0 | 0 | 0 | 0 | 0 | 0 | 1 | 0 | 1 | 1 | 3 |
| ENSMUSG00000022231 | Sema5a | ENSMUST00000067458 | 28.3857927 | 31 | 21.7152758 | 18.8276061 | 0 | 0 | 0 | 0 | 0 | 0 | 0 | 0 | 0 | 0 | 0 | 4 | 0 | 0 | 4 |
| ENSMUSG00000027200 | Sema6d | ENSMUST00000103239 | 28.2405526 | 30 | 20.2356766 | 21.170256 | 0 | 0 | 0 | 0 | 0 | 0 | 0 | 0 | 0 | 0 | 0 | 0 | 3 | 0 | 3 |
| ENSMUSG00000052917 | Senp7 | ENSMUST00000202000 | 23.2150679 | 34 | 22.9522558 | 20.2365309 | 0 | 0 | 0 | 0 | 0 | 0 | 0 | 0 | 0 | 0 | 0 | 4 | 1 | 0 | 5 |
| ENSMUSG00000051705 | Senp8 | ENSMUST00000051039 | 30.8459422 | 32 | 18.2943604 | 18.8101788 | 0 | 0 | 0 | 0 | 0 | 0 | 0 | 0 | 0 | 0 | 0 | 0 | 2 | 1 | 3 |
| ENSMUSG00000058013 | Sep11 | ENSMUST00000074733 | 23.4739179 | 30 | 24.1120977 | 21.9478357 | 0 | 0 | 0 | 0 | 0 | 0 | 0 | 0 | 0 | 0 | 0 | 0 | 1 | 4 | 5 |
| ENSMUSG00000029173 | Sepsecs | ENSMUST00000150709 | 27.5311925 | 30 | 20.0040908 | 22.9290243 | 0 | 0 | 0 | 0 | 0 | 0 | 0 | 0 | 0 | 0 | 2 | 1 | 2 | 0 | 5 |
| ENSMUSG00000019917 | Sept10 | ENSMUST00000165971 | 29.5343137 | 39 | 15.8088235 | 15.9313725 | 0 | 0 | 0 | 0 | 0 | 0 | 0 | 0 | 0 | 0 | 0 | 0 | 7 | 1 | 8 |
| ENSMUSG00000037072 | Sept15 | ENSMUST00000106211 | 32.9487179 | 30 | 18.4615385 | 18.3333333 | 0 | 0 | 0 | 0 | 0 | 0 | 0 | 0 | 0 | 0 | 0 | 0 | 0 | 2 | 2 |
| ENSMUSG00000026276 | Sept2 | ENSMUST00000027495 | 28.3320343 | 35 | 17.4201091 | 19.6414653 | 0 | 0 | 0 | 1 | 0 | 0 | 0 | 0 | 1 | 2 | 1 | 1 | 12 | 0 | 16 |
| ENSMUSG00000001833 | Sept7 | ENSMUST00000115272 | 30.6182532 | 38 | 15.8979392 | 15.8979392 | 0 | 0 | 0 | 0 | 0 | 0 | 0 | 0 | 0 | 0 | 4 | 0 | 5 | 1 | 10 |
| ENSMUSG00000036371 | Serbp1 | ENSMUST00000042990 | 29.0847064 | 33 | 16.9703128 | 21.1149016 | 2 | 1 | 0 | 1 | 0 | 0 | 0 | 0 | 1 | 2 | 2 | 3 | 17 | 4 | 31 |
| ENSMUSG00000074884 | Serf2 | ENSMUST00000139253 | 23.4446131 | 29 | 22.9514416 | 24.4688923 | 0 | 0 | 0 | 0 | 0 | 0 | 0 | 0 | 0 | 0 | 0 | 4 | 0 | 0 | 4 |

| | | | | | | | | | | | | | | | | | | | | |
|---|---|---|---|---|---|---|---|---|---|---|---|---|---|---|---|---|---|---|---|---|
| ENSMUSG00000017707 | Serinc3 | ENSMUST00000017851 | 30.0138313 | 32 | 15.7215307 | 21.8072845 | 0 | 0 | 0 | 0 | 0 | 0 | 0 | 0 | 0 | 0 | 2 | 0 | 3 | 0 | 5 |
| ENSMUSG00000021703 | Serinc5 | ENSMUST00000049488 | 26.0047281 | 29 | 21.6443394 | 23.1941161 | 0 | 0 | 0 | 0 | 0 | 0 | 0 | 0 | 0 | 0 | 1 | 0 | 0 | 0 | 1 |
| ENSMUSG00000026715 | Serpinc1 | ENSMUST00000064725 | 29.3235294 | 27 | 22.2647059 | 21.8823529 | 0 | 0 | 0 | 0 | 0 | 0 | 0 | 0 | 0 | 0 | 1 | 2 | 4 | 0 | 7 |
| ENSMUSG00000037411 | Serpine1 | ENSMUST00000041388 | 24.9101796 | 27 | 24.7904192 | 23.7724551 | 0 | 0 | 0 | 0 | 0 | 0 | 0 | 0 | 0 | 0 | 1 | 9 | 0 | 0 | 10 |
| ENSMUSG00000027834 | Serpini1 | ENSMUST00000029423 | 31.4139942 | 35 | 16.180758 | 17.3833819 | 1 | 0 | 0 | 3 | 0 | 0 | 0 | 0 | 3 | 6 | 0 | 24 | 1 | 1 | 33 |
| ENSMUSG00000049800 | Sertad2 | ENSMUST00000093292 | 26.6915539 | 30 | 21.6518899 | 21.9085394 | 3 | 1 | 0 | 4 | 0 | 0 | 0 | 1 | 3 | 8 | 0 | 0 | 1 | 1 | 14 |
| ENSMUSG00000055200 | Sertad3 | ENSMUST00000068641 | 19.2675159 | 32 | 21.1783439 | 27.7070064 | 0 | 0 | 0 | 0 | 0 | 0 | 0 | 0 | 0 | 0 | 2 | 1 | 3 | 0 | 6 |
| ENSMUSG00000032009 | Sesn3 | ENSMUST00000034507 | 22.9166667 | 39 | 17.1296296 | 20.6018519 | 0 | 0 | 0 | 0 | 0 | 0 | 0 | 0 | 0 | 0 | 0 | 0 | 5 | 0 | 5 |
| ENSMUSG00000042272 | Sestd1 | ENSMUST00000102660 | 27.5217199 | 32 | 20.1443086 | 20.1443086 | 0 | 0 | 0 | 1 | 0 | 0 | 0 | 0 | 1 | 2 | 1 | 0 | 3 | 0 | 6 |
| ENSMUSG00000056770 | Setd3 | ENSMUST00000147466 | 23.7665463 | 31 | 23.8267148 | 21.3598075 | 1 | 0 | 0 | 1 | 0 | 0 | 0 | 0 | 1 | 2 | 0 | 1 | 0 | 0 | 4 |
| ENSMUSG00000037111 | Setd7 | ENSMUST00000037141 | 25.8517034 | 29 | 23.3800935 | 21.760187 | 0 | 0 | 0 | 0 | 0 | 0 | 0 | 0 | 0 | 0 | 1 | 51 | 5 | 0 | 57 |
| ENSMUSG00000049327 | Setd8 | ENSMUST00000199798 | 29.5162278 | 21 | 25.7807716 | 23.2088181 | 0 | 0 | 0 | 0 | 0 | 0 | 0 | 0 | 0 | 0 | 0 | 1 | 0 | 0 | 1 |
| ENSMUSG00000002129 | Sf3a1 | ENSMUST00000002198 | 24.3557772 | 31 | 20.3657523 | 24.39734 | 0 | 0 | 0 | 0 | 0 | 0 | 0 | 0 | 0 | 0 | 0 | 0 | 4 | 0 | 4 |
| ENSMUSG00000025982 | Sf3b1 | ENSMUST00000027127 | 27.8060046 | 36 | 16.1662818 | 20.5080831 | 0 | 0 | 0 | 1 | 0 | 0 | 0 | 0 | 1 | 2 | 0 | 0 | 3 | 1 | 6 |
| ENSMUSG00000028820 | Sfpq | ENSMUST00000030623 | 28.3805031 | 36 | 15.0157233 | 20.2044025 | 0 | 0 | 0 | 0 | 0 | 0 | 0 | 0 | 0 | 0 | 0 | 0 | 0 | 1 | 1 |
| ENSMUSG00000027996 | Sfrp2 | ENSMUST00000029625 | 25.2796421 | 34 | 22.4832215 | 17.8970917 | 0 | 0 | 0 | 0 | 0 | 0 | 0 | 0 | 0 | 0 | 0 | 1 | 6 | 0 | 7 |
| ENSMUSG00000040848 | Sft2d2 | ENSMUST00000043338 | 23.1144542 | 30 | 22.5858914 | 24.5375076 | 0 | 0 | 0 | 0 | 0 | 0 | 0 | 0 | 0 | 0 | 0 | 1 | 0 | 2 | 3 |
| ENSMUSG00000029156 | Sgcb | ENSMUST00000081170 | 30.1529497 | 30 | 18.8638019 | 21.0123816 | 0 | 0 | 0 | 0 | 0 | 0 | 0 | 0 | 0 | 0 | 2 | 0 | 1 | 0 | 3 |
| ENSMUSG00000020354 | Sgcd | ENSMUST00000109220 | 32.6900585 | 31 | 17.1929825 | 18.7134503 | 0 | 0 | 0 | 0 | 0 | 0 | 0 | 0 | 0 | 0 | 0 | 0 | 2 | 0 | 2 |
| ENSMUSG00000028524 | Sgip1 | ENSMUST00000106882 | 30.9664694 | 31 | 18.4559031 | 19.4984503 | 1 | 1 | 0 | 2 | 0 | 0 | 0 | 0 | 2 | 4 | 0 | 1 | 0 | 0 | 7 |
| ENSMUSG00000019970 | Sgk1 | ENSMUST00000020145 | 23.6988848 | 35 | 19.4237918 | 22.2118959 | 0 | 0 | 0 | 0 | 0 | 0 | 0 | 0 | 0 | 0 | 3 | 0 | 5 | 0 | 8 |
| ENSMUSG00000050931 | Sgms2 | ENSMUST00000090246 | 25.7399321 | 32 | 21.4701601 | 21.2518195 | 1 | 0 | 0 | 1 | 0 | 0 | 0 | 0 | 1 | 2 | 6 | 3 | 11 | 3 | 26 |
| ENSMUSG00000020097 | Sgpl1 | ENSMUST00000150258 | 22.4706745 | 30 | 24.4501466 | 23.4970674 | 0 | 0 | 0 | 0 | 0 | 0 | 0 | 0 | 0 | 0 | 0 | 1 | 1 | 1 | 3 |
| ENSMUSG00000021054 | Sgpp1 | ENSMUST00000021450 | 29.2761394 | 35 | 16.2466488 | 19.5174263 | 0 | 0 | 0 | 0 | 0 | 0 | 0 | 0 | 0 | 0 | 1 | 0 | 1 | 0 | 2 |
| ENSMUSG00000005696 | Sh2d1a | ENSMUST00000189753 | 34.0909091 | 32 | 16.7207792 | 17.2077922 | 0 | 0 | 0 | 1 | 0 | 0 | 0 | 0 | 1 | 2 | 0 | 0 | 0 | 0 | 2 |
| ENSMUSG00000031246 | Sh3bgrl | ENSMUST00000033598 | 32.8280641 | 34 | 15.1580771 | 18.0597661 | 0 | 0 | 0 | 0 | 0 | 0 | 0 | 0 | 0 | 0 | 0 | 0 | 0 | 1 | 1 |
| ENSMUSG00000032261 | Sh3bgrl2 | ENSMUST00000113215 | 25.863747 | 30 | 21.8978102 | 22.43309 | 0 | 0 | 0 | 0 | 0 | 0 | 0 | 0 | 0 | 0 | 0 | 5 | 8 | 0 | 13 |
| ENSMUSG00000037062 | Sh3glb1 | ENSMUST00000198254 | 29.0429774 | 30 | 20.4253434 | 20.3810368 | 2 | 1 | 0 | 3 | 0 | 0 | 0 | 1 | 2 | 6 | 0 | 1 | 1 | 0 | 11 |
| ENSMUSG00000042708 | Shcbp1l | ENSMUST00000136614 | 24.4274809 | 31 | 22.9961832 | 21.6603053 | 1 | 0 | 0 | 1 | 0 | 0 | 0 | 0 | 1 | 2 | 0 | 1 | 3 | 0 | 7 |
| ENSMUSG00000042541 | Shfm1 | ENSMUST00000041111 | 31.6883117 | 36 | 15.7575758 | 16.5367965 | 0 | 0 | 1 | 0 | 0 | 0 | 0 | 0 | 0 | 1 | 0 | 0 | 0 | 0 | 1 |
| ENSMUSG00000050010 | Shisa3 | ENSMUST00000087241 | 30.7508939 | 29 | 19.4278903 | 20.7787048 | 1 | 1 | 1 | 1 | 0 | 0 | 0 | 0 | 1 | 3 | 0 | 1 | 0 | 0 | 6 |
| ENSMUSG00000022494 | Shisa9 | ENSMUST00000023138 | 28.5575992 | 29 | 22.8783479 | 19.8451113 | 0 | 0 | 0 | 0 | 0 | 0 | 0 | 0 | 0 | 0 | 1 | 0 | 1 | 0 | 2 |
| ENSMUSG00000090112 | Shprh | ENSMUST00000044053 | 29.8691681 | 35 | 17.649963 | 17.1068872 | 0 | 0 | 1 | 0 | 0 | 0 | 0 | 0 | 0 | 1 | 0 | 1 | 0 | 0 | 2 |
| ENSMUSG00000045180 | Shroom2 | ENSMUST00000101141 | 28.5856244 | 32 | 19.7747599 | 19.2779066 | 0 | 1 | 1 | 0 | 0 | 0 | 0 | 0 | 0 | 1 | 2 | 2 | 2 | 0 | 8 |
| ENSMUSG00000041362 | Shtn1 | ENSMUST00000163821 | 27.6237085 | 29 | 19.2495922 | 23.8716694 | 0 | 0 | 0 | 0 | 0 | 0 | 0 | 0 | 0 | 0 | 0 | 0 | 4 | 1 | 5 |
| ENSMUSG00000036840 | Siah1a | ENSMUST00000045296 | 28.7992495 | 38 | 14.6341463 | 18.9493433 | 0 | 0 | 0 | 2 | 0 | 0 | 0 | 0 | 2 | 4 | 0 | 0 | 3 | 2 | 9 |
| ENSMUSG00000040749 | Siah1b | ENSMUST00000037928 | 29.1115312 | 35 | 15.879017 | 20.0378072 | 0 | 0 | 0 | 0 | 0 | 0 | 0 | 0 | 0 | 0 | 0 | 0 | 1 | 0 | 1 |
| ENSMUSG00000027854 | Sike1 | ENSMUST00000029447 | 29.1729323 | 28 | 21.6240602 | 21.1278195 | 1 | 0 | 0 | 0 | 0 | 0 | 0 | 0 | 0 | 0 | 0 | 2 | 0 | 0 | 3 |
| ENSMUSG00000019913 | Sim1 | ENSMUST00000020071 | 33.9222615 | 32 | 16.1712742 | 17.8549158 | 2 | 1 | 0 | 4 | 0 | 0 | 0 | 1 | 3 | 8 | 0 | 2 | 0 | 0 | 13 |
| ENSMUSG00000020063 | Sirt1 | ENSMUST00000146028 | 31.0708402 | 32 | 17.4629325 | 19.4728171 | 0 | 0 | 0 | 0 | 0 | 0 | 0 | 0 | 0 | 0 | 1 | 0 | 0 | 0 | 1 |
| ENSMUSG00000034460 | Six4 | ENSMUST00000175693 | 27.7369439 | 31 | 20.4255319 | 20.3481625 | 0 | 0 | 0 | 0 | 0 | 0 | 0 | 0 | 0 | 0 | 8 | 2 | 15 | 3 | 28 |
| ENSMUSG00000020492 | Ska2 | ENSMUST00000020794 | 25.9484067 | 33 | 19.1578149 | 21.4339909 | 7 | 2 | 0 | 4 | 0 | 0 | 1 | 2 | 1 | 8 | 1 | 1 | 2 | 0 | 21 |
| ENSMUSG00000021965 | Ska3 | ENSMUST00000022536 | 27.4409044 | 29 | 20.6577595 | 22.4049332 | 0 | 0 | 0 | 0 | 0 | 0 | 0 | 0 | 0 | 0 | 0 | 2 | 0 | 0 | 2 |
| ENSMUSG00000059182 | Skap2 | ENSMUST00000204778 | 27.1673621 | 32 | 19.0078813 | 21.7431618 | 0 | 0 | 1 | 1 | 0 | 0 | 0 | 0 | 1 | 3 | 0 | 1 | 1 | 0 | 5 |
| ENSMUSG00000027660 | Skil | ENSMUST00000029194 | 27.2452069 | 34 | 17.3814329 | 20.938446 | 1 | 0 | 0 | 0 | 0 | 0 | 0 | 0 | 0 | 0 | 1 | 0 | 13 | 1 | 16 |
| ENSMUSG00000036309 | Skp1a | ENSMUST00000037324 | 27.2076372 | 36 | 17.8997613 | 19.2124105 | 0 | 0 | 0 | 0 | 0 | 0 | 0 | 0 | 0 | 0 | 0 | 1 | 4 | 1 | 6 |
| ENSMUSG00000054115 | Skp2 | ENSMUST00000110585 | 28.7766831 | 29 | 21.0796388 | 21.2438424 | 0 | 0 | 0 | 0 | 0 | 0 | 0 | 0 | 0 | 0 | 2 | 1 | 0 | 0 | 3 |
| ENSMUSG00000031684 | Slc10a7 | ENSMUST00000034111 | 26.0245036 | 29 | 21.7997465 | 22.7714406 | 0 | 0 | 0 | 0 | 0 | 0 | 0 | 0 | 0 | 0 | 1 | 0 | 0 | 0 | 1 |
| ENSMUSG00000024597 | Slc12a2 | ENSMUST00000115366 | 29.2057762 | 34 | 16.4620939 | 19.9277978 | 0 | 0 | 0 | 0 | 0 | 0 | 0 | 0 | 0 | 0 | 0 | 1 | 0 | 1 | 2 |
| ENSMUSG00000032902 | Slc16a1 | ENSMUST00000046212 | 27.2992701 | 34 | 18.2846715 | 20.1824818 | 1 | 0 | 0 | 0 | 0 | 0 | 0 | 0 | 0 | 0 | 3 | 1 | 8 | 1 | 14 |
| ENSMUSG00000019838 | Slc16a10 | ENSMUST00000092566 | 26.0654385 | 31 | 21.0613143 | 21.6387132 | 5 | 3 | 0 | 6 | 0 | 0 | 0 | 2 | 4 | 12 | 0 | 9 | 0 | 0 | 29 |
| ENSMUSG00000027896 | Slc16a4 | ENSMUST00000029502 | 37.3106061 | 34 | 15.1515152 | 13.6363636 | 0 | 0 | 0 | 1 | 0 | 0 | 0 | 0 | 1 | 2 | 0 | 0 | 0 | 0 | 2 |
| ENSMUSG00000020102 | Slc16a7 | ENSMUST00000063318 | 33.2041344 | 30 | 18.8630491 | 18.2816537 | 0 | 0 | 0 | 1 | 0 | 0 | 0 | 0 | 1 | 2 | 0 | 0 | 0 | 0 | 2 |
| ENSMUSG00000037455 | Slc18b1 | ENSMUST00000119597 | 23.6238532 | 33 | 24.7706422 | 18.7308869 | 0 | 0 | 0 | 0 | 0 | 0 | 0 | 0 | 0 | 0 | 1 | 0 | 0 | 0 | 1 |
| ENSMUSG00000040918 | Slc19a2 | ENSMUST00000044021 | 26.7273701 | 32 | 20.1928227 | 20.8891269 | 0 | 0 | 0 | 0 | 0 | 0 | 0 | 0 | 0 | 0 | 0 | 1 | 1 | 2 | 4 |
| ENSMUSG00000024935 | Slc1a1 | ENSMUST00000025875 | 27.4271845 | 30 | 21.8446602 | 21.1165049 | 0 | 0 | 0 | 0 | 0 | 0 | 0 | 0 | 0 | 0 | 0 | 1 | 0 | 0 | 1 |
| ENSMUSG00000005089 | Slc1a2 | ENSMUST00000080210 | 28.1513514 | 29 | 21.4810811 | 20.9297297 | 0 | 0 | 0 | 1 | 0 | 0 | 0 | 0 | 1 | 2 | 0 | 5 | 7 | 1 | 15 |
| ENSMUSG00000024354 | Slc23a1 | ENSMUST00000150877 | 20.323741 | 26 | 28.177458 | 25.6594724 | 0 | 0 | 0 | 0 | 0 | 0 | 0 | 0 | 0 | 0 | 0 | 1 | 0 | 0 | 1 |
| ENSMUSG00000027340 | Slc23a2 | ENSMUST00000028815 | 22.8372873 | 31 | 22.6935059 | 23.2207045 | 0 | 0 | 0 | 0 | 0 | 0 | 0 | 0 | 0 | 0 | 0 | 5 | 1 | 0 | 6 |
| ENSMUSG00000027010 | Slc25a12 | ENSMUST00000151937 | 24.7800587 | 33 | 21.2365591 | 21.2609971 | 1 | 0 | 0 | 0 | 0 | 0 | 0 | 0 | 0 | 0 | 2 | 8 | 1 | 0 | 12 |
| ENSMUSG00000022404 | Slc25a17 | ENSMUST00000023040 | 25.9887006 | 35 | 18.7853107 | 20.3389831 | 0 | 0 | 0 | 0 | 0 | 0 | 0 | 0 | 0 | 0 | 2 | 0 | 0 | 0 | 2 |
| ENSMUSG00000019082 | Slc25a22 | ENSMUST00000124266 | 18.503937 | 25 | 30.0688976 | 26.7716535 | 1 | 0 | 0 | 0 | 0 | 0 | 0 | 0 | 0 | 0 | 0 | 2 | 0 | 0 | 3 |
| ENSMUSG00000040322 | Slc25a24 | ENSMUST00000029477 | 29.32717 | 34 | 18.1818182 | 18.2331793 | 1 | 0 | 0 | 0 | 0 | 0 | 0 | 0 | 0 | 0 | 0 | 1 | 0 | 1 | 3 |
| ENSMUSG00000023912 | Slc25a27 | ENSMUST00000024705 | 26.2170088 | 26 | 24.2228739 | 23.2258065 | 0 | 0 | 1 | 0 | 0 | 0 | 0 | 0 | 0 | 1 | 0 | 0 | 0 | 0 | 1 |
| ENSMUSG00000022299 | Slc25a32 | ENSMUST00000022908 | 31.3659359 | 35 | 15.9640247 | 18.0438449 | 0 | 0 | 0 | 0 | 0 | 0 | 0 | 0 | 0 | 0 | 1 | 0 | 0 | 0 | 1 |
| ENSMUSG00000032449 | Slc25a36 | ENSMUST00000085206 | 28.8127295 | 33 | 18.7025704 | 19.3145655 | 1 | 0 | 0 | 0 | 0 | 0 | 0 | 0 | 0 | 0 | 0 | 1 | 6 | 0 | 8 |
| ENSMUSG00000054099 | Slc25a40 | ENSMUST00000196727 | 30.5168539 | 33 | 16.7640449 | 19.5955056 | 3 | 0 | 1 | 1 | 0 | 0 | 0 | 0 | 1 | 3 | 1 | 0 | 0 | 0 | 7 |
| ENSMUSG00000024259 | Slc25a46 | ENSMUST00000060396 | 29.6063513 | 34 | 16.0436652 | 20.3771088 | 2 | 0 | 0 | 3 | 0 | 0 | 0 | 0 | 3 | 6 | 0 | 0 | 0 | 3 | 11 |
| ENSMUSG00000045973 | Slc25a51 | ENSMUST00000107796 | 23.4177215 | 31 | 21.7902351 | 23.7492465 | 0 | 0 | 0 | 0 | 0 | 0 | 0 | 0 | 0 | 0 | 5 | 9 | 8 | 1 | 23 |
| ENSMUSG00000039908 | Slc26a11 | ENSMUST00000050880 | 28.4450063 | 29 | 18.9633375 | 23.8938053 | 2 | 2 | 1 | 0 | 0 | 0 | 0 | 0 | 0 | 1 | 0 | 0 | 0 | 0 | 5 |
| ENSMUSG00000037490 | Slc2a12 | ENSMUST00000042261 | 27.4881517 | 29 | 23.1279621 | 20.3317536 | 1 | 1 | 0 | 2 | 0 | 0 | 0 | 0 | 2 | 4 | 0 | 0 | 0 | 1 | 7 |
| ENSMUSG00000036298 | Slc2a13 | ENSMUST00000109283 | 29.6096904 | 32 | 20.2781516 | 18.5284881 | 1 | 1 | 0 | 2 | 0 | 0 | 0 | 1 | 1 | 4 | 0 | 0 | 0 | 0 | 6 |
| ENSMUSG00000003153 | Slc2a3 | ENSMUST00000032476 | 23.5626767 | 30 | 22.3374175 | 24.3166824 | 2 | 0 | 1 | 2 | 0 | 0 | 0 | 0 | 2 | 5 | 1 | 0 | 0 | 1 | 9 |
| ENSMUSG00000037434 | Slc30a1 | ENSMUST00000044954 | 27.8636884 | 33 | 16.9243986 | 22.0790378 | 1 | 0 | 0 | 0 | 0 | 0 | 0 | 0 | 0 | 0 | 1 | 1 | 4 | 4 | 11 |
| ENSMUSG00000026614 | Slc30a10 | ENSMUST00000061093 | 25.9805087 | 31 | 21.3453768 | 21.9871643 | 0 | 0 | 0 | 0 | 0 | 0 | 0 | 0 | 0 | 0 | 0 | 2 | 1 | 0 | 3 |
| ENSMUSG00000054414 | Slc30a7 | ENSMUST00000067485 | 26.2774672 | 31 | 21.3496294 | 20.9725653 | 2 | 0 | 0 | 0 | 0 | 0 | 0 | 0 | 0 | 0 | 0 | 0 | 2 | 4 | 8 |
| ENSMUSG00000027822 | Slc33a1 | ENSMUST00000029402 | 33.1882481 | 37 | 12.9488575 | 16.8661589 | 0 | 0 | 0 | 1 | 0 | 0 | 0 | 0 | 1 | 2 | 0 | 3 | 0 | 1 | 6 |
| ENSMUSG00000031156 | Slc35a2 | ENSMUST00000115660 | 21.7932752 | 29 | 23.2254047 | 26.0273973 | 1 | 1 | 0 | 0 | 0 | 0 | 0 | 0 | 0 | 0 | 0 | 0 | 3 | 0 | 5 |
| ENSMUSG00000027957 | Slc35a3 | ENSMUST00000029569 | 28.956229 | 36 | 16.3059163 | 18.5906686 | 0 | 0 | 0 | 0 | 0 | 0 | 0 | 0 | 0 | 0 | 2 | 9 | 5 | 0 | 16 |
| ENSMUSG00000022664 | Slc35a5 | ENSMUST00000023344 | 29.8756799 | 34 | 17.5213675 | 18.2595183 | 0 | 0 | 0 | 0 | 0 | 0 | 0 | 0 | 0 | 0 | 4 | 0 | 4 | 2 | 10 |
| ENSMUSG00000021432 | Slc35b3 | ENSMUST00000021870 | 31.629393 | 32 | 16.1341853 | 19.8083067 | 0 | 0 | 1 | 0 | 0 | 0 | 0 | 0 | 0 | 1 | 0 | 0 | 0 | 0 | 1 |
| ENSMUSG00000050473 | Slc35d3 | ENSMUST00000059805 | 30.227519 | 34 | 17.1180932 | 19.0682557 | 1 | 0 | 2 | 1 | 0 | 0 | 0 | 0 | 1 | 4 | 0 | 0 | 0 | 1 | 6 |
| ENSMUSG00000042202 | Slc35e2 | ENSMUST00000105608 | 25.6371161 | 32 | 21.0411675 | 21.6946199 | 0 | 0 | 0 | 1 | 0 | 0 | 0 | 0 | 1 | 2 | 4 | 2 | 25 | 0 | 33 |
| ENSMUSG00000060181 | Slc35e3 | ENSMUST00000079041 | 25.8445946 | 28 | 21.1993243 | 24.9577703 | 0 | 0 | 0 | 0 | 0 | 0 | 0 | 0 | 0 | 0 | 0 | 0 | 4 | 1 | 5 |
| ENSMUSG00000038602 | Slc35f1 | ENSMUST00000105473 | 30.7384615 | 31 | 19.0153846 | 18.8923077 | 1 | 0 | 0 | 0 | 0 | 0 | 0 | 0 | 0 | 0 | 5 | 0 | 5 | 0 | 11 |
| ENSMUSG00000021852 | Slc35f4 | ENSMUST00000123534 | 32.5564457 | 29 | 18.0626366 | 20.1019665 | 0 | 0 | 0 | 0 | 0 | 0 | 0 | 0 | 0 | 0 | 0 | 1 | 0 | 0 | 1 |
| ENSMUSG00000026342 | Slc35f5 | ENSMUST00000027580 | 29.7093649 | 32 | 17.007535 | 21.31324 | 0 | 0 | 0 | 0 | 0 | 0 | 0 | 0 | 0 | 0 | 0 | 0 | 1 | 0 | 1 |
| ENSMUSG00000043885 | Slc36a4 | ENSMUST00000061568 | 27.6585122 | 30 | 19.2960559 | 22.6160759 | 1 | 0 | 0 | 2 | 0 | 0 | 0 | 0 | 2 | 4 | 0 | 2 | 0 | 0 | 7 |
| ENSMUSG00000022462 | Slc38a2 | ENSMUST00000023099 | 28.503476 | 32 | 19.4657885 | 20.2341749 | 0 | 0 | 0 | 0 | 0 | 0 | 0 | 0 | 0 | 0 | 1 | 1 | 6 | 2 | 10 |
| ENSMUSG00000047789 | Slc38a9 | ENSMUST00000052514 | 25.9782455 | 29 | 21.8392428 | 22.9269671 | 0 | 0 | 0 | 0 | 0 | 0 | 0 | 0 | 0 | 0 | 0 | 2 | 15 | 1 | 18 |
| ENSMUSG00000025986 | Slc39a10 | ENSMUST00000027131 | 27.4802823 | 35 | 18.7629722 | 18.9290162 | 0 | 0 | 0 | 1 | 0 | 0 | 0 | 0 | 1 | 2 | 1 | 0 | 0 | 1 | 4 |
| ENSMUSG00000024270 | Slc39a6 | ENSMUST00000070726 | 28.0651341 | 33 | 16.3793103 | 22.4137931 | 0 | 0 | 0 | 0 | 0 | 0 | 0 | 0 | 0 | 0 | 0 | 2 | 0 | 0 | 2 |

| | | | | | | | | | | | | | | | | | | | | | |
|---|---|---|---|---|---|---|---|---|---|---|---|---|---|---|---|---|---|---|---|---|---|
| ENSMUSG00000048833 | Slc39a9 | ENSMUST00000085245 | 24.559956 | 30 | 22.909791 | 22.4422442 | 0 | 0 | 0 | 0 | 0 | 0 | 0 | 0 | 0 | 0 | 0 | 4 | 0 | 2 | 6 |
| ENSMUSG00000025993 | Slc40a1 | ENSMUST00000027137 | 30.4932735 | 34 | 17.1150972 | 18.6846039 | 0 | 0 | 0 | 0 | 0 | 0 | 0 | 0 | 0 | 0 | 0 | 0 | 0 | 1 | 1 |
| ENSMUSG00000028412 | Slc44a1 | ENSMUST00000107651 | 29.941618 | 34 | 16.8056714 | 19.3911593 | 0 | 0 | 0 | 0 | 0 | 0 | 0 | 0 | 0 | 0 | 0 | 0 | 1 | 0 | 1 |
| ENSMUSG00000028360 | Slc44a5 | ENSMUST00000089948 | 25.5501222 | 33 | 21.1491443 | 20.2322738 | 0 | 0 | 1 | 0 | 0 | 0 | 0 | 0 | 0 | 1 | 0 | 0 | 6 | 0 | 7 |
| ENSMUSG00000026435 | Slc45a3 | ENSMUST00000177943 | 23.2876712 | 28 | 24.4157937 | 24.0128928 | 3 | 2 | 3 | 3 | 0 | 0 | 0 | 0 | 3 | 9 | 0 | 1 | 0 | 0 | 15 |
| ENSMUSG00000026904 | Slc4a10 | ENSMUST00000112480 | 31.9389764 | 35 | 16.7322835 | 16.3877953 | 0 | 0 | 1 | 0 | 0 | 0 | 0 | 0 | 0 | 1 | 0 | 0 | 1 | 0 | 2 |
| ENSMUSG00000021733 | Slc4a7 | ENSMUST00000057015 | 29.4055482 | 37 | 16.1690885 | 17.7278732 | 5 | 3 | 0 | 5 | 0 | 0 | 0 | 1 | 4 | 10 | 1 | 0 | 0 | 0 | 19 |
| ENSMUSG00000089774 | Slc5a3 | ENSMUST00000113975 | 26.9612591 | 34 | 18.1840194 | 21.1622276 | 0 | 0 | 0 | 3 | 0 | 0 | 0 | 0 | 3 | 6 | 0 | 3 | 25 | 0 | 34 |
| ENSMUSG00000023945 | Slc5a7 | ENSMUST00000095712 | 33.0369357 | 32 | 17.6470588 | 16.8604651 | 3 | 2 | 0 | 0 | 0 | 0 | 0 | 0 | 0 | 0 | 0 | 2 | 0 | 0 | 7 |
| ENSMUSG00000030310 | Slc6a1 | ENSMUST00000032454 | 23.5986811 | 28 | 25.7654263 | 22.7508243 | 0 | 0 | 0 | 0 | 0 | 0 | 0 | 0 | 0 | 0 | 0 | 2 | 3 | 0 | 5 |
| ENSMUSG00000019894 | Slc6a15 | ENSMUST00000074204 | 29.7764228 | 34 | 16.2601626 | 19.5121951 | 1 | 0 | 0 | 0 | 0 | 0 | 0 | 0 | 0 | 0 | 0 | 0 | 0 | 0 | 1 |
| ENSMUSG00000041313 | Slc7a1 | ENSMUST00000048116 | 23.2007952 | 27 | 24.9701789 | 24.3936382 | 0 | 0 | 0 | 0 | 0 | 0 | 0 | 0 | 0 | 0 | 0 | 2 | 0 | 0 | 2 |
| ENSMUSG00000027737 | Slc7a11 | ENSMUST00000029297 | 29.0874525 | 32 | 18.6990766 | 20.0298751 | 0 | 0 | 0 | 0 | 0 | 0 | 0 | 0 | 0 | 0 | 0 | 2 | 2 | 0 | 4 |
| ENSMUSG00000069072 | Slc7a14 | ENSMUST00000091259 | 28.2245095 | 28 | 22.2418784 | 21.7915729 | 2 | 1 | 0 | 0 | 0 | 0 | 0 | 0 | 0 | 0 | 0 | 2 | 2 | 2 | 9 |
| ENSMUSG00000031596 | Slc7a2 | ENSMUST00000057784 | 27.8316592 | 31 | 19.8302736 | 21.1811569 | 1 | 0 | 0 | 2 | 0 | 0 | 0 | 0 | 2 | 4 | 8 | 1 | 8 | 2 | 24 |
| ENSMUSG00000031904 | Slc7a6 | ENSMUST00000034378 | 26.2826718 | 29 | 21.1035818 | 23.2333011 | 0 | 0 | 0 | 0 | 0 | 0 | 0 | 0 | 0 | 0 | 1 | 2 | 0 | 0 | 3 |
| ENSMUSG00000033106 | Slc7a6os | ENSMUST00000035925 | 27.3972603 | 27 | 22.9452055 | 22.6027397 | 0 | 0 | 0 | 0 | 0 | 0 | 0 | 0 | 0 | 0 | 0 | 1 | 0 | 0 | 1 |
| ENSMUSG00000060681 | Slc9a6 | ENSMUST00000077741 | 26.8511862 | 34 | 17.2897196 | 21.7109993 | 0 | 0 | 0 | 0 | 0 | 0 | 0 | 0 | 0 | 0 | 0 | 1 | 0 | 0 | 1 |
| ENSMUSG00000037341 | Slc9a7 | ENSMUST00000115393 | 29.8329356 | 32 | 22.1957041 | 16.2291169 | 0 | 0 | 0 | 0 | 0 | 0 | 0 | 0 | 0 | 0 | 0 | 1 | 0 | 0 | 1 |
| ENSMUSG00000037994 | Slc9b2 | ENSMUST00000051849 | 35.2121801 | 26 | 19.6631033 | 19.2743764 | 0 | 0 | 1 | 0 | 0 | 0 | 0 | 0 | 0 | 1 | 1 | 0 | 3 | 4 | 9 |
| ENSMUSG00000030235 | Slco1c1 | ENSMUST00000204998 | 27.3768044 | 29 | 22.6480836 | 21.1050274 | 0 | 0 | 0 | 0 | 0 | 0 | 0 | 0 | 0 | 0 | 0 | 1 | 0 | 1 | 2 |
| ENSMUSG00000040693 | Slco4c1 | ENSMUST00000071985 | 34.4041838 | 31 | 17.743743 | 16.7725065 | 1 | 0 | 1 | 0 | 0 | 0 | 0 | 0 | 0 | 1 | 0 | 0 | 1 | 0 | 3 |
| ENSMUSG00000025938 | Slco5a1 | ENSMUST00000188454 | 27.5427712 | 31 | 18.4433164 | 22.5982328 | 0 | 0 | 0 | 0 | 0 | 0 | 0 | 0 | 0 | 0 | 0 | 1 | 5 | 0 | 6 |
| ENSMUSG00000021597 | Slf1 | ENSMUST00000162944 | 31.1562224 | 38 | 12.886143 | 17.5639894 | 0 | 0 | 0 | 0 | 0 | 0 | 0 | 0 | 0 | 0 | 3 | 0 | 8 | 0 | 11 |
| ENSMUSG00000031558 | Slit2 | ENSMUST00000173107 | 32.9896907 | 32 | 17.3539519 | 17.7262314 | 0 | 0 | 0 | 1 | 0 | 0 | 0 | 0 | 1 | 2 | 0 | 0 | 0 | 0 | 2 |
| ENSMUSG00000036790 | Slitrk2 | ENSMUST00000166241 | 31.7704123 | 36 | 16.8552951 | 15.7437348 | 0 | 0 | 0 | 0 | 0 | 0 | 0 | 0 | 0 | 0 | 0 | 0 | 26 | 1 | 27 |
| ENSMUSG00000046699 | Slitrk4 | ENSMUST00000069926 | 33.8986045 | 35 | 14.8207031 | 15.8099276 | 1 | 0 | 0 | 1 | 0 | 0 | 0 | 0 | 1 | 2 | 2 | 41 | 1 | 3 | 50 |
| ENSMUSG00000045871 | Slitrk6 | ENSMUST00000078386 | 32.7365729 | 34 | 14.4927536 | 18.7553282 | 0 | 0 | 0 | 1 | 0 | 0 | 0 | 0 | 1 | 2 | 0 | 0 | 0 | 1 | 3 |
| ENSMUSG00000025060 | Slk | ENSMUST00000026043 | 27.1170314 | 33 | 18.8392008 | 20.7104345 | 0 | 0 | 0 | 0 | 0 | 0 | 0 | 0 | 0 | 0 | 0 | 1 | 0 | 0 | 1 |
| ENSMUSG00000016257 | Slmo2 | ENSMUST00000016401 | 29.0495315 | 31 | 17.5368139 | 22.2222222 | 0 | 0 | 1 | 1 | 0 | 0 | 0 | 0 | 1 | 3 | 0 | 0 | 0 | 0 | 3 |
| ENSMUSG00000027281 | Slx4ip | ENSMUST00000099311 | 29.5140806 | 30 | 19.7128658 | 20.6791828 | 0 | 0 | 0 | 1 | 0 | 0 | 0 | 0 | 1 | 2 | 0 | 1 | 0 | 0 | 3 |
| ENSMUSG00000024563 | Smad2 | ENSMUST00000168423 | 27.3726274 | 33 | 19.2807193 | 19.8801199 | 0 | 0 | 0 | 0 | 0 | 0 | 0 | 0 | 0 | 0 | 0 | 1 | 0 | 0 | 1 |
| ENSMUSG00000021540 | Smad5 | ENSMUST00000069557 | 25.8215962 | 31 | 20.0040825 | 22.7189222 | 0 | 0 | 0 | 0 | 0 | 0 | 0 | 0 | 0 | 0 | 0 | 0 | 0 | 2 | 2 |
| ENSMUSG00000025880 | Smad7 | ENSMUST00000026999 | 22.6751592 | 30 | 23.1210191 | 23.9490446 | 0 | 0 | 0 | 0 | 0 | 0 | 0 | 0 | 0 | 0 | 0 | 0 | 9 | 0 | 9 |
| ENSMUSG00000031715 | Smarca5 | ENSMUST00000043359 | 30.6338028 | 35 | 15.2973396 | 18.9749609 | 1 | 1 | 0 | 0 | 0 | 0 | 0 | 0 | 0 | 0 | 0 | 1 | 4 | 2 | 9 |
| ENSMUSG00000029920 | Smarcad1 | ENSMUST00000204620 | 31.6153029 | 30 | 18.0924548 | 20.5100956 | 0 | 0 | 0 | 0 | 0 | 0 | 0 | 0 | 0 | 0 | 1 | 0 | 0 | 0 | 1 |
| ENSMUSG00000032481 | Smarcc1 | ENSMUST00000199896 | 23.4974958 | 31 | 23.2053422 | 22.4123539 | 0 | 0 | 0 | 0 | 0 | 0 | 0 | 0 | 0 | 0 | 4 | 17 | 10 | 0 | 31 |
| ENSMUSG00000028312 | Smc2 | ENSMUST00000102915 | 28.7227414 | 37 | 16.1370717 | 18.2554517 | 0 | 0 | 0 | 0 | 0 | 0 | 0 | 0 | 0 | 0 | 0 | 0 | 3 | 0 | 3 |
| ENSMUSG00000024943 | Smc5 | ENSMUST00000087556 | 31.7134478 | 35 | 13.7410976 | 19.3548387 | 2 | 1 | 0 | 1 | 0 | 0 | 0 | 0 | 1 | 2 | 6 | 0 | 6 | 1 | 18 |
| ENSMUSG00000041846 | Smek1 | ENSMUST00000048305 | 29.6031746 | 35 | 16.5079365 | 19.3650794 | 0 | 0 | 0 | 0 | 0 | 0 | 0 | 0 | 0 | 0 | 0 | 1 | 6 | 5 | 12 |
| ENSMUSG00000020463 | Smek2 | ENSMUST00000020755 | 32.635797 | 31 | 17.4087266 | 18.9225289 | 0 | 0 | 0 | 0 | 0 | 0 | 0 | 0 | 0 | 0 | 0 | 0 | 1 | 0 | 1 |
| ENSMUSG00000020495 | Smg8 | ENSMUST00000143280 | 26.5129683 | 35 | 17.5792507 | 20.4610951 | 0 | 0 | 0 | 0 | 0 | 0 | 0 | 0 | 0 | 0 | 0 | 1 | 1 | 0 | 2 |
| ENSMUSG00000072704 | Smim10l1 | ENSMUST00000100864 | 34.2709798 | 30 | 16.3150492 | 19.737459 | 0 | 0 | 0 | 0 | 0 | 0 | 0 | 0 | 0 | 0 | 0 | 1 | 8 | 2 | 11 |
| ENSMUSG00000091264 | Smim13 | ENSMUST00000165561 | 28.8108958 | 32 | 18.2032478 | 20.9795705 | 0 | 0 | 0 | 0 | 0 | 0 | 0 | 0 | 0 | 0 | 0 | 1 | 4 | 1 | 6 |
| ENSMUSG00000037822 | Smim14 | ENSMUST00000139122 | 26.3409155 | 32 | 20.9772536 | 20.8368436 | 0 | 0 | 0 | 0 | 0 | 0 | 0 | 0 | 0 | 0 | 0 | 0 | 1 | 1 | 2 |
| ENSMUSG00000025024 | Smndc1 | ENSMUST00000025997 | 28.56 | 36 | 15.92 | 19.04 | 0 | 0 | 0 | 0 | 0 | 0 | 0 | 0 | 0 | 0 | 0 | 0 | 1 | 0 | 1 |
| ENSMUSG00000036061 | Smug1 | ENSMUST00000064067 | 23.5720375 | 31 | 23.1031543 | 22.3358909 | 1 | 0 | 0 | 0 | 0 | 0 | 0 | 0 | 0 | 0 | 0 | 1 | 0 | 0 | 2 |
| ENSMUSG00000042821 | Snai1 | ENSMUST00000052631 | 20.6566347 | 26 | 30.506156 | 22.8454172 | 0 | 0 | 0 | 0 | 0 | 0 | 0 | 0 | 0 | 0 | 0 | 2 | 0 | 0 | 2 |
| ENSMUSG00000022676 | Snai2 | ENSMUST00000023356 | 37.033667 | 29 | 17.1974522 | 16.7424932 | 0 | 0 | 0 | 0 | 0 | 0 | 0 | 0 | 0 | 0 | 0 | 0 | 0 | 1 | 1 |
| ENSMUSG00000027287 | Snap23 | ENSMUST00000028743 | 29.5715778 | 33 | 17.5548589 | 19.9582027 | 0 | 0 | 0 | 0 | 0 | 0 | 0 | 0 | 0 | 0 | 0 | 0 | 1 | 0 | 1 |
| ENSMUSG00000027273 | Snap25 | ENSMUST00000028727 | 29.4941634 | 33 | 21.4007782 | 16.0311284 | 0 | 0 | 0 | 0 | 0 | 0 | 0 | 0 | 0 | 0 | 1 | 2 | 2 | 0 | 5 |
| ENSMUSG00000022765 | Snap29 | ENSMUST00000023449 | 27.3159145 | 30 | 21.3776722 | 21.6547902 | 0 | 0 | 0 | 0 | 0 | 0 | 0 | 0 | 0 | 0 | 0 | 0 | 2 | 0 | 2 |
| ENSMUSG00000028483 | Snapc3 | ENSMUST00000123262 | 32.0726172 | 33 | 18.419062 | 16.7170953 | 0 | 0 | 0 | 0 | 0 | 0 | 0 | 0 | 0 | 0 | 1 | 1 | 4 | 0 | 6 |
| ENSMUSG00000001018 | Snapin | ENSMUST00000149884 | 28.3950617 | 30 | 22.2908093 | 18.9986283 | 1 | 0 | 0 | 1 | 0 | 0 | 0 | 0 | 1 | 2 | 0 | 0 | 1 | 0 | 4 |
| ENSMUSG00000044349 | Snhg11 | ENSMUST00000109488 | 26.4053537 | 30 | 22.08413 | 21.0133843 | 4 | 2 | 0 | 8 | 0 | 1 | 1 | 2 | 5 | 17 | 0 | 29 | 13 | 2 | 67 |
| ENSMUSG00000074088 | Snrnp40 | ENSMUST00000105994 | 24.9422633 | 33 | 19.630485 | 22.4018476 | 0 | 0 | 0 | 0 | 0 | 0 | 0 | 0 | 0 | 0 | 0 | 0 | 0 | 1 | 1 |
| ENSMUSG00000008333 | Snrpb2 | ENSMUST00000008477 | 29.1217257 | 34 | 16.7950693 | 20.4930663 | 0 | 0 | 1 | 0 | 0 | 0 | 0 | 0 | 0 | 1 | 0 | 0 | 0 | 0 | 1 |
| ENSMUSG00000002477 | Snrpd1 | ENSMUST00000002551 | 28.0555556 | 38 | 12.2222222 | 22.2222222 | 0 | 0 | 0 | 0 | 0 | 0 | 0 | 0 | 0 | 0 | 1 | 0 | 1 | 0 | 2 |
| ENSMUSG00000020180 | Snrpd3 | ENSMUST00000020397 | 22.3793272 | 33 | 22.4280839 | 22.1843003 | 0 | 0 | 0 | 0 | 0 | 0 | 0 | 0 | 0 | 0 | 0 | 0 | 4 | 0 | 4 |
| ENSMUSG00000090553 | Snrpe | ENSMUST00000164096 | 24.099723 | 34 | 22.7146814 | 18.8365651 | 0 | 0 | 0 | 0 | 0 | 0 | 0 | 0 | 0 | 0 | 0 | 1 | 0 | 0 | 1 |
| ENSMUSG00000057278 | Snrpg | ENSMUST00000089558 | 27.1559633 | 33 | 17.0642202 | 22.7522936 | 0 | 0 | 1 | 0 | 0 | 0 | 0 | 0 | 0 | 1 | 1 | 0 | 0 | 0 | 2 |
| ENSMUSG00000102252 | Snrpn | ENSMUST00000059305 | 32.4832215 | 35 | 17.3154362 | 14.8993289 | 0 | 0 | 0 | 0 | 0 | 0 | 0 | 0 | 0 | 0 | 0 | 1 | 0 | 0 | 1 |
| ENSMUSG00000041308 | Sntb2 | ENSMUST00000047425 | 26.2108262 | 29 | 21.474359 | 23.0769231 | 1 | 0 | 0 | 1 | 0 | 0 | 0 | 0 | 1 | 2 | 1 | 0 | 6 | 0 | 10 |
| ENSMUSG00000038301 | Snx10 | ENSMUST00000049152 | 30.8095515 | 32 | 17.2976121 | 19.5107746 | 1 | 0 | 0 | 0 | 0 | 0 | 0 | 0 | 0 | 0 | 0 | 0 | 0 | 1 | 2 |
| ENSMUSG00000046032 | Snx12 | ENSMUST00000120389 | 21.4642263 | 32 | 22.8508042 | 23.7382141 | 0 | 0 | 0 | 0 | 0 | 0 | 0 | 0 | 0 | 0 | 3 | 1 | 2 | 1 | 7 |
| ENSMUSG00000020590 | Snx13 | ENSMUST00000048519 | 29.3967443 | 32 | 18.8637089 | 19.4063198 | 2 | 0 | 0 | 1 | 0 | 0 | 0 | 0 | 1 | 2 | 0 | 0 | 1 | 3 | 8 |
| ENSMUSG00000032422 | Snx14 | ENSMUST00000126405 | 30.0153139 | 27 | 21.1842777 | 22.0520674 | 0 | 0 | 0 | 0 | 0 | 0 | 0 | 0 | 0 | 0 | 0 | 1 | 0 | 0 | 1 |
| ENSMUSG00000027534 | Snx16 | ENSMUST00000099223 | 31.0687023 | 35 | 16.870229 | 17.0992366 | 0 | 0 | 0 | 0 | 0 | 0 | 0 | 0 | 0 | 0 | 1 | 0 | 0 | 0 | 1 |
| ENSMUSG00000028136 | Snx27 | ENSMUST00000029783 | 24.5956127 | 30 | 22.8672723 | 22.4905828 | 0 | 0 | 0 | 0 | 0 | 0 | 0 | 0 | 0 | 0 | 2 | 3 | 6 | 0 | 11 |
| ENSMUSG00000026600 | Soat1 | ENSMUST00000189661 | 25.4820937 | 32 | 19.8806244 | 22.6354454 | 0 | 0 | 0 | 0 | 0 | 0 | 0 | 0 | 0 | 0 | 4 | 1 | 29 | 3 | 37 |
| ENSMUSG00000053113 | Socs3 | ENSMUST00000054002 | 23.7449118 | 22 | 26.2550882 | 28.0189959 | 1 | 0 | 0 | 1 | 0 | 0 | 0 | 0 | 1 | 2 | 0 | 2 | 0 | 0 | 5 |
| ENSMUSG00000048379 | Socs4 | ENSMUST00000065562 | 29.9343376 | 35 | 16.801854 | 18.3468521 | 0 | 0 | 0 | 0 | 0 | 0 | 0 | 0 | 0 | 0 | 3 | 25 | 6 | 4 | 38 |
| ENSMUSG00000056153 | Socs6 | ENSMUST00000070116 | 32.4956457 | 34 | 15.7999502 | 17.3426225 | 0 | 0 | 0 | 0 | 0 | 0 | 0 | 0 | 0 | 0 | 0 | 0 | 2 | 1 | 3 |
| ENSMUSG00000038485 | Socs7 | ENSMUST00000045540 | 23.0516975 | 27 | 22.8009259 | 27.6041667 | 1 | 0 | 0 | 1 | 0 | 0 | 0 | 0 | 1 | 2 | 0 | 2 | 0 | 1 | 6 |
| ENSMUSG00000006818 | Sod2 | ENSMUST00000007012 | 25.0339213 | 35 | 19.1316147 | 20.3867028 | 1 | 0 | 0 | 1 | 0 | 0 | 0 | 0 | 1 | 2 | 0 | 2 | 19 | 4 | 28 |
| ENSMUSG00000068747 | Sort1 | ENSMUST00000102632 | 23.5941893 | 28 | 24.6485473 | 24.2268041 | 1 | 0 | 0 | 1 | 0 | 0 | 0 | 0 | 1 | 2 | 0 | 2 | 1 | 0 | 6 |
| ENSMUSG00000024241 | Sos1 | ENSMUST00000068714 | 30.029352 | 33 | 18.2659743 | 18.8530142 | 0 | 0 | 0 | 0 | 0 | 0 | 0 | 0 | 0 | 0 | 1 | 0 | 2 | 0 | 3 |
| ENSMUSG00000034801 | Sos2 | ENSMUST00000035773 | 31.3190862 | 32 | 17.0228445 | 19.7494473 | 0 | 0 | 1 | 0 | 0 | 0 | 0 | 0 | 0 | 1 | 0 | 0 | 1 | 0 | 2 |
| ENSMUSG00000036169 | Sostdc1 | ENSMUST00000041407 | 28.0597015 | 33 | 20.6965174 | 18.1094527 | 0 | 0 | 2 | 0 | 0 | 0 | 0 | 0 | 0 | 2 | 0 | 0 | 0 | 1 | 3 |
| ENSMUSG00000063632 | Sox11 | ENSMUST00000079063 | 26.4769297 | 31 | 20.0517464 | 22.0209861 | 0 | 0 | 0 | 0 | 0 | 0 | 0 | 0 | 0 | 0 | 2 | 3 | 29 | 8 | 42 |
| ENSMUSG00000076431 | Sox4 | ENSMUST00000067230 | 28.4892086 | 31 | 19.3165468 | 21.3309353 | 0 | 0 | 0 | 0 | 0 | 0 | 0 | 0 | 0 | 0 | 0 | 1 | 3 | 4 | 8 |
| ENSMUSG00000041540 | Sox5 | ENSMUST00000038815 | 28.0223187 | 32 | 19.6941517 | 20.582765 | 1 | 0 | 0 | 1 | 0 | 0 | 0 | 0 | 1 | 2 | 5 | 3 | 19 | 2 | 32 |
| ENSMUSG00000051910 | Sox6 | ENSMUST00000206034 | 26.4472378 | 30 | 22.1964935 | 21.1048627 | 3 | 2 | 0 | 2 | 0 | 0 | 0 | 1 | 1 | 4 | 4 | 3 | 7 | 3 | 26 |
| ENSMUSG00000063060 | Sox7 | ENSMUST00000079652 | 25.3256151 | 30 | 24.409069 | 20.2122528 | 0 | 0 | 0 | 1 | 0 | 0 | 0 | 0 | 1 | 2 | 0 | 1 | 8 | 1 | 12 |
| ENSMUSG00000000567 | Sox9 | ENSMUST00000000579 | 28.6222222 | 35 | 17.6 | 18.7555556 | 1 | 0 | 0 | 2 | 0 | 0 | 0 | 0 | 2 | 4 | 1 | 3 | 7 | 9 | 25 |
| ENSMUSG00000001280 | Sp1 | ENSMUST00000001326 | 26.824274 | 34 | 19.7877885 | 19.7877885 | 0 | 0 | 0 | 0 | 0 | 0 | 0 | 0 | 0 | 0 | 1 | 3 | 3 | 3 | 10 |
| ENSMUSG00000027109 | Sp3 | ENSMUST00000102689 | 33.2519829 | 39 | 12.3245882 | 15.2532032 | 1 | 0 | 0 | 0 | 0 | 0 | 0 | 0 | 0 | 0 | 0 | 0 | 5 | 2 | 8 |
| ENSMUSG00000068859 | Sp9 | ENSMUST00000090813 | 29.8661174 | 28 | 20.8719533 | 21.489873 | 1 | 0 | 0 | 0 | 0 | 0 | 0 | 0 | 0 | 0 | 0 | 0 | 2 | 4 | 7 |
| ENSMUSG00000037617 | Spag1 | ENSMUST00000047348 | 29.3539326 | 37 | 18.6797753 | 14.6067416 | 1 | 0 | 0 | 2 | 0 | 0 | 0 | 0 | 2 | 4 | 0 | 0 | 0 | 0 | 5 |
| ENSMUSG00000018287 | Spag7 | ENSMUST00000018431 | 18.8235294 | 32 | 21.1764706 | 27.6470588 | 0 | 0 | 1 | 0 | 0 | 0 | 0 | 0 | 0 | 1 | 0 | 0 | 0 | 0 | 1 |
| ENSMUSG00000020859 | Spag9 | ENSMUST00000041956 | 30.0492611 | 33 | 17.6108374 | 19.7660099 | 0 | 0 | 0 | 0 | 0 | 0 | 0 | 0 | 0 | 0 | 0 | 2 | 0 | 0 | 2 |
| ENSMUSG00000024068 | Spast | ENSMUST00000024869 | 28.3827205 | 34 | 18.4219921 | 19.3859336 | 0 | 0 | 0 | 0 | 0 | 0 | 0 | 0 | 0 | 0 | 0 | 1 | 0 | 0 | 1 |

| | | | | | | | | | | | | | | | | | | | | | |
|---|---|---|---|---|---|---|---|---|---|---|---|---|---|---|---|---|---|---|---|---|---|
| ENSMUSG00000028188 | Spata1 | ENSMUST00000093951 | 30.4200869 | 29 | 21.7286335 | 18.7349107 | 0 | 0 | 0 | 0 | 0 | 0 | 0 | 0 | 0 | 0 | 0 | 28 | 1 | 0 | 29 |
| ENSMUSG00000034401 | Spata6 | ENSMUST00000084354 | 31.2958435 | 34 | 18.3374083 | 16.0146699 | 1 | 0 | 0 | 1 | 0 | 0 | 0 | 0 | 1 | 2 | 0 | 0 | 0 | 0 | 3 |
| ENSMUSG00000051934 | Spats2 | ENSMUST00000063517 | 24.3786982 | 32 | 22.7218935 | 21.0650888 | 0 | 0 | 0 | 0 | 0 | 0 | 0 | 0 | 0 | 0 | 0 | 0 | 4 | 1 | 5 |
| ENSMUSG00000054408 | Spcs3 | ENSMUST00000067476 | 28.5512367 | 33 | 17.4558304 | 20.7067138 | 0 | 0 | 0 | 0 | 0 | 0 | 0 | 0 | 0 | 0 | 2 | 0 | 1 | 0 | 3 |
| ENSMUSG00000043065 | Spice1 | ENSMUST00000050897 | 28.7092883 | 32 | 18.8781665 | 20.0844391 | 0 | 0 | 0 | 0 | 0 | 0 | 0 | 0 | 0 | 0 | 1 | 0 | 1 | 0 | 2 |
| ENSMUSG00000021395 | Spin1 | ENSMUST00000095797 | 24.3722304 | 33 | 20.6794682 | 21.5361891 | 0 | 0 | 0 | 0 | 0 | 0 | 0 | 0 | 0 | 0 | 2 | 7 | 6 | 1 | 16 |
| ENSMUSG00000071722 | Spin4 | ENSMUST00000096367 | 30.0170358 | 42 | 14.3781942 | 13.560477 | 0 | 0 | 0 | 1 | 0 | 0 | 0 | 0 | 1 | 2 | 0 | 1 | 2 | 1 | 6 |
| ENSMUSG00000044176 | Spink10 | ENSMUST00000162365 | 30.2600473 | 37 | 12.2931442 | 20.5673759 | 1 | 0 | 1 | 0 | 0 | 0 | 0 | 0 | 0 | 1 | 0 | 0 | 0 | 0 | 2 |
| ENSMUSG00000054162 | Spock3 | ENSMUST00000119068 | 34.8432056 | 38 | 13.3565621 | 14.0534262 | 1 | 0 | 0 | 3 | 0 | 0 | 0 | 0 | 3 | 6 | 3 | 1 | 1 | 0 | 12 |
| ENSMUSG00000057522 | Spop | ENSMUST00000107724 | 28.5623003 | 28 | 19.2332268 | 24.0255591 | 0 | 1 | 0 | 0 | 0 | 0 | 0 | 0 | 0 | 0 | 0 | 0 | 0 | 0 | 1 |
| ENSMUSG00000026771 | Spopl | ENSMUST00000132484 | 31.3186813 | 35 | 15.6985871 | 18.1514914 | 1 | 0 | 0 | 2 | 0 | 0 | 0 | 0 | 2 | 4 | 1 | 2 | 1 | 2 | 11 |
| ENSMUSG00000027366 | Sppl2a | ENSMUST00000028844 | 29.7042659 | 34 | 18.9217482 | 17.4823345 | 0 | 0 | 0 | 0 | 0 | 0 | 0 | 0 | 0 | 0 | 3 | 1 | 6 | 3 | 13 |
| ENSMUSG00000027351 | Spred1 | ENSMUST00000028829 | 29.3764434 | 34 | 17.3903002 | 19.4688222 | 1 | 0 | 1 | 0 | 0 | 0 | 0 | 0 | 0 | 1 | 0 | 1 | 1 | 2 | 6 |
| ENSMUSG00000045671 | Spred2 | ENSMUST00000093299 | 24.0791004 | 30 | 25.5913145 | 20.0465297 | 0 | 0 | 0 | 0 | 0 | 0 | 0 | 0 | 0 | 0 | 0 | 6 | 2 | 1 | 9 |
| ENSMUSG00000024427 | Spry4 | ENSMUST00000025295 | 23.6856981 | 29 | 23.9966083 | 23.7422272 | 2 | 0 | 0 | 1 | 0 | 0 | 0 | 0 | 1 | 2 | 0 | 0 | 6 | 0 | 10 |
| ENSMUSG00000021930 | Spryd7 | ENSMUST00000022497 | 26.9120914 | 36 | 17.9942875 | 19.2319898 | 1 | 0 | 0 | 1 | 0 | 0 | 0 | 0 | 1 | 2 | 4 | 2 | 3 | 2 | 14 |
| ENSMUSG00000021468 | Sptlc1 | ENSMUST00000021920 | 22.6027397 | 33 | 21.4041096 | 23.2020548 | 0 | 0 | 0 | 0 | 0 | 0 | 0 | 0 | 0 | 0 | 0 | 0 | 1 | 0 | 1 |
| ENSMUSG00000021036 | Sptlc2 | ENSMUST00000021424 | 26.7189758 | 30 | 21.8666116 | 21.4123477 | 0 | 0 | 0 | 0 | 0 | 0 | 0 | 0 | 0 | 0 | 1 | 2 | 2 | 0 | 5 |
| ENSMUSG00000044408 | Sptssa | ENSMUST00000056228 | 29.7731569 | 36 | 16.1625709 | 18.4310019 | 0 | 0 | 0 | 0 | 0 | 0 | 0 | 0 | 0 | 0 | 0 | 0 | 0 | 2 | 2 |
| ENSMUSG00000049516 | Spty2d1 | ENSMUST00000061639 | 28.1962681 | 33 | 17.6226676 | 20.7671044 | 0 | 0 | 0 | 0 | 0 | 0 | 0 | 0 | 0 | 0 | 1 | 0 | 1 | 0 | 2 |
| ENSMUSG00000032621 | Srek1 | ENSMUST00000074616 | 28.0656747 | 32 | 18.0092355 | 21.4982042 | 0 | 0 | 0 | 0 | 0 | 0 | 0 | 0 | 0 | 0 | 0 | 0 | 1 | 1 | 2 |
| ENSMUSG00000021716 | Srek1ip1 | ENSMUST00000022230 | 27.4623407 | 34 | 19.4090382 | 19.0614137 | 0 | 0 | 0 | 0 | 0 | 0 | 0 | 0 | 0 | 0 | 0 | 1 | 0 | 0 | 1 |
| ENSMUSG00000024528 | Srfbp1; LOC102641936 | ENSMUST00000025406 | 25.4545455 | 46 | 14.5454545 | 13.6363636 | 0 | 0 | 0 | 0 | 0 | 0 | 0 | 0 | 0 | 0 | 0 | 0 | 1 | 0 | 1 |
| ENSMUSG00000003161 | Sri | ENSMUST00000088786 | 31.5934066 | 32 | 18.2417582 | 18.4065934 | 0 | 0 | 0 | 0 | 0 | 0 | 0 | 0 | 0 | 0 | 0 | 1 | 0 | 0 | 1 |
| ENSMUSG00000014504 | Srp19 | ENSMUST00000142010 | 31.3924051 | 26 | 19.4936709 | 22.9113924 | 0 | 0 | 0 | 0 | 0 | 0 | 0 | 0 | 0 | 0 | 1 | 0 | 0 | 0 | 1 |
| ENSMUSG00000062604 | Srpk2 | ENSMUST00000088392 | 30.8754682 | 32 | 18.1882022 | 18.7734082 | 0 | 0 | 0 | 0 | 0 | 0 | 0 | 0 | 0 | 0 | 1 | 1 | 1 | 4 | 7 |
| ENSMUSG00000028809 | Srrm1 | ENSMUST00000105861 | 24.6813441 | 36 | 18.3082271 | 21.0892236 | 0 | 0 | 0 | 0 | 0 | 0 | 0 | 0 | 0 | 0 | 0 | 2 | 4 | 0 | 6 |
| ENSMUSG00000018379 | Srsf1 | ENSMUST00000139129 | 30.0072307 | 35 | 15.2566884 | 19.7637985 | 0 | 0 | 0 | 1 | 0 | 0 | 0 | 0 | 1 | 2 | 0 | 2 | 2 | 2 | 8 |
| ENSMUSG00000028676 | Srsf10; Gm5446; LOC632268 | ENSMUST00000126641 | 28.9860397 | 36 | 16.3115356 | 18.9566495 | 0 | 0 | 0 | 0 | 0 | 0 | 0 | 0 | 0 | 0 | 0 | 0 | 12 | 1 | 13 |
| ENSMUSG00000054679 | Srsf12 | ENSMUST00000067864 | 31.6072301 | 30 | 19.7361993 | 18.8080117 | 0 | 0 | 1 | 0 | 0 | 0 | 0 | 0 | 0 | 1 | 0 | 0 | 0 | 0 | 1 |
| ENSMUSG00000034120 | Srsf2 | ENSMUST00000092404 | 28.7453184 | 36 | 16.3857678 | 19.2883895 | 0 | 0 | 0 | 0 | 0 | 0 | 0 | 0 | 0 | 0 | 0 | 1 | 0 | 1 | 2 |
| ENSMUSG00000071172 | Srsf3 | ENSMUST00000130216 | 29.1560102 | 35 | 15.8056266 | 20.1023018 | 1 | 0 | 0 | 1 | 0 | 0 | 0 | 0 | 1 | 2 | 0 | 0 | 0 | 0 | 3 |
| ENSMUSG00000016921 | Srsf6 | ENSMUST00000130411 | 29.6531303 | 34 | 16.0321489 | 20.3891709 | 0 | 0 | 0 | 0 | 0 | 0 | 0 | 0 | 0 | 0 | 0 | 0 | 2 | 0 | 2 |
| ENSMUSG00000039086 | Ss18l1 | ENSMUST00000041126 | 25.2901864 | 33 | 19.2402392 | 22.968695 | 0 | 1 | 0 | 0 | 0 | 0 | 0 | 0 | 0 | 0 | 0 | 0 | 2 | 1 | 4 |
| ENSMUSG00000029911 | Ssbp1 | ENSMUST00000114779 | 26.8664564 | 33 | 20.0052576 | 20.1104101 | 1 | 0 | 0 | 0 | 0 | 0 | 0 | 0 | 0 | 0 | 0 | 0 | 2 | 1 | 4 |
| ENSMUSG00000003992 | Ssbp2 | ENSMUST00000004094 | 28.127286 | 30 | 21.6532553 | 19.7512802 | 0 | 0 | 0 | 0 | 0 | 0 | 0 | 0 | 0 | 0 | 0 | 3 | 1 | 2 | 6 |
| ENSMUSG00000021427 | Ssr1 | ENSMUST00000021864 | 26.5051571 | 30 | 20.4365555 | 23.3869033 | 0 | 0 | 0 | 0 | 0 | 0 | 0 | 0 | 0 | 0 | 1 | 5 | 13 | 5 | 24 |
| ENSMUSG00000027828 | Ssr3 | ENSMUST00000029414 | 27.9723303 | 34 | 17.3800259 | 21.054907 | 0 | 0 | 0 | 1 | 0 | 0 | 0 | 0 | 1 | 2 | 1 | 0 | 5 | 4 | 12 |
| ENSMUSG00000022403 | St13 | ENSMUST00000172107 | 28.5794497 | 36 | 17.3498035 | 18.1358787 | 0 | 0 | 0 | 0 | 0 | 0 | 0 | 0 | 0 | 0 | 0 | 1 | 15 | 2 | 18 |
| ENSMUSG00000033740 | St18 | ENSMUST00000043578 | 31.7948718 | 34 | 16.6239316 | 17.0940171 | 2 | 1 | 0 | 2 | 0 | 0 | 0 | 0 | 2 | 4 | 1 | 0 | 0 | 0 | 8 |
| ENSMUSG00000022747 | St3gal6 | ENSMUST00000137035 | 30.4520222 | 29 | 19.1514671 | 21.1736717 | 0 | 0 | 0 | 1 | 0 | 0 | 0 | 0 | 1 | 2 | 0 | 1 | 0 | 0 | 3 |
| ENSMUSG00000039037 | St6galnac5 | ENSMUST00000044278 | 27.3814898 | 29 | 19.3679458 | 23.8600451 | 1 | 1 | 0 | 0 | 0 | 0 | 0 | 0 | 0 | 0 | 1 | 1 | 0 | 0 | 4 |
| ENSMUSG00000029534 | St7 | ENSMUST00000144488 | 28.3213182 | 26 | 23.8414006 | 21.884655 | 0 | 0 | 0 | 0 | 0 | 0 | 0 | 0 | 0 | 0 | 0 | 1 | 0 | 0 | 1 |
| ENSMUSG00000045576 | St7l | ENSMUST00000059271 | 28.7615741 | 31 | 19.6180556 | 20.4861111 | 2 | 1 | 1 | 3 | 0 | 0 | 0 | 1 | 2 | 7 | 0 | 14 | 0 | 0 | 24 |
| ENSMUSG00000056812 | St8sia3 | ENSMUST00000025477 | 29.3563218 | 29 | 21.2413793 | 19.9310345 | 0 | 0 | 0 | 0 | 0 | 0 | 0 | 0 | 0 | 0 | 0 | 18 | 1 | 0 | 19 |
| ENSMUSG00000040710 | St8sia4 | ENSMUST00000189556 | 32.8976765 | 32 | 17.3927614 | 17.2252011 | 3 | 2 | 0 | 6 | 0 | 0 | 0 | 1 | 5 | 12 | 1 | 1 | 3 | 2 | 24 |
| ENSMUSG00000025862 | Stag2 | ENSMUST00000069619 | 32.7014218 | 36 | 15.0078989 | 15.9031069 | 0 | 1 | 1 | 0 | 0 | 0 | 0 | 0 | 0 | 1 | 0 | 0 | 1 | 0 | 3 |
| ENSMUSG00000026718 | Stam | ENSMUST00000102960 | 28.0279011 | 37 | 17.2479391 | 17.5015853 | 2 | 0 | 0 | 2 | 0 | 0 | 0 | 0 | 2 | 4 | 0 | 0 | 3 | 1 | 10 |
| ENSMUSG00000055371 | Stam2 | ENSMUST00000102759 | 27.0607553 | 34 | 18.7520525 | 20 | 0 | 0 | 0 | 1 | 0 | 0 | 0 | 0 | 1 | 2 | 0 | 0 | 1 | 0 | 3 |
| ENSMUSG00000024776 | Stambpl1 | ENSMUST00000054956 | 25.7692308 | 38 | 17.6923077 | 18.0769231 | 1 | 0 | 1 | 1 | 0 | 0 | 0 | 0 | 1 | 3 | 0 | 0 | 4 | 0 | 8 |
| ENSMUSG00000029254 | Stap1 | ENSMUST00000198435 | 34.0524535 | 31 | 17.6395939 | 17.7241963 | 0 | 0 | 1 | 0 | 0 | 0 | 0 | 0 | 0 | 1 | 0 | 0 | 0 | 0 | 1 |
| ENSMUSG00000003062 | Stard3nl | ENSMUST00000039694 | 24.248927 | 36 | 20.3862661 | 18.8841202 | 0 | 0 | 0 | 0 | 0 | 0 | 0 | 0 | 0 | 0 | 0 | 0 | 1 | 0 | 1 |
| ENSMUSG00000024378 | Stard4 | ENSMUST00000025236 | 26.9439849 | 31 | 19.2625857 | 23.2805483 | 0 | 0 | 0 | 0 | 0 | 0 | 0 | 0 | 0 | 0 | 1 | 0 | 0 | 0 | 1 |
| ENSMUSG00000025920 | Stau2 | ENSMUST00000127420 | 28.3690658 | 29 | 22.4349158 | 20.0612557 | 0 | 0 | 0 | 0 | 0 | 0 | 0 | 0 | 0 | 0 | 0 | 19 | 1 | 0 | 20 |
| ENSMUSG00000014813 | Stc1 | ENSMUST00000014957 | 28.1918565 | 31 | 22.2567288 | 18.4610076 | 0 | 0 | 0 | 0 | 0 | 0 | 0 | 0 | 0 | 0 | 0 | 2 | 5 | 0 | 7 |
| ENSMUSG00000020303 | Stc2 | ENSMUST00000020546 | 24.7796969 | 27 | 24.4977088 | 24.0394783 | 0 | 0 | 0 | 0 | 0 | 0 | 0 | 0 | 0 | 0 | 1 | 2 | 6 | 4 | 13 |
| ENSMUSG00000015653 | Steap2 | ENSMUST00000115426 | 31.5501166 | 31 | 19.0792541 | 18.3333333 | 0 | 0 | 0 | 2 | 0 | 0 | 0 | 0 | 2 | 4 | 1 | 3 | 1 | 0 | 9 |
| ENSMUSG00000031112 | Stk26 | ENSMUST00000033444 | 30.9080654 | 37 | 15.2284264 | 16.9204738 | 0 | 0 | 0 | 1 | 0 | 0 | 0 | 0 | 1 | 2 | 0 | 0 | 1 | 0 | 3 |
| ENSMUSG00000037885 | Stk35 | ENSMUST00000166282 | 20.9601082 | 29 | 26.7297724 | 23.4843363 | 0 | 0 | 0 | 0 | 0 | 0 | 0 | 0 | 0 | 0 | 0 | 7 | 9 | 0 | 16 |
| ENSMUSG00000018209 | Stk4 | ENSMUST00000018353 | 25.7514216 | 28 | 23.0165177 | 22.7728134 | 0 | 0 | 0 | 0 | 0 | 0 | 0 | 0 | 0 | 0 | 0 | 1 | 0 | 0 | 1 |
| ENSMUSG00000069631 | Strada | ENSMUST00000106867 | 29.6345077 | 31 | 19.1307211 | 20.546592 | 1 | 0 | 0 | 1 | 0 | 0 | 0 | 0 | 1 | 2 | 0 | 15 | 0 | 0 | 18 |
| ENSMUSG00000026915 | Strbp | ENSMUST00000028279 | 26.9171021 | 32 | 19.3757944 | 21.861319 | 0 | 0 | 0 | 0 | 0 | 0 | 0 | 0 | 0 | 0 | 3 | 3 | 41 | 5 | 52 |
| ENSMUSG00000024077 | Strn | ENSMUST00000145910 | 27.2010041 | 34 | 18.4687108 | 20.7638515 | 0 | 0 | 0 | 0 | 0 | 0 | 0 | 0 | 0 | 0 | 1 | 0 | 12 | 0 | 13 |
| ENSMUSG00000020954 | Strn3 | ENSMUST00000013130 | 32.1382843 | 35 | 15.6850192 | 17.2855314 | 1 | 0 | 0 | 1 | 0 | 0 | 0 | 0 | 1 | 2 | 0 | 1 | 1 | 0 | 5 |
| ENSMUSG00000032116 | Stt3a | ENSMUST00000120381 | 25.7541899 | 37 | 18.9385475 | 17.877095 | 0 | 0 | 0 | 1 | 0 | 0 | 0 | 0 | 1 | 2 | 0 | 6 | 20 | 5 | 33 |
| ENSMUSG00000032437 | Stt3b | ENSMUST00000035010 | 25.6259205 | 31 | 22.0913108 | 21.5022091 | 0 | 0 | 0 | 0 | 0 | 0 | 0 | 0 | 0 | 0 | 0 | 0 | 0 | 1 | 1 |
| ENSMUSG00000028879 | Stx12 | ENSMUST00000030698 | 27.6335878 | 32 | 18.0661578 | 22.1882952 | 0 | 0 | 0 | 0 | 0 | 0 | 0 | 0 | 0 | 0 | 1 | 0 | 0 | 0 | 1 |
| ENSMUSG00000027522 | Stx16 | ENSMUST00000087908 | 24.5997089 | 30 | 21.3245997 | 23.6171761 | 0 | 1 | 0 | 1 | 0 | 0 | 0 | 0 | 1 | 2 | 3 | 0 | 9 | 4 | 19 |
| ENSMUSG00000061455 | Stx17 | ENSMUST00000107720 | 27.9303837 | 30 | 21.9333194 | 20.3187251 | 0 | 0 | 0 | 0 | 0 | 0 | 0 | 0 | 0 | 0 | 0 | 1 | 2 | 0 | 3 |
| ENSMUSG00000029125 | Stx18 | ENSMUST00000114126 | 25.1599147 | 25 | 26.3681592 | 23.0987918 | 0 | 0 | 1 | 0 | 0 | 0 | 0 | 0 | 0 | 1 | 0 | 0 | 0 | 0 | 1 |
| ENSMUSG00000047854 | Stx19 | ENSMUST00000055557 | 30.5775076 | 31 | 20.1215805 | 18.1155015 | 5 | 2 | 0 | 7 | 0 | 0 | 1 | 2 | 4 | 14 | 0 | 0 | 0 | 2 | 23 |
| ENSMUSG00000030806 | Stx1b | ENSMUST00000106267 | 19.9178645 | 28 | 26.6940452 | 25.7260194 | 0 | 0 | 0 | 0 | 0 | 0 | 0 | 0 | 0 | 0 | 0 | 12 | 0 | 0 | 12 |
| ENSMUSG00000019998 | Stx7 | ENSMUST00000020174 | 25.3430186 | 38 | 17.1912833 | 19.6125908 | 2 | 1 | 0 | 1 | 0 | 0 | 0 | 1 | 0 | 2 | 0 | 0 | 0 | 0 | 5 |
| ENSMUSG00000027882 | Stxbp3 | ENSMUST00000138552 | 27.8294574 | 34 | 18.1395349 | 19.7674419 | 1 | 1 | 0 | 1 | 0 | 0 | 0 | 0 | 1 | 2 | 1 | 0 | 1 | 0 | 6 |
| ENSMUSG00000046314 | Stxbp6 | ENSMUST00000120531 | 28.3125371 | 34 | 18.7462864 | 18.9245395 | 0 | 0 | 0 | 1 | 0 | 0 | 0 | 0 | 1 | 2 | 3 | 20 | 3 | 0 | 28 |
| ENSMUSG00000022205 | Sub1 | ENSMUST00000022816 | 28.621049 | 33 | 17.2976728 | 20.7016325 | 0 | 0 | 0 | 1 | 0 | 0 | 0 | 0 | 1 | 2 | 0 | 0 | 1 | 0 | 3 |
| ENSMUSG00000022110 | Sucla2 | ENSMUST00000160507 | 31.2680115 | 35 | 16.426513 | 17.2910663 | 1 | 0 | 0 | 0 | 0 | 0 | 0 | 0 | 0 | 0 | 0 | 0 | 0 | 0 | 1 |
| ENSMUSG00000040297 | Suco | ENSMUST00000048377 | 28.1083481 | 37 | 15.8081705 | 19.5381883 | 0 | 1 | 0 | 1 | 0 | 0 | 0 | 0 | 1 | 2 | 1 | 0 | 12 | 1 | 17 |
| ENSMUSG00000026021 | Sumo1 | ENSMUST00000091374 | 28.1984334 | 35 | 17.6240209 | 19.5822454 | 0 | 0 | 0 | 0 | 0 | 0 | 0 | 0 | 0 | 0 | 0 | 0 | 0 | 2 | 2 |
| ENSMUSG00000020265 | Sumo3 | ENSMUST00000020501 | 23.8301978 | 30 | 21.0323203 | 24.9879402 | 0 | 0 | 0 | 0 | 0 | 0 | 0 | 0 | 0 | 0 | 0 | 1 | 1 | 0 | 2 |
| ENSMUSG00000036817 | Sun1 | ENSMUST00000058716 | 25.4392666 | 30 | 20.3972498 | 24.1405653 | 1 | 0 | 0 | 1 | 0 | 0 | 0 | 0 | 1 | 2 | 0 | 0 | 0 | 0 | 3 |
| ENSMUSG00000027751 | Supt20 | ENSMUST00000200439 | 28.6579213 | 28 | 24.1170535 | 19.3239152 | 0 | 1 | 0 | 0 | 0 | 0 | 0 | 0 | 0 | 0 | 0 | 0 | 0 | 0 | 1 |
| ENSMUSG00000021133 | Susd6 | ENSMUST00000068519 | 21.7192132 | 29 | 25.7612503 | 23.7402317 | 0 | 0 | 0 | 1 | 0 | 0 | 0 | 0 | 1 | 2 | 0 | 1 | 3 | 0 | 6 |
| ENSMUSG00000026646 | Suv39h2 | ENSMUST00000027956 | 29.0954416 | 32 | 19.7293447 | 19.5156695 | 0 | 0 | 0 | 0 | 0 | 0 | 0 | 0 | 0 | 0 | 0 | 0 | 0 | 1 | 1 |
| ENSMUSG00000045098 | Suv420h1 | ENSMUST00000113977 | 27.5607959 | 33 | 18.8651437 | 20.8548268 | 0 | 0 | 0 | 0 | 0 | 0 | 0 | 0 | 0 | 0 | 1 | 0 | 0 | 0 | 1 |
| ENSMUSG00000017548 | Suz12 | ENSMUST00000017692 | 27.056389 | 37 | 16.192447 | 20.0724263 | 0 | 0 | 0 | 0 | 0 | 0 | 0 | 0 | 0 | 0 | 0 | 1 | 1 | 0 | 2 |
| ENSMUSG00000052748 | Swt1 | ENSMUST00000064771 | 31.5789474 | 30 | 20.6578947 | 18.0263158 | 0 | 0 | 0 | 0 | 0 | 0 | 0 | 0 | 0 | 0 | 0 | 0 | 0 | 1 | 1 |
| ENSMUSG00000031357 | Syap1 | ENSMUST00000033723 | 32.5272068 | 34 | 17.0495768 | 16.0822249 | 0 | 0 | 1 | 0 | 0 | 0 | 0 | 0 | 0 | 1 | 0 | 0 | 0 | 0 | 1 |
| ENSMUSG00000020059 | Sycp3 | ENSMUST00000123244 | 37.2909699 | 31 | 13.8795987 | 17.8929766 | 0 | 0 | 0 | 0 | 0 | 0 | 0 | 0 | 0 | 0 | 2 | 0 | 2 | 0 | 4 |
| ENSMUSG00000032423 | Syncrip | ENSMUST00000069221 | 31.6494134 | 35 | 14.9482402 | 18.136646 | 2 | 1 | 0 | 0 | 0 | 0 | 0 | 0 | 0 | 0 | 0 | 0 | 2 | 0 | 5 |
| ENSMUSG00000090935 | Synj2bp | ENSMUST00000163402 | 32.0440636 | 31 | 16.5728274 | 20.3671971 | 0 | 0 | 0 | 0 | 0 | 0 | 0 | 0 | 0 | 0 | 1 | 3 | 9 | 3 | 16 |
| ENSMUSG00000021139 | Synj2bp-cox16 | ENSMUST00000166664 | 31.004902 | 34 | 18.9338235 | 15.6862745 | 0 | 0 | 0 | 0 | 0 | 0 | 0 | 0 | 0 | 0 | 1 | 0 | 3 | 2 | 6 |

| | | | | | | | | | | | | | | | | | | | | | |
|---|---|---|---|---|---|---|---|---|---|---|---|---|---|---|---|---|---|---|---|---|---|
| ENSMUSG00000056296 | Synpr | ENSMUST00000070323 | 30.5142084 | 37 | 15.2909337 | 17.5913396 | 0 | 1 | 0 | 0 | 0 | 0 | 0 | 0 | 0 | 0 | 0 | 0 | 3 | 0 | 4 |
| ENSMUSG00000020570 | Sypl | ENSMUST00000076698 | 28.7432351 | 35 | 17.8292243 | 18.7612748 | 0 | 0 | 0 | 0 | 0 | 0 | 0 | 0 | 0 | 0 | 2 | 14 | 0 | 0 | 16 |
| ENSMUSG00000016200 | Syt14 | ENSMUST00000195354 | 27.395456 | 31 | 20.464274 | 20.9252552 | 0 | 0 | 0 | 1 | 0 | 0 | 0 | 0 | 1 | 2 | 0 | 3 | 1 | 1 | 7 |
| ENSMUSG00000024261 | Syt4 | ENSMUST00000025110 | 31.0315789 | 33 | 15.7052632 | 19.8315789 | 2 | 0 | 1 | 2 | 0 | 0 | 0 | 0 | 2 | 5 | 1 | 0 | 2 | 3 | 13 |
| ENSMUSG00000040842 | Szrd1 | ENSMUST00000102487 | 21.1155378 | 24 | 27.0191959 | 27.7435712 | 0 | 0 | 0 | 0 | 0 | 0 | 0 | 0 | 0 | 0 | 3 | 2 | 4 | 0 | 9 |
| ENSMUSG00000035476 | Tab3 | ENSMUST00000048250 | 30.8556833 | 31 | 16.8326948 | 20.8429119 | 0 | 0 | 0 | 0 | 0 | 0 | 0 | 0 | 0 | 0 | 0 | 1 | 1 | 0 | 2 |
| ENSMUSG00000061762 | Tac1 | ENSMUST00000185187 | 27.8899083 | 32 | 18.7155963 | 21.2844037 | 1 | 1 | 0 | 2 | 0 | 0 | 0 | 1 | 1 | 4 | 0 | 0 | 0 | 0 | 6 |
| ENSMUSG00000028899 | Taf12 | ENSMUST00000030731 | 21.9072165 | 28 | 22.4548969 | 27.3518041 | 0 | 0 | 0 | 0 | 0 | 0 | 0 | 0 | 0 | 0 | 0 | 2 | 0 | 1 | 3 |
| ENSMUSG00000048100 | Taf13 | ENSMUST00000143054 | 28.4596577 | 31 | 18.8753056 | 21.5158924 | 0 | 0 | 0 | 1 | 0 | 0 | 0 | 0 | 1 | 2 | 2 | 0 | 0 | 0 | 4 |
| ENSMUSG00000072258 | Taf1a | ENSMUST00000195798 | 26.2721893 | 24 | 22.8402367 | 26.8639053 | 0 | 0 | 0 | 0 | 0 | 0 | 0 | 0 | 0 | 0 | 0 | 1 | 0 | 0 | 1 |
| ENSMUSG00000037343 | Taf2 | ENSMUST00000041733 | 28.3594394 | 34 | 17.2300082 | 20.1154163 | 0 | 0 | 0 | 0 | 0 | 0 | 0 | 0 | 0 | 0 | 0 | 0 | 3 | 1 | 4 |
| ENSMUSG00000023980 | Taf8 | ENSMUST00000067103 | 24.0458015 | 26 | 23.5005453 | 26.9356598 | 0 | 0 | 1 | 0 | 0 | 0 | 0 | 0 | 0 | 1 | 1 | 0 | 1 | 0 | 3 |
| ENSMUSG00000047242 | Taf9b | ENSMUST00000055497 | 28.4295469 | 34 | 19.9875854 | 17.7529485 | 0 | 0 | 0 | 1 | 0 | 0 | 0 | 0 | 1 | 2 | 0 | 0 | 0 | 1 | 3 |
| ENSMUSG00000052031 | Tagap1 | ENSMUST00000063683 | 25.4612546 | 31 | 20.5412054 | 22.6322263 | 0 | 0 | 0 | 0 | 0 | 0 | 0 | 0 | 0 | 0 | 1 | 0 | 0 | 0 | 1 |
| ENSMUSG00000064289 | Tank | ENSMUST00000112494 | 30.877428 | 35 | 16.8117883 | 17.0797053 | 0 | 0 | 0 | 0 | 0 | 0 | 0 | 0 | 0 | 0 | 0 | 0 | 1 | 0 | 1 |
| ENSMUSG00000017291 | Taok1 | ENSMUST00000017435 | 29.0752441 | 34 | 17.4956921 | 19.3337163 | 1 | 0 | 0 | 2 | 0 | 0 | 0 | 0 | 2 | 4 | 1 | 2 | 14 | 2 | 24 |
| ENSMUSG00000061288 | Taok3 | ENSMUST00000092889 | 25.0541516 | 30 | 27.5090253 | 17.0397112 | 0 | 0 | 0 | 0 | 0 | 0 | 0 | 0 | 0 | 0 | 0 | 58 | 0 | 0 | 58 |
| ENSMUSG00000041459 | Tardbp | ENSMUST00000084125 | 27.8959007 | 33 | 17.9622385 | 21.619323 | 0 | 0 | 0 | 0 | 0 | 0 | 0 | 0 | 0 | 0 | 0 | 1 | 3 | 0 | 4 |
| ENSMUSG00000039033 | Tasp1 | ENSMUST00000046656 | 28.8 | 35 | 14.4 | 22.2857143 | 0 | 0 | 0 | 0 | 0 | 0 | 0 | 0 | 0 | 0 | 0 | 1 | 0 | 0 | 1 |
| ENSMUSG00000048720 | Tbc1d12 | ENSMUST00000037302 | 31.6028226 | 34 | 16.6834677 | 17.641129 | 0 | 0 | 0 | 0 | 0 | 0 | 0 | 0 | 0 | 0 | 0 | 0 | 0 | 2 | 2 |
| ENSMUSG00000020130 | Tbc1d15 | ENSMUST00000020339 | 29.1343284 | 37 | 17.0149254 | 17.0149254 | 0 | 1 | 0 | 0 | 0 | 0 | 0 | 0 | 0 | 0 | 1 | 0 | 3 | 0 | 5 |
| ENSMUSG00000042203 | Tbc1d22b | ENSMUST00000048677 | 20.1768071 | 26 | 27.9771191 | 26.2090484 | 0 | 0 | 0 | 0 | 0 | 0 | 0 | 0 | 0 | 0 | 0 | 2 | 0 | 1 | 3 |
| ENSMUSG00000022364 | Tbc1d31 | ENSMUST00000161329 | 33.8095238 | 41 | 12.3809524 | 12.8571429 | 0 | 0 | 1 | 0 | 0 | 0 | 0 | 0 | 0 | 1 | 0 | 0 | 0 | 0 | 1 |
| ENSMUSG00000038122 | Tbc1d32 | ENSMUST00000099739 | 33.5564344 | 29 | 17.1585032 | 20.1703681 | 0 | 0 | 0 | 1 | 0 | 0 | 0 | 0 | 1 | 2 | 0 | 1 | 1 | 0 | 4 |
| ENSMUSG00000037287 | Tbcel | ENSMUST00000066148 | 23.2147937 | 30 | 22.3044097 | 24.2389758 | 0 | 0 | 0 | 0 | 0 | 0 | 0 | 0 | 0 | 0 | 2 | 10 | 1 | 0 | 13 |
| ENSMUSG00000027630 | Tbl1xr1 | ENSMUST00000193734 | 26.301281 | 35 | 18.4368413 | 20.5448354 | 0 | 0 | 0 | 1 | 0 | 0 | 0 | 0 | 1 | 2 | 1 | 1 | 13 | 4 | 21 |
| ENSMUSG00000014767 | Tbp | ENSMUST00000162505 | 27.3628838 | 30 | 21.2659538 | 20.8951363 | 3 | 2 | 1 | 2 | 0 | 0 | 0 | 0 | 2 | 5 | 1 | 1 | 4 | 4 | 20 |
| ENSMUSG00000071359 | Tbpl1 | ENSMUST00000127698 | 26.3293753 | 32 | 18.5338152 | 22.7155395 | 1 | 0 | 0 | 0 | 0 | 0 | 0 | 0 | 0 | 0 | 0 | 0 | 11 | 1 | 13 |
| ENSMUSG00000035033 | Tbr1 | ENSMUST00000048934 | 23.5083532 | 34 | 23.627685 | 18.5560859 | 0 | 0 | 0 | 0 | 0 | 0 | 0 | 0 | 0 | 0 | 0 | 2 | 2 | 0 | 4 |
| ENSMUSG00000032419 | Tbx18 | ENSMUST00000034991 | 28.9962825 | 34 | 17.0177613 | 20.2808757 | 1 | 0 | 0 | 0 | 0 | 0 | 0 | 0 | 0 | 0 | 3 | 1 | 5 | 1 | 11 |
| ENSMUSG00000046603 | Tcaim | ENSMUST00000136274 | 30.0330033 | 27 | 20.2970297 | 22.359736 | 0 | 1 | 0 | 1 | 0 | 0 | 0 | 0 | 1 | 2 | 0 | 0 | 0 | 0 | 3 |
| ENSMUSG00000033813 | Tcea1 | ENSMUST00000165720 | 30.0970874 | 34 | 17.7346278 | 17.9935275 | 0 | 0 | 0 | 0 | 0 | 0 | 0 | 0 | 0 | 0 | 0 | 1 | 0 | 0 | 1 |
| ENSMUSG00000049536 | Tceal1 | ENSMUST00000055104 | 31.6904652 | 33 | 17.3652695 | 17.779825 | 1 | 0 | 0 | 0 | 0 | 0 | 0 | 0 | 0 | 0 | 0 | 0 | 6 | 1 | 8 |
| ENSMUSG00000051579 | Tceal8 | ENSMUST00000163584 | 26.2908012 | 33 | 19.8813056 | 21.305638 | 0 | 0 | 0 | 0 | 0 | 0 | 0 | 0 | 0 | 0 | 0 | 1 | 1 | 0 | 2 |
| ENSMUSG00000051224 | Tceanc | ENSMUST00000112188 | 26.3766959 | 36 | 18.4756584 | 19.0742219 | 0 | 0 | 0 | 0 | 0 | 0 | 0 | 0 | 0 | 0 | 0 | 0 | 14 | 2 | 16 |
| ENSMUSG00000079658 | Tceb1; LOC102642819 | ENSMUST00000188641 | 30.6461932 | 33 | 16.8266155 | 19.7696737 | 0 | 0 | 1 | 0 | 0 | 0 | 0 | 0 | 0 | 1 | 0 | 0 | 1 | 0 | 2 |
| ENSMUSG00000053477 | Tcf4 | ENSMUST00000114985 | 30.5310471 | 33 | 16.8803063 | 19.6603962 | 1 | 0 | 0 | 2 | 0 | 0 | 0 | 0 | 2 | 4 | 4 | 1 | 17 | 4 | 31 |
| ENSMUSG00000024985 | Tcf7l2 | ENSMUST00000111656 | 25.9044216 | 33 | 22.1527468 | 18.5350603 | 0 | 0 | 0 | 0 | 0 | 0 | 0 | 0 | 0 | 0 | 2 | 1 | 7 | 0 | 10 |
| ENSMUSG00000027175 | Tcp11l1 | ENSMUST00000111118 | 22.7600963 | 31 | 25.7288045 | 20.0320941 | 0 | 0 | 1 | 0 | 0 | 0 | 0 | 0 | 0 | 1 | 0 | 3 | 2 | 0 | 6 |
| ENSMUSG00000014075 | Tctex1d2 | ENSMUST00000080316 | 30.7017544 | 39 | 12.8654971 | 17.5438596 | 0 | 0 | 0 | 1 | 0 | 0 | 0 | 0 | 1 | 2 | 0 | 0 | 0 | 0 | 2 |
| ENSMUSG00000034674 | Tdg; Tdg-ps | ENSMUST00000151390 | 24.0500241 | 32 | 20.4425204 | 24.001924 | 0 | 0 | 0 | 1 | 0 | 0 | 0 | 0 | 1 | 2 | 4 | 0 | 3 | 0 | 9 |
| ENSMUSG00000035958 | Tdp2 | ENSMUST00000038039 | 31.9848293 | 34 | 14.9178255 | 19.5954488 | 0 | 0 | 0 | 0 | 0 | 0 | 0 | 0 | 0 | 0 | 0 | 0 | 0 | 1 | 1 |
| ENSMUSG00000022019 | Tdrd3 | ENSMUST00000169504 | 28.8513514 | 36 | 17.5337838 | 17.2635135 | 2 | 1 | 0 | 3 | 0 | 0 | 0 | 1 | 2 | 6 | 0 | 1 | 0 | 0 | 10 |
| ENSMUSG00000060985 | Tdrd5 | ENSMUST00000121146 | 20.1438849 | 49 | 10.4316547 | 20.8633094 | 0 | 0 | 0 | 0 | 0 | 0 | 0 | 0 | 0 | 0 | 4 | 0 | 8 | 0 | 12 |
| ENSMUSG00000055320 | Tead1 | ENSMUST00000059768 | 25.3958512 | 30 | 24.1193077 | 20.044188 | 0 | 0 | 0 | 0 | 0 | 0 | 0 | 0 | 0 | 0 | 0 | 36 | 4 | 2 | 42 |
| ENSMUSG00000006386 | Tek | ENSMUST00000073939 | 29.6523517 | 32 | 18.7116564 | 20.0408998 | 1 | 0 | 0 | 0 | 0 | 0 | 0 | 0 | 0 | 0 | 0 | 0 | 0 | 0 | 1 |
| ENSMUSG00000025925 | Terf1 | ENSMUST00000188371 | 30.2061856 | 34 | 15.6701031 | 19.7938144 | 0 | 0 | 0 | 0 | 0 | 0 | 0 | 0 | 0 | 0 | 6 | 0 | 7 | 0 | 13 |
| ENSMUSG00000029552 | Tes | ENSMUST00000076654 | 27.6026743 | 35 | 19.4842407 | 17.9560649 | 0 | 0 | 0 | 0 | 0 | 0 | 0 | 0 | 0 | 0 | 0 | 1 | 0 | 1 | 2 |
| ENSMUSG00000047146 | Tet1 | ENSMUST00000050826 | 27.4428536 | 33 | 19.6181864 | 19.9070585 | 1 | 1 | 0 | 3 | 0 | 0 | 0 | 1 | 2 | 6 | 4 | 0 | 14 | 5 | 31 |
| ENSMUSG00000026049 | Tex30 | ENSMUST00000027215 | 32.8519856 | 40 | 11.5523466 | 15.1624549 | 1 | 0 | 0 | 1 | 0 | 0 | 0 | 0 | 1 | 2 | 0 | 0 | 0 | 0 | 3 |
| ENSMUSG00000090626 | Tex9 | ENSMUST00000085358 | 32.6740205 | 25 | 20.8824648 | 21.186763 | 0 | 0 | 0 | 1 | 0 | 0 | 0 | 0 | 1 | 2 | 0 | 0 | 7 | 0 | 9 |
| ENSMUSG00000003923 | Tfam | ENSMUST00000092430 | 27.7867528 | 30 | 19.450727 | 22.6494346 | 0 | 0 | 0 | 0 | 0 | 0 | 0 | 0 | 0 | 0 | 8 | 1 | 7 | 1 | 17 |
| ENSMUSG00000025927 | Tfap2b | ENSMUST00000027059 | 28.0917475 | 29 | 22.5821707 | 20.5249468 | 1 | 1 | 1 | 2 | 0 | 0 | 0 | 0 | 2 | 5 | 0 | 4 | 2 | 1 | 14 |
| ENSMUSG00000005718 | Tfap4 | ENSMUST00000005862 | 19.6188341 | 34 | 22.309417 | 24.103139 | 0 | 0 | 0 | 0 | 0 | 0 | 0 | 0 | 0 | 0 | 0 | 3 | 5 | 0 | 8 |
| ENSMUSG00000009733 | Tfcp2 | ENSMUST00000009877 | 23.8954574 | 30 | 22.7131301 | 23.2731798 | 0 | 0 | 0 | 0 | 0 | 0 | 0 | 0 | 0 | 0 | 0 | 1 | 7 | 2 | 10 |
| ENSMUSG00000026380 | Tfcp2l1 | ENSMUST00000027629 | 24.359969 | 30 | 22.898888 | 22.6273597 | 0 | 0 | 0 | 0 | 0 | 0 | 0 | 0 | 0 | 0 | 1 | 4 | 6 | 2 | 13 |
| ENSMUSG00000032411 | Tfdp2 | ENSMUST00000034982 | 26.8017621 | 28 | 22.061674 | 22.7488987 | 0 | 0 | 0 | 1 | 0 | 0 | 0 | 0 | 1 | 2 | 0 | 0 | 3 | 1 | 6 |
| ENSMUSG00000022757 | Tfg | ENSMUST00000065515 | 35.3075171 | 33 | 15.9453303 | 16.1731207 | 0 | 0 | 0 | 1 | 0 | 0 | 0 | 0 | 1 | 2 | 0 | 0 | 0 | 0 | 2 |
| ENSMUSG00000027082 | Tfpi | ENSMUST00000111711 | 29.6441397 | 32 | 19.4906954 | 19.0662749 | 1 | 0 | 1 | 2 | 0 | 0 | 0 | 0 | 2 | 5 | 0 | 0 | 0 | 1 | 7 |
| ENSMUSG00000022797 | Tfrc | ENSMUST00000023486 | 28.4784378 | 35 | 16.8022783 | 19.812856 | 0 | 0 | 0 | 0 | 0 | 0 | 0 | 0 | 0 | 0 | 0 | 0 | 3 | 1 | 4 |
| ENSMUSG00000022130 | Tgds | ENSMUST00000022727 | 27.7347115 | 31 | 20.1550388 | 20.6718346 | 0 | 0 | 1 | 0 | 0 | 0 | 0 | 0 | 0 | 1 | 0 | 0 | 0 | 0 | 1 |
| ENSMUSG00000029999 | Tgfa | ENSMUST00000032066 | 25.6376196 | 27 | 24.8671626 | 22.9543039 | 1 | 0 | 0 | 1 | 0 | 0 | 0 | 0 | 1 | 2 | 0 | 1 | 2 | 0 | 6 |
| ENSMUSG00000007613 | Tgfbr1 | ENSMUST00000007757 | 26.6971594 | 34 | 17.9104478 | 20.9436688 | 0 | 0 | 0 | 0 | 0 | 0 | 0 | 0 | 0 | 0 | 0 | 0 | 4 | 0 | 4 |
| ENSMUSG00000029287 | Tgfbr3 | ENSMUST00000031224 | 26.8165526 | 30 | 21.4727924 | 21.277289 | 0 | 0 | 0 | 1 | 0 | 0 | 0 | 0 | 1 | 2 | 0 | 2 | 4 | 2 | 10 |
| ENSMUSG00000056429 | Tgoln2; Tgoln1 | ENSMUST00000070524 | 25.4015791 | 33 | 20.5281786 | 21.2632725 | 1 | 0 | 0 | 0 | 0 | 0 | 0 | 0 | 0 | 0 | 0 | 0 | 4 | 3 | 8 |
| ENSMUSG00000028233 | Tgs1 | ENSMUST00000052712 | 26.889715 | 35 | 18.5254027 | 19.9504337 | 0 | 0 | 0 | 0 | 0 | 0 | 0 | 0 | 0 | 0 | 1 | 0 | 5 | 0 | 6 |
| ENSMUSG00000037214 | Thap1 | ENSMUST00000036807 | 43.2624113 | 33 | 11.3475177 | 12.0567376 | 0 | 0 | 0 | 0 | 0 | 0 | 0 | 0 | 0 | 0 | 0 | 0 | 0 | 1 | 1 |
| ENSMUSG00000020137 | Thap2 | ENSMUST00000020346 | 28.6430224 | 32 | 21.2798766 | 18.4271396 | 0 | 0 | 0 | 0 | 0 | 0 | 0 | 0 | 0 | 0 | 1 | 0 | 5 | 1 | 7 |
| ENSMUSG00000040152 | Thbs1 | ENSMUST00000039559 | 26.28327 | 32 | 21.0551331 | 20.3897338 | 1 | 0 | 0 | 1 | 0 | 0 | 0 | 0 | 1 | 2 | 0 | 6 | 3 | 0 | 12 |
| ENSMUSG00000024287 | Thoc1 | ENSMUST00000025137 | 32.1734475 | 32 | 19.1648822 | 17.0770878 | 0 | 0 | 0 | 0 | 0 | 0 | 0 | 0 | 0 | 0 | 1 | 0 | 0 | 0 | 1 |
| ENSMUSG00000037475 | Thoc2 | ENSMUST00000047037 | 28.7495547 | 37 | 17.4919843 | 16.4232276 | 0 | 0 | 0 | 2 | 0 | 0 | 0 | 0 | 2 | 4 | 0 | 1 | 1 | 0 | 6 |
| ENSMUSG00000053453 | Thoc7 | ENSMUST00000065865 | 28.8617886 | 36 | 17.0731707 | 18.2926829 | 0 | 0 | 0 | 1 | 0 | 0 | 0 | 0 | 1 | 2 | 0 | 0 | 0 | 0 | 2 |
| ENSMUSG00000030942 | Thumpd1 | ENSMUST00000033236 | 23.5406091 | 31 | 19.0989848 | 26.5862944 | 0 | 0 | 0 | 0 | 0 | 0 | 0 | 0 | 0 | 0 | 2 | 0 | 3 | 0 | 5 |
| ENSMUSG00000071337 | Tia1 | ENSMUST00000113712 | 29.3598862 | 32 | 18.3499289 | 19.886202 | 1 | 1 | 0 | 0 | 0 | 0 | 0 | 0 | 0 | 0 | 0 | 0 | 0 | 0 | 2 |
| ENSMUSG00000030846 | Tial1 | ENSMUST00000106226 | 29.4992743 | 35 | 16.1103048 | 19.0856313 | 1 | 0 | 0 | 1 | 0 | 0 | 0 | 0 | 1 | 2 | 0 | 0 | 20 | 0 | 23 |
| ENSMUSG00000046688 | Tifa | ENSMUST00000054483 | 26.5762171 | 34 | 17.7972865 | 21.707901 | 1 | 1 | 0 | 0 | 0 | 0 | 0 | 0 | 0 | 0 | 0 | 0 | 5 | 0 | 7 |
| ENSMUSG00000062580 | Timm17a | ENSMUST00000081104 | 29.7986577 | 28 | 21.1185682 | 21.163311 | 1 | 0 | 1 | 2 | 0 | 0 | 0 | 0 | 2 | 5 | 0 | 0 | 1 | 0 | 7 |
| ENSMUSG00000031158 | Timm17b | ENSMUST00000033498 | 25.4084507 | 26 | 23.6056338 | 24.9014085 | 0 | 0 | 0 | 0 | 0 | 0 | 0 | 0 | 0 | 0 | 0 | 2 | 3 | 0 | 5 |
| ENSMUSG00000024645 | Timm21 | ENSMUST00000025547 | 32.8611898 | 32 | 18.4135977 | 16.9971671 | 0 | 0 | 0 | 1 | 0 | 0 | 0 | 0 | 1 | 2 | 0 | 0 | 0 | 0 | 2 |
| ENSMUSG00000020044 | Timp3 | ENSMUST00000020234 | 27.7469478 | 30 | 21.281909 | 21.3651498 | 0 | 0 | 0 | 1 | 0 | 0 | 0 | 0 | 1 | 2 | 0 | 1 | 0 | 1 | 4 |
| ENSMUSG00000040843 | Tiprl | ENSMUST00000043235 | 27.4464389 | 30 | 19.3398958 | 23.2194557 | 1 | 0 | 0 | 0 | 0 | 0 | 0 | 0 | 0 | 0 | 1 | 0 | 0 | 3 | 5 |
| ENSMUSG00000024642 | Tle4 | ENSMUST00000052011 | 28.0385078 | 36 | 16.967509 | 18.7725632 | 1 | 0 | 1 | 1 | 0 | 0 | 0 | 0 | 1 | 3 | 0 | 0 | 2 | 0 | 6 |
| ENSMUSG00000028563 | Tm2d1 | ENSMUST00000125034 | 30.0215983 | 39 | 16.4146868 | 14.9028078 | 0 | 1 | 0 | 0 | 0 | 0 | 0 | 0 | 0 | 0 | 0 | 0 | 0 | 0 | 1 |
| ENSMUSG00000038623 | Tm6sf1 | ENSMUST00000126334 | 31.1520109 | 33 | 19.0865712 | 16.7689162 | 0 | 0 | 1 | 1 | 0 | 0 | 0 | 0 | 1 | 3 | 0 | 0 | 0 | 0 | 3 |
| ENSMUSG00000040234 | Tm7sf3 | ENSMUST00000037709 | 27.1499645 | 33 | 17.4840085 | 21.8905473 | 1 | 0 | 0 | 0 | 0 | 0 | 0 | 0 | 0 | 0 | 0 | 0 | 0 | 0 | 1 |
| ENSMUSG00000025016 | Tm9sf3 | ENSMUST00000025989 | 28.6761229 | 33 | 17.2576832 | 21.0638298 | 1 | 1 | 0 | 0 | 0 | 0 | 0 | 0 | 0 | 0 | 0 | 2 | 0 | 7 | 11 |
| ENSMUSG00000068040 | Tm9sf4 | ENSMUST00000089027 | 21.9298246 | 27 | 27.0175439 | 24.0350877 | 0 | 0 | 0 | 0 | 0 | 0 | 0 | 0 | 0 | 0 | 0 | 4 | 3 | 3 | 10 |
| ENSMUSG00000025591 | Tma16 | ENSMUST00000026681 | 30.7971014 | 29 | 17.3913043 | 22.4637681 | 1 | 0 | 0 | 1 | 0 | 0 | 0 | 0 | 1 | 2 | 0 | 0 | 0 | 0 | 3 |
| ENSMUSG00000042246 | Tmc7 | ENSMUST00000044195 | 25.2833479 | 33 | 19.8343505 | 21.8395815 | 2 | 0 | 1 | 1 | 0 | 0 | 0 | 0 | 1 | 3 | 0 | 0 | 6 | 1 | 12 |
| ENSMUSG00000052428 | Tmco1 | ENSMUST00000097473 | 27.1476613 | 28 | 22.5924794 | 22.7147661 | 0 | 0 | 0 | 0 | 0 | 0 | 0 | 0 | 0 | 0 | 0 | 2 | 0 | 0 | 2 |
| ENSMUSG00000038497 | Tmco3 | ENSMUST00000045229 | 27.2195122 | 33 | 18.3414634 | 21.1219512 | 3 | 1 | 0 | 0 | 0 | 0 | 0 | 0 | 0 | 0 | 0 | 0 | 0 | 7 | 11 |
| ENSMUSG00000021248 | Tmed10 | ENSMUST00000040766 | 25.6929638 | 31 | 21.9260839 | 21.0376688 | 0 | 0 | 0 | 0 | 0 | 0 | 0 | 0 | 0 | 0 | 0 | 1 | 0 | 0 | 1 |

| | | | | | | | | | | | | | | | | | | | | | |
|---|---|---|---|---|---|---|---|---|---|---|---|---|---|---|---|---|---|---|---|---|---|
| ENSMUSG00000029390 | Tmed2; Gm21540 | ENSMUST00000060226 | 25.2113759 | 35 | 18.3704842 | 20.9838586 | 0 | 0 | 0 | 2 | 0 | 0 | 0 | 0 | 2 | 4 | 2 | 0 | 0 | 0 | 6 |
| ENSMUSG00000004394 | Tmed4 | ENSMUST00000004508 | 25.8190328 | 30 | 22.074883 | 22.3088924 | 1 | 0 | 0 | 0 | 0 | 0 | 0 | 0 | 0 | 0 | 0 | 0 | 0 | 0 | 1 |
| ENSMUSG00000063406 | Tmed5 | ENSMUST00000002837 | 36.7536033 | 32 | 15.6829101 | 15.0995196 | 0 | 0 | 0 | 1 | 0 | 0 | 0 | 0 | 1 | 2 | 0 | 0 | 0 | 0 | 2 |
| ENSMUSG00000028347 | Tmeff1 | ENSMUST00000030032 | 31.7399618 | 36 | 14.2447419 | 18.0688337 | 1 | 0 | 0 | 0 | 0 | 0 | 0 | 0 | 0 | 0 | 0 | 0 | 0 | 0 | 1 |
| ENSMUSG00000029571 | Tmem106b | ENSMUST00000031556 | 29.6378717 | 33 | 18.5312563 | 18.3896419 | 1 | 0 | 0 | 0 | 0 | 0 | 0 | 0 | 0 | 0 | 1 | 1 | 1 | 1 | 5 |
| ENSMUSG00000042757 | Tmem108 | ENSMUST00000049452 | 26.4294049 | 31 | 23.1621937 | 19.6032672 | 0 | 0 | 0 | 1 | 0 | 0 | 0 | 0 | 1 | 2 | 0 | 0 | 0 | 0 | 2 |
| ENSMUSG00000050912 | Tmem123 | ENSMUST00000154371 | 29.400607 | 31 | 20.4097117 | 19.2336874 | 0 | 0 | 0 | 0 | 0 | 0 | 0 | 0 | 0 | 0 | 1 | 1 | 0 | 0 | 2 |
| ENSMUSG00000025933 | Tmem14a | ENSMUST00000027064 | 27.1157168 | 34 | 18.3074266 | 20.3799655 | 0 | 1 | 0 | 1 | 0 | 0 | 0 | 0 | 1 | 2 | 0 | 0 | 0 | 0 | 3 |
| ENSMUSG00000054871 | Tmem158 | ENSMUST00000068140 | 22.9665072 | 30 | 21.5311005 | 25.9968102 | 0 | 0 | 0 | 1 | 0 | 0 | 0 | 0 | 1 | 2 | 1 | 0 | 2 | 1 | 6 |
| ENSMUSG00000012422 | Tmem167 | ENSMUST00000161568 | 31.4260113 | 30 | 18.5867896 | 19.6364567 | 0 | 0 | 0 | 1 | 0 | 0 | 0 | 0 | 1 | 2 | 0 | 0 | 2 | 4 | 8 |
| ENSMUSG00000068732 | Tmem167b | ENSMUST00000090546 | 27.6396249 | 29 | 19.8532409 | 23.3999185 | 0 | 0 | 0 | 0 | 0 | 0 | 0 | 0 | 0 | 0 | 0 | 0 | 8 | 0 | 8 |
| ENSMUSG00000029569 | Tmem168 | ENSMUST00000031554 | 30.047619 | 36 | 15.952381 | 18.0952381 | 0 | 0 | 0 | 0 | 0 | 0 | 0 | 0 | 0 | 0 | 0 | 0 | 0 | 3 | 3 |
| ENSMUSG00000031953 | Tmem170 | ENSMUST00000034431 | 24.9318801 | 32 | 21.866485 | 20.7084469 | 0 | 0 | 0 | 0 | 0 | 0 | 0 | 0 | 0 | 0 | 4 | 0 | 7 | 0 | 11 |
| ENSMUSG00000087370 | Tmem170b | ENSMUST00000129449 | 26.859932 | 33 | 19.0208549 | 20.9584381 | 0 | 0 | 2 | 0 | 0 | 0 | 0 | 0 | 0 | 2 | 0 | 4 | 16 | 1 | 23 |
| ENSMUSG00000043061 | Tmem18 | ENSMUST00000057151 | 26.0746148 | 31 | 20.2351987 | 22.7493917 | 0 | 0 | 0 | 0 | 0 | 0 | 0 | 0 | 0 | 0 | 0 | 0 | 2 | 0 | 2 |
| ENSMUSG00000042305 | Tmem183a | ENSMUST00000049470 | 26.7259414 | 33 | 18.6192469 | 21.9665272 | 0 | 0 | 0 | 0 | 0 | 0 | 0 | 0 | 0 | 0 | 0 | 1 | 0 | 0 | 1 |
| ENSMUSG00000090213 | Tmem189 | ENSMUST00000006587 | 23.1245699 | 26 | 26.9098417 | 24.4322092 | 1 | 1 | 0 | 1 | 0 | 0 | 0 | 0 | 1 | 2 | 0 | 1 | 0 | 0 | 5 |
| ENSMUSG00000040195 | Tmem194 | ENSMUST00000048099 | 23.8402062 | 31 | 22.4226804 | 22.7448454 | 0 | 1 | 0 | 0 | 0 | 0 | 0 | 0 | 0 | 0 | 0 | 0 | 0 | 1 | 2 |
| ENSMUSG00000043015 | Tmem194b | ENSMUST00000188507 | 25.7744241 | 30 | 21.4853058 | 22.5178713 | 0 | 0 | 0 | 0 | 0 | 0 | 0 | 0 | 0 | 0 | 2 | 1 | 7 | 2 | 12 |
| ENSMUSG00000048004 | Tmem196 | ENSMUST00000058644 | 34.6031746 | 34 | 16.031746 | 15.5555556 | 0 | 1 | 0 | 2 | 0 | 0 | 0 | 1 | 1 | 4 | 0 | 0 | 0 | 0 | 5 |
| ENSMUSG00000024754 | Tmem2 | ENSMUST00000025663 | 25.6578947 | 32 | 18.6234818 | 23.6842105 | 0 | 0 | 2 | 0 | 0 | 0 | 0 | 0 | 0 | 2 | 1 | 0 | 3 | 0 | 6 |
| ENSMUSG00000029782 | Tmem209 | ENSMUST00000151187 | 24.8347107 | 28 | 26.6942149 | 20.4958678 | 0 | 0 | 0 | 0 | 0 | 0 | 0 | 0 | 0 | 0 | 1 | 0 | 0 | 0 | 1 |
| ENSMUSG00000046593 | Tmem215 | ENSMUST00000049655 | 30.5599369 | 33 | 16.3249211 | 19.7160883 | 1 | 0 | 0 | 0 | 0 | 0 | 0 | 0 | 0 | 0 | 0 | 1 | 11 | 0 | 13 |
| ENSMUSG00000027341 | Tmem230 | ENSMUST00000028816 | 27.5653924 | 32 | 21.1267606 | 18.8128773 | 0 | 0 | 0 | 0 | 0 | 0 | 0 | 0 | 0 | 0 | 0 | 0 | 0 | 1 | 1 |
| ENSMUSG00000049411 | Tmem241 | ENSMUST00000050228 | 24.0262245 | 31 | 21.673737 | 23.4091786 | 2 | 1 | 0 | 0 | 0 | 0 | 0 | 0 | 0 | 0 | 0 | 0 | 0 | 0 | 3 |
| ENSMUSG00000055296 | Tmem245 | ENSMUST00000068792 | 27.0189132 | 31 | 22.1104773 | 19.6712699 | 7 | 3 | 1 | 8 | 0 | 1 | 2 | 2 | 4 | 18 | 8 | 31 | 10 | 2 | 79 |
| ENSMUSG00000046675 | Tmem251 | ENSMUST00000057416 | 30.3529412 | 30 | 20.0392157 | 19.2941176 | 0 | 0 | 0 | 2 | 0 | 0 | 0 | 0 | 2 | 4 | 0 | 0 | 0 | 0 | 4 |
| ENSMUSG00000036339 | Tmem260 | ENSMUST00000111735 | 27.8710544 | 29 | 21.0879785 | 21.8603089 | 0 | 1 | 0 | 0 | 0 | 0 | 0 | 0 | 0 | 0 | 0 | 0 | 0 | 0 | 1 |
| ENSMUSG00000060935 | Tmem263 | ENSMUST00000095383 | 30.3963315 | 35 | 15.4602031 | 19.0959712 | 0 | 0 | 0 | 1 | 0 | 0 | 0 | 0 | 1 | 2 | 1 | 0 | 19 | 0 | 22 |
| ENSMUSG00000041353 | Tmem29 | ENSMUST00000163233 | 27.1794872 | 36 | 16.5384615 | 20.7692308 | 0 | 0 | 0 | 0 | 0 | 0 | 0 | 0 | 0 | 0 | 0 | 0 | 2 | 0 | 2 |
| ENSMUSG00000037720 | Tmem33 | ENSMUST00000037918 | 29.2971734 | 35 | 16.5011459 | 19.5760122 | 1 | 0 | 0 | 1 | 0 | 0 | 0 | 0 | 1 | 2 | 0 | 5 | 0 | 3 | 11 |
| ENSMUSG00000028420 | Tmem38b | ENSMUST00000030127 | 27.253778 | 36 | 16.6232413 | 20.0104221 | 1 | 0 | 1 | 0 | 0 | 0 | 0 | 0 | 0 | 1 | 0 | 0 | 4 | 0 | 6 |
| ENSMUSG00000053730 | Tmem39b | ENSMUST00000125445 | 21.4960058 | 25 | 29.7748729 | 23.7472767 | 0 | 0 | 0 | 0 | 0 | 0 | 0 | 0 | 0 | 0 | 0 | 1 | 0 | 0 | 1 |
| ENSMUSG00000047554 | Tmem41b | ENSMUST00000094097 | 28.7861272 | 33 | 19.460501 | 18.8439306 | 0 | 0 | 0 | 0 | 0 | 0 | 0 | 0 | 0 | 0 | 0 | 1 | 0 | 0 | 1 |
| ENSMUSG00000034620 | Tmem5 | ENSMUST00000140299 | 29.7023433 | 31 | 19.2843572 | 20.4243192 | 0 | 0 | 0 | 2 | 0 | 0 | 0 | 0 | 2 | 4 | 3 | 0 | 0 | 0 | 7 |
| ENSMUSG00000028221 | Tmem55a | ENSMUST00000029875 | 28.342246 | 37 | 17.1886937 | 17.723453 | 1 | 0 | 0 | 0 | 0 | 0 | 0 | 0 | 0 | 0 | 0 | 0 | 8 | 0 | 9 |
| ENSMUSG00000035953 | Tmem55b | ENSMUST00000160835 | 26.754386 | 26 | 22.9714912 | 24.122807 | 0 | 0 | 0 | 0 | 0 | 0 | 0 | 0 | 0 | 0 | 0 | 1 | 0 | 0 | 1 |
| ENSMUSG00000028132 | Tmem56 | ENSMUST00000029777 | 27.974339 | 35 | 17.6905132 | 19.226283 | 1 | 0 | 0 | 0 | 0 | 0 | 0 | 0 | 0 | 0 | 0 | 0 | 2 | 2 | 5 |
| ENSMUSG00000043252 | Tmem64 | ENSMUST00000062684 | 27.0937408 | 35 | 18.7775492 | 19.1007934 | 3 | 0 | 0 | 1 | 0 | 0 | 0 | 0 | 1 | 2 | 0 | 1 | 0 | 1 | 7 |
| ENSMUSG00000062373 | Tmem65 | ENSMUST00000072113 | 27.3484848 | 36 | 17.6136364 | 19.3560606 | 0 | 0 | 1 | 0 | 0 | 0 | 0 | 0 | 0 | 1 | 1 | 2 | 0 | 1 | 5 |
| ENSMUSG00000049488 | Tmem67 | ENSMUST00000131145 | 27.6880797 | 30 | 21.0924081 | 21.4015802 | 2 | 0 | 0 | 1 | 0 | 0 | 0 | 0 | 1 | 2 | 0 | 0 | 0 | 0 | 4 |
| ENSMUSG00000028232 | Tmem68 | ENSMUST00000154922 | 29.5387635 | 34 | 16.5358194 | 19.4798822 | 1 | 0 | 0 | 0 | 0 | 0 | 0 | 0 | 0 | 0 | 0 | 0 | 0 | 0 | 1 |
| ENSMUSG00000055900 | Tmem69 | ENSMUST00000106478 | 25.4221388 | 34 | 20.0750469 | 20.9193246 | 0 | 0 | 0 | 0 | 0 | 0 | 0 | 0 | 0 | 0 | 0 | 0 | 4 | 0 | 4 |
| ENSMUSG00000025940 | Tmem70 | ENSMUST00000065373 | 28.9411765 | 34 | 15.6470588 | 21.5294118 | 1 | 1 | 0 | 0 | 0 | 0 | 0 | 0 | 0 | 0 | 2 | 0 | 3 | 0 | 7 |
| ENSMUSG00000025505 | Tmem80 | ENSMUST00000026578 | 24.954792 | 28 | 22.3327306 | 24.4122966 | 0 | 0 | 0 | 0 | 0 | 0 | 0 | 0 | 0 | 0 | 0 | 0 | 0 | 1 | 1 |
| ENSMUSG00000033808 | Tmem87a | ENSMUST00000090042 | 26.4135021 | 33 | 18.649789 | 22.0253165 | 2 | 1 | 1 | 1 | 0 | 0 | 0 | 0 | 1 | 3 | 0 | 0 | 25 | 1 | 32 |
| ENSMUSG00000014353 | Tmem87b | ENSMUST00000110325 | 26.5935572 | 31 | 19.8766278 | 22.5154215 | 0 | 0 | 0 | 1 | 0 | 0 | 0 | 0 | 1 | 2 | 3 | 1 | 3 | 0 | 9 |
| ENSMUSG00000026411 | Tmem9 | ENSMUST00000063719 | 21.9251337 | 26 | 26.2923351 | 26.0249554 | 0 | 0 | 0 | 0 | 0 | 0 | 0 | 0 | 0 | 0 | 0 | 1 | 0 | 0 | 1 |
| ENSMUSG00000032186 | Tmod2 | ENSMUST00000098552 | 28.1648559 | 30 | 20.6896552 | 21.0793576 | 0 | 0 | 0 | 2 | 0 | 0 | 0 | 0 | 2 | 4 | 1 | 22 | 7 | 0 | 34 |
| ENSMUSG00000058587 | Tmod3 | ENSMUST00000072232 | 30 | 32 | 17.5 | 20.862069 | 0 | 0 | 0 | 0 | 0 | 0 | 0 | 0 | 0 | 0 | 0 | 0 | 1 | 0 | 1 |
| ENSMUSG00000019961 | Tmpo | ENSMUST00000072239 | 29.0813648 | 35 | 16.4829396 | 19.1076115 | 0 | 0 | 0 | 0 | 0 | 0 | 0 | 0 | 0 | 0 | 1 | 0 | 1 | 0 | 2 |
| ENSMUSG00000049775 | Tmsb4x | ENSMUST00000112172 | 30.6372549 | 29 | 17.8921569 | 22.0588235 | 0 | 0 | 0 | 0 | 0 | 0 | 0 | 0 | 0 | 0 | 0 | 1 | 0 | 0 | 1 |
| ENSMUSG00000030306 | Tmtc1 | ENSMUST00000060095 | 28.1506339 | 30 | 20.5630127 | 21.6815809 | 1 | 0 | 0 | 3 | 0 | 0 | 0 | 0 | 3 | 6 | 3 | 5 | 1 | 0 | 16 |
| ENSMUSG00000036676 | Tmtc3 | ENSMUST00000058154 | 31.6484969 | 35 | 18.004625 | 14.8992402 | 3 | 1 | 0 | 2 | 0 | 0 | 0 | 1 | 1 | 4 | 0 | 0 | 0 | 0 | 8 |
| ENSMUSG00000041594 | Tmtc4 | ENSMUST00000143189 | 28.7616511 | 31 | 19.4407457 | 20.9498447 | 2 | 0 | 0 | 0 | 0 | 0 | 0 | 0 | 0 | 0 | 0 | 0 | 0 | 0 | 2 |
| ENSMUSG00000021072 | Tmx1 | ENSMUST00000021471 | 27.4946921 | 36 | 16.507431 | 20.2760085 | 0 | 0 | 1 | 1 | 0 | 0 | 0 | 0 | 1 | 3 | 0 | 0 | 0 | 0 | 3 |
| ENSMUSG00000024614 | Tmx3 | ENSMUST00000025515 | 31.8788668 | 37 | 13.8391403 | 17.3559101 | 0 | 0 | 0 | 2 | 0 | 0 | 0 | 0 | 2 | 4 | 0 | 0 | 1 | 0 | 5 |
| ENSMUSG00000034723 | Tmx4 | ENSMUST00000038228 | 27.2580645 | 33 | 19.1705069 | 20.1382488 | 1 | 0 | 0 | 0 | 0 | 0 | 0 | 0 | 0 | 0 | 1 | 6 | 6 | 0 | 14 |
| ENSMUSG00000053475 | Tnfaip6 | ENSMUST00000065927 | 26.7132867 | 42 | 15.5244755 | 15.3846154 | 0 | 0 | 1 | 0 | 0 | 0 | 0 | 0 | 0 | 1 | 0 | 0 | 7 | 3 | 11 |
| ENSMUSG00000074345 | Tnfaip8l3 | ENSMUST00000098760 | 26.4125255 | 33 | 20.4220558 | 19.7413206 | 1 | 0 | 0 | 2 | 0 | 0 | 0 | 0 | 2 | 4 | 1 | 3 | 0 | 2 | 11 |
| ENSMUSG00000060548 | Tnfrsf19 | ENSMUST00000111234 | 27.0682425 | 29 | 21.3114754 | 22.722074 | 0 | 0 | 0 | 0 | 0 | 0 | 0 | 0 | 0 | 0 | 2 | 0 | 0 | 0 | 2 |
| ENSMUSG00000023915 | Tnfrsf21 | ENSMUST00000024708 | 22.6973684 | 38 | 21.7927632 | 17.6809211 | 0 | 0 | 0 | 0 | 0 | 0 | 0 | 0 | 0 | 0 | 0 | 17 | 1 | 3 | 21 |
| ENSMUSG00000035678 | Tnfsf9 | ENSMUST00000039490 | 29.6442688 | 36 | 16.6007905 | 17.3913043 | 1 | 0 | 1 | 1 | 0 | 0 | 0 | 0 | 1 | 3 | 0 | 0 | 0 | 0 | 4 |
| ENSMUSG00000018752 | Tnfsfm13 | ENSMUST00000180587 | 26.443769 | 27 | 25.3799392 | 20.9726444 | 0 | 0 | 0 | 0 | 0 | 0 | 0 | 0 | 0 | 0 | 0 | 1 | 0 | 0 | 1 |
| ENSMUSG00000031529 | Tnks | ENSMUST00000033929 | 26.7915309 | 30 | 21.7833876 | 21.5390879 | 0 | 0 | 0 | 0 | 0 | 0 | 0 | 0 | 0 | 0 | 0 | 3 | 0 | 0 | 3 |
| ENSMUSG00000009470 | Tnpo1 | ENSMUST00000109399 | 27.7452546 | 35 | 17.6157531 | 19.4074863 | 0 | 0 | 0 | 0 | 0 | 0 | 0 | 0 | 0 | 0 | 6 | 2 | 11 | 0 | 19 |
| ENSMUSG00000047888 | Tnrc6b | ENSMUST00000067689 | 25.6330749 | 31 | 21.4814815 | 22.1533161 | 0 | 0 | 0 | 0 | 0 | 0 | 0 | 0 | 0 | 0 | 1 | 28 | 14 | 4 | 47 |
| ENSMUSG00000037573 | Tob1 | ENSMUST00000041589 | 33.767927 | 39 | 11.7340287 | 15.3846154 | 0 | 0 | 0 | 1 | 0 | 0 | 0 | 0 | 1 | 2 | 0 | 0 | 1 | 6 | 9 |
| ENSMUSG00000025139 | Tollip | ENSMUST00000001950 | 23.4195402 | 30 | 22.5933908 | 24.3893678 | 1 | 0 | 0 | 0 | 0 | 0 | 0 | 0 | 0 | 0 | 1 | 0 | 0 | 0 | 2 |
| ENSMUSG00000020541 | Tom1l1 | ENSMUST00000020849 | 31.1314125 | 30 | 20.0281096 | 18.4820801 | 0 | 0 | 0 | 0 | 0 | 0 | 0 | 0 | 0 | 0 | 1 | 0 | 1 | 3 | 5 |
| ENSMUSG00000093904 | Tomm20 | ENSMUST00000179857 | 25.3504941 | 31 | 20.6389336 | 23.0751551 | 3 | 1 | 1 | 0 | 0 | 0 | 0 | 0 | 0 | 1 | 1 | 2 | 0 | 0 | 8 |
| ENSMUSG00000028998 | Tomm7 | ENSMUST00000030851 | 26.5822785 | 33 | 16.570771 | 23.4752589 | 0 | 0 | 0 | 0 | 0 | 0 | 0 | 0 | 0 | 0 | 0 | 1 | 0 | 0 | 1 |
| ENSMUSG00000020914 | Top2a | ENSMUST00000068031 | 26.4187867 | 31 | 19.9608611 | 22.7005871 | 0 | 0 | 1 | 1 | 0 | 0 | 0 | 0 | 1 | 3 | 0 | 0 | 0 | 0 | 3 |
| ENSMUSG00000026849 | Tor1a | ENSMUST00000028200 | 24.0625 | 30 | 21.5625 | 24.6875 | 0 | 0 | 0 | 1 | 0 | 0 | 0 | 0 | 1 | 2 | 0 | 0 | 0 | 0 | 2 |
| ENSMUSG00000026466 | Tor1aip1 | ENSMUST00000169241 | 28.2029235 | 31 | 20.9372313 | 19.6044712 | 1 | 0 | 0 | 1 | 0 | 0 | 0 | 0 | 1 | 2 | 0 | 2 | 0 | 0 | 5 |
| ENSMUSG00000050565 | Tor1aip2 | ENSMUST00000060404 | 26.9174258 | 32 | 20.0117543 | 21.3635028 | 0 | 0 | 0 | 0 | 0 | 0 | 0 | 0 | 0 | 0 | 2 | 1 | 5 | 1 | 9 |
| ENSMUSG00000041272 | Tox | ENSMUST00000039987 | 30.5984556 | 32 | 19.4015444 | 18.4362934 | 0 | 0 | 0 | 1 | 0 | 0 | 0 | 0 | 1 | 2 | 0 | 0 | 0 | 0 | 2 |
| ENSMUSG00000043668 | Tox3 | ENSMUST00000109621 | 33.2252836 | 33 | 15.9643436 | 18.0713128 | 2 | 1 | 1 | 0 | 0 | 0 | 0 | 0 | 0 | 1 | 0 | 0 | 0 | 1 | 5 |
| ENSMUSG00000000296 | Tpd52l1 | ENSMUST00000000305 | 27.3318872 | 33 | 21.0412148 | 18.4381779 | 0 | 1 | 0 | 0 | 0 | 0 | 0 | 0 | 0 | 0 | 0 | 0 | 0 | 0 | 1 |
| ENSMUSG00000029735 | Tpk1 | ENSMUST00000067888 | 30.3761931 | 30 | 20.0449186 | 19.0903987 | 0 | 0 | 0 | 0 | 0 | 0 | 0 | 0 | 0 | 0 | 0 | 0 | 0 | 1 | 1 |
| ENSMUSG00000032366 | Tpm1 | ENSMUST00000113690 | 26.2769879 | 29 | 22.2222222 | 22.538178 | 0 | 0 | 0 | 0 | 0 | 0 | 0 | 0 | 0 | 0 | 0 | 3 | 2 | 0 | 5 |
| ENSMUSG00000031799 | Tpm4 | ENSMUST00000003575 | 27.9210926 | 31 | 21.7754173 | 19.1957511 | 0 | 0 | 0 | 0 | 0 | 0 | 0 | 0 | 0 | 0 | 0 | 1 | 0 | 0 | 1 |
| ENSMUSG00000041763 | Tpp2 | ENSMUST00000190207 | 31.3854235 | 35 | 15.3644123 | 18.5817466 | 1 | 0 | 0 | 0 | 0 | 0 | 0 | 0 | 0 | 0 | 0 | 0 | 0 | 4 | 5 |
| ENSMUSG00000054226 | Tprkb | ENSMUST00000113753 | 26.371308 | 26 | 22.9254571 | 24.6835443 | 0 | 0 | 0 | 0 | 0 | 0 | 0 | 0 | 0 | 0 | 0 | 1 | 0 | 0 | 1 |
| ENSMUSG00000034118 | Tpst1 | ENSMUST00000040721 | 25 | 29 | 23.5294118 | 22.2426471 | 0 | 0 | 0 | 1 | 0 | 0 | 0 | 0 | 1 | 2 | 0 | 0 | 0 | 0 | 2 |
| ENSMUSG00000034292 | Traf3ip1 | ENSMUST00000047242 | 23.6220472 | 33 | 21.3254593 | 22.4409449 | 0 | 0 | 0 | 0 | 0 | 0 | 0 | 0 | 0 | 0 | 2 | 0 | 5 | 0 | 7 |
| ENSMUSG00000025935 | Tram1 | ENSMUST00000027068 | 27.0153846 | 38 | 17.9692308 | 17.4769231 | 0 | 0 | 0 | 0 | 0 | 0 | 0 | 0 | 0 | 0 | 0 | 0 | 10 | 0 | 10 |
| ENSMUSG00000044528 | Tram1l1 | ENSMUST00000058994 | 30.8255388 | 33 | 18.2594551 | 18.3407889 | 0 | 0 | 0 | 1 | 0 | 0 | 0 | 0 | 1 | 2 | 0 | 0 | 0 | 0 | 2 |
| ENSMUSG00000021711 | Trappc13 | ENSMUST00000144060 | 29.9145299 | 33 | 19.4067371 | 18.049271 | 0 | 0 | 1 | 0 | 0 | 0 | 0 | 0 | 0 | 1 | 0 | 0 | 7 | 0 | 8 |
| ENSMUSG00000079317 | Trappc2 | ENSMUST00000112192 | 29.9395161 | 35 | 17.5403226 | 17.641129 | 0 | 0 | 0 | 1 | 0 | 0 | 0 | 0 | 1 | 2 | 0 | 0 | 0 | 0 | 2 |
| ENSMUSG00000033382 | Trappc8 | ENSMUST00000025177 | 32.2391559 | 36 | 16.7643611 | 15.2403283 | 0 | 0 | 0 | 1 | 0 | 0 | 0 | 0 | 1 | 2 | 0 | 0 | 0 | 0 | 2 |
| ENSMUSG00000026723 | Trdmt1 | ENSMUST00000124488 | 29.8322684 | 31 | 19.0894569 | 19.8881789 | 1 | 0 | 0 | 2 | 0 | 0 | 0 | 0 | 2 | 4 | 4 | 0 | 3 | 0 | 12 |

| | | | | | | | | | | | | | | | | | | | | | |
|---|---|---|---|---|---|---|---|---|---|---|---|---|---|---|---|---|---|---|---|---|---|
| ENSMUSG00000038760 | Trhr | ENSMUST00000038856 | 34.3558282 | 32 | 15.9100204 | 17.9141104 | 1 | 1 | 1 | 0 | 0 | 0 | 0 | 0 | 0 | 1 | 0 | 0 | 0 | 0 | 3 |
| ENSMUSG00000057143 | Trim12c | ENSMUST00000059037 | 33.0529441 | 29 | 15.8832261 | 21.7219198 | 0 | 0 | 0 | 0 | 0 | 0 | 0 | 0 | 0 | 0 | 0 | 0 | 2 | 0 | 2 |
| ENSMUSG00000027993 | Trim2 | ENSMUST00000107692 | 29.5191069 | 31 | 18.484328 | 20.5667669 | 0 | 0 | 1 | 0 | 0 | 0 | 0 | 0 | 0 | 1 | 1 | 0 | 4 | 0 | 6 |
| ENSMUSG00000021712 | Trim23 | ENSMUST00000069187 | 32.4668705 | 36 | 13.8634047 | 17.1763507 | 0 | 0 | 0 | 2 | 0 | 0 | 0 | 0 | 2 | 4 | 0 | 0 | 0 | 2 | 6 |
| ENSMUSG00000033014 | Trim33 | ENSMUST00000029444 | 29.3670366 | 34 | 18.1463214 | 18.5984381 | 0 | 0 | 0 | 0 | 0 | 0 | 0 | 0 | 0 | 0 | 0 | 0 | 2 | 3 | 5 |
| ENSMUSG00000033949 | Trim36 | ENSMUST00000037011 | 27.8540976 | 34 | 17.6219801 | 20.0852676 | 1 | 0 | 0 | 1 | 0 | 0 | 0 | 0 | 1 | 2 | 0 | 0 | 3 | 0 | 6 |
| ENSMUSG00000032386 | Trip4 | ENSMUST00000179395 | 29.950827 | 31 | 18.2610639 | 21.1667412 | 0 | 0 | 1 | 2 | 0 | 0 | 0 | 0 | 2 | 5 | 0 | 1 | 0 | 1 | 7 |
| ENSMUSG00000055963 | Triqk | ENSMUST00000183345 | 32.9818781 | 33 | 16.4744646 | 17.660626 | 0 | 0 | 0 | 1 | 0 | 0 | 0 | 0 | 1 | 2 | 0 | 0 | 0 | 0 | 2 |
| ENSMUSG00000004127 | Trmt10a | ENSMUST00000040321 | 25.0521195 | 34 | 20.7435719 | 19.8054204 | 0 | 0 | 0 | 0 | 0 | 0 | 0 | 0 | 0 | 0 | 0 | 26 | 8 | 0 | 34 |
| ENSMUSG00000044763 | Trmt10c | ENSMUST00000059052 | 28.7932251 | 32 | 18.4897671 | 20.2540579 | 1 | 0 | 0 | 1 | 0 | 0 | 0 | 0 | 1 | 2 | 0 | 1 | 0 | 0 | 4 |
| ENSMUSG00000033439 | Trmt13 | ENSMUST00000183638 | 32.4977619 | 27 | 16.7412713 | 23.7242614 | 0 | 0 | 0 | 1 | 0 | 0 | 0 | 0 | 1 | 2 | 0 | 0 | 0 | 0 | 2 |
| ENSMUSG00000067369 | Trmt2b | ENSMUST00000128115 | 28.0264694 | 28 | 20.5916699 | 23.4332425 | 0 | 0 | 0 | 0 | 0 | 0 | 0 | 0 | 0 | 0 | 0 | 1 | 3 | 0 | 4 |
| ENSMUSG00000037376 | Trmt6 | ENSMUST00000039554 | 25.8649094 | 33 | 18.5337727 | 22.1581549 | 0 | 0 | 0 | 0 | 0 | 0 | 0 | 0 | 0 | 0 | 0 | 1 | 0 | 0 | 1 |
| ENSMUSG00000013736 | Trnt1 | ENSMUST00000113247 | 27.3569024 | 35 | 18.9393939 | 18.9393939 | 0 | 0 | 0 | 0 | 0 | 0 | 0 | 0 | 0 | 0 | 0 | 0 | 0 | 2 | 2 |
| ENSMUSG00000018199 | Trove2 | ENSMUST00000159879 | 31.8469505 | 33 | 16.7289988 | 18.1962025 | 0 | 0 | 0 | 1 | 0 | 0 | 0 | 0 | 1 | 2 | 2 | 0 | 7 | 0 | 11 |
| ENSMUSG00000028211 | Trp53inp1 | ENSMUST00000029865 | 28.280543 | 35 | 17.1719457 | 19.3665158 | 1 | 1 | 0 | 0 | 0 | 0 | 0 | 0 | 0 | 0 | 1 | 1 | 3 | 2 | 9 |
| ENSMUSG00000038375 | Trp53inp2 | ENSMUST00000043237 | 22.0078483 | 29 | 26.3570961 | 22.5310661 | 0 | 0 | 0 | 0 | 0 | 0 | 0 | 0 | 0 | 0 | 0 | 2 | 0 | 1 | 3 |
| ENSMUSG00000027365 | Trpm7 | ENSMUST00000103224 | 29.5003965 | 33 | 17.6843775 | 19.9841396 | 0 | 0 | 0 | 0 | 0 | 0 | 0 | 0 | 0 | 0 | 0 | 4 | 0 | 0 | 4 |
| ENSMUSG00000038679 | Trps1 | ENSMUST00000165201 | 30.7445551 | 34 | 16.8833361 | 18.5379031 | 1 | 1 | 0 | 1 | 0 | 0 | 0 | 0 | 1 | 2 | 4 | 2 | 9 | 3 | 22 |
| ENSMUSG00000025086 | Trub1 | ENSMUST00000026072 | 27.140825 | 29 | 21.1095306 | 22.8733997 | 0 | 0 | 0 | 0 | 0 | 0 | 0 | 0 | 0 | 0 | 0 | 1 | 0 | 0 | 1 |
| ENSMUSG00000027806 | Tsc22d2 | ENSMUST00000099090 | 32.6784344 | 32 | 16.8994628 | 18.2041443 | 1 | 0 | 0 | 0 | 0 | 0 | 0 | 0 | 0 | 0 | 0 | 1 | 6 | 0 | 8 |
| ENSMUSG00000060771 | Tsga10 | ENSMUST00000123082 | 29.9065421 | 35 | 16.3738318 | 19.1775701 | 0 | 0 | 0 | 0 | 0 | 0 | 0 | 0 | 0 | 0 | 0 | 0 | 4 | 0 | 4 |
| ENSMUSG00000049580 | Tsku | ENSMUST00000094161 | 21.5107914 | 27 | 26.9784173 | 24.7482014 | 0 | 0 | 0 | 0 | 0 | 0 | 0 | 0 | 0 | 0 | 0 | 0 | 13 | 1 | 14 |
| ENSMUSG00000026374 | Tsn | ENSMUST00000027623 | 25.921009 | 30 | 20.3451709 | 23.2824427 | 2 | 1 | 0 | 1 | 0 | 0 | 0 | 0 | 1 | 2 | 4 | 0 | 8 | 0 | 17 |
| ENSMUSG00000037824 | Tspan14 | ENSMUST00000047652 | 21.6367018 | 29 | 25.9144451 | 23.8065716 | 0 | 0 | 0 | 0 | 0 | 0 | 0 | 0 | 0 | 0 | 0 | 0 | 2 | 0 | 2 |
| ENSMUSG00000027858 | Tspan2 | ENSMUST00000029451 | 31.3432836 | 31 | 18.9253731 | 18.3880597 | 0 | 0 | 0 | 0 | 0 | 0 | 0 | 0 | 0 | 0 | 0 | 1 | 1 | 0 | 2 |
| ENSMUSG00000067377 | Tspan6 | ENSMUST00000087557 | 27.690583 | 34 | 19.8430493 | 18.0493274 | 0 | 0 | 0 | 0 | 0 | 0 | 0 | 0 | 0 | 0 | 0 | 0 | 1 | 0 | 1 |
| ENSMUSG00000058254 | Tspan7 | ENSMUST00000115526 | 26.9218701 | 31 | 21.4778789 | 20.6620646 | 4 | 3 | 0 | 6 | 0 | 1 | 2 | 2 | 2 | 13 | 0 | 26 | 3 | 1 | 50 |
| ENSMUSG00000047514 | Tspyl1 | ENSMUST00000061372 | 26.7385445 | 35 | 20.1078167 | 18.4366577 | 0 | 0 | 0 | 0 | 0 | 0 | 0 | 0 | 0 | 0 | 0 | 1 | 10 | 1 | 12 |
| ENSMUSG00000039485 | Tspyl4 | ENSMUST00000047935 | 24.0139211 | 30 | 23.0085073 | 22.5444702 | 0 | 0 | 0 | 0 | 0 | 0 | 0 | 0 | 0 | 0 | 0 | 0 | 0 | 1 | 1 |
| ENSMUSG00000027677 | Ttc14 | ENSMUST00000099153 | 32.0754717 | 32 | 16.7405344 | 18.8046093 | 2 | 0 | 0 | 2 | 0 | 0 | 0 | 0 | 2 | 4 | 2 | 1 | 0 | 10 | 19 |
| ENSMUSG00000056832 | Ttc26 | ENSMUST00000159145 | 27.3365142 | 29 | 21.8074656 | 21.3864721 | 1 | 0 | 0 | 1 | 0 | 0 | 0 | 0 | 1 | 2 | 1 | 0 | 3 | 0 | 7 |
| ENSMUSG00000022151 | Ttc33 | ENSMUST00000081640 | 27.2504092 | 33 | 17.1031097 | 22.7495908 | 0 | 0 | 0 | 0 | 0 | 0 | 0 | 0 | 0 | 0 | 1 | 1 | 0 | 0 | 2 |
| ENSMUSG00000038172 | Ttc39b | ENSMUST00000102823 | 27.1144279 | 33 | 19.563945 | 20.2809482 | 3 | 1 | 1 | 2 | 0 | 0 | 0 | 0 | 2 | 5 | 0 | 1 | 2 | 1 | 13 |
| ENSMUSG00000006288 | Ttc5 | ENSMUST00000006451 | 23.9754098 | 36 | 21.1065574 | 19.2622951 | 0 | 0 | 1 | 0 | 0 | 0 | 0 | 0 | 0 | 1 | 0 | 1 | 8 | 1 | 11 |
| ENSMUSG00000071660 | Ttc9c | ENSMUST00000096751 | 28.4750843 | 31 | 20.3072312 | 19.7826901 | 0 | 0 | 0 | 0 | 0 | 0 | 0 | 0 | 0 | 0 | 0 | 0 | 1 | 2 | 3 |
| ENSMUSG00000036745 | Ttll7 | ENSMUST00000037942 | 30.6716768 | 32 | 17.8017849 | 19.2578675 | 2 | 0 | 0 | 1 | 0 | 0 | 0 | 0 | 1 | 2 | 9 | 2 | 7 | 0 | 22 |
| ENSMUSG00000033790 | Tubgcp5 | ENSMUST00000206191 | 30.5515862 | 29 | 19.5484424 | 21.0917405 | 1 | 0 | 0 | 2 | 0 | 0 | 0 | 0 | 2 | 4 | 0 | 0 | 0 | 0 | 5 |
| ENSMUSG00000010054 | Tusc2 | ENSMUST00000010198 | 23.4166667 | 25 | 26.25 | 25.0833333 | 0 | 0 | 0 | 0 | 0 | 0 | 0 | 0 | 0 | 0 | 0 | 1 | 0 | 0 | 1 |
| ENSMUSG00000014177 | Tvp23b | ENSMUST00000014321 | 27.3866924 | 37 | 13.4040501 | 22.2757956 | 1 | 0 | 0 | 1 | 0 | 0 | 0 | 0 | 1 | 2 | 1 | 0 | 0 | 0 | 4 |
| ENSMUSG00000024098 | Twsg1 | ENSMUST00000024906 | 27.8347313 | 31 | 20.1304753 | 21.4041628 | 0 | 0 | 0 | 1 | 0 | 0 | 0 | 0 | 1 | 2 | 2 | 0 | 3 | 0 | 7 |
| ENSMUSG00000038344 | Txlng | ENSMUST00000041370 | 31.0630465 | 33 | 18.1316153 | 17.4413254 | 0 | 0 | 0 | 0 | 0 | 0 | 0 | 0 | 0 | 0 | 0 | 2 | 0 | 0 | 2 |
| ENSMUSG00000028367 | Txn1 | ENSMUST00000030051 | 30.8270677 | 31 | 19.5488722 | 18.2330827 | 0 | 0 | 0 | 0 | 0 | 0 | 0 | 0 | 0 | 0 | 0 | 0 | 0 | 1 | 1 |
| ENSMUSG00000021830 | Txndc16 | ENSMUST00000022377 | 28.8888889 | 32 | 18.968254 | 20.1587302 | 1 | 0 | 1 | 0 | 0 | 0 | 0 | 0 | 0 | 1 | 0 | 0 | 0 | 0 | 2 |
| ENSMUSG00000038393 | Txnip | ENSMUST00000074519 | 23.1905465 | 31 | 21.8611521 | 23.929099 | 0 | 0 | 0 | 0 | 0 | 0 | 0 | 0 | 0 | 0 | 0 | 0 | 2 | 1 | 3 |
| ENSMUSG00000024583 | Txnl1 | ENSMUST00000025476 | 31.2250333 | 35 | 15.5126498 | 18.1091877 | 0 | 0 | 0 | 1 | 0 | 0 | 0 | 0 | 1 | 2 | 1 | 0 | 0 | 0 | 3 |
| ENSMUSG00000047583 | Tyw3 | ENSMUST00000170461 | 27.478076 | 26 | 22.9869785 | 23.4121711 | 0 | 0 | 0 | 0 | 0 | 0 | 0 | 0 | 0 | 0 | 0 | 26 | 1 | 0 | 27 |
| ENSMUSG00000061613 | U2af1 | ENSMUST00000014684 | 27.1317829 | 36 | 14.7286822 | 22.4806202 | 0 | 0 | 0 | 0 | 0 | 0 | 0 | 0 | 0 | 0 | 0 | 0 | 0 | 4 | 4 |
| ENSMUSG00000032407 | U2surp | ENSMUST00000079659 | 28.2893277 | 36 | 16.7517195 | 18.6820501 | 0 | 0 | 0 | 0 | 0 | 0 | 0 | 0 | 0 | 0 | 2 | 1 | 10 | 3 | 16 |
| ENSMUSG00000035898 | Uba6 | ENSMUST00000039373 | 30.9247696 | 36 | 15.4623848 | 17.8412132 | 0 | 0 | 0 | 0 | 0 | 0 | 0 | 0 | 0 | 0 | 2 | 0 | 4 | 1 | 7 |
| ENSMUSG00000020390 | Ube2b | ENSMUST00000020657 | 27.9324895 | 34 | 18.8185654 | 19.3248945 | 0 | 0 | 0 | 0 | 0 | 0 | 0 | 0 | 0 | 0 | 0 | 0 | 1 | 0 | 1 |
| ENSMUSG00000001403 | Ube2c | ENSMUST00000001439 | 23.4042553 | 22 | 30.6633292 | 23.59199 | 0 | 0 | 0 | 0 | 0 | 0 | 0 | 0 | 0 | 0 | 0 | 2 | 0 | 0 | 2 |
| ENSMUSG00000078578 | Ube2d3 | ENSMUST00000197859 | 29.3103448 | 35 | 16.5652468 | 18.6950642 | 1 | 0 | 0 | 0 | 0 | 0 | 0 | 0 | 0 | 0 | 0 | 1 | 0 | 0 | 2 |
| ENSMUSG00000058317 | Ube2e2 | ENSMUST00000176555 | 21.905805 | 33 | 23.8773275 | 21.0295728 | 0 | 0 | 0 | 0 | 0 | 0 | 0 | 0 | 0 | 0 | 0 | 4 | 1 | 0 | 5 |
| ENSMUSG00000034343 | Ube2f | ENSMUST00000171165 | 25.6724207 | 27 | 24.227218 | 23.4443998 | 0 | 0 | 0 | 0 | 0 | 0 | 0 | 0 | 0 | 0 | 3 | 3 | 7 | 0 | 13 |
| ENSMUSG00000020794 | Ube2g1 | ENSMUST00000021148 | 27.4457429 | 36 | 18.196995 | 18.5308848 | 0 | 0 | 0 | 0 | 0 | 0 | 0 | 0 | 0 | 0 | 0 | 0 | 4 | 0 | 4 |
| ENSMUSG00000039159 | Ube2h | ENSMUST00000102993 | 24.0326975 | 29 | 22.7247956 | 23.8964578 | 0 | 0 | 0 | 0 | 0 | 0 | 0 | 0 | 0 | 0 | 0 | 2 | 10 | 0 | 12 |
| ENSMUSG00000028277 | Ube2j1 | ENSMUST00000124992 | 27.0079068 | 33 | 19.3091968 | 21.0153974 | 0 | 0 | 0 | 0 | 0 | 0 | 0 | 0 | 0 | 0 | 0 | 0 | 2 | 0 | 2 |
| ENSMUSG00000029203 | Ube2k | ENSMUST00000142407 | 27.0611538 | 35 | 17.6281258 | 20.5743996 | 1 | 0 | 0 | 0 | 0 | 0 | 0 | 0 | 0 | 0 | 2 | 0 | 8 | 0 | 11 |
| ENSMUSG00000074781 | Ube2n | ENSMUST00000099329 | 27.1667663 | 33 | 19.1870891 | 20.3526599 | 0 | 0 | 0 | 0 | 0 | 0 | 0 | 0 | 0 | 0 | 0 | 2 | 0 | 0 | 2 |
| ENSMUSG00000026429 | Ube2t | ENSMUST00000027687 | 27.7777778 | 43 | 13.4259259 | 15.7407407 | 1 | 0 | 0 | 0 | 0 | 0 | 0 | 0 | 0 | 0 | 0 | 0 | 1 | 0 | 2 |
| ENSMUSG00000022674 | Ube2v2 | ENSMUST00000115777 | 30.4032258 | 36 | 15.3763441 | 18.7096774 | 0 | 0 | 1 | 1 | 0 | 0 | 0 | 0 | 1 | 3 | 0 | 0 | 0 | 0 | 3 |
| ENSMUSG00000025939 | Ube2w | ENSMUST00000117146 | 28.0430397 | 38 | 15.870881 | 18.2246133 | 2 | 2 | 0 | 1 | 0 | 0 | 0 | 1 | 0 | 2 | 1 | 0 | 1 | 0 | 8 |
| ENSMUSG00000025326 | Ube3a | ENSMUST00000200758 | 29.0947623 | 37 | 17.2267635 | 16.833182 | 1 | 1 | 0 | 0 | 0 | 0 | 0 | 0 | 0 | 0 | 0 | 2 | 8 | 2 | 14 |
| ENSMUSG00000030870 | Ubfd1 | ENSMUST00000033158 | 21.1612548 | 34 | 22.4270776 | 22.2069345 | 0 | 0 | 0 | 0 | 0 | 0 | 0 | 0 | 0 | 0 | 1 | 0 | 9 | 2 | 12 |
| ENSMUSG00000038538 | Ubn2 | ENSMUST00000160583 | 28.1118748 | 33 | 18.4994273 | 20.532646 | 0 | 0 | 0 | 1 | 0 | 0 | 0 | 0 | 1 | 2 | 5 | 4 | 7 | 4 | 22 |
| ENSMUSG00000009741 | Ubp1 | ENSMUST00000009885 | 23.3480176 | 31 | 22.8524229 | 23.2929515 | 0 | 0 | 0 | 0 | 0 | 0 | 0 | 0 | 0 | 0 | 0 | 1 | 0 | 0 | 1 |
| ENSMUSG00000005312 | Ubqln1 | ENSMUST00000076454 | 27.8488372 | 33 | 17.3837209 | 21.627907 | 0 | 0 | 0 | 0 | 0 | 0 | 0 | 0 | 0 | 0 | 1 | 1 | 0 | 0 | 2 |
| ENSMUSG00000044308 | Ubr3 | ENSMUST00000055758 | 31.5767106 | 33 | 16.4045899 | 19.4220144 | 0 | 0 | 0 | 1 | 0 | 0 | 0 | 0 | 1 | 2 | 0 | 0 | 0 | 0 | 2 |
| ENSMUSG00000028243 | Ubxn2b | ENSMUST00000029907 | 28.0370502 | 31 | 20.1104382 | 20.3598147 | 0 | 0 | 0 | 0 | 0 | 0 | 0 | 0 | 0 | 0 | 0 | 1 | 0 | 0 | 1 |
| ENSMUSG00000053774 | Ubxn7 | ENSMUST00000115151 | 28.0072628 | 33 | 18.0889696 | 20.4266909 | 2 | 1 | 0 | 2 | 0 | 0 | 0 | 0 | 2 | 4 | 0 | 2 | 20 | 5 | 34 |
| ENSMUSG00000018189 | Uchl5 | ENSMUST00000018333 | 31.4363144 | 37 | 13.9566396 | 17.3441734 | 0 | 0 | 1 | 0 | 0 | 0 | 0 | 0 | 0 | 1 | 0 | 0 | 0 | 0 | 1 |
| ENSMUSG00000043262 | Uevld | ENSMUST00000094398 | 28.0193237 | 33 | 19.1626409 | 19.8067633 | 0 | 0 | 0 | 0 | 0 | 0 | 0 | 0 | 0 | 0 | 0 | 1 | 1 | 3 | 5 |
| ENSMUSG00000028381 | Ugcg | ENSMUST00000030074 | 27.2131148 | 36 | 17.2131148 | 19.5491803 | 0 | 0 | 1 | 1 | 0 | 0 | 0 | 0 | 1 | 3 | 4 | 2 | 15 | 2 | 26 |
| ENSMUSG00000029201 | Ugdh | ENSMUST00000031103 | 29.5238095 | 38 | 15.5102041 | 16.5986395 | 0 | 0 | 0 | 0 | 0 | 0 | 0 | 0 | 0 | 0 | 1 | 0 | 1 | 1 | 3 |
| ENSMUSG00000042104 | Uggt2 | ENSMUST00000156203 | 27.9157667 | 31 | 21.3822894 | 19.924406 | 0 | 0 | 0 | 0 | 0 | 0 | 0 | 0 | 0 | 0 | 0 | 1 | 0 | 0 | 1 |
| ENSMUSG00000032854 | Ugt8a | ENSMUST00000057944 | 32.0308483 | 34 | 16.2467866 | 17.2750643 | 1 | 0 | 0 | 0 | 0 | 0 | 0 | 0 | 0 | 0 | 0 | 0 | 0 | 0 | 1 |
| ENSMUSG00000026667 | Uhmk1 | ENSMUST00000123399 | 25.2772292 | 33 | 19.975695 | 21.9960504 | 2 | 2 | 0 | 2 | 0 | 0 | 0 | 1 | 1 | 4 | 6 | 5 | 2 | 0 | 21 |
| ENSMUSG00000024817 | Uhrf2 | ENSMUST00000025739 | 25.7575758 | 34 | 19.3473193 | 20.979021 | 1 | 0 | 0 | 0 | 0 | 0 | 0 | 0 | 0 | 0 | 0 | 0 | 0 | 0 | 1 |
| ENSMUSG00000025878 | Uimc1 | ENSMUST00000148702 | 28.8685524 | 26 | 19.7171381 | 25.0415973 | 1 | 0 | 0 | 1 | 0 | 0 | 0 | 0 | 1 | 2 | 0 | 0 | 0 | 0 | 3 |
| ENSMUSG00000059921 | Unc5c | ENSMUST00000106236 | 28.8255668 | 32 | 19.6473552 | 19.2537783 | 0 | 0 | 0 | 0 | 0 | 0 | 0 | 0 | 0 | 0 | 4 | 2 | 4 | 0 | 10 |
| ENSMUSG00000063626 | Unc5d | ENSMUST00000168630 | 30.6065113 | 30 | 19.6331185 | 19.4843827 | 0 | 0 | 0 | 0 | 0 | 0 | 0 | 0 | 0 | 0 | 1 | 2 | 5 | 0 | 8 |
| ENSMUSG00000031976 | Urb2 | ENSMUST00000173168 | 26.2172285 | 28 | 23.0604601 | 22.6859283 | 0 | 0 | 0 | 0 | 0 | 0 | 0 | 0 | 0 | 0 | 0 | 0 | 0 | 1 | 1 |
| ENSMUSG00000030421 | Uri1 | ENSMUST00000085513 | 28.0367695 | 34 | 19.6323047 | 18.3847669 | 0 | 0 | 0 | 0 | 0 | 0 | 0 | 0 | 0 | 0 | 4 | 0 | 5 | 0 | 9 |
| ENSMUSG00000025437 | Usp33 | ENSMUST00000123237 | 28.1048387 | 28 | 20.6451613 | 22.9032258 | 1 | 1 | 0 | 0 | 0 | 0 | 0 | 0 | 0 | 0 | 0 | 1 | 6 | 0 | 9 |
| ENSMUSG00000033364 | Usp37 | ENSMUST00000044260 | 25.4123711 | 32 | 20.3092784 | 22.0103093 | 0 | 0 | 0 | 1 | 0 | 0 | 0 | 0 | 1 | 2 | 0 | 1 | 22 | 1 | 26 |
| ENSMUSG00000040455 | Usp45 | ENSMUST00000137293 | 30.9979633 | 30 | 18.9409369 | 19.5723014 | 0 | 1 | 0 | 1 | 0 | 0 | 0 | 0 | 1 | 2 | 0 | 1 | 1 | 0 | 5 |
| ENSMUSG00000054814 | Usp46 | ENSMUST00000119154 | 26.7652805 | 32 | 19.7320681 | 21.0996372 | 1 | 0 | 0 | 2 | 0 | 0 | 0 | 0 | 2 | 4 | 0 | 1 | 1 | 4 | 11 |
| ENSMUSG00000039701 | Usp53 | ENSMUST00000090379 | 32.2117541 | 38 | 15.1637506 | 15.1188874 | 3 | 1 | 1 | 4 | 0 | 0 | 0 | 1 | 3 | 9 | 0 | 0 | 0 | 2 | 15 |
| ENSMUSG00000031010 | Usp9x | ENSMUST00000089302 | 28.4079084 | 36 | 17.0655567 | 18.8605619 | 0 | 0 | 0 | 0 | 0 | 0 | 0 | 0 | 0 | 0 | 2 | 0 | 7 | 0 | 9 |
| ENSMUSG00000079470 | Utp14b | ENSMUST00000053760 | 27.2841313 | 31 | 19.130253 | 22.099101 | 1 | 0 | 0 | 1 | 0 | 0 | 0 | 0 | 1 | 2 | 0 | 27 | 7 | 4 | 41 |
| ENSMUSG00000041747 | Utp15 | ENSMUST00000040972 | 23.2486865 | 33 | 21.0595447 | 22.1978984 | 1 | 0 | 0 | 2 | 0 | 0 | 0 | 0 | 2 | 4 | 0 | 2 | 4 | 0 | 11 |

| | | | | | | | | | | | | | | | | | | | | | |
|---|---|---|---|---|---|---|---|---|---|---|---|---|---|---|---|---|---|---|---|---|---|
| ENSMUSG00000054079 | Utp18 | ENSMUST00000066888 | 32.3076923 | 39 | 14.2011834 | 14.4378698 | 0 | 0 | 0 | 0 | 0 | 0 | 0 | 0 | 0 | 0 | 0 | 0 | 12 | 4 | 16 |
| ENSMUSG00000022313 | Utp23 | ENSMUST00000137116 | 25.9027443 | 32 | 20.8955224 | 21.2806933 | 0 | 0 | 0 | 0 | 0 | 0 | 0 | 0 | 0 | 0 | 0 | 14 | 0 | 0 | 14 |
| ENSMUSG00000068457 | Uty | ENSMUST00000154666 | 32.8559028 | 33 | 17.2526042 | 17.3828125 | 0 | 0 | 0 | 1 | 0 | 0 | 0 | 0 | 1 | 2 | 0 | 0 | 1 | 0 | 3 |
| ENSMUSG00000057363 | Uxs1 | ENSMUST00000126008 | 29.4333844 | 32 | 17.3047473 | 21.6232772 | 1 | 0 | 0 | 2 | 0 | 0 | 0 | 0 | 2 | 4 | 0 | 1 | 0 | 1 | 7 |
| ENSMUSG00000020894 | Vamp2 | ENSMUST00000021273 | 19.0337602 | 30 | 28.6961583 | 21.9441211 | 0 | 0 | 0 | 0 | 0 | 0 | 0 | 0 | 0 | 0 | 1 | 7 | 2 | 0 | 10 |
| ENSMUSG00000028955 | Vamp3 | ENSMUST00000030797 | 25.4913295 | 32 | 20.5780347 | 21.849711 | 0 | 0 | 0 | 0 | 0 | 0 | 0 | 0 | 0 | 0 | 1 | 0 | 3 | 0 | 4 |
| ENSMUSG00000026696 | Vamp4 | ENSMUST00000132158 | 26.1086198 | 33 | 18.5351271 | 22.1723966 | 0 | 0 | 0 | 0 | 0 | 0 | 0 | 0 | 0 | 0 | 4 | 0 | 9 | 0 | 13 |
| ENSMUSG00000046844 | Vat1l | ENSMUST00000049509 | 25.1074807 | 27 | 24.9785039 | 22.4419604 | 0 | 0 | 1 | 0 | 0 | 0 | 0 | 0 | 0 | 1 | 0 | 5 | 1 | 0 | 7 |
| ENSMUSG00000009621 | Vav2 | ENSMUST00000185188 | 20.4597701 | 26 | 26.1609195 | 27.5862069 | 0 | 0 | 0 | 0 | 0 | 0 | 0 | 0 | 0 | 0 | 8 | 0 | 9 | 0 | 17 |
| ENSMUSG00000033721 | Vav3 | ENSMUST00000046864 | 26.7592149 | 32 | 20.4404021 | 20.6797511 | 0 | 0 | 0 | 0 | 0 | 0 | 0 | 0 | 0 | 0 | 0 | 2 | 1 | 2 | 5 |
| ENSMUSG00000031197 | Vbp1 | ENSMUST00000033540 | 31.846473 | 37 | 14.8340249 | 16.5975104 | 1 | 1 | 0 | 2 | 0 | 0 | 0 | 0 | 2 | 4 | 0 | 0 | 1 | 0 | 7 |
| ENSMUSG00000045210 | Vcpip1 | ENSMUST00000057438 | 32.1331058 | 34 | 15.7679181 | 17.6450512 | 2 | 1 | 0 | 2 | 0 | 0 | 0 | 1 | 1 | 4 | 0 | 1 | 4 | 2 | 14 |
| ENSMUSG00000023951 | Vegfa | ENSMUST00000024747 | 27.4530831 | 29 | 21.5013405 | 21.7158177 | 3 | 1 | 0 | 1 | 0 | 0 | 0 | 0 | 1 | 2 | 0 | 3 | 0 | 1 | 10 |
| ENSMUSG00000027831 | Veph1 | ENSMUST00000029419 | 30.6738089 | 27 | 20.6089653 | 21.2574006 | 2 | 1 | 0 | 4 | 0 | 0 | 0 | 1 | 3 | 8 | 0 | 0 | 0 | 0 | 11 |
| ENSMUSG00000018377 | Vezf1 | ENSMUST00000018521 | 29.6143251 | 31 | 18.1473829 | 21.453168 | 0 | 0 | 0 | 0 | 0 | 0 | 0 | 0 | 0 | 0 | 0 | 2 | 0 | 0 | 2 |
| ENSMUSG00000036099 | Vezt | ENSMUST00000119818 | 27.5170107 | 32 | 20.1476185 | 20.620459 | 0 | 0 | 0 | 0 | 0 | 0 | 0 | 0 | 0 | 0 | 1 | 2 | 7 | 0 | 10 |
| ENSMUSG00000033933 | Vhl | ENSMUST00000035673 | 23.1589147 | 32 | 21.3178295 | 23.8856589 | 1 | 0 | 0 | 1 | 0 | 0 | 0 | 0 | 1 | 2 | 0 | 0 | 0 | 1 | 4 |
| ENSMUSG00000066735 | Vkorc1l1 | ENSMUST00000201855 | 28.2439248 | 30 | 20.3347088 | 21.7331499 | 0 | 0 | 0 | 1 | 0 | 0 | 0 | 0 | 1 | 2 | 5 | 4 | 9 | 1 | 21 |
| ENSMUSG00000073131 | Vma21 | ENSMUST00000114576 | 30.9124767 | 35 | 16.1745145 | 17.6908752 | 3 | 0 | 0 | 1 | 0 | 0 | 0 | 0 | 1 | 2 | 0 | 0 | 1 | 0 | 6 |
| ENSMUSG00000018171 | Vmp1 | ENSMUST00000018315 | 26.5419265 | 34 | 19.4733195 | 20.0970201 | 0 | 0 | 0 | 0 | 0 | 0 | 0 | 0 | 0 | 0 | 0 | 1 | 1 | 0 | 2 |
| ENSMUSG00000037788 | Vopp1 | ENSMUST00000145608 | 28.1937237 | 32 | 22.0657277 | 17.9639239 | 0 | 0 | 0 | 0 | 0 | 0 | 0 | 0 | 0 | 0 | 1 | 2 | 0 | 0 | 3 |
| ENSMUSG00000040325 | Vprbp | ENSMUST00000161758 | 30.6 | 29 | 22.3 | 17.7 | 0 | 1 | 0 | 1 | 0 | 0 | 0 | 0 | 1 | 2 | 0 | 1 | 0 | 0 | 4 |
| ENSMUSG00000035284 | Vps13c | ENSMUST00000077879 | 23.6734694 | 35 | 17.9591837 | 23.2653061 | 0 | 0 | 0 | 0 | 0 | 0 | 0 | 0 | 0 | 0 | 0 | 0 | 2 | 0 | 2 |
| ENSMUSG00000020078 | Vps26a | ENSMUST00000092473 | 27.7602524 | 33 | 18.8748686 | 20.4521556 | 1 | 0 | 0 | 1 | 0 | 0 | 0 | 0 | 1 | 2 | 0 | 0 | 3 | 0 | 6 |
| ENSMUSG00000029462 | Vps29 | ENSMUST00000155671 | 29.6314258 | 30 | 20.514064 | 20.1260912 | 1 | 0 | 0 | 3 | 0 | 0 | 0 | 0 | 3 | 6 | 1 | 0 | 1 | 2 | 11 |
| ENSMUSG00000031600 | Vps37a | ENSMUST00000098817 | 30.9940317 | 32 | 18.2547849 | 18.7281334 | 0 | 0 | 1 | 2 | 0 | 0 | 0 | 0 | 2 | 5 | 0 | 0 | 1 | 0 | 6 |
| ENSMUSG00000009907 | Vps4b | ENSMUST00000094646 | 28.9028837 | 33 | 19.5227047 | 18.2963208 | 4 | 2 | 0 | 5 | 0 | 0 | 0 | 1 | 4 | 10 | 0 | 1 | 3 | 0 | 20 |
| ENSMUSG00000021115 | Vrk1 | ENSMUST00000021539 | 24.5784148 | 32 | 20.7419899 | 22.9342327 | 0 | 0 | 0 | 0 | 0 | 0 | 0 | 0 | 0 | 0 | 1 | 0 | 4 | 0 | 5 |
| ENSMUSG00000048834 | Vstm2a | ENSMUST00000109645 | 31.0268841 | 36 | 17.0559718 | 16.0423094 | 0 | 0 | 0 | 1 | 0 | 0 | 0 | 0 | 1 | 2 | 0 | 0 | 0 | 0 | 2 |
| ENSMUSG00000050830 | Vwc2 | ENSMUST00000129670 | 31.5547025 | 32 | 19.6161228 | 17.2744722 | 0 | 0 | 0 | 2 | 0 | 0 | 0 | 0 | 2 | 4 | 0 | 0 | 1 | 0 | 5 |
| ENSMUSG00000045648 | Vwc2l | ENSMUST00000161937 | 29.3061473 | 30 | 21.1199026 | 19.7808886 | 0 | 0 | 0 | 0 | 0 | 0 | 0 | 0 | 0 | 0 | 1 | 3 | 0 | 0 | 4 |
| ENSMUSG00000024283 | Wac | ENSMUST00000167020 | 30.6334372 | 34 | 16.7878159 | 18.7608169 | 0 | 0 | 0 | 0 | 0 | 0 | 0 | 0 | 0 | 0 | 0 | 1 | 4 | 2 | 7 |
| ENSMUSG00000041408 | Wapl | ENSMUST00000048263 | 28.6960314 | 36 | 16.1796773 | 19.0143916 | 0 | 0 | 0 | 0 | 0 | 0 | 0 | 0 | 0 | 0 | 0 | 0 | 4 | 3 | 7 |
| ENSMUSG00000029684 | Wasl | ENSMUST00000031695 | 28.1854839 | 36 | 17.8225806 | 18.4274194 | 0 | 0 | 0 | 0 | 0 | 0 | 0 | 0 | 0 | 0 | 0 | 1 | 2 | 4 | 7 |
| ENSMUSG00000022023 | Wbp4 | ENSMUST00000022601 | 26.1460957 | 34 | 19.1939547 | 20.906801 | 1 | 0 | 1 | 0 | 0 | 0 | 0 | 0 | 0 | 1 | 0 | 0 | 3 | 0 | 5 |
| ENSMUSG00000037572 | Wdhd1 | ENSMUST00000111790 | 29.8377029 | 36 | 16.6042447 | 17.8526841 | 0 | 0 | 0 | 0 | 0 | 0 | 0 | 0 | 0 | 0 | 0 | 0 | 12 | 0 | 12 |
| ENSMUSG00000026019 | Wdr12 | ENSMUST00000027173 | 29.0368779 | 32 | 18.7134503 | 20.4797709 | 1 | 0 | 0 | 2 | 0 | 0 | 0 | 0 | 2 | 4 | 3 | 0 | 33 | 0 | 41 |
| ENSMUSG00000037890 | Wdr19 | ENSMUST00000203653 | 24.2935278 | 31 | 24.3846855 | 20.1002735 | 0 | 0 | 0 | 0 | 0 | 0 | 0 | 0 | 0 | 0 | 0 | 0 | 5 | 0 | 5 |
| ENSMUSG00000024400 | Wdr33 | ENSMUST00000025264 | 27.6104418 | 27 | 23.4437751 | 21.937751 | 0 | 0 | 0 | 0 | 0 | 0 | 0 | 0 | 0 | 0 | 1 | 0 | 0 | 0 | 1 |
| ENSMUSG00000038299 | Wdr36 | ENSMUST00000053663 | 28.9473684 | 39 | 16.0287081 | 16.507177 | 0 | 0 | 0 | 0 | 0 | 0 | 0 | 0 | 0 | 0 | 0 | 0 | 8 | 0 | 8 |
| ENSMUSG00000021147 | Wdr37 | ENSMUST00000176922 | 27.1070615 | 38 | 16.2056622 | 18.2232346 | 1 | 0 | 0 | 0 | 0 | 0 | 0 | 0 | 0 | 0 | 3 | 22 | 6 | 0 | 32 |
| ENSMUSG00000041057 | Wdr43 | ENSMUST00000047086 | 28.1021898 | 32 | 19.4160584 | 20.8029197 | 0 | 0 | 1 | 1 | 0 | 0 | 0 | 0 | 1 | 3 | 0 | 0 | 0 | 0 | 3 |
| ENSMUSG00000036769 | Wdr44 | ENSMUST00000101670 | 32.3416507 | 40 | 11.2284069 | 16.6026871 | 0 | 0 | 0 | 1 | 0 | 0 | 0 | 0 | 1 | 2 | 0 | 0 | 2 | 0 | 4 |
| ENSMUSG00000025173 | Wdr45b | ENSMUST00000026173 | 22.1340388 | 29 | 24.1622575 | 24.5149912 | 0 | 0 | 0 | 0 | 0 | 0 | 0 | 0 | 0 | 0 | 0 | 0 | 1 | 0 | 1 |
| ENSMUSG00000061559 | Wdr61 | ENSMUST00000118771 | 27.0701976 | 26 | 24.0857503 | 23.160992 | 0 | 0 | 0 | 0 | 0 | 0 | 0 | 0 | 0 | 0 | 0 | 1 | 0 | 0 | 1 |
| ENSMUSG00000039828 | Wdr70 | ENSMUST00000045766 | 34.2342342 | 31 | 18.9189189 | 16.2162162 | 0 | 0 | 0 | 0 | 0 | 0 | 0 | 0 | 0 | 0 | 0 | 0 | 0 | 1 | 1 |
| ENSMUSG00000027242 | Wdr76 | ENSMUST00000110602 | 27.5021758 | 32 | 19.5822454 | 20.9312446 | 1 | 1 | 0 | 1 | 0 | 0 | 0 | 0 | 1 | 2 | 0 | 1 | 10 | 2 | 17 |
| ENSMUSG00000020257 | Wdr82 | ENSMUST00000020490 | 23.9745121 | 30 | 22.0629231 | 23.5762644 | 0 | 0 | 0 | 0 | 0 | 0 | 0 | 0 | 0 | 0 | 0 | 1 | 4 | 0 | 5 |
| ENSMUSG00000078970 | Wdr92 | ENSMUST00000046955 | 26.5306122 | 31 | 21.4285714 | 21.2018141 | 0 | 0 | 2 | 0 | 0 | 0 | 0 | 0 | 0 | 2 | 0 | 0 | 0 | 0 | 2 |
| ENSMUSG00000041895 | Wipi1 | ENSMUST00000047186 | 24.789916 | 26 | 24.894958 | 24.1596639 | 0 | 0 | 0 | 0 | 0 | 0 | 0 | 0 | 0 | 0 | 0 | 12 | 0 | 0 | 12 |
| ENSMUSG00000005124 | Wisp1 | ENSMUST00000005255 | 28.6896184 | 30 | 20.7899653 | 21.0034694 | 0 | 0 | 0 | 0 | 0 | 0 | 0 | 0 | 0 | 0 | 1 | 2 | 2 | 0 | 5 |
| ENSMUSG00000028173 | Wls | ENSMUST00000200191 | 26.7322097 | 30 | 20.8333333 | 22.4719101 | 0 | 0 | 0 | 0 | 0 | 0 | 0 | 0 | 0 | 0 | 0 | 1 | 0 | 0 | 1 |
| ENSMUSG00000045962 | Wnk1 | ENSMUST00000060043 | 27.5301958 | 31 | 18.9504373 | 22.0741358 | 0 | 0 | 0 | 0 | 0 | 0 | 0 | 0 | 0 | 0 | 0 | 1 | 4 | 0 | 5 |
| ENSMUSG00000041245 | Wnk3 | ENSMUST00000149140 | 30.5007153 | 25 | 23.6623748 | 21.0586552 | 0 | 0 | 0 | 0 | 0 | 0 | 0 | 0 | 0 | 0 | 0 | 1 | 0 | 0 | 1 |
| ENSMUSG00000021994 | Wnt5a | ENSMUST00000063465 | 27.20213 | 33 | 19.8801864 | 20.213002 | 3 | 0 | 0 | 1 | 0 | 0 | 0 | 0 | 1 | 2 | 6 | 10 | 16 | 3 | 40 |
| ENSMUSG00000023147 | Wrb | ENSMUST00000023913 | 23.4736842 | 31 | 21.6842105 | 24.0526316 | 0 | 0 | 0 | 0 | 0 | 0 | 0 | 0 | 0 | 0 | 2 | 1 | 17 | 0 | 20 |
| ENSMUSG00000031583 | Wrn; LOC105242468 | ENSMUST00000033991 | 31.2852023 | 32 | 19.1500256 | 17.3067076 | 0 | 0 | 0 | 0 | 0 | 0 | 0 | 0 | 0 | 0 | 0 | 0 | 1 | 1 | 2 |
| ENSMUSG00000021400 | Wrnip1 | ENSMUST00000021832 | 30 | 36 | 13.1111111 | 20.6666667 | 0 | 0 | 1 | 0 | 0 | 0 | 0 | 0 | 0 | 1 | 0 | 0 | 0 | 2 | 3 |
| ENSMUSG00000017677 | Wsb1 | ENSMUST00000017821 | 30.1825994 | 36 | 15.3598281 | 18.7969925 | 0 | 1 | 0 | 0 | 0 | 0 | 0 | 0 | 0 | 0 | 0 | 0 | 1 | 0 | 2 |
| ENSMUSG00000016458 | Wt1 | ENSMUST00000111099 | 27.9874214 | 31 | 18.8679245 | 21.8553459 | 1 | 0 | 0 | 0 | 0 | 0 | 0 | 0 | 0 | 0 | 0 | 0 | 0 | 0 | 1 |
| ENSMUSG00000060475 | Wtap | ENSMUST00000159986 | 29.535865 | 33 | 16.5324127 | 20.6367472 | 0 | 0 | 0 | 0 | 0 | 0 | 0 | 0 | 0 | 0 | 0 | 1 | 0 | 0 | 1 |
| ENSMUSG00000041058 | Wwp1 | ENSMUST00000108246 | 30.7241278 | 33 | 18.7041923 | 17.1797127 | 0 | 0 | 1 | 0 | 0 | 0 | 0 | 0 | 0 | 1 | 0 | 0 | 0 | 0 | 1 |
| ENSMUSG00000027803 | Wwtr1 | ENSMUST00000029380 | 29.9969484 | 33 | 17.485505 | 19.8657309 | 0 | 0 | 0 | 0 | 0 | 0 | 0 | 0 | 0 | 0 | 0 | 1 | 1 | 3 | 5 |
| ENSMUSG00000025860 | Xiap | ENSMUST00000026978 | 29.4520548 | 35 | 16.9306507 | 18.9212329 | 0 | 0 | 0 | 1 | 0 | 0 | 0 | 0 | 1 | 2 | 1 | 9 | 1 | 0 | 13 |
| ENSMUSG00000015342 | Xk | ENSMUST00000015486 | 30.0485576 | 33 | 19.3373322 | 17.3379035 | 1 | 0 | 1 | 0 | 0 | 0 | 0 | 0 | 0 | 1 | 1 | 0 | 0 | 0 | 3 |
| ENSMUSG00000028329 | Xpa | ENSMUST00000132358 | 37.345679 | 23 | 16.9753086 | 23.1481481 | 0 | 0 | 0 | 1 | 0 | 0 | 0 | 0 | 1 | 2 | 0 | 0 | 0 | 0 | 2 |
| ENSMUSG00000020290 | Xpo1 | ENSMUST00000102869 | 28.4780579 | 33 | 16.43324 | 21.9421102 | 1 | 0 | 0 | 0 | 0 | 0 | 0 | 0 | 0 | 0 | 2 | 1 | 0 | 0 | 4 |
| ENSMUSG00000021952 | Xpo4 | ENSMUST00000174152 | 27.8206365 | 27 | 22.2275796 | 23.0472517 | 0 | 0 | 0 | 0 | 0 | 0 | 0 | 0 | 0 | 0 | 0 | 0 | 1 | 0 | 1 |
| ENSMUSG00000026469 | Xpr1 | ENSMUST00000027741 | 27.3686193 | 31 | 22.0380486 | 20.0414391 | 0 | 0 | 0 | 0 | 0 | 0 | 0 | 0 | 0 | 0 | 0 | 4 | 0 | 0 | 4 |
| ENSMUSG00000028933 | Xrcc2 | ENSMUST00000030773 | 24.5980009 | 30 | 22.468492 | 23.0334637 | 0 | 0 | 0 | 1 | 0 | 0 | 0 | 0 | 1 | 2 | 0 | 0 | 2 | 0 | 4 |
| ENSMUSG00000021615 | Xrcc4 | ENSMUST00000159199 | 26.7622951 | 32 | 22.3360656 | 19.057377 | 0 | 0 | 0 | 0 | 0 | 0 | 0 | 0 | 0 | 0 | 1 | 0 | 0 | 0 | 1 |
| ENSMUSG00000025436 | Xrcc6bp1 | ENSMUST00000168520 | 33.4710744 | 27 | 15.2892562 | 23.9669421 | 0 | 0 | 0 | 0 | 0 | 0 | 0 | 0 | 0 | 0 | 0 | 0 | 1 | 0 | 1 |
| ENSMUSG00000032410 | Xrn1 | ENSMUST00000185633 | 26.5919103 | 33 | 19.5835002 | 20.7849419 | 0 | 0 | 0 | 0 | 0 | 0 | 0 | 0 | 0 | 0 | 1 | 1 | 2 | 3 | 7 |
| ENSMUSG00000075054 | Yae1d1 | ENSMUST00000099735 | 32.2627737 | 29 | 18.0656934 | 20.2189781 | 0 | 0 | 0 | 0 | 0 | 0 | 0 | 0 | 0 | 0 | 0 | 0 | 0 | 2 | 2 |
| ENSMUSG00000022634 | Yaf2 | ENSMUST00000080299 | 30.0900901 | 36 | 17.7477477 | 16.1261261 | 0 | 0 | 0 | 0 | 0 | 0 | 0 | 0 | 0 | 0 | 1 | 0 | 0 | 0 | 1 |
| ENSMUSG00000020171 | Yeats4 | ENSMUST00000020382 | 18.8571429 | 32 | 22.8571429 | 26.6666667 | 0 | 0 | 0 | 0 | 0 | 0 | 0 | 0 | 0 | 0 | 0 | 0 | 1 | 0 | 1 |
| ENSMUSG00000014932 | Yes1 | ENSMUST00000072311 | 33.8555265 | 34 | 14.7084421 | 17.1018277 | 1 | 0 | 0 | 0 | 0 | 0 | 0 | 0 | 0 | 0 | 1 | 0 | 4 | 0 | 6 |
| ENSMUSG00000024072 | Yipf4 | ENSMUST00000024873 | 31.5654405 | 40 | 13.0025663 | 15.6544055 | 0 | 0 | 1 | 0 | 0 | 0 | 0 | 0 | 0 | 1 | 0 | 0 | 2 | 3 | 6 |
| ENSMUSG00000047694 | Yipf6 | ENSMUST00000054697 | 28.7168032 | 34 | 17.6966989 | 19.558203 | 1 | 0 | 0 | 0 | 0 | 0 | 0 | 0 | 0 | 0 | 2 | 0 | 7 | 0 | 10 |
| ENSMUSG00000002741 | Ykt6 | ENSMUST00000002818 | 21.8894009 | 27 | 22.1774194 | 28.6866359 | 0 | 0 | 0 | 0 | 0 | 0 | 0 | 0 | 0 | 0 | 0 | 1 | 0 | 0 | 1 |
| ENSMUSG00000026775 | Yme1l1 | ENSMUST00000028117 | 29.1610588 | 37 | 16.5096456 | 16.9582773 | 0 | 0 | 0 | 0 | 0 | 0 | 0 | 0 | 0 | 0 | 1 | 0 | 15 | 0 | 16 |
| ENSMUSG00000046404 | Yod1 | ENSMUST00000049813 | 29.3379227 | 36 | 17.0479854 | 18.1008301 | 3 | 1 | 0 | 3 | 0 | 0 | 0 | 0 | 3 | 6 | 0 | 0 | 26 | 1 | 37 |
| ENSMUSG00000018427 | Ypel2 | ENSMUST00000018571 | 25.9223766 | 29 | 22.3526593 | 22.975563 | 1 | 0 | 0 | 1 | 0 | 0 | 0 | 0 | 1 | 2 | 0 | 2 | 10 | 3 | 18 |
| ENSMUSG00000039770 | Ypel5 | ENSMUST00000045174 | 27.4780427 | 33 | 19.1551652 | 19.9498118 | 0 | 0 | 0 | 0 | 0 | 0 | 0 | 0 | 0 | 0 | 0 | 1 | 2 | 1 | 4 |
| ENSMUSG00000035851 | Ythdc1 | ENSMUST00000156363 | 30.0378788 | 29 | 18.9772727 | 21.6287879 | 0 | 0 | 0 | 0 | 0 | 0 | 0 | 0 | 0 | 0 | 1 | 0 | 2 | 0 | 3 |
| ENSMUSG00000034653 | Ythdc2 | ENSMUST00000201507 | 31.5955766 | 34 | 16.7851501 | 17.4960506 | 0 | 0 | 0 | 1 | 0 | 0 | 0 | 0 | 1 | 2 | 1 | 0 | 0 | 0 | 3 |
| ENSMUSG00000047213 | Ythdf3 | ENSMUST00000108345 | 30.3040209 | 38 | 15.2991174 | 16.7048055 | 0 | 0 | 0 | 0 | 0 | 0 | 0 | 0 | 0 | 0 | 0 | 0 | 0 | 2 | 2 |
| ENSMUSG00000020849 | Ywhae | ENSMUST00000067664 | 27.2804774 | 34 | 21.5686275 | 17.3913043 | 0 | 0 | 0 | 0 | 0 | 0 | 0 | 0 | 0 | 0 | 0 | 4 | 1 | 0 | 5 |
| ENSMUSG00000022285 | Ywhaz | ENSMUST00000022894 | 24.7946612 | 35 | 20.8932238 | 18.9938398 | 0 | 0 | 0 | 0 | 0 | 0 | 0 | 0 | 0 | 0 | 0 | 0 | 5 | 2 | 7 |
| ENSMUSG00000021264 | Yy1 | ENSMUST00000021692 | 31.5250768 | 36 | 15.1484135 | 17.8096213 | 0 | 0 | 0 | 1 | 0 | 0 | 0 | 0 | 1 | 2 | 0 | 0 | 0 | 0 | 2 |
| ENSMUSG00000091736 | Yy2 | ENSMUST00000065806 | 29.2134831 | 40 | 13.3509584 | 17.7792465 | 0 | 0 | 0 | 0 | 0 | 0 | 0 | 0 | 0 | 0 | 0 | 0 | 2 | 0 | 2 |
| ENSMUSG00000049090 | Zadh2 | ENSMUST00000060223 | 28.8990826 | 28 | 19.6330275 | 23.440367 | 0 | 0 | 0 | 0 | 0 | 0 | 0 | 0 | 0 | 0 | 0 | 0 | 0 | 2 | 2 |

| | | | | | | | | | | | | | | | | | | | | | |
|---|---|---|---|---|---|---|---|---|---|---|---|---|---|---|---|---|---|---|---|---|---|
| ENSMUSG00000033454 | Zbtb1 | ENSMUST00000042779 | 30.6574684 | 31 | 18.6683391 | 20.0794397 | 2 | 1 | 0 | 3 | 0 | 0 | 0 | 1 | 2 | 6 | 0 | 45 | 16 | 6 | 76 |
| ENSMUSG00000069114 | Zbtb10 | ENSMUST00000155203 | 29.2093023 | 35 | 16.3255814 | 19.372093 | 1 | 0 | 1 | 1 | 0 | 0 | 0 | 0 | 1 | 3 | 1 | 0 | 2 | 1 | 8 |
| ENSMUSG00000022601 | Zbtb11 | ENSMUST00000050248 | 31.0453401 | 37 | 14.4836272 | 17.6952141 | 1 | 1 | 0 | 1 | 0 | 0 | 0 | 0 | 1 | 2 | 0 | 0 | 7 | 1 | 12 |
| ENSMUSG00000049672 | Zbtb14 | ENSMUST00000062369 | 26.0038241 | 35 | 18.3556405 | 20.6022945 | 0 | 0 | 0 | 0 | 0 | 0 | 0 | 0 | 0 | 0 | 0 | 0 | 2 | 4 | 6 |
| ENSMUSG00000063659 | Zbtb18 | ENSMUST00000193480 | 27.3900907 | 31 | 19.5394278 | 21.6329379 | 1 | 0 | 0 | 2 | 0 | 0 | 0 | 0 | 2 | 4 | 0 | 5 | 4 | 2 | 16 |
| ENSMUSG00000022708 | Zbtb20 | ENSMUST00000114694 | 29.0175203 | 29 | 21.4644983 | 20.0938888 | 0 | 0 | 0 | 1 | 0 | 0 | 0 | 0 | 1 | 2 | 6 | 20 | 15 | 10 | 53 |
| ENSMUSG00000056459 | Zbtb25 | ENSMUST00000176102 | 32.0446097 | 34 | 15.9107807 | 17.9925651 | 0 | 0 | 0 | 0 | 0 | 0 | 0 | 0 | 0 | 0 | 0 | 0 | 1 | 2 | 3 |
| ENSMUSG00000050714 | Zbtb26 | ENSMUST00000067043 | 29.5227208 | 35 | 16.833381 | 18.9482709 | 2 | 1 | 1 | 5 | 0 | 0 | 0 | 1 | 4 | 11 | 0 | 0 | 7 | 2 | 23 |
| ENSMUSG00000048047 | Zbtb33 | ENSMUST00000049740 | 29.7796731 | 35 | 16.0270078 | 18.8343994 | 0 | 0 | 0 | 0 | 0 | 0 | 0 | 0 | 0 | 0 | 0 | 2 | 2 | 0 | 4 |
| ENSMUSG00000068966 | Zbtb34 | ENSMUST00000091037 | 23.462883 | 33 | 22.5992186 | 21.3859757 | 0 | 0 | 0 | 0 | 0 | 0 | 0 | 0 | 0 | 0 | 9 | 49 | 10 | 0 | 68 |
| ENSMUSG00000043467 | Zbtb37 | ENSMUST00000162226 | 27.6251544 | 32 | 19.5364433 | 20.9718219 | 0 | 0 | 0 | 0 | 0 | 0 | 0 | 0 | 0 | 0 | 4 | 14 | 32 | 3 | 53 |
| ENSMUSG00000033964 | Zbtb41 | ENSMUST00000039867 | 30.2100457 | 37 | 14.9406393 | 18.3013699 | 0 | 0 | 0 | 1 | 0 | 0 | 0 | 0 | 1 | 2 | 6 | 0 | 6 | 1 | 15 |
| ENSMUSG00000026788 | Zbtb43 | ENSMUST00000028125 | 24.2228739 | 29 | 22.9032258 | 24.2521994 | 0 | 0 | 0 | 0 | 0 | 0 | 0 | 0 | 0 | 0 | 0 | 0 | 1 | 2 | 3 |
| ENSMUSG00000047412 | Zbtb44 | ENSMUST00000115222 | 28.2004492 | 34 | 17.1673217 | 20.3677709 | 0 | 0 | 2 | 4 | 0 | 0 | 0 | 0 | 4 | 10 | 6 | 0 | 42 | 0 | 58 |
| ENSMUSG00000066798 | Zbtb6 | ENSMUST00000053098 | 27.9987273 | 36 | 16.9901368 | 18.5491569 | 2 | 0 | 0 | 0 | 0 | 0 | 0 | 0 | 0 | 0 | 1 | 0 | 13 | 1 | 17 |
| ENSMUSG00000043542 | Zc2hc1a | ENSMUST00000051064 | 29.458042 | 36 | 17.0891608 | 17.0017483 | 1 | 1 | 1 | 0 | 0 | 0 | 0 | 0 | 0 | 1 | 0 | 0 | 0 | 0 | 3 |
| ENSMUSG00000035164 | Zc3h12c | ENSMUST00000165519 | 29.7281497 | 34 | 16.1064016 | 19.8479977 | 0 | 0 | 0 | 1 | 0 | 0 | 0 | 0 | 1 | 2 | 1 | 0 | 5 | 2 | 10 |
| ENSMUSG00000027091 | Zc3h15 | ENSMUST00000081591 | 28.1045752 | 31 | 19.6078431 | 21.0784314 | 0 | 0 | 1 | 0 | 0 | 0 | 0 | 0 | 0 | 1 | 0 | 1 | 0 | 0 | 2 |
| ENSMUSG00000075600 | Zc3h3 | ENSMUST00000100538 | 16.2217659 | 23 | 38.1930185 | 22.587269 | 0 | 0 | 0 | 0 | 0 | 0 | 0 | 0 | 0 | 0 | 0 | 5 | 0 | 0 | 5 |
| ENSMUSG00000042851 | Zc3h6 | ENSMUST00000110320 | 32.0735444 | 39 | 14.6067416 | 14.7088866 | 1 | 0 | 1 | 0 | 0 | 0 | 0 | 0 | 0 | 1 | 0 | 0 | 0 | 1 | 3 |
| ENSMUSG00000027387 | Zc3h8 | ENSMUST00000028866 | 26.5822785 | 29 | 17.9023508 | 26.9439421 | 0 | 0 | 1 | 0 | 0 | 0 | 0 | 0 | 0 | 1 | 0 | 0 | 0 | 0 | 1 |
| ENSMUSG00000047749 | Zc3hav1l | ENSMUST00000058524 | 22.8581585 | 34 | 22.2715962 | 20.7607536 | 0 | 0 | 0 | 3 | 0 | 0 | 0 | 0 | 3 | 6 | 5 | 17 | 10 | 2 | 40 |
| ENSMUSG00000035062 | Zc4h2 | ENSMUST00000120620 | 26.0780287 | 34 | 20.2600958 | 19.4387406 | 0 | 0 | 0 | 0 | 0 | 0 | 0 | 0 | 0 | 0 | 0 | 2 | 0 | 1 | 3 |
| ENSMUSG00000018239 | Zcchc10 | ENSMUST00000018383 | 29.6610169 | 37 | 14.9717514 | 18.7853107 | 0 | 0 | 0 | 0 | 0 | 0 | 0 | 0 | 0 | 0 | 0 | 0 | 0 | 1 | 1 |
| ENSMUSG00000034610 | Zcchc11 | ENSMUST00000097925 | 34.2767296 | 38 | 13.6792453 | 13.6792453 | 0 | 0 | 0 | 0 | 0 | 0 | 0 | 0 | 0 | 0 | 0 | 0 | 1 | 0 | 1 |
| ENSMUSG00000029179 | Zcchc4 | ENSMUST00000031077 | 26.637869 | 30 | 20.7343413 | 22.2462203 | 0 | 0 | 1 | 1 | 0 | 0 | 0 | 0 | 1 | 3 | 0 | 1 | 1 | 0 | 5 |
| ENSMUSG00000035649 | Zcchc7 | ENSMUST00000142714 | 31.7375887 | 34 | 18.2624113 | 16.2529551 | 0 | 0 | 0 | 0 | 0 | 0 | 0 | 0 | 0 | 0 | 0 | 1 | 0 | 0 | 1 |
| ENSMUSG00000029427 | Zcchc8 | ENSMUST00000196282 | 25.0243902 | 32 | 20.2439024 | 22.3902439 | 1 | 1 | 0 | 0 | 0 | 0 | 0 | 0 | 0 | 0 | 0 | 0 | 1 | 3 | 6 |
| ENSMUSG00000021621 | Zcchc9 | ENSMUST00000022121 | 28.026534 | 35 | 17.5787728 | 19.5688226 | 0 | 1 | 0 | 1 | 0 | 0 | 0 | 1 | 0 | 2 | 0 | 0 | 0 | 0 | 3 |
| ENSMUSG00000027520 | Zdbf2 | ENSMUST00000114132 | 29.5184625 | 36 | 16.4759231 | 17.6635716 | 2 | 0 | 0 | 1 | 0 | 0 | 0 | 0 | 1 | 2 | 1 | 7 | 4 | 3 | 19 |
| ENSMUSG00000033906 | Zdhhc15 | ENSMUST00000042070 | 30.4018913 | 30 | 20.141844 | 19.2434988 | 0 | 0 | 0 | 0 | 0 | 0 | 0 | 0 | 0 | 0 | 0 | 3 | 0 | 0 | 3 |
| ENSMUSG00000035798 | Zdhhc17 | ENSMUST00000041723 | 28.2650326 | 33 | 17.617771 | 21.2179242 | 0 | 0 | 0 | 0 | 0 | 0 | 0 | 0 | 0 | 0 | 0 | 4 | 1 | 0 | 5 |
| ENSMUSG00000039470 | Zdhhc2 | ENSMUST00000049389 | 29.3859649 | 34 | 17.1539961 | 19.0545809 | 0 | 0 | 0 | 0 | 0 | 0 | 0 | 0 | 0 | 0 | 3 | 0 | 5 | 1 | 9 |
| ENSMUSG00000028403 | Zdhhc21 | ENSMUST00000030110 | 29.811619 | 33 | 17.7446977 | 19.404558 | 1 | 0 | 1 | 4 | 0 | 0 | 0 | 0 | 4 | 9 | 1 | 1 | 0 | 1 | 13 |
| ENSMUSG00000026872 | Zeb2 | ENSMUST00000176438 | 32.7261688 | 34 | 17.0613236 | 16.1708156 | 1 | 0 | 0 | 2 | 0 | 0 | 0 | 0 | 2 | 4 | 0 | 2 | 3 | 2 | 12 |
| ENSMUSG00000042213 | Zfand4 | ENSMUST00000112900 | 30.4668305 | 39 | 14.3734644 | 16.0933661 | 0 | 0 | 0 | 0 | 0 | 0 | 0 | 0 | 0 | 0 | 0 | 0 | 1 | 0 | 1 |
| ENSMUSG00000030629 | Zfand6 | ENSMUST00000069537 | 26.5013055 | 39 | 15.4046997 | 19.4516971 | 0 | 0 | 0 | 0 | 0 | 0 | 0 | 0 | 0 | 0 | 0 | 0 | 4 | 1 | 5 |
| ENSMUSG00000025255 | Zfhx4 | ENSMUST00000176383 | 30.9225092 | 34 | 16.900369 | 18.597786 | 1 | 0 | 1 | 1 | 0 | 0 | 0 | 0 | 1 | 3 | 0 | 0 | 2 | 0 | 6 |
| ENSMUSG00000055240 | Zfp101 | ENSMUST00000174512 | 28.1632653 | 30 | 22.8571429 | 19.4460641 | 0 | 0 | 0 | 0 | 0 | 0 | 0 | 0 | 0 | 0 | 0 | 0 | 0 | 1 | 1 |
| ENSMUSG00000051034 | Zfp11 | ENSMUST00000049778 | 29.0478752 | 33 | 18.6659494 | 19.6342119 | 4 | 2 | 0 | 5 | 0 | 1 | 1 | 2 | 2 | 11 | 0 | 0 | 1 | 2 | 20 |
| ENSMUSG00000087598 | Zfp111 | ENSMUST00000086006 | 27.4727932 | 34 | 20.1451028 | 18.6457074 | 0 | 0 | 0 | 0 | 0 | 0 | 0 | 0 | 0 | 0 | 0 | 15 | 9 | 4 | 28 |
| ENSMUSG00000068134 | Zfp120 | ENSMUST00000122859 | 34.4258065 | 28 | 16.8 | 20.4129032 | 0 | 1 | 0 | 1 | 0 | 0 | 0 | 0 | 1 | 2 | 0 | 1 | 0 | 0 | 4 |
| ENSMUSG00000092416 | Zfp141 | ENSMUST00000174407 | 30.7548347 | 32 | 18.0910792 | 19.588272 | 0 | 0 | 0 | 1 | 0 | 0 | 0 | 0 | 1 | 2 | 0 | 0 | 4 | 0 | 6 |
| ENSMUSG00000022811 | Zfp148 | ENSMUST00000165418 | 28.5561823 | 35 | 17.8380851 | 18.9967983 | 0 | 0 | 0 | 0 | 0 | 0 | 0 | 0 | 0 | 0 | 0 | 0 | 7 | 3 | 10 |
| ENSMUSG00000067942 | Zfp160 | ENSMUST00000088811 | 33.7426901 | 32 | 14.8538012 | 18.9473684 | 0 | 1 | 0 | 0 | 0 | 0 | 0 | 0 | 0 | 0 | 0 | 0 | 0 | 0 | 1 |
| ENSMUSG00000054737 | Zfp182 | ENSMUST00000115333 | 37.0666667 | 29 | 14.6666667 | 19.6444444 | 0 | 0 | 0 | 1 | 0 | 0 | 0 | 0 | 1 | 2 | 0 | 0 | 0 | 0 | 2 |
| ENSMUSG00000017421 | Zfp207 | ENSMUST00000123726 | 27.7210129 | 35 | 18.2141271 | 19.3247446 | 0 | 0 | 0 | 0 | 0 | 0 | 0 | 0 | 0 | 0 | 0 | 0 | 12 | 4 | 16 |
| ENSMUSG00000061544 | Zfp229 | ENSMUST00000182603 | 32.1688501 | 33 | 17.515769 | 17.5642892 | 0 | 0 | 0 | 1 | 0 | 0 | 0 | 0 | 1 | 2 | 2 | 0 | 8 | 0 | 12 |
| ENSMUSG00000051469 | Zfp24 | ENSMUST00000066497 | 29.8312711 | 32 | 17.4353206 | 20.4949381 | 0 | 0 | 0 | 1 | 0 | 0 | 0 | 0 | 1 | 2 | 0 | 0 | 0 | 3 | 5 |
| ENSMUSG00000030145 | Zfp248 | ENSMUST00000069292 | 30.7545973 | 32 | 20.0380469 | 16.7406468 | 0 | 0 | 0 | 0 | 0 | 0 | 0 | 0 | 0 | 0 | 0 | 32 | 0 | 0 | 32 |
| ENSMUSG00000063108 | Zfp26 | ENSMUST00000159569 | 24.7144949 | 41 | 16.6617862 | 17.8916545 | 1 | 1 | 0 | 0 | 0 | 0 | 0 | 0 | 0 | 0 | 3 | 0 | 10 | 1 | 16 |
| ENSMUSG00000060510 | Zfp266 | ENSMUST00000174462 | 27.995997 | 34 | 18.5389042 | 19.9892419 | 0 | 0 | 0 | 0 | 0 | 0 | 0 | 0 | 0 | 0 | 0 | 0 | 11 | 0 | 11 |
| ENSMUSG00000031365 | Zfp275 | ENSMUST00000033731 | 26.2295082 | 31 | 21.8650202 | 20.459868 | 1 | 1 | 0 | 2 | 0 | 0 | 0 | 0 | 2 | 4 | 0 | 1 | 0 | 1 | 8 |
| ENSMUSG00000049764 | Zfp280b | ENSMUST00000061617 | 22.867405 | 33 | 22.3286441 | 22.2089195 | 0 | 0 | 0 | 0 | 0 | 0 | 0 | 0 | 0 | 0 | 0 | 1 | 6 | 0 | 7 |
| ENSMUSG00000036916 | Zfp280c | ENSMUST00000114940 | 26.1336516 | 39 | 17.4224344 | 17.3031026 | 0 | 0 | 0 | 0 | 0 | 0 | 0 | 0 | 0 | 0 | 1 | 0 | 6 | 0 | 7 |
| ENSMUSG00000038535 | Zfp280d | ENSMUST00000184053 | 33.9309304 | 29 | 18.2864691 | 18.3808266 | 0 | 0 | 1 | 0 | 0 | 0 | 0 | 0 | 0 | 1 | 0 | 0 | 0 | 0 | 1 |
| ENSMUSG00000041483 | Zfp281 | ENSMUST00000047734 | 31.1121764 | 36 | 15.915628 | 17.0182167 | 0 | 0 | 0 | 0 | 0 | 0 | 0 | 0 | 0 | 0 | 0 | 2 | 2 | 0 | 4 |
| ENSMUSG00000031079 | Zfp300 | ENSMUST00000040667 | 29.3946932 | 33 | 19.0713101 | 18.6567164 | 0 | 0 | 0 | 1 | 0 | 0 | 0 | 0 | 1 | 2 | 0 | 2 | 0 | 0 | 4 |
| ENSMUSG00000046351 | Zfp322a | ENSMUST00000050101 | 30.1081917 | 32 | 16.9088099 | 20.8037094 | 3 | 1 | 0 | 2 | 0 | 0 | 0 | 0 | 2 | 4 | 0 | 0 | 5 | 0 | 13 |
| ENSMUSG00000029290 | Zfp326 | ENSMUST00000138615 | 35.0194553 | 21 | 16.3424125 | 27.3151751 | 1 | 0 | 0 | 2 | 0 | 0 | 0 | 0 | 2 | 4 | 0 | 0 | 0 | 0 | 5 |
| ENSMUSG00000057894 | Zfp329 | ENSMUST00000072222 | 29.395441 | 32 | 17.185332 | 21.0901883 | 3 | 1 | 0 | 3 | 0 | 0 | 0 | 0 | 3 | 6 | 4 | 16 | 12 | 1 | 43 |
| ENSMUSG00000031711 | Zfp330 | ENSMUST00000034147 | 27.3026316 | 33 | 18.2565789 | 21.7105263 | 1 | 0 | 0 | 0 | 0 | 0 | 0 | 0 | 0 | 0 | 0 | 0 | 0 | 0 | 1 |
| ENSMUSG00000044807 | Zfp354c | ENSMUST00000109135 | 30.2185164 | 32 | 18.1713629 | 19.5227142 | 0 | 0 | 0 | 0 | 0 | 0 | 0 | 0 | 0 | 0 | 0 | 2 | 0 | 0 | 2 |
| ENSMUSG00000044786 | Zfp36 | ENSMUST00000051241 | 23.9018088 | 32 | 21.9638243 | 21.7054264 | 0 | 0 | 1 | 0 | 0 | 0 | 0 | 0 | 0 | 1 | 0 | 0 | 0 | 0 | 1 |
| ENSMUSG00000044934 | Zfp367 | ENSMUST00000059817 | 30.3664921 | 35 | 13.8743455 | 21.1605585 | 0 | 0 | 0 | 0 | 0 | 0 | 0 | 0 | 0 | 0 | 0 | 0 | 2 | 2 | 4 |
| ENSMUSG00000021127 | Zfp36l1 | ENSMUST00000021552 | 27.7747403 | 32 | 22.6899945 | 17.222526 | 1 | 1 | 0 | 1 | 0 | 0 | 0 | 0 | 1 | 2 | 0 | 2 | 1 | 1 | 8 |
| ENSMUSG00000045817 | Zfp36l2 | ENSMUST00000060366 | 27.5521405 | 32 | 18.3863886 | 21.7343578 | 1 | 0 | 0 | 0 | 0 | 0 | 0 | 0 | 0 | 0 | 0 | 0 | 3 | 0 | 4 |
| ENSMUSG00000028389 | Zfp37 | ENSMUST00000068822 | 33.9901478 | 32 | 16.2561576 | 17.8571429 | 0 | 0 | 0 | 0 | 0 | 0 | 0 | 0 | 0 | 0 | 0 | 0 | 1 | 0 | 1 |
| ENSMUSG00000038346 | Zfp384 | ENSMUST00000046064 | 24.3668122 | 33 | 22.0960699 | 20.7860262 | 0 | 0 | 0 | 0 | 0 | 0 | 0 | 0 | 0 | 0 | 0 | 3 | 0 | 0 | 3 |
| ENSMUSG00000042063 | Zfp386 | ENSMUST00000183125 | 30.6818182 | 29 | 18.5724432 | 22.0525568 | 0 | 0 | 0 | 0 | 0 | 0 | 0 | 0 | 0 | 0 | 1 | 0 | 2 | 0 | 3 |
| ENSMUSG00000034522 | Zfp395 | ENSMUST00000066994 | 21.5769231 | 29 | 26.9230769 | 22.8846154 | 0 | 0 | 0 | 0 | 0 | 0 | 0 | 0 | 0 | 0 | 0 | 60 | 0 | 2 | 62 |
| ENSMUSG00000062519 | Zfp398 | ENSMUST00000114598 | 28.1487102 | 29 | 21.7147193 | 21.047041 | 0 | 0 | 1 | 0 | 0 | 0 | 0 | 0 | 0 | 1 | 6 | 3 | 14 | 3 | 27 |
| ENSMUSG00000002617 | Zfp40 | ENSMUST00000037057 | 34.6153846 | 30 | 18.1952663 | 17.2337278 | 0 | 0 | 0 | 0 | 0 | 0 | 0 | 0 | 0 | 0 | 0 | 0 | 0 | 2 | 2 |
| ENSMUSG00000059878 | Zfp422 | ENSMUST00000079749 | 25.2709146 | 35 | 20.4594712 | 18.8556567 | 0 | 0 | 0 | 0 | 0 | 0 | 0 | 0 | 0 | 0 | 5 | 3 | 18 | 0 | 26 |
| ENSMUSG00000068130 | Zfp442 | ENSMUST00000109916 | 33.9712919 | 31 | 15.430622 | 19.2583732 | 0 | 0 | 0 | 1 | 0 | 0 | 0 | 0 | 1 | 2 | 0 | 0 | 1 | 0 | 3 |
| ENSMUSG00000073176 | Zfp449 | ENSMUST00000101560 | 29.3313812 | 33 | 17.0815032 | 20.2537823 | 0 | 0 | 0 | 0 | 0 | 0 | 0 | 0 | 0 | 0 | 0 | 2 | 0 | 0 | 2 |
| ENSMUSG00000042197 | Zfp451 | ENSMUST00000044455 | 25.6109776 | 35 | 18.760016 | 20.7732372 | 7 | 3 | 0 | 4 | 0 | 0 | 0 | 1 | 3 | 8 | 11 | 4 | 21 | 4 | 58 |
| ENSMUSG00000051037 | Zfp455 | ENSMUST00000117110 | 33.0651649 | 34 | 16.9750603 | 16.0901046 | 0 | 0 | 0 | 1 | 0 | 0 | 0 | 0 | 1 | 2 | 0 | 1 | 0 | 0 | 3 |
| ENSMUSG00000039081 | Zfp503 | ENSMUST00000043409 | 34.3412527 | 28 | 16.5226782 | 21.1663067 | 1 | 1 | 2 | 0 | 0 | 0 | 0 | 0 | 0 | 2 | 0 | 0 | 0 | 0 | 4 |
| ENSMUSG00000049164 | Zfp518a | ENSMUST00000050092 | 30.9245484 | 36 | 14.7183847 | 18.0127524 | 0 | 0 | 0 | 0 | 0 | 0 | 0 | 0 | 0 | 0 | 2 | 1 | 4 | 0 | 7 |
| ENSMUSG00000023882 | Zfp54 | ENSMUST00000165230 | 29.8273155 | 35 | 18.2103611 | 17.4254317 | 0 | 0 | 0 | 1 | 0 | 0 | 0 | 0 | 1 | 2 | 0 | 0 | 1 | 0 | 3 |
| ENSMUSG00000045519 | Zfp560 | ENSMUST00000068079 | 27.5415896 | 37 | 19.3622921 | 16.4510166 | 1 | 0 | 0 | 1 | 0 | 0 | 0 | 0 | 1 | 2 | 1 | 7 | 1 | 0 | 12 |
| ENSMUSG00000030443 | Zfp583 | ENSMUST00000062765 | 28.6821705 | 35 | 22.9457364 | 13.3333333 | 0 | 0 | 0 | 0 | 0 | 0 | 0 | 0 | 0 | 0 | 0 | 12 | 0 | 0 | 12 |
| ENSMUSG00000057842 | Zfp595 | ENSMUST00000109735 | 28.3067422 | 32 | 19.4544519 | 20.0720535 | 0 | 0 | 0 | 0 | 0 | 0 | 0 | 0 | 0 | 0 | 1 | 0 | 2 | 0 | 3 |
| ENSMUSG00000039789 | Zfp597 | ENSMUST00000090522 | 28.978796 | 33 | 17.0363149 | 20.545942 | 0 | 0 | 0 | 0 | 0 | 0 | 0 | 0 | 0 | 0 | 0 | 0 | 8 | 3 | 11 |
| ENSMUSG00000037640 | Zfp60 | ENSMUST00000108336 | 30.7148682 | 31 | 18.1027453 | 20.3316119 | 0 | 0 | 0 | 1 | 0 | 0 | 0 | 0 | 1 | 2 | 0 | 1 | 1 | 0 | 4 |
| ENSMUSG00000030386 | Zfp606 | ENSMUST00000098822 | 33.2103321 | 31 | 15.8671587 | 20.3479178 | 0 | 0 | 0 | 0 | 0 | 0 | 0 | 0 | 0 | 0 | 0 | 0 | 4 | 1 | 5 |
| ENSMUSG00000052713 | Zfp608 | ENSMUST00000064763 | 30.6796117 | 37 | 16.1165049 | 16.6990291 | 0 | 0 | 0 | 0 | 0 | 0 | 0 | 0 | 0 | 0 | 0 | 0 | 2 | 1 | 3 |
| ENSMUSG00000030016 | Zfp638 | ENSMUST00000204751 | 30.0990776 | 32 | 18.8247352 | 19.3372053 | 0 | 0 | 0 | 0 | 0 | 0 | 0 | 0 | 0 | 0 | 0 | 0 | 1 | 0 | 1 |
| ENSMUSG00000047141 | Zfp654 | ENSMUST00000052588 | 33.1391364 | 35 | 15.3872515 | 16.5524332 | 1 | 0 | 0 | 0 | 0 | 0 | 0 | 0 | 0 | 0 | 0 | 0 | 1 | 0 | 2 |
| ENSMUSG00000034800 | Zfp661 | ENSMUST00000110366 | 28.024637 | 29 | 22.0853498 | 20.941487 | 0 | 0 | 0 | 0 | 0 | 0 | 0 | 0 | 0 | 0 | 0 | 1 | 0 | 0 | 1 |
| ENSMUSG00000058291 | Zfp68 | ENSMUST00000110905 | 32.3154233 | 30 | 17.5492849 | 19.6366448 | 1 | 0 | 0 | 1 | 0 | 0 | 0 | 0 | 1 | 2 | 2 | 0 | 0 | 1 | 6 |

| | | | | | | | | | | | | | | | | | | | | | |
|---|---|---|---|---|---|---|---|---|---|---|---|---|---|---|---|---|---|---|---|---|---|
| ENSMUSG00000062397 | Zfp706 | ENSMUST00000078976 | 25.6081946 | 34 | 19.8207426 | 20.9731114 | 0 | 0 | 0 | 0 | 0 | 0 | 0 | 0 | 0 | 0 | 2 | 6 | 2 | 0 | 10 |
| ENSMUSG00000058883 | Zfp708 | ENSMUST00000109742 | 38.3116883 | 29 | 15.7142857 | 16.4935065 | 0 | 0 | 0 | 0 | 0 | 0 | 0 | 0 | 0 | 0 | 0 | 1 | 1 | 0 | 2 |
| ENSMUSG00000100235 | Zfp708 | ENSMUST00000186303 | 34.948605 | 27 | 19.7136564 | 18.5389134 | 0 | 0 | 0 | 0 | 0 | 0 | 0 | 0 | 0 | 0 | 0 | 1 | 1 | 0 | 2 |
| ENSMUSG00000056019 | Zfp709 | ENSMUST00000188685 | 28.8888889 | 32 | 20.2284528 | 18.7123572 | 0 | 0 | 0 | 0 | 0 | 0 | 0 | 0 | 0 | 0 | 1 | 0 | 13 | 0 | 14 |
| ENSMUSG00000025529 | Zfp711 | ENSMUST00000071814 | 33.8837517 | 35 | 13.2100396 | 17.5693527 | 0 | 0 | 0 | 1 | 0 | 0 | 0 | 0 | 1 | 2 | 0 | 0 | 0 | 0 | 2 |
| ENSMUSG00000090641 | Zfp712 | ENSMUST00000167565 | 32.0571429 | 35 | 14.6285714 | 17.8285714 | 3 | 0 | 0 | 1 | 0 | 0 | 0 | 0 | 1 | 2 | 1 | 0 | 0 | 0 | 6 |
| ENSMUSG00000048280 | Zfp738 | ENSMUST00000137496 | 34.1617934 | 28 | 19.4444444 | 18.2261209 | 0 | 0 | 0 | 0 | 0 | 0 | 0 | 0 | 0 | 0 | 1 | 0 | 0 | 0 | 1 |
| ENSMUSG00000095432 | Zfp748 | ENSMUST00000181892 | 31.1135371 | 33 | 19.0502183 | 17.139738 | 0 | 0 | 0 | 0 | 0 | 0 | 0 | 0 | 0 | 0 | 0 | 0 | 1 | 0 | 1 |
| ENSMUSG00000057396 | Zfp759 | ENSMUST00000052716 | 33.0364372 | 31 | 17.0040486 | 19.1093117 | 0 | 0 | 0 | 0 | 0 | 0 | 0 | 0 | 0 | 0 | 1 | 0 | 0 | 0 | 1 |
| ENSMUSG00000067928 | Zfp760 | ENSMUST00000073312 | 32.9050279 | 33 | 15.9776536 | 18.2122905 | 0 | 0 | 0 | 0 | 0 | 0 | 0 | 0 | 0 | 0 | 0 | 0 | 4 | 2 | 6 |
| ENSMUSG00000067430 | Zfp763 | ENSMUST00000087654 | 30.3246753 | 34 | 15.7142857 | 20.1948052 | 0 | 0 | 0 | 0 | 0 | 0 | 0 | 0 | 0 | 0 | 1 | 0 | 2 | 4 | 7 |
| ENSMUSG00000040321 | Zfp770 | ENSMUST00000050668 | 30.5278884 | 33 | 16.4840637 | 19.6713147 | 0 | 0 | 0 | 0 | 0 | 0 | 0 | 0 | 0 | 0 | 0 | 2 | 1 | 0 | 3 |
| ENSMUSG00000074165 | Zfp788 | ENSMUST00000098508 | 29.0149893 | 34 | 16.2740899 | 20.7708779 | 0 | 0 | 0 | 0 | 0 | 0 | 0 | 0 | 0 | 0 | 0 | 1 | 10 | 2 | 13 |
| ENSMUSG00000095253 | Zfp799 | ENSMUST00000202988 | 26.8035944 | 33 | 20.3080873 | 19.8459564 | 2 | 1 | 0 | 1 | 0 | 0 | 0 | 0 | 1 | 2 | 4 | 2 | 4 | 0 | 15 |
| ENSMUSG00000039841 | Zfp800 | ENSMUST00000115320 | 34.0462428 | 39 | 13.1791908 | 14.1618497 | 0 | 0 | 0 | 2 | 0 | 0 | 0 | 0 | 2 | 4 | 0 | 0 | 0 | 0 | 4 |
| ENSMUSG00000003929 | Zfp81 | ENSMUST00000054072 | 31.0140788 | 33 | 17.3842073 | 18.7920832 | 2 | 0 | 0 | 1 | 0 | 0 | 0 | 0 | 1 | 2 | 1 | 0 | 1 | 0 | 6 |
| ENSMUSG00000046010 | Zfp830 | ENSMUST00000056677 | 28.8135593 | 35 | 16.9491525 | 18.8559322 | 1 | 0 | 0 | 1 | 0 | 0 | 0 | 0 | 1 | 2 | 0 | 0 | 0 | 0 | 3 |
| ENSMUSG00000046185 | Zfp84 | ENSMUST00000032802 | 28.4448383 | 29 | 20.248117 | 22.2862206 | 0 | 0 | 0 | 0 | 0 | 0 | 0 | 0 | 0 | 0 | 0 | 1 | 2 | 0 | 3 |
| ENSMUSG00000024298 | Zfp871 | ENSMUST00000159086 | 27.4390244 | 34 | 18.5815148 | 20.3145058 | 0 | 0 | 0 | 0 | 0 | 0 | 0 | 0 | 0 | 0 | 0 | 0 | 2 | 0 | 2 |
| ENSMUSG00000069206 | Zfp874a | ENSMUST00000180580 | 32.1357285 | 30 | 18.4298071 | 19.6606786 | 0 | 0 | 0 | 0 | 0 | 0 | 0 | 0 | 0 | 0 | 0 | 1 | 0 | 0 | 1 |
| ENSMUSG00000059839 | Zfp874b | ENSMUST00000019572 | 30.4310345 | 30 | 19.9137931 | 19.9568966 | 0 | 0 | 0 | 0 | 0 | 0 | 0 | 0 | 0 | 0 | 0 | 1 | 6 | 0 | 7 |
| ENSMUSG00000072623 | Zfp9 | ENSMUST00000161170 | 29.8951782 | 30 | 20.1677149 | 19.9161426 | 1 | 0 | 0 | 0 | 0 | 0 | 0 | 0 | 0 | 0 | 0 | 0 | 0 | 2 | 3 |
| ENSMUSG00000078861 | Zfp931 | ENSMUST00000108923 | 34.401508 | 34 | 14.7031103 | 16.9651272 | 0 | 0 | 1 | 0 | 0 | 0 | 0 | 0 | 0 | 1 | 0 | 0 | 0 | 0 | 1 |
| ENSMUSG00000059423 | Zfp933 | ENSMUST00000105718 | 27.2131148 | 32 | 19.9635701 | 21.0200364 | 3 | 1 | 0 | 6 | 0 | 0 | 1 | 2 | 3 | 12 | 2 | 0 | 4 | 0 | 22 |
| ENSMUSG00000060336 | Zfp937 | ENSMUST00000073782 | 32.1858864 | 28 | 18.8468158 | 20.9982788 | 0 | 0 | 0 | 2 | 0 | 0 | 0 | 0 | 2 | 4 | 0 | 0 | 1 | 0 | 5 |
| ENSMUSG00000062931 | Zfp938 | ENSMUST00000156218 | 36.5532382 | 29 | 17.3435785 | 17.5631175 | 0 | 0 | 0 | 0 | 0 | 0 | 0 | 0 | 0 | 0 | 0 | 1 | 0 | 0 | 1 |
| ENSMUSG00000060314 | Zfp941 | ENSMUST00000106052 | 26.091954 | 32 | 20.1149425 | 22.183908 | 11 | 5 | 0 | 29 | 7 | 8 | 9 | 10 | 10 | 73 | 0 | 0 | 0 | 1 | 90 |
| ENSMUSG00000053347 | Zfp943 | ENSMUST00000153985 | 28.7709497 | 36 | 19.273743 | 16.2942272 | 1 | 1 | 0 | 0 | 0 | 0 | 0 | 0 | 0 | 0 | 0 | 0 | 1 | 1 | 4 |
| ENSMUSG00000053390 | Zfp952 | ENSMUST00000087666 | 28.6226686 | 31 | 18.1492109 | 22.6685796 | 0 | 0 | 0 | 1 | 0 | 0 | 0 | 0 | 1 | 2 | 0 | 0 | 1 | 0 | 3 |
| ENSMUSG00000052446 | Zfp961 | ENSMUST00000131237 | 28.9512555 | 31 | 21.0487445 | 19.4239291 | 0 | 0 | 0 | 1 | 0 | 0 | 0 | 0 | 1 | 2 | 0 | 0 | 0 | 0 | 2 |
| ENSMUSG00000022201 | Zfr | ENSMUST00000122941 | 28.4142395 | 37 | 16.3754045 | 18.4466019 | 0 | 0 | 0 | 0 | 0 | 0 | 0 | 0 | 0 | 0 | 0 | 1 | 0 | 0 | 1 |
| ENSMUSG00000079509 | Zfx | ENSMUST00000088102 | 28.7154862 | 35 | 16.7827131 | 19.2557023 | 1 | 0 | 0 | 1 | 0 | 0 | 0 | 0 | 1 | 2 | 0 | 0 | 0 | 1 | 4 |
| ENSMUSG00000051278 | Zgrf1 | ENSMUST00000199888 | 29.1525424 | 28 | 20.2542373 | 22.5 | 0 | 0 | 0 | 0 | 0 | 0 | 0 | 0 | 0 | 0 | 4 | 1 | 1 | 0 | 6 |
| ENSMUSG00000032368 | Zic1 | ENSMUST00000034927 | 29.3933352 | 34 | 17.4024494 | 19.1113643 | 1 | 0 | 0 | 0 | 0 | 0 | 0 | 0 | 0 | 0 | 1 | 3 | 2 | 1 | 8 |
| ENSMUSG00000061524 | Zic2 | ENSMUST00000075888 | 26.1083744 | 37 | 21.5106732 | 15.5993432 | 0 | 0 | 0 | 0 | 0 | 0 | 0 | 0 | 0 | 0 | 0 | 2 | 7 | 3 | 12 |
| ENSMUSG00000067860 | Zic3 | ENSMUST00000088627 | 30.7462687 | 36 | 15.5721393 | 18.159204 | 0 | 0 | 0 | 0 | 0 | 0 | 0 | 0 | 0 | 0 | 1 | 0 | 2 | 0 | 3 |
| ENSMUSG00000036972 | Zic4 | ENSMUST00000066384 | 27.7995023 | 29 | 20.6541059 | 22.32492 | 0 | 0 | 0 | 0 | 0 | 0 | 0 | 0 | 0 | 0 | 1 | 1 | 0 | 0 | 2 |
| ENSMUSG00000041703 | Zic5 | ENSMUST00000039118 | 31.5996848 | 33 | 16.5090623 | 18.9519307 | 1 | 0 | 1 | 2 | 0 | 0 | 0 | 0 | 2 | 5 | 0 | 0 | 5 | 0 | 11 |
| ENSMUSG00000030393 | Zik1 | ENSMUST00000032551 | 28.2574568 | 35 | 17.2291994 | 19.2700157 | 0 | 0 | 0 | 0 | 0 | 0 | 0 | 0 | 0 | 0 | 0 | 0 | 1 | 1 | 2 |
| ENSMUSG00000021327 | Zkscan3 | ENSMUST00000117721 | 29.0600129 | 27 | 24.6504625 | 19.2944719 | 0 | 0 | 0 | 0 | 0 | 0 | 0 | 0 | 0 | 0 | 0 | 1 | 0 | 0 | 1 |
| ENSMUSG00000063894 | Zkscan8 | ENSMUST00000045228 | 28.4702759 | 34 | 19.9144417 | 18.0262576 | 0 | 0 | 0 | 0 | 0 | 0 | 0 | 0 | 0 | 0 | 0 | 2 | 5 | 0 | 7 |
| ENSMUSG00000052676 | Zmat1 | ENSMUST00000113185 | 32.3018868 | 34 | 15.245283 | 18.7169811 | 0 | 0 | 0 | 0 | 0 | 0 | 0 | 0 | 0 | 0 | 6 | 1 | 4 | 0 | 11 |
| ENSMUSG00000043207 | Zmpste24 | ENSMUST00000058754 | 28.3683911 | 33 | 21.0915293 | 17.7942013 | 0 | 0 | 0 | 0 | 0 | 0 | 0 | 0 | 0 | 0 | 0 | 2 | 0 | 0 | 2 |
| ENSMUSG00000021945 | Zmym2 | ENSMUST00000022511 | 28.6302395 | 35 | 16.8038922 | 19.1242515 | 1 | 0 | 1 | 0 | 0 | 0 | 0 | 0 | 0 | 1 | 2 | 0 | 2 | 0 | 6 |
| ENSMUSG00000040123 | Zmym5 | ENSMUST00000039812 | 27.9786231 | 30 | 18.8934297 | 23.1688148 | 0 | 0 | 0 | 0 | 0 | 0 | 0 | 0 | 0 | 0 | 4 | 0 | 4 | 0 | 8 |
| ENSMUSG00000036315 | Znrd1 | ENSMUST00000113669 | 21.5116279 | 35 | 28.4883721 | 14.5348837 | 0 | 0 | 0 | 0 | 0 | 0 | 0 | 0 | 0 | 0 | 0 | 1 | 0 | 0 | 1 |
| ENSMUSG00000058446 | Znrf2 | ENSMUST00000079869 | 26.9283882 | 33 | 19.1951075 | 21.3059775 | 1 | 0 | 0 | 1 | 0 | 0 | 0 | 0 | 1 | 2 | 4 | 0 | 8 | 1 | 16 |
| ENSMUSG00000041961 | Znrf3 | ENSMUST00000109867 | 26.7647059 | 31 | 21 | 20.8235294 | 2 | 1 | 0 | 1 | 0 | 0 | 0 | 0 | 1 | 2 | 0 | 0 | 20 | 2 | 27 |
| ENSMUSG00000028180 | Zranb2 | ENSMUST00000184802 | 31.708351 | 31 | 17.5922001 | 19.6693514 | 0 | 0 | 0 | 0 | 0 | 0 | 0 | 0 | 0 | 0 | 2 | 0 | 0 | 1 | 3 |
| ENSMUSG00000031370 | Zrsr2 | ENSMUST00000090840 | 29.778481 | 31 | 17.6898734 | 21.7721519 | 0 | 0 | 0 | 0 | 0 | 0 | 0 | 0 | 0 | 0 | 2 | 3 | 4 | 0 | 9 |
| ENSMUSG00000022228 | Zscan26 | ENSMUST00000032820 | 29.317852 | 30 | 20.7184325 | 19.5573295 | 0 | 0 | 0 | 0 | 0 | 0 | 0 | 0 | 0 | 0 | 0 | 0 | 18 | 0 | 18 |
| ENSMUSG00000032846 | Zswim6 | ENSMUST00000105097 | 31.0952119 | 33 | 17.3362686 | 18.1618052 | 2 | 1 | 0 | 1 | 0 | 0 | 0 | 0 | 1 | 2 | 0 | 3 | 2 | 0 | 10 |
| ENSMUSG00000039531 | Zufsp | ENSMUST00000048222 | 31.7777778 | 32 | 17.7777778 | 18 | 1 | 0 | 1 | 1 | 0 | 0 | 0 | 0 | 1 | 3 | 0 | 0 | 0 | 0 | 4 |
| ENSMUSG00000019923 | Zwint | ENSMUST00000105431 | 31.1674591 | 30 | 15.4780771 | 23.296355 | 0 | 1 | 0 | 0 | 0 | 0 | 0 | 0 | 0 | 0 | 0 | 0 | 0 | 0 | 1 |
| ENSMUSG00000073062 | Zxdb | ENSMUST00000101388 | 26.772743 | 40 | 16.464891 | 16.9491525 | 0 | 0 | 0 | 0 | 0 | 0 | 0 | 0 | 0 | 0 | 0 | 1 | 0 | 0 | 1 |
| ENSMUSG00000034636 | Zyg11b | ENSMUST00000043616 | 28.7738854 | 31 | 19.0605096 | 21.6401274 | 0 | 0 | 0 | 0 | 0 | 0 | 0 | 0 | 0 | 0 | 1 | 0 | 3 | 0 | 4 |

3'UTR sequences were retrieved from Ensembl BioMart and datasets were made non-redundant by Gene ID. The datasets were analyzed by scripts written in Perl v5.8.8 and using BioPerl modules as described in Bolognani et al, 2010, https://doi.org/10.1093/nar/gkp863

**Supplemental Table S5: RTddPCR validation of KHSRP-target mRNAs in neocortex.**
Values significantly different from *Khsrp-/-* mice are shown in bold blue font.

| mRNA | *Khsrp* $^{+/+}$ | *Khsrp* $^{+/-}$ | *Khsrp* $^{-/-}$ |
|---|---|---|---|
| *Arhgap5* | 70.39 ± 5.94 | 89.37 ± 22.07 | **139.5 ± 10.7**<br>***p = 0.0048*** |
| *Atrx* | 36.82 ± 5.43 | 34.30 ± 7.84 | **64.41 ± 2.38**<br>***p = 0.009*** |
| *Cacna2d1* | 29.46 ± 5.52 | 19.65 ± 6.08 | 30.93 ± 4.86 |
| *Cadps* | 70.24 ± 13.18 | 73.51 ± 22.82 | **149.4 ± 8.0**<br>***p = 0.0068*** |
| *Calb1* | 78.88 ± 14.73 | 91.78 ± 21.61 | 135.0 ± 21.0 |
| *Chm* | 30.37 ± 2.09 | 39.50 ± 8.35 | **56.52 ± 8.20**<br>***p = 0.045*** |
| *Cpeb2* | 33.61 ± 5.785 | 32.95 ± 11.52 | 43.79 ± 7.09 |
| *Crem* | 1.023 ± 0.076 | 1.814 ± 0.581 | 1.678 ± 0.234<br>*p = 0.07* |
| *Epha7* | 20.09 ± 3.00 | 43.57 ± 20.72 | **38.52 ± 3.01**<br>***p = 0.0081*** |
| *Fabp7* | 22.5 ± 2.47 | 23.4 ± 4.72 | 34.5 ± 6.4 |
| *Fubp1* | 43.94 ± 3.26 | 122.4 ± 28.9<br>*p = 0.542* | **140.1 ± 19.8**<br>***p = 0.010*** |
| *Fubp3* | 5.598 ± 0.893 | 5.720 ± 1.57 | 7.443 ± 0.821 |
| *Gap43* | 710.2 ± 44.0 | **1080 ± 125**<br>***p = 0.049*** | **1335 ± 180**<br>***p = 0.042*** |
| *Gdap1* | 24.78 ± 4.50 | 24.75 ± 6.63 | 43.34 ± 6.90 |
| *Gpm6a* | 739.0 ± 43.4 | 811.7 ± 175.6 | **1438 ± 97**<br>***p = 0.0021*** |
| *Gpm6b* | 215.1 ± 13.4 | 296.3 ± 67.4 | **353.5 ± 37.0**<br>***p = 0.0279*** |
| *Itgb8* | 9.46 ± 2.14 | 9.272 ± 3.568 | 12.75 ± 2.24 |
| *Kcn2* | 8.07 ± 0.66 | 7.547 ± 1.98 | 9.412 ± 0.865 |
| *Kif15* | 0.0997 ± 0.017 | 0.169 ± 0.096 | 0.207 ± 0.101 |
| *Ncam2* | 29.91 ± 1.83 | 32.45 ± 7.89 | 39.55 ± 6.13 |
| *Ndufs4* | 94.73 ± 6.70 | 128.5 ± 17.4 | 143.3 ± 19.8 |
| *Nrn1* | 1038 ± 40 | 995.7 ± 232.9 | 1393 ± 196 |
| *Ogt* | 20.07 ± 1.45 | 17.66 ± 3.46 | 22.12 ± 3.74 |
| *Pcdhb* | 4.390 ± 0.618 | 5.514 ± 0.988 | 6.535 ± 0.654<br>*p = 0.07* |
| *Rac1* | 135.6 ± 12.1 | 209.7 ± 42.3 | **269.5 ± 36.8**<br>***p = 0.031*** |
| *Reep1* | 23.72 ± 6.13 | 30.92 ± 11.57 | 29.75 ± 3.74 |
| *Rhot1* | 6.13 ± 0.91 | 8.80 ± 0.65 | 11.01 ± 1.67<br>*p = 0.069* |
| *Robo1* | 6.98 ± 1.01 | 8.773 ± 0.764 | 9.477 ± 1.517 |
| *Rtn4* | 227.5 ± 31.5 | 238.4 ± 44.9 | 307.1 ± 22.44<br>*p = 0.087* |
| *Scn3a* | 4.55 ± 0.73 | 4.407 ± 1.193 | 6.715 ± 0.852 |
| *Slc1a2* | 535.9 ± 75.9 | **789.9 ± 46.1**<br>***p = 0.0460*** | **885.6 ± 89.90**<br>***p = 0.037*** |
| *Slc4a10* | 45.12 ± 12.43 | 55.36 ± 0.57 | 61.40 ± 14.92 |
| *Snap25* | 673.6 ± 51.4 | **918.4 ± 27.2** | **1198 ± 110** |

| | | | |
|---|---|---|---|
| [illegible] | [illegible] | ***p* = 0.0018** | ***p* = 0.0027** |
| *Shh* | 0.754 ± 0.151 | 0.894 ± 0.272 | 0.831 ± 0.070 |
| *Syncrip* | 21.42 ± 3.50 | 28.99 ± 1.99 | 28.96 ± 0.84 |
| *Tor1a* | 15.68 ± 1.95 | 16.49 ± 3.30 | 20.92 ± 2.42 |
| *Usp33* | 24.77 ± 6.21 | 28.69 ± 13.91 | 35.38 ± 4.25 |
| *Wnt5b* | 0.400 ± 0.114 | 0.458 ± 0.113 | 0.663 ± 0.166 |

mRNA copy number per ng input RNA determined from RTddPCR and normalized to *Gapdh* mRNA is shown as mean ± SEM. These data correspond to the heat maps shown in Figure 1D with significant differences shown in blue font ($N \geq 3$; indicated p values from Student's *t* test vs. *Khsrp* $^{+/+}$; p values $<0.05$ are shown).

**Supplemental Table S6: RTddPCR validation of KHSRP-target mRNAs in hippocampus.**
Values significantly different from *Khsrp-/-* mice are shown in bold blue font.

| mRNA | *Khsrp* $^{+/+}$ | *Khsrp* $^{+/-}$ | *Khsrp* $^{-/-}$ |
|---|---|---|---|
| *Arhgap5* | 104.5 ± 6.3 | 122.3 ± 25.0 | **175.2 ± 22.7** ***p = 0.049*** |
| *Atrx* | 56.15 ± 1.6 | 84.16 ± 21.96 | **85.77 ± 3.47** ***p = 0.0010*** |
| *Cadps* | 69.01 ± 9.15 | 87.07 ± 14.54 | 71.80 ± 16.22 |
| *Chm* | 38.94 ± 1.65 | 70.60 ± 12.73 *p = 0.069* | **90.07 ± 10.55** ***p = 0.0096*** |
| *Epha7* | 101.2 ± 34.0 | 175.0 ± 67.6 | **270.9 ± 39.7** ***p = 0.031*** |
| *Fubp1* | 225.0 ± 53.1 | 362.8 ± 31.9 *p = 0.090* | **517.6 ± 32.0** ***p = 0.0041*** |
| *Gap43* | 326.5 ± 64.3 | **949.5 ± 145.9** ***p=0.0174*** | **1313 ± 297** ***p = 0.031*** |
| *Gpm6a* | 922.4 ± 79.3 | 1551 ± 216.0 *p = 0.052* | 1234 ± 131 |
| *Gpm6b* | 383.7 ± 27.3 | 461.3 ± 94.9 | **519.6 ± 35.7** ***p = 0.039*** |
| *Nrn1* | 1113 ± 188 | 1125 ± 338.6 | 2272 ± 644 |
| *Rac1* | 241.6 ± 82.2 | 362.9 ± 47.4 | 313.1 ± 37.8 |
| *Slc1a2* | 1084 ± 149 | 1461 ± 105.1 | **1772 ± 178** ***p = 0.041*** |
| *Slc4a10* | 54.5 ± 20.4 | 25.61 ± 7.67 | 99.77 ± 7.06 |
| *Snap25* | 764.8 ± 147.9 | 1624 ± 364 *p = 0.093* | **1833 ± 133** ***p = 0.0031*** |

mRNA copy number per ng input RNA determined from RTddPCR and normalized to *Gapdh* mRNA is shown as mean ± SEM. These data correspond to the heat maps shown in Figure 1E with significant differences shown in blue font (N ≥ 3; indicated p values from Student's t test vs. *Khsrp* $^{+/+}$; p values <0.05 are shown).

**Supplemental Table S7: RTddPCR validation of KHSRP-target mRNAs in cortical neuron cultures.**
Values significantly different from *Khsrp-/-* mice are shown in bold blue font.

| mRNA | Cell body preparations | | | Neurite preparations | | |
|---|---|---|---|---|---|---|
| | *Khsrp* $^{+/+}$ | *Khsrp* $^{+/-}$ | *Khsrp* $^{-/-}$ | *Khsrp* $^{+/+}$ | *Khsrp* $^{+/-}$ | *Khsrp* $^{-/-}$ |
| *Arhgap5* | 231 ± 23.6 | 221 ± 25.5 | **338 ± 20.4** ***p = 0.012*** | 248 ± 2.32 | 203 ± 17.3 | 318.9 ± 31.2 |
| *Atrx* | 127 ± 14.9 | 137 ± 9.9 | 157.5 ± 5.9 | 79.0 ± 5.0 | 96.9 ± 2.3 | **170.5 ± 31.5** ***p = 0.029*** |
| *Cadps* | 172 ± 11.8 | 142 ± 21.9 | 154 ± 12.0 | 71.2 ± 3.4 | 96.8 ± 11.8 | 107 ± 20.6 |
| *Chm* | 109 ± 10.0 | 124 ± 9.5 | **160 ± 12.0** ***p = 0.017*** | 69.6 ± 3.9 | 75.3 ± 5.6 | 96.6 ± 18.4 |
| *Epha7* | 153 ± 11.8 | 199 ± 53.6 | 316 ± 81.8 | 55.9 ± 8.95 | 63.3 ± 12.6 | 55.3 ± 13.0 |
| *Fubp1* | 289 ± 31.0 | 533 ± 113 | **3001 ± 947** ***p = 0.046*** | 85.6 ± 7.81 | **278 ± 47.2** ***p = 0.0159*** | **281 ± 45.1** ***p = 0.013*** |
| *Gap43* | 5085 ± 1106 | 8361 ± 459 | 7777 ± 441 | 53.1 ± 35.3 | **252 ± 54.7** ***p = 0.038*** | **301 ± 14** ***p = 0.0028*** |
| *Gpm6a* | 3129 ± 462 | 4180 ± 73.2 | **5094 ± 649** ***p = 0.049*** | 60.0 ± 2.4 | 68.0 ± 4.93 | 85.0 ± 19.8 |
| *Gpm6b* | 465 ± 49.8 | 499 ± 49.3 | 578 ± 40.9 | 109 ± 6.1 | 110 ± 11.4 | 142 ± 29.2 |
| *Slc1a2* | 780 ± 93 | 2382 ± 591 | **3745 ± 902** ***p = 0.039*** | 118 ± 3.15 | 160 ± 9.94 | 171 ± 28.9 |
| *Snap25* | 1548 ± 233 | 1562 ± 126 | 2041 ± 252 | 49.0 ± 2.9 | **68.8 ± 1.59** ***p = 0.010*** | **111 ± 19.8** ***p = 0.0104*** |

mRNA copy number per ng input RNA determined from RTddPCR and normalized to *GAPDH* is shown as mean ± SEM.
These data correspond to the heat maps shown in Figure 3C with significant differences shown in blue font (N ≥ 4; indicated p values from Student's *t* test vs. *Khsrp* $^{+/+}$ within each type of RNA preparation).

**Supplemental Table S8: Behaviorial analyses of KHSRP deficient mice**

| | Males | | | Females | | |
|---|---|---|---|---|---|---|
| **Behavior** | **Khsrp +/+** | **Khsrp +/-** | **Khsrp --/-** | **Khsrp +/+** | **Khsrp +/-** | **Khsrp --/-** |
| Total Time Grooming (s) | 15.9 | 18.8 | 17.8 | 21.2 | 14.8 | 20.9 |
| Latency to first Groom (s) | 185.8 | 186.4 | 164.1 | 134.4 | 115.4 | 113.2 |
| Average Weight (g) | 23.7 | 22.6 | 23.1 | 23.2 | 22.1 | 22.8 |
| **Clear Box Behaviors** | | | | | | |
| Sniffing | 100 | 100 | 100 | 100 | 100 | 100 |
| Abnormal Gait | 7 | 0 | 17 | 0 | 0 | 23 |
| Circling | 20 | 40 | 75* | 23 | 20 | 46 |
| Jumping | 27 | 20 | 25 | 0 | 20 | 23 |
| Freezing | 27 | 20 | 33 | 23 | 40 | 46 |
| Wild running | 0 | 0 | 17* | 0 | 0 | 15* |
| Defecation | 67 | 40 | 75 | 69 | 20 | 15* |
| Urination | 13 | 20 | 17 | 8 | 20 | 15 |
| Rearing Count | 95 | 103 | 100 | 87 | 80 | 103 |

Data are percentage of animals that displayed a response unless units are given in parenthesis.
* indicates significance of $p<0.05$ from Khsrp +/+ controls. Tukey post-hoc test after one way ANOVA
N= 15 Khsrp+/+ Males (M) and 13 WT Females (F), 5 Khsrp+/- M, 5 Khsrp+/- F, 12 Khsrp-/- M and 13 Khsrp-/- F.

**Supplemental Table S9: Primer sequences used for RTddPCR.**
All are listed as 5’ to 3’ direction with forward (sense) sequences on 1st lines and reverse (antisense) sequences on 2nd line of each row.

| mRNA | Forward and Reverse Primers (5’ à 3’) |
|---|---|
| *Argap5* | CTT CTC AGC TGT AAC CAG ATC C<br>GAA GAA AAC CCA CAA AGT AAA GGA |
| *Cacna2d1* | AGC GTT GCT GTT AAA TGA AGC<br>AGG ACA TGC TCA TTC TGG TG |
| *Cadps* | ACC ATT AAG TCT GAC CAC CAC<br>CTG AAC TAG TCA TCG AAG TAC TGC |
| *Calb1* | TTT TCC ATC ATC TCT CTG TCC AT<br>AGC TGC AGA ACT TGA TCC AG |
| *Chm* | TCG TTC TAA ATC TTC TCT TGC TGT<br>CTG TTC GAG TCA TTG AGT TAT GC |
| *Cpeb2* | CTC GTC ACA CAT CTG GTC ATC<br>CAT CAG TGC TCG ATT TGT TCA G |
| *Crem* | CAG CCA TCA CCA CAC CTT<br>CAC CAC CTA ACA TTG CTA CCA |
| *Epha7* | GGA GTT TCG GAC TTA AGC AGA T<br>CAG TAC TCG CTC CTT CAT GAC |
| *Fabp7* | CAT AAC AGC GAA CAG CAA CG<br>GAA GTG GGA TGG CAA AGA AAC |
| *Fubp1* | GCA CCA GCT ACA ACC CAA<br>GCC TTT GTA TAA TCA ACC TGT CC |
| *Fubp3* | CCA TTC CGA CAT CGA TCC AC<br>ATT CAG ATT GCC TCA GAG AGT T |
| *Gdap1* | CGC ATA AAC CAA GGC TCA TTG<br>GAC GCA CTC CTT CAG CTC |
| *Gpm6a* | ACT TGA TGT CTT CGT ACT TCT GC<br>GAC CTT CCA CTT GTT CAT TGT G |
| *Gpm6b* | GTA GTA GCC ATC ATG TAG ATC AGC<br>GAA CAT CTG CAA CAC GAA TGA G |
| *Itgb8* | TTC CCA GCC ACT AAA GCA C<br>ACT TCT CCT GTC CCT ATC TCC |
| *Kcnc2* | TTG TGT CTC CTT TAG TCT GTG C<br>AAC GTG TTT CCT GTT GAC GA |
| *Kif5* | CTT CAG CTG TTG ACT GTC TGG<br>CCT GGA GTC TCA GCA ATT AAG AG |
| *Ncam2* | TCA CCC ACA CTA AGC TCT ACT<br>TCC TCT CCT TCT ACC TGC TC |
| *Ndufs4* | CTG GTT TTG ATG TGC TCT TCT G<br>TCC GTC TGT AGA GTT CCA TCC |
| *Ogt* | GCC AAC TCA GCT AAC CCT T<br>GCA GTA GAA GTC AAT TAC CCC TT |
| *Pcdhb* | GTA GAC ATC AAC TTC CGA CCT C<br>CTA TGG TCA GTT CTT ATG CTA TGG A |
| *Prok2* | CTC TTA ACC CAG ATA CTG ACA GC<br>CTA CTG CTA CTT CTG CTG CTA C |
| *Rac1* | TCC ATC TAC CAT AAC ATT GGC A<br>GCT CAT CAG TTA CAC GAC CA |
| *Reep1* | ACA AGG GCG TCA TAG CTT C<br>TGT TCA TCC CAC ATT GTC TTC A |
| *Rhot1* | TGA CCA GCG ACA TAA TCA GTG<br>AGC CTC CGA CAT GAA GAA AG |
| *Rtn4* | GTT CAC ATG ACC AAG AGC AGA<br>GTG TGA TCC AAG CTA TCC AGA A |
| *Scn3a* | CTT AGC CTT CTC TTC CTG ACT TC |

| | |
|---|---|
| [illegible] | CAG CAC CTT TGA AGA TAG CGA |
| *Slc1a2* | AAA GAA TCG CCC ACC ACA T<br>CCA TGC TCC TCA TTC TCA CAG |
| *Slc4a10* | CCA TCT TCC ACA TCC TCT TCA<br>TCC CTC ATG ACC TTT TCA CAG |
| *Snap25* | CAA ATT TAA CCA CTT CCC AGC A<br>CAG AAT CGC CAG ATC GAC AG |
| *Ssh* | CGT AAG TCC TTC ACC AGC TTG<br>GAA TCC AAA GCT CAC ATC CAC |
| *Syncrip* | CAT AGC CAC CTC TAC CTC CA<br>CTA AGC CAC CAG ATC AGA AGA G |
| *Tora1a* | CAT GCA TCT TGT CCA TCT CAT C<br>CCA CGC CTC TAA CAT CAC AC |
| *Usp33* | TGC TGA TAA CCG TGG ACT AAC<br>GTG AAA CAG TTG ACC TAA ACA GC |
| *Wnt5b* | AAG GCA GTC TCT CGG CTA<br>CCA ACA CCA GTT TCG ACA GA |
| *Gap-43* | AAC GGA GAC TGC AGA AAG C<br>TCA GGC ATG TTC TTG GTC AG |
| *Nrn1* | ACGAACATCAAGACCGTGTG<br>GTTCGAATAAGCTGCCTTGG |
| *Hmgb1* | CATGGGCAAAGGAGATCC<br>CTCTGAGCACTTCTTGGAG |
| *Actb* | CTG AAC CCT AAG GCC AAC C<br>GTA CGA CCA GAG GCA TAC AG |
| *12s Ribosomal RNA* | GGC TAC ACC TTG ACC TAA CG<br>CCT TAC CCC TTC TCG CTA ATT C |
